# Chromatin signalling pathways and FANCE amplification affect ATR inhibitor sensitivity in metastatic breast cancer

**DOI:** 10.1101/2025.08.26.672363

**Authors:** Jogitha Selvarajah, Rayzel Fernandes, Marc Lorentzen, Emma Pei, Kyle Greenland, Zuza Kozik, Jyoti Choudhary, Charlotte Bevan

## Abstract

Metastatic breast cancer remains incurable, as many patients develop therapy resistance. Loss of ATM/p53 increases reliance on the ATR pathway, positioning ATR inhibitors (ATRi) as promising therapeutics. However, targeting a single pathway often leads to resistance due to tumour heterogeneity or alternative signalling mechanisms. Here, we combine chromatin-enrichment proteomics and phospho-proteomics with genome sequencing data in the ATRi-sensitive triple negative breast cancer (TNBC) cell line MDA-MB-453 to map adaptive responses to the ATR inhibitor AZD6738. We identify chromatin-associated activation of survival pathways, including AKT1, RPTOR (mTORC1), CDK4/5, and OGFR, alongside hyperphosphorylation of MKI67, SAFB2, and CHD4, indicating ATRi sensitivity. Complementary siRNA screening of DNA damage response (DDR) genes reveals that amplification of Fanconi anaemia (FA) pathway gene FANCE increases sensitivity to AZD6738. Analysis of breast cancer datasets highlights frequent FANCE amplification in metastatic patients, particularly in circulating tumour cells. Strikingly, pharmacological inhibition of the FA pathway (UBE2T/FANCL-IN-1) synergises with AZD6738. Together, our findings define adaptive resistance mechanisms to ATR inhibition and nominate FA pathway blockade as a rational combination strategy. Overall, our work provides fundamental insight into the complexity of DDR in metastatic breast cancer and offers a platform for mechanistic investigation, which can be exploited in cancer therapy.

## INTRODUCTION

Breast cancer is the most common cancer affecting women worldwide. Advances in early diagnosis and treatment strategies have significantly improved patient outcomes; however, therapeutic resistance and metastatic disease remain major clinical challenges. Despite initial treatment, 20–30% of patients develop metastases, which are largely incurable and associated with a poor five-year survival rate of approximately 25%^1^.

Triple-negative breast cancer (TNBC), a particularly aggressive subtype, is defined by the absence of oestrogen receptor (ER), progesterone receptor (PR), and HER2 expression. TNBC disproportionately affects younger women and women of African descent, and is often resistant to standard therapies. Up to 25% of TNBC patients relapse with metastatic disease after initial treatment^2^, underscoring the urgent need for novel therapeutic strategies to overcome resistance and prevent recurrence.

ATR (ataxia telangiectasia and Rad3-related) is a serine/threonine kinase in the phosphatidylinositol 3-kinase-related kinase (PIKK) family, which also includes ATM, DNA-PKcs, and mTOR. ATR is activated by replication fork stalling during DNA replication, typically due to replication stress or genotoxic insults^3, 4^. Upon sensing single-stranded DNA coated with replication protein A (RPA), ATR is recruited via ATRIP (ATR Interacting Protein) and activated by TopBP1 (DNA topoisomerase 2-binding protein 1). This triggers a downstream DNA damage response (DDR) cascade involving phosphorylation of H2AX (γH2AX) and activation of key effectors such as Chk1 and p53. These signalling events promote DNA repair, stabilise replication forks, and induce cell cycle arrest or apoptosis, depending on the extent of damage.

While ATR primarily responds to single-stranded DNA damage, ATM is activated by double-stranded DNA breaks. Cancer cells that lose ATM or p53 function and thus the ability to arrest the cell cycle and repair double-strand breaks often become dependent on ATR signalling to manage replication stress and maintain viability^5^. Therefore, inhibiting ATR in such contexts promotes unrestrained cell cycle progression despite unresolved replication stress, leading to replication fork collapse, mitotic catastrophe, and cell death^6^. Although this synthetic lethality framework has strong preclinical support, clinical trials using ATR inhibitors have not yet demonstrated clinical benefit, likely due to compensatory signalling mechanisms and tumour heterogeneity^7^. Understanding these resistance pathways is critical for improving patient selection and optimising therapeutic combinations.

AZD6738 is a selective ATR inhibitor currently in clinical development, including trials in breast cancer^8, 9, 10^. Since breast tumours frequently exhibit loss of ATM or CHEK2, many are hypothesised to rely on ATR to survive DNA damage^11^. However, clinical trials combining AZD6738 with the PARP inhibitor Olaparib in TNBC have thus far failed to demonstrate meaningful improvement in patient outcomes^12^. These findings highlight the complexity of ATR inhibitor responses and the need to identify additional determinants of sensitivity and resistance.

Chromatin-associated proteins play critical roles in coordinating transcription, cell cycle progression, DNA repair, and replication functions that are tightly regulated in response to DNA damage^13, 14, 15^. Upon ATR inhibition, alterations in chromatin-bound signalling may dictate cellular fate. To investigate this, we applied a chromatin enrichment proteomics approach to dissect chromatin-associated and DNA-bound protein dynamics following AZD6738 treatment. Compared to whole-cell proteomics, chromatin enrichment enables more sensitive detection of low-abundance nuclear proteins specifically involved in DNA damage responses^16^. We used tandem mass tag (TMT) based quantitative proteomics^17^ to compare untreated and AZD6738-treated samples, alongside phospho-proteomics to capture key phosphorylation events, which are central to DDR signalling. ATR inhibition in MDA-MB-453 cells, a model of metastatic TNBC, leads to robust upregulation of chromatin-bound proteins and phospho-sites involved in DNA repair and survival signalling. Complementary siRNA screen targeting genetically altered DDR genes revealed that amplification of Fanconi anaemia DNA repair pathway genes, FANCE, increases sensitivity to AZD6738. FANCE is a member of the Fanconi anaemia (FA) DNA repair pathway. The FA pathway plays a central role in repairing interstrand cross-links (ICLs) that block replication and transcription. ICL repair is initiated by the FANCM-MHF1/2 complex, which recruits and activates the multi-subunit FA core complex, including FANCA, FANCB, FANCC, FANCE, FANCF, FANCG, FANCL, and associated proteins^18, 19, 20^. This complex monoubiquitinates the FANCD2/FANCI heterodimer (ID2 complex), which orchestrates downstream repair via homologous recombination (HR) or NHEJ. While loss-of-function mutations in FA genes cause the inherited disorder Fanconi anaemia and predispose to cancer^21^, activation or amplification of FA genes in tumours has been associated with resistance to genotoxic therapy^22^. In our analysis of metastatic breast cancer samples, we found frequent co-amplification of FANCE and FANCG, especially in circulating tumour cells (CTCs) from blood, suggesting a possible role in promoting metastasis and therapeutic resistance. Notably, we demonstrate that pharmacologic inhibition of the FA pathway using UBE2T/FANCL-IN-1 synergises with AZD6738 in MDA-MB-453 cells. These findings indicate that targeting the FA pathway in combination with ATR inhibition could be an effective strategy to overcome resistance and potentially suppress metastatic progression in TNBC.

## RESULTS

### The MDA-MB-453 cell line is highly sensitive to the ATR inhibitor AZD6738

AZD6738 (Ceralasertib), an ATR-specific inhibitor in clinical trials for multiple cancers, was tested for its effects on cell viability on six metastatic breast cancer cell lines using a 4-day crystal violet assay (Figure 1A). T47D cells were resistant, whereas the other five lines were sensitive to varying degrees. The MDA-MB-453 cell line showed the greatest sensitivity (IC₅₀ = 0.59 µM). MDA-MB-453, a triple-negative breast cancer (TNBC)-like model (low HER2, ER⁻/PR⁻)^23^. The IC₅₀ values indicate that T47D is resistant to AZD6738, while the other five cell lines are sensitive to varying degrees. HCC1954, MDA-MB-436, and MDA-MB-231 are also models of TNBC; the other two cell lines, T47D and MCF-7, are ER-positive.

**Figure 1.**
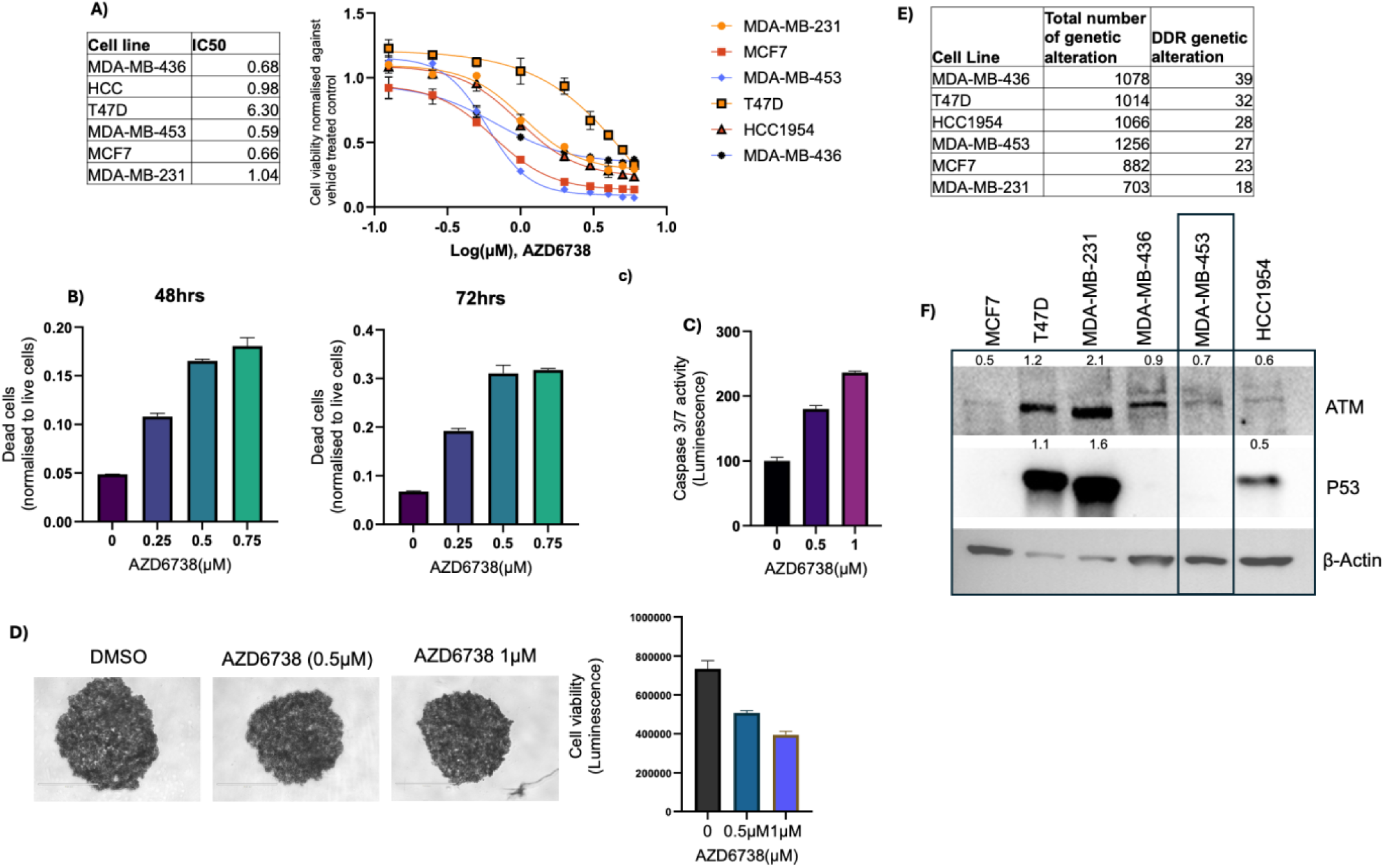
MDA-MB-453 cell line is highly sensitive to AZD6738. A) Six metastatic breast cancer cell lines were treated with increasing concentrations of AZD6738 for 4 days, and cell viability was measured by crystal violet (n=2). IC50 values were calculated on GraphPad Prism. B) MDA-MB-453 cells were treated with increasing concentrations of AZD6738 for 48 and 72 hrs, and CytoTox-Glo Cytotoxic Assay measured dead cells. Mean calculated from three wells ±SEM. C) MDA-MB-453 cells were treated with increasing concentrations of AZD6738 for 24 hours, and the Caspase-Glo 3,7 Assay System measured caspase 3/7 activity. Mean calculated from three wells ±SEM. D) MDA-MB-453 cells were seeded and left to form spheroids for 4 days, and treated with 0.5 and 1µM AZD6738 for 48 hours, and cell viability was measured with Promega CellTiter Glo. Mean calculated from three wells ±SEM (images representative of 3 experiments). E) Genome sequencing data were obtained from the Cell Model Passport database. The table shows the total number of genetically altered genes and the number of genetically altered genes in the DNA damage response signalling pathway for each cell line. F) Whole-cell lysates of the six breast cancer cell lines were analysed by Western blot for total ATM and p53. Using ImageJ, each blot was normalised to beta-actin.

To further examine the effects of ATR inhibition in MDA-MB-453 cells, a cytotoxicity assay was performed, confirming a dose- and time-dependent increase in cell death at 48 h and 72 h (Figure 1B). Caspase 3/7 activity assays demonstrated a dose-dependent induction of apoptosis within 24 h (Figure 1C). In 3D culture, AZD6738 treatment reduced spheroid size and viability (Figure 1D).

To explore how DDR signalling is altered in these cell lines, we obtained genomic sequencing data from the Cell Model Passport database^24^ and selected the altered DDR genes (Supplementary Table 1). We utilised the DDR list of genes compiled by Horizon Discovery, which comprises 240 DDR genes, including those involved in the DDR signalling pathway and DNA repair. Figure 1E shows the total number of genetic alterations in each cell line and the number of genetic alterations in DDR genes. Since ATM is known to be a synthetic lethality partner of ATR inhibitors^5^, we analysed the genetic alteration status. AZD76738-sensitive cell lines MCF7, HCC1954, and MDA-MB-436 all exhibit partial loss of ATM (decrease in copy number). We performed western blots of ATM protein expression levels in the six cell lines (Figure 1F), which shows that these cell lines have low ATM protein expression, compared to T47D and MDA-MB-231 with WT ATM. MDA-MB-453 cells have an uncharacterised missense mutation in ATM, but exhibit low ATM protein expression levels, similar to MCF7 and HCC1954. TP53 is also deleted in MDA-MB-453; therefore, the low ATM protein expression and the deleted TP53 may contribute to AZD6738 sensitivity in MDA-MB-453 cells.

### ATR inhibition remodels the chromatin and phospho-chromatin landscape in MDA-MB-453 cells

Having identified MDA-MB-453 as the most sensitive cell line to AZD6738, we next examined how ATR inhibition alters signalling pathways at the chromatin. ATR inhibition by AZD6738 induces DSB DNA damage; therefore, we focused on DNA-bound and chromatin-associated proteins. Chromatin fractions from untreated and AZD6738-treated cells were isolated, enriched, and labelled with tandem mass tags (TMT) for quantitative mass spectrometry (Figure 2A). To assess post-translational signalling changes, we also performed phospho-proteomics on the chromatin fraction. A portion of the trypsinised peptides was enriched for phospho-peptides using immobilised metal affinity chromatography (IMAC), enabling the identification of low-abundance phosphorylation events involved in DNA damage signalling^25^.

**Figure 2.**
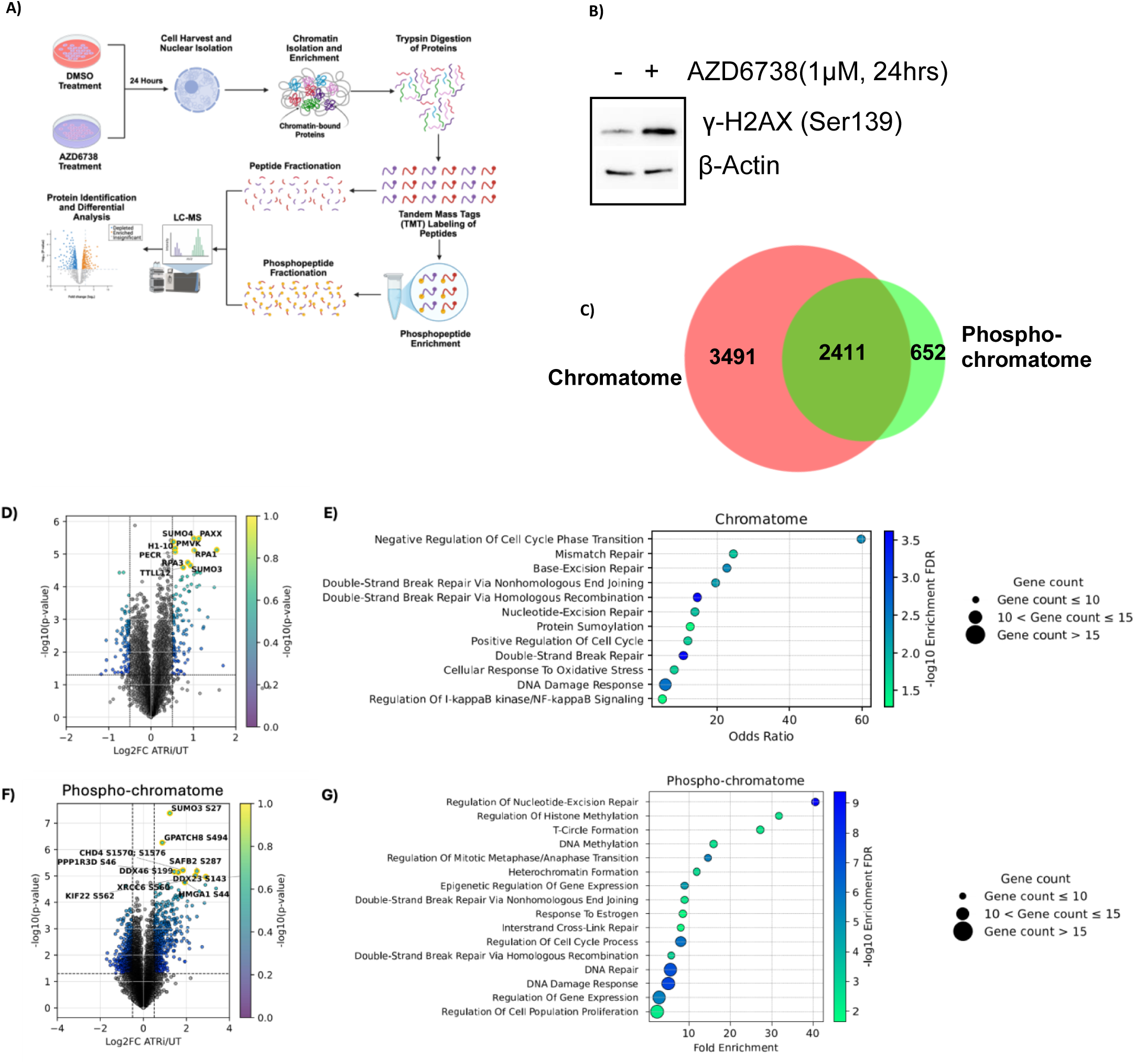
Chromatin proteomics and phospho-proteomics of AZD6738-treated MDA-MB-453 cells. Nuclear fractionation of untreated and AZD6738-treated (24 h) MDA-MB-453 cells was performed, followed by chromatin fractionation and enrichment, peptide digestion, TMT labelling and protein identification using LC-MS. Chromatin fraction peptides were further enriched for phospho-peptides by IMAC purification and were identified by LC-MS. A) Schematic diagram showing the experimental steps. B) Western blot of MDA-MB-453 whole cell lysate, untreated and treated with AZD6738 for 24 hours, and probed for phosphorylated H2AX. C) Venn diagram illustrating the total number of proteins that overlap between the proteins identified by chromatin proteomics and phospho-proteomics and the total number of proteins specific to chromatin proteomics or phospho-proteomics (Biovenn). The volcano plots show annotated selected significantly upregulated DDR proteins (D) chromatome and (F) phospho-chromatome. Enrichment dot plot: GOBP enrichment based on significantly upregulated (p < 0.05, log2FC > 0.3) proteins on the (E) chromatome of MDA-MB-453 cells in response to AZD6738 and (G) phospho-chromatome. Selected significantly enriched processes were plotted (BH FDR < 5%), three biological repeats.

Western blotting confirmed that the treatment conditions induced DNA damage, as shown by H2AX phosphorylation (Figure 2B). Chromatin enrichment proteomics identified >5,000 proteins, with 402 significantly upregulated and 300 downregulated after AZD6738 treatment (p < 0.05, fold change > 0.3 or < −0.3) (Supplementary Table 2). Phospho-proteomics identified >12,000 phospho-sites (3,063 proteins), with 942 upregulated and 1,307 downregulated (Supplementary Table 3). Approximately 650 proteins were detected only via phospho-proteomics, underscoring its value in capturing low-abundance chromatin-associated proteins (Figure 2C).

Volcano plots for both the chromatome and phospho-chromatome (Figures 2D and 2F) illustrate the distribution of proteins between untreated and AZD6738-treated samples. Among the most significantly upregulated proteins are SUMO and RPA family members; SUMOylation is activated by diverse forms of DNA damage^26^, whereas RPA proteins are well-established markers of replication stress and single-stranded DNA accumulation^27^. Consistently, pathway enrichment analyses (Figures 2E and 2G) revealed activation of DNA damage response, DNA repair, cell cycle, and chromatin regulation pathways.

To map these changes onto specific DNA repair mechanisms, we integrated our proteomics data with genomic information from the Cell Model Passport database^24^. We examined six repair pathways: NHEJ, HR, MMR, NER, FA, and BER, using curated protein lists from the MD Anderson Human DNA Repair Genes resource^28^ (Figure 3). Upregulated proteins and phospho-sites were detected in all pathways except BER, with NHEJ showing the highest enrichment of phospho-sites (Figure 3). Several low-abundance proteins, particularly in the FA pathway, were detected only through phospho-proteomics. Genomic profiling revealed no alterations in NHEJ or MMR genes, but multiple defects in HR and FA genes, with mostly amplifications in the FA pathway (BRCA2, FANCE, FANCG, BRIP1, FANCM, PALB2) (Supplementary Table 1). As the proteomics data indicate prominent upregulation of NHEJ by AZD6738 and with HR deficiency due to BRCA1 loss^29^, NHEJ appears to be the preferred DNA repair pathway under replication stress in these cells. This potential vulnerability could be exploited to enhance AZD6738 sensitivity.

**Figure 3.**
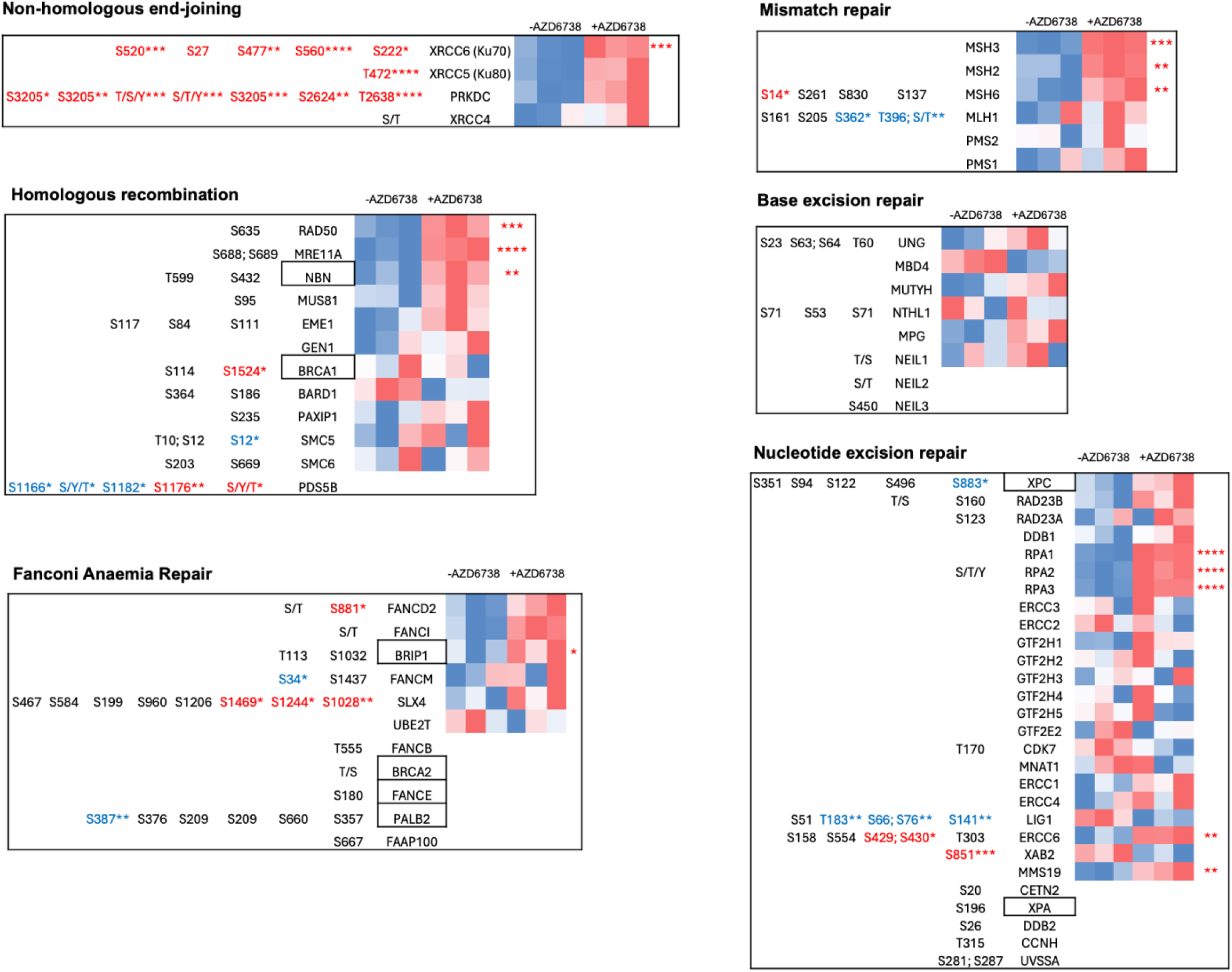
DNA repair pathways activated by AZD6738 treatment in MDA-MB-453 cells. A) Main DNA repair pathway genes identified by chromatin enrichment proteomics and phospho-proteomics were grouped and integrated with genetic alterations in the MDA-MB-453 cell line. Heat map showing untreated/AZD6738-treated chromatin proteomics data, with selected phospho-sites identified for each DNA repair pathway. Statistical significance shown in stars (Student t-test, n=3, p-value <0.05 *, <0.01**, 0.001***, 0.0001****) (red: upregulation, downregulation: blue, log2FC > 0.3). Genetically altered genes are boxed.

### AZD6738 induces distinct protein and phosphorylation level changes in chromatin-associated factors

To examine chromatin signalling changes linked to AZD6738 sensitivity or resistance, we identified the top 100 chromatin-associated proteins and phospho-sites most significantly upregulated in treated MDA-MB-453 cells. The upregulated chromatin-associated proteins were primarily involved in DNA damage response, transcription regulation, and chromatin remodelling. Notably, several proteins from cell survival pathways, including MAPK, mTOR, and CDK4/5, were also enriched (Figure 4A).

**Figure 4.**
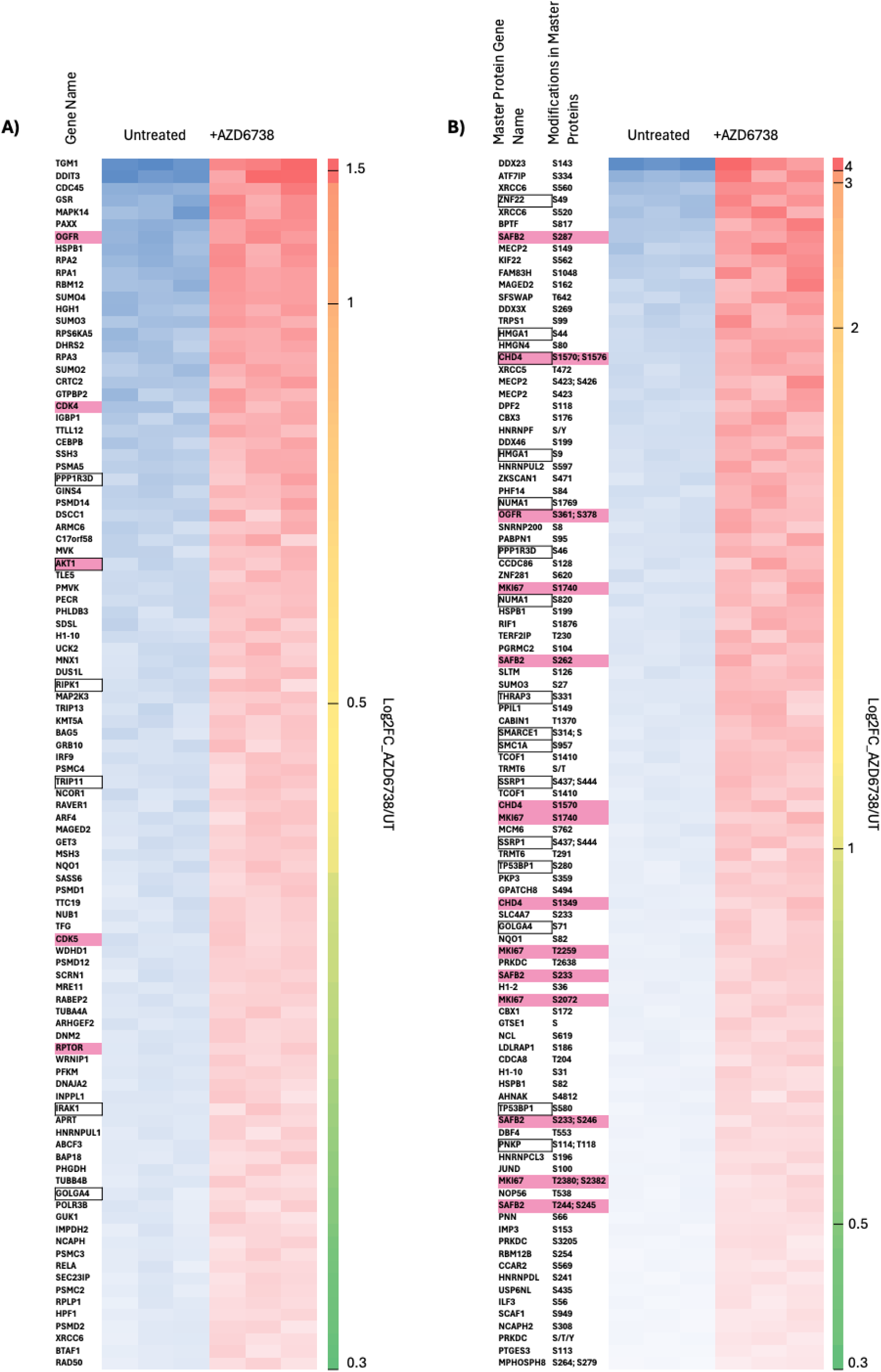
Top 100 upregulated chromatin proteins and phospho-sites. Top 100 statistically significant upregulated (p < 0.05, log2FC > 0.3) A) chromatin enrichment proteins and B) phospho-sites in untreated and AZD6738-treated (24 h) MDA-MB-453 cells. Genetically altered genes are boxed. Proteins/phospho-sites of interest are highlighted in pink.

The top 100 upregulated phospho-sites (Figure 4B) similarly included DNA repair and DDR proteins, transcription regulators, RNA metabolism proteins, proteasome, and chromatin remodelling proteins. Many phospho-sites remain uncharacterized, such as DDX23 phosphorylation at Ser143, which showed a >4-fold increase post-treatment. DDX23, an RNA helicase, is implicated in cancer progression, invasion, and metastasis in multiple malignancies^30, 31, 32^.

MKI67, CDH4, and SAFB2 exhibited multiple upregulated phospho-sites without changes in total protein abundance, suggesting phosphorylation-specific regulation. Ki-67 (MKI67) is a proliferation marker that promotes genome stability during mitosis^33^, while CDH4 participates in chromatin remodelling during DNA repair^34^. SAFB2 is a nuclear protein regulating chromatin organisation, transcription and RNA metabolism. SAFB2 is a multifunctional tumour suppressor; overexpression in breast cancer models suppresses proliferation, migration, and invasion whilst promoting apoptosis^35^. Hyperphosphorylation of these proteins could represent novel indicators of AZD6738 response.

AZD6738 treatment also enhanced HMGA1 phosphorylation at Ser44 and Ser9 without altering protein levels. The HMGA1 gene is amplified in MDA-MB-453 cells and is involved in transcription, heterochromatin organisation, and DNA replication, with established roles in TNBC progression, EMT, and metastasis^36, 37, 38, 39^. Interestingly, OGFR appeared in both the top 100 chromatin-enriched proteins and phospho-sites. OGFR, a receptor for opioid growth factor, regulates cell growth^40^ and may represent an adaptive survival pathway in response to AZD6738.

Collectively, these findings highlight distinct protein and phosphorylation level changes in chromatin - associated factors, many of which are previously uncharacterized, that may contribute to DNA damage adaptation and identify potential biomarkers or therapeutic targets for enhancing ATR inhibitor efficacy in breast cancer.

### mTOR pathway inhibition combines with AZD6738 in ATR inhibitor–sensitive breast cancer cells

Given the enrichment of mTOR pathway components among the top upregulated chromatin-associated proteins, we next investigated whether targeting mTOR could potentiate AZD6738 effects. Raptor (RPTOR) and AKT1 were among the top 100 upregulated chromatin-associated proteins (Figure 4A), and mTOR, Rictor, and AKT2 were also upregulated (Figure 5A). mTOR forms two complexes: mTORC1, containing Raptor, regulates cell proliferation and growth; mTORC2, containing Rictor, regulates cytoskeletal dynamics and cell survival. Although primarily cytoplasmic, mTOR can localise to chromatin, where it regulates transcription in prostate cancer cells^41, 42^. High chromatin mTOR levels correlate with poor prognosis in castration-resistant and metastatic prostate cancer ^41^.

**Figure 5.**
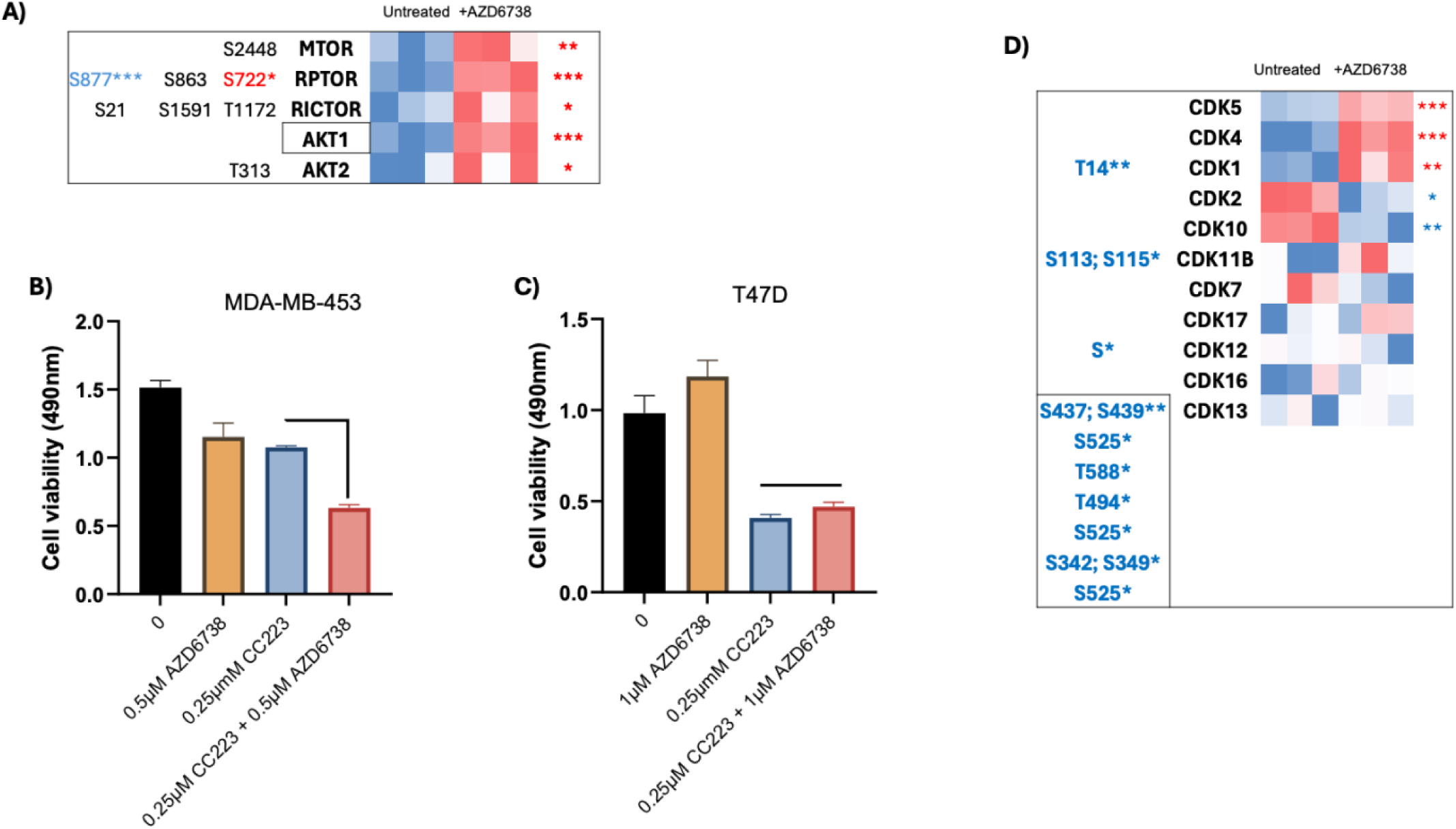
AZD6738 upregulates the mTOR signalling pathway proteins and CKD4/5 at the chromatin. A) Heat map showing proteins and phosphorylations of MTOR, RPTOR, RICTOR, AKT1, and AKT2 before and after AZD6738 treatment identified by chromatin proteomics and phospho-proteomics. B) MDA-MB-453 cells were untreated and treated with CC223, or AZD6738 alone and in combination for 24 hours, and cell viability was measured by CellTiter 96. Mean calculated from three wells ±SEM. B) T47D cells were treated with CC223, or AZD6738 alone or in combination for 72 hrs and cell viability was measured with CellTitre 96. Mean calculated from three wells ±SEM. D) Heatmap showing CDKs up/downregulation after AZD6738 treatment with phosphorylations identified by chromatin proteomics and phospho-proteomics. (blue: downregulated, red: upregulated log2FC > 0.3) (Student t-test n=3, p-value <0.05 *, <0.01**, 0.001***, 0.0001****). Genetically altered genes are boxed.

Previous studies show that mTOR inhibition can sensitise cells to DNA-damaging chemotherapy^43, 44^. We therefore treated cells with the second-generation mTOR kinase inhibitor CC223^45^ alone or with AZD6738 (Figure 5B). The combination significantly reduced MDA-MB-453 cell viability compared with either drug alone, while AZD6738-resistant T47D cells showed no added sensitivity (Figure 5C, Figure 7B). The mechanism by which chromatin-associated mTOR, Raptor, or AKT promotes survival after DNA damage remains unclear, but may involve transcriptional regulation. TORC2 has recently been reported to have a role in base excision repair in yeast^46^, supporting a potential DNA repair connection.

### ATR inhibition selectively enriches CDK4 and CDK5 at chromatin

We also observed selective enrichment of CDK4 and CDK5 at chromatin after AZD6738 treatment, pointing to another potential co-target. CDK4/6 inhibitors are established treatments for early and metastatic breast cancer^47^. CDKs regulate both cell cycle progression and transcription, with over 10 family members having diverse roles^48^. CDK5 participates in the DNA damage response^49^ and is linked to cell motility and EMT, key processes in metastasis^50, 51^. CDK5 inhibitors are in development, particularly for chemotherapy-resistant cancers^49^. CDK4, classically cytoplasmic and controlling G1-S progression, also influences migration and invasion^52^. Mouse studies have shown a chromatin-level transcriptional role for CDK4 in regulating genes involved in chromosome segregation^53^, suggesting a possible transcriptional function after DNA damage. When examining all other CDKs detected via chromatin enrichment and phospho-proteomics (Figure 5D), most were unaffected or downregulated by AZD6738, underscoring the specificity of CDK4 and CDK5 enrichment at chromatin.

### Amplified FANCE increases sensitivity to the ATR inhibitor AZD6738

After analysing global chromatin signalling pathways activated by AZD6738-induced DNA damage, we focused on the DNA damage response (DDR) pathway. We performed an siRNA screen targeting genetically altered DDR genes to assess how these changes affect AZD6738 sensitivity (Figure 6A). Genome sequencing data for MDA-MB-453 cells (Cell Model Passport^24^; 1152 altered genes) were compared with a DDR gene library from Horizon Discovery (240 genes), identifying 26 altered DDR genes in this cell line (Figure 6B). Mutations were classified as missense or nonsense; copy number alterations were categorised as deletion, loss, or gain/amplification (Figure 6C).

**Figure 6.**
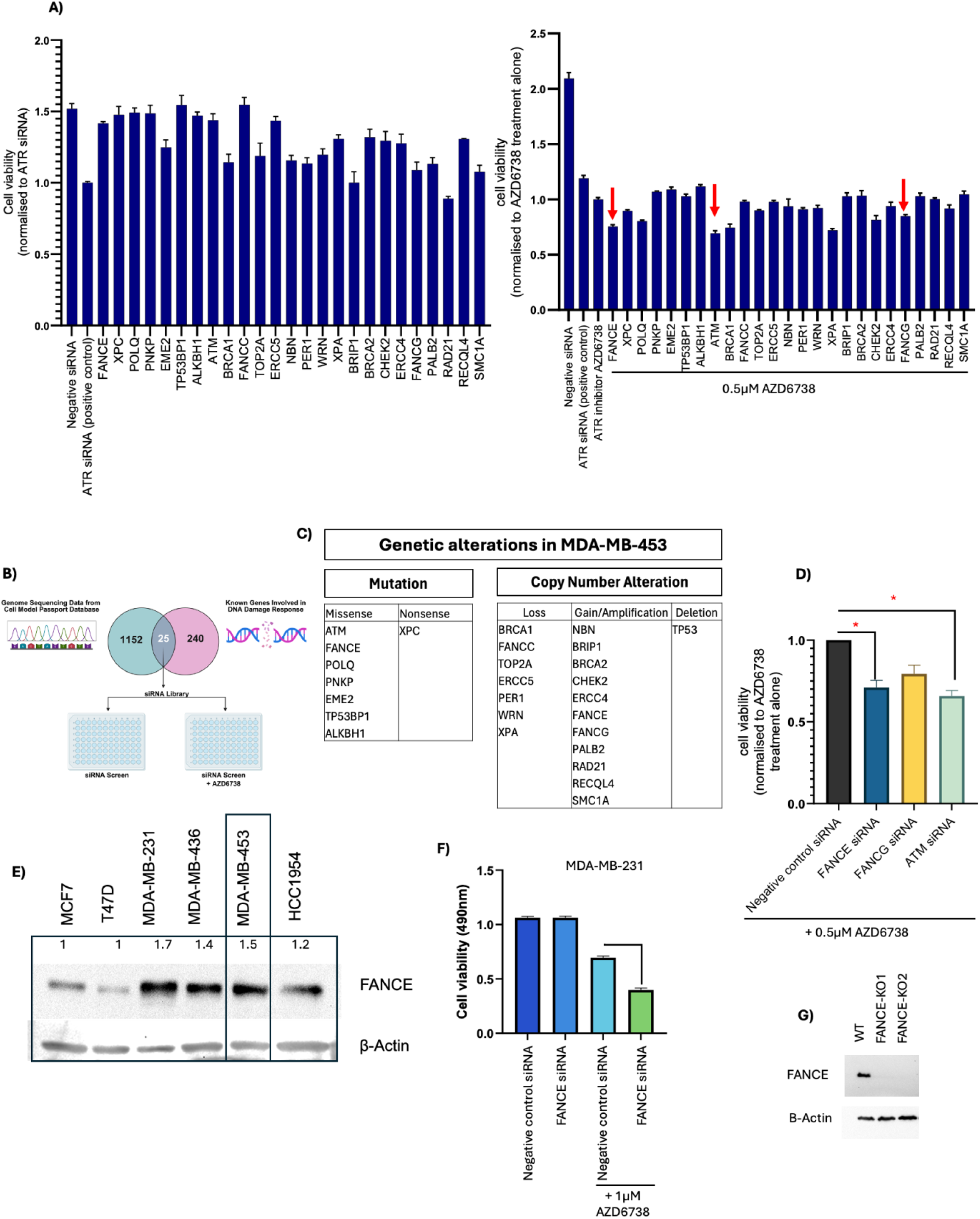
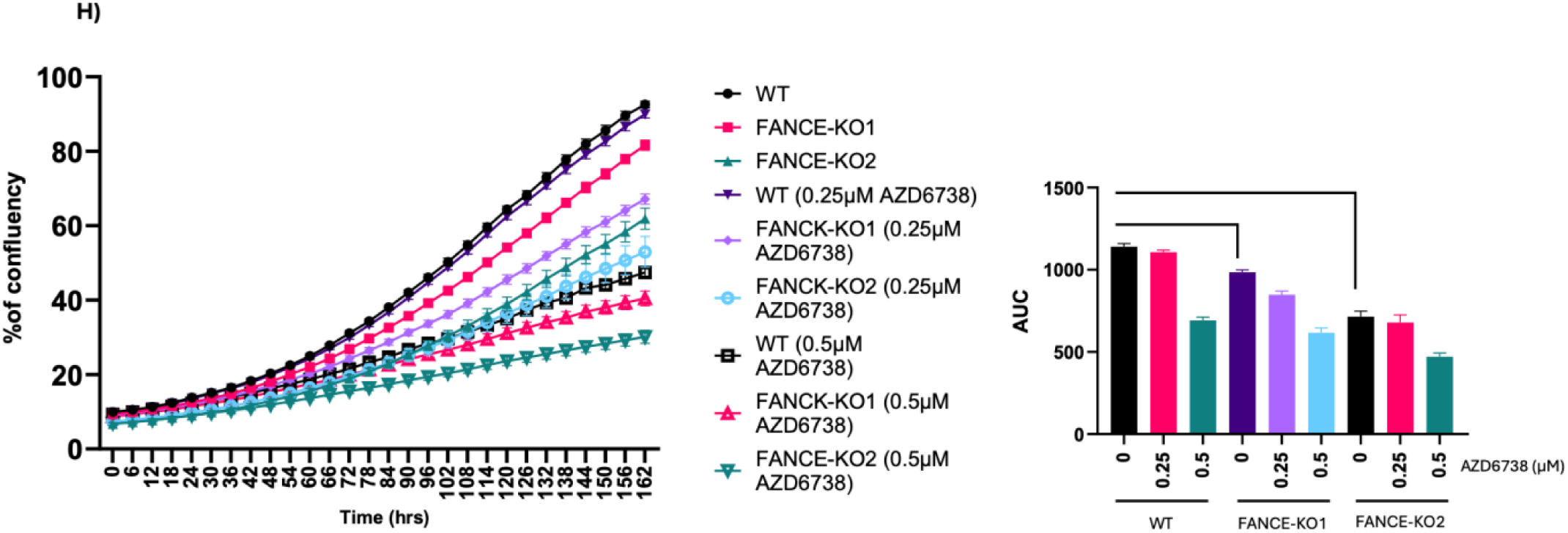
siRNA screen of genetically altered DDR target genes in MDA-MB-453 cells in the presence and absence of AZD6738. (A)The genome sequencing data for MDA-MB-453 cells were obtained from Cell Model Passport, and using a library of DDR signalling pathway genes (from Horizon Discovery), the DDR genes that are genetically altered in this cell line were selected for the siRNA screen. The 25 siRNA (ON-TARGETplus SMARTpool siRNA100nM) were reverse-transfected into MDA-MB-453 cells in 96-well plates (3 replicates per siRNA target) for 48 hours. The cells were then either untreated (DMSO) or treated with 0.5 µM of AZD6738 for 5 days, and cell viability was measured using CellTitre 96. Mean calculated from three wells ±SEM. Non-targeting control pool siRNA was used as a negative control, and ATR siRNA as a positive control. AZD6738 treatment alone also contains the non-targeting control pool siRNA. B) Schematic diagram of the experimental plan. C) The 25 genetically altered DDR target genes from MDA-MB-453 cells are categorised into Mutations (Missense, Nonsense) and Copy number alteration (Loss=decrease in copy number, Gain/Amplification=increase in copy number, Deletion=complete loss). D) Reverse transfection of MDA-MB-453 cells with FACE, FANCG and ATM siRNA, treated with AZD6738 for 5 days, normalised to AZD6738 treatment alone (n=2). Student T-test, p-value <0.05*. E) Whole-cell lysates of six breast cancer cell lines were analysed by Western blot for FANCE. Using ImageJ, each blot was normalised to beta-actin. F) MDA-MB-231 cells were reverse-transfected with FANCE siRNA for 48 hours and then untreated and treated with 1µM of AZD6738 for 5 days, and cell viability was measured by CellTiter 96. Mean calculated from three wells ±SEM. CRISPR-Cas9-mediated FANCE knockout reduces MDA-MB-453 cell growth and sensitises cells to AZD6738. G) Western blot analysis of FANCE protein levels in MDA-MB-453 cells following CRISPR-mediated gene knockout (KO). Β-actin was used as a loading control. H) MDA-MB-453 WT and FANCE-KO cell lines were untreated (DMSO) and treated with AZD6738 for 4 days, and cell viability was measured by crystal violet. Mean calculated from four wells ±SEM. I) MDA-MB-453 WT and FANCE-KO cell lines were either untreated (DMSO) or treated with AZD6738, and IncuCyte measured the percentage of cell confluency over 7 days, every 6 hours. Area under the curve (AUC) was measured, mean was calculated from four wells ±SEM.

The 25 (ON-TARGETplus SMARTpool) siRNAs were individually reverse-transfected into MDA-MB-453 cells, followed by AZD6738 treatment and viability measurement. TP53 was excluded from the screen as it is deleted in this line. As positive controls, GAPDH siRNA confirmed knockdown efficiency (90% mRNA reduction at 48 h), and ATR siRNA reduced viability by ∼30–40%. Knockdown of FANCE, FANCG, and ATM increased AZD6738 sensitivity (Figure 6A, D). ATM knockdown is known to increase sensitivity to ATR inhibitors, thus serving as a positive control and confirming the reliability of our screening approach. MDA-MB-453 cells carry amplifications of FANCE and FANCG. Since FANCE knockdown robustly reduced cell viability with AZD6738 treatment (Figure 6D), we prioritised FANCE for further study.

To assess whether FANCE amplification increases protein expression, we performed western blot analysis across six breast cancer cell lines (Figure 6E); indeed, we observed high protein levels in MDA-MB-453 cells. MDA-MB-231 cells had high FANCE protein levels without gene amplification; therefore, we tested whether its knockdown sensitised cells to AZD6738. Reverse transfection of FANCE siRNA, followed by AZD6738 treatment, robustly reduced cell viability (Figure 6F), confirming reproducibility in another metastatic/TNBC cell line.

We generated two FANCE-KO MDA-MB-453 lines using CRISPR–Cas9 (Figure 6G). InCucyte analysis showed reduced growth rate in FANCE-KO cells versus wild type (Figure 6H); consequently, FANCE-KO cell confluency was further reduced by AZD6738 treatment. Similarly, previously we observed that FANCG knockdown reduced viability (Figure 6A). As the FA pathway repairs ICLs, loss of FANCE or FANCG likely slows repair and reduces proliferation. These results indicate that amplification of FANCE and FANCG in MDA-MB-453 cells promotes cell growth or proliferation, potentially by sustaining DNA replication-associated DNA repair. Therefore, knockdown of these proteins further reduces cell viability in AZD6738-treated cells.

### Fanconi anaemia pathway inhibitor UBE2T/FANCL-IN-1 synergises with ATR inhibitor treatment

We next tested whether FA pathway inhibition could be combined with AZD6738. UBE2T/FANCL-IN-1 is a small molecule inhibitor^54^ that blocks the FA pathway by preventing monoubiquitination and activation of FANCD2, leading to ICL DNA repair. Combination treatment of AZD6738 with UBE2T/FANCL-IN-1 for seven days significantly reduced MDA-MB-453 cell confluency compared with either agent alone (Figure 7A). Synergy Finder analysis confirmed a synergistic interaction, with high Bliss and ZIP scores (Figure 7C, D). Like FANCE-KO cells, FA pathway inhibition alone reduced confluency, suggesting a growth-promoting role in MDA-MB-453.

**Figure 7.**
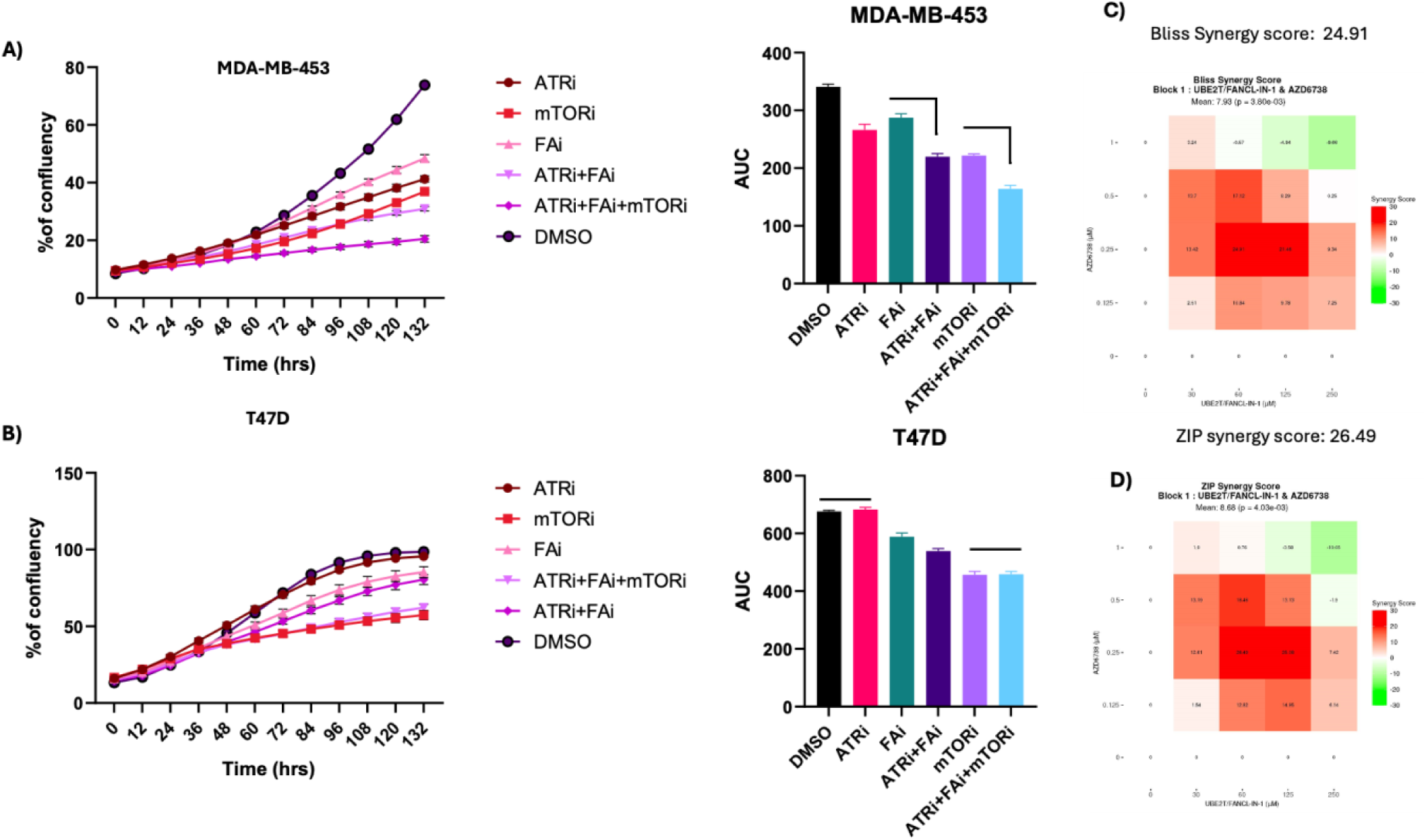
AZD6738 combination treatment with UBE2T-FANCL-IN-1 and CC223. A) MDA-MB-453 cells were untreated (DMSO) or treated with ATRi (AZD6738 0.5µM), mTORi (CC223, 0.25µM), FAi (250µM) alone or ATRi and FAi combined, or ATRi, FAi, mTORi combined. The percentage of cell confluency was measured using IncuCyte over 7 days, with readings taken every 12 hours. B) T47D cells were either untreated (DMSO) or treated with ATRi (AZD6738, 2 µM), mTORi (CC223, 0.25 µM), FAi (250 µM) alone, or ATRi and FAi combined, or ATRi, FAi, and mTORi combined. The percentage of cell confluency was measured using IncuCyte over 7 days, with readings taken every 12 hours. Area under the curve (AUC) was measured, mean calculated from four wells ±SEM. C) MDA-MB-453 cells were treated alone with AZD6738 or UBE2T/FANCL-IN-1 and combined with the concentration shown for 7 days. InCucyte measured the percentage of cell confluency at the end of treatment, and the significance of the combination was analysed using Syngergy Finder. C) Bliss synergy score and D) ZIP synergy score. (Between −10 and +10 is additive, 10+ is synergistic, and below −10 is antagonistic)

Next, we added mTOR pathway inhibitors to the FA pathway inhibitor/AZD6738 combination. In MDA-MB-453 cells, this combination increased AZD6738 sensitivity (Figure 7A). In contrast, in AZD6738-resistant T47D cells, FA+mTOR inhibition had no combinatorial effect with AZD6738 despite T47D’s sensitivity to mTOR inhibition (Figure 7B). Inhibiting the FA pathway, alone or with mTOR blockade, enhances AZD6738 efficacy in MDA-MB-453 cells, supporting FA inhibition as a rational strategy for combination therapy in ATR inhibitor–sensitive breast cancer.

### FANCE and FANCG amplifications are enriched in metastatic breast cancer

To determine whether our cell line findings match patient data, we analysed FANCE and FANCG status in cBioPortal^55, 56, 57^ (Figure 8A). Both were predominantly altered via amplification in metastatic breast cancer. Across 31 studies (6 metastatic, 26 non-metastatic), alterations occurred in 8% of metastatic versus 1% of non-metastatic studies (Figure 8B, C). Among 63 FANCE-amplified samples, mainly from the Metastatic Breast Cancer Project^58^ (2021), the majority were circulating tumour cells, with others from breast, liver, lymph nodes, bone, and bone marrow (Table 1). FANCG was co-amplified in over half of the cases; FANCA, FANCD2, UBE2T, and FANCI were often co-amplified, while FANCL, FANCF, and FANCM were commonly deleted. Although receptor status data were incomplete, FANCE-amplified samples tended toward ER+HER2–PR+. Since FANCE and FANCG amplifications are primarily found in metastasised patient samples (Figure 8A), they may be associated with chemoresistance, as cells with these amplifications may have escaped chemotherapy. More importantly, as FANCE amplification is primarily observed in blood samples, these results suggest that FANCE amplification, in combination with alterations in other genes, may promote cancer progression and metastasis. Together, our cell line and patient analyses demonstrate that FANCE amplification may promote resistance to ATR inhibition, be a feature of metastatic disease progression, and that targeting the FA pathway represents a promising therapeutic approach for metastatic breast cancer.

**Figure 8.**
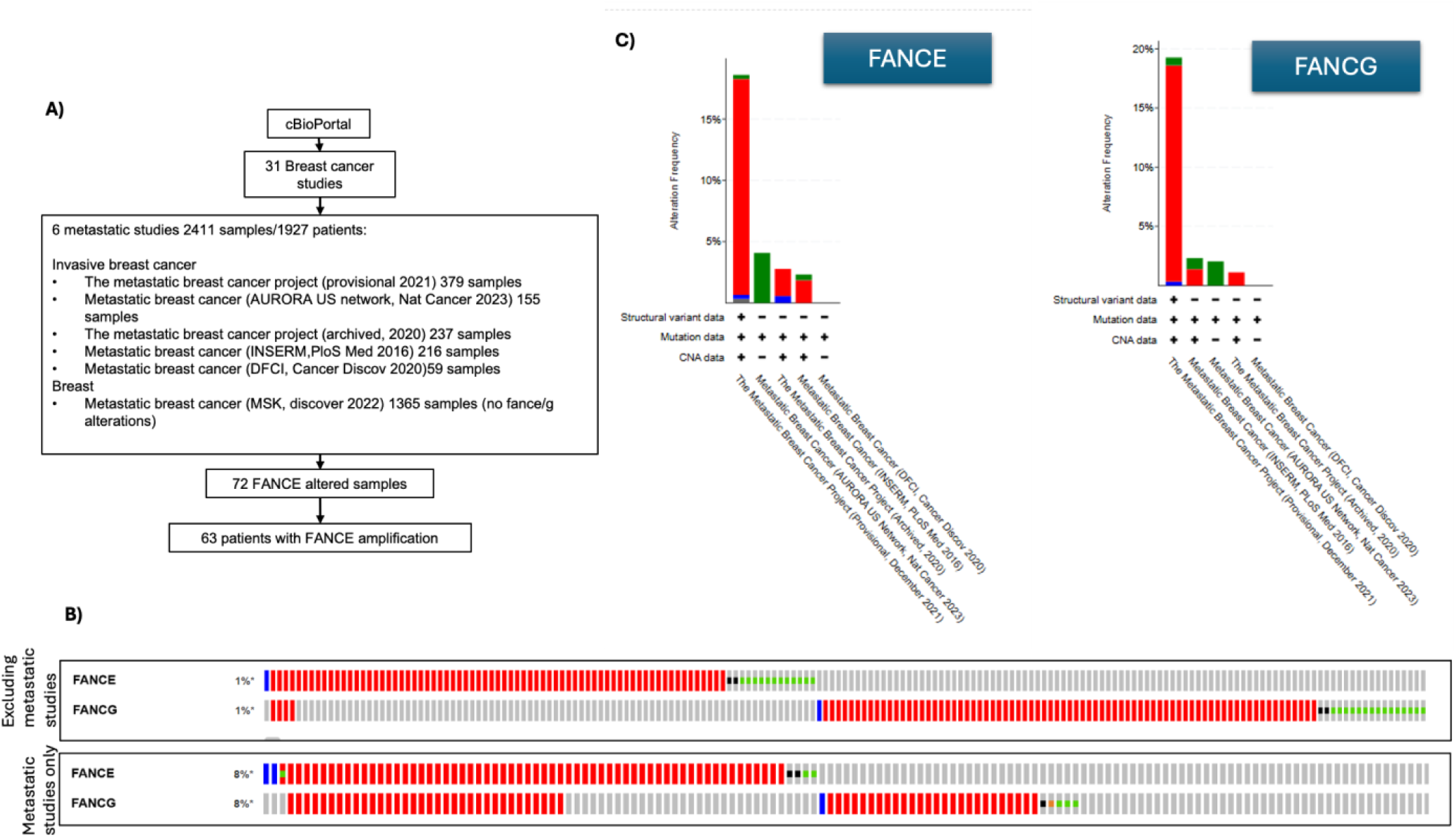
FANCE and FANCG are co-amplified in metastatic breast cancer patient samples. A) Schematic diagram showing analysis of genome sequencing data of metastatic patient samples from CBioPortal. B) Percentage of FANCE and FANCG genetic alterations in metastatic patient samples (6 metastatic studies), and non-metastatic samples (all studies excluding metastatic studies). C) FANCE and FANCG percentage and type of genetic alteration in the six metastatic breast cancer studies (Red=amplification, Blue=deep deletion, Green=mutation).

**Table 1.**
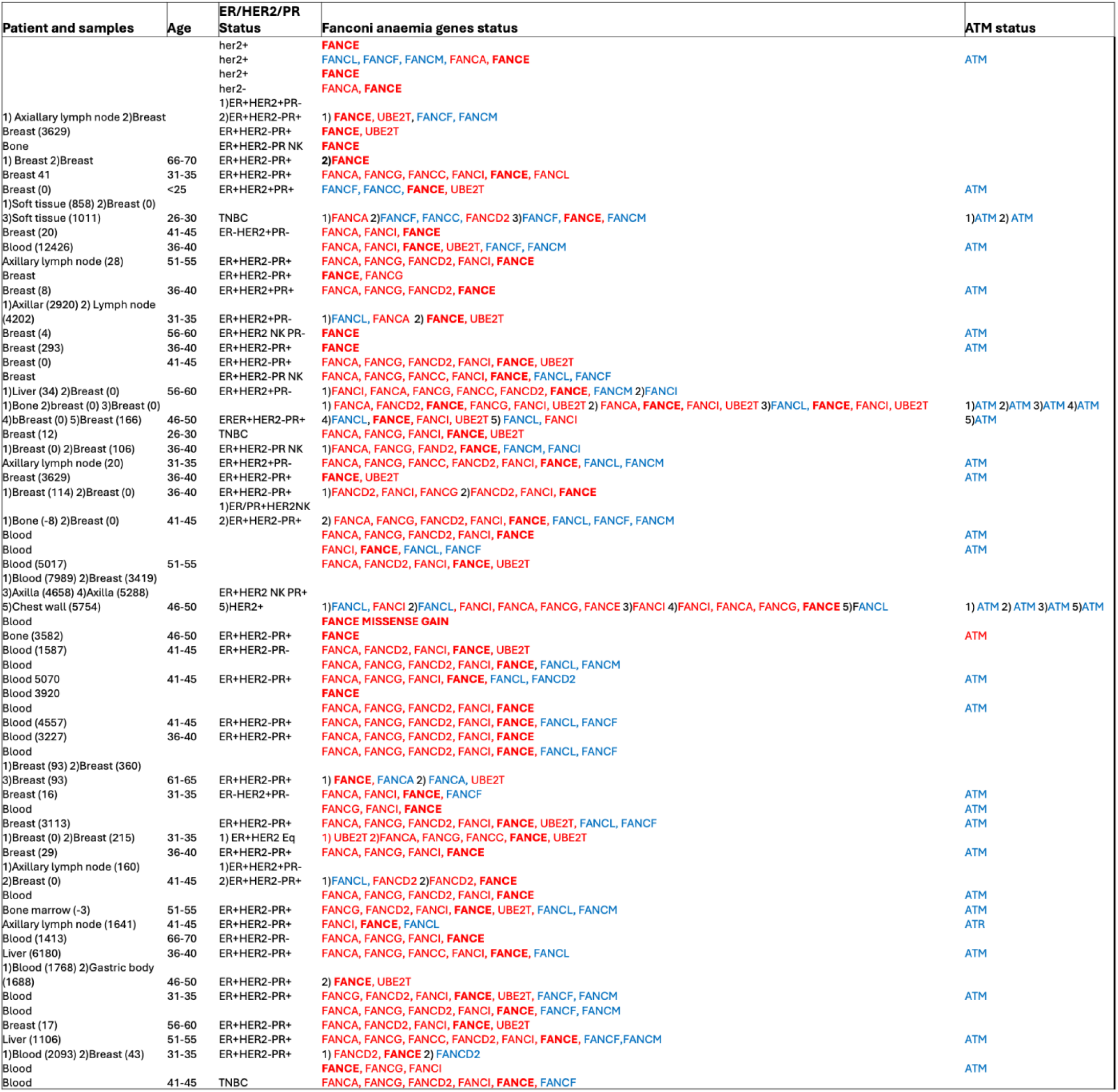
FANCE amplified 63 individual patient sample details, showing the type of metastatic sample and age. ER/HER2/PR status, genetically altered Fanconi anaemia genes and status of ATM (Red=amplification, Blue=deep deletion, Green=mutation). Numbers next to the sample type indicate the day following treatment on which samples are taken. (NK=not known)

## DISCUSSION

MDA-MB-453 breast cancer cells exhibit several genomic alterations that render them highly susceptible to replication stress and DNA damage. MYC amplification contributes to intrinsic replication stress, while p53 deletion compromises G1/S checkpoint control, and BRCA1 loss impairs replication fork repair. In this context, the cells become highly dependent on the ATR pathway to stabilise stalled forks and prevent mitotic catastrophe. In addition, AKT1 amplification exacerbates replication stress, and a missense mutation and low protein expression suggest compromised ATM function. Together, these features create a synthetic dependency on ATR, making MDA-MB-453 cells highly sensitive to the ATR inhibitor AZD6738, which leads to replication fork collapse, mitotic catastrophe, and ultimately cell death.

Using chromatin enrichment proteomics, we observed the upregulation of key markers associated with replication stress, including RPA1, RPA2, and RPA3, indicating single-stranded DNA accumulation. Furthermore, evidence of double-strand break repair through NHEJ was detected. Although HR is typically the preferred pathway for repair following replication fork collapse, HR is likely impaired in these cells due to BRCA1 deficiency. As a result, NHEJ may serve as an adaptive alternative repair mechanism to maintain cell survival^29^. Hyperphosphorylation of Ki-67 (MKI67) suggests failure of the G2/M checkpoint and inappropriate mitotic entry with unresolved DNA damage^33, 59^.

In addition, we observed upregulation of CDK4/5, AKT1, RPTOR, and OGFR signalling molecules likely involved in adaptive responses that promote resistance to ATR inhibition. Notably, inhibition of mTOR kinase, which targets both mTORC1 and AKT pathways, was effective in combination with AZD6738, supporting the therapeutic potential of dual-targeting strategies to overcome adaptive resistance.

To identify mechanisms that affect ATR inhibitor sensitivity, we performed a siRNA screen and found that FANCE knockdown increases ATR inhibitor sensitivity. As FANCE is amplified in this cell line, this suggests that the amplified FANCE may confer resistance to the ATR inhibitor. FANCE is part of the FA pathway, which is essential for the repair of interstrand crosslinks (ICLs), a particularly lethal form of DNA damage that blocks both transcription and replication. ICLs pose a particular problem in rapidly dividing cells; therefore, amplified genes in the FA pathway may give cancer cells an advantage in repairing the damage and progressing into cell division. This also explains why FANCE knockout or FANCG knockdown slows cell growth or proliferation. Our results show that treatments targeting the FA pathway could be an effective combination therapy with AZD6738 for metastatic cancer. There are other FA pathway inhibitors in development, such as the small molecule inhibitor that interferes with FANCM/RMI complex formation ^22^. In addition, amplified FANCE/FANCG could be predictive markers of chemoresistance or DDR-targeted treatments.

Clinically, our findings are relevant to patients with metastatic breast cancer, as amplifications of FANCE and FANCG are frequently observed in metastatic samples. Interestingly, these amplifications are particularly enriched in circulating tumour cells (CTCs) isolated from blood, suggesting a potential role in promoting metastasis. However, these amplifications often occur alongside deletions of other FA pathway components such as FANCM, FANCL (an E3 ubiquitin ligase), and BRCA2, raising important questions about how amplification of certain FA genes confers resistance and metastatic potential in the absence of other critical pathway members. We propose that the loss of core FA genes (e.g., FANCM, FANCL, BRCA2) may contribute to cancer initiation and genomic instability, while subsequent amplification of other FA components (e.g., FANCE, FANCG) during tumour progression or chemotherapy exposure may reflect clonal selection of resistant subpopulations. This shift could facilitate continued cell proliferation and metastasis, even in the face of DNA-damaging treatments. Further mechanistic studies are warranted to understand how FANCE and FANCG amplification drive resistance and metastasis. These findings highlight the complexity of DDR in advanced breast cancer, where multiple alterations, signalling interactions, and compensatory mechanisms shape the tumour’s response to treatment. Thus, when evaluating biomarkers for sensitivity to ATR inhibitors such as AZD6738, it is critical to consider not only single gene alterations but also the broader landscape of DDR and signalling pathway dysregulation.

In summary, metastatic breast cancer cells such as MDA-MB-453 exhibit extensive alterations in DNA damage response and repair mechanisms, driven by intrinsic replication stress, loss of key repair factors, and adaptive signalling changes. Chromatin proteomics and phospho-proteomics provide valuable insight into these mechanisms. The rewired DNA repair landscape in cancer cells not only promotes resistance to therapy but may also facilitate progression and metastasis. Our findings underscore the therapeutic potential of targeting ATR in combination with mTOR or FA pathway inhibitors and highlight the importance of understanding compensatory DDR mechanisms to overcome resistance and improve treatment strategies for metastatic breast cancer.

## MATERIALS AND METHODS

### Cell lines and treatments

Cell lines were originally purchased from ATCC. MCF7, MDA-MB-453, MDA-MB-436, and HCC-1954 were grown in DMEM supplemented with 10% fetal bovine serum and 2 mM L-glutamine. MDA-MB-231 and T47D were grown in RPMI supplemented with 10% fetal bovine serum and 2 mM L-glutamine. Cells were cultured at 37 °C in humidified incubators maintained at 5%CO2. These cell lines are derived initially from metastatic breast cancer patients.

### 3D cell culture and viability assay

Cells were seeded (3000 per well) in 96-well U-bottom plates with an Ultra-low attachment surface and left to grow spheroids for 4 days. Spheroids were treated with AZD6738 for 2 days, and 100 μL/well of CellTitre-Glo 3D Cell viability assay (Promega) solution was added. Cell viability was measured by luminescence.

### Compounds

AZD6738, CC223 and, UBE2T/FANCL-IN-1 were purchased from MedChemExpress. All inhibitors were solubilised in DMSO, and aliquots were stored at −80 °C.

### Cytotoxic assay

Cytotoxicity assay was performed using the CytoTox-Glo cytotoxic assay (Promega) on untreated and treated cells with AZD6738, following the manufacturer’s instructions. Dead cells were measured by luminescence, followed by the total cell number, which was measured after adding lysis reagent. Total cell number - Dead cell number = Viable cell number. Dead cells were normalised to viable or live cells.

### Apoptosis assay

The Caspase 3/7 activation assay was performed using the Caspase-Glo 3,7 Assay System (Promega) on untreated and treated cells with AZD6738, following the manufacturer’s instructions. Luminescence was measured, and the signal was normalised to viable cells measured by CellTitre 96.

### Cell proliferation/viability assay

Cells were seeded in 96-well plates, and treatments or siRNA transfection were carried out. At the end of the experiment, 20 μL per well of CellTitre 96 Aqueous One Solution Proliferation Assay (MTS) (Promega) solution was added, and the colour change was measured at 490 nm.

### Crystal violet staining

Cells were seeded (7000 per well) in 96-well plates and the next day treated with increasing concentrations of AZD6736 for 4 days. Cells were fixed in 3%w/v paraformaldyhyde, washed with PBS, stained (0.5% crystal violet, 20% methanol), washed with water, eluted with 1% acetic acid, and absorbance read at 570nm.

### Chromatin Enrichment

Flash-frozen cell pellets were thawed on ice and resuspended in a nuclear extraction buffer composed of 15 mM Tris-HCl (pH 7.5), 60 mM KCl, 15 mM NaCl, 5 mM MgCl₂, 1 mM CaCl₂, 250 mM sucrose, and 0.3% NP-40. The buffer was freshly supplemented with 1 mM DTT and a protease/phosphatase inhibitor cocktail (Halt™,Thermo Scientific). After a 5-minute incubation on ice, the nuclei were pelleted by centrifugation at 600 × g for 5 minutes at 4 °C. The nuclear pellets were washed once using extraction buffer lacking NP-40, then pelleted again and resuspended in ice-cold hypotonic buffer (3 mM EDTA, 0.2 mM EGTA, with fresh 1 mM DTT and protease/phosphatase inhibitors). Samples were incubated on ice for 30 minutes to solubilise chromatin. Chromatin was isolated by centrifugation at 1,700 × g for 5 minutes at 4 °C and washed twice with hypotonic buffer.

### MS Sample Preparation and TMT Labelling

Chromatin pellet was lysed by probe sonication in buffer containing 100 mM triethylammonium bicarbonate (TEAB), 1% sodium deoxycholate (SDC), 10% isopropanol, 50 mM NaCl, and 1:1,000 Pierce Universal Nuclease (Thermo), with inhibitors added freshly. Protein concentration was determined using the Quick Start™ Bradford Assay (Bio-Rad). Equal amounts of protein from each sample were pooled, reduced with 5 mM TCEP for 1 hour, then alkylated with 10 mM IAA for 30 minutes. Proteins were digested overnight at room temperature using trypsin (75 ng/μL final concentration; Thermo).

For downstream applications, 100 μg of digested protein was used for phosphopeptide enrichment, and 30 μg was used for total proteome analysis. TMT 10-plex reagents (Thermo) were used to label peptides as per manufacturer instructions. After labelling and pooling, SDC was precipitated by addition of 2% formic acid (v/v), followed by centrifugation at 10,000 rpm for 5 minutes. Supernatants were collected, dried in a vacuum concentrator, and stored for LC-MS.

### High-pH Reverse-Phase Fractionation and LC-MS Analysis

TMT-labelled peptides were separated by high-pH reversed-phase chromatography using a Waters XBridge C18 column (2.1 × 150 mm, 3.5 µm) on a Dionex UltiMate 3000 HPLC. The mobile phases consisted of 0.1% ammonium hydroxide in water (A) and 0.1% ammonium hydroxide in 100% acetonitrile (B). A gradient was run at 200 µL/min: 5 minutes at 5% B (isocratic), 40-minute linear gradient to 35% B, followed by a 5-minute ramp to 80% B, then held for 5 minutes. Fractions were collected every 42 seconds, pooled into 12 concatenated fractions, dried, and reconstituted in 0.1% TFA prior to LC-MS

### Phosphopeptide Enrichment

Peptides were reconstituted in 10 µL of binding buffer (20% isopropanol, 0.5% formic acid), and enriched using the High-Select Fe-NTA Magnetic Phosphopeptide Kit (Thermo). Resin was pre-washed and equilibrated in custom-made filter tips. Binding was carried out at room temperature for 30 minutes, followed by three washes in binding buffer, using centrifugation at 300 × g. Flow-through fractions were retained for total proteome analysis. Phosphopeptides were eluted in three steps with 40% acetonitrile and 400 mM ammonium hydroxide, vacuum-dried, and stored at −20 °C.

### LC-MS Acquisition

Peptides were separated using a Dionex UltiMate 3000 UHPLC coupled to an Orbitrap Ascend MS. An EASY-Spray C18 column (75 μm × 50 cm, 2 µm) maintained at 50 °C was used. Mobile phase A was 0.1% formic acid; phase B was 80% acetonitrile with 0.1% formic acid. The gradient was: 0–150 min to 38% B, 150–160 min to 95% B, held for 5 min, re-equilibrated to 5% B in 10 min, then held for 10 min. Full MS1 scans (m/z 400–1600) were acquired at 120,000 resolution with AGC target of 4 × 10⁵ and 50 ms max injection time. Top-speed mode (3 s cycle) selected ions for HCD fragmentation (38% NCE, 0.7 Th isolation). MS2 scans were collected in the ion trap at turbo scan rate with 32% NCE and max injection time of 35 ms.

For selected precursors, SPS-MS3 scans were acquired using the Orbitrap (45,000 resolution, 55% NCE, AGC 200%, 200 ms max injection time). Ions with charges +2 to +6 were selected, with dynamic exclusion enabled (repeat count = 1, exclusion time = 45 s, ±10 ppm window, isotope exclusion on).

### Database Search and Quantification

Spectra were analysed in Proteome Discoverer (version 3.0) using SequestHT and Comet search engines against the reviewed human UniProt database (canonical and isoforms). For total chromatome analysis, search parameters included a precursor tolerance of 20 ppm and fragment ion tolerance of 0.5 Da; for phosphopeptides, 10 ppm and 0.02 Da, respectively. Static modifications were set for carbamidomethylation (C) and TMT on lysines and N-termini. Dynamic modifications included methionine oxidation and deamidation (N/Q). Up to two missed cleavages were allowed, and peptide identification was filtered at 1% FDR using Percolator.

Reporter ion quantification was based on MS2 or MS3 scans, with centroid-based peak integration (15 ppm tolerance). Quantification used only unique peptides with S/N > 3. Protein abundances were normalised across TMT channels based on total signal; phosphopeptide data were further normalised against the phospho-enriched input.

### Data Analysis

P-values for differential expression were calculated using a two-sided Student’s *t*-test comparing ATR inhibitor-treated samples to untreated controls. Significantly regulated genes and phospho-sites were defined by thresholds of *p* < 0.05 and |log₂ fold change (log₂FC)| > 0.5.

Data visualisation and enrichment analysis were performed using Python (v3.10). Volcano plots were generated with **matplotlib, seaborn, numpy,** and **adjustText**. The plots display –log₁₀(*p*-value) versus log₂FC, with significantly changing features coloured according to statistical significance and effect size. Non-significant points were shown in grey. The top 10 most significant genes and phosphosites (ranked by lowest *p*-value) were annotated on the plots and highlighted with gold-edged markers.

Gene set enrichment analysis was conducted using the **gseapy** package (v1.0.6), leveraging the Enrichr API to test GO Biological Process (2023) gene set. Enrichment was assessed separately for upregulated and downregulated genes and phospho-site associated proteins. Selected 20 enriched terms were visualised as dot plots using **matplotlib**.

### siRNA transfections

In 96-well plates, siRNAs (100nM ON-TARGETplus pool) (Iorns et al 2009) were reverse-transfected with transfection reagent 2 (DharmaFECT transfection reagent, Horizon Discovery) in serum-free medium, following the manufacturer’s instructions ^60^. 7000 MDA-MB-435 cells were added per well (100 μL/well of media supplemented with 10% fetal bovine serum and 2mM L-Glutamine). After 48 hours, AZD6738 was added with fresh media, and after 5 days, cell viability was measured with CellTitre 96 (Promega). GAPDH siRNA was used as a positive control to assess knockdown efficiency. 50nM GAPDH transfected for 48hrs knocked down 90% of mRNA (Supplementary Figure 1). The non-targeting control pool siRNA was used as a negative control. MDA-MB-231 cells were reverse-transfected with siRNA, as for MDA-MB-453, except DharmaFECT transfection reagent 4 was used as recommended by the manufacturer. After 48hrs, AZD6738 was added to the media as the cells were fast-growing, for 5 days and cell viability was measured with CellTitre 98.

### Western blotting

Cells were lysed in RIPA buffer containing protease and phosphatase inhibitors, and protein concentration was measured using the Pierce BCA Protein Assay Kit (ThermoFisher). Proteins were separated on precast polyacrylamide gels (BIO-RAD) and blotted onto Immobilon-P PVDF membrane (Millipore). Non-specific binding was blocked with tris-buffered saline (TBS) with 0.1% Tween-20 and 5% w/v non-fat dried milk. Primary antibodies FANCE antibody was purchased from Cambridge Biosciences, ATM antibody was purchased from Bio-Techne, p53 antibody was purchased from Santa Cruz, and β-Actin and Phospho-H2A.X (Ser139) antibody were purchased from Cell Signalling Technology. Secondary antibodies, Anti-mouse IgG HRP-linked and Anti-rabbit IgG HRP-linked, were purchased from Cell Signalling Technology. Membranes were incubated in chemiluminescent substrate Immobilon Forte Western HRP (Millipore) and visualised by I-Bright. Bands were quantified using Image J.

### InCucyte analysis

MDA-MB-453 and T47D cells were seeded in 96-well plates and incubated overnight to allow attachment. The next day, cells were treated with indicated concentrations of AZD6738 and CC223 or equivalent volume of vehicle control (DMSO). Plates were then placed in an IncuCyte S3 live-cell analysis system (Sartorius) housed in a humidified incubator (37 °C, 5% CO₂). Phase-contrast images were acquired with a 10× objective every 6 h or 12 h over 7 days, or until cultures reached > 80% confluency. Percentage confluency was calculated using the IncuCyte software with the standard phase-contrast processing algorithm. Confluency data were exported as CSV files and analysed in GraphPad Prism. Results are mean of four wells ± SEM.

### Generation of MDA-MB-453 knockout cells

FANCE gene knockouts were generated using the CRISPR-Cas9 pLentiCRISPRv2 lentiviral vector system^61^. Synthesised DNA fragments encoding guide RNAs (gRNAs) targeting FANCE were cloned into the single-guide RNA (sgRNA) expression cassette within the vector using the HiFi DNA Assembly reaction (New England Biolabs). Two gRNAs were used: GCAAGCGACCCCAGTCGAA, targeting exon 1 ^62^, and AGGTTCCTCTGGCATATCCG, targeting exon 2 (designed using TrueDesign Genome Editor, Invitrogen).

Lentiviral particles were produced in HEK293T cells by co-transfecting the gRNA-containing pLentiCRISPRv2 constructs with the packaging plasmid psPAX2 and the envelope plasmid pMD2.G (Addgene lentivirus production protocol-https://www.addgene.org/protocols/lentivirus-production/) using FuGENE HD transfection reagent (Promega). Conditioned media containing lentivirus were harvested 72 hours post-transfection and used to transduce MDA-MB-453 cells in the presence of polybrene (8 µg/mL; Sigma-Aldrich). Transduced cells were selected with puromycin (1 µg/mL) until all untransduced and parallel cultures of control cells were killed (approximately 10 days).

### Statistical analysis

Statistical analyses were carried out using GraphPad Prism v10. Pairwise comparisons were performed using the Student’s T-test. For combination therapy synergy analyses online Synergy finder was used. Cells were seeded (5000), the next day treated with AZD6738 and UBE2T/FANCL-IN-1 alone and in combination for 7 days and the percentage of cells was measured by IncuCyte on the 7^th^ day. A file containing the percentage of cell confluency was uploaded to the SynergyFinder 2.0 web application (synergy finder.fimm.fi). Bliss and ZIP synergy scores were calculated using default parameters.

## Data and Material Availability

The data presented in this study and in the Supplementary information are available from the corresponding author upon request.

## Acknowledgments

Jogitha Selvarajah holds a Daphne Jackson Fellowship jointly funded by Imperial College London and the Medical Research Council (MRC). We would like to thank Professor Laki Buluwela for providing vectors, reagents and support with CRISPR experiments. We would also like to thank Prof Simak Ali for providing all six breast cancer cell lines and for helpful suggestions on the manuscript.

---

**Supplementary Figure 1.**
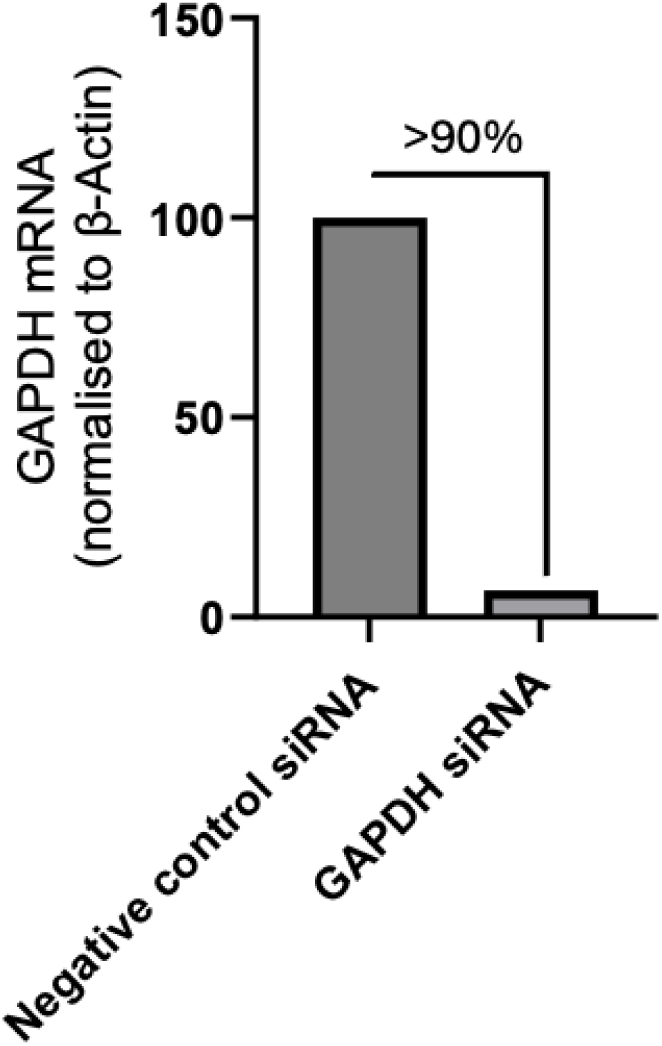
GAPDH siRNA was used as a positive control to assess knockdown efficiency. 50nM GAPDH siRNA was reverse-transfected for 48 hours in MDA-MB-453 cells, along with a Negative control siRNA (non-targeting control pool siRNA). Total GAPDH mRNA was measured with qRT-PCR and normalised to actin mRNA.

