## Supplementary Table 1 for "Chromatin signalling pathways and FANCE amplification affect ATR inhibitor sensitivity in metastatic breast cancer"

| Symbol | Cell Line | Type | Protein mutation | cDNA mutation | Effect | CNA call |
|---|---|---|---|---|---|---|
| NBN | HCC1954 | snp | p.T253I | c.758C>T | missense | undefined |
| NUDT1 | HCC1954 | snp | p.V106M | c.316G>A | missense | undefined |
| RNF168 | HCC1954 | snp | p.A344T | c.1030G>A | missense | undefined |
| OGG1 | HCC1954 | snp | p.G348E | c.1043G>A | missense | undefined |
| GEN1 | HCC1954 | snp | p.S873R | c.2619T>G | missense | undefined |
| SMC6 | HCC1954 | snp | p.A292V | c.875C>T | missense | undefined |
| XAB2 | HCC1954 | snp | p.R575Q | c.1724G>A | missense | undefined |
| TP53 | HCC1954 | snp | p.Y163C | c.488A>G | missense | undefined |
| FANCA | HCC1954 | snp | p.A1346T | c.4036G>A | missense | undefined |
| TDG | HCC1954 | snp | p.K201T | c.602A>C | missense | undefined |
| POLL | HCC1954 | snp | p.R57W | c.169C>T | missense | undefined |
| DCLRE1C | HCC1954 | snp | p.E632K | c.1894G>A | missense | undefined |
| BRCA1 | HCC1954 | frameshift | p.V1830fs*20 | c.5488_5489delGT | frameshift | undefined |
| FANCF | HCC1954 | CNA | undefined | undefined | undefined | Loss |
| POLE | HCC1954 | CNA | undefined | undefined | undefined | Loss |
| ATM | HCC1954 | CNA | undefined | undefined | undefined | Loss |
| ATRX | HCC1954 | CNA | undefined | undefined | undefined | Loss |
| ERCC2 | HCC1954 | CNA | undefined | undefined | undefined | Gain |
| FANCG | HCC1954 | CNA | undefined | undefined | undefined | Loss |
| UBE2A | HCC1954 | CNA | undefined | undefined | undefined | Loss |
| WRN | HCC1954 | CNA | undefined | undefined | undefined | Loss |
| NBN | HCC1954 | CNA | undefined | undefined | undefined | Amplification |
| RAD21 | HCC1954 | CNA | undefined | undefined | undefined | Amplification |
| BRIP1 | HCC1954 | CNA | undefined | undefined | undefined | Gain |
| FANCA | HCC1954 | CNA | undefined | undefined | undefined | Gain |
| MEN1 | HCC1954 | CNA | undefined | undefined | undefined | Gain |
| PMS2 | HCC1954 | CNA | undefined | undefined | undefined | Gain |
| POLD1 | HCC1954 | CNA | undefined | undefined | undefined | Gain |
| MSH5 | MCF7 | snp | p.S3F | c.8C>T | missense | undefined |
| NBN | MCF7 | snp | p.R43* | c.127C>T | nonsense | undefined |
| NBN | MCF7 | snp | p.N321Y | c.961A>T | missense | undefined |
| NBN | MCF7 | snp | p.P325H | c.974C>A | missense | undefined |
| XRCC4 | MCF7 | snp | p.K197R | c.590A>G | missense | undefined |
| FANCM | MCF7 | snp | p.R1931* | c.5791C>T | nonsense | undefined |
| PER1 | MCF7 | snp | p.A1196V | c.3587C>T | missense | undefined |
| PALB2 | MCF7 | snp | p.V560L | c.1678G>C | missense | undefined |
| ERCC6 | MCF7 | snp | p.V83I | c.247G>A | missense | undefined |
| ERCC6 | MCF7 | snp | p.S1099R | c.3297T>A | missense | undefined |
| ATM | MCF7 | CNA | undefined | undefined | undefined | Loss |
| ATRX | MCF7 | CNA | undefined | undefined | undefined | Loss |
| ERCC5 | MCF7 | CNA | undefined | undefined | undefined | Loss |
| FANCF | MCF7 | CNA | undefined | undefined | undefined | Loss |
| FANCG | MCF7 | CNA | undefined | undefined | undefined | Loss |
| FANCA | MCF7 | CNA | undefined | undefined | undefined | Gain |
| MUTYH | MCF7 | CNA | undefined | undefined | undefined | Loss |
| SMC1A | MCF7 | CNA | undefined | undefined | undefined | Loss |
| WRN | MCF7 | CNA | undefined | undefined | undefined | Loss |

| | | | | | | |
|---|---|---|---|---|---|---|
| BRIP1 | MCF7 | CNA | undefined | undefined | undefined | Amplification |
| NBN | MCF7 | CNA | undefined | undefined | undefined | Amplification |
| DDB2 | MCF7 | CNA | undefined | undefined | undefined | Gain |
| NPM1 | MCF7 | CNA | undefined | undefined | undefined | Gain |
| FANCC | MDA-MB-231 | snp | p.V449M | c.1345G>A | missense | undefined |
| MSH3 | MDA-MB-231 | snp | p.G896R | c.2686G>A | missense | undefined |
| CCNO | MDA-MB-231 | snp | p.P45H | c.134C>A | missense | undefined |
| PER1 | MDA-MB-231 | snp | p.E189D | c.567G>T | missense | undefined |
| TP53 | MDA-MB-231 | snp | p.R280K | c.839G>A | missense | undefined |
| NTHL1 | MDA-MB-231 | snp | p.R100C | c.298C>T | missense | undefined |
| TP53BP1 | MDA-MB-231 | snp | p.I1179V | c.3535A>G | missense | undefined |
| ATM | MDA-MB-231 | snp | p.N1005I | c.3014A>T | missense | undefined |
| MMS19 | MDA-MB-231 | snp | p.E859K | c.2575G>A | missense | undefined |
| MSH4 | MDA-MB-231 | snp | p.Q815R | c.2444A>G | missense | undefined |
| RECQL | MDA-MB-231 | splice_variant | p.? | c.1667_1667+3delAgta | ess_splice | undefined |
| NBN | MDA-MB-231 | CNA | undefined | undefined | undefined | Gain |
| WRN | MDA-MB-231 | CNA | undefined | undefined | undefined | Loss |
| ERCC2 | MDA-MB-231 | CNA | undefined | undefined | undefined | Gain |
| PER1 | MDA-MB-231 | CNA | undefined | undefined | undefined | Gain |
| RAD21 | MDA-MB-231 | CNA | undefined | undefined | undefined | Gain |
| TCEA1 | MDA-MB-231 | CNA | undefined | undefined | undefined | Gain |
| TP53 | MDA-MB-231 | CNA | undefined | undefined | undefined | Gain |
| BRCA1 | MDA-MB-436 | splice_variant | p.? | c.5340+1G>A | ess_splice | undefined |
| TP53 | MDA-MB-436 | snp | p.E204A | c.611A>C | missense | undefined |
| FANCI | MDA-MB-436 | snp | p.S812G | c.2434A>G | missense | undefined |
| SOD1 | MDA-MB-436 | snp | p.I150V | c.448A>G | missense | undefined |
| PMS1 | MDA-MB-436 | snp | p.E537K | c.1609G>A | missense | undefined |
| MLH3 | MDA-MB-436 | snp | p.F92L | c.276C>G | missense | undefined |
| DNA2 | MDA-MB-436 | snp | p.H461R | c.1382A>G | missense | undefined |
| PRKDC | MDA-MB-436 | frameshift | p.D723fs*8 | c.2167delG | frameshift | undefined |
| TP53 | MDA-MB-436 | frameshift | p.E204fs*7 | c.610_611insCGTGTGG | frameshift | undefined |
| BRCA1 | MDA-MB-436 | CNA | undefined | undefined | undefined | Loss |
| ERCC2 | MDA-MB-436 | CNA | undefined | undefined | undefined | Loss |
| ERCC4 | MDA-MB-436 | CNA | undefined | undefined | undefined | Loss |
| MLH1 | MDA-MB-436 | CNA | undefined | undefined | undefined | Loss |
| NBN | MDA-MB-436 | CNA | undefined | undefined | undefined | Loss |
| PALB2 | MDA-MB-436 | CNA | undefined | undefined | undefined | Loss |
| ATM | MDA-MB-436 | CNA | undefined | undefined | undefined | Loss |
| ATR | MDA-MB-436 | CNA | undefined | undefined | undefined | Loss |
| BRIP1 | MDA-MB-436 | CNA | undefined | undefined | undefined | Loss |
| CHEK2 | MDA-MB-436 | CNA | undefined | undefined | undefined | Loss |
| FANCE | MDA-MB-436 | CNA | undefined | undefined | undefined | Loss |
| MEN1 | MDA-MB-436 | CNA | undefined | undefined | undefined | Gain |
| MUTYH | MDA-MB-436 | CNA | undefined | undefined | undefined | Gain |
| POLD1 | MDA-MB-436 | CNA | undefined | undefined | undefined | Loss |
| SMC1A | MDA-MB-436 | CNA | undefined | undefined | undefined | Gain |
| TCEA1 | MDA-MB-436 | CNA | undefined | undefined | undefined | Gain |
| TOP2A | MDA-MB-436 | CNA | undefined | undefined | undefined | Loss |

| | | | | | | |
|---|---|---|---|---|---|---|
| WRN | MDA-MB-436 | CNA | undefined | undefined | undefined | Loss |
| XPA | MDA-MB-436 | CNA | undefined | undefined | undefined | Loss |
| ERCC5 | MDA-MB-436 | CNA | undefined | undefined | undefined | Amplification |
| NPM1 | MDA-MB-436 | CNA | undefined | undefined | undefined | Amplification |
| PMS2 | MDA-MB-436 | CNA | undefined | undefined | undefined | Amplification |
| RAD21 | MDA-MB-436 | CNA | undefined | undefined | undefined | Amplification |
| RECQL4 | MDA-MB-436 | CNA | undefined | undefined | undefined | Amplification |
| DDB2 | MDA-MB-436 | CNA | undefined | undefined | undefined | Gain |
| FANCC | MDA-MB-436 | CNA | undefined | undefined | undefined | Gain |
| FANCD2 | MDA-MB-436 | CNA | undefined | undefined | undefined | Gain |
| POLQ | MDA-MB-436 | CNA | undefined | undefined | undefined | Gain |
| UBE2A | MDA-MB-436 | CNA | undefined | undefined | undefined | Gain |
| XPC | MDA-MB-436 | CNA | undefined | undefined | undefined | Gain |
| FANCE | MDA-MB-453 | snp | p.S337L | c.1010C>T | missense | undefined |
| XPC | MDA-MB-453 | snp | p.Q742* | c.2224C>T | nonsense | undefined |
| POLQ | MDA-MB-453 | snp | p.W1760C | c.5280G>T | missense | undefined |
| PNKP | MDA-MB-453 | snp | p.P20S | c.58C>T | missense | undefined |
| EME2 | MDA-MB-453 | snp | p.R192H | c.575G>A | missense | undefined |
| TP53BP1 | MDA-MB-453 | snp | p.T387S | c.1159A>T | missense | undefined |
| ALKBH1 | MDA-MB-453 | snp | p.L14V | c.40C>G | missense | undefined |
| ATM | MDA-MB-453 | snp | p.E1856Q | c.5566G>C | missense | undefined |
| BRCA1 | MDA-MB-453 | CNA | undefined | undefined | undefined | Loss |
| FANCC | MDA-MB-453 | CNA | undefined | undefined | undefined | Loss |
| TOP2A | MDA-MB-453 | CNA | undefined | undefined | undefined | Loss |
| ERCC5 | MDA-MB-453 | CNA | undefined | undefined | undefined | Loss |
| NBN | MDA-MB-453 | CNA | undefined | undefined | undefined | Gain |
| PER1 | MDA-MB-453 | CNA | undefined | undefined | undefined | Loss |
| TP53 | MDA-MB-453 | CNA | undefined | undefined | undefined | Deletion |
| WRN | MDA-MB-453 | CNA | undefined | undefined | undefined | Loss |
| XPA | MDA-MB-453 | CNA | undefined | undefined | undefined | Loss |
| BRIP1 | MDA-MB-453 | CNA | undefined | undefined | undefined | Amplification |
| BRCA2 | MDA-MB-453 | CNA | undefined | undefined | undefined | Gain |
| CHEK2 | MDA-MB-453 | CNA | undefined | undefined | undefined | Gain |
| ERCC4 | MDA-MB-453 | CNA | undefined | undefined | undefined | Gain |
| FANCE | MDA-MB-453 | CNA | undefined | undefined | undefined | Gain |
| FANCG | MDA-MB-453 | CNA | undefined | undefined | undefined | Gain |
| PALB2 | MDA-MB-453 | CNA | undefined | undefined | undefined | Gain |
| RAD21 | MDA-MB-453 | CNA | undefined | undefined | undefined | Gain |
| RECQL4 | MDA-MB-453 | CNA | undefined | undefined | undefined | Gain |
| SMC1A | MDA-MB-453 | CNA | undefined | undefined | undefined | Gain |
| TCEA1 | T47D | snp | p.G155A | c.464G>C | missense | undefined |
| TP53 | T47D | snp | p.L194F | c.580C>T | missense | undefined |
| POLQ | T47D | snp | p.E964A | c.2891A>C | missense | undefined |
| TDG | T47D | snp | p.K201T | c.602A>C | missense | undefined |
| BLM | T47D | CNA | undefined | undefined | undefined | Loss |
| CHEK2 | T47D | CNA | undefined | undefined | undefined | Loss |
| FANCG | T47D | CNA | undefined | undefined | undefined | Loss |
| MUTYH | T47D | CNA | undefined | undefined | undefined | Loss |

| | | | | | | |
|---|---|---|---|---|---|---|
| PER1 | T47D | CNA | undefined | undefined | undefined | Loss |
| TP53 | T47D | CNA | undefined | undefined | undefined | Loss |
| ATRX | T47D | CNA | undefined | undefined | undefined | Loss |
| BRCA2 | T47D | CNA | undefined | undefined | undefined | Loss |
| DDB2 | T47D | CNA | undefined | undefined | undefined | Gain |
| ERCC2 | T47D | CNA | undefined | undefined | undefined | Loss |
| ERCC3 | T47D | CNA | undefined | undefined | undefined | Loss |
| FANCC | T47D | CNA | undefined | undefined | undefined | Loss |
| MSH2 | T47D | CNA | undefined | undefined | undefined | Loss |
| MSH6 | T47D | CNA | undefined | undefined | undefined | Loss |
| NBN | T47D | CNA | undefined | undefined | undefined | Gain |
| POLD1 | T47D | CNA | undefined | undefined | undefined | Loss |
| UBE2A | T47D | CNA | undefined | undefined | undefined | Loss |
| WRN | T47D | CNA | undefined | undefined | undefined | Loss |
| XPA | T47D | CNA | undefined | undefined | undefined | Loss |
| ABL1 | T47D | CNA | undefined | undefined | undefined | Gain |
| ATR | T47D | CNA | undefined | undefined | undefined | Gain |
| FANCA | T47D | CNA | undefined | undefined | undefined | Gain |
| FANCE | T47D | CNA | undefined | undefined | undefined | Gain |
| FANCF | T47D | CNA | undefined | undefined | undefined | Gain |
| PMS2 | T47D | CNA | undefined | undefined | undefined | Gain |
| POLE | T47D | CNA | undefined | undefined | undefined | Gain |
| RAD21 | T47D | CNA | undefined | undefined | undefined | Gain |
| TCEA1 | T47D | CNA | undefined | undefined | undefined | Gain |
