## Supplementary Table 2 for "Chromatin signalling pathways and FANCE amplification affect ATR inhibitor sensitivity in metastatic breast cancer"

| Gene name | Untreated_R1 | Untreated_R2 | Untreated_R3 | AZD6738_R1 | AZD6738_R2 | AZD6738_R3 | p-value_ATRi/UT | Log2FC_ATRi/UT |
|---|---|---|---|---|---|---|---|---|
| PPIA | 0.17 | 0.18 | 0.18 | -0.21 | -0.19 | -0.21 | 1.3E-06 | -0.38 |
| SUMO4 | -0.64 | -0.56 | -0.57 | 0.43 | 0.41 | 0.41 | 3.3E-06 | 1.01 |
| PAXX | -0.65 | -0.70 | -0.66 | 0.40 | 0.49 | 0.48 | 3.3E-06 | 1.13 |
| MSH3 | -0.25 | -0.26 | -0.23 | 0.20 | 0.22 | 0.21 | 3.8E-06 | 0.46 |
| H1-10 | -0.31 | -0.29 | -0.27 | 0.26 | 0.22 | 0.24 | 4.3E-06 | 0.53 |
| PMVK | -0.32 | -0.28 | -0.34 | 0.25 | 0.25 | 0.27 | 6.7E-06 | 0.57 |
| TGM1 | -0.97 | -1.03 | -0.92 | 0.52 | 0.56 | 0.65 | 7.5E-06 | 1.55 |
| RPA1 | -0.54 | -0.61 | -0.64 | 0.45 | 0.39 | 0.42 | 7.7E-06 | 1.02 |
| PECR | -0.29 | -0.32 | -0.34 | 0.28 | 0.23 | 0.26 | 8.6E-06 | 0.57 |
| PFKM | -0.20 | -0.23 | -0.20 | 0.18 | 0.20 | 0.17 | 9.3E-06 | 0.39 |
| WDHD1 | -0.24 | -0.21 | -0.22 | 0.20 | 0.17 | 0.21 | 1.2E-05 | 0.42 |
| MAP2K3 | -0.26 | -0.25 | -0.30 | 0.21 | 0.22 | 0.25 | 1.5E-05 | 0.50 |
| EGLN1 | -0.17 | -0.16 | -0.14 | 0.14 | 0.14 | 0.15 | 1.6E-05 | 0.30 |
| RPA3 | -0.46 | -0.55 | -0.49 | 0.42 | 0.34 | 0.35 | 1.8E-05 | 0.87 |
| APRT | -0.19 | -0.21 | -0.19 | 0.20 | 0.17 | 0.15 | 2.0E-05 | 0.37 |
| GPD1L | -0.14 | -0.12 | -0.16 | 0.12 | 0.13 | 0.13 | 2.1E-05 | 0.27 |
| SUMO3 | -0.48 | -0.56 | -0.57 | 0.44 | 0.39 | 0.34 | 2.2E-05 | 0.93 |
| RABEP2 | -0.20 | -0.25 | -0.20 | 0.19 | 0.19 | 0.19 | 2.6E-05 | 0.41 |
| TTLL12 | -0.41 | -0.44 | -0.43 | 0.38 | 0.34 | 0.27 | 2.6E-05 | 0.76 |
| HPF1 | -0.15 | -0.17 | -0.19 | 0.14 | 0.14 | 0.17 | 2.7E-05 | 0.32 |
| PSMD12 | -0.19 | -0.25 | -0.22 | 0.19 | 0.18 | 0.21 | 2.9E-05 | 0.42 |
| IPO8 | -0.10 | -0.12 | -0.12 | 0.09 | 0.11 | 0.11 | 2.9E-05 | 0.22 |
| ABCF3 | -0.18 | -0.18 | -0.22 | 0.15 | 0.19 | 0.17 | 3.3E-05 | 0.36 |
| MRE11 | -0.23 | -0.20 | -0.23 | 0.16 | 0.19 | 0.22 | 3.3E-05 | 0.41 |
| OGFR | -0.64 | -0.73 | -0.58 | 0.40 | 0.51 | 0.43 | 3.6E-05 | 1.10 |
| TGOLN2 | 0.29 | 0.28 | 0.29 | -0.40 | -0.38 | -0.30 | 3.6E-05 | -0.65 |
| RPA2 | -0.61 | -0.69 | -0.60 | 0.50 | 0.35 | 0.46 | 3.6E-05 | 1.07 |
| SCD | 0.28 | 0.36 | 0.33 | -0.42 | -0.37 | -0.47 | 3.7E-05 | -0.75 |
| CDC45 | -0.70 | -0.72 | -0.67 | 0.41 | 0.40 | 0.58 | 3.8E-05 | 1.16 |
| NACC1 | -0.14 | -0.13 | -0.12 | 0.12 | 0.14 | 0.10 | 4.2E-05 | 0.25 |
| RBM12 | -0.54 | -0.55 | -0.69 | 0.45 | 0.39 | 0.41 | 4.2E-05 | 1.01 |
| VAT1 | -0.14 | -0.16 | -0.15 | 0.11 | 0.13 | 0.16 | 4.6E-05 | 0.28 |
| CRTC2 | -0.50 | -0.50 | -0.48 | 0.28 | 0.37 | 0.44 | 4.8E-05 | 0.85 |
| SSH3 | -0.37 | -0.46 | -0.42 | 0.26 | 0.36 | 0.35 | 5.0E-05 | 0.74 |
| IRF9 | -0.28 | -0.24 | -0.26 | 0.18 | 0.26 | 0.21 | 5.0E-05 | 0.48 |
| HSPH1 | -0.10 | -0.11 | -0.12 | 0.10 | 0.08 | 0.12 | 5.5E-05 | 0.21 |
| PSMA5 | -0.41 | -0.42 | -0.39 | 0.24 | 0.36 | 0.36 | 6.2E-05 | 0.73 |
| GINS4 | -0.40 | -0.44 | -0.32 | 0.31 | 0.28 | 0.33 | 6.3E-05 | 0.69 |
| WRNIP1 | -0.19 | -0.21 | -0.22 | 0.22 | 0.15 | 0.18 | 6.5E-05 | 0.39 |
| LRRFIP2 | 0.10 | 0.09 | 0.11 | -0.12 | -0.09 | -0.12 | 7.0E-05 | -0.21 |
| DNM2 | -0.18 | -0.23 | -0.22 | 0.22 | 0.16 | 0.18 | 7.1E-05 | 0.40 |
| HGH1 | -0.65 | -0.53 | -0.48 | 0.42 | 0.34 | 0.44 | 8.4E-05 | 0.95 |
| DNAJC7 | -0.08 | -0.09 | -0.08 | 0.07 | 0.07 | 0.10 | 8.8E-05 | 0.16 |

| | | | | | | | | |
|---|---|---|---|---|---|---|---|---|
| **MAGED2** | -0.22 | -0.24 | -0.29 | 0.19 | 0.25 | 0.20 | 9.1E-05 | 0.46 |
| **GSR** | -0.67 | -0.61 | -0.76 | 0.53 | 0.35 | 0.50 | 9.2E-05 | 1.14 |
| **SUMO2** | -0.41 | -0.48 | -0.59 | 0.33 | 0.41 | 0.36 | 9.6E-05 | 0.86 |
| **CLP1** | -0.10 | -0.11 | -0.13 | 0.12 | 0.11 | 0.09 | 9.8E-05 | 0.22 |
| **TLE5** | -0.31 | -0.32 | -0.34 | 0.19 | 0.32 | 0.27 | 9.9E-05 | 0.58 |
| **ARHGEF2** | -0.25 | -0.19 | -0.21 | 0.22 | 0.16 | 0.17 | 9.9E-05 | 0.40 |
| **HSPB1** | -0.67 | -0.71 | -0.55 | 0.49 | 0.34 | 0.49 | 1.0E-04 | 1.08 |
| **DDIT3** | -1.00 | -0.88 | -0.93 | 0.38 | 0.65 | 0.64 | 1.0E-04 | 1.49 |
| **POLD1** | -0.13 | -0.10 | -0.13 | 0.09 | 0.12 | 0.12 | 1.1E-04 | 0.23 |
| **PSMD11** | -0.13 | -0.14 | -0.15 | 0.10 | 0.16 | 0.13 | 1.1E-04 | 0.27 |
| **RPTOR** | -0.19 | -0.25 | -0.20 | 0.17 | 0.16 | 0.22 | 1.1E-04 | 0.40 |
| **GID8** | -0.09 | -0.08 | -0.12 | 0.09 | 0.10 | 0.09 | 1.2E-04 | 0.19 |
| **AARS1** | -0.11 | -0.17 | -0.13 | 0.11 | 0.13 | 0.13 | 1.2E-04 | 0.26 |
| **IGBP1** | -0.48 | -0.35 | -0.51 | 0.31 | 0.33 | 0.37 | 1.2E-04 | 0.78 |
| **PLD1** | -0.12 | -0.09 | -0.11 | 0.12 | 0.09 | 0.09 | 1.2E-04 | 0.21 |
| **NUB1** | -0.23 | -0.18 | -0.27 | 0.21 | 0.16 | 0.21 | 1.3E-04 | 0.42 |
| **RPS6KA5** | -0.63 | -0.51 | -0.43 | 0.36 | 0.35 | 0.44 | 1.4E-04 | 0.90 |
| **DHRS2** | -0.59 | -0.56 | -0.39 | 0.39 | 0.35 | 0.39 | 1.4E-04 | 0.89 |
| **FASN** | -0.08 | -0.12 | -0.12 | 0.11 | 0.10 | 0.09 | 1.6E-04 | 0.20 |
| **CANX** | 0.14 | 0.10 | 0.13 | -0.14 | -0.15 | -0.11 | 1.6E-04 | -0.25 |
| **DNAJA2** | -0.17 | -0.22 | -0.21 | 0.21 | 0.18 | 0.14 | 1.7E-04 | 0.38 |
| **AKT1** | -0.28 | -0.39 | -0.32 | 0.27 | 0.21 | 0.32 | 1.7E-04 | 0.59 |
| **SET** | 0.11 | 0.08 | 0.13 | -0.11 | -0.13 | -0.11 | 1.8E-04 | -0.22 |
| **PARP14** | -0.09 | -0.08 | -0.07 | 0.06 | 0.07 | 0.09 | 1.8E-04 | 0.15 |
| **RACGAP1** | 0.12 | 0.20 | 0.13 | -0.18 | -0.16 | -0.16 | 1.8E-04 | -0.32 |
| **ALCAM** | 0.17 | 0.28 | 0.26 | -0.28 | -0.32 | -0.25 | 1.8E-04 | -0.52 |
| **MNX1** | -0.22 | -0.30 | -0.34 | 0.24 | 0.27 | 0.20 | 1.8E-04 | 0.52 |
| **PSMD14** | -0.38 | -0.39 | -0.39 | 0.24 | 0.26 | 0.41 | 1.9E-04 | 0.69 |
| **DOP1B** | -0.08 | -0.07 | -0.08 | 0.07 | 0.06 | 0.09 | 1.9E-04 | 0.15 |
| **NCOR1** | -0.32 | -0.21 | -0.24 | 0.22 | 0.24 | 0.19 | 2.0E-04 | 0.47 |
| **SMC4** | -0.12 | -0.14 | -0.11 | 0.11 | 0.08 | 0.14 | 2.0E-04 | 0.23 |
| **KMT5A** | -0.27 | -0.32 | -0.22 | 0.25 | 0.18 | 0.25 | 2.1E-04 | 0.50 |
| **TUBA4A** | -0.20 | -0.21 | -0.24 | 0.18 | 0.24 | 0.14 | 2.2E-04 | 0.40 |
| **CLCN2** | 0.09 | 0.07 | 0.08 | -0.07 | -0.07 | -0.10 | 2.2E-04 | -0.16 |
| **DNMT1** | 0.10 | 0.06 | 0.09 | -0.10 | -0.09 | -0.08 | 2.2E-04 | -0.18 |
| **NCAPH** | -0.17 | -0.19 | -0.17 | 0.11 | 0.17 | 0.19 | 2.2E-04 | 0.33 |
| **VARS1** | -0.16 | -0.16 | -0.15 | 0.10 | 0.14 | 0.18 | 2.3E-04 | 0.30 |
| **ARFGEF1** | -0.14 | -0.13 | -0.15 | 0.09 | 0.16 | 0.14 | 2.3E-04 | 0.27 |
| **TFG** | -0.18 | -0.22 | -0.27 | 0.23 | 0.16 | 0.19 | 2.3E-04 | 0.42 |
| **DTYMK** | -0.10 | -0.15 | -0.13 | 0.11 | 0.10 | 0.14 | 2.4E-04 | 0.24 |
| **PHLDB3** | -0.36 | -0.22 | -0.34 | 0.23 | 0.24 | 0.28 | 2.4E-04 | 0.56 |
| **MRPL33** | 0.18 | 0.15 | 0.14 | -0.16 | -0.15 | -0.22 | 2.4E-04 | -0.33 |
| **HMGCR** | 0.14 | 0.23 | 0.26 | -0.23 | -0.23 | -0.27 | 2.4E-04 | -0.45 |
| **CYB5A** | 0.17 | 0.24 | 0.19 | -0.26 | -0.18 | -0.25 | 2.4E-04 | -0.43 |
| **PSMC3** | -0.16 | -0.19 | -0.17 | 0.13 | 0.20 | 0.13 | 2.4E-04 | 0.33 |
| **CDK5** | -0.27 | -0.21 | -0.19 | 0.24 | 0.15 | 0.19 | 2.5E-04 | 0.42 |
| **SASS6** | -0.19 | -0.27 | -0.24 | 0.15 | 0.22 | 0.23 | 2.5E-04 | 0.43 |
| **C19orf47** | 0.08 | 0.05 | 0.09 | -0.08 | -0.07 | -0.08 | 2.6E-04 | -0.15 |

| | | | | | | | | |
|---|---|---|---|---|---|---|---|---|
| TRIM28 | 0.05 | 0.05 | 0.06 | -0.06 | -0.08 | -0.04 | 2.7E-04 | -0.12 |
| MRPL51 | 0.08 | 0.09 | 0.08 | -0.06 | -0.09 | -0.11 | 2.7E-04 | -0.17 |
| XRCC6 | -0.12 | -0.18 | -0.19 | 0.18 | 0.12 | 0.14 | 2.8E-04 | 0.31 |
| SCRN1 | -0.16 | -0.28 | -0.24 | 0.19 | 0.21 | 0.18 | 3.0E-04 | 0.42 |
| CEBPB | -0.51 | -0.44 | -0.32 | 0.27 | 0.31 | 0.39 | 3.0E-04 | 0.75 |
| SEPTIN9 | 0.10 | 0.09 | 0.05 | -0.09 | -0.09 | -0.08 | 3.0E-04 | -0.17 |
| CTSD | -0.07 | -0.09 | -0.07 | 0.08 | 0.08 | 0.05 | 3.0E-04 | 0.15 |
| CDK4 | -0.52 | -0.52 | -0.35 | 0.40 | 0.27 | 0.37 | 3.3E-04 | 0.81 |
| ZKSCAN1 | -0.12 | -0.12 | -0.17 | 0.15 | 0.09 | 0.13 | 3.4E-04 | 0.26 |
| PSMC4 | -0.23 | -0.27 | -0.27 | 0.14 | 0.24 | 0.27 | 3.4E-04 | 0.47 |
| TNPO1 | -0.12 | -0.15 | -0.09 | 0.12 | 0.10 | 0.12 | 3.5E-04 | 0.23 |
| PSMC2 | -0.15 | -0.22 | -0.14 | 0.12 | 0.18 | 0.15 | 3.5E-04 | 0.32 |
| GET3 | -0.18 | -0.30 | -0.26 | 0.17 | 0.22 | 0.24 | 3.5E-04 | 0.46 |
| TRIP11 | -0.22 | -0.25 | -0.29 | 0.14 | 0.27 | 0.24 | 3.6E-04 | 0.47 |
| CEBPA | 0.19 | 0.14 | 0.14 | -0.21 | -0.13 | -0.18 | 3.6E-04 | -0.33 |
| GOLGA4 | -0.19 | -0.23 | -0.13 | 0.14 | 0.18 | 0.16 | 3.6E-04 | 0.34 |
| RPLP1 | -0.12 | -0.21 | -0.18 | 0.14 | 0.17 | 0.14 | 3.7E-04 | 0.32 |
| TTC19 | -0.19 | -0.20 | -0.30 | 0.21 | 0.17 | 0.21 | 3.7E-04 | 0.43 |
| SETX | 0.04 | 0.05 | 0.05 | -0.03 | -0.06 | -0.05 | 3.8E-04 | -0.10 |
| RAD50 | -0.13 | -0.17 | -0.18 | 0.13 | 0.19 | 0.11 | 3.9E-04 | 0.30 |
| DUS1L | -0.23 | -0.33 | -0.30 | 0.16 | 0.26 | 0.29 | 3.9E-04 | 0.52 |
| SLC26A6 | 0.15 | 0.30 | 0.32 | -0.33 | -0.31 | -0.32 | 4.0E-04 | -0.58 |
| IMPDH2 | -0.14 | -0.24 | -0.16 | 0.16 | 0.17 | 0.14 | 4.0E-04 | 0.34 |
| HELLS | 0.17 | 0.18 | 0.17 | -0.27 | -0.19 | -0.15 | 4.1E-04 | -0.38 |
| CBX5 | 0.04 | 0.03 | 0.04 | -0.03 | -0.04 | -0.05 | 4.1E-04 | -0.07 |
| PSPC1 | 0.05 | 0.07 | 0.05 | -0.04 | -0.07 | -0.06 | 4.2E-04 | -0.11 |
| SNX3 | -0.09 | -0.10 | -0.12 | 0.09 | 0.13 | 0.07 | 4.2E-04 | 0.20 |
| H3-3A | -0.09 | -0.14 | -0.12 | 0.09 | 0.10 | 0.14 | 4.3E-04 | 0.23 |
| HSP90AB1 | -0.08 | -0.09 | -0.12 | 0.08 | 0.11 | 0.09 | 4.3E-04 | 0.19 |
| BTAF1 | -0.14 | -0.17 | -0.18 | 0.19 | 0.10 | 0.14 | 4.4E-04 | 0.31 |
| SLC25A17 | 0.23 | 0.17 | 0.14 | -0.21 | -0.16 | -0.25 | 4.4E-04 | -0.39 |
| EIPR1 | -0.09 | -0.17 | -0.16 | 0.12 | 0.12 | 0.14 | 4.5E-04 | 0.27 |
| PHGDH | -0.16 | -0.23 | -0.17 | 0.14 | 0.22 | 0.14 | 4.5E-04 | 0.36 |
| SNRPD2 | 0.10 | 0.08 | 0.11 | -0.14 | -0.08 | -0.09 | 4.6E-04 | -0.20 |
| FASTKD2 | 0.13 | 0.17 | 0.16 | -0.23 | -0.17 | -0.13 | 4.6E-04 | -0.33 |
| SEC23IP | -0.13 | -0.20 | -0.18 | 0.11 | 0.19 | 0.16 | 4.7E-04 | 0.32 |
| ARMC6 | -0.42 | -0.37 | -0.26 | 0.23 | 0.25 | 0.36 | 4.7E-04 | 0.63 |
| COX18 | 0.13 | 0.25 | 0.17 | -0.23 | -0.21 | -0.19 | 4.7E-04 | -0.40 |
| MRPS36 | 0.17 | 0.27 | 0.18 | -0.23 | -0.21 | -0.29 | 4.7E-04 | -0.45 |
| NQO1 | -0.20 | -0.23 | -0.31 | 0.18 | 0.27 | 0.19 | 4.8E-04 | 0.46 |
| UBA1 | -0.09 | -0.12 | -0.12 | 0.07 | 0.11 | 0.12 | 4.8E-04 | 0.21 |
| CFAP299 | 0.23 | 0.29 | 0.38 | -0.30 | -0.39 | -0.47 | 4.9E-04 | -0.69 |
| RAVER1 | -0.24 | -0.19 | -0.33 | 0.18 | 0.22 | 0.25 | 4.9E-04 | 0.47 |
| TUBB4B | -0.15 | -0.20 | -0.21 | 0.22 | 0.14 | 0.13 | 5.0E-04 | 0.35 |
| MAPK14 | -0.56 | -0.60 | -0.88 | 0.50 | 0.37 | 0.50 | 5.1E-04 | 1.14 |
| GTPBP2 | -0.62 | -0.38 | -0.44 | 0.43 | 0.34 | 0.31 | 5.2E-04 | 0.84 |
| SUZ12 | 0.08 | 0.10 | 0.07 | -0.10 | -0.06 | -0.10 | 5.3E-04 | -0.17 |
| TTC39A | -0.17 | -0.09 | -0.14 | 0.14 | 0.12 | 0.10 | 5.3E-04 | 0.25 |

| | | | | | | | | |
|---|---|---|---|---|---|---|---|---|
| CMPK1 | -0.11 | -0.11 | -0.18 | 0.14 | 0.11 | 0.11 | 5.3E-04 | 0.25 |
| DSCC1 | -0.34 | -0.35 | -0.41 | 0.37 | 0.17 | 0.33 | 5.3E-04 | 0.66 |
| CBX3 | 0.13 | 0.11 | 0.09 | -0.10 | -0.10 | -0.16 | 5.5E-04 | -0.23 |
| RPS25 | -0.12 | -0.11 | -0.06 | 0.09 | 0.08 | 0.10 | 5.5E-04 | 0.19 |
| PSMC5 | -0.11 | -0.15 | -0.15 | 0.16 | 0.13 | 0.08 | 5.5E-04 | 0.26 |
| MUC5B | 0.22 | 0.30 | 0.31 | -0.36 | -0.24 | -0.43 | 5.6E-04 | -0.62 |
| MVK | -0.39 | -0.39 | -0.23 | 0.24 | 0.25 | 0.32 | 5.7E-04 | 0.61 |
| HCCS | 0.18 | 0.21 | 0.31 | -0.31 | -0.22 | -0.32 | 5.7E-04 | -0.52 |
| JPT1 | 0.30 | 0.36 | 0.16 | -0.34 | -0.31 | -0.38 | 5.8E-04 | -0.62 |
| TRIP13 | -0.23 | -0.36 | -0.22 | 0.19 | 0.27 | 0.21 | 5.8E-04 | 0.50 |
| FBXW11 | 0.20 | 0.16 | 0.24 | -0.28 | -0.16 | -0.27 | 5.8E-04 | -0.44 |
| NR2C2 | -0.10 | -0.09 | -0.13 | 0.09 | 0.13 | 0.07 | 5.8E-04 | 0.20 |
| INPPL1 | -0.21 | -0.21 | -0.18 | 0.23 | 0.19 | 0.11 | 5.9E-04 | 0.38 |
| EXOC4 | -0.08 | -0.12 | -0.14 | 0.08 | 0.11 | 0.13 | 5.9E-04 | 0.22 |
| GRB10 | -0.23 | -0.24 | -0.31 | 0.28 | 0.15 | 0.23 | 5.9E-04 | 0.48 |
| WDR90 | 0.12 | 0.15 | 0.13 | -0.09 | -0.18 | -0.17 | 6.0E-04 | -0.28 |
| GPC4 | 0.12 | 0.18 | 0.15 | -0.19 | -0.11 | -0.19 | 6.1E-04 | -0.31 |
| GCN1 | -0.11 | -0.17 | -0.11 | 0.09 | 0.13 | 0.14 | 6.3E-04 | 0.25 |
| ITSN2 | -0.18 | -0.11 | -0.16 | 0.12 | 0.18 | 0.11 | 6.4E-04 | 0.29 |
| IRAK1 | -0.19 | -0.20 | -0.20 | 0.11 | 0.24 | 0.16 | 6.5E-04 | 0.37 |
| PIH1D2 | 0.24 | 0.23 | 0.34 | -0.36 | -0.23 | -0.42 | 6.6E-04 | -0.61 |
| BAG5 | -0.35 | -0.25 | -0.19 | 0.24 | 0.18 | 0.25 | 6.7E-04 | 0.48 |
| CTNNB1 | 0.12 | 0.12 | 0.17 | -0.12 | -0.14 | -0.20 | 6.7E-04 | -0.29 |
| HNRNPUL1 | -0.16 | -0.23 | -0.19 | 0.20 | 0.10 | 0.21 | 7.0E-04 | 0.36 |
| RELA | -0.16 | -0.24 | -0.12 | 0.15 | 0.16 | 0.15 | 7.0E-04 | 0.32 |
| BAP18 | -0.18 | -0.17 | -0.22 | 0.13 | 0.14 | 0.23 | 7.0E-04 | 0.36 |
| DHX30 | 0.04 | 0.07 | 0.09 | -0.06 | -0.07 | -0.08 | 7.1E-04 | -0.13 |
| SLC43A3 | 0.09 | 0.17 | 0.17 | -0.12 | -0.17 | -0.18 | 7.1E-04 | -0.30 |
| GPD2 | 0.13 | 0.14 | 0.08 | -0.15 | -0.14 | -0.09 | 7.2E-04 | -0.24 |
| PSMD2 | -0.12 | -0.19 | -0.19 | 0.17 | 0.18 | 0.10 | 7.3E-04 | 0.32 |
| UCK2 | -0.21 | -0.32 | -0.33 | 0.21 | 0.32 | 0.18 | 7.3E-04 | 0.53 |
| ARF4 | -0.21 | -0.29 | -0.25 | 0.12 | 0.26 | 0.25 | 7.3E-04 | 0.46 |
| C17orf58 | -0.39 | -0.30 | -0.36 | 0.29 | 0.38 | 0.17 | 7.3E-04 | 0.63 |
| RIPK1 | -0.25 | -0.28 | -0.32 | 0.28 | 0.30 | 0.13 | 7.4E-04 | 0.52 |
| PPP1R3D | -0.30 | -0.46 | -0.43 | 0.22 | 0.29 | 0.41 | 7.4E-04 | 0.70 |
| TWF2 | 0.09 | 0.06 | 0.09 | -0.08 | -0.12 | -0.06 | 7.5E-04 | -0.17 |
| RAB7A | 0.06 | 0.11 | 0.12 | -0.12 | -0.10 | -0.10 | 7.6E-04 | -0.20 |
| MRPL3 | 0.08 | 0.05 | 0.08 | -0.07 | -0.06 | -0.10 | 7.9E-04 | -0.15 |
| PSMD1 | -0.16 | -0.30 | -0.24 | 0.16 | 0.18 | 0.25 | 8.1E-04 | 0.43 |
| POLR3B | -0.22 | -0.19 | -0.14 | 0.17 | 0.11 | 0.21 | 8.1E-04 | 0.34 |
| DPM1 | 0.16 | 0.13 | 0.12 | -0.20 | -0.16 | -0.10 | 8.3E-04 | -0.29 |
| MRPL14 | 0.12 | 0.17 | 0.17 | -0.20 | -0.11 | -0.20 | 8.4E-04 | -0.32 |
| SDSL | -0.39 | -0.26 | -0.24 | 0.22 | 0.31 | 0.19 | 8.4E-04 | 0.54 |
| HSPA8 | -0.11 | -0.09 | -0.08 | 0.07 | 0.07 | 0.12 | 8.5E-04 | 0.18 |
| CCDC124 | 0.18 | 0.17 | 0.15 | -0.17 | -0.14 | -0.27 | 8.5E-04 | -0.36 |
| GUK1 | -0.20 | -0.21 | -0.14 | 0.22 | 0.15 | 0.11 | 8.5E-04 | 0.34 |
| YJU2 | 0.13 | 0.06 | 0.15 | -0.12 | -0.13 | -0.11 | 8.5E-04 | -0.24 |
| APC | -0.71 | -0.79 | -0.98 | 0.42 | 0.35 | 0.76 | 8.6E-04 | 1.34 |

| | | | | | | | | |
|---|---|---|---|---|---|---|---|---|
| **TMOD2** | -0.24 | -0.38 | -0.40 | 0.22 | 0.23 | 0.37 | 8.7E-04 | 0.61 |
| **FAM133B** | 0.25 | 0.16 | 0.15 | -0.15 | -0.24 | -0.25 | 9.1E-04 | -0.40 |
| **RNF14** | -0.26 | -0.21 | -0.33 | 0.25 | 0.28 | 0.14 | 9.1E-04 | 0.49 |
| **CASKIN2** | -0.35 | -0.35 | -0.19 | 0.18 | 0.26 | 0.29 | 9.1E-04 | 0.54 |
| **SYNE1** | 0.12 | 0.23 | 0.23 | -0.27 | -0.23 | -0.17 | 9.1E-04 | -0.42 |
| **FIS1** | 0.14 | 0.21 | 0.25 | -0.22 | -0.18 | -0.31 | 9.2E-04 | -0.44 |
| **NMRAL1** | -0.21 | -0.32 | -0.36 | 0.26 | 0.31 | 0.16 | 9.3E-04 | 0.54 |
| **HERC5** | -0.12 | -0.20 | -0.24 | 0.14 | 0.15 | 0.20 | 9.4E-04 | 0.35 |
| **DHFR** | -0.23 | -0.25 | -0.27 | 0.13 | 0.20 | 0.31 | 9.4E-04 | 0.46 |
| **FUBP1** | 0.11 | 0.14 | 0.13 | -0.18 | -0.15 | -0.08 | 9.4E-04 | -0.26 |
| **CHORDC1** | 0.13 | 0.09 | 0.10 | -0.16 | -0.10 | -0.09 | 9.4E-04 | -0.22 |
| **OGDH** | 0.08 | 0.18 | 0.18 | -0.14 | -0.16 | -0.20 | 9.5E-04 | -0.32 |
| **ASMTL** | -0.18 | -0.16 | -0.19 | 0.09 | 0.22 | 0.16 | 9.6E-04 | 0.33 |
| **UBE2V1** | 0.38 | 0.21 | 0.33 | -0.27 | -0.47 | -0.44 | 9.7E-04 | -0.70 |
| **PCK2** | -0.08 | -0.15 | -0.18 | 0.11 | 0.13 | 0.14 | 9.8E-04 | 0.27 |
| **ARFGEF2** | -0.09 | -0.19 | -0.13 | 0.10 | 0.14 | 0.13 | 9.8E-04 | 0.26 |
| **CAD** | -0.12 | -0.21 | -0.14 | 0.10 | 0.18 | 0.14 | 1.0E-03 | 0.29 |
| **ODR4** | 0.06 | 0.10 | 0.08 | -0.08 | -0.07 | -0.11 | 1.0E-03 | -0.17 |
| **GALK1** | -0.24 | -0.26 | -0.22 | 0.11 | 0.23 | 0.28 | 1.0E-03 | 0.45 |
| **TXN** | 0.24 | 0.15 | 0.22 | -0.30 | -0.27 | -0.15 | 1.0E-03 | -0.44 |
| **LUC7L** | 0.11 | 0.12 | 0.16 | -0.09 | -0.18 | -0.16 | 1.0E-03 | -0.27 |
| **RPUSD3** | 0.09 | 0.08 | 0.06 | -0.10 | -0.10 | -0.05 | 1.0E-03 | -0.16 |
| **EPM2AIP1** | -0.18 | -0.35 | -0.26 | 0.26 | 0.17 | 0.23 | 1.0E-03 | 0.48 |
| **SNX6** | -0.08 | -0.13 | -0.16 | 0.10 | 0.09 | 0.15 | 1.0E-03 | 0.24 |
| **STAT1** | -0.12 | -0.16 | -0.21 | 0.10 | 0.16 | 0.18 | 1.1E-03 | 0.31 |
| **PGK1** | 0.10 | 0.13 | 0.07 | -0.13 | -0.08 | -0.10 | 1.1E-03 | -0.20 |
| **TNPO2** | -0.11 | -0.13 | -0.09 | 0.10 | 0.14 | 0.06 | 1.1E-03 | 0.21 |
| **NUDCD2** | -0.40 | -0.38 | -0.52 | 0.30 | 0.21 | 0.48 | 1.1E-03 | 0.76 |
| **RHOA** | 0.16 | 0.30 | 0.27 | -0.26 | -0.23 | -0.39 | 1.1E-03 | -0.54 |
| **PLAA** | -0.10 | -0.11 | -0.13 | 0.13 | 0.12 | 0.06 | 1.1E-03 | 0.22 |
| **UBR7** | -0.33 | -0.37 | -0.18 | 0.28 | 0.18 | 0.27 | 1.1E-03 | 0.53 |
| **TRIM16** | -0.19 | -0.19 | -0.32 | 0.16 | 0.25 | 0.18 | 1.1E-03 | 0.43 |
| **FBRSL1** | 0.22 | 0.15 | 0.13 | -0.20 | -0.24 | -0.13 | 1.1E-03 | -0.36 |
| **FKBP8** | 0.11 | 0.10 | 0.19 | -0.14 | -0.13 | -0.17 | 1.1E-03 | -0.28 |
| **PARP10** | -0.28 | -0.58 | -0.39 | 0.34 | 0.34 | 0.28 | 1.1E-03 | 0.74 |
| **MICOS10** | 0.14 | 0.26 | 0.30 | -0.36 | -0.25 | -0.24 | 1.1E-03 | -0.52 |
| **TOR1AIP2** | 0.25 | 0.28 | 0.18 | -0.32 | -0.18 | -0.36 | 1.1E-03 | -0.52 |
| **DCD** | -0.38 | -0.51 | -0.65 | 0.50 | 0.36 | 0.26 | 1.1E-03 | 0.89 |
| **CEBPZOS** | 0.19 | 0.37 | 0.20 | -0.29 | -0.27 | -0.39 | 1.1E-03 | -0.57 |
| **HNRNPD** | 0.06 | 0.05 | 0.10 | -0.09 | -0.07 | -0.06 | 1.2E-03 | -0.14 |
| **CLTC** | -0.12 | -0.12 | -0.06 | 0.11 | 0.08 | 0.09 | 1.2E-03 | 0.19 |
| **EIF5** | 0.19 | 0.16 | 0.12 | -0.16 | -0.12 | -0.24 | 1.2E-03 | -0.33 |
| **SLC25A30** | 0.13 | 0.34 | 0.24 | -0.32 | -0.25 | -0.27 | 1.2E-03 | -0.52 |
| **ISG20** | -0.13 | -0.20 | -0.23 | 0.23 | 0.12 | 0.15 | 1.2E-03 | 0.35 |
| **PPIL1** | 0.06 | 0.04 | 0.05 | -0.05 | -0.07 | -0.03 | 1.2E-03 | -0.10 |
| **AFG2A** | 0.07 | 0.07 | 0.11 | -0.12 | -0.06 | -0.09 | 1.2E-03 | -0.17 |
| **EIF4A1** | -0.23 | -0.19 | -0.13 | 0.11 | 0.21 | 0.15 | 1.2E-03 | 0.34 |
| **GINS3** | -0.28 | -0.25 | -0.35 | 0.22 | 0.15 | 0.35 | 1.2E-03 | 0.53 |

| | | | | | | | | |
|---|---|---|---|---|---|---|---|---|
| RPF1 | 0.05 | 0.07 | 0.05 | -0.06 | -0.03 | -0.07 | 1.2E-03 | -0.11 |
| TRIM14 | -0.18 | -0.23 | -0.25 | 0.16 | 0.13 | 0.28 | 1.2E-03 | 0.41 |
| RNH1 | -0.14 | -0.11 | -0.13 | 0.12 | 0.06 | 0.16 | 1.2E-03 | 0.24 |
| DDX39A | 0.13 | 0.17 | 0.10 | -0.13 | -0.11 | -0.20 | 1.3E-03 | -0.28 |
| VASN | 0.10 | 0.27 | 0.22 | -0.20 | -0.22 | -0.26 | 1.3E-03 | -0.42 |
| CISD2 | 0.12 | 0.15 | 0.13 | -0.21 | -0.10 | -0.13 | 1.3E-03 | -0.28 |
| ACBD3 | -0.17 | -0.18 | -0.14 | 0.20 | 0.14 | 0.08 | 1.3E-03 | 0.30 |
| CEBPG | -0.17 | -0.26 | -0.32 | 0.15 | 0.28 | 0.20 | 1.3E-03 | 0.46 |
| GBF1 | -0.18 | -0.13 | -0.08 | 0.10 | 0.14 | 0.11 | 1.3E-03 | 0.24 |
| FECH | -0.07 | -0.08 | -0.11 | 0.09 | 0.05 | 0.10 | 1.3E-03 | 0.17 |
| CLCA2 | 0.11 | 0.12 | 0.19 | -0.14 | -0.12 | -0.21 | 1.3E-03 | -0.30 |
| PPFIA1 | -0.16 | -0.08 | -0.20 | 0.12 | 0.15 | 0.14 | 1.3E-03 | 0.28 |
| PVR | 0.21 | 0.15 | 0.36 | -0.34 | -0.25 | -0.29 | 1.3E-03 | -0.54 |
| UHRF1 | 0.09 | 0.13 | 0.09 | -0.15 | -0.12 | -0.07 | 1.3E-03 | -0.21 |
| UBA6 | -0.13 | -0.19 | -0.13 | 0.09 | 0.19 | 0.13 | 1.3E-03 | 0.28 |
| SLC25A29 | 0.11 | 0.29 | 0.29 | -0.24 | -0.28 | -0.31 | 1.3E-03 | -0.51 |
| UTRN | -0.22 | -0.17 | -0.12 | 0.14 | 0.11 | 0.21 | 1.4E-03 | 0.32 |
| GATD1 | -0.16 | -0.11 | -0.08 | 0.14 | 0.08 | 0.10 | 1.4E-03 | 0.23 |
| CBX4 | -0.19 | -0.17 | -0.18 | 0.21 | 0.19 | 0.08 | 1.4E-03 | 0.34 |
| RAMAC | 0.17 | 0.13 | 0.13 | -0.09 | -0.17 | -0.22 | 1.4E-03 | -0.31 |
| H2AC18 | -0.09 | -0.12 | -0.10 | 0.06 | 0.08 | 0.14 | 1.4E-03 | 0.20 |
| NUSAP1 | 0.06 | 0.15 | 0.12 | -0.11 | -0.11 | -0.14 | 1.4E-03 | -0.22 |
| VPS52 | -0.10 | -0.17 | -0.12 | 0.10 | 0.09 | 0.17 | 1.4E-03 | 0.25 |
| NFKB1 | -0.26 | -0.34 | -0.31 | 0.29 | 0.12 | 0.34 | 1.4E-03 | 0.55 |
| YME1L1 | 0.05 | 0.14 | 0.13 | -0.13 | -0.10 | -0.12 | 1.4E-03 | -0.22 |
| METAP2 | 0.22 | 0.25 | 0.24 | -0.29 | -0.17 | -0.40 | 1.4E-03 | -0.52 |
| ABHD12 | 0.18 | 0.12 | 0.26 | -0.25 | -0.15 | -0.25 | 1.4E-03 | -0.41 |
| TRAPPC11 | -0.19 | -0.12 | -0.24 | 0.10 | 0.19 | 0.18 | 1.5E-03 | 0.34 |
| TCAF1 | 0.07 | 0.10 | 0.12 | -0.12 | -0.06 | -0.13 | 1.5E-03 | -0.21 |
| SRP72 | -0.08 | -0.15 | -0.13 | 0.12 | 0.14 | 0.07 | 1.5E-03 | 0.23 |
| EXD2 | 0.09 | 0.12 | 0.15 | -0.15 | -0.08 | -0.17 | 1.5E-03 | -0.25 |
| GABARAPL2 | 0.21 | 0.10 | 0.11 | -0.19 | -0.12 | -0.16 | 1.5E-03 | -0.30 |
| XRCC5 | -0.12 | -0.17 | -0.17 | 0.11 | 0.10 | 0.21 | 1.5E-03 | 0.29 |
| MSH2 | -0.12 | -0.12 | -0.23 | 0.13 | 0.16 | 0.14 | 1.5E-03 | 0.30 |
| ATIC | -0.18 | -0.22 | -0.09 | 0.13 | 0.14 | 0.17 | 1.5E-03 | 0.31 |
| HSPA1B | -0.06 | -0.05 | -0.07 | 0.03 | 0.07 | 0.07 | 1.5E-03 | 0.12 |
| CTBP1 | -0.18 | -0.24 | -0.15 | 0.12 | 0.14 | 0.24 | 1.5E-03 | 0.35 |
| JUN | -0.56 | -0.63 | -0.51 | 0.24 | 0.32 | 0.63 | 1.5E-03 | 0.96 |
| MOB2 | -0.26 | -0.14 | -0.30 | 0.14 | 0.23 | 0.22 | 1.5E-03 | 0.43 |
| TMEM141 | 0.15 | 0.34 | 0.26 | -0.30 | -0.23 | -0.40 | 1.6E-03 | -0.56 |
| DNAJC15 | 0.19 | 0.33 | 0.27 | -0.45 | -0.30 | -0.23 | 1.6E-03 | -0.59 |
| ERO1A | -0.03 | -0.06 | -0.08 | 0.06 | 0.04 | 0.06 | 1.6E-03 | 0.11 |
| KRT7 | -0.17 | -0.08 | -0.17 | 0.15 | 0.14 | 0.10 | 1.6E-03 | 0.27 |
| MRPS16 | 0.16 | 0.09 | 0.09 | -0.16 | -0.09 | -0.12 | 1.6E-03 | -0.23 |
| MICU2 | 0.07 | 0.07 | 0.05 | -0.04 | -0.09 | -0.08 | 1.6E-03 | -0.13 |
| EPS8L1 | -0.13 | -0.10 | -0.07 | 0.13 | 0.09 | 0.06 | 1.6E-03 | 0.19 |
| IKBKG | -0.39 | -0.45 | -0.22 | 0.20 | 0.37 | 0.27 | 1.6E-03 | 0.63 |
| NOL6 | 0.10 | 0.10 | 0.07 | -0.08 | -0.07 | -0.14 | 1.6E-03 | -0.19 |

| | | | | | | | | |
|---|---|---|---|---|---|---|---|---|
| CWC22 | 0.07 | 0.05 | 0.03 | -0.05 | -0.06 | -0.04 | 1.6E-03 | -0.10 |
| PSMD13 | -0.11 | -0.17 | -0.17 | 0.16 | 0.17 | 0.07 | 1.7E-03 | 0.29 |
| USP9X | -0.09 | -0.18 | -0.09 | 0.10 | 0.12 | 0.12 | 1.7E-03 | 0.23 |
| H2BC26 | -0.54 | -0.80 | -0.94 | 0.54 | 0.26 | 0.65 | 1.7E-03 | 1.24 |
| NUP43 | -0.17 | -0.17 | -0.30 | 0.21 | 0.13 | 0.21 | 1.7E-03 | 0.40 |
| PACSIN2 | 0.10 | 0.08 | 0.11 | -0.14 | -0.12 | -0.06 | 1.7E-03 | -0.20 |
| WARS1 | -0.16 | -0.15 | -0.28 | 0.12 | 0.21 | 0.19 | 1.7E-03 | 0.37 |
| HSPA4L | -0.04 | -0.08 | -0.05 | 0.07 | 0.05 | 0.04 | 1.7E-03 | 0.11 |
| MBTPS1 | 0.09 | 0.19 | 0.20 | -0.18 | -0.13 | -0.24 | 1.7E-03 | -0.34 |
| PSMF1 | -0.24 | -0.16 | -0.10 | 0.16 | 0.13 | 0.16 | 1.7E-03 | 0.32 |
| PRKDC | -0.07 | -0.12 | -0.12 | 0.07 | 0.09 | 0.14 | 1.8E-03 | 0.20 |
| RAB8A | 0.14 | 0.19 | 0.17 | -0.13 | -0.17 | -0.28 | 1.8E-03 | -0.36 |
| GMFB | 0.20 | 0.21 | 0.24 | -0.35 | -0.14 | -0.28 | 1.8E-03 | -0.47 |
| KRT8 | -0.24 | -0.12 | -0.22 | 0.23 | 0.12 | 0.15 | 1.8E-03 | 0.36 |
| FUNDC2 | 0.15 | 0.19 | 0.37 | -0.32 | -0.26 | -0.27 | 1.8E-03 | -0.52 |
| NBN | -0.24 | -0.14 | -0.26 | 0.14 | 0.26 | 0.15 | 1.8E-03 | 0.40 |
| SNX1 | -0.06 | -0.07 | -0.11 | 0.05 | 0.08 | 0.10 | 1.8E-03 | 0.16 |
| BCL2L13 | 0.14 | 0.40 | 0.34 | -0.44 | -0.29 | -0.40 | 1.8E-03 | -0.67 |
| TFAP4 | 0.14 | 0.11 | 0.23 | -0.24 | -0.16 | -0.15 | 1.8E-03 | -0.34 |
| ZNF106 | 0.06 | 0.13 | 0.15 | -0.08 | -0.14 | -0.15 | 1.8E-03 | -0.23 |
| H1-0 | -0.22 | -0.28 | -0.24 | 0.22 | 0.10 | 0.31 | 1.8E-03 | 0.46 |
| SLC25A40 | 0.06 | 0.17 | 0.17 | -0.16 | -0.12 | -0.17 | 1.8E-03 | -0.28 |
| ARFGEF3 | -0.16 | -0.13 | -0.07 | 0.09 | 0.10 | 0.15 | 1.8E-03 | 0.24 |
| LYPLA1 | -0.13 | -0.06 | -0.10 | 0.06 | 0.08 | 0.12 | 1.8E-03 | 0.19 |
| ISYNA1 | -0.22 | -0.23 | -0.12 | 0.13 | 0.14 | 0.23 | 1.9E-03 | 0.35 |
| PTEN | -0.43 | -0.38 | -0.27 | 0.19 | 0.22 | 0.43 | 1.9E-03 | 0.64 |
| CDH3 | 0.08 | 0.09 | 0.16 | -0.09 | -0.15 | -0.12 | 1.9E-03 | -0.23 |
| USP5 | -0.08 | -0.17 | -0.14 | 0.16 | 0.12 | 0.08 | 1.9E-03 | 0.25 |
| USP22 | 0.20 | 0.10 | 0.13 | -0.22 | -0.11 | -0.16 | 1.9E-03 | -0.31 |
| DST | -0.22 | -0.19 | -0.13 | 0.10 | 0.14 | 0.23 | 1.9E-03 | 0.34 |
| MRPL2 | 0.12 | 0.14 | 0.07 | -0.11 | -0.09 | -0.17 | 1.9E-03 | -0.23 |
| GDF15 | -1.04 | -0.72 | -0.49 | 0.40 | 0.44 | 0.59 | 1.9E-03 | 1.23 |
| TDRKH | 0.08 | 0.11 | 0.18 | -0.15 | -0.10 | -0.15 | 1.9E-03 | -0.26 |
| CALML5 | 0.19 | 0.32 | 0.25 | -0.45 | -0.23 | -0.27 | 1.9E-03 | -0.57 |
| POR | 0.07 | 0.03 | 0.07 | -0.06 | -0.06 | -0.05 | 1.9E-03 | -0.12 |
| ARHGEF16 | -0.19 | -0.23 | -0.22 | 0.12 | 0.15 | 0.29 | 1.9E-03 | 0.40 |
| CMC2 | 0.14 | 0.15 | 0.09 | -0.18 | -0.08 | -0.16 | 1.9E-03 | -0.27 |
| CCDC51 | 0.07 | 0.21 | 0.16 | -0.18 | -0.16 | -0.15 | 1.9E-03 | -0.31 |
| MRPL32 | 0.11 | 0.14 | 0.13 | -0.17 | -0.07 | -0.18 | 2.0E-03 | -0.27 |
| LCLAT1 | 0.16 | 0.18 | 0.13 | -0.15 | -0.12 | -0.26 | 2.0E-03 | -0.33 |
| BMP2K | -0.25 | -0.15 | -0.15 | 0.18 | 0.10 | 0.21 | 2.0E-03 | 0.34 |
| UGGT1 | 0.16 | 0.14 | 0.07 | -0.18 | -0.10 | -0.13 | 2.0E-03 | -0.26 |
| RNGTT | -0.23 | -0.23 | -0.33 | 0.19 | 0.33 | 0.13 | 2.0E-03 | 0.48 |
| GATA2 | 0.10 | 0.09 | 0.14 | -0.17 | -0.13 | -0.07 | 2.0E-03 | -0.23 |
| HYOU1 | -0.08 | -0.13 | -0.20 | 0.15 | 0.11 | 0.11 | 2.0E-03 | 0.26 |
| MSH6 | -0.11 | -0.18 | -0.19 | 0.08 | 0.14 | 0.20 | 2.0E-03 | 0.30 |
| ORC5 | -0.12 | -0.19 | -0.14 | 0.14 | 0.07 | 0.19 | 2.1E-03 | 0.28 |
| MYBL1 | 0.04 | 0.10 | 0.06 | -0.08 | -0.05 | -0.09 | 2.1E-03 | -0.14 |

| | | | | | | | | |
|---|---|---|---|---|---|---|---|---|
| **PABPN1** | 0.10 | 0.06 | 0.09 | -0.07 | -0.07 | -0.13 | 2.1E-03 | -0.17 |
| **COPG1** | -0.13 | -0.18 | -0.08 | 0.15 | 0.12 | 0.08 | 2.1E-03 | 0.25 |
| **EEF1AKMT4-ECE2** | -0.52 | -0.52 | -0.40 | 0.24 | 0.57 | 0.25 | 2.1E-03 | 0.84 |
| **STK38L** | -0.17 | -0.16 | -0.33 | 0.18 | 0.17 | 0.22 | 2.1E-03 | 0.41 |
| **PDXDC1** | -0.18 | -0.24 | -0.29 | 0.18 | 0.30 | 0.12 | 2.1E-03 | 0.44 |
| **BRPF3** | 0.07 | 0.10 | 0.04 | -0.10 | -0.06 | -0.07 | 2.1E-03 | -0.15 |
| **POLR3A** | -0.17 | -0.15 | -0.08 | 0.08 | 0.12 | 0.17 | 2.1E-03 | 0.26 |
| **CORO7** | -0.19 | -0.22 | -0.20 | 0.11 | 0.28 | 0.13 | 2.1E-03 | 0.38 |
| **KPNA1** | -0.08 | -0.08 | -0.06 | 0.11 | 0.05 | 0.05 | 2.1E-03 | 0.14 |
| **PCCA** | 0.33 | 0.24 | 0.17 | -0.31 | -0.20 | -0.42 | 2.1E-03 | -0.56 |
| **CDC42EP4** | 0.12 | 0.17 | 0.17 | -0.10 | -0.16 | -0.25 | 2.1E-03 | -0.32 |
| **HBA1** | -0.13 | -0.26 | -0.13 | 0.12 | 0.16 | 0.17 | 2.1E-03 | 0.32 |
| **HP1BP3** | -0.05 | -0.12 | -0.11 | 0.08 | 0.07 | 0.11 | 2.2E-03 | 0.18 |
| **IGSF3** | 0.11 | 0.23 | 0.37 | -0.30 | -0.29 | -0.28 | 2.2E-03 | -0.52 |
| **BLVRA** | -0.17 | -0.32 | -0.14 | 0.16 | 0.22 | 0.17 | 2.2E-03 | 0.39 |
| **RPS3** | -0.10 | -0.13 | -0.07 | 0.13 | 0.08 | 0.06 | 2.2E-03 | 0.19 |
| **GINS1** | -0.12 | -0.28 | -0.20 | 0.20 | 0.13 | 0.19 | 2.2E-03 | 0.38 |
| **ARPC5L** | 0.05 | 0.12 | 0.11 | -0.08 | -0.10 | -0.13 | 2.2E-03 | -0.20 |
| **AK6** | -0.33 | -0.50 | -0.25 | 0.28 | 0.19 | 0.38 | 2.2E-03 | 0.64 |
| **COG5** | -0.07 | -0.11 | -0.17 | 0.09 | 0.11 | 0.11 | 2.2E-03 | 0.22 |
| **TIMMDC1** | 0.11 | 0.23 | 0.20 | -0.15 | -0.17 | -0.29 | 2.2E-03 | -0.38 |
| **PC** | -0.04 | -0.11 | -0.09 | 0.08 | 0.06 | 0.08 | 2.2E-03 | 0.15 |
| **SGTA** | -0.10 | -0.16 | -0.11 | 0.11 | 0.07 | 0.17 | 2.2E-03 | 0.24 |
| **PEX6** | -0.08 | -0.11 | -0.16 | 0.14 | 0.07 | 0.11 | 2.2E-03 | 0.22 |
| **CDC42SE1** | 0.14 | 0.31 | 0.42 | -0.31 | -0.34 | -0.47 | 2.3E-03 | -0.67 |
| **PTMA** | 0.49 | 0.40 | 0.13 | -0.39 | -0.47 | -0.52 | 2.3E-03 | -0.80 |
| **RMI1** | -0.49 | -0.51 | -0.34 | 0.41 | 0.14 | 0.46 | 2.3E-03 | 0.78 |
| **ZNHIT6** | -0.18 | -0.12 | -0.10 | 0.17 | 0.08 | 0.11 | 2.3E-03 | 0.25 |
| **NHLRC2** | -0.16 | -0.14 | -0.09 | 0.07 | 0.12 | 0.16 | 2.3E-03 | 0.24 |
| **SNX4** | -0.15 | -0.15 | -0.17 | 0.06 | 0.16 | 0.21 | 2.3E-03 | 0.30 |
| **KIF22** | -0.13 | -0.13 | -0.13 | 0.07 | 0.10 | 0.19 | 2.3E-03 | 0.24 |
| **ZDHHC3** | 0.09 | 0.07 | 0.06 | -0.04 | -0.07 | -0.11 | 2.3E-03 | -0.15 |
| **TES** | -0.13 | -0.09 | -0.05 | 0.10 | 0.10 | 0.06 | 2.3E-03 | 0.18 |
| **RGS19** | 0.05 | 0.10 | 0.08 | -0.09 | -0.05 | -0.11 | 2.3E-03 | -0.16 |
| **PSMD4** | -0.11 | -0.13 | -0.22 | 0.10 | 0.14 | 0.18 | 2.3E-03 | 0.29 |
| **SHTN1** | -0.06 | -0.11 | -0.06 | 0.06 | 0.06 | 0.09 | 2.3E-03 | 0.14 |
| **GARS1** | -0.09 | -0.14 | -0.24 | 0.16 | 0.13 | 0.14 | 2.4E-03 | 0.30 |
| **MRPS2** | 0.17 | 0.09 | 0.09 | -0.17 | -0.09 | -0.12 | 2.4E-03 | -0.24 |
| **NHERF1** | 0.27 | 0.34 | 0.15 | -0.23 | -0.28 | -0.42 | 2.4E-03 | -0.56 |
| **DCAF8** | -0.11 | -0.18 | -0.17 | 0.19 | 0.16 | 0.07 | 2.4E-03 | 0.29 |
| **HBD** | -0.11 | -0.20 | -0.10 | 0.09 | 0.16 | 0.11 | 2.4E-03 | 0.26 |
| **ALDH16A1** | -0.11 | -0.16 | -0.13 | 0.18 | 0.06 | 0.13 | 2.4E-03 | 0.26 |
| **WAC** | -0.20 | -0.07 | -0.19 | 0.13 | 0.15 | 0.12 | 2.4E-03 | 0.29 |
| **TUBA1A** | -0.11 | -0.17 | -0.22 | 0.09 | 0.20 | 0.16 | 2.4E-03 | 0.32 |
| **PSMC6** | -0.17 | -0.23 | -0.13 | 0.08 | 0.17 | 0.21 | 2.4E-03 | 0.33 |
| **APOBEC3B** | 0.22 | 0.09 | 0.26 | -0.26 | -0.26 | -0.15 | 2.4E-03 | -0.41 |
| **OXA1L** | 0.06 | 0.15 | 0.10 | -0.15 | -0.10 | -0.08 | 2.4E-03 | -0.21 |
| **SOAT1** | 0.13 | 0.17 | 0.10 | -0.11 | -0.11 | -0.22 | 2.4E-03 | -0.28 |

| | | | | | | | | |
|---|---|---|---|---|---|---|---|---|
| CACTIN | 0.07 | 0.09 | 0.08 | -0.10 | -0.04 | -0.12 | 2.4E-03 | -0.17 |
| MRPL27 | 0.13 | 0.17 | 0.06 | -0.14 | -0.10 | -0.17 | 2.5E-03 | -0.26 |
| CHTF18 | -0.12 | -0.26 | -0.26 | 0.18 | 0.12 | 0.25 | 2.5E-03 | 0.40 |
| GAK | -0.24 | -0.13 | -0.10 | 0.14 | 0.12 | 0.17 | 2.5E-03 | 0.30 |
| MRPL48 | 0.12 | 0.11 | 0.09 | -0.05 | -0.14 | -0.15 | 2.5E-03 | -0.22 |
| NEURL4 | 0.08 | 0.13 | 0.05 | -0.10 | -0.07 | -0.10 | 2.5E-03 | -0.17 |
| AGRN | -0.15 | -0.18 | -0.21 | 0.16 | 0.08 | 0.24 | 2.5E-03 | 0.34 |
| ZC3H14 | 0.08 | 0.06 | 0.03 | -0.06 | -0.05 | -0.08 | 2.5E-03 | -0.12 |
| PDAP1 | 0.06 | 0.12 | 0.09 | -0.08 | -0.14 | -0.08 | 2.5E-03 | -0.19 |
| ATP5F1D | 0.10 | 0.31 | 0.31 | -0.35 | -0.23 | -0.29 | 2.5E-03 | -0.53 |
| DNAJB1 | -0.07 | -0.11 | -0.13 | 0.11 | 0.13 | 0.05 | 2.6E-03 | 0.20 |
| ITIH3 | -0.21 | -0.25 | -0.16 | 0.28 | 0.12 | 0.14 | 2.6E-03 | 0.38 |
| CSRP1 | 0.14 | 0.15 | 0.05 | -0.11 | -0.15 | -0.11 | 2.6E-03 | -0.23 |
| ACOX3 | -0.10 | -0.19 | -0.26 | 0.20 | 0.17 | 0.11 | 2.6E-03 | 0.34 |
| ECE1 | 0.14 | 0.19 | 0.41 | -0.31 | -0.30 | -0.31 | 2.6E-03 | -0.55 |
| AKR1A1 | -0.14 | -0.15 | -0.10 | 0.18 | 0.12 | 0.06 | 2.6E-03 | 0.25 |
| TDP1 | 0.09 | 0.15 | 0.15 | -0.07 | -0.20 | -0.16 | 2.6E-03 | -0.27 |
| TCEA3 | 0.16 | 0.09 | 0.19 | -0.23 | -0.11 | -0.15 | 2.6E-03 | -0.31 |
| SAMHD1 | -0.21 | -0.24 | -0.30 | 0.23 | 0.31 | 0.09 | 2.6E-03 | 0.46 |
| RPS12 | -0.22 | -0.24 | -0.12 | 0.19 | 0.09 | 0.23 | 2.6E-03 | 0.36 |
| RPS2 | -0.08 | -0.12 | -0.11 | 0.14 | 0.11 | 0.04 | 2.6E-03 | 0.20 |
| RBM3 | 0.06 | 0.11 | 0.18 | -0.13 | -0.11 | -0.15 | 2.7E-03 | -0.24 |
| IKBKB | -0.24 | -0.25 | -0.27 | 0.07 | 0.27 | 0.30 | 2.7E-03 | 0.47 |
| FNBP1L | 0.10 | 0.10 | 0.09 | -0.16 | -0.06 | -0.09 | 2.7E-03 | -0.20 |
| MBD4 | 0.03 | 0.06 | 0.07 | -0.07 | -0.05 | -0.04 | 2.7E-03 | -0.11 |
| AFF4 | 0.14 | 0.11 | 0.09 | -0.08 | -0.10 | -0.19 | 2.7E-03 | -0.24 |
| FOSL2 | -0.18 | -0.17 | -0.17 | 0.14 | 0.07 | 0.24 | 2.7E-03 | 0.32 |
| CYRIB | 0.22 | 0.19 | 0.14 | -0.23 | -0.11 | -0.30 | 2.7E-03 | -0.40 |
| CIC | -0.26 | -0.27 | -0.10 | 0.21 | 0.18 | 0.15 | 2.7E-03 | 0.39 |
| SMAP2 | -0.38 | -0.20 | -0.27 | 0.25 | 0.13 | 0.33 | 2.7E-03 | 0.52 |
| PGAM5 | 0.07 | 0.05 | 0.06 | -0.03 | -0.07 | -0.10 | 2.7E-03 | -0.13 |
| RDX | 0.05 | 0.11 | 0.12 | -0.07 | -0.09 | -0.14 | 2.7E-03 | -0.19 |
| OPA3 | 0.33 | 0.22 | 0.38 | -0.60 | -0.29 | -0.32 | 2.7E-03 | -0.71 |
| AP2B1 | -0.08 | -0.05 | -0.04 | 0.07 | 0.05 | 0.04 | 2.8E-03 | 0.11 |
| ERCC6 | -0.16 | -0.40 | -0.29 | 0.30 | 0.17 | 0.22 | 2.8E-03 | 0.51 |
| CDC34 | -0.42 | -0.56 | -0.32 | 0.16 | 0.35 | 0.47 | 2.8E-03 | 0.76 |
| PYCR3 | -0.14 | -0.21 | -0.26 | 0.08 | 0.24 | 0.21 | 2.8E-03 | 0.38 |
| EIF4G2 | -0.12 | -0.10 | -0.05 | 0.12 | 0.06 | 0.07 | 2.8E-03 | 0.18 |
| MLST8 | -0.11 | -0.14 | -0.18 | 0.15 | 0.18 | 0.06 | 2.8E-03 | 0.28 |
| EIF4B | 0.06 | 0.15 | 0.17 | -0.10 | -0.14 | -0.19 | 2.8E-03 | -0.28 |
| MAPK3 | -0.19 | -0.13 | -0.07 | 0.09 | 0.14 | 0.12 | 2.9E-03 | 0.24 |
| DDX28 | 0.06 | 0.08 | 0.06 | -0.10 | -0.04 | -0.06 | 2.9E-03 | -0.13 |
| UCHL5 | -0.13 | -0.17 | -0.18 | 0.09 | 0.11 | 0.23 | 2.9E-03 | 0.30 |
| NIBAN2 | 0.10 | 0.10 | 0.07 | -0.11 | -0.13 | -0.04 | 2.9E-03 | -0.18 |
| MRPL58 | 0.10 | 0.15 | 0.10 | -0.14 | -0.07 | -0.18 | 2.9E-03 | -0.24 |
| DLD | 0.18 | 0.33 | 0.22 | -0.32 | -0.17 | -0.41 | 2.9E-03 | -0.54 |
| EIF2AK4 | -0.14 | -0.07 | -0.13 | 0.15 | 0.10 | 0.06 | 2.9E-03 | 0.21 |
| PTGES3 | -0.15 | -0.23 | -0.17 | 0.23 | 0.07 | 0.18 | 2.9E-03 | 0.35 |

| | | | | | | | | |
|---|---|---|---|---|---|---|---|---|
| **STMN1** | 0.41 | 0.37 | 0.11 | -0.30 | -0.40 | -0.46 | 2.9E-03 | -0.68 |
| **BRF1** | -0.55 | -0.65 | -0.23 | 0.36 | 0.34 | 0.36 | 2.9E-03 | 0.83 |
| **GMPPA** | -0.13 | -0.22 | -0.17 | 0.13 | 0.24 | 0.10 | 3.0E-03 | 0.33 |
| **SYNRG** | -0.14 | -0.15 | -0.12 | 0.19 | 0.13 | 0.06 | 3.0E-03 | 0.26 |
| **DSTN** | 0.14 | 0.11 | 0.04 | -0.09 | -0.09 | -0.13 | 3.0E-03 | -0.20 |
| **C1D** | 0.14 | 0.17 | 0.18 | -0.26 | -0.08 | -0.21 | 3.0E-03 | -0.35 |
| **COP1** | 0.08 | 0.12 | 0.11 | -0.15 | -0.05 | -0.13 | 3.0E-03 | -0.21 |
| **RCN2** | -0.16 | -0.07 | -0.19 | 0.15 | 0.14 | 0.09 | 3.0E-03 | 0.27 |
| **NME2** | -0.09 | -0.15 | -0.19 | 0.09 | 0.11 | 0.19 | 3.0E-03 | 0.27 |
| **NOMO2** | 0.07 | 0.11 | 0.08 | -0.08 | -0.05 | -0.14 | 3.1E-03 | -0.18 |
| **GBA1** | 0.06 | 0.03 | 0.11 | -0.07 | -0.06 | -0.08 | 3.1E-03 | -0.14 |
| **KIFC1** | -0.07 | -0.09 | -0.06 | 0.04 | 0.06 | 0.11 | 3.1E-03 | 0.14 |
| **ADNP** | 0.07 | 0.05 | 0.06 | -0.07 | -0.10 | -0.03 | 3.1E-03 | -0.13 |
| **EED** | 0.10 | 0.04 | 0.09 | -0.10 | -0.04 | -0.10 | 3.1E-03 | -0.16 |
| **ATP6V0A1** | -0.03 | -0.08 | -0.10 | 0.07 | 0.06 | 0.07 | 3.1E-03 | 0.14 |
| **FAM234A** | 0.09 | 0.28 | 0.24 | -0.29 | -0.16 | -0.26 | 3.1E-03 | -0.44 |
| **CDC42** | 0.08 | 0.21 | 0.24 | -0.25 | -0.14 | -0.22 | 3.1E-03 | -0.38 |
| **AIFM1** | 0.09 | 0.18 | 0.07 | -0.12 | -0.10 | -0.15 | 3.1E-03 | -0.23 |
| **BDH1** | 0.09 | 0.38 | 0.29 | -0.35 | -0.27 | -0.32 | 3.1E-03 | -0.57 |
| **SRP68** | -0.11 | -0.18 | -0.06 | 0.12 | 0.12 | 0.08 | 3.1E-03 | 0.23 |
| **RBM15B** | -0.05 | -0.07 | -0.03 | 0.07 | 0.04 | 0.04 | 3.1E-03 | 0.10 |
| **MAD1L1** | -0.14 | -0.14 | -0.06 | 0.10 | 0.07 | 0.15 | 3.1E-03 | 0.22 |
| **EEPD1** | 0.08 | 0.31 | 0.24 | -0.26 | -0.29 | -0.20 | 3.2E-03 | -0.46 |
| **GART** | -0.10 | -0.15 | -0.10 | 0.07 | 0.08 | 0.17 | 3.2E-03 | 0.22 |
| **RFC1** | -0.06 | -0.04 | -0.08 | 0.06 | 0.08 | 0.03 | 3.2E-03 | 0.12 |
| **EXOSC8** | 0.13 | 0.17 | 0.08 | -0.13 | -0.09 | -0.20 | 3.2E-03 | -0.26 |
| **CREB3L4** | 0.07 | 0.07 | 0.06 | -0.06 | -0.04 | -0.11 | 3.2E-03 | -0.13 |
| **SNX32** | 0.08 | 0.22 | 0.12 | -0.13 | -0.20 | -0.14 | 3.2E-03 | -0.30 |
| **COG1** | -0.11 | -0.20 | -0.17 | 0.19 | 0.06 | 0.17 | 3.3E-03 | 0.30 |
| **EPS15L1** | -0.21 | -0.08 | -0.11 | 0.14 | 0.10 | 0.13 | 3.3E-03 | 0.26 |
| **TK1** | -0.09 | -0.16 | -0.23 | 0.13 | 0.20 | 0.10 | 3.3E-03 | 0.30 |
| **SLC16A2** | 0.10 | 0.17 | 0.31 | -0.19 | -0.21 | -0.28 | 3.3E-03 | -0.42 |
| **YARS1** | -0.09 | -0.21 | -0.26 | 0.16 | 0.21 | 0.12 | 3.3E-03 | 0.35 |
| **RNF168** | 0.05 | 0.17 | 0.16 | -0.17 | -0.13 | -0.11 | 3.3E-03 | -0.26 |
| **PSMD5** | -0.14 | -0.16 | -0.11 | 0.07 | 0.10 | 0.20 | 3.3E-03 | 0.26 |
| **PPP1R7** | -0.23 | -0.15 | -0.08 | 0.16 | 0.12 | 0.13 | 3.4E-03 | 0.29 |
| **TBCD** | -0.06 | -0.16 | -0.16 | 0.08 | 0.16 | 0.11 | 3.4E-03 | 0.25 |
| **GTF2I** | -0.06 | -0.04 | -0.07 | 0.08 | 0.03 | 0.05 | 3.4E-03 | 0.11 |
| **FAM210B** | 0.09 | 0.31 | 0.20 | -0.27 | -0.18 | -0.25 | 3.4E-03 | -0.43 |
| **TCIRG1** | 0.06 | 0.07 | 0.06 | -0.03 | -0.10 | -0.08 | 3.4E-03 | -0.13 |
| **SDE2** | 0.08 | 0.11 | 0.12 | -0.07 | -0.17 | -0.09 | 3.4E-03 | -0.22 |
| **HDGFL2** | 0.15 | 0.09 | 0.14 | -0.18 | -0.06 | -0.17 | 3.4E-03 | -0.26 |
| **FAM111B** | 0.12 | 0.05 | 0.07 | -0.06 | -0.08 | -0.12 | 3.4E-03 | -0.17 |
| **GTF3C3** | -0.07 | -0.05 | -0.13 | 0.10 | 0.06 | 0.08 | 3.4E-03 | 0.16 |
| **RIC8A** | -0.23 | -0.31 | -0.19 | 0.12 | 0.17 | 0.32 | 3.5E-03 | 0.44 |
| **C1orf167** | 0.15 | 0.14 | 0.05 | -0.12 | -0.08 | -0.15 | 3.5E-03 | -0.23 |
| **ATP5PF** | 0.07 | 0.40 | 0.35 | -0.37 | -0.34 | -0.34 | 3.5E-03 | -0.63 |
| **UBR5** | -0.26 | -0.25 | -0.08 | 0.17 | 0.16 | 0.19 | 3.5E-03 | 0.37 |

| | | | | | | | | |
|---|---|---|---|---|---|---|---|---|
| **MMS19** | -0.24 | -0.26 | -0.14 | 0.10 | 0.17 | 0.28 | 3.5E-03 | 0.40 |
| **H1-3** | -0.09 | -0.13 | -0.10 | 0.12 | 0.04 | 0.14 | 3.6E-03 | 0.21 |
| **SART3** | -0.10 | -0.15 | -0.14 | 0.07 | 0.19 | 0.09 | 3.6E-03 | 0.25 |
| **RNF213** | -0.12 | -0.14 | -0.09 | 0.10 | 0.17 | 0.05 | 3.6E-03 | 0.22 |
| **ZFP36L2** | 0.12 | 0.08 | 0.15 | -0.20 | -0.11 | -0.08 | 3.6E-03 | -0.25 |
| **TRAPPC8** | -0.12 | -0.15 | -0.28 | 0.17 | 0.20 | 0.12 | 3.6E-03 | 0.35 |
| **TIMM44** | -0.10 | -0.17 | -0.12 | 0.17 | 0.05 | 0.12 | 3.6E-03 | 0.24 |
| **PET100** | 0.21 | 0.14 | 0.18 | -0.30 | -0.10 | -0.22 | 3.6E-03 | -0.38 |
| **ITIH2** | -0.07 | -0.20 | -0.13 | 0.10 | 0.16 | 0.11 | 3.6E-03 | 0.26 |
| **RBM14** | -0.12 | -0.13 | -0.25 | 0.21 | 0.10 | 0.14 | 3.7E-03 | 0.32 |
| **SHROOM2** | -0.28 | -0.23 | -0.09 | 0.16 | 0.21 | 0.15 | 3.7E-03 | 0.37 |
| **YTHDF1** | -0.14 | -0.11 | -0.08 | 0.11 | 0.05 | 0.15 | 3.7E-03 | 0.21 |
| **FOXP1** | -0.30 | -0.33 | -0.23 | 0.12 | 0.19 | 0.39 | 3.7E-03 | 0.52 |
| **TXNL1** | -0.14 | -0.16 | -0.11 | 0.10 | 0.20 | 0.07 | 3.7E-03 | 0.26 |
| **GDI1** | 0.10 | 0.25 | 0.13 | -0.25 | -0.14 | -0.16 | 3.7E-03 | -0.35 |
| **EXOSC10** | 0.09 | 0.04 | 0.07 | -0.11 | -0.06 | -0.05 | 3.7E-03 | -0.14 |
| **ZFYVE21** | 0.13 | 0.20 | 0.26 | -0.33 | -0.13 | -0.22 | 3.8E-03 | -0.43 |
| **ATP5PD** | 0.08 | 0.25 | 0.15 | -0.24 | -0.15 | -0.16 | 3.8E-03 | -0.35 |
| **PANK4** | -0.24 | -0.22 | -0.28 | 0.28 | 0.06 | 0.28 | 3.8E-03 | 0.45 |
| **MMP24OS** | 0.20 | 0.23 | 0.08 | -0.19 | -0.27 | -0.13 | 3.8E-03 | -0.37 |
| **MYH10** | 0.04 | 0.04 | 0.09 | -0.06 | -0.07 | -0.05 | 3.8E-03 | -0.12 |
| **PPFIBP1** | -0.23 | -0.08 | -0.24 | 0.11 | 0.18 | 0.20 | 3.8E-03 | 0.34 |
| **ACTR5** | -0.05 | -0.15 | -0.17 | 0.11 | 0.11 | 0.12 | 3.8E-03 | 0.24 |
| **SPOUT1** | 0.14 | 0.11 | 0.09 | -0.11 | -0.07 | -0.19 | 3.8E-03 | -0.24 |
| **RDH11** | -0.10 | -0.10 | -0.19 | 0.13 | 0.06 | 0.16 | 3.8E-03 | 0.25 |
| **POLR3E** | -0.07 | -0.10 | -0.19 | 0.12 | 0.08 | 0.12 | 3.9E-03 | 0.23 |
| **SNRPB2** | 0.10 | 0.05 | 0.07 | -0.11 | -0.04 | -0.09 | 3.9E-03 | -0.16 |
| **MYBL2** | -0.22 | -0.14 | -0.20 | 0.06 | 0.19 | 0.23 | 3.9E-03 | 0.35 |
| **CYB5R1** | 0.03 | 0.08 | 0.12 | -0.09 | -0.06 | -0.09 | 3.9E-03 | -0.16 |
| **FLII** | 0.09 | 0.11 | 0.09 | -0.14 | -0.04 | -0.13 | 3.9E-03 | -0.20 |
| **SDHA** | 0.11 | 0.42 | 0.28 | -0.28 | -0.31 | -0.43 | 3.9E-03 | -0.61 |
| **PXDNL** | -0.20 | -0.22 | -0.45 | 0.31 | 0.22 | 0.19 | 3.9E-03 | 0.53 |
| **RAB3GAP2** | -0.08 | -0.15 | -0.12 | 0.07 | 0.17 | 0.08 | 3.9E-03 | 0.22 |
| **PCYT2** | -0.25 | -0.18 | -0.17 | 0.07 | 0.27 | 0.19 | 3.9E-03 | 0.38 |
| **ATAD2** | 0.06 | 0.06 | 0.05 | -0.07 | -0.09 | -0.02 | 4.0E-03 | -0.12 |
| **RIGI** | -0.17 | -0.30 | -0.19 | 0.24 | 0.24 | 0.08 | 4.0E-03 | 0.40 |
| **LRBA** | -0.04 | -0.15 | -0.12 | 0.07 | 0.11 | 0.11 | 4.0E-03 | 0.20 |
| **PSMD3** | -0.11 | -0.12 | -0.11 | 0.17 | 0.09 | 0.05 | 4.0E-03 | 0.22 |
| **C9orf78** | 0.05 | 0.15 | 0.18 | -0.11 | -0.13 | -0.18 | 4.0E-03 | -0.27 |
| **NEDD4L** | -0.16 | -0.25 | -0.09 | 0.15 | 0.19 | 0.10 | 4.1E-03 | 0.32 |
| **CPEB4** | -0.17 | -0.17 | -0.10 | 0.20 | 0.05 | 0.14 | 4.1E-03 | 0.28 |
| **ACACA** | -0.10 | -0.23 | -0.11 | 0.09 | 0.16 | 0.15 | 4.1E-03 | 0.28 |
| **CCDC97** | -0.11 | -0.20 | -0.14 | 0.14 | 0.20 | 0.06 | 4.1E-03 | 0.28 |
| **FHIP2A** | -0.31 | -0.26 | -0.13 | 0.12 | 0.19 | 0.29 | 4.1E-03 | 0.43 |
| **KRT18** | -0.27 | -0.12 | -0.23 | 0.25 | 0.09 | 0.19 | 4.2E-03 | 0.38 |
| **KIDINS220** | 0.05 | 0.09 | 0.10 | -0.05 | -0.13 | -0.08 | 4.2E-03 | -0.17 |
| **TMEM204** | 0.23 | 0.25 | 0.10 | -0.17 | -0.17 | -0.34 | 4.2E-03 | -0.42 |
| **CKB** | -0.06 | -0.11 | -0.04 | 0.09 | 0.04 | 0.07 | 4.2E-03 | 0.14 |

| | | | | | | | | |
|---|---|---|---|---|---|---|---|---|
| ATP5F1B | 0.05 | 0.21 | 0.21 | -0.21 | -0.16 | -0.16 | 4.2E-03 | -0.34 |
| TOM1 | -0.12 | -0.03 | -0.10 | 0.07 | 0.07 | 0.10 | 4.2E-03 | 0.17 |
| FAM210A | 0.22 | 0.56 | 0.25 | -0.57 | -0.30 | -0.54 | 4.2E-03 | -0.82 |
| TBL1XR1 | -0.11 | -0.17 | -0.16 | 0.05 | 0.20 | 0.15 | 4.2E-03 | 0.28 |
| SLC4A2 | 0.07 | 0.19 | 0.19 | -0.20 | -0.10 | -0.21 | 4.2E-03 | -0.32 |
| KLHDC7A | 0.05 | 0.11 | 0.18 | -0.11 | -0.10 | -0.16 | 4.3E-03 | -0.24 |
| CLPTM1 | 0.14 | 0.15 | 0.28 | -0.31 | -0.12 | -0.24 | 4.3E-03 | -0.42 |
| RUVBL2 | -0.05 | -0.08 | -0.08 | 0.10 | 0.03 | 0.07 | 4.3E-03 | 0.14 |
| CWF19L2 | 0.11 | 0.07 | 0.09 | -0.14 | -0.04 | -0.10 | 4.3E-03 | -0.18 |
| DNAAF5 | -0.33 | -0.24 | -0.13 | 0.18 | 0.13 | 0.28 | 4.3E-03 | 0.43 |
| LPCAT3 | 0.10 | 0.12 | 0.30 | -0.23 | -0.19 | -0.19 | 4.4E-03 | -0.38 |
| MCM3 | 0.15 | 0.04 | 0.10 | -0.09 | -0.09 | -0.14 | 4.4E-03 | -0.21 |
| SEPTIN5 | -0.35 | -0.11 | -0.27 | 0.27 | 0.15 | 0.20 | 4.4E-03 | 0.45 |
| UGP2 | 0.47 | 0.59 | 0.20 | -0.43 | -0.57 | -0.90 | 4.4E-03 | -1.06 |
| NOTCH2 | -0.27 | -0.19 | -0.12 | 0.08 | 0.22 | 0.21 | 4.4E-03 | 0.36 |
| COG2 | -0.15 | -0.23 | -0.12 | 0.08 | 0.23 | 0.14 | 4.4E-03 | 0.32 |
| SLC29A1 | 0.08 | 0.33 | 0.41 | -0.35 | -0.41 | -0.29 | 4.4E-03 | -0.62 |
| AK1 | -0.08 | -0.21 | -0.11 | 0.11 | 0.10 | 0.16 | 4.4E-03 | 0.26 |
| PAIP2 | 0.10 | 0.14 | 0.27 | -0.24 | -0.13 | -0.22 | 4.4E-03 | -0.37 |
| MCM2 | 0.21 | 0.09 | 0.15 | -0.10 | -0.16 | -0.25 | 4.4E-03 | -0.32 |
| COMMD10 | -0.07 | -0.12 | -0.09 | 0.13 | 0.03 | 0.10 | 4.4E-03 | 0.18 |
| TEDC2 | 0.12 | 0.28 | 0.23 | -0.37 | -0.25 | -0.14 | 4.4E-03 | -0.47 |
| HSF1 | -0.18 | -0.26 | -0.45 | 0.26 | 0.15 | 0.32 | 4.5E-03 | 0.54 |
| DBP | 0.21 | 0.07 | 0.12 | -0.20 | -0.14 | -0.11 | 4.5E-03 | -0.29 |
| RAPGEF6 | -0.24 | -0.47 | -0.20 | 0.21 | 0.32 | 0.19 | 4.5E-03 | 0.54 |
| NBR1 | -0.17 | -0.11 | -0.08 | 0.10 | 0.16 | 0.06 | 4.5E-03 | 0.23 |
| ZC3H10 | -0.09 | -0.06 | -0.03 | 0.06 | 0.04 | 0.08 | 4.5E-03 | 0.12 |
| RABEP1 | -0.05 | -0.09 | -0.08 | 0.08 | 0.03 | 0.10 | 4.5E-03 | 0.14 |
| TOPORS | 0.07 | 0.13 | 0.08 | -0.05 | -0.14 | -0.13 | 4.5E-03 | -0.20 |
| ALG8 | 0.13 | 0.14 | 0.20 | -0.14 | -0.11 | -0.28 | 4.5E-03 | -0.33 |
| ATP1B3 | 0.05 | 0.28 | 0.28 | -0.23 | -0.25 | -0.24 | 4.5E-03 | -0.44 |
| EIF5B | 0.23 | 0.15 | 0.12 | -0.25 | -0.09 | -0.24 | 4.5E-03 | -0.36 |
| LRPAP1 | 0.08 | 0.18 | 0.09 | -0.17 | -0.13 | -0.08 | 4.5E-03 | -0.24 |
| KIN | 0.10 | 0.07 | 0.06 | -0.05 | -0.07 | -0.13 | 4.6E-03 | -0.16 |
| DNAJB2 | -0.40 | -0.20 | -0.22 | 0.17 | 0.16 | 0.35 | 4.6E-03 | 0.50 |
| PNPLA7 | -0.05 | -0.11 | -0.17 | 0.09 | 0.09 | 0.12 | 4.6E-03 | 0.21 |
| STAM2 | -0.24 | -0.26 | -0.16 | 0.18 | 0.08 | 0.30 | 4.6E-03 | 0.40 |
| CXXC1 | -0.08 | -0.06 | -0.12 | 0.05 | 0.13 | 0.06 | 4.6E-03 | 0.17 |
| NFAT5 | 0.06 | 0.12 | 0.21 | -0.15 | -0.17 | -0.11 | 4.6E-03 | -0.27 |
| PLEKHG3 | 0.08 | 0.17 | 0.14 | -0.20 | -0.07 | -0.18 | 4.6E-03 | -0.28 |
| GSTK1 | -0.20 | -0.24 | -0.30 | 0.08 | 0.20 | 0.34 | 4.6E-03 | 0.46 |
| RBBP6 | 0.07 | 0.08 | 0.07 | -0.05 | -0.05 | -0.13 | 4.6E-03 | -0.15 |
| UBE2C | -0.08 | -0.23 | -0.15 | 0.09 | 0.16 | 0.16 | 4.7E-03 | 0.29 |
| TOMM22 | -0.03 | -0.05 | -0.02 | 0.02 | 0.05 | 0.03 | 4.7E-03 | 0.06 |
| ATP5PO | 0.05 | 0.28 | 0.21 | -0.23 | -0.19 | -0.21 | 4.7E-03 | -0.39 |
| LRWD1 | -0.07 | -0.09 | -0.07 | 0.10 | 0.02 | 0.11 | 4.7E-03 | 0.16 |
| STAG2 | -0.05 | -0.10 | -0.12 | 0.09 | 0.04 | 0.12 | 4.7E-03 | 0.17 |
| CHST14 | 0.12 | 0.25 | 0.10 | -0.16 | -0.24 | -0.13 | 4.7E-03 | -0.33 |

| | | | | | | | | |
|---|---|---|---|---|---|---|---|---|
| **EXOC6** | -0.21 | -0.23 | -0.08 | 0.15 | 0.09 | 0.21 | 4.7E-03 | 0.32 |
| **MT2A** | 0.15 | 0.31 | 0.13 | -0.32 | -0.15 | -0.23 | 4.7E-03 | -0.43 |
| **SEL1L** | 0.17 | 0.14 | 0.11 | -0.14 | -0.08 | -0.25 | 4.7E-03 | -0.30 |
| **DDX52** | 0.06 | 0.04 | 0.06 | -0.06 | -0.02 | -0.08 | 4.7E-03 | -0.11 |
| **HOOK1** | -0.30 | -0.28 | -0.09 | 0.24 | 0.16 | 0.17 | 4.7E-03 | 0.41 |
| **NOMO1** | 0.15 | 0.11 | 0.10 | -0.05 | -0.18 | -0.16 | 4.8E-03 | -0.25 |
| **MST1R** | -0.22 | -0.14 | -0.12 | 0.16 | 0.06 | 0.20 | 4.8E-03 | 0.30 |
| **PTPRD** | 0.27 | 0.05 | 0.25 | -0.26 | -0.20 | -0.20 | 4.8E-03 | -0.41 |
| **MOSPD1** | 0.08 | 0.14 | 0.13 | -0.11 | -0.07 | -0.19 | 4.8E-03 | -0.24 |
| **COG4** | -0.15 | -0.26 | -0.09 | 0.15 | 0.19 | 0.11 | 4.8E-03 | 0.31 |
| **ATAD5** | 0.16 | 0.11 | 0.19 | -0.27 | -0.16 | -0.09 | 4.8E-03 | -0.32 |
| **PNP** | 0.25 | 0.26 | 0.15 | -0.39 | -0.12 | -0.29 | 4.9E-03 | -0.49 |
| **NSUN4** | 0.11 | 0.04 | 0.09 | -0.12 | -0.04 | -0.10 | 4.9E-03 | -0.17 |
| **DLST** | 0.10 | 0.19 | 0.11 | -0.08 | -0.13 | -0.22 | 4.9E-03 | -0.28 |
| **MAPK1** | 0.09 | 0.10 | 0.06 | -0.11 | -0.03 | -0.11 | 4.9E-03 | -0.17 |
| **PRPF19** | 0.07 | 0.03 | 0.08 | -0.08 | -0.04 | -0.08 | 4.9E-03 | -0.12 |
| **KHSRP** | 0.11 | 0.08 | 0.08 | -0.15 | -0.10 | -0.04 | 4.9E-03 | -0.19 |
| **SGPL1** | -0.12 | -0.20 | -0.09 | 0.19 | 0.08 | 0.10 | 5.0E-03 | 0.26 |
| **VIPAS39** | -0.03 | -0.12 | -0.07 | 0.07 | 0.07 | 0.07 | 5.0E-03 | 0.14 |
| **CDK1** | -0.10 | -0.09 | -0.13 | 0.14 | 0.03 | 0.12 | 5.0E-03 | 0.21 |
| **EPPK1** | -0.07 | -0.08 | -0.04 | 0.06 | 0.09 | 0.03 | 5.0E-03 | 0.13 |
| **MGAT4B** | 0.08 | 0.09 | 0.18 | -0.12 | -0.08 | -0.18 | 5.0E-03 | -0.24 |
| **ACSF3** | -0.05 | -0.12 | -0.16 | 0.06 | 0.11 | 0.12 | 5.0E-03 | 0.21 |
| **IPO4** | -0.27 | -0.44 | -0.21 | 0.20 | 0.16 | 0.39 | 5.0E-03 | 0.55 |
| **LRATD2** | 0.02 | 0.06 | 0.07 | -0.07 | -0.04 | -0.04 | 5.0E-03 | -0.10 |
| **COG3** | -0.19 | -0.23 | -0.21 | 0.04 | 0.23 | 0.27 | 5.0E-03 | 0.39 |
| **MFSD10** | -0.03 | -0.04 | -0.03 | 0.06 | 0.02 | 0.02 | 5.0E-03 | 0.07 |
| **MRPL4** | 0.10 | 0.15 | 0.10 | -0.18 | -0.05 | -0.16 | 5.0E-03 | -0.24 |
| **RBM7** | 0.07 | 0.06 | 0.06 | -0.04 | -0.11 | -0.05 | 5.0E-03 | -0.13 |
| **OCIAD2** | 0.09 | 0.14 | 0.15 | -0.19 | -0.05 | -0.18 | 5.0E-03 | -0.26 |
| **QRICH1** | -0.26 | -0.11 | -0.15 | 0.14 | 0.10 | 0.22 | 5.0E-03 | 0.33 |
| **EARS2** | -0.06 | -0.15 | -0.22 | 0.12 | 0.16 | 0.11 | 5.0E-03 | 0.27 |
| **TRIR** | 0.10 | 0.11 | 0.26 | -0.20 | -0.13 | -0.20 | 5.1E-03 | -0.34 |
| **RPS27A** | -0.10 | -0.18 | -0.09 | 0.10 | 0.17 | 0.07 | 5.1E-03 | 0.23 |
| **LDAF1** | -0.33 | -0.27 | -0.09 | 0.23 | 0.14 | 0.22 | 5.1E-03 | 0.42 |
| **PNN** | 0.09 | 0.06 | 0.05 | -0.07 | -0.03 | -0.10 | 5.1E-03 | -0.13 |
| **ITGA3** | 0.06 | 0.08 | 0.19 | -0.13 | -0.14 | -0.10 | 5.1E-03 | -0.24 |
| **MARCHF6** | 0.12 | 0.23 | 0.43 | -0.41 | -0.23 | -0.32 | 5.1E-03 | -0.58 |
| **GRIPAP1** | -0.07 | -0.12 | -0.17 | 0.16 | 0.06 | 0.11 | 5.1E-03 | 0.23 |
| **SETD2** | 0.04 | 0.04 | 0.03 | -0.03 | -0.06 | -0.02 | 5.1E-03 | -0.07 |
| **NUBP1** | -0.28 | -0.23 | -0.13 | 0.07 | 0.26 | 0.22 | 5.1E-03 | 0.40 |
| **FA2H** | 0.18 | 0.07 | 0.13 | -0.16 | -0.07 | -0.20 | 5.1E-03 | -0.27 |
| **ILK** | -0.13 | -0.22 | -0.17 | 0.05 | 0.19 | 0.21 | 5.2E-03 | 0.32 |
| **VPS28** | -0.10 | -0.04 | -0.04 | 0.05 | 0.05 | 0.07 | 5.2E-03 | 0.11 |
| **HSPA9** | -0.07 | -0.09 | -0.18 | 0.13 | 0.06 | 0.12 | 5.2E-03 | 0.22 |
| **TMEM186** | -0.10 | -0.14 | -0.14 | 0.17 | 0.04 | 0.13 | 5.2E-03 | 0.24 |
| **H1-2** | -0.11 | -0.16 | -0.24 | 0.21 | 0.07 | 0.18 | 5.2E-03 | 0.32 |
| **SLC25A10** | 0.04 | 0.16 | 0.09 | -0.12 | -0.10 | -0.10 | 5.2E-03 | -0.20 |

| | | | | | | | | |
|---|---|---|---|---|---|---|---|---|
| ZBTB7C | -0.14 | -0.24 | -0.11 | 0.11 | 0.10 | 0.22 | 5.2E-03 | 0.31 |
| MRPL12 | 0.16 | 0.24 | 0.06 | -0.14 | -0.15 | -0.22 | 5.2E-03 | -0.32 |
| LYPD3 | 0.03 | 0.14 | 0.12 | -0.13 | -0.12 | -0.07 | 5.3E-03 | -0.20 |
| AIMP1 | -0.10 | -0.12 | -0.07 | 0.03 | 0.13 | 0.12 | 5.3E-03 | 0.19 |
| WDR55 | -0.10 | -0.15 | -0.07 | 0.13 | 0.04 | 0.14 | 5.3E-03 | 0.21 |
| SHARPIN | -0.28 | -0.61 | -0.30 | 0.28 | 0.20 | 0.44 | 5.3E-03 | 0.70 |
| ATP5F1E | 0.12 | 0.27 | 0.19 | -0.30 | -0.10 | -0.29 | 5.3E-03 | -0.43 |
| GIGYF2 | -0.29 | -0.28 | -0.10 | 0.21 | 0.11 | 0.25 | 5.3E-03 | 0.41 |
| LUC7L3 | 0.32 | 0.44 | 0.26 | -0.38 | -0.28 | -0.74 | 5.3E-03 | -0.81 |
| ATP5PB | 0.08 | 0.25 | 0.22 | -0.31 | -0.14 | -0.20 | 5.3E-03 | -0.40 |
| MRPS18B | 0.12 | 0.05 | 0.16 | -0.08 | -0.10 | -0.18 | 5.4E-03 | -0.23 |
| GRHL1 | -0.10 | -0.13 | -0.05 | 0.05 | 0.08 | 0.13 | 5.4E-03 | 0.18 |
| ALDH5A1 | -0.02 | -0.06 | -0.09 | 0.07 | 0.04 | 0.05 | 5.4E-03 | 0.11 |
| BAD | 0.13 | 0.49 | 0.24 | -0.27 | -0.44 | -0.41 | 5.4E-03 | -0.66 |
| CLIC3 | -0.18 | -0.26 | -0.26 | 0.11 | 0.34 | 0.13 | 5.4E-03 | 0.43 |
| CERT1 | -0.08 | -0.05 | -0.05 | 0.04 | 0.10 | 0.04 | 5.4E-03 | 0.12 |
| MCM5 | 0.22 | 0.08 | 0.17 | -0.19 | -0.10 | -0.25 | 5.4E-03 | -0.34 |
| GALNT11 | 0.04 | 0.15 | 0.17 | -0.12 | -0.10 | -0.17 | 5.4E-03 | -0.25 |
| LSS | -0.11 | -0.09 | -0.12 | 0.03 | 0.10 | 0.16 | 5.5E-03 | 0.21 |
| SAR1B | -0.22 | -0.44 | -0.67 | 0.44 | 0.28 | 0.26 | 5.6E-03 | 0.77 |
| CD81 | 0.07 | 0.06 | 0.11 | -0.04 | -0.08 | -0.13 | 5.6E-03 | -0.16 |
| HUWE1 | -0.11 | -0.22 | -0.09 | 0.13 | 0.17 | 0.08 | 5.6E-03 | 0.26 |
| KRT19 | -0.22 | -0.07 | -0.19 | 0.17 | 0.08 | 0.17 | 5.6E-03 | 0.30 |
| ARHGEF11 | -0.25 | -0.08 | -0.23 | 0.09 | 0.20 | 0.20 | 5.6E-03 | 0.35 |
| STARD3 | 0.08 | 0.09 | 0.15 | -0.05 | -0.12 | -0.18 | 5.6E-03 | -0.22 |
| RHOD | 0.06 | 0.21 | 0.12 | -0.16 | -0.09 | -0.17 | 5.6E-03 | -0.27 |
| SNRPA | 0.19 | 0.26 | 0.25 | -0.22 | -0.17 | -0.47 | 5.6E-03 | -0.52 |
| RAPH1 | -0.16 | -0.10 | -0.11 | 0.10 | 0.05 | 0.18 | 5.6E-03 | 0.23 |
| KIF4A | -0.02 | -0.05 | -0.09 | 0.04 | 0.05 | 0.06 | 5.7E-03 | 0.10 |
| AURKA | -0.11 | -0.02 | -0.10 | 0.07 | 0.07 | 0.08 | 5.7E-03 | 0.15 |
| RRM2 | -0.23 | -0.13 | -0.18 | 0.05 | 0.24 | 0.18 | 5.7E-03 | 0.34 |
| OAS2 | -0.10 | -0.05 | -0.03 | 0.06 | 0.06 | 0.05 | 5.7E-03 | 0.12 |
| TPM3 | 0.06 | 0.07 | 0.19 | -0.09 | -0.12 | -0.13 | 5.7E-03 | -0.22 |
| COG6 | -0.28 | -0.33 | -0.11 | 0.28 | 0.11 | 0.21 | 5.8E-03 | 0.44 |
| CKAP2 | -0.17 | -0.09 | -0.06 | 0.08 | 0.08 | 0.14 | 5.8E-03 | 0.21 |
| KIF14 | 0.06 | 0.04 | 0.09 | -0.05 | -0.10 | -0.04 | 5.8E-03 | -0.13 |
| DHX8 | 0.07 | 0.07 | 0.06 | -0.07 | -0.02 | -0.11 | 5.8E-03 | -0.14 |
| HSPBP1 | -0.01 | -0.04 | -0.04 | 0.02 | 0.02 | 0.04 | 5.8E-03 | 0.06 |
| GRHPR | -0.12 | -0.10 | -0.23 | 0.07 | 0.13 | 0.20 | 5.9E-03 | 0.29 |
| AGPAT5 | -0.09 | -0.05 | -0.11 | 0.12 | 0.03 | 0.08 | 5.9E-03 | 0.16 |
| GNB2 | 0.06 | 0.14 | 0.11 | -0.18 | -0.06 | -0.10 | 5.9E-03 | -0.22 |
| ATE1 | -0.15 | -0.21 | -0.24 | 0.10 | 0.11 | 0.30 | 5.9E-03 | 0.37 |
| RAB23 | -0.07 | -0.04 | -0.03 | 0.03 | 0.07 | 0.04 | 5.9E-03 | 0.09 |
| NUP58 | 0.16 | 0.15 | 0.18 | -0.16 | -0.09 | -0.31 | 5.9E-03 | -0.35 |
| EXOSC9 | 0.09 | 0.10 | 0.04 | -0.06 | -0.05 | -0.12 | 5.9E-03 | -0.15 |
| PSMA7 | -0.10 | -0.15 | -0.21 | 0.18 | 0.05 | 0.19 | 5.9E-03 | 0.30 |
| ANXA7 | -0.21 | -0.29 | -0.16 | 0.17 | 0.08 | 0.31 | 6.0E-03 | 0.41 |
| DDOST | 0.04 | 0.07 | 0.13 | -0.12 | -0.06 | -0.08 | 6.0E-03 | -0.17 |

| | | | | | | | | |
|---|---|---|---|---|---|---|---|---|
| HM13 | 0.10 | 0.16 | 0.27 | -0.18 | -0.13 | -0.30 | 6.0E-03 | -0.37 |
| ABHD16A | 0.08 | 0.10 | 0.10 | -0.14 | -0.03 | -0.14 | 6.0E-03 | -0.20 |
| TBC1D22A | -0.35 | -0.31 | -0.11 | 0.14 | 0.20 | 0.31 | 6.0E-03 | 0.47 |
| MOV10 | -0.08 | -0.04 | -0.05 | 0.07 | 0.02 | 0.07 | 6.0E-03 | 0.11 |
| STX5 | -0.08 | -0.08 | -0.10 | 0.14 | 0.04 | 0.06 | 6.0E-03 | 0.16 |
| KPNA6 | -0.09 | -0.07 | -0.09 | 0.08 | 0.13 | 0.03 | 6.1E-03 | 0.16 |
| CDS2 | 0.10 | 0.10 | 0.15 | -0.22 | -0.08 | -0.10 | 6.1E-03 | -0.25 |
| TP53RK | -0.42 | -0.33 | -0.11 | 0.26 | 0.16 | 0.29 | 6.1E-03 | 0.52 |
| LPP | 0.09 | 0.10 | 0.20 | -0.10 | -0.23 | -0.12 | 6.1E-03 | -0.28 |
| CSDE1 | -0.10 | -0.10 | -0.13 | 0.07 | 0.06 | 0.18 | 6.1E-03 | 0.21 |
| CNPY3 | 0.18 | 0.20 | 0.07 | -0.12 | -0.13 | -0.26 | 6.2E-03 | -0.32 |
| ARHGAP1 | -0.16 | -0.06 | -0.17 | 0.06 | 0.15 | 0.16 | 6.2E-03 | 0.25 |
| STAT2 | -0.11 | -0.21 | -0.14 | 0.05 | 0.17 | 0.19 | 6.2E-03 | 0.29 |
| RHOG | 0.08 | 0.20 | 0.13 | -0.09 | -0.13 | -0.23 | 6.2E-03 | -0.29 |
| PSMB2 | -0.21 | -0.25 | -0.06 | 0.13 | 0.14 | 0.18 | 6.2E-03 | 0.32 |
| TCOF1 | 0.19 | 0.08 | 0.11 | -0.22 | -0.08 | -0.13 | 6.2E-03 | -0.27 |
| KLRG2 | -0.14 | -0.07 | -0.14 | 0.11 | 0.04 | 0.16 | 6.2E-03 | 0.22 |
| GRWD1 | -0.02 | -0.02 | -0.05 | 0.04 | 0.04 | 0.02 | 6.2E-03 | 0.06 |
| DENND4C | -0.32 | -0.23 | -0.16 | 0.11 | 0.33 | 0.15 | 6.2E-03 | 0.43 |
| LETM1 | 0.06 | 0.34 | 0.22 | -0.31 | -0.19 | -0.25 | 6.3E-03 | -0.45 |
| GOSR2 | 0.19 | 0.18 | 0.06 | -0.21 | -0.09 | -0.18 | 6.3E-03 | -0.30 |
| SIGIRR | -0.15 | -0.22 | -0.10 | 0.08 | 0.11 | 0.22 | 6.3E-03 | 0.29 |
| SRPK1 | 0.03 | 0.07 | 0.12 | -0.07 | -0.07 | -0.10 | 6.3E-03 | -0.15 |
| RPS6KB2 | -0.19 | -0.18 | -0.09 | 0.04 | 0.19 | 0.17 | 6.3E-03 | 0.29 |
| TIPIN | 0.23 | 0.14 | 0.04 | -0.16 | -0.15 | -0.15 | 6.3E-03 | -0.29 |
| AARS2 | -0.22 | -0.15 | -0.12 | 0.16 | 0.05 | 0.22 | 6.3E-03 | 0.31 |
| PTDSS1 | 0.03 | 0.11 | 0.08 | -0.10 | -0.09 | -0.05 | 6.3E-03 | -0.15 |
| WRN | 0.11 | 0.17 | 0.41 | -0.32 | -0.22 | -0.32 | 6.3E-03 | -0.52 |
| DHX16 | 0.09 | 0.02 | 0.06 | -0.04 | -0.05 | -0.09 | 6.3E-03 | -0.12 |
| BCL7B | 0.19 | 0.09 | 0.06 | -0.15 | -0.07 | -0.15 | 6.3E-03 | -0.24 |
| NBEAL2 | -0.17 | -0.21 | -0.05 | 0.13 | 0.10 | 0.16 | 6.4E-03 | 0.27 |
| SDHB | 0.08 | 0.39 | 0.27 | -0.23 | -0.28 | -0.40 | 6.4E-03 | -0.55 |
| FANCI | -0.04 | -0.18 | -0.17 | 0.10 | 0.14 | 0.12 | 6.4E-03 | 0.25 |
| RABGEF1 | -0.27 | -0.21 | -0.33 | 0.29 | 0.32 | 0.05 | 6.4E-03 | 0.49 |
| GATB | -0.12 | -0.15 | -0.31 | 0.15 | 0.11 | 0.24 | 6.4E-03 | 0.36 |
| SH3GLB1 | -0.15 | -0.16 | -0.23 | 0.26 | 0.15 | 0.06 | 6.4E-03 | 0.33 |
| ERG28 | 0.15 | 0.16 | 0.07 | -0.18 | -0.06 | -0.18 | 6.4E-03 | -0.27 |
| ZNF281 | -0.06 | -0.13 | -0.09 | 0.11 | 0.03 | 0.12 | 6.4E-03 | 0.18 |
| LUC7L2 | 0.19 | 0.23 | 0.09 | -0.13 | -0.14 | -0.32 | 6.5E-03 | -0.37 |
| APTX | -0.11 | -0.06 | -0.10 | 0.08 | 0.04 | 0.13 | 6.5E-03 | 0.17 |
| PSMC1 | -0.16 | -0.17 | -0.06 | 0.09 | 0.18 | 0.08 | 6.5E-03 | 0.25 |
| GPKOW | 0.10 | 0.10 | 0.05 | -0.07 | -0.05 | -0.14 | 6.6E-03 | -0.17 |
| PSMB5 | -0.17 | -0.19 | -0.16 | 0.12 | 0.07 | 0.27 | 6.6E-03 | 0.33 |
| ZMYM4 | 0.07 | 0.07 | 0.05 | -0.02 | -0.09 | -0.10 | 6.6E-03 | -0.13 |
| ZNF276 | -0.06 | -0.04 | -0.12 | 0.09 | 0.05 | 0.06 | 6.7E-03 | 0.14 |
| NR2F1 | 0.07 | 0.11 | 0.13 | -0.05 | -0.10 | -0.18 | 6.7E-03 | -0.21 |
| SERINC3 | 0.04 | 0.14 | 0.12 | -0.15 | -0.11 | -0.06 | 6.7E-03 | -0.21 |
| OAS1 | 0.10 | 0.22 | 0.07 | -0.10 | -0.14 | -0.20 | 6.7E-03 | -0.28 |

| | | | | | | | | |
|---|---|---|---|---|---|---|---|---|
| **DBI** | 0.32 | 0.34 | 0.03 | -0.27 | -0.28 | -0.31 | 6.7E-03 | -0.52 |
| **DCTN1** | -0.13 | -0.13 | -0.02 | 0.09 | 0.08 | 0.09 | 6.7E-03 | 0.18 |
| **ATP2A2** | 0.05 | 0.08 | 0.14 | -0.10 | -0.05 | -0.13 | 6.7E-03 | -0.18 |
| **SDHC** | 0.05 | 0.35 | 0.38 | -0.37 | -0.40 | -0.24 | 6.7E-03 | -0.59 |
| **SLC26A11** | 0.04 | 0.13 | 0.15 | -0.08 | -0.18 | -0.09 | 6.7E-03 | -0.23 |
| **PINX1** | 0.09 | 0.08 | 0.10 | -0.17 | -0.06 | -0.06 | 6.7E-03 | -0.19 |
| **ELOC** | -0.17 | -0.23 | -0.25 | 0.10 | 0.12 | 0.33 | 6.8E-03 | 0.40 |
| **ERC1** | -0.22 | -0.06 | -0.13 | 0.15 | 0.13 | 0.08 | 6.8E-03 | 0.26 |
| **ZNF830** | 0.13 | 0.09 | 0.08 | -0.09 | -0.05 | -0.18 | 6.8E-03 | -0.21 |
| **ACTA2** | -0.24 | -0.39 | -0.26 | 0.30 | 0.06 | 0.36 | 6.8E-03 | 0.53 |
| **GFUS** | -0.09 | -0.19 | -0.27 | 0.10 | 0.13 | 0.24 | 6.8E-03 | 0.34 |
| **SLC25A32** | 0.13 | 0.26 | 0.08 | -0.18 | -0.11 | -0.25 | 6.9E-03 | -0.34 |
| **CDK10** | 0.26 | 0.25 | 0.32 | -0.24 | -0.22 | -0.59 | 6.9E-03 | -0.63 |
| **MECP2** | -0.07 | -0.09 | -0.16 | 0.06 | 0.08 | 0.15 | 6.9E-03 | 0.20 |
| **RNF40** | -0.05 | -0.04 | -0.13 | 0.05 | 0.08 | 0.08 | 6.9E-03 | 0.15 |
| **ARL6IP4** | 0.26 | 0.14 | 0.18 | -0.18 | -0.12 | -0.37 | 6.9E-03 | -0.41 |
| **RFC2** | -0.03 | -0.04 | -0.03 | 0.05 | 0.03 | 0.01 | 6.9E-03 | 0.07 |
| **WASHC5** | -0.17 | -0.15 | -0.03 | 0.08 | 0.13 | 0.11 | 7.0E-03 | 0.22 |
| **ORC4** | -0.06 | -0.08 | -0.09 | 0.02 | 0.11 | 0.10 | 7.0E-03 | 0.15 |
| **ICA1** | -0.05 | -0.11 | -0.07 | 0.10 | 0.09 | 0.03 | 7.0E-03 | 0.15 |
| **SLC25A21** | 0.10 | 0.37 | 0.16 | -0.30 | -0.16 | -0.30 | 7.0E-03 | -0.47 |
| **GAA** | -0.05 | -0.08 | -0.06 | 0.05 | 0.11 | 0.03 | 7.0E-03 | 0.13 |
| **ALDH4A1** | -0.05 | -0.15 | -0.23 | 0.12 | 0.14 | 0.12 | 7.0E-03 | 0.27 |
| **GPATCH8** | 0.08 | 0.09 | 0.05 | -0.05 | -0.05 | -0.13 | 7.0E-03 | -0.15 |
| **VEZF1** | -0.29 | -0.31 | -0.30 | 0.27 | 0.04 | 0.40 | 7.1E-03 | 0.54 |
| **LDLRAP1** | -0.25 | -0.38 | -0.19 | 0.16 | 0.13 | 0.38 | 7.1E-03 | 0.49 |
| **TMX2** | 0.12 | 0.17 | 0.20 | -0.14 | -0.10 | -0.31 | 7.1E-03 | -0.35 |
| **TIA1** | 0.07 | 0.08 | 0.02 | -0.04 | -0.09 | -0.06 | 7.1E-03 | -0.12 |
| **KTI12** | -0.18 | -0.06 | -0.12 | 0.05 | 0.12 | 0.15 | 7.1E-03 | 0.23 |
| **CHUK** | -0.20 | -0.13 | -0.17 | 0.16 | 0.24 | 0.04 | 7.1E-03 | 0.31 |
| **DNAJC9** | 0.13 | 0.15 | 0.10 | -0.11 | -0.07 | -0.24 | 7.2E-03 | -0.27 |
| **POLDIP3** | 0.05 | 0.07 | 0.03 | -0.09 | -0.04 | -0.03 | 7.2E-03 | -0.10 |
| **IPO7** | -0.03 | -0.09 | -0.10 | 0.08 | 0.10 | 0.03 | 7.2E-03 | 0.14 |
| **MRPL17** | 0.09 | 0.07 | 0.05 | -0.03 | -0.09 | -0.12 | 7.2E-03 | -0.15 |
| **MCFD2** | 0.22 | 0.33 | 0.12 | -0.23 | -0.16 | -0.41 | 7.2E-03 | -0.49 |
| **TLE3** | -0.10 | -0.15 | -0.22 | 0.05 | 0.16 | 0.20 | 7.2E-03 | 0.29 |
| **TELO2** | -0.11 | -0.23 | -0.07 | 0.13 | 0.14 | 0.10 | 7.2E-03 | 0.26 |
| **DDX47** | 0.10 | 0.09 | 0.04 | -0.04 | -0.08 | -0.12 | 7.2E-03 | -0.15 |
| **HLA-B** | 0.04 | 0.15 | 0.21 | -0.19 | -0.16 | -0.10 | 7.3E-03 | -0.28 |
| **ATL3** | -0.05 | -0.18 | -0.22 | 0.13 | 0.10 | 0.17 | 7.3E-03 | 0.28 |
| **MAX** | -0.16 | -0.15 | -0.10 | 0.18 | 0.03 | 0.17 | 7.3E-03 | 0.27 |
| **STIM1** | 0.15 | 0.10 | 0.11 | -0.05 | -0.13 | -0.22 | 7.3E-03 | -0.25 |
| **ALG12** | -0.36 | -0.37 | -0.39 | 0.35 | 0.03 | 0.47 | 7.4E-03 | 0.65 |
| **COPB1** | -0.19 | -0.24 | -0.10 | 0.25 | 0.08 | 0.14 | 7.4E-03 | 0.33 |
| **DHX37** | 0.11 | 0.11 | 0.08 | -0.08 | -0.06 | -0.19 | 7.4E-03 | -0.21 |
| **CORO1B** | 0.04 | 0.05 | 0.08 | -0.09 | -0.07 | -0.02 | 7.4E-03 | -0.12 |
| **SIKE1** | 0.07 | 0.05 | 0.17 | -0.12 | -0.07 | -0.13 | 7.5E-03 | -0.20 |
| **RAB8B** | 0.17 | 0.28 | 0.27 | -0.32 | -0.11 | -0.46 | 7.5E-03 | -0.54 |

| | | | | | | | | |
|---|---|---|---|---|---|---|---|---|
| **GPANK1** | 0.07 | 0.06 | 0.08 | -0.12 | -0.09 | -0.02 | 7.5E-03 | -0.15 |
| **PPOX** | -0.24 | -0.15 | -0.21 | 0.12 | 0.08 | 0.31 | 7.5E-03 | 0.37 |
| **SPECC1L** | 0.05 | 0.10 | 0.20 | -0.09 | -0.14 | -0.15 | 7.6E-03 | -0.25 |
| **VAPB** | 0.03 | 0.08 | 0.10 | -0.08 | -0.03 | -0.10 | 7.6E-03 | -0.14 |
| **GCC1** | -0.23 | -0.28 | -0.07 | 0.13 | 0.25 | 0.14 | 7.6E-03 | 0.36 |
| **IRF7** | -0.10 | -0.08 | -0.03 | 0.08 | 0.09 | 0.03 | 7.6E-03 | 0.14 |
| **STARD10** | -0.04 | -0.19 | -0.14 | 0.14 | 0.09 | 0.10 | 7.6E-03 | 0.23 |
| **LRRC57** | 0.03 | 0.17 | 0.13 | -0.10 | -0.11 | -0.16 | 7.6E-03 | -0.23 |
| **GALE** | -0.28 | -0.27 | -0.39 | 0.17 | 0.11 | 0.46 | 7.7E-03 | 0.56 |
| **NLN** | -0.08 | -0.15 | -0.11 | 0.03 | 0.16 | 0.13 | 7.7E-03 | 0.22 |
| **KDM4B** | -0.09 | -0.05 | -0.05 | 0.08 | 0.02 | 0.08 | 7.7E-03 | 0.12 |
| **COMT** | -0.26 | -0.35 | -0.16 | 0.24 | 0.07 | 0.33 | 7.8E-03 | 0.47 |
| **JADE3** | 0.11 | 0.20 | 0.15 | -0.14 | -0.09 | -0.29 | 7.8E-03 | -0.32 |
| **MCM6** | 0.31 | 0.12 | 0.19 | -0.18 | -0.17 | -0.40 | 7.8E-03 | -0.46 |
| **FASTKD3** | 0.18 | 0.16 | 0.17 | -0.05 | -0.28 | -0.28 | 7.8E-03 | -0.37 |
| **SSR3** | 0.08 | 0.09 | 0.06 | -0.03 | -0.08 | -0.14 | 7.9E-03 | -0.16 |
| **IFIT3** | -0.24 | -0.38 | -0.15 | 0.13 | 0.15 | 0.34 | 7.9E-03 | 0.46 |
| **SLC35A4** | 0.25 | 0.20 | 0.17 | -0.38 | -0.28 | -0.08 | 7.9E-03 | -0.45 |
| **TEX2** | -0.18 | -0.20 | -0.11 | 0.23 | 0.03 | 0.18 | 7.9E-03 | 0.31 |
| **HMCES** | -0.60 | -0.81 | -0.81 | 0.42 | 0.06 | 0.86 | 7.9E-03 | 1.18 |
| **CLK3** | 0.06 | 0.25 | 0.18 | -0.27 | -0.14 | -0.14 | 7.9E-03 | -0.34 |
| **SMARCAL1** | -0.61 | -0.57 | -0.44 | 0.09 | 0.32 | 0.70 | 7.9E-03 | 0.91 |
| **PTPRK** | -0.15 | -0.09 | -0.08 | 0.16 | 0.06 | 0.07 | 8.0E-03 | 0.20 |
| **GGH** | -0.09 | -0.08 | -0.10 | 0.12 | 0.01 | 0.12 | 8.0E-03 | 0.18 |
| **DHX40** | 0.09 | 0.01 | 0.09 | -0.07 | -0.05 | -0.08 | 8.0E-03 | -0.13 |
| **TMED4** | -0.10 | -0.10 | -0.16 | 0.03 | 0.12 | 0.17 | 8.0E-03 | 0.22 |
| **ZMPSTE24** | 0.21 | 0.24 | 0.17 | -0.30 | -0.07 | -0.37 | 8.1E-03 | -0.45 |
| **SLC50A1** | -0.24 | -0.06 | -0.14 | 0.16 | 0.09 | 0.15 | 8.1E-03 | 0.27 |
| **MICU1** | 0.10 | 0.04 | 0.03 | -0.06 | -0.04 | -0.06 | 8.1E-03 | -0.11 |
| **ZBTB1** | 0.05 | 0.16 | 0.11 | -0.16 | -0.14 | -0.05 | 8.1E-03 | -0.22 |
| **RPL38** | -0.25 | -0.45 | -0.19 | 0.23 | 0.11 | 0.38 | 8.2E-03 | 0.54 |
| **SASH1** | -0.09 | -0.26 | -0.15 | 0.07 | 0.17 | 0.20 | 8.2E-03 | 0.32 |
| **SON** | 0.07 | 0.03 | 0.05 | -0.08 | -0.04 | -0.03 | 8.2E-03 | -0.10 |
| **GEMIN5** | 0.02 | 0.05 | 0.05 | -0.04 | -0.06 | -0.02 | 8.2E-03 | -0.08 |
| **CTSB** | -0.19 | -0.39 | -0.21 | 0.28 | 0.07 | 0.30 | 8.2E-03 | 0.48 |
| **TFAM** | -0.02 | -0.07 | -0.11 | 0.07 | 0.05 | 0.07 | 8.2E-03 | 0.13 |
| **SLC35A3** | 0.12 | 0.16 | 0.18 | -0.10 | -0.12 | -0.30 | 8.2E-03 | -0.33 |
| **MTPAP** | 0.07 | 0.11 | 0.09 | -0.08 | -0.05 | -0.17 | 8.3E-03 | -0.19 |
| **CASC3** | -0.16 | -0.06 | -0.07 | 0.12 | 0.10 | 0.05 | 8.3E-03 | 0.19 |
| **RCOR3** | 0.05 | 0.02 | 0.02 | -0.01 | -0.04 | -0.04 | 8.4E-03 | -0.06 |
| **MRPL53** | 0.15 | 0.18 | 0.05 | -0.14 | -0.08 | -0.21 | 8.4E-03 | -0.27 |
| **USP48** | -0.04 | -0.16 | -0.18 | 0.16 | 0.08 | 0.11 | 8.4E-03 | 0.24 |
| **YY1** | -0.03 | -0.05 | -0.09 | 0.08 | 0.04 | 0.04 | 8.4E-03 | 0.11 |
| **ATP5MG** | 0.08 | 0.35 | 0.26 | -0.35 | -0.15 | -0.35 | 8.5E-03 | -0.51 |
| **GNA11** | 0.03 | 0.13 | 0.11 | -0.07 | -0.08 | -0.15 | 8.5E-03 | -0.19 |
| **TOP3A** | -0.36 | -0.36 | -0.16 | 0.21 | 0.10 | 0.40 | 8.5E-03 | 0.53 |
| **MTFR1** | 0.07 | 0.11 | 0.06 | -0.13 | -0.02 | -0.11 | 8.5E-03 | -0.17 |
| **ERCC6** | -0.11 | -0.02 | -0.07 | 0.06 | 0.06 | 0.07 | 8.6E-03 | 0.13 |

| | | | | | | | | |
|---|---|---|---|---|---|---|---|---|
| RAC1 | 0.03 | 0.18 | 0.15 | -0.18 | -0.09 | -0.13 | 8.6E-03 | -0.25 |
| ZFPL1 | -0.13 | -0.10 | -0.04 | 0.12 | 0.09 | 0.03 | 8.7E-03 | 0.17 |
| IQANK1 | -0.17 | -0.09 | -0.08 | 0.04 | 0.11 | 0.16 | 8.7E-03 | 0.21 |
| MYL6 | 0.08 | 0.03 | 0.09 | -0.04 | -0.06 | -0.11 | 8.7E-03 | -0.14 |
| DIP2B | 0.06 | 0.04 | 0.13 | -0.11 | -0.06 | -0.07 | 8.7E-03 | -0.16 |
| PIR | -0.14 | -0.15 | -0.23 | 0.23 | 0.03 | 0.19 | 8.7E-03 | 0.32 |
| ATP5F1C | 0.07 | 0.23 | 0.23 | -0.33 | -0.16 | -0.14 | 8.8E-03 | -0.39 |
| ZNF275 | 0.48 | 0.04 | 0.39 | -0.51 | -0.42 | -0.29 | 8.8E-03 | -0.71 |
| DDX21 | 0.11 | 0.09 | 0.02 | -0.09 | -0.05 | -0.09 | 8.8E-03 | -0.15 |
| UBR4 | -0.06 | -0.16 | -0.12 | 0.05 | 0.17 | 0.10 | 8.8E-03 | 0.22 |
| PAIP1 | 0.06 | 0.15 | 0.26 | -0.10 | -0.18 | -0.25 | 8.8E-03 | -0.33 |
| LZTS2 | 0.13 | 0.06 | 0.13 | -0.07 | -0.08 | -0.20 | 8.8E-03 | -0.22 |
| NCAPG | -0.09 | -0.13 | -0.14 | 0.02 | 0.16 | 0.15 | 8.8E-03 | 0.23 |
| DNMT3B | 0.20 | 0.11 | 0.05 | -0.17 | -0.08 | -0.15 | 8.8E-03 | -0.26 |
| TRAPPC5 | -0.01 | -0.08 | -0.07 | 0.04 | 0.05 | 0.06 | 8.9E-03 | 0.10 |
| GEMIN4 | -0.02 | -0.14 | -0.10 | 0.10 | 0.07 | 0.07 | 8.9E-03 | 0.17 |
| STX4 | 0.05 | 0.12 | 0.17 | -0.05 | -0.15 | -0.16 | 8.9E-03 | -0.23 |
| MPST | -0.20 | -0.24 | -0.05 | 0.15 | 0.18 | 0.10 | 9.0E-03 | 0.30 |
| RSRC2 | 0.11 | 0.09 | 0.26 | -0.26 | -0.10 | -0.17 | 9.0E-03 | -0.33 |
| AP1G1 | -0.09 | -0.07 | -0.02 | 0.08 | 0.03 | 0.06 | 9.0E-03 | 0.12 |
| IFT140 | -0.17 | -0.20 | -0.25 | 0.20 | 0.03 | 0.30 | 9.0E-03 | 0.39 |
| PCNT | -0.10 | -0.05 | -0.04 | 0.03 | 0.05 | 0.10 | 9.0E-03 | 0.12 |
| SMC2 | -0.12 | -0.11 | -0.19 | 0.04 | 0.21 | 0.12 | 9.1E-03 | 0.26 |
| CEP170B | -0.28 | -0.15 | -0.07 | 0.16 | 0.19 | 0.10 | 9.1E-03 | 0.31 |
| TMEM230 | 0.24 | 0.24 | 0.26 | -0.47 | -0.08 | -0.38 | 9.1E-03 | -0.56 |
| BRD1 | -0.02 | -0.05 | -0.06 | 0.02 | 0.04 | 0.07 | 9.1E-03 | 0.09 |
| GIT1 | -0.21 | -0.12 | -0.19 | 0.27 | 0.07 | 0.10 | 9.1E-03 | 0.32 |
| ABCB8 | 0.03 | 0.10 | 0.18 | -0.13 | -0.11 | -0.10 | 9.1E-03 | -0.21 |
| UBL5 | 0.09 | 0.10 | 0.17 | -0.19 | -0.04 | -0.17 | 9.1E-03 | -0.25 |
| TGS1 | -0.15 | -0.13 | -0.08 | 0.03 | 0.18 | 0.11 | 9.2E-03 | 0.23 |
| ENTPD2 | 0.07 | 0.27 | 0.36 | -0.36 | -0.15 | -0.34 | 9.2E-03 | -0.52 |
| TRIOBP | -0.02 | -0.08 | -0.08 | 0.08 | 0.03 | 0.05 | 9.2E-03 | 0.12 |
| ABHD14B | -0.31 | -0.10 | -0.11 | 0.15 | 0.15 | 0.16 | 9.2E-03 | 0.32 |
| GTF2F1 | 0.24 | 0.05 | 0.15 | -0.22 | -0.10 | -0.17 | 9.2E-03 | -0.31 |
| ZCCHC10 | 0.04 | 0.15 | 0.17 | -0.15 | -0.07 | -0.20 | 9.2E-03 | -0.26 |
| CMBL | -0.05 | -0.10 | -0.21 | 0.11 | 0.08 | 0.13 | 9.4E-03 | 0.23 |
| CNBP | 0.10 | 0.21 | 0.29 | -0.38 | -0.11 | -0.24 | 9.4E-03 | -0.44 |
| LONP2 | 0.02 | 0.09 | 0.11 | -0.05 | -0.08 | -0.10 | 9.4E-03 | -0.15 |
| ZNF385A | 0.07 | 0.15 | 0.09 | -0.19 | -0.07 | -0.07 | 9.4E-03 | -0.22 |
| PAAF1 | -0.46 | -0.49 | -0.18 | 0.32 | 0.10 | 0.45 | 9.4E-03 | 0.66 |
| ASCC3 | -0.09 | -0.11 | -0.05 | 0.12 | 0.03 | 0.08 | 9.4E-03 | 0.16 |
| RAB2B | 0.11 | 0.15 | 0.07 | -0.21 | -0.06 | -0.10 | 9.5E-03 | -0.23 |
| CCDC88C | -0.76 | -0.35 | -0.45 | 0.33 | 0.13 | 0.62 | 9.5E-03 | 0.88 |
| PCYT1A | -0.19 | -0.21 | -0.24 | 0.31 | 0.22 | 0.02 | 9.5E-03 | 0.40 |
| COA6 | 0.19 | 0.24 | 0.07 | -0.13 | -0.13 | -0.30 | 9.6E-03 | -0.35 |
| TMEM115 | 0.10 | 0.28 | 0.33 | -0.22 | -0.19 | -0.47 | 9.6E-03 | -0.53 |
| BIRC6 | -0.05 | -0.23 | -0.18 | 0.13 | 0.20 | 0.08 | 9.6E-03 | 0.29 |
| CHKB | -0.40 | -0.37 | -0.13 | 0.35 | 0.09 | 0.28 | 9.6E-03 | 0.55 |

| | | | | | | | | |
|---|---|---|---|---|---|---|---|---|
| STAT6 | -0.29 | -0.22 | -0.24 | 0.04 | 0.38 | 0.20 | 9.6E-03 | 0.46 |
| NOLC1 | 0.16 | 0.12 | 0.06 | -0.14 | -0.05 | -0.19 | 9.6E-03 | -0.24 |
| MARCKS | 0.10 | 0.28 | 0.09 | -0.20 | -0.11 | -0.24 | 9.7E-03 | -0.34 |
| PRPF39 | -0.39 | -0.34 | -0.24 | 0.24 | 0.06 | 0.47 | 9.7E-03 | 0.58 |
| COX15 | 0.03 | 0.10 | 0.12 | -0.04 | -0.10 | -0.12 | 9.7E-03 | -0.17 |
| POC1B | 0.10 | 0.14 | 0.10 | -0.16 | -0.19 | -0.03 | 9.7E-03 | -0.24 |
| AHCY | -0.02 | -0.08 | -0.09 | 0.05 | 0.08 | 0.04 | 9.7E-03 | 0.12 |
| TRAK2 | 0.10 | 0.06 | 0.12 | -0.16 | -0.10 | -0.04 | 9.8E-03 | -0.20 |
| VPS39 | -0.06 | -0.11 | -0.02 | 0.08 | 0.05 | 0.06 | 9.8E-03 | 0.13 |
| PLXNB2 | 0.04 | 0.06 | 0.15 | -0.10 | -0.11 | -0.06 | 9.9E-03 | -0.17 |
| IDE | -0.12 | -0.03 | -0.13 | 0.06 | 0.14 | 0.07 | 9.9E-03 | 0.18 |
| CCDC14 | 0.10 | 0.13 | 0.17 | -0.19 | -0.03 | -0.22 | 9.9E-03 | -0.28 |
| RAD54L2 | -0.13 | -0.31 | -0.17 | 0.08 | 0.16 | 0.28 | 9.9E-03 | 0.37 |
| ABCB7 | 0.02 | 0.25 | 0.31 | -0.29 | -0.21 | -0.19 | 9.9E-03 | -0.43 |
| ALDH3A2 | -0.09 | -0.18 | -0.22 | 0.08 | 0.11 | 0.25 | 9.9E-03 | 0.30 |
| LAP3 | -0.03 | -0.15 | -0.20 | 0.14 | 0.12 | 0.09 | 9.9E-03 | 0.25 |
| NSUN2 | -0.10 | -0.13 | -0.11 | 0.16 | 0.12 | 0.01 | 1.0E-02 | 0.21 |
| SSR4 | -0.05 | -0.06 | -0.07 | 0.10 | 0.01 | 0.06 | 1.0E-02 | 0.12 |
| CALCOCO2 | -0.08 | -0.06 | -0.08 | 0.12 | 0.04 | 0.03 | 1.0E-02 | 0.14 |
| POLR3H | -0.18 | -0.16 | -0.08 | 0.22 | 0.04 | 0.12 | 1.0E-02 | 0.27 |
| PRR3 | 0.18 | 0.14 | 0.17 | -0.29 | -0.04 | -0.23 | 1.0E-02 | -0.35 |
| TMEM87A | 0.11 | 0.15 | 0.40 | -0.36 | -0.18 | -0.26 | 1.0E-02 | -0.48 |
| TNIP2 | 0.08 | 0.15 | 0.37 | -0.20 | -0.20 | -0.33 | 1.0E-02 | -0.44 |
| SYNJ2BP | 0.04 | 0.12 | 0.16 | -0.18 | -0.07 | -0.10 | 1.0E-02 | -0.22 |
| OS9 | 0.04 | 0.13 | 0.20 | -0.09 | -0.11 | -0.20 | 1.0E-02 | -0.26 |
| MTUS1 | 0.03 | 0.13 | 0.20 | -0.17 | -0.11 | -0.12 | 1.0E-02 | -0.25 |
| CCDC42 | 0.10 | 0.06 | 0.12 | -0.18 | -0.06 | -0.06 | 1.0E-02 | -0.20 |
| ACBD6 | -0.71 | -0.30 | -0.33 | 0.14 | 0.51 | 0.33 | 1.0E-02 | 0.77 |
| MYH9 | 0.06 | 0.03 | 0.14 | -0.06 | -0.08 | -0.11 | 1.0E-02 | -0.16 |
| GRK2 | -0.14 | -0.19 | -0.11 | 0.10 | 0.05 | 0.24 | 1.0E-02 | 0.27 |
| RPL8 | -0.03 | -0.10 | -0.11 | 0.10 | 0.09 | 0.03 | 1.0E-02 | 0.15 |
| ALKBH2 | -0.15 | -0.09 | -0.06 | 0.16 | 0.07 | 0.06 | 1.0E-02 | 0.19 |
| KDM6B | -0.20 | -0.14 | -0.07 | 0.05 | 0.12 | 0.20 | 1.0E-02 | 0.26 |
| PEBP1 | 0.22 | 0.19 | 0.03 | -0.20 | -0.09 | -0.22 | 1.0E-02 | -0.32 |
| MTOR | -0.07 | -0.15 | -0.08 | 0.12 | 0.13 | 0.02 | 1.0E-02 | 0.19 |
| ZNF32 | 0.03 | 0.05 | 0.10 | -0.03 | -0.07 | -0.09 | 1.0E-02 | -0.12 |
| BAZ2B | -0.09 | -0.11 | -0.09 | 0.04 | 0.05 | 0.17 | 1.0E-02 | 0.19 |
| ZNF652 | -0.05 | -0.08 | -0.02 | 0.03 | 0.06 | 0.06 | 1.0E-02 | 0.09 |
| BCAM | 0.02 | 0.26 | 0.40 | -0.29 | -0.27 | -0.27 | 1.0E-02 | -0.50 |
| IKBIP | 0.07 | 0.15 | 0.35 | -0.22 | -0.29 | -0.16 | 1.0E-02 | -0.41 |
| FAM3C | 0.08 | 0.22 | 0.18 | -0.20 | -0.07 | -0.29 | 1.0E-02 | -0.35 |
| TMA7 | 0.23 | 0.28 | 0.05 | -0.28 | -0.10 | -0.28 | 1.0E-02 | -0.41 |
| CKAP2L | -0.16 | -0.24 | -0.20 | 0.02 | 0.28 | 0.21 | 1.0E-02 | 0.37 |
| BLVRB | -0.16 | -0.28 | -0.09 | 0.06 | 0.19 | 0.21 | 1.0E-02 | 0.33 |
| COA1 | 0.12 | 0.26 | 0.14 | -0.08 | -0.20 | -0.33 | 1.0E-02 | -0.38 |
| PUF60 | 0.07 | 0.05 | 0.01 | -0.06 | -0.03 | -0.05 | 1.0E-02 | -0.09 |
| GSTZ1 | -0.09 | -0.16 | -0.18 | 0.02 | 0.17 | 0.18 | 1.0E-02 | 0.27 |
| BCLAF3 | -0.23 | -0.25 | -0.19 | 0.11 | 0.09 | 0.37 | 1.0E-02 | 0.41 |

| | | | | | | | | |
|---|---|---|---|---|---|---|---|---|
| TIGD2 | -0.18 | -0.37 | -0.24 | 0.12 | 0.13 | 0.39 | 1.0E-02 | 0.47 |
| TRIM4 | -0.04 | -0.03 | -0.03 | 0.06 | 0.00 | 0.05 | 1.0E-02 | 0.07 |
| SEC63 | 0.05 | 0.06 | 0.13 | -0.13 | -0.04 | -0.09 | 1.0E-02 | -0.16 |
| DBNL | 0.07 | 0.13 | 0.07 | -0.06 | -0.06 | -0.17 | 1.1E-02 | -0.19 |
| PEX1 | -0.07 | -0.08 | -0.10 | 0.13 | 0.01 | 0.10 | 1.1E-02 | 0.16 |
| OSBP | 0.17 | 0.19 | 0.14 | -0.11 | -0.12 | -0.35 | 1.1E-02 | -0.36 |
| SWAP70 | 0.08 | 0.17 | 0.03 | -0.07 | -0.12 | -0.12 | 1.1E-02 | -0.20 |
| MRPL40 | 0.06 | 0.09 | 0.03 | -0.07 | -0.10 | -0.02 | 1.1E-02 | -0.12 |
| PML | -0.09 | -0.08 | -0.12 | 0.10 | 0.15 | 0.02 | 1.1E-02 | 0.19 |
| TARS1 | -0.10 | -0.12 | -0.17 | 0.02 | 0.17 | 0.17 | 1.1E-02 | 0.25 |
| SNX2 | -0.17 | -0.24 | -0.16 | 0.09 | 0.09 | 0.31 | 1.1E-02 | 0.36 |
| AHNAK | -0.14 | -0.04 | -0.09 | 0.12 | 0.08 | 0.05 | 1.1E-02 | 0.17 |
| ARL3 | -0.08 | -0.04 | -0.17 | 0.09 | 0.07 | 0.10 | 1.1E-02 | 0.18 |
| HMGA1 | -0.03 | -0.08 | -0.11 | 0.06 | 0.04 | 0.11 | 1.1E-02 | 0.14 |
| ZNF585B | 0.04 | 0.09 | 0.11 | -0.08 | -0.04 | -0.15 | 1.1E-02 | -0.17 |
| TRAPPC3 | -0.08 | -0.09 | -0.16 | 0.17 | 0.12 | 0.03 | 1.1E-02 | 0.22 |
| GMEB1 | -0.28 | -0.31 | -0.15 | 0.06 | 0.36 | 0.19 | 1.1E-02 | 0.45 |
| ZNF207 | 0.13 | 0.26 | 0.08 | -0.16 | -0.10 | -0.28 | 1.1E-02 | -0.34 |
| CRMP1 | -0.30 | -0.10 | -0.18 | 0.09 | 0.14 | 0.27 | 1.1E-02 | 0.36 |
| DEGS1 | 0.13 | 0.06 | 0.16 | -0.05 | -0.13 | -0.22 | 1.1E-02 | -0.25 |
| HSPA14 | -0.18 | -0.10 | -0.04 | 0.11 | 0.11 | 0.07 | 1.1E-02 | 0.20 |
| FUCA2 | -0.17 | -0.12 | -0.06 | 0.18 | 0.07 | 0.07 | 1.1E-02 | 0.22 |
| ZNF496 | -0.24 | -0.23 | -0.33 | 0.01 | 0.34 | 0.30 | 1.1E-02 | 0.49 |
| STAT3 | -0.07 | -0.15 | -0.15 | 0.14 | 0.17 | 0.02 | 1.1E-02 | 0.23 |
| VPS18 | -0.10 | -0.22 | -0.06 | 0.15 | 0.12 | 0.07 | 1.1E-02 | 0.24 |
| OTUB1 | -0.13 | -0.11 | -0.33 | 0.24 | 0.16 | 0.10 | 1.1E-02 | 0.36 |
| METTL13 | -0.28 | -0.37 | -0.23 | 0.01 | 0.36 | 0.33 | 1.1E-02 | 0.53 |
| LIG1 | 0.14 | 0.20 | 0.02 | -0.18 | -0.09 | -0.13 | 1.1E-02 | -0.25 |
| LTV1 | -0.26 | -0.35 | -0.05 | 0.19 | 0.18 | 0.19 | 1.1E-02 | 0.41 |
| PTPA | 0.14 | 0.09 | 0.13 | -0.24 | -0.10 | -0.06 | 1.1E-02 | -0.25 |
| STT3A | 0.04 | 0.07 | 0.18 | -0.08 | -0.09 | -0.14 | 1.1E-02 | -0.20 |
| ADAM15 | 0.12 | 0.13 | 0.09 | -0.17 | -0.02 | -0.18 | 1.1E-02 | -0.24 |
| TOPBP1 | -0.06 | -0.21 | -0.14 | 0.05 | 0.14 | 0.18 | 1.1E-02 | 0.26 |
| RFC5 | -0.09 | -0.08 | -0.11 | 0.01 | 0.13 | 0.12 | 1.1E-02 | 0.18 |
| STX6 | -0.27 | -0.28 | -0.17 | 0.12 | 0.10 | 0.38 | 1.1E-02 | 0.44 |
| ATP1B1 | 0.01 | 0.16 | 0.15 | -0.12 | -0.15 | -0.09 | 1.1E-02 | -0.23 |
| ATAD3A | 0.06 | 0.17 | 0.11 | -0.05 | -0.13 | -0.20 | 1.1E-02 | -0.23 |
| SRD5A3 | 0.06 | 0.31 | 0.37 | -0.41 | -0.14 | -0.38 | 1.1E-02 | -0.55 |
| TMCO1 | -0.09 | -0.09 | -0.04 | 0.04 | 0.04 | 0.12 | 1.1E-02 | 0.14 |
| DAXX | -0.06 | -0.07 | -0.13 | 0.09 | 0.13 | 0.03 | 1.1E-02 | 0.17 |
| MB21D2 | 0.13 | 0.13 | 0.11 | -0.13 | -0.04 | -0.24 | 1.1E-02 | -0.26 |
| RALGAPB | -0.10 | -0.17 | -0.06 | 0.12 | 0.04 | 0.15 | 1.1E-02 | 0.21 |
| PCGF1 | 0.12 | 0.03 | 0.08 | -0.07 | -0.04 | -0.13 | 1.1E-02 | -0.16 |
| RALA | 0.03 | 0.17 | 0.10 | -0.14 | -0.07 | -0.12 | 1.1E-02 | -0.21 |
| ARF5 | -0.09 | -0.18 | -0.08 | 0.04 | 0.17 | 0.10 | 1.1E-02 | 0.22 |
| HLA-C | 0.07 | 0.21 | 0.23 | -0.09 | -0.19 | -0.31 | 1.1E-02 | -0.37 |
| ATP5F1A | 0.03 | 0.18 | 0.25 | -0.24 | -0.15 | -0.14 | 1.1E-02 | -0.32 |
| AFDN | -0.16 | -0.10 | -0.05 | 0.04 | 0.10 | 0.14 | 1.1E-02 | 0.20 |

| | | | | | | | | |
|---|---|---|---|---|---|---|---|---|
| **CCDC71L** | -0.51 | -0.34 | -0.10 | 0.33 | 0.26 | 0.16 | 1.1E-02 | 0.57 |
| **PTPN1** | 0.04 | 0.14 | 0.17 | -0.09 | -0.09 | -0.21 | 1.1E-02 | -0.25 |
| **TRAPPC9** | -0.19 | -0.06 | -0.08 | 0.15 | 0.07 | 0.09 | 1.1E-02 | 0.21 |
| **ITPRID2** | 0.03 | 0.07 | 0.10 | -0.03 | -0.06 | -0.11 | 1.1E-02 | -0.13 |
| **THNSL1** | -0.10 | -0.07 | -0.18 | 0.03 | 0.16 | 0.12 | 1.1E-02 | 0.22 |
| **PROM2** | 0.17 | 0.10 | 0.39 | -0.37 | -0.31 | -0.14 | 1.1E-02 | -0.49 |
| **FANCD2** | -0.09 | -0.18 | -0.16 | 0.03 | 0.13 | 0.22 | 1.1E-02 | 0.27 |
| **SNTB2** | -0.22 | -0.12 | -0.09 | 0.09 | 0.07 | 0.22 | 1.1E-02 | 0.27 |
| **OSBPL8** | 0.04 | 0.19 | 0.07 | -0.10 | -0.10 | -0.14 | 1.1E-02 | -0.21 |
| **RPS9** | -0.08 | -0.15 | -0.02 | 0.10 | 0.06 | 0.07 | 1.1E-02 | 0.16 |
| **LIN37** | -0.16 | -0.08 | -0.08 | 0.03 | 0.11 | 0.15 | 1.1E-02 | 0.20 |
| **ATP6V0C** | -0.11 | -0.08 | -0.13 | 0.08 | 0.03 | 0.18 | 1.1E-02 | 0.20 |
| **ZNF777** | -0.22 | -0.15 | -0.09 | 0.13 | 0.04 | 0.23 | 1.1E-02 | 0.29 |
| **FNBP4** | 0.15 | 0.14 | 0.07 | -0.07 | -0.09 | -0.23 | 1.2E-02 | -0.25 |
| **HDHD5** | -0.04 | -0.14 | -0.20 | 0.07 | 0.11 | 0.15 | 1.2E-02 | 0.24 |
| **RNF20** | -0.06 | -0.12 | -0.14 | 0.02 | 0.12 | 0.15 | 1.2E-02 | 0.20 |
| **NVL** | 0.07 | 0.10 | 0.10 | -0.08 | -0.03 | -0.17 | 1.2E-02 | -0.19 |
| **RCC1L** | 0.09 | 0.05 | 0.03 | -0.02 | -0.08 | -0.09 | 1.2E-02 | -0.12 |
| **UMPS** | -0.08 | -0.22 | -0.06 | 0.13 | 0.09 | 0.11 | 1.2E-02 | 0.23 |
| **MCM4** | 0.19 | 0.07 | 0.15 | -0.05 | -0.18 | -0.25 | 1.2E-02 | -0.30 |
| **PCM1** | 0.05 | 0.08 | 0.10 | -0.02 | -0.13 | -0.09 | 1.2E-02 | -0.15 |
| **FOXP4** | -0.53 | -0.49 | -0.35 | 0.22 | 0.10 | 0.66 | 1.2E-02 | 0.78 |
| **ITPR3** | 0.04 | 0.05 | 0.05 | -0.07 | -0.01 | -0.07 | 1.2E-02 | -0.10 |
| **GSDMD** | -0.24 | -0.24 | -0.04 | 0.10 | 0.15 | 0.19 | 1.2E-02 | 0.32 |
| **ZNF768** | 0.05 | 0.04 | 0.12 | -0.03 | -0.11 | -0.09 | 1.2E-02 | -0.14 |
| **SLC12A2** | 0.02 | 0.11 | 0.16 | -0.08 | -0.09 | -0.15 | 1.2E-02 | -0.20 |
| **CYP1B1** | -0.39 | -0.20 | -0.08 | 0.18 | 0.15 | 0.24 | 1.2E-02 | 0.41 |
| **ACSS2** | -0.31 | -0.30 | -0.26 | 0.03 | 0.23 | 0.44 | 1.2E-02 | 0.53 |
| **RHOT1** | 0.13 | 0.08 | 0.05 | -0.11 | -0.03 | -0.14 | 1.2E-02 | -0.18 |
| **ACY1** | -0.37 | -0.32 | -0.19 | 0.14 | 0.11 | 0.45 | 1.2E-02 | 0.53 |
| **NDUFA8** | -0.06 | -0.06 | -0.04 | 0.09 | 0.01 | 0.05 | 1.2E-02 | 0.11 |
| **APP** | 0.14 | 0.11 | 0.08 | -0.02 | -0.17 | -0.18 | 1.2E-02 | -0.23 |
| **HK1** | -0.14 | -0.36 | -0.34 | 0.10 | 0.17 | 0.40 | 1.2E-02 | 0.51 |
| **ABT1** | 0.10 | 0.09 | 0.10 | -0.03 | -0.10 | -0.19 | 1.2E-02 | -0.21 |
| **EEIG1** | 0.18 | 0.02 | 0.15 | -0.18 | -0.12 | -0.09 | 1.2E-02 | -0.24 |
| **MGST2** | 0.09 | 0.09 | 0.18 | -0.06 | -0.11 | -0.22 | 1.2E-02 | -0.25 |
| **PREP** | -0.08 | -0.26 | -0.11 | 0.11 | 0.10 | 0.19 | 1.2E-02 | 0.28 |
| **EIF2B3** | -0.06 | -0.13 | -0.16 | 0.10 | 0.19 | 0.04 | 1.2E-02 | 0.23 |
| **SEPTIN11** | 0.14 | 0.19 | 0.03 | -0.20 | -0.11 | -0.09 | 1.2E-02 | -0.26 |
| **TMT1A** | -0.32 | -0.41 | -0.15 | 0.05 | 0.31 | 0.35 | 1.2E-02 | 0.53 |
| **GRPEL1** | 0.16 | 0.16 | 0.01 | -0.10 | -0.13 | -0.12 | 1.2E-02 | -0.22 |
| **EPB41L4A** | 0.03 | 0.10 | 0.15 | -0.14 | -0.05 | -0.11 | 1.2E-02 | -0.19 |
| **IFIH1** | -0.14 | -0.12 | -0.21 | 0.02 | 0.23 | 0.16 | 1.2E-02 | 0.29 |
| **KDF1** | -0.26 | -0.23 | -0.06 | 0.22 | 0.19 | 0.07 | 1.2E-02 | 0.34 |
| **AKR7A3** | -0.10 | -0.11 | -0.26 | 0.13 | 0.22 | 0.07 | 1.2E-02 | 0.30 |
| **CLPTM1L** | 0.15 | 0.06 | 0.10 | -0.05 | -0.10 | -0.20 | 1.2E-02 | -0.22 |
| **KIF23** | 0.05 | 0.07 | 0.15 | -0.14 | -0.11 | -0.04 | 1.2E-02 | -0.18 |
| **PDIA6** | 0.28 | 0.33 | 0.02 | -0.16 | -0.30 | -0.31 | 1.2E-02 | -0.47 |

| | | | | | | | | |
|---|---|---|---|---|---|---|---|---|
| SUGP2 | 0.05 | 0.03 | 0.03 | -0.06 | -0.05 | -0.01 | 1.2E-02 | -0.08 |
| LENG9 | -0.13 | -0.17 | -0.04 | 0.08 | 0.17 | 0.06 | 1.2E-02 | 0.22 |
| SLC1A5 | 0.04 | 0.29 | 0.49 | -0.48 | -0.24 | -0.37 | 1.2E-02 | -0.64 |
| FAM32A | 0.04 | 0.02 | 0.08 | -0.07 | -0.03 | -0.05 | 1.2E-02 | -0.10 |
| PEX3 | 0.17 | 0.17 | 0.13 | -0.22 | -0.03 | -0.29 | 1.2E-02 | -0.34 |
| ZDHHC20 | 0.01 | 0.16 | 0.18 | -0.10 | -0.17 | -0.13 | 1.2E-02 | -0.25 |
| RACK1 | -0.14 | -0.20 | -0.06 | 0.21 | 0.06 | 0.10 | 1.2E-02 | 0.26 |
| SH3GLB2 | -0.23 | -0.14 | -0.16 | 0.03 | 0.15 | 0.27 | 1.2E-02 | 0.32 |
| JUP | -0.32 | -0.27 | -0.18 | 0.31 | 0.32 | 0.01 | 1.2E-02 | 0.47 |
| TAF10 | -0.09 | -0.18 | -0.13 | 0.07 | 0.07 | 0.22 | 1.2E-02 | 0.25 |
| MYC | 0.10 | 0.04 | 0.20 | -0.07 | -0.15 | -0.15 | 1.2E-02 | -0.23 |
| PPFIBP2 | -0.13 | -0.17 | -0.08 | 0.10 | 0.04 | 0.21 | 1.3E-02 | 0.24 |
| SCAF1 | 0.03 | 0.04 | 0.02 | -0.04 | -0.03 | -0.01 | 1.3E-02 | -0.05 |
| TMOD3 | -0.07 | -0.13 | -0.06 | 0.11 | 0.02 | 0.12 | 1.3E-02 | 0.17 |
| ABCD1 | 0.05 | 0.13 | 0.07 | -0.13 | -0.03 | -0.12 | 1.3E-02 | -0.17 |
| MARCKSL1 | 0.01 | 0.25 | 0.26 | -0.18 | -0.16 | -0.27 | 1.3E-02 | -0.38 |
| COASY | -0.46 | -0.23 | -0.23 | 0.25 | 0.07 | 0.42 | 1.3E-02 | 0.55 |
| SYNE4 | 0.11 | 0.03 | 0.15 | -0.14 | -0.04 | -0.14 | 1.3E-02 | -0.21 |
| PRKAR2A | -0.21 | -0.06 | -0.15 | 0.21 | 0.10 | 0.06 | 1.3E-02 | 0.26 |
| MRPS35 | 0.21 | 0.18 | 0.07 | -0.26 | -0.06 | -0.20 | 1.3E-02 | -0.33 |
| LRRCC1 | 0.10 | 0.15 | 0.22 | -0.07 | -0.16 | -0.31 | 1.3E-02 | -0.34 |
| MOB4 | -0.02 | -0.09 | -0.03 | 0.03 | 0.05 | 0.06 | 1.3E-02 | 0.10 |
| NSD1 | 0.11 | 0.07 | 0.03 | -0.11 | -0.07 | -0.03 | 1.3E-02 | -0.14 |
| MORC3 | -0.16 | -0.11 | -0.07 | 0.19 | 0.07 | 0.05 | 1.3E-02 | 0.21 |
| NUCB1 | 0.12 | 0.19 | 0.03 | -0.07 | -0.17 | -0.12 | 1.3E-02 | -0.24 |
| PPDPF | 0.10 | 0.34 | 0.35 | -0.24 | -0.20 | -0.56 | 1.3E-02 | -0.60 |
| EIF2B5 | -0.05 | -0.17 | -0.15 | 0.19 | 0.10 | 0.05 | 1.3E-02 | 0.24 |
| MORF4L1 | 0.04 | 0.11 | 0.14 | -0.16 | -0.03 | -0.13 | 1.3E-02 | -0.21 |
| DENND2D | -0.05 | -0.04 | -0.12 | 0.09 | 0.03 | 0.07 | 1.3E-02 | 0.13 |
| TRIM33 | -0.13 | -0.06 | -0.16 | 0.09 | 0.04 | 0.19 | 1.3E-02 | 0.22 |
| CLPB | 0.08 | 0.22 | 0.12 | -0.24 | -0.05 | -0.19 | 1.3E-02 | -0.30 |
| DERL1 | 0.06 | 0.08 | 0.22 | -0.12 | -0.09 | -0.20 | 1.3E-02 | -0.26 |
| UXS1 | 0.07 | 0.12 | 0.10 | -0.05 | -0.07 | -0.19 | 1.3E-02 | -0.20 |
| NAT1 | -0.20 | -0.46 | -0.15 | 0.32 | 0.23 | 0.11 | 1.3E-02 | 0.49 |
| CTCF | -0.07 | -0.14 | -0.09 | 0.07 | 0.04 | 0.17 | 1.3E-02 | 0.19 |
| AP2A2 | -0.20 | -0.09 | -0.04 | 0.09 | 0.08 | 0.13 | 1.3E-02 | 0.21 |
| AK2 | 0.15 | 0.18 | 0.10 | -0.28 | -0.04 | -0.16 | 1.3E-02 | -0.30 |
| PLCG1 | 0.16 | 0.12 | 0.32 | -0.15 | -0.17 | -0.41 | 1.3E-02 | -0.45 |
| EIF1AX | 0.06 | 0.06 | 0.10 | -0.01 | -0.12 | -0.10 | 1.3E-02 | -0.15 |
| CEPT1 | 0.08 | 0.17 | 0.12 | -0.07 | -0.10 | -0.25 | 1.3E-02 | -0.26 |
| PBX1 | -0.08 | -0.14 | -0.10 | 0.05 | 0.06 | 0.19 | 1.3E-02 | 0.21 |
| RRBP1 | 0.05 | 0.17 | 0.17 | -0.05 | -0.16 | -0.22 | 1.3E-02 | -0.27 |
| DECR1 | -0.03 | -0.05 | -0.11 | 0.03 | 0.06 | 0.10 | 1.3E-02 | 0.13 |
| EXT2 | 0.09 | 0.12 | 0.24 | -0.10 | -0.12 | -0.29 | 1.3E-02 | -0.32 |
| SLC25A28 | 0.01 | 0.29 | 0.25 | -0.16 | -0.27 | -0.22 | 1.3E-02 | -0.40 |
| TAF5 | -0.02 | -0.09 | -0.11 | 0.11 | 0.03 | 0.08 | 1.3E-02 | 0.15 |
| RNPEP | -0.15 | -0.09 | -0.26 | 0.20 | 0.04 | 0.19 | 1.3E-02 | 0.30 |
| DGAT1 | 0.10 | 0.18 | 0.05 | -0.15 | -0.05 | -0.17 | 1.3E-02 | -0.23 |

| | | | | | | | | |
|---|---|---|---|---|---|---|---|---|
| **KIAA0319L** | 0.10 | 0.16 | 0.15 | -0.05 | -0.14 | -0.28 | 1.3E-02 | -0.29 |
| **FAM83H** | -0.08 | -0.04 | -0.01 | 0.04 | 0.05 | 0.04 | 1.3E-02 | 0.09 |
| **NASP** | -0.04 | -0.07 | -0.07 | 0.10 | 0.01 | 0.06 | 1.3E-02 | 0.12 |
| **PKP3** | -0.12 | -0.08 | -0.02 | 0.10 | 0.06 | 0.05 | 1.3E-02 | 0.14 |
| **MMUT** | -0.03 | -0.11 | -0.21 | 0.08 | 0.12 | 0.11 | 1.3E-02 | 0.22 |
| **HDGF** | 0.21 | 0.18 | 0.01 | -0.12 | -0.13 | -0.21 | 1.3E-02 | -0.29 |
| **NNT** | 0.01 | 0.14 | 0.09 | -0.09 | -0.07 | -0.09 | 1.3E-02 | -0.16 |
| **SF3A2** | 0.09 | 0.04 | 0.03 | -0.09 | -0.03 | -0.05 | 1.3E-02 | -0.11 |
| **ITPR2** | 0.02 | 0.01 | 0.03 | -0.01 | -0.01 | -0.04 | 1.4E-02 | -0.04 |
| **PHF23** | 0.04 | 0.20 | 0.08 | -0.08 | -0.10 | -0.15 | 1.4E-02 | -0.22 |
| **NOP2** | 0.08 | 0.07 | 0.08 | -0.08 | -0.02 | -0.14 | 1.4E-02 | -0.15 |
| **XAB2** | 0.07 | 0.04 | 0.13 | -0.09 | -0.03 | -0.13 | 1.4E-02 | -0.16 |
| **STAG3** | -0.09 | -0.07 | -0.04 | 0.08 | 0.01 | 0.10 | 1.4E-02 | 0.13 |
| **CBX2** | 0.05 | 0.05 | 0.09 | -0.12 | -0.07 | -0.02 | 1.4E-02 | -0.13 |
| **SPTLC2** | -0.04 | -0.07 | -0.19 | 0.10 | 0.07 | 0.12 | 1.4E-02 | 0.20 |
| **KIAA1217** | -0.11 | -0.02 | -0.06 | 0.08 | 0.07 | 0.03 | 1.4E-02 | 0.12 |
| **TMEM165** | 0.07 | 0.23 | 0.19 | -0.16 | -0.09 | -0.32 | 1.4E-02 | -0.36 |
| **PURB** | -0.22 | -0.03 | -0.15 | 0.10 | 0.09 | 0.18 | 1.4E-02 | 0.25 |
| **RCC1** | -0.06 | -0.14 | -0.15 | 0.15 | 0.01 | 0.15 | 1.4E-02 | 0.22 |
| **RO60** | 0.07 | 0.09 | 0.09 | -0.09 | -0.02 | -0.17 | 1.4E-02 | -0.18 |
| **NDUFS4** | 0.05 | 0.03 | 0.08 | -0.07 | -0.01 | -0.09 | 1.4E-02 | -0.12 |
| **UGDH** | -0.08 | -0.11 | -0.01 | 0.06 | 0.09 | 0.05 | 1.4E-02 | 0.13 |
| **DHX36** | 0.03 | 0.07 | 0.08 | -0.03 | -0.11 | -0.04 | 1.4E-02 | -0.12 |
| **SUN1** | -0.06 | -0.10 | -0.09 | 0.00 | 0.12 | 0.12 | 1.4E-02 | 0.17 |
| **ERCC6L** | 0.08 | 0.12 | 0.12 | -0.15 | -0.01 | -0.18 | 1.4E-02 | -0.22 |
| **MRPS9** | 0.18 | 0.09 | 0.05 | -0.17 | -0.05 | -0.13 | 1.4E-02 | -0.22 |
| **TIAL1** | 0.03 | 0.08 | 0.07 | -0.11 | -0.05 | -0.03 | 1.4E-02 | -0.12 |
| **AKR7A2** | -0.05 | -0.13 | -0.09 | 0.11 | 0.01 | 0.13 | 1.4E-02 | 0.17 |
| **NRK** | 0.03 | 0.08 | 0.16 | -0.11 | -0.12 | -0.05 | 1.4E-02 | -0.18 |
| **NCDN** | -0.10 | -0.06 | -0.22 | 0.07 | 0.10 | 0.18 | 1.4E-02 | 0.24 |
| **ZNF512** | -0.06 | -0.07 | -0.17 | 0.07 | 0.06 | 0.14 | 1.4E-02 | 0.19 |
| **MTERF3** | 0.13 | 0.13 | 0.05 | -0.13 | -0.03 | -0.18 | 1.4E-02 | -0.21 |
| **MRPS18A** | 0.09 | 0.05 | 0.02 | -0.04 | -0.09 | -0.03 | 1.4E-02 | -0.11 |
| **TGFB3** | 0.19 | 0.20 | 0.01 | -0.21 | -0.11 | -0.13 | 1.4E-02 | -0.28 |
| **H2AZ1** | -0.09 | -0.22 | -0.22 | 0.07 | 0.10 | 0.29 | 1.4E-02 | 0.33 |
| **CPXM1** | 0.40 | 0.37 | 0.28 | -0.25 | -0.33 | -0.89 | 1.4E-02 | -0.84 |
| **SLC25A19** | 0.04 | 0.14 | 0.09 | -0.05 | -0.08 | -0.17 | 1.4E-02 | -0.19 |
| **SNRNP70** | 0.18 | 0.25 | 0.21 | -0.12 | -0.18 | -0.48 | 1.4E-02 | -0.47 |
| **RTN3** | -0.08 | -0.08 | -0.12 | 0.01 | 0.09 | 0.16 | 1.4E-02 | 0.18 |
| **RPS6KB1** | -0.15 | -0.15 | -0.01 | 0.07 | 0.12 | 0.10 | 1.4E-02 | 0.20 |
| **DVL2** | -0.41 | -0.33 | -0.24 | 0.47 | 0.02 | 0.27 | 1.4E-02 | 0.58 |
| **DNAJC8** | 0.11 | 0.04 | 0.08 | -0.15 | -0.05 | -0.05 | 1.4E-02 | -0.16 |
| **TRIM21** | -0.15 | -0.32 | -0.27 | 0.40 | 0.12 | 0.10 | 1.4E-02 | 0.45 |
| **CAB39** | 0.19 | 0.22 | 0.24 | -0.15 | -0.15 | -0.50 | 1.4E-02 | -0.48 |
| **UTP14A** | 0.15 | 0.13 | 0.14 | -0.15 | -0.04 | -0.29 | 1.4E-02 | -0.30 |
| **CIT** | 0.06 | 0.06 | 0.17 | -0.05 | -0.12 | -0.15 | 1.4E-02 | -0.20 |
| **ARHGEF39** | 0.12 | 0.18 | 0.07 | -0.24 | -0.11 | -0.06 | 1.4E-02 | -0.26 |
| **TASOR2** | 0.03 | 0.04 | 0.08 | -0.08 | -0.06 | -0.01 | 1.4E-02 | -0.10 |

| | | | | | | | | |
|---|---|---|---|---|---|---|---|---|
| SNRPC | 0.07 | 0.09 | 0.29 | -0.25 | -0.13 | -0.14 | 1.4E-02 | -0.33 |
| DHODH | 0.09 | 0.16 | 0.22 | -0.12 | -0.10 | -0.33 | 1.4E-02 | -0.34 |
| IPO13 | -0.43 | -0.15 | -0.13 | 0.27 | 0.19 | 0.14 | 1.4E-02 | 0.44 |
| TNIK | 0.00 | 0.18 | 0.14 | -0.10 | -0.12 | -0.13 | 1.5E-02 | -0.23 |
| HERC2 | 0.04 | 0.13 | 0.08 | -0.06 | -0.05 | -0.15 | 1.5E-02 | -0.17 |
| CLPP | -0.01 | -0.07 | -0.11 | 0.06 | 0.07 | 0.05 | 1.5E-02 | 0.12 |
| CDCA8 | -0.11 | -0.09 | -0.05 | 0.03 | 0.07 | 0.14 | 1.5E-02 | 0.16 |
| MRPL30 | 0.13 | 0.10 | 0.19 | -0.29 | -0.14 | -0.05 | 1.5E-02 | -0.30 |
| TLK2 | -0.11 | -0.14 | -0.07 | 0.16 | 0.01 | 0.12 | 1.5E-02 | 0.20 |
| CHPF | -0.03 | -0.02 | -0.06 | 0.06 | 0.05 | 0.01 | 1.5E-02 | 0.08 |
| PDE7A | -0.13 | -0.12 | -0.10 | 0.14 | 0.18 | 0.00 | 1.5E-02 | 0.22 |
| LIN9 | -0.04 | -0.06 | -0.05 | 0.07 | 0.06 | 0.00 | 1.5E-02 | 0.09 |
| TMA16 | -0.06 | -0.09 | -0.16 | 0.10 | 0.16 | 0.02 | 1.5E-02 | 0.20 |
| TSR1 | -0.10 | -0.20 | -0.05 | 0.16 | 0.11 | 0.05 | 1.5E-02 | 0.22 |
| EI24 | 0.24 | 0.10 | 0.17 | -0.30 | -0.04 | -0.25 | 1.5E-02 | -0.37 |
| SCP2 | -0.03 | -0.10 | -0.18 | 0.06 | 0.12 | 0.10 | 1.5E-02 | 0.19 |
| ING4 | 0.04 | 0.08 | 0.08 | -0.01 | -0.09 | -0.10 | 1.5E-02 | -0.13 |
| TIMM13 | 0.12 | 0.09 | 0.04 | -0.11 | -0.02 | -0.14 | 1.5E-02 | -0.17 |
| RNPC3 | 0.19 | 0.12 | 0.07 | -0.08 | -0.09 | -0.25 | 1.5E-02 | -0.26 |
| ARPP19 | 0.18 | 0.23 | 0.07 | -0.30 | -0.20 | -0.06 | 1.5E-02 | -0.35 |
| TFRC | 0.03 | 0.10 | 0.25 | -0.19 | -0.13 | -0.12 | 1.5E-02 | -0.27 |
| MT-ND4 | -0.11 | -0.04 | -0.10 | 0.04 | 0.05 | 0.15 | 1.5E-02 | 0.16 |
| EXOC8 | -0.14 | -0.23 | -0.27 | 0.04 | 0.15 | 0.34 | 1.5E-02 | 0.39 |
| ZNF597 | 0.20 | 0.02 | 0.10 | -0.10 | -0.15 | -0.11 | 1.5E-02 | -0.22 |
| EIF2AK3 | 0.06 | 0.12 | 0.26 | -0.25 | -0.16 | -0.09 | 1.5E-02 | -0.31 |
| COPA | -0.07 | -0.10 | -0.02 | 0.07 | 0.02 | 0.09 | 1.5E-02 | 0.13 |
| HOOK2 | -0.20 | -0.18 | -0.01 | 0.12 | 0.12 | 0.10 | 1.5E-02 | 0.24 |
| MAP2K2 | -0.19 | -0.17 | -0.08 | 0.13 | 0.23 | 0.03 | 1.5E-02 | 0.27 |
| MAP4 | 0.15 | 0.19 | 0.03 | -0.19 | -0.05 | -0.17 | 1.5E-02 | -0.26 |
| SELENOH | -0.13 | -0.13 | -0.30 | 0.04 | 0.20 | 0.25 | 1.5E-02 | 0.35 |
| MRPS5 | 0.15 | 0.10 | 0.06 | -0.19 | -0.03 | -0.13 | 1.5E-02 | -0.22 |
| TSEN34 | -0.07 | -0.16 | -0.05 | 0.08 | 0.04 | 0.14 | 1.5E-02 | 0.18 |
| PSME4 | -0.26 | -0.14 | -0.16 | 0.01 | 0.23 | 0.24 | 1.5E-02 | 0.35 |
| SPAG5 | -0.09 | -0.10 | -0.03 | 0.02 | 0.07 | 0.12 | 1.5E-02 | 0.15 |
| INO80C | -0.06 | -0.08 | -0.17 | 0.15 | 0.10 | 0.03 | 1.5E-02 | 0.20 |
| GGT1 | 0.01 | 0.36 | 0.40 | -0.49 | -0.24 | -0.27 | 1.5E-02 | -0.59 |
| MGST1 | 0.12 | 0.14 | 0.07 | -0.16 | -0.02 | -0.19 | 1.5E-02 | -0.23 |
| C1orf131 | -0.05 | -0.08 | -0.08 | 0.12 | 0.01 | 0.08 | 1.5E-02 | 0.14 |
| CDK5RAP2 | -0.07 | -0.05 | -0.06 | 0.07 | 0.00 | 0.10 | 1.5E-02 | 0.12 |
| MYH14 | 0.03 | 0.06 | 0.14 | -0.07 | -0.06 | -0.12 | 1.5E-02 | -0.16 |
| SNX7 | -0.20 | -0.05 | -0.16 | 0.16 | 0.18 | 0.03 | 1.5E-02 | 0.26 |
| HERPUD1 | 0.10 | 0.13 | 0.25 | -0.21 | -0.29 | -0.05 | 1.5E-02 | -0.34 |
| ITIH4 | -0.17 | -0.07 | -0.13 | 0.10 | 0.03 | 0.20 | 1.5E-02 | 0.23 |
| LARS1 | -0.17 | -0.15 | -0.05 | 0.14 | 0.17 | 0.03 | 1.6E-02 | 0.24 |
| MAVS | 0.00 | 0.19 | 0.17 | -0.14 | -0.14 | -0.12 | 1.6E-02 | -0.25 |
| RAB21 | 0.03 | 0.09 | 0.20 | -0.15 | -0.08 | -0.13 | 1.6E-02 | -0.22 |
| MYORG | 0.05 | 0.02 | 0.10 | -0.05 | -0.04 | -0.10 | 1.6E-02 | -0.12 |
| KMT2C | -0.02 | -0.10 | -0.09 | 0.10 | 0.07 | 0.03 | 1.6E-02 | 0.13 |

| | | | | | | | | |
|---|---|---|---|---|---|---|---|---|
| **CDC5L** | 0.07 | 0.02 | 0.05 | -0.07 | -0.01 | -0.07 | 1.6E-02 | -0.10 |
| **ASF1B** | -0.07 | -0.13 | -0.06 | 0.07 | 0.15 | 0.02 | 1.6E-02 | 0.18 |
| **GTPBP10** | 0.06 | 0.11 | 0.08 | -0.09 | -0.02 | -0.16 | 1.6E-02 | -0.17 |
| **GLCE** | 0.13 | 0.19 | 0.08 | -0.12 | -0.06 | -0.27 | 1.6E-02 | -0.28 |
| **PEX13** | 0.05 | 0.07 | 0.10 | -0.11 | -0.13 | -0.01 | 1.6E-02 | -0.16 |
| **NSL1** | -0.07 | -0.08 | -0.13 | 0.02 | 0.08 | 0.16 | 1.6E-02 | 0.18 |
| **GUCY1A2** | 0.19 | 0.26 | 0.13 | -0.35 | -0.03 | -0.32 | 1.6E-02 | -0.43 |
| **SCAF11** | 0.14 | 0.11 | 0.09 | -0.22 | -0.12 | -0.03 | 1.6E-02 | -0.24 |
| **ARFRP1** | -0.07 | -0.05 | -0.21 | 0.12 | 0.11 | 0.07 | 1.6E-02 | 0.21 |
| **PRR15** | 0.10 | 0.21 | 0.05 | -0.06 | -0.14 | -0.20 | 1.6E-02 | -0.25 |
| **MPG** | -0.09 | -0.12 | -0.04 | 0.08 | 0.02 | 0.14 | 1.6E-02 | 0.17 |
| **GALNT7** | 0.03 | 0.12 | 0.29 | -0.14 | -0.19 | -0.18 | 1.6E-02 | -0.31 |
| **SF1** | 0.08 | 0.13 | 0.07 | -0.13 | -0.01 | -0.17 | 1.6E-02 | -0.20 |
| **CFL1** | 0.22 | 0.16 | 0.06 | -0.06 | -0.17 | -0.27 | 1.6E-02 | -0.31 |
| **DICER1** | 0.03 | 0.14 | 0.19 | -0.09 | -0.09 | -0.22 | 1.6E-02 | -0.25 |
| **RAD1** | -0.19 | -0.16 | -0.06 | 0.03 | 0.20 | 0.14 | 1.6E-02 | 0.26 |
| **RPL36** | -0.09 | -0.36 | -0.17 | 0.09 | 0.18 | 0.26 | 1.6E-02 | 0.38 |
| **SRSF7** | 0.06 | 0.07 | 0.01 | -0.05 | -0.02 | -0.07 | 1.6E-02 | -0.09 |
| **ERP29** | 0.31 | 0.27 | 0.06 | -0.13 | -0.22 | -0.43 | 1.6E-02 | -0.47 |
| **THOC2** | 0.07 | 0.07 | 0.05 | -0.05 | -0.02 | -0.12 | 1.6E-02 | -0.13 |
| **EIF4E** | 0.05 | 0.13 | 0.08 | -0.16 | -0.08 | -0.03 | 1.6E-02 | -0.18 |
| **EIF2A** | 0.13 | 0.10 | 0.09 | -0.07 | -0.06 | -0.23 | 1.6E-02 | -0.23 |
| **SPIRE1** | -0.29 | -0.24 | -0.02 | 0.16 | 0.12 | 0.20 | 1.6E-02 | 0.34 |
| **SRSF3** | 0.12 | 0.10 | 0.09 | -0.03 | -0.09 | -0.21 | 1.6E-02 | -0.21 |
| **REXO4** | -0.05 | -0.04 | -0.15 | 0.10 | 0.08 | 0.05 | 1.6E-02 | 0.15 |
| **DTNA** | -0.28 | -0.20 | -0.09 | 0.12 | 0.07 | 0.31 | 1.6E-02 | 0.36 |
| **REPS1** | -0.13 | -0.07 | -0.09 | 0.16 | 0.10 | 0.01 | 1.6E-02 | 0.19 |
| **MCTS1** | 0.13 | 0.13 | 0.07 | -0.17 | -0.01 | -0.18 | 1.6E-02 | -0.23 |
| **RAB2A** | 0.02 | 0.07 | 0.13 | -0.13 | -0.05 | -0.07 | 1.6E-02 | -0.16 |
| **CCNB2** | -0.07 | -0.06 | -0.17 | 0.06 | 0.15 | 0.06 | 1.6E-02 | 0.18 |
| **PSMA1** | -0.12 | -0.31 | -0.28 | 0.02 | 0.25 | 0.32 | 1.7E-02 | 0.43 |
| **PTBP3** | 0.07 | 0.12 | 0.15 | -0.24 | -0.07 | -0.07 | 1.7E-02 | -0.24 |
| **ACADSB** | -0.08 | -0.09 | -0.11 | 0.00 | 0.16 | 0.11 | 1.7E-02 | 0.18 |
| **TMPRSS11A** | 0.08 | 0.08 | 0.32 | -0.18 | -0.26 | -0.13 | 1.7E-02 | -0.34 |
| **AUP1** | 0.08 | 0.11 | 0.31 | -0.14 | -0.14 | -0.31 | 1.7E-02 | -0.36 |
| **NUBP2** | -0.20 | -0.17 | -0.21 | 0.07 | 0.08 | 0.35 | 1.7E-02 | 0.36 |
| **CCDC85C** | -0.35 | -0.08 | -0.15 | 0.24 | 0.16 | 0.10 | 1.7E-02 | 0.36 |
| **ZSCAN26** | 0.05 | 0.03 | 0.14 | -0.12 | -0.08 | -0.04 | 1.7E-02 | -0.15 |
| **SIX4** | -0.11 | -0.03 | -0.04 | 0.04 | 0.05 | 0.08 | 1.7E-02 | 0.11 |
| **IFIT2** | -0.08 | -0.23 | -0.07 | 0.18 | 0.08 | 0.09 | 1.7E-02 | 0.24 |
| **PRDM15** | -0.16 | -0.25 | -0.11 | 0.16 | 0.02 | 0.27 | 1.7E-02 | 0.32 |
| **PPIL3** | 0.07 | 0.04 | 0.05 | -0.03 | -0.03 | -0.11 | 1.7E-02 | -0.11 |
| **RMND5A** | -0.09 | -0.10 | -0.28 | 0.22 | 0.08 | 0.12 | 1.7E-02 | 0.30 |
| **RGL2** | -0.22 | -0.05 | -0.20 | 0.08 | 0.08 | 0.24 | 1.7E-02 | 0.29 |
| **FCF1** | 0.40 | 0.25 | 0.05 | -0.30 | -0.14 | -0.44 | 1.7E-02 | -0.52 |
| **POLE3** | -0.06 | -0.28 | -0.23 | 0.18 | 0.05 | 0.26 | 1.7E-02 | 0.36 |
| **H1-5** | -0.03 | -0.06 | -0.12 | 0.03 | 0.06 | 0.11 | 1.7E-02 | 0.14 |
| **SLC27A4** | 0.08 | 0.10 | 0.16 | -0.11 | -0.04 | -0.22 | 1.7E-02 | -0.24 |

| | | | | | | | | |
|---|---|---|---|---|---|---|---|---|
| **RPN1** | 0.05 | 0.07 | 0.11 | -0.02 | -0.08 | -0.14 | 1.7E-02 | -0.15 |
| **ALDH18A1** | 0.11 | 0.12 | 0.18 | -0.10 | -0.07 | -0.29 | 1.7E-02 | -0.29 |
| **SSX2IP** | 0.02 | 0.18 | 0.14 | -0.20 | -0.07 | -0.10 | 1.7E-02 | -0.24 |
| **CDKAL1** | 0.06 | 0.17 | 0.26 | -0.27 | -0.05 | -0.25 | 1.7E-02 | -0.36 |
| **TRNT1** | -0.10 | -0.05 | -0.25 | 0.07 | 0.13 | 0.15 | 1.7E-02 | 0.25 |
| **CKS2** | -0.36 | -0.49 | -0.69 | 0.54 | -0.05 | 0.56 | 1.7E-02 | 0.86 |
| **QNG1** | -0.13 | -0.09 | -0.19 | 0.01 | 0.16 | 0.21 | 1.7E-02 | 0.26 |
| **ABCF2** | -0.07 | -0.11 | -0.05 | 0.04 | 0.14 | 0.03 | 1.7E-02 | 0.15 |
| **UQCC1** | -0.10 | -0.09 | -0.07 | 0.13 | 0.00 | 0.11 | 1.7E-02 | 0.17 |
| **PRPF18** | 0.14 | 0.14 | 0.08 | -0.16 | -0.02 | -0.22 | 1.7E-02 | -0.25 |
| **MPC1** | 0.02 | 0.20 | 0.16 | -0.13 | -0.08 | -0.22 | 1.7E-02 | -0.27 |
| **SLC12A4** | 0.02 | 0.09 | 0.21 | -0.13 | -0.08 | -0.14 | 1.7E-02 | -0.22 |
| **TMEM245** | 0.02 | 0.15 | 0.19 | -0.18 | -0.05 | -0.17 | 1.7E-02 | -0.25 |
| **SRRT** | 0.17 | 0.24 | 0.12 | -0.17 | -0.07 | -0.38 | 1.8E-02 | -0.38 |
| **ATP13A3** | 0.04 | 0.11 | 0.22 | -0.23 | -0.09 | -0.10 | 1.8E-02 | -0.26 |
| **SCYL3** | -0.09 | -0.20 | -0.21 | 0.08 | 0.07 | 0.28 | 1.8E-02 | 0.31 |
| **PLEKHA7** | -0.10 | -0.11 | -0.14 | 0.02 | 0.21 | 0.08 | 1.8E-02 | 0.22 |
| **ATP2A3** | 0.09 | 0.10 | 0.20 | -0.16 | -0.03 | -0.25 | 1.8E-02 | -0.28 |
| **STRBP** | 0.08 | 0.05 | 0.05 | -0.05 | -0.13 | -0.02 | 1.8E-02 | -0.13 |
| **ITK** | 0.61 | 0.18 | 0.45 | -1.05 | -0.23 | -0.64 | 1.8E-02 | -1.05 |
| **SUPV3L1** | 0.16 | 0.07 | 0.02 | -0.11 | -0.06 | -0.11 | 1.8E-02 | -0.18 |
| **IPO9** | -0.12 | -0.18 | -0.07 | 0.02 | 0.20 | 0.11 | 1.8E-02 | 0.24 |
| **RAB13** | 0.03 | 0.09 | 0.12 | -0.15 | -0.05 | -0.06 | 1.8E-02 | -0.16 |
| **DAG1** | -0.34 | -0.23 | -0.34 | 0.09 | 0.12 | 0.51 | 1.8E-02 | 0.54 |
| **STEAP3** | 0.02 | 0.18 | 0.33 | -0.28 | -0.13 | -0.21 | 1.8E-02 | -0.38 |
| **POLR3D** | -0.09 | -0.13 | -0.18 | 0.07 | 0.06 | 0.24 | 1.8E-02 | 0.26 |
| **ADI1** | -0.13 | -0.16 | -0.31 | 0.31 | 0.16 | 0.04 | 1.8E-02 | 0.37 |
| **SOX11** | 0.20 | 0.50 | 0.05 | -0.45 | -0.22 | -0.31 | 1.8E-02 | -0.58 |
| **KLC2** | 0.03 | 0.03 | 0.05 | -0.05 | -0.06 | 0.00 | 1.8E-02 | -0.08 |
| **XPC** | -0.04 | -0.09 | -0.14 | 0.04 | 0.06 | 0.15 | 1.8E-02 | 0.18 |
| **CAND1** | 0.24 | 0.17 | 0.00 | -0.20 | -0.10 | -0.17 | 1.8E-02 | -0.30 |
| **TBCA** | 0.31 | 0.30 | 0.01 | -0.15 | -0.37 | -0.23 | 1.8E-02 | -0.46 |
| **XPO5** | -0.06 | -0.18 | -0.03 | 0.07 | 0.09 | 0.10 | 1.8E-02 | 0.18 |
| **RAD23B** | -0.03 | -0.05 | -0.08 | 0.06 | 0.01 | 0.08 | 1.8E-02 | 0.10 |
| **MYEF2** | -0.10 | -0.10 | -0.06 | 0.16 | 0.03 | 0.05 | 1.8E-02 | 0.16 |
| **AACS** | -0.12 | -0.24 | -0.43 | 0.20 | 0.37 | 0.08 | 1.8E-02 | 0.48 |
| **MRPL23** | 0.09 | 0.07 | 0.16 | -0.13 | -0.02 | -0.20 | 1.8E-02 | -0.23 |
| **AK4** | -0.04 | -0.10 | -0.24 | 0.08 | 0.14 | 0.12 | 1.8E-02 | 0.24 |
| **UBE3C** | -0.19 | -0.11 | -0.28 | 0.13 | 0.31 | 0.05 | 1.8E-02 | 0.36 |
| **CPSF7** | -0.03 | -0.08 | -0.13 | 0.12 | 0.02 | 0.08 | 1.8E-02 | 0.16 |
| **NUDT8** | 0.13 | 0.05 | 0.07 | -0.15 | -0.02 | -0.11 | 1.8E-02 | -0.17 |
| **COX11** | 0.06 | 0.01 | 0.12 | -0.07 | -0.05 | -0.07 | 1.8E-02 | -0.12 |
| **RHEB** | -0.05 | -0.19 | -0.20 | 0.21 | 0.15 | 0.03 | 1.8E-02 | 0.28 |
| **ADSL** | 0.24 | 0.23 | 0.10 | -0.08 | -0.19 | -0.41 | 1.8E-02 | -0.42 |
| **RBM26** | -0.03 | -0.07 | -0.11 | 0.09 | 0.09 | 0.02 | 1.8E-02 | 0.13 |
| **HTRA2** | 0.15 | 0.07 | 0.33 | -0.31 | -0.08 | -0.27 | 1.8E-02 | -0.40 |
| **STUB1** | -0.14 | -0.09 | -0.02 | 0.11 | 0.05 | 0.07 | 1.8E-02 | 0.16 |
| **MAFK** | -0.06 | -0.06 | -0.06 | 0.00 | 0.09 | 0.08 | 1.8E-02 | 0.12 |

| | | | | | | | | |
|---|---|---|---|---|---|---|---|---|
| **FLVCR1** | 0.02 | 0.13 | 0.24 | -0.18 | -0.18 | -0.08 | 1.8E-02 | -0.27 |
| **CS** | -0.01 | -0.12 | -0.17 | 0.11 | 0.08 | 0.08 | 1.8E-02 | 0.19 |
| **FBLN1** | -0.36 | -0.50 | -0.20 | 0.21 | 0.06 | 0.53 | 1.9E-02 | 0.63 |
| **LGALS3** | 0.01 | 0.11 | 0.19 | -0.13 | -0.08 | -0.12 | 1.9E-02 | -0.22 |
| **TEDC1** | 0.09 | 0.02 | 0.21 | -0.13 | -0.09 | -0.13 | 1.9E-02 | -0.23 |
| **SEC24C** | 0.08 | 0.15 | 0.19 | -0.18 | -0.03 | -0.26 | 1.9E-02 | -0.30 |
| **CHAF1A** | 0.07 | 0.02 | 0.04 | -0.07 | -0.04 | -0.01 | 1.9E-02 | -0.09 |
| **TRADD** | -0.21 | -0.20 | -0.01 | 0.13 | 0.09 | 0.15 | 1.9E-02 | 0.26 |
| **KHDRBS3** | -0.11 | -0.03 | -0.06 | 0.09 | 0.01 | 0.08 | 1.9E-02 | 0.13 |
| **POLRMT** | 0.15 | 0.15 | 0.05 | -0.14 | -0.03 | -0.22 | 1.9E-02 | -0.25 |
| **KCTD10** | 0.16 | 0.12 | 0.24 | -0.03 | -0.25 | -0.33 | 1.9E-02 | -0.37 |
| **RHBDD2** | 0.03 | 0.06 | 0.15 | -0.06 | -0.06 | -0.14 | 1.9E-02 | -0.17 |
| **MIER1** | 0.02 | 0.05 | 0.01 | -0.04 | -0.02 | -0.02 | 1.9E-02 | -0.05 |
| **SF3B3** | 0.04 | 0.00 | 0.05 | -0.04 | -0.03 | -0.02 | 1.9E-02 | -0.06 |
| **S100A11** | -0.03 | -0.27 | -0.39 | 0.25 | 0.14 | 0.19 | 1.9E-02 | 0.42 |
| **MIDEAS** | -0.12 | -0.12 | -0.10 | 0.12 | 0.00 | 0.19 | 1.9E-02 | 0.22 |
| **EPS15** | -0.25 | -0.15 | -0.06 | 0.03 | 0.20 | 0.18 | 1.9E-02 | 0.29 |
| **ACSF2** | -0.04 | -0.04 | -0.16 | 0.08 | 0.09 | 0.05 | 1.9E-02 | 0.15 |
| **IVD** | -0.05 | -0.11 | -0.27 | 0.10 | 0.11 | 0.16 | 1.9E-02 | 0.27 |
| **SYNE3** | -0.28 | -0.33 | -0.06 | 0.20 | 0.06 | 0.31 | 1.9E-02 | 0.42 |
| **UBE2L3** | 0.27 | 0.14 | 0.11 | -0.05 | -0.21 | -0.35 | 1.9E-02 | -0.38 |
| **PCBD1** | 0.05 | 0.29 | 0.08 | -0.19 | -0.18 | -0.12 | 1.9E-02 | -0.31 |
| **TTC13** | 0.05 | 0.05 | 0.11 | -0.02 | -0.07 | -0.13 | 1.9E-02 | -0.15 |
| **NAE1** | -0.08 | -0.12 | -0.06 | 0.12 | 0.00 | 0.11 | 1.9E-02 | 0.16 |
| **KIF2C** | -0.11 | -0.01 | -0.12 | 0.05 | 0.10 | 0.07 | 1.9E-02 | 0.15 |
| **ACTBL2** | -0.35 | -0.29 | -0.47 | 0.47 | -0.05 | 0.41 | 1.9E-02 | 0.65 |
| **HRAS** | 0.01 | 0.14 | 0.26 | -0.17 | -0.10 | -0.19 | 1.9E-02 | -0.29 |
| **ZC3H18** | 0.15 | 0.16 | 0.10 | -0.14 | -0.03 | -0.30 | 1.9E-02 | -0.30 |
| **EIF4G1** | -0.09 | -0.06 | 0.00 | 0.06 | 0.04 | 0.04 | 1.9E-02 | 0.10 |
| **EPS8** | -0.18 | -0.18 | -0.15 | 0.18 | -0.01 | 0.27 | 1.9E-02 | 0.31 |
| **NDUFA9** | -0.06 | -0.05 | -0.17 | 0.11 | 0.03 | 0.13 | 1.9E-02 | 0.18 |
| **UQCR11** | -0.16 | -0.35 | -0.46 | 0.32 | 0.01 | 0.42 | 1.9E-02 | 0.58 |
| **NDUFA11** | -0.08 | -0.09 | -0.20 | 0.14 | 0.01 | 0.18 | 1.9E-02 | 0.24 |
| **FAM162A** | 0.06 | 0.30 | 0.17 | -0.12 | -0.15 | -0.36 | 1.9E-02 | -0.38 |
| **TIMM17B** | 0.02 | 0.15 | 0.18 | -0.20 | -0.06 | -0.13 | 2.0E-02 | -0.24 |
| **PLIN3** | 0.13 | 0.03 | 0.14 | -0.09 | -0.19 | -0.04 | 2.0E-02 | -0.21 |
| **CROCC** | -0.24 | -0.17 | -0.05 | 0.11 | 0.06 | 0.23 | 2.0E-02 | 0.29 |
| **COQ10B** | 0.23 | 0.10 | 0.12 | -0.21 | -0.03 | -0.28 | 2.0E-02 | -0.32 |
| **MRPS12** | 0.20 | 0.16 | 0.06 | -0.28 | -0.05 | -0.14 | 2.0E-02 | -0.30 |
| **TMEM43** | -0.16 | -0.15 | -0.11 | 0.06 | 0.05 | 0.26 | 2.0E-02 | 0.27 |
| **PLG** | -0.05 | -0.04 | -0.01 | 0.01 | 0.06 | 0.03 | 2.0E-02 | 0.07 |
| **ISG15** | -0.20 | -0.08 | -0.07 | 0.03 | 0.17 | 0.11 | 2.0E-02 | 0.22 |
| **NDUFAF5** | -0.19 | -0.16 | -0.30 | 0.21 | 0.00 | 0.33 | 2.0E-02 | 0.39 |
| **VPS33A** | -0.14 | -0.09 | -0.04 | 0.10 | 0.13 | 0.01 | 2.0E-02 | 0.17 |
| **HNRNPR** | 0.04 | 0.02 | 0.08 | -0.03 | -0.03 | -0.08 | 2.0E-02 | -0.09 |
| **CUSTOS** | 0.07 | 0.10 | 0.11 | -0.11 | -0.01 | -0.17 | 2.0E-02 | -0.18 |
| **USP3** | -0.28 | -0.29 | -0.23 | 0.38 | -0.03 | 0.29 | 2.0E-02 | 0.48 |
| **CHMP1A** | 0.05 | 0.06 | 0.03 | -0.08 | 0.00 | -0.07 | 2.0E-02 | -0.10 |

| | | | | | | | | |
|---|---|---|---|---|---|---|---|---|
| PICALM | -0.16 | -0.11 | -0.09 | 0.01 | 0.10 | 0.21 | 2.0E-02 | 0.22 |
| CEP78 | 0.04 | 0.11 | 0.31 | -0.21 | -0.20 | -0.12 | 2.0E-02 | -0.32 |
| S100P | -0.13 | -0.27 | -0.12 | 0.17 | 0.25 | 0.02 | 2.0E-02 | 0.32 |
| CDCA7 | 0.25 | 0.21 | 0.37 | -0.07 | -0.39 | -0.63 | 2.0E-02 | -0.64 |
| CHMP1B | -0.14 | -0.14 | -0.11 | 0.06 | 0.04 | 0.25 | 2.0E-02 | 0.25 |
| SOX12 | 0.21 | 0.00 | 0.16 | -0.10 | -0.12 | -0.20 | 2.0E-02 | -0.27 |
| SRSF10 | -0.09 | -0.13 | -0.10 | 0.09 | 0.01 | 0.19 | 2.0E-02 | 0.20 |
| ARL8B | -0.04 | -0.07 | -0.19 | 0.11 | 0.11 | 0.06 | 2.0E-02 | 0.19 |
| KIF18A | -0.23 | -0.14 | -0.02 | 0.12 | 0.15 | 0.07 | 2.0E-02 | 0.24 |
| GOPC | -0.14 | -0.19 | -0.20 | 0.01 | 0.15 | 0.30 | 2.0E-02 | 0.33 |
| FBRS | -0.20 | -0.02 | -0.17 | 0.07 | 0.18 | 0.11 | 2.0E-02 | 0.25 |
| AIP | -0.02 | -0.08 | -0.14 | 0.11 | 0.09 | 0.03 | 2.0E-02 | 0.16 |
| WDR1 | 0.13 | 0.15 | 0.00 | -0.05 | -0.14 | -0.12 | 2.0E-02 | -0.20 |
| MRPS26 | 0.14 | 0.07 | 0.05 | -0.17 | -0.04 | -0.07 | 2.0E-02 | -0.18 |
| TSEN54 | -0.13 | -0.25 | -0.19 | 0.11 | 0.04 | 0.33 | 2.0E-02 | 0.35 |
| RFC3 | -0.10 | -0.04 | -0.21 | 0.07 | 0.07 | 0.17 | 2.0E-02 | 0.22 |
| DPM3 | 0.19 | 0.08 | 0.12 | -0.28 | -0.09 | -0.06 | 2.1E-02 | -0.27 |
| CADM1 | -0.01 | 0.25 | 0.30 | -0.13 | -0.28 | -0.23 | 2.1E-02 | -0.39 |
| MIA2 | -0.05 | -0.08 | -0.25 | 0.09 | 0.15 | 0.10 | 2.1E-02 | 0.24 |
| MRPS27 | 0.16 | 0.13 | 0.04 | -0.20 | -0.03 | -0.14 | 2.1E-02 | -0.23 |
| MRM3 | 0.03 | 0.07 | 0.02 | -0.04 | -0.06 | -0.02 | 2.1E-02 | -0.08 |
| MRPL50 | 0.13 | 0.14 | 0.03 | -0.06 | -0.07 | -0.19 | 2.1E-02 | -0.21 |
| POU2F1 | 0.03 | 0.13 | 0.13 | -0.19 | -0.06 | -0.07 | 2.1E-02 | -0.20 |
| ENPP1 | 0.05 | 0.11 | 0.28 | -0.27 | -0.12 | -0.11 | 2.1E-02 | -0.31 |
| CDC37 | 0.06 | 0.02 | 0.14 | -0.05 | -0.07 | -0.11 | 2.1E-02 | -0.15 |
| MKI67 | 0.04 | 0.07 | 0.05 | -0.10 | -0.07 | 0.00 | 2.1E-02 | -0.11 |
| ADAM22 | 0.14 | 0.23 | 0.17 | -0.09 | -0.13 | -0.42 | 2.1E-02 | -0.39 |
| CCDC59 | -0.12 | -0.18 | -0.26 | 0.32 | 0.04 | 0.12 | 2.1E-02 | 0.35 |
| ORMDL2 | 0.07 | 0.10 | 0.10 | -0.01 | -0.11 | -0.18 | 2.1E-02 | -0.19 |
| INTS3 | 0.11 | 0.02 | 0.03 | -0.07 | -0.03 | -0.08 | 2.1E-02 | -0.11 |
| SRRM2 | 0.07 | 0.11 | 0.09 | -0.04 | -0.06 | -0.19 | 2.1E-02 | -0.18 |
| LRCH1 | -0.14 | -0.03 | -0.05 | 0.11 | 0.04 | 0.06 | 2.1E-02 | 0.14 |
| SLC7A6 | 0.03 | 0.22 | 0.28 | -0.28 | -0.29 | -0.06 | 2.1E-02 | -0.38 |
| INO80 | -0.06 | -0.11 | -0.12 | 0.15 | 0.00 | 0.11 | 2.1E-02 | 0.18 |
| SRFBP1 | 0.09 | 0.03 | 0.01 | -0.05 | -0.04 | -0.04 | 2.1E-02 | -0.09 |
| PFKFB3 | -0.26 | -0.25 | -0.30 | 0.21 | 0.45 | -0.01 | 2.1E-02 | 0.48 |
| CCZ1B | -0.07 | -0.08 | -0.17 | 0.19 | 0.03 | 0.09 | 2.1E-02 | 0.21 |
| CEP131 | 0.00 | 0.11 | 0.14 | -0.05 | -0.11 | -0.11 | 2.1E-02 | -0.17 |
| WDR74 | 0.04 | 0.03 | 0.08 | -0.05 | -0.01 | -0.09 | 2.1E-02 | -0.10 |
| VDAC1 | -0.04 | -0.06 | -0.13 | 0.11 | 0.02 | 0.08 | 2.1E-02 | 0.14 |
| ADK | -0.05 | -0.10 | -0.23 | 0.18 | 0.06 | 0.10 | 2.1E-02 | 0.24 |
| CGREF1 | 0.06 | 0.18 | 0.12 | -0.04 | -0.12 | -0.24 | 2.1E-02 | -0.25 |
| ATP13A1 | 0.03 | 0.04 | 0.14 | -0.05 | -0.07 | -0.11 | 2.1E-02 | -0.15 |
| TRIM24 | 0.07 | 0.06 | 0.07 | 0.00 | -0.12 | -0.11 | 2.1E-02 | -0.14 |
| SPTLC1 | -0.07 | -0.15 | -0.27 | 0.11 | 0.05 | 0.26 | 2.1E-02 | 0.31 |
| SPCS2 | 0.06 | 0.05 | 0.26 | -0.16 | -0.11 | -0.14 | 2.2E-02 | -0.26 |
| AKAP5 | -0.20 | -0.07 | -0.09 | 0.13 | 0.02 | 0.18 | 2.2E-02 | 0.23 |
| TRAF3IP2 | -0.44 | -0.64 | -0.35 | 0.54 | 0.52 | -0.08 | 2.2E-02 | 0.81 |

| | | | | | | | | |
|---|---|---|---|---|---|---|---|---|
| CHCHD2 | 0.10 | 0.06 | 0.12 | -0.15 | -0.15 | 0.00 | 2.2E-02 | -0.19 |
| CDC42BPA | -0.04 | -0.08 | -0.12 | 0.13 | 0.01 | 0.10 | 2.2E-02 | 0.16 |
| GFPT1 | -0.07 | -0.18 | -0.23 | 0.05 | 0.27 | 0.09 | 2.2E-02 | 0.30 |
| H2AC21 | 0.10 | 0.02 | 0.14 | -0.03 | -0.15 | -0.10 | 2.2E-02 | -0.18 |
| TOMM40 | -0.07 | -0.11 | -0.08 | 0.11 | -0.01 | 0.14 | 2.2E-02 | 0.17 |
| JTB | 0.09 | 0.12 | 0.13 | -0.04 | -0.08 | -0.25 | 2.2E-02 | -0.24 |
| PSMD9 | -0.08 | -0.35 | -0.15 | 0.13 | 0.28 | 0.09 | 2.2E-02 | 0.36 |
| COMMD7 | -0.22 | -0.06 | -0.09 | 0.05 | 0.20 | 0.08 | 2.2E-02 | 0.24 |
| ANKHD1 | -0.12 | -0.12 | -0.09 | 0.22 | 0.06 | 0.03 | 2.2E-02 | 0.21 |
| AMMECR1 | -0.12 | -0.03 | -0.07 | 0.07 | 0.02 | 0.12 | 2.2E-02 | 0.15 |
| BICDL2 | -0.22 | -0.16 | -0.03 | 0.11 | 0.06 | 0.21 | 2.2E-02 | 0.26 |
| POLR2A | 0.06 | 0.01 | 0.04 | -0.06 | -0.05 | -0.01 | 2.2E-02 | -0.08 |
| KDM2A | 0.11 | 0.03 | 0.04 | -0.10 | -0.03 | -0.05 | 2.2E-02 | -0.12 |
| SPCS1 | 0.14 | 0.20 | 0.18 | -0.18 | -0.04 | -0.38 | 2.2E-02 | -0.37 |
| RPL36AL | -0.11 | -0.21 | -0.11 | 0.24 | 0.02 | 0.11 | 2.2E-02 | 0.27 |
| NUCKS1 | 0.34 | 0.24 | 0.04 | -0.41 | -0.10 | -0.25 | 2.2E-02 | -0.46 |
| CCDC61 | 0.04 | 0.11 | 0.37 | -0.15 | -0.24 | -0.24 | 2.2E-02 | -0.38 |
| TRPS1 | 0.03 | 0.08 | 0.03 | -0.03 | -0.10 | -0.03 | 2.2E-02 | -0.10 |
| TNKS1BP1 | -0.07 | -0.01 | -0.10 | 0.07 | 0.02 | 0.09 | 2.2E-02 | 0.12 |
| ZNF318 | -0.12 | -0.02 | -0.12 | 0.08 | 0.14 | 0.02 | 2.2E-02 | 0.17 |
| RNMT | 0.15 | 0.17 | 0.15 | -0.13 | -0.05 | -0.36 | 2.2E-02 | -0.33 |
| RPN2 | 0.03 | 0.08 | 0.21 | -0.15 | -0.07 | -0.14 | 2.2E-02 | -0.23 |
| C1GALT1 | -0.09 | -0.28 | -0.14 | 0.21 | 0.02 | 0.22 | 2.2E-02 | 0.32 |
| ZNF202 | -0.13 | -0.10 | -0.01 | 0.10 | 0.03 | 0.10 | 2.2E-02 | 0.16 |
| CAP1 | 0.05 | 0.14 | 0.01 | -0.06 | -0.06 | -0.10 | 2.2E-02 | -0.14 |
| RLF | 0.04 | 0.07 | 0.07 | -0.13 | -0.03 | -0.03 | 2.2E-02 | -0.12 |
| CETN3 | 0.08 | 0.09 | 0.06 | -0.17 | -0.06 | -0.02 | 2.2E-02 | -0.16 |
| AATF | -0.01 | -0.11 | -0.12 | 0.05 | 0.04 | 0.13 | 2.2E-02 | 0.16 |
| TMEM120A | 0.08 | 0.05 | 0.12 | -0.14 | -0.13 | 0.00 | 2.2E-02 | -0.17 |
| TMTC3 | 0.06 | 0.07 | 0.14 | -0.07 | -0.04 | -0.18 | 2.2E-02 | -0.19 |
| C5orf24 | -0.06 | -0.14 | -0.17 | 0.21 | 0.02 | 0.10 | 2.2E-02 | 0.23 |
| MDH1 | 0.29 | 0.33 | -0.01 | -0.25 | -0.13 | -0.37 | 2.2E-02 | -0.45 |
| CENPB | -0.05 | -0.12 | -0.12 | 0.15 | 0.00 | 0.12 | 2.2E-02 | 0.19 |
| GNL3L | 0.06 | 0.06 | 0.17 | -0.13 | -0.03 | -0.17 | 2.3E-02 | -0.20 |
| ENO1 | 0.22 | 0.26 | -0.01 | -0.15 | -0.13 | -0.26 | 2.3E-02 | -0.34 |
| SLC25A3 | 0.02 | 0.15 | 0.04 | -0.09 | -0.07 | -0.07 | 2.3E-02 | -0.15 |
| CFAP61 | 0.08 | 0.17 | 0.24 | -0.13 | -0.08 | -0.36 | 2.3E-02 | -0.35 |
| ABCD3 | 0.05 | 0.06 | 0.10 | -0.05 | -0.02 | -0.15 | 2.3E-02 | -0.14 |
| CRKL | -0.03 | -0.14 | -0.24 | 0.15 | 0.18 | 0.04 | 2.3E-02 | 0.26 |
| SLC37A4 | 0.10 | 0.20 | 0.24 | -0.29 | -0.01 | -0.34 | 2.3E-02 | -0.39 |
| ALDH7A1 | -0.04 | -0.11 | -0.29 | 0.16 | 0.14 | 0.09 | 2.3E-02 | 0.28 |
| WASH4P | -0.09 | -0.24 | -0.11 | 0.10 | 0.04 | 0.24 | 2.3E-02 | 0.27 |
| ATF1 | 0.14 | -0.01 | 0.19 | -0.08 | -0.15 | -0.13 | 2.3E-02 | -0.23 |
| HMGCL | -0.02 | -0.11 | -0.11 | 0.08 | 0.13 | 0.01 | 2.3E-02 | 0.16 |
| GPT2 | -0.03 | -0.12 | -0.28 | 0.10 | 0.18 | 0.11 | 2.3E-02 | 0.27 |
| GRHL2 | 0.00 | -0.13 | -0.13 | 0.10 | 0.06 | 0.08 | 2.3E-02 | 0.17 |
| RPS5 | -0.06 | -0.16 | -0.11 | 0.20 | 0.08 | 0.02 | 2.3E-02 | 0.21 |
| SLC7A11 | -0.46 | -0.14 | -0.13 | 0.12 | 0.17 | 0.31 | 2.3E-02 | 0.44 |

| | | | | | | | | |
|---|---|---|---|---|---|---|---|---|
| ERAL1 | 0.14 | 0.08 | 0.04 | -0.15 | -0.01 | -0.12 | 2.3E-02 | -0.18 |
| FAM110C | -0.31 | -0.19 | -0.02 | 0.20 | 0.09 | 0.15 | 2.3E-02 | 0.32 |
| EWSR1 | 0.08 | 0.02 | 0.07 | -0.09 | 0.00 | -0.09 | 2.3E-02 | -0.12 |
| UBL3 | -0.01 | 0.18 | 0.29 | -0.16 | -0.15 | -0.24 | 2.3E-02 | -0.33 |
| TNK1 | 0.08 | 0.03 | 0.14 | -0.15 | -0.11 | -0.02 | 2.3E-02 | -0.18 |
| EXOSC2 | 0.06 | 0.08 | 0.23 | -0.05 | -0.15 | -0.22 | 2.3E-02 | -0.26 |
| SOX13 | 0.09 | 0.08 | 0.07 | -0.11 | 0.00 | -0.16 | 2.3E-02 | -0.17 |
| ALG1 | -0.04 | -0.09 | -0.15 | 0.01 | 0.12 | 0.14 | 2.3E-02 | 0.18 |
| IQGAP2 | -0.08 | -0.06 | 0.00 | 0.04 | 0.04 | 0.05 | 2.3E-02 | 0.09 |
| MTCH1 | -0.01 | -0.02 | -0.03 | 0.03 | 0.00 | 0.03 | 2.3E-02 | 0.04 |
| SLC7A5 | -0.43 | -0.24 | -0.10 | 0.12 | 0.13 | 0.39 | 2.3E-02 | 0.47 |
| CLU | -0.06 | -0.05 | -0.06 | 0.11 | 0.05 | 0.01 | 2.3E-02 | 0.11 |
| SLMAP | -0.01 | 0.15 | 0.14 | -0.08 | -0.12 | -0.10 | 2.3E-02 | -0.19 |
| LMNA | -0.05 | -0.09 | -0.05 | 0.05 | 0.01 | 0.11 | 2.3E-02 | 0.12 |
| NUP62 | 0.20 | 0.16 | 0.01 | -0.11 | -0.08 | -0.22 | 2.3E-02 | -0.26 |
| CCDC167 | 0.06 | 0.12 | 0.11 | -0.04 | -0.06 | -0.21 | 2.3E-02 | -0.20 |
| EMC8 | 0.09 | 0.11 | 0.14 | -0.13 | -0.01 | -0.24 | 2.3E-02 | -0.24 |
| PLCB3 | 0.01 | 0.08 | 0.11 | -0.03 | -0.06 | -0.12 | 2.4E-02 | -0.14 |
| RPS14 | -0.08 | -0.13 | -0.08 | 0.18 | 0.02 | 0.06 | 2.4E-02 | 0.18 |
| XAF1 | 0.02 | 0.41 | 0.27 | -0.23 | -0.18 | -0.49 | 2.4E-02 | -0.53 |
| BRD3 | 0.04 | 0.04 | 0.09 | -0.02 | -0.12 | -0.04 | 2.4E-02 | -0.12 |
| APOL2 | -0.12 | -0.20 | -0.01 | 0.11 | 0.13 | 0.06 | 2.4E-02 | 0.21 |
| TRAPPC12 | -0.01 | -0.12 | -0.10 | 0.03 | 0.12 | 0.08 | 2.4E-02 | 0.15 |
| RSBN1 | 0.06 | 0.09 | 0.03 | -0.05 | -0.02 | -0.13 | 2.4E-02 | -0.13 |
| ARMC10 | 0.05 | 0.05 | 0.18 | -0.03 | -0.13 | -0.15 | 2.4E-02 | -0.20 |
| CMAS | -0.05 | -0.07 | -0.10 | 0.12 | 0.00 | 0.09 | 2.4E-02 | 0.14 |
| ABCE1 | 0.11 | 0.16 | 0.05 | -0.10 | -0.03 | -0.23 | 2.4E-02 | -0.23 |
| ACTR8 | -0.11 | -0.14 | -0.06 | 0.07 | 0.02 | 0.19 | 2.4E-02 | 0.20 |
| EHF | -0.04 | -0.10 | -0.11 | 0.00 | 0.11 | 0.12 | 2.4E-02 | 0.16 |
| SHPRH | -0.10 | 0.00 | -0.08 | 0.06 | 0.03 | 0.08 | 2.4E-02 | 0.12 |
| OBSL1 | 0.13 | 0.03 | 0.09 | -0.12 | -0.01 | -0.14 | 2.4E-02 | -0.17 |
| CHAF1B | 0.10 | 0.07 | 0.03 | -0.14 | -0.03 | -0.05 | 2.4E-02 | -0.14 |
| SLC1A4 | -0.03 | 0.25 | 0.40 | -0.26 | -0.30 | -0.21 | 2.4E-02 | -0.46 |
| TOE1 | 0.04 | 0.06 | 0.17 | -0.12 | -0.03 | -0.15 | 2.4E-02 | -0.19 |
| ZNF143 | -0.11 | -0.03 | -0.10 | 0.04 | 0.15 | 0.04 | 2.4E-02 | 0.16 |
| CKAP5 | 0.04 | 0.04 | 0.04 | -0.03 | -0.01 | -0.09 | 2.4E-02 | -0.08 |
| NUBPL | -0.06 | -0.06 | -0.25 | 0.15 | 0.07 | 0.11 | 2.4E-02 | 0.24 |
| FLAD1 | -0.15 | -0.41 | -0.08 | 0.22 | 0.10 | 0.21 | 2.4E-02 | 0.39 |
| LRP10 | 0.07 | 0.10 | 0.28 | -0.12 | -0.10 | -0.31 | 2.4E-02 | -0.33 |
| POLR1D | 0.15 | 0.17 | 0.02 | -0.14 | -0.21 | -0.03 | 2.4E-02 | -0.24 |
| RHBDF1 | 0.03 | 0.02 | 0.07 | -0.01 | -0.06 | -0.06 | 2.4E-02 | -0.09 |
| UQCR10 | -0.11 | -0.09 | -0.32 | 0.15 | 0.05 | 0.25 | 2.4E-02 | 0.32 |
| TMUB1 | 0.03 | 0.07 | 0.17 | -0.04 | -0.09 | -0.17 | 2.4E-02 | -0.19 |
| LIN7A | 0.04 | 0.08 | 0.19 | -0.19 | -0.03 | -0.13 | 2.4E-02 | -0.22 |
| MPC2 | 0.02 | 0.24 | 0.10 | -0.10 | -0.09 | -0.21 | 2.4E-02 | -0.26 |
| SMC3 | -0.02 | -0.06 | -0.09 | 0.06 | 0.01 | 0.09 | 2.4E-02 | 0.11 |
| SPIN4 | -0.19 | -0.20 | -0.13 | 0.16 | -0.01 | 0.30 | 2.4E-02 | 0.32 |
| TOR1A | -0.06 | -0.19 | -0.20 | 0.21 | 0.00 | 0.18 | 2.4E-02 | 0.28 |

| | | | | | | | | |
|---|---|---|---|---|---|---|---|---|
| **AQR** | 0.09 | 0.03 | 0.04 | -0.05 | -0.02 | -0.10 | 2.4E-02 | -0.11 |
| **MPV17L2** | 0.17 | 0.17 | 0.00 | -0.09 | -0.20 | -0.09 | 2.4E-02 | -0.24 |
| **CPT2** | -0.02 | -0.10 | -0.25 | 0.11 | 0.10 | 0.12 | 2.4E-02 | 0.24 |
| **PHF20** | 0.13 | 0.15 | 0.11 | -0.20 | 0.01 | -0.25 | 2.4E-02 | -0.28 |
| **RYBP** | 0.07 | 0.13 | 0.02 | -0.12 | -0.09 | -0.03 | 2.4E-02 | -0.15 |
| **ZNF665** | 0.20 | 0.22 | 0.16 | -0.12 | -0.11 | -0.47 | 2.5E-02 | -0.43 |
| **RICTOR** | -0.25 | -0.11 | -0.05 | 0.18 | 0.04 | 0.15 | 2.5E-02 | 0.26 |
| **TPM4** | 0.15 | 0.13 | 0.13 | -0.13 | -0.03 | -0.31 | 2.5E-02 | -0.29 |
| **MICAL3** | 0.03 | 0.10 | 0.11 | -0.07 | -0.03 | -0.16 | 2.5E-02 | -0.17 |
| **UQCC6** | -0.03 | 0.25 | 0.42 | -0.33 | -0.18 | -0.31 | 2.5E-02 | -0.49 |
| **TAGLN2** | 0.19 | 0.23 | 0.01 | -0.17 | -0.06 | -0.26 | 2.5E-02 | -0.31 |
| **HECTD3** | -0.16 | -0.20 | 0.00 | 0.14 | 0.11 | 0.07 | 2.5E-02 | 0.23 |
| **PIK3R4** | -0.29 | -0.17 | -0.08 | 0.20 | 0.01 | 0.25 | 2.5E-02 | 0.33 |
| **SMC1A** | -0.02 | -0.05 | -0.05 | 0.00 | 0.04 | 0.08 | 2.5E-02 | 0.08 |
| **G3BP1** | -0.23 | -0.15 | -0.05 | 0.22 | 0.03 | 0.12 | 2.5E-02 | 0.27 |
| **XPOT** | -0.28 | -0.09 | -0.12 | 0.17 | 0.02 | 0.23 | 2.5E-02 | 0.30 |
| **ATP1A1** | -0.01 | 0.13 | 0.24 | -0.17 | -0.14 | -0.10 | 2.5E-02 | -0.26 |
| **CPNE3** | -0.14 | -0.20 | -0.01 | 0.07 | 0.09 | 0.16 | 2.5E-02 | 0.22 |
| **RNF185** | -0.22 | -0.04 | -0.08 | 0.14 | 0.12 | 0.05 | 2.5E-02 | 0.21 |
| **TIMM8B** | 0.08 | 0.03 | 0.01 | -0.04 | -0.06 | -0.03 | 2.5E-02 | -0.08 |
| **TFB1M** | 0.07 | 0.02 | 0.08 | -0.11 | -0.01 | -0.05 | 2.5E-02 | -0.12 |
| **HAUS7** | -0.33 | -0.15 | -0.21 | 0.16 | 0.02 | 0.39 | 2.5E-02 | 0.42 |
| **BUB3** | 0.13 | 0.15 | 0.02 | -0.06 | -0.07 | -0.19 | 2.5E-02 | -0.20 |
| **FAM98C** | -0.17 | -0.28 | -0.12 | 0.33 | 0.03 | 0.13 | 2.5E-02 | 0.36 |
| **DDX31** | -0.02 | -0.07 | -0.08 | 0.07 | 0.01 | 0.09 | 2.5E-02 | 0.11 |
| **DOT1L** | 0.04 | 0.08 | 0.08 | -0.02 | -0.05 | -0.14 | 2.5E-02 | -0.14 |
| **SPAG1** | -0.02 | -0.10 | -0.19 | 0.04 | 0.12 | 0.12 | 2.5E-02 | 0.19 |
| **MUS81** | -0.03 | -0.04 | -0.10 | 0.03 | 0.10 | 0.03 | 2.5E-02 | 0.11 |
| **IPO5** | 0.03 | 0.10 | 0.05 | -0.01 | -0.06 | -0.11 | 2.5E-02 | -0.12 |
| **PIGBOS1** | 0.17 | 0.32 | 0.18 | -0.42 | -0.01 | -0.41 | 2.5E-02 | -0.50 |
| **CLSTN1** | 0.11 | 0.19 | 0.20 | 0.00 | -0.29 | -0.29 | 2.5E-02 | -0.36 |
| **TNC** | -0.04 | -0.06 | -0.10 | 0.11 | 0.00 | 0.07 | 2.5E-02 | 0.13 |
| **EPS8L2** | -0.02 | -0.02 | -0.08 | 0.04 | 0.03 | 0.05 | 2.5E-02 | 0.08 |
| **IARS1** | -0.08 | -0.08 | -0.04 | 0.09 | 0.11 | -0.01 | 2.5E-02 | 0.13 |
| **MRPS11** | 0.17 | 0.10 | 0.03 | -0.13 | -0.02 | -0.18 | 2.5E-02 | -0.21 |
| **ZNG1F** | -0.17 | -0.14 | -0.28 | 0.01 | 0.14 | 0.34 | 2.6E-02 | 0.36 |
| **BOLA2** | -0.01 | -0.06 | -0.10 | 0.07 | 0.02 | 0.08 | 2.6E-02 | 0.11 |
| **S100A10** | -0.08 | -0.04 | -0.25 | 0.09 | 0.12 | 0.11 | 2.6E-02 | 0.23 |
| **ARID5B** | 0.02 | 0.12 | 0.06 | -0.03 | -0.12 | -0.06 | 2.6E-02 | -0.14 |
| **POLD3** | -0.14 | -0.06 | -0.23 | 0.01 | 0.16 | 0.21 | 2.6E-02 | 0.27 |
| **GTF2A1L** | -0.33 | -0.22 | -0.18 | 0.22 | -0.02 | 0.39 | 2.6E-02 | 0.44 |
| **MYO18A** | 0.02 | 0.01 | 0.08 | -0.03 | -0.04 | -0.05 | 2.6E-02 | -0.08 |
| **OPA1** | 0.10 | 0.06 | 0.00 | -0.05 | -0.05 | -0.07 | 2.6E-02 | -0.11 |
| **NCAPD2** | -0.10 | -0.12 | -0.05 | 0.00 | 0.15 | 0.09 | 2.6E-02 | 0.17 |
| **PPA1** | 0.23 | 0.18 | 0.01 | -0.16 | -0.06 | -0.26 | 2.6E-02 | -0.30 |
| **UTP18** | 0.06 | 0.03 | 0.03 | -0.03 | -0.01 | -0.09 | 2.6E-02 | -0.08 |
| **SCAMP1** | 0.06 | 0.15 | 0.33 | -0.35 | -0.22 | -0.07 | 2.6E-02 | -0.39 |
| **SKIC3** | -0.05 | -0.21 | -0.16 | 0.20 | 0.01 | 0.15 | 2.6E-02 | 0.26 |

| | | | | | | | | |
|---|---|---|---|---|---|---|---|---|
| EIF4E2 | -0.28 | -0.01 | -0.20 | 0.13 | 0.09 | 0.21 | 2.6E-02 | 0.30 |
| IGHMBP2 | -0.15 | -0.05 | -0.03 | 0.10 | 0.09 | 0.02 | 2.6E-02 | 0.15 |
| SQOR | 0.05 | 0.07 | 0.08 | -0.15 | -0.06 | -0.01 | 2.6E-02 | -0.14 |
| STAU1 | 0.06 | 0.02 | 0.05 | -0.03 | -0.01 | -0.09 | 2.6E-02 | -0.09 |
| SLC25A11 | 0.05 | 0.09 | 0.01 | -0.03 | -0.04 | -0.10 | 2.6E-02 | -0.11 |
| CUL3 | 0.15 | 0.17 | -0.01 | -0.14 | -0.06 | -0.14 | 2.6E-02 | -0.21 |
| SDF4 | -0.08 | -0.01 | -0.04 | 0.05 | 0.06 | 0.01 | 2.6E-02 | 0.08 |
| CHMP6 | 0.03 | 0.21 | 0.07 | -0.16 | -0.06 | -0.12 | 2.6E-02 | -0.21 |
| SPINT2 | 0.08 | 0.15 | 0.28 | -0.36 | -0.04 | -0.21 | 2.6E-02 | -0.37 |
| MDN1 | 0.09 | 0.05 | 0.10 | -0.02 | -0.06 | -0.17 | 2.6E-02 | -0.16 |
| SZRD1 | 0.17 | 0.25 | 0.23 | -0.36 | -0.46 | 0.00 | 2.6E-02 | -0.49 |
| HRNR | -1.65 | -1.39 | -0.48 | 1.20 | 0.09 | 0.28 | 2.6E-02 | 1.70 |
| CEP63 | -0.15 | -0.09 | -0.08 | 0.02 | 0.07 | 0.21 | 2.6E-02 | 0.21 |
| VPS13A | -0.03 | -0.12 | -0.20 | 0.08 | 0.05 | 0.18 | 2.6E-02 | 0.22 |
| G6PC3 | 0.05 | 0.15 | 0.27 | -0.06 | -0.17 | -0.32 | 2.7E-02 | -0.34 |
| PIK3C2A | -0.11 | -0.09 | 0.00 | 0.09 | 0.07 | 0.03 | 2.7E-02 | 0.13 |
| COG7 | -0.30 | -0.30 | 0.01 | 0.14 | 0.17 | 0.20 | 2.7E-02 | 0.36 |
| CHD2 | 0.00 | -0.04 | -0.05 | 0.05 | 0.02 | 0.02 | 2.7E-02 | 0.06 |
| NXF1 | 0.04 | 0.03 | 0.04 | -0.02 | -0.01 | -0.08 | 2.7E-02 | -0.08 |
| KLC1 | -0.02 | -0.09 | -0.16 | 0.08 | 0.03 | 0.14 | 2.7E-02 | 0.18 |
| SURF4 | 0.09 | 0.06 | 0.21 | -0.22 | -0.15 | -0.02 | 2.7E-02 | -0.25 |
| SLFN11 | -0.04 | -0.10 | -0.12 | 0.00 | 0.15 | 0.11 | 2.7E-02 | 0.17 |
| LARS2 | 0.00 | -0.12 | -0.17 | 0.06 | 0.12 | 0.09 | 2.7E-02 | 0.19 |
| NOPCHAP1 | 0.18 | 0.02 | 0.10 | -0.15 | -0.03 | -0.15 | 2.7E-02 | -0.21 |
| VPS11 | -0.17 | -0.13 | -0.13 | 0.27 | 0.01 | 0.09 | 2.7E-02 | 0.26 |
| EIF1B | -0.06 | -0.12 | -0.22 | 0.19 | 0.01 | 0.16 | 2.7E-02 | 0.25 |
| HARS1 | 0.24 | 0.24 | -0.02 | -0.25 | -0.09 | -0.20 | 2.7E-02 | -0.34 |
| NOP53 | 0.18 | 0.12 | 0.06 | -0.15 | -0.01 | -0.24 | 2.7E-02 | -0.25 |
| SPC24 | -0.06 | -0.13 | -0.07 | 0.15 | 0.00 | 0.09 | 2.7E-02 | 0.16 |
| SORT1 | 0.05 | 0.17 | 0.32 | -0.24 | -0.06 | -0.35 | 2.7E-02 | -0.40 |
| EME1 | -0.14 | -0.12 | 0.00 | 0.07 | 0.13 | 0.04 | 2.7E-02 | 0.17 |
| SIRT5 | -0.11 | -0.15 | -0.01 | 0.09 | 0.13 | 0.03 | 2.7E-02 | 0.17 |
| PRIM2 | -0.11 | -0.22 | -0.16 | 0.07 | 0.04 | 0.31 | 2.7E-02 | 0.30 |
| RAB14 | 0.01 | 0.07 | 0.15 | -0.06 | -0.06 | -0.12 | 2.7E-02 | -0.16 |
| TOMM7 | -0.10 | -0.09 | 0.01 | 0.07 | 0.06 | 0.05 | 2.7E-02 | 0.12 |
| VAMP8 | 0.00 | 0.14 | 0.37 | -0.22 | -0.20 | -0.19 | 2.7E-02 | -0.37 |
| EDC4 | -0.13 | -0.06 | -0.04 | 0.05 | 0.14 | 0.03 | 2.7E-02 | 0.15 |
| GLG1 | 0.07 | 0.05 | 0.10 | -0.15 | -0.09 | 0.00 | 2.7E-02 | -0.15 |
| SUPT20H | -0.22 | -0.08 | -0.25 | 0.28 | -0.01 | 0.19 | 2.7E-02 | 0.34 |
| MRPL21 | 0.17 | 0.21 | 0.00 | -0.12 | -0.07 | -0.24 | 2.7E-02 | -0.27 |
| SNRPG | 0.03 | 0.08 | 0.00 | -0.03 | -0.05 | -0.05 | 2.7E-02 | -0.08 |
| TMEM87B | 0.07 | 0.21 | 0.37 | -0.44 | -0.05 | -0.34 | 2.7E-02 | -0.49 |
| CDH4 | -0.24 | -0.47 | -0.12 | 0.17 | 0.07 | 0.42 | 2.7E-02 | 0.50 |
| INTS8 | -0.03 | -0.03 | -0.06 | 0.02 | 0.08 | 0.01 | 2.7E-02 | 0.08 |
| B3GNT9 | 0.00 | 0.11 | 0.14 | -0.04 | -0.12 | -0.11 | 2.8E-02 | -0.17 |
| RDH13 | 0.01 | 0.05 | 0.14 | -0.09 | -0.04 | -0.09 | 2.8E-02 | -0.14 |
| RIMOC1 | -0.21 | -0.23 | -0.24 | 0.30 | 0.32 | -0.06 | 2.8E-02 | 0.41 |
| TBC1D4 | 0.20 | 0.24 | 0.36 | -0.28 | -0.09 | -0.68 | 2.8E-02 | -0.61 |

| | | | | | | | | |
|---|---|---|---|---|---|---|---|---|
| **SEC11A** | 0.05 | 0.03 | 0.21 | -0.10 | -0.07 | -0.14 | 2.8E-02 | -0.20 |
| **TRIP4** | -0.11 | -0.14 | -0.11 | 0.23 | 0.01 | 0.08 | 2.8E-02 | 0.23 |
| **ARL4A** | 0.15 | 0.17 | 0.10 | -0.13 | -0.03 | -0.32 | 2.8E-02 | -0.30 |
| **SEC22B** | 0.02 | 0.01 | 0.06 | -0.02 | -0.04 | -0.02 | 2.8E-02 | -0.06 |
| **TOP1MT** | 0.06 | 0.17 | 0.37 | -0.21 | -0.10 | -0.44 | 2.8E-02 | -0.45 |
| **PSMG2** | 0.25 | 0.20 | 0.02 | -0.29 | -0.23 | -0.04 | 2.8E-02 | -0.35 |
| **TMEM33** | 0.02 | 0.10 | 0.11 | -0.16 | -0.06 | -0.03 | 2.8E-02 | -0.16 |
| **PWP2** | -0.04 | -0.09 | -0.09 | 0.05 | 0.01 | 0.14 | 2.8E-02 | 0.14 |
| **C11orf98** | 0.12 | 0.13 | 0.09 | -0.05 | -0.06 | -0.26 | 2.8E-02 | -0.23 |
| **PEX19** | 0.09 | 0.13 | 0.14 | -0.04 | -0.09 | -0.28 | 2.8E-02 | -0.26 |
| **LUM** | -0.03 | -0.22 | -0.29 | 0.20 | 0.22 | 0.03 | 2.8E-02 | 0.33 |
| **SEPTIN8** | 0.08 | 0.18 | 0.01 | -0.14 | -0.09 | -0.06 | 2.8E-02 | -0.18 |
| **MAP2K7** | -0.19 | -0.11 | -0.14 | -0.02 | 0.19 | 0.22 | 2.8E-02 | 0.27 |
| **STEEP1** | 0.10 | 0.11 | 0.04 | -0.04 | -0.05 | -0.19 | 2.8E-02 | -0.18 |
| **TWNK** | 0.09 | 0.06 | 0.02 | -0.05 | -0.02 | -0.12 | 2.8E-02 | -0.12 |
| **SLC30A4** | 0.06 | 0.21 | 0.38 | -0.49 | -0.10 | -0.24 | 2.8E-02 | -0.49 |
| **RIF1** | 0.15 | 0.12 | 0.15 | -0.32 | -0.14 | -0.02 | 2.8E-02 | -0.30 |
| **LMAN1** | 0.01 | 0.11 | 0.29 | -0.15 | -0.11 | -0.22 | 2.8E-02 | -0.29 |
| **MACO1** | 0.06 | 0.09 | 0.14 | -0.01 | -0.11 | -0.20 | 2.8E-02 | -0.20 |
| **ASNS** | -0.32 | -0.19 | -0.33 | 0.03 | 0.13 | 0.50 | 2.8E-02 | 0.50 |
| **BCOR** | 0.10 | 0.06 | 0.00 | -0.04 | -0.04 | -0.09 | 2.8E-02 | -0.11 |
| **PBDC1** | -0.11 | -0.05 | -0.13 | 0.07 | 0.01 | 0.19 | 2.8E-02 | 0.19 |
| **SUCLG2** | 0.01 | -0.13 | -0.21 | 0.10 | 0.11 | 0.09 | 2.8E-02 | 0.21 |
| **FLNB** | -0.10 | -0.06 | -0.03 | 0.11 | 0.05 | 0.01 | 2.8E-02 | 0.12 |
| **COPS2** | -0.09 | -0.06 | -0.01 | 0.08 | 0.01 | 0.05 | 2.8E-02 | 0.10 |
| **ABCA3** | 0.02 | 0.13 | 0.25 | -0.05 | -0.18 | -0.23 | 2.8E-02 | -0.28 |
| **FAIM** | -0.25 | -0.34 | -0.22 | 0.34 | 0.36 | -0.06 | 2.9E-02 | 0.48 |
| **TOP1** | -0.01 | -0.05 | -0.09 | 0.07 | 0.01 | 0.07 | 2.9E-02 | 0.10 |
| **RAD51AP1** | 0.10 | 0.12 | 0.15 | -0.27 | -0.16 | 0.00 | 2.9E-02 | -0.27 |
| **FYCO1** | -0.27 | -0.34 | -0.03 | 0.09 | 0.12 | 0.32 | 2.9E-02 | 0.39 |
| **NME6** | -0.12 | -0.07 | -0.26 | 0.16 | 0.21 | 0.01 | 2.9E-02 | 0.28 |
| **SF3B2** | 0.07 | 0.01 | 0.06 | -0.01 | -0.07 | -0.06 | 2.9E-02 | -0.09 |
| **IER3IP1** | 0.02 | 0.15 | 0.29 | -0.29 | -0.08 | -0.16 | 2.9E-02 | -0.33 |
| **EXOC7** | -0.09 | -0.11 | -0.05 | 0.17 | 0.03 | 0.04 | 2.9E-02 | 0.16 |
| **KLF12** | 0.23 | 0.10 | 0.07 | -0.27 | -0.03 | -0.15 | 2.9E-02 | -0.28 |
| **PRPH** | -0.11 | -0.14 | -0.21 | 0.21 | -0.03 | 0.22 | 2.9E-02 | 0.29 |
| **MICALL1** | -0.09 | -0.19 | -0.19 | 0.06 | 0.05 | 0.30 | 2.9E-02 | 0.29 |
| **UBE2J1** | 0.01 | 0.14 | 0.31 | -0.12 | -0.14 | -0.27 | 2.9E-02 | -0.33 |
| **MUTYH** | -0.28 | -0.26 | -0.04 | 0.06 | 0.12 | 0.31 | 2.9E-02 | 0.36 |
| **NUDC** | 0.08 | 0.04 | 0.03 | -0.09 | 0.00 | -0.08 | 2.9E-02 | -0.11 |
| **BRD2** | 0.07 | 0.06 | 0.05 | -0.02 | -0.03 | -0.14 | 2.9E-02 | -0.13 |
| **GMPS** | -0.05 | -0.06 | -0.08 | -0.01 | 0.11 | 0.07 | 2.9E-02 | 0.12 |
| **CYBC1** | 0.06 | 0.09 | 0.34 | -0.11 | -0.16 | -0.31 | 2.9E-02 | -0.35 |
| **DYNC1H1** | -0.01 | -0.06 | -0.11 | 0.03 | 0.09 | 0.04 | 2.9E-02 | 0.11 |
| **CDK2** | 0.13 | 0.13 | 0.08 | -0.27 | -0.08 | -0.03 | 2.9E-02 | -0.24 |
| **GLI3** | 0.04 | 0.21 | 0.23 | -0.11 | -0.10 | -0.36 | 2.9E-02 | -0.35 |
| **ELMO2** | -0.21 | -0.11 | -0.08 | 0.04 | 0.07 | 0.24 | 2.9E-02 | 0.25 |
| **MAFG** | -0.02 | -0.18 | -0.05 | 0.10 | 0.07 | 0.06 | 2.9E-02 | 0.16 |

| | | | | | | | | |
|---|---|---|---|---|---|---|---|---|
| TAF2 | -0.06 | -0.08 | -0.12 | 0.01 | 0.07 | 0.17 | 2.9E-02 | 0.17 |
| PRRC2C | -0.01 | 0.14 | 0.09 | -0.09 | -0.05 | -0.09 | 2.9E-02 | -0.15 |
| COX6A1 | 0.10 | 0.12 | 0.18 | -0.16 | -0.01 | -0.29 | 2.9E-02 | -0.28 |
| TAF6L | -0.18 | -0.09 | -0.04 | 0.18 | 0.04 | 0.06 | 2.9E-02 | 0.20 |
| ARFIP1 | -0.15 | -0.23 | -0.03 | 0.04 | 0.11 | 0.21 | 2.9E-02 | 0.25 |
| OSBP2 | 0.29 | 0.41 | 0.19 | -0.29 | -0.13 | -0.79 | 3.0E-02 | -0.70 |
| ATAD1 | 0.00 | -0.03 | -0.08 | 0.04 | 0.05 | 0.02 | 3.0E-02 | 0.07 |
| PTP4A2 | 0.31 | 0.31 | 0.07 | -0.42 | -0.02 | -0.44 | 3.0E-02 | -0.52 |
| SLC30A9 | -0.03 | 0.16 | 0.26 | -0.13 | -0.19 | -0.14 | 3.0E-02 | -0.29 |
| FASTKD5 | -0.02 | -0.03 | -0.06 | 0.00 | 0.05 | 0.06 | 3.0E-02 | 0.07 |
| SVIL | 0.00 | 0.11 | 0.08 | -0.10 | -0.02 | -0.08 | 3.0E-02 | -0.13 |
| TMEM237 | 0.03 | 0.20 | 0.20 | -0.31 | -0.06 | -0.14 | 3.0E-02 | -0.31 |
| LIN54 | -0.20 | -0.02 | -0.12 | 0.06 | 0.07 | 0.18 | 3.0E-02 | 0.21 |
| ERGIC2 | 0.05 | 0.02 | 0.01 | -0.02 | -0.04 | -0.02 | 3.0E-02 | -0.05 |
| CHEK1 | 0.13 | 0.18 | 0.05 | -0.27 | -0.04 | -0.10 | 3.0E-02 | -0.25 |
| NDUFB5 | 0.06 | 0.04 | 0.01 | -0.02 | -0.03 | -0.07 | 3.0E-02 | -0.07 |
| USP14 | 0.21 | 0.19 | 0.00 | -0.08 | -0.26 | -0.13 | 3.0E-02 | -0.29 |
| RTKN | -0.41 | -0.37 | -0.04 | 0.28 | 0.04 | 0.34 | 3.0E-02 | 0.49 |
| MRPS15 | 0.15 | 0.12 | 0.02 | -0.19 | -0.03 | -0.10 | 3.0E-02 | -0.20 |
| MTDH | 0.03 | 0.09 | 0.07 | -0.02 | -0.04 | -0.15 | 3.0E-02 | -0.14 |
| SPINT1 | 0.19 | 0.09 | 0.24 | -0.42 | -0.11 | -0.09 | 3.0E-02 | -0.38 |
| SEC24D | 0.13 | 0.22 | 0.14 | -0.19 | -0.01 | -0.38 | 3.0E-02 | -0.36 |
| TXLNG | -0.08 | -0.04 | 0.00 | 0.04 | 0.05 | 0.03 | 3.0E-02 | 0.08 |
| EPHB3 | 0.04 | 0.04 | 0.10 | 0.00 | -0.11 | -0.07 | 3.0E-02 | -0.12 |
| BCORL1 | 0.07 | 0.12 | 0.01 | -0.09 | -0.10 | -0.02 | 3.0E-02 | -0.14 |
| ZNF611 | 0.12 | 0.17 | 0.27 | -0.29 | 0.00 | -0.38 | 3.0E-02 | -0.41 |
| PACS1 | -0.33 | -0.06 | -0.21 | 0.19 | 0.02 | 0.29 | 3.0E-02 | 0.37 |
| AFG3L2 | 0.05 | 0.10 | 0.07 | 0.00 | -0.09 | -0.15 | 3.0E-02 | -0.15 |
| ATXN7L3B | -0.05 | -0.18 | -0.18 | 0.15 | 0.22 | -0.01 | 3.0E-02 | 0.26 |
| PRPS2 | -0.29 | -0.21 | -0.14 | 0.04 | 0.10 | 0.40 | 3.0E-02 | 0.39 |
| VPS16 | 0.01 | -0.11 | -0.10 | 0.09 | 0.06 | 0.04 | 3.0E-02 | 0.13 |
| ZC3H13 | 0.11 | 0.11 | 0.10 | -0.05 | -0.05 | -0.25 | 3.0E-02 | -0.22 |
| SGSM3 | -0.06 | -0.08 | -0.01 | 0.07 | 0.07 | 0.01 | 3.0E-02 | 0.10 |
| SLBP | -0.09 | -0.10 | -0.26 | 0.15 | 0.24 | 0.01 | 3.0E-02 | 0.28 |
| TMED10 | 0.05 | 0.06 | 0.08 | -0.06 | 0.00 | -0.14 | 3.0E-02 | -0.13 |
| PRPF38B | 0.16 | 0.16 | 0.07 | -0.14 | -0.01 | -0.28 | 3.0E-02 | -0.27 |
| ADH5 | -0.02 | -0.14 | -0.26 | 0.19 | 0.14 | 0.05 | 3.1E-02 | 0.27 |
| HNRNPH1 | 0.06 | 0.04 | 0.12 | -0.13 | 0.00 | -0.10 | 3.1E-02 | -0.15 |
| MRPL11 | 0.11 | 0.11 | -0.02 | -0.07 | -0.06 | -0.09 | 3.1E-02 | -0.14 |
| MTREX | 0.10 | 0.09 | 0.00 | -0.02 | -0.09 | -0.09 | 3.1E-02 | -0.13 |
| ETFA | -0.02 | -0.07 | -0.20 | 0.08 | 0.07 | 0.11 | 3.1E-02 | 0.18 |
| STAMBP | 0.07 | 0.14 | 0.09 | -0.04 | -0.06 | -0.23 | 3.1E-02 | -0.21 |
| NF1 | -0.12 | -0.02 | -0.07 | 0.12 | 0.03 | 0.04 | 3.1E-02 | 0.13 |
| ITGB1 | 0.04 | 0.09 | 0.25 | -0.17 | -0.21 | -0.04 | 3.1E-02 | -0.27 |
| ZDHHC13 | -0.15 | -0.10 | -0.21 | -0.03 | 0.19 | 0.24 | 3.1E-02 | 0.29 |
| SEPTIN7 | 0.26 | 0.25 | -0.05 | -0.20 | -0.13 | -0.21 | 3.1E-02 | -0.34 |
| ATG16L1 | -0.14 | -0.25 | -0.05 | 0.24 | 0.11 | 0.03 | 3.1E-02 | 0.28 |
| AKAP9 | -0.14 | -0.12 | 0.00 | 0.12 | 0.08 | 0.04 | 3.1E-02 | 0.16 |

| | | | | | | | | |
|---|---|---|---|---|---|---|---|---|
| BRMS1L | 0.02 | 0.05 | 0.01 | -0.03 | 0.00 | -0.04 | 3.1E-02 | -0.05 |
| CCDC93 | -0.01 | -0.11 | -0.12 | 0.13 | 0.03 | 0.05 | 3.1E-02 | 0.15 |
| ZBTB9 | -0.17 | -0.32 | -0.09 | 0.24 | -0.01 | 0.26 | 3.1E-02 | 0.36 |
| RSL24D1 | 0.07 | 0.09 | 0.03 | -0.06 | -0.01 | -0.14 | 3.1E-02 | -0.14 |
| TMEM181 | 0.03 | 0.03 | 0.05 | -0.05 | 0.00 | -0.07 | 3.1E-02 | -0.07 |
| SENP2 | -0.11 | -0.05 | -0.03 | 0.02 | 0.04 | 0.12 | 3.1E-02 | 0.13 |
| F11R | -0.02 | 0.16 | 0.33 | -0.20 | -0.23 | -0.13 | 3.1E-02 | -0.35 |
| CDC42SE2 | 0.14 | 0.22 | 0.30 | 0.01 | -0.42 | -0.42 | 3.1E-02 | -0.50 |
| TMEM201 | -0.07 | -0.22 | -0.07 | 0.10 | 0.02 | 0.19 | 3.1E-02 | 0.23 |
| GOLM2 | 0.00 | 0.05 | 0.09 | -0.03 | -0.04 | -0.08 | 3.1E-02 | -0.10 |
| HOXC10 | 0.07 | 0.08 | 0.04 | -0.14 | -0.04 | -0.02 | 3.1E-02 | -0.13 |
| ARHGAP11A | -0.14 | -0.22 | -0.06 | -0.01 | 0.17 | 0.21 | 3.1E-02 | 0.26 |
| HEXIM1 | -0.06 | -0.14 | -0.02 | 0.08 | 0.02 | 0.11 | 3.1E-02 | 0.15 |
| SARNP | 0.05 | 0.00 | 0.04 | -0.05 | -0.02 | -0.02 | 3.1E-02 | -0.06 |
| CLUH | -0.04 | -0.06 | -0.05 | 0.09 | 0.07 | -0.01 | 3.1E-02 | 0.10 |
| RNF169 | 0.05 | 0.10 | 0.11 | -0.16 | -0.13 | 0.01 | 3.1E-02 | -0.18 |
| FAM136A | 0.11 | 0.08 | 0.05 | -0.01 | -0.07 | -0.18 | 3.2E-02 | -0.17 |
| ABCF1 | 0.07 | 0.07 | 0.08 | -0.15 | 0.01 | -0.10 | 3.2E-02 | -0.15 |
| TMEM134 | 0.04 | 0.09 | 0.18 | -0.07 | -0.05 | -0.23 | 3.2E-02 | -0.22 |
| C6orf47 | 0.05 | 0.02 | 0.07 | -0.03 | -0.02 | -0.10 | 3.2E-02 | -0.10 |
| ACO2 | 0.00 | -0.08 | -0.16 | 0.09 | 0.09 | 0.04 | 3.2E-02 | 0.15 |
| PIP5K1A | -0.05 | -0.09 | -0.13 | 0.15 | -0.01 | 0.11 | 3.2E-02 | 0.18 |
| C2orf49 | 0.05 | 0.06 | 0.13 | 0.00 | -0.13 | -0.13 | 3.2E-02 | -0.17 |
| SRPK2 | 0.06 | 0.05 | 0.07 | -0.03 | -0.02 | -0.13 | 3.2E-02 | -0.12 |
| MED6 | 0.18 | 0.02 | 0.12 | -0.11 | -0.03 | -0.21 | 3.2E-02 | -0.23 |
| NDRG3 | 0.09 | 0.15 | 0.03 | -0.09 | -0.03 | -0.19 | 3.2E-02 | -0.19 |
| NAT14 | -0.21 | -0.26 | -0.21 | 0.04 | 0.08 | 0.44 | 3.2E-02 | 0.41 |
| SRP9 | 0.06 | 0.15 | 0.03 | -0.15 | -0.02 | -0.11 | 3.2E-02 | -0.17 |
| SCPEP1 | -0.16 | -0.15 | -0.09 | 0.26 | 0.07 | 0.01 | 3.2E-02 | 0.25 |
| SF3A1 | 0.10 | 0.02 | 0.01 | -0.03 | -0.05 | -0.07 | 3.2E-02 | -0.09 |
| BRIP1 | -0.05 | -0.45 | -0.21 | 0.23 | 0.07 | 0.28 | 3.2E-02 | 0.43 |
| TBP | -0.04 | -0.06 | -0.14 | 0.01 | 0.08 | 0.13 | 3.2E-02 | 0.15 |
| CPNE1 | -0.08 | -0.06 | -0.03 | 0.04 | 0.10 | 0.01 | 3.2E-02 | 0.10 |
| PMPCA | -0.02 | -0.03 | -0.07 | 0.00 | 0.07 | 0.05 | 3.2E-02 | 0.08 |
| PRR14 | -0.12 | -0.08 | -0.03 | 0.00 | 0.08 | 0.14 | 3.2E-02 | 0.15 |
| ERBIN | -0.17 | -0.04 | -0.07 | 0.04 | 0.06 | 0.16 | 3.2E-02 | 0.18 |
| MLLT6 | 0.04 | 0.09 | 0.15 | -0.03 | -0.07 | -0.21 | 3.2E-02 | -0.20 |
| TCP1 | -0.15 | -0.10 | -0.08 | 0.10 | -0.01 | 0.21 | 3.2E-02 | 0.21 |
| RPS13 | -0.06 | -0.07 | -0.02 | 0.08 | 0.08 | -0.01 | 3.2E-02 | 0.10 |
| EAF1 | 0.20 | -0.02 | 0.15 | -0.09 | -0.10 | -0.18 | 3.2E-02 | -0.24 |
| CEP85 | -0.18 | -0.09 | -0.20 | 0.15 | -0.02 | 0.28 | 3.2E-02 | 0.29 |
| MRPS24 | 0.20 | 0.15 | 0.00 | -0.24 | -0.08 | -0.08 | 3.2E-02 | -0.25 |
| DAZAP1 | 0.02 | 0.12 | 0.27 | -0.26 | -0.07 | -0.14 | 3.2E-02 | -0.29 |
| LEMD3 | 0.06 | 0.03 | 0.12 | -0.15 | -0.04 | -0.04 | 3.2E-02 | -0.14 |
| ZC3H4 | 0.07 | 0.12 | 0.06 | 0.00 | -0.09 | -0.18 | 3.2E-02 | -0.18 |
| P2RX6 | -0.29 | -0.15 | -0.06 | 0.05 | 0.10 | 0.28 | 3.2E-02 | 0.31 |
| TTI1 | -0.14 | -0.25 | -0.01 | 0.13 | 0.05 | 0.17 | 3.2E-02 | 0.25 |
| NELFCD | -0.15 | -0.17 | -0.11 | -0.04 | 0.20 | 0.21 | 3.3E-02 | 0.27 |

| | | | | | | | | |
|---|---|---|---|---|---|---|---|---|
| DPP3 | -0.09 | -0.06 | 0.00 | 0.01 | 0.07 | 0.07 | 3.3E-02 | 0.10 |
| AGK | 0.00 | 0.07 | 0.07 | -0.07 | -0.02 | -0.04 | 3.3E-02 | -0.09 |
| RNF6 | 0.09 | 0.07 | 0.14 | -0.09 | -0.23 | -0.01 | 3.3E-02 | -0.21 |
| VAPA | 0.06 | 0.11 | 0.09 | -0.09 | 0.00 | -0.19 | 3.3E-02 | -0.18 |
| CCNQ | 0.16 | 0.31 | 0.22 | -0.16 | -0.11 | -0.61 | 3.3E-02 | -0.53 |
| PLEKHG4 | -0.26 | -0.19 | 0.01 | 0.11 | 0.09 | 0.17 | 3.3E-02 | 0.27 |
| CDKN2AIPNL | -0.05 | -0.05 | -0.20 | 0.03 | 0.15 | 0.10 | 3.3E-02 | 0.19 |
| EMSY | 0.00 | -0.04 | -0.03 | 0.05 | 0.02 | 0.01 | 3.3E-02 | 0.05 |
| TMEM128 | 0.09 | 0.09 | 0.18 | -0.03 | -0.10 | -0.27 | 3.3E-02 | -0.25 |
| MDC1 | -0.11 | -0.10 | -0.12 | -0.01 | 0.10 | 0.21 | 3.3E-02 | 0.21 |
| EFTUD2 | 0.05 | 0.00 | 0.06 | -0.04 | -0.01 | -0.07 | 3.3E-02 | -0.08 |
| WBP4 | 0.12 | 0.06 | 0.01 | -0.11 | -0.08 | -0.02 | 3.3E-02 | -0.13 |
| RPP38 | 0.11 | -0.02 | 0.16 | -0.06 | -0.09 | -0.13 | 3.3E-02 | -0.18 |
| GNPNAT1 | 0.28 | 0.25 | 0.04 | -0.09 | -0.16 | -0.45 | 3.3E-02 | -0.43 |
| SKP1 | 0.04 | 0.01 | 0.06 | -0.04 | -0.06 | 0.00 | 3.3E-02 | -0.07 |
| SFXN1 | -0.02 | 0.15 | 0.19 | -0.14 | -0.05 | -0.16 | 3.3E-02 | -0.23 |
| GALNT6 | 0.03 | 0.03 | 0.11 | -0.02 | -0.05 | -0.10 | 3.3E-02 | -0.11 |
| MPP2 | 0.01 | 0.17 | 0.18 | -0.22 | -0.02 | -0.18 | 3.3E-02 | -0.26 |
| TOR1B | -0.33 | -0.05 | -0.09 | 0.09 | 0.15 | 0.18 | 3.3E-02 | 0.30 |
| KIF18B | -0.24 | -0.15 | -0.04 | 0.08 | 0.05 | 0.25 | 3.3E-02 | 0.27 |
| GRSF1 | 0.03 | 0.04 | 0.12 | -0.13 | -0.05 | -0.02 | 3.3E-02 | -0.13 |
| PPP4R1 | -0.13 | -0.21 | -0.03 | 0.20 | 0.02 | 0.11 | 3.3E-02 | 0.23 |
| SLC39A13 | -0.14 | -0.01 | -0.31 | 0.13 | 0.17 | 0.09 | 3.4E-02 | 0.28 |
| PTCD1 | 0.18 | 0.21 | 0.01 | -0.21 | -0.03 | -0.22 | 3.4E-02 | -0.29 |
| TBX3 | -0.02 | 0.23 | 0.12 | -0.15 | -0.10 | -0.13 | 3.4E-02 | -0.24 |
| ZNF99 | 0.01 | -0.35 | -0.24 | 0.13 | 0.13 | 0.24 | 3.4E-02 | 0.36 |
| M6PR | 0.04 | 0.14 | 0.60 | -0.34 | -0.22 | -0.49 | 3.4E-02 | -0.60 |
| NIFK | 0.08 | 0.04 | 0.02 | -0.02 | -0.03 | -0.11 | 3.4E-02 | -0.10 |
| TOLLIP | -0.19 | -0.03 | -0.14 | 0.02 | 0.10 | 0.21 | 3.4E-02 | 0.23 |
| NCL | 0.06 | 0.08 | 0.03 | -0.08 | 0.01 | -0.11 | 3.4E-02 | -0.12 |
| POF1B | 0.01 | -0.19 | -0.28 | 0.20 | 0.11 | 0.09 | 3.4E-02 | 0.29 |
| CASP8 | -0.26 | -0.48 | -0.01 | 0.11 | 0.26 | 0.24 | 3.4E-02 | 0.45 |
| KIF11 | -0.04 | -0.15 | -0.05 | 0.01 | 0.12 | 0.09 | 3.4E-02 | 0.15 |
| INTS6 | -0.01 | -0.04 | -0.09 | 0.02 | 0.04 | 0.08 | 3.4E-02 | 0.09 |
| METTL14 | 0.17 | 0.17 | 0.01 | -0.08 | -0.07 | -0.24 | 3.4E-02 | -0.25 |
| MRPL20 | 0.06 | 0.00 | 0.15 | -0.06 | -0.09 | -0.07 | 3.4E-02 | -0.15 |
| ALDH6A1 | 0.00 | -0.09 | -0.17 | 0.06 | 0.07 | 0.11 | 3.4E-02 | 0.16 |
| LRRIQ1 | 0.15 | -0.01 | 0.13 | -0.12 | -0.16 | -0.03 | 3.4E-02 | -0.20 |
| POLR3F | -0.16 | -0.17 | -0.10 | 0.22 | -0.04 | 0.19 | 3.4E-02 | 0.27 |
| NR2F2 | -0.14 | -0.19 | -0.12 | 0.04 | 0.05 | 0.31 | 3.4E-02 | 0.28 |
| FBL | 0.00 | -0.06 | -0.12 | 0.05 | 0.04 | 0.08 | 3.4E-02 | 0.12 |
| USP42 | 0.09 | 0.02 | 0.05 | 0.00 | -0.09 | -0.09 | 3.4E-02 | -0.11 |
| SNRPA1 | 0.05 | 0.00 | 0.05 | -0.06 | -0.01 | -0.03 | 3.4E-02 | -0.07 |
| SLC25A6 | -0.20 | -0.25 | -0.15 | 0.08 | 0.03 | 0.39 | 3.4E-02 | 0.37 |
| ENSA | 0.16 | 0.11 | 0.14 | -0.29 | 0.02 | -0.20 | 3.4E-02 | -0.29 |
| PTBP1 | 0.06 | 0.04 | 0.07 | -0.13 | -0.06 | 0.00 | 3.4E-02 | -0.12 |
| RTCB | -0.08 | -0.08 | -0.01 | 0.10 | 0.01 | 0.05 | 3.5E-02 | 0.11 |
| VRK1 | 0.03 | 0.02 | 0.01 | -0.01 | -0.01 | -0.05 | 3.5E-02 | -0.04 |

| | | | | | | | | |
|---|---|---|---|---|---|---|---|---|
| SCGN | 0.64 | -0.18 | 0.59 | -0.74 | -0.54 | -0.38 | 3.5E-02 | -0.90 |
| TSC22D3 | -0.01 | -0.23 | -0.14 | 0.19 | 0.05 | 0.10 | 3.5E-02 | 0.24 |
| GADD45GIP1 | 0.06 | 0.07 | 0.00 | -0.08 | -0.04 | -0.02 | 3.5E-02 | -0.09 |
| PTCD3 | 0.17 | 0.13 | -0.01 | -0.18 | -0.05 | -0.10 | 3.5E-02 | -0.21 |
| TERF2IP | -0.05 | -0.02 | -0.07 | -0.01 | 0.07 | 0.07 | 3.5E-02 | 0.09 |
| PCGF3 | -0.15 | -0.06 | -0.03 | 0.12 | 0.10 | 0.01 | 3.5E-02 | 0.16 |
| CAPN2 | -0.13 | -0.08 | -0.03 | -0.01 | 0.11 | 0.12 | 3.5E-02 | 0.15 |
| GPATCH11 | 0.00 | 0.09 | 0.13 | -0.14 | -0.06 | -0.04 | 3.5E-02 | -0.15 |
| MRPL38 | 0.05 | 0.06 | 0.03 | -0.10 | -0.04 | 0.00 | 3.5E-02 | -0.09 |
| OVOL1 | 0.30 | 0.02 | 0.21 | -0.26 | -0.03 | -0.35 | 3.5E-02 | -0.39 |
| SLC12A7 | 0.01 | 0.10 | 0.27 | -0.24 | -0.12 | -0.09 | 3.5E-02 | -0.28 |
| MANEAL | 0.05 | 0.06 | 0.12 | -0.03 | -0.04 | -0.18 | 3.5E-02 | -0.16 |
| MBOAT2 | 0.06 | 0.12 | 0.11 | -0.09 | -0.01 | -0.22 | 3.5E-02 | -0.20 |
| GNAI3 | -0.18 | -0.21 | -0.16 | 0.17 | -0.04 | 0.33 | 3.5E-02 | 0.34 |
| PANX1 | -0.13 | -0.21 | -0.18 | -0.05 | 0.27 | 0.23 | 3.5E-02 | 0.32 |
| ZNF451 | -0.06 | -0.04 | -0.05 | 0.00 | 0.04 | 0.11 | 3.5E-02 | 0.10 |
| CCDC22 | -0.08 | -0.19 | -0.01 | 0.03 | 0.12 | 0.11 | 3.5E-02 | 0.19 |
| CNNM3 | 0.03 | 0.12 | 0.17 | -0.01 | -0.14 | -0.21 | 3.5E-02 | -0.23 |
| TARBP2 | 0.06 | 0.05 | 0.10 | -0.16 | -0.02 | -0.04 | 3.5E-02 | -0.14 |
| GLRX5 | -0.03 | -0.06 | -0.22 | 0.09 | 0.06 | 0.13 | 3.5E-02 | 0.20 |
| UBE2G2 | 0.12 | -0.01 | 0.09 | -0.04 | -0.08 | -0.09 | 3.5E-02 | -0.14 |
| CLNS1A | -0.09 | -0.39 | -0.60 | 0.03 | 0.27 | 0.50 | 3.5E-02 | 0.62 |
| APH1A | -0.20 | -0.21 | -0.14 | -0.01 | 0.12 | 0.35 | 3.5E-02 | 0.34 |
| SLC20A1 | 0.01 | 0.23 | 0.25 | -0.36 | -0.10 | -0.12 | 3.5E-02 | -0.36 |
| SLIT3 | -0.39 | -0.07 | -0.11 | 0.21 | 0.06 | 0.21 | 3.5E-02 | 0.35 |
| TP53I11 | 0.11 | 0.07 | 0.14 | -0.20 | 0.02 | -0.18 | 3.5E-02 | -0.23 |
| CHD3 | -0.09 | -0.07 | -0.04 | 0.11 | -0.01 | 0.10 | 3.6E-02 | 0.13 |
| POLQ | 0.17 | 0.26 | 0.33 | -0.29 | -0.04 | -0.67 | 3.6E-02 | -0.59 |
| MFAP1 | 0.05 | 0.02 | 0.09 | -0.09 | 0.00 | -0.08 | 3.6E-02 | -0.11 |
| SUV39H2 | -0.08 | -0.12 | -0.38 | 0.26 | 0.04 | 0.19 | 3.6E-02 | 0.36 |
| NUDT12 | -0.01 | -0.18 | -0.28 | 0.21 | 0.04 | 0.16 | 3.6E-02 | 0.29 |
| RAB1B | 0.06 | 0.11 | 0.11 | -0.18 | 0.02 | -0.14 | 3.6E-02 | -0.20 |
| DNAJA3 | -0.09 | -0.06 | -0.01 | 0.00 | 0.08 | 0.08 | 3.6E-02 | 0.11 |
| LTN1 | -0.01 | -0.13 | -0.07 | 0.11 | 0.06 | 0.02 | 3.6E-02 | 0.14 |
| ALB | -0.15 | -0.16 | -0.24 | 0.36 | 0.04 | 0.06 | 3.6E-02 | 0.34 |
| DCAF5 | -0.17 | -0.10 | 0.00 | 0.14 | 0.08 | 0.04 | 3.6E-02 | 0.18 |
| BPNT2 | 0.02 | 0.14 | 0.22 | -0.02 | -0.25 | -0.16 | 3.6E-02 | -0.27 |
| GAPDH | 0.23 | 0.18 | -0.02 | -0.18 | -0.06 | -0.20 | 3.6E-02 | -0.28 |
| NAA40 | 0.27 | 0.05 | 0.22 | -0.44 | -0.11 | -0.10 | 3.6E-02 | -0.40 |
| BZW1 | -0.09 | -0.16 | -0.02 | 0.05 | 0.04 | 0.16 | 3.6E-02 | 0.17 |
| TOMM70 | 0.01 | 0.06 | 0.21 | -0.10 | -0.08 | -0.13 | 3.6E-02 | -0.20 |
| MRPS23 | 0.19 | 0.13 | 0.01 | -0.23 | -0.03 | -0.13 | 3.6E-02 | -0.24 |
| TRIQK | -0.37 | -0.52 | -0.17 | 0.07 | 0.11 | 0.60 | 3.6E-02 | 0.62 |
| MTPN | 0.00 | -0.20 | -0.29 | 0.05 | 0.19 | 0.18 | 3.6E-02 | 0.30 |
| ZMYM3 | 0.12 | 0.04 | 0.05 | -0.14 | -0.07 | -0.01 | 3.6E-02 | -0.14 |
| ERLEC1 | 0.11 | 0.10 | 0.13 | -0.03 | -0.08 | -0.29 | 3.6E-02 | -0.25 |
| ORC3 | -0.11 | -0.17 | -0.02 | 0.05 | 0.04 | 0.18 | 3.6E-02 | 0.19 |
| HTT | -0.17 | -0.17 | -0.09 | 0.03 | 0.06 | 0.29 | 3.6E-02 | 0.27 |

| | | | | | | | | |
|---|---|---|---|---|---|---|---|---|
| RCL1 | -0.02 | -0.02 | -0.12 | 0.04 | 0.06 | 0.06 | 3.7E-02 | 0.11 |
| DHX15 | 0.08 | 0.03 | 0.00 | -0.02 | -0.04 | -0.06 | 3.7E-02 | -0.08 |
| RRAGC | -0.16 | -0.04 | -0.06 | 0.13 | 0.00 | 0.10 | 3.7E-02 | 0.17 |
| STIM2 | 0.03 | 0.03 | 0.19 | -0.10 | -0.05 | -0.12 | 3.7E-02 | -0.17 |
| PPP1R12A | 0.04 | 0.15 | 0.09 | -0.05 | -0.05 | -0.22 | 3.7E-02 | -0.20 |
| PHF6 | -0.06 | -0.05 | -0.12 | 0.02 | 0.05 | 0.15 | 3.7E-02 | 0.15 |
| FOXK2 | 0.00 | -0.05 | -0.04 | 0.00 | 0.04 | 0.05 | 3.7E-02 | 0.06 |
| ASB7 | 0.43 | 0.59 | 0.20 | -0.63 | -0.11 | -1.24 | 3.7E-02 | -1.07 |
| LENG8 | 0.08 | 0.08 | 0.00 | -0.08 | -0.01 | -0.09 | 3.7E-02 | -0.11 |
| DIAPH1 | -0.05 | -0.01 | -0.06 | 0.04 | 0.07 | 0.01 | 3.7E-02 | 0.08 |
| EEF2K | -0.23 | -0.37 | -0.06 | 0.24 | 0.00 | 0.30 | 3.7E-02 | 0.40 |
| MRPS33 | 0.35 | 0.18 | -0.04 | -0.25 | -0.12 | -0.23 | 3.7E-02 | -0.37 |
| PDS5B | -0.02 | -0.04 | -0.03 | 0.02 | 0.00 | 0.06 | 3.7E-02 | 0.06 |
| RMI2 | -0.29 | -0.17 | -0.03 | 0.24 | 0.02 | 0.16 | 3.7E-02 | 0.30 |
| TKT | 0.20 | 0.13 | -0.01 | -0.05 | -0.15 | -0.16 | 3.7E-02 | -0.23 |
| SLC25A20 | 0.00 | 0.03 | 0.10 | -0.05 | -0.04 | -0.04 | 3.8E-02 | -0.09 |
| XPR1 | 0.10 | 0.02 | 0.10 | 0.00 | -0.15 | -0.09 | 3.8E-02 | -0.15 |
| MRPL28 | 0.01 | 0.03 | 0.09 | -0.02 | -0.07 | -0.05 | 3.8E-02 | -0.08 |
| TXNL4A | 0.05 | 0.04 | 0.12 | -0.02 | -0.05 | -0.15 | 3.8E-02 | -0.14 |
| PCCB | 0.06 | 0.10 | 0.36 | -0.35 | -0.06 | -0.22 | 3.8E-02 | -0.38 |
| ZBTB14 | -0.07 | -0.12 | -0.32 | 0.03 | 0.14 | 0.27 | 3.8E-02 | 0.31 |
| ATP2B4 | -0.03 | -0.04 | -0.02 | -0.01 | 0.06 | 0.05 | 3.8E-02 | 0.06 |
| SRPRB | 0.09 | 0.05 | 0.00 | -0.04 | -0.02 | -0.08 | 3.8E-02 | -0.09 |
| NUP50 | 0.06 | 0.05 | 0.01 | -0.08 | -0.01 | -0.04 | 3.8E-02 | -0.08 |
| GPR108 | -0.01 | 0.20 | 0.26 | -0.27 | -0.20 | -0.05 | 3.8E-02 | -0.32 |
| AKT2 | -0.31 | -0.31 | -0.04 | 0.28 | -0.01 | 0.27 | 3.8E-02 | 0.40 |
| RPP40 | 0.14 | 0.12 | 0.05 | -0.03 | -0.07 | -0.24 | 3.8E-02 | -0.21 |
| MRPL47 | 0.09 | 0.14 | 0.00 | -0.11 | -0.02 | -0.11 | 3.8E-02 | -0.15 |
| PPP1CA | 0.08 | 0.07 | 0.20 | -0.23 | 0.00 | -0.16 | 3.8E-02 | -0.25 |
| HEATR3 | -0.07 | -0.17 | -0.20 | 0.25 | 0.16 | -0.03 | 3.8E-02 | 0.27 |
| SLC4A1AP | -0.03 | -0.09 | -0.27 | 0.05 | 0.14 | 0.16 | 3.8E-02 | 0.24 |
| POLR1B | 0.06 | 0.00 | 0.12 | -0.10 | -0.03 | -0.07 | 3.8E-02 | -0.13 |
| XPO1 | -0.04 | -0.10 | -0.09 | 0.16 | 0.02 | 0.04 | 3.8E-02 | 0.15 |
| TMEM41A | -0.01 | 0.08 | 0.13 | -0.10 | -0.07 | -0.05 | 3.8E-02 | -0.14 |
| KRT12 | -0.67 | -0.39 | -0.17 | 0.11 | 0.11 | 0.64 | 3.8E-02 | 0.70 |
| KLHL7 | 0.10 | 0.13 | 0.07 | -0.21 | 0.02 | -0.15 | 3.8E-02 | -0.22 |
| CBFB | -0.06 | -0.09 | -0.10 | 0.01 | 0.04 | 0.18 | 3.8E-02 | 0.16 |
| SMAD2 | 0.08 | 0.06 | 0.06 | 0.02 | -0.12 | -0.12 | 3.8E-02 | -0.14 |
| S100PBP | 0.08 | 0.27 | 0.22 | -0.48 | -0.07 | -0.14 | 3.8E-02 | -0.42 |
| FEN1 | -0.01 | -0.05 | -0.01 | 0.01 | 0.04 | 0.02 | 3.8E-02 | 0.05 |
| SEPTIN2 | 0.19 | 0.26 | -0.03 | -0.19 | -0.07 | -0.21 | 3.8E-02 | -0.30 |
| ATP6V1H | -0.01 | -0.10 | -0.08 | 0.10 | 0.00 | 0.07 | 3.9E-02 | 0.12 |
| VPS51 | -0.22 | -0.34 | -0.02 | 0.23 | 0.02 | 0.24 | 3.9E-02 | 0.36 |
| OVOL2 | -0.02 | -0.03 | -0.11 | 0.03 | 0.09 | 0.03 | 3.9E-02 | 0.10 |
| GALNT2 | 0.02 | 0.08 | 0.06 | -0.01 | -0.05 | -0.12 | 3.9E-02 | -0.11 |
| FAM50A | 0.10 | 0.15 | 0.01 | -0.13 | -0.01 | -0.15 | 3.9E-02 | -0.18 |
| ZNHIT2 | -0.11 | -0.25 | -0.05 | 0.10 | 0.02 | 0.24 | 3.9E-02 | 0.26 |
| SFXN5 | 0.04 | 0.01 | 0.14 | -0.04 | -0.09 | -0.07 | 3.9E-02 | -0.13 |

| | | | | | | | | |
|---|---|---|---|---|---|---|---|---|
| **RAB10** | 0.00 | 0.07 | 0.06 | -0.03 | -0.02 | -0.08 | 3.9E-02 | -0.09 |
| **TMEM214** | 0.04 | 0.02 | 0.02 | 0.00 | -0.03 | -0.05 | 3.9E-02 | -0.05 |
| **STK24** | 0.12 | 0.15 | 0.00 | -0.09 | -0.02 | -0.17 | 3.9E-02 | -0.19 |
| **PSMG1** | 0.05 | 0.14 | 0.26 | -0.17 | -0.33 | -0.02 | 3.9E-02 | -0.33 |
| **MRPL46** | -0.04 | -0.06 | -0.13 | 0.03 | 0.04 | 0.15 | 3.9E-02 | 0.15 |
| **MARS1** | -0.16 | -0.10 | -0.02 | 0.13 | 0.12 | 0.00 | 3.9E-02 | 0.18 |
| **LRRFIP1** | 0.05 | 0.12 | 0.01 | -0.05 | -0.03 | -0.11 | 3.9E-02 | -0.12 |
| **YAP1** | 0.01 | 0.07 | 0.10 | -0.11 | -0.07 | -0.01 | 3.9E-02 | -0.12 |
| **PPP5C** | 0.04 | 0.11 | 0.34 | -0.30 | -0.05 | -0.24 | 3.9E-02 | -0.36 |
| **NEU1** | 0.10 | 0.04 | 0.22 | 0.00 | -0.20 | -0.21 | 3.9E-02 | -0.26 |
| **SCN4A** | 0.12 | 0.12 | 0.32 | -0.21 | -0.03 | -0.44 | 3.9E-02 | -0.41 |
| **USP40** | 0.00 | -0.10 | -0.19 | 0.12 | 0.10 | 0.04 | 3.9E-02 | 0.18 |
| **POLA2** | -0.06 | -0.43 | -0.11 | 0.21 | 0.18 | 0.11 | 4.0E-02 | 0.36 |
| **LMAN2L** | 0.03 | 0.09 | 0.18 | -0.23 | -0.05 | -0.06 | 4.0E-02 | -0.21 |
| **PSME1** | 0.07 | 0.19 | -0.01 | -0.08 | -0.12 | -0.08 | 4.0E-02 | -0.18 |
| **ACOT7** | -0.01 | -0.09 | -0.25 | 0.13 | 0.12 | 0.05 | 4.0E-02 | 0.22 |
| **PTPRF** | 0.00 | 0.07 | 0.24 | -0.16 | -0.11 | -0.09 | 4.0E-02 | -0.23 |
| **ATP5ME** | 0.22 | 0.53 | 0.02 | -0.47 | -0.08 | -0.48 | 4.0E-02 | -0.60 |
| **SLX4** | -0.13 | -0.06 | -0.21 | 0.17 | -0.02 | 0.20 | 4.0E-02 | 0.25 |
| **SAMM50** | -0.06 | -0.11 | -0.11 | 0.04 | 0.01 | 0.20 | 4.0E-02 | 0.18 |
| **PAG1** | 0.13 | 0.14 | 0.09 | -0.08 | -0.03 | -0.30 | 4.0E-02 | -0.25 |
| **CIB1** | -0.41 | -0.63 | -0.23 | 0.13 | 0.04 | 0.70 | 4.0E-02 | 0.71 |
| **HBE1** | -0.04 | -0.10 | -0.20 | 0.02 | 0.20 | 0.08 | 4.0E-02 | 0.21 |
| **AP1M2** | -0.13 | -0.11 | 0.00 | 0.12 | 0.02 | 0.07 | 4.0E-02 | 0.15 |
| **LNPK** | 0.08 | 0.15 | 0.01 | -0.14 | -0.11 | -0.01 | 4.0E-02 | -0.17 |
| **UBE2V2** | 0.29 | 0.28 | 0.02 | -0.33 | -0.01 | -0.41 | 4.0E-02 | -0.45 |
| **DPY30** | -0.02 | -0.14 | -0.15 | 0.12 | -0.01 | 0.16 | 4.0E-02 | 0.20 |
| **BDP1** | -0.02 | -0.18 | -0.22 | 0.21 | 0.00 | 0.17 | 4.0E-02 | 0.27 |
| **GVINP1** | 0.07 | 0.08 | 0.28 | -0.24 | -0.01 | -0.25 | 4.0E-02 | -0.31 |
| **NEPRO** | 0.10 | 0.04 | 0.04 | -0.04 | -0.02 | -0.13 | 4.0E-02 | -0.12 |
| **TRMT10C** | 0.11 | 0.04 | 0.04 | -0.13 | -0.02 | -0.04 | 4.0E-02 | -0.12 |
| **CDV3** | 0.18 | 0.19 | -0.03 | -0.18 | -0.05 | -0.16 | 4.0E-02 | -0.24 |
| **ZBTB17** | -0.13 | -0.09 | -0.06 | -0.02 | 0.10 | 0.17 | 4.0E-02 | 0.18 |
| **TRMT2B** | 0.26 | 0.13 | -0.02 | -0.07 | -0.21 | -0.15 | 4.0E-02 | -0.27 |
| **ECT2** | 0.01 | 0.08 | 0.18 | -0.05 | -0.07 | -0.19 | 4.0E-02 | -0.19 |
| **MRPS31** | 0.19 | 0.11 | -0.02 | -0.10 | -0.06 | -0.15 | 4.0E-02 | -0.20 |
| **TAPBP** | 0.02 | 0.14 | 0.05 | -0.12 | -0.01 | -0.09 | 4.0E-02 | -0.14 |
| **CRYBG1** | -0.11 | -0.04 | -0.01 | 0.05 | 0.03 | 0.08 | 4.0E-02 | 0.10 |
| **ARL6IP6** | -0.13 | -0.16 | -0.09 | 0.04 | 0.03 | 0.26 | 4.0E-02 | 0.24 |
| **NAA20** | -0.03 | 0.13 | 0.26 | -0.11 | -0.14 | -0.17 | 4.1E-02 | -0.26 |
| **UBR1** | -0.13 | -0.15 | -0.03 | 0.16 | 0.13 | -0.01 | 4.1E-02 | 0.20 |
| **NUFIP1** | 0.05 | 0.09 | 0.15 | -0.11 | 0.00 | -0.22 | 4.1E-02 | -0.21 |
| **BYSL** | 0.05 | 0.04 | 0.13 | -0.05 | -0.03 | -0.16 | 4.1E-02 | -0.15 |
| **RAB6D** | 0.09 | 0.23 | 0.39 | -0.26 | -0.06 | -0.61 | 4.1E-02 | -0.55 |
| **NCOR2** | -0.05 | -0.13 | -0.25 | 0.03 | 0.08 | 0.25 | 4.1E-02 | 0.26 |
| **ATP4A** | 0.06 | 0.11 | 0.27 | -0.16 | -0.03 | -0.32 | 4.1E-02 | -0.32 |
| **LAD1** | -0.11 | 0.01 | -0.10 | 0.03 | 0.09 | 0.06 | 4.1E-02 | 0.13 |
| **DYRK1A** | -0.18 | -0.37 | 0.00 | 0.13 | 0.10 | 0.24 | 4.1E-02 | 0.34 |

| | | | | | | | | |
|---|---|---|---|---|---|---|---|---|
| NSRP1 | 0.11 | 0.19 | 0.02 | -0.11 | -0.02 | -0.22 | 4.1E-02 | -0.22 |
| HAUS3 | -0.07 | -0.06 | -0.16 | 0.09 | 0.00 | 0.18 | 4.1E-02 | 0.18 |
| MBD2 | -0.07 | -0.07 | -0.08 | -0.02 | 0.07 | 0.14 | 4.1E-02 | 0.14 |
| STT3B | 0.01 | 0.02 | 0.13 | -0.05 | -0.07 | -0.05 | 4.1E-02 | -0.11 |
| NSF | 0.01 | 0.07 | 0.20 | -0.04 | -0.11 | -0.16 | 4.1E-02 | -0.20 |
| EIF3C | 0.04 | 0.00 | 0.15 | -0.08 | -0.04 | -0.08 | 4.1E-02 | -0.14 |
| ARMCX2 | 0.10 | 0.08 | 0.24 | -0.29 | 0.00 | -0.19 | 4.1E-02 | -0.30 |
| DDX20 | -0.06 | -0.16 | -0.02 | 0.03 | 0.05 | 0.14 | 4.1E-02 | 0.15 |
| TXNDC5 | 0.08 | 0.31 | 0.02 | -0.22 | -0.07 | -0.18 | 4.1E-02 | -0.29 |
| SSR1 | -0.13 | -0.21 | -0.06 | 0.05 | 0.04 | 0.26 | 4.1E-02 | 0.25 |
| ZC3HAV1 | -0.13 | -0.08 | -0.06 | 0.19 | 0.05 | 0.01 | 4.1E-02 | 0.18 |
| SDAD1 | 0.13 | 0.13 | 0.04 | -0.13 | 0.01 | -0.21 | 4.2E-02 | -0.21 |
| SLC35A1 | 0.03 | 0.05 | 0.21 | -0.03 | -0.13 | -0.17 | 4.2E-02 | -0.21 |
| DDR1 | -0.27 | -0.20 | 0.00 | 0.16 | 0.21 | 0.03 | 4.2E-02 | 0.29 |
| ARPC3 | 0.05 | 0.09 | 0.14 | 0.01 | -0.11 | -0.20 | 4.2E-02 | -0.19 |
| DEGS2 | 0.08 | 0.01 | 0.18 | -0.03 | -0.16 | -0.11 | 4.2E-02 | -0.19 |
| SORD | 0.19 | 0.23 | -0.04 | -0.19 | -0.16 | -0.08 | 4.2E-02 | -0.27 |
| FKBP15 | 0.30 | 0.31 | 0.35 | -0.12 | -0.28 | -0.98 | 4.2E-02 | -0.78 |
| SNRNP200 | 0.06 | 0.00 | 0.02 | -0.02 | -0.02 | -0.04 | 4.2E-02 | -0.06 |
| MANBA | -0.07 | -0.11 | -0.21 | 0.01 | 0.08 | 0.25 | 4.2E-02 | 0.24 |
| SCYL2 | 0.04 | 0.08 | 0.15 | 0.00 | -0.10 | -0.20 | 4.2E-02 | -0.19 |
| P3H4 | 0.35 | 0.34 | 0.18 | -0.18 | -0.16 | -0.86 | 4.2E-02 | -0.69 |
| PLPP3 | 0.40 | 0.22 | 0.25 | -0.82 | -0.03 | -0.35 | 4.2E-02 | -0.69 |
| SKIC2 | -0.04 | -0.16 | -0.04 | 0.12 | 0.01 | 0.09 | 4.2E-02 | 0.16 |
| BFAR | 0.18 | 0.06 | 0.29 | 0.01 | -0.35 | -0.30 | 4.2E-02 | -0.39 |
| IBTK | 0.16 | 0.08 | 0.76 | -0.16 | -0.56 | -0.83 | 4.2E-02 | -0.85 |
| MEAK7 | 0.04 | 0.16 | 0.12 | -0.13 | 0.00 | -0.23 | 4.2E-02 | -0.23 |
| SUPT6H | 0.16 | 0.06 | 0.03 | -0.02 | -0.07 | -0.18 | 4.3E-02 | -0.18 |
| ESYT1 | -0.14 | -0.09 | 0.01 | 0.08 | 0.09 | 0.03 | 4.3E-02 | 0.14 |
| CEP95 | 0.02 | 0.04 | 0.16 | -0.10 | -0.10 | -0.03 | 4.3E-02 | -0.15 |
| PHF14 | 0.08 | 0.03 | 0.11 | -0.15 | -0.09 | 0.01 | 4.3E-02 | -0.15 |
| TRRAP | -0.05 | -0.07 | -0.09 | 0.12 | -0.02 | 0.10 | 4.3E-02 | 0.14 |
| PNPLA6 | 0.06 | 0.03 | 0.18 | -0.01 | -0.16 | -0.14 | 4.3E-02 | -0.19 |
| ZBTB8OS | -0.03 | -0.19 | -0.04 | 0.04 | 0.09 | 0.11 | 4.3E-02 | 0.16 |
| MBIP | -0.09 | -0.10 | -0.04 | 0.04 | 0.01 | 0.16 | 4.3E-02 | 0.14 |
| TUBB2A | -0.12 | -0.15 | -0.32 | 0.36 | 0.12 | 0.00 | 4.3E-02 | 0.36 |
| DNAJA4 | -0.09 | -0.10 | -0.17 | 0.09 | 0.24 | -0.01 | 4.3E-02 | 0.23 |
| KIAA1328 | 0.21 | 0.15 | 0.28 | -0.59 | -0.15 | -0.07 | 4.3E-02 | -0.49 |
| MIPEP | -0.03 | -0.04 | -0.23 | 0.11 | 0.08 | 0.08 | 4.3E-02 | 0.19 |
| HMGB1 | 0.12 | 0.08 | -0.01 | -0.11 | -0.07 | -0.03 | 4.3E-02 | -0.13 |
| ST7L | -0.03 | -0.12 | -0.05 | 0.10 | -0.01 | 0.10 | 4.3E-02 | 0.13 |
| RECQL | -0.08 | -0.01 | -0.06 | 0.08 | 0.06 | 0.00 | 4.3E-02 | 0.09 |
| EPB41L2 | 0.06 | 0.03 | 0.19 | -0.19 | -0.07 | -0.05 | 4.3E-02 | -0.20 |
| FKBP5 | -0.11 | -0.02 | -0.09 | 0.00 | 0.13 | 0.06 | 4.3E-02 | 0.14 |
| ACIN1 | 0.16 | 0.21 | 0.15 | -0.07 | -0.08 | -0.46 | 4.4E-02 | -0.38 |
| MSTO1 | -0.07 | -0.36 | -0.44 | 0.38 | -0.05 | 0.33 | 4.4E-02 | 0.51 |
| GLRX3 | -0.04 | -0.21 | -0.13 | -0.02 | 0.15 | 0.20 | 4.4E-02 | 0.24 |
| MRPL22 | 0.10 | 0.06 | 0.09 | 0.02 | -0.11 | -0.17 | 4.4E-02 | -0.17 |

| | | | | | | | | |
|---|---|---|---|---|---|---|---|---|
| **ARFIP2** | -0.06 | -0.14 | -0.20 | 0.27 | 0.06 | 0.03 | 4.4E-02 | 0.25 |
| **RTF1** | 0.22 | 0.06 | 0.05 | -0.24 | -0.06 | -0.07 | 4.4E-02 | -0.23 |
| **PHRF1** | 0.07 | 0.00 | 0.08 | -0.09 | -0.01 | -0.06 | 4.4E-02 | -0.10 |
| **FXR1** | -0.10 | -0.03 | -0.14 | 0.14 | 0.13 | -0.02 | 4.4E-02 | 0.17 |
| **NAA16** | 0.14 | 0.03 | 0.14 | 0.01 | -0.15 | -0.20 | 4.4E-02 | -0.21 |
| **FLOT2** | -0.01 | -0.01 | -0.01 | 0.02 | 0.00 | 0.00 | 4.4E-02 | 0.01 |
| **RBM8A** | 0.06 | 0.03 | 0.10 | -0.03 | -0.02 | -0.15 | 4.4E-02 | -0.13 |
| **RPS6KA3** | -0.04 | -0.10 | -0.30 | 0.20 | 0.15 | 0.03 | 4.4E-02 | 0.27 |
| **VBP1** | -0.08 | -0.09 | -0.12 | 0.20 | 0.08 | -0.02 | 4.4E-02 | 0.18 |
| **LMTK2** | -0.16 | -0.05 | -0.03 | 0.04 | 0.04 | 0.14 | 4.4E-02 | 0.15 |
| **TOR1AIP1** | 0.02 | 0.04 | 0.11 | -0.11 | 0.00 | -0.08 | 4.4E-02 | -0.12 |
| **BRIX1** | 0.05 | 0.02 | 0.05 | 0.01 | -0.07 | -0.08 | 4.4E-02 | -0.09 |
| **C3orf38** | -0.01 | -0.07 | -0.04 | 0.07 | 0.06 | -0.01 | 4.4E-02 | 0.08 |
| **MRI1** | -0.24 | -0.05 | -0.38 | 0.01 | 0.18 | 0.36 | 4.4E-02 | 0.41 |
| **SLC25A44** | -0.09 | -0.09 | -0.12 | -0.01 | 0.07 | 0.21 | 4.4E-02 | 0.19 |
| **FGFR2** | 0.11 | 0.22 | 0.15 | -0.41 | 0.00 | -0.16 | 4.4E-02 | -0.35 |
| **MTO1** | 0.01 | 0.06 | 0.02 | -0.03 | -0.05 | -0.01 | 4.4E-02 | -0.06 |
| **NME2P1** | -0.08 | -0.19 | -0.10 | 0.21 | -0.04 | 0.16 | 4.4E-02 | 0.24 |
| **PIP4K2A** | 0.07 | 0.11 | 0.18 | -0.21 | 0.03 | -0.24 | 4.4E-02 | -0.26 |
| **ALG3** | 0.04 | 0.07 | 0.13 | 0.01 | -0.11 | -0.17 | 4.4E-02 | -0.17 |
| **PRIM1** | 0.01 | -0.19 | -0.30 | 0.22 | 0.04 | 0.15 | 4.4E-02 | 0.30 |
| **ZNF22** | 0.12 | 0.06 | 0.08 | -0.10 | 0.01 | -0.19 | 4.4E-02 | -0.18 |
| **NAT10** | 0.02 | 0.02 | 0.05 | -0.01 | -0.02 | -0.07 | 4.5E-02 | -0.07 |
| **TENT4B** | 0.11 | 0.11 | 0.03 | 0.01 | -0.12 | -0.17 | 4.5E-02 | -0.18 |
| **SP110** | 0.00 | 0.08 | 0.05 | 0.00 | -0.08 | -0.05 | 4.5E-02 | -0.09 |
| **MPRIP** | 0.06 | 0.04 | 0.05 | 0.02 | -0.08 | -0.10 | 4.5E-02 | -0.10 |
| **HIRIP3** | 0.15 | 0.11 | 0.16 | -0.34 | 0.02 | -0.17 | 4.5E-02 | -0.30 |
| **APLP2** | 0.09 | 0.12 | 0.14 | 0.00 | -0.10 | -0.29 | 4.5E-02 | -0.25 |
| **TRPM7** | 0.01 | 0.06 | 0.03 | -0.06 | 0.00 | -0.04 | 4.5E-02 | -0.07 |
| **RANBP10** | -0.07 | -0.07 | -0.07 | 0.12 | 0.09 | -0.03 | 4.5E-02 | 0.13 |
| **ZNF692** | -0.08 | -0.07 | -0.07 | -0.03 | 0.12 | 0.12 | 4.5E-02 | 0.14 |
| **UBXN7** | -0.14 | -0.19 | -0.06 | 0.25 | -0.02 | 0.12 | 4.5E-02 | 0.25 |
| **HERC1** | -0.13 | -0.16 | 0.02 | 0.11 | 0.06 | 0.08 | 4.5E-02 | 0.17 |
| **NLE1** | 0.07 | 0.03 | 0.03 | -0.01 | -0.03 | -0.10 | 4.5E-02 | -0.09 |
| **RPL35** | -0.04 | -0.07 | -0.01 | 0.08 | 0.05 | 0.00 | 4.5E-02 | 0.08 |
| **SLC29A2** | 0.04 | 0.12 | 0.49 | -0.17 | -0.49 | -0.18 | 4.5E-02 | -0.50 |
| **CALU** | 0.18 | 0.18 | -0.04 | -0.16 | -0.08 | -0.12 | 4.5E-02 | -0.23 |
| **MAPRE1** | 0.06 | 0.11 | 0.00 | -0.06 | -0.10 | -0.01 | 4.5E-02 | -0.11 |
| **GALNT10** | 0.00 | 0.12 | 0.12 | -0.01 | -0.09 | -0.16 | 4.5E-02 | -0.17 |
| **H4C1** | -0.06 | -0.11 | -0.02 | 0.12 | 0.04 | 0.02 | 4.5E-02 | 0.12 |
| **STOML2** | 0.04 | 0.08 | 0.02 | -0.11 | -0.02 | -0.02 | 4.5E-02 | -0.10 |
| **DDX3X** | 0.10 | 0.10 | 0.15 | -0.06 | -0.04 | -0.31 | 4.5E-02 | -0.25 |
| **AKAP11** | -0.15 | -0.25 | -0.07 | 0.11 | 0.00 | 0.30 | 4.5E-02 | 0.29 |
| **ZCCHC7** | 0.14 | 0.19 | 0.02 | -0.24 | 0.00 | -0.15 | 4.5E-02 | -0.25 |
| **ALG6** | 0.01 | 0.09 | 0.07 | -0.02 | -0.04 | -0.12 | 4.5E-02 | -0.11 |
| **USP36** | 0.03 | 0.04 | 0.05 | -0.03 | 0.00 | -0.09 | 4.5E-02 | -0.08 |
| **ZNF460** | -0.03 | -0.25 | -0.33 | 0.35 | 0.05 | 0.11 | 4.5E-02 | 0.37 |
| **RBFOX2** | -0.13 | -0.30 | -0.19 | 0.11 | 0.00 | 0.40 | 4.5E-02 | 0.37 |

| | | | | | | | | |
|---|---|---|---|---|---|---|---|---|
| ZRANB2 | 0.29 | 0.11 | 0.04 | -0.20 | -0.02 | -0.30 | 4.5E-02 | -0.32 |
| GANAB | -0.01 | -0.07 | -0.13 | 0.08 | 0.10 | 0.01 | 4.5E-02 | 0.13 |
| TKFC | -0.07 | -0.10 | 0.00 | 0.04 | 0.10 | 0.01 | 4.5E-02 | 0.11 |
| TOM1L1 | -0.02 | -0.05 | -0.09 | 0.01 | 0.11 | 0.04 | 4.5E-02 | 0.11 |
| ACD | -0.22 | -0.10 | -0.13 | 0.16 | -0.04 | 0.26 | 4.6E-02 | 0.28 |
| OCLN | -0.05 | 0.01 | -0.07 | 0.05 | 0.04 | 0.02 | 4.6E-02 | 0.07 |
| NAPG | 0.08 | 0.12 | 0.13 | -0.01 | -0.08 | -0.27 | 4.6E-02 | -0.23 |
| PELO | -0.08 | -0.16 | 0.00 | 0.06 | 0.05 | 0.11 | 4.6E-02 | 0.15 |
| DCLK1 | -0.35 | -0.21 | 0.00 | 0.25 | 0.03 | 0.19 | 4.6E-02 | 0.35 |
| SYF2 | 0.15 | 0.21 | 0.01 | -0.17 | -0.01 | -0.26 | 4.6E-02 | -0.27 |
| RPP14 | -0.19 | -0.51 | -0.24 | 0.22 | -0.05 | 0.53 | 4.6E-02 | 0.55 |
| CREB3L2 | -0.06 | -0.22 | -0.11 | 0.23 | 0.14 | -0.02 | 4.6E-02 | 0.25 |
| BAG6 | -0.04 | -0.14 | -0.08 | 0.01 | 0.18 | 0.05 | 4.6E-02 | 0.17 |
| ACSL1 | -0.01 | -0.09 | -0.23 | 0.15 | 0.12 | 0.03 | 4.6E-02 | 0.21 |
| DDX56 | 0.06 | 0.03 | 0.06 | -0.04 | 0.00 | -0.11 | 4.6E-02 | -0.10 |
| MRPL18 | 0.07 | 0.13 | 0.20 | -0.32 | 0.00 | -0.13 | 4.6E-02 | -0.28 |
| ZNF296 | 0.03 | -0.30 | -0.19 | 0.13 | 0.09 | 0.17 | 4.6E-02 | 0.28 |
| ODF2 | 0.07 | 0.03 | 0.06 | -0.13 | 0.00 | -0.05 | 4.6E-02 | -0.11 |
| TUBGCP6 | 0.20 | 0.22 | 0.24 | -0.19 | -0.03 | -0.61 | 4.6E-02 | -0.50 |
| PQBP1 | 0.11 | 0.03 | 0.05 | 0.01 | -0.09 | -0.13 | 4.6E-02 | -0.14 |
| C3orf33 | 0.04 | 0.10 | 0.31 | -0.33 | -0.18 | -0.04 | 4.7E-02 | -0.33 |
| TMEM223 | 0.17 | 0.19 | 0.06 | -0.02 | -0.35 | -0.12 | 4.7E-02 | -0.30 |
| HDAC1 | -0.08 | -0.15 | -0.05 | 0.04 | 0.03 | 0.19 | 4.7E-02 | 0.18 |
| DCUN1D5 | -0.19 | -0.14 | 0.03 | 0.09 | 0.06 | 0.12 | 4.7E-02 | 0.19 |
| UQCRC2 | 0.00 | 0.03 | 0.03 | 0.00 | -0.02 | -0.04 | 4.7E-02 | -0.04 |
| COMTD1 | -0.05 | 0.00 | -0.06 | 0.03 | 0.01 | 0.07 | 4.7E-02 | 0.07 |
| CRK | -0.10 | -0.01 | -0.27 | 0.09 | 0.17 | 0.07 | 4.7E-02 | 0.24 |
| CHID1 | 0.07 | 0.04 | 0.14 | -0.02 | -0.06 | -0.20 | 4.7E-02 | -0.18 |
| JARID2 | 0.24 | 0.17 | 0.17 | -0.54 | -0.05 | -0.14 | 4.7E-02 | -0.44 |
| SRSF1 | 0.17 | 0.19 | 0.16 | -0.05 | -0.10 | -0.48 | 4.7E-02 | -0.38 |
| PIBF1 | 0.03 | 0.13 | 0.07 | -0.14 | -0.12 | 0.01 | 4.7E-02 | -0.16 |
| AFG2B | 0.06 | 0.00 | 0.09 | -0.10 | -0.03 | -0.03 | 4.7E-02 | -0.10 |
| RNF126 | -0.34 | -0.52 | -0.28 | -0.10 | 0.65 | 0.25 | 4.7E-02 | 0.65 |
| ACTRT1 | -0.01 | 0.13 | 0.20 | -0.10 | -0.04 | -0.22 | 4.7E-02 | -0.23 |
| STXBP5 | -0.07 | -0.02 | -0.16 | 0.12 | 0.11 | 0.00 | 4.7E-02 | 0.16 |
| SLC22A18 | 0.09 | 0.09 | 0.26 | -0.37 | -0.03 | -0.13 | 4.7E-02 | -0.32 |
| HSPE1 | 0.43 | 0.55 | -0.17 | -0.26 | -0.46 | -0.44 | 4.7E-02 | -0.66 |
| L2HGDH | 0.01 | -0.14 | -0.18 | 0.10 | 0.16 | 0.02 | 4.7E-02 | 0.20 |
| NAGA | -0.10 | -0.10 | 0.02 | 0.04 | 0.05 | 0.08 | 4.7E-02 | 0.11 |
| ANGEL2 | -0.08 | 0.00 | -0.16 | 0.11 | 0.07 | 0.03 | 4.7E-02 | 0.15 |
| NHERF2 | -0.01 | 0.08 | 0.22 | -0.15 | -0.10 | -0.07 | 4.8E-02 | -0.20 |
| ZNF420 | 0.12 | 0.12 | 0.03 | -0.06 | -0.02 | -0.20 | 4.8E-02 | -0.18 |
| TMEM14C | -0.49 | -0.43 | -0.51 | 0.22 | -0.13 | 0.82 | 4.8E-02 | 0.78 |
| ACO1 | -0.22 | -0.17 | 0.00 | 0.22 | 0.09 | 0.03 | 4.8E-02 | 0.25 |
| MYRIP | 0.38 | 1.04 | -0.22 | -1.12 | -0.41 | -0.80 | 4.8E-02 | -1.18 |
| TFAP2A | 0.01 | -0.09 | -0.17 | 0.08 | 0.04 | 0.11 | 4.8E-02 | 0.16 |
| CENPU | 0.02 | 0.03 | 0.02 | -0.03 | -0.06 | 0.01 | 4.8E-02 | -0.05 |
| EXOC5 | -0.12 | -0.17 | 0.01 | 0.04 | 0.13 | 0.08 | 4.8E-02 | 0.17 |

| | | | | | | | | |
|---|---|---|---|---|---|---|---|---|
| RBM25 | 0.18 | 0.20 | 0.09 | -0.10 | -0.04 | -0.42 | 4.8E-02 | -0.35 |
| NFKB2 | -0.15 | -0.25 | -0.15 | 0.31 | 0.22 | -0.07 | 4.8E-02 | 0.34 |
| AKAP17A | 0.01 | 0.06 | 0.03 | -0.04 | 0.00 | -0.07 | 4.8E-02 | -0.07 |
| ANP32A | 0.20 | 0.23 | 0.00 | -0.19 | -0.01 | -0.30 | 4.8E-02 | -0.31 |
| ZNF280C | -0.01 | -0.11 | -0.06 | 0.01 | 0.10 | 0.04 | 4.8E-02 | 0.11 |
| QPCTL | 0.00 | 0.09 | 0.12 | -0.08 | -0.01 | -0.13 | 4.8E-02 | -0.14 |
| SEC61B | 0.07 | 0.10 | 0.11 | 0.03 | -0.15 | -0.19 | 4.8E-02 | -0.19 |
| FN3KRP | 0.18 | 0.19 | 0.00 | 0.00 | -0.22 | -0.21 | 4.9E-02 | -0.27 |
| SAR1A | -0.02 | -0.22 | -0.30 | 0.06 | 0.32 | 0.08 | 4.9E-02 | 0.33 |
| DDX1 | -0.07 | -0.06 | 0.01 | 0.05 | 0.03 | 0.04 | 4.9E-02 | 0.08 |
| NUDT4B | -0.07 | -0.33 | -0.16 | 0.34 | 0.04 | 0.08 | 4.9E-02 | 0.34 |
| UNC119B | -0.05 | -0.40 | -0.11 | 0.27 | 0.15 | 0.05 | 4.9E-02 | 0.34 |
| STXBP3 | -0.03 | 0.12 | 0.18 | -0.16 | -0.08 | -0.06 | 4.9E-02 | -0.19 |
| SLC33A1 | -0.01 | 0.11 | 0.27 | -0.04 | -0.22 | -0.16 | 4.9E-02 | -0.27 |
| DSP | -0.14 | -0.06 | -0.03 | 0.15 | 0.06 | 0.01 | 4.9E-02 | 0.15 |
| SESN2 | -0.20 | -0.36 | -0.01 | 0.01 | 0.21 | 0.25 | 4.9E-02 | 0.35 |
| NIPA2 | -0.13 | -0.05 | -0.01 | 0.07 | 0.01 | 0.10 | 4.9E-02 | 0.12 |
| DR1 | 0.03 | -0.15 | -0.16 | 0.05 | 0.12 | 0.09 | 4.9E-02 | 0.18 |
| ZCCHC8 | 0.07 | 0.08 | 0.06 | 0.00 | -0.17 | -0.06 | 4.9E-02 | -0.14 |
| PTDSS2 | 0.02 | 0.09 | 0.23 | -0.08 | -0.06 | -0.24 | 4.9E-02 | -0.24 |
| GZF1 | 0.00 | 0.03 | 0.06 | -0.03 | -0.06 | -0.01 | 4.9E-02 | -0.06 |
| PITPNB | 0.14 | 0.04 | 0.07 | -0.12 | -0.16 | 0.02 | 4.9E-02 | -0.17 |
| SLC25A22 | -0.02 | 0.09 | 0.11 | -0.05 | -0.09 | -0.05 | 4.9E-02 | -0.12 |
| DNAJC1 | 0.18 | 0.20 | -0.04 | -0.16 | -0.05 | -0.18 | 4.9E-02 | -0.24 |
| KRT17 | 0.01 | -0.21 | -0.31 | 0.21 | 0.19 | 0.03 | 5.0E-02 | 0.31 |
| CC2D1A | 0.02 | 0.04 | 0.15 | -0.13 | -0.07 | -0.02 | 5.0E-02 | -0.14 |
| TATDN1 | 0.16 | 0.19 | 0.00 | -0.26 | -0.06 | -0.08 | 5.0E-02 | -0.25 |
| RUVBL1 | 0.00 | -0.09 | -0.22 | 0.10 | 0.03 | 0.15 | 5.0E-02 | 0.20 |
| TRMT1L | -0.03 | -0.01 | -0.06 | 0.08 | 0.03 | 0.00 | 5.0E-02 | 0.07 |
| NPM3 | -0.16 | -0.18 | 0.04 | 0.10 | 0.09 | 0.08 | 5.0E-02 | 0.19 |
| ATP7B | -0.15 | -0.14 | 0.01 | 0.02 | 0.09 | 0.16 | 5.0E-02 | 0.18 |
| LMNB2 | -0.01 | -0.09 | -0.05 | 0.03 | 0.01 | 0.10 | 5.0E-02 | 0.10 |
| DIDO1 | 0.02 | 0.08 | 0.01 | -0.04 | -0.06 | -0.01 | 5.0E-02 | -0.07 |
| TFAP2C | -0.08 | -0.11 | -0.03 | 0.02 | 0.03 | 0.16 | 5.0E-02 | 0.14 |
| ELOVL5 | 0.08 | 0.06 | 0.43 | -0.14 | -0.42 | -0.14 | 5.0E-02 | -0.42 |
| CUL4B | -0.05 | -0.03 | -0.09 | 0.08 | -0.01 | 0.10 | 5.1E-02 | 0.11 |
| DNASE1L1 | -0.21 | -0.08 | 0.00 | 0.06 | 0.05 | 0.15 | 5.1E-02 | 0.18 |
| TSPAN4 | 0.08 | 0.02 | 0.44 | -0.08 | -0.29 | -0.30 | 5.1E-02 | -0.41 |
| BAK1 | -0.25 | -0.49 | -0.32 | 0.08 | 0.01 | 0.66 | 5.1E-02 | 0.60 |
| PLP2 | -0.08 | 0.20 | 0.55 | -0.40 | -0.30 | -0.20 | 5.1E-02 | -0.53 |
| OSBPL3 | 0.06 | 0.05 | 0.09 | -0.12 | 0.03 | -0.13 | 5.1E-02 | -0.14 |
| KHDRBS1 | 0.05 | 0.10 | 0.02 | -0.01 | -0.04 | -0.13 | 5.1E-02 | -0.12 |
| EEF1A2 | 0.07 | 0.04 | 0.06 | -0.02 | -0.02 | -0.14 | 5.1E-02 | -0.11 |
| PNISR | 0.08 | 0.09 | 0.08 | -0.13 | 0.03 | -0.18 | 5.1E-02 | -0.17 |
| VAMP4 | -0.04 | 0.14 | 0.41 | -0.12 | -0.27 | -0.24 | 5.1E-02 | -0.38 |
| MAEA | -0.09 | -0.19 | -0.04 | 0.20 | 0.08 | 0.00 | 5.1E-02 | 0.20 |
| C16orf87 | -0.01 | 0.04 | 0.03 | -0.03 | -0.03 | -0.01 | 5.1E-02 | -0.04 |
| TNRC18 | 0.07 | 0.13 | 0.01 | -0.16 | -0.03 | -0.04 | 5.1E-02 | -0.15 |

| | | | | | | | | |
|---|---|---|---|---|---|---|---|---|
| **XRN2** | 0.08 | 0.06 | 0.02 | -0.05 | 0.00 | -0.13 | 5.1E-02 | -0.11 |
| **SUCLA2** | -0.01 | -0.08 | -0.24 | 0.17 | 0.08 | 0.05 | 5.1E-02 | 0.21 |
| **ACOX1** | 0.02 | -0.14 | -0.14 | 0.04 | 0.06 | 0.14 | 5.1E-02 | 0.16 |
| **MRPL9** | 0.07 | 0.09 | 0.04 | -0.17 | 0.00 | -0.05 | 5.1E-02 | -0.14 |
| **CEBPZ** | 0.04 | 0.03 | 0.01 | -0.03 | 0.00 | -0.06 | 5.1E-02 | -0.06 |
| **RBMX** | -0.01 | -0.07 | -0.06 | 0.00 | 0.03 | 0.09 | 5.1E-02 | 0.09 |
| **CHTOP** | -0.14 | -0.14 | -0.01 | 0.18 | 0.00 | 0.08 | 5.1E-02 | 0.18 |
| **PUM1** | -0.09 | -0.10 | 0.02 | 0.09 | 0.02 | 0.06 | 5.1E-02 | 0.12 |
| **NUP42** | 0.07 | 0.04 | 0.00 | -0.02 | -0.02 | -0.07 | 5.2E-02 | -0.07 |
| **AP3M2** | 0.01 | 0.10 | 0.10 | -0.14 | 0.01 | -0.10 | 5.2E-02 | -0.15 |
| **CA12** | 0.00 | 0.10 | 0.72 | -0.55 | -0.42 | -0.22 | 5.2E-02 | -0.67 |
| **HMGN1** | -0.03 | -0.07 | -0.07 | -0.02 | 0.10 | 0.09 | 5.2E-02 | 0.11 |
| **PARP1** | 0.00 | -0.02 | -0.05 | 0.02 | 0.01 | 0.04 | 5.2E-02 | 0.05 |
| **GNA13** | -0.02 | 0.09 | 0.24 | -0.15 | -0.13 | -0.07 | 5.2E-02 | -0.22 |
| **SYS1** | 0.21 | 0.28 | 0.03 | -0.07 | -0.12 | -0.44 | 5.2E-02 | -0.38 |
| **RAD18** | 0.05 | 0.03 | 0.04 | -0.10 | -0.03 | 0.00 | 5.2E-02 | -0.08 |
| **DNMBP** | 0.20 | -0.03 | 0.41 | -0.43 | -0.13 | -0.17 | 5.2E-02 | -0.44 |
| **TMEM199** | 0.10 | 0.17 | 0.36 | -0.52 | 0.01 | -0.28 | 5.2E-02 | -0.47 |
| **NAALADL2** | 0.01 | 0.08 | 0.15 | -0.01 | -0.09 | -0.17 | 5.2E-02 | -0.17 |
| **NOL12** | 0.19 | 0.06 | 0.12 | 0.02 | -0.17 | -0.29 | 5.2E-02 | -0.27 |
| **NCAM2** | -0.07 | 0.21 | 0.42 | -0.29 | -0.31 | -0.11 | 5.2E-02 | -0.43 |
| **KDM6A** | -0.08 | -0.20 | -0.08 | 0.21 | -0.03 | 0.15 | 5.2E-02 | 0.23 |
| **GPR180** | 0.04 | 0.11 | 0.03 | -0.05 | 0.00 | -0.14 | 5.2E-02 | -0.13 |
| **SNX16** | -0.17 | -0.07 | -0.34 | 0.35 | 0.13 | 0.00 | 5.2E-02 | 0.35 |
| **CHD1L** | -0.07 | -0.01 | -0.02 | 0.02 | 0.06 | 0.01 | 5.2E-02 | 0.06 |
| **TMEM126A** | 0.03 | 0.03 | 0.04 | 0.01 | -0.07 | -0.03 | 5.2E-02 | -0.06 |
| **TTC9** | 0.11 | 0.18 | -0.01 | -0.15 | -0.01 | -0.14 | 5.2E-02 | -0.19 |
| **CPSF2** | -0.07 | -0.10 | -0.09 | 0.11 | -0.04 | 0.16 | 5.2E-02 | 0.16 |
| **JPH1** | 0.07 | 0.08 | 0.07 | 0.03 | -0.11 | -0.16 | 5.2E-02 | -0.16 |
| **LAS1L** | -0.01 | -0.02 | -0.01 | 0.03 | -0.01 | 0.02 | 5.2E-02 | 0.03 |
| **PTRHD1** | -0.06 | -0.08 | -0.21 | 0.13 | -0.02 | 0.20 | 5.3E-02 | 0.22 |
| **NSD2** | 0.10 | 0.09 | 0.06 | -0.21 | -0.08 | 0.01 | 5.3E-02 | -0.18 |
| **TRIM32** | -0.28 | -0.20 | 0.05 | 0.15 | 0.10 | 0.12 | 5.3E-02 | 0.27 |
| **MYADM** | 0.00 | 0.14 | 0.10 | -0.17 | -0.03 | -0.05 | 5.3E-02 | -0.16 |
| **POT1** | -0.19 | -0.31 | -0.31 | 0.33 | -0.13 | 0.42 | 5.3E-02 | 0.47 |
| **SRSF4** | 0.16 | 0.15 | 0.07 | -0.03 | -0.08 | -0.34 | 5.3E-02 | -0.28 |
| **ARCN1** | -0.06 | -0.02 | -0.03 | 0.08 | 0.02 | 0.01 | 5.3E-02 | 0.07 |
| **RAP1GAP** | -0.20 | -0.05 | -0.05 | 0.03 | 0.18 | 0.06 | 5.3E-02 | 0.19 |
| **KRT16** | -0.27 | -0.20 | -0.32 | 0.15 | -0.06 | 0.51 | 5.3E-02 | 0.46 |
| **SFN** | 0.13 | 0.14 | 0.00 | -0.05 | -0.05 | -0.21 | 5.3E-02 | -0.19 |
| **ARFGAP1** | -0.25 | -0.04 | -0.14 | -0.02 | 0.25 | 0.14 | 5.3E-02 | 0.27 |
| **PABPC1** | -0.05 | -0.03 | -0.03 | 0.03 | 0.08 | -0.01 | 5.3E-02 | 0.07 |
| **BBX** | 0.00 | 0.14 | 0.42 | -0.39 | -0.07 | -0.24 | 5.3E-02 | -0.42 |
| **BCS1L** | -0.03 | 0.13 | 0.11 | -0.06 | -0.05 | -0.12 | 5.3E-02 | -0.14 |
| **PRPF40A** | 0.18 | 0.16 | 0.09 | -0.05 | -0.06 | -0.40 | 5.3E-02 | -0.32 |
| **VIRMA** | 0.08 | 0.15 | 0.11 | -0.02 | -0.07 | -0.29 | 5.4E-02 | -0.24 |
| **POP1** | 0.06 | 0.11 | 0.08 | -0.04 | -0.02 | -0.22 | 5.4E-02 | -0.18 |
| **LLGL2** | -0.09 | -0.08 | 0.02 | 0.06 | 0.06 | 0.03 | 5.4E-02 | 0.10 |

| | | | | | | | | |
|---|---|---|---|---|---|---|---|---|
| EMC3 | 0.02 | 0.13 | 0.08 | -0.06 | -0.01 | -0.19 | 5.4E-02 | -0.17 |
| TMTC2 | 0.02 | 0.03 | 0.12 | -0.03 | -0.03 | -0.12 | 5.4E-02 | -0.12 |
| SMG6 | 0.10 | 0.18 | 0.07 | -0.28 | -0.12 | 0.01 | 5.4E-02 | -0.24 |
| GTF2F2 | 0.18 | 0.06 | -0.01 | -0.12 | -0.08 | -0.05 | 5.4E-02 | -0.16 |
| TIMM22 | 0.06 | 0.13 | 0.16 | -0.04 | -0.05 | -0.30 | 5.4E-02 | -0.25 |
| LIMS1 | -0.03 | -0.21 | -0.22 | -0.01 | 0.13 | 0.28 | 5.4E-02 | 0.28 |
| EIF6 | 0.05 | 0.09 | 0.19 | -0.23 | 0.02 | -0.15 | 5.4E-02 | -0.23 |
| FDFT1 | -0.06 | -0.11 | -0.07 | 0.07 | -0.02 | 0.18 | 5.4E-02 | 0.16 |
| RAB1A | 0.00 | 0.05 | 0.23 | -0.05 | -0.14 | -0.13 | 5.4E-02 | -0.20 |
| DNAJC19 | 0.06 | 0.15 | 0.02 | -0.15 | -0.09 | -0.01 | 5.4E-02 | -0.16 |
| ALDOA | 0.25 | 0.27 | -0.08 | -0.22 | -0.21 | -0.10 | 5.5E-02 | -0.32 |
| IDH2 | -0.02 | -0.07 | -0.16 | 0.06 | 0.16 | 0.01 | 5.5E-02 | 0.16 |
| HAUS1 | 0.07 | 0.12 | 0.07 | 0.04 | -0.16 | -0.16 | 5.5E-02 | -0.18 |
| TAX1BP1 | -0.22 | -0.04 | -0.01 | 0.11 | 0.06 | 0.08 | 5.5E-02 | 0.17 |
| RINT1 | 0.08 | 0.06 | 0.03 | -0.07 | 0.02 | -0.12 | 5.5E-02 | -0.11 |
| FAM98B | -0.16 | -0.13 | 0.01 | 0.15 | 0.01 | 0.09 | 5.5E-02 | 0.18 |
| UBE2NL | 0.40 | 0.21 | 0.09 | -0.54 | 0.05 | -0.44 | 5.5E-02 | -0.54 |
| ALG9 | 0.01 | 0.06 | 0.22 | -0.19 | -0.12 | -0.03 | 5.5E-02 | -0.21 |
| SAP18 | 0.09 | 0.04 | 0.05 | -0.01 | -0.03 | -0.15 | 5.5E-02 | -0.12 |
| ERCC1 | -0.28 | -0.06 | -0.20 | 0.10 | 0.00 | 0.34 | 5.5E-02 | 0.32 |
| DENND11 | -0.38 | 0.00 | -0.13 | 0.11 | 0.21 | 0.11 | 5.5E-02 | 0.31 |
| MRTFB | -0.09 | 0.00 | -0.05 | 0.04 | 0.08 | 0.01 | 5.5E-02 | 0.09 |
| GPATCH1 | 0.04 | 0.01 | 0.14 | -0.02 | -0.07 | -0.11 | 5.5E-02 | -0.13 |
| DCTPP1 | 0.05 | 0.16 | 0.00 | -0.12 | -0.09 | -0.02 | 5.5E-02 | -0.15 |
| H2BC12 | -0.03 | -0.11 | -0.07 | -0.02 | 0.12 | 0.10 | 5.5E-02 | 0.14 |
| MAP3K1 | -0.03 | 0.26 | 0.12 | -0.10 | -0.20 | -0.09 | 5.5E-02 | -0.24 |
| GRAMD4 | -0.11 | -0.05 | -0.01 | 0.03 | 0.11 | 0.02 | 5.5E-02 | 0.11 |
| PHKB | -0.28 | -0.28 | -0.08 | 0.36 | -0.07 | 0.23 | 5.5E-02 | 0.39 |
| SLC16A6 | -0.03 | 0.10 | 0.30 | -0.17 | -0.15 | -0.11 | 5.6E-02 | -0.27 |
| SHMT2 | 0.02 | -0.08 | -0.14 | 0.07 | 0.06 | 0.06 | 5.6E-02 | 0.13 |
| PITRM1 | 0.01 | -0.05 | -0.13 | 0.07 | 0.06 | 0.04 | 5.6E-02 | 0.11 |
| RRP7A | 0.06 | 0.07 | 0.05 | -0.11 | 0.03 | -0.11 | 5.6E-02 | -0.12 |
| HASPIN | -0.04 | -0.05 | -0.09 | 0.13 | -0.01 | 0.05 | 5.6E-02 | 0.11 |
| TRMU | -0.01 | -0.13 | -0.14 | 0.19 | 0.04 | 0.02 | 5.6E-02 | 0.18 |
| HEATR5B | -0.13 | -0.09 | -0.04 | -0.04 | 0.15 | 0.14 | 5.6E-02 | 0.17 |
| NOS1AP | -0.12 | -0.01 | -0.04 | 0.10 | 0.07 | 0.00 | 5.6E-02 | 0.12 |
| ACADVL | -0.03 | -0.08 | -0.19 | -0.01 | 0.10 | 0.17 | 5.6E-02 | 0.19 |
| MIA3 | 0.00 | -0.07 | -0.22 | 0.11 | 0.10 | 0.05 | 5.6E-02 | 0.18 |
| CDIPT | -0.04 | 0.14 | 0.34 | -0.11 | -0.26 | -0.15 | 5.6E-02 | -0.32 |
| ERN1 | 0.02 | -0.28 | -0.29 | 0.17 | 0.26 | 0.02 | 5.6E-02 | 0.33 |
| LYPLAL1 | -0.28 | -0.32 | -0.04 | 0.04 | 0.08 | 0.40 | 5.6E-02 | 0.38 |
| IMMP2L | 0.01 | 0.17 | 0.16 | -0.26 | -0.13 | 0.00 | 5.6E-02 | -0.24 |
| JUNB | -0.16 | -0.15 | -0.09 | 0.28 | -0.03 | 0.10 | 5.6E-02 | 0.25 |
| TBC1D10A | -0.15 | -0.08 | -0.01 | 0.13 | 0.09 | 0.00 | 5.6E-02 | 0.15 |
| GPX4 | -0.05 | -0.18 | -0.17 | -0.01 | 0.08 | 0.27 | 5.6E-02 | 0.25 |
| EMD | 0.03 | 0.02 | 0.03 | 0.01 | -0.04 | -0.06 | 5.6E-02 | -0.06 |
| ANKMY2 | 0.20 | 0.11 | 0.20 | -0.46 | 0.02 | -0.18 | 5.6E-02 | -0.38 |
| STRN4 | -0.02 | 0.05 | 0.07 | -0.04 | -0.04 | -0.03 | 5.6E-02 | -0.07 |

| | | | | | | | | |
|---|---|---|---|---|---|---|---|---|
| **PRPF6** | -0.01 | -0.07 | -0.05 | 0.00 | 0.03 | 0.09 | 5.6E-02 | 0.08 |
| **DDX23** | 0.08 | 0.05 | 0.04 | 0.01 | -0.05 | -0.14 | 5.6E-02 | -0.12 |
| **YIF1A** | 0.00 | 0.06 | 0.11 | -0.05 | -0.11 | -0.02 | 5.7E-02 | -0.11 |
| **U2SURP** | 0.13 | 0.07 | 0.06 | -0.05 | -0.01 | -0.22 | 5.7E-02 | -0.18 |
| **BARD1** | 0.02 | 0.15 | 0.11 | -0.23 | -0.04 | -0.04 | 5.7E-02 | -0.20 |
| **CCNE2** | 0.05 | 0.06 | 0.27 | -0.26 | -0.18 | 0.00 | 5.7E-02 | -0.27 |
| **DHX35** | 0.02 | 0.06 | 0.09 | -0.03 | -0.01 | -0.14 | 5.7E-02 | -0.12 |
| **ZNF445** | -0.05 | -0.17 | -0.15 | 0.07 | -0.01 | 0.26 | 5.7E-02 | 0.23 |
| **MRTO4** | 0.14 | 0.01 | 0.08 | -0.01 | -0.07 | -0.17 | 5.7E-02 | -0.16 |
| **PTGR3** | -0.11 | -0.21 | -0.20 | -0.08 | 0.21 | 0.30 | 5.7E-02 | 0.32 |
| **NSMCE4A** | 0.15 | 0.12 | -0.02 | -0.16 | -0.03 | -0.08 | 5.7E-02 | -0.18 |
| **LMX1B** | -0.28 | -0.16 | 0.01 | 0.15 | 0.19 | 0.02 | 5.7E-02 | 0.26 |
| **BSG** | -0.03 | 0.12 | 0.17 | -0.02 | -0.14 | -0.14 | 5.7E-02 | -0.19 |
| **RRS1** | 0.03 | 0.02 | 0.15 | -0.09 | -0.01 | -0.11 | 5.7E-02 | -0.14 |
| **CCT7** | 0.13 | 0.11 | -0.02 | -0.09 | -0.02 | -0.12 | 5.8E-02 | -0.15 |
| **MRPS17** | 0.18 | 0.09 | 0.01 | -0.19 | 0.01 | -0.13 | 5.8E-02 | -0.20 |
| **HK2** | -0.01 | -0.28 | -0.21 | -0.03 | 0.24 | 0.21 | 5.8E-02 | 0.31 |
| **DBT** | 0.07 | 0.17 | 0.01 | -0.02 | -0.07 | -0.18 | 5.8E-02 | -0.18 |
| **TRIO** | 0.02 | 0.04 | 0.23 | -0.11 | -0.18 | -0.04 | 5.8E-02 | -0.21 |
| **ELOF1** | 0.11 | 0.12 | 0.14 | -0.33 | 0.01 | -0.11 | 5.8E-02 | -0.27 |
| **POLR2G** | 0.03 | 0.03 | 0.10 | -0.08 | -0.11 | 0.01 | 5.8E-02 | -0.11 |
| **STK38** | -0.04 | -0.08 | -0.43 | 0.20 | 0.18 | 0.09 | 5.8E-02 | 0.34 |
| **ATG9B** | -0.15 | -0.07 | -0.54 | 0.36 | 0.04 | 0.19 | 5.8E-02 | 0.45 |
| **SFXN3** | -0.03 | 0.10 | 0.15 | -0.09 | -0.04 | -0.10 | 5.8E-02 | -0.15 |
| **DNAJA1** | -0.07 | -0.09 | -0.14 | 0.21 | 0.08 | -0.03 | 5.8E-02 | 0.19 |
| **APPL1** | -0.16 | -0.03 | -0.02 | 0.02 | 0.07 | 0.10 | 5.8E-02 | 0.13 |
| **ZBTB7A** | 0.06 | 0.08 | 0.04 | -0.13 | -0.09 | 0.02 | 5.8E-02 | -0.12 |
| **SLK** | 0.01 | -0.07 | -0.09 | 0.03 | 0.09 | 0.02 | 5.8E-02 | 0.10 |
| **ANXA9** | -0.17 | -0.29 | 0.03 | 0.08 | 0.09 | 0.21 | 5.8E-02 | 0.27 |
| **MRPL16** | 0.01 | 0.05 | 0.43 | -0.25 | -0.13 | -0.21 | 5.8E-02 | -0.36 |
| **SEC24B** | -0.03 | 0.09 | 0.18 | -0.13 | -0.06 | -0.08 | 5.8E-02 | -0.17 |
| **NT5C3A** | 0.02 | 0.08 | 0.06 | -0.13 | -0.02 | -0.02 | 5.8E-02 | -0.11 |
| **LDHA** | 0.30 | 0.19 | -0.08 | -0.16 | -0.19 | -0.14 | 5.9E-02 | -0.30 |
| **RHBDF2** | 0.08 | 0.15 | 0.10 | 0.03 | -0.14 | -0.27 | 5.9E-02 | -0.24 |
| **PPIG** | 0.24 | 0.31 | 0.07 | -0.16 | -0.04 | -0.58 | 5.9E-02 | -0.47 |
| **SAE1** | -0.02 | -0.04 | -0.10 | 0.10 | 0.00 | 0.05 | 5.9E-02 | 0.10 |
| **NPAT** | 0.15 | 0.33 | 0.11 | -0.28 | 0.04 | -0.50 | 5.9E-02 | -0.44 |
| **WBP11** | 0.01 | -0.11 | -0.13 | 0.08 | 0.00 | 0.12 | 5.9E-02 | 0.15 |
| **PNPT1** | 0.14 | 0.10 | 0.01 | -0.01 | -0.07 | -0.19 | 5.9E-02 | -0.17 |
| **VPS8** | -0.08 | -0.39 | -0.08 | 0.02 | 0.28 | 0.17 | 5.9E-02 | 0.34 |
| **UAP1L1** | -0.04 | -0.17 | -0.22 | 0.03 | 0.30 | 0.04 | 5.9E-02 | 0.27 |
| **PDILT** | -0.02 | 0.09 | 0.16 | -0.13 | -0.06 | -0.05 | 5.9E-02 | -0.15 |
| **WBP2** | 0.00 | 0.07 | 0.26 | -0.06 | -0.09 | -0.24 | 5.9E-02 | -0.24 |
| **RPS6KA4** | -0.39 | -0.20 | 0.00 | 0.02 | 0.19 | 0.27 | 5.9E-02 | 0.35 |
| **PMPCB** | 0.36 | 0.29 | -0.12 | -0.19 | -0.28 | -0.19 | 5.9E-02 | -0.40 |
| **EPHA4** | -0.05 | 0.24 | 0.35 | -0.41 | -0.18 | -0.08 | 5.9E-02 | -0.40 |
| **EFNA1** | 0.04 | 0.06 | 0.19 | -0.01 | -0.09 | -0.22 | 5.9E-02 | -0.20 |
| **SPIN2A** | 0.11 | 0.02 | 0.05 | -0.07 | -0.13 | 0.01 | 5.9E-02 | -0.13 |

| | | | | | | | | |
|---|---|---|---|---|---|---|---|---|
| **JAGN1** | 0.03 | 0.17 | 0.63 | -0.78 | -0.21 | -0.23 | 5.9E-02 | -0.68 |
| **CAPN13** | 0.22 | 0.19 | 0.06 | -0.08 | -0.05 | -0.44 | 5.9E-02 | -0.35 |
| **LSM14B** | 0.01 | 0.12 | 0.07 | -0.13 | 0.01 | -0.10 | 5.9E-02 | -0.14 |
| **INF2** | -0.14 | -0.08 | -0.01 | -0.01 | 0.10 | 0.13 | 6.0E-02 | 0.15 |
| **GTPBP8** | 0.15 | 0.04 | 0.08 | -0.02 | -0.05 | -0.23 | 6.0E-02 | -0.19 |
| **RBM47** | 0.02 | 0.03 | 0.13 | -0.11 | -0.01 | -0.06 | 6.0E-02 | -0.12 |
| **SMPD2** | -0.17 | -0.18 | 0.05 | 0.10 | 0.09 | 0.09 | 6.0E-02 | 0.20 |
| **STXBP1** | -0.04 | 0.10 | 0.21 | -0.14 | -0.10 | -0.07 | 6.0E-02 | -0.20 |
| **ETFB** | 0.02 | -0.07 | -0.14 | 0.06 | 0.09 | 0.03 | 6.0E-02 | 0.13 |
| **ITCH** | 0.01 | 0.08 | 0.26 | -0.14 | -0.03 | -0.24 | 6.0E-02 | -0.25 |
| **NOP14** | 0.04 | -0.01 | 0.07 | -0.03 | -0.01 | -0.06 | 6.0E-02 | -0.07 |
| **NKIRAS2** | -0.13 | -0.13 | -0.13 | 0.29 | 0.04 | 0.00 | 6.0E-02 | 0.24 |
| **LEMD2** | -0.06 | -0.11 | -0.03 | 0.01 | 0.02 | 0.16 | 6.0E-02 | 0.13 |
| **LIMK2** | -0.13 | -0.06 | -0.09 | -0.03 | 0.08 | 0.19 | 6.0E-02 | 0.17 |
| **KDM1A** | 0.00 | -0.04 | -0.06 | 0.04 | 0.00 | 0.05 | 6.0E-02 | 0.07 |
| **FMR1** | -0.07 | -0.10 | -0.06 | 0.03 | 0.18 | 0.00 | 6.0E-02 | 0.15 |
| **HEXB** | -0.08 | -0.26 | -0.15 | 0.31 | -0.05 | 0.15 | 6.0E-02 | 0.30 |
| **COX19** | -0.03 | 0.14 | 0.12 | -0.06 | -0.05 | -0.14 | 6.0E-02 | -0.16 |
| **NAP1L1** | 0.10 | 0.05 | 0.06 | -0.15 | 0.03 | -0.11 | 6.0E-02 | -0.15 |
| **FAR1** | 0.11 | 0.04 | 0.05 | -0.05 | 0.00 | -0.17 | 6.0E-02 | -0.14 |
| **DPH5** | -0.17 | -0.41 | -0.01 | 0.29 | 0.16 | 0.03 | 6.0E-02 | 0.36 |
| **GYS1** | -0.14 | -0.10 | -0.01 | 0.01 | 0.17 | 0.06 | 6.1E-02 | 0.16 |
| **SHCBP1** | -0.07 | -0.04 | -0.02 | -0.01 | 0.09 | 0.03 | 6.1E-02 | 0.08 |
| **BSPRY** | 0.01 | 0.08 | 0.61 | -0.27 | -0.18 | -0.51 | 6.1E-02 | -0.55 |
| **TMEM70** | -0.11 | -0.12 | -0.03 | 0.11 | -0.03 | 0.15 | 6.1E-02 | 0.16 |
| **IRF2BP2** | 0.25 | 0.19 | -0.07 | -0.09 | -0.13 | -0.21 | 6.1E-02 | -0.27 |
| **GUCD1** | -0.01 | -0.03 | -0.06 | 0.01 | 0.01 | 0.08 | 6.1E-02 | 0.07 |
| **CFDP1** | 0.35 | 0.17 | -0.05 | -0.10 | -0.14 | -0.33 | 6.1E-02 | -0.35 |
| **WFS1** | 0.01 | 0.02 | 0.13 | -0.02 | -0.07 | -0.09 | 6.1E-02 | -0.11 |
| **EYA3** | -0.04 | 0.10 | 0.17 | -0.07 | -0.11 | -0.07 | 6.1E-02 | -0.16 |
| **BHLHE40** | 0.09 | 0.02 | 0.16 | -0.19 | -0.13 | 0.02 | 6.1E-02 | -0.19 |
| **HELZ** | -0.14 | -0.07 | 0.02 | 0.09 | 0.04 | 0.04 | 6.1E-02 | 0.12 |
| **BCL2L1** | 0.07 | 0.01 | 0.09 | 0.01 | -0.07 | -0.12 | 6.1E-02 | -0.12 |
| **ATG9A** | 0.27 | 0.06 | 0.25 | -0.23 | 0.02 | -0.52 | 6.1E-02 | -0.44 |
| **NPEPPS** | -0.04 | -0.03 | -0.21 | 0.14 | 0.09 | 0.02 | 6.1E-02 | 0.18 |
| **ARVCF** | 0.01 | 0.04 | 0.18 | -0.15 | -0.06 | -0.05 | 6.1E-02 | -0.16 |
| **NCKIPSD** | -0.16 | -0.22 | 0.01 | -0.01 | 0.13 | 0.20 | 6.1E-02 | 0.23 |
| **GEMIN2** | 0.02 | 0.05 | 0.05 | -0.06 | 0.02 | -0.09 | 6.2E-02 | -0.08 |
| **DOCK7** | -0.08 | -0.03 | -0.04 | 0.10 | 0.05 | -0.01 | 6.2E-02 | 0.09 |
| **VPS35** | 0.01 | -0.04 | -0.05 | 0.03 | 0.04 | 0.01 | 6.2E-02 | 0.05 |
| **PRDX1** | 0.32 | 0.24 | -0.07 | -0.15 | -0.10 | -0.36 | 6.2E-02 | -0.37 |
| **FKBP3** | 0.03 | 0.21 | 0.25 | 0.02 | -0.20 | -0.42 | 6.2E-02 | -0.37 |
| **TMED1** | 0.05 | 0.01 | 0.19 | -0.01 | -0.10 | -0.17 | 6.2E-02 | -0.18 |
| **CORO1C** | 0.02 | 0.07 | 0.04 | -0.09 | -0.06 | 0.02 | 6.2E-02 | -0.09 |
| **NGRN** | 0.02 | 0.15 | 0.13 | -0.02 | -0.25 | -0.06 | 6.2E-02 | -0.21 |
| **SUSD2** | -0.06 | 0.16 | 0.39 | -0.26 | -0.25 | -0.09 | 6.2E-02 | -0.36 |
| **WAPL** | -0.07 | -0.03 | -0.07 | -0.03 | 0.08 | 0.10 | 6.2E-02 | 0.11 |
| **NUP54** | 0.16 | 0.16 | 0.10 | -0.06 | -0.03 | -0.40 | 6.2E-02 | -0.31 |

| | | | | | | | | |
|---|---|---|---|---|---|---|---|---|
| **TCAF2** | 0.02 | 0.18 | 0.17 | -0.02 | -0.09 | -0.32 | 6.2E-02 | -0.27 |
| **TBC1D17** | -0.14 | -0.13 | -0.03 | 0.17 | 0.14 | -0.04 | 6.2E-02 | 0.19 |
| **PRKCSH** | 0.20 | 0.44 | -0.10 | -0.27 | -0.26 | -0.16 | 6.3E-02 | -0.41 |
| **SDC1** | -0.02 | 0.11 | 0.11 | -0.02 | -0.11 | -0.08 | 6.3E-02 | -0.14 |
| **GGCT** | 0.18 | 0.29 | -0.08 | -0.12 | -0.15 | -0.18 | 6.3E-02 | -0.28 |
| **CBR3** | -0.38 | -0.12 | -0.02 | 0.15 | 0.24 | 0.04 | 6.3E-02 | 0.32 |
| **GTF3C5** | 0.00 | -0.09 | -0.15 | 0.16 | 0.05 | 0.01 | 6.3E-02 | 0.16 |
| **BLTP3B** | -0.10 | -0.03 | -0.16 | -0.03 | 0.17 | 0.12 | 6.3E-02 | 0.18 |
| **TUBGCP2** | -0.23 | -0.15 | -0.12 | 0.02 | 0.03 | 0.37 | 6.3E-02 | 0.31 |
| **FERMT2** | 0.11 | 0.19 | 0.05 | -0.18 | 0.04 | -0.26 | 6.3E-02 | -0.25 |
| **POLR3C** | -0.06 | -0.19 | -0.16 | 0.19 | 0.22 | -0.07 | 6.3E-02 | 0.25 |
| **EIF2S3** | 0.03 | 0.08 | 0.14 | -0.06 | 0.00 | -0.20 | 6.3E-02 | -0.17 |
| **USP10** | -0.11 | -0.05 | -0.08 | 0.19 | 0.02 | 0.01 | 6.4E-02 | 0.15 |
| **TPM1** | 0.08 | 0.03 | 0.05 | -0.07 | -0.11 | 0.02 | 6.4E-02 | -0.10 |
| **MGMT** | 0.11 | 0.14 | -0.01 | -0.18 | -0.06 | -0.03 | 6.4E-02 | -0.17 |
| **BOLA3** | 0.08 | 0.16 | 0.06 | -0.12 | 0.02 | -0.26 | 6.4E-02 | -0.22 |
| **DHX29** | 0.08 | 0.12 | 0.05 | -0.13 | 0.04 | -0.18 | 6.4E-02 | -0.17 |
| **LASP1** | -0.04 | -0.04 | -0.13 | 0.10 | -0.02 | 0.13 | 6.4E-02 | 0.14 |
| **HSD17B12** | 0.00 | -0.10 | -0.19 | 0.09 | 0.01 | 0.16 | 6.4E-02 | 0.18 |
| **MRPL45** | 0.12 | 0.08 | -0.01 | 0.01 | -0.10 | -0.12 | 6.4E-02 | -0.14 |
| **STAB1** | -0.35 | -0.47 | -0.34 | -0.05 | 0.07 | 0.75 | 6.4E-02 | 0.64 |
| **RAB9A** | 0.00 | 0.02 | 0.03 | 0.00 | -0.04 | -0.02 | 6.4E-02 | -0.03 |
| **NCOA5** | 0.01 | -0.05 | -0.06 | 0.01 | 0.04 | 0.05 | 6.4E-02 | 0.07 |
| **ZC3H11A** | 0.05 | 0.00 | 0.07 | -0.09 | -0.03 | -0.01 | 6.4E-02 | -0.08 |
| **SLC25A4** | -0.06 | 0.00 | -0.16 | 0.08 | 0.01 | 0.12 | 6.5E-02 | 0.14 |
| **RBM10** | 0.08 | 0.09 | 0.03 | -0.05 | 0.01 | -0.16 | 6.5E-02 | -0.14 |
| **KMT5C** | 0.03 | 0.03 | 0.04 | -0.06 | -0.07 | 0.02 | 6.5E-02 | -0.07 |
| **CDYL** | -0.05 | -0.06 | -0.08 | 0.00 | 0.03 | 0.16 | 6.5E-02 | 0.13 |
| **PACSIN3** | 0.00 | 0.07 | 0.21 | -0.04 | -0.18 | -0.07 | 6.5E-02 | -0.19 |
| **PAXBP1** | 0.02 | 0.03 | 0.05 | -0.08 | -0.03 | 0.01 | 6.5E-02 | -0.07 |
| **GET4** | 0.02 | -0.48 | -0.20 | 0.26 | 0.21 | 0.05 | 6.5E-02 | 0.40 |
| **MOGS** | 0.03 | 0.07 | 0.04 | -0.03 | 0.00 | -0.12 | 6.5E-02 | -0.10 |
| **IKZF5** | -0.11 | -0.10 | -0.07 | 0.05 | -0.01 | 0.22 | 6.5E-02 | 0.18 |
| **TBC1D15** | -0.16 | -0.16 | 0.02 | 0.17 | 0.06 | 0.03 | 6.5E-02 | 0.18 |
| **STAU2** | 0.05 | 0.05 | 0.06 | -0.07 | 0.02 | -0.11 | 6.5E-02 | -0.10 |
| **MAF1** | -0.18 | -0.25 | -0.29 | 0.38 | -0.14 | 0.32 | 6.5E-02 | 0.42 |
| **NELFB** | -0.10 | 0.01 | -0.11 | 0.04 | 0.01 | 0.14 | 6.5E-02 | 0.13 |
| **ECI2** | 0.01 | -0.13 | -0.16 | 0.09 | 0.16 | 0.01 | 6.6E-02 | 0.18 |
| **MRPL15** | 0.22 | 0.21 | -0.03 | -0.01 | -0.18 | -0.29 | 6.6E-02 | -0.29 |
| **ANXA2** | -0.02 | -0.07 | -0.03 | 0.10 | 0.02 | 0.00 | 6.6E-02 | 0.08 |
| **TXNDC17** | 0.10 | 0.02 | 0.28 | -0.12 | -0.33 | -0.03 | 6.6E-02 | -0.29 |
| **SH3GL1** | -0.03 | -0.04 | -0.11 | 0.00 | 0.13 | 0.04 | 6.6E-02 | 0.12 |
| **AMOTL1** | -0.16 | -0.15 | -0.01 | 0.08 | -0.01 | 0.22 | 6.6E-02 | 0.20 |
| **FN3K** | -0.01 | 0.05 | 0.18 | -0.07 | -0.05 | -0.11 | 6.6E-02 | -0.15 |
| **EXOC3** | -0.04 | -0.07 | -0.14 | -0.02 | 0.17 | 0.07 | 6.6E-02 | 0.16 |
| **SRRM1** | 0.11 | 0.14 | 0.02 | -0.07 | 0.00 | -0.22 | 6.6E-02 | -0.18 |
| **RPS16** | -0.06 | -0.10 | -0.04 | 0.16 | 0.03 | 0.00 | 6.6E-02 | 0.13 |
| **IRF2BP1** | -0.21 | -0.12 | 0.03 | 0.04 | 0.12 | 0.10 | 6.6E-02 | 0.19 |

| | | | | | | | | |
|---|---|---|---|---|---|---|---|---|
| **CEP120** | -0.23 | -0.23 | 0.05 | 0.20 | 0.12 | 0.04 | 6.6E-02 | 0.26 |
| **IFT20** | -0.29 | -0.23 | -0.02 | 0.28 | -0.05 | 0.22 | 6.6E-02 | 0.33 |
| **NUP107** | -0.02 | -0.17 | -0.02 | 0.02 | 0.09 | 0.08 | 6.6E-02 | 0.13 |
| **RNF139** | 0.12 | 0.01 | 0.22 | -0.28 | -0.01 | -0.10 | 6.6E-02 | -0.25 |
| **VDR** | 0.01 | 0.08 | 0.10 | 0.01 | -0.08 | -0.13 | 6.6E-02 | -0.13 |
| **UTP4** | 0.01 | 0.06 | 0.11 | -0.13 | 0.01 | -0.07 | 6.6E-02 | -0.12 |
| **GGA2** | -0.56 | -0.27 | -0.04 | -0.03 | 0.24 | 0.45 | 6.6E-02 | 0.51 |
| **MRPS6** | 0.13 | 0.07 | -0.01 | -0.14 | -0.01 | -0.06 | 6.6E-02 | -0.13 |
| **VPS29** | -0.03 | -0.03 | -0.07 | 0.10 | 0.01 | 0.02 | 6.6E-02 | 0.09 |
| **DNAJC30** | 0.03 | 0.01 | 0.17 | -0.13 | -0.02 | -0.08 | 6.6E-02 | -0.14 |
| **DDX27** | 0.01 | 0.02 | 0.04 | -0.01 | -0.01 | -0.07 | 6.7E-02 | -0.05 |
| **ACTR3** | 0.06 | 0.11 | 0.03 | 0.03 | -0.13 | -0.13 | 6.7E-02 | -0.15 |
| **RNASEH2A** | 0.01 | 0.03 | 0.08 | -0.10 | -0.01 | -0.02 | 6.7E-02 | -0.08 |
| **MLLT1** | 0.07 | 0.06 | -0.02 | -0.04 | -0.06 | -0.02 | 6.7E-02 | -0.08 |
| **AP1S2** | -0.45 | -0.27 | 0.07 | 0.21 | 0.19 | 0.12 | 6.7E-02 | 0.39 |
| **SYNE2** | 0.05 | 0.03 | 0.03 | -0.08 | -0.06 | 0.02 | 6.7E-02 | -0.08 |
| **SEMA3F** | 0.09 | 0.00 | 0.31 | -0.26 | -0.03 | -0.18 | 6.7E-02 | -0.29 |
| **ECHS1** | 0.01 | -0.06 | -0.18 | 0.08 | 0.10 | 0.04 | 6.7E-02 | 0.15 |
| **DAAM1** | -0.04 | 0.09 | 0.20 | -0.10 | -0.08 | -0.09 | 6.7E-02 | -0.17 |
| **AFP** | 0.03 | -0.44 | -0.29 | 0.37 | 0.09 | 0.10 | 6.7E-02 | 0.42 |
| **CSNK1A1** | 0.04 | 0.04 | 0.07 | 0.01 | -0.04 | -0.13 | 6.7E-02 | -0.11 |
| **UQCRFS1** | 0.05 | 0.02 | 0.02 | 0.01 | -0.04 | -0.07 | 6.7E-02 | -0.06 |
| **NFX1** | -0.19 | -0.07 | 0.00 | 0.14 | 0.10 | 0.01 | 6.7E-02 | 0.17 |
| **TYK2** | 0.04 | 0.04 | 0.09 | -0.16 | -0.01 | -0.02 | 6.7E-02 | -0.12 |
| **ETAA1** | -0.19 | -0.11 | -0.14 | -0.08 | 0.29 | 0.17 | 6.7E-02 | 0.28 |
| **DECR2** | -0.22 | -0.09 | -0.06 | 0.13 | -0.04 | 0.23 | 6.8E-02 | 0.23 |
| **KANK1** | -0.02 | -0.03 | -0.16 | 0.06 | 0.12 | 0.02 | 6.8E-02 | 0.14 |
| **VPS53** | -0.08 | 0.02 | -0.19 | 0.06 | 0.09 | 0.07 | 6.8E-02 | 0.16 |
| **YLPM1** | 0.08 | 0.12 | 0.07 | 0.01 | -0.07 | -0.25 | 6.8E-02 | -0.20 |
| **IDH3G** | -0.09 | -0.04 | 0.01 | 0.08 | 0.01 | 0.04 | 6.8E-02 | 0.08 |
| **RAB3GAP1** | -0.19 | -0.22 | 0.04 | 0.06 | 0.08 | 0.18 | 6.8E-02 | 0.23 |
| **SNW1** | 0.11 | 0.04 | -0.01 | -0.03 | -0.03 | -0.08 | 6.8E-02 | -0.09 |
| **TBRG1** | -0.39 | -0.18 | -0.55 | 0.05 | 0.01 | 0.71 | 6.8E-02 | 0.63 |
| **LEPROT** | -0.06 | 0.00 | -0.21 | 0.13 | 0.09 | 0.02 | 6.8E-02 | 0.17 |
| **CSNK1G1** | -0.56 | -0.15 | -0.40 | 0.26 | -0.13 | 0.63 | 6.8E-02 | 0.62 |
| **DAP3** | 0.14 | 0.12 | -0.02 | -0.18 | -0.02 | -0.07 | 6.8E-02 | -0.17 |
| **DENR** | 0.17 | 0.17 | -0.05 | -0.15 | -0.02 | -0.15 | 6.8E-02 | -0.21 |
| **EIF3D** | -0.03 | -0.07 | -0.05 | 0.07 | 0.11 | -0.03 | 6.8E-02 | 0.10 |
| **PBK** | -0.37 | -0.31 | 0.09 | 0.17 | 0.15 | 0.16 | 6.8E-02 | 0.36 |
| **MTARC2** | 0.01 | 0.02 | 0.09 | -0.03 | -0.02 | -0.07 | 6.9E-02 | -0.08 |
| **TNS3** | -0.30 | -0.40 | 0.00 | 0.04 | 0.42 | 0.10 | 6.9E-02 | 0.42 |
| **COA7** | -0.02 | 0.11 | 0.06 | -0.06 | -0.01 | -0.09 | 6.9E-02 | -0.11 |
| **MACROH2A2** | -0.06 | -0.11 | -0.09 | 0.05 | -0.02 | 0.20 | 6.9E-02 | 0.16 |
| **NKAP** | 0.13 | 0.08 | 0.07 | -0.06 | 0.00 | -0.26 | 6.9E-02 | -0.20 |
| **CENPP** | 0.07 | -0.03 | 0.14 | -0.05 | -0.06 | -0.07 | 6.9E-02 | -0.12 |
| **H1-4** | 0.01 | -0.03 | -0.10 | 0.05 | 0.04 | 0.03 | 6.9E-02 | 0.08 |
| **BOD1L1** | 0.19 | 0.01 | 0.14 | -0.16 | 0.03 | -0.26 | 6.9E-02 | -0.25 |
| **PCNP** | 0.11 | 0.13 | -0.03 | -0.08 | -0.03 | -0.13 | 6.9E-02 | -0.14 |

| | | | | | | | | |
|---|---|---|---|---|---|---|---|---|
| EIF4ENIF1 | -0.71 | -0.20 | -0.04 | 0.05 | 0.36 | 0.28 | 6.9E-02 | 0.55 |
| GTF2H1 | -0.12 | -0.11 | -0.15 | 0.31 | 0.02 | 0.01 | 7.0E-02 | 0.24 |
| PELP1 | 0.02 | 0.03 | 0.04 | 0.00 | -0.01 | -0.08 | 7.0E-02 | -0.06 |
| MPHOSPH8 | 0.05 | 0.02 | 0.04 | -0.09 | 0.01 | -0.04 | 7.0E-02 | -0.08 |
| TMEM45B | -0.17 | -0.04 | -0.07 | 0.20 | 0.07 | -0.01 | 7.0E-02 | 0.18 |
| ANKZF1 | -0.03 | -0.16 | -0.04 | -0.01 | 0.10 | 0.13 | 7.0E-02 | 0.15 |
| ADNP2 | 0.00 | -0.10 | -0.09 | -0.02 | 0.08 | 0.12 | 7.0E-02 | 0.13 |
| DNTTIP1 | -0.13 | -0.10 | 0.01 | 0.00 | 0.07 | 0.13 | 7.0E-02 | 0.14 |
| SUGP1 | -0.05 | -0.09 | -0.04 | -0.01 | 0.04 | 0.14 | 7.0E-02 | 0.11 |
| ZNF711 | -0.09 | -0.02 | -0.01 | 0.00 | 0.06 | 0.05 | 7.0E-02 | 0.08 |
| NUP93 | 0.19 | 0.16 | 0.14 | -0.03 | -0.08 | -0.49 | 7.0E-02 | -0.36 |
| CACYBP | 0.25 | 0.25 | -0.08 | -0.09 | -0.13 | -0.29 | 7.0E-02 | -0.31 |
| RABL3 | 0.03 | 0.04 | 0.16 | 0.01 | -0.12 | -0.14 | 7.0E-02 | -0.16 |
| BOP1 | 0.05 | 0.07 | 0.06 | -0.10 | 0.04 | -0.14 | 7.0E-02 | -0.13 |
| GLE1 | 0.07 | 0.03 | 0.02 | 0.02 | -0.07 | -0.07 | 7.0E-02 | -0.08 |
| SREK1 | 0.03 | 0.00 | 0.06 | -0.06 | -0.01 | -0.02 | 7.0E-02 | -0.06 |
| KRAS | -0.04 | 0.27 | 0.09 | -0.12 | -0.07 | -0.19 | 7.0E-02 | -0.23 |
| KAT6A | 0.01 | 0.01 | 0.05 | -0.05 | -0.01 | -0.01 | 7.0E-02 | -0.04 |
| WRAP73 | 0.08 | 0.10 | 0.05 | 0.04 | -0.15 | -0.14 | 7.0E-02 | -0.16 |
| ZNF579 | -0.12 | -0.11 | 0.02 | 0.00 | 0.08 | 0.12 | 7.0E-02 | 0.13 |
| ADAM17 | -0.26 | -0.08 | -0.19 | 0.28 | -0.09 | 0.26 | 7.0E-02 | 0.32 |
| MRPL44 | 0.03 | 0.03 | 0.00 | -0.01 | -0.04 | 0.00 | 7.0E-02 | -0.04 |
| ZNF16 | 0.01 | 0.03 | 0.35 | -0.16 | -0.10 | -0.19 | 7.0E-02 | -0.28 |
| ZNF467 | 0.05 | 0.11 | -0.02 | -0.03 | -0.09 | -0.04 | 7.1E-02 | -0.10 |
| GOLGA2 | -0.09 | -0.04 | -0.04 | -0.03 | 0.10 | 0.09 | 7.1E-02 | 0.11 |
| NSDHL | 0.03 | -0.09 | -0.13 | 0.09 | 0.02 | 0.07 | 7.1E-02 | 0.12 |
| SMARCE1 | 0.09 | 0.09 | -0.02 | -0.09 | -0.01 | -0.08 | 7.1E-02 | -0.11 |
| NAA50 | 0.20 | 0.13 | -0.04 | -0.05 | -0.08 | -0.20 | 7.1E-02 | -0.21 |
| MAT2A | -0.24 | -0.10 | -0.08 | -0.04 | 0.28 | 0.11 | 7.1E-02 | 0.26 |
| GPALPP1 | 0.15 | 0.20 | -0.03 | -0.17 | 0.00 | -0.18 | 7.1E-02 | -0.22 |
| CDC40 | 0.13 | 0.06 | 0.01 | -0.12 | 0.01 | -0.09 | 7.1E-02 | -0.13 |
| NSMCE2 | -0.03 | -0.05 | -0.04 | 0.01 | 0.01 | 0.10 | 7.1E-02 | 0.07 |
| VAMP7 | -0.04 | 0.11 | 0.41 | -0.33 | -0.12 | -0.14 | 7.1E-02 | -0.36 |
| PLRG1 | 0.11 | 0.11 | 0.04 | -0.21 | 0.04 | -0.13 | 7.1E-02 | -0.19 |
| RALBP1 | -0.02 | -0.15 | -0.06 | 0.01 | 0.16 | 0.04 | 7.1E-02 | 0.15 |
| MPHOSPH6 | 0.13 | 0.07 | 0.09 | 0.03 | -0.11 | -0.25 | 7.1E-02 | -0.20 |
| DDX60 | -0.15 | -0.09 | -0.07 | -0.06 | 0.18 | 0.15 | 7.1E-02 | 0.19 |
| SPNS1 | 0.06 | 0.08 | 0.41 | -0.49 | -0.07 | -0.14 | 7.1E-02 | -0.42 |
| NCBP1 | 0.01 | 0.05 | 0.11 | -0.14 | -0.01 | -0.04 | 7.1E-02 | -0.12 |
| GPI | -0.01 | -0.08 | -0.03 | 0.02 | 0.00 | 0.09 | 7.2E-02 | 0.08 |
| TARBP1 | -0.02 | -0.38 | -0.17 | 0.32 | -0.01 | 0.15 | 7.2E-02 | 0.34 |
| ITGA6 | -0.04 | 0.11 | 0.23 | -0.17 | -0.08 | -0.08 | 7.2E-02 | -0.21 |
| SMAP1 | 0.04 | 0.09 | 0.09 | -0.01 | -0.20 | -0.04 | 7.2E-02 | -0.16 |
| EXO1 | 0.01 | 0.09 | 0.19 | -0.18 | 0.03 | -0.18 | 7.2E-02 | -0.21 |
| NARS2 | 0.00 | -0.06 | -0.25 | 0.07 | 0.07 | 0.14 | 7.2E-02 | 0.19 |
| TAF12 | -0.02 | -0.07 | -0.12 | 0.11 | -0.03 | 0.12 | 7.2E-02 | 0.14 |
| E4F1 | -0.12 | -0.06 | -0.07 | -0.05 | 0.12 | 0.16 | 7.2E-02 | 0.16 |
| TMEM259 | -0.34 | -0.22 | 0.04 | 0.03 | 0.14 | 0.27 | 7.2E-02 | 0.32 |

| | | | | | | | | |
|---|---|---|---|---|---|---|---|---|
| **EPB41L5** | -0.06 | 0.00 | -0.02 | 0.05 | 0.03 | 0.00 | 7.2E-02 | 0.05 |
| **PHLDB2** | 0.04 | -0.22 | -0.26 | 0.22 | 0.01 | 0.14 | 7.2E-02 | 0.27 |
| **HMGCS1** | -0.02 | -0.03 | -0.11 | 0.04 | 0.11 | 0.00 | 7.2E-02 | 0.11 |
| **PARK7** | 0.16 | 0.16 | -0.05 | -0.05 | -0.10 | -0.15 | 7.2E-02 | -0.19 |
| **EXOSC7** | 0.13 | 0.06 | 0.02 | -0.01 | -0.04 | -0.17 | 7.2E-02 | -0.14 |
| **EXTL3** | 0.02 | 0.05 | 0.05 | 0.02 | -0.05 | -0.09 | 7.2E-02 | -0.08 |
| **TMEM63B** | -0.20 | -0.12 | -0.13 | 0.18 | -0.09 | 0.28 | 7.3E-02 | 0.27 |
| **YARS2** | -0.03 | -0.05 | -0.13 | 0.02 | 0.01 | 0.16 | 7.3E-02 | 0.14 |
| **ST14** | 0.05 | 0.03 | 0.34 | -0.32 | -0.13 | -0.05 | 7.3E-02 | -0.31 |
| **VRK2** | -0.10 | -0.09 | -0.02 | 0.07 | -0.02 | 0.15 | 7.3E-02 | 0.13 |
| **CCAR1** | 0.25 | 0.30 | 0.19 | -0.04 | -0.16 | -0.80 | 7.3E-02 | -0.58 |
| **GCDH** | 0.05 | -0.18 | -0.18 | 0.04 | 0.12 | 0.12 | 7.3E-02 | 0.20 |
| **SGO2** | -0.43 | 0.02 | -0.21 | 0.06 | 0.33 | 0.10 | 7.3E-02 | 0.37 |
| **BID** | -0.18 | -0.15 | 0.00 | 0.22 | 0.02 | 0.05 | 7.3E-02 | 0.20 |
| **C18orf21** | 0.16 | 0.14 | 0.01 | 0.00 | -0.09 | -0.28 | 7.3E-02 | -0.23 |
| **ARHGAP8** | -0.20 | -0.21 | -0.01 | -0.03 | 0.12 | 0.27 | 7.3E-02 | 0.26 |
| **CGN** | -0.03 | 0.07 | 0.09 | -0.08 | -0.03 | -0.04 | 7.3E-02 | -0.10 |
| **TSNAX** | -0.18 | -0.27 | -0.10 | 0.41 | -0.02 | 0.06 | 7.4E-02 | 0.33 |
| **BANP** | -0.08 | -0.11 | -0.08 | 0.03 | -0.01 | 0.22 | 7.4E-02 | 0.17 |
| **RBM27** | -0.03 | 0.11 | 0.14 | -0.01 | -0.11 | -0.12 | 7.4E-02 | -0.15 |
| **VDAC3** | 0.00 | -0.08 | -0.16 | 0.16 | 0.03 | 0.03 | 7.4E-02 | 0.15 |
| **PGAM1** | 0.06 | 0.10 | -0.01 | -0.01 | -0.05 | -0.10 | 7.4E-02 | -0.10 |
| **MRPL42** | 0.18 | 0.27 | -0.04 | -0.22 | 0.00 | -0.27 | 7.4E-02 | -0.30 |
| **SEC31A** | 0.01 | -0.04 | -0.15 | 0.07 | 0.07 | 0.03 | 7.4E-02 | 0.12 |
| **PUM2** | 0.05 | -0.01 | 0.20 | -0.13 | -0.03 | -0.11 | 7.4E-02 | -0.17 |
| **RPS11** | -0.03 | -0.06 | -0.05 | -0.01 | 0.11 | 0.02 | 7.5E-02 | 0.09 |
| **MRPL39** | 0.15 | 0.11 | 0.02 | 0.02 | -0.10 | -0.22 | 7.5E-02 | -0.19 |
| **KNOP1** | 0.08 | 0.08 | 0.01 | 0.01 | -0.06 | -0.14 | 7.5E-02 | -0.12 |
| **CACNA1S** | -0.12 | 0.41 | 0.39 | -0.31 | -0.06 | -0.56 | 7.5E-02 | -0.54 |
| **RBBP5** | 0.07 | 0.04 | -0.01 | 0.00 | -0.05 | -0.04 | 7.5E-02 | -0.06 |
| **CSE1L** | 0.01 | -0.04 | -0.10 | 0.03 | 0.07 | 0.03 | 7.5E-02 | 0.08 |
| **POLB** | -0.14 | -0.16 | -0.07 | 0.28 | -0.05 | 0.10 | 7.5E-02 | 0.23 |
| **UBE2S** | 0.00 | -0.30 | -0.32 | -0.02 | 0.14 | 0.37 | 7.5E-02 | 0.37 |
| **ATP8B1** | -0.04 | 0.00 | -0.05 | 0.06 | -0.01 | 0.04 | 7.5E-02 | 0.06 |
| **PPP6C** | -0.05 | -0.16 | -0.24 | 0.33 | 0.00 | 0.05 | 7.5E-02 | 0.28 |
| **KANSL3** | 0.03 | 0.06 | 0.13 | -0.18 | -0.06 | 0.01 | 7.5E-02 | -0.15 |
| **SC5D** | 0.00 | 0.05 | 0.21 | -0.07 | -0.18 | -0.04 | 7.5E-02 | -0.19 |
| **DCTN2** | -0.06 | -0.16 | 0.01 | 0.03 | 0.04 | 0.12 | 7.5E-02 | 0.14 |
| **POLR2D** | 0.16 | 0.08 | -0.02 | -0.12 | 0.00 | -0.12 | 7.6E-02 | -0.15 |
| **SEH1L** | 0.07 | 0.00 | 0.18 | -0.03 | -0.05 | -0.19 | 7.6E-02 | -0.17 |
| **ASH1L** | 0.11 | -0.06 | 0.24 | -0.10 | -0.10 | -0.14 | 7.6E-02 | -0.21 |
| **DAD1** | -0.02 | 0.10 | 0.11 | 0.00 | -0.12 | -0.08 | 7.6E-02 | -0.13 |
| **GPX1** | -0.28 | -0.02 | -0.09 | 0.03 | 0.06 | 0.24 | 7.6E-02 | 0.24 |
| **RPL23** | -0.05 | -0.10 | -0.04 | 0.16 | 0.02 | 0.01 | 7.6E-02 | 0.12 |
| **PDCD7** | 0.12 | 0.02 | 0.12 | -0.15 | 0.04 | -0.18 | 7.6E-02 | -0.18 |
| **TPCN1** | -0.21 | -0.16 | -0.01 | 0.19 | -0.05 | 0.19 | 7.6E-02 | 0.24 |
| **ZNF319** | -0.33 | -0.33 | 0.02 | 0.11 | 0.01 | 0.39 | 7.6E-02 | 0.38 |
| **COIL** | 0.13 | 0.17 | 0.01 | -0.12 | 0.02 | -0.26 | 7.6E-02 | -0.22 |

| | | | | | | | | |
|---|---|---|---|---|---|---|---|---|
| TUBB | -0.06 | -0.21 | -0.38 | 0.44 | 0.04 | 0.04 | 7.6E-02 | 0.39 |
| PPHLN1 | 0.10 | 0.07 | 0.01 | -0.04 | 0.00 | -0.15 | 7.6E-02 | -0.12 |
| SDC2 | -0.03 | 0.19 | 0.12 | 0.00 | -0.14 | -0.18 | 7.6E-02 | -0.20 |
| MACROH2A1 | -0.06 | -0.09 | -0.02 | -0.02 | 0.05 | 0.13 | 7.6E-02 | 0.11 |
| MORC2 | -0.39 | -0.27 | -0.27 | 0.56 | -0.19 | 0.29 | 7.6E-02 | 0.53 |
| TMED3 | -0.01 | 0.06 | 0.13 | -0.13 | -0.03 | -0.03 | 7.6E-02 | -0.12 |
| MYO5B | -0.01 | -0.04 | -0.04 | 0.06 | 0.00 | 0.01 | 7.6E-02 | 0.05 |
| WDR76 | -0.02 | -0.07 | -0.13 | 0.12 | -0.04 | 0.12 | 7.6E-02 | 0.14 |
| TFIP11 | 0.02 | 0.00 | 0.07 | -0.06 | 0.00 | -0.04 | 7.6E-02 | -0.06 |
| IGF2R | 0.01 | 0.03 | 0.15 | -0.03 | -0.04 | -0.15 | 7.6E-02 | -0.14 |
| CDCA7L | 0.07 | 0.24 | 0.25 | -0.55 | -0.15 | 0.01 | 7.7E-02 | -0.41 |
| BAX | -0.13 | -0.18 | 0.04 | 0.05 | 0.07 | 0.12 | 7.7E-02 | 0.17 |
| AP2S1 | -0.15 | -0.07 | -0.04 | -0.03 | 0.09 | 0.18 | 7.7E-02 | 0.17 |
| RPS6 | -0.09 | -0.12 | 0.00 | 0.14 | 0.00 | 0.05 | 7.7E-02 | 0.13 |
| PAFAH1B3 | -0.13 | -0.10 | 0.01 | -0.01 | 0.12 | 0.08 | 7.7E-02 | 0.14 |
| PES1 | 0.07 | 0.02 | 0.05 | -0.03 | 0.01 | -0.11 | 7.7E-02 | -0.09 |
| CCP110 | -0.07 | -0.06 | 0.00 | -0.01 | 0.05 | 0.10 | 7.7E-02 | 0.09 |
| RAB5IF | 0.06 | 0.13 | 0.05 | -0.02 | -0.03 | -0.21 | 7.7E-02 | -0.16 |
| SLC12A9 | -0.03 | 0.07 | 0.25 | -0.12 | -0.04 | -0.16 | 7.7E-02 | -0.20 |
| MRPL41 | 0.16 | 0.21 | -0.07 | -0.05 | -0.12 | -0.17 | 7.7E-02 | -0.22 |
| RGS10 | 0.05 | 0.15 | 0.14 | 0.03 | -0.31 | -0.11 | 7.7E-02 | -0.24 |
| TMEM39B | 0.14 | 0.14 | 0.02 | -0.16 | 0.05 | -0.22 | 7.7E-02 | -0.21 |
| HSP90AA1 | 0.00 | -0.02 | -0.13 | 0.06 | 0.05 | 0.03 | 7.8E-02 | 0.10 |
| CCDC47 | 0.07 | 0.10 | 0.10 | -0.06 | 0.01 | -0.25 | 7.8E-02 | -0.19 |
| CNTROB | 0.10 | 0.09 | -0.04 | -0.07 | -0.05 | -0.04 | 7.8E-02 | -0.11 |
| GOT2 | 0.36 | 0.27 | -0.13 | -0.21 | -0.26 | -0.14 | 7.8E-02 | -0.37 |
| TMEM39A | -0.02 | 0.09 | 0.35 | -0.14 | -0.29 | -0.07 | 7.8E-02 | -0.31 |
| LRRC40 | 0.12 | -0.03 | 0.08 | -0.11 | -0.05 | -0.03 | 7.8E-02 | -0.12 |
| SLC27A2 | 0.01 | 0.02 | 0.09 | -0.06 | 0.01 | -0.06 | 7.8E-02 | -0.08 |
| DEPDC1B | -0.16 | -0.11 | -0.02 | 0.07 | -0.02 | 0.20 | 7.8E-02 | 0.18 |
| UFSP2 | 0.04 | 0.25 | 0.09 | -0.14 | 0.01 | -0.32 | 7.8E-02 | -0.27 |
| CHMP4B | -0.04 | -0.04 | -0.01 | 0.07 | -0.01 | 0.03 | 7.8E-02 | 0.06 |
| ZC3H7A | -0.15 | -0.08 | 0.01 | 0.04 | 0.14 | 0.03 | 7.8E-02 | 0.14 |
| MARK2 | -0.05 | -0.13 | -0.01 | 0.02 | 0.02 | 0.13 | 7.8E-02 | 0.12 |
| AIF1L | 0.06 | 0.05 | 0.29 | 0.03 | -0.26 | -0.24 | 7.8E-02 | -0.29 |
| RAB3A | -0.03 | -0.18 | -0.11 | 0.01 | 0.03 | 0.23 | 7.8E-02 | 0.20 |
| DPF2 | 0.12 | 0.07 | 0.06 | -0.23 | -0.01 | -0.02 | 7.8E-02 | -0.17 |
| PHIP | -0.04 | -0.09 | -0.05 | 0.02 | 0.00 | 0.15 | 7.8E-02 | 0.12 |
| NR1H2 | -0.30 | -0.29 | -0.01 | 0.28 | -0.08 | 0.28 | 7.8E-02 | 0.36 |
| PARP2 | -0.07 | -0.03 | 0.00 | 0.03 | 0.00 | 0.07 | 7.8E-02 | 0.07 |
| RRP9 | 0.07 | 0.07 | 0.00 | -0.01 | -0.02 | -0.11 | 7.9E-02 | -0.09 |
| SGO1 | -0.06 | -0.17 | -0.04 | -0.02 | 0.08 | 0.18 | 7.9E-02 | 0.17 |
| HSP90B1 | -0.01 | -0.03 | -0.13 | 0.09 | 0.04 | 0.01 | 7.9E-02 | 0.10 |
| SLC25A13 | -0.01 | 0.06 | 0.05 | -0.01 | -0.03 | -0.07 | 7.9E-02 | -0.07 |
| DIMT1 | 0.07 | 0.03 | 0.10 | -0.08 | 0.03 | -0.16 | 7.9E-02 | -0.13 |
| MBTD1 | -0.11 | -0.07 | -0.11 | 0.11 | -0.06 | 0.21 | 7.9E-02 | 0.18 |
| BCLAF1 | 0.08 | 0.10 | 0.03 | -0.08 | 0.03 | -0.18 | 7.9E-02 | -0.14 |
| SREK1IP1 | 0.25 | 0.09 | 0.10 | -0.41 | -0.11 | 0.00 | 7.9E-02 | -0.32 |

| | | | | | | | | |
|---|---|---|---|---|---|---|---|---|
| RNASEL | -0.07 | -0.20 | 0.02 | 0.03 | 0.13 | 0.06 | 7.9E-02 | 0.16 |
| POGK | 0.02 | -0.08 | -0.18 | 0.05 | 0.13 | 0.04 | 7.9E-02 | 0.15 |
| ACOT9 | 0.02 | -0.14 | -0.20 | 0.20 | 0.10 | 0.00 | 7.9E-02 | 0.21 |
| SP1 | 0.11 | 0.09 | -0.03 | -0.09 | -0.06 | -0.02 | 7.9E-02 | -0.12 |
| TIMM21 | -0.22 | -0.18 | -0.10 | 0.13 | -0.07 | 0.36 | 7.9E-02 | 0.30 |
| ETFDH | 0.03 | 0.05 | 0.17 | 0.02 | -0.19 | -0.11 | 7.9E-02 | -0.18 |
| IARS2 | 0.02 | -0.06 | -0.10 | 0.07 | 0.02 | 0.04 | 7.9E-02 | 0.09 |
| RPS27 | -0.13 | -0.15 | 0.04 | 0.11 | 0.08 | 0.03 | 7.9E-02 | 0.15 |
| GLUD1 | 0.18 | 0.14 | -0.06 | -0.07 | -0.06 | -0.17 | 7.9E-02 | -0.19 |
| DROSHA | 0.00 | 0.01 | 0.05 | -0.04 | -0.01 | -0.01 | 8.0E-02 | -0.04 |
| TERF1 | 0.00 | -0.04 | -0.05 | -0.01 | 0.04 | 0.05 | 8.0E-02 | 0.06 |
| EXPH5 | 0.00 | 0.03 | 0.05 | -0.05 | -0.01 | -0.01 | 8.0E-02 | -0.05 |
| MRPL13 | 0.13 | 0.13 | -0.05 | -0.08 | -0.06 | -0.09 | 8.0E-02 | -0.14 |
| PPP6R3 | -0.06 | -0.06 | -0.20 | -0.04 | 0.20 | 0.13 | 8.0E-02 | 0.20 |
| TPT1 | -0.03 | -0.05 | -0.21 | 0.02 | 0.19 | 0.05 | 8.0E-02 | 0.18 |
| RBM17 | 0.03 | 0.00 | 0.05 | 0.00 | -0.05 | -0.03 | 8.0E-02 | -0.05 |
| ARHGAP17 | -0.08 | -0.03 | -0.11 | -0.04 | 0.10 | 0.15 | 8.0E-02 | 0.14 |
| MCM7 | 0.15 | -0.02 | 0.07 | -0.07 | -0.02 | -0.12 | 8.0E-02 | -0.13 |
| DCAKD | -0.01 | -0.14 | -0.09 | -0.03 | 0.15 | 0.10 | 8.0E-02 | 0.15 |
| ZCCHC17 | 0.18 | 0.03 | 0.00 | -0.05 | -0.05 | -0.13 | 8.0E-02 | -0.14 |
| TBCB | -0.17 | -0.14 | -0.15 | -0.08 | 0.34 | 0.12 | 8.0E-02 | 0.28 |
| GTPBP4 | 0.12 | 0.05 | 0.04 | -0.04 | 0.00 | -0.19 | 8.0E-02 | -0.14 |
| PRR15L | -0.11 | -0.04 | -0.08 | -0.01 | 0.04 | 0.18 | 8.0E-02 | 0.14 |
| TUSC3 | 0.05 | 0.07 | 0.11 | -0.06 | 0.02 | -0.21 | 8.1E-02 | -0.16 |
| SCMH1 | 0.00 | -0.08 | -0.04 | 0.01 | 0.08 | 0.03 | 8.1E-02 | 0.07 |
| BCAP29 | 0.10 | 0.10 | -0.04 | -0.06 | -0.03 | -0.09 | 8.1E-02 | -0.11 |
| XPNPEP1 | 0.02 | -0.27 | -0.26 | 0.09 | 0.33 | 0.01 | 8.1E-02 | 0.31 |
| RANBP1 | -0.10 | -0.12 | 0.01 | 0.11 | -0.02 | 0.10 | 8.1E-02 | 0.13 |
| GOLGB1 | 0.05 | 0.10 | 0.12 | 0.05 | -0.14 | -0.22 | 8.1E-02 | -0.19 |
| SLC25A35 | -0.08 | 0.01 | -0.06 | 0.08 | 0.04 | 0.01 | 8.1E-02 | 0.08 |
| MYO1E | -0.01 | 0.03 | 0.14 | -0.09 | -0.05 | -0.03 | 8.1E-02 | -0.11 |
| PARG | 0.02 | 0.04 | 0.26 | -0.24 | -0.05 | -0.07 | 8.1E-02 | -0.23 |
| GOLT1B | 0.04 | -0.02 | 0.28 | -0.11 | -0.13 | -0.11 | 8.1E-02 | -0.21 |
| MYO5C | -0.05 | -0.05 | -0.01 | 0.01 | 0.01 | 0.10 | 8.1E-02 | 0.08 |
| WTAP | 0.15 | 0.11 | 0.05 | 0.00 | -0.05 | -0.30 | 8.1E-02 | -0.23 |
| S100A16 | -0.17 | -0.14 | 0.00 | 0.04 | 0.02 | 0.22 | 8.1E-02 | 0.19 |
| SAFB | 0.06 | 0.02 | -0.01 | -0.02 | -0.03 | -0.02 | 8.1E-02 | -0.05 |
| SMARCAD1 | 0.04 | -0.13 | -0.40 | 0.09 | 0.16 | 0.16 | 8.2E-02 | 0.30 |
| CUL4A | -0.06 | -0.15 | -0.08 | 0.07 | -0.03 | 0.22 | 8.2E-02 | 0.18 |
| NOTCH3 | -0.05 | 0.11 | 0.20 | -0.11 | -0.14 | -0.04 | 8.2E-02 | -0.18 |
| ALDH1B1 | -0.02 | -0.04 | -0.15 | 0.02 | 0.14 | 0.03 | 8.2E-02 | 0.14 |
| ZNF845 | 0.22 | 0.08 | -0.04 | -0.04 | -0.14 | -0.12 | 8.2E-02 | -0.19 |
| CAMK2G | -0.21 | -0.28 | 0.01 | 0.31 | 0.10 | -0.01 | 8.2E-02 | 0.29 |
| TNPO3 | -0.22 | -0.21 | 0.01 | 0.14 | -0.03 | 0.24 | 8.2E-02 | 0.25 |
| SLC25A33 | 0.01 | 0.08 | 0.12 | -0.17 | -0.08 | 0.02 | 8.2E-02 | -0.15 |
| ZNF678 | 0.28 | 0.07 | 0.17 | -0.04 | -0.08 | -0.52 | 8.2E-02 | -0.39 |
| HSPA4 | 0.00 | 0.07 | 0.05 | -0.07 | 0.02 | -0.08 | 8.2E-02 | -0.08 |
| DDX54 | 0.10 | 0.03 | 0.01 | 0.00 | -0.04 | -0.10 | 8.2E-02 | -0.09 |

| | | | | | | | | |
|---|---|---|---|---|---|---|---|---|
| PCNX3 | -0.02 | 0.05 | 0.07 | -0.06 | -0.01 | -0.04 | 8.2E-02 | -0.07 |
| PPIL2 | 0.09 | 0.04 | -0.01 | -0.08 | 0.00 | -0.04 | 8.3E-02 | -0.08 |
| PAXIP1 | -0.05 | -0.12 | -0.05 | 0.04 | -0.01 | 0.18 | 8.3E-02 | 0.14 |
| SMAD1 | 0.15 | 0.13 | 0.17 | -0.24 | 0.09 | -0.39 | 8.3E-02 | -0.33 |
| POLR1E | -0.01 | -0.11 | -0.07 | -0.01 | 0.13 | 0.05 | 8.3E-02 | 0.12 |
| GLO1 | 0.07 | 0.19 | -0.04 | -0.09 | -0.08 | -0.08 | 8.3E-02 | -0.16 |
| CENPN | 0.09 | 0.11 | 0.07 | -0.25 | -0.03 | 0.00 | 8.3E-02 | -0.18 |
| SERPINF1 | -0.21 | -0.43 | -0.02 | 0.15 | -0.03 | 0.39 | 8.3E-02 | 0.39 |
| FAF2 | -0.03 | 0.05 | 0.15 | -0.03 | -0.10 | -0.07 | 8.3E-02 | -0.13 |
| SNRPE | 0.00 | -0.07 | -0.14 | 0.04 | 0.01 | 0.15 | 8.3E-02 | 0.14 |
| VPS26A | -0.01 | -0.13 | -0.07 | 0.16 | 0.05 | -0.01 | 8.4E-02 | 0.14 |
| SEC23A | 0.10 | 0.16 | 0.02 | -0.06 | 0.00 | -0.25 | 8.4E-02 | -0.20 |
| RPGRIP1L | 0.14 | 0.27 | 0.02 | -0.23 | 0.05 | -0.35 | 8.4E-02 | -0.32 |
| OR2J2 | -0.03 | 0.07 | 0.14 | -0.12 | -0.05 | -0.03 | 8.4E-02 | -0.13 |
| UFM1 | 0.05 | 0.06 | 0.23 | -0.19 | 0.04 | -0.24 | 8.4E-02 | -0.24 |
| ATP5MF | -0.55 | -0.33 | -0.81 | 0.06 | -0.15 | 1.00 | 8.4E-02 | 0.87 |
| SACM1L | 0.01 | -0.08 | -0.03 | 0.04 | 0.01 | 0.05 | 8.4E-02 | 0.07 |
| FKBP4 | 0.12 | 0.09 | -0.04 | -0.03 | -0.07 | -0.09 | 8.4E-02 | -0.12 |
| BAZ2A | 0.05 | 0.01 | 0.04 | -0.08 | -0.03 | 0.01 | 8.4E-02 | -0.07 |
| PPP2R5E | -0.10 | -0.05 | -0.05 | 0.16 | -0.03 | 0.06 | 8.5E-02 | 0.13 |
| RABGAP1 | -0.37 | -0.58 | 0.06 | 0.38 | -0.03 | 0.31 | 8.5E-02 | 0.52 |
| CRLF3 | -0.26 | -0.03 | -0.07 | 0.17 | -0.03 | 0.17 | 8.5E-02 | 0.22 |
| TRAPPC13 | -0.04 | -0.27 | -0.06 | 0.02 | 0.25 | 0.06 | 8.5E-02 | 0.23 |
| HMGN5 | 0.13 | 0.06 | 0.08 | -0.25 | -0.07 | 0.02 | 8.5E-02 | -0.19 |
| UTP6 | 0.07 | 0.06 | 0.00 | -0.02 | -0.01 | -0.11 | 8.5E-02 | -0.09 |
| SLC12A6 | -0.04 | 0.21 | 0.11 | -0.02 | -0.12 | -0.19 | 8.5E-02 | -0.20 |
| TMEM161A | -0.06 | -0.07 | -0.04 | 0.08 | -0.04 | 0.12 | 8.5E-02 | 0.11 |
| ABCA1 | 0.31 | 0.30 | 0.11 | -0.26 | 0.05 | -0.78 | 8.5E-02 | -0.57 |
| DDRGK1 | -0.07 | 0.12 | 0.28 | -0.07 | -0.15 | -0.16 | 8.6E-02 | -0.24 |
| CAPN5 | -0.09 | -0.12 | -0.18 | 0.28 | 0.14 | -0.08 | 8.6E-02 | 0.24 |
| SLC25A1 | 0.03 | 0.03 | 0.02 | -0.01 | 0.00 | -0.08 | 8.6E-02 | -0.06 |
| GMIP | -0.54 | -0.20 | 0.06 | 0.11 | 0.16 | 0.26 | 8.6E-02 | 0.41 |
| MED7 | 0.04 | 0.00 | 0.54 | -0.07 | -0.39 | -0.32 | 8.6E-02 | -0.46 |
| PARD3 | -0.21 | -0.09 | 0.00 | 0.04 | 0.02 | 0.20 | 8.6E-02 | 0.19 |
| DPP7 | 0.02 | 0.02 | 0.22 | -0.09 | -0.02 | -0.18 | 8.6E-02 | -0.18 |
| DIXDC1 | 0.07 | -0.04 | 0.24 | -0.06 | -0.16 | -0.09 | 8.6E-02 | -0.20 |
| FUBP3 | 0.01 | 0.02 | 0.17 | -0.13 | -0.01 | -0.08 | 8.6E-02 | -0.14 |
| SAMD1 | -0.07 | -0.12 | -0.04 | -0.05 | 0.15 | 0.10 | 8.6E-02 | 0.14 |
| POC1A | 0.07 | -0.04 | 0.26 | -0.11 | -0.15 | -0.07 | 8.7E-02 | -0.20 |
| WDR75 | -0.02 | -0.06 | -0.04 | -0.03 | 0.06 | 0.07 | 8.7E-02 | 0.08 |
| C2CD2L | -0.06 | -0.17 | 0.02 | 0.08 | 0.09 | 0.02 | 8.7E-02 | 0.13 |
| G3BP2 | -0.01 | 0.14 | 0.16 | 0.02 | -0.12 | -0.23 | 8.7E-02 | -0.21 |
| PWWP3A | -0.02 | 0.06 | 0.09 | -0.09 | -0.01 | -0.05 | 8.7E-02 | -0.09 |
| FAM8A1 | -0.02 | 0.12 | 0.21 | -0.09 | -0.01 | -0.25 | 8.7E-02 | -0.22 |
| SEC11C | 0.03 | 0.12 | 0.16 | -0.06 | 0.00 | -0.30 | 8.7E-02 | -0.23 |
| ZNF696 | 0.21 | -0.03 | 0.56 | -0.72 | -0.02 | -0.32 | 8.7E-02 | -0.60 |
| SIX1 | 0.03 | 0.14 | 0.05 | -0.04 | -0.21 | 0.00 | 8.7E-02 | -0.16 |
| TEX264 | 0.03 | -0.08 | -0.26 | 0.11 | 0.06 | 0.11 | 8.7E-02 | 0.20 |

| | | | | | | | | |
|---|---|---|---|---|---|---|---|---|
| HNRNPL | -0.02 | -0.11 | -0.09 | 0.05 | -0.02 | 0.17 | 8.7E-02 | 0.14 |
| FN1 | -0.21 | -0.09 | -0.29 | -0.12 | 0.22 | 0.37 | 8.7E-02 | 0.35 |
| TTC38 | -0.16 | -0.38 | -0.08 | 0.38 | -0.09 | 0.19 | 8.7E-02 | 0.37 |
| RAP1GDS1 | -0.04 | -0.11 | -0.03 | 0.13 | 0.06 | -0.03 | 8.7E-02 | 0.12 |
| CPSF3 | 0.02 | 0.04 | 0.11 | 0.03 | -0.08 | -0.13 | 8.8E-02 | -0.12 |
| RNF31 | -0.34 | -0.30 | -0.07 | 0.18 | 0.47 | -0.10 | 8.8E-02 | 0.42 |
| PIN4 | 0.33 | 0.10 | -0.04 | -0.02 | -0.21 | -0.24 | 8.8E-02 | -0.28 |
| CGGBP1 | -0.01 | 0.10 | 0.02 | -0.02 | -0.03 | -0.06 | 8.8E-02 | -0.08 |
| SMPD4 | -0.05 | -0.04 | -0.04 | 0.01 | 0.00 | 0.12 | 8.8E-02 | 0.09 |
| SLC38A2 | 0.03 | 0.08 | 0.21 | 0.04 | -0.15 | -0.27 | 8.8E-02 | -0.23 |
| CCT2 | 0.03 | 0.07 | 0.11 | -0.17 | 0.04 | -0.09 | 8.8E-02 | -0.14 |
| BRD4 | 0.05 | -0.01 | 0.12 | 0.01 | -0.08 | -0.10 | 8.8E-02 | -0.11 |
| PDE4DIP | -0.17 | -0.13 | 0.03 | 0.17 | 0.05 | 0.03 | 8.8E-02 | 0.17 |
| SCAMP4 | -0.07 | 0.12 | 0.36 | -0.27 | -0.15 | -0.08 | 8.8E-02 | -0.31 |
| RUSF1 | 0.10 | -0.03 | 0.21 | -0.21 | -0.01 | -0.11 | 8.8E-02 | -0.21 |
| RPL18 | 0.02 | 0.01 | 0.12 | -0.07 | 0.01 | -0.10 | 8.8E-02 | -0.11 |
| SBF1 | -0.06 | 0.12 | 0.23 | -0.03 | -0.13 | -0.16 | 8.8E-02 | -0.21 |
| MTFP1 | 0.03 | 0.01 | 0.11 | 0.00 | -0.05 | -0.13 | 8.8E-02 | -0.11 |
| PFKP | -0.03 | -0.09 | -0.34 | 0.30 | 0.06 | 0.02 | 8.9E-02 | 0.28 |
| TAF9 | -0.02 | -0.07 | 0.00 | 0.07 | 0.02 | 0.00 | 8.9E-02 | 0.06 |
| SYTL1 | -0.14 | -0.08 | 0.01 | 0.06 | -0.01 | 0.14 | 8.9E-02 | 0.14 |
| NRM | -0.09 | -0.25 | -0.07 | -0.01 | 0.06 | 0.31 | 8.9E-02 | 0.25 |
| ZBTB11 | 0.01 | 0.11 | 0.02 | -0.10 | -0.05 | 0.00 | 8.9E-02 | -0.10 |
| ZBTB7B | -0.12 | -0.14 | 0.05 | 0.07 | 0.03 | 0.09 | 8.9E-02 | 0.14 |
| TANC1 | -0.03 | -0.10 | -0.13 | 0.18 | -0.05 | 0.09 | 8.9E-02 | 0.16 |
| ECH1 | -0.01 | 0.02 | 0.12 | -0.05 | -0.04 | -0.06 | 8.9E-02 | -0.09 |
| MRPS28 | 0.10 | 0.10 | 0.08 | -0.28 | 0.00 | -0.03 | 8.9E-02 | -0.20 |
| PPP1R8 | 0.17 | -0.02 | 0.06 | -0.05 | -0.04 | -0.13 | 8.9E-02 | -0.14 |
| TRIM39 | -0.12 | -0.17 | -0.23 | -0.11 | 0.18 | 0.36 | 8.9E-02 | 0.31 |
| GTF2A1 | 0.07 | 0.01 | 0.17 | -0.04 | -0.02 | -0.22 | 8.9E-02 | -0.17 |
| PRPF4B | 0.12 | 0.09 | 0.01 | -0.07 | 0.02 | -0.18 | 9.0E-02 | -0.15 |
| LNPEP | -0.02 | 0.06 | 0.11 | -0.09 | -0.07 | 0.00 | 9.0E-02 | -0.10 |
| PPP1R10 | 0.06 | 0.07 | 0.08 | 0.06 | -0.15 | -0.14 | 9.0E-02 | -0.15 |
| RRM1 | 0.13 | 0.21 | -0.04 | -0.19 | 0.01 | -0.18 | 9.0E-02 | -0.22 |
| TAOK3 | -0.08 | -0.21 | -0.04 | -0.02 | 0.07 | 0.25 | 9.0E-02 | 0.21 |
| SMIM20 | 0.00 | 0.05 | 0.18 | -0.02 | -0.18 | -0.05 | 9.0E-02 | -0.16 |
| NME7 | -0.23 | -0.13 | 0.00 | -0.04 | 0.14 | 0.21 | 9.0E-02 | 0.22 |
| SEC23B | 0.07 | 0.13 | 0.06 | -0.02 | -0.02 | -0.26 | 9.0E-02 | -0.19 |
| UFD1 | 0.08 | 0.09 | 0.00 | -0.11 | 0.03 | -0.11 | 9.0E-02 | -0.12 |
| SNX27 | -0.04 | -0.05 | -0.02 | 0.06 | 0.08 | -0.03 | 9.0E-02 | 0.08 |
| MBNL1 | -0.11 | -0.05 | -0.08 | -0.03 | 0.06 | 0.20 | 9.0E-02 | 0.16 |
| FLNA | 0.08 | 0.10 | 0.02 | -0.01 | -0.02 | -0.20 | 9.0E-02 | -0.15 |
| DDX46 | 0.14 | 0.11 | 0.07 | 0.00 | -0.05 | -0.33 | 9.1E-02 | -0.23 |
| INTS1 | 0.01 | -0.12 | -0.05 | 0.02 | 0.02 | 0.11 | 9.1E-02 | 0.11 |
| DHTKD1 | -0.02 | -0.01 | -0.23 | 0.07 | 0.07 | 0.08 | 9.1E-02 | 0.16 |
| RAB18 | 0.01 | 0.01 | 0.06 | -0.06 | 0.01 | -0.03 | 9.1E-02 | -0.05 |
| CDK11B | 0.01 | -0.10 | -0.10 | 0.04 | 0.14 | 0.00 | 9.1E-02 | 0.12 |
| USP54 | -0.27 | -0.05 | -0.04 | 0.20 | 0.15 | -0.03 | 9.1E-02 | 0.23 |

| | | | | | | | | |
|---|---|---|---|---|---|---|---|---|
| SHISA5 | 0.08 | -0.04 | 0.35 | -0.27 | -0.11 | -0.09 | 9.1E-02 | -0.29 |
| NFRKB | -0.02 | -0.06 | -0.04 | 0.00 | 0.01 | 0.11 | 9.1E-02 | 0.08 |
| NUDT5 | 0.17 | 0.17 | -0.03 | 0.03 | -0.17 | -0.20 | 9.1E-02 | -0.21 |
| BIRC5 | -0.02 | -0.11 | -0.12 | 0.00 | 0.02 | 0.20 | 9.1E-02 | 0.16 |
| KIF20A | 0.05 | 0.06 | 0.10 | -0.19 | -0.07 | 0.03 | 9.1E-02 | -0.15 |
| TSHR | 0.23 | 0.27 | 0.10 | -0.06 | -0.05 | -0.66 | 9.2E-02 | -0.46 |
| AP2M1 | -0.16 | -0.14 | -0.01 | 0.06 | -0.02 | 0.24 | 9.2E-02 | 0.20 |
| TRAM1 | 0.00 | 0.11 | 0.03 | -0.08 | 0.00 | -0.07 | 9.2E-02 | -0.09 |
| PEX2 | 0.04 | 0.00 | 0.20 | -0.16 | -0.07 | -0.02 | 9.2E-02 | -0.16 |
| ZBED4 | -0.48 | -0.19 | 0.04 | 0.05 | 0.13 | 0.32 | 9.2E-02 | 0.38 |
| UNG | -0.47 | -0.33 | 0.10 | 0.17 | 0.32 | 0.05 | 9.2E-02 | 0.41 |
| SLC25A12 | -0.04 | 0.00 | -0.04 | 0.05 | -0.02 | 0.04 | 9.2E-02 | 0.05 |
| CUX1 | 0.01 | 0.04 | 0.07 | -0.07 | -0.08 | 0.02 | 9.3E-02 | -0.08 |
| ZWINT | -0.22 | -0.24 | 0.00 | 0.02 | 0.03 | 0.33 | 9.3E-02 | 0.28 |
| PFKL | -0.14 | -0.15 | 0.03 | 0.00 | 0.07 | 0.16 | 9.3E-02 | 0.16 |
| THRAP3 | 0.06 | 0.08 | 0.01 | -0.02 | 0.00 | -0.13 | 9.3E-02 | -0.10 |
| UBTF | 0.04 | 0.05 | 0.04 | -0.11 | 0.03 | -0.06 | 9.3E-02 | -0.09 |
| HMGN4 | -0.05 | -0.11 | -0.01 | 0.08 | -0.03 | 0.11 | 9.3E-02 | 0.11 |
| NUP188 | 0.15 | 0.16 | 0.07 | 0.01 | -0.07 | -0.39 | 9.3E-02 | -0.27 |
| BCR | -0.08 | -0.18 | 0.02 | 0.00 | 0.16 | 0.07 | 9.3E-02 | 0.16 |
| NUDT16L1 | 0.08 | -0.03 | 0.13 | -0.05 | -0.02 | -0.12 | 9.3E-02 | -0.12 |
| IRF2BPL | -0.33 | 0.04 | -0.23 | 0.32 | 0.11 | 0.00 | 9.3E-02 | 0.32 |
| ABHD11 | -0.01 | -0.08 | -0.13 | -0.04 | 0.10 | 0.14 | 9.3E-02 | 0.14 |
| CBX6 | 0.07 | 0.25 | 0.00 | -0.12 | -0.25 | 0.00 | 9.3E-02 | -0.23 |
| ESRP2 | -0.15 | -0.09 | 0.03 | 0.10 | 0.01 | 0.08 | 9.3E-02 | 0.13 |
| TMEM30A | 0.07 | 0.09 | 0.03 | -0.01 | -0.01 | -0.18 | 9.4E-02 | -0.13 |
| SNRNP40 | 0.07 | -0.01 | 0.02 | -0.04 | -0.02 | -0.02 | 9.4E-02 | -0.05 |
| TSPO | -0.10 | -0.08 | -0.22 | 0.07 | -0.04 | 0.31 | 9.4E-02 | 0.25 |
| ARHGEF1 | -0.08 | -0.07 | 0.00 | 0.01 | 0.13 | 0.01 | 9.4E-02 | 0.10 |
| VPS37B | -0.16 | -0.03 | -0.02 | 0.03 | 0.02 | 0.14 | 9.4E-02 | 0.13 |
| GTPBP6 | -0.05 | -0.13 | 0.01 | 0.10 | -0.01 | 0.08 | 9.4E-02 | 0.11 |
| NKIRAS1 | -0.07 | -0.26 | -0.08 | -0.03 | 0.07 | 0.31 | 9.4E-02 | 0.26 |
| PTPN2 | 0.01 | 0.00 | 0.19 | -0.06 | -0.12 | -0.05 | 9.4E-02 | -0.14 |
| AGPAT2 | -0.15 | 0.00 | -0.06 | -0.03 | 0.10 | 0.13 | 9.4E-02 | 0.14 |
| RANGAP1 | -0.08 | -0.11 | -0.13 | 0.23 | -0.08 | 0.13 | 9.4E-02 | 0.20 |
| PUM3 | 0.01 | -0.04 | -0.05 | 0.03 | 0.00 | 0.04 | 9.4E-02 | 0.05 |
| SNF8 | -0.05 | 0.02 | -0.08 | 0.05 | 0.03 | 0.01 | 9.5E-02 | 0.07 |
| MED14 | 0.06 | 0.12 | 0.03 | 0.05 | -0.14 | -0.15 | 9.5E-02 | -0.15 |
| CNN2 | 0.01 | 0.13 | 0.08 | -0.10 | 0.03 | -0.17 | 9.5E-02 | -0.15 |
| TTC17 | 0.04 | 0.14 | 0.13 | 0.06 | -0.14 | -0.27 | 9.5E-02 | -0.22 |
| BANF1 | -0.12 | -0.24 | -0.15 | 0.13 | -0.09 | 0.38 | 9.5E-02 | 0.31 |
| CCDC12 | 0.09 | 0.04 | -0.02 | -0.08 | 0.00 | -0.05 | 9.5E-02 | -0.08 |
| C2CD5 | -0.16 | -0.01 | -0.01 | 0.06 | 0.06 | 0.05 | 9.5E-02 | 0.11 |
| TMEM50B | -0.05 | -0.07 | -0.33 | 0.24 | -0.05 | 0.19 | 9.5E-02 | 0.28 |
| TAF6 | 0.02 | -0.05 | -0.15 | 0.08 | 0.03 | 0.05 | 9.5E-02 | 0.12 |
| BICRA | 0.16 | 0.04 | 0.05 | -0.09 | 0.03 | -0.21 | 9.6E-02 | -0.17 |
| HADHB | 0.01 | -0.06 | -0.08 | 0.09 | 0.01 | 0.02 | 9.6E-02 | 0.08 |
| SRP14 | 0.06 | 0.10 | -0.02 | -0.07 | -0.01 | -0.07 | 9.6E-02 | -0.09 |

| | | | | | | | | |
|---|---|---|---|---|---|---|---|---|
| **WWP2** | 0.02 | 0.03 | 0.10 | 0.02 | -0.07 | -0.11 | 9.6E-02 | -0.10 |
| **ANKRD28** | -0.18 | -0.35 | -0.14 | 0.38 | 0.31 | -0.17 | 9.6E-02 | 0.40 |
| **PDLIM5** | 0.18 | 0.29 | -0.11 | -0.08 | -0.16 | -0.19 | 9.6E-02 | -0.27 |
| **CDK7** | 0.06 | 0.10 | 0.07 | 0.03 | -0.07 | -0.22 | 9.6E-02 | -0.16 |
| **RAB11FIP1** | -0.10 | -0.03 | -0.01 | -0.01 | 0.05 | 0.09 | 9.6E-02 | 0.09 |
| **YWHAE** | 0.02 | 0.03 | 0.02 | -0.02 | 0.01 | -0.06 | 9.6E-02 | -0.04 |
| **PTGES2** | 0.02 | -0.06 | -0.22 | 0.11 | 0.09 | 0.03 | 9.6E-02 | 0.16 |
| **SLC39A7** | -0.05 | -0.06 | -0.03 | 0.07 | -0.03 | 0.08 | 9.6E-02 | 0.08 |
| **MT-ND6** | -0.16 | -0.24 | -0.16 | 0.35 | -0.15 | 0.26 | 9.6E-02 | 0.34 |
| **SIPA1L2** | -0.20 | -0.12 | 0.04 | 0.03 | 0.15 | 0.07 | 9.6E-02 | 0.18 |
| **MBOAT7** | 0.03 | 0.04 | 0.02 | 0.02 | -0.04 | -0.08 | 9.6E-02 | -0.07 |
| **NARS1** | 0.05 | 0.10 | -0.02 | -0.04 | -0.01 | -0.09 | 9.6E-02 | -0.09 |
| **FUT3** | -0.19 | -0.04 | -0.11 | 0.17 | -0.08 | 0.20 | 9.6E-02 | 0.21 |
| **UNC13D** | 0.01 | 0.05 | 0.07 | -0.09 | -0.09 | 0.03 | 9.7E-02 | -0.10 |
| **APEH** | 0.00 | -0.05 | -0.10 | 0.00 | 0.03 | 0.11 | 9.7E-02 | 0.09 |
| **LAMTOR4** | -0.14 | -0.04 | -0.01 | 0.00 | 0.13 | 0.04 | 9.7E-02 | 0.12 |
| **CTNNA1** | 0.02 | 0.08 | 0.04 | -0.08 | -0.09 | 0.03 | 9.7E-02 | -0.09 |
| **LONP1** | 0.02 | -0.07 | -0.15 | 0.03 | 0.13 | 0.02 | 9.7E-02 | 0.13 |
| **FAM120C** | -0.07 | -0.18 | -0.02 | -0.04 | 0.12 | 0.16 | 9.7E-02 | 0.17 |
| **RPRD1B** | 0.04 | 0.04 | 0.09 | 0.02 | -0.15 | -0.05 | 9.7E-02 | -0.12 |
| **GSE1** | -0.16 | -0.11 | -0.01 | -0.01 | 0.21 | 0.04 | 9.7E-02 | 0.17 |
| **SCFD2** | 0.10 | 0.07 | 0.04 | 0.00 | -0.02 | -0.21 | 9.7E-02 | -0.15 |
| **MSL1** | -0.12 | -0.07 | -0.05 | 0.03 | -0.01 | 0.21 | 9.8E-02 | 0.15 |
| **FAM174A** | -0.07 | 0.28 | 0.26 | 0.04 | -0.29 | -0.33 | 9.8E-02 | -0.35 |
| **CNOT1** | -0.10 | -0.14 | 0.03 | 0.06 | 0.01 | 0.12 | 9.8E-02 | 0.13 |
| **DNMT3A** | 0.09 | 0.02 | 0.05 | -0.11 | -0.10 | 0.04 | 9.8E-02 | -0.11 |
| **GK** | 0.03 | -0.11 | -0.05 | 0.04 | 0.05 | 0.03 | 9.8E-02 | 0.08 |
| **ITGA2** | -0.08 | 0.13 | 0.49 | -0.42 | -0.19 | -0.10 | 9.8E-02 | -0.42 |
| **THOC6** | 0.09 | 0.07 | 0.01 | 0.02 | -0.05 | -0.15 | 9.8E-02 | -0.12 |
| **EMC1** | 0.00 | 0.01 | 0.13 | -0.05 | -0.03 | -0.06 | 9.8E-02 | -0.09 |
| **GFM1** | 0.02 | -0.08 | -0.24 | 0.17 | 0.06 | 0.03 | 9.9E-02 | 0.19 |
| **MEF2D** | 0.15 | -0.07 | 0.19 | -0.17 | -0.04 | -0.10 | 9.9E-02 | -0.19 |
| **COQ8B** | 0.24 | 0.42 | -0.07 | -0.54 | -0.27 | 0.01 | 9.9E-02 | -0.46 |
| **OSBPL9** | -0.09 | -0.08 | -0.02 | 0.06 | -0.03 | 0.15 | 9.9E-02 | 0.12 |
| **RSRC1** | -0.06 | -0.05 | 0.02 | 0.01 | 0.02 | 0.06 | 9.9E-02 | 0.06 |
| **BAHD1** | 0.08 | 0.03 | 0.04 | -0.02 | -0.15 | 0.00 | 9.9E-02 | -0.11 |
| **DDX6** | 0.03 | 0.06 | 0.00 | -0.03 | 0.01 | -0.07 | 9.9E-02 | -0.06 |
| **NOM1** | 0.06 | 0.09 | -0.01 | 0.00 | -0.04 | -0.10 | 9.9E-02 | -0.09 |
| **UHRF2** | -0.09 | -0.05 | 0.01 | 0.01 | 0.02 | 0.09 | 9.9E-02 | 0.08 |
| **ST13** | 0.00 | -0.04 | -0.02 | 0.03 | -0.01 | 0.04 | 9.9E-02 | 0.04 |
| **SQSTM1** | -0.14 | -0.09 | 0.00 | 0.08 | 0.15 | -0.03 | 9.9E-02 | 0.14 |
| **LCMT1** | -0.09 | -0.13 | 0.02 | 0.12 | -0.02 | 0.09 | 9.9E-02 | 0.13 |
| **RRP8** | -0.05 | -0.08 | -0.07 | 0.10 | -0.06 | 0.14 | 1.0E-01 | 0.13 |
| **PHB2** | -0.01 | -0.01 | -0.03 | 0.05 | 0.00 | 0.00 | 1.0E-01 | 0.04 |
| **BUD13** | 0.00 | -0.09 | -0.06 | -0.01 | 0.03 | 0.11 | 1.0E-01 | 0.10 |
| **MVP** | 0.09 | 0.13 | 0.36 | 0.13 | -0.48 | -0.42 | 1.0E-01 | -0.45 |
| **PAK1IP1** | 0.08 | 0.07 | 0.06 | 0.03 | -0.06 | -0.21 | 1.0E-01 | -0.15 |
| **RIBC2** | 0.03 | 0.11 | 0.11 | -0.21 | -0.12 | 0.05 | 1.0E-01 | -0.17 |

| | | | | | | | | |
|---|---|---|---|---|---|---|---|---|
| **PHF3** | 0.03 | 0.05 | 0.04 | 0.03 | -0.11 | -0.05 | 1.0E-01 | -0.09 |
| **AFF1** | 0.18 | 0.03 | 0.07 | -0.15 | -0.23 | 0.05 | 1.0E-01 | -0.20 |
| **RRP1B** | 0.01 | 0.01 | 0.03 | 0.01 | -0.02 | -0.04 | 1.0E-01 | -0.03 |
| **MRPL54** | 0.01 | 0.08 | 0.20 | 0.02 | -0.26 | -0.10 | 1.0E-01 | -0.21 |
| **NSA2** | 0.06 | 0.01 | 0.12 | 0.04 | -0.13 | -0.12 | 1.0E-01 | -0.13 |
| **EDC3** | -0.30 | -0.10 | -0.01 | 0.23 | -0.04 | 0.15 | 1.0E-01 | 0.25 |
| **DHX33** | -0.06 | -0.12 | 0.02 | 0.11 | 0.01 | 0.03 | 1.0E-01 | 0.10 |
| **CD59** | -0.13 | -0.17 | -0.01 | 0.07 | -0.04 | 0.24 | 1.0E-01 | 0.20 |
| **ACADM** | 0.04 | -0.09 | -0.29 | 0.11 | 0.12 | 0.06 | 1.0E-01 | 0.21 |
| **PAICS** | 0.20 | 0.21 | -0.10 | -0.14 | -0.08 | -0.15 | 1.0E-01 | -0.22 |
| **HMG20A** | -0.03 | -0.10 | -0.02 | -0.02 | 0.05 | 0.12 | 1.0E-01 | 0.10 |
| **PIP4P1** | -0.08 | -0.07 | 0.00 | -0.02 | 0.04 | 0.12 | 1.0E-01 | 0.10 |
| **MED13** | 0.32 | 0.09 | 0.25 | 0.14 | -0.66 | -0.39 | 1.0E-01 | -0.52 |
| **NDUFB4** | -0.13 | -0.09 | -0.02 | 0.16 | -0.05 | 0.10 | 1.0E-01 | 0.15 |
| **BECN1** | -0.42 | 0.06 | -0.15 | 0.20 | 0.16 | 0.06 | 1.0E-01 | 0.31 |
| **SUMF2** | -0.05 | -0.17 | -0.03 | 0.02 | 0.20 | 0.01 | 1.0E-01 | 0.16 |
| **POLR1C** | -0.06 | -0.16 | 0.00 | -0.03 | 0.09 | 0.13 | 1.0E-01 | 0.14 |
| **TST** | -0.04 | -0.09 | -0.11 | 0.21 | 0.00 | 0.00 | 1.0E-01 | 0.15 |
| **ZNF397** | -0.22 | 0.39 | 0.41 | -0.39 | -0.25 | -0.16 | 1.0E-01 | -0.46 |
| **TACC1** | -0.07 | 0.12 | 0.22 | -0.17 | -0.03 | -0.12 | 1.0E-01 | -0.20 |
| **TXNRD1** | 0.03 | 0.00 | 0.36 | -0.29 | -0.11 | -0.07 | 1.0E-01 | -0.28 |
| **TOX4** | 0.06 | 0.09 | 0.01 | 0.03 | -0.12 | -0.08 | 1.0E-01 | -0.11 |
| **RPLP0** | -0.07 | -0.08 | -0.02 | 0.14 | 0.03 | -0.02 | 1.0E-01 | 0.11 |
| **WDR26** | -0.01 | -0.14 | -0.07 | 0.11 | 0.15 | -0.04 | 1.0E-01 | 0.15 |
| **CD93** | 0.04 | 0.21 | 0.21 | -0.13 | 0.04 | -0.47 | 1.0E-01 | -0.34 |
| **CRAT** | 0.04 | -0.11 | -0.09 | 0.01 | 0.07 | 0.08 | 1.0E-01 | 0.11 |
| **FCHO2** | 0.04 | 0.09 | 0.00 | -0.10 | -0.02 | -0.01 | 1.0E-01 | -0.08 |
| **CYB561** | -0.06 | 0.08 | 0.27 | -0.16 | -0.12 | -0.06 | 1.0E-01 | -0.21 |
| **TM9SF2** | 0.07 | 0.12 | 0.11 | 0.09 | -0.22 | -0.21 | 1.0E-01 | -0.22 |
| **GNG10** | 0.08 | 0.04 | 0.14 | 0.00 | -0.03 | -0.27 | 1.0E-01 | -0.19 |
| **KRI1** | 0.06 | 0.10 | 0.05 | 0.02 | -0.04 | -0.21 | 1.0E-01 | -0.15 |
| **MIEF1** | -0.01 | -0.25 | -0.25 | 0.11 | -0.06 | 0.36 | 1.0E-01 | 0.31 |
| **PDCD6** | -0.03 | -0.10 | 0.01 | 0.04 | 0.01 | 0.07 | 1.0E-01 | 0.08 |
| **PPP2R5A** | -0.13 | -0.08 | 0.03 | 0.09 | 0.00 | 0.08 | 1.0E-01 | 0.12 |
| **LLGL1** | 0.01 | 0.03 | 0.07 | -0.06 | 0.02 | -0.08 | 1.0E-01 | -0.08 |
| **ATF7IP** | 0.05 | -0.20 | -0.21 | 0.00 | 0.22 | 0.10 | 1.0E-01 | 0.22 |
| **SMU1** | 0.02 | 0.01 | 0.04 | -0.05 | 0.01 | -0.02 | 1.0E-01 | -0.04 |
| **MRPL19** | 0.19 | 0.15 | -0.03 | 0.01 | -0.10 | -0.28 | 1.0E-01 | -0.23 |
| **STX7** | -0.08 | 0.10 | 0.33 | -0.14 | -0.19 | -0.08 | 1.0E-01 | -0.26 |
| **RPS7** | -0.01 | -0.02 | -0.05 | 0.01 | 0.06 | 0.00 | 1.0E-01 | 0.05 |
| **SBNO1** | 0.30 | 0.07 | 0.04 | -0.38 | 0.00 | -0.10 | 1.0E-01 | -0.30 |
| **PCNA** | 0.13 | 0.01 | 0.12 | -0.06 | 0.02 | -0.25 | 1.0E-01 | -0.18 |
| **CLK4** | 0.02 | 0.17 | 0.11 | -0.14 | 0.05 | -0.25 | 1.0E-01 | -0.21 |
| **P3H3** | 0.13 | -0.05 | 0.17 | -0.04 | -0.06 | -0.18 | 1.0E-01 | -0.17 |
| **HNRNPK** | 0.02 | 0.00 | 0.10 | -0.10 | -0.03 | 0.00 | 1.0E-01 | -0.09 |
| **PLCB1** | 0.10 | -0.07 | 0.20 | -0.11 | -0.06 | -0.09 | 1.0E-01 | -0.17 |
| **NECTIN1** | -0.28 | -0.24 | 0.09 | 0.13 | 0.04 | 0.19 | 1.0E-01 | 0.26 |
| **ATXN2** | -0.19 | -0.06 | -0.11 | 0.31 | 0.04 | -0.03 | 1.0E-01 | 0.23 |

| | | | | | | | | |
|---|---|---|---|---|---|---|---|---|
| HSD17B4 | 0.00 | -0.03 | -0.07 | -0.01 | 0.06 | 0.03 | 1.0E-01 | 0.06 |
| KDM2B | 0.01 | 0.04 | 0.03 | -0.03 | -0.07 | 0.01 | 1.0E-01 | -0.06 |
| TM7SF2 | -0.01 | 0.06 | 0.03 | -0.03 | 0.01 | -0.07 | 1.0E-01 | -0.06 |
| DDX39B | 0.01 | 0.02 | 0.16 | -0.12 | 0.01 | -0.10 | 1.0E-01 | -0.13 |
| MRPL55 | -0.14 | -0.07 | -0.04 | 0.03 | -0.02 | 0.22 | 1.0E-01 | 0.17 |
| EIF2AK1 | 0.16 | 0.23 | 0.00 | -0.38 | -0.05 | -0.02 | 1.0E-01 | -0.28 |
| NELFA | -0.05 | 0.03 | -0.17 | 0.06 | 0.03 | 0.09 | 1.1E-01 | 0.12 |
| EP300 | -0.05 | -0.09 | -0.07 | -0.06 | 0.13 | 0.12 | 1.1E-01 | 0.13 |
| PEX10 | 0.08 | 0.01 | 0.16 | -0.24 | 0.01 | -0.05 | 1.1E-01 | -0.18 |
| NDUFV2 | 0.03 | 0.03 | 0.00 | 0.00 | -0.01 | -0.06 | 1.1E-01 | -0.04 |
| CHD9 | 0.04 | 0.05 | 0.10 | -0.19 | -0.03 | 0.01 | 1.1E-01 | -0.13 |
| XRCC4 | -0.15 | -0.14 | 0.04 | 0.01 | 0.05 | 0.17 | 1.1E-01 | 0.16 |
| JPT2 | 0.17 | 0.23 | -0.09 | -0.20 | -0.08 | -0.07 | 1.1E-01 | -0.22 |
| ARPC1B | -0.11 | -0.04 | 0.01 | 0.10 | 0.04 | 0.00 | 1.1E-01 | 0.09 |
| TMEM30B | -0.10 | -0.01 | 0.00 | 0.03 | 0.05 | 0.02 | 1.1E-01 | 0.07 |
| PCNX1 | 0.09 | -0.05 | 0.23 | -0.16 | -0.03 | -0.11 | 1.1E-01 | -0.19 |
| PRDX2 | 0.28 | 0.23 | -0.12 | -0.06 | -0.16 | -0.25 | 1.1E-01 | -0.28 |
| EXOC2 | -0.09 | -0.24 | 0.04 | 0.13 | 0.11 | 0.01 | 1.1E-01 | 0.18 |
| COBL | -0.26 | -0.09 | 0.04 | 0.03 | 0.11 | 0.14 | 1.1E-01 | 0.20 |
| LRRC47 | 0.00 | 0.07 | 0.03 | -0.03 | 0.00 | -0.09 | 1.1E-01 | -0.07 |
| WDR25 | 0.07 | 0.09 | 0.01 | -0.07 | -0.14 | 0.03 | 1.1E-01 | -0.11 |
| ZMYND8 | 0.10 | 0.03 | 0.00 | -0.10 | -0.02 | -0.01 | 1.1E-01 | -0.09 |
| VTA1 | -0.07 | -0.04 | -0.12 | 0.19 | -0.04 | 0.05 | 1.1E-01 | 0.15 |
| WDR37 | -0.16 | -0.22 | 0.05 | 0.12 | 0.00 | 0.17 | 1.1E-01 | 0.20 |
| PEX26 | 0.07 | -0.22 | -0.53 | 0.21 | 0.30 | 0.01 | 1.1E-01 | 0.40 |
| CAGE1 | -0.26 | -0.07 | -0.27 | 0.04 | -0.05 | 0.47 | 1.1E-01 | 0.35 |
| ABCB9 | -0.02 | 0.04 | 0.14 | -0.11 | -0.03 | -0.03 | 1.1E-01 | -0.11 |
| RBM34 | 0.10 | 0.10 | -0.03 | -0.05 | 0.00 | -0.13 | 1.1E-01 | -0.12 |
| OCIAD1 | 0.04 | -0.01 | 0.06 | -0.02 | -0.06 | 0.00 | 1.1E-01 | -0.06 |
| YWHAQ | -0.01 | -0.08 | -0.08 | 0.13 | -0.02 | 0.05 | 1.1E-01 | 0.10 |
| GRAMD1A | 0.09 | 0.02 | 0.05 | 0.04 | -0.09 | -0.12 | 1.1E-01 | -0.11 |
| NAMPT | -0.14 | -0.02 | -0.02 | 0.07 | -0.01 | 0.11 | 1.1E-01 | 0.12 |
| SENP3 | 0.02 | -0.03 | -0.10 | 0.03 | 0.03 | 0.04 | 1.1E-01 | 0.07 |
| POLR2B | 0.10 | 0.00 | 0.01 | -0.04 | -0.06 | -0.01 | 1.1E-01 | -0.07 |
| GPS2 | -0.27 | -0.16 | 0.06 | 0.21 | 0.04 | 0.07 | 1.1E-01 | 0.23 |
| TMX3 | 0.14 | 0.10 | -0.03 | 0.02 | -0.13 | -0.12 | 1.1E-01 | -0.14 |
| PLCB4 | -0.13 | -0.06 | 0.03 | 0.06 | 0.06 | 0.03 | 1.1E-01 | 0.10 |
| ATP5IF1 | 0.31 | 0.15 | -0.09 | -0.28 | -0.05 | -0.12 | 1.1E-01 | -0.28 |
| FXR2 | -0.11 | -0.04 | -0.08 | 0.09 | 0.18 | -0.05 | 1.1E-01 | 0.15 |
| GGA3 | -0.18 | -0.19 | 0.05 | 0.02 | 0.05 | 0.21 | 1.1E-01 | 0.20 |
| ZNF148 | -0.05 | -0.13 | -0.01 | 0.13 | -0.03 | 0.08 | 1.1E-01 | 0.12 |
| FAM83B | 0.19 | -0.03 | 0.18 | -0.07 | 0.00 | -0.32 | 1.1E-01 | -0.24 |
| SETDB1 | 0.05 | -0.17 | -0.13 | 0.08 | 0.01 | 0.12 | 1.1E-01 | 0.15 |
| ST3GAL2 | -0.16 | -0.12 | -0.10 | -0.04 | 0.04 | 0.33 | 1.1E-01 | 0.24 |
| ALYREF | 0.00 | 0.01 | 0.14 | -0.08 | -0.07 | -0.01 | 1.1E-01 | -0.10 |
| AKAP1 | -0.02 | 0.07 | 0.07 | 0.00 | -0.04 | -0.09 | 1.1E-01 | -0.08 |
| DUS2 | -0.12 | -0.05 | -0.43 | 0.11 | 0.40 | -0.04 | 1.1E-01 | 0.36 |
| IMP4 | 0.00 | 0.02 | 0.14 | -0.10 | 0.00 | -0.07 | 1.1E-01 | -0.11 |

| | | | | | | | | |
|---|---|---|---|---|---|---|---|---|
| SCAF8 | -0.16 | -0.11 | -0.23 | -0.02 | 0.01 | 0.42 | 1.1E-01 | 0.30 |
| NAA25 | 0.37 | 0.02 | 0.25 | 0.15 | -0.61 | -0.45 | 1.1E-01 | -0.52 |
| ZNRF2 | 0.09 | 0.04 | 0.01 | -0.11 | 0.02 | -0.06 | 1.1E-01 | -0.09 |
| ILF2 | 0.04 | 0.05 | -0.02 | -0.03 | -0.04 | 0.00 | 1.1E-01 | -0.05 |
| SHOC2 | -0.13 | -0.08 | -0.05 | -0.05 | 0.07 | 0.21 | 1.1E-01 | 0.16 |
| DNAJB14 | 0.06 | 0.07 | 0.24 | -0.38 | -0.04 | -0.01 | 1.1E-01 | -0.27 |
| FLCN | -0.11 | -0.16 | -0.08 | 0.15 | -0.10 | 0.25 | 1.1E-01 | 0.22 |
| HSDL2 | 0.02 | -0.03 | -0.24 | 0.07 | 0.08 | 0.08 | 1.1E-01 | 0.16 |
| SMARCA2 | -0.02 | -0.10 | -0.13 | 0.01 | 0.00 | 0.21 | 1.1E-01 | 0.16 |
| PTPN23 | 0.00 | 0.03 | 0.11 | -0.04 | 0.00 | -0.11 | 1.1E-01 | -0.09 |
| LMBRD2 | -0.05 | 0.12 | 0.15 | -0.01 | -0.07 | -0.18 | 1.1E-01 | -0.16 |
| SEC62 | 0.05 | 0.14 | 0.01 | 0.04 | -0.14 | -0.13 | 1.1E-01 | -0.14 |
| ELFN1 | 0.01 | -0.08 | -0.02 | 0.04 | 0.01 | 0.03 | 1.1E-01 | 0.06 |
| TUFM | 0.04 | -0.09 | -0.17 | 0.11 | 0.09 | 0.00 | 1.1E-01 | 0.14 |
| SCAMP2 | -0.04 | 0.09 | 0.27 | -0.11 | -0.26 | -0.01 | 1.1E-01 | -0.24 |
| ZNF691 | 0.03 | 0.03 | 0.02 | -0.01 | 0.01 | -0.08 | 1.1E-01 | -0.06 |
| TBC1D16 | -0.17 | -0.32 | -0.37 | -0.02 | 0.66 | -0.04 | 1.1E-01 | 0.49 |
| ZNF367 | 0.01 | 0.09 | 0.29 | -0.25 | 0.07 | -0.30 | 1.1E-01 | -0.29 |
| PRCC | 0.07 | 0.03 | 0.01 | -0.11 | -0.01 | 0.00 | 1.1E-01 | -0.08 |
| IGFBP5 | 0.04 | 0.06 | 0.15 | -0.20 | 0.07 | -0.15 | 1.1E-01 | -0.18 |
| CLASP1 | -0.01 | 0.17 | 0.04 | 0.02 | -0.10 | -0.14 | 1.1E-01 | -0.14 |
| HERPUD2 | -0.02 | 0.05 | 0.19 | -0.05 | -0.03 | -0.16 | 1.1E-01 | -0.15 |
| NECAP2 | 0.02 | 0.13 | 0.03 | 0.03 | -0.10 | -0.13 | 1.1E-01 | -0.13 |
| AGPS | -0.05 | 0.02 | -0.14 | 0.00 | 0.07 | 0.08 | 1.1E-01 | 0.11 |
| LSM6 | -0.04 | -0.20 | -0.39 | 0.37 | -0.14 | 0.25 | 1.1E-01 | 0.37 |
| ACBD5 | -0.02 | 0.06 | 0.09 | -0.03 | 0.00 | -0.11 | 1.1E-01 | -0.10 |
| BRWD3 | -0.02 | 0.07 | 0.16 | -0.17 | -0.07 | 0.01 | 1.1E-01 | -0.15 |
| SLC19A1 | -0.17 | 0.11 | 0.75 | -0.48 | -0.15 | -0.44 | 1.1E-01 | -0.59 |
| PTPN11 | -0.01 | 0.13 | 0.21 | -0.31 | 0.03 | -0.10 | 1.1E-01 | -0.24 |
| REEP5 | 0.25 | 0.26 | -0.13 | -0.22 | -0.17 | -0.07 | 1.1E-01 | -0.28 |
| TACSTD2 | -0.04 | 0.05 | 0.23 | -0.10 | -0.13 | -0.04 | 1.1E-01 | -0.17 |
| VPS25 | -0.09 | 0.02 | -0.06 | 0.09 | 0.01 | 0.02 | 1.1E-01 | 0.08 |
| SDF2L1 | 0.04 | 0.08 | 0.01 | 0.03 | -0.05 | -0.12 | 1.1E-01 | -0.10 |
| RAB5A | -0.19 | -0.11 | -0.02 | 0.02 | 0.00 | 0.26 | 1.1E-01 | 0.20 |
| PRKD2 | 0.02 | -0.11 | -0.26 | 0.21 | 0.12 | -0.03 | 1.1E-01 | 0.22 |
| ZNF324 | 0.08 | -0.02 | 0.08 | -0.11 | 0.02 | -0.05 | 1.1E-01 | -0.10 |
| ADIPOR1 | -0.03 | 0.07 | 0.16 | -0.13 | -0.08 | 0.00 | 1.1E-01 | -0.14 |
| STARD3NL | -0.02 | 0.09 | 0.26 | 0.04 | -0.21 | -0.23 | 1.1E-01 | -0.24 |
| NKTR | 0.05 | 0.08 | 0.06 | -0.01 | 0.00 | -0.19 | 1.1E-01 | -0.13 |
| ATAD3B | 0.12 | 0.14 | 0.00 | 0.02 | -0.06 | -0.26 | 1.1E-01 | -0.19 |
| WWP1 | -0.02 | 0.06 | 0.23 | -0.23 | -0.06 | -0.02 | 1.1E-01 | -0.19 |
| IWS1 | 0.06 | 0.00 | 0.08 | -0.13 | -0.02 | 0.00 | 1.1E-01 | -0.09 |
| LARP4 | -0.09 | -0.04 | 0.01 | 0.03 | 0.00 | 0.08 | 1.2E-01 | 0.08 |
| XXYLT1 | 0.08 | -0.03 | 0.11 | -0.06 | -0.12 | 0.00 | 1.2E-01 | -0.11 |
| FAM172A | 0.02 | 0.10 | 0.10 | -0.21 | -0.07 | 0.04 | 1.2E-01 | -0.16 |
| TAF4 | -0.01 | -0.06 | -0.06 | 0.01 | 0.00 | 0.11 | 1.2E-01 | 0.08 |
| ELP1 | -0.40 | -0.35 | 0.05 | 0.25 | -0.10 | 0.38 | 1.2E-01 | 0.41 |
| ZNF668 | 0.05 | 0.10 | 0.19 | 0.09 | -0.30 | -0.19 | 1.2E-01 | -0.25 |

| | | | | | | | | |
|---|---|---|---|---|---|---|---|---|
| HDGFL3 | -0.18 | -0.11 | -0.17 | -0.05 | 0.03 | 0.40 | 1.2E-01 | 0.28 |
| LRPPRC | 0.09 | 0.08 | -0.04 | -0.01 | -0.05 | -0.07 | 1.2E-01 | -0.09 |
| UBQLN2 | -0.15 | -0.07 | 0.00 | -0.03 | 0.17 | 0.06 | 1.2E-01 | 0.14 |
| IGFBP2 | -0.42 | -0.43 | 0.15 | 0.17 | 0.04 | 0.33 | 1.2E-01 | 0.41 |
| MANEA | -0.08 | 0.24 | 0.35 | -0.38 | 0.07 | -0.37 | 1.2E-01 | -0.39 |
| DLG5 | -0.04 | -0.04 | 0.01 | 0.00 | 0.02 | 0.04 | 1.2E-01 | 0.04 |
| ARHGAP39 | -0.06 | -0.06 | -0.17 | -0.08 | 0.16 | 0.17 | 1.2E-01 | 0.18 |
| GATA3 | 0.09 | 0.03 | 0.00 | 0.01 | -0.10 | -0.04 | 1.2E-01 | -0.08 |
| OASL | -0.15 | -0.20 | 0.01 | 0.24 | 0.11 | -0.06 | 1.2E-01 | 0.21 |
| RPL7L1 | 0.07 | 0.07 | 0.06 | -0.07 | 0.05 | -0.20 | 1.2E-01 | -0.14 |
| CAPZA2 | 0.01 | 0.07 | 0.10 | -0.02 | 0.00 | -0.18 | 1.2E-01 | -0.13 |
| PMF1 | -0.15 | -0.18 | -0.03 | -0.07 | 0.09 | 0.29 | 1.2E-01 | 0.23 |
| VTN | -0.16 | -0.25 | -0.04 | 0.35 | 0.09 | -0.07 | 1.2E-01 | 0.27 |
| CAPZA1 | 0.00 | 0.04 | 0.11 | -0.01 | -0.02 | -0.14 | 1.2E-01 | -0.10 |
| HAT1 | -0.18 | -0.05 | -0.07 | -0.03 | 0.25 | 0.04 | 1.2E-01 | 0.18 |
| ZNFX1 | -0.01 | -0.02 | -0.14 | -0.02 | 0.07 | 0.11 | 1.2E-01 | 0.11 |
| MED1 | 0.03 | 0.12 | 0.03 | 0.04 | -0.09 | -0.14 | 1.2E-01 | -0.12 |
| PPL | -0.05 | -0.05 | -0.05 | 0.10 | 0.09 | -0.05 | 1.2E-01 | 0.10 |
| RSF1 | -0.03 | -0.05 | -0.03 | -0.02 | 0.02 | 0.10 | 1.2E-01 | 0.07 |
| HNRNPDL | 0.07 | -0.01 | 0.04 | -0.04 | -0.08 | 0.01 | 1.2E-01 | -0.07 |
| MRPS10 | 0.19 | 0.13 | -0.04 | -0.22 | 0.03 | -0.12 | 1.2E-01 | -0.20 |
| UBN2 | -0.10 | -0.04 | -0.01 | -0.02 | 0.03 | 0.12 | 1.2E-01 | 0.09 |
| ZNF512B | -0.03 | 0.02 | -0.08 | 0.00 | 0.05 | 0.04 | 1.2E-01 | 0.06 |
| CYC1 | 0.02 | 0.06 | 0.09 | -0.01 | 0.00 | -0.18 | 1.2E-01 | -0.12 |
| FARSA | -0.07 | -0.11 | 0.02 | 0.10 | 0.06 | -0.01 | 1.2E-01 | 0.10 |
| IGSF8 | -0.04 | 0.05 | 0.32 | -0.22 | -0.11 | -0.06 | 1.2E-01 | -0.24 |
| CCNYL1 | -0.24 | -0.09 | 0.05 | 0.11 | 0.04 | 0.10 | 1.2E-01 | 0.18 |
| NSMCE3 | 0.45 | 0.67 | -0.39 | -0.38 | -0.46 | -0.36 | 1.2E-01 | -0.64 |
| HIBADH | 0.04 | -0.05 | -0.17 | 0.07 | 0.07 | 0.04 | 1.2E-01 | 0.12 |
| LACTB | -0.01 | -0.01 | -0.08 | 0.07 | 0.03 | -0.01 | 1.2E-01 | 0.06 |
| MAD2L1 | -0.28 | -0.10 | 0.01 | 0.07 | -0.01 | 0.26 | 1.2E-01 | 0.23 |
| KCNK10 | 0.05 | -0.17 | -0.61 | 0.05 | 0.13 | 0.37 | 1.2E-01 | 0.42 |
| MCCC1 | 0.02 | -0.10 | -0.10 | 0.12 | 0.06 | -0.02 | 1.2E-01 | 0.11 |
| PIAS1 | -0.01 | -0.08 | -0.11 | 0.17 | -0.03 | 0.04 | 1.2E-01 | 0.13 |
| ARV1 | 0.12 | 0.22 | -0.05 | -0.01 | -0.24 | -0.09 | 1.2E-01 | -0.21 |
| POLR2M | -0.14 | -0.15 | 0.05 | 0.01 | 0.14 | 0.07 | 1.2E-01 | 0.15 |
| RPL7 | -0.04 | -0.07 | -0.01 | 0.11 | -0.01 | 0.02 | 1.2E-01 | 0.08 |
| TMEM168 | 0.08 | 0.07 | 0.10 | -0.15 | 0.08 | -0.21 | 1.2E-01 | -0.18 |
| H3C15 | 0.00 | -0.08 | -0.03 | -0.03 | 0.08 | 0.06 | 1.2E-01 | 0.07 |
| CSK | -0.06 | 0.03 | -0.28 | 0.08 | 0.03 | 0.17 | 1.2E-01 | 0.20 |
| PIGG | -0.06 | -0.07 | -0.07 | 0.18 | 0.04 | -0.04 | 1.2E-01 | 0.13 |
| RBM12B | 0.02 | 0.06 | 0.08 | -0.05 | 0.03 | -0.15 | 1.2E-01 | -0.11 |
| CLASRP | -0.12 | -0.08 | -0.23 | 0.29 | -0.12 | 0.18 | 1.2E-01 | 0.26 |
| MMAA | -0.03 | -0.04 | -0.19 | -0.05 | 0.18 | 0.11 | 1.2E-01 | 0.16 |
| INTS14 | 0.09 | 0.03 | 0.00 | -0.02 | 0.00 | -0.10 | 1.2E-01 | -0.08 |
| RFXANK | -0.15 | -0.07 | -0.06 | -0.06 | 0.08 | 0.23 | 1.2E-01 | 0.18 |
| RMDN3 | 0.02 | 0.01 | 0.02 | 0.01 | -0.02 | -0.03 | 1.2E-01 | -0.03 |
| EIF3E | -0.11 | -0.16 | 0.04 | 0.13 | 0.00 | 0.08 | 1.2E-01 | 0.14 |

| | | | | | | | | |
|---|---|---|---|---|---|---|---|---|
| TRMT112 | 0.00 | -0.04 | -0.25 | 0.14 | 0.14 | -0.02 | 1.2E-01 | 0.19 |
| CHST12 | 0.06 | 0.09 | 0.27 | 0.02 | -0.08 | -0.45 | 1.2E-01 | -0.31 |
| RSBN1L | 0.06 | 0.12 | -0.03 | -0.10 | 0.02 | -0.09 | 1.2E-01 | -0.11 |
| GOLGA8O | -3.28 | -2.87 | -2.09 | 1.49 | -2.71 | 1.36 | 1.2E-01 | 2.79 |
| NIPSNAP1 | 0.05 | -0.16 | -0.19 | 0.02 | 0.05 | 0.20 | 1.2E-01 | 0.19 |
| JMJD6 | 0.48 | 0.20 | -0.17 | -0.34 | -0.06 | -0.29 | 1.2E-01 | -0.40 |
| ODF2L | 0.03 | -0.02 | 0.12 | -0.08 | -0.04 | -0.02 | 1.2E-01 | -0.09 |
| EMC7 | -0.04 | 0.02 | -0.06 | 0.00 | 0.03 | 0.05 | 1.2E-01 | 0.05 |
| ZNF581 | -0.05 | -0.04 | -0.04 | 0.03 | -0.03 | 0.11 | 1.2E-01 | 0.08 |
| MIOS | 0.05 | 0.10 | 0.04 | -0.15 | 0.06 | -0.12 | 1.2E-01 | -0.14 |
| SPPL2A | -0.14 | 0.07 | 0.87 | -0.61 | -0.05 | -0.75 | 1.2E-01 | -0.74 |
| ITPKB | -0.27 | -0.14 | -0.07 | -0.03 | 0.02 | 0.39 | 1.2E-01 | 0.28 |
| TMX4 | 0.03 | 0.04 | 0.13 | -0.19 | -0.06 | 0.03 | 1.2E-01 | -0.14 |
| KDELR3 | -0.20 | -0.29 | 0.01 | 0.08 | -0.05 | 0.36 | 1.2E-01 | 0.29 |
| TECR | 0.00 | 0.02 | 0.16 | 0.00 | -0.12 | -0.08 | 1.2E-01 | -0.12 |
| MTM1 | 0.12 | 0.42 | -0.11 | -0.08 | -0.31 | -0.17 | 1.2E-01 | -0.33 |
| AR | 0.09 | -0.14 | -0.29 | 0.10 | 0.10 | 0.10 | 1.2E-01 | 0.22 |
| LRRC14 | -0.24 | -0.01 | -0.31 | -0.08 | 0.41 | 0.11 | 1.2E-01 | 0.33 |
| ZNF770 | -0.26 | -0.28 | 0.02 | -0.10 | 0.34 | 0.17 | 1.2E-01 | 0.31 |
| SSB | -0.02 | -0.03 | -0.06 | 0.04 | 0.09 | -0.03 | 1.2E-01 | 0.07 |
| KAT7 | 0.01 | 0.00 | 0.05 | -0.05 | -0.04 | 0.01 | 1.2E-01 | -0.05 |
| ERMP1 | 0.04 | 0.05 | 0.04 | 0.04 | -0.07 | -0.11 | 1.2E-01 | -0.09 |
| A2M | -0.09 | 0.06 | -0.42 | 0.11 | 0.17 | 0.11 | 1.2E-01 | 0.28 |
| AAAS | -0.08 | -0.11 | 0.02 | 0.06 | -0.02 | 0.12 | 1.2E-01 | 0.11 |
| DHRS4 | 0.02 | -0.08 | -0.05 | 0.01 | 0.01 | 0.09 | 1.2E-01 | 0.07 |
| SLC35A5 | 0.00 | 0.18 | 0.06 | -0.17 | -0.15 | 0.05 | 1.2E-01 | -0.17 |
| DLGAP4 | -0.05 | -0.14 | -0.05 | 0.08 | -0.06 | 0.19 | 1.2E-01 | 0.15 |
| TAP1 | -0.01 | 0.05 | 0.10 | -0.03 | 0.01 | -0.13 | 1.2E-01 | -0.10 |
| ANAPC2 | 0.05 | -0.22 | -0.10 | 0.02 | 0.07 | 0.16 | 1.2E-01 | 0.17 |
| UBTD2 | 0.13 | 0.04 | 0.15 | 0.09 | -0.31 | -0.16 | 1.2E-01 | -0.24 |
| ZNF609 | -0.06 | 0.02 | -0.11 | 0.04 | 0.10 | 0.00 | 1.2E-01 | 0.09 |
| SVIP | -0.22 | -0.06 | 0.03 | 0.12 | 0.05 | 0.05 | 1.2E-01 | 0.15 |
| ZNF316 | 0.01 | 0.05 | 0.14 | -0.20 | -0.02 | -0.01 | 1.2E-01 | -0.14 |
| NUP98 | 0.02 | -0.04 | -0.09 | 0.06 | 0.03 | 0.01 | 1.2E-01 | 0.07 |
| HMGB3 | 0.18 | 0.12 | -0.05 | -0.15 | 0.03 | -0.17 | 1.2E-01 | -0.18 |
| SLX9 | -0.05 | -0.09 | -0.02 | 0.15 | -0.02 | 0.02 | 1.2E-01 | 0.10 |
| UBAP2L | -0.12 | -0.03 | 0.01 | 0.07 | 0.08 | -0.01 | 1.2E-01 | 0.09 |
| BCAP31 | 0.07 | 0.03 | 0.01 | -0.10 | 0.00 | -0.01 | 1.2E-01 | -0.07 |
| NPM1 | 0.05 | 0.02 | 0.02 | 0.01 | -0.01 | -0.10 | 1.2E-01 | -0.07 |
| SCAMP3 | -0.05 | 0.09 | 0.37 | -0.34 | -0.16 | 0.00 | 1.2E-01 | -0.30 |
| CWC25 | 0.18 | 0.05 | -0.02 | -0.14 | 0.02 | -0.12 | 1.3E-01 | -0.15 |
| SFXN4 | -0.17 | -0.01 | 0.00 | 0.05 | 0.04 | 0.07 | 1.3E-01 | 0.11 |
| RNPS1 | 0.08 | 0.02 | 0.12 | 0.04 | -0.07 | -0.22 | 1.3E-01 | -0.15 |
| AKAP8 | 0.03 | 0.03 | 0.02 | 0.02 | -0.03 | -0.07 | 1.3E-01 | -0.05 |
| NUP88 | 0.10 | 0.08 | -0.01 | 0.00 | -0.02 | -0.15 | 1.3E-01 | -0.11 |
| CHCHD4 | 0.25 | 0.31 | -0.12 | -0.27 | 0.03 | -0.33 | 1.3E-01 | -0.34 |
| YPEL5 | -0.05 | -0.03 | -0.01 | -0.02 | 0.06 | 0.04 | 1.3E-01 | 0.06 |
| VCP | 0.02 | 0.05 | 0.05 | 0.01 | -0.01 | -0.12 | 1.3E-01 | -0.08 |

| | | | | | | | | |
|---|---|---|---|---|---|---|---|---|
| ZFHX3 | -0.05 | -0.08 | -0.02 | -0.02 | 0.02 | 0.15 | 1.3E-01 | 0.10 |
| TOM1L2 | -0.16 | -0.03 | -0.06 | -0.06 | 0.11 | 0.18 | 1.3E-01 | 0.16 |
| UQCRB | -0.01 | -0.01 | -0.11 | 0.07 | 0.07 | -0.01 | 1.3E-01 | 0.08 |
| VPS45 | -0.01 | 0.03 | 0.10 | -0.08 | 0.01 | -0.06 | 1.3E-01 | -0.08 |
| DBN1 | 0.04 | 0.20 | 0.16 | 0.04 | -0.09 | -0.44 | 1.3E-01 | -0.30 |
| CEP68 | -0.04 | -0.13 | -0.06 | -0.03 | 0.04 | 0.19 | 1.3E-01 | 0.14 |
| GTPBP1 | -0.11 | -0.06 | 0.04 | 0.03 | 0.07 | 0.03 | 1.3E-01 | 0.08 |
| DCXR | 0.01 | -0.03 | -0.18 | 0.12 | 0.05 | 0.01 | 1.3E-01 | 0.12 |
| GNG5 | -0.04 | 0.05 | 0.21 | -0.07 | -0.12 | -0.05 | 1.3E-01 | -0.15 |
| NEMF | 0.04 | 0.01 | 0.08 | 0.01 | -0.03 | -0.12 | 1.3E-01 | -0.09 |
| IL1R1 | 0.08 | -0.10 | 0.45 | -0.27 | -0.16 | -0.12 | 1.3E-01 | -0.33 |
| KMO | 0.08 | 0.04 | 0.02 | -0.14 | -0.01 | 0.00 | 1.3E-01 | -0.09 |
| CEP128 | 0.05 | 0.10 | 0.01 | -0.13 | 0.05 | -0.10 | 1.3E-01 | -0.11 |
| RXYLT1 | 0.03 | -0.02 | 0.14 | -0.02 | -0.04 | -0.10 | 1.3E-01 | -0.10 |
| MADD | -0.13 | -0.11 | 0.03 | 0.16 | 0.01 | 0.03 | 1.3E-01 | 0.14 |
| ERLIN2 | 0.00 | -0.06 | -0.10 | 0.14 | -0.01 | 0.02 | 1.3E-01 | 0.10 |
| DCAF7 | -0.13 | -0.05 | -0.05 | 0.01 | -0.02 | 0.22 | 1.3E-01 | 0.15 |
| GDAP2 | -0.11 | -0.17 | 0.06 | 0.07 | 0.11 | 0.02 | 1.3E-01 | 0.14 |
| UTP20 | 0.04 | 0.01 | 0.03 | 0.02 | -0.03 | -0.07 | 1.3E-01 | -0.05 |
| SLC46A1 | -0.04 | 0.07 | 0.40 | -0.38 | -0.16 | -0.01 | 1.3E-01 | -0.32 |
| MT-CYB | -0.06 | -0.18 | -0.43 | 0.01 | -0.02 | 0.51 | 1.3E-01 | 0.39 |
| PKP2 | -0.19 | -0.07 | 0.01 | -0.03 | 0.09 | 0.15 | 1.3E-01 | 0.15 |
| NUDT9 | -0.06 | 0.04 | -0.20 | 0.01 | 0.08 | 0.12 | 1.3E-01 | 0.15 |
| VPS33B | -0.25 | -0.02 | -0.02 | 0.15 | 0.13 | -0.03 | 1.3E-01 | 0.18 |
| ARRDC1 | -0.02 | 0.06 | 0.06 | -0.01 | -0.01 | -0.08 | 1.3E-01 | -0.06 |
| SFSWAP | 0.04 | 0.01 | 0.04 | 0.03 | -0.04 | -0.07 | 1.3E-01 | -0.06 |
| REPIN1 | -0.01 | 0.10 | 0.00 | -0.04 | -0.02 | -0.04 | 1.3E-01 | -0.07 |
| FKBP10 | 0.03 | 0.00 | 0.00 | -0.02 | -0.02 | 0.00 | 1.3E-01 | -0.02 |
| MRPL24 | 0.12 | 0.08 | -0.05 | -0.04 | -0.03 | -0.10 | 1.3E-01 | -0.11 |
| PGRMC2 | 0.08 | 0.05 | 0.03 | 0.00 | 0.00 | -0.17 | 1.3E-01 | -0.11 |
| TBC1D24 | -0.18 | -0.29 | 0.07 | 0.17 | -0.04 | 0.21 | 1.3E-01 | 0.25 |
| SNIP1 | 0.03 | 0.07 | 0.00 | -0.08 | 0.01 | -0.03 | 1.3E-01 | -0.06 |
| RALGAPA1 | -0.40 | -0.23 | 0.07 | -0.07 | 0.22 | 0.29 | 1.3E-01 | 0.34 |
| VPS13B | -0.26 | -0.12 | 0.07 | 0.01 | 0.11 | 0.16 | 1.3E-01 | 0.20 |
| ULK2 | 0.15 | 0.35 | 0.18 | -0.21 | 0.10 | -0.85 | 1.3E-01 | -0.55 |
| PRDX4 | -0.19 | 0.30 | 0.31 | -0.22 | -0.21 | -0.10 | 1.3E-01 | -0.32 |
| DHX38 | 0.06 | -0.01 | 0.04 | -0.04 | 0.01 | -0.07 | 1.3E-01 | -0.06 |
| TASP1 | -0.02 | 0.08 | 0.11 | -0.09 | 0.03 | -0.13 | 1.3E-01 | -0.12 |
| KIAA1191 | 0.02 | -0.02 | -0.30 | 0.07 | 0.09 | 0.11 | 1.3E-01 | 0.19 |
| SETD1A | -0.22 | -0.07 | -0.17 | -0.15 | 0.30 | 0.23 | 1.3E-01 | 0.28 |
| ZSWIM6 | -3.02 | -2.71 | -2.05 | 1.73 | -2.63 | 1.00 | 1.3E-01 | 2.63 |
| MRPL49 | 0.13 | 0.11 | -0.06 | -0.01 | -0.09 | -0.11 | 1.3E-01 | -0.13 |
| GEN1 | -0.39 | -0.28 | 0.12 | 0.04 | 0.11 | 0.28 | 1.3E-01 | 0.32 |
| JUND | -0.09 | -0.17 | -0.01 | 0.09 | -0.06 | 0.21 | 1.3E-01 | 0.17 |
| STK26 | -0.15 | -0.13 | 0.01 | -0.06 | 0.13 | 0.17 | 1.3E-01 | 0.16 |
| U2AF1 | 0.04 | -0.04 | 0.17 | -0.05 | -0.07 | -0.08 | 1.3E-01 | -0.12 |
| IBA57 | -0.04 | -0.20 | 0.00 | -0.02 | 0.09 | 0.14 | 1.3E-01 | 0.15 |
| DHCR24 | 0.05 | 0.03 | 0.00 | -0.03 | 0.01 | -0.07 | 1.3E-01 | -0.06 |

| | | | | | | | | |
|---|---|---|---|---|---|---|---|---|
| **UQCC3** | -0.06 | 0.15 | 0.27 | -0.33 | -0.01 | -0.08 | 1.3E-01 | -0.26 |
| **TPD52** | -0.09 | 0.15 | 0.17 | -0.12 | -0.02 | -0.13 | 1.3E-01 | -0.17 |
| **MMGT1** | 0.17 | 0.19 | 0.01 | -0.16 | 0.08 | -0.37 | 1.3E-01 | -0.27 |
| **MRPL37** | 0.03 | 0.06 | 0.03 | 0.03 | -0.12 | -0.04 | 1.3E-01 | -0.09 |
| **MAGI1** | -0.21 | -0.02 | -0.02 | 0.00 | 0.05 | 0.17 | 1.3E-01 | 0.15 |
| **TCF7L2** | -0.01 | -0.09 | -0.02 | -0.03 | 0.05 | 0.09 | 1.3E-01 | 0.08 |
| **MBD1** | -0.01 | -0.09 | -0.03 | 0.04 | -0.03 | 0.10 | 1.3E-01 | 0.08 |
| **DNAJC10** | 0.00 | -0.03 | -0.05 | -0.01 | 0.07 | 0.01 | 1.3E-01 | 0.05 |
| **ARL2** | 0.01 | -0.44 | -0.08 | -0.04 | 0.22 | 0.23 | 1.3E-01 | 0.31 |
| **RAP2B** | -0.04 | -0.07 | -0.10 | 0.20 | 0.02 | -0.03 | 1.3E-01 | 0.13 |
| **ANXA1** | 0.03 | -0.28 | -0.09 | 0.18 | 0.15 | -0.03 | 1.3E-01 | 0.21 |
| **NFIC** | 0.01 | -0.03 | -0.12 | -0.01 | 0.08 | 0.06 | 1.3E-01 | 0.09 |
| **RECQL4** | -0.02 | 0.03 | 0.15 | -0.08 | -0.01 | -0.08 | 1.3E-01 | -0.11 |
| **WDFY1** | -0.02 | 0.01 | 0.25 | -0.18 | -0.05 | -0.06 | 1.3E-01 | -0.18 |
| **RELL1** | 0.13 | 0.03 | 0.07 | -0.24 | 0.03 | -0.05 | 1.3E-01 | -0.16 |
| **FHOD1** | -0.01 | -0.05 | -0.04 | 0.05 | 0.07 | -0.03 | 1.3E-01 | 0.07 |
| **TCAIM** | -0.28 | 0.04 | -0.08 | 0.18 | 0.00 | 0.10 | 1.3E-01 | 0.20 |
| **EEA1** | -0.04 | -0.03 | -0.08 | 0.03 | -0.03 | 0.15 | 1.3E-01 | 0.10 |
| **DHX9** | 0.01 | -0.04 | -0.04 | 0.03 | -0.01 | 0.05 | 1.3E-01 | 0.05 |
| **GTF3C2** | 0.01 | -0.04 | -0.14 | 0.11 | 0.01 | 0.03 | 1.3E-01 | 0.11 |
| **MKRN1** | -0.08 | -0.09 | -0.05 | 0.20 | 0.02 | -0.03 | 1.3E-01 | 0.14 |
| **CLTB** | -0.15 | -0.09 | 0.02 | 0.16 | -0.02 | 0.06 | 1.3E-01 | 0.14 |
| **HACL1** | -0.06 | 0.02 | -0.22 | 0.06 | 0.16 | 0.01 | 1.3E-01 | 0.16 |
| **VARS2** | -0.20 | -0.19 | 0.09 | 0.08 | 0.13 | 0.05 | 1.3E-01 | 0.18 |
| **ZCCHC3** | -0.11 | -0.02 | 0.00 | 0.05 | -0.01 | 0.09 | 1.3E-01 | 0.09 |
| **SORL1** | -0.02 | 0.01 | 0.17 | -0.08 | -0.05 | -0.05 | 1.3E-01 | -0.11 |
| **PATJ** | -0.18 | -0.07 | 0.02 | -0.02 | 0.07 | 0.15 | 1.3E-01 | 0.14 |
| **SIRPA** | -0.65 | -0.46 | -0.33 | 1.01 | -0.04 | -0.21 | 1.3E-01 | 0.74 |
| **CARMIL1** | 0.05 | 0.09 | 0.08 | -0.23 | 0.05 | -0.07 | 1.3E-01 | -0.16 |
| **CBLL1** | 0.05 | 0.17 | 0.02 | 0.01 | -0.03 | -0.24 | 1.3E-01 | -0.17 |
| **TTC27** | -0.53 | -0.32 | 0.13 | 0.31 | 0.27 | -0.04 | 1.3E-01 | 0.42 |
| **KDM5C** | -0.48 | -0.18 | 0.04 | 0.37 | -0.06 | 0.17 | 1.3E-01 | 0.37 |
| **ZYG11B** | -0.04 | 0.19 | 0.07 | -0.08 | 0.00 | -0.17 | 1.3E-01 | -0.16 |
| **LSM5** | -0.12 | -0.11 | 0.02 | -0.01 | 0.02 | 0.18 | 1.3E-01 | 0.13 |
| **WDR12** | -0.02 | 0.06 | 0.06 | -0.05 | 0.02 | -0.08 | 1.3E-01 | -0.07 |
| **SP3** | 0.01 | -0.03 | -0.07 | 0.03 | -0.01 | 0.06 | 1.3E-01 | 0.06 |
| **TUBD1** | 0.10 | 0.02 | 0.07 | 0.06 | -0.12 | -0.15 | 1.3E-01 | -0.13 |
| **DDAH2** | -0.04 | 0.10 | 0.19 | -0.22 | 0.01 | -0.07 | 1.3E-01 | -0.17 |
| **MYG1** | -0.17 | -0.05 | -0.01 | -0.05 | 0.16 | 0.10 | 1.3E-01 | 0.15 |
| **MYO1C** | -0.10 | -0.04 | 0.01 | 0.00 | 0.02 | 0.11 | 1.3E-01 | 0.09 |
| **EIF2S1** | 0.00 | 0.02 | 0.15 | -0.06 | 0.02 | -0.15 | 1.3E-01 | -0.12 |
| **CLGN** | 0.05 | 0.05 | 0.04 | -0.06 | 0.04 | -0.13 | 1.4E-01 | -0.10 |
| **VANGL1** | -0.04 | 0.06 | 0.12 | -0.08 | -0.07 | 0.00 | 1.4E-01 | -0.10 |
| **EIF4H** | 0.05 | 0.19 | -0.04 | -0.03 | -0.03 | -0.16 | 1.4E-01 | -0.15 |
| **ING5** | 0.06 | 0.07 | 0.00 | -0.05 | 0.03 | -0.12 | 1.4E-01 | -0.09 |
| **UPF2** | -0.04 | -0.04 | 0.01 | 0.00 | 0.05 | 0.02 | 1.4E-01 | 0.04 |
| **KPNA3** | 0.04 | 0.06 | 0.09 | -0.05 | 0.04 | -0.20 | 1.4E-01 | -0.13 |
| **PMM2** | 0.17 | 0.22 | -0.09 | -0.23 | 0.01 | -0.13 | 1.4E-01 | -0.22 |

| | | | | | | | | |
|---|---|---|---|---|---|---|---|---|
| **RPS8** | -0.01 | -0.05 | -0.02 | 0.05 | 0.06 | -0.03 | 1.4E-01 | 0.06 |
| **MLEC** | 0.09 | 0.13 | 0.02 | 0.02 | -0.03 | -0.26 | 1.4E-01 | -0.17 |
| **RSPRY1** | -0.11 | -0.03 | -0.01 | 0.09 | -0.04 | 0.08 | 1.4E-01 | 0.09 |
| **NDUFS7** | -0.03 | 0.01 | -0.07 | 0.07 | 0.01 | 0.01 | 1.4E-01 | 0.05 |
| **WDR18** | -0.05 | -0.01 | -0.03 | 0.08 | 0.03 | -0.02 | 1.4E-01 | 0.06 |
| **SMS** | 0.05 | -0.18 | -0.08 | 0.11 | 0.06 | 0.02 | 1.4E-01 | 0.13 |
| **BAZ1A** | -0.03 | -0.03 | -0.10 | -0.03 | 0.04 | 0.13 | 1.4E-01 | 0.10 |
| **FH** | 0.24 | 0.28 | -0.16 | -0.12 | -0.16 | -0.15 | 1.4E-01 | -0.26 |
| **PPIF** | 0.17 | 0.07 | -0.04 | -0.16 | 0.00 | -0.06 | 1.4E-01 | -0.14 |
| **NTPCR** | 0.02 | -0.02 | 0.10 | -0.04 | -0.05 | -0.02 | 1.4E-01 | -0.07 |
| **ABLIM1** | -0.12 | -0.02 | -0.02 | 0.05 | -0.02 | 0.13 | 1.4E-01 | 0.10 |
| **FBXL20** | 0.02 | 0.01 | 0.18 | -0.13 | 0.04 | -0.14 | 1.4E-01 | -0.15 |
| **BTF3** | 0.24 | 0.10 | 0.09 | 0.13 | -0.23 | -0.44 | 1.4E-01 | -0.33 |
| **LRRC8E** | -0.01 | 0.04 | 0.12 | -0.08 | 0.03 | -0.11 | 1.4E-01 | -0.11 |
| **C8orf82** | 0.06 | -0.03 | 0.29 | -0.01 | -0.09 | -0.28 | 1.4E-01 | -0.23 |
| **ICE2** | -0.05 | -0.23 | -0.28 | -0.03 | -0.01 | 0.47 | 1.4E-01 | 0.33 |
| **HMGXB4** | -0.04 | -0.02 | -0.19 | 0.20 | -0.01 | 0.03 | 1.4E-01 | 0.15 |
| **IQGAP1** | -0.05 | -0.04 | -0.01 | 0.00 | 0.00 | 0.10 | 1.4E-01 | 0.07 |
| **MICA** | -0.17 | -0.18 | 0.08 | 0.01 | 0.08 | 0.15 | 1.4E-01 | 0.17 |
| **CCAR2** | 0.02 | -0.06 | -0.13 | 0.02 | 0.12 | 0.01 | 1.4E-01 | 0.11 |
| **STX18** | -0.03 | 0.02 | 0.18 | -0.02 | -0.08 | -0.09 | 1.4E-01 | -0.12 |
| **ADRM1** | 0.12 | -0.34 | -0.24 | 0.03 | 0.20 | 0.14 | 1.4E-01 | 0.28 |
| **PDCD6IP** | 0.02 | 0.03 | 0.07 | 0.01 | -0.01 | -0.13 | 1.4E-01 | -0.08 |
| **SOD1** | 0.59 | 0.51 | -0.37 | -0.72 | -0.18 | -0.31 | 1.4E-01 | -0.65 |
| **RB1** | -0.02 | -0.01 | -0.15 | -0.03 | 0.08 | 0.12 | 1.4E-01 | 0.12 |
| **IDH3B** | -0.11 | -0.20 | -0.32 | 0.54 | -0.05 | -0.02 | 1.4E-01 | 0.37 |
| **PSMB4** | -0.08 | -0.23 | -0.08 | 0.07 | -0.08 | 0.33 | 1.4E-01 | 0.24 |
| **PSME3** | 0.04 | 0.12 | -0.02 | -0.11 | 0.00 | -0.05 | 1.4E-01 | -0.10 |
| **CZIB** | -0.01 | 0.01 | 0.22 | -0.15 | -0.05 | -0.04 | 1.4E-01 | -0.15 |
| **GPATCH4** | 0.00 | 0.04 | 0.04 | -0.07 | 0.02 | -0.04 | 1.4E-01 | -0.05 |
| **EIF3J** | -0.02 | 0.06 | 0.06 | -0.08 | 0.00 | -0.03 | 1.4E-01 | -0.07 |
| **RPL9** | -0.02 | -0.06 | -0.07 | 0.13 | 0.07 | -0.05 | 1.4E-01 | 0.10 |
| **GNS** | 0.14 | -0.05 | 0.07 | -0.01 | -0.06 | -0.09 | 1.4E-01 | -0.11 |
| **DHRS13** | 0.08 | -0.04 | 0.20 | -0.12 | 0.03 | -0.19 | 1.4E-01 | -0.18 |
| **MRPL1** | 0.10 | 0.12 | -0.06 | -0.02 | -0.04 | -0.11 | 1.4E-01 | -0.11 |
| **SLC4A7** | -0.11 | 0.12 | 0.40 | -0.33 | -0.07 | -0.11 | 1.4E-01 | -0.31 |
| **SMCHD1** | 0.00 | -0.01 | -0.05 | 0.01 | 0.00 | 0.04 | 1.4E-01 | 0.04 |
| **UCKL1** | -0.18 | -0.40 | 0.11 | 0.06 | 0.22 | 0.11 | 1.4E-01 | 0.29 |
| **GRTP1** | 0.08 | 0.10 | 0.10 | -0.19 | -0.23 | 0.11 | 1.4E-01 | -0.20 |
| **TXK** | -0.05 | -0.11 | -0.09 | -0.07 | 0.07 | 0.22 | 1.4E-01 | 0.16 |
| **RAB4A** | 0.00 | -0.01 | 0.26 | -0.12 | -0.14 | -0.02 | 1.4E-01 | -0.17 |
| **BMS1** | 0.04 | 0.03 | 0.01 | 0.00 | 0.00 | -0.08 | 1.4E-01 | -0.05 |
| **PAK2** | 0.00 | 0.00 | -0.12 | 0.03 | 0.06 | 0.02 | 1.4E-01 | 0.08 |
| **PIGO** | -0.10 | -0.06 | 0.03 | 0.03 | 0.01 | 0.08 | 1.4E-01 | 0.08 |
| **DCTN6** | 0.00 | 0.06 | 0.08 | -0.11 | -0.09 | 0.04 | 1.4E-01 | -0.10 |
| **SPPL2B** | -0.08 | -0.08 | 0.02 | -0.01 | 0.03 | 0.12 | 1.4E-01 | 0.09 |
| **ACTR2** | 0.04 | 0.08 | 0.01 | -0.01 | 0.00 | -0.13 | 1.4E-01 | -0.09 |
| **KLHDC1** | -0.12 | 0.07 | -0.24 | 0.08 | 0.02 | 0.15 | 1.4E-01 | 0.18 |

| | | | | | | | | |
|---|---|---|---|---|---|---|---|---|
| ALG5 | 0.08 | -0.23 | -0.20 | 0.20 | -0.03 | 0.12 | 1.4E-01 | 0.22 |
| RRAS | -0.07 | 0.08 | 0.27 | -0.18 | -0.13 | -0.02 | 1.4E-01 | -0.20 |
| EIF2B4 | -0.06 | -0.11 | -0.04 | 0.17 | 0.08 | -0.06 | 1.4E-01 | 0.13 |
| YY1AP1 | 0.02 | -0.30 | -0.08 | 0.17 | -0.05 | 0.18 | 1.4E-01 | 0.22 |
| CEP43 | -0.32 | 0.12 | -0.20 | 0.15 | 0.09 | 0.09 | 1.4E-01 | 0.25 |
| PPP1R13L | -0.19 | -0.10 | 0.05 | -0.02 | 0.12 | 0.11 | 1.4E-01 | 0.15 |
| CASD1 | 0.08 | 0.14 | 0.07 | 0.04 | -0.04 | -0.33 | 1.4E-01 | -0.21 |
| USP28 | 0.06 | -0.71 | -0.14 | 0.37 | 0.16 | 0.04 | 1.4E-01 | 0.45 |
| NADSYN1 | -0.04 | 0.02 | 0.41 | -0.32 | -0.11 | -0.07 | 1.4E-01 | -0.30 |
| PARL | 0.14 | 0.08 | -0.04 | 0.02 | -0.09 | -0.13 | 1.4E-01 | -0.13 |
| UBQLN4 | 0.04 | 0.00 | 0.04 | 0.01 | -0.02 | -0.07 | 1.4E-01 | -0.05 |
| WLS | -0.05 | 0.10 | 0.14 | -0.15 | -0.07 | 0.01 | 1.4E-01 | -0.13 |
| ARFGAP2 | -0.26 | -0.28 | -0.01 | -0.09 | 0.08 | 0.44 | 1.4E-01 | 0.33 |
| NADK2 | -0.03 | -0.12 | 0.01 | 0.07 | -0.02 | 0.08 | 1.4E-01 | 0.09 |
| STRIP1 | -0.12 | -0.05 | 0.03 | 0.02 | 0.09 | 0.03 | 1.4E-01 | 0.09 |
| MAGT1 | -0.01 | 0.05 | 0.13 | 0.01 | -0.04 | -0.16 | 1.4E-01 | -0.12 |
| APOC3 | -0.10 | -0.03 | -0.26 | 0.00 | 0.00 | 0.33 | 1.4E-01 | 0.24 |
| PSME3IP1 | 0.06 | -0.02 | 0.19 | -0.22 | -0.03 | -0.01 | 1.4E-01 | -0.16 |
| RPL18A | -0.03 | -0.09 | -0.06 | 0.18 | 0.01 | -0.02 | 1.4E-01 | 0.12 |
| PARP9 | -0.11 | -0.03 | -0.01 | 0.06 | 0.11 | -0.03 | 1.4E-01 | 0.09 |
| TBL3 | 0.02 | 0.01 | 0.08 | -0.10 | 0.01 | -0.02 | 1.4E-01 | -0.07 |
| THOC5 | 0.12 | 0.10 | 0.02 | 0.05 | -0.06 | -0.26 | 1.4E-01 | -0.17 |
| TOMM34 | 0.02 | -0.05 | -0.06 | 0.07 | 0.00 | 0.01 | 1.4E-01 | 0.06 |
| TOP2A | 0.10 | 0.07 | 0.00 | -0.16 | -0.05 | 0.03 | 1.4E-01 | -0.11 |
| HMGN2 | -0.04 | -0.05 | -0.05 | -0.03 | 0.02 | 0.13 | 1.4E-01 | 0.09 |
| LMCD1 | 0.04 | 0.08 | 0.19 | -0.32 | 0.07 | -0.11 | 1.4E-01 | -0.22 |
| PPP3CA | 0.13 | -0.03 | 0.15 | 0.05 | -0.22 | -0.10 | 1.4E-01 | -0.17 |
| MON2 | -0.15 | -0.06 | -0.02 | -0.04 | 0.19 | 0.05 | 1.5E-01 | 0.14 |
| UGT2B4 | -2.13 | -1.55 | -0.15 | 0.75 | -0.98 | 1.23 | 1.5E-01 | 1.61 |
| PCYOX1 | 0.05 | -0.09 | -0.20 | 0.12 | -0.01 | 0.09 | 1.5E-01 | 0.15 |
| YIPF5 | 0.01 | 0.05 | 0.10 | -0.04 | 0.03 | -0.17 | 1.5E-01 | -0.11 |
| EFNA5 | -0.04 | 0.19 | 0.22 | -0.26 | -0.29 | 0.10 | 1.5E-01 | -0.27 |
| HINT1 | 0.04 | 0.18 | -0.03 | -0.12 | 0.00 | -0.09 | 1.5E-01 | -0.13 |
| KCTD18 | 0.01 | -0.01 | 0.12 | -0.07 | -0.04 | -0.02 | 1.5E-01 | -0.08 |
| SLC16A1 | -0.05 | 0.13 | 0.08 | -0.04 | -0.11 | -0.02 | 1.5E-01 | -0.11 |
| SLC39A11 | 0.08 | -0.02 | 0.14 | -0.11 | 0.04 | -0.16 | 1.5E-01 | -0.14 |
| API5 | 0.12 | 0.08 | -0.05 | -0.01 | -0.05 | -0.10 | 1.5E-01 | -0.10 |
| NDUFB6 | 0.05 | 0.06 | -0.01 | 0.00 | -0.01 | -0.10 | 1.5E-01 | -0.07 |
| ARMC1 | 0.05 | 0.02 | 0.01 | -0.09 | 0.01 | -0.01 | 1.5E-01 | -0.06 |
| ALKBH5 | -0.03 | 0.03 | 0.14 | -0.02 | -0.04 | -0.10 | 1.5E-01 | -0.10 |
| NAIP | -0.12 | -0.15 | 0.03 | -0.05 | 0.15 | 0.11 | 1.5E-01 | 0.15 |
| PAM16 | -0.19 | -0.28 | 0.04 | 0.07 | -0.05 | 0.33 | 1.5E-01 | 0.26 |
| RPL5 | 0.05 | 0.06 | 0.18 | 0.02 | -0.03 | -0.31 | 1.5E-01 | -0.20 |
| LAMTOR1 | 0.00 | 0.08 | 0.14 | -0.02 | 0.01 | -0.23 | 1.5E-01 | -0.15 |
| PIGN | 0.10 | 0.08 | 0.12 | 0.12 | -0.21 | -0.25 | 1.5E-01 | -0.21 |
| SSBP1 | 0.00 | -0.12 | -0.63 | 0.34 | -0.10 | 0.31 | 1.5E-01 | 0.43 |
| DEDD | -0.04 | -0.19 | 0.00 | 0.04 | -0.01 | 0.17 | 1.5E-01 | 0.14 |
| EFNB1 | -0.08 | 0.10 | 0.59 | -0.42 | -0.57 | 0.08 | 1.5E-01 | -0.51 |

| | | | | | | | | |
|---|---|---|---|---|---|---|---|---|
| **TRAPPC6B** | -0.02 | 0.02 | -0.24 | 0.09 | 0.03 | 0.10 | 1.5E-01 | 0.15 |
| **MBLAC2** | -0.03 | 0.03 | 0.15 | -0.01 | -0.05 | -0.12 | 1.5E-01 | -0.11 |
| **ARL6IP5** | 0.04 | -0.07 | -0.14 | 0.09 | 0.09 | -0.02 | 1.5E-01 | 0.11 |
| **LRRC8D** | -0.28 | -0.06 | 0.03 | 0.04 | 0.03 | 0.20 | 1.5E-01 | 0.19 |
| **POMT1** | 0.03 | -0.01 | 0.18 | -0.07 | 0.02 | -0.17 | 1.5E-01 | -0.14 |
| **PIP4K2B** | 0.12 | 0.13 | -0.06 | -0.12 | 0.01 | -0.10 | 1.5E-01 | -0.13 |
| **AQP3** | 0.02 | 0.05 | 0.27 | -0.35 | -0.12 | 0.06 | 1.5E-01 | -0.25 |
| **ZNF532** | -0.62 | -0.66 | -0.53 | 1.20 | -0.16 | -0.28 | 1.5E-01 | 0.86 |
| **GCFC2** | -0.09 | 0.07 | -0.34 | 0.11 | 0.03 | 0.17 | 1.5E-01 | 0.22 |
| **MPND** | -0.16 | -0.11 | -0.01 | 0.05 | -0.05 | 0.25 | 1.5E-01 | 0.18 |
| **GLMN** | -0.08 | 0.01 | -0.11 | 0.04 | -0.03 | 0.15 | 1.5E-01 | 0.11 |
| **HMOX2** | 0.25 | 0.08 | -0.07 | -0.06 | -0.04 | -0.21 | 1.5E-01 | -0.19 |
| **HNRNPA1L2** | 0.05 | -0.01 | 0.05 | -0.08 | -0.05 | 0.03 | 1.5E-01 | -0.07 |
| **KIAA1671** | -0.13 | 0.04 | -0.06 | 0.05 | 0.07 | 0.02 | 1.5E-01 | 0.09 |
| **ZNF292** | -0.01 | -0.07 | -0.10 | 0.01 | -0.02 | 0.18 | 1.5E-01 | 0.12 |
| **CCDC90B** | 0.07 | 0.13 | -0.05 | -0.07 | 0.00 | -0.10 | 1.5E-01 | -0.11 |
| **APEX1** | 0.13 | 0.19 | -0.09 | -0.09 | -0.03 | -0.13 | 1.5E-01 | -0.16 |
| **TPX2** | -0.06 | -0.07 | -0.06 | -0.02 | 0.00 | 0.19 | 1.5E-01 | 0.12 |
| **MPHOSPH10** | 0.07 | 0.02 | 0.01 | 0.03 | -0.05 | -0.08 | 1.5E-01 | -0.06 |
| **ABCB6** | -0.07 | 0.07 | 0.21 | -0.07 | -0.10 | -0.06 | 1.5E-01 | -0.15 |
| **RPP30** | 0.11 | 0.02 | 0.06 | -0.05 | 0.04 | -0.20 | 1.5E-01 | -0.13 |
| **CXADR** | -0.15 | -0.10 | -0.07 | 0.19 | -0.13 | 0.20 | 1.5E-01 | 0.20 |
| **SMARCA5** | 0.00 | -0.05 | -0.04 | -0.02 | 0.02 | 0.08 | 1.5E-01 | 0.06 |
| **NACAD** | -0.15 | -0.11 | -0.06 | 0.19 | -0.13 | 0.21 | 1.5E-01 | 0.20 |
| **POU2F3** | -0.07 | 0.03 | 0.34 | -0.17 | -0.15 | -0.05 | 1.5E-01 | -0.23 |
| **LETMD1** | 0.04 | 0.08 | 0.01 | -0.04 | -0.13 | 0.03 | 1.5E-01 | -0.09 |
| **SRSF8** | 0.29 | -0.17 | 0.40 | 0.01 | -0.54 | -0.20 | 1.5E-01 | -0.42 |
| **IFFO1** | -0.11 | -0.10 | 0.05 | 0.00 | 0.09 | 0.07 | 1.5E-01 | 0.10 |
| **PRKACA** | -0.24 | -0.22 | 0.06 | 0.22 | -0.09 | 0.20 | 1.5E-01 | 0.25 |
| **MTERF4** | 0.14 | -0.01 | 0.03 | -0.12 | -0.10 | 0.03 | 1.5E-01 | -0.12 |
| **NDUFS2** | 0.00 | -0.01 | -0.05 | 0.04 | -0.02 | 0.03 | 1.5E-01 | 0.04 |
| **PRXL2A** | -0.10 | -0.23 | 0.07 | 0.13 | 0.00 | 0.10 | 1.5E-01 | 0.17 |
| **PIEZO1** | -0.02 | 0.06 | 0.17 | -0.19 | 0.01 | -0.05 | 1.5E-01 | -0.15 |
| **LRCH3** | -0.01 | 0.00 | 0.21 | -0.03 | -0.05 | -0.15 | 1.5E-01 | -0.14 |
| **PHF19** | 0.05 | 0.03 | -0.02 | 0.00 | -0.03 | -0.03 | 1.5E-01 | -0.04 |
| **POLDIP2** | 0.15 | 0.03 | -0.03 | -0.08 | -0.06 | -0.01 | 1.5E-01 | -0.10 |
| **TMED9** | -0.03 | -0.10 | 0.01 | 0.08 | -0.02 | 0.04 | 1.5E-01 | 0.08 |
| **PLK2** | -0.07 | 0.03 | 0.35 | -0.07 | -0.19 | -0.12 | 1.5E-01 | -0.23 |
| **GAPVD1** | 0.26 | -0.13 | 0.30 | -0.38 | -0.25 | 0.06 | 1.5E-01 | -0.33 |
| **RRP36** | 0.10 | 0.04 | -0.02 | 0.01 | -0.06 | -0.08 | 1.5E-01 | -0.08 |
| **PICK1** | -0.11 | -0.15 | 0.05 | 0.15 | -0.01 | 0.05 | 1.5E-01 | 0.14 |
| **NFYC** | -0.02 | -0.11 | -0.06 | -0.05 | 0.05 | 0.17 | 1.5E-01 | 0.12 |
| **SLC39A10** | -0.07 | 0.10 | 0.19 | -0.17 | -0.10 | 0.01 | 1.5E-01 | -0.16 |
| **IST1** | 0.17 | 0.11 | -0.07 | -0.06 | -0.01 | -0.16 | 1.5E-01 | -0.15 |
| **CDC7** | -0.28 | 0.01 | -0.15 | -0.10 | 0.31 | 0.13 | 1.6E-01 | 0.25 |
| **SLC25A25** | -0.01 | 0.03 | 0.06 | -0.03 | -0.07 | 0.01 | 1.6E-01 | -0.06 |
| **TPR** | 0.08 | 0.05 | -0.03 | -0.08 | -0.01 | -0.02 | 1.6E-01 | -0.07 |
| **SRSF6** | 0.02 | -0.02 | 0.05 | 0.00 | -0.04 | -0.02 | 1.6E-01 | -0.04 |

| | | | | | | | | |
|---|---|---|---|---|---|---|---|---|
| **OAT** | -0.03 | 0.01 | -0.12 | 0.00 | 0.03 | 0.11 | 1.6E-01 | 0.09 |
| **NPNT** | -0.02 | 0.05 | 0.10 | -0.13 | -0.02 | 0.00 | 1.6E-01 | -0.09 |
| **JADE1** | 0.06 | 0.00 | 0.06 | -0.09 | 0.05 | -0.10 | 1.6E-01 | -0.09 |
| **GARRE1** | 0.00 | 0.02 | 0.16 | -0.12 | 0.04 | -0.11 | 1.6E-01 | -0.12 |
| **NUDCD1** | -0.20 | -0.19 | -0.10 | -0.18 | 0.38 | 0.19 | 1.6E-01 | 0.29 |
| **SREBF2** | 0.00 | 0.03 | 0.05 | -0.02 | -0.08 | 0.02 | 1.6E-01 | -0.06 |
| **MTERF1** | 0.07 | 0.05 | 0.04 | -0.13 | 0.07 | -0.12 | 1.6E-01 | -0.11 |
| **EIF3F** | -0.04 | -0.02 | -0.10 | 0.16 | 0.02 | -0.03 | 1.6E-01 | 0.10 |
| **ZDHHC5** | -0.04 | 0.04 | 0.09 | -0.02 | -0.03 | -0.04 | 1.6E-01 | -0.07 |
| **ANKRD54** | -0.16 | -0.15 | 0.04 | 0.20 | -0.04 | 0.07 | 1.6E-01 | 0.17 |
| **RPL30** | -0.06 | -0.06 | 0.03 | 0.06 | 0.03 | -0.01 | 1.6E-01 | 0.06 |
| **PPIH** | 0.02 | -0.06 | -0.05 | 0.00 | 0.07 | 0.01 | 1.6E-01 | 0.06 |
| **LRP6** | -0.10 | 0.39 | 0.10 | -0.37 | -0.05 | -0.09 | 1.6E-01 | -0.30 |
| **RGPD8** | 0.09 | -0.10 | -0.38 | 0.15 | 0.13 | 0.04 | 1.6E-01 | 0.24 |
| **SAFB2** | 0.01 | -0.03 | -0.04 | 0.01 | 0.00 | 0.05 | 1.6E-01 | 0.04 |
| **DUSP23** | 0.02 | 0.07 | 0.05 | -0.08 | -0.13 | 0.06 | 1.6E-01 | -0.10 |
| **MT-CO2** | -0.11 | -0.08 | 0.03 | 0.01 | 0.00 | 0.14 | 1.6E-01 | 0.11 |
| **RAB12** | -0.15 | -0.07 | -0.11 | 0.07 | -0.09 | 0.30 | 1.6E-01 | 0.20 |
| **ZBTB33** | 0.03 | 0.02 | 0.10 | -0.14 | -0.07 | 0.05 | 1.6E-01 | -0.11 |
| **COX14** | 0.10 | 0.08 | -0.05 | -0.02 | -0.09 | -0.03 | 1.6E-01 | -0.09 |
| **NES** | -0.24 | -0.10 | 0.07 | 0.12 | 0.00 | 0.12 | 1.6E-01 | 0.17 |
| **LSM12** | -0.02 | -0.10 | -0.09 | 0.20 | -0.05 | 0.03 | 1.6E-01 | 0.13 |
| **PRPS1** | 0.21 | 0.30 | -0.12 | -0.10 | -0.01 | -0.39 | 1.6E-01 | -0.30 |
| **SLC24A5** | 0.19 | 0.06 | 0.02 | -0.03 | 0.02 | -0.31 | 1.6E-01 | -0.20 |
| **ZNF326** | -0.05 | 0.04 | 0.15 | -0.07 | -0.05 | -0.03 | 1.6E-01 | -0.10 |
| **COX5B** | 0.00 | -0.04 | -0.09 | 0.11 | -0.03 | 0.04 | 1.6E-01 | 0.08 |
| **RHPN1** | -0.37 | 0.09 | -0.10 | 0.14 | 0.06 | 0.12 | 1.6E-01 | 0.23 |
| **WARS2** | 0.01 | -0.02 | -0.11 | 0.10 | 0.01 | 0.01 | 1.6E-01 | 0.08 |
| **MYL6B** | 0.05 | -0.02 | 0.12 | 0.01 | -0.13 | -0.05 | 1.6E-01 | -0.10 |
| **CLMN** | 0.00 | -0.02 | -0.18 | -0.03 | 0.08 | 0.13 | 1.6E-01 | 0.13 |
| **ADAM10** | -0.02 | 0.00 | 0.20 | -0.08 | -0.03 | -0.09 | 1.6E-01 | -0.13 |
| **SP100** | -0.09 | 0.21 | 0.14 | 0.00 | -0.23 | -0.08 | 1.6E-01 | -0.19 |
| **MISP** | -0.20 | -0.10 | 0.05 | 0.10 | -0.03 | 0.15 | 1.6E-01 | 0.16 |
| **SIRT3** | -0.04 | -0.02 | -0.17 | -0.03 | 0.20 | 0.03 | 1.6E-01 | 0.15 |
| **NCS1** | -0.23 | -0.25 | 0.06 | 0.04 | -0.03 | 0.33 | 1.6E-01 | 0.26 |
| **AHR** | -0.21 | -0.12 | 0.04 | -0.07 | 0.15 | 0.17 | 1.6E-01 | 0.18 |
| **RAI14** | -0.01 | 0.03 | 0.04 | -0.05 | 0.01 | -0.03 | 1.6E-01 | -0.04 |
| **PYM1** | 0.05 | 0.06 | 0.09 | 0.09 | -0.15 | -0.18 | 1.6E-01 | -0.15 |
| **PYGB** | -0.06 | -0.11 | -0.06 | 0.01 | 0.24 | -0.04 | 1.6E-01 | 0.15 |
| **PRDX6** | 0.02 | -0.04 | -0.11 | -0.02 | 0.05 | 0.10 | 1.6E-01 | 0.09 |
| **IDI1** | -0.12 | 0.07 | -0.36 | -0.03 | 0.21 | 0.16 | 1.6E-01 | 0.25 |
| **LEO1** | 0.01 | -0.06 | -0.02 | 0.01 | 0.06 | 0.00 | 1.6E-01 | 0.05 |
| **PRR12** | -0.09 | -0.04 | -0.05 | 0.03 | -0.05 | 0.17 | 1.6E-01 | 0.11 |
| **RAP1A** | -0.20 | -0.07 | -0.05 | -0.01 | -0.02 | 0.30 | 1.6E-01 | 0.20 |
| **ACAT1** | 0.05 | -0.06 | -0.16 | 0.07 | 0.06 | 0.02 | 1.6E-01 | 0.11 |
| **AARSD1** | -0.24 | -0.41 | 0.13 | -0.01 | 0.29 | 0.12 | 1.6E-01 | 0.31 |
| **MEAF6** | -0.11 | -0.12 | 0.01 | 0.07 | -0.06 | 0.19 | 1.6E-01 | 0.14 |
| **ZNF428** | 0.06 | 0.12 | 0.05 | -0.28 | -0.01 | 0.03 | 1.6E-01 | -0.17 |

| | | | | | | | | |
|---|---|---|---|---|---|---|---|---|
| AURKAIP1 | 0.29 | 0.18 | -0.11 | -0.31 | 0.04 | -0.17 | 1.6E-01 | -0.26 |
| CDCA5 | -0.16 | -0.12 | 0.02 | -0.04 | 0.04 | 0.23 | 1.6E-01 | 0.16 |
| HSPA1L | 0.07 | 0.13 | 0.20 | -0.30 | 0.17 | -0.37 | 1.6E-01 | -0.29 |
| CCT4 | 0.11 | 0.11 | -0.06 | -0.07 | 0.00 | -0.11 | 1.6E-01 | -0.11 |
| PPIL4 | 0.10 | 0.03 | -0.02 | 0.00 | -0.03 | -0.08 | 1.6E-01 | -0.07 |
| TAOK1 | 0.01 | -0.01 | 0.06 | -0.02 | 0.00 | -0.04 | 1.7E-01 | -0.04 |
| USP20 | 0.09 | 0.20 | -0.07 | -0.04 | -0.16 | -0.04 | 1.7E-01 | -0.15 |
| CARS1 | -0.12 | 0.04 | -0.04 | 0.02 | 0.07 | 0.04 | 1.7E-01 | 0.08 |
| MAL2 | -0.23 | -0.20 | 0.05 | 0.03 | -0.02 | 0.31 | 1.7E-01 | 0.23 |
| LGALS8 | -0.15 | 0.00 | 0.00 | 0.04 | 0.00 | 0.09 | 1.7E-01 | 0.09 |
| PEX11B | 0.10 | 0.05 | 0.11 | -0.29 | 0.08 | -0.09 | 1.7E-01 | -0.18 |
| OSBPL2 | -0.02 | -0.02 | -0.11 | 0.03 | 0.14 | -0.03 | 1.7E-01 | 0.10 |
| TP53BP1 | 0.01 | -0.05 | -0.14 | -0.05 | 0.10 | 0.11 | 1.7E-01 | 0.11 |
| SPDL1 | -0.11 | -0.06 | -0.23 | 0.12 | 0.33 | -0.13 | 1.7E-01 | 0.24 |
| LAGE3 | -0.20 | -0.03 | 0.01 | 0.13 | -0.03 | 0.10 | 1.7E-01 | 0.14 |
| FAM219A | 0.13 | 0.30 | 0.11 | -0.21 | -0.69 | 0.16 | 1.7E-01 | -0.43 |
| RPUSD4 | 0.02 | 0.04 | 0.02 | -0.09 | -0.01 | 0.02 | 1.7E-01 | -0.05 |
| GTF3C4 | -0.02 | -0.04 | -0.01 | -0.02 | 0.05 | 0.04 | 1.7E-01 | 0.04 |
| BROX | 0.05 | 0.10 | 0.05 | 0.04 | -0.03 | -0.23 | 1.7E-01 | -0.14 |
| DTX3L | -0.22 | -0.13 | 0.07 | 0.18 | -0.01 | 0.07 | 1.7E-01 | 0.18 |
| SF3B1 | 0.05 | -0.02 | 0.02 | -0.01 | -0.03 | -0.01 | 1.7E-01 | -0.04 |
| CNPY2 | -0.03 | 0.22 | 0.05 | 0.05 | -0.15 | -0.17 | 1.7E-01 | -0.17 |
| HS6ST2 | -0.03 | 0.28 | 0.14 | 0.14 | -0.30 | -0.35 | 1.7E-01 | -0.30 |
| SLC2A8 | -0.16 | 0.08 | 0.48 | -0.22 | -0.16 | -0.15 | 1.7E-01 | -0.31 |
| SPECC1 | 0.01 | 0.06 | 0.03 | -0.06 | -0.09 | 0.04 | 1.7E-01 | -0.07 |
| VMA21 | -0.16 | -0.41 | 0.06 | 0.03 | 0.01 | 0.36 | 1.7E-01 | 0.30 |
| SLC9A7 | -0.09 | 0.08 | 0.22 | -0.09 | -0.09 | -0.06 | 1.7E-01 | -0.15 |
| SCARB2 | -0.18 | -0.11 | -0.01 | 0.14 | -0.10 | 0.21 | 1.7E-01 | 0.18 |
| TSFM | 0.00 | -0.02 | -0.24 | 0.08 | 0.17 | -0.03 | 1.7E-01 | 0.16 |
| GOLGA5 | 0.16 | 0.15 | -0.11 | -0.11 | -0.07 | -0.06 | 1.7E-01 | -0.15 |
| ACLY | 0.09 | 0.05 | -0.02 | -0.01 | -0.01 | -0.12 | 1.7E-01 | -0.08 |
| USP39 | 0.07 | 0.06 | -0.03 | -0.01 | -0.02 | -0.06 | 1.7E-01 | -0.06 |
| SHANK2 | 0.03 | -0.02 | 0.12 | -0.03 | 0.00 | -0.10 | 1.7E-01 | -0.08 |
| PGD | 0.13 | 0.14 | -0.09 | -0.05 | -0.04 | -0.12 | 1.7E-01 | -0.13 |
| THOC7 | 0.10 | 0.09 | 0.05 | 0.06 | -0.05 | -0.29 | 1.7E-01 | -0.17 |
| FRA10AC1 | -0.02 | 0.06 | 0.08 | 0.04 | -0.07 | -0.12 | 1.7E-01 | -0.09 |
| LMNB1 | 0.01 | -0.04 | -0.05 | 0.00 | 0.01 | 0.07 | 1.7E-01 | 0.05 |
| NSMCE1 | -0.04 | -0.07 | -0.02 | 0.02 | -0.03 | 0.14 | 1.7E-01 | 0.09 |
| CMTM4 | 0.03 | -0.13 | -0.06 | 0.13 | 0.03 | -0.01 | 1.7E-01 | 0.10 |
| TIMM29 | -0.06 | -0.05 | 0.02 | 0.01 | 0.00 | 0.08 | 1.7E-01 | 0.06 |
| DDX42 | 0.06 | 0.08 | 0.11 | 0.07 | -0.06 | -0.29 | 1.7E-01 | -0.18 |
| FPGS | -0.39 | -0.21 | 0.14 | 0.01 | 0.19 | 0.16 | 1.7E-01 | 0.28 |
| DYNLL2 | 0.05 | 0.02 | 0.10 | -0.17 | 0.07 | -0.09 | 1.7E-01 | -0.12 |
| SPATS2L | -0.27 | -0.20 | -0.30 | 0.49 | -0.34 | 0.37 | 1.7E-01 | 0.43 |
| NCOA6 | -0.28 | -0.40 | -0.21 | 0.33 | -0.36 | 0.60 | 1.7E-01 | 0.49 |
| MEIS2 | -0.28 | -0.15 | 0.08 | 0.02 | 0.02 | 0.25 | 1.7E-01 | 0.21 |
| PRKRIP1 | 0.10 | 0.03 | -0.03 | -0.04 | -0.06 | 0.00 | 1.7E-01 | -0.07 |
| CTBP2 | 0.01 | -0.08 | -0.15 | 0.19 | 0.06 | -0.05 | 1.7E-01 | 0.14 |

| | | | | | | | | |
|---|---|---|---|---|---|---|---|---|
| **CNEP1R1** | 0.09 | 0.10 | -0.03 | -0.14 | 0.04 | -0.07 | 1.7E-01 | -0.11 |
| **CSNK2A1** | -0.12 | -0.11 | 0.07 | 0.07 | 0.04 | 0.04 | 1.7E-01 | 0.10 |
| **COQ7** | 0.06 | 0.00 | 0.04 | -0.11 | 0.03 | -0.03 | 1.7E-01 | -0.07 |
| **STX12** | -0.07 | 0.04 | 0.30 | -0.18 | -0.09 | -0.05 | 1.7E-01 | -0.19 |
| **LZTR1** | -0.05 | 0.20 | 0.06 | -0.03 | -0.04 | -0.16 | 1.7E-01 | -0.14 |
| **SERBP1** | -0.05 | -0.05 | -0.04 | 0.15 | 0.01 | -0.03 | 1.7E-01 | 0.09 |
| **LAMTOR2** | 0.00 | 0.02 | 0.12 | -0.12 | 0.03 | -0.07 | 1.7E-01 | -0.10 |
| **NCAPD3** | -0.56 | -0.60 | 0.27 | 0.10 | 0.10 | 0.39 | 1.7E-01 | 0.50 |
| **SLF1** | -0.19 | -0.12 | -0.01 | 0.01 | -0.04 | 0.30 | 1.7E-01 | 0.20 |
| **PIGS** | 0.02 | -0.05 | -0.04 | 0.07 | 0.00 | 0.00 | 1.7E-01 | 0.05 |
| **GRK6** | -0.01 | 0.08 | 0.01 | -0.04 | 0.01 | -0.07 | 1.7E-01 | -0.06 |
| **ASCC2** | -0.08 | -0.07 | 0.03 | 0.09 | 0.00 | 0.02 | 1.7E-01 | 0.08 |
| **DEK** | 0.04 | 0.06 | -0.02 | -0.06 | 0.02 | -0.05 | 1.7E-01 | -0.06 |
| **PIGQ** | 0.03 | 0.00 | 0.10 | -0.07 | -0.10 | 0.04 | 1.7E-01 | -0.09 |
| **SLC9A2** | -0.01 | 0.02 | 0.14 | -0.13 | -0.01 | -0.01 | 1.7E-01 | -0.10 |
| **EIF3L** | 0.06 | 0.01 | 0.08 | 0.03 | -0.02 | -0.16 | 1.7E-01 | -0.10 |
| **PRICKLE3** | 0.06 | 0.02 | 0.16 | -0.26 | -0.10 | 0.08 | 1.7E-01 | -0.17 |
| **SMC1B** | 0.00 | -0.04 | -0.07 | -0.03 | 0.03 | 0.12 | 1.7E-01 | 0.08 |
| **SNRNP35** | -0.06 | -0.05 | -0.13 | 0.19 | -0.10 | 0.12 | 1.7E-01 | 0.15 |
| **GOLGA3** | -0.08 | -0.03 | -0.06 | 0.17 | 0.03 | -0.05 | 1.7E-01 | 0.11 |
| **ZNF114** | 0.14 | 0.15 | 0.07 | 0.09 | -0.09 | -0.46 | 1.7E-01 | -0.27 |
| **NDUFB7** | -0.01 | 0.04 | 0.06 | 0.02 | -0.03 | -0.09 | 1.8E-01 | -0.07 |
| **MYL12B** | 0.03 | -0.03 | 0.12 | -0.03 | -0.09 | -0.01 | 1.8E-01 | -0.09 |
| **ZNF616** | 0.11 | 0.06 | 0.07 | 0.06 | -0.05 | -0.28 | 1.8E-01 | -0.17 |
| **RBM45** | 0.02 | -0.02 | 0.12 | -0.07 | -0.08 | 0.02 | 1.8E-01 | -0.08 |
| **AGAP1** | -0.08 | 0.10 | 0.16 | -0.13 | -0.06 | -0.02 | 1.8E-01 | -0.13 |
| **MFSD14A** | -0.05 | 0.15 | 0.10 | -0.16 | 0.05 | -0.13 | 1.8E-01 | -0.15 |
| **HPCAL1** | -0.11 | -0.04 | -0.01 | 0.08 | -0.06 | 0.12 | 1.8E-01 | 0.10 |
| **HACD2** | -0.08 | -0.12 | 0.04 | 0.12 | -0.02 | 0.04 | 1.8E-01 | 0.10 |
| **CUL1** | 0.10 | 0.02 | -0.01 | -0.10 | 0.00 | -0.01 | 1.8E-01 | -0.07 |
| **TMEM179B** | -0.02 | -0.01 | 0.38 | -0.32 | -0.07 | -0.05 | 1.8E-01 | -0.26 |
| **MTF1** | -0.12 | -0.23 | -0.17 | 0.26 | -0.23 | 0.35 | 1.8E-01 | 0.30 |
| **USE1** | 0.03 | -0.03 | 0.23 | 0.04 | -0.14 | -0.17 | 1.8E-01 | -0.16 |
| **SYNM** | 0.43 | -0.21 | 0.39 | -0.77 | -0.16 | 0.00 | 1.8E-01 | -0.51 |
| **CDKN2AIP** | -0.03 | 0.05 | 0.04 | -0.01 | -0.01 | -0.05 | 1.8E-01 | -0.04 |
| **TRAM2** | -0.12 | -0.11 | 0.02 | 0.03 | -0.03 | 0.19 | 1.8E-01 | 0.13 |
| **FGB** | -0.20 | -0.11 | -0.22 | -0.21 | 0.17 | 0.44 | 1.8E-01 | 0.31 |
| **MRPL57** | -0.02 | -0.09 | -0.12 | 0.06 | -0.07 | 0.21 | 1.8E-01 | 0.14 |
| **PCMT1** | -0.02 | -0.06 | -0.20 | -0.09 | 0.20 | 0.13 | 1.8E-01 | 0.17 |
| **RAB22A** | -0.17 | -0.06 | 0.01 | 0.01 | -0.01 | 0.20 | 1.8E-01 | 0.14 |
| **ARPC1A** | 0.05 | 0.04 | 0.05 | 0.06 | -0.08 | -0.13 | 1.8E-01 | -0.09 |
| **ARHGAP10** | 0.08 | -0.02 | 0.14 | 0.06 | -0.09 | -0.19 | 1.8E-01 | -0.14 |
| **PRPSAP1** | 0.05 | 0.03 | 0.05 | -0.05 | -0.14 | 0.05 | 1.8E-01 | -0.09 |
| **SLC25A15** | 0.03 | 0.11 | -0.03 | -0.03 | 0.00 | -0.08 | 1.8E-01 | -0.08 |
| **COPS4** | 0.00 | -0.06 | -0.28 | 0.24 | 0.12 | -0.08 | 1.8E-01 | 0.20 |
| **SNU13** | -0.15 | -0.21 | 0.04 | -0.04 | 0.04 | 0.26 | 1.8E-01 | 0.19 |
| **ZNF593** | 0.21 | 0.02 | 0.02 | 0.08 | -0.15 | -0.21 | 1.8E-01 | -0.17 |
| **CALR** | 0.13 | 0.15 | -0.08 | -0.03 | -0.04 | -0.15 | 1.8E-01 | -0.14 |

| | | | | | | | | |
|---|---|---|---|---|---|---|---|---|
| **MTHFD1L** | 0.01 | 0.01 | -0.27 | 0.03 | 0.09 | 0.10 | 1.8E-01 | 0.16 |
| **ERCC4** | -0.16 | -0.07 | 0.04 | 0.09 | -0.03 | 0.11 | 1.8E-01 | 0.12 |
| **SNRPD3** | 0.04 | 0.03 | 0.05 | 0.06 | -0.07 | -0.11 | 1.8E-01 | -0.08 |
| **ZNF629** | -0.11 | 0.00 | -0.01 | -0.02 | 0.03 | 0.11 | 1.8E-01 | 0.08 |
| **AGPAT3** | -0.01 | 0.05 | 0.05 | 0.03 | -0.06 | -0.06 | 1.8E-01 | -0.06 |
| **GATM** | 0.24 | 0.40 | -0.15 | -0.55 | -0.17 | 0.04 | 1.8E-01 | -0.39 |
| **MLYCD** | 0.10 | 0.04 | 0.00 | -0.15 | -0.01 | 0.01 | 1.8E-01 | -0.10 |
| **VRK3** | 0.10 | 0.08 | -0.05 | 0.01 | -0.09 | -0.06 | 1.8E-01 | -0.09 |
| **SPATS2** | 0.01 | 0.01 | 0.22 | 0.02 | -0.05 | -0.24 | 1.8E-01 | -0.17 |
| **HOMER3** | -0.24 | -0.09 | 0.05 | -0.02 | 0.04 | 0.22 | 1.8E-01 | 0.18 |
| **TMEM135** | 0.00 | 0.16 | -0.01 | -0.01 | -0.11 | -0.04 | 1.8E-01 | -0.10 |
| **SHROOM3** | -0.01 | 0.02 | 0.20 | -0.22 | -0.03 | 0.00 | 1.8E-01 | -0.15 |
| **CLPX** | 0.21 | 0.04 | -0.06 | -0.09 | -0.03 | -0.10 | 1.8E-01 | -0.14 |
| **GRHL3** | 0.00 | -0.05 | 0.38 | -0.07 | -0.28 | -0.06 | 1.8E-01 | -0.25 |
| **ZNF639** | 0.04 | -0.09 | -0.07 | 0.01 | 0.09 | 0.02 | 1.8E-01 | 0.08 |
| **TEFM** | -0.03 | -0.05 | 0.02 | 0.04 | -0.01 | 0.03 | 1.8E-01 | 0.04 |
| **SF3B4** | 0.05 | 0.04 | 0.01 | -0.12 | 0.03 | -0.02 | 1.8E-01 | -0.07 |
| **GALNT1** | -0.18 | -0.18 | 0.10 | 0.11 | 0.09 | 0.02 | 1.8E-01 | 0.16 |
| **PLEKHF2** | -0.04 | -0.09 | -0.05 | 0.16 | -0.08 | 0.09 | 1.8E-01 | 0.12 |
| **PPP4R3A** | 0.05 | -0.06 | -0.15 | 0.02 | 0.02 | 0.12 | 1.8E-01 | 0.11 |
| **INTS11** | 0.01 | -0.11 | -0.02 | 0.04 | -0.02 | 0.09 | 1.8E-01 | 0.08 |
| **SERF2** | 0.06 | 0.09 | 0.07 | -0.27 | 0.05 | -0.04 | 1.8E-01 | -0.15 |
| **FGFR4** | 0.00 | 0.09 | 0.10 | 0.07 | -0.09 | -0.19 | 1.8E-01 | -0.13 |
| **TMEM59** | -0.08 | 0.09 | 0.22 | -0.08 | 0.01 | -0.21 | 1.8E-01 | -0.17 |
| **SYVN1** | -0.01 | -0.01 | 0.21 | -0.08 | 0.01 | -0.16 | 1.8E-01 | -0.14 |
| **ENAM** | 0.01 | -0.11 | 0.00 | 0.00 | 0.05 | 0.05 | 1.8E-01 | 0.07 |
| **WDR82** | 0.02 | -0.01 | 0.13 | -0.08 | -0.11 | 0.04 | 1.8E-01 | -0.10 |
| **SETD3** | 0.02 | 0.10 | 0.07 | -0.24 | -0.01 | 0.03 | 1.8E-01 | -0.14 |
| **ERH** | 0.04 | 0.02 | 0.10 | 0.01 | 0.00 | -0.18 | 1.8E-01 | -0.11 |
| **GRB2** | -0.03 | 0.02 | 0.15 | -0.01 | -0.13 | -0.03 | 1.8E-01 | -0.10 |
| **SRSF11** | -0.07 | 0.03 | 0.32 | -0.23 | -0.04 | -0.07 | 1.8E-01 | -0.21 |
| **REV3L** | 0.02 | 0.03 | 0.04 | -0.10 | 0.00 | 0.01 | 1.8E-01 | -0.06 |
| **RPS21** | 0.02 | 0.03 | 0.23 | 0.06 | -0.09 | -0.31 | 1.8E-01 | -0.21 |
| **RPL31** | 0.04 | 0.00 | 0.12 | 0.01 | 0.00 | -0.19 | 1.8E-01 | -0.12 |
| **AGFG1** | -0.05 | -0.01 | -0.14 | -0.01 | 0.20 | -0.01 | 1.8E-01 | 0.13 |
| **GPR39** | -0.10 | 0.11 | 0.19 | -0.05 | -0.11 | -0.06 | 1.8E-01 | -0.14 |
| **FAM83D** | -0.10 | 0.00 | -0.12 | 0.20 | 0.05 | -0.06 | 1.8E-01 | 0.14 |
| **PSMA6** | 0.08 | -0.13 | -0.18 | -0.02 | 0.10 | 0.13 | 1.8E-01 | 0.15 |
| **CCDC134** | 0.04 | 0.03 | 0.05 | -0.06 | 0.05 | -0.13 | 1.8E-01 | -0.09 |
| **LIG3** | -0.04 | 0.00 | -0.04 | 0.07 | 0.03 | -0.03 | 1.8E-01 | 0.05 |
| **SFT2D3** | 0.26 | 0.10 | -0.12 | -0.11 | -0.10 | -0.08 | 1.8E-01 | -0.18 |
| **IMPDH1** | -0.33 | -0.21 | 0.14 | 0.00 | 0.13 | 0.20 | 1.8E-01 | 0.24 |
| **COPS5** | 0.14 | 0.08 | 0.02 | -0.22 | 0.11 | -0.16 | 1.8E-01 | -0.17 |
| **LENG1** | -0.09 | 0.07 | 0.23 | -0.15 | -0.04 | -0.07 | 1.8E-01 | -0.16 |
| **SMNDC1** | 0.12 | 0.03 | -0.03 | -0.10 | -0.01 | -0.02 | 1.8E-01 | -0.08 |
| **BABAM1** | 0.03 | -0.06 | -0.12 | -0.04 | 0.06 | 0.11 | 1.8E-01 | 0.10 |
| **SNAP91** | -1.18 | -1.74 | 0.40 | -0.21 | 0.21 | 0.94 | 1.9E-01 | 1.15 |
| **ARHGAP35** | -0.06 | -0.08 | -0.20 | -0.08 | 0.32 | 0.03 | 1.9E-01 | 0.20 |

| | | | | | | | | |
|---|---|---|---|---|---|---|---|---|
| LSM7 | -0.02 | -0.16 | -0.13 | 0.05 | -0.09 | 0.31 | 1.9E-01 | 0.20 |
| CCDC138 | 0.24 | 0.15 | -0.09 | 0.08 | -0.18 | -0.27 | 1.9E-01 | -0.23 |
| RING1 | 0.04 | -0.02 | 0.02 | 0.00 | -0.02 | -0.02 | 1.9E-01 | -0.03 |
| UBE2Z | 0.11 | 0.00 | 0.05 | -0.17 | 0.05 | -0.06 | 1.9E-01 | -0.11 |
| MYOM1 | -0.12 | 0.17 | 0.15 | -0.09 | -0.07 | -0.08 | 1.9E-01 | -0.15 |
| HNRNPA2B1 | -0.04 | -0.12 | 0.01 | 0.11 | -0.05 | 0.08 | 1.9E-01 | 0.10 |
| CKAP4 | -0.01 | 0.03 | 0.12 | 0.04 | -0.07 | -0.13 | 1.9E-01 | -0.10 |
| PHF1 | -0.11 | 0.01 | -0.11 | -0.08 | 0.13 | 0.13 | 1.9E-01 | 0.13 |
| PTK7 | -0.13 | 0.09 | 0.36 | -0.21 | -0.18 | -0.01 | 1.9E-01 | -0.24 |
| DNAJC11 | -0.02 | 0.02 | -0.07 | 0.03 | 0.00 | 0.05 | 1.9E-01 | 0.05 |
| P2RX4 | 0.00 | -0.03 | -0.16 | 0.17 | 0.04 | -0.03 | 1.9E-01 | 0.12 |
| MARCHF5 | 0.11 | -0.02 | 0.10 | 0.04 | -0.04 | -0.21 | 1.9E-01 | -0.13 |
| MT-ND3 | -0.10 | -0.04 | -0.25 | 0.29 | -0.16 | 0.19 | 1.9E-01 | 0.24 |
| PPP1R9B | -0.08 | 0.01 | -0.01 | 0.01 | 0.01 | 0.05 | 1.9E-01 | 0.05 |
| RFX1 | -0.10 | -0.13 | 0.06 | 0.03 | 0.01 | 0.11 | 1.9E-01 | 0.10 |
| SRSF2 | 0.04 | 0.02 | 0.15 | 0.03 | -0.02 | -0.25 | 1.9E-01 | -0.15 |
| CCDC43 | 0.11 | 0.03 | 0.17 | -0.39 | -0.04 | 0.06 | 1.9E-01 | -0.23 |
| THOC1 | 0.12 | 0.11 | -0.04 | 0.04 | -0.06 | -0.20 | 1.9E-01 | -0.14 |
| CRELD1 | 0.00 | -0.01 | 0.09 | -0.06 | -0.02 | -0.01 | 1.9E-01 | -0.06 |
| NDUFS3 | 0.00 | -0.05 | -0.06 | 0.11 | 0.00 | -0.01 | 1.9E-01 | 0.07 |
| MIB1 | -0.07 | 0.06 | 0.19 | -0.01 | -0.13 | -0.07 | 1.9E-01 | -0.13 |
| ATXN7 | -0.06 | -0.01 | -0.04 | 0.12 | -0.01 | 0.00 | 1.9E-01 | 0.07 |
| UBIAD1 | 0.01 | 0.09 | 0.10 | 0.08 | -0.08 | -0.22 | 1.9E-01 | -0.14 |
| PURA | 0.03 | -0.01 | 0.12 | -0.01 | 0.01 | -0.14 | 1.9E-01 | -0.09 |
| COPS7B | -0.04 | 0.05 | 0.16 | 0.04 | -0.12 | -0.11 | 1.9E-01 | -0.12 |
| AGO2 | -0.07 | -0.01 | 0.02 | 0.01 | 0.02 | 0.04 | 1.9E-01 | 0.04 |
| KATNB1 | -0.10 | 0.06 | -0.13 | 0.11 | 0.07 | -0.02 | 1.9E-01 | 0.11 |
| BAIAP2L1 | -0.25 | -0.09 | 0.05 | 0.18 | -0.07 | 0.14 | 1.9E-01 | 0.18 |
| ILF3 | 0.04 | 0.00 | 0.00 | -0.04 | -0.01 | 0.00 | 1.9E-01 | -0.03 |
| SUV39H1 | 0.02 | -0.01 | 0.07 | -0.07 | -0.04 | 0.02 | 1.9E-01 | -0.06 |
| HMBS | -0.26 | -0.08 | -0.01 | -0.05 | 0.02 | 0.32 | 1.9E-01 | 0.21 |
| DSG2 | -0.02 | 0.01 | -0.02 | 0.01 | 0.00 | 0.02 | 1.9E-01 | 0.02 |
| ZNF638 | 0.00 | 0.00 | -0.07 | 0.06 | 0.00 | 0.01 | 1.9E-01 | 0.05 |
| NIPAL3 | -0.12 | 0.09 | 0.36 | -0.22 | 0.00 | -0.19 | 1.9E-01 | -0.25 |
| CDC16 | -0.02 | -0.02 | -0.08 | -0.02 | 0.01 | 0.12 | 1.9E-01 | 0.07 |
| NPTN | -0.04 | 0.04 | 0.15 | -0.10 | -0.11 | 0.03 | 1.9E-01 | -0.11 |
| IQSEC2 | -0.12 | -0.29 | 0.07 | 0.17 | -0.07 | 0.18 | 1.9E-01 | 0.21 |
| CYCS | 0.25 | 0.21 | -0.17 | -0.12 | -0.07 | -0.16 | 1.9E-01 | -0.21 |
| LMO7 | -0.03 | 0.08 | 0.08 | -0.05 | -0.12 | 0.03 | 1.9E-01 | -0.09 |
| SPDEF | 0.00 | 0.03 | 0.08 | 0.02 | -0.12 | -0.01 | 1.9E-01 | -0.07 |
| RFC4 | -0.11 | -0.07 | 0.04 | -0.02 | 0.10 | 0.05 | 1.9E-01 | 0.09 |
| AXIN1 | 0.09 | -0.03 | 0.11 | 0.03 | -0.18 | -0.05 | 1.9E-01 | -0.12 |
| SMAD5 | 0.01 | 0.09 | -0.02 | -0.04 | -0.04 | -0.01 | 1.9E-01 | -0.05 |
| TUFT1 | -0.40 | -0.20 | -0.30 | 0.24 | -0.37 | 0.68 | 1.9E-01 | 0.48 |
| HOMEZ | -0.13 | -0.14 | 0.06 | 0.16 | -0.01 | 0.04 | 1.9E-01 | 0.13 |
| CDC73 | 0.00 | -0.04 | -0.05 | 0.06 | 0.05 | -0.03 | 1.9E-01 | 0.05 |
| POGLUT1 | -0.05 | 0.15 | 0.03 | -0.09 | -0.04 | -0.02 | 1.9E-01 | -0.10 |
| B3GALT6 | -0.12 | -0.08 | 0.03 | 0.15 | -0.01 | 0.00 | 1.9E-01 | 0.11 |

| | | | | | | | | |
|---|---|---|---|---|---|---|---|---|
| **BST2** | -0.02 | 0.02 | 0.11 | -0.09 | -0.02 | 0.00 | 1.9E-01 | -0.07 |
| **DIABLO** | 0.09 | 0.09 | -0.04 | -0.11 | -0.05 | 0.02 | 1.9E-01 | -0.09 |
| **MRPS7** | 0.16 | 0.06 | -0.05 | -0.12 | 0.02 | -0.08 | 1.9E-01 | -0.12 |
| **POLR2E** | -0.02 | -0.09 | -0.07 | -0.03 | -0.01 | 0.20 | 1.9E-01 | 0.12 |
| **IDH3A** | 0.02 | -0.03 | -0.16 | -0.03 | 0.11 | 0.08 | 1.9E-01 | 0.11 |
| **REEP6** | 0.15 | 0.16 | -0.01 | -0.36 | -0.07 | 0.07 | 1.9E-01 | -0.22 |
| **UBP1** | -0.12 | -0.16 | 0.04 | 0.17 | -0.08 | 0.12 | 1.9E-01 | 0.15 |
| **TRPV2** | 0.08 | 0.01 | 0.21 | -0.36 | -0.08 | 0.08 | 1.9E-01 | -0.22 |
| **ESYT2** | -0.07 | -0.08 | 0.01 | 0.10 | -0.05 | 0.07 | 1.9E-01 | 0.08 |
| **SEMA3C** | 0.06 | 0.09 | -0.01 | -0.07 | 0.05 | -0.13 | 1.9E-01 | -0.09 |
| **UPF1** | -0.06 | -0.03 | 0.00 | 0.06 | 0.06 | -0.04 | 1.9E-01 | 0.06 |
| **PDK2** | -0.12 | -0.02 | 0.00 | 0.00 | 0.00 | 0.13 | 1.9E-01 | 0.09 |
| **NDUFC1** | 0.03 | -0.02 | -0.20 | 0.10 | 0.04 | 0.03 | 1.9E-01 | 0.12 |
| **NSD3** | 0.08 | 0.03 | -0.01 | -0.10 | -0.03 | 0.02 | 1.9E-01 | -0.07 |
| **POLE** | -0.05 | 0.00 | -0.05 | 0.09 | -0.04 | 0.04 | 1.9E-01 | 0.07 |
| **STX8** | -0.11 | -0.10 | 0.01 | -0.03 | 0.01 | 0.20 | 1.9E-01 | 0.13 |
| **DTX2** | 0.01 | -0.45 | -0.05 | 0.06 | -0.03 | 0.35 | 2.0E-01 | 0.29 |
| **PALS1** | -0.18 | -0.06 | 0.02 | -0.02 | 0.02 | 0.19 | 2.0E-01 | 0.14 |
| **UBXN6** | -0.36 | -0.27 | 0.18 | 0.17 | 0.06 | 0.12 | 2.0E-01 | 0.27 |
| **KDM5A** | 0.03 | -0.06 | -0.04 | -0.01 | 0.03 | 0.05 | 2.0E-01 | 0.05 |
| **ANXA11** | -0.07 | -0.04 | -0.01 | 0.01 | 0.12 | -0.02 | 2.0E-01 | 0.07 |
| **OGFOD3** | -0.06 | 0.05 | 0.31 | -0.19 | 0.07 | -0.25 | 2.0E-01 | -0.23 |
| **RBM42** | -0.03 | -0.63 | 0.04 | 0.28 | -0.02 | 0.19 | 2.0E-01 | 0.36 |
| **GATAD2B** | 0.02 | 0.04 | 0.02 | -0.10 | -0.01 | 0.02 | 2.0E-01 | -0.06 |
| **EIF5A** | 0.10 | 0.11 | -0.07 | -0.03 | -0.02 | -0.11 | 2.0E-01 | -0.10 |
| **AMACR** | -0.17 | -0.17 | 0.07 | 0.08 | -0.05 | 0.20 | 2.0E-01 | 0.17 |
| **LNX1** | 0.06 | 0.01 | 0.15 | -0.27 | 0.00 | 0.01 | 2.0E-01 | -0.16 |
| **STX16** | -0.02 | 0.14 | 0.09 | 0.03 | -0.01 | -0.26 | 2.0E-01 | -0.16 |
| **BRCC3** | -0.04 | -0.22 | 0.03 | 0.14 | 0.12 | -0.05 | 2.0E-01 | 0.15 |
| **SRCIN1** | -0.05 | 0.02 | -0.03 | 0.00 | 0.01 | 0.04 | 2.0E-01 | 0.04 |
| **LIN7C** | -0.25 | -0.12 | 0.06 | -0.01 | 0.01 | 0.26 | 2.0E-01 | 0.19 |
| **OLA1** | 0.16 | 0.08 | -0.07 | -0.02 | -0.02 | -0.16 | 2.0E-01 | -0.12 |
| **ZNF131** | -0.04 | 0.03 | -0.45 | 0.27 | -0.03 | 0.11 | 2.0E-01 | 0.27 |
| **RBBP8NL** | -0.07 | 0.00 | -0.43 | -0.13 | 0.35 | 0.17 | 2.0E-01 | 0.30 |
| **NMD3** | -0.03 | 0.02 | 0.25 | -0.05 | 0.00 | -0.22 | 2.0E-01 | -0.17 |
| **BRI3BP** | 0.10 | 0.02 | -0.03 | -0.03 | -0.02 | -0.05 | 2.0E-01 | -0.06 |
| **TMEM62** | -0.08 | 0.25 | 0.34 | -0.12 | -0.71 | 0.09 | 2.0E-01 | -0.42 |
| **TCHP** | 0.08 | 0.28 | -0.10 | -0.23 | -0.01 | -0.08 | 2.0E-01 | -0.19 |
| **PHF21A** | -0.09 | -0.10 | 0.04 | 0.01 | -0.01 | 0.13 | 2.0E-01 | 0.10 |
| **RPS18** | -0.11 | -0.09 | 0.02 | 0.17 | 0.02 | -0.03 | 2.0E-01 | 0.11 |
| **WDR6** | 0.06 | -0.04 | 0.08 | -0.06 | -0.06 | 0.02 | 2.0E-01 | -0.07 |
| **STK35** | -0.12 | 0.00 | -0.07 | -0.07 | 0.06 | 0.18 | 2.0E-01 | 0.12 |
| **INPP5A** | -0.05 | 0.06 | 0.08 | 0.00 | -0.06 | -0.05 | 2.0E-01 | -0.07 |
| **COX8A** | -0.11 | -0.15 | 0.02 | 0.21 | -0.08 | 0.08 | 2.0E-01 | 0.15 |
| **PREB** | -0.01 | 0.01 | 0.08 | -0.08 | -0.03 | 0.02 | 2.0E-01 | -0.06 |
| **PKP4** | -0.15 | -0.06 | 0.04 | 0.03 | -0.01 | 0.13 | 2.0E-01 | 0.11 |
| **PLSCR1** | -0.03 | 0.02 | 0.12 | -0.02 | -0.05 | -0.04 | 2.0E-01 | -0.07 |
| **DARS1** | -0.07 | -0.03 | -0.02 | 0.07 | 0.10 | -0.06 | 2.0E-01 | 0.08 |

| | | | | | | | | |
|---|---|---|---|---|---|---|---|---|
| FAM50B | 0.08 | -0.02 | 0.07 | 0.05 | -0.13 | -0.06 | 2.0E-01 | -0.09 |
| GLA | 0.02 | -0.06 | -0.23 | -0.02 | 0.23 | 0.02 | 2.0E-01 | 0.17 |
| NRF1 | 0.05 | -0.17 | -0.10 | -0.03 | 0.05 | 0.18 | 2.0E-01 | 0.14 |
| TAF11 | -0.01 | 0.00 | -0.05 | 0.01 | -0.01 | 0.07 | 2.0E-01 | 0.04 |
| DCTN4 | -0.03 | 0.09 | 0.04 | -0.08 | 0.01 | -0.04 | 2.0E-01 | -0.07 |
| EEF2 | 0.18 | 0.19 | -0.12 | -0.04 | -0.03 | -0.23 | 2.0E-01 | -0.18 |
| CD9 | -0.04 | -0.21 | -0.15 | 0.18 | -0.18 | 0.32 | 2.0E-01 | 0.24 |
| FAM120B | 0.05 | 0.03 | 0.11 | 0.07 | -0.07 | -0.22 | 2.0E-01 | -0.13 |
| TXNDC15 | -0.01 | 0.45 | 0.17 | -0.80 | 0.29 | -0.49 | 2.0E-01 | -0.54 |
| PNKP | -0.02 | 0.05 | 0.07 | 0.03 | -0.05 | -0.09 | 2.0E-01 | -0.07 |
| MCOLN1 | 0.24 | 0.06 | -0.01 | 0.05 | -0.37 | -0.05 | 2.0E-01 | -0.22 |
| YTHDC2 | 0.02 | 0.02 | 0.03 | 0.00 | 0.01 | -0.08 | 2.0E-01 | -0.05 |
| DDB1 | -0.02 | -0.03 | -0.04 | -0.02 | 0.00 | 0.11 | 2.0E-01 | 0.06 |
| ABCC6 | -0.05 | 0.08 | 0.12 | -0.12 | 0.02 | -0.06 | 2.0E-01 | -0.10 |
| TIAM1 | -0.13 | 0.24 | 0.16 | -0.14 | 0.01 | -0.19 | 2.0E-01 | -0.19 |
| SMURF2 | -0.06 | 0.24 | 0.22 | 0.06 | -0.06 | -0.52 | 2.0E-01 | -0.31 |
| HNRNPH3 | -0.02 | -0.03 | 0.01 | -0.01 | 0.02 | 0.03 | 2.0E-01 | 0.02 |
| BCKDHB | 0.14 | 0.18 | -0.10 | -0.03 | -0.03 | -0.19 | 2.0E-01 | -0.16 |
| EIF3A | -0.02 | 0.01 | 0.05 | -0.02 | 0.00 | -0.03 | 2.0E-01 | -0.03 |
| FAM241B | -0.22 | 0.28 | 0.38 | -0.04 | -0.10 | -0.47 | 2.0E-01 | -0.35 |
| SP6 | -0.01 | 0.12 | 0.04 | -0.09 | 0.06 | -0.13 | 2.1E-01 | -0.10 |
| MYBBP1A | 0.00 | 0.02 | 0.04 | 0.02 | -0.02 | -0.06 | 2.1E-01 | -0.04 |
| PMS1 | -0.19 | -0.15 | 0.07 | -0.07 | 0.10 | 0.20 | 2.1E-01 | 0.17 |
| PCBP2 | 0.04 | 0.00 | 0.00 | -0.04 | 0.02 | -0.03 | 2.1E-01 | -0.03 |
| NUMA1 | -0.04 | -0.05 | -0.08 | -0.04 | 0.00 | 0.20 | 2.1E-01 | 0.11 |
| PUSL1 | 0.01 | -0.04 | -0.12 | -0.05 | 0.13 | 0.06 | 2.1E-01 | 0.10 |
| ACTB | -0.11 | -0.04 | 0.01 | 0.13 | 0.02 | -0.03 | 2.1E-01 | 0.09 |
| NSUN3 | 0.04 | -0.03 | 0.25 | -0.30 | 0.00 | -0.02 | 2.1E-01 | -0.19 |
| C20orf96 | 0.16 | 0.09 | 0.02 | -0.15 | 0.12 | -0.30 | 2.1E-01 | -0.19 |
| PDPR | 0.15 | -0.01 | 0.10 | 0.08 | -0.08 | -0.28 | 2.1E-01 | -0.17 |
| EPHA5 | -0.08 | -0.07 | -0.12 | -0.08 | 0.02 | 0.29 | 2.1E-01 | 0.17 |
| NRDE2 | 0.04 | -0.01 | 0.12 | 0.05 | -0.06 | -0.17 | 2.1E-01 | -0.11 |
| BCL7A | 0.12 | 0.06 | -0.03 | -0.17 | -0.01 | 0.00 | 2.1E-01 | -0.11 |
| PLOD1 | 0.10 | 0.00 | 0.02 | 0.05 | -0.13 | -0.06 | 2.1E-01 | -0.09 |
| COX20 | -0.10 | 0.02 | 0.35 | -0.15 | -0.06 | -0.13 | 2.1E-01 | -0.21 |
| DDX5 | 0.04 | -0.03 | 0.09 | -0.07 | -0.06 | 0.03 | 2.1E-01 | -0.07 |
| ZSCAN29 | -0.01 | 0.06 | 0.01 | 0.02 | -0.03 | -0.05 | 2.1E-01 | -0.04 |
| RABL6 | -0.04 | -0.08 | -0.02 | 0.01 | -0.04 | 0.14 | 2.1E-01 | 0.09 |
| ARMCX3 | 0.03 | 0.04 | 0.04 | 0.02 | 0.00 | -0.14 | 2.1E-01 | -0.07 |
| GTF3C1 | 0.00 | 0.00 | -0.06 | 0.06 | -0.01 | 0.01 | 2.1E-01 | 0.04 |
| NOL9 | 0.00 | 0.01 | 0.03 | 0.02 | -0.01 | -0.05 | 2.1E-01 | -0.03 |
| TEX10 | -0.01 | -0.04 | -0.01 | -0.03 | 0.05 | 0.05 | 2.1E-01 | 0.04 |
| OXSM | 0.05 | -0.03 | -0.23 | 0.11 | 0.00 | 0.09 | 2.1E-01 | 0.13 |
| RAB11A | -0.07 | 0.04 | 0.23 | -0.16 | -0.04 | -0.04 | 2.1E-01 | -0.14 |
| OIP5 | 0.09 | 0.01 | 0.25 | -0.47 | 0.07 | -0.05 | 2.1E-01 | -0.27 |
| UQCRQ | -0.01 | 0.01 | -0.08 | 0.08 | -0.01 | 0.01 | 2.1E-01 | 0.06 |
| ABL1 | 0.07 | -0.19 | -0.36 | -0.03 | 0.40 | -0.01 | 2.1E-01 | 0.28 |
| ATP9B | -0.09 | 0.06 | 0.22 | -0.12 | -0.03 | -0.07 | 2.1E-01 | -0.14 |

| | | | | | | | | |
|---|---|---|---|---|---|---|---|---|
| AK3 | -0.05 | 0.00 | 0.30 | -0.19 | -0.01 | -0.09 | 2.1E-01 | -0.18 |
| SNAP29 | 0.00 | 0.01 | 0.17 | 0.06 | -0.09 | -0.18 | 2.1E-01 | -0.13 |
| MKRN2 | -0.11 | -0.11 | 0.06 | -0.01 | 0.05 | 0.10 | 2.1E-01 | 0.10 |
| PPP2R5D | -0.08 | -0.09 | -0.03 | 0.17 | 0.12 | -0.11 | 2.1E-01 | 0.13 |
| CBR4 | 0.07 | -0.11 | -0.19 | 0.04 | -0.02 | 0.17 | 2.1E-01 | 0.14 |
| ZBTB41 | 0.06 | 0.08 | -0.01 | 0.03 | -0.02 | -0.15 | 2.1E-01 | -0.09 |
| RANBP9 | -0.02 | -0.16 | -0.08 | 0.26 | 0.04 | -0.08 | 2.1E-01 | 0.16 |
| DUT | 0.24 | 0.09 | -0.11 | -0.03 | -0.14 | -0.10 | 2.1E-01 | -0.16 |
| KMT5B | 0.05 | 0.08 | 0.05 | -0.23 | -0.02 | 0.05 | 2.1E-01 | -0.12 |
| SETDB2 | -0.23 | -0.03 | -0.15 | -0.02 | -0.07 | 0.41 | 2.1E-01 | 0.24 |
| CUL7 | 0.11 | 0.08 | 0.03 | 0.11 | -0.11 | -0.24 | 2.1E-01 | -0.16 |
| PIK3C2B | -0.28 | -0.15 | 0.13 | 0.07 | 0.09 | 0.09 | 2.1E-01 | 0.18 |
| AHCYL1 | 0.11 | 0.04 | 0.08 | 0.00 | 0.04 | -0.30 | 2.1E-01 | -0.16 |
| ANAPC4 | -0.01 | -0.19 | 0.71 | -0.45 | -0.37 | -0.02 | 2.1E-01 | -0.45 |
| TC2N | -0.08 | -0.13 | 0.06 | -0.01 | 0.07 | 0.07 | 2.1E-01 | 0.09 |
| NOP58 | 0.01 | -0.03 | -0.03 | 0.05 | 0.01 | -0.02 | 2.1E-01 | 0.03 |
| NUP205 | 0.10 | 0.07 | 0.10 | 0.06 | -0.01 | -0.37 | 2.1E-01 | -0.20 |
| CCDC127 | -0.08 | -0.10 | 0.00 | 0.16 | -0.09 | 0.10 | 2.1E-01 | 0.12 |
| NFIX | 0.11 | 0.07 | -0.06 | -0.06 | 0.01 | -0.09 | 2.1E-01 | -0.08 |
| ERGIC1 | 0.05 | 0.03 | 0.06 | -0.03 | 0.05 | -0.17 | 2.1E-01 | -0.10 |
| UBR2 | -0.24 | -0.08 | 0.02 | -0.10 | 0.10 | 0.24 | 2.2E-01 | 0.18 |
| SF3A3 | 0.11 | 0.02 | -0.01 | 0.04 | -0.06 | -0.12 | 2.2E-01 | -0.09 |
| AP1AR | -0.06 | 0.09 | 0.14 | 0.04 | -0.13 | -0.10 | 2.2E-01 | -0.12 |
| FKBP11 | -0.08 | 0.04 | 0.31 | -0.19 | -0.19 | 0.04 | 2.2E-01 | -0.21 |
| CARS2 | -0.04 | -0.14 | -0.11 | -0.10 | 0.05 | 0.30 | 2.2E-01 | 0.18 |
| DCUN1D1 | 0.21 | 0.12 | -0.13 | -0.06 | -0.09 | -0.09 | 2.2E-01 | -0.15 |
| TFCP2 | -0.04 | -0.05 | 0.01 | 0.08 | -0.02 | 0.01 | 2.2E-01 | 0.05 |
| SEC13 | 0.02 | 0.07 | 0.02 | 0.02 | 0.00 | -0.13 | 2.2E-01 | -0.08 |
| RHOT2 | -0.01 | 0.03 | 0.08 | 0.04 | -0.07 | -0.07 | 2.2E-01 | -0.07 |
| TMC4 | -0.11 | -0.09 | 0.05 | 0.04 | -0.02 | 0.12 | 2.2E-01 | 0.09 |
| MRPL34 | 0.23 | 0.13 | -0.09 | 0.08 | -0.15 | -0.24 | 2.2E-01 | -0.20 |
| ZC3H8 | 0.03 | 0.08 | 0.07 | -0.23 | 0.02 | 0.02 | 2.2E-01 | -0.12 |
| CDPF1 | -0.23 | -0.12 | 0.05 | 0.07 | -0.07 | 0.25 | 2.2E-01 | 0.18 |
| MIGA1 | -0.06 | 0.06 | 0.15 | -0.03 | 0.00 | -0.13 | 2.2E-01 | -0.11 |
| SERINC5 | -0.07 | 0.04 | 0.26 | -0.11 | -0.18 | 0.01 | 2.2E-01 | -0.17 |
| TWF1 | 0.03 | -0.06 | 0.25 | -0.04 | -0.18 | -0.03 | 2.2E-01 | -0.15 |
| DRG2 | 0.03 | -0.12 | -0.17 | -0.10 | 0.11 | 0.21 | 2.2E-01 | 0.16 |
| EPN2 | -0.22 | 0.00 | 0.01 | 0.17 | 0.00 | 0.02 | 2.2E-01 | 0.13 |
| FAM91A1 | -0.01 | 0.07 | 0.11 | -0.01 | 0.03 | -0.22 | 2.2E-01 | -0.12 |
| CAPG | -0.01 | 0.09 | 0.05 | -0.14 | 0.06 | -0.07 | 2.2E-01 | -0.09 |
| HMGB2 | 0.09 | 0.11 | -0.04 | -0.06 | 0.04 | -0.16 | 2.2E-01 | -0.11 |
| RUNX1 | -0.20 | -0.18 | 0.11 | -0.03 | 0.15 | 0.11 | 2.2E-01 | 0.16 |
| ABCC1 | -0.11 | 0.09 | 0.25 | -0.02 | -0.22 | -0.05 | 2.2E-01 | -0.18 |
| CSTF3 | 0.01 | -0.05 | -0.08 | 0.04 | -0.05 | 0.12 | 2.2E-01 | 0.08 |
| JMY | -0.08 | -0.22 | 0.03 | 0.19 | -0.09 | 0.13 | 2.2E-01 | 0.16 |
| SNX18 | -0.06 | -0.16 | 0.06 | 0.03 | 0.01 | 0.11 | 2.2E-01 | 0.10 |
| TNK2 | -0.07 | 0.01 | 0.30 | -0.07 | -0.05 | -0.17 | 2.2E-01 | -0.17 |
| SH3BP4 | 0.00 | 0.07 | 0.02 | -0.01 | 0.02 | -0.09 | 2.2E-01 | -0.06 |

| | | | | | | | | |
|---|---|---|---|---|---|---|---|---|
| CCDC71 | -0.01 | -0.06 | -0.13 | 0.08 | -0.09 | 0.19 | 2.2E-01 | 0.13 |
| TTC14 | 0.12 | 0.04 | 0.26 | 0.15 | -0.12 | -0.60 | 2.2E-01 | -0.33 |
| ZNF274 | -0.15 | -0.10 | -0.18 | -0.08 | -0.04 | 0.44 | 2.2E-01 | 0.25 |
| DERL2 | -0.20 | 0.17 | 0.33 | -0.18 | -0.12 | -0.09 | 2.2E-01 | -0.23 |
| CNOT11 | -0.04 | -0.02 | -0.16 | 0.07 | 0.20 | -0.08 | 2.2E-01 | 0.14 |
| HNRNPM | -0.05 | 0.01 | -0.01 | 0.03 | 0.03 | -0.01 | 2.2E-01 | 0.03 |
| PEG10 | -0.01 | 0.01 | 0.05 | -0.01 | -0.01 | -0.03 | 2.2E-01 | -0.03 |
| EXOC1 | 0.01 | -0.09 | -0.02 | -0.01 | 0.09 | 0.00 | 2.2E-01 | 0.06 |
| ETHE1 | 0.15 | 0.13 | -0.06 | -0.26 | 0.03 | -0.03 | 2.2E-01 | -0.16 |
| RBM28 | 0.04 | 0.03 | -0.01 | 0.00 | 0.00 | -0.06 | 2.2E-01 | -0.03 |
| GMEB2 | -0.42 | -0.24 | 0.19 | 0.23 | 0.11 | 0.02 | 2.2E-01 | 0.28 |
| ASPM | -0.18 | -0.15 | 0.10 | 0.15 | -0.02 | 0.07 | 2.2E-01 | 0.14 |
| CARD19 | -0.03 | 0.02 | 0.16 | -0.12 | -0.02 | -0.02 | 2.2E-01 | -0.10 |
| EBP | -0.10 | -0.26 | 0.00 | 0.01 | -0.07 | 0.36 | 2.2E-01 | 0.22 |
| ZNF146 | -0.01 | -0.03 | -0.16 | -0.04 | 0.20 | 0.01 | 2.2E-01 | 0.13 |
| TMED5 | -0.02 | 0.14 | 0.07 | -0.24 | 0.05 | -0.04 | 2.2E-01 | -0.14 |
| SLC30A1 | -0.20 | -0.09 | 0.07 | 0.00 | 0.01 | 0.19 | 2.2E-01 | 0.14 |
| RP9 | 0.08 | -0.01 | 0.00 | -0.04 | 0.00 | -0.03 | 2.2E-01 | -0.05 |
| VDAC2 | 0.00 | 0.00 | -0.08 | 0.07 | 0.01 | -0.01 | 2.2E-01 | 0.05 |
| ZNF174 | 0.27 | 0.03 | 0.00 | -0.39 | 0.07 | -0.07 | 2.2E-01 | -0.23 |
| DACH1 | 0.04 | 0.02 | 0.06 | -0.12 | -0.07 | 0.06 | 2.2E-01 | -0.08 |
| SURF1 | -0.02 | 0.04 | 0.07 | -0.05 | 0.03 | -0.07 | 2.2E-01 | -0.06 |
| ELAC2 | 0.02 | -0.08 | -0.22 | 0.26 | -0.06 | 0.03 | 2.2E-01 | 0.17 |
| ATP6V1G2 | -0.16 | 0.05 | -0.05 | 0.09 | -0.03 | 0.09 | 2.2E-01 | 0.10 |
| ESPN | -0.05 | 0.18 | 0.24 | 0.19 | -0.34 | -0.33 | 2.2E-01 | -0.28 |
| TBRG4 | 0.00 | 0.01 | -0.15 | 0.10 | 0.04 | -0.01 | 2.2E-01 | 0.09 |
| ANAPC1 | -0.04 | -0.03 | -0.14 | -0.03 | -0.02 | 0.22 | 2.2E-01 | 0.13 |
| NDE1 | -0.09 | 0.09 | 0.16 | -0.02 | -0.13 | -0.03 | 2.2E-01 | -0.12 |
| AAMP | -0.12 | -0.12 | 0.07 | 0.07 | 0.11 | -0.03 | 2.2E-01 | 0.11 |
| TMEM11 | 0.01 | -0.03 | -0.24 | 0.23 | 0.05 | -0.05 | 2.2E-01 | 0.16 |
| ULBP2 | 0.00 | 0.01 | 0.10 | -0.09 | -0.09 | 0.05 | 2.2E-01 | -0.08 |
| USP8 | 0.22 | -0.19 | 0.31 | -0.03 | -0.08 | -0.33 | 2.2E-01 | -0.26 |
| CUL5 | -0.16 | -0.03 | 0.04 | 0.02 | 0.01 | 0.10 | 2.3E-01 | 0.09 |
| ZBTB42 | 0.07 | 0.03 | 0.00 | -0.11 | -0.02 | 0.03 | 2.3E-01 | -0.07 |
| ASH2L | 0.00 | 0.01 | -0.09 | 0.02 | -0.01 | 0.07 | 2.3E-01 | 0.05 |
| PRRC2A | -0.07 | -0.01 | 0.00 | 0.07 | 0.02 | -0.02 | 2.3E-01 | 0.05 |
| SLC27A1 | -0.04 | -0.15 | -0.03 | -0.08 | 0.05 | 0.22 | 2.3E-01 | 0.14 |
| TARDBP | 0.02 | 0.03 | 0.00 | -0.05 | -0.02 | 0.02 | 2.3E-01 | -0.03 |
| ERI3 | 0.09 | 0.09 | -0.06 | -0.06 | -0.07 | 0.01 | 2.3E-01 | -0.08 |
| ECHDC1 | -0.14 | -0.21 | 0.11 | 0.07 | 0.00 | 0.13 | 2.3E-01 | 0.15 |
| RAB20 | 0.03 | 0.03 | 0.11 | -0.22 | 0.03 | 0.01 | 2.3E-01 | -0.12 |
| STOM | -0.11 | 0.04 | -0.07 | -0.03 | 0.04 | 0.12 | 2.3E-01 | 0.09 |
| B4GALT7 | 0.01 | 0.03 | 0.09 | 0.04 | -0.03 | -0.15 | 2.3E-01 | -0.09 |
| HEATR1 | 0.02 | 0.00 | 0.01 | 0.00 | 0.01 | -0.05 | 2.3E-01 | -0.03 |
| AUH | 0.09 | -0.14 | -0.24 | -0.03 | 0.04 | 0.24 | 2.3E-01 | 0.18 |
| ZSCAN21 | 0.11 | 0.18 | -0.07 | 0.02 | -0.02 | -0.26 | 2.3E-01 | -0.16 |
| CDK5RAP3 | -0.02 | 0.00 | 0.13 | 0.00 | -0.03 | -0.09 | 2.3E-01 | -0.08 |
| PAX9 | -0.01 | 0.00 | 0.17 | 0.05 | -0.12 | -0.12 | 2.3E-01 | -0.12 |

| | | | | | | | | |
|---|---|---|---|---|---|---|---|---|
| UPF3B | 0.01 | -0.02 | 0.11 | -0.09 | 0.01 | -0.03 | 2.3E-01 | -0.07 |
| SRA1 | 0.22 | 0.23 | -0.17 | 0.01 | -0.13 | -0.24 | 2.3E-01 | -0.22 |
| IMUP | -0.05 | -0.09 | 0.63 | -0.59 | 0.04 | -0.24 | 2.3E-01 | -0.42 |
| SRPRA | 0.01 | -0.02 | 0.10 | 0.00 | -0.09 | -0.01 | 2.3E-01 | -0.06 |
| ZNF518A | -0.08 | 0.11 | 0.17 | -0.13 | -0.13 | 0.04 | 2.3E-01 | -0.13 |
| HIGD1A | -0.10 | 0.32 | 0.05 | -0.01 | -0.07 | -0.27 | 2.3E-01 | -0.21 |
| ZNF800 | 0.01 | -0.01 | -0.09 | 0.08 | 0.00 | 0.00 | 2.3E-01 | 0.06 |
| DPCD | -0.09 | -0.16 | 0.00 | 0.06 | -0.09 | 0.23 | 2.3E-01 | 0.15 |
| CLK2 | -0.12 | 0.18 | 0.14 | 0.02 | -0.13 | -0.13 | 2.3E-01 | -0.15 |
| LPCAT1 | -0.01 | -0.15 | -0.04 | -0.08 | 0.06 | 0.20 | 2.3E-01 | 0.13 |
| RCOR1 | 0.06 | 0.01 | -0.01 | -0.01 | -0.06 | 0.01 | 2.3E-01 | -0.04 |
| ANLN | -0.07 | 0.07 | 0.13 | 0.01 | -0.11 | -0.04 | 2.3E-01 | -0.10 |
| DHRS7B | 0.03 | -0.02 | 0.03 | -0.03 | 0.01 | -0.02 | 2.3E-01 | -0.03 |
| NAA35 | 0.05 | -0.08 | -0.17 | 0.05 | 0.17 | -0.04 | 2.3E-01 | 0.13 |
| CDS1 | 0.11 | 0.12 | -0.03 | 0.03 | -0.01 | -0.25 | 2.3E-01 | -0.14 |
| CRTAP | -0.10 | 0.03 | -0.05 | -0.02 | 0.03 | 0.10 | 2.3E-01 | 0.07 |
| CETN2 | 0.00 | 0.00 | 0.02 | -0.03 | 0.01 | 0.00 | 2.3E-01 | -0.01 |
| ZPR1 | -0.02 | 0.02 | -0.15 | -0.05 | 0.08 | 0.10 | 2.3E-01 | 0.10 |
| STIP1 | 0.02 | -0.05 | -0.06 | 0.10 | 0.03 | -0.03 | 2.3E-01 | 0.06 |
| EIF2S2 | 0.03 | 0.02 | 0.07 | 0.02 | 0.01 | -0.17 | 2.3E-01 | -0.09 |
| LSM14A | 0.05 | 0.02 | -0.02 | 0.01 | -0.05 | -0.01 | 2.3E-01 | -0.04 |
| TPI1 | -0.20 | -0.05 | -0.04 | 0.13 | -0.14 | 0.25 | 2.3E-01 | 0.18 |
| EMC4 | -0.07 | -0.05 | -0.04 | 0.18 | -0.07 | 0.03 | 2.3E-01 | 0.10 |
| PPP2R1B | -0.04 | -0.14 | 0.04 | 0.01 | 0.00 | 0.12 | 2.3E-01 | 0.09 |
| PPWD1 | -0.01 | -0.03 | -0.13 | -0.05 | 0.02 | 0.18 | 2.3E-01 | 0.11 |
| CCNT2 | -0.07 | -0.12 | -0.03 | 0.24 | -0.06 | 0.01 | 2.3E-01 | 0.13 |
| NDUFAB1 | 0.00 | 0.00 | -0.16 | 0.13 | -0.04 | 0.05 | 2.3E-01 | 0.10 |
| NACA2 | 0.16 | -0.01 | 0.05 | 0.09 | -0.09 | -0.24 | 2.3E-01 | -0.15 |
| DHCR7 | -0.01 | 0.03 | 0.10 | -0.08 | 0.05 | -0.10 | 2.3E-01 | -0.08 |
| RPRD1A | -0.16 | -0.13 | 0.02 | 0.10 | -0.11 | 0.23 | 2.3E-01 | 0.16 |
| HNRNPF | -0.02 | -0.04 | 0.01 | -0.01 | 0.00 | 0.05 | 2.3E-01 | 0.03 |
| MAIP1 | -0.03 | -0.06 | 0.03 | 0.02 | 0.02 | 0.02 | 2.3E-01 | 0.04 |
| EIF4G3 | -0.13 | -0.02 | 0.00 | -0.06 | 0.06 | 0.13 | 2.3E-01 | 0.09 |
| FAM111A | 0.01 | 0.03 | 0.00 | -0.03 | 0.02 | -0.03 | 2.3E-01 | -0.03 |
| MGAT1 | -0.16 | -0.05 | 0.05 | 0.10 | 0.05 | -0.02 | 2.3E-01 | 0.10 |
| NUP133 | 0.03 | -0.04 | -0.10 | -0.02 | 0.06 | 0.05 | 2.3E-01 | 0.06 |
| YJU2B | -0.01 | 0.03 | 0.15 | -0.22 | 0.01 | 0.01 | 2.3E-01 | -0.13 |
| UBA2 | -0.07 | 0.04 | -0.20 | 0.18 | -0.06 | 0.07 | 2.3E-01 | 0.14 |
| RAB5C | -0.08 | 0.02 | 0.25 | -0.11 | -0.03 | -0.09 | 2.3E-01 | -0.14 |
| PA2G4 | 0.04 | 0.04 | 0.06 | 0.07 | -0.06 | -0.16 | 2.3E-01 | -0.09 |
| EMC10 | -0.03 | 0.05 | 0.05 | 0.00 | 0.00 | -0.06 | 2.3E-01 | -0.04 |
| SLC25A14 | -0.07 | 0.07 | 0.10 | -0.04 | -0.05 | -0.02 | 2.4E-01 | -0.07 |
| LRRC31 | -0.20 | -0.12 | 0.11 | 0.08 | 0.06 | 0.04 | 2.4E-01 | 0.13 |
| MICOS13 | 0.08 | 0.03 | -0.02 | 0.03 | -0.04 | -0.09 | 2.4E-01 | -0.06 |
| RBFA | 0.09 | -0.02 | 0.18 | -0.32 | 0.05 | -0.02 | 2.4E-01 | -0.18 |
| CCT5 | -0.02 | 0.25 | -0.02 | -0.08 | 0.03 | -0.19 | 2.4E-01 | -0.15 |
| BLOC1S4 | 0.05 | -0.04 | -0.35 | 0.08 | -0.05 | 0.24 | 2.4E-01 | 0.21 |
| MAP4K3 | -0.04 | 0.03 | -0.14 | 0.04 | 0.13 | -0.03 | 2.4E-01 | 0.09 |

| | | | | | | | | |
|---|---|---|---|---|---|---|---|---|
| ANKRD22 | 0.01 | 0.10 | -0.02 | 0.01 | -0.03 | -0.09 | 2.4E-01 | -0.06 |
| CYB5B | 0.07 | 0.12 | -0.01 | 0.10 | -0.20 | -0.11 | 2.4E-01 | -0.13 |
| ARPC4 | 0.02 | 0.08 | -0.01 | -0.09 | -0.05 | 0.04 | 2.4E-01 | -0.07 |
| CENPO | -0.05 | 0.02 | -0.07 | -0.02 | 0.01 | 0.09 | 2.4E-01 | 0.06 |
| KARS1 | -0.03 | 0.03 | 0.07 | -0.03 | 0.00 | -0.04 | 2.4E-01 | -0.05 |
| RUFY1 | -0.20 | -0.13 | 0.11 | 0.04 | 0.08 | 0.07 | 2.4E-01 | 0.13 |
| DDX51 | -0.08 | -0.05 | 0.03 | 0.04 | -0.03 | 0.08 | 2.4E-01 | 0.06 |
| UBE2A | 0.13 | 0.12 | -0.07 | 0.05 | -0.10 | -0.15 | 2.4E-01 | -0.12 |
| TMEM208 | -0.27 | -0.21 | -0.08 | -0.09 | -0.10 | 0.56 | 2.4E-01 | 0.31 |
| DHRS7 | 0.07 | -0.06 | -0.15 | 0.04 | 0.07 | 0.02 | 2.4E-01 | 0.09 |
| ZNF7 | -0.18 | -0.05 | 0.06 | 0.08 | -0.01 | 0.08 | 2.4E-01 | 0.10 |
| UBE2K | 0.08 | -0.21 | -0.14 | 0.06 | 0.22 | -0.05 | 2.4E-01 | 0.17 |
| CPSF4 | -0.06 | -0.11 | -0.05 | 0.23 | -0.12 | 0.09 | 2.4E-01 | 0.14 |
| KIF21A | 0.01 | -0.04 | -0.15 | 0.06 | 0.15 | -0.06 | 2.4E-01 | 0.11 |
| USP30 | 0.06 | -0.04 | 0.05 | 0.00 | -0.06 | -0.02 | 2.4E-01 | -0.05 |
| MYO1D | -0.04 | 0.03 | 0.10 | -0.06 | -0.05 | 0.02 | 2.4E-01 | -0.06 |
| MAN1B1 | -0.03 | 0.02 | 0.09 | 0.00 | -0.04 | -0.04 | 2.4E-01 | -0.05 |
| CAMLG | 0.19 | 0.13 | -0.12 | 0.00 | -0.11 | -0.13 | 2.4E-01 | -0.14 |
| LCOR | -0.01 | -0.03 | -0.06 | -0.01 | 0.11 | -0.01 | 2.4E-01 | 0.06 |
| GPRIN1 | -0.13 | 0.10 | 0.23 | -0.05 | -0.14 | -0.05 | 2.4E-01 | -0.15 |
| POM121 | -0.02 | -0.02 | -0.06 | -0.03 | 0.01 | 0.12 | 2.4E-01 | 0.07 |
| FNBP1 | 0.02 | -0.02 | 0.09 | 0.04 | -0.07 | -0.07 | 2.4E-01 | -0.07 |
| ADGRB2 | 0.04 | -0.02 | 0.10 | 0.04 | -0.04 | -0.13 | 2.4E-01 | -0.08 |
| NPC1L1 | -0.23 | -0.10 | 0.00 | 0.06 | -0.12 | 0.32 | 2.4E-01 | 0.20 |
| CNN3 | 0.12 | -0.25 | 0.51 | -0.07 | -0.26 | -0.21 | 2.4E-01 | -0.31 |
| EPB41L4B | 0.00 | 0.02 | 0.10 | -0.15 | -0.04 | 0.05 | 2.4E-01 | -0.09 |
| PGM1 | -0.14 | -0.14 | -0.05 | -0.11 | 0.01 | 0.36 | 2.4E-01 | 0.20 |
| TMEM54 | -0.02 | 0.14 | 0.07 | 0.06 | -0.03 | -0.25 | 2.4E-01 | -0.14 |
| KANK2 | -0.15 | 0.06 | -0.13 | -0.05 | 0.04 | 0.20 | 2.4E-01 | 0.14 |
| MTRES1 | 0.29 | -0.05 | -0.03 | -0.09 | -0.12 | -0.03 | 2.4E-01 | -0.15 |
| INTS6L | 0.02 | -0.05 | 0.25 | -0.21 | 0.01 | -0.05 | 2.4E-01 | -0.16 |
| EIF3B | -0.10 | -0.16 | -0.01 | -0.08 | 0.02 | 0.29 | 2.4E-01 | 0.16 |
| RPL24 | -0.05 | -0.08 | 0.04 | 0.05 | 0.06 | -0.02 | 2.4E-01 | 0.06 |
| POLR2I | 0.25 | 0.16 | -0.16 | -0.09 | -0.02 | -0.19 | 2.4E-01 | -0.18 |
| SRSF5 | 0.09 | 0.03 | 0.04 | -0.19 | 0.09 | -0.08 | 2.4E-01 | -0.11 |
| MREG | -0.12 | -0.17 | 0.06 | 0.07 | -0.07 | 0.19 | 2.4E-01 | 0.14 |
| ORC1 | -0.09 | -0.11 | 0.02 | -0.05 | 0.01 | 0.20 | 2.4E-01 | 0.11 |
| NUP37 | 0.08 | 0.03 | 0.02 | 0.03 | 0.00 | -0.16 | 2.4E-01 | -0.08 |
| RHBDD1 | -0.04 | -0.10 | -0.04 | 0.17 | -0.11 | 0.10 | 2.4E-01 | 0.12 |
| TAF15 | 0.17 | 0.13 | -0.09 | 0.05 | -0.08 | -0.22 | 2.5E-01 | -0.16 |
| GAS2L1 | -0.11 | 0.21 | 0.11 | -0.06 | 0.01 | -0.20 | 2.5E-01 | -0.15 |
| FTSJ3 | 0.01 | -0.02 | -0.02 | 0.03 | 0.01 | -0.01 | 2.5E-01 | 0.02 |
| COX6C | 0.05 | 0.01 | 0.04 | 0.06 | -0.06 | -0.11 | 2.5E-01 | -0.07 |
| PIGU | 0.08 | 0.03 | -0.02 | 0.01 | -0.09 | -0.01 | 2.5E-01 | -0.06 |
| METTL15 | 0.07 | 0.06 | 0.04 | -0.23 | -0.06 | 0.09 | 2.5E-01 | -0.12 |
| SENP7 | 0.04 | 0.02 | 0.03 | -0.07 | -0.08 | 0.06 | 2.5E-01 | -0.06 |
| BRD8 | 0.09 | 0.06 | -0.03 | 0.06 | -0.12 | -0.07 | 2.5E-01 | -0.09 |
| TAF5L | -0.23 | 0.00 | -0.29 | 0.52 | -0.06 | -0.10 | 2.5E-01 | 0.30 |

| | | | | | | | | |
|---|---|---|---|---|---|---|---|---|
| ZC3HC1 | 0.03 | -0.05 | 0.13 | 0.01 | -0.09 | -0.05 | 2.5E-01 | -0.08 |
| HGSNAT | 0.17 | -0.11 | 0.12 | -0.15 | -0.02 | -0.04 | 2.5E-01 | -0.13 |
| PDK3 | -0.02 | -0.04 | -0.03 | -0.06 | 0.06 | 0.09 | 2.5E-01 | 0.06 |
| NUP155 | 0.07 | 0.00 | 0.03 | 0.04 | -0.01 | -0.15 | 2.5E-01 | -0.08 |
| RPL6 | 0.04 | 0.02 | 0.08 | 0.06 | -0.03 | -0.19 | 2.5E-01 | -0.10 |
| MCUB | 0.06 | 0.10 | -0.02 | 0.06 | -0.07 | -0.15 | 2.5E-01 | -0.10 |
| DDX50 | 0.09 | 0.04 | -0.04 | 0.01 | -0.06 | -0.04 | 2.5E-01 | -0.06 |
| SETD5 | 0.04 | -0.17 | -0.15 | 0.05 | -0.10 | 0.28 | 2.5E-01 | 0.17 |
| RGS3 | -0.11 | -0.02 | -0.02 | -0.08 | 0.13 | 0.08 | 2.5E-01 | 0.09 |
| VTI1B | -0.05 | 0.06 | 0.12 | 0.04 | -0.11 | -0.07 | 2.5E-01 | -0.09 |
| MRS2 | 0.01 | 0.01 | 0.02 | 0.03 | -0.04 | -0.04 | 2.5E-01 | -0.03 |
| RTN4RL2 | -0.11 | 0.09 | 0.21 | -0.12 | 0.01 | -0.11 | 2.5E-01 | -0.14 |
| RRP12 | -0.01 | -0.02 | -0.03 | 0.06 | 0.03 | -0.03 | 2.5E-01 | 0.04 |
| DSC2 | -0.04 | -0.02 | -0.01 | 0.09 | -0.02 | 0.00 | 2.5E-01 | 0.05 |
| VPS13D | -0.14 | -0.08 | 0.00 | -0.11 | 0.07 | 0.23 | 2.5E-01 | 0.14 |
| ZC3H3 | -0.01 | 0.02 | 0.08 | 0.00 | 0.00 | -0.10 | 2.5E-01 | -0.06 |
| PDHB | 0.06 | 0.13 | -0.04 | 0.02 | 0.00 | -0.19 | 2.5E-01 | -0.11 |
| RASSF7 | -0.28 | -0.06 | 0.08 | -0.05 | 0.12 | 0.14 | 2.5E-01 | 0.16 |
| MTG1 | 0.05 | 0.08 | -0.04 | -0.03 | -0.08 | 0.02 | 2.5E-01 | -0.06 |
| NUP214 | 0.07 | 0.04 | -0.02 | 0.01 | 0.00 | -0.11 | 2.5E-01 | -0.06 |
| LRRC45 | 0.10 | 0.03 | 0.16 | 0.10 | -0.03 | -0.42 | 2.5E-01 | -0.21 |
| ACTL6A | 0.02 | -0.07 | -0.02 | 0.05 | -0.03 | 0.05 | 2.5E-01 | 0.05 |
| MCMBP | 0.01 | -0.08 | -0.02 | 0.05 | 0.07 | -0.04 | 2.5E-01 | 0.06 |
| BCAS2 | 0.03 | -0.01 | 0.03 | -0.05 | 0.03 | -0.04 | 2.5E-01 | -0.04 |
| SLC9A6 | -0.33 | -0.01 | 0.05 | 0.04 | 0.02 | 0.17 | 2.5E-01 | 0.17 |
| RBBP9 | -0.17 | -0.18 | 0.11 | -0.02 | 0.19 | 0.04 | 2.5E-01 | 0.15 |
| KDM5B | -0.03 | -0.01 | -0.01 | 0.01 | -0.03 | 0.06 | 2.5E-01 | 0.04 |
| B3GAT3 | 0.05 | 0.03 | 0.07 | 0.09 | -0.09 | -0.17 | 2.5E-01 | -0.10 |
| NOL8 | 0.03 | -0.05 | -0.04 | 0.01 | 0.01 | 0.04 | 2.5E-01 | 0.04 |
| SMN1 | -0.01 | 0.03 | 0.20 | -0.30 | 0.03 | 0.01 | 2.5E-01 | -0.16 |
| HNRNPU | 0.01 | -0.03 | -0.03 | 0.01 | -0.01 | 0.05 | 2.5E-01 | 0.03 |
| EIF4A2 | -0.07 | 0.05 | 0.15 | -0.03 | -0.03 | -0.08 | 2.5E-01 | -0.09 |
| SIDT2 | -0.05 | 0.04 | 0.14 | 0.00 | -0.01 | -0.13 | 2.6E-01 | -0.09 |
| PRRG1 | 0.04 | 0.04 | 0.01 | -0.13 | 0.01 | 0.02 | 2.6E-01 | -0.06 |
| TMTC4 | 0.02 | 0.01 | 0.07 | -0.09 | 0.07 | -0.09 | 2.6E-01 | -0.07 |
| PARN | -0.06 | 0.04 | -0.04 | 0.05 | 0.03 | -0.01 | 2.6E-01 | 0.04 |
| TXNDC12 | -0.24 | -0.05 | -0.03 | -0.11 | 0.03 | 0.33 | 2.6E-01 | 0.19 |
| ISCA2 | -0.15 | 0.10 | -0.15 | -0.02 | 0.09 | 0.10 | 2.6E-01 | 0.12 |
| OSBPL7 | -0.02 | 0.00 | -0.06 | 0.02 | 0.08 | -0.04 | 2.6E-01 | 0.05 |
| MTHFD2 | 0.04 | -0.06 | -0.12 | -0.05 | 0.07 | 0.11 | 2.6E-01 | 0.09 |
| ISCU | -0.29 | -0.26 | 0.07 | 0.09 | -0.17 | 0.43 | 2.6E-01 | 0.28 |
| TAP2 | -0.02 | 0.03 | 0.05 | 0.03 | -0.05 | -0.04 | 2.6E-01 | -0.04 |
| RNF121 | 0.01 | -0.07 | -0.15 | 0.11 | 0.19 | -0.12 | 2.6E-01 | 0.14 |
| SMARCC2 | 0.01 | -0.06 | -0.06 | 0.00 | -0.03 | 0.12 | 2.6E-01 | 0.07 |
| KRT1 | -0.23 | -0.18 | -0.60 | 0.52 | -0.55 | 0.57 | 2.6E-01 | 0.52 |
| CTTN | -0.01 | 0.10 | 0.04 | 0.04 | -0.02 | -0.15 | 2.6E-01 | -0.09 |
| TRMT2A | -0.03 | -0.08 | -0.01 | 0.08 | 0.09 | -0.07 | 2.6E-01 | 0.07 |
| TEAD3 | 0.05 | 0.03 | -0.01 | 0.04 | -0.06 | -0.07 | 2.6E-01 | -0.05 |

| | | | | | | | | |
|---|---|---|---|---|---|---|---|---|
| SRP19 | -0.05 | -0.08 | 0.02 | -0.02 | 0.13 | 0.00 | 2.6E-01 | 0.07 |
| CPSF6 | 0.07 | 0.02 | 0.06 | 0.09 | -0.10 | -0.14 | 2.6E-01 | -0.10 |
| MFN2 | -0.10 | -0.09 | 0.04 | 0.09 | -0.06 | 0.11 | 2.6E-01 | 0.09 |
| RIOX1 | -0.01 | 0.12 | 0.00 | -0.09 | -0.06 | 0.03 | 2.6E-01 | -0.08 |
| ZNF598 | -0.03 | 0.01 | 0.08 | -0.01 | -0.05 | -0.01 | 2.6E-01 | -0.04 |
| PSMD6 | -0.10 | -0.16 | 0.03 | -0.11 | 0.22 | 0.09 | 2.6E-01 | 0.14 |
| NAPA | 0.01 | 0.06 | 0.08 | 0.09 | -0.08 | -0.19 | 2.6E-01 | -0.11 |
| RBM39 | 0.08 | 0.05 | -0.02 | -0.10 | 0.07 | -0.10 | 2.6E-01 | -0.08 |
| TCF20 | 0.02 | 0.04 | 0.02 | -0.09 | -0.02 | 0.04 | 2.6E-01 | -0.05 |
| TRPT1 | -0.06 | 0.09 | 0.11 | -0.13 | 0.05 | -0.07 | 2.6E-01 | -0.10 |
| U2AF2 | 0.04 | 0.01 | -0.01 | -0.01 | 0.01 | -0.03 | 2.6E-01 | -0.02 |
| CABIN1 | 0.04 | -0.04 | -0.11 | -0.04 | 0.08 | 0.07 | 2.6E-01 | 0.08 |
| IQSEC1 | 0.01 | 0.03 | -0.01 | -0.03 | -0.02 | 0.02 | 2.6E-01 | -0.03 |
| RAB24 | -0.06 | 0.04 | 0.15 | -0.12 | -0.03 | 0.00 | 2.6E-01 | -0.09 |
| PARD6G | -0.04 | -0.10 | -0.01 | -0.09 | 0.10 | 0.12 | 2.6E-01 | 0.09 |
| C1orf122 | -0.05 | -0.04 | -0.07 | -0.01 | -0.05 | 0.20 | 2.6E-01 | 0.10 |
| ATP6AP1 | 0.03 | -0.06 | -0.09 | 0.13 | -0.02 | 0.00 | 2.6E-01 | 0.08 |
| MEA1 | 0.06 | 0.12 | 0.02 | 0.06 | -0.30 | -0.01 | 2.6E-01 | -0.15 |
| SFPQ | 0.05 | -0.01 | 0.05 | 0.01 | 0.01 | -0.11 | 2.6E-01 | -0.06 |
| RTF2 | 0.15 | 0.19 | -0.11 | -0.28 | 0.05 | -0.06 | 2.6E-01 | -0.18 |
| SCRIB | -0.15 | 0.00 | 0.03 | 0.01 | 0.02 | 0.07 | 2.6E-01 | 0.08 |
| MPV17L | 0.19 | 0.06 | -0.02 | 0.14 | -0.15 | -0.28 | 2.7E-01 | -0.18 |
| ATAD2B | -0.07 | -0.06 | -0.03 | 0.07 | -0.09 | 0.15 | 2.7E-01 | 0.09 |
| EEF1B2 | 0.01 | 0.03 | 0.03 | 0.05 | -0.05 | -0.08 | 2.7E-01 | -0.05 |
| NFS1 | -0.05 | 0.00 | -0.05 | 0.12 | 0.01 | -0.04 | 2.7E-01 | 0.06 |
| DLAT | -0.03 | 0.03 | -0.11 | 0.09 | -0.04 | 0.05 | 2.7E-01 | 0.07 |
| CCPG1 | -0.05 | 0.05 | 0.13 | -0.05 | 0.03 | -0.13 | 2.7E-01 | -0.09 |
| CDC20 | -0.06 | 0.12 | 0.06 | -0.06 | -0.12 | 0.03 | 2.7E-01 | -0.09 |
| SPTBN1 | -0.10 | -0.03 | 0.01 | 0.15 | -0.02 | -0.01 | 2.7E-01 | 0.08 |
| TNRC6B | -0.51 | -0.29 | 0.19 | 0.28 | -0.19 | 0.32 | 2.7E-01 | 0.34 |
| RAP2A | -0.06 | 0.15 | 0.11 | -0.16 | -0.19 | 0.11 | 2.7E-01 | -0.15 |
| PDIA5 | 0.05 | 0.09 | 0.00 | -0.06 | 0.07 | -0.15 | 2.7E-01 | -0.09 |
| ARHGAP26 | -0.20 | 0.02 | 0.01 | 0.13 | 0.00 | 0.02 | 2.7E-01 | 0.11 |
| HSBP1 | -0.19 | -0.11 | 0.10 | -0.03 | 0.10 | 0.11 | 2.7E-01 | 0.12 |
| TBC1D10B | 0.04 | 0.11 | -0.02 | 0.02 | 0.01 | -0.18 | 2.7E-01 | -0.09 |
| RNASEH2B | -0.06 | 0.06 | -0.13 | -0.03 | 0.08 | 0.08 | 2.7E-01 | 0.08 |
| BRDT | -0.17 | -0.32 | 0.11 | 0.25 | -0.14 | 0.18 | 2.7E-01 | 0.22 |
| MAP7D1 | -0.14 | 0.04 | -0.02 | 0.09 | 0.00 | 0.01 | 2.7E-01 | 0.08 |
| PSMB1 | -0.13 | 0.11 | 0.29 | -0.30 | 0.04 | -0.08 | 2.7E-01 | -0.20 |
| TRAF3 | 0.14 | -0.11 | 0.13 | -0.02 | -0.15 | -0.01 | 2.7E-01 | -0.12 |
| MAD2L2 | -0.01 | 0.05 | 0.05 | -0.06 | -0.10 | 0.06 | 2.7E-01 | -0.07 |
| POMGNT1 | -0.04 | 0.03 | 0.12 | -0.07 | 0.04 | -0.09 | 2.7E-01 | -0.08 |
| ATP6V1B2 | 0.01 | 0.04 | 0.00 | 0.00 | 0.01 | -0.05 | 2.7E-01 | -0.03 |
| AIMP2 | -0.10 | -0.03 | 0.02 | 0.06 | 0.10 | -0.05 | 2.7E-01 | 0.07 |
| MYO19 | 0.02 | -0.02 | 0.06 | 0.01 | -0.03 | -0.03 | 2.7E-01 | -0.04 |
| QKI | 0.07 | 0.00 | 0.07 | -0.18 | 0.06 | -0.03 | 2.7E-01 | -0.09 |
| MAFB | 0.15 | -0.01 | -0.03 | -0.06 | -0.08 | 0.01 | 2.7E-01 | -0.08 |
| SMARCC1 | 0.08 | 0.04 | -0.04 | -0.09 | 0.01 | -0.01 | 2.7E-01 | -0.06 |

| | | | | | | | | |
|---|---|---|---|---|---|---|---|---|
| MSI1 | 0.10 | 0.12 | -0.05 | -0.14 | 0.08 | -0.13 | 2.7E-01 | -0.11 |
| KATNA1 | -0.16 | 0.16 | -0.40 | 0.29 | 0.01 | 0.01 | 2.7E-01 | 0.24 |
| CCDC86 | 0.01 | 0.01 | 0.06 | -0.10 | -0.01 | 0.03 | 2.7E-01 | -0.05 |
| TAF1 | -0.01 | -0.02 | -0.04 | 0.01 | -0.04 | 0.10 | 2.7E-01 | 0.05 |
| BUB1B | 0.02 | 0.00 | -0.09 | -0.01 | 0.06 | 0.02 | 2.7E-01 | 0.05 |
| DISP2 | 0.01 | 0.22 | 0.06 | 0.01 | 0.07 | -0.44 | 2.7E-01 | -0.22 |
| IFIT5 | -0.05 | 0.02 | -0.09 | 0.04 | 0.12 | -0.05 | 2.7E-01 | 0.07 |
| CDR2L | -0.02 | 0.02 | -0.09 | 0.08 | -0.03 | 0.03 | 2.7E-01 | 0.06 |
| CAPZB | -0.03 | -0.03 | 0.21 | 0.00 | -0.07 | -0.10 | 2.7E-01 | -0.11 |
| PYCR1 | -0.12 | -0.05 | 0.02 | 0.11 | -0.07 | 0.09 | 2.7E-01 | 0.09 |
| COPS6 | 0.05 | -0.08 | -0.14 | 0.10 | 0.12 | -0.08 | 2.7E-01 | 0.11 |
| MIS18BP1 | -0.01 | -0.07 | -0.01 | -0.03 | 0.01 | 0.11 | 2.7E-01 | 0.06 |
| FLYWCH1 | -0.01 | 0.04 | 0.07 | -0.14 | 0.02 | 0.01 | 2.7E-01 | -0.07 |
| SYNGR1 | -0.09 | -0.05 | 0.54 | -0.56 | 0.02 | -0.09 | 2.7E-01 | -0.35 |
| NIN | -0.26 | -0.35 | -0.06 | 0.68 | -0.17 | -0.10 | 2.7E-01 | 0.36 |
| PALS2 | -0.06 | 0.06 | 0.11 | -0.09 | -0.06 | 0.03 | 2.7E-01 | -0.08 |
| TAF9B | -0.03 | -0.03 | -0.05 | 0.09 | -0.07 | 0.08 | 2.7E-01 | 0.07 |
| HARS2 | 0.03 | -0.01 | -0.15 | -0.03 | 0.08 | 0.07 | 2.7E-01 | 0.08 |
| HADHA | 0.07 | -0.01 | -0.01 | 0.00 | -0.01 | -0.05 | 2.7E-01 | -0.04 |
| POLE4 | -0.02 | -0.09 | 0.03 | 0.07 | 0.03 | -0.02 | 2.7E-01 | 0.06 |
| FOXO3 | -0.11 | -0.11 | 0.05 | -0.04 | 0.02 | 0.18 | 2.8E-01 | 0.11 |
| ACOT2 | -0.05 | -0.08 | -0.05 | -0.05 | -0.02 | 0.22 | 2.8E-01 | 0.11 |
| NDUFA13 | -0.01 | -0.03 | -0.09 | 0.16 | -0.01 | -0.04 | 2.8E-01 | 0.08 |
| CLK1 | 0.10 | 0.01 | 0.07 | 0.07 | -0.02 | -0.25 | 2.8E-01 | -0.13 |
| ALDOC | 0.11 | 0.10 | 0.06 | -0.03 | -0.43 | 0.11 | 2.8E-01 | -0.20 |
| RAB4B | -0.06 | 0.01 | 0.21 | 0.01 | -0.09 | -0.10 | 2.8E-01 | -0.11 |
| PIGA | -0.02 | 0.06 | 0.05 | -0.04 | 0.04 | -0.10 | 2.8E-01 | -0.06 |
| EXOSC3 | 0.06 | -0.07 | -0.29 | 0.27 | -0.08 | 0.05 | 2.8E-01 | 0.18 |
| CYFIP2 | -0.03 | 0.03 | 0.10 | 0.03 | -0.07 | -0.06 | 2.8E-01 | -0.06 |
| ARGLU1 | 0.01 | -0.08 | 0.01 | 0.02 | 0.00 | 0.04 | 2.8E-01 | 0.04 |
| GSK3B | 0.05 | 0.10 | -0.06 | -0.01 | -0.01 | -0.07 | 2.8E-01 | -0.06 |
| ELMO3 | -0.05 | 0.08 | 0.15 | -0.21 | -0.06 | 0.06 | 2.8E-01 | -0.12 |
| BRPF1 | 0.02 | -0.01 | 0.05 | -0.09 | 0.01 | 0.01 | 2.8E-01 | -0.04 |
| YBX2 | -0.02 | 0.01 | 0.11 | 0.00 | -0.01 | -0.09 | 2.8E-01 | -0.06 |
| CCDC120 | -0.39 | -0.04 | 0.11 | 0.01 | 0.08 | 0.18 | 2.8E-01 | 0.19 |
| CDK17 | -0.15 | -0.01 | 0.00 | -0.08 | 0.11 | 0.10 | 2.8E-01 | 0.10 |
| CARM1 | -0.08 | -0.39 | 0.06 | 0.29 | -0.15 | 0.16 | 2.8E-01 | 0.23 |
| HDAC3 | 0.10 | -0.63 | 0.00 | 0.21 | 0.20 | -0.04 | 2.8E-01 | 0.30 |
| PPP1CC | -0.07 | -0.09 | -0.03 | -0.11 | 0.07 | 0.20 | 2.8E-01 | 0.12 |
| PPRC1 | 0.26 | -0.07 | 0.01 | 0.01 | -0.05 | -0.19 | 2.8E-01 | -0.14 |
| LRRC1 | -0.22 | -0.04 | 0.07 | 0.00 | 0.03 | 0.12 | 2.8E-01 | 0.12 |
| TUBG1 | 0.09 | -0.10 | 0.16 | -0.11 | -0.02 | -0.04 | 2.8E-01 | -0.10 |
| BPNT1 | 0.07 | 0.26 | -0.11 | -0.20 | 0.06 | -0.13 | 2.8E-01 | -0.17 |
| SRGAP2 | -0.03 | 0.04 | 0.05 | 0.02 | -0.03 | -0.07 | 2.8E-01 | -0.04 |
| KCTD15 | 0.02 | -0.01 | 0.11 | -0.14 | 0.04 | -0.03 | 2.8E-01 | -0.08 |
| FUS | 0.04 | -0.01 | 0.02 | -0.06 | -0.02 | 0.02 | 2.8E-01 | -0.03 |
| TDRD7 | 1.52 | -0.30 | -0.83 | -0.72 | -0.77 | -0.80 | 2.8E-01 | -0.89 |
| RFT1 | 0.01 | -0.03 | 0.21 | -0.23 | -0.03 | 0.04 | 2.8E-01 | -0.14 |

| | | | | | | | | |
|---|---|---|---|---|---|---|---|---|
| NHP2 | 0.04 | 0.04 | 0.06 | 0.02 | 0.04 | -0.22 | 2.8E-01 | -0.11 |
| DNAJC2 | -0.03 | -0.06 | -0.11 | 0.24 | -0.06 | -0.01 | 2.8E-01 | 0.12 |
| MYCBP2 | -0.11 | 0.03 | -0.03 | -0.02 | 0.11 | 0.01 | 2.8E-01 | 0.07 |
| SSNA1 | 0.05 | -0.14 | -0.10 | 0.18 | -0.06 | 0.04 | 2.8E-01 | 0.12 |
| UBFD1 | 0.34 | 0.25 | -0.27 | -0.15 | -0.25 | -0.04 | 2.8E-01 | -0.25 |
| AAK1 | -0.21 | -0.15 | 0.11 | 0.18 | -0.08 | 0.12 | 2.8E-01 | 0.16 |
| STK40 | 0.05 | 0.02 | 0.37 | -0.28 | -0.69 | 0.28 | 2.8E-01 | -0.38 |
| INPP4B | 0.38 | -0.35 | 0.50 | -0.84 | -0.34 | 0.21 | 2.8E-01 | -0.50 |
| POP4 | -0.05 | -0.05 | 0.00 | 0.09 | -0.07 | 0.07 | 2.8E-01 | 0.06 |
| PDHX | 0.01 | 0.07 | -0.01 | 0.03 | -0.05 | -0.05 | 2.8E-01 | -0.05 |
| NDUFAF4 | -0.01 | 0.01 | -0.16 | 0.16 | -0.01 | -0.01 | 2.8E-01 | 0.10 |
| TRIM25 | -0.10 | 0.01 | -0.04 | 0.14 | -0.04 | 0.00 | 2.8E-01 | 0.08 |
| NUP153 | 0.03 | -0.01 | -0.11 | 0.07 | 0.00 | 0.02 | 2.8E-01 | 0.06 |
| KDM3B | -0.07 | -0.05 | 0.04 | 0.00 | 0.02 | 0.05 | 2.8E-01 | 0.05 |
| MBOAT1 | 0.08 | 0.11 | 0.03 | -0.29 | 0.13 | -0.09 | 2.8E-01 | -0.15 |
| MED10 | 0.12 | 0.13 | -0.04 | 0.04 | 0.01 | -0.30 | 2.8E-01 | -0.15 |
| TNFRSF12A | -0.06 | 0.10 | 0.13 | 0.00 | -0.23 | 0.03 | 2.8E-01 | -0.12 |
| DYNC1LI2 | 0.05 | -0.01 | 0.00 | 0.00 | -0.03 | -0.01 | 2.8E-01 | -0.03 |
| BCKDHA | 0.11 | 0.10 | -0.07 | 0.05 | -0.08 | -0.12 | 2.8E-01 | -0.09 |
| KAT2A | -0.03 | -0.01 | -0.12 | 0.18 | -0.01 | -0.04 | 2.8E-01 | 0.10 |
| SNAP23 | -0.02 | 0.04 | 0.04 | 0.03 | -0.04 | -0.05 | 2.8E-01 | -0.04 |
| SETMAR | -0.27 | -0.19 | 0.15 | 0.11 | -0.06 | 0.19 | 2.8E-01 | 0.18 |
| ZNF865 | -0.02 | -0.04 | -0.08 | -0.09 | 0.05 | 0.17 | 2.8E-01 | 0.09 |
| TMEM94 | -0.25 | -0.04 | 0.09 | 0.02 | 0.09 | 0.07 | 2.8E-01 | 0.12 |
| CUX1 | 0.01 | -0.04 | 0.16 | -0.02 | 0.00 | -0.11 | 2.8E-01 | -0.09 |
| ACSL3 | 0.02 | -0.01 | -0.09 | 0.00 | 0.07 | 0.01 | 2.8E-01 | 0.05 |
| SAXO1 | -0.02 | -0.12 | 0.45 | -0.13 | -0.31 | 0.02 | 2.8E-01 | -0.25 |
| AVEN | -0.17 | -0.01 | 0.02 | 0.00 | -0.01 | 0.16 | 2.9E-01 | 0.10 |
| RBMS1 | -0.05 | -0.07 | -0.06 | 0.23 | -0.08 | 0.01 | 2.9E-01 | 0.11 |
| ARHGDIA | 0.00 | -0.07 | -0.02 | 0.05 | -0.05 | 0.09 | 2.9E-01 | 0.06 |
| MAPKAPK2 | -0.01 | 0.04 | -0.30 | 0.24 | -0.03 | 0.01 | 2.9E-01 | 0.17 |
| SSBP3 | 0.06 | 0.07 | -0.01 | 0.08 | -0.13 | -0.08 | 2.9E-01 | -0.08 |
| HAX1 | -0.03 | -0.01 | 0.12 | -0.06 | 0.00 | -0.04 | 2.9E-01 | -0.06 |
| BZW2 | 0.17 | 0.06 | -0.09 | -0.10 | 0.01 | -0.07 | 2.9E-01 | -0.10 |
| NDFIP1 | -0.07 | 0.05 | 0.18 | -0.15 | -0.11 | 0.07 | 2.9E-01 | -0.12 |
| MED12 | 0.01 | 0.00 | -0.18 | 0.16 | -0.03 | 0.02 | 2.9E-01 | 0.10 |
| RIPK2 | -0.36 | -0.27 | 0.20 | -0.08 | 0.13 | 0.27 | 2.9E-01 | 0.25 |
| GLS | -0.22 | -0.05 | 0.01 | 0.03 | -0.09 | 0.27 | 2.9E-01 | 0.16 |
| KRT10 | -0.09 | -0.20 | -0.53 | 0.10 | -0.38 | 0.73 | 2.9E-01 | 0.43 |
| WDR11 | 0.00 | -0.01 | 0.08 | 0.01 | 0.00 | -0.08 | 2.9E-01 | -0.05 |
| HDAC2 | -0.15 | -0.14 | 0.10 | -0.04 | 0.05 | 0.15 | 2.9E-01 | 0.12 |
| H3Y2 | 0.33 | -0.48 | 0.60 | -0.44 | -0.14 | -0.20 | 2.9E-01 | -0.41 |
| FDPS | 0.01 | 0.01 | -0.08 | 0.04 | 0.01 | 0.00 | 2.9E-01 | 0.04 |
| MMAB | 0.06 | -0.01 | -0.23 | 0.06 | 0.10 | 0.00 | 2.9E-01 | 0.11 |
| ZCCHC9 | 0.14 | 0.08 | -0.06 | 0.01 | 0.01 | -0.22 | 2.9E-01 | -0.12 |
| IMP3 | -0.04 | 0.04 | -0.08 | 0.01 | 0.06 | 0.00 | 2.9E-01 | 0.05 |
| ARHGEF12 | -0.08 | 0.06 | 0.12 | -0.05 | -0.04 | -0.02 | 2.9E-01 | -0.07 |
| RPS3A | -0.08 | -0.13 | 0.06 | 0.15 | 0.01 | -0.02 | 2.9E-01 | 0.09 |

| | | | | | | | | |
|---|---|---|---|---|---|---|---|---|
| CDK2AP2 | -0.10 | -0.17 | -0.06 | -0.14 | 0.00 | 0.39 | 2.9E-01 | 0.20 |
| KAT2B | -0.18 | 0.10 | -0.35 | 0.41 | -0.07 | -0.03 | 2.9E-01 | 0.25 |
| ATP2C1 | 0.04 | 0.03 | 0.02 | 0.06 | -0.04 | -0.12 | 2.9E-01 | -0.06 |
| PLGRKT | 0.07 | 0.02 | 0.01 | -0.02 | 0.04 | -0.13 | 2.9E-01 | -0.06 |
| P4HA1 | 0.03 | -0.03 | -0.10 | 0.08 | 0.01 | -0.01 | 2.9E-01 | 0.06 |
| PDCD11 | 0.06 | 0.03 | -0.02 | 0.02 | -0.01 | -0.08 | 2.9E-01 | -0.04 |
| UTP3 | -0.01 | -0.03 | 0.14 | 0.01 | -0.07 | -0.05 | 2.9E-01 | -0.07 |
| SLC5A6 | -0.16 | 0.11 | 0.33 | -0.31 | -0.07 | 0.03 | 2.9E-01 | -0.21 |
| HBS1L | 0.03 | 0.06 | -0.03 | 0.01 | 0.00 | -0.07 | 2.9E-01 | -0.04 |
| PXMP4 | 0.06 | -0.09 | -0.08 | 0.02 | -0.02 | 0.09 | 2.9E-01 | 0.07 |
| NEDD4 | -0.12 | 0.00 | -0.01 | 0.11 | 0.06 | -0.07 | 2.9E-01 | 0.08 |
| AURKB | -0.09 | 0.02 | -0.01 | -0.02 | 0.01 | 0.08 | 2.9E-01 | 0.05 |
| MIGA2 | -0.17 | -0.20 | 0.05 | 0.00 | -0.09 | 0.34 | 2.9E-01 | 0.18 |
| PIGK | 0.07 | 0.06 | 0.04 | 0.11 | -0.07 | -0.24 | 2.9E-01 | -0.12 |
| FGFR1OP2 | -0.12 | 0.15 | 0.10 | -0.06 | -0.06 | -0.04 | 2.9E-01 | -0.10 |
| COQ9 | 0.05 | -0.02 | -0.17 | 0.00 | 0.11 | 0.01 | 2.9E-01 | 0.09 |
| FAM98A | -0.04 | -0.01 | -0.04 | 0.06 | 0.08 | -0.06 | 2.9E-01 | 0.05 |
| KMT2D | 0.12 | -0.01 | 0.01 | -0.02 | 0.04 | -0.14 | 2.9E-01 | -0.08 |
| CIAO2A | 0.07 | -0.17 | -0.10 | 0.12 | -0.10 | 0.15 | 2.9E-01 | 0.13 |
| CIMAP1B | -0.26 | 0.04 | -0.20 | 0.12 | -0.22 | 0.40 | 3.0E-01 | 0.24 |
| MIS18A | 0.08 | -0.04 | 0.09 | 0.01 | -0.19 | 0.02 | 3.0E-01 | -0.10 |
| SPOP | 0.01 | 0.18 | 0.01 | -0.26 | 0.12 | -0.11 | 3.0E-01 | -0.15 |
| FDXR | 0.00 | 0.01 | -0.12 | -0.04 | 0.04 | 0.09 | 3.0E-01 | 0.07 |
| DDX49 | 0.08 | 0.02 | -0.03 | -0.06 | 0.03 | -0.04 | 3.0E-01 | -0.05 |
| RALY | 0.04 | 0.00 | 0.01 | -0.06 | -0.01 | 0.02 | 3.0E-01 | -0.03 |
| CRACR2B | -0.08 | -0.03 | 0.03 | 0.05 | -0.02 | 0.03 | 3.0E-01 | 0.05 |
| TLN2 | 0.04 | -0.02 | 0.03 | -0.02 | -0.04 | 0.02 | 3.0E-01 | -0.03 |
| GORASP2 | 0.06 | 0.04 | 0.04 | -0.23 | 0.03 | 0.04 | 3.0E-01 | -0.10 |
| RAP1B | 0.02 | 0.10 | 0.00 | -0.18 | 0.00 | 0.04 | 3.0E-01 | -0.09 |
| BET1L | 0.00 | 0.09 | 0.04 | 0.05 | 0.00 | -0.19 | 3.0E-01 | -0.09 |
| SLC37A1 | 0.05 | 0.17 | 0.02 | 0.18 | -0.29 | -0.19 | 3.0E-01 | -0.18 |
| CCDC6 | 0.03 | 0.01 | 0.02 | 0.02 | 0.00 | -0.08 | 3.0E-01 | -0.04 |
| TINF2 | 0.00 | -0.04 | -0.09 | -0.08 | 0.08 | 0.12 | 3.0E-01 | 0.08 |
| UBE2T | 0.08 | 0.17 | -0.03 | -0.32 | 0.06 | 0.00 | 3.0E-01 | -0.16 |
| SLC25A51 | 0.07 | -0.04 | 0.07 | -0.02 | 0.03 | -0.12 | 3.0E-01 | -0.07 |
| KIAA2013 | -0.07 | 0.10 | 0.07 | -0.01 | -0.01 | -0.09 | 3.0E-01 | -0.07 |
| FAM3A | -0.01 | -0.03 | 0.01 | 0.01 | 0.03 | -0.01 | 3.0E-01 | 0.02 |
| CSNK1D | 0.10 | 0.03 | -0.01 | 0.08 | -0.14 | -0.08 | 3.0E-01 | -0.09 |
| PHC3 | 0.01 | 0.00 | -0.05 | 0.00 | 0.00 | 0.04 | 3.0E-01 | 0.03 |
| USP12 | -0.01 | -0.03 | 0.21 | -0.20 | -0.01 | 0.01 | 3.0E-01 | -0.12 |
| DOCK5 | -0.16 | -0.33 | 0.12 | -0.11 | 0.06 | 0.32 | 3.0E-01 | 0.21 |
| RPS20 | 0.02 | -0.01 | 0.10 | 0.06 | -0.05 | -0.13 | 3.0E-01 | -0.07 |
| MDH2 | 0.17 | 0.11 | -0.13 | -0.08 | -0.06 | -0.04 | 3.0E-01 | -0.11 |
| CEP250 | -0.10 | -0.15 | 0.08 | 0.01 | -0.01 | 0.15 | 3.0E-01 | 0.10 |
| PHF20L1 | 0.01 | -0.01 | -0.10 | -0.02 | 0.01 | 0.10 | 3.0E-01 | 0.06 |
| SLC43A2 | -0.11 | 0.06 | 0.22 | -0.04 | -0.16 | 0.00 | 3.0E-01 | -0.13 |
| CCNY | -0.15 | 0.01 | -0.02 | -0.06 | 0.02 | 0.17 | 3.0E-01 | 0.10 |
| TERF2 | -0.03 | -0.08 | 0.02 | -0.04 | 0.04 | 0.08 | 3.0E-01 | 0.06 |

| | | | | | | | | |
|---|---|---|---|---|---|---|---|---|
| SLC18B1 | -0.11 | 0.01 | 0.30 | -0.09 | -0.10 | -0.05 | 3.0E-01 | -0.15 |
| MPPE1 | 0.38 | 0.17 | -0.23 | -0.44 | 0.01 | -0.05 | 3.0E-01 | -0.27 |
| ATF2 | 0.02 | 0.01 | 0.07 | -0.15 | 0.00 | 0.04 | 3.0E-01 | -0.07 |
| ANKRD20A4P | -0.05 | 0.07 | -0.13 | 0.04 | 0.04 | 0.02 | 3.0E-01 | 0.07 |
| SLC27A3 | -0.03 | 0.01 | 0.09 | -0.07 | -0.03 | 0.02 | 3.0E-01 | -0.05 |
| INO80B | 0.09 | 0.01 | 0.01 | -0.01 | -0.15 | 0.05 | 3.0E-01 | -0.08 |
| SRI | 0.00 | 0.02 | 0.09 | -0.02 | 0.05 | -0.16 | 3.0E-01 | -0.08 |
| JMJD1C | -0.12 | 0.01 | -0.04 | -0.09 | 0.07 | 0.15 | 3.0E-01 | 0.10 |
| PRPF4 | 0.15 | 0.00 | -0.03 | -0.02 | 0.01 | -0.12 | 3.0E-01 | -0.08 |
| ERCC2 | 0.04 | 0.10 | -0.01 | 0.05 | -0.02 | -0.18 | 3.0E-01 | -0.09 |
| NFATC3 | -0.02 | 0.05 | -0.30 | -0.03 | 0.01 | 0.24 | 3.0E-01 | 0.16 |
| HLTF | -0.04 | -0.03 | 0.03 | 0.01 | 0.01 | 0.02 | 3.0E-01 | 0.02 |
| CASK | -0.12 | -0.03 | 0.04 | 0.02 | -0.03 | 0.11 | 3.0E-01 | 0.07 |
| FAM120A | 0.01 | 0.00 | 0.06 | -0.04 | 0.03 | -0.05 | 3.0E-01 | -0.04 |
| UMAD1 | -0.07 | 0.02 | 0.18 | -0.07 | -0.08 | 0.00 | 3.0E-01 | -0.09 |
| SYNGR2 | -0.04 | -0.05 | 0.35 | -0.24 | -0.20 | 0.10 | 3.0E-01 | -0.20 |
| KRIT1 | 0.01 | -0.02 | 0.13 | -0.08 | 0.06 | -0.12 | 3.0E-01 | -0.08 |
| BTF3L4 | 0.20 | 0.10 | -0.10 | 0.06 | -0.06 | -0.23 | 3.0E-01 | -0.14 |
| VAC14 | -0.20 | -0.10 | 0.11 | 0.03 | 0.00 | 0.13 | 3.0E-01 | 0.11 |
| PHB1 | 0.00 | 0.01 | 0.02 | 0.02 | -0.02 | -0.04 | 3.0E-01 | -0.02 |
| ZNF444 | 0.13 | 0.05 | -0.08 | -0.05 | -0.05 | -0.02 | 3.0E-01 | -0.07 |
| EIF3K | 0.02 | -0.01 | 0.11 | -0.07 | 0.07 | -0.12 | 3.1E-01 | -0.08 |
| KDM4D | -0.15 | -0.38 | 0.15 | 0.07 | -0.08 | 0.29 | 3.1E-01 | 0.22 |
| CLN3 | -0.09 | 0.02 | 0.24 | -0.12 | -0.08 | 0.00 | 3.1E-01 | -0.12 |
| PYCARD | 0.16 | -0.07 | 0.11 | 0.12 | -0.24 | -0.13 | 3.1E-01 | -0.15 |
| ANKRD11 | -0.27 | -0.11 | 0.12 | 0.11 | -0.07 | 0.17 | 3.1E-01 | 0.16 |
| DNLZ | -0.17 | -0.14 | -0.13 | -0.21 | 0.50 | 0.01 | 3.1E-01 | 0.25 |
| AZGP1 | 0.01 | 0.04 | 0.02 | 0.03 | -0.09 | 0.00 | 3.1E-01 | -0.04 |
| TOP2B | 0.03 | -0.03 | -0.10 | -0.03 | 0.01 | 0.11 | 3.1E-01 | 0.07 |
| DCAF13 | -0.05 | -0.11 | 0.03 | -0.06 | 0.03 | 0.14 | 3.1E-01 | 0.08 |
| ABCA2 | -0.01 | -0.16 | 0.05 | -0.01 | 0.04 | 0.08 | 3.1E-01 | 0.08 |
| CMTR1 | -0.10 | 0.07 | 0.17 | 0.03 | -0.11 | -0.09 | 3.1E-01 | -0.10 |
| COQ5 | 0.12 | -0.14 | -0.18 | 0.13 | 0.05 | -0.01 | 3.1E-01 | 0.12 |
| ZNF346 | 0.05 | -0.04 | 0.07 | -0.07 | -0.04 | 0.03 | 3.1E-01 | -0.05 |
| NUP160 | -0.07 | -0.14 | 0.04 | -0.05 | 0.01 | 0.19 | 3.1E-01 | 0.10 |
| MAP1S | 0.06 | -0.02 | 0.04 | -0.02 | 0.04 | -0.10 | 3.1E-01 | -0.05 |
| WDR41 | -0.08 | -0.05 | -0.23 | 0.12 | -0.22 | 0.37 | 3.1E-01 | 0.21 |
| MAN1A2 | -0.02 | 0.02 | 0.11 | 0.07 | -0.10 | -0.09 | 3.1E-01 | -0.08 |
| NUDT21 | -0.02 | 0.01 | 0.10 | 0.04 | -0.05 | -0.08 | 3.1E-01 | -0.06 |
| PWP1 | 0.05 | 0.04 | 0.01 | 0.06 | -0.03 | -0.13 | 3.1E-01 | -0.07 |
| TTC9C | 0.12 | -0.14 | -0.31 | 0.28 | 0.07 | -0.10 | 3.1E-01 | 0.19 |
| NOP10 | -0.18 | 0.08 | 0.49 | 0.00 | 0.01 | -0.62 | 3.1E-01 | -0.33 |
| POLR1G | -0.01 | 0.03 | 0.05 | -0.02 | 0.03 | -0.09 | 3.1E-01 | -0.05 |
| TFE3 | -0.61 | -0.52 | 0.32 | 0.44 | -0.26 | 0.27 | 3.1E-01 | 0.42 |
| RAB27B | -0.05 | 0.01 | 0.12 | -0.04 | -0.03 | -0.02 | 3.1E-01 | -0.06 |
| EEF1E1 | -0.03 | -0.02 | -0.06 | 0.09 | 0.11 | -0.09 | 3.1E-01 | 0.07 |
| CD46 | -0.27 | -0.07 | 0.02 | 0.03 | -0.13 | 0.34 | 3.1E-01 | 0.19 |
| CLIC1 | 0.09 | 0.05 | -0.02 | -0.17 | 0.02 | 0.02 | 3.1E-01 | -0.08 |

| | | | | | | | | |
|---|---|---|---|---|---|---|---|---|
| MACF1 | -0.05 | 0.06 | 0.06 | 0.00 | 0.00 | -0.08 | 3.1E-01 | -0.05 |
| MLH1 | -0.08 | -0.06 | 0.06 | -0.01 | 0.06 | 0.03 | 3.1E-01 | 0.05 |
| MRM1 | 0.01 | 0.02 | 0.10 | -0.19 | 0.02 | 0.03 | 3.1E-01 | -0.09 |
| MYO5A | 0.14 | -0.11 | 0.12 | 0.04 | -0.06 | -0.15 | 3.1E-01 | -0.11 |
| BNIP1 | -0.03 | -0.11 | 0.01 | 0.06 | -0.07 | 0.12 | 3.1E-01 | 0.08 |
| RTN4IP1 | 0.02 | 0.00 | -0.10 | 0.01 | 0.06 | 0.00 | 3.1E-01 | 0.05 |
| HAGH | 0.05 | 0.12 | -0.06 | -0.14 | 0.04 | -0.03 | 3.1E-01 | -0.08 |
| ZFP64 | 0.07 | -0.12 | -0.16 | -0.11 | 0.10 | 0.19 | 3.1E-01 | 0.13 |
| TEX11 | -0.11 | 0.10 | 0.38 | -0.51 | -0.29 | 0.24 | 3.1E-01 | -0.30 |
| ZBED5 | 0.06 | 0.09 | 0.13 | 0.18 | -0.08 | -0.48 | 3.1E-01 | -0.22 |
| SNAP47 | -0.15 | -0.11 | 0.11 | 0.09 | 0.03 | 0.01 | 3.1E-01 | 0.09 |
| PRUNE1 | -0.44 | -0.10 | 0.15 | 0.16 | 0.22 | -0.09 | 3.1E-01 | 0.23 |
| GALNT4 | 0.21 | 0.06 | -0.09 | -0.14 | 0.08 | -0.15 | 3.1E-01 | -0.13 |
| ZNF787 | -0.07 | -0.10 | 0.05 | 0.03 | -0.05 | 0.13 | 3.2E-01 | 0.08 |
| MRRF | -0.03 | 0.09 | 0.03 | -0.09 | -0.08 | 0.06 | 3.2E-01 | -0.07 |
| PPP1R21 | 0.03 | 0.12 | 0.00 | -0.07 | -0.19 | 0.09 | 3.2E-01 | -0.10 |
| E2F3 | 0.09 | -0.01 | 0.05 | 0.07 | -0.19 | -0.03 | 3.2E-01 | -0.09 |
| ARPC5 | 0.07 | 0.14 | -0.10 | -0.03 | -0.07 | -0.03 | 3.2E-01 | -0.08 |
| RBMX2 | -0.02 | -0.04 | 0.17 | -0.02 | -0.11 | -0.01 | 3.2E-01 | -0.09 |
| NDUFA3 | 0.04 | -0.05 | -0.08 | 0.09 | 0.03 | -0.04 | 3.2E-01 | 0.06 |
| HAUS4 | -0.29 | -0.13 | 0.16 | 0.04 | 0.07 | 0.10 | 3.2E-01 | 0.15 |
| HIRA | 0.04 | -0.03 | -0.12 | -0.04 | 0.05 | 0.09 | 3.2E-01 | 0.07 |
| INTS10 | 0.06 | 0.05 | 0.01 | 0.04 | 0.01 | -0.18 | 3.2E-01 | -0.08 |
| PAWR | -0.02 | 0.15 | -0.02 | -0.03 | 0.01 | -0.11 | 3.2E-01 | -0.08 |
| SLC25A24 | -0.01 | -0.01 | 0.08 | -0.02 | -0.06 | 0.01 | 3.2E-01 | -0.04 |
| MCM10 | -0.06 | -0.21 | 0.06 | 0.08 | -0.08 | 0.18 | 3.2E-01 | 0.13 |
| GLT8D1 | 0.02 | 0.15 | 0.03 | 0.14 | -0.10 | -0.27 | 3.2E-01 | -0.14 |
| EHMT2 | 0.04 | 0.01 | 0.04 | -0.13 | 0.00 | 0.04 | 3.2E-01 | -0.06 |
| TRA2B | -0.01 | -0.05 | -0.01 | 0.05 | -0.05 | 0.07 | 3.2E-01 | 0.04 |
| PMS2 | -0.01 | 0.00 | -0.47 | -0.09 | 0.46 | -0.04 | 3.2E-01 | 0.27 |
| CHIC2 | 0.05 | -0.11 | 0.23 | -0.12 | 0.02 | -0.09 | 3.2E-01 | -0.12 |
| CTIF | -0.12 | -0.22 | 0.02 | -0.19 | 0.10 | 0.33 | 3.2E-01 | 0.19 |
| MRPL43 | 0.12 | 0.10 | -0.10 | -0.06 | 0.02 | -0.10 | 3.2E-01 | -0.09 |
| NACA | 0.12 | -0.02 | 0.04 | 0.07 | -0.03 | -0.21 | 3.2E-01 | -0.10 |
| ELAVL4 | 0.04 | -0.32 | 0.05 | 0.03 | 0.09 | 0.07 | 3.2E-01 | 0.14 |
| POLR2K | 0.01 | 0.13 | -0.02 | -0.17 | -0.01 | 0.04 | 3.2E-01 | -0.09 |
| LMAN2 | 0.03 | 0.02 | 0.01 | -0.02 | 0.04 | -0.09 | 3.2E-01 | -0.04 |
| GALK2 | -0.13 | -0.01 | -0.08 | -0.04 | 0.28 | -0.06 | 3.2E-01 | 0.13 |
| RPS19BP1 | 0.04 | 0.04 | -0.01 | 0.02 | 0.01 | -0.09 | 3.2E-01 | -0.04 |
| ASPH | -0.03 | -0.07 | 0.01 | 0.12 | -0.03 | -0.01 | 3.2E-01 | 0.06 |
| PBXIP1 | 0.01 | 0.03 | 0.09 | 0.06 | -0.18 | -0.02 | 3.2E-01 | -0.08 |
| KLHL22 | -0.16 | -0.03 | 0.01 | 0.03 | -0.08 | 0.19 | 3.2E-01 | 0.11 |
| ANAPC7 | -0.03 | -0.05 | -0.02 | -0.08 | 0.06 | 0.11 | 3.2E-01 | 0.06 |
| KDELR1 | 0.02 | 0.01 | -0.09 | 0.03 | 0.01 | 0.02 | 3.2E-01 | 0.04 |
| PFDN4 | 0.01 | 0.06 | 0.00 | 0.02 | -0.10 | 0.01 | 3.2E-01 | -0.05 |
| RPL22 | 0.06 | 0.02 | 0.02 | 0.05 | 0.00 | -0.16 | 3.2E-01 | -0.07 |
| ATP11C | -0.08 | 0.04 | 0.16 | -0.12 | -0.03 | 0.01 | 3.2E-01 | -0.09 |
| SYAP1 | -0.02 | 0.15 | -0.03 | -0.06 | -0.01 | -0.04 | 3.2E-01 | -0.07 |

| | | | | | | | | |
|---|---|---|---|---|---|---|---|---|
| COX7A2 | 0.01 | -0.02 | -0.10 | 0.13 | -0.01 | -0.02 | 3.2E-01 | 0.07 |
| GTF2H5 | 0.07 | 0.12 | 0.04 | 0.21 | -0.23 | -0.27 | 3.2E-01 | -0.18 |
| MARF1 | -0.08 | 0.16 | 0.09 | -0.06 | -0.22 | 0.08 | 3.2E-01 | -0.13 |
| SFMBT2 | 0.07 | -0.15 | 0.27 | -0.06 | -0.09 | -0.08 | 3.2E-01 | -0.14 |
| LYAR | 0.01 | 0.10 | -0.01 | 0.06 | -0.08 | -0.08 | 3.2E-01 | -0.06 |
| NEK2 | 0.12 | -0.15 | 0.18 | -0.05 | -0.08 | -0.06 | 3.2E-01 | -0.11 |
| PNO1 | 0.02 | -0.01 | 0.05 | -0.09 | 0.00 | 0.02 | 3.2E-01 | -0.04 |
| NME1 | 0.03 | -0.12 | -0.01 | 0.07 | 0.03 | -0.01 | 3.2E-01 | 0.06 |
| GCNT1 | 0.15 | -0.11 | 0.16 | -0.02 | -0.29 | 0.05 | 3.2E-01 | -0.15 |
| ATP6V1G1 | 0.14 | 0.18 | -0.13 | -0.01 | 0.02 | -0.24 | 3.3E-01 | -0.14 |
| POTED | -0.46 | -0.22 | -0.16 | 0.46 | -0.59 | 0.57 | 3.3E-01 | 0.43 |
| MCRS1 | 0.00 | -0.08 | -0.04 | 0.01 | -0.06 | 0.15 | 3.3E-01 | 0.07 |
| DBF4 | -0.20 | -0.18 | 0.10 | -0.13 | 0.25 | 0.10 | 3.3E-01 | 0.16 |
| AP1M1 | -0.03 | 0.00 | 0.14 | 0.00 | 0.01 | -0.12 | 3.3E-01 | -0.07 |
| HLF | 0.06 | -0.10 | 0.18 | -0.01 | -0.14 | -0.02 | 3.3E-01 | -0.10 |
| PPM1G | 0.12 | -0.18 | -0.13 | -0.04 | 0.07 | 0.13 | 3.3E-01 | 0.12 |
| PUS1 | -0.11 | -0.04 | 0.06 | 0.04 | -0.01 | 0.06 | 3.3E-01 | 0.06 |
| CHP1 | -0.13 | 0.02 | 0.30 | -0.06 | -0.17 | -0.03 | 3.3E-01 | -0.15 |
| CDC42BPG | -0.08 | -0.02 | 0.03 | 0.00 | 0.00 | 0.07 | 3.3E-01 | 0.04 |
| DLG3 | -0.08 | 0.02 | -0.02 | 0.06 | 0.05 | -0.04 | 3.3E-01 | 0.05 |
| FLYWCH2 | 0.04 | 0.15 | -0.08 | -0.06 | 0.00 | -0.06 | 3.3E-01 | -0.08 |
| GEMIN8 | 0.03 | 0.09 | -0.05 | 0.00 | -0.02 | -0.06 | 3.3E-01 | -0.05 |
| EPCAM | -0.02 | 0.01 | 0.15 | -0.17 | -0.09 | 0.09 | 3.3E-01 | -0.10 |
| ERGIC3 | 0.07 | 0.06 | -0.05 | -0.08 | -0.05 | 0.04 | 3.3E-01 | -0.06 |
| SLC2A10 | -0.15 | 0.15 | 0.20 | -0.20 | 0.06 | -0.11 | 3.3E-01 | -0.15 |
| ZBTB44 | 0.07 | -0.41 | 0.04 | 0.09 | 0.02 | 0.12 | 3.3E-01 | 0.18 |
| PRKACB | -0.17 | -0.08 | -0.05 | -0.20 | 0.10 | 0.33 | 3.3E-01 | 0.17 |
| PRSS8 | -0.25 | -0.35 | 0.24 | 0.10 | 0.00 | 0.16 | 3.3E-01 | 0.21 |
| SPEN | 0.01 | 0.00 | 0.04 | 0.01 | -0.07 | 0.01 | 3.3E-01 | -0.03 |
| C8orf33 | 0.00 | 0.01 | 0.05 | -0.04 | 0.04 | -0.05 | 3.3E-01 | -0.03 |
| METTL18 | -0.04 | 0.02 | -0.03 | 0.03 | -0.04 | 0.06 | 3.3E-01 | 0.04 |
| PIK3C3 | -0.52 | 0.13 | -0.05 | -0.11 | 0.11 | 0.31 | 3.3E-01 | 0.25 |
| RBL2 | -0.83 | -1.08 | 0.65 | 0.06 | 0.15 | 0.32 | 3.3E-01 | 0.60 |
| TMED2 | -0.13 | -0.02 | 0.02 | 0.15 | -0.01 | -0.03 | 3.3E-01 | 0.08 |
| ZNF92 | -0.22 | -0.15 | -0.01 | -0.07 | -0.13 | 0.46 | 3.3E-01 | 0.21 |
| ZNF654 | -0.06 | -0.01 | 0.00 | -0.05 | 0.08 | 0.03 | 3.3E-01 | 0.04 |
| EEF1G | 0.08 | 0.12 | -0.07 | -0.08 | 0.08 | -0.15 | 3.3E-01 | -0.10 |
| COPZ1 | -0.14 | -0.08 | 0.05 | 0.16 | 0.09 | -0.10 | 3.3E-01 | 0.11 |
| CNOT2 | 0.17 | 0.19 | -0.14 | -0.19 | 0.14 | -0.25 | 3.3E-01 | -0.18 |
| RASEF | -0.12 | -0.15 | 0.05 | 0.12 | -0.13 | 0.19 | 3.4E-01 | 0.13 |
| BAG1 | -0.06 | -0.02 | 0.03 | 0.01 | 0.00 | 0.04 | 3.4E-01 | 0.03 |
| TCEA1 | 0.06 | -0.01 | -0.01 | -0.02 | -0.01 | 0.00 | 3.4E-01 | -0.03 |
| KIAA1522 | -0.01 | -0.03 | 0.11 | -0.07 | 0.00 | -0.01 | 3.4E-01 | -0.05 |
| HDAC6 | 0.06 | -0.12 | -0.04 | 0.02 | 0.07 | -0.01 | 3.4E-01 | 0.06 |
| EZH2 | 0.05 | 0.02 | 0.00 | -0.11 | 0.02 | 0.02 | 3.4E-01 | -0.05 |
| SLC25A5 | 0.01 | 0.03 | -0.15 | 0.07 | 0.00 | 0.02 | 3.4E-01 | 0.07 |
| DHX34 | -0.06 | -0.07 | 0.02 | -0.04 | 0.00 | 0.14 | 3.4E-01 | 0.07 |
| TMC5 | -0.06 | 0.08 | 0.05 | -0.02 | 0.00 | -0.06 | 3.4E-01 | -0.05 |

| | | | | | | | | |
|---|---|---|---|---|---|---|---|---|
| IMMT | 0.00 | -0.03 | 0.01 | 0.03 | 0.00 | -0.01 | 3.4E-01 | 0.02 |
| AP3D1 | 0.05 | -0.02 | 0.03 | 0.01 | 0.00 | -0.06 | 3.4E-01 | -0.03 |
| GTF2IRD1 | -0.07 | -0.10 | 0.04 | 0.04 | -0.07 | 0.14 | 3.4E-01 | 0.08 |
| FUT8 | 0.05 | 0.06 | 0.07 | 0.13 | -0.05 | -0.29 | 3.4E-01 | -0.13 |
| CYSRT1 | -0.08 | -0.05 | -0.24 | 0.45 | -0.03 | -0.16 | 3.4E-01 | 0.21 |
| NDFIP2 | -0.02 | -0.02 | 0.26 | -0.34 | -0.03 | 0.08 | 3.4E-01 | -0.17 |
| RPL10A | 0.00 | -0.01 | 0.09 | 0.05 | -0.03 | -0.11 | 3.4E-01 | -0.06 |
| MIS12 | -0.05 | -0.06 | -0.08 | -0.12 | 0.03 | 0.25 | 3.4E-01 | 0.12 |
| NDUFA12 | 0.07 | 0.01 | 0.04 | 0.10 | -0.08 | -0.15 | 3.4E-01 | -0.08 |
| MARVELD2 | -0.19 | -0.03 | 0.07 | 0.07 | -0.03 | 0.09 | 3.4E-01 | 0.09 |
| POLR2F | 0.12 | 0.03 | -0.06 | -0.01 | -0.06 | -0.02 | 3.4E-01 | -0.06 |
| CCT6A | 0.10 | 0.08 | -0.07 | -0.07 | 0.05 | -0.11 | 3.4E-01 | -0.08 |
| PLPBP | -0.08 | 0.01 | 0.02 | 0.00 | 0.02 | 0.03 | 3.4E-01 | 0.03 |
| SIN3A | 0.01 | -0.03 | -0.08 | 0.13 | -0.06 | 0.02 | 3.4E-01 | 0.06 |
| NDUFA4 | -0.01 | -0.03 | -0.04 | 0.12 | -0.05 | 0.02 | 3.4E-01 | 0.05 |
| PEX5 | 0.01 | 0.06 | 0.04 | 0.05 | 0.01 | -0.18 | 3.4E-01 | -0.08 |
| PLCH1 | 0.17 | -0.28 | 0.46 | -0.31 | -0.43 | 0.17 | 3.4E-01 | -0.30 |
| TSPAN13 | -0.03 | -0.12 | 0.39 | -0.28 | -0.08 | 0.02 | 3.4E-01 | -0.20 |
| ABCC3 | -0.29 | 0.00 | 0.08 | 0.08 | -0.02 | 0.12 | 3.4E-01 | 0.13 |
| SHANK1 | -0.83 | -0.20 | 1.38 | -1.98 | 0.13 | -0.70 | 3.4E-01 | -0.97 |
| KANSL2 | -0.23 | 0.38 | 0.11 | -0.20 | 0.04 | -0.20 | 3.4E-01 | -0.21 |
| STRN | 0.00 | 0.06 | 0.12 | -0.03 | 0.11 | -0.31 | 3.4E-01 | -0.14 |
| MYT1 | -0.28 | 0.08 | 0.00 | 0.04 | 0.06 | 0.06 | 3.4E-01 | 0.12 |
| PRAF2 | -0.15 | -0.05 | 0.01 | -0.13 | 0.09 | 0.21 | 3.4E-01 | 0.12 |
| MEN1 | 0.03 | -0.07 | -0.06 | 0.04 | -0.06 | 0.10 | 3.4E-01 | 0.06 |
| TUT7 | -0.04 | 0.20 | 0.01 | -0.13 | 0.11 | -0.18 | 3.4E-01 | -0.12 |
| KTN1 | -0.06 | 0.06 | 0.09 | 0.00 | 0.00 | -0.09 | 3.5E-01 | -0.06 |
| EFHD1 | -0.08 | 0.04 | 0.21 | 0.07 | -0.19 | -0.07 | 3.5E-01 | -0.12 |
| SNRNP27 | 0.26 | 0.14 | -0.17 | 0.07 | -0.10 | -0.27 | 3.5E-01 | -0.17 |
| PPP1R12C | -0.17 | -0.01 | 0.05 | 0.10 | -0.01 | 0.02 | 3.5E-01 | 0.08 |
| HCFC2 | 0.10 | -0.06 | 0.04 | -0.10 | 0.00 | 0.01 | 3.5E-01 | -0.06 |
| CLOCK | -0.06 | -0.03 | 0.04 | -0.01 | 0.01 | 0.06 | 3.5E-01 | 0.04 |
| PPID | 0.01 | 0.01 | -0.07 | 0.03 | 0.02 | -0.01 | 3.5E-01 | 0.03 |
| KIFBP | -0.02 | -0.14 | 0.00 | 0.18 | 0.05 | -0.10 | 3.5E-01 | 0.10 |
| RPL7A | 0.02 | -0.05 | -0.03 | 0.00 | 0.08 | -0.02 | 3.5E-01 | 0.04 |
| MTA1 | -0.03 | -0.05 | -0.04 | -0.10 | 0.06 | 0.14 | 3.5E-01 | 0.08 |
| EPHA2 | -0.05 | 0.05 | 0.14 | 0.11 | -0.09 | -0.20 | 3.5E-01 | -0.11 |
| SUPT3H | -0.24 | 0.31 | 0.18 | -0.23 | -0.11 | 0.01 | 3.5E-01 | -0.19 |
| NUDT6 | 0.06 | 0.11 | 0.05 | -0.27 | 0.22 | -0.23 | 3.5E-01 | -0.17 |
| CINP | 0.13 | 0.19 | -0.12 | -0.26 | 0.11 | -0.11 | 3.5E-01 | -0.15 |
| CDK12 | 0.03 | 0.01 | 0.02 | 0.03 | -0.01 | -0.08 | 3.5E-01 | -0.04 |
| LARP7 | -0.02 | 0.00 | 0.09 | -0.02 | 0.03 | -0.09 | 3.5E-01 | -0.05 |
| EEFSEC | 0.01 | 0.05 | 0.00 | 0.03 | -0.01 | -0.09 | 3.5E-01 | -0.04 |
| RAD21 | -0.01 | -0.05 | 0.01 | -0.03 | 0.03 | 0.05 | 3.5E-01 | 0.03 |
| SNX8 | -0.04 | -0.03 | -0.06 | 0.06 | 0.17 | -0.11 | 3.5E-01 | 0.09 |
| PRDX5 | 0.04 | -0.05 | -0.22 | -0.14 | 0.22 | 0.11 | 3.5E-01 | 0.14 |
| EIF3I | -0.14 | -0.11 | 0.11 | 0.08 | -0.02 | 0.07 | 3.5E-01 | 0.09 |
| ELOVL1 | 0.26 | -0.43 | 0.54 | -0.41 | 0.10 | -0.36 | 3.5E-01 | -0.35 |

| | | | | | | | | |
|---|---|---|---|---|---|---|---|---|
| **DCP2** | -0.23 | 0.04 | 0.01 | -0.06 | 0.05 | 0.16 | 3.5E-01 | 0.11 |
| **USP7** | -0.01 | -0.02 | 0.13 | 0.04 | -0.01 | -0.14 | 3.5E-01 | -0.07 |
| **RREB1** | 0.02 | 0.01 | 0.02 | 0.01 | -0.09 | 0.02 | 3.5E-01 | -0.04 |
| **LRSAM1** | 0.05 | -0.03 | 0.11 | -0.19 | 0.04 | 0.01 | 3.5E-01 | -0.09 |
| **S100A14** | -0.11 | -0.05 | 0.07 | 0.02 | 0.00 | 0.07 | 3.5E-01 | 0.06 |
| **NGDN** | 0.03 | -0.02 | 0.04 | 0.04 | -0.03 | -0.07 | 3.5E-01 | -0.04 |
| **FNDC3B** | 0.06 | 0.06 | 0.01 | 0.11 | -0.07 | -0.18 | 3.5E-01 | -0.09 |
| **DNAJC21** | -0.11 | 0.12 | 0.11 | -0.05 | -0.10 | 0.01 | 3.5E-01 | -0.09 |
| **YWHAG** | 0.00 | -0.01 | 0.07 | -0.02 | 0.04 | -0.09 | 3.5E-01 | -0.05 |
| **FDX1** | -0.06 | -0.09 | -0.04 | 0.26 | -0.06 | -0.05 | 3.5E-01 | 0.11 |
| **TRIM37** | -0.05 | -0.12 | 0.06 | 0.00 | -0.02 | 0.12 | 3.5E-01 | 0.07 |
| **SLC25A36** | -0.08 | 0.14 | 0.11 | -0.18 | 0.12 | -0.14 | 3.5E-01 | -0.12 |
| **MCUR1** | 0.08 | 0.15 | -0.06 | -0.02 | 0.07 | -0.26 | 3.5E-01 | -0.12 |
| **ANKRD50** | 0.00 | -0.07 | 0.25 | -0.20 | 0.10 | -0.14 | 3.5E-01 | -0.14 |
| **TRA2A** | 0.02 | -0.14 | -0.21 | 0.40 | -0.01 | -0.15 | 3.5E-01 | 0.19 |
| **POGZ** | 0.04 | 0.02 | 0.02 | -0.11 | -0.05 | 0.07 | 3.5E-01 | -0.06 |
| **BIN3** | -0.04 | -0.02 | -0.03 | 0.14 | -0.06 | 0.00 | 3.6E-01 | 0.06 |
| **PLK1** | -0.04 | -0.04 | 0.04 | -0.02 | 0.03 | 0.05 | 3.6E-01 | 0.03 |
| **N4BP3** | -0.28 | -0.10 | 0.02 | -0.24 | 0.38 | 0.12 | 3.6E-01 | 0.21 |
| **PSMA4** | 0.18 | -0.04 | 0.01 | -0.20 | 0.05 | -0.02 | 3.6E-01 | -0.10 |
| **MALSU1** | -0.08 | 0.06 | -0.25 | -0.18 | 0.18 | 0.22 | 3.6E-01 | 0.16 |
| **DTL** | -0.12 | -0.21 | 0.14 | 0.11 | 0.08 | -0.03 | 3.6E-01 | 0.12 |
| **RCN1** | 0.05 | 0.17 | -0.10 | 0.01 | -0.03 | -0.14 | 3.6E-01 | -0.09 |
| **WDR77** | -0.13 | -0.23 | 0.08 | -0.17 | 0.25 | 0.12 | 3.6E-01 | 0.16 |
| **AP1G2** | -0.13 | -0.12 | 0.05 | 0.26 | -0.05 | -0.04 | 3.6E-01 | 0.12 |
| **VAMP5** | -0.15 | 0.08 | 0.34 | -0.42 | 0.11 | -0.07 | 3.6E-01 | -0.22 |
| **SNRPD1** | 0.03 | -0.01 | 0.04 | -0.05 | -0.05 | 0.04 | 3.6E-01 | -0.04 |
| **CCNL2** | 0.14 | 0.06 | -0.05 | -0.13 | 0.12 | -0.17 | 3.6E-01 | -0.11 |
| **TIMM23** | 0.10 | 0.00 | 0.08 | 0.15 | -0.28 | -0.10 | 3.6E-01 | -0.13 |
| **GNAS** | -0.06 | 0.05 | 0.08 | -0.06 | -0.01 | 0.00 | 3.6E-01 | -0.05 |
| **OTUD3** | -0.18 | -0.06 | 0.06 | -0.09 | 0.06 | 0.17 | 3.6E-01 | 0.11 |
| **C6orf132** | -0.01 | 0.12 | -0.01 | -0.11 | 0.04 | -0.03 | 3.6E-01 | -0.06 |
| **SPRYD4** | 0.09 | -0.07 | -0.12 | 0.03 | 0.04 | 0.03 | 3.6E-01 | 0.06 |
| **DBR1** | 0.05 | 0.04 | 0.00 | -0.05 | 0.08 | -0.13 | 3.6E-01 | -0.06 |
| **MTX2** | -0.01 | -0.01 | 0.09 | 0.01 | 0.01 | -0.10 | 3.6E-01 | -0.05 |
| **ANO8** | -0.12 | -0.02 | 0.37 | -0.28 | -0.06 | 0.02 | 3.6E-01 | -0.18 |
| **RPL35A** | 0.03 | -0.03 | 0.07 | -0.01 | 0.03 | -0.09 | 3.6E-01 | -0.05 |
| **RETREG3** | -0.11 | 0.03 | -0.07 | -0.11 | 0.14 | 0.09 | 3.6E-01 | 0.09 |
| **PRSS23** | 0.05 | 0.09 | -0.04 | -0.07 | 0.08 | -0.12 | 3.6E-01 | -0.07 |
| **THOP1** | -0.16 | -0.03 | -0.15 | -0.29 | 0.20 | 0.33 | 3.6E-01 | 0.20 |
| **C19orf53** | 0.05 | 0.06 | -0.03 | 0.03 | 0.00 | -0.12 | 3.6E-01 | -0.05 |
| **KIFAP3** | 0.04 | -0.33 | 0.00 | -0.07 | 0.31 | -0.01 | 3.6E-01 | 0.17 |
| **PTPRA** | 0.00 | 0.00 | 0.14 | 0.05 | -0.24 | 0.02 | 3.6E-01 | -0.11 |
| **ESRRA** | 0.03 | -0.35 | 0.05 | 0.24 | 0.03 | -0.05 | 3.6E-01 | 0.16 |
| **ORC2** | -0.04 | -0.04 | 0.00 | -0.01 | -0.03 | 0.11 | 3.7E-01 | 0.05 |
| **ADGRG1** | -0.08 | -0.08 | 0.35 | -0.05 | -0.14 | -0.07 | 3.7E-01 | -0.15 |
| **ARL15** | -0.11 | 0.03 | 0.25 | -0.11 | -0.16 | 0.06 | 3.7E-01 | -0.13 |
| **LETM2** | 0.30 | -0.35 | 0.48 | -0.94 | 0.21 | -0.13 | 3.7E-01 | -0.43 |

| | | | | | | | | |
|---|---|---|---|---|---|---|---|---|
| **ACAA1** | -0.01 | 0.00 | -0.04 | -0.05 | 0.04 | 0.05 | 3.7E-01 | 0.03 |
| **GPAT4** | 0.09 | 0.14 | 0.05 | -0.10 | 0.24 | -0.54 | 3.7E-01 | -0.23 |
| **KRT9** | -0.19 | -0.10 | -0.46 | 0.57 | -0.57 | 0.38 | 3.7E-01 | 0.38 |
| **PDXK** | -0.16 | 0.16 | 0.18 | -0.06 | 0.05 | -0.21 | 3.7E-01 | -0.13 |
| **TMEM63A** | -0.08 | -0.06 | 0.02 | -0.07 | 0.02 | 0.15 | 3.7E-01 | 0.07 |
| **PWWP2A** | -0.02 | -0.03 | -0.05 | -0.09 | 0.05 | 0.12 | 3.7E-01 | 0.06 |
| **POLR2H** | -0.06 | -0.12 | 0.01 | -0.01 | -0.08 | 0.24 | 3.7E-01 | 0.10 |
| **RBM4** | -0.01 | 0.05 | 0.08 | 0.11 | -0.14 | -0.10 | 3.7E-01 | -0.08 |
| **PRR36** | -0.12 | 0.10 | 0.22 | -0.12 | -0.25 | 0.12 | 3.7E-01 | -0.15 |
| **MTF2** | -0.05 | -0.05 | -0.03 | -0.07 | 0.00 | 0.19 | 3.7E-01 | 0.08 |
| **LSG1** | -0.05 | 0.05 | 0.08 | 0.01 | -0.09 | 0.00 | 3.7E-01 | -0.05 |
| **COQ2** | -0.01 | 0.05 | -0.19 | 0.14 | -0.05 | 0.03 | 3.7E-01 | 0.09 |
| **PI4K2A** | -0.08 | -0.17 | 0.04 | -0.11 | 0.02 | 0.26 | 3.7E-01 | 0.12 |
| **KEAP1** | -0.12 | -0.05 | 0.39 | -0.21 | 0.03 | -0.12 | 3.7E-01 | -0.18 |
| **SMARCA4** | 0.01 | -0.02 | -0.05 | -0.03 | -0.01 | 0.09 | 3.7E-01 | 0.04 |
| **ACAP1** | -0.04 | 0.04 | -0.31 | -0.18 | 0.10 | 0.31 | 3.7E-01 | 0.18 |
| **INO80D** | 0.02 | 0.03 | -0.14 | 0.00 | 0.05 | 0.04 | 3.7E-01 | 0.06 |
| **CLCN7** | -0.04 | 0.04 | 0.06 | 0.04 | -0.09 | -0.03 | 3.7E-01 | -0.05 |
| **PAK4** | -0.16 | -0.13 | 0.12 | 0.04 | -0.03 | 0.13 | 3.7E-01 | 0.10 |
| **ASS1** | 0.05 | 0.07 | -0.04 | 0.01 | 0.02 | -0.12 | 3.7E-01 | -0.06 |
| **NOA1** | 0.10 | 0.05 | -0.03 | -0.19 | -0.01 | 0.07 | 3.7E-01 | -0.09 |
| **HDLBP** | -0.05 | -0.07 | 0.03 | 0.10 | 0.06 | -0.07 | 3.7E-01 | 0.06 |
| **SHB** | 0.02 | -0.18 | 0.39 | -0.22 | 0.00 | -0.10 | 3.7E-01 | -0.18 |
| **UBA3** | 0.04 | -0.13 | -0.08 | 0.22 | -0.03 | -0.06 | 3.7E-01 | 0.10 |
| **TRAM1L1** | 0.10 | 0.10 | -0.06 | -0.15 | 0.13 | -0.15 | 3.7E-01 | -0.11 |
| **PSMD7** | -0.04 | -0.15 | -0.23 | -0.36 | 0.35 | 0.28 | 3.8E-01 | 0.23 |
| **STRAP** | 0.14 | 0.02 | -0.02 | 0.12 | -0.11 | -0.19 | 3.8E-01 | -0.10 |
| **LRRC59** | -0.02 | 0.00 | 0.00 | -0.02 | 0.02 | 0.03 | 3.8E-01 | 0.02 |
| **NDUFB11** | 0.01 | 0.07 | -0.02 | 0.05 | -0.05 | -0.07 | 3.8E-01 | -0.04 |
| **KIF13B** | -0.22 | -0.33 | 0.15 | 0.41 | -0.15 | 0.01 | 3.8E-01 | 0.22 |
| **CCDC50** | 0.06 | 0.08 | -0.03 | -0.15 | 0.09 | -0.06 | 3.8E-01 | -0.08 |
| **RAB29** | 0.04 | -0.05 | 0.07 | -0.03 | -0.03 | 0.00 | 3.8E-01 | -0.04 |
| **C2CD3** | -0.21 | 0.01 | 0.48 | -0.40 | 0.05 | -0.09 | 3.8E-01 | -0.24 |
| **TRNAU1AP** | -0.27 | -0.10 | 0.14 | -0.02 | 0.01 | 0.19 | 3.8E-01 | 0.14 |
| **EIF4A3** | 0.04 | -0.05 | -0.03 | 0.00 | 0.00 | 0.04 | 3.8E-01 | 0.03 |
| **SYNCRIP** | 0.00 | -0.07 | 0.02 | 0.04 | -0.02 | 0.03 | 3.8E-01 | 0.03 |
| **SLIRP** | -0.03 | -0.05 | -0.07 | -0.04 | -0.06 | 0.23 | 3.8E-01 | 0.09 |
| **RAN** | 0.03 | -0.02 | -0.04 | 0.00 | 0.00 | 0.03 | 3.8E-01 | 0.02 |
| **WDR46** | 0.02 | 0.00 | 0.08 | 0.10 | -0.09 | -0.13 | 3.8E-01 | -0.07 |
| **DAZL** | -0.01 | -0.12 | -0.22 | 0.03 | -0.22 | 0.44 | 3.8E-01 | 0.20 |
| **CEP295** | -0.01 | -0.21 | 0.00 | -0.11 | 0.01 | 0.28 | 3.8E-01 | 0.13 |
| **HNRNPA0** | 0.01 | -0.01 | -0.05 | -0.04 | 0.02 | 0.07 | 3.8E-01 | 0.03 |
| **MESD** | 0.19 | 0.14 | -0.17 | 0.00 | -0.08 | -0.11 | 3.8E-01 | -0.11 |
| **STK33** | 0.02 | -0.06 | -0.08 | 0.08 | 0.14 | -0.11 | 3.8E-01 | 0.08 |
| **RNF170** | -0.06 | 0.04 | 0.14 | 0.07 | -0.11 | -0.09 | 3.8E-01 | -0.08 |
| **CRIP1** | 0.31 | 0.00 | -0.08 | 0.14 | -0.12 | -0.32 | 3.8E-01 | -0.18 |
| **IGF1R** | -0.03 | 0.02 | 0.08 | -0.13 | -0.02 | 0.05 | 3.8E-01 | -0.06 |
| **MYDGF** | -0.02 | -0.04 | -0.15 | 0.07 | -0.16 | 0.26 | 3.8E-01 | 0.13 |

| | | | | | | | | |
|---|---|---|---|---|---|---|---|---|
| TAPT1 | 0.02 | 0.02 | 0.08 | 0.08 | 0.00 | -0.23 | 3.8E-01 | -0.09 |
| CRNKL1 | 0.07 | 0.03 | -0.04 | 0.00 | 0.00 | -0.05 | 3.8E-01 | -0.04 |
| SQLE | -0.05 | 0.02 | -0.07 | 0.02 | 0.12 | -0.06 | 3.8E-01 | 0.06 |
| PLPP6 | -0.12 | 0.13 | 0.12 | -0.09 | -0.13 | 0.06 | 3.8E-01 | -0.10 |
| SBNO2 | 0.06 | 0.01 | 0.19 | 0.12 | 0.05 | -0.54 | 3.8E-01 | -0.21 |
| WDR43 | 0.03 | 0.02 | -0.01 | 0.00 | 0.02 | -0.06 | 3.8E-01 | -0.03 |
| OGT | 0.03 | 0.01 | 0.07 | 0.05 | -0.20 | 0.02 | 3.8E-01 | -0.08 |
| YWHAZ | 0.06 | 0.09 | -0.05 | 0.04 | 0.01 | -0.16 | 3.8E-01 | -0.07 |
| ELF2 | 0.03 | -0.01 | 0.11 | -0.20 | -0.10 | 0.13 | 3.8E-01 | -0.10 |
| CLASP2 | -0.01 | -0.02 | 0.10 | -0.02 | -0.08 | 0.03 | 3.8E-01 | -0.05 |
| ELMOD2 | 0.12 | -0.20 | -0.07 | 0.02 | 0.03 | 0.07 | 3.8E-01 | 0.09 |
| RPL13 | 0.00 | 0.03 | 0.05 | 0.01 | 0.05 | -0.15 | 3.8E-01 | -0.06 |
| P3H1 | -0.06 | 0.01 | 0.01 | 0.01 | -0.01 | 0.04 | 3.8E-01 | 0.03 |
| H3C1 | 0.04 | -0.07 | -0.04 | -0.03 | 0.08 | 0.01 | 3.8E-01 | 0.04 |
| WASHC2A | 0.08 | 0.12 | -0.02 | -0.19 | 0.19 | -0.23 | 3.8E-01 | -0.14 |
| PFN1 | 0.12 | 0.12 | -0.12 | -0.11 | 0.05 | -0.08 | 3.8E-01 | -0.09 |
| NDUFC2 | 0.03 | 0.05 | -0.01 | 0.05 | -0.02 | -0.11 | 3.8E-01 | -0.05 |
| ORC6 | -0.13 | -0.15 | 0.10 | -0.11 | 0.13 | 0.12 | 3.8E-01 | 0.11 |
| RPS10 | -0.05 | -0.09 | 0.04 | 0.08 | 0.08 | -0.07 | 3.9E-01 | 0.06 |
| FARS2 | -0.03 | 0.03 | -0.11 | 0.03 | -0.06 | 0.13 | 3.9E-01 | 0.07 |
| KPNA2 | 0.00 | -0.06 | 0.15 | -0.02 | -0.07 | -0.01 | 3.9E-01 | -0.06 |
| METTL16 | 0.00 | -0.09 | -0.01 | -0.02 | 0.14 | -0.03 | 3.9E-01 | 0.06 |
| SEPTIN6 | 0.25 | 0.19 | -0.18 | -0.42 | 0.15 | -0.10 | 3.9E-01 | -0.21 |
| RPL27A | 0.00 | -0.06 | 0.15 | -0.05 | 0.00 | -0.06 | 3.9E-01 | -0.06 |
| UTP23 | 0.14 | 0.13 | -0.13 | -0.01 | 0.02 | -0.17 | 3.9E-01 | -0.10 |
| NSFL1C | -0.02 | 0.05 | -0.11 | 0.06 | -0.02 | 0.03 | 3.9E-01 | 0.05 |
| LSM2 | 0.05 | -0.04 | -0.11 | -0.01 | -0.02 | 0.12 | 3.9E-01 | 0.06 |
| CTPS2 | -0.01 | -0.11 | 0.03 | -0.01 | 0.11 | -0.02 | 3.9E-01 | 0.06 |
| OSBPL6 | -0.17 | 0.13 | 0.18 | -0.05 | -0.06 | -0.06 | 3.9E-01 | -0.10 |
| HINT2 | 0.08 | 0.00 | -0.25 | 0.09 | 0.01 | 0.04 | 3.9E-01 | 0.10 |
| ASCC1 | 0.07 | 0.03 | 0.05 | 0.17 | -0.13 | -0.23 | 3.9E-01 | -0.12 |
| ZGPAT | -0.10 | -0.03 | 0.02 | -0.02 | -0.04 | 0.16 | 3.9E-01 | 0.07 |
| RBM22 | 0.09 | -0.03 | 0.02 | 0.06 | -0.11 | -0.04 | 3.9E-01 | -0.06 |
| INO80E | 0.03 | -0.03 | 0.12 | 0.09 | -0.16 | -0.06 | 3.9E-01 | -0.08 |
| RAB33B | -0.07 | 0.08 | 0.07 | -0.02 | 0.01 | -0.07 | 3.9E-01 | -0.05 |
| LSM4 | 0.09 | -0.12 | 0.16 | -0.09 | -0.12 | 0.05 | 3.9E-01 | -0.10 |
| SMC5 | -0.04 | -0.06 | 0.03 | 0.05 | -0.06 | 0.07 | 3.9E-01 | 0.05 |
| SEC61A2 | -0.06 | -0.06 | 0.28 | 0.02 | -0.12 | -0.10 | 3.9E-01 | -0.12 |
| CIR1 | 0.25 | 0.12 | -0.19 | 0.04 | -0.11 | -0.16 | 3.9E-01 | -0.14 |
| MRPS21 | 0.10 | 0.15 | -0.09 | -0.30 | 0.05 | 0.03 | 3.9E-01 | -0.13 |
| DEAF1 | -0.01 | -0.03 | -0.07 | -0.11 | 0.15 | 0.06 | 3.9E-01 | 0.07 |
| PMFBP1 | 0.13 | 0.05 | -0.04 | -0.25 | 0.06 | 0.02 | 3.9E-01 | -0.10 |
| CHCHD6 | 0.02 | 0.06 | 0.01 | 0.02 | 0.04 | -0.15 | 3.9E-01 | -0.06 |
| GPRC5C | -0.01 | 0.02 | -0.07 | 0.05 | 0.04 | -0.04 | 3.9E-01 | 0.04 |
| CDC42BPB | -0.26 | -0.23 | 0.22 | 0.05 | -0.01 | 0.17 | 3.9E-01 | 0.16 |
| CNP | -0.01 | 0.03 | -0.08 | 0.05 | 0.02 | -0.02 | 3.9E-01 | 0.04 |
| BCL7C | 0.04 | 0.07 | -0.04 | -0.08 | 0.05 | -0.04 | 4.0E-01 | -0.05 |
| CC2D1B | -0.02 | -0.03 | 0.16 | -0.15 | 0.03 | -0.01 | 4.0E-01 | -0.08 |

| | | | | | | | | |
|---|---|---|---|---|---|---|---|---|
| MFSD1 | -0.07 | -0.11 | 0.40 | -0.08 | -0.24 | 0.02 | 4.0E-01 | -0.17 |
| PPME1 | 0.12 | 0.20 | -0.15 | -0.15 | 0.11 | -0.18 | 4.0E-01 | -0.13 |
| ZBTB21 | -0.03 | 0.08 | -0.20 | 0.16 | -0.05 | 0.02 | 4.0E-01 | 0.10 |
| MTA2 | 0.00 | -0.04 | -0.02 | -0.07 | 0.06 | 0.06 | 4.0E-01 | 0.04 |
| OAF | 0.01 | 0.02 | 0.15 | -0.33 | 0.13 | -0.02 | 4.0E-01 | -0.13 |
| INTS15 | -0.03 | -0.19 | 0.06 | 0.00 | -0.06 | 0.19 | 4.0E-01 | 0.10 |
| RPL19 | -0.02 | 0.01 | 0.11 | 0.04 | 0.00 | -0.14 | 4.0E-01 | -0.06 |
| INIP | 0.11 | -0.01 | -0.03 | -0.08 | 0.05 | -0.06 | 4.0E-01 | -0.06 |
| THYN1 | -0.09 | 0.02 | -0.02 | -0.04 | -0.01 | 0.14 | 4.0E-01 | 0.06 |
| ERP44 | 0.00 | 0.00 | -0.04 | -0.01 | 0.07 | -0.01 | 4.0E-01 | 0.03 |
| MRPS30 | 0.01 | -0.06 | 0.00 | 0.07 | 0.00 | -0.02 | 4.0E-01 | 0.03 |
| RPL34 | 0.00 | -0.08 | -0.02 | 0.16 | -0.02 | -0.05 | 4.0E-01 | 0.07 |
| L1RE1 | -0.13 | 0.31 | 0.03 | 0.04 | 0.00 | -0.33 | 4.0E-01 | -0.17 |
| NME3 | 0.02 | -0.07 | 0.00 | -0.03 | 0.05 | 0.02 | 4.0E-01 | 0.03 |
| MTX3 | -0.10 | -0.09 | 0.08 | 0.10 | -0.06 | 0.06 | 4.0E-01 | 0.07 |
| CDK16 | -0.21 | -0.17 | 0.17 | -0.04 | 0.10 | 0.10 | 4.0E-01 | 0.12 |
| TXNDC9 | -0.53 | -0.12 | 0.09 | 0.35 | 0.38 | -0.41 | 4.0E-01 | 0.30 |
| RCC2 | 0.05 | -0.01 | -0.15 | -0.04 | 0.05 | 0.09 | 4.0E-01 | 0.07 |
| FAM114A2 | -0.08 | -0.18 | 0.12 | 0.02 | 0.10 | 0.01 | 4.0E-01 | 0.09 |
| CHSY1 | 0.02 | 0.01 | 0.08 | 0.12 | -0.07 | -0.18 | 4.0E-01 | -0.08 |
| HCAR1 | -0.16 | 0.06 | 0.28 | -0.23 | -0.06 | 0.05 | 4.0E-01 | -0.14 |
| EPHB4 | -0.14 | 0.20 | 0.09 | -0.12 | 0.06 | -0.11 | 4.0E-01 | -0.11 |
| CCNK | 0.00 | -0.01 | -0.07 | 0.10 | 0.02 | -0.05 | 4.0E-01 | 0.05 |
| CRBN | 0.03 | -0.03 | -0.16 | -0.11 | 0.15 | 0.08 | 4.0E-01 | 0.09 |
| EBNA1BP2 | 0.02 | -0.02 | 0.03 | 0.03 | -0.03 | -0.04 | 4.0E-01 | -0.02 |
| BMAL1 | -0.07 | -0.04 | 0.05 | 0.01 | -0.02 | 0.06 | 4.0E-01 | 0.04 |
| F8A1 | -0.17 | 0.15 | -0.19 | -0.02 | -0.01 | 0.19 | 4.0E-01 | 0.12 |
| SYBU | 0.15 | 0.07 | -0.11 | -0.02 | -0.01 | -0.09 | 4.0E-01 | -0.08 |
| NXT2 | 0.13 | 0.00 | 0.01 | 0.16 | -0.14 | -0.20 | 4.0E-01 | -0.11 |
| VTCN1 | -0.13 | 0.11 | 0.14 | -0.11 | 0.02 | -0.05 | 4.0E-01 | -0.09 |
| ROGDI | -0.22 | -0.13 | 0.11 | 0.26 | 0.05 | -0.13 | 4.0E-01 | 0.14 |
| WNK1 | 0.06 | 0.09 | -0.03 | -0.11 | 0.13 | -0.18 | 4.0E-01 | -0.09 |
| SUPT7L | 0.02 | 0.06 | 0.06 | -0.22 | 0.15 | -0.10 | 4.0E-01 | -0.10 |
| VCL | 0.05 | -0.02 | 0.05 | -0.03 | -0.14 | 0.07 | 4.0E-01 | -0.06 |
| HNRNPAB | 0.05 | -0.02 | 0.01 | 0.01 | -0.01 | -0.03 | 4.0E-01 | -0.02 |
| RRP15 | 0.11 | 0.03 | -0.04 | -0.02 | 0.05 | -0.14 | 4.0E-01 | -0.07 |
| TIPRL | -0.03 | -0.04 | -0.10 | -0.16 | 0.20 | 0.10 | 4.0E-01 | 0.10 |
| RBM6 | 0.03 | 0.00 | 0.02 | 0.05 | -0.07 | -0.03 | 4.0E-01 | -0.03 |
| TMEM260 | -0.12 | -0.16 | 0.10 | 0.11 | -0.11 | 0.15 | 4.0E-01 | 0.11 |
| MYO10 | 0.05 | -0.16 | -0.06 | -0.05 | -0.04 | 0.22 | 4.1E-01 | 0.10 |
| MRPL10 | 0.03 | 0.01 | -0.01 | 0.03 | -0.03 | -0.03 | 4.1E-01 | -0.02 |
| RPL37A | 0.02 | -0.06 | -0.17 | 0.26 | 0.08 | -0.17 | 4.1E-01 | 0.13 |
| ZC3H15 | -0.01 | -0.09 | -0.05 | 0.23 | -0.05 | -0.06 | 4.1E-01 | 0.09 |
| CTR9 | 0.04 | 0.04 | -0.03 | 0.02 | 0.00 | -0.09 | 4.1E-01 | -0.04 |
| B2M | -0.05 | -0.05 | 0.24 | 0.03 | -0.18 | -0.03 | 4.1E-01 | -0.11 |
| RPL26 | 0.09 | 0.00 | 0.03 | 0.04 | 0.04 | -0.22 | 4.1E-01 | -0.08 |
| GHITM | 0.00 | -0.02 | 0.01 | 0.00 | 0.02 | -0.01 | 4.1E-01 | 0.01 |
| RER1 | -0.03 | 0.01 | 0.07 | 0.02 | -0.04 | -0.03 | 4.1E-01 | -0.03 |

| | | | | | | | | |
|---|---|---|---|---|---|---|---|---|
| REEP3 | 0.02 | 0.00 | -0.14 | -0.05 | 0.16 | -0.01 | 4.1E-01 | 0.07 |
| CEP192 | -0.02 | -0.04 | -0.01 | 0.01 | 0.09 | -0.05 | 4.1E-01 | 0.04 |
| CENPL | 0.05 | 0.01 | -0.23 | -0.05 | 0.00 | 0.20 | 4.1E-01 | 0.11 |
| WDR48 | -0.15 | -0.05 | 0.39 | -0.17 | -0.12 | 0.01 | 4.1E-01 | -0.16 |
| CBS | -0.11 | -0.02 | 0.03 | 0.00 | -0.04 | 0.13 | 4.1E-01 | 0.06 |
| INTS7 | 0.02 | -0.04 | -0.02 | -0.03 | 0.01 | 0.06 | 4.1E-01 | 0.03 |
| TUBGCP4 | -0.12 | -0.08 | 0.09 | 0.08 | -0.04 | 0.06 | 4.1E-01 | 0.07 |
| CHD1 | 0.06 | 0.02 | -0.03 | -0.05 | -0.03 | 0.04 | 4.1E-01 | -0.03 |
| EXOSC4 | 0.08 | -0.10 | 0.10 | -0.03 | 0.01 | -0.08 | 4.1E-01 | -0.06 |
| STRN3 | -0.03 | -0.04 | -0.02 | 0.10 | -0.10 | 0.08 | 4.1E-01 | 0.06 |
| CSTF1 | 0.08 | 0.00 | -0.03 | 0.00 | -0.04 | -0.01 | 4.1E-01 | -0.03 |
| GPR89B | -0.12 | 0.14 | 0.10 | -0.04 | 0.03 | -0.13 | 4.1E-01 | -0.09 |
| FBXO38 | -0.01 | 0.02 | 0.06 | -0.11 | 0.00 | 0.04 | 4.1E-01 | -0.05 |
| ARNT | -0.18 | 0.03 | 0.04 | 0.05 | -0.02 | 0.06 | 4.1E-01 | 0.07 |
| SURF6 | 0.05 | 0.09 | -0.05 | 0.03 | 0.02 | -0.16 | 4.1E-01 | -0.07 |
| CENPE | -0.12 | 0.08 | 0.25 | 0.12 | -0.04 | -0.36 | 4.1E-01 | -0.16 |
| NECTIN2 | -0.05 | 0.12 | 0.03 | -0.10 | -0.10 | 0.08 | 4.1E-01 | -0.07 |
| COPE | -0.05 | 0.00 | 0.01 | 0.06 | -0.03 | 0.01 | 4.1E-01 | 0.03 |
| TMF1 | -0.02 | -0.04 | 0.16 | -0.10 | 0.07 | -0.10 | 4.1E-01 | -0.08 |
| TBC1D31 | -0.12 | -0.05 | 0.06 | 0.12 | -0.04 | 0.01 | 4.1E-01 | 0.07 |
| OMA1 | -0.08 | 0.01 | 0.00 | -0.02 | -0.02 | 0.10 | 4.1E-01 | 0.04 |
| SRSF9 | 0.12 | 0.03 | -0.05 | 0.07 | -0.08 | -0.10 | 4.1E-01 | -0.07 |
| PIP4K2C | 0.11 | -0.04 | -0.20 | 0.03 | 0.08 | 0.00 | 4.1E-01 | 0.08 |
| EEF1A1 | 0.04 | 0.00 | -0.15 | 0.09 | 0.08 | -0.08 | 4.1E-01 | 0.07 |
| WDR70 | 0.15 | 0.04 | -0.09 | -0.11 | 0.02 | -0.03 | 4.1E-01 | -0.07 |
| MED26 | 0.01 | 0.00 | 0.10 | 0.06 | -0.23 | 0.02 | 4.1E-01 | -0.09 |
| PPP4C | -0.10 | -0.07 | -0.06 | -0.07 | -0.11 | 0.35 | 4.1E-01 | 0.14 |
| MAPRE3 | -0.30 | 0.01 | 0.05 | -0.13 | 0.09 | 0.23 | 4.1E-01 | 0.14 |
| VKORC1 | -0.09 | -0.10 | 0.01 | -0.09 | -0.04 | 0.27 | 4.1E-01 | 0.11 |
| NCAPH2 | -0.40 | -0.45 | 1.13 | -0.42 | -0.24 | -0.48 | 4.1E-01 | -0.48 |
| GOSR1 | 0.01 | 0.01 | 0.09 | 0.11 | -0.08 | -0.15 | 4.1E-01 | -0.07 |
| CCDC137 | -0.04 | -0.02 | -0.02 | -0.05 | -0.01 | 0.13 | 4.1E-01 | 0.05 |
| EP400 | -0.02 | 0.03 | -0.05 | 0.02 | -0.01 | 0.02 | 4.2E-01 | 0.02 |
| VPS50 | 0.10 | -0.04 | 0.03 | -0.13 | 0.07 | -0.04 | 4.2E-01 | -0.06 |
| PAPSS1 | 0.01 | 0.03 | -0.09 | -0.03 | 0.05 | 0.03 | 4.2E-01 | 0.04 |
| ELOA | -0.16 | -0.06 | 0.03 | -0.07 | 0.27 | -0.06 | 4.2E-01 | 0.11 |
| EIF2AK2 | -0.01 | -0.02 | 0.08 | 0.02 | -0.01 | -0.07 | 4.2E-01 | -0.04 |
| MT-ND5 | -0.04 | -0.01 | 0.02 | 0.00 | 0.02 | 0.01 | 4.2E-01 | 0.02 |
| GORAB | -0.19 | 0.15 | 0.20 | -0.01 | -0.18 | -0.01 | 4.2E-01 | -0.12 |
| SLC3A2 | -0.11 | 0.08 | 0.14 | -0.12 | 0.02 | -0.02 | 4.2E-01 | -0.08 |
| MNAT1 | -0.10 | 0.11 | 0.19 | 0.18 | -0.38 | -0.09 | 4.2E-01 | -0.16 |
| NUP210 | 0.01 | -0.02 | -0.14 | 0.22 | 0.01 | -0.10 | 4.2E-01 | 0.09 |
| ARHGEF17 | -0.25 | -0.07 | 0.14 | -0.01 | 0.03 | 0.12 | 4.2E-01 | 0.11 |
| RPL23A | 0.02 | -0.02 | -0.07 | 0.09 | 0.03 | -0.05 | 4.2E-01 | 0.04 |
| RHOB | -0.08 | 0.00 | 0.19 | -0.11 | -0.05 | 0.03 | 4.2E-01 | -0.08 |
| HEBP1 | 0.07 | 0.08 | -0.08 | -0.02 | -0.02 | -0.02 | 4.2E-01 | -0.05 |
| YIPF3 | 0.01 | -0.09 | 0.21 | 0.02 | -0.11 | -0.05 | 4.2E-01 | -0.09 |
| SH3BP1 | -0.01 | -0.03 | -0.02 | -0.07 | 0.06 | 0.07 | 4.2E-01 | 0.04 |

| | | | | | | | | |
|---|---|---|---|---|---|---|---|---|
| NOP9 | 0.02 | 0.01 | -0.12 | -0.05 | 0.09 | 0.04 | 4.2E-01 | 0.06 |
| WWC3 | -0.06 | -0.12 | 0.05 | 0.17 | 0.01 | -0.07 | 4.2E-01 | 0.08 |
| S100A6 | 0.29 | -0.05 | -0.09 | -0.02 | -0.13 | -0.04 | 4.2E-01 | -0.11 |
| RAB3D | -0.13 | -0.03 | 0.02 | 0.21 | -0.10 | 0.01 | 4.2E-01 | 0.09 |
| TALDO1 | 0.11 | 0.09 | -0.11 | -0.05 | 0.02 | -0.08 | 4.2E-01 | -0.07 |
| NR2C1 | -0.06 | 0.03 | -0.19 | 0.30 | -0.07 | -0.07 | 4.2E-01 | 0.12 |
| PDPK1 | -0.19 | 0.11 | -0.54 | -0.47 | 0.49 | 0.29 | 4.2E-01 | 0.31 |
| ST7 | -0.11 | -0.04 | 0.05 | 0.05 | -0.06 | 0.09 | 4.2E-01 | 0.06 |
| HGS | -0.15 | -0.06 | 0.07 | -0.10 | 0.09 | 0.12 | 4.2E-01 | 0.08 |
| MYSM1 | -0.13 | 0.00 | -0.02 | -0.12 | 0.04 | 0.20 | 4.2E-01 | 0.09 |
| UBE2E1 | 0.06 | 0.01 | 0.06 | -0.24 | -0.06 | 0.14 | 4.2E-01 | -0.10 |
| PPP1R18 | 0.11 | -0.07 | -0.26 | -0.13 | 0.12 | 0.18 | 4.2E-01 | 0.13 |
| CAPRIN1 | -0.08 | -0.07 | -0.01 | 0.25 | -0.09 | -0.03 | 4.2E-01 | 0.10 |
| URB2 | 0.08 | 0.01 | -0.04 | -0.01 | 0.02 | -0.07 | 4.2E-01 | -0.04 |
| TFAP2B | 0.05 | 0.01 | 0.00 | -0.02 | -0.08 | 0.05 | 4.3E-01 | -0.04 |
| BRCA1 | -0.01 | -0.07 | 0.23 | -0.01 | 0.05 | -0.23 | 4.3E-01 | -0.11 |
| EML4 | 0.11 | -0.02 | -0.01 | -0.11 | -0.03 | 0.05 | 4.3E-01 | -0.06 |
| NDUFA6 | 0.02 | 0.09 | -0.04 | -0.09 | 0.05 | -0.04 | 4.3E-01 | -0.05 |
| SLC31A1 | 0.05 | -0.22 | 0.39 | -0.26 | 0.04 | -0.09 | 4.3E-01 | -0.17 |
| VEZT | 0.05 | -0.03 | -0.08 | 0.06 | 0.01 | -0.01 | 4.3E-01 | 0.04 |
| CPSF1 | -0.04 | -0.04 | 0.01 | 0.08 | -0.07 | 0.05 | 4.3E-01 | 0.04 |
| SEC24A | 0.08 | 0.09 | 0.03 | 0.21 | -0.09 | -0.39 | 4.3E-01 | -0.15 |
| TIMM10 | -0.01 | 0.03 | 0.02 | 0.00 | 0.02 | -0.06 | 4.3E-01 | -0.03 |
| PPIB | -0.06 | 0.30 | -0.09 | -0.09 | -0.06 | -0.04 | 4.3E-01 | -0.11 |
| ATCAY | -0.46 | -0.28 | 0.34 | 0.20 | -0.02 | 0.07 | 4.3E-01 | 0.22 |
| BRWD1 | 0.01 | 0.01 | 0.03 | -0.08 | -0.01 | 0.04 | 4.3E-01 | -0.03 |
| TRIM27 | 0.05 | 0.05 | -0.02 | -0.08 | 0.08 | -0.09 | 4.3E-01 | -0.05 |
| KLHL1 | 0.04 | -0.15 | -0.07 | 0.19 | -0.17 | 0.12 | 4.3E-01 | 0.11 |
| TJP1 | -0.09 | 0.03 | -0.01 | 0.10 | -0.01 | -0.03 | 4.3E-01 | 0.04 |
| C17orf75 | -0.01 | 0.01 | 0.08 | 0.04 | 0.01 | -0.15 | 4.3E-01 | -0.06 |
| EIF3G | -0.02 | -0.01 | 0.08 | 0.00 | 0.02 | -0.08 | 4.3E-01 | -0.04 |
| UQCRC1 | -0.05 | -0.01 | 0.13 | -0.07 | 0.00 | -0.01 | 4.3E-01 | -0.05 |
| KRT80 | 0.13 | 0.23 | -0.07 | 0.14 | 0.09 | -0.70 | 4.3E-01 | -0.25 |
| PRMT7 | -0.32 | -0.15 | 0.22 | -0.01 | 0.12 | 0.07 | 4.3E-01 | 0.14 |
| TMEM104 | -0.16 | -0.01 | 0.05 | 0.14 | -0.04 | 0.00 | 4.3E-01 | 0.07 |
| UBA52 | 0.06 | 0.03 | 0.02 | -0.05 | 0.11 | -0.18 | 4.3E-01 | -0.07 |
| TRMT6 | 0.01 | 0.09 | 0.00 | -0.06 | 0.10 | -0.16 | 4.3E-01 | -0.07 |
| CAAP1 | 0.02 | 0.05 | -0.01 | 0.04 | -0.01 | -0.09 | 4.3E-01 | -0.04 |
| MRPL35 | 0.02 | 0.05 | -0.19 | 0.10 | -0.02 | 0.02 | 4.3E-01 | 0.07 |
| NCBP2 | 0.08 | -0.05 | -0.11 | 0.03 | 0.04 | 0.00 | 4.3E-01 | 0.05 |
| RPL36A | 0.07 | 0.00 | -0.30 | 0.25 | -0.02 | -0.06 | 4.3E-01 | 0.13 |
| PDIK1L | -0.01 | 0.07 | 0.05 | -0.20 | -0.02 | 0.09 | 4.3E-01 | -0.08 |
| NDUFA10 | 0.03 | -0.01 | -0.12 | 0.11 | 0.03 | -0.06 | 4.3E-01 | 0.06 |
| NEMP2 | -0.11 | -0.06 | 0.03 | 0.19 | -0.12 | 0.05 | 4.3E-01 | 0.09 |
| MAP7 | -0.03 | 0.06 | 0.04 | 0.07 | -0.05 | -0.10 | 4.3E-01 | -0.05 |
| SNRPB | 0.00 | -0.01 | -0.03 | -0.03 | 0.00 | 0.06 | 4.3E-01 | 0.02 |
| RRAGB | 0.02 | -0.07 | 0.12 | -0.03 | 0.02 | -0.07 | 4.3E-01 | -0.05 |
| AEN | -0.11 | -0.09 | -0.05 | -0.26 | 0.29 | 0.15 | 4.3E-01 | 0.14 |

| | | | | | | | | |
|---|---|---|---|---|---|---|---|---|
| NKAPD1 | 0.14 | -0.05 | 0.02 | 0.11 | -0.10 | -0.14 | 4.3E-01 | -0.08 |
| GSPT1 | 0.07 | 0.07 | -0.07 | -0.07 | 0.05 | -0.06 | 4.3E-01 | -0.05 |
| AGR2 | 0.09 | 0.06 | -0.02 | -0.26 | 0.00 | 0.10 | 4.3E-01 | -0.10 |
| TMEM192 | -0.07 | 0.09 | 0.07 | 0.08 | -0.09 | -0.09 | 4.4E-01 | -0.07 |
| MPP3 | 0.02 | -0.01 | 0.12 | -0.14 | 0.17 | -0.21 | 4.4E-01 | -0.11 |
| EXOSC6 | 0.05 | 0.07 | -0.05 | 0.07 | -0.04 | -0.13 | 4.4E-01 | -0.06 |
| RBPJ | 0.03 | 0.02 | -0.12 | 0.04 | 0.01 | 0.02 | 4.4E-01 | 0.04 |
| GTSE1 | -0.15 | -0.12 | 0.08 | -0.15 | 0.25 | 0.06 | 4.4E-01 | 0.12 |
| GPC1 | -0.15 | 0.07 | 0.25 | -0.15 | -0.17 | 0.10 | 4.4E-01 | -0.13 |
| GNPTAB | 0.05 | -0.22 | 0.00 | -0.11 | 0.19 | 0.05 | 4.4E-01 | 0.10 |
| TLN1 | -0.03 | -0.01 | -0.03 | -0.04 | -0.02 | 0.11 | 4.4E-01 | 0.04 |
| EHD4 | 0.00 | 0.03 | -0.11 | 0.04 | 0.08 | -0.04 | 4.4E-01 | 0.05 |
| NFXL1 | -0.10 | 0.06 | 0.13 | -0.12 | -0.01 | 0.02 | 4.4E-01 | -0.07 |
| SIN3B | 0.04 | 0.01 | -0.11 | 0.04 | 0.01 | 0.00 | 4.4E-01 | 0.04 |
| PABPC4 | -0.08 | -0.09 | 0.07 | 0.06 | 0.10 | -0.07 | 4.4E-01 | 0.06 |
| IK | 0.12 | 0.06 | -0.09 | 0.00 | 0.00 | -0.10 | 4.4E-01 | -0.06 |
| TUBGCP3 | -0.04 | 0.14 | -0.32 | 0.01 | -0.03 | 0.19 | 4.4E-01 | 0.13 |
| MIER3 | 0.09 | -0.36 | 0.66 | -0.02 | 0.28 | -1.39 | 4.4E-01 | -0.51 |
| NANS | -0.03 | -0.03 | 0.14 | 0.00 | -0.09 | 0.00 | 4.4E-01 | -0.06 |
| PLD2 | -0.10 | -0.12 | 0.10 | 0.11 | 0.00 | -0.02 | 4.4E-01 | 0.07 |
| MARVELD3 | -0.03 | -0.08 | 0.04 | 0.10 | -0.04 | 0.00 | 4.4E-01 | 0.05 |
| DAGLB | -0.20 | 0.10 | 0.26 | -0.10 | 0.07 | -0.19 | 4.4E-01 | -0.13 |
| PDIA4 | 0.14 | 0.17 | -0.18 | -0.02 | -0.03 | -0.12 | 4.4E-01 | -0.10 |
| DNAJC4 | 0.01 | 0.03 | 0.03 | 0.03 | -0.15 | 0.03 | 4.4E-01 | -0.05 |
| DNAJC3 | 0.04 | -0.03 | -0.06 | -0.03 | 0.05 | 0.04 | 4.4E-01 | 0.03 |
| HSD17B7 | 0.04 | -0.03 | -0.07 | 0.04 | 0.04 | -0.03 | 4.4E-01 | 0.03 |
| NPLOC4 | 0.05 | 0.05 | -0.06 | -0.03 | -0.02 | 0.00 | 4.4E-01 | -0.03 |
| ZMAT2 | 0.12 | -0.03 | -0.01 | 0.07 | -0.08 | -0.10 | 4.5E-01 | -0.06 |
| TJP3 | -0.12 | 0.02 | 0.23 | 0.03 | -0.13 | -0.05 | 4.5E-01 | -0.09 |
| NKRF | 0.03 | -0.03 | -0.03 | 0.06 | 0.00 | -0.02 | 4.5E-01 | 0.03 |
| NUTM2B | -0.06 | -0.04 | 0.20 | -0.03 | -0.03 | -0.06 | 4.5E-01 | -0.07 |
| NDUFA5 | 0.04 | 0.02 | -0.01 | 0.05 | -0.02 | -0.09 | 4.5E-01 | -0.04 |
| SFXN2 | -0.03 | 0.08 | 0.02 | -0.10 | -0.05 | 0.07 | 4.5E-01 | -0.05 |
| PAFAH1B1 | 0.04 | 0.12 | -0.05 | 0.07 | 0.00 | -0.21 | 4.5E-01 | -0.08 |
| CCS | -0.04 | -0.09 | 0.02 | 0.08 | 0.11 | -0.11 | 4.5E-01 | 0.06 |
| GNAI2 | -0.13 | -0.11 | 0.11 | 0.05 | -0.04 | 0.09 | 4.5E-01 | 0.07 |
| PGM3 | -0.24 | -0.08 | 0.02 | 0.30 | 0.19 | -0.29 | 4.5E-01 | 0.16 |
| HCFC1 | 0.03 | 0.03 | -0.02 | -0.03 | -0.03 | 0.03 | 4.5E-01 | -0.02 |
| B3GLCT | -0.04 | 0.03 | -0.08 | -0.07 | 0.01 | 0.14 | 4.5E-01 | 0.06 |
| RAB19 | -0.05 | -0.05 | 0.22 | -0.04 | -0.13 | 0.02 | 4.5E-01 | -0.09 |
| CYFIP1 | -0.03 | 0.01 | 0.05 | -0.05 | 0.02 | -0.01 | 4.5E-01 | -0.02 |
| MISP3 | -0.08 | 0.06 | 0.09 | -0.05 | 0.04 | -0.07 | 4.5E-01 | -0.05 |
| YEATS2 | 0.04 | -0.13 | -0.02 | 0.14 | -0.09 | 0.04 | 4.5E-01 | 0.07 |
| RPS19 | -0.04 | -0.05 | 0.03 | 0.08 | 0.03 | -0.06 | 4.5E-01 | 0.04 |
| EFCAB14 | -0.04 | 0.02 | 0.08 | -0.02 | 0.05 | -0.10 | 4.5E-01 | -0.05 |
| CYP51A1 | 0.02 | 0.02 | -0.29 | 0.33 | -0.03 | -0.12 | 4.5E-01 | 0.14 |
| FKBP2 | 0.17 | 0.06 | -0.08 | 0.10 | -0.03 | -0.25 | 4.5E-01 | -0.11 |
| AP3S1 | 0.04 | -0.01 | 0.09 | -0.06 | 0.13 | -0.20 | 4.5E-01 | -0.08 |

| | | | | | | | | |
|---|---|---|---|---|---|---|---|---|
| CSNK2B | 0.02 | -0.04 | -0.05 | 0.10 | 0.06 | -0.08 | 4.5E-01 | 0.05 |
| LIMA1 | -0.04 | 0.04 | 0.04 | 0.01 | -0.04 | -0.01 | 4.5E-01 | -0.02 |
| RMND1 | 0.01 | 0.01 | -0.06 | 0.02 | 0.06 | -0.04 | 4.5E-01 | 0.03 |
| CRIP2 | 0.02 | 0.07 | -0.22 | 0.07 | -0.04 | 0.08 | 4.5E-01 | 0.08 |
| ARF3 | -0.02 | 0.00 | -0.03 | -0.04 | 0.10 | 0.00 | 4.5E-01 | 0.04 |
| YTHDC1 | 0.00 | -0.01 | -0.01 | 0.06 | -0.01 | -0.02 | 4.5E-01 | 0.02 |
| TRIP12 | -0.02 | -0.01 | -0.03 | -0.05 | 0.00 | 0.10 | 4.5E-01 | 0.04 |
| PLOD3 | 0.04 | 0.02 | 0.02 | 0.08 | -0.03 | -0.14 | 4.5E-01 | -0.05 |
| BAZ1B | 0.02 | -0.06 | -0.02 | -0.02 | -0.02 | 0.10 | 4.5E-01 | 0.04 |
| ELF1 | -0.12 | -0.12 | 0.08 | -0.10 | 0.02 | 0.21 | 4.5E-01 | 0.09 |
| PLD3 | -0.02 | -0.09 | 0.06 | 0.03 | 0.03 | -0.01 | 4.5E-01 | 0.04 |
| ALDH9A1 | 0.10 | -0.03 | -0.19 | 0.02 | 0.09 | 0.00 | 4.5E-01 | 0.07 |
| NFIA | 0.07 | 0.00 | -0.20 | 0.08 | 0.10 | -0.07 | 4.5E-01 | 0.08 |
| DKC1 | 0.04 | 0.02 | -0.01 | 0.04 | 0.00 | -0.11 | 4.5E-01 | -0.04 |
| PSIP1 | -0.04 | -0.04 | 0.01 | -0.02 | -0.03 | 0.12 | 4.6E-01 | 0.04 |
| NDUFA7 | 0.02 | 0.00 | -0.07 | 0.05 | 0.02 | -0.03 | 4.6E-01 | 0.03 |
| GSN | 0.07 | 0.01 | 0.01 | 0.03 | 0.06 | -0.19 | 4.6E-01 | -0.07 |
| POLR3G | -0.27 | -0.17 | 0.16 | 0.31 | -0.18 | 0.06 | 4.6E-01 | 0.16 |
| RAD17 | 0.02 | 0.07 | -0.02 | 0.08 | -0.11 | -0.06 | 4.6E-01 | -0.05 |
| LRRC8A | -0.01 | -0.07 | -0.13 | 0.07 | 0.29 | -0.21 | 4.6E-01 | 0.12 |
| RALGAPA2 | -0.36 | -0.24 | 0.86 | -0.27 | -0.07 | -0.37 | 4.6E-01 | -0.33 |
| RPS17 | -0.10 | -0.09 | 0.08 | 0.15 | -0.02 | -0.03 | 4.6E-01 | 0.07 |
| LSM8 | 0.11 | -0.04 | 0.05 | -0.15 | -0.15 | 0.15 | 4.6E-01 | -0.09 |
| ATXN2L | -0.04 | 0.05 | 0.03 | 0.00 | 0.01 | -0.06 | 4.6E-01 | -0.03 |
| PTPN13 | -0.29 | -0.06 | 0.16 | 0.06 | 0.13 | -0.04 | 4.6E-01 | 0.11 |
| CBX8 | 0.00 | -0.03 | -0.02 | 0.02 | -0.05 | 0.07 | 4.6E-01 | 0.03 |
| NOP16 | -0.02 | -0.04 | -0.03 | 0.16 | 0.00 | -0.08 | 4.6E-01 | 0.06 |
| FANCM | -0.10 | -0.07 | 0.06 | 0.05 | -0.10 | 0.14 | 4.6E-01 | 0.07 |
| TMEM175 | -0.13 | -0.10 | 0.45 | -0.15 | -0.28 | 0.10 | 4.6E-01 | -0.18 |
| CSNK1E | -0.10 | -0.09 | 0.10 | 0.05 | 0.05 | -0.02 | 4.6E-01 | 0.05 |
| CCDC28B | -0.08 | 0.03 | 0.16 | -0.03 | 0.06 | -0.15 | 4.6E-01 | -0.08 |
| CEBPD | 0.05 | -0.30 | 0.03 | 0.28 | -0.03 | -0.09 | 4.6E-01 | 0.13 |
| PCF11 | -0.01 | -0.02 | -0.03 | 0.05 | 0.06 | -0.06 | 4.6E-01 | 0.03 |
| CIRBP | 0.00 | -0.03 | 0.07 | 0.03 | 0.00 | -0.08 | 4.6E-01 | -0.03 |
| GPC3 | -0.02 | 0.09 | 0.05 | -0.26 | 0.07 | 0.04 | 4.6E-01 | -0.09 |
| ABCG1 | -0.03 | 0.02 | 0.14 | -0.24 | 0.15 | -0.08 | 4.6E-01 | -0.10 |
| WDR59 | 0.09 | 0.10 | -0.48 | 0.20 | -0.02 | 0.03 | 4.6E-01 | 0.16 |
| GABPA | 0.01 | -0.03 | -0.06 | -0.02 | 0.13 | -0.04 | 4.6E-01 | 0.05 |
| SPRYD7 | -0.07 | 0.04 | 0.13 | -0.14 | 0.05 | -0.02 | 4.6E-01 | -0.07 |
| TRIM56 | -0.11 | -0.05 | 0.09 | 0.00 | 0.01 | 0.07 | 4.6E-01 | 0.05 |
| ASPSCR1 | -0.02 | -0.04 | 0.03 | 0.00 | 0.03 | 0.00 | 4.6E-01 | 0.02 |
| C11orf68 | 0.08 | 0.02 | 0.00 | 0.12 | -0.05 | -0.20 | 4.6E-01 | -0.08 |
| ABLIM3 | 0.07 | 0.10 | -0.08 | -0.10 | 0.08 | -0.08 | 4.6E-01 | -0.06 |
| BRAP | -0.05 | -0.07 | 0.00 | -0.13 | 0.04 | 0.19 | 4.6E-01 | 0.08 |
| ARID1B | -0.06 | -0.06 | 0.05 | -0.03 | -0.01 | 0.10 | 4.6E-01 | 0.04 |
| CISD3 | -0.13 | -0.05 | 0.32 | -0.07 | -0.04 | -0.08 | 4.7E-01 | -0.11 |
| MNT | -0.07 | -0.16 | 0.10 | 0.10 | -0.10 | 0.11 | 4.7E-01 | 0.08 |
| SYMPK | 0.01 | 0.01 | -0.06 | 0.06 | 0.00 | -0.01 | 4.7E-01 | 0.03 |

| | | | | | | | | |
|---|---|---|---|---|---|---|---|---|
| CPNE2 | 0.06 | 0.10 | 0.02 | -0.19 | -0.37 | 0.28 | 4.7E-01 | -0.16 |
| STK10 | 0.00 | -0.51 | -0.10 | -0.03 | -0.41 | 0.71 | 4.7E-01 | 0.29 |
| ZNF384 | 0.01 | -0.08 | 0.00 | -0.01 | -0.04 | 0.11 | 4.7E-01 | 0.04 |
| RNF113A | 0.16 | -0.02 | -0.06 | -0.04 | -0.07 | 0.01 | 4.7E-01 | -0.06 |
| ZNF740 | -0.07 | -0.05 | 0.04 | -0.04 | -0.03 | 0.14 | 4.7E-01 | 0.05 |
| NDUFAF3 | -0.07 | -0.05 | 0.06 | -0.01 | -0.01 | 0.08 | 4.7E-01 | 0.04 |
| ECSIT | -0.06 | 0.00 | 0.14 | 0.01 | -0.04 | -0.05 | 4.7E-01 | -0.05 |
| CIZ1 | -0.04 | -0.06 | 0.00 | -0.12 | 0.07 | 0.13 | 4.7E-01 | 0.06 |
| CSTF2 | 0.02 | -0.07 | 0.16 | 0.12 | -0.13 | -0.12 | 4.7E-01 | -0.08 |
| RPL4 | -0.05 | -0.03 | 0.18 | 0.02 | 0.00 | -0.14 | 4.7E-01 | -0.07 |
| CSNK1G2-AS1 | -0.38 | -0.44 | 0.28 | 0.51 | -0.32 | 0.08 | 4.7E-01 | 0.27 |
| VWA8 | 0.04 | -0.01 | 0.01 | 0.02 | 0.02 | -0.08 | 4.7E-01 | -0.03 |
| C18orf32 | 0.08 | 0.22 | -0.16 | 0.10 | -0.20 | -0.10 | 4.7E-01 | -0.11 |
| KRT2 | -0.09 | -0.31 | -0.84 | -0.25 | -0.71 | 1.16 | 4.7E-01 | 0.48 |
| RAE1 | 0.06 | 0.02 | -0.03 | 0.05 | -0.03 | -0.08 | 4.7E-01 | -0.04 |
| AFTPH | -0.19 | -0.15 | 0.17 | -0.03 | 0.16 | 0.00 | 4.7E-01 | 0.10 |
| PCIF1 | -0.09 | 0.00 | 0.19 | -0.10 | 0.02 | -0.03 | 4.7E-01 | -0.07 |
| CHMP4A | 0.10 | 0.05 | -0.06 | -0.02 | 0.06 | -0.14 | 4.7E-01 | -0.06 |
| MAGED1 | -0.11 | -0.02 | 0.02 | 0.14 | 0.03 | -0.09 | 4.7E-01 | 0.06 |
| ERBB3 | -0.12 | -0.14 | 0.13 | 0.05 | -0.07 | 0.13 | 4.7E-01 | 0.08 |
| ZNF219 | -0.21 | -0.10 | 0.15 | 0.02 | -0.04 | 0.14 | 4.7E-01 | 0.09 |
| GNL3 | 0.08 | 0.04 | -0.05 | 0.01 | 0.04 | -0.14 | 4.7E-01 | -0.05 |
| BORCS7 | -0.15 | -0.03 | 0.32 | -0.02 | -0.11 | -0.07 | 4.7E-01 | -0.12 |
| TOMM40L | -0.04 | -0.01 | 0.00 | 0.02 | -0.05 | 0.07 | 4.7E-01 | 0.03 |
| KIF5B | 0.04 | -0.01 | -0.06 | 0.01 | 0.04 | -0.01 | 4.7E-01 | 0.03 |
| NTMT1 | 0.02 | 0.08 | 0.03 | -0.21 | 0.19 | -0.16 | 4.7E-01 | -0.10 |
| PTPRS | -0.03 | 0.06 | 0.13 | 0.11 | -0.38 | 0.05 | 4.7E-01 | -0.13 |
| GDAP1 | -0.06 | -0.07 | 0.06 | 0.08 | -0.03 | 0.01 | 4.7E-01 | 0.04 |
| ATR | -0.06 | -0.11 | 0.04 | -0.10 | 0.21 | 0.00 | 4.7E-01 | 0.08 |
| SPIN1 | 0.00 | 0.05 | -0.14 | -0.03 | 0.04 | 0.06 | 4.7E-01 | 0.05 |
| MUL1 | -0.01 | 0.02 | 0.07 | -0.10 | -0.09 | 0.10 | 4.7E-01 | -0.06 |
| FEM1B | 0.00 | -0.04 | 0.17 | -0.25 | 0.10 | -0.02 | 4.7E-01 | -0.10 |
| GOLGA1 | -0.04 | -0.05 | -0.03 | -0.14 | 0.10 | 0.14 | 4.8E-01 | 0.07 |
| PTTG1IP | 0.13 | -0.11 | 0.12 | 0.14 | -0.09 | -0.24 | 4.8E-01 | -0.11 |
| CASP7 | 0.04 | 0.08 | -0.02 | -0.21 | 0.01 | 0.09 | 4.8E-01 | -0.07 |
| CGAS | 0.01 | -0.01 | 0.07 | -0.01 | -0.12 | 0.06 | 4.8E-01 | -0.04 |
| BPTF | 0.04 | 0.01 | 0.01 | -0.13 | 0.00 | 0.06 | 4.8E-01 | -0.04 |
| RSU1 | 0.03 | -0.08 | -0.06 | -0.08 | 0.00 | 0.16 | 4.8E-01 | 0.06 |
| FAM76B | 0.08 | 0.07 | -0.03 | 0.14 | -0.07 | -0.21 | 4.8E-01 | -0.08 |
| AGAP3 | 0.01 | -0.06 | 0.11 | 0.04 | -0.09 | -0.03 | 4.8E-01 | -0.05 |
| NUDCD3 | -0.13 | 0.12 | -0.19 | -0.17 | 0.08 | 0.24 | 4.8E-01 | 0.12 |
| PTRH1 | -0.09 | -0.06 | 0.08 | 0.05 | 0.02 | 0.00 | 4.8E-01 | 0.04 |
| PGAP3 | -0.19 | 0.00 | 0.02 | -0.16 | 0.17 | 0.11 | 4.8E-01 | 0.10 |
| MAGOHB | 0.06 | 0.00 | -0.01 | 0.04 | -0.05 | -0.03 | 4.8E-01 | -0.03 |
| TTF1 | -0.05 | 0.01 | 0.00 | 0.03 | -0.04 | 0.04 | 4.8E-01 | 0.02 |
| UNC45A | -0.02 | 0.00 | 0.06 | 0.04 | -0.03 | -0.06 | 4.8E-01 | -0.03 |
| ARAP1 | 0.06 | -0.24 | 0.06 | -0.01 | 0.07 | 0.04 | 4.8E-01 | 0.08 |
| RETSAT | 0.03 | 0.01 | -0.02 | 0.01 | -0.04 | 0.01 | 4.8E-01 | -0.02 |

| | | | | | | | | |
|---|---|---|---|---|---|---|---|---|
| FARP2 | -0.09 | 0.09 | -0.08 | 0.00 | 0.06 | 0.01 | 4.8E-01 | 0.05 |
| RPRD2 | 0.03 | -0.01 | -0.07 | 0.02 | -0.04 | 0.06 | 4.8E-01 | 0.03 |
| ACP1 | 0.18 | -0.21 | 0.25 | -0.46 | 0.22 | -0.11 | 4.8E-01 | -0.19 |
| CCT8 | 0.10 | 0.07 | -0.08 | 0.01 | 0.03 | -0.13 | 4.8E-01 | -0.06 |
| ATM | -0.03 | -0.05 | -0.04 | -0.05 | 0.22 | -0.08 | 4.8E-01 | 0.07 |
| SUB1 | 0.20 | 0.03 | -0.11 | -0.12 | -0.10 | 0.08 | 4.8E-01 | -0.09 |
| FARSB | -0.02 | -0.07 | 0.01 | 0.13 | 0.00 | -0.07 | 4.8E-01 | 0.05 |
| CHD4 | 0.03 | 0.02 | -0.01 | -0.07 | -0.02 | 0.04 | 4.8E-01 | -0.03 |
| SLC20A2 | -0.11 | 0.09 | 0.09 | -0.05 | -0.04 | 0.00 | 4.8E-01 | -0.05 |
| FOXK1 | 0.02 | 0.01 | 0.03 | -0.10 | 0.07 | -0.03 | 4.8E-01 | -0.04 |
| CAPN7 | 0.01 | 0.03 | 0.01 | 0.03 | 0.01 | -0.10 | 4.8E-01 | -0.03 |
| OTUD6B | -0.04 | -0.06 | 0.05 | 0.04 | 0.00 | 0.00 | 4.8E-01 | 0.03 |
| LSM3 | 0.03 | -0.11 | -0.02 | 0.01 | -0.07 | 0.14 | 4.8E-01 | 0.06 |
| CRB3 | -0.23 | 0.05 | 0.04 | 0.10 | -0.08 | 0.08 | 4.8E-01 | 0.08 |
| PRC1 | -0.07 | -0.02 | 0.03 | -0.04 | 0.07 | 0.02 | 4.8E-01 | 0.03 |
| RBMXL1 | 0.05 | 0.03 | -0.03 | -0.05 | 0.05 | -0.05 | 4.8E-01 | -0.03 |
| YBX1 | -0.11 | -0.04 | 0.27 | -0.06 | -0.01 | -0.08 | 4.8E-01 | -0.09 |
| CHMP2B | 0.02 | 0.04 | -0.01 | -0.07 | 0.06 | -0.05 | 4.8E-01 | -0.03 |
| PDHA1 | 0.03 | 0.08 | -0.06 | -0.05 | 0.03 | -0.03 | 4.8E-01 | -0.04 |
| AMFR | -0.07 | -0.14 | 0.36 | -0.04 | -0.13 | -0.04 | 4.9E-01 | -0.12 |
| ANXA5 | 0.01 | -0.03 | 0.05 | 0.01 | -0.03 | -0.01 | 4.9E-01 | -0.02 |
| UACA | -0.10 | -0.05 | 0.07 | -0.06 | 0.02 | 0.11 | 4.9E-01 | 0.05 |
| RPS24 | -0.06 | 0.11 | -0.16 | 0.08 | 0.06 | -0.05 | 4.9E-01 | 0.07 |
| CNTRL | -0.12 | -0.26 | 0.16 | -0.16 | 0.17 | 0.14 | 4.9E-01 | 0.12 |
| SAP30BP | 0.06 | 0.07 | -0.07 | 0.01 | -0.04 | -0.03 | 4.9E-01 | -0.04 |
| CYB5R3 | -0.01 | 0.03 | -0.08 | -0.03 | -0.01 | 0.09 | 4.9E-01 | 0.04 |
| EMC9 | -0.13 | -0.06 | 0.10 | 0.06 | -0.04 | 0.05 | 4.9E-01 | 0.06 |
| MVD | -0.01 | -0.06 | -0.08 | -0.03 | -0.12 | 0.26 | 4.9E-01 | 0.09 |
| BOK | 0.06 | -0.18 | -0.03 | -0.10 | 0.20 | 0.01 | 4.9E-01 | 0.09 |
| BCKDK | 0.14 | 0.04 | -0.10 | 0.05 | -0.09 | -0.07 | 4.9E-01 | -0.06 |
| NDST1 | -0.15 | 0.04 | 0.24 | -0.13 | -0.12 | 0.08 | 4.9E-01 | -0.10 |
| CD2BP2 | 0.02 | 0.11 | -0.04 | 0.09 | -0.01 | -0.20 | 4.9E-01 | -0.07 |
| ATF7 | 0.01 | 0.07 | 0.01 | -0.13 | -0.10 | 0.13 | 4.9E-01 | -0.06 |
| CISD1 | -0.14 | -0.07 | 0.06 | -0.06 | -0.06 | 0.24 | 4.9E-01 | 0.09 |
| SURF2 | -0.37 | 0.13 | -0.06 | 0.32 | -0.22 | 0.09 | 4.9E-01 | 0.16 |
| DRAP1 | 0.11 | 0.01 | -0.04 | -0.09 | 0.10 | -0.11 | 4.9E-01 | -0.06 |
| PHAX | -0.04 | 0.08 | -0.13 | -0.03 | 0.06 | 0.04 | 4.9E-01 | 0.05 |
| ZNF644 | -0.07 | -0.01 | 0.03 | -0.04 | 0.00 | 0.09 | 4.9E-01 | 0.04 |
| CAST | -0.14 | 0.19 | 0.08 | 0.12 | -0.06 | -0.26 | 4.9E-01 | -0.11 |
| ATP6V0D1 | 0.03 | -0.02 | -0.09 | 0.13 | -0.01 | -0.05 | 4.9E-01 | 0.05 |
| TADA3 | -0.29 | -0.04 | 0.09 | -0.11 | 0.31 | -0.04 | 4.9E-01 | 0.13 |
| SPTBN2 | -0.10 | -0.02 | 0.05 | 0.10 | 0.01 | -0.05 | 4.9E-01 | 0.05 |
| THAP12 | -0.03 | 0.09 | 0.00 | 0.03 | 0.01 | -0.11 | 4.9E-01 | -0.04 |
| CTDSP1 | -0.15 | 0.19 | 0.09 | -0.06 | -0.22 | 0.10 | 4.9E-01 | -0.10 |
| FYTTD1 | -0.01 | 0.01 | -0.06 | -0.07 | 0.10 | 0.02 | 4.9E-01 | 0.04 |
| RAB5B | -0.15 | -0.12 | 0.12 | -0.03 | -0.06 | 0.20 | 4.9E-01 | 0.09 |
| OTUD4 | -0.05 | -0.26 | 0.15 | -0.07 | 0.11 | 0.09 | 4.9E-01 | 0.10 |
| EMG1 | -0.03 | -0.03 | 0.11 | -0.02 | -0.06 | 0.01 | 4.9E-01 | -0.04 |

| | | | | | | | | |
|---|---|---|---|---|---|---|---|---|
| UNC13C | 0.07 | -0.10 | 0.15 | 0.06 | -0.22 | 0.01 | 5.0E-01 | -0.09 |
| PPP2R2A | -0.04 | -0.06 | 0.04 | 0.00 | 0.09 | -0.04 | 5.0E-01 | 0.04 |
| AGO1 | -0.08 | -0.02 | 0.06 | -0.01 | 0.03 | 0.02 | 5.0E-01 | 0.03 |
| ITM2B | -0.16 | -0.26 | 0.17 | 0.09 | -0.19 | 0.26 | 5.0E-01 | 0.14 |
| NEFL | -0.69 | 0.11 | -0.04 | -0.35 | -0.05 | 0.66 | 5.0E-01 | 0.29 |
| ZNF280D | 0.02 | 0.02 | 0.01 | -0.09 | -0.04 | 0.07 | 5.0E-01 | -0.04 |
| WIZ | 0.01 | -0.05 | -0.03 | -0.04 | -0.02 | 0.12 | 5.0E-01 | 0.04 |
| KMT2A | 0.02 | 0.03 | 0.00 | -0.09 | -0.01 | 0.05 | 5.0E-01 | -0.03 |
| USO1 | -0.10 | 0.08 | -0.12 | 0.19 | 0.05 | -0.13 | 5.0E-01 | 0.08 |
| APOOL | -0.04 | -0.06 | 0.04 | 0.05 | -0.05 | 0.05 | 5.0E-01 | 0.03 |
| NCLN | -0.01 | -0.05 | 0.12 | -0.02 | 0.01 | -0.06 | 5.0E-01 | -0.04 |
| TMED7 | -0.19 | -0.23 | 0.17 | 0.26 | -0.22 | 0.13 | 5.0E-01 | 0.14 |
| SECTM1 | -0.34 | -0.12 | 0.21 | -0.03 | -0.05 | 0.24 | 5.0E-01 | 0.14 |
| CENPQ | 0.07 | 0.11 | -0.09 | -0.15 | 0.05 | -0.01 | 5.0E-01 | -0.06 |
| TMEM126B | -0.01 | -0.01 | -0.04 | 0.06 | -0.07 | 0.06 | 5.0E-01 | 0.03 |
| RPL13A | 0.07 | 0.04 | -0.01 | 0.12 | -0.01 | -0.24 | 5.0E-01 | -0.08 |
| PRRC2B | -0.13 | -0.06 | 0.11 | 0.01 | 0.06 | 0.01 | 5.0E-01 | 0.05 |
| FAM20B | -0.01 | 0.03 | -0.15 | 0.20 | -0.09 | -0.01 | 5.0E-01 | 0.08 |
| EPB41 | -0.15 | 0.01 | 0.06 | 0.04 | 0.01 | 0.01 | 5.0E-01 | 0.05 |
| CERS6 | 0.01 | -0.12 | 0.03 | 0.04 | -0.07 | 0.10 | 5.0E-01 | 0.05 |
| SPG7 | -0.03 | 0.06 | 0.01 | 0.01 | 0.00 | -0.04 | 5.0E-01 | -0.02 |
| CAPNS1 | -0.04 | -0.26 | 0.12 | 0.17 | -0.15 | 0.12 | 5.0E-01 | 0.11 |
| WDR89 | 0.09 | -0.01 | -0.19 | 0.10 | 0.03 | -0.05 | 5.0E-01 | 0.07 |
| RBM19 | 0.05 | -0.02 | 0.00 | 0.01 | -0.03 | -0.01 | 5.1E-01 | -0.02 |
| PRKAG1 | -0.04 | 0.07 | -0.13 | -0.04 | 0.14 | -0.01 | 5.1E-01 | 0.06 |
| KANSL1 | -0.12 | -0.02 | -0.03 | -0.17 | 0.03 | 0.27 | 5.1E-01 | 0.10 |
| SLC30A6 | -0.03 | 0.00 | -0.10 | 0.18 | 0.09 | -0.17 | 5.1E-01 | 0.08 |
| DYNLL1 | 0.01 | -0.02 | 0.04 | -0.06 | 0.00 | 0.03 | 5.1E-01 | -0.02 |
| ZNF24 | 0.02 | -0.03 | -0.02 | 0.04 | -0.03 | 0.02 | 5.1E-01 | 0.02 |
| CMSS1 | -0.03 | -0.08 | 0.06 | 0.00 | 0.05 | 0.00 | 5.1E-01 | 0.03 |
| DNAJC5 | -0.11 | -0.04 | 0.04 | 0.05 | -0.11 | 0.15 | 5.1E-01 | 0.07 |
| ZNF687 | 0.02 | 0.01 | 0.02 | -0.11 | -0.02 | 0.07 | 5.1E-01 | -0.04 |
| PPIP5K1 | 0.10 | -0.18 | -0.17 | -0.20 | 0.01 | 0.36 | 5.1E-01 | 0.14 |
| NDUFB1 | -0.01 | 0.06 | -0.02 | 0.00 | -0.04 | 0.00 | 5.1E-01 | -0.02 |
| NAA10 | 0.01 | 0.05 | -0.14 | 0.01 | 0.07 | -0.02 | 5.1E-01 | 0.05 |
| EMC6 | -0.03 | 0.01 | 0.12 | 0.08 | 0.01 | -0.21 | 5.1E-01 | -0.07 |
| EEF1D | -0.05 | -0.07 | 0.05 | 0.03 | -0.07 | 0.09 | 5.1E-01 | 0.04 |
| TRIM26 | 0.06 | -0.07 | 0.06 | -0.01 | 0.03 | -0.07 | 5.1E-01 | -0.04 |
| SLCO2A1 | -0.10 | 0.05 | 0.14 | 0.05 | 0.01 | -0.17 | 5.1E-01 | -0.07 |
| APBA3 | -0.06 | 0.02 | 0.11 | -0.11 | 0.06 | -0.03 | 5.1E-01 | -0.05 |
| RAB40C | 0.07 | -0.07 | 0.19 | -0.51 | 0.06 | 0.15 | 5.1E-01 | -0.16 |
| DDX59 | 0.12 | 0.13 | -0.13 | -0.18 | -0.13 | 0.15 | 5.1E-01 | -0.10 |
| CLCC1 | 0.06 | 0.05 | -0.04 | -0.16 | -0.02 | 0.09 | 5.1E-01 | -0.06 |
| CEP350 | -0.04 | -0.03 | 0.02 | -0.06 | 0.03 | 0.07 | 5.1E-01 | 0.03 |
| CTNNBL1 | 0.02 | -0.03 | 0.04 | -0.04 | -0.01 | 0.02 | 5.1E-01 | -0.02 |
| GUF1 | 0.07 | -0.03 | -0.12 | -0.04 | 0.08 | 0.02 | 5.1E-01 | 0.05 |
| MTA3 | 0.01 | -0.07 | 0.02 | -0.03 | 0.00 | 0.07 | 5.1E-01 | 0.03 |
| NUTF2 | 0.06 | 0.08 | -0.03 | -0.19 | 0.17 | -0.12 | 5.1E-01 | -0.08 |

| | | | | | | | | |
|---|---|---|---|---|---|---|---|---|
| CSTF2T | -0.03 | -0.01 | -0.04 | -0.02 | -0.07 | 0.16 | 5.1E-01 | 0.05 |
| MLLT11 | 0.04 | -0.18 | 0.04 | 0.09 | -0.07 | 0.07 | 5.1E-01 | 0.06 |
| YEATS4 | 0.00 | 0.01 | -0.08 | -0.06 | 0.01 | 0.11 | 5.1E-01 | 0.04 |
| SRBD1 | -0.01 | -0.01 | -0.01 | -0.04 | 0.05 | 0.02 | 5.1E-01 | 0.02 |
| DNAJC25 | -0.02 | -0.04 | 0.14 | -0.10 | -0.07 | 0.08 | 5.1E-01 | -0.06 |
| ZNHIT1 | 0.06 | 0.07 | -0.03 | -0.23 | 0.05 | 0.06 | 5.1E-01 | -0.07 |
| CEP170 | -0.04 | -0.05 | 0.02 | 0.11 | -0.07 | 0.02 | 5.1E-01 | 0.04 |
| CHMP2A | 0.04 | -0.02 | 0.00 | 0.01 | -0.04 | 0.00 | 5.2E-01 | -0.02 |
| GNG12 | -0.02 | 0.04 | 0.04 | -0.09 | -0.07 | 0.09 | 5.2E-01 | -0.04 |
| SBDS | 0.07 | 0.03 | -0.05 | -0.02 | 0.01 | -0.03 | 5.2E-01 | -0.03 |
| NUP85 | -0.02 | 0.00 | 0.04 | 0.00 | -0.01 | -0.01 | 5.2E-01 | -0.01 |
| GPS1 | -0.04 | -0.11 | 0.07 | -0.06 | 0.05 | 0.09 | 5.2E-01 | 0.05 |
| ELOB | 0.02 | 0.00 | -0.09 | 0.13 | 0.00 | -0.06 | 5.2E-01 | 0.05 |
| PKN2 | -0.03 | -0.09 | 0.05 | -0.03 | 0.10 | -0.02 | 5.2E-01 | 0.04 |
| SELENOS | 0.08 | -0.05 | 0.06 | -0.10 | 0.12 | -0.13 | 5.2E-01 | -0.06 |
| HIBCH | 0.08 | -0.01 | -0.16 | 0.02 | 0.08 | -0.03 | 5.2E-01 | 0.06 |
| RPL29 | 0.05 | -0.05 | 0.08 | 0.11 | -0.04 | -0.17 | 5.2E-01 | -0.06 |
| NECTIN4 | -0.21 | -0.04 | 0.11 | 0.02 | -0.06 | 0.14 | 5.2E-01 | 0.08 |
| MEPCE | -0.02 | 0.06 | 0.07 | 0.04 | 0.08 | -0.26 | 5.2E-01 | -0.08 |
| INTS13 | -0.01 | -0.05 | 0.00 | 0.02 | -0.07 | 0.10 | 5.2E-01 | 0.04 |
| MRPS25 | 0.10 | -0.10 | -0.09 | -0.01 | -0.03 | 0.12 | 5.2E-01 | 0.06 |
| SGF29 | -0.02 | 0.00 | -0.09 | 0.17 | 0.07 | -0.15 | 5.2E-01 | 0.07 |
| NOVA1 | -0.07 | -0.08 | 0.06 | 0.06 | -0.11 | 0.13 | 5.2E-01 | 0.06 |
| SKIC8 | -0.01 | 0.15 | -0.01 | -0.08 | 0.18 | -0.27 | 5.2E-01 | -0.10 |
| GTF2E2 | -0.09 | 0.07 | 0.10 | -0.11 | 0.01 | 0.02 | 5.2E-01 | -0.05 |
| SMARCD3 | 0.08 | -0.02 | 0.03 | -0.21 | 0.14 | -0.06 | 5.2E-01 | -0.07 |
| TUBE1 | -0.08 | -0.28 | 0.17 | 0.18 | 0.06 | -0.10 | 5.2E-01 | 0.11 |
| ETV6 | -0.02 | -0.02 | 0.01 | 0.06 | 0.00 | -0.04 | 5.2E-01 | 0.02 |
| ZNF622 | -0.06 | -0.09 | 0.06 | -0.05 | 0.15 | -0.02 | 5.2E-01 | 0.05 |
| NPHP4 | -0.16 | 0.02 | 0.29 | -0.02 | -0.26 | 0.07 | 5.2E-01 | -0.12 |
| REEP4 | 0.07 | -0.06 | 0.04 | -0.08 | 0.04 | -0.02 | 5.2E-01 | -0.04 |
| CFAP20 | 0.11 | -0.01 | -0.02 | 0.10 | -0.05 | -0.15 | 5.2E-01 | -0.06 |
| ELL | -0.27 | -0.07 | 0.10 | 0.27 | -0.25 | 0.13 | 5.2E-01 | 0.13 |
| AASS | 0.26 | -0.05 | -0.07 | -0.31 | 0.04 | 0.07 | 5.2E-01 | -0.11 |
| PRADC1 | -0.06 | -0.16 | 0.04 | 0.31 | -0.12 | -0.06 | 5.2E-01 | 0.10 |
| HSPA13 | -0.10 | -0.12 | 0.12 | 0.02 | -0.02 | 0.08 | 5.2E-01 | 0.06 |
| SLC25A23 | -0.11 | 0.07 | -0.03 | -0.01 | 0.05 | 0.01 | 5.2E-01 | 0.04 |
| MBD3 | 0.08 | -0.07 | 0.07 | -0.15 | 0.02 | 0.04 | 5.2E-01 | -0.06 |
| CORO2A | -0.04 | 0.00 | 0.01 | 0.00 | 0.05 | -0.03 | 5.2E-01 | 0.02 |
| BTBD9 | 0.05 | -0.02 | 0.09 | 0.22 | -0.19 | -0.20 | 5.2E-01 | -0.10 |
| ZNF592 | 0.04 | -0.03 | 0.03 | -0.06 | -0.01 | 0.03 | 5.2E-01 | -0.02 |
| UBE2I | 0.28 | 0.11 | -0.21 | 0.13 | 0.04 | -0.49 | 5.2E-01 | -0.17 |
| PREX1 | -0.12 | 0.08 | -0.08 | 0.09 | 0.14 | -0.14 | 5.3E-01 | 0.07 |
| NINJ1 | -0.09 | -0.05 | 0.04 | -0.11 | 0.03 | 0.17 | 5.3E-01 | 0.06 |
| RPF2 | -0.06 | -0.13 | 0.09 | 0.12 | -0.09 | 0.05 | 5.3E-01 | 0.06 |
| ESS2 | -0.05 | -0.04 | 0.04 | -0.03 | 0.07 | 0.00 | 5.3E-01 | 0.03 |
| INTS9 | -0.01 | 0.00 | -0.05 | -0.04 | -0.02 | 0.11 | 5.3E-01 | 0.04 |
| PLA2G4E | 0.24 | 0.19 | -0.21 | -0.71 | 0.15 | 0.13 | 5.3E-01 | -0.22 |

| | | | | | | | | |
|---|---|---|---|---|---|---|---|---|
| ATP6V1E1 | 0.04 | -0.01 | -0.06 | 0.06 | -0.01 | -0.01 | 5.3E-01 | 0.02 |
| TARS2 | -0.01 | -0.03 | 0.00 | 0.07 | -0.02 | -0.02 | 5.3E-01 | 0.02 |
| SMIM26 | 0.11 | 0.03 | -0.05 | -0.11 | -0.18 | 0.16 | 5.3E-01 | -0.08 |
| NUP35 | -0.02 | 0.00 | 0.09 | 0.09 | -0.07 | -0.10 | 5.3E-01 | -0.05 |
| UBL4A | -0.23 | -0.08 | 0.14 | 0.03 | -0.11 | 0.20 | 5.3E-01 | 0.10 |
| HSD17B10 | 0.05 | -0.01 | -0.12 | -0.08 | 0.07 | 0.07 | 5.3E-01 | 0.05 |
| MLF2 | -0.16 | -0.12 | 0.05 | 0.28 | -0.29 | 0.16 | 5.3E-01 | 0.13 |
| ITFG1 | -0.08 | -0.05 | 0.22 | -0.01 | -0.11 | 0.00 | 5.3E-01 | -0.07 |
| ZNF574 | -0.07 | -0.09 | 0.07 | -0.09 | 0.03 | 0.13 | 5.3E-01 | 0.06 |
| EPC1 | -0.05 | 0.06 | 0.08 | -0.21 | 0.03 | 0.07 | 5.3E-01 | -0.07 |
| WDR3 | 0.06 | 0.01 | -0.03 | 0.05 | -0.05 | -0.04 | 5.3E-01 | -0.03 |
| NOC4L | 0.02 | -0.09 | -0.01 | 0.16 | -0.04 | -0.05 | 5.3E-01 | 0.05 |
| KPNA4 | -0.05 | 0.03 | -0.04 | 0.10 | 0.02 | -0.07 | 5.3E-01 | 0.04 |
| MTMR1 | -0.04 | -0.05 | 0.04 | 0.05 | -0.07 | 0.06 | 5.3E-01 | 0.03 |
| YWHAH | -0.07 | 0.04 | -0.02 | -0.04 | 0.08 | 0.01 | 5.3E-01 | 0.03 |
| SCO1 | -0.13 | -0.15 | 0.10 | 0.18 | -0.22 | 0.17 | 5.3E-01 | 0.11 |
| SND1 | -0.02 | -0.04 | 0.00 | 0.13 | -0.01 | -0.06 | 5.3E-01 | 0.04 |
| ZRSR2 | -0.01 | -0.01 | 0.06 | 0.04 | -0.04 | -0.04 | 5.3E-01 | -0.02 |
| WASF2 | -0.02 | 0.09 | -0.01 | -0.09 | -0.03 | 0.06 | 5.3E-01 | -0.04 |
| RAB35 | -0.06 | 0.05 | 0.05 | -0.04 | 0.01 | -0.01 | 5.3E-01 | -0.02 |
| SLC25A16 | 0.03 | 0.05 | -0.03 | -0.01 | 0.06 | -0.12 | 5.3E-01 | -0.04 |
| MTIF3 | 0.08 | 0.01 | -0.01 | -0.20 | 0.02 | 0.08 | 5.3E-01 | -0.06 |
| PPM1L | 0.07 | 0.06 | -0.04 | 0.05 | 0.06 | -0.22 | 5.3E-01 | -0.07 |
| MFSD11 | -0.03 | 0.07 | 0.12 | 0.29 | -0.31 | -0.21 | 5.3E-01 | -0.13 |
| DLG1 | -0.10 | 0.08 | 0.08 | -0.08 | 0.00 | 0.01 | 5.3E-01 | -0.04 |
| PKNOX2 | 0.12 | -0.05 | -0.01 | -0.05 | 0.05 | -0.07 | 5.4E-01 | -0.04 |
| SAAL1 | 0.13 | 0.03 | -0.08 | -0.09 | 0.10 | -0.11 | 5.4E-01 | -0.06 |
| CLIP1 | -0.10 | -0.16 | 0.15 | -0.04 | 0.04 | 0.09 | 5.4E-01 | 0.07 |
| TRIM3 | -0.04 | -0.03 | 0.04 | 0.03 | 0.01 | -0.02 | 5.4E-01 | 0.02 |
| QDPR | -0.01 | 0.03 | -0.11 | -0.10 | 0.02 | 0.15 | 5.4E-01 | 0.06 |
| GHDC | 0.00 | 0.01 | 0.05 | 0.03 | 0.04 | -0.14 | 5.4E-01 | -0.04 |
| ACAD8 | 0.04 | 0.14 | -0.09 | -0.03 | 0.08 | -0.15 | 5.4E-01 | -0.06 |
| ITPR1 | 0.02 | 0.00 | -0.01 | 0.00 | 0.00 | -0.01 | 5.4E-01 | -0.01 |
| SMARCD2 | -0.02 | -0.02 | 0.02 | 0.01 | -0.03 | 0.04 | 5.4E-01 | 0.02 |
| PPP1CB | -0.08 | -0.06 | 0.06 | 0.13 | 0.00 | -0.06 | 5.4E-01 | 0.05 |
| CHAMP1 | 0.02 | 0.00 | -0.05 | -0.04 | 0.03 | 0.04 | 5.4E-01 | 0.02 |
| FOXJ3 | 0.03 | 0.15 | -0.07 | -0.28 | 0.13 | -0.01 | 5.4E-01 | -0.09 |
| YTHDF3 | -0.08 | 0.12 | 0.02 | 0.05 | -0.06 | -0.06 | 5.4E-01 | -0.05 |
| LDLR | 0.06 | -0.03 | 0.09 | -0.31 | 0.16 | -0.01 | 5.4E-01 | -0.09 |
| RRM2B | -0.53 | 0.09 | 0.06 | 0.08 | 0.42 | -0.30 | 5.4E-01 | 0.19 |
| SIX5 | -0.04 | 0.13 | 0.15 | -0.86 | 0.15 | 0.22 | 5.4E-01 | -0.24 |
| GDI2 | 0.04 | 0.02 | 0.00 | -0.07 | -0.11 | 0.11 | 5.4E-01 | -0.05 |
| EAPP | 0.06 | 0.00 | 0.02 | 0.14 | -0.09 | -0.15 | 5.4E-01 | -0.06 |
| PRKAA1 | -0.02 | 0.08 | -0.19 | 0.10 | 0.12 | -0.12 | 5.4E-01 | 0.07 |
| GTF2A2 | 0.10 | -0.04 | -0.14 | 0.02 | -0.02 | 0.07 | 5.4E-01 | 0.05 |
| CPT1A | 0.12 | -0.03 | -0.01 | 0.04 | 0.03 | -0.15 | 5.4E-01 | -0.05 |
| EFR3A | -0.06 | 0.07 | 0.04 | -0.05 | -0.07 | 0.06 | 5.4E-01 | -0.04 |
| THOC3 | 0.06 | 0.08 | -0.07 | 0.03 | 0.02 | -0.13 | 5.4E-01 | -0.05 |

| | | | | | | | | |
|---|---|---|---|---|---|---|---|---|
| TM9SF3 | -0.03 | -0.02 | 0.00 | 0.10 | 0.02 | -0.07 | 5.4E-01 | 0.03 |
| PRMT1 | 0.09 | 0.04 | -0.07 | -0.07 | 0.06 | -0.06 | 5.4E-01 | -0.04 |
| GAS2L3 | -0.27 | 0.38 | 0.05 | 0.03 | -0.23 | -0.05 | 5.4E-01 | -0.13 |
| CTNND1 | -0.05 | -0.01 | 0.03 | 0.03 | -0.01 | 0.01 | 5.4E-01 | 0.02 |
| HNRNPH2 | 0.00 | -0.05 | 0.02 | 0.01 | 0.05 | -0.03 | 5.4E-01 | 0.02 |
| ATP13A2 | -0.05 | -0.07 | 0.05 | -0.08 | 0.02 | 0.12 | 5.4E-01 | 0.04 |
| SLC35F2 | -0.07 | -0.10 | 0.10 | 0.02 | 0.06 | -0.02 | 5.4E-01 | 0.04 |
| THEM6 | -0.05 | -0.04 | 0.06 | 0.02 | -0.01 | 0.02 | 5.5E-01 | 0.02 |
| METTL2B | -0.02 | 0.02 | -0.08 | -0.08 | 0.15 | 0.00 | 5.5E-01 | 0.05 |
| NSUN5 | -0.04 | -0.02 | 0.03 | 0.05 | 0.00 | -0.02 | 5.5E-01 | 0.02 |
| IFIT1 | 0.04 | -0.09 | -0.11 | 0.12 | 0.20 | -0.21 | 5.5E-01 | 0.09 |
| DPF3 | 0.03 | 0.03 | -0.04 | -0.04 | 0.01 | 0.00 | 5.5E-01 | -0.02 |
| SLC2A1 | -0.21 | -0.11 | 0.17 | -0.07 | 0.02 | 0.16 | 5.5E-01 | 0.09 |
| WDR36 | 0.04 | -0.02 | -0.05 | 0.00 | 0.02 | 0.00 | 5.5E-01 | 0.02 |
| COPS3 | 0.03 | 0.02 | 0.01 | -0.13 | 0.10 | -0.04 | 5.5E-01 | -0.04 |
| USP16 | -0.15 | -0.01 | 0.29 | -0.20 | 0.01 | 0.01 | 5.5E-01 | -0.10 |
| HOXC11 | 0.06 | 0.01 | -0.01 | 0.09 | -0.07 | -0.09 | 5.5E-01 | -0.04 |
| NDUFB10 | 0.04 | 0.02 | -0.15 | 0.14 | -0.04 | -0.03 | 5.5E-01 | 0.05 |
| HTATSF1 | 0.10 | 0.05 | -0.06 | 0.10 | 0.00 | -0.22 | 5.5E-01 | -0.07 |
| CANT1 | 0.01 | 0.09 | -0.03 | 0.10 | -0.07 | -0.11 | 5.5E-01 | -0.05 |
| KIF1C | -0.07 | -0.09 | 0.27 | 0.00 | -0.14 | -0.01 | 5.5E-01 | -0.08 |
| NONO | 0.03 | -0.01 | 0.02 | -0.03 | 0.06 | -0.06 | 5.5E-01 | -0.03 |
| PPIE | 0.02 | -0.04 | -0.04 | 0.11 | -0.08 | 0.02 | 5.5E-01 | 0.04 |
| QSOX1 | -0.08 | -0.06 | 0.08 | 0.01 | -0.04 | 0.10 | 5.5E-01 | 0.04 |
| NDUFS6 | 0.08 | 0.04 | -0.06 | 0.09 | -0.06 | -0.11 | 5.5E-01 | -0.05 |
| DOCK6 | -0.37 | 0.09 | 0.08 | 0.06 | -0.08 | 0.15 | 5.5E-01 | 0.11 |
| GAR1 | 0.10 | 0.03 | -0.04 | 0.14 | -0.07 | -0.19 | 5.5E-01 | -0.07 |
| ZFX | 0.02 | -0.01 | -0.04 | 0.01 | -0.04 | 0.06 | 5.5E-01 | 0.02 |
| PSMB6 | 0.02 | -0.02 | 0.08 | -0.01 | -0.20 | 0.11 | 5.5E-01 | -0.06 |
| GMCL1 | 0.05 | -0.05 | -0.08 | 0.01 | -0.07 | 0.13 | 5.5E-01 | 0.05 |
| PIGW | -0.07 | -0.11 | 0.09 | -0.08 | 0.03 | 0.13 | 5.5E-01 | 0.06 |
| PSEN2 | -0.02 | 0.13 | 0.00 | 0.09 | -0.27 | 0.04 | 5.5E-01 | -0.08 |
| CRYZL1 | -0.29 | 0.17 | -0.11 | -0.18 | 0.28 | 0.03 | 5.5E-01 | 0.12 |
| BAP1 | -0.09 | -0.13 | 0.10 | -0.14 | 0.10 | 0.14 | 5.5E-01 | 0.07 |
| APMAP | 0.10 | 0.08 | -0.11 | -0.05 | -0.08 | 0.04 | 5.5E-01 | -0.05 |
| HNRNPUL2 | 0.04 | -0.02 | 0.00 | -0.02 | 0.00 | 0.00 | 5.5E-01 | -0.01 |
| RC3H1 | -0.08 | -0.09 | 0.26 | -0.10 | -0.08 | 0.04 | 5.5E-01 | -0.08 |
| PRMT5 | 0.11 | -0.01 | -0.28 | 0.18 | 0.12 | -0.17 | 5.6E-01 | 0.10 |
| APOO | 0.01 | 0.00 | -0.04 | 0.04 | 0.03 | -0.04 | 5.6E-01 | 0.02 |
| MAU2 | -0.04 | -0.02 | 0.00 | -0.06 | -0.02 | 0.14 | 5.6E-01 | 0.04 |
| GCAT | 0.11 | 0.03 | -0.07 | -0.02 | 0.08 | -0.15 | 5.6E-01 | -0.05 |
| ESRP1 | -0.03 | -0.02 | 0.01 | -0.07 | 0.08 | 0.03 | 5.6E-01 | 0.03 |
| SCAI | 0.06 | -0.01 | -0.03 | -0.02 | 0.00 | -0.01 | 5.6E-01 | -0.02 |
| ANO10 | 0.06 | 0.08 | -0.05 | 0.14 | -0.09 | -0.16 | 5.6E-01 | -0.06 |
| CROT | 0.00 | -0.09 | 0.01 | -0.04 | 0.16 | -0.06 | 5.6E-01 | 0.05 |
| PITX1 | -0.06 | -0.05 | 0.01 | 0.18 | -0.13 | 0.03 | 5.6E-01 | 0.06 |
| SNAPIN | -0.02 | -0.14 | 0.05 | 0.03 | -0.12 | 0.18 | 5.6E-01 | 0.07 |
| RFTN1 | -0.09 | -0.04 | 0.07 | -0.02 | 0.10 | -0.03 | 5.6E-01 | 0.04 |

| | | | | | | | | |
|---|---|---|---|---|---|---|---|---|
| ANK3 | -0.09 | 0.01 | 0.03 | 0.03 | 0.05 | -0.04 | 5.6E-01 | 0.03 |
| NDUFAF2 | -0.01 | 0.01 | -0.03 | 0.04 | 0.01 | -0.03 | 5.6E-01 | 0.02 |
| HNRNPA1 | 0.07 | -0.08 | -0.77 | 0.88 | -0.52 | -0.20 | 5.6E-01 | 0.32 |
| DNASE2 | -0.04 | 0.33 | -0.16 | -0.17 | -0.06 | 0.04 | 5.6E-01 | -0.10 |
| ANKRD17 | 0.01 | -0.13 | 0.00 | 0.13 | -0.17 | 0.13 | 5.6E-01 | 0.07 |
| RPL3 | -0.01 | -0.05 | 0.02 | 0.06 | 0.04 | -0.06 | 5.6E-01 | 0.03 |
| LARGE1 | -0.03 | 0.01 | 0.05 | -0.03 | 0.05 | -0.06 | 5.6E-01 | -0.03 |
| DAP | 0.10 | 0.21 | -0.12 | 0.25 | 0.05 | -0.68 | 5.6E-01 | -0.19 |
| RTCA | -0.08 | 0.13 | -0.25 | -0.17 | 0.29 | 0.01 | 5.6E-01 | 0.11 |
| ALG10 | -0.09 | -0.01 | 0.19 | -0.15 | 0.00 | 0.04 | 5.6E-01 | -0.06 |
| MYO1B | -0.07 | -0.02 | 0.15 | -0.01 | -0.05 | -0.01 | 5.6E-01 | -0.04 |
| TMEM167A | -0.03 | -0.01 | 0.08 | -0.02 | -0.06 | 0.04 | 5.6E-01 | -0.03 |
| RPL37 | -0.02 | -0.02 | 0.13 | 0.05 | 0.05 | -0.20 | 5.6E-01 | -0.06 |
| PPAN | 0.03 | -0.04 | 0.03 | 0.01 | 0.00 | -0.04 | 5.6E-01 | -0.02 |
| RTRAF | 0.01 | -0.03 | -0.04 | 0.07 | 0.06 | -0.09 | 5.6E-01 | 0.03 |
| KNL1 | -0.07 | -0.06 | 0.08 | 0.05 | -0.01 | 0.00 | 5.6E-01 | 0.03 |
| RHBDD3 | -0.10 | 0.05 | 0.12 | 0.01 | 0.04 | -0.14 | 5.7E-01 | -0.05 |
| EXOSC1 | 0.05 | 0.12 | -0.06 | 0.22 | -0.20 | -0.17 | 5.7E-01 | -0.09 |
| NLRX1 | 0.00 | -0.02 | 0.12 | -0.08 | 0.18 | -0.24 | 5.7E-01 | -0.08 |
| SPTY2D1 | -0.01 | 0.03 | 0.04 | 0.10 | -0.11 | -0.06 | 5.7E-01 | -0.04 |
| HNRNPA3 | -0.05 | -0.07 | 0.20 | -0.10 | -0.02 | 0.02 | 5.7E-01 | -0.06 |
| GOLGA7 | -0.24 | 0.01 | 0.12 | 0.01 | 0.05 | 0.03 | 5.7E-01 | 0.07 |
| DDX19A | -0.04 | 0.01 | 0.05 | -0.01 | -0.05 | 0.03 | 5.7E-01 | -0.02 |
| UTP25 | 0.06 | -0.11 | 0.00 | 0.01 | -0.02 | 0.06 | 5.7E-01 | 0.03 |
| WDR83 | 0.06 | -0.06 | 0.05 | 0.08 | -0.08 | -0.05 | 5.7E-01 | -0.04 |
| ARID4B | -0.03 | 0.05 | 0.02 | 0.04 | -0.08 | 0.00 | 5.7E-01 | -0.03 |
| AHCTF1 | 0.01 | -0.02 | -0.02 | -0.01 | -0.03 | 0.06 | 5.7E-01 | 0.02 |
| MITF | -0.03 | -0.17 | 0.09 | -0.12 | 0.09 | 0.12 | 5.7E-01 | 0.07 |
| PXMP2 | -0.10 | -0.11 | 0.10 | -0.10 | 0.02 | 0.17 | 5.7E-01 | 0.06 |
| PODXL | -0.08 | -0.01 | 0.05 | -0.03 | 0.03 | 0.04 | 5.7E-01 | 0.03 |
| CCNB1 | 0.09 | 0.17 | -0.17 | -0.14 | 0.04 | -0.02 | 5.7E-01 | -0.07 |
| RANBP2 | 0.06 | 0.00 | 0.00 | 0.08 | 0.01 | -0.16 | 5.7E-01 | -0.05 |
| PRDM10 | 0.01 | 0.02 | -0.07 | -0.03 | -0.01 | 0.08 | 5.7E-01 | 0.03 |
| TASOR | -0.01 | -0.04 | 0.01 | -0.01 | -0.03 | 0.08 | 5.7E-01 | 0.02 |
| PAF1 | 0.13 | 0.05 | -0.10 | 0.01 | 0.05 | -0.15 | 5.7E-01 | -0.06 |
| HSPA5 | 0.05 | 0.00 | -0.09 | -0.01 | 0.05 | 0.00 | 5.7E-01 | 0.03 |
| TAMM41 | -0.23 | -0.30 | 0.31 | -0.04 | 0.10 | 0.08 | 5.7E-01 | 0.12 |
| ZNHIT3 | -0.09 | -0.03 | 0.20 | -0.08 | 0.06 | -0.09 | 5.7E-01 | -0.06 |
| CHD8 | 0.07 | -0.02 | 0.00 | -0.13 | -0.01 | 0.07 | 5.7E-01 | -0.04 |
| F11 | 0.00 | -0.09 | 0.04 | -0.01 | -0.04 | 0.10 | 5.7E-01 | 0.03 |
| ATP9A | -0.01 | 0.23 | -0.11 | -0.20 | 0.05 | 0.01 | 5.7E-01 | -0.08 |
| PSEN1 | -0.08 | 0.04 | 0.09 | -0.05 | -0.04 | 0.04 | 5.7E-01 | -0.04 |
| RPS28 | -0.05 | -0.07 | 0.20 | -0.17 | 0.01 | 0.05 | 5.7E-01 | -0.07 |
| ING2 | 0.04 | 0.09 | -0.07 | -0.09 | 0.07 | -0.06 | 5.7E-01 | -0.04 |
| RPL32 | -0.14 | -0.06 | 0.32 | -0.01 | 0.01 | -0.17 | 5.7E-01 | -0.09 |
| RPL27 | -0.01 | -0.03 | -0.05 | 0.20 | -0.02 | -0.11 | 5.8E-01 | 0.06 |
| CNNM4 | -0.05 | 0.01 | 0.11 | 0.08 | -0.03 | -0.13 | 5.8E-01 | -0.05 |
| ZNF557 | 0.10 | -0.01 | 0.00 | -0.22 | 0.04 | 0.07 | 5.8E-01 | -0.06 |

| | | | | | | | | |
|---|---|---|---|---|---|---|---|---|
| MED25 | 0.14 | -0.07 | 0.03 | -0.27 | 0.06 | 0.07 | 5.8E-01 | -0.08 |
| UQCC2 | -0.02 | 0.17 | -0.05 | 0.13 | 0.00 | -0.28 | 5.8E-01 | -0.08 |
| MPZL1 | 0.02 | -0.02 | 0.12 | -0.27 | -0.12 | 0.23 | 5.8E-01 | -0.09 |
| GATAD1 | -0.02 | -0.01 | 0.06 | -0.06 | -0.02 | 0.05 | 5.8E-01 | -0.02 |
| LANCL2 | 0.13 | -0.07 | -0.18 | -0.09 | 0.04 | 0.14 | 5.8E-01 | 0.07 |
| CENPM | -0.11 | 0.00 | 0.02 | 0.03 | -0.12 | 0.18 | 5.8E-01 | 0.06 |
| DUS3L | -0.13 | 0.00 | 0.07 | 0.00 | -0.01 | 0.06 | 5.8E-01 | 0.04 |
| NDUFS1 | 0.00 | -0.01 | 0.03 | 0.02 | -0.02 | -0.02 | 5.8E-01 | -0.01 |
| PDLIM7 | 0.02 | -0.05 | 0.07 | 0.05 | -0.10 | -0.01 | 5.8E-01 | -0.03 |
| RPL12 | 0.01 | -0.03 | 0.06 | 0.04 | -0.01 | -0.07 | 5.8E-01 | -0.02 |
| GPR107 | -0.13 | 0.06 | 0.13 | 0.01 | -0.05 | -0.03 | 5.8E-01 | -0.05 |
| C11orf52 | -0.05 | 0.12 | 0.01 | -0.17 | -0.05 | 0.12 | 5.8E-01 | -0.06 |
| VPS4B | 0.05 | -0.07 | -0.05 | 0.07 | 0.07 | -0.09 | 5.8E-01 | 0.04 |
| TMPO | 0.02 | -0.04 | 0.00 | 0.00 | -0.03 | 0.05 | 5.8E-01 | 0.02 |
| FAAH | 0.06 | 0.07 | -0.02 | 0.22 | -0.24 | -0.13 | 5.8E-01 | -0.08 |
| BNIP3 | -0.03 | -0.03 | 0.11 | -0.01 | -0.10 | 0.05 | 5.8E-01 | -0.04 |
| G2E3 | 0.06 | -0.23 | 0.06 | 0.13 | -0.01 | -0.04 | 5.8E-01 | 0.07 |
| CASP3 | 0.00 | -0.01 | -0.08 | -0.14 | 0.20 | 0.01 | 5.8E-01 | 0.06 |
| RNF114 | -0.16 | 0.03 | 0.05 | 0.04 | -0.05 | 0.08 | 5.8E-01 | 0.05 |
| C7orf50 | -0.04 | 0.03 | 0.06 | 0.04 | 0.01 | -0.10 | 5.8E-01 | -0.03 |
| PTPRG | -0.09 | 0.07 | 0.07 | -0.03 | 0.02 | -0.05 | 5.8E-01 | -0.04 |
| SCO2 | -0.13 | -0.04 | 0.07 | 0.04 | -0.12 | 0.15 | 5.8E-01 | 0.06 |
| MYOF | -0.08 | 0.00 | 0.13 | 0.01 | -0.04 | -0.03 | 5.8E-01 | -0.04 |
| PLEKHA6 | -0.03 | 0.03 | 0.04 | 0.05 | -0.09 | 0.00 | 5.8E-01 | -0.03 |
| CWC15 | 0.05 | -0.07 | -0.05 | -0.04 | -0.03 | 0.13 | 5.9E-01 | 0.04 |
| TVP23C | -0.02 | 0.00 | 0.06 | -0.09 | 0.06 | -0.02 | 5.9E-01 | -0.03 |
| PGRMC1 | -0.10 | 0.08 | -0.08 | 0.05 | -0.11 | 0.14 | 5.9E-01 | 0.06 |
| STK16 | -0.09 | 0.04 | 0.10 | 0.01 | -0.02 | -0.05 | 5.9E-01 | -0.03 |
| HNRNPC | 0.01 | -0.02 | -0.03 | -0.04 | -0.02 | 0.10 | 5.9E-01 | 0.03 |
| ELAPOR1 | -0.03 | -0.08 | 0.04 | -0.06 | -0.02 | 0.14 | 5.9E-01 | 0.04 |
| ATP6V1D | -0.01 | 0.07 | 0.00 | 0.10 | -0.08 | -0.08 | 5.9E-01 | -0.04 |
| KIF2A | 0.05 | 0.01 | -0.02 | -0.07 | 0.08 | -0.06 | 5.9E-01 | -0.03 |
| ZBTB40 | 0.00 | 0.01 | -0.06 | -0.05 | 0.00 | 0.09 | 5.9E-01 | 0.03 |
| B3GNT2 | -0.06 | 0.07 | -0.05 | 0.00 | 0.05 | -0.01 | 5.9E-01 | 0.03 |
| PI4KA | -0.08 | -0.04 | 0.06 | 0.01 | -0.06 | 0.10 | 5.9E-01 | 0.04 |
| ZFP2 | 0.13 | -0.07 | 0.01 | 0.04 | 0.04 | -0.18 | 5.9E-01 | -0.06 |
| SDC4 | -0.08 | 0.09 | 0.03 | 0.02 | -0.03 | -0.04 | 5.9E-01 | -0.03 |
| TRAP1 | 0.10 | 0.01 | -0.19 | 0.00 | 0.07 | 0.00 | 5.9E-01 | 0.05 |
| HADH | 0.09 | -0.05 | -0.10 | 0.03 | -0.03 | 0.04 | 5.9E-01 | 0.04 |
| DDX10 | 0.01 | -0.02 | -0.01 | -0.01 | -0.02 | 0.05 | 5.9E-01 | 0.01 |
| CENPI | 0.03 | -0.08 | 0.00 | 0.04 | -0.08 | 0.09 | 5.9E-01 | 0.03 |
| SUDS3 | 0.02 | -0.04 | -0.06 | 0.14 | -0.14 | 0.06 | 6.0E-01 | 0.05 |
| ZBTB4 | -0.10 | 0.01 | 0.05 | 0.03 | -0.04 | 0.05 | 6.0E-01 | 0.03 |
| OAS3 | 0.05 | 0.04 | -0.05 | -0.08 | 0.08 | -0.05 | 6.0E-01 | -0.03 |
| POLR3K | -0.10 | 0.00 | 0.25 | -0.53 | 0.07 | 0.18 | 6.0E-01 | -0.14 |
| UBQLN1 | -0.13 | -0.09 | 0.11 | -0.14 | 0.16 | 0.06 | 6.0E-01 | 0.07 |
| THBS1 | -0.01 | -0.05 | 0.03 | -0.01 | -0.01 | 0.05 | 6.0E-01 | 0.02 |
| OXR1 | -0.01 | 0.01 | 0.07 | -0.19 | -0.02 | 0.12 | 6.0E-01 | -0.05 |

| | | | | | | | | |
|---|---|---|---|---|---|---|---|---|
| DMBT1 | 0.02 | -0.12 | 0.02 | -0.12 | 0.07 | 0.12 | 6.0E-01 | 0.05 |
| PRPF31 | 0.10 | -0.06 | -0.09 | 0.02 | 0.04 | -0.01 | 6.0E-01 | 0.03 |
| RBM15 | -0.02 | -0.05 | 0.04 | 0.04 | -0.02 | 0.01 | 6.0E-01 | 0.02 |
| CCT3 | 0.04 | 0.16 | -0.13 | -0.03 | 0.01 | -0.07 | 6.0E-01 | -0.05 |
| ABCB10 | -0.39 | -0.30 | 0.39 | -0.07 | -0.03 | 0.24 | 6.0E-01 | 0.15 |
| PARP16 | -0.10 | -0.14 | 0.11 | -0.02 | -0.13 | 0.25 | 6.0E-01 | 0.08 |
| UBE4A | -0.10 | -0.02 | 0.21 | -0.19 | 0.09 | -0.02 | 6.0E-01 | -0.07 |
| HAUS8 | -0.36 | -0.21 | 0.33 | -0.04 | 0.05 | 0.12 | 6.0E-01 | 0.12 |
| MFF | 0.00 | 0.06 | -0.02 | -0.07 | -0.05 | 0.07 | 6.0E-01 | -0.03 |
| KAT8 | 0.02 | -0.02 | -0.03 | 0.00 | -0.03 | 0.06 | 6.0E-01 | 0.02 |
| WRAP53 | -0.27 | 0.13 | 0.01 | 0.06 | -0.10 | 0.13 | 6.0E-01 | 0.08 |
| SIPA1L3 | -0.07 | 0.05 | -0.03 | -0.04 | 0.03 | 0.04 | 6.0E-01 | 0.03 |
| CALM3 | 0.01 | 0.03 | -0.02 | 0.03 | -0.06 | 0.01 | 6.0E-01 | -0.02 |
| ABI2 | -0.10 | -0.21 | 0.43 | -0.06 | -0.12 | -0.03 | 6.0E-01 | -0.11 |
| UBE3B | -0.15 | -0.03 | 0.11 | -0.04 | 0.02 | 0.09 | 6.0E-01 | 0.05 |
| TAF8 | 0.00 | 0.03 | -0.08 | 0.01 | -0.07 | 0.11 | 6.0E-01 | 0.03 |
| MGST3 | 0.04 | -0.04 | 0.03 | 0.00 | 0.00 | -0.03 | 6.0E-01 | -0.01 |
| DARS2 | 0.15 | 0.07 | -0.14 | -0.11 | 0.04 | -0.02 | 6.0E-01 | -0.06 |
| RDH16 | -0.13 | 0.07 | 0.14 | -0.14 | 0.02 | 0.03 | 6.0E-01 | -0.06 |
| TTC3 | -0.17 | 0.10 | -0.04 | -0.01 | -0.10 | 0.19 | 6.0E-01 | 0.06 |
| ZDHHC21 | -0.25 | -0.12 | 0.19 | -0.01 | -0.13 | 0.25 | 6.0E-01 | 0.10 |
| ATP6V1C1 | 0.03 | 0.02 | -0.02 | 0.04 | -0.06 | -0.01 | 6.0E-01 | -0.02 |
| ADCK1 | 0.03 | -0.06 | -0.05 | -0.11 | 0.18 | 0.00 | 6.0E-01 | 0.05 |
| CTPS1 | -0.45 | 0.89 | -0.31 | -0.15 | -0.22 | -0.23 | 6.1E-01 | -0.24 |
| C19orf18 | -0.04 | 0.15 | 0.03 | 0.05 | 0.21 | -0.49 | 6.1E-01 | -0.12 |
| RAB6A | -0.07 | -0.01 | 0.03 | 0.05 | -0.07 | 0.06 | 6.1E-01 | 0.03 |
| TRAF2 | 0.00 | -0.01 | 0.05 | 0.01 | 0.05 | -0.10 | 6.1E-01 | -0.03 |
| BRD9 | 0.10 | 0.02 | -0.27 | 0.19 | 0.01 | -0.10 | 6.1E-01 | 0.08 |
| MOCS3 | -0.12 | 0.07 | -0.02 | -0.04 | -0.02 | 0.12 | 6.1E-01 | 0.04 |
| FBXO11 | -0.19 | -0.06 | 0.11 | 0.17 | -0.18 | 0.10 | 6.1E-01 | 0.08 |
| UNC93B1 | 0.18 | 0.10 | -0.18 | 0.15 | -0.19 | -0.10 | 6.1E-01 | -0.08 |
| RPL17 | -0.05 | -0.07 | 0.04 | 0.17 | -0.04 | -0.07 | 6.1E-01 | 0.05 |
| DMAC2 | -0.14 | 0.03 | 0.05 | 0.03 | -0.05 | 0.08 | 6.1E-01 | 0.04 |
| CENPK | -0.01 | -0.07 | -0.01 | -0.03 | -0.11 | 0.20 | 6.1E-01 | 0.05 |
| PYCR2 | 0.09 | 0.06 | -0.09 | 0.07 | -0.02 | -0.13 | 6.1E-01 | -0.04 |
| SMPDL3B | -0.22 | -0.01 | 0.13 | -0.05 | 0.00 | 0.12 | 6.1E-01 | 0.06 |
| ZDHHC17 | 0.02 | 0.07 | -0.01 | 0.15 | -0.04 | -0.21 | 6.1E-01 | -0.06 |
| NIT2 | 0.18 | -0.03 | -0.08 | -0.08 | 0.11 | -0.13 | 6.1E-01 | -0.06 |
| SLC38A10 | -0.06 | 0.05 | 0.05 | 0.01 | 0.03 | -0.09 | 6.1E-01 | -0.03 |
| NAA15 | 0.05 | -0.03 | -0.07 | 0.08 | 0.05 | -0.08 | 6.1E-01 | 0.03 |
| STX10 | -0.05 | -0.07 | 0.20 | 0.07 | -0.13 | -0.04 | 6.1E-01 | -0.06 |
| MAN2A1 | 0.04 | 0.04 | -0.01 | 0.16 | -0.10 | -0.15 | 6.1E-01 | -0.05 |
| HSD17B8 | 0.11 | -0.02 | -0.16 | 0.03 | 0.05 | -0.02 | 6.1E-01 | 0.04 |
| DLGAP5 | 0.08 | -0.10 | -0.04 | -0.06 | 0.03 | 0.08 | 6.2E-01 | 0.04 |
| GTF2H4 | 0.01 | 0.02 | -0.09 | 0.12 | 0.01 | -0.08 | 6.2E-01 | 0.04 |
| NR2F6 | -0.01 | -0.03 | 0.00 | 0.07 | 0.02 | -0.06 | 6.2E-01 | 0.02 |
| CFAP298 | -0.06 | 0.04 | -0.06 | -0.07 | -0.04 | 0.17 | 6.2E-01 | 0.05 |
| PCGF2 | 0.15 | -0.19 | 0.12 | 0.05 | -0.14 | -0.02 | 6.2E-01 | -0.06 |

| | | | | | | | | |
|---|---|---|---|---|---|---|---|---|
| TXNRD2 | 0.18 | 0.05 | -0.16 | 0.01 | -0.06 | -0.03 | 6.2E-01 | -0.05 |
| SUGT1 | -0.02 | 0.03 | -0.09 | 0.15 | 0.04 | -0.13 | 6.2E-01 | 0.05 |
| SCML2 | -0.01 | -0.01 | 0.06 | -0.10 | -0.01 | 0.06 | 6.2E-01 | -0.03 |
| ATF6 | -0.16 | -0.13 | 0.15 | 0.15 | 0.10 | -0.16 | 6.2E-01 | 0.07 |
| MANF | -0.03 | -0.04 | 0.01 | 0.03 | -0.11 | 0.13 | 6.2E-01 | 0.04 |
| GATAD2A | 0.02 | 0.00 | 0.01 | -0.07 | 0.02 | 0.02 | 6.2E-01 | -0.02 |
| ATP2B1 | -0.06 | -0.06 | 0.18 | -0.04 | -0.03 | 0.00 | 6.2E-01 | -0.04 |
| MARK4 | -0.33 | -0.15 | 0.61 | 0.04 | -0.35 | -0.05 | 6.2E-01 | -0.17 |
| NDUFV3 | 0.02 | 0.02 | -0.10 | 0.12 | -0.02 | -0.05 | 6.2E-01 | 0.03 |
| ZNF236 | -0.04 | 0.03 | -0.05 | -0.06 | -0.04 | 0.14 | 6.2E-01 | 0.04 |
| STXBP4 | -0.23 | 0.00 | 0.35 | -0.35 | -0.05 | 0.16 | 6.2E-01 | -0.12 |
| ACTR1A | 0.09 | 0.08 | -0.09 | 0.10 | 0.04 | -0.24 | 6.2E-01 | -0.06 |
| NFYB | 0.16 | 0.03 | -0.09 | -0.31 | 0.10 | 0.07 | 6.2E-01 | -0.08 |
| TMEM209 | 0.07 | -0.07 | -0.11 | 0.26 | -0.11 | -0.07 | 6.3E-01 | 0.07 |
| POC5 | -0.09 | -0.05 | 0.24 | -0.10 | -0.28 | 0.21 | 6.3E-01 | -0.09 |
| ANP32B | 0.04 | 0.18 | -0.15 | -0.14 | 0.11 | -0.08 | 6.3E-01 | -0.06 |
| MTG2 | 0.10 | 0.07 | -0.12 | 0.03 | -0.08 | -0.02 | 6.3E-01 | -0.04 |
| ERCC3 | -0.01 | 0.05 | -0.16 | 0.23 | -0.03 | -0.12 | 6.3E-01 | 0.06 |
| RUNDC1 | -0.08 | -0.08 | 0.10 | 0.07 | -0.03 | 0.02 | 6.3E-01 | 0.04 |
| CNTNAP2 | -0.55 | 0.17 | 0.51 | -0.16 | -0.13 | -0.06 | 6.3E-01 | -0.16 |
| TBK1 | -0.03 | 0.03 | 0.05 | 0.04 | 0.05 | -0.14 | 6.3E-01 | -0.03 |
| COX4I1 | 0.05 | 0.03 | -0.14 | 0.07 | 0.00 | -0.03 | 6.3E-01 | 0.03 |
| RAD23A | -0.09 | -0.03 | 0.06 | -0.10 | 0.10 | 0.06 | 6.3E-01 | 0.04 |
| MORF4L2 | -0.05 | 0.06 | 0.02 | -0.03 | 0.01 | -0.02 | 6.3E-01 | -0.02 |
| ACOT13 | 0.15 | 0.21 | -0.26 | -0.18 | -0.01 | 0.03 | 6.3E-01 | -0.08 |
| NDUFAF1 | 0.02 | 0.05 | -0.03 | 0.05 | 0.00 | -0.09 | 6.3E-01 | -0.02 |
| COL23A1 | -0.25 | -0.14 | 0.13 | -0.09 | -0.26 | 0.47 | 6.3E-01 | 0.13 |
| CCDC77 | -0.12 | 0.03 | 0.03 | -0.08 | 0.03 | 0.10 | 6.3E-01 | 0.04 |
| SMAP | -0.16 | -0.27 | -0.55 | -0.77 | -0.44 | 1.18 | 6.3E-01 | 0.32 |
| GALNT3 | -0.01 | -0.06 | -0.01 | 0.21 | -0.10 | -0.05 | 6.3E-01 | 0.05 |
| ATP5MJ | 0.06 | 0.04 | -0.02 | -0.29 | 0.04 | 0.13 | 6.3E-01 | -0.07 |
| DGKZ | 0.02 | -0.08 | 0.17 | -0.28 | -0.18 | 0.27 | 6.3E-01 | -0.10 |
| AP2A1 | -0.03 | -0.01 | 0.01 | 0.09 | 0.00 | -0.06 | 6.3E-01 | 0.02 |
| NDUFB8 | 0.11 | -0.08 | -0.32 | 0.50 | -0.13 | -0.25 | 6.3E-01 | 0.14 |
| FNDC3A | 0.03 | -0.08 | -0.03 | 0.16 | 0.03 | -0.13 | 6.3E-01 | 0.05 |
| RPL14 | -0.04 | -0.07 | 0.07 | 0.06 | 0.00 | -0.02 | 6.3E-01 | 0.03 |
| NOVA2 | 0.04 | -0.09 | -0.02 | -0.10 | 0.01 | 0.15 | 6.3E-01 | 0.04 |
| NUFIP2 | -0.05 | 0.05 | 0.05 | 0.10 | -0.05 | -0.10 | 6.3E-01 | -0.04 |
| CENPH | -0.02 | -0.04 | 0.01 | -0.05 | -0.03 | 0.11 | 6.3E-01 | 0.03 |
| COX7A2L | 0.07 | 0.05 | -0.06 | 0.12 | -0.02 | -0.17 | 6.3E-01 | -0.05 |
| MCRIP2 | -0.07 | 0.04 | 0.12 | -0.14 | -0.25 | 0.24 | 6.3E-01 | -0.08 |
| ECI1 | 0.24 | 0.18 | -0.31 | -0.04 | -0.25 | 0.09 | 6.3E-01 | -0.10 |
| IQGAP3 | -0.06 | -0.01 | 0.04 | 0.00 | -0.01 | 0.04 | 6.3E-01 | 0.02 |
| SUPT16H | 0.00 | -0.04 | -0.01 | -0.08 | 0.02 | 0.11 | 6.3E-01 | 0.03 |
| SERPINH1 | 0.05 | -0.01 | -0.07 | 0.04 | 0.01 | -0.02 | 6.4E-01 | 0.02 |
| PRKCI | -0.08 | -0.12 | 0.08 | -0.08 | 0.25 | -0.09 | 6.4E-01 | 0.07 |
| PLS3 | -0.08 | 0.07 | -0.09 | -0.08 | -0.07 | 0.23 | 6.4E-01 | 0.06 |
| NCSTN | -0.02 | 0.04 | -0.16 | -0.24 | 0.11 | 0.22 | 6.4E-01 | 0.08 |

| | | | | | | | | |
|---|---|---|---|---|---|---|---|---|
| **FOXRED2** | 0.09 | -0.01 | 0.01 | -0.35 | 0.03 | 0.18 | 6.4E-01 | -0.08 |
| **ZNF28** | 0.06 | -0.15 | 0.17 | 0.01 | -0.26 | 0.11 | 6.4E-01 | -0.07 |
| **MED24** | 0.05 | 0.07 | -0.07 | 0.09 | -0.08 | -0.07 | 6.4E-01 | -0.04 |
| **UBXN4** | 0.08 | -0.02 | -0.15 | 0.01 | -0.08 | 0.13 | 6.4E-01 | 0.05 |
| **POLG** | -0.02 | 0.01 | 0.05 | 0.06 | -0.06 | -0.04 | 6.4E-01 | -0.02 |
| **CEP135** | -0.08 | -0.03 | 0.15 | -0.06 | -0.01 | 0.01 | 6.4E-01 | -0.04 |
| **CEP290** | 0.00 | 0.00 | 0.06 | -0.13 | -0.10 | 0.15 | 6.4E-01 | -0.05 |
| **DDX24** | 0.05 | 0.05 | -0.06 | 0.02 | 0.02 | -0.09 | 6.4E-01 | -0.03 |
| **ZMYND11** | 0.03 | 0.01 | -0.01 | -0.03 | -0.06 | 0.06 | 6.4E-01 | -0.02 |
| **BAIAP2** | -0.02 | -0.06 | 0.02 | -0.13 | 0.09 | 0.09 | 6.4E-01 | 0.04 |
| **NFKBIL1** | -0.13 | -0.21 | 0.22 | 0.01 | -0.06 | 0.13 | 6.4E-01 | 0.07 |
| **PEX14** | -0.06 | 0.04 | 0.10 | -0.21 | -0.14 | 0.22 | 6.4E-01 | -0.07 |
| **DCTN3** | 0.01 | 0.02 | 0.00 | 0.00 | 0.05 | -0.08 | 6.4E-01 | -0.02 |
| **SRP54** | -0.01 | 0.01 | -0.05 | -0.01 | 0.11 | -0.06 | 6.4E-01 | 0.03 |
| **IRF6** | 0.02 | -0.01 | -0.05 | 0.09 | -0.06 | 0.01 | 6.4E-01 | 0.02 |
| **NCBP3** | -0.02 | -0.03 | 0.08 | 0.00 | -0.04 | 0.01 | 6.4E-01 | -0.02 |
| **GNL2** | 0.06 | -0.03 | -0.01 | 0.03 | -0.04 | -0.02 | 6.4E-01 | -0.02 |
| **TPD52L2** | -0.03 | 0.03 | 0.02 | -0.01 | 0.03 | -0.04 | 6.4E-01 | -0.01 |
| **RAB11FIP5** | -0.13 | 0.05 | 0.03 | -0.01 | 0.05 | 0.00 | 6.4E-01 | 0.03 |
| **AAR2** | 0.15 | -0.06 | -0.02 | -0.04 | 0.14 | -0.20 | 6.4E-01 | -0.06 |
| **XRCC1** | 0.01 | 0.07 | -0.15 | 0.12 | 0.02 | -0.09 | 6.5E-01 | 0.04 |
| **CHD6** | -0.08 | 0.01 | 0.12 | -0.15 | 0.04 | 0.04 | 6.5E-01 | -0.04 |
| **PKMYT1** | 0.07 | 0.10 | -0.38 | -0.17 | 0.32 | -0.04 | 6.5E-01 | 0.11 |
| **DNAH17** | -0.02 | -0.04 | 0.00 | -0.13 | 0.05 | 0.14 | 6.5E-01 | 0.04 |
| **EIF2B1** | 0.04 | -0.04 | -0.07 | 0.12 | 0.09 | -0.15 | 6.5E-01 | 0.05 |
| **NDUFS5** | 0.07 | 0.06 | -0.23 | 0.07 | 0.10 | -0.09 | 6.5E-01 | 0.06 |
| **APOE** | -0.10 | -0.12 | 0.14 | -0.05 | 0.12 | -0.01 | 6.5E-01 | 0.05 |
| **C1QBP** | 0.02 | 0.00 | -0.04 | -0.05 | 0.06 | 0.02 | 6.5E-01 | 0.02 |
| **MFSD4B** | -0.11 | 0.07 | 0.12 | 0.20 | -0.18 | -0.13 | 6.5E-01 | -0.07 |
| **CHMP5** | -0.19 | -0.06 | 0.16 | 0.03 | -0.02 | 0.05 | 6.5E-01 | 0.05 |
| **DCP1A** | -0.20 | -0.03 | 0.12 | 0.17 | 0.02 | -0.11 | 6.5E-01 | 0.06 |
| **SCAF4** | -0.01 | 0.02 | 0.02 | -0.02 | -0.05 | 0.05 | 6.5E-01 | -0.02 |
| **RARS1** | -0.03 | -0.01 | 0.09 | 0.00 | 0.07 | -0.13 | 6.5E-01 | -0.04 |
| **SYTL2** | -0.14 | -0.04 | 0.11 | -0.04 | 0.01 | 0.07 | 6.5E-01 | 0.04 |
| **SPINDOC** | -0.52 | -0.38 | 0.40 | -0.52 | 0.14 | 0.50 | 6.5E-01 | 0.20 |
| **LANCL1** | -0.01 | -0.10 | 0.05 | 0.11 | -0.11 | 0.05 | 6.5E-01 | 0.04 |
| **TMEM205** | -0.09 | -0.04 | 0.07 | 0.10 | -0.10 | 0.05 | 6.5E-01 | 0.04 |
| **MPV17** | -0.17 | -0.01 | 0.11 | 0.08 | -0.07 | 0.05 | 6.5E-01 | 0.05 |
| **F2** | -0.01 | -0.18 | 0.07 | -0.18 | 0.19 | 0.07 | 6.5E-01 | 0.06 |
| **TBC1D20** | -0.12 | 0.07 | 0.10 | 0.02 | -0.07 | 0.00 | 6.5E-01 | -0.04 |
| **ACOT8** | 0.05 | 0.02 | -0.15 | 0.10 | 0.09 | -0.13 | 6.5E-01 | 0.05 |
| **COX5A** | 0.04 | 0.02 | -0.03 | 0.08 | -0.08 | -0.04 | 6.5E-01 | -0.02 |
| **KIF16B** | -0.02 | -0.05 | 0.05 | 0.00 | 0.02 | 0.00 | 6.5E-01 | 0.01 |
| **BRK1** | -0.12 | -0.02 | 0.09 | -0.01 | 0.02 | 0.03 | 6.5E-01 | 0.03 |
| **IDH1** | 0.04 | -0.02 | -0.06 | -0.01 | 0.09 | -0.04 | 6.6E-01 | 0.02 |
| **VPS72** | -0.02 | -0.06 | 0.03 | 0.05 | -0.10 | 0.09 | 6.6E-01 | 0.03 |
| **BAG2** | 0.01 | -0.06 | 0.01 | 0.00 | 0.10 | -0.06 | 6.6E-01 | 0.02 |
| **UTP11** | 0.04 | 0.02 | -0.04 | -0.03 | 0.02 | -0.02 | 6.6E-01 | -0.01 |

| | | | | | | | | |
|---|---|---|---|---|---|---|---|---|
| ZNF524 | -0.02 | 0.03 | 0.05 | -0.22 | -0.02 | 0.15 | 6.6E-01 | -0.05 |
| CTDSPL2 | 0.04 | 0.03 | -0.04 | -0.06 | 0.01 | 0.02 | 6.6E-01 | -0.02 |
| ING1 | -0.02 | -0.02 | 0.12 | 0.19 | -0.31 | -0.01 | 6.6E-01 | -0.07 |
| PON2 | -0.05 | -0.04 | 0.15 | 0.01 | -0.10 | 0.03 | 6.6E-01 | -0.04 |
| MVB12A | -0.06 | 0.07 | -0.05 | -0.03 | 0.03 | 0.03 | 6.6E-01 | 0.02 |
| RBX1 | -0.10 | -0.04 | 0.08 | 0.00 | -0.04 | 0.08 | 6.6E-01 | 0.03 |
| SH3BGRL3 | -0.16 | -0.21 | 0.47 | 0.04 | -0.10 | -0.16 | 6.6E-01 | -0.11 |
| AP3B1 | 0.10 | -0.06 | -0.01 | -0.03 | 0.03 | -0.05 | 6.6E-01 | -0.03 |
| CNNM2 | -0.02 | 0.08 | 0.04 | 0.26 | -0.09 | -0.33 | 6.6E-01 | -0.08 |
| MCU | 0.05 | 0.08 | -0.10 | -0.01 | 0.02 | -0.06 | 6.6E-01 | -0.03 |
| GPATCH2 | -0.01 | 0.03 | 0.02 | -0.02 | 0.06 | -0.08 | 6.6E-01 | -0.02 |
| PLEC | -0.09 | 0.06 | 0.07 | 0.04 | 0.03 | -0.12 | 6.6E-01 | -0.03 |
| CFAP36 | -0.29 | 0.10 | -0.22 | -0.66 | 0.29 | 0.49 | 6.6E-01 | 0.18 |
| DNAJC17 | 0.01 | 0.01 | 0.00 | -0.09 | 0.01 | 0.05 | 6.6E-01 | -0.02 |
| ZC3H7B | -0.11 | 0.02 | 0.15 | 0.06 | -0.01 | -0.12 | 6.6E-01 | -0.04 |
| VPS36 | 0.00 | 0.03 | 0.01 | -0.01 | 0.07 | -0.09 | 6.6E-01 | -0.02 |
| ATXN10 | -0.04 | -0.03 | 0.05 | 0.04 | -0.02 | 0.00 | 6.6E-01 | 0.02 |
| SNX5 | -0.03 | 0.08 | -0.10 | 0.03 | 0.07 | -0.06 | 6.6E-01 | 0.03 |
| BLZF1 | -0.03 | -0.03 | 0.10 | -0.05 | -0.04 | 0.05 | 6.6E-01 | -0.03 |
| RLIM | -0.25 | -0.03 | 0.39 | -0.33 | 0.40 | -0.42 | 6.6E-01 | -0.15 |
| SIPA1L1 | -0.09 | 0.01 | 0.05 | -0.05 | 0.04 | 0.05 | 6.6E-01 | 0.02 |
| ERLIN1 | 0.05 | 0.01 | -0.03 | 0.02 | 0.03 | -0.08 | 6.6E-01 | -0.02 |
| SAP130 | -0.01 | -0.04 | 0.07 | 0.04 | -0.02 | -0.05 | 6.6E-01 | -0.02 |
| DPAGT1 | 0.07 | -0.06 | -0.06 | 0.10 | -0.09 | 0.03 | 6.6E-01 | 0.03 |
| HNRNPA1L3 | -0.07 | -0.05 | -0.02 | -0.28 | 0.11 | 0.25 | 6.6E-01 | 0.08 |
| TUSC1 | -0.10 | 0.06 | 0.08 | -0.07 | 0.04 | -0.02 | 6.7E-01 | -0.03 |
| UBE2M | 0.08 | -0.01 | -0.02 | -0.13 | 0.09 | -0.02 | 6.7E-01 | -0.03 |
| MED31 | -0.01 | -0.05 | 0.09 | -0.03 | 0.04 | -0.05 | 6.7E-01 | -0.02 |
| ZBTB45 | -0.15 | -0.01 | 0.26 | -0.38 | 0.15 | 0.05 | 6.7E-01 | -0.09 |
| SPTAN1 | -0.09 | -0.02 | 0.06 | 0.10 | -0.02 | -0.04 | 6.7E-01 | 0.03 |
| USP46 | -0.03 | -0.03 | 0.14 | -0.03 | 0.19 | -0.28 | 6.7E-01 | -0.07 |
| KPNB1 | 0.04 | 0.00 | -0.03 | -0.01 | 0.00 | -0.01 | 6.7E-01 | -0.01 |
| ARRB1 | -0.04 | 0.21 | -0.10 | 0.10 | -0.18 | -0.03 | 6.7E-01 | -0.06 |
| WDR33 | -0.01 | -0.01 | -0.03 | 0.15 | -0.06 | -0.05 | 6.7E-01 | 0.03 |
| ARID4A | -0.04 | -0.07 | 0.05 | -0.07 | -0.03 | 0.15 | 6.7E-01 | 0.03 |
| PRPF8 | 0.03 | -0.04 | 0.02 | 0.00 | -0.02 | 0.01 | 6.7E-01 | -0.01 |
| LAMTOR3 | 0.03 | 0.08 | -0.06 | 0.10 | -0.02 | -0.15 | 6.7E-01 | -0.04 |
| ZMYM2 | 0.02 | -0.03 | -0.03 | -0.08 | 0.01 | 0.10 | 6.7E-01 | 0.03 |
| PPA2 | 0.10 | 0.01 | -0.18 | 0.02 | 0.08 | -0.05 | 6.7E-01 | 0.04 |
| UAP1 | -0.04 | -0.09 | 0.09 | 0.02 | 0.00 | 0.02 | 6.7E-01 | 0.02 |
| HEATR5A | 0.05 | -0.05 | 0.05 | -0.17 | 0.13 | -0.02 | 6.7E-01 | -0.04 |
| FAR2 | -0.01 | -0.06 | 0.03 | 0.03 | 0.07 | -0.07 | 6.7E-01 | 0.02 |
| FHDC1 | -0.09 | -0.28 | 0.47 | -0.21 | 0.02 | -0.03 | 6.7E-01 | -0.11 |
| TSPAN15 | 0.01 | -0.13 | 0.20 | -0.38 | 0.05 | 0.17 | 6.7E-01 | -0.09 |
| PRPF40B | 0.03 | 0.00 | 0.06 | 0.23 | 0.01 | -0.40 | 6.7E-01 | -0.08 |
| NFU1 | 0.06 | 0.10 | -0.27 | 0.15 | -0.08 | 0.00 | 6.7E-01 | 0.06 |
| RXRA | -0.03 | -0.10 | 0.22 | 0.23 | -0.46 | 0.03 | 6.8E-01 | -0.10 |
| UFL1 | -0.04 | -0.03 | 0.04 | 0.01 | -0.04 | 0.05 | 6.8E-01 | 0.02 |

| | | | | | | | | |
|---|---|---|---|---|---|---|---|---|
| MAD2L1BP | -0.05 | 0.03 | 0.07 | -0.15 | 0.05 | 0.04 | 6.8E-01 | -0.03 |
| SCYL1 | -0.02 | -0.03 | 0.02 | 0.03 | -0.06 | 0.05 | 6.8E-01 | 0.02 |
| RFX5 | -0.10 | 0.00 | 0.06 | -0.01 | 0.00 | 0.04 | 6.8E-01 | 0.02 |
| MGAT2 | -0.11 | 0.05 | 0.11 | 0.10 | -0.08 | -0.10 | 6.8E-01 | -0.04 |
| NOL7 | 0.04 | 0.03 | -0.04 | 0.01 | 0.02 | -0.06 | 6.8E-01 | -0.02 |
| RPL28 | 0.02 | -0.02 | 0.03 | 0.06 | 0.01 | -0.11 | 6.8E-01 | -0.02 |
| COPS8 | -0.04 | 0.05 | 0.04 | -0.11 | -0.05 | 0.11 | 6.8E-01 | -0.03 |
| MCAT | 0.07 | -0.03 | -0.01 | 0.05 | 0.00 | -0.10 | 6.8E-01 | -0.02 |
| DDX55 | 0.02 | 0.00 | 0.00 | 0.05 | -0.01 | -0.06 | 6.8E-01 | -0.01 |
| CHD7 | -0.10 | 0.10 | 0.06 | -0.17 | -0.11 | 0.18 | 6.8E-01 | -0.06 |
| PHF12 | 0.05 | 0.02 | -0.04 | -0.05 | 0.04 | -0.02 | 6.8E-01 | -0.02 |
| TSPYL1 | -0.07 | 0.03 | 0.06 | 0.06 | -0.05 | -0.05 | 6.9E-01 | -0.02 |
| SLC12A8 | -0.30 | -0.02 | 0.40 | 0.11 | -0.18 | -0.13 | 6.9E-01 | -0.10 |
| ESCO1 | 0.04 | -0.04 | -0.04 | -0.07 | 0.04 | 0.06 | 6.9E-01 | 0.02 |
| HMOX1 | -0.17 | -0.06 | 0.16 | 0.01 | 0.05 | -0.01 | 6.9E-01 | 0.04 |
| POLR1F | -0.08 | -0.09 | 0.09 | 0.13 | 0.07 | -0.14 | 6.9E-01 | 0.04 |
| ERBB2 | -0.18 | 0.20 | -0.11 | -0.05 | 0.03 | 0.08 | 6.9E-01 | 0.05 |
| PLCD1 | -0.20 | 0.12 | -0.01 | 0.04 | -0.10 | 0.13 | 6.9E-01 | 0.05 |
| PCID2 | 0.01 | -0.01 | 0.00 | 0.00 | 0.01 | -0.01 | 6.9E-01 | 0.00 |
| PLEKHJ1 | -0.05 | 0.10 | 0.03 | -0.33 | -0.13 | 0.30 | 6.9E-01 | -0.08 |
| MMTAG2 | 0.11 | -0.06 | 0.01 | 0.14 | -0.09 | -0.13 | 6.9E-01 | -0.04 |
| TRIM59 | 0.01 | -0.03 | 0.04 | -0.06 | 0.00 | 0.03 | 6.9E-01 | -0.01 |
| CNOT8 | 0.07 | -0.26 | 0.28 | 0.26 | -0.10 | -0.38 | 6.9E-01 | -0.10 |
| INTS2 | 0.02 | -0.03 | -0.02 | -0.08 | 0.02 | 0.09 | 6.9E-01 | 0.02 |
| ATP2C2 | -0.08 | 0.00 | 0.04 | 0.01 | 0.08 | -0.06 | 6.9E-01 | 0.02 |
| BLM | 0.04 | -0.17 | 0.04 | -0.08 | -0.05 | 0.20 | 6.9E-01 | 0.05 |
| NEIL1 | -0.09 | 0.05 | -0.02 | 0.06 | 0.10 | -0.12 | 6.9E-01 | 0.03 |
| PARVA | 0.06 | 0.03 | -0.18 | -0.14 | 0.05 | 0.15 | 6.9E-01 | 0.05 |
| PSMB7 | -0.06 | 0.19 | -0.28 | 0.19 | -0.18 | 0.07 | 6.9E-01 | 0.07 |
| CKMT1A | -0.03 | 0.02 | -0.05 | 0.14 | -0.07 | -0.04 | 6.9E-01 | 0.03 |
| EVPL | 0.01 | 0.00 | 0.02 | 0.07 | 0.01 | -0.12 | 6.9E-01 | -0.02 |
| BLTP3A | -0.17 | -0.06 | 0.08 | 0.06 | -0.27 | 0.29 | 6.9E-01 | 0.08 |
| MPZL3 | -0.27 | 0.01 | 0.16 | 0.01 | 0.00 | 0.05 | 6.9E-01 | 0.05 |
| DNAJB6 | 0.04 | 0.00 | -0.01 | 0.09 | -0.07 | -0.05 | 6.9E-01 | -0.02 |
| EPRS1 | -0.03 | 0.00 | -0.02 | -0.03 | 0.14 | -0.07 | 6.9E-01 | 0.03 |
| WDR7 | -0.24 | 0.00 | 0.14 | 0.08 | -0.10 | 0.08 | 6.9E-01 | 0.05 |
| CDCA2 | 0.00 | 0.05 | -0.03 | -0.08 | 0.00 | 0.05 | 6.9E-01 | -0.02 |
| SMC6 | -0.01 | -0.02 | 0.04 | -0.03 | -0.01 | 0.02 | 6.9E-01 | -0.01 |
| DNAJB11 | 0.04 | 0.00 | -0.02 | 0.02 | 0.04 | -0.10 | 6.9E-01 | -0.02 |
| TMLHE | 0.13 | -0.08 | -0.01 | 0.02 | 0.03 | -0.10 | 7.0E-01 | -0.03 |
| SARAF | 0.14 | -0.02 | -0.23 | 0.17 | 0.08 | -0.17 | 7.0E-01 | 0.06 |
| PARP12 | -0.03 | 0.02 | 0.04 | -0.04 | 0.05 | -0.04 | 7.0E-01 | -0.02 |
| MAPK12 | 0.05 | 0.18 | -0.39 | 0.02 | 0.15 | -0.10 | 7.0E-01 | 0.08 |
| HNRNPLL | 0.04 | -0.12 | -0.01 | 0.23 | -0.05 | -0.12 | 7.0E-01 | 0.05 |
| H2AX | 0.02 | -0.05 | -0.02 | -0.14 | 0.05 | 0.13 | 7.0E-01 | 0.03 |
| MCEE | 0.08 | 0.01 | -0.05 | -0.14 | 0.01 | 0.07 | 7.0E-01 | -0.03 |
| XPO4 | 0.02 | -0.06 | 0.10 | 0.10 | 0.04 | -0.21 | 7.0E-01 | -0.04 |
| PTPN12 | -0.01 | -0.04 | 0.11 | -0.16 | 0.18 | -0.10 | 7.0E-01 | -0.05 |

| | | | | | | | | |
|---|---|---|---|---|---|---|---|---|
| PDZD2 | -0.10 | -0.06 | 0.21 | -0.09 | -0.03 | 0.04 | 7.0E-01 | -0.04 |
| GLYR1 | 0.06 | 0.03 | -0.13 | 0.03 | 0.03 | -0.03 | 7.0E-01 | 0.03 |
| CNOT10 | 0.09 | -0.22 | 0.21 | -0.21 | -0.05 | 0.13 | 7.0E-01 | -0.07 |
| RRP1 | 0.01 | 0.00 | 0.01 | -0.01 | 0.03 | -0.02 | 7.0E-01 | -0.01 |
| NHS | -0.08 | 0.02 | 0.10 | -0.14 | 0.07 | 0.01 | 7.0E-01 | -0.03 |
| RSL1D1 | 0.04 | 0.02 | -0.05 | 0.01 | 0.02 | -0.05 | 7.0E-01 | -0.01 |
| SCOC | -0.03 | -0.13 | 0.22 | -0.13 | -0.12 | 0.15 | 7.0E-01 | -0.06 |
| ZNF785 | -0.08 | -0.01 | 0.18 | 0.25 | 0.12 | -0.62 | 7.0E-01 | -0.12 |
| LTA4H | -0.06 | 0.04 | -0.02 | -0.01 | -0.07 | 0.11 | 7.0E-01 | 0.03 |
| PRKAR1A | -0.06 | 0.00 | 0.09 | -0.04 | -0.02 | 0.02 | 7.0E-01 | -0.02 |
| CEP57 | -0.03 | -0.04 | 0.05 | 0.00 | -0.03 | 0.05 | 7.0E-01 | 0.01 |
| ZNF142 | -0.07 | -0.05 | 0.18 | -0.13 | -0.02 | 0.08 | 7.0E-01 | -0.04 |
| PRKCZ | -0.17 | 0.10 | 0.01 | 0.07 | -0.06 | 0.03 | 7.0E-01 | 0.04 |
| INTS12 | 0.08 | 0.08 | -0.13 | -0.03 | 0.02 | -0.04 | 7.0E-01 | -0.03 |
| CHCHD3 | -0.01 | -0.01 | 0.05 | 0.06 | -0.05 | -0.04 | 7.0E-01 | -0.02 |
| ANO2 | -0.33 | 0.07 | 0.13 | 0.13 | -0.08 | 0.02 | 7.0E-01 | 0.06 |
| DCUN1D3 | -0.11 | 0.05 | 0.10 | -0.02 | -0.03 | 0.00 | 7.1E-01 | -0.03 |
| TRIM41 | 0.11 | 0.16 | -0.19 | 0.17 | -0.01 | -0.31 | 7.1E-01 | -0.07 |
| HOXA10 | -0.03 | 0.17 | -0.32 | -0.22 | -0.05 | 0.35 | 7.1E-01 | 0.09 |
| CHERP | 0.04 | -0.01 | -0.02 | -0.01 | 0.01 | -0.02 | 7.1E-01 | -0.01 |
| GNA12 | -0.09 | 0.02 | 0.02 | -0.07 | 0.08 | 0.02 | 7.1E-01 | 0.02 |
| EPHX2 | -0.02 | 0.03 | 0.02 | 0.07 | -0.11 | 0.00 | 7.1E-01 | -0.02 |
| REX1BD | -0.25 | 0.11 | -0.12 | 0.22 | -0.50 | 0.37 | 7.1E-01 | 0.12 |
| MLKL | -0.48 | 0.14 | 0.40 | 0.49 | -0.28 | -0.67 | 7.1E-01 | -0.17 |
| CERS2 | -0.17 | -0.04 | 0.13 | -0.06 | -0.06 | 0.17 | 7.1E-01 | 0.05 |
| UTY | 0.01 | 0.04 | 0.00 | -0.20 | 0.10 | 0.04 | 7.1E-01 | -0.04 |
| CSNK1G3 | -0.21 | -0.02 | 0.13 | -0.02 | -0.10 | 0.18 | 7.1E-01 | 0.05 |
| ARHGAP32 | 0.02 | -0.14 | 0.01 | -0.25 | 0.12 | 0.19 | 7.1E-01 | 0.06 |
| RAB25 | -0.19 | -0.05 | 0.14 | -0.15 | 0.06 | 0.15 | 7.1E-01 | 0.05 |
| ZFP62 | -0.02 | 0.09 | -0.11 | -0.01 | 0.00 | 0.04 | 7.1E-01 | 0.02 |
| CDC27 | 0.02 | -0.05 | -0.03 | -0.14 | 0.06 | 0.12 | 7.1E-01 | 0.03 |
| PHC2 | -0.01 | 0.03 | -0.01 | -0.06 | 0.02 | 0.02 | 7.1E-01 | -0.01 |
| POLR2L | 0.09 | -0.08 | -0.04 | -0.02 | 0.00 | 0.05 | 7.1E-01 | 0.02 |
| TIMELESS | 0.03 | -0.01 | 0.00 | -0.03 | -0.05 | 0.06 | 7.1E-01 | -0.01 |
| COPB2 | 0.00 | -0.01 | -0.02 | 0.06 | 0.03 | -0.07 | 7.1E-01 | 0.02 |
| BMI1 | -0.03 | -0.07 | 0.05 | 0.12 | -0.11 | 0.03 | 7.1E-01 | 0.03 |
| RDH14 | 0.10 | 0.08 | -0.13 | 0.13 | -0.11 | -0.10 | 7.1E-01 | -0.04 |
| SSRP1 | 0.04 | -0.02 | 0.00 | -0.07 | 0.02 | 0.02 | 7.1E-01 | -0.01 |
| LGALS1 | -0.06 | 0.10 | 0.04 | -0.26 | 0.37 | -0.29 | 7.1E-01 | -0.09 |
| SMIM7 | 0.06 | -0.09 | 0.06 | -0.06 | -0.08 | 0.10 | 7.1E-01 | -0.03 |
| PFDN2 | 0.05 | 0.02 | -0.05 | 0.01 | 0.00 | -0.03 | 7.1E-01 | -0.01 |
| TNNI3K | 0.07 | -0.09 | 0.08 | 0.13 | 0.08 | -0.33 | 7.1E-01 | -0.06 |
| ARF6 | -0.07 | 0.04 | 0.00 | 0.03 | 0.06 | -0.07 | 7.1E-01 | 0.02 |
| CCDC9B | 0.13 | 0.07 | -0.15 | 0.02 | 0.08 | -0.19 | 7.1E-01 | -0.05 |
| PLEKHH3 | -0.13 | 0.00 | 0.05 | -0.05 | -0.10 | 0.20 | 7.1E-01 | 0.04 |
| MED23 | 0.09 | -0.01 | -0.05 | 0.03 | -0.01 | -0.04 | 7.2E-01 | -0.02 |
| RAP2C | -0.15 | -0.07 | 0.15 | -0.06 | -0.01 | 0.12 | 7.2E-01 | 0.04 |
| VPS37C | -0.06 | 0.00 | 0.04 | -0.02 | 0.04 | 0.00 | 7.2E-01 | 0.01 |

| | | | | | | | | |
|---|---|---|---|---|---|---|---|---|
| CCDC112 | 0.07 | -0.11 | 0.08 | -0.04 | 0.01 | -0.01 | 7.2E-01 | -0.03 |
| BCAR3 | -0.07 | -0.01 | 0.01 | -0.16 | 0.07 | 0.14 | 7.2E-01 | 0.04 |
| VTI1A | -0.20 | -0.15 | 0.22 | 0.17 | -0.17 | 0.06 | 7.2E-01 | 0.06 |
| NOL10 | 0.08 | 0.03 | -0.08 | 0.08 | -0.05 | -0.08 | 7.2E-01 | -0.03 |
| STK3 | -0.12 | -0.05 | 0.08 | -0.11 | 0.23 | -0.06 | 7.2E-01 | 0.05 |
| MED8 | -0.08 | -0.06 | 0.00 | 0.36 | -0.18 | -0.11 | 7.2E-01 | 0.07 |
| CNOT9 | -0.01 | 0.11 | -0.06 | -0.13 | 0.05 | 0.03 | 7.2E-01 | -0.03 |
| TCEANC2 | -0.18 | -0.06 | 0.13 | 0.17 | -0.20 | 0.10 | 7.2E-01 | 0.06 |
| TFPT | -0.02 | 0.17 | -0.10 | -0.13 | 0.09 | -0.03 | 7.2E-01 | -0.04 |
| SUN2 | -0.01 | 0.00 | 0.02 | -0.03 | 0.01 | 0.01 | 7.2E-01 | -0.01 |
| PPP2CA | 0.01 | 0.07 | -0.05 | 0.05 | -0.01 | -0.08 | 7.2E-01 | -0.02 |
| TAF3 | -0.01 | -0.03 | 0.01 | 0.00 | -0.06 | 0.09 | 7.2E-01 | 0.02 |
| GOLIM4 | -0.06 | 0.00 | 0.08 | 0.06 | -0.07 | -0.03 | 7.2E-01 | -0.02 |
| MYCBP | -0.01 | 0.01 | 0.02 | 0.07 | -0.07 | -0.03 | 7.2E-01 | -0.02 |
| MELK | -0.14 | 0.08 | 0.11 | -0.08 | 0.10 | -0.08 | 7.2E-01 | -0.04 |
| MSL3 | 0.00 | 0.01 | -0.07 | -0.13 | 0.00 | 0.17 | 7.2E-01 | 0.03 |
| SMCO4 | -0.08 | -0.20 | 0.16 | -0.12 | -0.06 | 0.24 | 7.2E-01 | 0.06 |
| PIGT | 0.01 | -0.03 | 0.05 | 0.11 | -0.08 | -0.08 | 7.2E-01 | -0.03 |
| TAF7 | 0.04 | 0.00 | -0.08 | -0.10 | 0.09 | 0.04 | 7.2E-01 | 0.02 |
| SARS2 | 0.08 | -0.04 | -0.08 | 0.05 | 0.03 | -0.05 | 7.2E-01 | 0.02 |
| HSPD1 | 0.10 | 0.07 | -0.13 | -0.06 | -0.02 | 0.03 | 7.2E-01 | -0.03 |
| PRPF38A | 0.11 | 0.01 | -0.08 | 0.11 | -0.03 | -0.15 | 7.2E-01 | -0.04 |
| KAT14 | 0.14 | -0.06 | -0.16 | -0.02 | 0.13 | -0.07 | 7.2E-01 | 0.04 |
| TONSL | 0.03 | 0.01 | -0.07 | 0.00 | 0.09 | -0.06 | 7.3E-01 | 0.02 |
| RAI1 | 0.00 | 0.03 | -0.06 | -0.05 | -0.01 | 0.09 | 7.3E-01 | 0.02 |
| VPS41 | -0.03 | -0.14 | 0.10 | -0.03 | 0.18 | -0.10 | 7.3E-01 | 0.04 |
| ZFR | 0.05 | 0.00 | -0.03 | 0.05 | -0.04 | -0.04 | 7.3E-01 | -0.01 |
| SLC35B2 | -0.01 | -0.01 | 0.05 | 0.08 | -0.03 | -0.08 | 7.3E-01 | -0.02 |
| PDIA3 | 0.05 | 0.01 | -0.10 | 0.06 | 0.04 | -0.07 | 7.3E-01 | 0.02 |
| NDUFA2 | 0.01 | -0.02 | -0.01 | 0.08 | -0.03 | -0.03 | 7.3E-01 | 0.01 |
| YWHAB | 0.03 | 0.03 | -0.10 | 0.08 | -0.02 | -0.03 | 7.3E-01 | 0.02 |
| ADCY3 | -0.25 | 0.15 | 0.16 | -0.05 | -0.02 | -0.02 | 7.3E-01 | -0.05 |
| FKRP | 0.06 | -0.05 | 0.03 | 0.19 | -0.11 | -0.15 | 7.3E-01 | -0.04 |
| CARHSP1 | 0.05 | 0.02 | -0.16 | -0.18 | 0.03 | 0.21 | 7.3E-01 | 0.05 |
| AP1S1 | -0.11 | -0.08 | 0.12 | 0.17 | -0.07 | -0.05 | 7.3E-01 | 0.04 |
| CD47 | -0.19 | -0.02 | 0.05 | -0.23 | -0.10 | 0.39 | 7.3E-01 | 0.07 |
| PLXNA3 | -0.07 | -0.06 | 0.09 | 0.02 | 0.06 | -0.05 | 7.3E-01 | 0.02 |
| PDCD10 | -0.04 | 0.09 | -0.19 | -0.04 | -0.22 | 0.33 | 7.3E-01 | 0.07 |
| ABCC5 | -0.09 | 0.00 | 0.12 | -0.06 | 0.00 | 0.02 | 7.3E-01 | -0.02 |
| PIGH | 0.00 | -0.05 | 0.09 | -0.11 | 0.03 | 0.05 | 7.3E-01 | -0.02 |
| LBR | -0.01 | -0.01 | 0.04 | 0.04 | -0.03 | -0.03 | 7.3E-01 | -0.01 |
| LARP1 | -0.02 | -0.01 | 0.04 | 0.02 | 0.04 | -0.08 | 7.3E-01 | -0.02 |
| URB1 | 0.03 | -0.02 | -0.02 | -0.02 | -0.01 | 0.05 | 7.3E-01 | 0.01 |
| WDFY3 | -0.01 | 0.03 | 0.00 | -0.06 | 0.05 | -0.01 | 7.3E-01 | -0.01 |
| ZMAT5 | 0.03 | -0.01 | -0.05 | 0.06 | -0.09 | 0.06 | 7.3E-01 | 0.02 |
| CTSA | 0.19 | -0.17 | -0.17 | 0.35 | -0.14 | -0.14 | 7.3E-01 | 0.07 |
| SP2 | -0.04 | -0.08 | 0.16 | 0.00 | -0.10 | 0.05 | 7.3E-01 | -0.03 |
| ARL8A | 0.02 | 0.03 | -0.15 | -0.18 | 0.27 | -0.03 | 7.3E-01 | 0.05 |

| | | | | | | | | |
|---|---|---|---|---|---|---|---|---|
| SAP30L | 0.10 | 0.08 | -0.14 | -0.12 | 0.02 | 0.04 | 7.3E-01 | -0.03 |
| COLGALT1 | 0.00 | -0.08 | 0.05 | -0.05 | 0.05 | 0.02 | 7.3E-01 | 0.02 |
| PTMS | -0.12 | -0.02 | 0.18 | 0.06 | -0.18 | 0.04 | 7.3E-01 | -0.04 |
| SEC22A | 0.04 | 0.00 | -0.01 | 0.13 | -0.02 | -0.17 | 7.3E-01 | -0.03 |
| GOLPH3 | 0.02 | 0.06 | -0.03 | 0.11 | 0.04 | -0.22 | 7.3E-01 | -0.04 |
| NMT1 | 0.07 | 0.00 | -0.03 | 0.10 | -0.01 | -0.14 | 7.3E-01 | -0.03 |
| NDUFV1 | 0.03 | 0.00 | -0.06 | 0.07 | -0.01 | -0.04 | 7.4E-01 | 0.02 |
| CWC27 | 0.15 | -0.06 | -0.15 | -0.09 | 0.11 | 0.02 | 7.4E-01 | 0.04 |
| ZFYVE19 | -0.01 | 0.03 | 0.00 | -0.13 | 0.08 | 0.02 | 7.4E-01 | -0.02 |
| DIS3 | -0.05 | -0.02 | 0.10 | 0.07 | 0.01 | -0.13 | 7.4E-01 | -0.03 |
| PRDM2 | -0.04 | -0.06 | 0.06 | -0.03 | -0.02 | 0.08 | 7.4E-01 | 0.02 |
| MTHFD1 | 0.03 | 0.03 | -0.08 | 0.00 | 0.04 | -0.03 | 7.4E-01 | 0.01 |
| PALMD | 0.06 | -0.19 | -0.02 | 0.17 | -0.34 | 0.23 | 7.4E-01 | 0.07 |
| XYLT2 | -0.16 | -0.01 | 0.12 | 0.03 | 0.01 | 0.00 | 7.4E-01 | 0.03 |
| PRODH | 0.06 | 0.07 | -0.18 | 0.00 | 0.05 | -0.01 | 7.4E-01 | 0.03 |
| PPT1 | 0.01 | -0.06 | -0.01 | 0.17 | -0.16 | 0.03 | 7.4E-01 | 0.04 |
| VAV3 | -0.23 | 0.04 | 0.09 | -0.08 | 0.26 | -0.12 | 7.4E-01 | 0.06 |
| GALNT14 | 0.01 | 0.04 | -0.02 | 0.06 | 0.04 | -0.14 | 7.4E-01 | -0.02 |
| STX17 | -0.07 | -0.01 | 0.12 | 0.05 | 0.01 | -0.10 | 7.4E-01 | -0.03 |
| UBAP2 | -0.27 | -0.17 | 0.30 | 0.11 | 0.10 | -0.14 | 7.4E-01 | 0.07 |
| MON1B | -0.13 | 0.03 | -0.01 | -0.29 | 0.13 | 0.22 | 7.4E-01 | 0.06 |
| EDF1 | 0.00 | 0.00 | -0.07 | 0.21 | -0.06 | -0.11 | 7.4E-01 | 0.04 |
| ILKAP | -0.09 | -0.03 | 0.07 | -0.02 | -0.08 | 0.13 | 7.4E-01 | 0.03 |
| C1orf198 | -0.09 | 0.07 | -0.03 | -0.09 | 0.14 | -0.01 | 7.4E-01 | 0.03 |
| DDX41 | 0.06 | -0.02 | -0.02 | 0.06 | 0.01 | -0.09 | 7.4E-01 | -0.02 |
| COQ8A | 0.06 | -0.14 | -0.01 | 0.25 | -0.02 | -0.18 | 7.4E-01 | 0.05 |
| CENATAC | -0.12 | 0.02 | 0.15 | -0.20 | -0.09 | 0.20 | 7.5E-01 | -0.05 |
| PHF8 | -0.08 | 0.10 | -0.05 | 0.04 | -0.04 | 0.03 | 7.5E-01 | 0.02 |
| TLE1 | 0.07 | -0.15 | 0.04 | -0.04 | 0.02 | 0.06 | 7.5E-01 | 0.03 |
| ZC3H11B | -0.13 | -0.11 | 0.30 | 0.15 | -0.22 | -0.06 | 7.5E-01 | -0.06 |
| ANXA6 | -0.04 | -0.13 | 0.12 | 0.06 | -0.08 | 0.05 | 7.5E-01 | 0.03 |
| PCMTD1 | -0.20 | -0.02 | 0.14 | -0.12 | 0.13 | 0.04 | 7.5E-01 | 0.04 |
| IRX5 | -0.46 | 0.08 | 0.07 | -0.57 | 0.24 | 0.39 | 7.5E-01 | 0.12 |
| SDHAF2 | 0.09 | 0.06 | -0.29 | 0.01 | -0.23 | 0.27 | 7.5E-01 | 0.07 |
| PDE12 | 0.01 | -0.18 | 0.10 | 0.06 | -0.14 | 0.12 | 7.5E-01 | 0.04 |
| ATP6V0A2 | -0.01 | -0.02 | 0.00 | 0.12 | -0.05 | -0.05 | 7.5E-01 | 0.02 |
| SUPT5H | 0.06 | -0.04 | 0.00 | -0.06 | -0.07 | 0.10 | 7.5E-01 | -0.02 |
| BTBD2 | 0.03 | -0.02 | -0.06 | -0.11 | 0.17 | -0.03 | 7.5E-01 | 0.03 |
| SF3B5 | -0.04 | -0.10 | 0.09 | -0.10 | 0.00 | 0.14 | 7.5E-01 | 0.03 |
| SS18L1 | 0.04 | 0.06 | -0.17 | -0.10 | 0.17 | -0.03 | 7.5E-01 | 0.04 |
| LUZP1 | -0.12 | -0.04 | 0.10 | 0.11 | -0.01 | -0.06 | 7.5E-01 | 0.03 |
| TMCC1 | -0.02 | -0.01 | 0.06 | -0.09 | 0.09 | -0.03 | 7.5E-01 | -0.02 |
| KRT5 | -0.01 | 0.09 | -0.41 | 0.19 | -0.65 | 0.50 | 7.5E-01 | 0.13 |
| UBAC2 | -0.13 | -0.06 | 0.11 | -0.16 | 0.08 | 0.13 | 7.5E-01 | 0.04 |
| HEXA | 0.05 | -0.05 | -0.03 | -0.09 | 0.14 | -0.01 | 7.5E-01 | 0.02 |
| FBXL6 | 0.04 | -0.05 | 0.03 | -0.06 | 0.00 | 0.04 | 7.5E-01 | -0.01 |
| LLPH | 0.00 | -0.05 | 0.08 | 0.07 | -0.08 | -0.02 | 7.5E-01 | -0.02 |
| TFB2M | 0.08 | 0.01 | -0.06 | -0.03 | 0.01 | 0.01 | 7.5E-01 | -0.01 |

| | | | | | | | | |
|---|---|---|---|---|---|---|---|---|
| **MYO9B** | 0.02 | 0.00 | -0.01 | 0.07 | -0.03 | -0.06 | 7.5E-01 | -0.01 |
| **OCRL** | -0.04 | -0.14 | 0.13 | -0.06 | 0.02 | 0.08 | 7.5E-01 | 0.03 |
| **OXCT1** | 0.11 | 0.07 | -0.25 | -0.03 | 0.06 | 0.01 | 7.6E-01 | 0.04 |
| **CTNND2** | -0.02 | -0.03 | 0.09 | -0.15 | 0.00 | 0.11 | 7.6E-01 | -0.03 |
| **TLCD4** | -0.08 | 0.00 | 0.05 | -0.03 | 0.01 | 0.04 | 7.6E-01 | 0.01 |
| **SNRNP48** | 0.05 | 0.11 | -0.12 | -0.05 | -0.01 | 0.02 | 7.6E-01 | -0.02 |
| **CMC1** | -0.03 | -0.01 | 0.09 | -0.43 | 0.11 | 0.18 | 7.6E-01 | -0.07 |
| **MYO6** | -0.02 | 0.01 | 0.00 | 0.03 | -0.02 | 0.00 | 7.6E-01 | 0.01 |
| **CYBRD1** | -0.04 | -0.11 | 0.20 | -0.13 | -0.02 | 0.08 | 7.6E-01 | -0.04 |
| **MAK16** | 0.14 | -0.01 | -0.20 | 0.10 | 0.04 | -0.09 | 7.6E-01 | 0.04 |
| **ESF1** | 0.05 | 0.00 | -0.07 | 0.01 | -0.01 | 0.02 | 7.6E-01 | 0.01 |
| **SEC16A** | 0.01 | 0.08 | -0.05 | 0.09 | 0.02 | -0.16 | 7.6E-01 | -0.03 |
| **ASAP2** | -0.02 | -0.12 | 0.00 | -0.37 | 0.19 | 0.23 | 7.6E-01 | 0.06 |
| **N4BP2L2** | 0.03 | 0.07 | -0.07 | -0.04 | -0.08 | 0.08 | 7.6E-01 | -0.02 |
| **YIPF6** | -0.12 | -0.08 | 0.12 | 0.03 | -0.17 | 0.18 | 7.6E-01 | 0.04 |
| **RPSA2** | 0.01 | 0.03 | 0.00 | 0.16 | -0.02 | -0.21 | 7.6E-01 | -0.04 |
| **VAMP3** | -0.11 | 0.06 | 0.01 | -0.04 | -0.06 | 0.13 | 7.6E-01 | 0.03 |
| **OFD1** | -0.03 | -0.02 | 0.04 | 0.01 | 0.01 | -0.01 | 7.6E-01 | 0.01 |
| **REST** | 0.20 | -0.09 | -0.06 | 0.10 | -0.07 | -0.10 | 7.6E-01 | -0.04 |
| **NDUFS8** | 0.09 | 0.05 | -0.11 | 0.14 | -0.04 | -0.16 | 7.6E-01 | -0.03 |
| **AHSA1** | 0.00 | -0.04 | 0.02 | 0.05 | 0.02 | -0.05 | 7.6E-01 | 0.01 |
| **COPG2** | 0.08 | -0.32 | 0.12 | -0.17 | 0.06 | 0.15 | 7.6E-01 | 0.06 |
| **CUL2** | 0.00 | -0.06 | 0.03 | 0.09 | 0.05 | -0.11 | 7.6E-01 | 0.02 |
| **RFX7** | -0.07 | -0.01 | 0.12 | 0.09 | 0.01 | -0.16 | 7.6E-01 | -0.03 |
| **LSR** | -0.11 | 0.04 | 0.04 | 0.03 | -0.02 | 0.02 | 7.6E-01 | 0.02 |
| **CLDN3** | -0.12 | 0.06 | 0.09 | 0.05 | -0.13 | 0.03 | 7.6E-01 | -0.03 |
| **GTF2H3** | -0.14 | 0.00 | 0.06 | 0.01 | -0.19 | 0.22 | 7.6E-01 | 0.04 |
| **PRORP** | 0.13 | 0.05 | -0.15 | 0.02 | -0.05 | -0.02 | 7.6E-01 | -0.03 |
| **TMEM67** | -0.11 | 0.18 | -0.13 | -0.02 | 0.01 | 0.05 | 7.6E-01 | 0.03 |
| **CEP55** | 0.08 | 0.06 | -0.11 | 0.14 | -0.05 | -0.15 | 7.6E-01 | -0.03 |
| **FBXO28** | 0.01 | -0.03 | 0.05 | -0.13 | -0.01 | 0.11 | 7.6E-01 | -0.02 |
| **MTIF2** | 0.19 | -0.05 | -0.10 | -0.06 | -0.12 | 0.11 | 7.6E-01 | -0.04 |
| **NT5DC2** | -0.48 | -0.18 | 0.67 | -0.57 | 0.10 | 0.09 | 7.7E-01 | -0.13 |
| **RPP25** | -0.09 | -0.16 | 0.18 | -0.01 | 0.12 | -0.07 | 7.7E-01 | 0.04 |
| **CHMP3** | 0.08 | 0.04 | -0.09 | -0.02 | 0.04 | -0.05 | 7.7E-01 | -0.02 |
| **PHPT1** | 0.05 | -0.03 | -0.01 | 0.03 | 0.01 | -0.06 | 7.7E-01 | -0.01 |
| **TBC1D30** | -0.03 | -0.03 | 0.04 | -0.03 | 0.03 | 0.01 | 7.7E-01 | 0.01 |
| **PHF2** | 0.06 | 0.00 | -0.04 | -0.03 | 0.02 | -0.01 | 7.7E-01 | -0.01 |
| **CHPF2** | -0.06 | 0.01 | 0.08 | 0.08 | 0.01 | -0.13 | 7.7E-01 | -0.02 |
| **BABAM2** | 0.15 | -0.20 | 0.09 | -0.01 | 0.06 | -0.12 | 7.7E-01 | -0.04 |
| **ARID2** | 0.00 | 0.00 | -0.01 | -0.01 | -0.03 | 0.05 | 7.7E-01 | 0.01 |
| **NUMB** | -0.10 | 0.07 | 0.06 | -0.05 | 0.04 | -0.02 | 7.7E-01 | -0.02 |
| **SP4** | -0.02 | -0.07 | 0.06 | -0.06 | 0.12 | -0.03 | 7.7E-01 | 0.02 |
| **NCKAP1** | -0.02 | -0.01 | 0.06 | -0.11 | 0.07 | 0.01 | 7.7E-01 | -0.02 |
| **MGME1** | 0.09 | 0.10 | -0.16 | 0.04 | -0.01 | -0.08 | 7.7E-01 | -0.03 |
| **PLCD3** | -0.11 | -0.03 | 0.10 | 0.07 | -0.10 | 0.06 | 7.7E-01 | 0.03 |
| **CLINT1** | -0.02 | 0.01 | 0.04 | 0.11 | -0.02 | -0.12 | 7.7E-01 | -0.02 |
| **BRD7** | 0.03 | 0.00 | -0.01 | -0.07 | -0.02 | 0.07 | 7.7E-01 | -0.01 |

| | | | | | | | | |
|---|---|---|---|---|---|---|---|---|
| **KDM4A** | 0.10 | 0.13 | -0.35 | 0.13 | 0.07 | -0.15 | 7.7E-01 | 0.05 |
| **CHMP4C** | -0.12 | -0.03 | 0.19 | 0.04 | -0.13 | 0.02 | 7.7E-01 | -0.03 |
| **CENPT** | -0.02 | 0.00 | -0.03 | -0.13 | -0.02 | 0.19 | 7.7E-01 | 0.03 |
| **ETF1** | 0.08 | 0.02 | -0.06 | 0.21 | -0.11 | -0.17 | 7.7E-01 | -0.04 |
| **SAPCD2** | -0.06 | 0.05 | -0.01 | -0.05 | 0.04 | 0.03 | 7.8E-01 | 0.01 |
| **CREB1** | -0.05 | -0.08 | 0.10 | 0.02 | -0.09 | 0.10 | 7.8E-01 | 0.02 |
| **RPS15A** | -0.05 | -0.07 | 0.09 | -0.03 | 0.02 | 0.03 | 7.8E-01 | 0.02 |
| **MED20** | 0.00 | -0.05 | 0.00 | 0.18 | -0.03 | -0.12 | 7.8E-01 | 0.03 |
| **EIF2D** | -0.09 | 0.06 | 0.06 | -0.11 | 0.05 | 0.02 | 7.8E-01 | -0.02 |
| **CCDC149** | -0.23 | 0.00 | 0.14 | 0.21 | -0.15 | -0.01 | 7.8E-01 | 0.05 |
| **TBCC** | -0.14 | 0.06 | 0.13 | 0.21 | -0.41 | 0.07 | 7.8E-01 | -0.06 |
| **MUC1** | 0.06 | -0.09 | 0.06 | 0.10 | -0.14 | 0.00 | 7.8E-01 | -0.03 |
| **ATP5MC3** | -0.24 | 0.16 | 0.12 | -0.06 | -0.14 | 0.10 | 7.8E-01 | -0.04 |
| **ZNF580** | -0.09 | -0.03 | 0.06 | -0.07 | -0.10 | 0.21 | 7.8E-01 | 0.03 |
| **PPP1R13B** | -0.11 | 0.04 | 0.10 | -0.20 | 0.09 | 0.04 | 7.8E-01 | -0.03 |
| **B4GAT1** | 0.07 | -0.12 | 0.08 | 0.15 | -0.16 | -0.05 | 7.8E-01 | -0.03 |
| **PNKD** | -0.06 | -0.06 | 0.14 | -0.07 | -0.02 | 0.05 | 7.8E-01 | -0.02 |
| **MTX1** | -0.03 | -0.06 | 0.07 | 0.01 | 0.00 | 0.01 | 7.8E-01 | 0.01 |
| **PRDX3** | -0.04 | 0.10 | -0.02 | 0.21 | -0.13 | -0.15 | 7.8E-01 | -0.04 |
| **SMTN** | 0.03 | -0.01 | -0.04 | 0.05 | 0.06 | -0.09 | 7.8E-01 | 0.02 |
| **LRRC41** | -0.28 | 0.08 | 0.12 | 0.10 | -0.10 | 0.04 | 7.8E-01 | 0.04 |
| **USP15** | -0.11 | 0.01 | 0.14 | -0.16 | 0.00 | 0.10 | 7.8E-01 | -0.03 |
| **YTHDF2** | -0.01 | 0.02 | 0.03 | 0.19 | -0.13 | -0.13 | 7.8E-01 | -0.03 |
| **RP2** | -0.08 | 0.05 | 0.05 | 0.00 | 0.01 | -0.03 | 7.8E-01 | -0.01 |
| **NIPBL** | 0.01 | -0.02 | 0.03 | -0.08 | -0.04 | 0.09 | 7.8E-01 | -0.02 |
| **GPRC5A** | -0.04 | -0.06 | 0.01 | -0.26 | 0.14 | 0.16 | 7.8E-01 | 0.04 |
| **NEK1** | -0.06 | 0.02 | 0.08 | -0.20 | 0.22 | -0.09 | 7.8E-01 | -0.04 |
| **XPNPEP3** | 0.05 | 0.02 | -0.12 | 0.01 | -0.14 | 0.16 | 7.8E-01 | 0.03 |
| **FRG1** | 0.02 | -0.01 | 0.01 | -0.07 | -0.01 | 0.06 | 7.9E-01 | -0.01 |
| **KMT2B** | -0.06 | 0.02 | 0.01 | -0.03 | -0.03 | 0.08 | 7.9E-01 | 0.01 |
| **LEPROTL1** | -0.12 | 0.02 | 0.14 | -0.33 | 0.09 | 0.13 | 7.9E-01 | -0.05 |
| **DNAJB12** | 0.06 | -0.02 | -0.02 | 0.01 | -0.07 | 0.04 | 7.9E-01 | -0.01 |
| **KRR1** | 0.09 | 0.01 | -0.08 | 0.06 | -0.01 | -0.08 | 7.9E-01 | -0.02 |
| **FLOT1** | -0.06 | -0.01 | 0.08 | -0.05 | 0.03 | 0.00 | 7.9E-01 | -0.01 |
| **POLR2C** | 0.12 | 0.03 | -0.11 | 0.13 | -0.06 | -0.12 | 7.9E-01 | -0.03 |
| **RBBP7** | 0.01 | -0.03 | 0.04 | -0.10 | 0.02 | 0.06 | 7.9E-01 | -0.01 |
| **TSC22D1** | -0.03 | 0.11 | -0.04 | -0.43 | 0.25 | 0.05 | 7.9E-01 | -0.06 |
| **RCBTB1** | 0.01 | -0.23 | 0.14 | -0.06 | 0.14 | -0.04 | 7.9E-01 | 0.04 |
| **DNAJC16** | 0.03 | -0.01 | -0.06 | 0.12 | -0.01 | -0.08 | 7.9E-01 | 0.02 |
| **TOMM20** | 0.03 | -0.03 | -0.01 | 0.06 | 0.00 | -0.05 | 7.9E-01 | 0.01 |
| **TCERG1** | 0.09 | -0.05 | -0.07 | -0.04 | 0.00 | 0.06 | 7.9E-01 | 0.02 |
| **SEC61A1** | -0.02 | -0.01 | 0.08 | 0.29 | -0.22 | -0.18 | 7.9E-01 | -0.05 |
| **C14orf93** | 0.01 | 0.08 | -0.05 | -0.15 | -0.13 | 0.21 | 7.9E-01 | -0.03 |
| **QSOX2** | -0.07 | -0.06 | 0.09 | 0.11 | -0.14 | 0.06 | 7.9E-01 | 0.03 |
| **TMBIM6** | -0.01 | 0.00 | 0.04 | -0.05 | 0.14 | -0.12 | 7.9E-01 | -0.02 |
| **SLC35F6** | -0.09 | 0.00 | 0.12 | 0.04 | 0.04 | -0.12 | 7.9E-01 | -0.02 |
| **G6PD** | 0.09 | -0.02 | -0.13 | 0.18 | -0.02 | -0.13 | 7.9E-01 | 0.03 |
| **TRPM4** | -0.04 | 0.01 | 0.02 | -0.01 | -0.03 | 0.05 | 7.9E-01 | 0.01 |

| | | | | | | | | |
|---|---|---|---|---|---|---|---|---|
| TENT2 | -0.04 | 0.01 | 0.05 | -0.01 | -0.07 | 0.05 | 7.9E-01 | -0.01 |
| MAP3K20 | -0.01 | -0.19 | 0.14 | -0.03 | 0.10 | -0.04 | 7.9E-01 | 0.03 |
| DGKE | 0.01 | 0.01 | 0.01 | 0.03 | 0.09 | -0.15 | 7.9E-01 | -0.02 |
| MEIKIN | -0.09 | 0.07 | 0.06 | -0.25 | -0.05 | 0.22 | 7.9E-01 | -0.04 |
| PMEPA1 | -0.11 | -0.18 | 0.33 | 0.01 | -0.30 | 0.16 | 7.9E-01 | -0.06 |
| SNX17 | 0.03 | -0.08 | 0.07 | -0.08 | -0.05 | 0.10 | 7.9E-01 | -0.02 |
| MRGBP | -0.05 | 0.03 | 0.04 | -0.06 | 0.01 | 0.03 | 7.9E-01 | -0.01 |
| ZCWPW1 | -0.13 | 0.04 | 0.04 | -0.09 | 0.09 | 0.03 | 7.9E-01 | 0.02 |
| CLTA | -0.03 | 0.02 | 0.03 | 0.10 | -0.03 | -0.10 | 8.0E-01 | -0.02 |
| PSMA3 | 0.14 | -0.21 | 0.00 | 0.21 | -0.10 | -0.07 | 8.0E-01 | 0.04 |
| ANKLE2 | -0.06 | 0.01 | 0.05 | 0.00 | -0.01 | -0.01 | 8.0E-01 | -0.01 |
| LIMCH1 | -0.04 | 0.01 | 0.01 | 0.04 | 0.01 | -0.03 | 8.0E-01 | 0.01 |
| ARPC2 | 0.03 | 0.05 | -0.07 | -0.03 | 0.00 | 0.02 | 8.0E-01 | -0.01 |
| ZCRB1 | -0.04 | -0.05 | 0.06 | 0.00 | -0.06 | 0.08 | 8.0E-01 | 0.01 |
| MRPS34 | 0.02 | -0.10 | 0.10 | -0.12 | -0.04 | 0.12 | 8.0E-01 | -0.02 |
| DPP9 | 0.03 | -0.12 | 0.00 | 0.29 | -0.08 | -0.17 | 8.0E-01 | 0.04 |
| ADAR | 0.03 | -0.01 | -0.01 | -0.04 | 0.01 | 0.02 | 8.0E-01 | -0.01 |
| MRPS14 | -0.03 | 0.10 | -0.11 | -0.02 | -0.05 | 0.10 | 8.0E-01 | 0.02 |
| TAOK2 | 0.00 | -0.04 | 0.06 | 0.01 | 0.02 | -0.05 | 8.0E-01 | -0.01 |
| HGFAC | 0.00 | 0.00 | -0.06 | -0.04 | -0.13 | 0.20 | 8.0E-01 | 0.03 |
| METAP1 | -0.01 | -0.05 | 0.08 | 0.00 | -0.06 | 0.04 | 8.0E-01 | -0.01 |
| POMGNT2 | -0.07 | -0.05 | 0.15 | -0.11 | 0.02 | 0.05 | 8.0E-01 | -0.02 |
| ADSS2 | 0.03 | -0.15 | 0.07 | -0.11 | -0.02 | 0.16 | 8.0E-01 | 0.03 |
| PAFAH1B2 | 0.01 | -0.04 | 0.06 | -0.29 | 0.21 | 0.00 | 8.0E-01 | -0.04 |
| RECQL5 | -0.02 | 0.04 | -0.01 | 0.00 | 0.00 | -0.02 | 8.0E-01 | 0.00 |
| EHMT1 | 0.04 | 0.00 | -0.02 | -0.10 | 0.02 | 0.06 | 8.0E-01 | -0.01 |
| GEMIN7 | -0.20 | -0.01 | 0.15 | 0.01 | -0.06 | 0.09 | 8.0E-01 | 0.03 |
| TSN | -0.20 | 0.03 | 0.20 | -0.15 | -0.14 | 0.19 | 8.0E-01 | -0.04 |
| WDR5 | 0.04 | 0.04 | -0.11 | 0.05 | 0.01 | -0.04 | 8.1E-01 | 0.01 |
| MAP2K1 | 0.04 | -0.01 | -0.06 | 0.02 | 0.08 | -0.08 | 8.1E-01 | 0.01 |
| SIL1 | 0.22 | 0.08 | -0.26 | 0.05 | -0.17 | 0.03 | 8.1E-01 | -0.04 |
| CENPX | -0.11 | 0.07 | 0.07 | -0.06 | -0.13 | 0.14 | 8.1E-01 | -0.03 |
| RIOX2 | -0.04 | 0.01 | 0.01 | 0.10 | -0.02 | -0.06 | 8.1E-01 | 0.01 |
| HINT3 | -0.17 | -0.14 | 0.17 | -0.31 | 0.03 | 0.32 | 8.1E-01 | 0.06 |
| FIBP | -0.10 | 0.00 | 0.07 | -0.02 | 0.09 | -0.05 | 8.1E-01 | 0.02 |
| HTATIP2 | 0.00 | 0.14 | -0.19 | -0.06 | 0.10 | 0.00 | 8.1E-01 | 0.03 |
| AKAP8L | -0.03 | 0.00 | 0.02 | 0.03 | 0.02 | -0.04 | 8.1E-01 | 0.01 |
| KIAA1143 | -0.19 | 0.19 | 0.04 | -0.40 | 0.25 | 0.02 | 8.1E-01 | -0.06 |
| ADCY6 | -0.15 | -0.07 | 0.15 | 0.17 | 0.05 | -0.19 | 8.1E-01 | 0.04 |
| CENPF | -0.01 | 0.00 | 0.00 | 0.02 | -0.03 | 0.02 | 8.1E-01 | 0.00 |
| TIMM10B | 0.02 | -0.05 | -0.01 | 0.15 | -0.02 | -0.10 | 8.1E-01 | 0.02 |
| CDK13 | -0.01 | 0.07 | -0.08 | 0.00 | 0.02 | -0.01 | 8.1E-01 | 0.01 |
| DYNC1LI1 | 0.00 | 0.01 | -0.03 | -0.09 | 0.10 | 0.00 | 8.1E-01 | 0.01 |
| YIF1B | -0.11 | 0.01 | 0.11 | -0.04 | 0.02 | -0.01 | 8.1E-01 | -0.02 |
| NDC80 | -0.12 | -0.04 | 0.05 | -0.32 | 0.03 | 0.32 | 8.1E-01 | 0.05 |
| TMPO | -0.02 | 0.03 | -0.04 | 0.07 | -0.09 | 0.03 | 8.1E-01 | 0.01 |
| POLR1A | -0.01 | -0.08 | 0.07 | -0.05 | 0.03 | 0.04 | 8.1E-01 | 0.01 |
| SF3B6 | 0.14 | -0.05 | -0.06 | 0.11 | -0.02 | -0.14 | 8.1E-01 | -0.02 |

| | | | | | | | | |
|---|---|---|---|---|---|---|---|---|
| TRAPPC4 | 0.07 | -0.05 | -0.07 | -0.10 | 0.19 | -0.06 | 8.1E-01 | 0.03 |
| TSG101 | 0.01 | 0.01 | -0.01 | -0.05 | 0.05 | -0.02 | 8.1E-01 | -0.01 |
| TTC39C | -0.27 | -0.01 | 0.31 | 0.13 | -0.25 | -0.01 | 8.1E-01 | -0.05 |
| RAB17 | -0.10 | 0.00 | 0.11 | -0.06 | -0.04 | 0.07 | 8.1E-01 | -0.02 |
| COMMD4 | -0.15 | -0.05 | 0.15 | 0.02 | -0.05 | 0.06 | 8.1E-01 | 0.02 |
| CD2AP | 0.01 | 0.02 | -0.05 | -0.04 | 0.00 | 0.05 | 8.1E-01 | 0.01 |
| SMARCB1 | 0.11 | 0.05 | -0.23 | -0.13 | 0.08 | 0.08 | 8.1E-01 | 0.03 |
| LRIF1 | -0.03 | -0.01 | 0.04 | -0.02 | 0.00 | 0.01 | 8.1E-01 | -0.01 |
| ASF1A | 0.14 | 0.09 | -0.20 | -0.09 | 0.09 | -0.06 | 8.1E-01 | -0.03 |
| ABCC11 | -0.18 | 0.10 | 0.04 | 0.06 | -0.02 | -0.02 | 8.1E-01 | 0.02 |
| LYPLA2 | -0.05 | -0.09 | 0.11 | 0.05 | 0.00 | -0.03 | 8.2E-01 | 0.02 |
| MZT2B | -0.04 | 0.08 | -0.07 | 0.11 | -0.02 | -0.07 | 8.2E-01 | 0.02 |
| SAMD4B | 0.01 | -0.06 | 0.06 | -0.03 | -0.02 | 0.04 | 8.2E-01 | -0.01 |
| ACP2 | 0.00 | -0.09 | 0.07 | -0.05 | 0.01 | 0.05 | 8.2E-01 | 0.01 |
| SLC30A7 | -0.09 | -0.11 | 0.14 | 0.15 | -0.08 | -0.04 | 8.2E-01 | 0.03 |
| ASXL3 | 0.09 | 0.03 | -0.10 | 0.02 | 0.06 | -0.11 | 8.2E-01 | -0.02 |
| ATP6V1F | 0.02 | 0.00 | -0.05 | 0.10 | -0.10 | 0.02 | 8.2E-01 | 0.01 |
| CEP41 | -0.11 | 0.14 | -0.10 | -0.22 | 0.08 | 0.18 | 8.2E-01 | 0.04 |
| ARFGAP3 | 0.04 | 0.01 | -0.07 | -0.02 | 0.07 | -0.03 | 8.2E-01 | 0.01 |
| NMNAT1 | -0.03 | 0.15 | -0.19 | 0.19 | -0.06 | -0.10 | 8.2E-01 | 0.03 |
| RPL15 | 0.02 | -0.04 | 0.04 | 0.10 | 0.05 | -0.19 | 8.2E-01 | -0.02 |
| KRTCAP3 | -0.17 | -0.04 | 0.15 | 0.12 | -0.15 | 0.05 | 8.2E-01 | 0.03 |
| SUCLG1 | 0.11 | 0.10 | -0.26 | -0.02 | 0.04 | 0.01 | 8.2E-01 | 0.03 |
| RPL10 | -0.04 | -0.01 | 0.06 | 0.04 | -0.01 | -0.04 | 8.2E-01 | -0.01 |
| HMG20B | -0.02 | -0.01 | 0.04 | -0.07 | -0.01 | 0.06 | 8.3E-01 | -0.01 |
| CAMSAP3 | -0.03 | 0.02 | 0.01 | -0.01 | 0.00 | 0.00 | 8.3E-01 | 0.00 |
| COQ3 | 0.17 | 0.27 | -0.61 | 0.06 | 0.07 | -0.11 | 8.3E-01 | 0.07 |
| MFSD6 | -0.18 | 0.11 | 0.10 | -0.20 | 0.01 | 0.12 | 8.3E-01 | -0.03 |
| CHCHD1 | 0.22 | -0.07 | -0.13 | -0.19 | 0.11 | 0.01 | 8.3E-01 | -0.03 |
| FOXA1 | 0.01 | -0.04 | 0.04 | -0.06 | -0.03 | 0.07 | 8.3E-01 | -0.01 |
| SRCAP | 0.03 | 0.00 | -0.02 | -0.08 | -0.05 | 0.10 | 8.3E-01 | -0.01 |
| TIMM50 | -0.02 | 0.03 | 0.02 | 0.12 | -0.20 | 0.04 | 8.3E-01 | -0.02 |
| STXBP2 | -0.11 | -0.03 | 0.11 | -0.01 | 0.03 | -0.01 | 8.3E-01 | 0.01 |
| MCCC2 | 0.09 | 0.00 | -0.07 | 0.12 | -0.06 | -0.10 | 8.3E-01 | -0.02 |
| MTFR1L | -0.07 | 0.03 | 0.05 | -0.08 | 0.06 | 0.00 | 8.3E-01 | -0.01 |
| SLC6A9 | -0.19 | 0.01 | 0.14 | 0.01 | -0.04 | 0.05 | 8.3E-01 | 0.02 |
| MED17 | -0.03 | -0.03 | 0.08 | 0.05 | -0.04 | -0.03 | 8.3E-01 | -0.01 |
| MED4 | -0.15 | -0.06 | 0.23 | 0.08 | -0.14 | 0.00 | 8.3E-01 | -0.03 |
| GTF2E1 | -0.06 | -0.05 | 0.07 | -0.15 | 0.04 | 0.13 | 8.3E-01 | 0.02 |
| TMEM63C | -0.16 | 0.00 | 0.19 | -0.17 | -0.01 | 0.12 | 8.3E-01 | -0.03 |
| PGAP2 | -0.07 | 0.08 | -0.04 | 0.04 | -0.08 | 0.06 | 8.3E-01 | 0.01 |
| CERS5 | -0.37 | -0.18 | 0.31 | -0.56 | 0.32 | 0.24 | 8.3E-01 | 0.08 |
| REXO1 | 0.03 | 0.01 | -0.03 | 0.03 | -0.06 | 0.02 | 8.3E-01 | -0.01 |
| YKT6 | -0.05 | -0.02 | 0.10 | 0.12 | -0.12 | -0.03 | 8.3E-01 | -0.02 |
| SLTM | 0.04 | 0.00 | -0.05 | -0.02 | -0.03 | 0.06 | 8.3E-01 | 0.01 |
| CYP20A1 | 0.04 | 0.05 | -0.12 | -0.03 | 0.07 | -0.02 | 8.3E-01 | 0.01 |
| CCNT1 | -0.07 | -0.02 | 0.02 | 0.27 | -0.12 | -0.13 | 8.3E-01 | 0.03 |
| SELENBP1 | 0.05 | 0.05 | -0.08 | -0.11 | 0.08 | 0.00 | 8.3E-01 | -0.02 |

| | | | | | | | | |
|---|---|---|---|---|---|---|---|---|
| ZNF185 | -0.26 | 0.09 | 0.10 | -0.20 | 0.09 | 0.14 | 8.3E-01 | 0.04 |
| POLD2 | -0.02 | 0.03 | -0.05 | -0.16 | 0.11 | 0.08 | 8.3E-01 | 0.02 |
| MTHFS | -0.08 | -0.01 | 0.05 | 0.01 | -0.12 | 0.14 | 8.3E-01 | 0.02 |
| MIB2 | -0.10 | 0.04 | 0.08 | 0.12 | -0.18 | 0.02 | 8.3E-01 | -0.02 |
| RNF130 | -0.38 | -0.10 | 0.49 | -0.11 | 0.01 | -0.06 | 8.3E-01 | -0.06 |
| ARHGDIB | -0.09 | -0.08 | 0.14 | 0.00 | 0.02 | 0.00 | 8.4E-01 | 0.02 |
| BACH1 | -0.09 | 0.12 | -0.01 | 0.12 | 0.04 | -0.21 | 8.4E-01 | -0.03 |
| POLR2J | 0.14 | 0.04 | -0.16 | 0.01 | 0.00 | -0.05 | 8.4E-01 | -0.02 |
| KAT6B | 0.02 | 0.03 | -0.04 | -0.09 | -0.01 | 0.08 | 8.4E-01 | -0.01 |
| ZBTB2 | 0.15 | -0.17 | 0.05 | -0.24 | 0.08 | 0.09 | 8.4E-01 | -0.03 |
| TJP2 | -0.09 | 0.04 | 0.02 | 0.06 | 0.00 | -0.04 | 8.4E-01 | 0.01 |
| VAV2 | -0.07 | 0.06 | 0.03 | -0.13 | 0.06 | 0.04 | 8.4E-01 | -0.02 |
| MGAT4A | -0.10 | 0.06 | 0.00 | -0.16 | 0.10 | 0.08 | 8.4E-01 | 0.02 |
| IFI30 | -0.07 | -0.10 | 0.20 | -0.12 | 0.13 | -0.06 | 8.4E-01 | -0.03 |
| CD151 | -0.02 | -0.02 | 0.07 | -0.31 | 0.14 | 0.09 | 8.4E-01 | -0.03 |
| RBBP4 | -0.08 | -0.11 | 0.15 | 0.00 | 0.00 | 0.02 | 8.4E-01 | 0.02 |
| C1orf174 | -0.07 | -0.03 | 0.03 | -0.27 | 0.09 | 0.20 | 8.4E-01 | 0.03 |
| NDC1 | 0.05 | -0.01 | -0.05 | 0.04 | -0.01 | -0.02 | 8.4E-01 | 0.01 |
| PARD6A | -0.11 | 0.05 | 0.08 | -0.09 | 0.08 | -0.02 | 8.4E-01 | -0.02 |
| EMC2 | 0.02 | -0.04 | -0.01 | 0.12 | -0.01 | -0.10 | 8.4E-01 | 0.01 |
| CAPN1 | 0.16 | 0.05 | -0.19 | 0.05 | -0.05 | -0.05 | 8.4E-01 | -0.02 |
| TIMM9 | 0.00 | 0.07 | -0.05 | 0.08 | 0.00 | -0.11 | 8.4E-01 | -0.01 |
| GNPAT | 0.04 | 0.06 | -0.13 | -0.12 | 0.07 | 0.07 | 8.4E-01 | 0.02 |
| THUMPD1 | 0.05 | 0.06 | -0.09 | -0.14 | 0.13 | -0.03 | 8.4E-01 | -0.02 |
| NBAS | 0.01 | -0.04 | 0.04 | 0.02 | -0.03 | 0.00 | 8.5E-01 | -0.01 |
| EZR | 0.00 | -0.03 | 0.04 | 0.04 | -0.08 | 0.03 | 8.5E-01 | -0.01 |
| SART1 | 0.07 | -0.01 | -0.08 | 0.09 | -0.03 | -0.04 | 8.5E-01 | 0.01 |
| MFN1 | -0.08 | 0.03 | 0.07 | -0.06 | -0.09 | 0.12 | 8.5E-01 | -0.02 |
| SLC9A1 | -0.11 | -0.04 | 0.12 | 0.01 | -0.05 | 0.05 | 8.5E-01 | 0.02 |
| KIF20B | -0.02 | -0.04 | 0.04 | 0.07 | -0.07 | 0.02 | 8.5E-01 | 0.01 |
| PKM | 0.05 | 0.05 | -0.08 | -0.04 | 0.02 | 0.00 | 8.5E-01 | -0.01 |
| SARS1 | 0.03 | 0.04 | -0.08 | 0.02 | -0.04 | 0.03 | 8.5E-01 | 0.01 |
| NPDC1 | -0.06 | -0.13 | 0.11 | 0.30 | -0.18 | -0.09 | 8.5E-01 | 0.03 |
| EFL1 | -0.23 | 0.06 | 0.11 | 0.14 | -0.17 | 0.05 | 8.5E-01 | 0.03 |
| RBM4B | 0.06 | -0.03 | -0.01 | 0.29 | -0.14 | -0.23 | 8.5E-01 | -0.03 |
| BEND3 | -0.02 | -0.06 | 0.06 | -0.07 | 0.05 | 0.03 | 8.5E-01 | 0.01 |
| STAG1 | 0.06 | -0.02 | -0.03 | -0.01 | -0.04 | 0.04 | 8.5E-01 | -0.01 |
| FOXRED1 | 0.02 | -0.01 | -0.02 | 0.03 | -0.07 | 0.04 | 8.5E-01 | 0.01 |
| ELF3 | -0.11 | 0.03 | 0.05 | -0.16 | 0.11 | 0.07 | 8.5E-01 | 0.02 |
| GPAA1 | 0.04 | 0.00 | -0.02 | 0.07 | -0.06 | -0.02 | 8.5E-01 | -0.01 |
| UQCRHL | -0.01 | -0.02 | 0.00 | 0.12 | -0.10 | -0.01 | 8.5E-01 | 0.01 |
| TACO1 | 0.14 | 0.01 | -0.13 | -0.01 | 0.08 | -0.11 | 8.5E-01 | -0.02 |
| FASTK | 0.04 | -0.10 | 0.02 | -0.14 | -0.04 | 0.19 | 8.5E-01 | 0.02 |
| KNSTRN | -0.08 | -0.07 | 0.17 | -0.11 | -0.12 | 0.17 | 8.5E-01 | -0.02 |
| CXXC5 | -0.03 | -0.05 | 0.10 | -0.12 | 0.01 | 0.08 | 8.5E-01 | -0.02 |
| MGP | 0.04 | -0.06 | 0.04 | -0.34 | 0.11 | 0.15 | 8.5E-01 | -0.03 |
| EPHX1 | 0.12 | -0.01 | -0.14 | 0.00 | 0.03 | -0.01 | 8.5E-01 | 0.02 |
| ANXA4 | -0.02 | -0.05 | 0.09 | -0.02 | 0.04 | -0.04 | 8.5E-01 | -0.01 |

| | | | | | | | | |
|---|---|---|---|---|---|---|---|---|
| **TTC1** | 0.03 | -0.07 | 0.06 | -0.28 | 0.15 | 0.07 | 8.5E-01 | -0.03 |
| **PPP2R5C** | 0.01 | -0.05 | 0.06 | 0.04 | -0.10 | 0.05 | 8.5E-01 | -0.01 |
| **SMG1** | -0.12 | 0.04 | 0.09 | -0.07 | 0.01 | 0.04 | 8.6E-01 | -0.01 |
| **ZNF646** | 0.02 | -0.04 | 0.02 | -0.02 | -0.01 | 0.03 | 8.6E-01 | 0.00 |
| **TEF** | -0.09 | 0.08 | -0.01 | 0.08 | -0.02 | -0.04 | 8.6E-01 | 0.01 |
| **RMC1** | 0.05 | 0.11 | -0.14 | -0.12 | 0.06 | 0.02 | 8.6E-01 | -0.02 |
| **CDAN1** | 0.01 | -0.05 | -0.02 | -0.16 | -0.10 | 0.29 | 8.6E-01 | 0.03 |
| **ME2** | 0.13 | 0.10 | -0.22 | -0.07 | 0.04 | -0.01 | 8.6E-01 | -0.02 |
| **SYTL4** | -0.06 | 0.02 | 0.05 | -0.11 | 0.01 | 0.08 | 8.6E-01 | -0.01 |
| **GNB1L** | 0.10 | -0.12 | -0.08 | 0.35 | -0.01 | -0.33 | 8.6E-01 | 0.04 |
| **NOSIP** | 0.01 | 0.03 | -0.03 | -0.07 | 0.03 | 0.02 | 8.6E-01 | -0.01 |
| **PBRM1** | 0.06 | -0.01 | -0.06 | 0.04 | -0.03 | 0.00 | 8.6E-01 | 0.01 |
| **DNAAF10** | -0.03 | -0.06 | 0.11 | -0.22 | 0.06 | 0.11 | 8.6E-01 | -0.02 |
| **ATPAF2** | -0.01 | 0.14 | -0.21 | 0.07 | -0.25 | 0.20 | 8.6E-01 | 0.03 |
| **TMCO3** | -0.07 | 0.01 | 0.03 | 0.17 | -0.02 | -0.13 | 8.6E-01 | 0.02 |
| **DNM1L** | 0.02 | 0.16 | -0.16 | -0.03 | 0.17 | -0.21 | 8.6E-01 | -0.03 |
| **GBE1** | 0.09 | -0.14 | 0.07 | -0.12 | 0.05 | 0.03 | 8.6E-01 | -0.02 |
| **VASP** | 0.12 | -0.05 | -0.11 | -0.10 | 0.12 | -0.01 | 8.6E-01 | 0.02 |
| **CIAPIN1** | 0.07 | 0.03 | -0.13 | 0.03 | 0.01 | -0.02 | 8.6E-01 | 0.01 |
| **TADA2B** | 0.09 | -0.05 | -0.03 | 0.20 | 0.06 | -0.34 | 8.6E-01 | -0.03 |
| **VPS13C** | -0.05 | -0.08 | 0.15 | -0.31 | 0.02 | 0.21 | 8.6E-01 | -0.03 |
| **NAA30** | 0.11 | 0.03 | -0.21 | -0.26 | 0.15 | 0.12 | 8.6E-01 | 0.03 |
| **TSC2** | -0.26 | 0.00 | 0.17 | -0.15 | -0.14 | 0.30 | 8.6E-01 | 0.03 |
| **FIP1L1** | 0.02 | 0.01 | -0.01 | 0.11 | -0.06 | -0.06 | 8.6E-01 | -0.01 |
| **SENP1** | 0.03 | 0.02 | -0.04 | 0.07 | 0.01 | -0.10 | 8.7E-01 | -0.01 |
| **TBCE** | 0.07 | 0.00 | -0.09 | 0.03 | 0.04 | -0.06 | 8.7E-01 | 0.01 |
| **AIDA** | -0.05 | 0.19 | -0.24 | 0.29 | 0.04 | -0.32 | 8.7E-01 | 0.04 |
| **INTS4** | -0.01 | 0.01 | 0.01 | 0.02 | -0.07 | 0.04 | 8.7E-01 | -0.01 |
| **CENPS** | 0.00 | 0.00 | -0.02 | -0.03 | -0.05 | 0.10 | 8.7E-01 | 0.01 |
| **RFXAP** | -0.04 | -0.04 | 0.09 | 0.02 | -0.01 | -0.03 | 8.7E-01 | -0.01 |
| **SMAD3** | -0.03 | 0.15 | -0.15 | -0.02 | 0.07 | -0.03 | 8.7E-01 | 0.02 |
| **PEX16** | 0.03 | -0.04 | -0.02 | 0.14 | -0.03 | -0.10 | 8.7E-01 | 0.01 |
| **COTL1** | -0.08 | 0.13 | -0.03 | 0.01 | 0.18 | -0.24 | 8.7E-01 | -0.02 |
| **MSL2** | -0.02 | -0.12 | 0.16 | -0.05 | -0.07 | 0.08 | 8.7E-01 | -0.02 |
| **BICRAL** | 0.09 | 0.03 | -0.17 | -0.11 | -0.08 | 0.20 | 8.7E-01 | 0.02 |
| **ENO2** | -0.15 | 0.10 | 0.07 | 0.15 | -0.03 | -0.16 | 8.7E-01 | -0.02 |
| **EIF2B2** | -0.03 | -0.04 | 0.04 | 0.12 | 0.03 | -0.14 | 8.7E-01 | 0.01 |
| **SELENOT** | 0.00 | -0.02 | 0.04 | 0.20 | -0.03 | -0.21 | 8.7E-01 | -0.02 |
| **TM9SF1** | -0.01 | 0.01 | 0.01 | 0.16 | -0.03 | -0.17 | 8.7E-01 | -0.02 |
| **UBE2D3** | 0.07 | 0.03 | -0.09 | 0.01 | 0.05 | -0.07 | 8.7E-01 | -0.01 |
| **TBC1D9B** | 0.06 | -0.17 | 0.13 | -0.30 | 0.07 | 0.16 | 8.7E-01 | -0.03 |
| **NTHL1** | 0.11 | 0.01 | -0.15 | 0.10 | -0.04 | -0.05 | 8.7E-01 | 0.02 |
| **TMEM143** | -0.13 | -0.07 | 0.21 | -0.06 | 0.01 | 0.01 | 8.7E-01 | -0.02 |
| **C22orf39** | -0.19 | 0.04 | 0.17 | -0.30 | 0.02 | 0.20 | 8.7E-01 | -0.03 |
| **NCAPG2** | -0.04 | -0.03 | 0.08 | 0.02 | -0.01 | -0.03 | 8.7E-01 | -0.01 |
| **RNFT2** | -0.09 | -0.02 | 0.04 | 0.21 | -0.29 | 0.10 | 8.7E-01 | 0.03 |
| **ERI1** | -0.06 | -0.02 | 0.09 | -0.06 | -0.03 | 0.07 | 8.8E-01 | -0.01 |
| **ILVBL** | -0.03 | 0.06 | -0.04 | -0.06 | 0.03 | 0.04 | 8.8E-01 | 0.01 |

| | | | | | | | | |
|---|---|---|---|---|---|---|---|---|
| TRABD | 0.00 | 0.03 | -0.03 | -0.08 | 0.06 | 0.01 | 8.8E-01 | -0.01 |
| DIPK2A | -0.16 | -0.06 | 0.18 | -0.06 | 0.08 | -0.01 | 8.8E-01 | 0.02 |
| RPS23 | -0.03 | 0.03 | 0.02 | 0.01 | 0.08 | -0.11 | 8.8E-01 | -0.01 |
| POLG2 | -0.04 | -0.01 | 0.03 | 0.13 | -0.04 | -0.08 | 8.8E-01 | 0.01 |
| NIP7 | 0.09 | -0.05 | -0.03 | 0.14 | -0.03 | -0.14 | 8.8E-01 | -0.02 |
| FAM76A | -0.04 | 0.01 | -0.01 | 0.22 | -0.05 | -0.16 | 8.8E-01 | 0.02 |
| SLC39A3 | -0.03 | -0.08 | 0.09 | 0.14 | -0.08 | -0.04 | 8.8E-01 | 0.01 |
| ZW10 | 0.03 | -0.04 | -0.01 | 0.07 | 0.00 | -0.06 | 8.8E-01 | 0.01 |
| ATRX | 0.01 | 0.03 | -0.03 | -0.03 | -0.02 | 0.04 | 8.8E-01 | 0.00 |
| USP24 | 0.15 | 0.14 | -0.47 | 0.33 | 0.13 | -0.48 | 8.8E-01 | 0.05 |
| UBE2O | 0.14 | -0.06 | -0.11 | -0.13 | 0.09 | 0.05 | 8.8E-01 | 0.02 |
| METTL17 | -0.06 | 0.03 | 0.05 | -0.04 | -0.15 | 0.15 | 8.8E-01 | -0.02 |
| TYW1 | 0.00 | 0.00 | -0.05 | 0.00 | 0.23 | -0.22 | 8.8E-01 | 0.02 |
| CRYZ | 0.16 | 0.09 | -0.24 | -0.02 | -0.07 | 0.05 | 8.8E-01 | -0.02 |
| GFER | 0.06 | 0.07 | -0.16 | -0.01 | 0.13 | -0.10 | 8.8E-01 | 0.02 |
| TM9SF4 | -0.04 | -0.01 | 0.03 | 0.12 | -0.05 | -0.06 | 8.9E-01 | 0.01 |
| STK11IP | -0.02 | -0.11 | 0.10 | -0.05 | -0.06 | 0.13 | 8.9E-01 | 0.01 |
| ICE1 | 0.08 | -0.39 | 0.22 | -0.11 | -0.10 | 0.22 | 8.9E-01 | 0.03 |
| HERC4 | 0.09 | -0.36 | 0.18 | -0.14 | -0.10 | 0.24 | 8.9E-01 | 0.03 |
| AHSG | 0.17 | -0.04 | -0.22 | -0.28 | 0.30 | -0.01 | 8.9E-01 | 0.03 |
| XPO7 | 0.06 | -0.19 | 0.09 | -0.03 | 0.16 | -0.12 | 8.9E-01 | 0.02 |
| COX6B1 | 0.01 | -0.02 | 0.02 | 0.04 | -0.02 | -0.02 | 8.9E-01 | 0.00 |
| ANKFY1 | -0.52 | -0.26 | 0.66 | -0.69 | 0.13 | 0.23 | 8.9E-01 | -0.07 |
| NF2 | -0.10 | -0.01 | 0.12 | -0.07 | 0.07 | -0.03 | 8.9E-01 | -0.01 |
| DNTTIP2 | -0.03 | -0.01 | 0.04 | -0.02 | 0.01 | 0.02 | 8.9E-01 | 0.00 |
| OSTM1 | -0.06 | 0.01 | 0.03 | 0.04 | -0.16 | 0.13 | 8.9E-01 | 0.01 |
| ATP6AP2 | 0.10 | -0.09 | -0.09 | 0.35 | -0.10 | -0.25 | 8.9E-01 | 0.03 |
| HELZ2 | -0.06 | 0.01 | 0.04 | 0.04 | 0.02 | -0.05 | 8.9E-01 | 0.01 |
| ACAD9 | -0.01 | 0.05 | -0.05 | 0.05 | 0.02 | -0.06 | 8.9E-01 | 0.01 |
| DLX1 | -0.01 | -0.29 | 0.24 | 0.13 | -0.12 | 0.00 | 8.9E-01 | 0.02 |
| NELFE | 0.01 | 0.05 | -0.05 | 0.06 | 0.02 | -0.09 | 8.9E-01 | -0.01 |
| NAPRT | 0.04 | -0.11 | 0.08 | -0.22 | 0.12 | 0.06 | 8.9E-01 | -0.02 |
| PLPP2 | 0.15 | -0.14 | -0.03 | 0.03 | -0.08 | 0.07 | 8.9E-01 | 0.01 |
| SLC25A46 | -0.39 | 0.39 | 0.00 | -0.17 | -0.01 | 0.08 | 8.9E-01 | -0.03 |
| QARS1 | -0.06 | 0.01 | 0.03 | 0.03 | 0.08 | -0.10 | 9.0E-01 | 0.01 |
| COX7C | -0.03 | -0.03 | 0.06 | -0.03 | -0.02 | 0.05 | 9.0E-01 | 0.01 |
| CLSPN | -0.01 | -0.06 | 0.08 | -0.14 | -0.03 | 0.14 | 9.0E-01 | -0.01 |
| EFNB2 | -0.15 | -0.03 | 0.19 | 0.07 | 0.00 | -0.11 | 9.0E-01 | -0.02 |
| GNB1 | -0.09 | -0.01 | 0.10 | 0.03 | -0.11 | 0.05 | 9.0E-01 | -0.01 |
| ARMT1 | -0.04 | 0.07 | -0.04 | -0.05 | 0.12 | -0.06 | 9.0E-01 | 0.01 |
| ANKIB1 | -0.15 | 0.04 | 0.12 | 0.09 | -0.05 | -0.07 | 9.0E-01 | -0.01 |
| NFIL3 | 0.01 | -0.04 | 0.04 | -0.08 | -0.06 | 0.12 | 9.0E-01 | -0.01 |
| NAP1L4 | -0.02 | -0.02 | 0.03 | 0.01 | 0.10 | -0.10 | 9.0E-01 | 0.01 |
| CCDC186 | -0.05 | -0.08 | 0.14 | 0.03 | -0.04 | -0.01 | 9.0E-01 | -0.01 |
| PIGC | -0.03 | 0.05 | -0.04 | -0.06 | 0.01 | 0.06 | 9.0E-01 | 0.01 |
| DAPK3 | -0.12 | 0.09 | 0.00 | -0.09 | 0.10 | 0.00 | 9.0E-01 | 0.01 |
| NDUFB9 | -0.07 | -0.05 | 0.12 | 0.04 | 0.00 | -0.06 | 9.0E-01 | -0.01 |
| USF2 | -0.05 | 0.00 | 0.03 | 0.01 | -0.08 | 0.07 | 9.0E-01 | 0.01 |

| | | | | | | | | |
|---|---|---|---|---|---|---|---|---|
| SLC7A1 | -0.22 | 0.00 | 0.18 | -0.08 | 0.03 | 0.05 | 9.0E-01 | 0.02 |
| MAZ | 0.03 | -0.05 | 0.00 | -0.20 | 0.16 | 0.05 | 9.0E-01 | 0.01 |
| PATZ1 | -0.02 | 0.01 | 0.00 | -0.04 | -0.06 | 0.11 | 9.0E-01 | 0.01 |
| PLEKHN1 | -0.13 | -0.05 | 0.16 | -0.01 | -0.03 | 0.06 | 9.0E-01 | 0.01 |
| PRPF3 | 0.04 | -0.01 | -0.04 | -0.02 | -0.01 | 0.03 | 9.1E-01 | 0.00 |
| ZNF135 | -0.16 | -0.04 | 0.16 | -0.08 | 0.19 | -0.11 | 9.1E-01 | 0.02 |
| RTN4R | -0.15 | 0.07 | 0.10 | -0.03 | -0.07 | 0.08 | 9.1E-01 | -0.01 |
| NOL11 | 0.04 | -0.01 | -0.03 | -0.01 | 0.03 | -0.01 | 9.1E-01 | 0.00 |
| HIP1R | -0.13 | 0.07 | 0.04 | 0.07 | 0.00 | -0.06 | 9.1E-01 | 0.01 |
| PSME2 | -0.11 | 0.01 | 0.08 | -0.06 | 0.02 | 0.05 | 9.1E-01 | 0.01 |
| AK9 | -0.10 | 0.12 | -0.06 | -0.23 | 0.10 | 0.14 | 9.1E-01 | 0.02 |
| NIPSNAP2 | 0.23 | 0.04 | -0.27 | 0.04 | -0.07 | -0.02 | 9.1E-01 | -0.02 |
| CBX1 | 0.05 | 0.03 | -0.08 | -0.13 | 0.08 | 0.02 | 9.1E-01 | -0.01 |
| TRMT61A | -0.14 | -0.03 | 0.13 | -0.22 | 0.10 | 0.13 | 9.1E-01 | 0.02 |
| CNDP2 | -0.06 | -0.01 | 0.03 | 0.22 | 0.02 | -0.24 | 9.1E-01 | 0.02 |
| TRIP6 | -0.06 | 0.04 | 0.01 | 0.01 | 0.05 | -0.05 | 9.1E-01 | 0.01 |
| CASZ1 | 0.03 | 0.00 | -0.03 | 0.00 | -0.03 | 0.02 | 9.1E-01 | 0.00 |
| EVI5L | -0.08 | 0.05 | 0.04 | -0.08 | -0.14 | 0.19 | 9.1E-01 | -0.01 |
| UBE3A | 0.14 | -0.13 | -0.04 | 0.31 | 0.14 | -0.58 | 9.1E-01 | -0.03 |
| ITGB4 | -0.20 | 0.01 | 0.16 | -0.02 | -0.04 | 0.07 | 9.1E-01 | 0.01 |
| FMN1 | -0.02 | 0.08 | -0.05 | 0.08 | -0.09 | -0.01 | 9.1E-01 | -0.01 |
| SEPHS1 | 0.53 | -0.17 | -0.40 | 0.09 | -0.17 | -0.07 | 9.1E-01 | -0.03 |
| ELAVL1 | 0.07 | -0.11 | 0.05 | -0.14 | -0.01 | 0.12 | 9.1E-01 | -0.01 |
| ANAPC5 | -0.03 | -0.08 | 0.11 | -0.06 | -0.02 | 0.07 | 9.1E-01 | -0.01 |
| RASAL1 | -0.11 | 0.06 | 0.06 | 0.07 | 0.00 | -0.08 | 9.1E-01 | -0.01 |
| C1GALT1C1 | -0.16 | -0.12 | 0.24 | -0.10 | 0.05 | 0.06 | 9.1E-01 | 0.02 |
| ZNF48 | -0.01 | 0.02 | -0.02 | -0.01 | 0.07 | -0.05 | 9.1E-01 | 0.00 |
| SMARCD1 | 0.05 | 0.03 | -0.10 | -0.09 | 0.03 | 0.07 | 9.1E-01 | 0.01 |
| SPRED2 | 0.06 | -0.09 | 0.02 | 0.01 | -0.04 | 0.04 | 9.1E-01 | 0.01 |
| GLB1 | 0.01 | -0.07 | 0.05 | 0.11 | -0.01 | -0.09 | 9.1E-01 | 0.01 |
| ITPRIPL2 | 0.00 | 0.01 | 0.00 | -0.08 | 0.09 | -0.02 | 9.1E-01 | -0.01 |
| GTF2H2 | -0.01 | -0.05 | 0.06 | 0.17 | -0.03 | -0.17 | 9.1E-01 | -0.01 |
| ISY1 | 0.03 | -0.05 | 0.02 | 0.03 | -0.06 | 0.04 | 9.1E-01 | 0.00 |
| RPL11 | 0.02 | 0.00 | -0.07 | 0.24 | -0.04 | -0.20 | 9.1E-01 | 0.02 |
| EHD1 | 0.03 | -0.01 | -0.02 | 0.00 | 0.06 | -0.07 | 9.1E-01 | 0.00 |
| AP1B1 | -0.08 | 0.02 | 0.07 | 0.15 | -0.07 | -0.11 | 9.1E-01 | -0.01 |
| SLC35E1 | -0.08 | -0.03 | 0.12 | -0.15 | 0.08 | 0.05 | 9.1E-01 | -0.01 |
| GNL1 | 0.04 | 0.02 | -0.08 | -0.16 | 0.13 | 0.05 | 9.1E-01 | 0.01 |
| DOCK10 | -0.12 | -0.04 | 0.17 | -0.05 | -0.01 | 0.04 | 9.1E-01 | -0.01 |
| INPP5K | -0.02 | 0.00 | 0.01 | -0.01 | 0.10 | -0.08 | 9.2E-01 | 0.01 |
| EIF3H | 0.02 | 0.03 | -0.04 | -0.01 | 0.03 | -0.03 | 9.2E-01 | 0.00 |
| TMEM109 | -0.04 | -0.05 | 0.09 | 0.01 | -0.09 | 0.06 | 9.2E-01 | -0.01 |
| PCBP4 | 0.10 | -0.09 | -0.06 | -0.26 | 0.05 | 0.21 | 9.2E-01 | 0.02 |
| COA3 | -0.06 | 0.05 | 0.02 | -0.13 | -0.02 | 0.13 | 9.2E-01 | -0.01 |
| MATR3 | 0.03 | -0.01 | -0.02 | -0.04 | -0.04 | 0.07 | 9.2E-01 | 0.00 |
| TBL2 | -0.01 | 0.01 | 0.01 | 0.12 | -0.01 | -0.13 | 9.2E-01 | -0.01 |
| MED16 | -0.02 | 0.06 | -0.05 | 0.08 | 0.01 | -0.09 | 9.2E-01 | 0.01 |
| RBM5 | 0.04 | -0.01 | -0.04 | -0.04 | 0.02 | 0.02 | 9.2E-01 | 0.00 |

| | | | | | | | | |
|---|---|---|---|---|---|---|---|---|
| GSTO1 | 0.08 | 0.02 | -0.09 | -0.10 | 0.10 | -0.01 | 9.2E-01 | -0.01 |
| MICALL2 | -0.11 | 0.05 | 0.05 | -0.06 | 0.00 | 0.07 | 9.2E-01 | 0.01 |
| SLC38A1 | -0.04 | -0.12 | 0.12 | -0.18 | -0.07 | 0.25 | 9.2E-01 | 0.02 |
| PLOD2 | -0.14 | -0.02 | 0.12 | 0.08 | -0.19 | 0.11 | 9.2E-01 | 0.01 |
| CENPC | 0.01 | 0.02 | -0.02 | -0.08 | 0.06 | 0.01 | 9.2E-01 | 0.00 |
| EIF1AD | 0.12 | -0.03 | -0.13 | -0.07 | -0.17 | 0.25 | 9.2E-01 | 0.02 |
| SCLT1 | 0.04 | -0.06 | 0.02 | -0.05 | 0.04 | 0.01 | 9.2E-01 | 0.00 |
| RPS26 | -0.07 | 0.00 | 0.05 | -0.12 | 0.12 | 0.01 | 9.2E-01 | 0.01 |
| RAB43 | -0.03 | -0.10 | 0.11 | 0.06 | -0.09 | 0.03 | 9.2E-01 | 0.01 |
| CLDND1 | -0.14 | 0.05 | 0.08 | -0.04 | -0.02 | 0.07 | 9.2E-01 | 0.01 |
| ATP6V1A | 0.00 | 0.02 | -0.04 | 0.09 | -0.05 | -0.04 | 9.2E-01 | 0.01 |
| ISG20L2 | 0.01 | -0.05 | 0.05 | -0.03 | 0.00 | 0.03 | 9.2E-01 | 0.00 |
| ZMAT4 | -0.03 | -0.15 | 0.15 | -0.15 | 0.01 | 0.15 | 9.2E-01 | 0.01 |
| HOMER1 | -0.12 | 0.06 | 0.03 | -0.19 | 0.04 | 0.16 | 9.2E-01 | 0.01 |
| DNAJC13 | -0.08 | 0.03 | 0.04 | 0.05 | -0.06 | 0.02 | 9.2E-01 | 0.01 |
| MACROD1 | 0.13 | 0.04 | -0.16 | -0.15 | 0.07 | 0.05 | 9.2E-01 | -0.01 |
| SUMO1P1 | -0.61 | -0.33 | 0.79 | 0.02 | -0.03 | -0.27 | 9.3E-01 | -0.04 |
| NOP56 | 0.04 | -0.01 | -0.03 | 0.01 | 0.04 | -0.06 | 9.3E-01 | 0.00 |
| GMPPB | 0.15 | -0.14 | -0.05 | -0.15 | 0.18 | -0.03 | 9.3E-01 | 0.01 |
| RPS15 | 0.05 | -0.04 | 0.00 | 0.14 | 0.06 | -0.22 | 9.3E-01 | -0.01 |
| MTCH2 | -0.06 | -0.02 | 0.08 | 0.06 | -0.05 | -0.02 | 9.3E-01 | -0.01 |
| CEP152 | -0.01 | -0.01 | 0.02 | -0.08 | 0.01 | 0.07 | 9.3E-01 | 0.00 |
| SLC30A5 | 0.02 | 0.02 | -0.06 | 0.19 | -0.02 | -0.16 | 9.3E-01 | 0.01 |
| RPL21 | -0.08 | -0.07 | 0.13 | 0.07 | 0.01 | -0.08 | 9.3E-01 | 0.01 |
| PPP1R15B | -0.07 | 0.06 | 0.01 | -0.14 | 0.02 | 0.10 | 9.3E-01 | -0.01 |
| PEF1 | 0.01 | -0.02 | 0.02 | 0.01 | 0.04 | -0.05 | 9.3E-01 | 0.00 |
| CENPW | 0.09 | 0.04 | -0.13 | -0.06 | 0.01 | 0.04 | 9.3E-01 | -0.01 |
| TPCN2 | -0.23 | 0.09 | 0.13 | -0.04 | 0.14 | -0.14 | 9.3E-01 | -0.01 |
| DDX17 | 0.02 | -0.03 | 0.01 | -0.02 | -0.02 | 0.04 | 9.3E-01 | 0.00 |
| RANBP6 | -0.06 | -0.02 | 0.07 | 0.01 | 0.05 | -0.05 | 9.3E-01 | 0.00 |
| ISOC1 | -0.04 | 0.07 | -0.05 | 0.05 | 0.06 | -0.11 | 9.3E-01 | 0.01 |
| NEDD8 | -0.01 | 0.05 | -0.04 | 0.12 | -0.09 | -0.04 | 9.3E-01 | -0.01 |
| SPAG9 | -0.19 | 0.05 | 0.11 | -0.08 | 0.13 | -0.05 | 9.3E-01 | 0.01 |
| SIX2 | -0.09 | -0.20 | 0.24 | -0.06 | -0.20 | 0.25 | 9.3E-01 | 0.02 |
| S100A13 | -0.03 | -0.02 | 0.04 | 0.12 | -0.08 | -0.04 | 9.3E-01 | 0.01 |
| MYO7B | 0.00 | -0.22 | 0.22 | 0.02 | 0.14 | -0.21 | 9.3E-01 | -0.01 |
| EXOSC5 | -0.06 | 0.04 | -0.02 | 0.24 | -0.20 | -0.04 | 9.4E-01 | 0.01 |
| ZNF649 | -0.13 | -0.07 | 0.20 | 0.10 | -0.18 | 0.05 | 9.4E-01 | -0.01 |
| DSN1 | -0.01 | 0.02 | -0.01 | -0.09 | 0.11 | -0.03 | 9.4E-01 | -0.01 |
| CAT | 0.03 | 0.08 | -0.11 | -0.08 | 0.05 | 0.02 | 9.4E-01 | -0.01 |
| SREBF1 | 0.01 | -0.06 | 0.03 | 0.15 | -0.04 | -0.10 | 9.4E-01 | 0.01 |
| NOC2L | 0.07 | 0.00 | -0.08 | 0.00 | 0.04 | -0.04 | 9.4E-01 | 0.00 |
| HMGN3 | 0.02 | -0.01 | -0.02 | -0.07 | 0.01 | 0.06 | 9.4E-01 | 0.00 |
| SCCPDH | 0.08 | -0.03 | -0.07 | -0.10 | 0.11 | -0.01 | 9.4E-01 | 0.01 |
| ADARB1 | 0.02 | -0.06 | 0.04 | -0.09 | 0.04 | 0.04 | 9.4E-01 | 0.00 |
| GALNT9 | -0.41 | -0.09 | 0.41 | -0.08 | -0.03 | 0.08 | 9.4E-01 | 0.02 |
| PDF | -0.04 | -0.02 | 0.06 | 0.00 | 0.08 | -0.09 | 9.4E-01 | 0.00 |
| TRUB1 | 0.00 | -0.04 | 0.02 | 0.10 | 0.09 | -0.18 | 9.4E-01 | 0.01 |

| | | | | | | | | |
|---|---|---|---|---|---|---|---|---|
| RPP25L | -0.07 | -0.02 | 0.07 | 0.09 | -0.09 | 0.00 | 9.4E-01 | 0.01 |
| PTRH2 | -0.08 | -0.10 | 0.18 | 0.13 | -0.18 | 0.02 | 9.4E-01 | -0.01 |
| ARID1A | 0.00 | 0.02 | -0.03 | -0.12 | 0.01 | 0.12 | 9.4E-01 | 0.01 |
| GCA | -0.01 | -0.04 | 0.06 | -0.09 | -0.02 | 0.11 | 9.4E-01 | -0.01 |
| ZNF14 | 0.03 | 0.01 | -0.05 | 0.14 | -0.09 | -0.04 | 9.4E-01 | 0.01 |
| MED22 | 0.00 | -0.08 | 0.08 | 0.18 | -0.16 | -0.05 | 9.4E-01 | -0.01 |
| BAG3 | -0.04 | 0.04 | 0.00 | 0.00 | -0.08 | 0.07 | 9.4E-01 | 0.00 |
| KCNMA1 | -0.11 | 0.06 | 0.05 | 0.04 | -0.07 | 0.01 | 9.4E-01 | -0.01 |
| DHX57 | -0.17 | -0.06 | 0.23 | 0.00 | -0.02 | -0.01 | 9.4E-01 | -0.01 |
| LMF2 | 0.08 | -0.03 | -0.04 | -0.01 | -0.03 | 0.03 | 9.4E-01 | 0.00 |
| BPHL | 0.07 | 0.01 | -0.09 | -0.04 | 0.02 | 0.02 | 9.4E-01 | 0.00 |
| SENP5 | -0.02 | -0.08 | 0.08 | -0.10 | -0.08 | 0.17 | 9.4E-01 | 0.01 |
| METTL8 | 0.07 | -0.04 | -0.04 | 0.04 | -0.05 | 0.00 | 9.4E-01 | 0.00 |
| ABCA12 | -0.07 | 0.01 | 0.06 | -0.01 | 0.04 | -0.02 | 9.4E-01 | 0.00 |
| XBP1 | -0.23 | -0.02 | 0.24 | -0.21 | 0.11 | 0.05 | 9.4E-01 | -0.01 |
| RPS4X | -0.01 | -0.01 | 0.01 | 0.02 | 0.06 | -0.08 | 9.4E-01 | 0.00 |
| TPRN | -0.08 | 0.00 | 0.08 | -0.03 | -0.01 | 0.03 | 9.5E-01 | 0.00 |
| CLCN3 | -0.09 | -0.06 | 0.14 | -0.17 | 0.06 | 0.08 | 9.5E-01 | -0.01 |
| TMEM131L | -0.14 | -0.17 | 0.29 | -0.09 | 0.02 | 0.02 | 9.5E-01 | -0.01 |
| LRCH4 | -0.26 | 0.04 | 0.22 | 0.10 | -0.08 | -0.06 | 9.5E-01 | -0.01 |
| SLC35D1 | -0.07 | 0.02 | 0.03 | -0.22 | 0.05 | 0.16 | 9.5E-01 | 0.01 |
| COX7B | -0.09 | -0.14 | 0.18 | 0.23 | -0.28 | 0.04 | 9.5E-01 | 0.01 |
| ATL2 | -0.01 | 0.04 | -0.03 | 0.12 | -0.02 | -0.12 | 9.5E-01 | -0.01 |
| ZNF124 | -0.37 | 0.19 | 0.17 | -0.11 | 0.04 | 0.01 | 9.5E-01 | -0.01 |
| CHTF8 | 0.07 | -0.08 | 0.01 | -0.25 | 0.08 | 0.14 | 9.5E-01 | -0.01 |
| PIK3R2 | -0.01 | 0.01 | 0.00 | -0.04 | 0.03 | 0.02 | 9.5E-01 | 0.00 |
| SIRT6 | -0.10 | -0.04 | 0.14 | -0.05 | 0.00 | 0.04 | 9.5E-01 | -0.01 |
| GLTP | -0.10 | 0.16 | -0.07 | 0.26 | -0.12 | -0.19 | 9.5E-01 | -0.01 |
| PRKRA | -0.06 | -0.10 | 0.16 | -0.01 | 0.03 | -0.04 | 9.5E-01 | -0.01 |
| TMX1 | 0.13 | 0.03 | -0.16 | -0.06 | -0.02 | 0.06 | 9.5E-01 | -0.01 |
| CAMK2D | 0.07 | -0.06 | -0.01 | 0.10 | 0.03 | -0.14 | 9.5E-01 | -0.01 |
| GIPC1 | -0.07 | 0.01 | 0.06 | 0.04 | 0.00 | -0.05 | 9.5E-01 | 0.00 |
| PPM1N | -1.47 | -0.49 | 1.23 | -1.01 | 0.14 | -0.03 | 9.5E-01 | -0.06 |
| TRUB2 | 0.04 | -0.06 | 0.01 | 0.04 | -0.02 | -0.01 | 9.5E-01 | 0.00 |
| CDYL2 | 0.00 | -0.02 | 0.03 | -0.05 | 0.07 | -0.03 | 9.5E-01 | 0.00 |
| RC3H2 | 0.07 | -0.03 | -0.04 | 0.01 | -0.25 | 0.21 | 9.5E-01 | -0.01 |
| TXLNA | 0.06 | -0.01 | -0.06 | 0.07 | 0.02 | -0.08 | 9.5E-01 | 0.00 |
| CRABP2 | -0.01 | 0.13 | -0.14 | -0.07 | 0.16 | -0.09 | 9.5E-01 | 0.01 |
| PPP2CB | -0.13 | -0.10 | 0.22 | -0.05 | -0.02 | 0.05 | 9.5E-01 | -0.01 |
| DDX18 | 0.03 | -0.03 | -0.01 | 0.03 | -0.01 | -0.02 | 9.5E-01 | 0.00 |
| MUC15 | -0.06 | -0.03 | 0.09 | -0.01 | -0.13 | 0.13 | 9.5E-01 | -0.01 |
| EZH1 | -0.03 | -0.07 | 0.08 | -0.12 | 0.01 | 0.11 | 9.5E-01 | 0.00 |
| INPP5J | 0.23 | -0.01 | -0.27 | -0.34 | 0.36 | -0.11 | 9.6E-01 | -0.01 |
| NOC3L | 0.07 | 0.01 | -0.10 | 0.12 | 0.02 | -0.14 | 9.6E-01 | 0.01 |
| MCM3AP | 0.00 | -0.03 | 0.02 | -0.05 | -0.01 | 0.06 | 9.6E-01 | 0.00 |
| PLEKHA5 | -0.16 | 0.05 | 0.11 | -0.15 | -0.07 | 0.20 | 9.6E-01 | -0.01 |
| CELF1 | 0.00 | -0.07 | 0.07 | -0.13 | 0.02 | 0.10 | 9.6E-01 | 0.00 |
| CDCP1 | -0.12 | 0.03 | 0.09 | -0.03 | -0.03 | 0.05 | 9.6E-01 | 0.00 |

| | | | | | | | | |
|---|---|---|---|---|---|---|---|---|
| PSMD10 | -0.28 | -0.37 | 0.57 | -0.22 | -0.03 | 0.11 | 9.6E-01 | -0.02 |
| INTS5 | 0.02 | -0.05 | 0.04 | 0.05 | -0.06 | 0.00 | 9.6E-01 | 0.00 |
| STX3 | -0.07 | -0.05 | 0.12 | -0.07 | -0.05 | 0.10 | 9.6E-01 | 0.00 |
| POLR1D | 0.06 | -0.04 | -0.02 | -0.03 | 0.09 | -0.07 | 9.6E-01 | 0.00 |
| KDM1B | 0.01 | 0.04 | -0.04 | -0.05 | 0.01 | 0.04 | 9.6E-01 | 0.00 |
| HID1 | 0.00 | -0.02 | 0.01 | 0.11 | -0.08 | -0.03 | 9.6E-01 | 0.00 |
| RNF214 | -0.02 | -0.10 | 0.10 | 0.16 | -0.03 | -0.13 | 9.6E-01 | 0.01 |
| SOGA1 | -0.10 | 0.01 | 0.08 | 0.08 | -0.05 | -0.03 | 9.6E-01 | 0.00 |
| TMBIM1 | -0.13 | 0.05 | 0.08 | 0.03 | -0.12 | 0.07 | 9.6E-01 | 0.00 |
| ANO1 | -0.18 | -0.08 | 0.25 | -0.06 | -0.07 | 0.09 | 9.6E-01 | -0.01 |
| MAN2A2 | -0.11 | 0.03 | 0.08 | 0.04 | -0.05 | 0.01 | 9.6E-01 | 0.00 |
| P4HB | 0.05 | 0.00 | -0.05 | 0.03 | -0.03 | 0.00 | 9.6E-01 | 0.00 |
| NDUFB3 | 0.03 | 0.03 | -0.07 | 0.13 | -0.08 | -0.06 | 9.6E-01 | 0.00 |
| PDS5A | 0.03 | 0.00 | -0.03 | -0.01 | 0.01 | 0.00 | 9.6E-01 | 0.00 |
| UTP15 | 0.02 | -0.04 | 0.01 | 0.07 | -0.03 | -0.04 | 9.6E-01 | 0.00 |
| ZBTB48 | 0.01 | 0.05 | -0.06 | -0.11 | 0.09 | 0.01 | 9.6E-01 | 0.00 |
| DRG1 | -0.03 | -0.05 | 0.07 | 0.18 | -0.04 | -0.16 | 9.6E-01 | -0.01 |
| HHAT | 0.16 | -0.01 | -0.17 | 0.24 | -0.07 | -0.21 | 9.6E-01 | -0.01 |
| PTCD2 | 0.00 | -0.09 | 0.08 | -0.26 | 0.09 | 0.14 | 9.7E-01 | -0.01 |
| SERINC1 | -0.10 | 0.02 | 0.09 | -0.04 | 0.00 | 0.03 | 9.7E-01 | 0.00 |
| USF1 | -0.02 | -0.01 | 0.02 | 0.04 | -0.10 | 0.06 | 9.7E-01 | 0.00 |
| BUD31 | -0.02 | -0.06 | 0.07 | 0.12 | -0.17 | 0.05 | 9.7E-01 | 0.00 |
| BCAT2 | 0.13 | 0.01 | -0.14 | -0.01 | 0.06 | -0.07 | 9.7E-01 | 0.00 |
| RAB27A | -0.10 | -0.02 | 0.12 | 0.13 | -0.08 | -0.06 | 9.7E-01 | 0.00 |
| PPP2R1A | 0.02 | -0.02 | -0.01 | -0.01 | -0.01 | 0.02 | 9.7E-01 | 0.00 |
| DYNC1I2 | 0.03 | 0.05 | -0.08 | 0.01 | -0.01 | 0.00 | 9.7E-01 | 0.00 |
| TRIM13 | -0.05 | -0.02 | 0.06 | 0.05 | -0.07 | 0.02 | 9.7E-01 | 0.00 |
| DIP2A | -0.02 | 0.00 | 0.02 | -0.06 | 0.09 | -0.03 | 9.7E-01 | 0.00 |
| CPD | -0.05 | -0.04 | 0.09 | 0.08 | -0.02 | -0.05 | 9.7E-01 | 0.00 |
| PHF10 | 0.15 | 0.00 | -0.17 | 0.03 | 0.06 | -0.09 | 9.7E-01 | 0.00 |
| ZNF184 | 0.09 | -0.14 | 0.04 | 0.10 | -0.17 | 0.05 | 9.7E-01 | 0.00 |
| RXRB | 0.04 | -0.04 | 0.00 | 0.02 | -0.03 | 0.02 | 9.7E-01 | 0.00 |
| VPS4A | 0.00 | -0.01 | 0.01 | 0.06 | 0.02 | -0.09 | 9.7E-01 | 0.00 |
| OSBPL5 | -0.03 | -0.01 | 0.05 | 0.05 | -0.07 | 0.01 | 9.7E-01 | 0.00 |
| TAX1BP3 | 0.01 | 0.05 | -0.07 | 0.03 | -0.09 | 0.06 | 9.7E-01 | 0.00 |
| PEX12 | -0.26 | -0.13 | 0.36 | 0.10 | -0.14 | -0.02 | 9.7E-01 | -0.01 |
| CEP97 | 0.29 | -0.08 | -0.29 | -0.27 | -0.16 | 0.37 | 9.7E-01 | 0.01 |
| ACTR6 | 0.00 | -0.16 | 0.15 | -0.11 | -0.13 | 0.21 | 9.7E-01 | -0.01 |
| PATL1 | -0.05 | 0.02 | 0.03 | -0.09 | 0.03 | 0.05 | 9.7E-01 | 0.00 |
| NEMP1 | -0.03 | -0.09 | 0.12 | -0.02 | -0.05 | 0.06 | 9.7E-01 | 0.00 |
| PTPN6 | -0.01 | 0.09 | -0.08 | 0.03 | -0.02 | -0.01 | 9.7E-01 | 0.00 |
| ZFP91 | -0.01 | 0.00 | -0.01 | -0.28 | 0.11 | 0.14 | 9.7E-01 | 0.00 |
| CCDC9 | -0.03 | -0.01 | 0.04 | -0.01 | 0.04 | -0.04 | 9.8E-01 | 0.00 |
| PPP1R16A | -0.18 | -0.10 | 0.26 | -0.09 | 0.02 | 0.05 | 9.8E-01 | 0.00 |
| CDSN | 0.15 | -0.02 | -0.13 | 0.09 | -0.04 | -0.07 | 9.8E-01 | 0.00 |
| ZFC3H1 | -0.10 | -0.08 | 0.17 | -0.01 | -0.08 | 0.08 | 9.8E-01 | 0.00 |
| MME | -0.05 | -0.03 | 0.08 | -0.03 | -0.06 | 0.08 | 9.8E-01 | 0.00 |
| TMEM9 | -0.19 | -0.12 | 0.26 | 0.06 | -0.23 | 0.15 | 9.8E-01 | 0.01 |

| | | | | | | | | |
|---|---|---|---|---|---|---|---|---|
| ACTN4 | -0.07 | -0.02 | 0.10 | 0.01 | -0.01 | -0.01 | 9.8E-01 | 0.00 |
| TRAF4 | 0.01 | -0.01 | 0.00 | 0.09 | 0.00 | -0.09 | 9.8E-01 | 0.00 |
| SNX9 | -0.05 | -0.03 | 0.07 | 0.08 | 0.00 | -0.08 | 9.8E-01 | 0.00 |
| RPLP2 | -0.03 | -0.02 | 0.04 | 0.13 | -0.01 | -0.12 | 9.8E-01 | 0.00 |
| MYO1F | -0.10 | -0.17 | 0.23 | 0.11 | -0.21 | 0.08 | 9.8E-01 | 0.00 |
| VPS26B | -0.10 | 0.02 | 0.07 | 0.07 | -0.02 | -0.05 | 9.8E-01 | 0.00 |
| PRR14L | -0.12 | 0.04 | 0.08 | 0.01 | 0.09 | -0.10 | 9.8E-01 | 0.00 |
| MARK3 | -0.05 | 0.00 | 0.05 | 0.02 | 0.01 | -0.03 | 9.8E-01 | 0.00 |
| INCENP | 0.00 | -0.01 | 0.01 | -0.13 | 0.06 | 0.07 | 9.8E-01 | 0.00 |
| OSBPL10 | -0.11 | 0.04 | 0.06 | -0.08 | -0.01 | 0.08 | 9.8E-01 | 0.00 |
| RELCH | -0.07 | 0.13 | -0.08 | -0.09 | -0.12 | 0.20 | 9.8E-01 | 0.00 |
| MED30 | 0.27 | -0.14 | -0.24 | 0.47 | -0.22 | -0.33 | 9.8E-01 | 0.01 |
| LPGAT1 | 0.07 | 0.02 | -0.09 | 0.05 | 0.09 | -0.16 | 9.8E-01 | 0.00 |
| LDB1 | -0.21 | -0.12 | 0.30 | -0.13 | -0.06 | 0.16 | 9.8E-01 | 0.00 |
| SMIM1 | -0.15 | 0.01 | 0.13 | 0.14 | -0.16 | 0.00 | 9.8E-01 | 0.00 |
| ACTN1 | -0.10 | -0.03 | 0.12 | -0.03 | -0.06 | 0.09 | 9.8E-01 | 0.00 |
| AHDC1 | 0.05 | -0.04 | -0.01 | -0.08 | -0.06 | 0.14 | 9.8E-01 | 0.00 |
| PHF5A | 0.10 | -0.01 | -0.09 | 0.08 | -0.06 | -0.03 | 9.8E-01 | 0.00 |
| NCEH1 | 0.09 | 0.06 | -0.16 | -0.03 | -0.09 | 0.11 | 9.9E-01 | 0.00 |
| RALB | -0.03 | 0.03 | 0.00 | 0.00 | -0.12 | 0.12 | 9.9E-01 | 0.00 |
| CCSAP | -0.04 | 0.09 | -0.07 | -0.13 | 0.22 | -0.10 | 9.9E-01 | 0.00 |
| DEPDC1 | 0.08 | 0.07 | -0.17 | -0.27 | 0.14 | 0.10 | 9.9E-01 | 0.00 |
| HACD3 | -0.05 | -0.08 | 0.12 | 0.03 | -0.03 | -0.01 | 9.9E-01 | 0.00 |
| HS2ST1 | -0.02 | 0.03 | -0.01 | 0.15 | -0.04 | -0.12 | 9.9E-01 | 0.00 |
| ABHD10 | 0.08 | 0.07 | -0.16 | -0.02 | -0.11 | 0.11 | 9.9E-01 | 0.00 |
| CCDC106 | 0.07 | 0.04 | -0.11 | -0.04 | 0.02 | 0.01 | 9.9E-01 | 0.00 |
| GATD3B | 0.09 | -0.02 | -0.08 | -0.07 | 0.17 | -0.11 | 9.9E-01 | 0.00 |
| DGCR8 | 0.07 | 0.00 | -0.08 | 0.14 | -0.10 | -0.05 | 9.9E-01 | 0.00 |
| ACAD11 | -0.16 | 0.00 | 0.14 | -0.14 | 0.08 | 0.05 | 9.9E-01 | 0.00 |
| SCFD1 | -0.02 | -0.01 | 0.01 | 0.19 | -0.09 | -0.11 | 9.9E-01 | 0.00 |
| RAB7B | -0.07 | -0.11 | 0.15 | -0.17 | -0.09 | 0.24 | 9.9E-01 | 0.00 |
| ENAH | -0.08 | -0.01 | 0.08 | 0.00 | -0.09 | 0.08 | 9.9E-01 | 0.00 |
| ABI1 | 0.14 | 0.07 | -0.22 | 0.15 | -0.04 | -0.14 | 9.9E-01 | 0.00 |
| MCL1 | -0.10 | -0.04 | 0.13 | -0.09 | 0.05 | 0.03 | 9.9E-01 | 0.00 |
| AGTRAP | -0.14 | -0.01 | 0.14 | -0.04 | 0.03 | 0.01 | 9.9E-01 | 0.00 |
| RMDN1 | 0.22 | -0.09 | -0.16 | 0.06 | -0.05 | -0.03 | 9.9E-01 | 0.00 |
| ZDHHC18 | -0.23 | 0.17 | 0.04 | -0.12 | -0.02 | 0.13 | 9.9E-01 | 0.00 |
| GRB7 | -0.04 | 0.12 | -0.09 | 0.10 | -0.02 | -0.08 | 9.9E-01 | 0.00 |
| SLU7 | 0.09 | -0.04 | -0.06 | 0.04 | -0.04 | -0.01 | 9.9E-01 | 0.00 |
| LAMTOR5 | -0.74 | -0.36 | 0.84 | -0.06 | -0.21 | 0.00 | 9.9E-01 | 0.00 |
| SLC39A1 | -0.30 | 0.11 | 0.14 | -0.06 | -0.26 | 0.28 | 9.9E-01 | 0.00 |
| MLPH | -0.13 | 0.01 | 0.12 | -0.07 | 0.02 | 0.05 | 9.9E-01 | 0.00 |
| TOMM5 | 0.01 | -0.02 | 0.00 | 0.01 | 0.03 | -0.04 | 9.9E-01 | 0.00 |
| TMEM185A | -0.11 | -0.05 | 0.16 | -0.07 | -0.07 | 0.14 | 1.0E+00 | 0.00 |
| MGAT5 | 0.01 | -0.24 | 0.19 | 0.01 | -0.31 | 0.26 | 1.0E+00 | 0.00 |
| UBN1 | 0.02 | 0.00 | -0.02 | -0.05 | 0.01 | 0.04 | 1.0E+00 | 0.00 |
| MSMO1 | 0.15 | 0.14 | -0.31 | 0.11 | -0.04 | -0.09 | 1.0E+00 | 0.00 |
| DMAP1 | 0.04 | 0.00 | -0.05 | -0.07 | 0.04 | 0.03 | 1.0E+00 | 0.00 |

| | | | | | | | | |
|---|---|---|---|---|---|---|---|---|
| **PTGFRN** | 0.00 | 0.10 | -0.11 | 0.05 | -0.13 | 0.07 | 1.0E+00 | 0.00 |
| **LARP4B** | -0.06 | 0.00 | 0.05 | 0.03 | 0.00 | -0.04 | 1.0E+00 | 0.00 |
| **NR3C1** | -0.04 | -0.06 | 0.07 | -0.28 | 0.09 | 0.17 | 1.0E+00 | 0.00 |
| **PCBP1** | 0.01 | -0.01 | 0.00 | 0.01 | -0.07 | 0.05 | 1.0E+00 | 0.00 |
| **CSNK2A2** | 0.02 | 0.15 | -0.19 | 0.13 | -0.12 | -0.02 | 1.0E+00 | 0.00 |
| **NFIB** | 0.05 | 0.02 | -0.08 | 0.02 | -0.04 | 0.02 | 1.0E+00 | 0.00 |
| **MRPS22** | 0.09 | 0.00 | -0.09 | -0.05 | -0.07 | 0.11 | 1.0E+00 | 0.00 |
