## Supplementary Table 3 for "Chromatin signalling pathways and FANCE amplification affect ATR inhibitor sensitivity in metastatic breast cancer"

| Master Protein Gene Name | Modifications in Master Proteins | Untreated_R1 | Untreated_R2 | Untreated_R3 | AZD6738_R1 | AZD6738_R2 | AZD6738_R3 | p-value ATRi/UT |
|---|---|---|---|---|---|---|---|---|
| **AAK1** | **S678** | -0.25 | -0.40 | 0.42 | 0.07 | 0.05 | -0.02 | 0.685 |
| **AAK1** | **T620; S624** | -0.03 | -0.01 | -0.14 | -0.04 | 0.27 | -0.09 | 0.428 |
| **AAK1** | **T** | -0.03 | 0.13 | -0.05 | -0.03 | 0.02 | -0.05 | 0.596 |
| **AATF** | **S203** | 0.07 | 0.02 | -0.03 | 0.05 | -0.01 | -0.10 | 0.504 |
| **AATF** | **S203** | 0.02 | 0.04 | 0.03 | -0.01 | 0.01 | -0.09 | 0.108 |
| **AATF** | **S/Y/T** | 0.34 | 0.48 | 0.17 | -0.21 | -0.19 | -1.05 | 0.052 |
| **AATF** | **S/Y/T** | 0.44 | 0.53 | 0.33 | -0.21 | -0.32 | -1.81 | 0.08 |
| **ABCA12** | **S/T/Y** | -0.68 | -0.04 | 0.23 | 0.11 | 0.00 | 0.20 | 0.384 |
| **ABCA12** | **S2595** | -0.66 | 0.21 | 0.66 | -0.11 | -0.28 | -0.17 | 0.551 |
| **ABCA2** | **S1327** | -0.28 | -0.12 | -0.19 | 0.11 | -0.11 | 0.46 | 0.111 |
| **ABCA2** | **T2412** | -0.07 | -0.09 | -0.21 | 0.09 | -0.24 | 0.42 | 0.334 |
| **ABCA2** | **S** | -0.35 | 0.31 | 0.05 | -0.21 | 0.15 | -0.04 | 0.879 |
| **ABCA2** | **T2412; S2419** | 0.12 | 0.21 | -0.16 | -0.03 | -0.17 | -0.01 | 0.368 |
| **ABCB6** | **S61** | -0.08 | 0.00 | 0.30 | -0.65 | 0.21 | 0.04 | 0.512 |
| **ABCC1** | **S915; S918** | -0.15 | 0.09 | -0.03 | -0.03 | -0.06 | 0.15 | 0.607 |
| **ABCC1** | **S/Y** | -0.23 | 0.11 | 0.10 | 0.04 | 0.19 | -0.27 | 0.983 |
| **ABCC1** | **S918; S919; S/Y** | -0.07 | -0.16 | 0.25 | -0.02 | 0.00 | -0.03 | 0.862 |
| **ABCC1** | **S930** | -0.04 | 0.14 | 0.11 | -0.04 | -0.14 | -0.05 | 0.084 |
| **ABCC3** | **S** | -0.02 | 0.12 | 0.27 | -0.24 | -0.16 | -0.03 | 0.064 |
| **ABCC5** | **S803** | 0.22 | 0.57 | 0.16 | -0.86 | -0.19 | -0.33 | 0.032 |
| **ABCD1** | **S691** | -0.11 | 0.35 | 0.25 | -0.22 | 0.15 | -0.62 | 0.208 |
| **ABCD1** | **S58** | 0.42 | 0.24 | 0.16 | -0.70 | -0.11 | -0.29 | 0.028 |
| **ABCD3** | **S40** | 0.27 | 0.17 | 0.19 | -0.23 | -0.13 | -0.39 | 0.005 |
| **ABCF1** | **S166** | 0.24 | -0.24 | -0.14 | -0.01 | 0.06 | 0.04 | 0.64 |
| **ABCF1** | **S109** | 0.02 | 0.16 | 0.05 | -0.14 | 0.21 | -0.37 | 0.382 |
| **ABCF1** | **T108** | 0.18 | 0.22 | 0.20 | -0.29 | 0.00 | -0.44 | 0.027 |
| **ABCF3** | **S283** | -0.18 | -0.25 | 0.04 | 0.12 | 0.03 | 0.20 | 0.071 |
| **ABI1** | **S222** | 0.12 | 0.06 | 0.05 | -0.12 | -0.18 | 0.04 | 0.071 |
| **ABI1** | **S231** | 0.07 | 0.32 | 0.33 | -0.18 | -0.07 | -0.70 | 0.058 |
| **ABI1** | **S183** | 0.12 | 0.33 | 0.41 | -0.43 | 0.28 | -1.43 | 0.182 |
| **ABI1** | **T/S** | 0.23 | 0.65 | 0.33 | -0.23 | -0.62 | -1.01 | 0.017 |
| **ABI2** | **T361** | -0.24 | 0.45 | 0.21 | 0.01 | -0.37 | -0.22 | 0.223 |
| **ABI2** | **S227** | -0.02 | 0.46 | 0.56 | -0.28 | -0.19 | -1.09 | 0.064 |
| **ABL1** | **S917** | -0.07 | 0.05 | -0.08 | 0.03 | -0.01 | 0.07 | 0.243 |
| **ABL1** | **S569** | 0.29 | -0.40 | 0.19 | -0.45 | 0.17 | 0.02 | 0.718 |
| **ABL1** | **T852** | 0.16 | 0.30 | -0.04 | -0.39 | -0.19 | 0.06 | 0.127 |
| **ABL2** | **S620** | -0.29 | -0.33 | -0.20 | 0.09 | 0.09 | 0.48 | 0.022 |
| **ABL2** | **S631** | -0.32 | 0.22 | -0.25 | -0.10 | 0.11 | 0.25 | 0.365 |

| | | | | | | | | |
|---|---|---|---|---|---|---|---|---|
| ABLIM1 | S458 | -0.06 | -0.06 | -0.18 | -0.01 | 0.21 | 0.08 | 0.063 |
| ABLIM1 | S458; S/T/Y | 0.02 | 0.16 | -0.13 | -0.23 | 0.10 | 0.04 | 0.737 |
| ABLIM1 | S431 | 0.01 | 0.05 | 0.05 | -0.09 | 0.02 | -0.05 | 0.097 |
| ABLIM1 | S677 | 0.36 | -0.14 | -0.12 | -0.18 | -0.20 | 0.19 | 0.668 |
| ABLIM1 | S431; S435 | 0.06 | 0.06 | 0.05 | -0.14 | -0.05 | 0.00 | 0.05 |
| ABLIM1 | S655 | 0.02 | 0.20 | 0.22 | -0.04 | -0.11 | -0.37 | 0.053 |
| ABLIM1 | S435 | -0.03 | 0.31 | 0.16 | -0.38 | 0.11 | -0.29 | 0.136 |
| ABLIM3 | S567 | -0.06 | -0.21 | -0.36 | 0.09 | 0.10 | 0.33 | 0.031 |
| ABLIM3 | S373 | -0.14 | -0.07 | -0.30 | 0.08 | -0.17 | 0.47 | 0.205 |
| ABLIM3 | T279; T/S | -0.19 | 0.11 | 0.02 | 0.20 | -0.28 | 0.08 | 0.902 |
| ABLIM3 | S503 | 0.17 | -0.13 | -0.06 | 0.02 | -0.02 | 0.01 | 0.924 |
| ABLIM3 | S388 | -0.07 | 0.03 | 0.14 | -0.04 | 0.14 | -0.24 | 0.555 |
| ABLIM3 | S/T | -0.42 | 0.16 | 0.48 | -0.47 | -0.14 | 0.15 | 0.515 |
| ABLIM3 | S282 | 0.03 | 0.29 | 0.11 | -0.23 | 0.02 | -0.32 | 0.066 |
| ABLIM3 | T543 | -0.08 | 0.17 | 0.32 | 0.26 | -0.10 | -0.87 | 0.35 |
| ABLIM3 | S567 | 0.37 | 0.46 | -0.22 | -0.19 | -0.31 | -0.34 | 0.093 |
| ABT1 | S32 | -0.30 | -0.19 | -0.45 | 0.11 | 0.19 | 0.44 | 0.011 |
| ABT1 | S32 | -0.25 | 0.02 | -0.54 | 0.14 | -0.03 | 0.47 | 0.107 |
| ACACA | S25 | -0.01 | -0.19 | 0.07 | 0.06 | 0.09 | -0.03 | 0.373 |
| ACACA | S835 | 0.05 | -0.41 | 0.23 | -0.04 | 0.04 | 0.05 | 0.779 |
| ACAP2 | S521 | 0.20 | -0.30 | -0.35 | 0.42 | -0.41 | 0.21 | 0.506 |
| ACAP2 | S379 | 0.07 | 0.35 | 0.14 | -0.35 | 0.26 | -0.74 | 0.202 |
| ACAP3 | S813 | -0.35 | -0.12 | 0.13 | 0.40 | -0.19 | -0.01 | 0.463 |
| ACBD4 | S166 | -0.02 | -0.02 | 0.15 | -0.12 | 0.02 | -0.02 | 0.324 |
| ACBD5 | T421 | -0.22 | -0.05 | 0.25 | -0.05 | 0.09 | -0.07 | 0.98 |
| ACBD5 | S215 | 0.06 | 0.02 | -0.08 | 0.07 | 0.01 | -0.09 | 0.913 |
| ACBD5 | S418 | -0.05 | 0.17 | 0.06 | 0.11 | 0.05 | -0.41 | 0.455 |
| ACBD5 | S262 | 0.28 | 0.15 | -0.06 | -0.29 | 0.08 | -0.26 | 0.14 |
| ACBD5 | S262 | 0.11 | 0.27 | 0.00 | 0.06 | -0.15 | -0.38 | 0.131 |
| ACD | S313 | -1.02 | -0.94 | -0.87 | 0.03 | 0.73 | 0.82 | 0.004 |
| ACD | S313 | -0.38 | -0.19 | -0.27 | 0.32 | 0.12 | 0.25 | 0.003 |
| ACD | S/T | -0.26 | -0.16 | -0.30 | 0.29 | 0.06 | 0.26 | 0.007 |
| ACD | S339; S349 | 0.00 | -0.13 | -0.20 | 0.02 | 0.07 | 0.21 | 0.065 |
| ACD | S339 | -0.14 | 0.23 | -0.03 | -0.16 | -0.34 | 0.32 | 0.741 |
| ACIN1 | T765 | -1.08 | -1.26 | -0.62 | 0.63 | 0.56 | 0.52 | 0.001 |
| ACIN1 | S328 | -0.51 | -0.51 | -0.22 | 0.23 | 0.35 | 0.37 | 0.002 |
| ACIN1 | S410; T/S | -0.44 | -0.17 | -0.31 | 0.11 | 0.02 | 0.56 | 0.043 |
| ACIN1 | S710 | -0.15 | 0.00 | -0.34 | 0.35 | 0.06 | -0.01 | 0.111 |
| ACIN1 | T976 | -0.38 | 0.16 | -0.21 | -0.08 | 0.09 | 0.31 | 0.264 |
| ACIN1 | S410; T414 | -0.14 | -0.09 | -0.12 | 0.17 | 0.01 | 0.14 | 0.011 |
| ACIN1 | T/S | -0.51 | 0.01 | 0.23 | -0.19 | 0.16 | 0.16 | 0.621 |
| ACIN1 | S710 | -0.02 | 0.07 | -0.24 | 0.18 | 0.16 | -0.22 | 0.549 |
| ACIN1 | S/T/Y | -0.06 | 0.00 | 0.00 | 0.19 | 0.05 | -0.20 | 0.794 |
| ACIN1 | S424; S425; S453 | 0.08 | -0.04 | 0.03 | -0.03 | -0.14 | 0.07 | 0.462 |
| ACIN1 | S424; S425; S453 | 0.26 | 0.22 | -0.49 | 0.04 | 0.32 | -0.64 | 0.814 |
| ACIN1 | S424; S425; S453 | 0.26 | 0.22 | -0.49 | 0.04 | 0.32 | -0.64 | 0.814 |
| ACIN1 | S838 | 0.04 | 0.02 | 0.13 | 0.06 | -0.02 | -0.27 | 0.237 |
| ACIN1 | S424; S425; S453 | 0.06 | 0.09 | 0.12 | -0.21 | -0.24 | 0.14 | 0.198 |

| | | | | | | | | |
|---|---|---|---|---|---|---|---|---|
| ACIN1 | S208; S216 | 0.10 | 0.07 | 0.12 | -0.18 | 0.12 | -0.28 | 0.157 |
| ACIN1 | T414 | 0.04 | 0.15 | 0.13 | -0.08 | -0.09 | -0.18 | 0.009 |
| ACIN1 | S522 | 0.18 | 0.19 | -0.02 | -0.19 | -0.07 | -0.14 | 0.03 |
| ACIN1 | S/T | 0.21 | -0.29 | 0.33 | -0.71 | -0.04 | 0.24 | 0.494 |
| ACIN1 | S838 | 0.07 | 0.15 | 0.16 | 0.08 | -0.14 | -0.40 | 0.119 |
| ACIN1 | T682 | 0.15 | 0.14 | 0.11 | -0.02 | -0.07 | -0.37 | 0.06 |
| ACIN1 | S424; S425 | 0.33 | 0.04 | 0.02 | -0.01 | -0.40 | -0.08 | 0.131 |
| ACIN1 | S990 | 0.21 | 0.06 | 0.19 | -0.08 | -0.08 | -0.37 | 0.037 |
| ACIN1 | S700 | 0.08 | 0.07 | 0.27 | -0.25 | 0.19 | -0.51 | 0.203 |
| ACIN1 | S490 | 0.19 | 0.36 | -0.18 | 0.19 | -0.08 | -0.73 | 0.35 |
| ACIN1 | S/T | 0.13 | 0.16 | 0.20 | -0.17 | -0.17 | -0.20 | 1E-04 |
| ACIN1 | S208 | 0.18 | 0.27 | 0.05 | -0.26 | -0.07 | -0.25 | 0.015 |
| ACIN1 | S490 | 0.18 | 0.29 | 0.04 | -0.04 | -0.11 | -0.48 | 0.071 |
| ACIN1 | S240; S243 | 0.25 | 0.14 | 0.14 | -0.33 | -0.06 | -0.22 | 0.012 |
| ACIN1 | S/T | 0.10 | 0.66 | -0.33 | -0.56 | 0.16 | -0.41 | 0.311 |
| ACIN1 | S657 | 0.14 | 0.19 | 0.31 | -0.13 | -0.09 | -0.56 | 0.04 |
| ACIN1 | S863 | 0.28 | 0.29 | 0.10 | -0.24 | -0.25 | -0.32 | 0.002 |
| ACIN1 | S714 | 0.33 | 0.15 | 0.24 | -0.35 | -0.18 | -0.36 | 0.002 |
| ACIN1 | S655; S657 | 0.30 | 0.38 | 0.05 | -0.11 | -0.29 | -0.54 | 0.024 |
| ACIN1 | S216 | 0.28 | 0.33 | 0.25 | -0.31 | -0.09 | -0.72 | 0.024 |
| ACIN1 | S1004 | 0.22 | 0.30 | 0.34 | -0.35 | 0.02 | -0.88 | 0.06 |
| ACIN1 | S1004 | 0.28 | 0.26 | 0.38 | -0.29 | -0.20 | -0.74 | 0.014 |
| ACIN1 | S838 | 0.58 | 0.10 | 0.32 | -0.36 | -0.27 | -0.79 | 0.019 |
| ACLY | S481 | 0.18 | 0.33 | -0.10 | -0.13 | -0.09 | -0.27 | 0.091 |
| ACLY | S455 | 0.39 | 0.41 | 0.12 | -0.48 | -0.14 | -0.63 | 0.014 |
| ACSL1 | S620 | -0.35 | -0.18 | 0.01 | 0.11 | 0.10 | 0.23 | 0.046 |
| ACSS2 | S30 | -0.07 | -0.14 | -0.22 | 0.16 | 0.13 | 0.10 | 0.005 |
| ACSS2 | S267 | -0.26 | 0.03 | 0.01 | -0.15 | -0.04 | 0.34 | 0.529 |
| ACSS2 | S/T | 0.11 | -0.06 | -0.07 | 0.09 | -0.05 | -0.03 | 0.892 |
| ACTB | S52 | 0.07 | 0.21 | 0.16 | -0.17 | -0.04 | -0.28 | 0.018 |
| ACTB | S199 | 0.19 | 0.31 | 0.32 | -0.19 | -0.28 | -0.58 | 0.007 |
| ACTL6A | S233 | 0.15 | 0.14 | -0.06 | -0.22 | 0.14 | -0.20 | 0.274 |
| ACTN4 | S159 | -0.45 | 0.33 | -0.26 | 0.20 | 0.02 | 0.03 | 0.438 |
| ACTR3B | T119 | -0.20 | -0.39 | -0.26 | -0.21 | -0.05 | 0.77 | 0.215 |
| ACTR8 | S132 | -0.08 | -0.09 | -0.18 | 0.11 | 0.02 | 0.18 | 0.017 |
| ACTR8 | S132 | 0.09 | 0.08 | 0.09 | -0.11 | -0.10 | -0.07 | 2E-04 |
| ADAM17 | S791 | -0.17 | -0.11 | -0.13 | 0.20 | -0.01 | 0.18 | 0.02 |
| ADAM22 | S857 | 0.17 | 0.38 | 0.43 | -0.90 | -0.30 | -0.18 | 0.029 |
| ADAR | S/T | -0.27 | 0.46 | -0.61 | 0.48 | -0.20 | -0.20 | 0.686 |
| ADAR | S629 | -0.08 | -0.07 | 0.17 | -0.14 | 0.20 | -0.11 | 0.875 |
| ADAR | T601 | 0.10 | 0.15 | 0.25 | 0.17 | 0.07 | -1.14 | 0.337 |
| ADARB1 | S26 | 0.14 | 0.12 | 0.12 | -0.27 | 0.17 | -0.37 | 0.159 |
| ADCY6 | S54 | -0.04 | 0.05 | 0.07 | -0.11 | -0.12 | 0.13 | 0.53 |
| ADNP | S409 | -0.17 | -0.29 | -0.21 | 0.13 | 0.13 | 0.32 | 0.005 |
| ADNP | S709 | -0.13 | 0.21 | -0.35 | -0.05 | -0.02 | 0.25 | 0.484 |
| ADNP | S921 | 0.00 | -0.08 | -0.11 | 0.01 | -0.02 | 0.18 | 0.175 |
| ADNP | S889; S891; S | 0.22 | -0.05 | -0.09 | -0.01 | -0.09 | -0.01 | 0.573 |
| ADNP | S889; S891; S | 0.22 | -0.05 | -0.09 | -0.01 | -0.09 | -0.01 | 0.573 |

| | | | | | | | | |
|---|---|---|---|---|---|---|---|---|
| **ADNP** | **S876; S878** | 0.00 | 0.01 | 0.13 | -0.04 | -0.08 | -0.02 | 0.118 |
| **ADNP** | **S953** | -0.06 | 0.38 | -0.19 | -0.25 | 0.07 | -0.04 | 0.593 |
| **ADNP** | **S886; S** | -0.01 | 0.13 | 0.14 | 0.01 | -0.05 | -0.24 | 0.116 |
| **ADNP** | **S875** | 0.07 | 0.07 | 0.13 | -0.19 | -0.09 | -0.01 | 0.027 |
| **ADNP** | **S953** | 0.15 | 0.14 | -0.02 | -0.05 | -0.06 | -0.20 | 0.055 |
| **ADNP** | **S982** | 0.27 | -0.13 | 0.16 | -0.25 | -0.29 | 0.14 | 0.273 |
| **ADNP** | **S970** | -0.03 | 0.08 | 0.27 | -0.08 | -0.17 | -0.12 | 0.063 |
| **ADNP** | **S953** | 0.10 | 0.20 | 0.07 | -0.10 | -0.09 | -0.22 | 0.01 |
| **ADNP** | **S921** | 0.31 | 0.21 | -0.18 | -0.27 | 0.08 | -0.26 | 0.238 |
| **ADNP** | **S953** | -0.05 | 0.29 | 0.12 | 0.01 | -0.22 | -0.23 | 0.1 |
| **ADNP** | **S/Y** | 0.19 | 0.10 | 0.10 | -0.14 | -0.08 | -0.22 | 0.006 |
| **ADNP** | **S608** | 0.14 | 0.37 | -0.10 | 0.13 | -0.48 | -0.22 | 0.214 |
| **ADNP** | **S709** | 0.06 | 0.64 | 0.12 | -0.48 | 0.03 | -0.81 | 0.086 |
| **ADNP** | **S970** | 0.25 | 0.49 | 0.41 | -0.55 | -0.45 | -0.59 | 4E-04 |
| **ADNP** | **S970** | 0.47 | 0.39 | 0.30 | -0.60 | -0.39 | -0.62 | 5E-04 |
| **ADNP2** | **T866** | 0.11 | 0.06 | 0.13 | -0.10 | -0.22 | 0.00 | 0.037 |
| **AEBP2** | **S141** | -0.03 | -0.07 | 0.30 | -0.06 | -0.10 | -0.09 | 0.293 |
| **AEBP2** | **S18; S24** | 0.12 | 0.05 | 0.06 | 0.03 | -0.25 | -0.05 | 0.127 |
| **AEBP2** | **S206** | -0.04 | 0.18 | 0.21 | -0.13 | -0.22 | -0.05 | 0.058 |
| **AEBP2** | **S371** | 0.20 | 0.11 | 0.05 | -0.07 | -0.21 | -0.12 | 0.013 |
| **AFAP1** | **S668** | -0.26 | 0.22 | 0.44 | -0.27 | 0.07 | -0.39 | 0.258 |
| **AFDN** | **S/T** | -0.22 | -0.69 | -0.29 | 0.48 | 0.27 | 0.14 | 0.017 |
| **AFDN** | **S1182** | -0.52 | -0.01 | -0.17 | 0.57 | 0.09 | -0.21 | 0.229 |
| **AFDN** | **S1182** | -0.15 | -0.10 | 0.00 | 0.15 | 0.05 | 0.03 | 0.043 |
| **AFDN** | **S1182; S** | -0.05 | -0.04 | -0.13 | 0.13 | -0.13 | 0.18 | 0.26 |
| **AFDN** | **S1083** | -0.11 | -0.03 | -0.05 | -0.06 | 0.25 | -0.04 | 0.334 |
| **AFDN** | **S1512** | -0.07 | 0.03 | -0.12 | 0.10 | -0.08 | 0.12 | 0.269 |
| **AFDN** | **S1779** | -0.09 | 0.15 | 0.00 | 0.03 | 0.10 | -0.23 | 0.682 |
| **AFF1** | **S588** | -0.06 | -0.17 | -0.48 | 0.16 | -0.04 | 0.43 | 0.085 |
| **AFF1** | **S588** | 0.23 | 0.25 | 0.33 | -0.34 | -0.17 | -0.51 | 0.004 |
| **AFF4** | **S/T** | 0.05 | -0.22 | -0.38 | 0.32 | 0.06 | 0.07 | 0.091 |
| **AFF4** | **T674; S/Y** | -0.02 | -0.09 | -0.24 | 0.34 | -0.09 | 0.04 | 0.218 |
| **AFF4** | **S222** | -0.08 | 0.15 | -0.13 | 0.09 | 0.00 | -0.06 | 0.78 |
| **AFF4** | **S703** | -0.04 | -0.06 | 0.12 | 0.14 | -0.10 | -0.08 | 0.845 |
| **AFF4** | **S487** | 0.16 | 0.03 | -0.16 | -0.35 | -0.05 | 0.28 | 0.827 |
| **AFF4** | **S/Y** | -0.24 | -0.25 | 0.52 | 0.01 | -0.09 | -0.09 | 0.805 |
| **AFF4** | **S549** | 0.15 | 0.23 | -0.02 | 0.04 | 0.00 | -0.52 | 0.221 |
| **AFF4** | **S814** | 0.22 | 0.03 | 0.15 | 0.05 | -0.17 | -0.36 | 0.083 |
| **AFF4** | **S1043** | 0.28 | 0.28 | -0.10 | -0.17 | -0.48 | 0.04 | 0.144 |
| **AFF4** | **S387; S392** | 0.37 | 0.22 | -0.10 | -0.39 | -0.13 | -0.10 | 0.093 |
| **AFF4** | **S703** | 0.35 | 0.26 | -0.08 | -0.01 | -0.25 | -0.41 | 0.084 |
| **AFF4** | **S694** | 0.30 | 0.19 | -0.04 | 0.11 | 0.10 | -0.98 | 0.337 |
| **AFF4** | **S836** | 0.32 | 0.06 | 0.23 | -0.14 | -0.23 | -0.35 | 0.01 |
| **AFF4** | **T674** | 0.30 | 0.22 | 0.08 | 0.03 | -0.35 | -0.44 | 0.045 |
| **AFF4** | **S487** | 0.54 | 0.04 | 0.16 | -0.47 | -0.12 | -0.39 | 0.035 |
| **AFG2B** | **T368** | -0.18 | -0.05 | -0.17 | 0.15 | 0.45 | -0.35 | 0.411 |
| **AGAP1** | **S521** | -0.13 | -0.06 | 0.14 | -0.02 | 0.09 | -0.04 | 0.796 |
| **AGAP3** | **S538** | 0.06 | 0.16 | -0.04 | -0.32 | 0.09 | -0.01 | 0.35 |

| | | | | | | | | |
|---|---|---|---|---|---|---|---|---|
| AGAP3 | T/S | 0.20 | -0.07 | 0.41 | -0.27 | -0.03 | -0.41 | 0.082 |
| AGFG1 | S181 | 0.04 | -0.01 | -0.04 | 0.03 | -0.01 | -0.01 | 0.72 |
| AHCTF1 | S1142 | -0.06 | -0.33 | -0.88 | 0.23 | -0.19 | 0.72 | 0.132 |
| AHCTF1 | S2226 | -0.28 | -0.51 | -0.21 | 0.01 | 0.16 | 0.57 | 0.038 |
| AHCTF1 | S1513 | -0.20 | -0.07 | -0.73 | 0.48 | 0.11 | 0.14 | 0.07 |
| AHCTF1 | S1160 | -0.12 | -0.28 | -0.24 | 0.10 | 0.04 | 0.39 | 0.032 |
| AHCTF1 | S/T | -0.63 | -0.21 | 0.23 | -0.08 | 0.07 | 0.40 | 0.311 |
| AHCTF1 | S2212 | 0.03 | -0.44 | -0.10 | 0.09 | 0.14 | 0.19 | 0.101 |
| AHCTF1 | S1797 | -0.26 | -0.59 | 0.30 | 0.27 | -0.25 | 0.28 | 0.413 |
| AHCTF1 | S1513 | -0.11 | -0.06 | -0.26 | 0.05 | 0.08 | 0.24 | 0.034 |
| AHCTF1 | S1884 | -0.10 | -0.09 | -0.21 | 0.12 | 0.12 | 0.12 | 0.003 |
| AHCTF1 | T1369 | -0.31 | 0.08 | -0.12 | 0.14 | 0.14 | 0.02 | 0.155 |
| AHCTF1 | T1369 | 0.00 | -0.27 | -0.04 | -0.36 | 0.17 | 0.37 | 0.523 |
| AHCTF1 | S1884 | 0.01 | -0.08 | -0.18 | 0.12 | -0.12 | 0.20 | 0.247 |
| AHCTF1 | S1541 | 0.10 | -0.10 | -0.24 | -0.04 | 0.24 | 0.00 | 0.326 |
| AHCTF1 | S | 0.01 | -0.06 | -0.01 | 0.01 | 0.11 | -0.07 | 0.526 |
| AHCTF1 | S2065 | 0.12 | -0.12 | -0.06 | -0.05 | -0.12 | 0.20 | 0.773 |
| AHCTF1 | S1944 | 0.02 | -0.06 | -0.02 | -0.02 | -0.09 | 0.15 | 0.695 |
| AHCTF1 | S1914 | -0.19 | 0.02 | 0.13 | -0.01 | -0.07 | 0.09 | 0.898 |
| AHCTF1 | S1214 | 0.03 | -0.04 | 0.04 | -0.06 | -0.04 | 0.06 | 0.643 |
| AHCTF1 | S1232 | -0.01 | 0.02 | 0.02 | -0.11 | 0.19 | -0.14 | 0.797 |
| AHCTF1 | S1297 | 0.13 | -0.06 | 0.03 | 0.04 | -0.17 | 0.01 | 0.438 |
| AHCTF1 | S1297 | 0.03 | 0.06 | 0.04 | 0.08 | -0.11 | -0.12 | 0.214 |
| AHCTF1 | S1232 | 0.15 | 0.14 | -0.08 | -0.25 | -0.12 | 0.11 | 0.309 |
| AHCTF1 | S1297 | 0.16 | 0.07 | -0.02 | -0.09 | 0.06 | -0.22 | 0.187 |
| AHCTF1 | S1214; S1222 | 0.00 | 0.03 | 0.19 | -0.13 | -0.10 | -0.02 | 0.074 |
| AHCTF1 | S1222 | 0.05 | 0.07 | 0.24 | -0.29 | -0.09 | -0.05 | 0.052 |
| AHCTF1 | T1210; S1216; S1222 | 0.07 | 0.06 | 0.30 | -0.23 | -0.03 | -0.25 | 0.04 |
| AHCTF1 | S1283 | 0.03 | 0.11 | 0.31 | 0.06 | -0.44 | -0.18 | 0.11 |
| AHCTF1 | S2120 | 0.09 | 0.23 | 0.16 | -0.05 | -0.11 | -0.41 | 0.042 |
| AHCTF1 | S1160 | 0.34 | 0.21 | 0.20 | -0.06 | -0.08 | -0.91 | 0.101 |
| AHCTF1 | S1283 | 0.39 | 0.31 | 0.19 | -0.46 | -0.12 | -0.61 | 0.011 |
| AHCTF1 | S1283 | 0.34 | 0.31 | 0.23 | -0.53 | 0.10 | -0.84 | 0.061 |
| AHCTF1 | S1283 | 0.38 | 0.42 | 0.27 | -0.59 | -0.06 | -0.91 | 0.025 |
| AHCYL1 | S/T/Y | -0.22 | -0.24 | 0.10 | -0.38 | 0.35 | 0.23 | 0.499 |
| AHCYL1 | S/T/Y | -0.22 | -0.24 | 0.10 | -0.38 | 0.35 | 0.23 | 0.499 |
| AHCYL1 | S84; S85 | 0.19 | -0.11 | -0.09 | -0.18 | -0.06 | 0.21 | 0.965 |
| AHDC1 | S369 | -0.48 | -0.14 | -0.16 | 0.33 | 0.04 | 0.25 | 0.029 |
| AHDC1 | S1187 | -0.18 | -0.35 | -0.13 | 0.40 | -0.22 | 0.31 | 0.134 |
| AHDC1 | S896 | 0.08 | -0.15 | -0.18 | 0.18 | -0.31 | 0.28 | 0.551 |
| AHDC1 | S1064 | 0.06 | 0.22 | -0.53 | 0.20 | -0.02 | -0.06 | 0.633 |
| AHDC1 | S596 | -0.01 | -0.05 | -0.13 | 0.10 | -0.07 | 0.14 | 0.196 |
| AHDC1 | S1187 | 0.28 | -0.38 | -0.20 | 0.26 | -0.75 | 0.44 | 0.851 |
| AHDC1 | S596 | -0.06 | -0.05 | -0.01 | 0.01 | 0.03 | 0.08 | 0.037 |
| AHDC1 | S1399; S1403 | 0.04 | -0.01 | -0.16 | 0.14 | -0.08 | 0.05 | 0.428 |
| AHDC1 | S829 | 0.05 | -0.14 | 0.07 | 0.24 | 0.06 | -0.36 | 0.942 |
| AHDC1 | S1399 | 0.07 | 0.43 | -0.35 | -0.44 | -0.30 | 0.35 | 0.613 |
| AHDC1 | S1507 | 0.14 | 0.20 | -0.07 | -0.23 | 0.17 | -0.28 | 0.297 |

| | | | | | | | | |
|---|---|---|---|---|---|---|---|---|
| AHDC1 | T99 | 0.17 | 0.08 | 0.23 | -0.14 | -0.22 | -0.18 | 0.003 |
| AHNAK | S4486 | -1.03 | -1.08 | -0.65 | 0.46 | 0.69 | 0.50 | 7E-04 |
| AHNAK | S5552 | -0.57 | -0.46 | -0.45 | 0.47 | 0.72 | -0.26 | 0.053 |
| AHNAK | S4425 | -0.29 | -0.58 | -0.49 | 0.46 | 0.13 | 0.41 | 0.004 |
| AHNAK | S5739 | -0.28 | -0.41 | -0.45 | 0.47 | 0.16 | 0.25 | 0.003 |
| AHNAK | T | -0.40 | -0.16 | -0.54 | 0.39 | 0.24 | 0.22 | 0.006 |
| AHNAK | S5731; S5739 | -0.47 | -0.25 | -0.36 | 0.13 | 0.16 | 0.53 | 0.012 |
| AHNAK | S4812 | -0.34 | -0.37 | -0.32 | 0.30 | 0.17 | 0.36 | 4E-04 |
| AHNAK | S3894 | -0.05 | -0.91 | -0.19 | 0.40 | -0.30 | 0.57 | 0.184 |
| AHNAK | S5749; S5752; S5763 | -0.28 | -0.33 | -0.25 | 0.23 | 0.13 | 0.35 | 0.001 |
| AHNAK | T490 | -0.52 | -0.10 | -0.26 | 0.16 | 0.42 | 0.12 | 0.028 |
| AHNAK | S5749; S5752 | -0.23 | -0.32 | -0.28 | 0.25 | 0.14 | 0.30 | 6E-04 |
| AHNAK | T4430 | -0.30 | -0.40 | -0.14 | 0.37 | 0.04 | 0.26 | 0.015 |
| AHNAK | T4766 | -0.15 | -0.12 | -0.56 | -0.01 | 0.37 | 0.28 | 0.057 |
| AHNAK | S539 | -0.19 | -0.16 | -0.37 | -0.07 | 0.03 | 0.57 | 0.115 |
| AHNAK | S4953 | -0.28 | 0.09 | -0.61 | 0.68 | 0.17 | -0.46 | 0.362 |
| AHNAK | S5745; S5749; S5752 | -0.18 | -0.29 | -0.18 | 0.01 | 0.14 | 0.39 | 0.03 |
| AHNAK | S5448 | 0.11 | -0.37 | -0.40 | 0.08 | 0.24 | 0.21 | 0.084 |
| AHNAK | S4480 | -0.07 | -0.32 | -0.25 | 0.03 | 0.07 | 0.41 | 0.054 |
| AHNAK | S177 | -0.09 | -0.21 | -0.30 | 0.15 | 0.10 | 0.28 | 0.01 |
| AHNAK | T/S | -0.14 | -0.52 | 0.01 | 0.41 | -0.17 | 0.23 | 0.184 |
| AHNAK | T4564 | -0.19 | -0.07 | -0.25 | 0.10 | 0.24 | 0.11 | 0.01 |
| AHNAK | T4100 | -0.25 | -0.04 | -0.21 | 0.27 | 0.04 | 0.12 | 0.027 |
| AHNAK | S5841 | -0.12 | -0.07 | -0.23 | 0.17 | 0.03 | 0.17 | 0.016 |
| AHNAK | T3716 | -0.03 | -0.16 | -0.19 | 0.12 | 0.18 | 0.04 | 0.017 |
| AHNAK | S93 | -0.09 | -0.05 | -0.22 | 0.22 | -0.03 | 0.13 | 0.064 |
| AHNAK | T5794 | -0.11 | -0.14 | -0.09 | 0.15 | 0.14 | 0.02 | 0.009 |
| AHNAK | S210; S216 | -0.19 | -0.05 | -0.10 | 0.10 | -0.07 | 0.26 | 0.109 |
| AHNAK | S135 | -0.10 | -0.20 | -0.01 | 0.10 | 0.09 | 0.10 | 0.021 |
| AHNAK | S41 | -0.10 | -0.13 | -0.07 | 0.03 | 0.14 | 0.11 | 0.008 |
| AHNAK | S216 | -0.11 | -0.05 | -0.13 | 0.12 | -0.01 | 0.15 | 0.026 |
| AHNAK | S5866 | -0.16 | -0.15 | 0.00 | 0.39 | 0.05 | -0.22 | 0.396 |
| AHNAK | S1042 | -0.36 | 0.11 | 0.01 | -0.03 | 0.10 | 0.11 | 0.377 |
| AHNAK | S5752 | -0.06 | -0.02 | -0.14 | 0.11 | -0.02 | 0.11 | 0.065 |
| AHNAK | S5854 | -0.05 | -0.10 | -0.04 | -0.04 | 0.17 | 0.04 | 0.126 |
| AHNAK | S | -0.14 | 0.26 | -0.32 | 0.03 | 0.08 | 0.02 | 0.566 |
| AHNAK | S5731 | 0.01 | 0.07 | -0.19 | -0.04 | 0.09 | 0.04 | 0.49 |
| AHNAK | S4803 | 0.02 | 0.06 | -0.19 | 0.04 | -0.19 | 0.22 | 0.689 |
| AHNAK | S5782 | -0.10 | -0.07 | 0.07 | 0.17 | 0.06 | -0.16 | 0.614 |
| AHNAK | S5589 | -0.17 | 0.20 | -0.11 | 0.05 | -0.11 | 0.11 | 0.758 |
| AHNAK | S5110 | 0.02 | 0.01 | -0.08 | 0.02 | -0.10 | 0.11 | 0.724 |
| AHNAK | S93 | 0.00 | -0.04 | 0.00 | 0.10 | 0.05 | -0.12 | 0.747 |
| AHNAK | S4360 | -0.12 | 0.23 | -0.16 | 0.06 | -0.14 | 0.08 | 0.904 |
| AHNAK | S5867; S5870 | 0.00 | -0.06 | 0.05 | 0.06 | 0.04 | -0.10 | 0.934 |
| AHNAK | S | -0.18 | 0.27 | -0.13 | 0.17 | -0.27 | 0.06 | 0.999 |
| AHNAK | S613 | -0.10 | 0.20 | -0.11 | 0.22 | 0.04 | -0.32 | 0.952 |
| AHNAK | S230 | 0.06 | -0.16 | 0.08 | 0.04 | 0.29 | -0.40 | 0.944 |
| AHNAK | S93 | 0.00 | 0.06 | 0.07 | -0.04 | 0.09 | -0.20 | 0.331 |

| | | | | | | | | |
|---|---|---|---|---|---|---|---|---|
| AHNAK | S793 | 0.03 | 0.11 | -0.04 | 0.15 | 0.09 | -0.43 | 0.636 |
| AHNAK | S5870 | 0.04 | 0.02 | 0.05 | 0.11 | 0.11 | -0.41 | 0.591 |
| AHNAK | S5863; S5865 | 0.04 | 0.03 | 0.09 | -0.07 | -0.03 | -0.06 | 0.009 |
| AHNAK | S201 | -0.16 | 0.14 | 0.24 | -0.22 | 0.03 | -0.08 | 0.312 |
| AHNAK | S1170 | 0.31 | -0.09 | -0.01 | -0.31 | 0.14 | -0.13 | 0.407 |
| AHNAK | S115 | 0.10 | 0.14 | 0.05 | -0.17 | -0.19 | 0.03 | 0.045 |
| AHNAK | S5400 | 0.00 | 0.45 | -0.18 | -0.14 | -0.02 | -0.22 | 0.333 |
| AHNAK | S5866; S5867; S5870 | 0.06 | 0.06 | 0.20 | -0.03 | 0.05 | -0.42 | 0.187 |
| AHNAK | S511 | 0.14 | 0.21 | 0.20 | -0.18 | -0.03 | -0.45 | 0.031 |
| AHNAK | S93 | 0.25 | 0.12 | 0.20 | -0.21 | -0.12 | -0.34 | 0.005 |
| AHNAK | S511 | 0.22 | 0.10 | 0.28 | -0.28 | -0.07 | -0.36 | 0.012 |
| AHNAK | T/S/Y | 0.43 | -0.06 | 0.10 | -0.56 | 0.54 | -1.05 | 0.356 |
| AHNAK | S5620 | -0.04 | 0.50 | 0.16 | -0.09 | -0.01 | -0.84 | 0.167 |
| AHNAK | S5841 | 0.20 | 0.26 | 0.19 | 0.00 | 0.04 | -1.04 | 0.195 |
| AHNAK | S5110 | 0.24 | 0.42 | 0.29 | -0.32 | 0.06 | -1.28 | 0.107 |
| AIF1L | S133 | -0.06 | -0.12 | -0.03 | 0.05 | 0.03 | 0.12 | 0.02 |
| AIFM1 | S524 | -0.03 | 0.06 | -0.14 | -0.15 | 0.02 | 0.21 | 0.634 |
| AJM1 | S856 | 0.17 | -0.52 | -0.39 | 0.43 | 0.22 | -0.14 | 0.202 |
| AJM1 | S109 | 0.20 | 0.07 | 0.02 | 0.36 | -0.20 | -0.67 | 0.427 |
| AJM1 | T/S | 0.16 | 0.15 | 0.16 | -0.14 | -0.25 | -0.14 | 1E-03 |
| AJUBA | S119 | -0.04 | -0.17 | -0.20 | -0.14 | 0.19 | 0.29 | 0.142 |
| AKAP1 | T533 | -0.23 | -0.15 | -0.46 | 0.49 | -0.11 | 0.25 | 0.068 |
| AKAP1 | T533 | -0.10 | -0.26 | -0.21 | 0.17 | -0.02 | 0.32 | 0.034 |
| AKAP1 | S309 | -0.18 | -0.27 | -0.07 | 0.03 | 0.05 | 0.35 | 0.059 |
| AKAP1 | S309 | -0.42 | -0.24 | 0.09 | 0.66 | -0.13 | -0.24 | 0.416 |
| AKAP1 | S309 | -0.21 | -0.33 | 0.05 | 0.14 | -0.21 | 0.42 | 0.259 |
| AKAP1 | S592 | -0.53 | -0.21 | 0.39 | -0.07 | 0.20 | 0.05 | 0.569 |
| AKAP1 | S429 | -0.18 | 0.02 | -0.03 | 0.11 | -0.13 | 0.18 | 0.346 |
| AKAP1 | T70 | 0.02 | 0.02 | 0.03 | 0.03 | 0.10 | -0.21 | 0.628 |
| AKAP1 | T533 | -0.04 | 0.32 | -0.13 | 0.05 | 0.10 | -0.38 | 0.569 |
| AKAP1 | T533 | 0.27 | 0.18 | -0.08 | -0.13 | -0.25 | -0.05 | 0.086 |
| AKAP10 | S187 | -0.12 | -0.17 | 0.01 | 0.05 | -0.11 | 0.29 | 0.246 |
| AKAP11 | S456 | -0.07 | 0.03 | -0.25 | 0.15 | -0.26 | 0.30 | 0.438 |
| AKAP11 | S456 | -0.19 | 0.17 | -0.17 | -0.01 | -0.37 | 0.42 | 0.782 |
| AKAP11 | S | 0.07 | -0.06 | -0.06 | 0.09 | -0.09 | 0.04 | 0.689 |
| AKAP11 | S1242 | -0.04 | 0.04 | 0.06 | -0.09 | 0.15 | -0.12 | 0.696 |
| AKAP13 | S2728 | -0.35 | 0.23 | 0.43 | -0.53 | -0.27 | 0.23 | 0.42 |
| AKAP17A | S511; S515 | 0.12 | -0.19 | -0.07 | 0.01 | 0.12 | -0.01 | 0.448 |
| AKAP17A | S499 | 0.08 | -0.06 | -0.07 | 0.03 | -0.01 | 0.03 | 0.493 |
| AKAP17A | S537 | 0.08 | 0.04 | 0.06 | -0.03 | -0.11 | -0.05 | 0.009 |
| AKAP17A | S537 | 0.09 | 0.16 | 0.03 | 0.00 | -0.02 | -0.30 | 0.128 |
| AKAP17A | S633 | 0.22 | 0.16 | 0.22 | -0.03 | -0.13 | -0.60 | 0.063 |
| AKAP5 | S178 | -0.33 | -0.39 | -0.21 | 0.28 | 0.04 | 0.41 | 0.01 |
| AKAP8 | S323; S328; S339 | -0.10 | -0.16 | 0.07 | 0.06 | 0.15 | -0.04 | 0.234 |
| AKAP8 | S323; S328; S339 | 0.02 | -0.02 | 0.16 | -0.03 | 0.04 | -0.20 | 0.261 |
| AKAP8 | S323; S328; S339 | -0.01 | 0.00 | 0.19 | -0.07 | 0.00 | -0.12 | 0.16 |
| AKAP8 | S339 | 0.15 | 0.09 | 0.11 | -0.02 | -0.01 | -0.38 | 0.109 |
| AKAP8 | S323; S328 | 0.12 | 0.00 | 0.42 | -0.10 | -0.34 | -0.23 | 0.047 |

| | | | | | | | | |
|---|---|---|---|---|---|---|---|---|
| AKAP8 | S328 | 0.20 | 0.18 | 0.26 | -0.20 | -0.08 | -0.51 | 0.021 |
| AKAP8L | S283 | -0.12 | 0.06 | -0.03 | 0.00 | 0.14 | -0.06 | 0.515 |
| AKAP8L | S283 | -0.03 | -0.01 | 0.00 | 0.00 | 0.15 | -0.13 | 0.865 |
| AKAP9 | S1327 | -0.15 | -0.32 | 0.13 | 0.41 | -0.17 | -0.02 | 0.433 |
| AKAP9 | T155 | -0.12 | -0.07 | -0.08 | 0.09 | 0.21 | -0.05 | 0.081 |
| AKAP9 | S/T/Y | -0.06 | -0.02 | -0.18 | -0.01 | 0.25 | -0.02 | 0.173 |
| AKAP9 | S31 | -0.02 | 0.00 | -0.06 | -0.03 | 0.26 | -0.19 | 0.789 |
| AKAP9 | S2885 | -0.06 | 0.00 | 0.08 | -0.04 | -0.19 | 0.18 | 0.867 |
| AKAP9 | S3576 | 0.02 | 0.02 | 0.37 | -0.11 | -0.04 | -0.35 | 0.111 |
| AKAP9 | S3842 | 0.20 | 0.27 | 0.09 | -0.23 | 0.00 | -0.45 | 0.042 |
| AKR1A1 | S211 | -0.20 | -0.25 | -0.33 | 0.31 | 0.08 | 0.26 | 0.004 |
| AKT1S1 | T/S | 0.13 | -0.05 | -0.01 | 0.16 | 0.04 | -0.33 | 0.709 |
| AKT2 | T313 | -0.05 | 0.31 | 0.17 | -0.03 | -0.19 | -0.30 | 0.069 |
| ALDH18A1 | S427 | 0.02 | 0.10 | -0.19 | 0.03 | 0.10 | -0.08 | 0.701 |
| ALDH4A1 | S44 | -0.17 | -0.06 | -0.22 | 0.14 | -0.07 | 0.31 | 0.079 |
| ALDOA | S36 | 0.16 | 0.23 | 0.01 | -0.18 | -0.30 | 0.00 | 0.055 |
| ALG3 | S | 0.19 | 0.10 | 0.12 | -0.21 | 0.10 | -0.37 | 0.1 |
| ALG5 | S/T | -0.31 | -0.28 | -0.15 | 0.25 | 0.07 | 0.30 | 0.007 |
| ALKBH5 | S384 | -0.13 | -0.01 | -0.05 | -0.13 | 0.26 | 0.03 | 0.377 |
| ALKBH5 | S371; S/Y | -0.02 | -0.24 | 0.09 | 0.06 | 0.05 | 0.02 | 0.355 |
| ALKBH5 | S64; S69 | 0.10 | 0.02 | 0.03 | -0.06 | -0.01 | -0.09 | 0.049 |
| ALKBH5 | S64; S69 | 0.23 | 0.11 | -0.15 | -0.12 | -0.04 | -0.07 | 0.293 |
| ALKBH5 | S384 | 0.15 | 0.02 | 0.06 | -0.10 | -0.01 | -0.15 | 0.044 |
| ALKBH5 | S64 | -0.12 | 0.04 | 0.27 | -0.15 | 0.22 | -0.36 | 0.462 |
| ALKBH5 | S64 | 0.10 | 0.10 | 0.10 | -0.04 | -0.06 | -0.23 | 0.025 |
| ALKBH5 | S64 | 0.20 | 0.03 | 0.15 | -0.05 | 0.21 | -0.73 | 0.331 |
| ALKBH5 | S64 | 0.16 | 0.19 | 0.22 | -0.13 | -0.22 | -0.32 | 0.002 |
| ALMS1 | S2805 | 0.09 | -0.12 | -0.15 | 0.19 | 0.07 | -0.11 | 0.389 |
| ALYREF | S239 | -0.13 | -0.06 | -0.14 | 0.16 | -0.08 | 0.22 | 0.091 |
| ALYREF | S239 | -0.03 | -0.01 | -0.10 | 0.08 | 0.03 | 0.03 | 0.052 |
| ALYREF | S194 | 0.30 | 0.10 | 0.17 | -0.71 | -0.26 | 0.17 | 0.154 |
| AMFR | S509 | 0.14 | 0.06 | -0.11 | 0.15 | -0.05 | -0.24 | 0.614 |
| AMOTL1 | S241 | 0.09 | -0.30 | -0.32 | 0.07 | -0.09 | 0.41 | 0.199 |
| AMOTL1 | S241 | -0.08 | -0.09 | -0.14 | 0.12 | -0.01 | 0.17 | 0.024 |
| AMOTL1 | S720 | -0.12 | -0.14 | 0.00 | -0.06 | 0.17 | 0.12 | 0.118 |
| AMOTL1 | S809 | -0.01 | -0.09 | 0.16 | 0.02 | 0.02 | -0.12 | 0.604 |
| AMOTL1 | T/S | -0.14 | 0.15 | 0.19 | -0.37 | 0.09 | 0.01 | 0.439 |
| AMOTL1 | S906; T/S | 0.04 | -0.24 | 0.36 | -0.40 | 0.23 | -0.13 | 0.58 |
| AMOTL1 | S872 | 0.41 | 0.10 | 0.26 | -0.23 | -0.16 | -0.61 | 0.023 |
| AMOTL1 | S906 | 0.42 | 0.33 | 0.22 | -0.97 | -0.12 | -0.29 | 0.042 |
| AMOTL2 | S540 | -0.48 | -0.19 | -0.01 | -0.17 | 0.28 | 0.40 | 0.147 |
| AMOTL2 | S236 | -0.23 | -0.05 | -0.09 | -0.15 | 0.26 | 0.19 | 0.182 |
| ANAPC1 | S | -0.03 | 0.23 | 0.09 | -0.38 | 0.20 | -0.20 | 0.304 |
| ANAPC2 | S470 | 0.02 | 0.04 | 0.12 | -0.16 | 0.09 | -0.13 | 0.19 |
| ANAPC4 | S777 | 0.15 | 0.11 | -0.01 | -0.28 | -0.15 | 0.12 | 0.221 |
| ANAPC5 | S | -0.04 | 0.04 | 0.07 | -0.13 | 0.06 | -0.01 | 0.446 |
| ANK3 | S4342 | -0.14 | -0.08 | 0.01 | 0.00 | -0.04 | 0.22 | 0.224 |
| ANK3 | T4258 | -0.40 | -0.07 | 0.58 | -0.32 | -0.13 | 0.11 | 0.659 |

| | | | | | | | | |
|---|---|---|---|---|---|---|---|---|
| ANKIB1 | S737 | -0.03 | 0.05 | 0.03 | 0.04 | -0.08 | -0.02 | 0.426 |
| ANKLE2 | S866 | 0.45 | -0.07 | 0.07 | -0.30 | -0.34 | 0.05 | 0.153 |
| ANKRD11 | S408; T/S | -0.01 | -0.11 | -0.08 | 0.08 | -0.10 | 0.19 | 0.257 |
| ANKRD11 | S834 | 0.19 | 0.18 | 0.15 | -0.13 | 0.02 | -0.53 | 0.075 |
| ANKRD12 | S1143 | -0.42 | 0.18 | -0.30 | 0.25 | 0.25 | -0.10 | 0.225 |
| ANKRD12 | S1799 | -0.55 | 0.25 | 0.05 | 0.18 | -0.20 | 0.12 | 0.688 |
| ANKRD12 | S568 | 0.08 | -0.10 | 0.37 | 0.07 | -0.26 | -0.26 | 0.2 |
| ANKRD12 | S1799 | -0.19 | 0.27 | 0.30 | 0.10 | -0.37 | -0.24 | 0.233 |
| ANKRD17 | S2401 | 0.23 | 0.13 | -0.36 | 0.03 | -0.30 | 0.16 | 0.887 |
| ANKRD17 | S2067 | 0.16 | 0.10 | 0.09 | -0.16 | 0.01 | -0.23 | 0.03 |
| ANKRD17 | S19 | 0.17 | 0.21 | 0.29 | 0.01 | -0.16 | -0.74 | 0.089 |
| ANKRD17 | S | 0.14 | 0.09 | 0.47 | -0.09 | -0.32 | -0.49 | 0.033 |
| ANKRD24 | S935 | -0.39 | -0.13 | -0.62 | -0.23 | -0.08 | 0.90 | 0.205 |
| ANKRD24 | S935 | -0.22 | -0.16 | -0.09 | -0.35 | -0.14 | 0.69 | 0.518 |
| ANKRD26 | S631 | 0.05 | -0.04 | -0.02 | 0.03 | -0.03 | 0.00 | 0.938 |
| ANKRD34A | S392 | 0.04 | -0.97 | 0.09 | -0.47 | 0.47 | 0.36 | 0.433 |
| ANKRD54 | S63 | -0.13 | -0.12 | 0.00 | 0.20 | 0.20 | -0.20 | 0.35 |
| ANKS1A | S647 | 0.41 | 0.07 | -0.13 | -0.19 | -0.03 | -0.22 | 0.19 |
| ANKS1A | S628 | 0.45 | 0.10 | -0.17 | -0.28 | -0.09 | -0.14 | 0.189 |
| ANKS1A | S663 | 0.45 | -0.07 | 0.16 | -0.26 | 0.10 | -0.61 | 0.161 |
| ANKZF1 | T607 | -0.05 | 0.13 | 0.01 | -0.13 | 0.15 | -0.14 | 0.57 |
| ANO8 | S669 | -0.04 | -0.32 | -0.16 | 0.13 | 0.05 | 0.26 | 0.032 |
| ANO8 | S641 | -0.47 | 0.18 | 0.12 | 0.04 | 0.06 | -0.02 | 0.708 |
| ANO8 | S641 | -0.10 | 0.42 | -0.03 | -0.63 | 0.20 | -0.07 | 0.419 |
| ANXA2 | S127 | -0.16 | 0.31 | -0.37 | -0.12 | 0.14 | 0.09 | 0.635 |
| ANXA2 | S184 | 0.07 | 0.06 | -0.23 | -0.07 | 0.02 | 0.12 | 0.622 |
| ANXA2 | S12 | 0.09 | 0.02 | -0.04 | 0.08 | -0.01 | -0.15 | 0.515 |
| ANXA2 | S85 | 0.03 | 0.21 | -0.09 | 0.06 | -0.07 | -0.17 | 0.365 |
| AP1S1 | S147 | -0.39 | 0.04 | -0.06 | 0.02 | -0.19 | 0.44 | 0.384 |
| AP2A1 | S655 | -0.06 | -0.08 | -0.05 | 0.09 | 0.10 | -0.02 | 0.034 |
| AP2M1 | T156 | 0.20 | 0.47 | -0.46 | -0.49 | 0.12 | -0.10 | 0.527 |
| AP2M1 | T156 | -0.12 | 0.36 | 0.08 | -0.17 | -0.07 | -0.15 | 0.18 |
| AP3B1 | S276 | 0.06 | -0.04 | -0.02 | -0.07 | 0.11 | -0.04 | 0.992 |
| AP3D1 | S829 | 0.35 | 0.12 | 0.12 | -0.28 | 0.03 | -0.53 | 0.064 |
| AP3D1 | T/S | 0.23 | 0.23 | 0.12 | -0.11 | 0.07 | -0.77 | 0.147 |
| APBA3 | S9 | -0.02 | -0.27 | -0.15 | 0.47 | -0.13 | -0.02 | 0.277 |
| APBB2 | S | -0.19 | 0.01 | -0.07 | -0.09 | -0.19 | 0.43 | 0.544 |
| APBB2 | S412 | -0.19 | 0.01 | -0.07 | -0.09 | -0.19 | 0.43 | 0.544 |
| APC | S2674 | 0.15 | 0.13 | -0.34 | -0.44 | -0.15 | 0.45 | 0.931 |
| APC | T/S/Y | 0.28 | -0.04 | 0.08 | 0.03 | 0.27 | -0.92 | 0.456 |
| APC | S2350 | 0.24 | 0.24 | 0.35 | -0.52 | 0.18 | -0.89 | 0.095 |
| APC2 | S1767 | -0.06 | 0.02 | -0.23 | 0.06 | -0.22 | 0.35 | 0.441 |
| API5 | S464 | 0.04 | 0.02 | -0.05 | -0.03 | -0.10 | 0.12 | 0.931 |
| API5 | S464 | 0.15 | 0.12 | -0.13 | -0.09 | -0.04 | -0.04 | 0.33 |
| API5 | S464 | 0.15 | 0.18 | 0.20 | -0.20 | -0.07 | -0.36 | 0.01 |
| API5 | S464 | 0.28 | 0.15 | 0.26 | -0.35 | 0.05 | -0.60 | 0.051 |
| APIP | S87 | 0.09 | -0.04 | -0.11 | 0.02 | -0.26 | 0.24 | 0.897 |
| APLP2 | S213 | -0.07 | -0.06 | 0.24 | 0.09 | -0.06 | -0.19 | 0.517 |

| | | | | | | | | |
|---|---|---|---|---|---|---|---|---|
| APOO | S44 | 0.00 | -0.25 | -0.47 | 0.25 | -0.22 | 0.48 | 0.172 |
| APOOL | S220 | 0.06 | -0.14 | 0.04 | 0.09 | -0.08 | 0.01 | 0.831 |
| APOOL | S203 | 0.11 | -0.06 | -0.05 | -0.06 | 0.03 | 0.03 | 0.998 |
| APTX | S132; S135 | -0.02 | -0.09 | 0.23 | 0.03 | -0.23 | 0.05 | 0.533 |
| AR | S310 | -0.48 | -0.03 | -0.22 | 0.10 | 0.16 | 0.34 | 0.042 |
| AR | S/T | -0.12 | -0.12 | -0.08 | 0.09 | 0.14 | 0.06 | 0.001 |
| ARAF | S582 | 0.04 | -0.06 | -0.05 | 0.01 | -0.05 | 0.09 | 0.46 |
| ARFGAP1 | S/T | -0.44 | 0.14 | -0.44 | 0.37 | -0.37 | 0.44 | 0.294 |
| ARFGAP1 | S/T | 0.01 | -0.02 | -0.34 | 0.05 | 0.10 | 0.15 | 0.136 |
| ARFGEF1 | S52 | -0.17 | -0.05 | -0.11 | 0.34 | -0.10 | 0.02 | 0.219 |
| ARFGEF1 | S52 | 0.07 | -0.12 | -0.04 | 0.10 | 0.08 | -0.11 | 0.561 |
| ARFGEF1 | S1079 | 0.04 | -0.01 | 0.07 | -0.17 | 0.21 | -0.17 | 0.558 |
| ARFGEF2 | S1528 | -0.03 | 0.09 | 0.01 | -0.36 | 0.12 | 0.10 | 0.686 |
| ARFGEF3 | S1651 | -0.03 | -0.05 | -0.21 | 0.04 | 0.14 | 0.07 | 0.049 |
| ARFGEF3 | T1849; S1856; S1857 | -0.20 | 0.04 | -0.11 | -0.01 | -0.03 | 0.26 | 0.234 |
| ARFGEF3 | T1849; S1851; S1852 | -0.20 | 0.04 | -0.11 | -0.01 | -0.03 | 0.26 | 0.234 |
| ARFGEF3 | S1061; S1066 | -0.17 | -0.03 | -0.08 | -0.02 | -0.11 | 0.34 | 0.312 |
| ARFGEF3 | S/T | 0.07 | -0.06 | -0.25 | -0.11 | -0.07 | 0.35 | 0.472 |
| ARFGEF3 | S592 | -0.15 | 0.09 | -0.15 | 0.17 | -0.12 | 0.12 | 0.35 |
| ARFGEF3 | S632 | -0.24 | 0.15 | -0.02 | 0.20 | 0.00 | -0.13 | 0.734 |
| ARFGEF3 | S1856; S1857 | -0.12 | 0.05 | -0.02 | 0.03 | -0.15 | 0.18 | 0.658 |
| ARFGEF3 | S2079 | 0.02 | 0.21 | -0.27 | -0.09 | -0.11 | 0.18 | 0.977 |
| ARFGEF3 | S/T | 0.05 | -0.14 | 0.14 | -0.20 | 0.02 | 0.11 | 0.762 |
| ARFGEF3 | S1066 | 0.00 | -0.02 | 0.09 | -0.07 | -0.07 | 0.07 | 0.444 |
| ARFGEF3 | S1066 | -0.21 | 0.04 | 0.27 | -0.23 | -0.06 | 0.12 | 0.616 |
| ARFGEF3 | S1991 | 0.31 | -0.21 | 0.00 | 0.20 | -0.10 | -0.29 | 0.663 |
| ARFGEF3 | S/T | -0.15 | 0.26 | 0.15 | -0.35 | 0.21 | -0.25 | 0.362 |
| ARGLU1 | S94 | 0.07 | 0.03 | -0.27 | -0.12 | 0.27 | -0.03 | 0.583 |
| ARGLU1 | S60 | 0.16 | -0.11 | -0.02 | -0.03 | 0.02 | -0.03 | 0.764 |
| ARHGAP1 | S51 | -0.09 | -0.09 | -0.07 | 0.18 | 0.11 | -0.06 | 0.091 |
| ARHGAP10 | S591 | 0.03 | 0.11 | -0.09 | -0.12 | -0.07 | 0.12 | 0.704 |
| ARHGAP11A | S675 | -0.29 | -0.34 | -0.40 | 0.17 | 0.30 | 0.34 | 5E-04 |
| ARHGAP11A | S787 | -0.14 | -0.24 | -0.08 | 0.07 | 0.21 | 0.14 | 0.01 |
| ARHGAP11A | T508 | -0.29 | -0.10 | -0.09 | -0.07 | 0.17 | 0.30 | 0.081 |
| ARHGAP11A | S875 | -0.06 | -0.23 | -0.11 | 0.01 | 0.14 | 0.21 | 0.03 |
| ARHGAP11A | T567 | -0.15 | -0.14 | -0.07 | 0.38 | -0.01 | -0.08 | 0.204 |
| ARHGAP12 | S240 | 0.31 | -0.84 | -0.74 | 0.30 | 0.18 | 0.32 | 0.135 |
| ARHGAP12 | S103 | -0.09 | -0.02 | 0.16 | -0.07 | 0.14 | -0.15 | 0.694 |
| ARHGAP17 | S575 | 0.12 | 0.05 | -0.13 | -0.17 | -0.06 | 0.16 | 0.781 |
| ARHGAP17 | S676 | 0.36 | 0.12 | -0.22 | 0.26 | -0.85 | 0.02 | 0.504 |
| ARHGAP17 | S676 | 0.29 | 0.41 | 0.12 | -0.28 | -0.67 | -0.14 | 0.024 |
| ARHGAP21 | S1605 | -0.05 | -0.09 | 0.13 | -0.12 | -0.07 | 0.17 | 0.959 |
| ARHGAP21 | S1433 | 0.26 | -0.01 | -0.25 | 0.06 | -0.13 | 0.00 | 0.895 |
| ARHGAP29 | S559 | 0.11 | -0.09 | -0.23 | -0.14 | -0.18 | 0.42 | 0.654 |
| ARHGAP32 | S892 | -0.28 | -0.27 | -0.19 | 0.36 | -0.05 | 0.28 | 0.028 |
| ARHGAP32 | T1236 | -0.31 | -0.10 | -0.10 | 0.13 | 0.27 | 0.04 | 0.032 |
| ARHGAP32 | S952 | -0.20 | -0.03 | -0.01 | -0.04 | 0.29 | -0.06 | 0.333 |
| ARHGAP32 | S821 | -0.01 | 0.09 | -0.31 | 0.08 | -0.05 | 0.15 | 0.363 |

| | | | | | | | | |
|---|---|---|---|---|---|---|---|---|
| ARHGAP32 | S/T | -0.02 | 0.16 | 0.18 | -0.16 | -0.01 | -0.18 | 0.059 |
| ARHGAP35 | S975 | 0.15 | 0.09 | -0.18 | 0.05 | -0.26 | 0.10 | 0.737 |
| ARHGAP35 | S1179 | 0.02 | 0.18 | -0.04 | -0.08 | 0.03 | -0.14 | 0.23 |
| ARHGAP39 | S286 | -0.11 | -0.08 | -0.29 | 0.23 | -0.19 | 0.33 | 0.17 |
| ARHGAP39 | S/T | -0.28 | -0.18 | -0.07 | -0.49 | 0.60 | 0.16 | 0.449 |
| ARHGAP39 | S173 | -0.03 | -0.07 | -0.14 | 0.11 | 0.19 | -0.09 | 0.172 |
| ARHGAP39 | S407 | -0.23 | 0.36 | 0.24 | -0.22 | -0.05 | -0.22 | 0.209 |
| ARHGAP5 | S765 | -0.10 | -0.10 | 0.06 | 0.00 | 0.22 | -0.10 | 0.485 |
| ARHGAP5 | T/S | -0.07 | 0.31 | -0.23 | -0.34 | 0.03 | 0.18 | 0.849 |
| ARHGDIA | T160 | -0.02 | -0.10 | -0.09 | 0.05 | 0.00 | 0.14 | 0.053 |
| ARHGEF11 | S663; T668 | 0.12 | 0.08 | -0.40 | 0.03 | -0.54 | 0.48 | 0.876 |
| ARHGEF11 | S1155 | -0.07 | 0.11 | -0.07 | -0.02 | -0.03 | 0.07 | 0.797 |
| ARHGEF11 | T590 | -0.01 | 0.05 | -0.04 | 0.10 | -0.09 | -0.02 | 0.912 |
| ARHGEF11 | S1295 | -0.11 | 0.09 | 0.03 | 0.04 | 0.10 | -0.17 | 0.908 |
| ARHGEF11 | S1155 | -0.16 | 0.18 | 0.00 | -0.01 | -0.06 | 0.03 | 0.864 |
| ARHGEF11 | T668 | 0.28 | -0.41 | 0.08 | 0.17 | 0.43 | -1.00 | 0.818 |
| ARHGEF12 | S1331 | -0.32 | -0.15 | -0.13 | 0.09 | -0.18 | 0.52 | 0.181 |
| ARHGEF12 | S190 | -0.11 | -0.20 | -0.06 | -0.20 | 0.19 | 0.29 | 0.231 |
| ARHGEF12 | S1288 | -0.02 | 0.13 | 0.22 | 0.04 | -0.16 | -0.27 | 0.107 |
| ARHGEF12 | T/S/Y | 0.00 | 0.30 | 0.16 | -0.47 | -0.20 | 0.09 | 0.132 |
| ARHGEF16 | S208 | -0.04 | 0.06 | -0.51 | 0.15 | -0.15 | 0.35 | 0.283 |
| ARHGEF16 | S227; S230; S/T | -0.19 | 0.15 | -0.10 | 0.05 | 0.00 | 0.07 | 0.455 |
| ARHGEF17 | S142 | -0.11 | 0.04 | -0.01 | 0.03 | 0.02 | 0.02 | 0.34 |
| ARHGEF18 | S1291 | 0.35 | 0.23 | 0.04 | -0.16 | -0.08 | -0.54 | 0.05 |
| ARHGEF2 | S956 | -0.27 | -0.12 | -0.13 | 0.27 | 0.16 | 0.03 | 0.019 |
| ARHGEF2 | T/S | -0.10 | -0.14 | -0.03 | 0.00 | 0.03 | 0.21 | 0.083 |
| ARHGEF2 | T/S | -0.13 | -0.10 | 0.02 | 0.14 | 0.18 | -0.15 | 0.332 |
| ARHGEF2 | S956; S960 | 0.06 | 0.01 | -0.10 | 0.02 | 0.06 | -0.06 | 0.762 |
| ARHGEF26 | S392 | -0.06 | 0.08 | -0.04 | -0.01 | -0.07 | 0.10 | 0.86 |
| ARHGEF26 | S725 | 0.04 | 0.05 | -0.04 | -0.07 | -0.06 | 0.07 | 0.614 |
| ARHGEF26 | S124 | -0.14 | 0.18 | 0.04 | -0.03 | -0.12 | 0.04 | 0.554 |
| ARHGEF26 | S80 | 0.14 | 0.27 | 0.12 | 0.00 | -0.29 | -0.34 | 0.03 |
| ARHGEF5 | S1126 | -0.13 | -0.07 | 0.02 | 0.09 | -0.12 | 0.19 | 0.333 |
| ARHGEF5 | S1405 | 0.05 | 0.03 | -0.23 | -0.09 | 0.10 | 0.10 | 0.477 |
| ARHGEF6 | S684 | -0.31 | -0.15 | 0.18 | -0.46 | -0.01 | 0.52 | 0.737 |
| ARHGEF7 | S249 | 0.03 | 0.03 | 0.02 | 0.02 | -0.11 | 0.00 | 0.246 |
| ARHGEF7 | S518 | 0.05 | 0.14 | 0.05 | -0.24 | 0.24 | -0.32 | 0.348 |
| ARID1A | T1888 | -0.77 | -0.03 | -0.29 | 0.24 | 0.46 | 0.07 | 0.063 |
| ARID1A | T1888 | -0.18 | -0.22 | -0.46 | 0.08 | 0.23 | 0.39 | 0.014 |
| ARID1A | S1184 | -0.06 | -0.37 | -0.19 | 0.06 | 0.42 | 0.02 | 0.075 |
| ARID1A | S1751 | 0.03 | -0.14 | -0.34 | 0.05 | 0.26 | 0.06 | 0.101 |
| ARID1A | S696 | 0.01 | -0.01 | -0.10 | 0.02 | -0.07 | 0.13 | 0.438 |
| ARID1A | S696; S702 | 0.04 | 0.06 | -0.02 | -0.14 | -0.19 | 0.22 | 0.641 |
| ARID1A | S1755 | 0.00 | 0.09 | 0.02 | -0.02 | 0.06 | -0.17 | 0.326 |
| ARID1A | S1755 | 0.06 | 0.05 | 0.05 | -0.08 | -0.03 | -0.04 | 0.002 |
| ARID1A | S1944 | -0.05 | 0.12 | 0.46 | -0.82 | -0.32 | 0.27 | 0.25 |
| ARID1B | S1625 | 0.02 | -0.15 | -0.17 | -0.14 | -0.13 | 0.46 | 0.483 |
| ARID2 | S1391 | -0.50 | 0.00 | -0.01 | 0.13 | -0.19 | 0.40 | 0.297 |

| | | | | | | | | |
|---|---|---|---|---|---|---|---|---|
| ARID2 | S689 | 0.18 | 0.04 | -0.27 | -0.06 | 0.19 | -0.16 | 0.97 |
| ARID2 | S1300 | -0.33 | 0.51 | -0.29 | -0.33 | -0.02 | 0.24 | 0.998 |
| ARID2 | S1496 | -0.13 | 0.00 | 0.31 | -0.03 | -0.25 | 0.04 | 0.416 |
| ARID2 | T1726 | 0.13 | 0.26 | -0.08 | -0.18 | -0.26 | 0.05 | 0.165 |
| ARID3A | S88 | -0.46 | -0.06 | 0.16 | -0.09 | -0.11 | 0.41 | 0.485 |
| ARID3A | S77; S81; S88 | -0.11 | 0.00 | -0.12 | 0.21 | 0.36 | -0.49 | 0.711 |
| ARID4A | S1140; S1145 | 0.03 | -0.15 | -0.06 | -0.16 | 0.01 | 0.29 | 0.486 |
| ARID4A | S1109 | 0.09 | -0.03 | -0.04 | -0.05 | -0.12 | 0.14 | 0.896 |
| ARID4A | S159; S160 | 0.16 | 0.05 | 0.13 | -0.25 | -0.06 | -0.07 | 0.031 |
| ARID4A | S557; S561 | 0.01 | 0.08 | 0.26 | -0.44 | -0.04 | 0.04 | 0.195 |
| ARID4A | S | 0.14 | 0.21 | 0.21 | -0.22 | -0.09 | -0.34 | 0.006 |
| ARID4B | S790; T793 | -0.04 | -0.14 | -0.22 | 0.08 | -0.13 | 0.35 | 0.193 |
| ARID4B | S1152; S1153; S1155 | -0.07 | -0.13 | -0.05 | 0.08 | -0.06 | 0.20 | 0.122 |
| ARID4B | S483 | -0.03 | 0.05 | -0.22 | -0.04 | 0.00 | 0.20 | 0.345 |
| ARID4B | S675 | -0.23 | -0.18 | 0.20 | 0.18 | 0.13 | -0.17 | 0.545 |
| ARID4B | S790; T793 | 0.13 | 0.04 | -0.12 | -0.12 | 0.04 | 0.01 | 0.667 |
| ARID4B | S675 | 0.47 | 0.02 | -0.50 | 0.23 | 0.02 | -0.50 | 0.83 |
| ARID4B | S/T | -0.05 | 0.08 | 0.16 | -0.12 | -0.19 | 0.09 | 0.27 |
| ARID4B | S790 | 0.13 | 0.17 | 0.02 | -0.05 | -0.14 | -0.17 | 0.017 |
| ARID5A | T313 | -0.13 | 0.08 | -0.05 | -0.20 | -0.13 | 0.36 | 0.825 |
| ARID5A | S23 | 0.35 | 0.31 | 0.14 | -0.31 | -0.21 | -0.49 | 0.004 |
| ARID5B | T539 | -0.06 | -0.12 | 0.05 | -0.20 | -0.14 | 0.38 | 0.78 |
| ARID5B | S1032 | 0.01 | 0.08 | -0.01 | 0.09 | -0.05 | -0.14 | 0.445 |
| ARID5B | S264 | 0.00 | 0.02 | 0.15 | 0.02 | 0.03 | -0.24 | 0.324 |
| ARID5B | T539 | 0.03 | 0.36 | -0.08 | 0.08 | -0.17 | -0.31 | 0.242 |
| ARK2N | S145 | 0.09 | 0.01 | -0.16 | 0.00 | 0.00 | 0.05 | 0.63 |
| ARK2N | S66 | 0.10 | -0.03 | 0.08 | -0.04 | 0.04 | -0.16 | 0.222 |
| ARK2N | S66 | 0.38 | -0.01 | -0.15 | -0.33 | 0.12 | -0.12 | 0.43 |
| ARK2N | S145 | 0.07 | 0.01 | 0.31 | -0.25 | -0.11 | -0.10 | 0.054 |
| ARL6IP4 | S141 | -0.16 | -0.20 | -0.04 | 0.50 | 0.08 | -0.35 | 0.445 |
| ARL6IP4 | S55 | 0.17 | 0.14 | 0.10 | -0.10 | -0.15 | -0.20 | 0.001 |
| ARL6IP6 | S80 | -0.04 | 0.00 | -0.07 | -0.06 | 0.01 | 0.14 | 0.388 |
| ARL6IP6 | S80 | -0.01 | -0.03 | -0.02 | -0.12 | 0.00 | 0.15 | 0.724 |
| ARL6IP6 | S80 | -0.02 | 0.06 | 0.06 | -0.11 | -0.03 | 0.03 | 0.246 |
| ARL6IP6 | S60 | 0.10 | 0.07 | 0.21 | -0.19 | -0.20 | -0.03 | 0.021 |
| ARL6IP6 | S80 | 0.13 | 0.22 | 0.12 | -0.28 | 0.01 | -0.28 | 0.028 |
| ARMCX3 | S110 | -0.08 | 0.00 | -0.04 | 0.18 | 0.07 | -0.16 | 0.527 |
| ARMCX3 | S119 | -0.04 | -0.14 | 0.17 | 0.22 | -0.19 | -0.07 | 0.962 |
| ARMCX3 | S119 | 0.01 | -0.08 | 0.12 | 0.13 | -0.08 | -0.13 | 0.699 |
| ARMCX3 | S110 | 0.05 | -0.13 | 0.29 | -0.23 | 0.10 | -0.14 | 0.365 |
| ARMCX3 | S67 | 0.19 | 0.15 | 0.12 | -0.05 | -0.02 | -0.50 | 0.091 |
| ARPP19 | S23 | 0.24 | 0.36 | -0.10 | -0.22 | -0.19 | -0.21 | 0.055 |
| ARRB1 | S412 | 0.11 | 0.29 | 0.03 | -0.10 | -0.03 | -0.40 | 0.081 |
| ARRB1 | S412 | 0.15 | 0.24 | 0.13 | -0.10 | -0.19 | -0.31 | 0.006 |
| ARV1 | T22 | -0.20 | 0.02 | -0.14 | -0.16 | 0.19 | 0.23 | 0.235 |
| ARVCF | S332; S335 | -0.18 | -0.16 | -0.09 | 0.07 | -0.13 | 0.40 | 0.173 |
| ARVCF | S864 | 0.01 | -0.07 | 0.03 | -0.01 | -0.10 | 0.13 | 0.819 |
| ARVCF | T642 | 0.13 | 0.12 | 0.33 | -0.21 | -0.12 | -0.37 | 0.012 |

| | | | | | | | | |
|---|---|---|---|---|---|---|---|---|
| **ASCC2** | **S713** | -0.34 | -0.15 | 0.05 | 0.19 | -0.01 | 0.18 | 0.11 |
| **ASCC3** | **S2195** | -0.09 | -0.09 | -0.24 | 0.13 | 0.12 | 0.13 | 0.005 |
| **ASCL5** | **S** | -0.31 | 0.04 | -0.03 | -0.12 | 0.11 | 0.24 | 0.295 |
| **ASF1A** | **S175** | 0.02 | 0.15 | 0.00 | -0.14 | 0.01 | -0.05 | 0.141 |
| **ASH1L** | **S1689** | -0.25 | -0.04 | -0.02 | 0.10 | -0.36 | 0.43 | 0.541 |
| **ASH1L** | **S449** | -0.05 | -0.01 | -0.06 | -0.02 | 0.00 | 0.14 | 0.192 |
| **ASH1L** | **S2563** | 0.06 | -0.06 | 0.11 | -0.28 | -0.04 | 0.17 | 0.588 |
| **ASH1L** | **T2561** | 0.06 | -0.06 | 0.11 | -0.28 | -0.04 | 0.17 | 0.588 |
| **ASH1L** | **S1630** | -0.31 | 0.28 | 0.12 | -0.54 | -0.11 | 0.35 | 0.694 |
| **ASH2L** | **S101** | -0.09 | -0.21 | 0.04 | 0.01 | -0.07 | 0.26 | 0.287 |
| **ASH2L** | **S101** | -0.16 | -0.21 | 0.17 | -0.30 | 0.02 | 0.36 | 0.691 |
| **ASH2L** | **S101** | -0.06 | 0.08 | 0.06 | -0.08 | -0.04 | 0.02 | 0.295 |
| **ASH2L** | **S101** | -0.05 | 0.07 | 0.07 | -0.02 | -0.03 | -0.05 | 0.202 |
| **ASPH** | **S29** | 0.28 | 0.33 | 0.21 | -0.24 | -0.20 | -0.60 | 0.009 |
| **ASPH** | **S29** | 0.24 | 0.40 | 0.20 | -0.41 | -0.18 | -0.49 | 0.004 |
| **ASPM** | **T/S** | -0.55 | -1.39 | -0.70 | 0.38 | 0.51 | 0.67 | 0.007 |
| **ASPM** | **S367** | -0.24 | -0.43 | -0.40 | 0.34 | 0.36 | 0.13 | 0.003 |
| **ASPM** | **T/S** | -0.11 | -0.55 | -0.21 | 0.12 | 0.31 | 0.26 | 0.022 |
| **ASPM** | **S392** | -0.26 | -0.13 | -0.18 | 0.23 | 0.11 | 0.15 | 0.003 |
| **ASPM** | **S605** | -0.29 | -0.30 | 0.06 | 0.15 | -0.12 | 0.38 | 0.165 |
| **ASPM** | **S1103** | -0.08 | -0.18 | -0.14 | 0.17 | 0.13 | 0.07 | 0.004 |
| **ASPM** | **S577** | -0.08 | -0.31 | 0.10 | 0.06 | -0.10 | 0.27 | 0.344 |
| **ASPM** | **S1103** | 0.02 | -0.16 | -0.13 | 0.05 | 0.15 | 0.05 | 0.062 |
| **ASPM** | **S425** | 0.11 | -0.17 | -0.14 | -0.18 | 0.19 | 0.14 | 0.484 |
| **ASPM** | **S14** | 0.01 | -0.05 | 0.01 | 0.04 | -0.05 | 0.04 | 0.543 |
| **ASPM** | **T279** | -0.52 | 0.22 | 0.24 | 0.42 | -0.35 | -0.26 | 0.917 |
| **ASPM** | **S425** | 0.28 | -0.42 | 0.18 | 0.10 | -0.37 | 0.08 | 0.796 |
| **ASPM** | **S425** | -0.27 | -0.08 | 0.43 | -0.28 | 0.14 | -0.07 | 0.711 |
| **ASPSCR1** | **S500** | -0.24 | 0.22 | 0.16 | 0.54 | -0.63 | -0.37 | 0.63 |
| **ASXL1** | **S503** | 0.10 | 0.20 | 0.14 | -0.16 | -0.24 | -0.09 | 0.004 |
| **ATAD1** | **S322** | -0.08 | -0.01 | -0.07 | 0.01 | -0.01 | 0.14 | 0.127 |
| **ATAD1** | **S322** | -0.05 | -0.03 | 0.08 | 0.04 | -0.09 | 0.03 | 0.911 |
| **ATAD2** | **S1302** | -0.28 | -0.28 | -0.49 | 0.17 | 0.01 | 0.59 | 0.032 |
| **ATAD2** | **S1243** | -0.21 | -0.10 | -0.48 | 0.16 | 0.21 | 0.28 | 0.015 |
| **ATAD2** | **S** | -0.27 | -0.16 | -0.27 | 0.12 | -0.22 | 0.59 | 0.17 |
| **ATAD2** | **S/T** | -0.11 | -0.17 | -0.30 | 0.02 | 0.27 | 0.20 | 0.019 |
| **ATAD2** | **S** | -0.19 | -0.21 | -0.20 | 0.14 | -0.10 | 0.43 | 0.083 |
| **ATAD2** | **S141** | -0.10 | -0.05 | -0.03 | -0.07 | 0.08 | 0.15 | 0.173 |
| **ATAD2** | **S87; S88** | -0.18 | 0.11 | -0.06 | -0.02 | -0.10 | 0.21 | 0.596 |
| **ATAD2** | **S1243** | -0.21 | -0.02 | 0.14 | -0.11 | 0.35 | -0.24 | 0.891 |
| **ATAD2** | **S** | -0.20 | -0.04 | 0.18 | -0.29 | 0.15 | 0.13 | 0.918 |
| **ATAD2** | **T142** | -0.01 | -0.02 | 0.00 | -0.07 | 0.01 | 0.08 | 0.773 |
| **ATAD2** | **T1152; T1158** | 0.31 | -0.11 | -0.20 | 0.01 | 0.07 | -0.14 | 0.908 |
| **ATAD2** | **S1283** | 0.08 | 0.10 | -0.16 | -0.09 | 0.17 | -0.15 | 0.816 |
| **ATAD2** | **S87; S88** | -0.21 | 0.18 | 0.08 | -0.01 | -0.10 | 0.01 | 0.694 |
| **ATAD2** | **S1283** | 0.10 | 0.05 | -0.04 | -0.03 | 0.02 | -0.11 | 0.238 |
| **ATAD2** | **S1131** | -0.07 | 0.02 | 0.18 | -0.15 | 0.27 | -0.33 | 0.592 |
| **ATAD2** | **S1190** | 0.02 | 0.09 | 0.09 | -0.21 | -0.14 | 0.12 | 0.222 |

| | | | | | | | | |
|---|---|---|---|---|---|---|---|---|
| ATAD2 | S/T/Y | 0.04 | 0.21 | 0.01 | -0.18 | 0.02 | -0.13 | 0.104 |
| ATAD2 | S1139 | 0.08 | -0.12 | 0.23 | 0.18 | 0.17 | -0.76 | 0.575 |
| ATAD2 | S1200 | 0.05 | -0.07 | 0.35 | -0.22 | -0.07 | -0.11 | 0.139 |
| ATAD2 | S | 0.09 | 0.06 | 0.20 | -0.23 | -0.04 | -0.14 | 0.022 |
| ATAD2 | S969 | 0.04 | 0.05 | 0.26 | -0.30 | 0.27 | -0.46 | 0.295 |
| ATAD2 | Y/S/T | 0.05 | 0.17 | 0.17 | -0.24 | 0.06 | -0.28 | 0.072 |
| ATAD2 | T1152 | 0.06 | 0.11 | 0.22 | -0.43 | 0.17 | -0.25 | 0.179 |
| ATAD2 | S1243; S1255 | 0.10 | 0.14 | 0.21 | -0.31 | -0.03 | -0.18 | 0.021 |
| ATAD2 | S88 | 0.06 | 0.08 | 0.30 | -0.26 | -0.09 | -0.17 | 0.024 |
| ATAD2 | S342 | 0.24 | 0.19 | 0.12 | -0.43 | 0.13 | -0.41 | 0.086 |
| ATAD2 | S1200 | 0.33 | 0.09 | 0.19 | -0.34 | -0.05 | -0.35 | 0.019 |
| ATAD2 | S1233 | 0.23 | 0.14 | 0.34 | -0.31 | 0.08 | -0.75 | 0.087 |
| ATAD2 | S337; S342 | 0.36 | 0.27 | 0.21 | -0.37 | -0.25 | -0.42 | 6E-04 |
| ATAD2B | S1379 | -0.30 | -0.40 | -0.42 | 0.22 | 0.54 | 0.10 | 0.008 |
| ATAD2B | S61 | -0.41 | -0.18 | -0.16 | 0.21 | -0.08 | 0.45 | 0.063 |
| ATAD2B | S16 | 0.05 | -0.11 | -0.66 | 0.40 | -0.33 | 0.36 | 0.3 |
| ATAD2B | S372 | -0.15 | -0.16 | -0.30 | 0.32 | -0.04 | 0.23 | 0.035 |
| ATAD2B | S1298 | -0.22 | -0.10 | -0.20 | 0.05 | -0.23 | 0.53 | 0.266 |
| ATAD2B | S118 | -0.23 | -0.17 | 0.00 | 0.14 | 0.20 | 0.01 | 0.047 |
| ATAD2B | S1273 | -0.15 | -0.25 | -0.01 | -0.13 | 0.00 | 0.43 | 0.264 |
| ATAD2B | S81 | -0.06 | -0.21 | -0.11 | 0.24 | -0.03 | 0.13 | 0.058 |
| ATAD2B | T/S | -0.02 | -0.21 | -0.08 | -0.04 | 0.02 | 0.29 | 0.178 |
| ATAD2B | S1276 | 0.07 | -0.14 | -0.20 | 0.08 | 0.15 | 0.01 | 0.134 |
| ATAD2B | S1338 | -0.09 | -0.05 | -0.12 | 0.19 | -0.14 | 0.17 | 0.214 |
| ATAD2B | S1233 | -0.15 | -0.04 | 0.00 | 0.11 | -0.04 | 0.10 | 0.137 |
| ATAD2B | S1347 | 0.07 | -0.03 | 0.03 | -0.06 | 0.02 | -0.04 | 0.287 |
| ATAD2B | S1298 | -0.01 | 0.02 | 0.03 | 0.21 | 0.06 | -0.38 | 0.787 |
| ATAD2B | S86; S | -0.26 | 0.18 | 0.12 | 0.06 | -0.55 | 0.28 | 0.779 |
| ATAD2B | S318 | -0.17 | -0.02 | 0.30 | 0.17 | -0.25 | -0.11 | 0.61 |
| ATAD2B | S/T | 0.27 | 0.05 | -0.19 | -0.04 | -0.20 | 0.06 | 0.539 |
| ATAD2B | S1238 | -0.01 | 0.06 | 0.10 | -0.09 | -0.21 | 0.12 | 0.328 |
| ATAD2B | S1347 | 0.04 | 0.16 | -0.04 | -0.16 | 0.14 | -0.18 | 0.365 |
| ATAD2B | S1347 | 0.23 | 0.37 | -0.08 | -0.33 | 0.16 | -0.55 | 0.173 |
| ATAD2B | S/T | -0.04 | 0.44 | 0.27 | -0.19 | -0.62 | -0.11 | 0.067 |
| ATAD5 | T590 | 0.09 | -0.53 | -0.15 | 0.12 | -0.09 | 0.40 | 0.214 |
| ATAD5 | S86 | 0.17 | 0.02 | -0.38 | -0.15 | 0.33 | -0.10 | 0.694 |
| ATAD5 | S602 | -0.01 | -0.12 | 0.19 | -0.14 | 0.06 | -0.01 | 0.665 |
| ATAD5 | S306 | 0.02 | 0.05 | 0.00 | 0.04 | -0.17 | 0.04 | 0.505 |
| ATAD5 | S/T | 0.01 | 0.05 | 0.04 | -0.05 | 0.00 | -0.04 | 0.031 |
| ATAD5 | S354 | 0.04 | 0.09 | 0.04 | -0.15 | -0.15 | 0.11 | 0.233 |
| ATAD5 | S817 | 0.05 | 0.04 | 0.14 | -0.12 | -0.18 | 0.05 | 0.1 |
| ATAD5 | S356 | 0.23 | 0.12 | 0.13 | -0.33 | -0.26 | 0.02 | 0.035 |
| ATAD5 | S1222 | 0.33 | 0.23 | -0.01 | -0.28 | -0.19 | -0.19 | 0.018 |
| ATAD5 | S70 | 0.59 | 0.06 | -0.01 | -0.26 | -0.26 | -0.33 | 0.061 |
| ATAD5 | S306 | 0.20 | 0.26 | 0.40 | -0.43 | -0.22 | -0.44 | 0.002 |
| ATF2 | S328 | 0.00 | -0.25 | -0.42 | 0.29 | -0.05 | 0.29 | 0.072 |
| ATF2 | S136 | -0.13 | -0.19 | -0.08 | 0.15 | 0.08 | 0.13 | 0.003 |
| ATF2 | S328 | 0.16 | -0.10 | -0.22 | 0.12 | -0.19 | 0.17 | 0.618 |

| | | | | | | | | |
|---|---|---|---|---|---|---|---|---|
| ATF2 | T/S | -0.07 | 0.15 | 0.07 | -0.14 | 0.21 | -0.27 | 0.491 |
| ATF2 | T71 | 0.06 | 0.04 | 0.19 | -0.23 | -0.11 | 0.01 | 0.071 |
| ATF2 | S136 | 0.14 | 0.15 | 0.13 | -0.06 | -0.12 | -0.29 | 0.012 |
| ATF2 | T/S | 0.14 | 0.23 | 0.18 | -0.22 | -0.04 | -0.40 | 0.02 |
| ATF6 | S94 | 0.10 | 0.04 | -0.02 | -0.55 | -0.15 | 0.41 | 0.648 |
| ATF7 | S423 | 0.06 | -0.51 | -0.44 | 0.58 | -0.34 | 0.31 | 0.21 |
| ATF7 | T101 | 0.00 | 0.14 | -0.92 | 0.22 | -0.06 | 0.32 | 0.301 |
| ATF7 | S423 | -0.08 | -0.14 | -0.30 | 0.41 | -0.30 | 0.26 | 0.257 |
| ATF7 | T53 | -0.31 | -0.05 | 0.18 | -0.09 | 0.20 | 0.01 | 0.574 |
| ATF7 | S423 | 0.37 | 0.62 | -0.80 | -0.97 | 0.09 | 0.03 | 0.573 |
| ATF7 | S413 | 0.04 | 0.05 | 0.37 | -0.13 | 0.13 | -0.66 | 0.222 |
| ATF7IP | S334 | -2.67 | -2.35 | -2.56 | 1.06 | 0.62 | 0.90 | 3E-05 |
| ATF7IP | S113 | -0.36 | 0.09 | -0.09 | -0.06 | 0.16 | 0.19 | 0.231 |
| ATF7IP | S673 | -0.10 | 0.08 | -0.26 | 0.05 | 0.03 | 0.16 | 0.174 |
| ATF7IP | S113 | 0.13 | 0.01 | -0.02 | 0.02 | 0.06 | -0.22 | 0.428 |
| ATF7IP | S559 | 0.12 | 0.00 | 0.09 | -0.12 | -0.01 | -0.10 | 0.048 |
| ATG2A | S/T | -0.20 | -0.21 | -0.24 | -0.08 | 0.02 | 0.54 | 0.126 |
| ATG2B | S497 | -0.21 | -0.06 | -0.11 | 0.06 | 0.23 | 0.05 | 0.029 |
| ATG2B | S/T | -0.03 | 0.00 | -0.05 | -0.15 | 0.14 | 0.08 | 0.62 |
| ATG2B | S255 | 0.26 | -0.03 | 0.30 | -0.31 | -0.05 | -0.27 | 0.041 |
| ATG9A | S828 | -0.01 | 0.09 | -0.06 | -0.08 | -0.24 | 0.25 | 0.87 |
| ATN1 | S661 | -0.47 | -0.04 | 0.08 | -0.20 | 0.35 | 0.15 | 0.355 |
| ATN1 | S34 | 0.25 | -0.18 | -0.39 | 0.17 | -0.30 | 0.30 | 0.568 |
| ATN1 | S645 | 0.00 | -0.06 | -0.13 | 0.13 | -0.07 | 0.11 | 0.173 |
| ATN1 | S34 | 0.09 | 0.01 | -0.10 | -0.02 | -0.05 | 0.06 | 0.995 |
| ATN1 | S632 | 0.12 | 0.03 | 0.18 | -0.19 | -0.16 | -0.02 | 0.028 |
| ATN1 | T653; S661 | 0.25 | 0.06 | 0.06 | -0.20 | 0.06 | -0.30 | 0.09 |
| ATN1 | S101 | 0.08 | 0.20 | 0.12 | -0.19 | -0.09 | -0.17 | 0.004 |
| ATN1 | S101 | 0.24 | 0.32 | -0.04 | -0.37 | -0.08 | -0.18 | 0.052 |
| ATP11C | S445 | 0.15 | 0.12 | 0.00 | 0.10 | -0.01 | -0.43 | 0.28 |
| ATP13A1 | S899 | -0.10 | -0.09 | 0.13 | 0.01 | -0.15 | 0.17 | 0.82 |
| ATP13A1 | S899 | 0.15 | 0.23 | 0.31 | -0.24 | -0.22 | -0.38 | 0.002 |
| ATP13A2 | S151 | 0.12 | 0.18 | 0.10 | 0.03 | -0.43 | -0.08 | 0.104 |
| ATP13A3 | S817 | -0.03 | 0.08 | 0.15 | 0.08 | -0.12 | -0.18 | 0.214 |
| ATP13A3 | S817 | 0.35 | -0.04 | -0.03 | -0.03 | -0.09 | -0.23 | 0.208 |
| ATP1A1 | S16 | 0.10 | 0.07 | 0.14 | -0.11 | -0.09 | -0.13 | 8E-04 |
| ATP1A1 | S16 | 0.21 | 0.18 | 0.22 | -0.36 | -0.05 | -0.33 | 0.01 |
| ATP1A1 | S16 | 0.15 | 0.33 | 0.18 | -0.35 | -0.15 | -0.30 | 0.004 |
| ATP2A2 | T441 | -0.07 | -0.11 | 0.07 | -0.06 | 0.02 | 0.14 | 0.426 |
| ATP2A2 | S663 | -0.09 | 0.03 | 0.15 | -0.05 | 0.02 | -0.08 | 0.415 |
| ATP2A2 | S378 | 0.23 | 0.01 | 0.08 | -0.13 | 0.31 | -0.75 | 0.397 |
| ATP5F1A | T/S | 0.32 | 0.17 | 0.37 | -0.33 | -0.14 | -0.68 | 0.017 |
| ATP6AP1 | S465 | 0.05 | -0.16 | -0.63 | 0.48 | -0.05 | 0.08 | 0.178 |
| ATP6V0A2 | S695; S700 | -0.24 | -0.09 | 0.14 | -0.02 | -0.11 | 0.26 | 0.542 |
| ATP6V0A2 | S695; S700 | -0.10 | 0.10 | 0.03 | 0.01 | -0.07 | 0.01 | 0.662 |
| ATP6V0A2 | S695 | -0.01 | 0.09 | -0.04 | 0.00 | -0.13 | 0.08 | 0.686 |
| ATP6V0A2 | S695 | 0.04 | 0.14 | 0.07 | -0.03 | -0.02 | -0.24 | 0.081 |
| ATP7A | S357 | -0.15 | -0.06 | -0.21 | -0.44 | 0.44 | 0.24 | 0.466 |

| | | | | | | | | |
|---|---|---|---|---|---|---|---|---|
| ATR | T1989 | 0.33 | 0.04 | -0.68 | 0.33 | -0.41 | 0.12 | 0.767 |
| ATR | T1989 | 0.30 | -0.08 | -0.35 | 0.12 | -0.13 | 0.05 | 0.792 |
| ATRX | S871; S875; S876 | -0.54 | -0.27 | -0.30 | 0.36 | 0.30 | 0.20 | 0.002 |
| ATRX | S26 | -0.23 | -0.30 | -0.30 | 0.27 | 0.02 | 0.39 | 0.011 |
| ATRX | S1348; S1352 | -0.08 | -0.11 | -0.28 | 0.08 | -0.12 | 0.40 | 0.162 |
| ATRX | S | 0.11 | -0.41 | -0.07 | 0.01 | 0.13 | 0.16 | 0.24 |
| ATRX | S1992; S1995 | -0.09 | -0.09 | -0.17 | 0.15 | -0.08 | 0.23 | 0.087 |
| ATRX | S92 | 0.05 | -0.24 | -0.12 | 0.14 | -0.06 | 0.19 | 0.17 |
| ATRX | S92 | 0.05 | -0.24 | -0.12 | 0.14 | -0.06 | 0.19 | 0.17 |
| ATRX | S1992; S1996 | -0.07 | 0.14 | -0.36 | -0.13 | 0.11 | 0.23 | 0.4 |
| ATRX | T674; S675; S677 | -0.05 | -0.07 | -0.10 | 0.10 | -0.05 | 0.15 | 0.081 |
| ATRX | S1527 | -0.05 | -0.04 | -0.06 | 0.03 | -0.09 | 0.19 | 0.308 |
| ATRX | S930 | -0.01 | -0.06 | -0.08 | 0.05 | 0.17 | -0.09 | 0.293 |
| ATRX | S598 | 0.11 | -0.19 | -0.10 | -0.19 | 0.42 | -0.17 | 0.744 |
| ATRX | S92; S112 | -0.04 | -0.01 | -0.05 | 0.09 | 0.06 | -0.06 | 0.234 |
| ATRX | S92; T102; S112 | 0.00 | -0.06 | -0.03 | 0.09 | 0.09 | -0.09 | 0.372 |
| ATRX | S677 | -0.01 | -0.03 | -0.05 | 0.00 | -0.08 | 0.16 | 0.457 |
| ATRX | S677 | -0.09 | -0.01 | 0.02 | -0.05 | -0.05 | 0.17 | 0.573 |
| ATRX | T674; S675; S677 | -0.05 | 0.01 | -0.04 | -0.10 | -0.09 | 0.24 | 0.715 |
| ATRX | S925 | -0.05 | 0.05 | -0.06 | -0.05 | -0.04 | 0.15 | 0.594 |
| ATRX | S92; S108; S112 | -0.02 | 0.13 | -0.19 | 0.13 | 0.01 | -0.09 | 0.725 |
| ATRX | T674; S677 | 0.00 | -0.01 | -0.04 | -0.03 | 0.01 | 0.06 | 0.33 |
| ATRX | S92; S120 | -0.02 | -0.04 | 0.03 | -0.07 | -0.04 | 0.14 | 0.783 |
| ATRX | S/T | 0.37 | -0.31 | -0.15 | -0.12 | -0.13 | 0.21 | 0.94 |
| ATRX | S92; T117 | 0.05 | 0.05 | -0.12 | 0.01 | -0.01 | 0.01 | 0.865 |
| ATRX | S675; S677 | 0.00 | 0.02 | -0.03 | 0.01 | -0.03 | 0.04 | 0.762 |
| ATRX | S1348; S1352 | 0.05 | 0.04 | -0.10 | 0.12 | -0.09 | -0.03 | 0.97 |
| ATRX | S871 | -0.08 | -0.11 | 0.16 | -0.03 | 0.16 | -0.14 | 0.997 |
| ATRX | S1012; S1013; T1016 | -0.11 | 0.04 | 0.06 | 0.02 | -0.12 | 0.09 | 0.985 |
| ATRX | S92 | 0.07 | 0.02 | -0.09 | -0.07 | 0.04 | 0.02 | 0.913 |
| ATRX | S889 | 0.15 | -0.05 | -0.08 | 0.14 | 0.07 | -0.27 | 0.854 |
| ATRX | S92 | 0.09 | 0.11 | -0.11 | 0.02 | 0.01 | -0.12 | 0.513 |
| ATRX | S889 | 0.25 | -0.11 | -0.06 | 0.23 | -0.05 | -0.34 | 0.695 |
| ATRX | S92 | 0.12 | 0.07 | -0.07 | -0.02 | -0.07 | -0.04 | 0.233 |
| ATRX | S849; S850 | 0.11 | 0.07 | -0.04 | -0.08 | 0.00 | -0.06 | 0.15 |
| ATRX | S1061 | 0.17 | 0.00 | -0.02 | -0.07 | -0.15 | 0.04 | 0.252 |
| ATRX | S1348 | 0.01 | 0.09 | 0.09 | -0.01 | -0.12 | -0.08 | 0.031 |
| ATRX | S729 | 0.10 | 0.00 | 0.09 | -0.22 | 0.03 | -0.03 | 0.163 |
| ATRX | S703 | -0.11 | 0.23 | 0.11 | -0.35 | 0.15 | -0.12 | 0.358 |
| ATRX | S1996 | 0.08 | 0.13 | 0.07 | -0.19 | -0.09 | -0.02 | 0.023 |
| ATRX | S729 | 0.08 | 0.05 | 0.17 | -0.11 | -0.05 | -0.18 | 0.016 |
| ATRX | S/T | -0.09 | 0.10 | 0.32 | -0.28 | -0.01 | -0.10 | 0.169 |
| ATRX | S92 | 0.04 | 0.22 | 0.06 | 0.16 | -0.18 | -0.39 | 0.227 |
| ATRX | S1011; S/Y/T | 0.13 | 0.13 | 0.12 | -0.17 | -0.11 | -0.14 | 8E-05 |
| ATRX | S92 | 0.21 | 0.32 | -0.22 | -0.46 | 0.28 | -0.33 | 0.384 |
| ATRX | S598 | 0.19 | 0.06 | 0.18 | -0.21 | -0.06 | -0.22 | 0.011 |
| ATRX | S784 | 0.26 | 0.16 | 0.07 | -0.25 | -0.10 | -0.22 | 0.008 |
| ATRX | S1527 | 0.21 | 0.24 | 0.07 | -0.15 | 0.04 | -0.56 | 0.101 |

| | | | | | | | | |
|---|---|---|---|---|---|---|---|---|
| **ATRX** | **S784; S788** | 0.14 | 0.14 | 0.27 | -0.31 | -0.01 | -0.35 | 0.026 |
| **ATRX** | **S1948** | 0.32 | 0.17 | 0.15 | -0.46 | -0.03 | -0.30 | 0.025 |
| **ATRX** | **T977; S978** | 0.25 | 0.30 | 0.20 | -0.28 | -0.11 | -0.55 | 0.013 |
| **ATRX** | **S784** | 0.32 | 0.19 | 0.26 | -0.39 | -0.12 | -0.43 | 0.005 |
| **ATRX** | **S814; S815** | 0.22 | 0.30 | 0.30 | -0.33 | -0.05 | -0.72 | 0.031 |
| **ATRX** | **S1352** | 0.39 | 0.51 | 0.44 | -0.68 | -0.26 | -1.14 | 0.012 |
| **ATXN1** | **S88** | -0.23 | -0.12 | -0.08 | 0.07 | -0.08 | 0.36 | 0.13 |
| **ATXN1** | **S406** | -0.08 | 0.00 | -0.18 | -0.01 | 0.11 | 0.14 | 0.071 |
| **ATXN1** | **S775** | -0.11 | -0.06 | -0.06 | 0.10 | 0.16 | -0.05 | 0.082 |
| **ATXN2** | **T/S** | -0.04 | -0.02 | -0.04 | 0.13 | -0.10 | 0.06 | 0.437 |
| **ATXN2** | **S889** | -0.14 | 0.29 | 0.07 | 0.08 | 0.08 | -0.52 | 0.46 |
| **ATXN2** | **S889** | -0.14 | 0.29 | 0.07 | 0.08 | 0.08 | -0.52 | 0.46 |
| **ATXN2** | **S772** | 0.15 | 0.20 | 0.10 | -0.10 | -0.15 | -0.26 | 0.005 |
| **ATXN2** | **S784** | 0.13 | 0.22 | 0.18 | -0.27 | -0.23 | -0.12 | 0.002 |
| **ATXN2L** | **S594** | -0.32 | 0.12 | -0.37 | 0.56 | 0.04 | -0.25 | 0.342 |
| **ATXN2L** | **S111** | -0.03 | -0.24 | -0.08 | -0.13 | 0.03 | 0.37 | 0.266 |
| **ATXN2L** | **S111** | -0.05 | -0.12 | -0.09 | 0.22 | -0.10 | 0.10 | 0.174 |
| **ATXN2L** | **S559** | 0.01 | 0.08 | -0.05 | 0.19 | 0.00 | -0.27 | 0.807 |
| **ATXN2L** | **S594** | 0.08 | 0.07 | -0.01 | -0.07 | -0.33 | 0.19 | 0.502 |
| **ATXN2L** | **S680** | -0.22 | 0.19 | 0.20 | -0.16 | -0.22 | 0.14 | 0.487 |
| **ATXN2L** | **S424** | -0.05 | 0.18 | 0.07 | -0.37 | 0.17 | -0.07 | 0.403 |
| **ATXN2L** | **S594** | 0.51 | 0.30 | -0.32 | 0.05 | -0.41 | -0.39 | 0.226 |
| **ATXN2L** | **S339** | -0.13 | 0.26 | 0.43 | -0.13 | -0.07 | -0.56 | 0.122 |
| **ATXN7L2** | **S575** | -0.05 | 0.19 | -0.04 | -0.17 | 0.12 | -0.09 | 0.545 |
| **ATXN7L3** | **S281** | 0.11 | -0.19 | -0.01 | -0.04 | -0.21 | 0.29 | 0.811 |
| **ATXN7L3** | **S131** | 0.08 | -0.01 | 0.03 | -0.02 | -0.02 | -0.07 | 0.078 |
| **AXIN1** | **S493** | 0.04 | -0.02 | -0.17 | 0.18 | -0.27 | 0.19 | 0.649 |
| **B3GAT3** | **S329** | -0.16 | 0.42 | -0.27 | 0.29 | -0.37 | -0.08 | 0.861 |
| **BACH1** | **S688** | -0.11 | -0.64 | 0.04 | 0.07 | -0.06 | 0.47 | 0.202 |
| **BAD** | **S25** | -0.51 | -0.17 | 0.35 | -0.42 | 0.25 | 0.26 | 0.689 |
| **BAG3** | **T406** | 0.02 | 0.04 | -0.08 | 0.08 | -0.17 | 0.09 | 0.927 |
| **BAG3** | **S173** | -0.02 | -0.06 | 0.36 | -0.23 | 0.14 | -0.28 | 0.313 |
| **BAG3** | **S173** | -0.21 | 0.04 | 0.44 | -0.07 | 0.05 | -0.40 | 0.374 |
| **BAG6** | **S113** | -0.51 | -0.60 | 0.56 | 0.35 | -0.58 | 0.28 | 0.697 |
| **BAG6** | **S973** | 0.18 | -0.02 | -0.12 | 0.01 | 0.12 | -0.19 | 0.8 |
| **BAG6** | **S1117** | -0.10 | 0.26 | 0.05 | -0.27 | -0.06 | 0.07 | 0.339 |
| **BAHCC1** | **S1141** | -0.35 | -0.13 | 0.05 | -0.08 | 0.18 | 0.24 | 0.163 |
| **BAHCC1** | **S1718** | 0.19 | 0.35 | 0.02 | -0.19 | -0.09 | -0.41 | 0.035 |
| **BAHCC1** | **S2075** | 0.31 | 0.57 | 0.16 | -0.66 | -0.17 | -0.66 | 0.014 |
| **BAHD1** | **S44** | -0.30 | -0.18 | 0.27 | 0.26 | -0.54 | 0.28 | 0.848 |
| **BAHD1** | **S101; S105** | 0.02 | -0.04 | -0.08 | -0.08 | -0.03 | 0.20 | 0.516 |
| **BAHD1** | **T588** | 0.12 | 0.04 | -0.13 | -0.22 | 0.05 | 0.11 | 0.84 |
| **BAHD1** | **S** | 0.03 | -0.08 | 0.13 | 0.00 | -0.17 | 0.08 | 0.574 |
| **BAHD1** | **S101** | 0.07 | 0.14 | 0.09 | -0.24 | -0.08 | -0.01 | 0.044 |
| **BAHD1** | **S679** | -0.43 | 0.34 | 0.56 | -0.49 | 0.10 | -0.44 | 0.287 |
| **BAIAP2** | **S/T/Y** | -0.13 | -0.13 | -0.07 | -0.03 | 0.21 | 0.11 | 0.041 |
| **BAIAP2** | **S325** | 0.12 | -0.01 | 0.04 | -0.05 | -0.05 | -0.05 | 0.065 |
| **BAIAP2L1** | **S261** | -0.72 | 0.34 | -0.23 | 0.34 | -0.16 | 0.15 | 0.41 |

| | | | | | | | | |
|---|---|---|---|---|---|---|---|---|
| **BAIAP2L1** | **S261** | -0.56 | -0.03 | 0.13 | 0.38 | -0.40 | 0.25 | 0.515 |
| **BAIAP2L1** | **S261** | -0.04 | -0.13 | -0.05 | -0.11 | 0.16 | 0.13 | 0.219 |
| **BAIAP2L1** | **S414** | -0.34 | 0.04 | 0.35 | -0.23 | 0.05 | 0.04 | 0.791 |
| **BAIAP2L1** | **T405** | -0.90 | 0.18 | 0.83 | -0.30 | -0.38 | -0.04 | 0.621 |
| **BAIAP2L1** | **S354** | -0.14 | 0.45 | 0.27 | -0.41 | 0.10 | -0.52 | 0.143 |
| **BANP** | **S145** | -0.59 | -0.14 | -0.29 | 0.25 | 0.00 | 0.51 | 0.041 |
| **BAP1** | **S583; S597** | 0.16 | 0.18 | -0.18 | -0.09 | -0.13 | 0.02 | 0.381 |
| **BAP18** | **S96** | -0.04 | 0.15 | -0.24 | -0.02 | -0.17 | 0.26 | 0.723 |
| **BAP18** | **S96** | 0.18 | 0.14 | -0.03 | -0.15 | 0.03 | -0.21 | 0.101 |
| **BAP18** | **S96** | 0.20 | 0.13 | 0.01 | -0.02 | -0.08 | -0.29 | 0.064 |
| **BARD1** | **S186** | -0.03 | -0.01 | -0.06 | 0.19 | 0.11 | -0.24 | 0.725 |
| **BARD1** | **S186** | -0.09 | 0.06 | 0.01 | 0.13 | 0.03 | -0.15 | 0.928 |
| **BARD1** | **S364** | -0.05 | 0.13 | 0.35 | -0.87 | 0.14 | 0.04 | 0.332 |
| **BAZ1A** | **S286** | -0.54 | -0.93 | -0.66 | 0.61 | 0.66 | 0.08 | 0.006 |
| **BAZ1A** | **S1402** | -0.45 | -0.62 | -0.51 | 0.40 | 0.53 | 0.20 | 0.001 |
| **BAZ1A** | **S1320** | -0.65 | -0.54 | 0.01 | 0.06 | -0.01 | 0.71 | 0.103 |
| **BAZ1A** | **T771** | -0.22 | -0.20 | -0.25 | 0.26 | -0.11 | 0.38 | 0.056 |
| **BAZ1A** | **T731** | -0.09 | -0.18 | -0.33 | 0.25 | 0.19 | 0.08 | 0.011 |
| **BAZ1A** | **T731** | 0.04 | -0.13 | -0.26 | -0.01 | 0.15 | 0.15 | 0.108 |
| **BAZ1A** | **S1413** | -0.01 | -0.06 | -0.19 | 0.02 | -0.11 | 0.30 | 0.295 |
| **BAZ1A** | **T731** | 0.02 | -0.07 | -0.18 | -0.05 | 0.08 | 0.16 | 0.181 |
| **BAZ1A** | **S1413** | -0.09 | 0.06 | -0.03 | -0.05 | -0.11 | 0.20 | 0.77 |
| **BAZ1A** | **S1413** | 0.11 | 0.05 | -0.05 | -0.18 | 0.04 | 0.01 | 0.375 |
| **BAZ1A** | **S/Y** | -0.19 | 0.01 | 0.33 | 0.24 | -0.33 | -0.17 | 0.59 |
| **BAZ1A** | **S1371** | 0.23 | -0.25 | 0.21 | -0.24 | -0.04 | 0.00 | 0.417 |
| **BAZ1A** | **T731** | 0.03 | 0.13 | 0.34 | -0.35 | 0.04 | -0.29 | 0.073 |
| **BAZ1A** | **T1547** | 0.15 | 0.13 | 0.52 | -0.16 | -0.48 | -0.41 | 0.018 |
| **BAZ1B** | **S152; S158; S161** | -0.14 | -0.25 | -0.35 | -0.18 | 0.31 | 0.42 | 0.094 |
| **BAZ1B** | **T710; S716; S** | -0.19 | -0.27 | -0.24 | 0.02 | 0.21 | 0.35 | 0.013 |
| **BAZ1B** | **S361** | -0.11 | -0.20 | -0.28 | 0.15 | 0.12 | 0.25 | 0.004 |
| **BAZ1B** | **S** | 0.16 | -0.30 | -0.25 | 0.24 | -0.24 | 0.27 | 0.373 |
| **BAZ1B** | **S** | 0.16 | -0.30 | -0.25 | 0.24 | -0.24 | 0.27 | 0.373 |
| **BAZ1B** | **T833** | 0.15 | -0.24 | -0.29 | -0.18 | 0.42 | 0.01 | 0.406 |
| **BAZ1B** | **S1468** | -0.14 | -0.17 | 0.01 | -0.23 | -0.16 | 0.53 | 0.584 |
| **BAZ1B** | **S1342** | 0.04 | -0.15 | -0.08 | 0.00 | -0.03 | 0.19 | 0.264 |
| **BAZ1B** | **S152; S158; S161** | 0.13 | -0.13 | -0.19 | 0.16 | 0.02 | -0.02 | 0.381 |
| **BAZ1B** | **S361** | -0.13 | 0.14 | -0.11 | -0.01 | -0.17 | 0.24 | 0.734 |
| **BAZ1B** | **S329** | -0.08 | -0.03 | 0.02 | -0.16 | 0.27 | -0.07 | 0.758 |
| **BAZ1B** | **S** | -0.10 | -0.08 | 0.13 | -0.04 | 0.23 | -0.17 | 0.896 |
| **BAZ1B** | **S1468** | -0.03 | 0.02 | -0.01 | -0.11 | -0.03 | 0.15 | 0.919 |
| **BAZ1B** | **S** | -0.09 | -0.27 | 0.25 | -0.46 | 0.51 | -0.18 | 0.987 |
| **BAZ1B** | **S1342** | 0.08 | -0.06 | 0.02 | -0.06 | 0.05 | -0.03 | 0.599 |
| **BAZ1B** | **S330** | 0.01 | 0.08 | -0.06 | -0.17 | 0.24 | -0.15 | 0.834 |
| **BAZ1B** | **S705; S708** | 0.03 | -0.08 | 0.10 | -0.15 | 0.01 | 0.07 | 0.634 |
| **BAZ1B** | **S705; S/T** | 0.08 | -0.06 | 0.03 | -0.11 | -0.11 | 0.15 | 0.677 |
| **BAZ1B** | **S705; S708** | 0.04 | -0.05 | 0.09 | -0.10 | -0.04 | 0.05 | 0.415 |
| **BAZ1B** | **S158** | 0.07 | 0.04 | -0.01 | -0.17 | 0.02 | 0.05 | 0.416 |
| **BAZ1B** | **S947** | 0.06 | 0.07 | 0.04 | -0.13 | 0.05 | -0.10 | 0.101 |

| | | | | | | | | |
|---|---|---|---|---|---|---|---|---|
| BAZ1B | S705; S708; T710 | 0.08 | 0.09 | 0.07 | -0.15 | 0.08 | -0.19 | 0.12 |
| BAZ1B | S1124 | 0.20 | 0.00 | 0.18 | -0.20 | -0.06 | -0.17 | 0.025 |
| BAZ1B | S | 0.24 | 0.09 | 0.04 | -0.14 | 0.09 | -0.42 | 0.152 |
| BAZ1B | S705; S708 | 0.06 | -0.05 | 0.39 | -0.28 | -0.03 | -0.19 | 0.12 |
| BAZ1B | S705; S708 | 0.29 | 0.10 | 0.07 | -0.33 | 0.03 | -0.26 | 0.058 |
| BAZ1B | S158; S | 0.19 | 0.15 | 0.18 | -0.27 | 0.06 | -0.42 | 0.056 |
| BAZ1B | S705; S716 | 0.29 | 0.35 | -0.12 | -0.27 | 0.09 | -0.54 | 0.154 |
| BAZ1B | S349 | 0.25 | 0.18 | 0.14 | -0.24 | -0.09 | -0.34 | 0.006 |
| BAZ1B | S158 | 0.27 | 0.27 | 0.11 | -0.36 | -0.01 | -0.43 | 0.027 |
| BAZ1B | S158 | 0.26 | 0.20 | 0.19 | -0.30 | -0.05 | -0.44 | 0.015 |
| BAZ1B | S | 0.22 | 0.27 | 0.20 | -0.20 | -0.06 | -0.61 | 0.034 |
| BAZ1B | S312 | 0.34 | 0.34 | -0.05 | -0.62 | 0.21 | -0.52 | 0.146 |
| BAZ1B | S708 | 0.30 | 0.12 | 0.48 | -0.16 | -0.34 | -0.76 | 0.025 |
| BAZ1B | S708 | 0.37 | 0.37 | 0.21 | -0.29 | -0.24 | -0.76 | 0.013 |
| BAZ1B | S708 | 0.40 | 0.42 | 0.37 | -0.64 | -0.17 | -0.93 | 0.012 |
| BAZ1B | S708 | 0.44 | 0.50 | 0.38 | -0.60 | -0.31 | -1.11 | 0.009 |
| BAZ2A | S135 | -0.16 | -0.20 | -0.43 | 0.44 | -0.40 | 0.47 | 0.22 |
| BAZ2A | T1050; S1051 | -0.28 | -0.27 | -0.13 | 0.18 | 0.04 | 0.35 | 0.015 |
| BAZ2A | S/T | -0.21 | -0.08 | -0.27 | 0.10 | -0.36 | 0.59 | 0.348 |
| BAZ2A | S509 | -0.14 | -0.22 | -0.08 | 0.35 | -0.13 | 0.14 | 0.138 |
| BAZ2A | S1542 | -0.01 | -0.12 | -0.24 | 0.08 | 0.00 | 0.25 | 0.078 |
| BAZ2A | S1051 | -0.11 | -0.07 | -0.07 | 0.03 | -0.06 | 0.25 | 0.165 |
| BAZ2A | S1051 | -0.09 | -0.07 | -0.07 | 0.09 | -0.03 | 0.16 | 0.056 |
| BAZ2A | S1770 | 0.04 | -0.15 | -0.09 | 0.02 | -0.02 | 0.16 | 0.2 |
| BAZ2A | S496; S509 | 0.10 | -0.09 | -0.17 | 0.02 | -0.15 | 0.24 | 0.534 |
| BAZ2A | T1621 | -0.03 | 0.03 | -0.13 | 0.09 | -0.21 | 0.20 | 0.625 |
| BAZ2A | T548 | 0.06 | -0.10 | -0.03 | 0.02 | 0.21 | -0.19 | 0.81 |
| BAZ2A | S/Y/T | -0.13 | 0.14 | -0.08 | 0.02 | 0.10 | -0.08 | 0.761 |
| BAZ2A | S1778; S1783 | -0.04 | -0.10 | 0.10 | -0.06 | 0.05 | 0.05 | 0.687 |
| BAZ2A | S1397 | 0.07 | -0.07 | -0.02 | 0.02 | 0.08 | -0.09 | 0.843 |
| BAZ2A | S1559 | 0.10 | 0.00 | -0.11 | -0.04 | 0.05 | 0.00 | 0.903 |
| BAZ2A | S1397 | -0.11 | 0.23 | -0.15 | -0.08 | 0.10 | -0.03 | 0.974 |
| BAZ2A | S1397 | 0.03 | 0.02 | -0.04 | 0.05 | 0.03 | -0.11 | 0.806 |
| BAZ2A | S1397 | 0.10 | 0.01 | 0.07 | -0.01 | -0.15 | -0.03 | 0.073 |
| BAZ2A | S1184 | 0.01 | 0.13 | 0.18 | -0.48 | -0.06 | 0.11 | 0.239 |
| BAZ2A | S1783 | 0.29 | 0.17 | 0.16 | -0.24 | -0.14 | -0.35 | 0.003 |
| BAZ2A | S1783 | 0.25 | 0.08 | 0.39 | -0.11 | -0.33 | -0.47 | 0.017 |
| BAZ2B | T1283 | -0.51 | -0.25 | 0.11 | 0.33 | -0.14 | 0.29 | 0.184 |
| BAZ2B | T1283 | -0.18 | -0.14 | -0.04 | 0.11 | 0.10 | 0.12 | 0.005 |
| BAZ2B | S1680 | 0.02 | -0.11 | 0.02 | 0.06 | -0.02 | 0.02 | 0.383 |
| BAZ2B | T2014 | -0.08 | 0.13 | 0.18 | -0.23 | -0.01 | -0.03 | 0.191 |
| BBX | S40 | -0.56 | -0.04 | -0.13 | 0.16 | 0.03 | 0.37 | 0.085 |
| BBX | S159 | -0.21 | -0.01 | 0.16 | 0.11 | -0.27 | 0.17 | 0.888 |
| BBX | S844 | -0.05 | 0.18 | 0.09 | 0.01 | -0.17 | -0.09 | 0.132 |
| BBX | S/Y/T | 0.35 | 0.41 | -0.24 | -0.39 | -0.31 | -0.04 | 0.147 |
| BBX | S844 | 0.21 | 0.37 | 0.18 | -0.22 | 0.03 | -0.87 | 0.092 |
| BCAR1 | S355 | -0.05 | 0.06 | -0.03 | 0.01 | -0.22 | 0.20 | 0.957 |
| BCAR3 | S338 | 0.04 | -0.27 | -0.58 | 0.26 | -0.04 | 0.38 | 0.097 |

| | | | | | | | | |
|---|---|---|---|---|---|---|---|---|
| **BCAR3** | **S338** | -0.42 | -0.35 | 0.17 | 0.19 | -0.03 | 0.28 | 0.172 |
| **BCKDHA** | **S337** | 0.07 | 0.14 | 0.11 | -0.11 | -0.29 | 0.05 | 0.094 |
| **BCKDHA** | **S347** | 0.24 | 0.35 | -0.21 | -0.08 | -0.23 | -0.20 | 0.176 |
| **BCKDK** | **S31** | 0.05 | 0.12 | 0.13 | -0.13 | -0.07 | -0.13 | 0.003 |
| **BCKDK** | **S31** | 0.23 | 0.25 | 0.10 | -0.28 | -0.20 | -0.19 | 0.002 |
| **BCL2L12** | **S273** | -0.15 | -0.12 | 0.04 | 0.10 | -0.05 | 0.15 | 0.161 |
| **BCL2L13** | **S291** | -0.18 | 0.04 | -0.28 | -0.04 | 0.15 | 0.24 | 0.11 |
| **BCL2L13** | **S/T** | -0.19 | 0.31 | -0.04 | -0.19 | -0.14 | 0.17 | 0.697 |
| **BCL2L13** | **T374; S** | 0.00 | 0.12 | 0.01 | 0.03 | -0.37 | 0.15 | 0.552 |
| **BCL2L13** | **S420** | 0.03 | 0.12 | 0.17 | -0.17 | -0.13 | -0.07 | 0.009 |
| **BCL2L13** | **S303** | 0.11 | 0.29 | 0.31 | -0.44 | -0.27 | -0.18 | 0.006 |
| **BCL2L13** | **S291** | 0.05 | 0.31 | 0.36 | -0.42 | -0.16 | -0.32 | 0.011 |
| **BCL2L13** | **S420; S426** | 0.18 | 0.52 | 0.16 | -0.48 | -0.85 | 0.06 | 0.071 |
| **BCL3** | **S41** | -0.17 | -0.13 | -0.28 | 0.22 | -0.08 | 0.33 | 0.054 |
| **BCL3** | **S374** | 0.01 | 0.27 | 0.30 | -0.41 | -0.07 | -0.23 | 0.034 |
| **BCL6** | **S** | -0.36 | -0.01 | -0.08 | 0.13 | -0.07 | 0.30 | 0.153 |
| **BCL7A** | **S173** | -0.23 | -0.18 | -0.22 | 0.07 | -0.07 | 0.49 | 0.093 |
| **BCL7B** | **T/S** | -0.25 | 0.14 | -0.34 | -0.10 | 0.33 | 0.11 | 0.24 |
| **BCL7B** | **S134** | -0.17 | 0.05 | 0.10 | 0.27 | -0.22 | -0.09 | 0.971 |
| **BCL7B** | **S122** | 0.45 | -0.30 | -0.10 | 0.03 | -0.03 | -0.16 | 0.778 |
| **BCL7C** | **S114; T118; S126** | -0.39 | -0.30 | -0.39 | 0.15 | 0.09 | 0.57 | 0.015 |
| **BCL7C** | **S/T** | -0.36 | -0.16 | -0.40 | 0.33 | -0.23 | 0.55 | 0.096 |
| **BCL7C** | **S114; S126** | -0.19 | -0.11 | -0.14 | 0.05 | 0.04 | 0.29 | 0.029 |
| **BCL7C** | **S114; S126** | -0.10 | -0.30 | 0.13 | 0.15 | 0.08 | -0.01 | 0.291 |
| **BCL7C** | **S126** | -0.03 | -0.06 | -0.08 | 0.01 | -0.03 | 0.17 | 0.163 |
| **BCL7C** | **S126** | 0.02 | 0.06 | 0.04 | -0.08 | -0.08 | 0.03 | 0.091 |
| **BCL9L** | **S21** | -0.14 | -0.07 | -0.28 | 0.22 | -0.06 | 0.26 | 0.064 |
| **BCL9L** | **S88** | 0.10 | -0.03 | -0.36 | 0.03 | 0.10 | 0.10 | 0.267 |
| **BCL9L** | **S88** | 0.13 | 0.08 | -0.51 | -0.09 | 0.03 | 0.25 | 0.522 |
| **BCLAF1** | **T494; S496** | -0.76 | -0.58 | -0.47 | 0.27 | 0.40 | 0.57 | 0.001 |
| **BCLAF1** | **S198** | -0.07 | -0.03 | -0.21 | 0.12 | 0.06 | 0.12 | 0.024 |
| **BCLAF1** | **S658** | -0.04 | -0.16 | -0.10 | 0.11 | -0.03 | 0.19 | 0.064 |
| **BCLAF1** | **S385; S389** | -0.12 | 0.08 | -0.28 | 0.04 | -0.18 | 0.37 | 0.383 |
| **BCLAF1** | **S319** | -0.06 | -0.03 | -0.16 | 0.22 | 0.07 | -0.07 | 0.157 |
| **BCLAF1** | **S397; T402** | -0.18 | -0.10 | 0.05 | -0.12 | 0.16 | 0.16 | 0.274 |
| **BCLAF1** | **S422** | -0.12 | -0.08 | -0.03 | 0.04 | -0.03 | 0.20 | 0.113 |
| **BCLAF1** | **S512** | 0.09 | 0.00 | -0.30 | 0.15 | -0.03 | 0.04 | 0.391 |
| **BCLAF1** | **S290** | -0.07 | -0.02 | -0.06 | -0.01 | 0.03 | 0.11 | 0.088 |
| **BCLAF1** | **S385** | 0.01 | 0.01 | -0.14 | -0.02 | -0.01 | 0.13 | 0.354 |
| **BCLAF1** | **S385; S389** | 0.07 | -0.01 | -0.16 | 0.04 | -0.09 | 0.13 | 0.573 |
| **BCLAF1** | **T840** | -0.03 | -0.01 | 0.01 | 0.09 | 0.03 | -0.10 | 0.82 |
| **BCLAF1** | **S385** | 0.03 | -0.09 | 0.03 | -0.01 | -0.17 | 0.18 | 0.934 |
| **BCLAF1** | **S496** | -0.09 | -0.02 | 0.08 | 0.23 | 0.02 | -0.27 | 0.978 |
| **BCLAF1** | **S183** | 0.06 | 0.01 | -0.07 | -0.07 | 0.01 | 0.06 | 0.992 |
| **BCLAF1** | **S385** | 0.08 | 0.04 | -0.11 | 0.05 | -0.02 | -0.05 | 0.89 |
| **BCLAF1** | **S648** | 0.07 | -0.01 | -0.02 | -0.02 | 0.12 | -0.16 | 0.729 |
| **BCLAF1** | **S177** | 0.13 | 0.05 | -0.06 | 0.01 | -0.05 | -0.09 | 0.24 |
| **BCLAF1** | **S177** | 0.07 | 0.07 | 0.04 | -0.03 | -0.07 | -0.08 | 0.002 |

| | | | | | | | | |
|---|---|---|---|---|---|---|---|---|
| BCLAF1 | S450 | 0.06 | 0.09 | 0.03 | -0.10 | -0.03 | -0.06 | 0.009 |
| BCLAF1 | S285; S290 | -0.04 | 0.07 | 0.16 | 0.04 | -0.24 | -0.02 | 0.255 |
| BCLAF1 | S177 | 0.09 | 0.10 | 0.00 | -0.03 | -0.02 | -0.15 | 0.058 |
| BCLAF1 | S102; S104 | 0.02 | 0.08 | 0.12 | -0.05 | -0.02 | -0.18 | 0.047 |
| BCLAF1 | S762; S763 | 0.12 | 0.16 | -0.07 | -0.21 | 0.26 | -0.34 | 0.434 |
| BCLAF1 | S/T | 0.15 | 0.10 | -0.01 | -0.02 | 0.04 | -0.31 | 0.218 |
| BCLAF1 | S290 | 0.04 | 0.12 | 0.09 | -0.04 | -0.13 | -0.10 | 0.008 |
| BCLAF1 | S198 | 0.08 | 0.15 | 0.03 | -0.06 | 0.05 | -0.29 | 0.143 |
| BCLAF1 | S496 | 0.10 | 0.14 | 0.05 | -0.07 | -0.04 | -0.21 | 0.029 |
| BCLAF1 | S320 | 0.14 | 0.14 | 0.08 | -0.15 | -0.01 | -0.25 | 0.023 |
| BCLAF1 | S414 | 0.14 | 0.26 | -0.05 | -0.02 | 0.02 | -0.46 | 0.209 |
| BCLAF1 | S319 | 0.14 | 0.24 | 0.00 | -0.13 | -0.11 | -0.19 | 0.02 |
| BCLAF1 | S496 | 0.18 | 0.15 | 0.05 | -0.12 | 0.03 | -0.37 | 0.083 |
| BCLAF1 | S/Y/T | -0.02 | 0.36 | 0.02 | -0.45 | 0.18 | -0.22 | 0.264 |
| BCLAF1 | S496 | 0.19 | 0.13 | 0.07 | -0.15 | 0.05 | -0.37 | 0.086 |
| BCLAF1 | S884 | 0.22 | 0.12 | 0.05 | -0.19 | 0.17 | -0.50 | 0.207 |
| BCLAF1 | S658 | 0.12 | 0.15 | 0.15 | -0.13 | 0.05 | -0.43 | 0.087 |
| BCLAF1 | S183 | 0.18 | 0.13 | 0.17 | -0.10 | -0.21 | -0.22 | 0.001 |
| BCLAF1 | S397 | 0.02 | 0.22 | 0.21 | -0.34 | 0.12 | -0.35 | 0.118 |
| BCLAF1 | S285; S290 | 0.09 | 0.18 | 0.22 | -0.16 | -0.10 | -0.31 | 0.008 |
| BCLAF1 | S222 | 0.20 | 0.21 | 0.11 | -0.19 | -0.01 | -0.41 | 0.034 |
| BCLAF1 | S177 | 0.05 | 0.14 | 0.33 | -0.22 | 0.09 | -0.55 | 0.12 |
| BCLAF1 | S531 | 0.14 | 0.17 | 0.26 | -0.24 | 0.00 | -0.44 | 0.034 |
| BCLAF1 | S760 | 0.17 | 0.15 | 0.31 | -0.27 | -0.01 | -0.51 | 0.034 |
| BCLAF1 | S397 | 0.24 | 0.17 | 0.20 | -0.16 | 0.16 | -0.94 | 0.188 |
| BCLAF1 | S276 | 0.26 | 0.31 | 0.30 | -0.33 | 0.03 | -0.97 | 0.071 |
| BCLAF1 | S397 | 0.12 | 0.70 | 0.12 | -0.77 | 0.26 | -1.24 | 0.137 |
| BCLAF3 | S139 | 0.01 | -0.12 | -0.07 | 0.11 | -0.10 | 0.14 | 0.243 |
| BCLAF3 | S402 | 0.10 | -0.02 | -0.08 | 0.13 | -0.06 | -0.08 | 0.989 |
| BCOR | S/T | -0.20 | -0.05 | 0.03 | 0.04 | -0.16 | 0.29 | 0.434 |
| BCOR | S1410 | 0.14 | -0.07 | -0.01 | 0.02 | -0.12 | 0.03 | 0.623 |
| BCOR | S/T | -0.06 | 0.15 | -0.01 | 0.14 | -0.16 | -0.09 | 0.578 |
| BCOR | S423 | 0.09 | 0.11 | 0.13 | 0.02 | -0.13 | -0.26 | 0.045 |
| BCOR | S423 | -0.01 | 0.11 | 0.26 | -0.11 | -0.22 | -0.08 | 0.044 |
| BCOR | S1139 | 0.24 | 0.19 | -0.05 | 0.03 | -0.35 | -0.13 | 0.123 |
| BCOR | S1127 | 0.00 | 0.14 | 0.26 | -0.15 | -0.15 | -0.16 | 0.018 |
| BCOR | S1410 | 0.32 | 0.08 | 0.08 | -0.24 | -0.20 | -0.14 | 0.015 |
| BCOR | T1129 | 0.13 | 0.19 | 0.19 | -0.09 | -0.47 | -0.06 | 0.049 |
| BCORL1 | S155 | -0.18 | -0.05 | -0.16 | 0.16 | 0.27 | -0.09 | 0.097 |
| BCORL1 | S1033 | 0.02 | -0.23 | 0.08 | 0.00 | 0.25 | -0.18 | 0.682 |
| BCORL1 | S1033 | -0.06 | -0.42 | 0.39 | -0.26 | -0.07 | 0.25 | 0.992 |
| BCORL1 | S1753 | 0.03 | -0.10 | 0.11 | -0.02 | 0.15 | -0.20 | 0.777 |
| BCORL1 | S1450 | 0.09 | 0.18 | -0.11 | -0.06 | -0.22 | 0.08 | 0.381 |
| BCORL1 | S1192 | 0.30 | -0.18 | 0.04 | -0.52 | 0.22 | -0.01 | 0.57 |
| BCR | S354 | 0.38 | 0.05 | 0.07 | -0.26 | -0.01 | -0.34 | 0.067 |
| BDP1 | T1025 | -0.06 | -0.31 | -0.21 | 0.19 | -0.05 | 0.34 | 0.057 |
| BDP1 | T1131 | -0.10 | -0.10 | -0.26 | 0.17 | 0.11 | 0.14 | 0.007 |
| BDP1 | S/T | -0.41 | 0.17 | -0.35 | 0.15 | -0.50 | 0.61 | 0.487 |

| | | | | | | | | |
|---|---|---|---|---|---|---|---|---|
| BDP1 | S1524 | -0.23 | 0.13 | 0.03 | 0.06 | -0.11 | 0.10 | 0.777 |
| BDP1 | S1781 | -0.10 | 0.07 | -0.08 | -0.38 | -0.09 | 0.44 | 0.93 |
| BDP1 | S | 0.09 | 0.01 | 0.32 | -0.85 | -0.10 | 0.25 | 0.333 |
| BEND3 | S164 | 0.04 | -0.06 | 0.07 | -0.08 | -0.07 | 0.09 | 0.64 |
| BET1 | S50 | 0.09 | 0.15 | -0.04 | 0.01 | -0.03 | -0.20 | 0.166 |
| BET1L | S9 | 0.03 | 0.17 | -0.23 | 0.05 | -0.22 | 0.14 | 0.994 |
| BHLHE40 | S207 | 0.24 | 0.13 | -0.16 | -0.10 | -0.08 | -0.08 | 0.266 |
| BICD2 | S582 | 0.18 | -0.18 | 0.22 | 0.32 | -0.72 | -0.05 | 0.54 |
| BICDL2 | S36 | -0.17 | -0.19 | -0.14 | 0.08 | 0.18 | 0.18 | 0.001 |
| BICDL2 | T/S | -0.09 | -0.30 | 0.02 | -0.07 | 0.09 | 0.29 | 0.186 |
| BICRA | S1256 | -0.21 | -0.28 | -0.13 | 0.37 | -0.14 | 0.27 | 0.085 |
| BICRA | S1413 | 0.14 | -0.05 | -0.12 | 0.04 | -0.10 | 0.07 | 0.896 |
| BICRA | T921 | -0.03 | -0.11 | 0.13 | 0.02 | -0.06 | 0.04 | 0.941 |
| BICRAL | S934 | -0.09 | 0.07 | 0.10 | -0.20 | -0.05 | 0.13 | 0.577 |
| BICRAL | S623 | 0.05 | -0.02 | 0.11 | 0.01 | -0.19 | 0.03 | 0.285 |
| BICRAL | S | 0.22 | 0.12 | 0.07 | -0.18 | -0.15 | -0.14 | 0.003 |
| BLM | S304 | -0.17 | 0.16 | 0.01 | 0.03 | -0.18 | 0.11 | 0.913 |
| BLTP1 | T3996 | -0.02 | -0.23 | -0.20 | 0.29 | 0.29 | -0.24 | 0.236 |
| BLTP1 | S2755 | -0.10 | -0.04 | -0.11 | 0.02 | 0.12 | 0.10 | 0.01 |
| BLTP1 | S | 0.00 | -0.09 | 0.30 | -0.15 | 0.03 | -0.13 | 0.3 |
| BLTP1 | S3562 | -0.17 | 0.14 | 0.22 | -0.07 | 0.23 | -0.49 | 0.512 |
| BLTP1 | S2368 | 0.25 | 0.44 | -0.39 | -0.59 | 0.01 | 0.01 | 0.419 |
| BLTP1 | S2603 | 0.66 | 0.02 | 0.01 | -0.32 | -0.25 | -0.39 | 0.066 |
| BLTP3B | S989 | 0.14 | 0.31 | 0.33 | -0.29 | -0.15 | -0.55 | 0.011 |
| BLVRB | S/T | 0.03 | -0.08 | 0.11 | -0.02 | 0.08 | -0.14 | 0.625 |
| BLZF1 | S/T | -0.14 | 0.03 | 0.20 | -0.05 | -0.01 | -0.06 | 0.52 |
| BLZF1 | S15 | 0.09 | 0.18 | 0.25 | -0.25 | -0.01 | -0.36 | 0.027 |
| BMAL1 | T527 | -0.22 | -0.22 | -0.07 | 0.04 | 0.14 | 0.25 | 0.016 |
| BMAL1 | T527 | 0.05 | -0.14 | -0.04 | 0.00 | -0.10 | 0.19 | 0.516 |
| BMAL1 | S512; S513 | -0.12 | -0.17 | 0.38 | 0.29 | -0.57 | -0.01 | 0.703 |
| BMAL1 | S512; S513 | 0.20 | 0.02 | 0.28 | 0.25 | -0.29 | -0.70 | 0.224 |
| BMI1 | S251 | -0.05 | -0.08 | 0.09 | -0.04 | -0.03 | 0.10 | 0.737 |
| BMI1 | S316 | 0.08 | -0.05 | 0.04 | 0.07 | -0.10 | -0.05 | 0.441 |
| BMI1 | S253; S255 | 0.02 | -0.05 | 0.14 | -0.14 | 0.03 | -0.02 | 0.33 |
| BMI1 | S/T | 0.39 | 0.18 | 0.16 | -0.25 | -0.17 | -0.50 | 0.012 |
| BMP2K | S1029 | -0.13 | -0.02 | -0.20 | 0.18 | 0.15 | -0.01 | 0.045 |
| BMP2K | S1107; S1111 | -0.08 | -0.22 | 0.12 | -0.32 | 0.22 | 0.19 | 0.657 |
| BMP2K | S/T | -0.01 | -0.04 | -0.06 | 0.23 | -0.10 | -0.05 | 0.571 |
| BMP2K | T1000 | 0.23 | 0.24 | 0.35 | -0.39 | -0.09 | -0.57 | 0.013 |
| BMS1 | S731 | -0.84 | -1.13 | -0.48 | 0.57 | 0.36 | 0.59 | 0.003 |
| BMS1 | S820 | -0.27 | -0.28 | -0.31 | 0.15 | 0.22 | 0.34 | 7E-04 |
| BMS1 | S552 | 0.04 | -0.05 | -0.05 | 0.02 | 0.04 | 0.00 | 0.299 |
| BNIP2 | S114 | -0.03 | 0.09 | -0.12 | 0.14 | 0.03 | -0.13 | 0.737 |
| BNIP3 | S66 | -0.11 | -0.20 | -0.03 | 0.22 | 0.06 | 0.01 | 0.063 |
| BNIP3 | S86 | -0.04 | -0.03 | -0.02 | 0.05 | -0.10 | 0.13 | 0.46 |
| BNIP3 | S92; S/T | 0.07 | -0.09 | -0.02 | 0.11 | -0.12 | 0.04 | 0.788 |
| BOD1L1 | S482; S484 | 0.01 | -0.15 | -0.19 | 0.20 | -0.41 | 0.40 | 0.533 |
| BOD1L1 | S482 | -0.13 | -0.11 | 0.08 | 0.02 | -0.08 | 0.20 | 0.394 |

| | | | | | | | | |
|---|---|---|---|---|---|---|---|---|
| BOD1L1 | S1531 | 0.09 | 0.00 | 0.08 | -0.07 | -0.04 | -0.07 | 0.016 |
| BOD1L1 | S2986 | 0.37 | 0.16 | 0.07 | -0.30 | 0.01 | -0.49 | 0.055 |
| BOK | S | 0.04 | 0.15 | 0.14 | 0.03 | -0.06 | -0.36 | 0.116 |
| BOP1 | S126; S127 | -0.01 | 0.15 | 0.16 | -0.25 | 0.15 | -0.26 | 0.206 |
| BPTF | S817 | -1.82 | -1.79 | -1.89 | 0.56 | 0.75 | 1.00 | 4E-05 |
| BPTF | S1827 | -0.31 | -0.35 | -0.43 | 0.26 | 0.60 | -0.07 | 0.033 |
| BPTF | S1559 | -0.09 | -0.35 | -0.46 | 0.06 | 0.43 | 0.21 | 0.027 |
| BPTF | S1617 | -0.27 | -0.26 | -0.41 | 0.10 | -0.37 | 0.80 | 0.224 |
| BPTF | S1310 | -0.18 | -0.46 | -0.12 | 0.10 | 0.42 | 0.08 | 0.04 |
| BPTF | T739 | -0.21 | -0.14 | -0.30 | -0.04 | 0.12 | 0.44 | 0.058 |
| BPTF | T739; S742 | -0.24 | -0.15 | -0.11 | 0.14 | -0.01 | 0.30 | 0.034 |
| BPTF | S572 | -0.02 | -0.07 | -0.39 | 0.17 | -0.15 | 0.35 | 0.195 |
| BPTF | S1310 | 0.02 | -0.58 | 0.05 | -0.07 | 0.15 | 0.28 | 0.275 |
| BPTF | T739; S742 | 0.04 | -0.34 | -0.17 | -0.09 | 0.19 | 0.27 | 0.151 |
| BPTF | S763 | -0.14 | -0.29 | -0.02 | 0.03 | -0.06 | 0.39 | 0.158 |
| BPTF | T739 | -0.05 | -0.07 | -0.17 | 0.21 | -0.08 | 0.13 | 0.126 |
| BPTF | S572 | 0.03 | -0.10 | -0.23 | -0.06 | -0.09 | 0.37 | 0.352 |
| BPTF | T739; S742 | 0.03 | -0.06 | -0.22 | 0.05 | -0.05 | 0.21 | 0.229 |
| BPTF | S742 | -0.03 | -0.30 | 0.09 | 0.01 | 0.00 | 0.18 | 0.334 |
| BPTF | S216 | 0.00 | -0.07 | -0.12 | 0.07 | -0.03 | 0.13 | 0.103 |
| BPTF | T739 | 0.02 | -0.02 | -0.20 | 0.27 | -0.02 | -0.10 | 0.419 |
| BPTF | S650 | 0.03 | 0.07 | -0.17 | -0.19 | 0.23 | -0.01 | 0.815 |
| BPTF | S572 | 0.05 | -0.11 | 0.01 | -0.05 | -0.03 | 0.12 | 0.688 |
| BPTF | S1251 | -0.15 | 0.17 | -0.04 | -0.14 | -0.30 | 0.35 | 0.927 |
| BPTF | S1251 | -0.09 | 0.08 | 0.03 | -0.21 | 0.10 | 0.07 | 0.858 |
| BPTF | S/T | -0.03 | -0.02 | 0.11 | -0.13 | -0.16 | 0.20 | 0.702 |
| BPTF | S650 | 0.19 | 0.10 | -0.27 | 0.11 | 0.21 | -0.46 | 0.845 |
| BPTF | S1300 | 0.06 | 0.04 | -0.02 | -0.12 | 0.09 | -0.06 | 0.412 |
| BPTF | S2370 | 0.07 | -0.08 | 0.10 | -0.03 | 0.18 | -0.28 | 0.649 |
| BPTF | S2370 | 0.06 | 0.06 | 0.00 | -0.05 | 0.06 | -0.14 | 0.231 |
| BPTF | S/T | 0.04 | 0.05 | 0.13 | -0.07 | 0.13 | -0.33 | 0.294 |
| BPTF | S2471 | 0.02 | 0.02 | 0.19 | -0.11 | 0.03 | -0.19 | 0.121 |
| BPTF | S719 | 0.07 | 0.17 | 0.08 | -0.16 | -0.01 | -0.19 | 0.023 |
| BPTF | S/T | -0.13 | 0.42 | -0.03 | 0.37 | -0.63 | -0.25 | 0.489 |
| BPTF | S1231 | 0.15 | 0.14 | 0.14 | -0.30 | 0.13 | -0.35 | 0.11 |
| BPTF | S763 | 0.20 | 0.16 | 0.30 | -0.41 | -0.04 | -0.39 | 0.017 |
| BPTF | S1673 | 0.13 | 0.24 | 0.36 | -0.39 | -0.20 | -0.30 | 0.004 |
| BRAF | S446 | -0.03 | -0.27 | 0.21 | -0.02 | -0.41 | 0.37 | 0.98 |
| BRAP | S117 | 0.04 | 0.02 | -0.17 | -0.10 | 0.14 | 0.06 | 0.513 |
| BRAT1 | S742 | -0.20 | -0.52 | -0.50 | 0.39 | -0.03 | 0.52 | 0.024 |
| BRCA1 | S1524 | -0.40 | -0.22 | -0.19 | 0.01 | 0.17 | 0.46 | 0.029 |
| BRCA1 | S1542 | -0.28 | -0.04 | 0.27 | 0.12 | -0.14 | 0.02 | 0.936 |
| BRCA1 | S114 | 0.02 | -0.05 | 0.09 | -0.14 | 0.01 | 0.06 | 0.596 |
| BRCA2 | T/S | 0.04 | 0.24 | 0.42 | -0.54 | 0.02 | -0.42 | 0.054 |
| BRD1 | S1055; T/S | -0.66 | -0.32 | -0.05 | 0.12 | 0.11 | 0.52 | 0.057 |
| BRD1 | S18 | -0.20 | -0.23 | -0.06 | 0.11 | -0.24 | 0.48 | 0.26 |
| BRD1 | S128 | 0.05 | 0.01 | 0.05 | -0.02 | 0.01 | -0.10 | 0.103 |
| BRD1 | S852 | -0.07 | 0.05 | 0.20 | -0.02 | -0.09 | -0.09 | 0.192 |

| | | | | | | | | |
|---|---|---|---|---|---|---|---|---|
| **BRD1** | **S696** | 0.47 | -0.22 | 0.11 | 0.03 | 0.03 | -0.66 | 0.35 |
| **BRD1** | **T1051** | 0.15 | 0.11 | 0.25 | -0.18 | -0.02 | -0.42 | 0.037 |
| **BRD2** | **S/T** | 0.13 | -0.05 | 0.04 | 0.07 | -0.02 | -0.19 | 0.412 |
| **BRD2** | **S301** | 0.25 | -0.15 | -0.04 | 0.30 | 0.02 | -0.53 | 0.745 |
| **BRD2** | **S633** | 0.26 | 0.15 | -0.02 | -0.38 | 0.00 | -0.09 | 0.114 |
| **BRD2** | **S633** | 0.16 | 0.20 | 0.04 | -0.08 | -0.03 | -0.35 | 0.058 |
| **BRD3** | **S560; S563** | -0.21 | -0.14 | -0.28 | 0.21 | 0.11 | 0.23 | 0.002 |
| **BRD3** | **S558; S563** | -0.12 | 0.06 | -0.08 | 0.44 | -0.01 | -0.45 | 0.885 |
| **BRD3** | **T250** | 0.01 | -0.04 | 0.02 | 0.20 | -0.19 | -0.03 | 0.988 |
| **BRD3** | **T250** | 0.26 | -0.24 | 0.15 | 0.24 | -0.23 | -0.29 | 0.537 |
| **BRD3** | **S680** | 0.05 | 0.06 | 0.13 | -0.19 | 0.07 | -0.14 | 0.125 |
| **BRD3** | **S563** | 0.15 | 0.07 | 0.16 | -0.08 | -0.05 | -0.30 | 0.033 |
| **BRD3** | **S563** | 0.21 | 0.09 | 0.15 | -0.18 | 0.04 | -0.40 | 0.069 |
| **BRD3** | **S263** | 0.25 | 0.16 | 0.17 | -0.26 | -0.06 | -0.38 | 0.012 |
| **BRD4** | **S470** | 0.00 | -0.05 | -0.05 | 0.10 | -0.01 | 0.01 | 0.165 |
| **BRD4** | **S/T** | -0.05 | -0.06 | 0.00 | -0.03 | -0.05 | 0.17 | 0.41 |
| **BRD4** | **T296** | 0.03 | -0.06 | 0.00 | -0.10 | 0.08 | 0.04 | 0.781 |
| **BRD4** | **S1126** | -0.04 | -0.02 | 0.03 | -0.15 | 0.07 | 0.09 | 0.881 |
| **BRD4** | **S601** | 0.09 | 0.01 | 0.10 | -0.04 | -0.16 | -0.02 | 0.05 |
| **BRD4** | **S1117** | 0.09 | 0.17 | 0.27 | 0.00 | -0.09 | -0.57 | 0.098 |
| **BRD4** | **S1126** | -0.03 | 0.22 | 0.55 | -0.12 | -0.22 | -0.69 | 0.072 |
| **BRD4** | **S601** | 0.39 | 0.42 | 0.23 | -0.34 | -0.14 | -1.08 | 0.041 |
| **BRD7** | **S/T** | -0.01 | -0.19 | -0.20 | 0.22 | 0.04 | 0.09 | 0.036 |
| **BRD7** | **S/T** | 0.10 | -0.03 | -0.18 | 0.12 | 0.03 | -0.07 | 0.546 |
| **BRD7** | **S279** | 0.01 | 0.01 | -0.11 | 0.04 | -0.02 | 0.07 | 0.293 |
| **BRD7** | **S279; S289** | 0.15 | 0.13 | -0.05 | -0.22 | -0.09 | 0.06 | 0.208 |
| **BRD7** | **S289** | 0.28 | 0.12 | 0.11 | -0.18 | -0.09 | -0.32 | 0.013 |
| **BRD8** | **S387** | 0.10 | -0.08 | -0.09 | 0.08 | -0.15 | 0.11 | 0.752 |
| **BRD9** | **S566** | -0.07 | -0.07 | -0.05 | 0.09 | -0.13 | 0.20 | 0.303 |
| **BRD9** | **T103** | -0.12 | -0.01 | -0.02 | 0.01 | -0.11 | 0.22 | 0.423 |
| **BRD9** | **S588** | 0.21 | 0.09 | 0.11 | 0.01 | -0.28 | -0.21 | 0.036 |
| **BRF1** | **S358; T365** | 0.04 | -0.21 | -0.18 | -0.06 | 0.16 | 0.20 | 0.13 |
| **BRF1** | **T365** | 0.13 | -0.10 | -0.07 | 0.17 | -0.15 | -0.01 | 0.892 |
| **BRF1** | **S553** | 0.22 | -0.18 | 0.01 | -0.23 | -0.38 | 0.40 | 0.76 |
| **BRF1** | **S553** | 0.16 | 0.02 | 0.22 | -0.29 | 0.04 | -0.21 | 0.064 |
| **BRF1** | **S553** | 0.22 | 0.00 | 0.16 | -0.27 | -0.53 | 0.26 | 0.275 |
| **BRIP1** | **S1032** | -0.17 | -0.05 | -0.24 | -0.02 | 0.08 | 0.33 | 0.076 |
| **BRIP1** | **T113** | -0.25 | -0.41 | 0.43 | 0.41 | 0.03 | -0.48 | 0.863 |
| **BRME1** | **S370** | 0.30 | -0.10 | 0.30 | -0.43 | -0.09 | -0.12 | 0.092 |
| **BRMS1L** | **S197** | 0.21 | -0.02 | 0.08 | 0.00 | -0.07 | -0.25 | 0.12 |
| **BRMS1L** | **S197** | 0.19 | 0.13 | 0.03 | -0.16 | 0.00 | -0.23 | 0.041 |
| **BRPF1** | **S844** | -0.28 | -0.20 | -0.28 | 0.07 | 0.08 | 0.46 | 0.026 |
| **BRPF1** | **S460; S462** | -0.03 | -0.18 | -0.25 | 0.05 | -0.03 | 0.36 | 0.104 |
| **BRPF1** | **S77** | -0.08 | -0.24 | 0.02 | -0.12 | 0.25 | 0.11 | 0.24 |
| **BRPF1** | **S452** | -0.12 | -0.03 | 0.09 | -0.12 | 0.01 | 0.14 | 0.751 |
| **BRPF1** | **S460** | 0.06 | -0.07 | -0.03 | -0.09 | 0.05 | 0.08 | 0.703 |
| **BRPF1** | **S460** | 0.04 | 0.02 | -0.01 | 0.04 | -0.08 | -0.02 | 0.395 |
| **BRPF1** | **S120** | -0.03 | 0.04 | 0.06 | -0.02 | 0.03 | -0.10 | 0.311 |

| | | | | | | | | |
|---|---|---|---|---|---|---|---|---|
| **BRPF1** | **S120** | 0.07 | 0.04 | -0.02 | -0.01 | 0.16 | -0.26 | 0.61 |
| **BRPF1** | **S860** | 0.18 | -0.15 | 0.03 | 0.31 | -0.41 | -0.08 | 0.751 |
| **BRPF1** | **S460; S462** | 0.10 | 0.08 | -0.07 | -0.18 | 0.07 | -0.02 | 0.428 |
| **BRPF1** | **S577** | -0.04 | 0.13 | 0.11 | -0.10 | 0.04 | -0.16 | 0.152 |
| **BRPF1** | **T145** | 0.14 | -0.09 | 0.16 | -0.19 | 0.10 | -0.16 | 0.287 |
| **BRPF1** | **S577** | 0.04 | 0.07 | 0.16 | -0.21 | 0.10 | -0.21 | 0.152 |
| **BRPF1** | **S860** | 0.16 | 0.03 | 0.07 | 0.19 | -0.36 | -0.16 | 0.299 |
| **BRPF1** | **S84; S** | 0.17 | 0.02 | 0.10 | -0.07 | -0.53 | 0.19 | 0.335 |
| **BRPF1** | **T1043; S1044; T1046** | 0.02 | 0.31 | 0.28 | -0.15 | -0.28 | -0.32 | 0.012 |
| **BRPF3** | **S900** | -0.07 | -0.27 | -0.34 | -0.01 | 0.26 | 0.30 | 0.03 |
| **BRPF3** | **S886** | -0.21 | -0.11 | -0.02 | 0.29 | -0.05 | 0.05 | 0.143 |
| **BRPF3** | **S962** | -0.07 | -0.10 | -0.09 | 0.13 | -0.11 | 0.21 | 0.168 |
| **BRPF3** | **S740** | -0.02 | -0.10 | -0.12 | 0.03 | 0.06 | 0.14 | 0.024 |
| **BRPF3** | **S798** | -0.15 | 0.02 | -0.01 | 0.16 | 0.11 | -0.16 | 0.525 |
| **BRPF3** | **S387** | 0.16 | -0.22 | -0.05 | -0.15 | -0.02 | 0.23 | 0.751 |
| **BRPF3** | **S713** | 0.05 | -0.01 | -0.10 | 0.04 | -0.06 | 0.07 | 0.522 |
| **BRPF3** | **S959; S962; S965** | 0.02 | 0.04 | 0.08 | -0.12 | 0.03 | -0.06 | 0.114 |
| **BRPF3** | **S17; S19** | -0.01 | 0.07 | 0.12 | 0.01 | -0.10 | -0.09 | 0.083 |
| **BRPF3** | **S962; S965** | 0.04 | 0.04 | 0.10 | -0.25 | -0.09 | 0.13 | 0.314 |
| **BRPF3** | **S959; S962** | 0.04 | 0.06 | 0.13 | -0.25 | -0.28 | 0.23 | 0.356 |
| **BRPF3** | **S713** | 0.20 | 0.17 | 0.00 | -0.25 | -0.14 | -0.04 | 0.039 |
| **BRWD1** | **S1943** | -0.17 | -0.19 | -0.27 | 0.12 | 0.03 | 0.38 | 0.024 |
| **BRWD1** | **S2063** | -0.24 | -0.27 | -0.01 | 0.14 | -0.26 | 0.48 | 0.263 |
| **BRWD1** | **S1289; S/T** | 0.09 | -0.15 | -0.24 | -0.01 | -0.20 | 0.40 | 0.469 |
| **BRWD1** | **S2131** | 0.13 | -0.09 | -0.25 | 0.02 | 0.07 | 0.08 | 0.305 |
| **BRWD1** | **S1475** | 0.07 | -0.35 | 0.07 | -0.28 | 0.05 | 0.33 | 0.666 |
| **BRWD1** | **S696** | 0.39 | -0.17 | -0.38 | -0.26 | 0.26 | 0.01 | 0.841 |
| **BRWD1** | **S1943** | -0.07 | -0.12 | 0.13 | 0.15 | -0.17 | 0.05 | 0.834 |
| **BRWD1** | **T/S** | 0.05 | -0.04 | -0.03 | 0.12 | -0.11 | 0.00 | 0.905 |
| **BRWD1** | **S1678** | 0.11 | -0.02 | -0.05 | -0.06 | 0.08 | -0.07 | 0.711 |
| **BRWD1** | **S2166** | 0.02 | -0.08 | 0.14 | 0.01 | -0.08 | -0.01 | 0.495 |
| **BRWD1** | **S1907** | 0.35 | -0.34 | 0.04 | -0.20 | 0.05 | -0.01 | 0.763 |
| **BRWD1** | **S2158** | -0.04 | -0.04 | 0.19 | -0.02 | -0.11 | 0.00 | 0.401 |
| **BRWD1** | **S/T** | -0.08 | 0.05 | 0.14 | -0.11 | -0.10 | 0.08 | 0.417 |
| **BRWD1** | **S2158; S2162; S2166** | 0.13 | 0.08 | 0.05 | 0.04 | -0.01 | -0.34 | 0.183 |
| **BRWD1** | **S2162; S/T** | 0.11 | 0.11 | 0.11 | -0.31 | -0.43 | 0.28 | 0.296 |
| **BRWD1** | **S1788** | 0.25 | 0.16 | 0.01 | -0.20 | -0.01 | -0.29 | 0.046 |
| **BRWD1** | **S** | 0.01 | 0.24 | 0.18 | -0.37 | -0.36 | 0.17 | 0.155 |
| **BRWD1** | **S2051** | 0.24 | 0.25 | 0.21 | -0.40 | -0.15 | -0.29 | 0.002 |
| **BRWD1** | **S/T** | -0.04 | 0.47 | 0.29 | -0.31 | -0.99 | 0.16 | 0.166 |
| **BRWD3** | **S1763** | 0.10 | -0.16 | -0.01 | -0.20 | 0.04 | 0.19 | 0.83 |
| **BRWD3** | **S1579** | 0.05 | -0.02 | 0.14 | -0.01 | -0.09 | -0.09 | 0.078 |
| **BRWD3** | **S1763** | 0.03 | -0.38 | 0.30 | -1.11 | 0.36 | 0.30 | 0.806 |
| **BRWD3** | **S703** | 0.20 | -0.03 | 0.00 | 0.04 | 0.16 | -0.45 | 0.517 |
| **BRWD3** | **S884; S885; S886** | 0.05 | 0.10 | 0.16 | -0.13 | -0.04 | -0.18 | 0.015 |
| **BRWD3** | **S1579** | 0.20 | 0.18 | 0.00 | -0.23 | 0.02 | -0.23 | 0.06 |
| **BRWD3** | **S** | 0.02 | 0.36 | 0.43 | 0.20 | -0.60 | -0.85 | 0.114 |
| **BST2** | **S131** | 0.19 | 0.03 | 0.20 | -0.19 | 0.06 | -0.38 | 0.093 |

| | | | | | | | | |
|---|---|---|---|---|---|---|---|---|
| **BTBD9** | **S595; S** | 0.02 | -0.02 | 0.03 | 0.18 | 0.04 | -0.29 | 0.833 |
| **BTBD9** | **S599** | 0.40 | 0.13 | 0.09 | -0.44 | -0.07 | -0.26 | 0.034 |
| **BUB1B** | **S670** | -0.04 | -0.03 | -0.02 | 0.09 | -0.05 | 0.04 | 0.249 |
| **BUB1B** | **S543** | -0.22 | 0.25 | 0.27 | -0.29 | 0.10 | -0.22 | 0.306 |
| **BUB3** | **S211** | 0.08 | -0.01 | 0.07 | 0.16 | -0.17 | -0.15 | 0.423 |
| **BUB3** | **S211** | -0.18 | 0.17 | 0.18 | 0.24 | 0.16 | -0.85 | 0.603 |
| **BUD13** | **S567** | -0.25 | -0.13 | -0.20 | 0.31 | -0.22 | 0.36 | 0.142 |
| **BUD13** | **S214** | 0.08 | -0.27 | -0.39 | 0.55 | -0.41 | 0.18 | 0.398 |
| **BUD13** | **S** | 0.16 | -0.43 | -0.22 | 0.26 | 0.04 | 0.08 | 0.191 |
| **BUD13** | **S57** | -0.22 | 0.01 | -0.04 | 0.06 | 0.09 | 0.07 | 0.089 |
| **BUD13** | **S226** | 0.03 | -0.03 | -0.16 | 0.08 | -0.17 | 0.22 | 0.48 |
| **BUD13** | **S248** | -0.05 | -0.19 | 0.13 | -0.14 | -0.03 | 0.23 | 0.72 |
| **BUD13** | **S163** | -0.09 | 0.00 | 0.03 | 0.01 | -0.07 | 0.11 | 0.596 |
| **BUD13** | **S214** | 0.14 | -0.20 | 0.04 | -0.01 | -0.21 | 0.19 | 0.973 |
| **BUD13** | **S258** | 0.03 | -0.03 | 0.10 | 0.10 | -0.06 | -0.16 | 0.433 |
| **BUD13** | **S364** | 0.05 | -0.01 | 0.09 | 0.02 | -0.08 | -0.08 | 0.101 |
| **BUD13** | **S271** | -0.09 | 0.06 | 0.14 | -0.31 | 0.21 | -0.08 | 0.599 |
| **BUD13** | **T159; S163** | 0.00 | -0.03 | 0.16 | -0.11 | -0.18 | 0.13 | 0.406 |
| **BUD13** | **S175** | -0.17 | -0.03 | 0.33 | -0.37 | -0.08 | 0.20 | 0.599 |
| **BUD13** | **S139** | 0.16 | -0.04 | 0.07 | -0.15 | -0.07 | 0.00 | 0.131 |
| **BUD13** | **S28** | 0.17 | -0.16 | 0.18 | -0.19 | -0.01 | -0.04 | 0.316 |
| **BUD13** | **S354** | 0.12 | 0.02 | 0.09 | -0.04 | -0.11 | -0.08 | 0.012 |
| **BUD13** | **S** | 0.06 | 0.03 | 0.18 | -0.06 | -0.15 | -0.09 | 0.026 |
| **BUD13** | **S325** | 0.12 | 0.11 | 0.11 | -0.13 | 0.02 | -0.28 | 0.047 |
| **BUD13** | **S/T** | -0.08 | 0.03 | 0.37 | -0.12 | -0.10 | -0.19 | 0.152 |
| **BUD13** | **S407** | 0.05 | 0.19 | 0.10 | -0.10 | 0.08 | -0.39 | 0.16 |
| **BUD13** | **T/S** | 0.11 | 0.10 | 0.24 | -0.17 | -0.09 | -0.26 | 0.008 |
| **BUD13** | **S248** | 0.17 | 0.11 | 0.24 | -0.16 | -0.09 | -0.36 | 0.013 |
| **BUD13** | **S271** | 0.18 | 0.31 | 0.16 | -0.25 | -0.02 | -0.56 | 0.04 |
| **BUD13** | **S226** | 0.12 | 0.31 | 0.23 | -0.38 | -0.07 | -0.36 | 0.013 |
| **BUD13** | **S404; S407** | 0.15 | 0.26 | 0.30 | -0.38 | -0.15 | -0.33 | 0.004 |
| **BUD13** | **S210; S211** | 0.11 | 0.04 | 0.52 | -0.46 | 0.02 | -0.48 | 0.076 |
| **BUD13** | **S364; S372; S375** | 0.31 | 0.33 | 0.37 | -0.40 | -0.13 | -0.87 | 0.022 |
| **BUD13** | **S364; S372; S375** | 0.22 | 0.45 | 0.35 | -0.42 | -0.21 | -0.80 | 0.011 |
| **BYSL** | **S98** | -0.17 | -0.12 | -0.08 | 0.20 | -0.06 | 0.18 | 0.057 |
| **BYSL** | **S98** | -0.01 | -0.06 | -0.02 | 0.03 | -0.06 | 0.12 | 0.339 |
| **C11orf52** | **S62** | -0.01 | -0.18 | 0.07 | 0.11 | 0.25 | -0.30 | 0.761 |
| **C11orf52** | **Y103** | 0.07 | 0.02 | 0.03 | -0.24 | -0.18 | 0.25 | 0.575 |
| **C11orf52** | **Y/T** | 0.04 | 0.40 | -0.01 | -0.53 | -0.55 | 0.35 | 0.3 |
| **C14orf93** | **S268; S272; T273** | 0.13 | 0.01 | -0.10 | -0.10 | -0.01 | 0.05 | 0.712 |
| **C14orf93** | **S119** | -0.05 | 0.20 | 0.05 | -0.13 | -0.13 | 0.03 | 0.201 |
| **C14orf93** | **S285** | 0.15 | 0.10 | 0.09 | 0.23 | 0.20 | -1.20 | 0.48 |
| **C14orf93** | **S224** | 0.27 | -0.01 | 0.52 | -0.35 | -0.03 | -0.72 | 0.067 |
| **C16orf87** | **S50** | 0.15 | 0.21 | 0.16 | -0.22 | -0.07 | -0.33 | 0.008 |
| **C19orf47** | **S151; T155** | -0.19 | -0.55 | -0.21 | 0.26 | -0.22 | 0.61 | 0.115 |
| **C19orf47** | **S306** | 0.11 | -0.17 | 0.10 | 0.01 | -0.34 | 0.22 | 0.819 |
| **C19orf47** | **S306** | 0.19 | -0.02 | -0.07 | -0.06 | -0.10 | 0.05 | 0.492 |
| **C19orf47** | **S124** | 0.11 | 0.06 | 0.00 | -0.11 | -0.10 | 0.04 | 0.111 |

| | | | | | | | | |
|---|---|---|---|---|---|---|---|---|
| C19orf47 | S151; S/T | -0.07 | 0.07 | 0.21 | 0.16 | -0.25 | -0.18 | 0.349 |
| C19orf47 | S280 | 0.20 | 0.22 | 0.21 | -0.23 | -0.13 | -0.37 | 0.003 |
| C1orf131 | S279 | -0.18 | -0.19 | -0.32 | -0.08 | 0.16 | 0.46 | 0.064 |
| C1orf131 | T142 | 0.11 | -0.08 | -0.02 | -0.05 | 0.02 | 0.02 | 0.93 |
| C1orf159 | S342 | -0.48 | -0.52 | -0.62 | 0.47 | -0.78 | 0.98 | 0.217 |
| C1orf174 | S148 | 0.03 | 0.16 | 0.37 | -0.64 | -0.01 | -0.10 | 0.121 |
| C1orf174 | S148 | -0.04 | 0.13 | 0.41 | -1.28 | 0.14 | 0.12 | 0.354 |
| C1orf198 | T211 | -0.05 | -0.06 | -0.21 | 0.09 | -0.12 | 0.29 | 0.205 |
| C1orf198 | S175 | 0.04 | -0.03 | 0.02 | 0.00 | 0.01 | -0.05 | 0.449 |
| C1orf21 | S95 | 0.12 | -0.16 | -0.27 | 0.27 | -0.12 | 0.08 | 0.335 |
| C1orf216 | S126 | -0.08 | -0.05 | 0.28 | -0.37 | 0.15 | -0.01 | 0.551 |
| C1orf226 | S223 | -0.16 | -0.08 | -0.09 | -0.07 | 0.02 | 0.32 | 0.169 |
| C1QBP | S/T | -0.17 | -0.06 | 0.05 | 0.02 | -0.04 | 0.18 | 0.282 |
| C2CD3 | S1829 | -0.02 | 0.00 | -0.19 | 0.13 | 0.05 | 0.01 | 0.139 |
| C2CD3 | S2132 | 0.00 | 0.09 | -0.16 | -0.46 | 0.22 | 0.20 | 0.962 |
| C2CD3 | S2018; S2020 | 0.15 | -0.04 | -0.04 | -0.06 | -0.02 | 0.00 | 0.455 |
| C2CD3 | S1606 | 0.25 | -0.36 | 0.19 | -0.18 | 0.21 | -0.22 | 0.724 |
| C2CD3 | S2114 | -0.08 | 0.17 | 0.11 | -0.11 | -0.09 | -0.02 | 0.148 |
| C2CD5 | S284; T285 | -0.02 | -0.11 | -0.08 | 0.29 | 0.00 | -0.12 | 0.35 |
| C2CD5 | S/T | 0.15 | -0.20 | -0.10 | 0.06 | 0.05 | 0.01 | 0.428 |
| C2CD5 | S659; S661; S662 | 0.08 | -0.06 | -0.08 | -0.03 | -0.18 | 0.23 | 0.835 |
| C2CD5 | S659; S662 | -0.08 | -0.02 | 0.07 | 0.09 | -0.09 | 0.02 | 0.762 |
| C2CD5 | S852 | -0.10 | -0.02 | 0.13 | 0.08 | -0.08 | -0.01 | 0.918 |
| C2CD5 | S659 | 0.00 | 0.22 | 0.12 | -0.10 | 0.01 | -0.32 | 0.101 |
| C2CD5 | S281 | 0.16 | 0.04 | 0.20 | -0.21 | -0.41 | 0.12 | 0.137 |
| C2CD5 | S323 | 0.09 | 0.08 | 0.38 | -0.51 | 0.22 | -0.49 | 0.16 |
| C2CD5 | S323 | 0.00 | 0.35 | 0.39 | -0.36 | 0.05 | -0.71 | 0.08 |
| C2orf49 | S182; S/T | 0.41 | -0.21 | -0.16 | 0.28 | 0.19 | -0.84 | 0.758 |
| C2orf49 | S182 | 0.20 | 0.18 | -0.11 | -0.33 | 0.28 | -0.36 | 0.376 |
| C2orf49 | S189 | 0.21 | 0.12 | 0.19 | -0.13 | -0.14 | -0.33 | 0.006 |
| C4orf19 | S279 | -0.24 | 0.01 | -0.52 | 0.39 | -0.31 | 0.41 | 0.221 |
| C5orf24 | S37 | -0.31 | -0.27 | -0.72 | 0.34 | 0.14 | 0.48 | 0.012 |
| C6orf132 | S906 | -0.19 | -0.03 | -0.28 | -0.09 | 0.28 | 0.22 | 0.087 |
| C6orf132 | S/T | -0.78 | 0.22 | -0.01 | -0.65 | 0.67 | 0.05 | 0.687 |
| C6orf132 | S969 | 0.03 | 0.16 | -0.18 | -0.10 | 0.15 | -0.10 | 0.902 |
| C6orf132 | S969 | 0.27 | -0.06 | 0.38 | -0.10 | -0.16 | -0.50 | 0.066 |
| C6orf47 | S112 | 0.29 | 0.21 | 0.01 | -0.17 | 0.04 | -0.53 | 0.105 |
| C7orf50 | S59 | -0.16 | -0.04 | -0.11 | 0.11 | 0.05 | 0.13 | 0.009 |
| C7orf50 | S36 | -0.17 | 0.15 | -0.17 | -0.02 | 0.00 | 0.18 | 0.389 |
| C7orf50 | S175 | -0.08 | 0.05 | -0.01 | 0.05 | -0.10 | 0.09 | 0.722 |
| C9orf152 | S88 | -0.03 | -0.09 | -0.15 | 0.22 | -0.07 | 0.09 | 0.146 |
| C9orf152 | S88 | 0.17 | 0.14 | -0.12 | -0.05 | 0.18 | -0.41 | 0.463 |
| C9orf78 | S261 | 0.27 | 0.35 | 0.04 | -0.29 | -0.06 | -0.49 | 0.032 |
| C9orf78 | S261 | 0.24 | 0.27 | 0.23 | -0.33 | -0.08 | -0.52 | 0.013 |
| CAAP1 | S312 | -0.22 | -0.12 | -0.20 | 0.22 | 0.05 | 0.20 | 0.005 |
| CAAP1 | S203 | -0.16 | -0.03 | -0.13 | 0.06 | 0.06 | 0.17 | 0.021 |
| CAAP1 | S/T | -0.07 | -0.26 | 0.06 | 0.45 | -0.08 | -0.21 | 0.55 |
| CAAP1 | S203 | -0.14 | 0.00 | 0.03 | 0.07 | 0.14 | -0.13 | 0.54 |

| | | | | | | | | |
|---|---|---|---|---|---|---|---|---|
| CAAP1 | S203 | 0.03 | 0.11 | -0.09 | -0.01 | 0.08 | -0.13 | 0.661 |
| CAAP1 | S92 | 0.08 | 0.09 | -0.05 | -0.02 | -0.01 | -0.10 | 0.184 |
| CAAP1 | S203 | 0.13 | -0.07 | 0.09 | -0.02 | 0.11 | -0.28 | 0.432 |
| CAAP1 | S89 | 0.05 | 0.13 | 0.08 | -0.07 | -0.07 | -0.14 | 0.005 |
| CAAP1 | S203 | 0.10 | 0.07 | 0.09 | 0.04 | -0.02 | -0.32 | 0.172 |
| CAAP1 | S203 | 0.11 | 0.18 | 0.05 | -0.09 | 0.09 | -0.41 | 0.181 |
| CABIN1 | T1370 | -0.65 | -0.73 | -0.74 | 0.45 | 0.56 | 0.41 | 3E-05 |
| CABIN1 | S1439 | -0.54 | -0.35 | -0.55 | 0.30 | 0.23 | 0.53 | 0.002 |
| CABIN1 | S433 | -0.23 | -0.51 | -0.41 | 0.20 | 0.35 | 0.34 | 0.002 |
| CABIN1 | S/T | -0.04 | 0.06 | -0.23 | 0.02 | 0.16 | 0.01 | 0.236 |
| CABIN1 | S1439 | -0.07 | 0.19 | 0.08 | -0.16 | 0.00 | -0.07 | 0.169 |
| CABIN1 | S2094 | 0.10 | 0.08 | 0.03 | -0.10 | -0.04 | -0.09 | 0.005 |
| CABIN1 | T1927 | 0.11 | 0.08 | 0.11 | -0.24 | 0.03 | -0.13 | 0.059 |
| CABIN1 | S2094 | 0.08 | 0.26 | 0.05 | -0.21 | 0.03 | -0.28 | 0.067 |
| CABLES1 | T415 | -0.07 | -0.15 | 0.00 | 0.03 | 0.05 | 0.12 | 0.055 |
| CABLES1 | S/T | -0.20 | -0.17 | 0.30 | 0.12 | 0.02 | -0.14 | 0.9 |
| CABLES1 | S291 | -0.10 | 0.19 | 0.01 | -0.07 | -0.01 | -0.04 | 0.442 |
| CABLES2 | S130 | 0.25 | 0.14 | -0.25 | -0.21 | 0.28 | -0.34 | 0.603 |
| CAD | S1038 | 0.05 | -0.07 | 0.04 | 0.06 | 0.14 | -0.24 | 0.853 |
| CAD | S1859 | -0.09 | -0.09 | 0.23 | -0.11 | -0.08 | 0.11 | 0.752 |
| CAD | S1859 | 0.19 | 0.22 | 0.18 | -0.26 | -0.02 | -0.45 | 0.024 |
| CADM1 | S434 | 0.13 | 0.22 | 0.11 | -0.42 | 0.00 | -0.11 | 0.063 |
| CADPS2 | S58 | 0.08 | 0.11 | -0.07 | -0.05 | -0.11 | 0.03 | 0.282 |
| CAMK2D | S330 | -0.11 | -0.47 | -0.49 | 0.57 | -0.24 | 0.38 | 0.095 |
| CAMK2D | T287 | 0.02 | -0.16 | -0.12 | 0.02 | -0.01 | 0.22 | 0.137 |
| CAMK2D | S/T | 0.18 | -0.14 | -0.13 | 0.28 | -0.04 | -0.22 | 0.842 |
| CAMK2D | T337 | -0.45 | 0.19 | 0.21 | -0.03 | 0.01 | -0.03 | 0.994 |
| CAMK2G | S419 | -0.76 | 0.02 | 0.30 | -0.25 | -0.06 | 0.45 | 0.639 |
| CAMK2G | S419 | 0.08 | -0.12 | -0.08 | 0.05 | -0.16 | 0.20 | 0.582 |
| CAMK2G | S419 | -0.06 | 0.03 | 0.00 | 0.01 | 0.02 | 0.00 | 0.629 |
| CAMSAP1 | S862 | -0.18 | -0.29 | 0.03 | -0.19 | 0.41 | 0.11 | 0.266 |
| CAMSAP2 | S1319 | -0.18 | 0.13 | -0.24 | -0.20 | 0.05 | 0.34 | 0.459 |
| CAMSAP2 | S1148 | 0.25 | 0.36 | 0.31 | -0.59 | -0.22 | -0.39 | 0.003 |
| CAMSAP2 | S599 | 0.22 | 0.46 | 0.29 | -0.85 | 0.03 | -0.61 | 0.043 |
| CAMSAP3 | S585 | -0.11 | -0.32 | -0.22 | 0.31 | -0.21 | 0.39 | 0.128 |
| CAMSAP3 | S814 | -0.04 | -0.15 | 0.20 | 0.13 | -0.17 | -0.01 | 0.899 |
| CAMSAP3 | S/T | 0.10 | 0.08 | -0.12 | 0.06 | -0.23 | 0.09 | 0.737 |
| CAMSAP3 | S341; S/T | 0.09 | -0.02 | 0.25 | -0.04 | -0.39 | 0.04 | 0.202 |
| CANX | S583 | 0.02 | 0.15 | -0.04 | -0.13 | -0.22 | 0.19 | 0.499 |
| CANX | S554 | 0.13 | 0.16 | 0.12 | -0.15 | 0.04 | -0.36 | 0.064 |
| CANX | S554 | 0.08 | 0.11 | 0.20 | -0.07 | 0.06 | -0.48 | 0.147 |
| CANX | S583 | 0.18 | 0.16 | 0.21 | -0.26 | -0.26 | -0.11 | 0.002 |
| CANX | S554; S564 | 0.27 | 0.20 | 0.31 | -0.35 | -0.13 | -0.49 | 0.006 |
| CANX | S554; S564 | 0.33 | 0.29 | 0.30 | -0.54 | -0.30 | -0.36 | 6E-04 |
| CAP2 | S301 | -0.06 | -0.19 | -0.13 | 0.10 | 0.00 | 0.24 | 0.038 |
| CAP2 | S312 | -0.16 | 0.12 | -0.23 | 0.01 | -0.14 | 0.31 | 0.429 |
| CARMIL1 | S880 | 0.26 | 0.30 | -0.18 | -0.11 | 0.18 | -0.65 | 0.33 |
| CARMIL2 | S274; S275 | -0.18 | 0.27 | -0.19 | -0.17 | -0.04 | 0.23 | 0.86 |

| | | | | | | | | |
|---|---|---|---|---|---|---|---|---|
| CASC3 | S363 | 0.11 | -0.13 | -0.20 | 0.27 | -0.28 | 0.15 | 0.563 |
| CASC3 | S363 | -0.20 | -0.02 | 0.01 | 0.04 | -0.22 | 0.32 | 0.522 |
| CASC3 | S125 | 0.11 | 0.02 | 0.09 | 0.07 | -0.05 | -0.26 | 0.207 |
| CASC3 | S66 | 0.15 | 0.14 | 0.08 | -0.05 | -0.20 | -0.16 | 0.007 |
| CASC3 | S45 | -0.01 | 0.32 | 0.30 | -0.17 | -0.30 | -0.28 | 0.017 |
| CASC3 | S363 | 0.14 | 0.24 | 0.33 | 0.04 | -0.16 | -0.92 | 0.123 |
| CASK | S313 | -0.20 | 0.00 | -0.28 | -0.08 | 0.21 | 0.27 | 0.095 |
| CASKIN2 | S471 | -0.36 | 0.33 | -0.26 | -0.04 | 0.19 | 0.04 | 0.516 |
| CASKIN2 | S842 | 0.20 | -0.08 | 0.17 | -0.18 | 0.19 | -0.41 | 0.307 |
| CASKIN2 | S403 | 0.07 | 0.12 | 0.35 | 0.01 | 0.16 | -1.11 | 0.294 |
| CASP8AP2 | S875 | 0.49 | -0.01 | 0.11 | -0.28 | -0.55 | 0.03 | 0.11 |
| CASP8AP2 | S875 | 0.44 | 0.00 | 0.17 | -0.41 | -0.40 | 0.02 | 0.071 |
| CASZ1 | S981 | -0.11 | -0.16 | -0.15 | 0.11 | 0.03 | 0.23 | 0.011 |
| CASZ1 | S57 | -0.07 | -0.20 | 0.01 | 0.20 | -0.26 | 0.25 | 0.431 |
| CASZ1 | T/Y/S | -0.03 | 0.05 | -0.25 | 0.11 | 0.29 | -0.25 | 0.522 |
| CASZ1 | S130 | -0.01 | -0.08 | -0.07 | 0.06 | 0.03 | 0.07 | 0.015 |
| CASZ1 | S130 | 0.09 | -0.10 | -0.15 | 0.05 | 0.02 | 0.07 | 0.244 |
| CASZ1 | S130 | 0.05 | -0.01 | -0.11 | 0.05 | 0.05 | -0.03 | 0.427 |
| CASZ1 | S130 | 0.10 | -0.09 | -0.08 | -0.27 | 0.01 | 0.27 | 0.871 |
| CASZ1 | S130 | 0.07 | -0.04 | -0.05 | -0.02 | 0.04 | 0.00 | 0.791 |
| CASZ1 | S130 | 0.01 | 0.02 | -0.04 | 0.08 | 0.01 | -0.08 | 0.849 |
| CASZ1 | S130 | 0.05 | -0.07 | 0.06 | 0.05 | -0.04 | -0.06 | 0.63 |
| CASZ1 | S130 | 0.05 | -0.02 | 0.03 | 0.16 | -0.05 | -0.20 | 0.686 |
| CASZ1 | T/S | -0.18 | 0.18 | 0.06 | -0.03 | -0.10 | 0.04 | 0.681 |
| CASZ1 | S130 | 0.02 | -0.07 | 0.13 | 0.08 | 0.01 | -0.20 | 0.553 |
| CASZ1 | S130 | 0.00 | 0.13 | -0.03 | 0.22 | -0.12 | -0.25 | 0.618 |
| CASZ1 | S151 | 0.05 | 0.09 | 0.00 | -0.13 | -0.02 | 0.01 | 0.154 |
| CASZ1 | S/T | 0.09 | -0.03 | 0.13 | -0.12 | -0.21 | 0.10 | 0.256 |
| CASZ1 | S720 | 0.10 | 0.13 | 0.03 | -0.08 | -0.08 | -0.12 | 0.004 |
| CASZ1 | T1378 | 0.17 | 0.16 | -0.07 | -0.23 | 0.08 | -0.16 | 0.204 |
| CASZ1 | S130 | 0.14 | 0.01 | 0.13 | -0.16 | -0.07 | -0.08 | 0.019 |
| CASZ1 | T1378 | 0.13 | 0.28 | -0.13 | -0.17 | 0.07 | -0.26 | 0.247 |
| CASZ1 | S130 | 0.00 | 0.42 | -0.04 | 0.15 | -0.01 | -0.79 | 0.347 |
| CASZ1 | S741 | 0.48 | 0.13 | 0.44 | -0.43 | -0.13 | -1.00 | 0.036 |
| CBFA2T2 | S264 | -0.07 | 0.23 | 0.17 | -0.70 | 0.20 | -0.02 | 0.379 |
| CBX1 | S172 | -0.47 | -0.33 | -0.43 | 0.27 | 0.39 | 0.29 | 2E-04 |
| CBX1 | S89 | -0.06 | -0.26 | 0.18 | 0.06 | -0.21 | 0.22 | 0.705 |
| CBX1 | S89 | 0.05 | 0.00 | -0.13 | 0.01 | -0.02 | 0.08 | 0.446 |
| CBX1 | S89 | 0.17 | 0.08 | -0.21 | -0.08 | 0.08 | -0.06 | 0.813 |
| CBX2 | S302 | 0.08 | 0.17 | 0.00 | 0.07 | -0.16 | -0.19 | 0.142 |
| CBX3 | S176 | -1.16 | -0.99 | -1.10 | 0.58 | 0.76 | 0.48 | 5E-05 |
| CBX3 | T173; S176 | -0.58 | -0.47 | -0.70 | 0.45 | 0.27 | 0.51 | 5E-04 |
| CBX3 | S93; S95 | 0.01 | -0.11 | -0.13 | 0.00 | -0.02 | 0.23 | 0.175 |
| CBX3 | S93 | 0.09 | 0.04 | -0.07 | -0.04 | -0.02 | -0.01 | 0.426 |
| CBX3 | S95; S99; S102 | 0.09 | 0.10 | 0.02 | -0.14 | -0.06 | -0.03 | 0.025 |
| CBX3 | S95; S97 | 0.14 | 0.01 | 0.05 | -0.15 | -0.08 | 0.00 | 0.068 |
| CBX3 | S95 | 0.18 | 0.14 | 0.05 | -0.17 | -0.02 | -0.23 | 0.021 |
| CBX3 | S95 | 0.25 | 0.29 | 0.19 | -0.31 | -0.12 | -0.49 | 0.007 |

| | | | | | | | | |
|---|---|---|---|---|---|---|---|---|
| **CBX3** | **S99; S102** | 0.24 | 0.42 | 0.21 | -0.29 | -0.22 | -0.61 | 0.008 |
| **CBX4** | **S415** | -0.16 | 0.01 | 0.36 | -0.06 | -0.11 | -0.11 | 0.349 |
| **CBX5** | **S14** | 0.10 | 0.15 | 0.15 | -0.14 | -0.16 | -0.13 | 1E-04 |
| **CBX5** | **S95** | 0.23 | 0.05 | 0.13 | -0.11 | -0.16 | -0.18 | 0.006 |
| **CBX5** | **S13** | 0.01 | 0.01 | 0.43 | -0.25 | -0.14 | -0.18 | 0.076 |
| **CBX5** | **S97** | 0.11 | 0.16 | 0.27 | -0.26 | -0.16 | -0.20 | 0.002 |
| **CBX6** | **S316** | 0.11 | 0.01 | -0.33 | 0.09 | -0.24 | 0.27 | 0.607 |
| **CBX6** | **S301** | 0.07 | -0.07 | -0.11 | 0.30 | -0.38 | 0.10 | 0.834 |
| **CBX6** | **S280** | 0.18 | 0.14 | -0.04 | -0.13 | -0.22 | 0.03 | 0.112 |
| **CBX6** | **S280; S/T** | 0.02 | 0.03 | 0.27 | -0.35 | -0.11 | 0.06 | 0.17 |
| **CBX6** | **S280** | 0.18 | 0.21 | 0.17 | -0.29 | -0.19 | -0.16 | 6E-04 |
| **CBX8** | **T264** | -0.42 | -0.06 | -0.27 | 0.32 | 0.07 | 0.22 | 0.024 |
| **CBX8** | **S265** | -0.14 | -0.15 | -0.16 | 0.17 | 0.16 | 0.07 | 9E-04 |
| **CBX8** | **S110** | -0.11 | -0.11 | -0.15 | 0.13 | 0.14 | 0.08 | 5E-04 |
| **CBX8** | **S110** | 0.15 | -0.04 | -0.25 | 0.18 | -0.06 | -0.02 | 0.582 |
| **CBX8** | **S191** | 0.22 | 0.17 | 0.12 | -0.25 | 0.02 | -0.38 | 0.037 |
| **CBX8** | **S130** | 0.10 | -0.03 | 0.44 | -0.16 | 0.30 | -1.13 | 0.323 |
| **CC2D1A** | **T92** | -0.30 | -0.14 | 0.01 | 0.13 | 0.16 | 0.08 | 0.041 |
| **CC2D1A** | **T120** | -0.24 | -0.22 | 0.43 | -0.09 | -0.25 | 0.23 | 0.927 |
| **CC2D1A** | **T120** | 0.51 | -0.08 | 0.06 | -0.85 | -0.09 | 0.12 | 0.276 |
| **CC2D1A** | **S455** | 0.36 | 0.49 | -0.13 | -0.55 | -0.10 | -0.34 | 0.068 |
| **CC2D1B** | **S528** | 0.06 | -0.15 | 0.10 | -0.10 | 0.13 | -0.05 | 0.929 |
| **CCAR1** | **T627** | 0.08 | 0.32 | 0.19 | 0.31 | -0.55 | -0.67 | 0.185 |
| **CCAR2** | **S569** | -0.20 | -0.27 | -0.23 | 0.19 | 0.17 | 0.24 | 1E-04 |
| **CCAR2** | **S** | -0.21 | -0.01 | -0.05 | 0.27 | -0.03 | -0.01 | 0.22 |
| **CCAR2** | **S/T** | 0.13 | -0.09 | -0.23 | 0.14 | 0.26 | -0.30 | 0.665 |
| **CCAR2** | **S626** | 0.26 | -0.18 | -0.23 | 0.15 | 0.01 | -0.08 | 0.668 |
| **CCDC106** | **S130** | 0.08 | 0.12 | 0.22 | -0.13 | -0.10 | -0.24 | 0.007 |
| **CCDC120** | **S280** | -0.20 | -0.42 | -0.93 | 0.79 | -0.03 | 0.20 | 0.064 |
| **CCDC120** | **S256** | -0.13 | -0.29 | -0.17 | 0.05 | 0.07 | 0.39 | 0.038 |
| **CCDC137** | **S19** | 0.16 | 0.16 | 0.27 | -0.26 | -0.04 | -0.40 | 0.018 |
| **CCDC138** | **S49** | 0.26 | -0.03 | 0.01 | -0.41 | -0.15 | 0.22 | 0.397 |
| **CCDC138** | **S149** | 0.29 | 0.57 | 0.29 | -0.77 | -0.06 | -0.93 | 0.027 |
| **CCDC14** | **S798** | -0.07 | -0.05 | 0.17 | 0.12 | -0.28 | 0.06 | 0.755 |
| **CCDC168** | **S6050** | 0.13 | 0.10 | 0.17 | -0.25 | 0.05 | -0.26 | 0.05 |
| **CCDC186** | **S740** | 0.04 | -0.01 | -0.07 | 0.05 | 0.25 | -0.32 | 0.989 |
| **CCDC6** | **S323** | -0.24 | -0.13 | -0.36 | 0.17 | 0.12 | 0.32 | 0.008 |
| **CCDC6** | **S240; S244** | -0.17 | -0.02 | 0.28 | -0.06 | -0.17 | 0.08 | 0.632 |
| **CCDC6** | **S244** | 0.22 | 0.18 | 0.10 | -0.38 | -0.09 | -0.11 | 0.023 |
| **CCDC6** | **S419** | 0.33 | 0.25 | -0.05 | -0.16 | -0.18 | -0.31 | 0.035 |
| **CCDC61** | **T285** | 0.07 | 0.12 | -0.09 | 0.10 | -0.20 | -0.03 | 0.506 |
| **CCDC61** | **S336** | 0.36 | 0.03 | -0.10 | -0.86 | -0.21 | 0.42 | 0.468 |
| **CCDC61** | **S447** | 0.16 | 0.18 | 0.08 | 0.07 | -0.04 | -0.57 | 0.186 |
| **CCDC61** | **S376** | 0.26 | 0.22 | 0.30 | -0.29 | -0.09 | -0.61 | 0.018 |
| **CCDC61** | **S473** | 0.00 | 0.48 | 0.22 | 0.21 | -0.68 | -0.60 | 0.136 |
| **CCDC71** | **S97** | -0.11 | -0.20 | 0.05 | 0.12 | 0.13 | -0.01 | 0.129 |
| **CCDC71** | **S453** | -0.15 | 0.10 | -0.06 | -0.15 | 0.08 | 0.14 | 0.621 |
| **CCDC86** | **S128** | -0.77 | -1.03 | -0.84 | 0.36 | 0.73 | 0.50 | 4E-04 |

| | | | | | | | | |
|---|---|---|---|---|---|---|---|---|
| **CCDC86** | **S80; S91** | -0.39 | -0.28 | -0.01 | 0.12 | 0.11 | 0.34 | 0.039 |
| **CCDC86** | **S47** | -0.32 | -0.14 | -0.03 | 0.24 | -0.09 | 0.25 | 0.102 |
| **CCDC86** | **S160** | -0.63 | -0.06 | 0.20 | -0.26 | 0.11 | 0.42 | 0.461 |
| **CCDC86** | **S160** | -0.08 | -0.06 | -0.21 | -0.03 | -0.05 | 0.37 | 0.212 |
| **CCDC86** | **S91** | -0.19 | -0.03 | -0.14 | 0.32 | -0.21 | 0.16 | 0.276 |
| **CCDC86** | **S47** | -0.04 | -0.21 | -0.01 | 0.00 | -0.15 | 0.34 | 0.385 |
| **CCDC86** | **S102; S110** | -0.14 | -0.06 | 0.05 | -0.04 | 0.09 | 0.08 | 0.255 |
| **CCDC86** | **T182; S/Y/T** | 0.23 | -0.11 | -0.30 | -0.08 | -0.14 | 0.30 | 0.702 |
| **CCDC86** | **S102; S110; S113** | -0.05 | -0.06 | 0.00 | -0.01 | -0.04 | 0.14 | 0.306 |
| **CCDC86** | **S113** | -0.05 | -0.08 | 0.02 | -0.04 | 0.01 | 0.13 | 0.298 |
| **CCDC86** | **S102** | 0.08 | 0.28 | -0.53 | 0.26 | -0.01 | -0.24 | 0.851 |
| **CCDC86** | **S47** | 0.00 | 0.01 | -0.07 | -0.03 | -0.05 | 0.12 | 0.589 |
| **CCDC86** | **S102; S110; S113** | 0.02 | -0.04 | 0.02 | -0.03 | -0.12 | 0.14 | 0.944 |
| **CCDC86** | **S102** | 0.06 | -0.08 | 0.04 | -0.05 | -0.04 | 0.06 | 0.842 |
| **CCDC86** | **S80; S91** | -0.03 | -0.01 | 0.06 | 0.02 | -0.04 | 0.01 | 0.749 |
| **CCDC86** | **S91** | -0.03 | -0.04 | 0.11 | 0.02 | 0.00 | -0.07 | 0.654 |
| **CCDC86** | **S102; S/Y** | 0.01 | -0.02 | 0.08 | -0.07 | 0.00 | 0.00 | 0.337 |
| **CCDC86** | **S58** | 0.05 | -0.08 | 0.09 | -0.02 | -0.16 | 0.10 | 0.66 |
| **CCDC86** | **S102; S110** | 0.03 | -0.09 | 0.12 | -0.22 | 0.01 | 0.13 | 0.702 |
| **CCDC86** | **S102** | 0.07 | 0.04 | 0.02 | -0.06 | -0.04 | -0.03 | 0.005 |
| **CCDC86** | **S91** | 0.01 | 0.09 | 0.14 | -0.17 | -0.03 | -0.06 | 0.044 |
| **CCDC86** | **S18** | 0.01 | 0.07 | 0.15 | -0.31 | 0.11 | -0.08 | 0.251 |
| **CCDC88C** | **S227** | -0.13 | -0.37 | -0.07 | 0.04 | 0.06 | 0.36 | 0.066 |
| **CCDC9** | **S376; T385; T/S** | -0.57 | 0.15 | -0.14 | 0.18 | 0.23 | 0.01 | 0.217 |
| **CCDC9** | **S376; T389; S390** | -0.27 | -0.17 | -0.02 | 0.08 | 0.05 | 0.27 | 0.045 |
| **CCDC9** | **S376; T385** | 0.02 | -0.18 | -0.17 | 0.13 | 0.04 | 0.13 | 0.043 |
| **CCDC9** | **S386** | -0.20 | 0.04 | -0.14 | 0.08 | 0.08 | 0.11 | 0.06 |
| **CCDC9** | **T389; S390** | -0.15 | -0.08 | -0.02 | 0.14 | -0.08 | 0.16 | 0.135 |
| **CCDC9** | **S386** | 0.01 | -0.01 | -0.08 | 0.02 | 0.02 | 0.03 | 0.178 |
| **CCDC9** | **S80** | 0.02 | 0.01 | -0.06 | 0.10 | -0.06 | -0.02 | 0.795 |
| **CCDC9** | **T95** | 0.00 | -0.06 | 0.06 | -0.04 | 0.08 | -0.04 | 0.964 |
| **CCDC9** | **S60** | 0.23 | 0.03 | -0.09 | 0.02 | 0.03 | -0.25 | 0.389 |
| **CCDC9** | **S521** | -0.10 | 0.09 | 0.29 | 0.37 | -0.38 | -0.47 | 0.437 |
| **CCDC93** | **S305** | -0.42 | -0.01 | 0.16 | 0.15 | 0.07 | -0.02 | 0.423 |
| **CCDC97** | **S337** | -0.05 | -0.32 | -0.35 | 0.09 | -0.10 | 0.54 | 0.12 |
| **CCHCR1** | **S/T** | -0.32 | 0.14 | 0.17 | -0.01 | -0.25 | 0.19 | 0.939 |
| **CCM2** | **S384** | -0.21 | -0.14 | -0.15 | 0.30 | 0.18 | -0.05 | 0.042 |
| **CCNE2** | **T/S** | 0.29 | -0.05 | 0.12 | -0.06 | -0.38 | -0.01 | 0.142 |
| **CCNE2** | **S21** | 0.35 | 0.15 | 0.20 | -0.55 | -0.17 | -0.16 | 0.022 |
| **CCNH** | **T315** | -0.05 | -0.08 | -0.10 | 0.05 | 0.07 | 0.09 | 0.001 |
| **CCNK** | **S340** | -0.03 | 0.01 | -0.17 | 0.00 | -0.06 | 0.22 | 0.301 |
| **CCNK** | **S324; S** | -0.05 | -0.06 | -0.04 | 0.13 | -0.07 | 0.07 | 0.197 |
| **CCNK** | **S328** | 0.08 | -0.07 | 0.01 | 0.01 | -0.07 | 0.03 | 0.714 |
| **CCNK** | **S340** | 0.17 | 0.02 | -0.10 | 0.06 | -0.07 | -0.10 | 0.503 |
| **CCNK** | **S/T/Y** | 0.50 | -0.05 | 0.28 | -0.25 | 0.04 | -0.91 | 0.127 |
| **CCNL1** | **S65** | -0.45 | -0.56 | -0.20 | 0.14 | 0.22 | 0.54 | 0.012 |
| **CCNL1** | **S65** | 0.06 | -0.16 | -0.06 | 0.03 | -0.17 | 0.25 | 0.539 |
| **CCNL1** | **S/Y** | 0.06 | -0.17 | 0.04 | 0.02 | 0.01 | 0.02 | 0.635 |

| | | | | | | | | |
|---|---|---|---|---|---|---|---|---|
| **CCNL1** | **S352** | 0.10 | 0.17 | -0.16 | 0.23 | -0.10 | -0.31 | 0.641 |
| **CCNL1** | **S374** | 0.13 | -0.06 | 0.13 | 0.11 | 0.06 | -0.46 | 0.44 |
| **CCNL1** | **S445** | 0.15 | 0.16 | 0.18 | -0.06 | -0.06 | -0.47 | 0.055 |
| **CCNL2** | **S369** | -0.03 | -0.16 | -0.15 | 0.15 | -0.08 | 0.22 | 0.109 |
| **CCNL2** | **S369** | -0.13 | 0.19 | -0.28 | 0.03 | 0.15 | -0.01 | 0.411 |
| **CCNL2** | **S369** | 0.22 | 0.04 | -0.13 | -0.03 | 0.07 | -0.21 | 0.475 |
| **CCNL2** | **S330** | 0.17 | -0.02 | 0.03 | -0.11 | -0.06 | -0.03 | 0.102 |
| **CCNL2** | **S348** | 0.21 | -0.02 | 0.05 | 0.00 | -0.19 | -0.07 | 0.126 |
| **CCNT2** | **S601** | -0.28 | -0.20 | -0.17 | 0.45 | -0.13 | 0.19 | 0.086 |
| **CCNT2** | **S480** | 0.17 | 0.07 | 0.03 | -0.13 | 0.19 | -0.43 | 0.316 |
| **CCNY** | **S25** | -0.26 | -0.09 | -0.17 | -0.03 | 0.05 | 0.40 | 0.089 |
| **CCNY** | **S/Y/T** | -0.11 | -0.03 | -0.23 | -0.12 | 0.28 | 0.15 | 0.162 |
| **CCNY** | **S326** | -0.08 | -0.02 | 0.09 | 0.03 | -0.01 | -0.02 | 0.871 |
| **CCNYL1** | **S122** | -0.24 | -0.15 | -0.24 | 0.33 | 0.00 | 0.20 | 0.018 |
| **CCP110** | **S170** | -0.01 | 0.04 | 0.21 | -0.07 | -0.08 | -0.12 | 0.074 |
| **CCSAP** | **S240** | -0.04 | -0.06 | -0.16 | 0.05 | -0.05 | 0.23 | 0.143 |
| **CCSER2** | **T430** | 0.10 | 0.02 | 0.12 | -0.01 | 0.03 | -0.30 | 0.186 |
| **CCT8** | **Y304** | 0.27 | 0.03 | 0.08 | -0.04 | -0.09 | -0.32 | 0.07 |
| **CCZ1B** | **S266** | 0.08 | -0.12 | 0.17 | -0.26 | -0.06 | 0.13 | 0.492 |
| **CD2AP** | **S510; T512** | -0.05 | -0.12 | 0.00 | -0.07 | -0.15 | 0.33 | 0.572 |
| **CD2AP** | **S510** | 0.03 | 0.06 | 0.11 | -0.20 | 0.06 | -0.08 | 0.143 |
| **CD2AP** | **S256** | 0.01 | -0.11 | 0.30 | -0.10 | 0.01 | -0.17 | 0.318 |
| **CD2AP** | **S510; S514** | 0.12 | 0.10 | 0.27 | -0.15 | 0.01 | -0.46 | 0.07 |
| **CD2AP** | **S510** | 0.22 | 0.31 | 0.22 | -0.35 | -0.02 | -0.59 | 0.027 |
| **CD2BP2** | **S195** | 0.10 | 0.02 | 0.16 | 0.17 | -0.22 | -0.30 | 0.249 |
| **CD2BP2** | **S195** | 0.13 | 0.13 | 0.13 | -0.02 | 0.00 | -0.46 | 0.124 |
| **CD2BP2** | **S49** | 0.24 | -0.09 | 0.30 | -0.05 | -0.11 | -0.40 | 0.104 |
| **CDC16** | **S560** | -0.01 | 0.11 | -0.07 | -0.38 | 0.38 | -0.14 | 0.838 |
| **CDC20** | **T/S** | -0.03 | 0.15 | -0.05 | -0.06 | 0.03 | -0.05 | 0.506 |
| **CDC20** | **T106** | -0.12 | 0.12 | 0.06 | 0.02 | 0.36 | -0.62 | 0.748 |
| **CDC26** | **S42** | 0.23 | -0.27 | -0.06 | -0.07 | -0.11 | 0.21 | 0.824 |
| **CDC40** | **S43** | 0.12 | 0.14 | -0.33 | -0.22 | 0.05 | 0.17 | 0.909 |
| **CDC42BPG** | **S1550** | 0.22 | -0.11 | -0.50 | 0.12 | -0.79 | 0.62 | 0.821 |
| **CDC42BPG** | **S** | -0.01 | -0.13 | 0.00 | 0.17 | 0.02 | -0.07 | 0.336 |
| **CDC42BPG** | **S480** | -0.12 | 0.03 | -0.02 | 0.03 | -0.08 | 0.14 | 0.463 |
| **CDC42EP4** | **S292** | 0.03 | -0.07 | -0.04 | 0.07 | -0.05 | 0.05 | 0.326 |
| **CDC45** | **S144; S148; S151** | -0.32 | -0.16 | -0.26 | 0.35 | 0.26 | -0.02 | 0.021 |
| **CDC45** | **S148; S151** | 0.16 | -0.06 | 0.00 | 0.01 | -0.08 | -0.04 | 0.379 |
| **CDC5L** | **T385** | -0.27 | -0.22 | -0.22 | 0.33 | -0.41 | 0.53 | 0.251 |
| **CDC5L** | **T355** | -0.07 | -0.05 | -0.12 | 0.02 | 0.01 | 0.19 | 0.071 |
| **CDC5L** | **T352; T355** | 0.14 | -0.14 | -0.13 | -0.09 | -0.02 | 0.21 | 0.586 |
| **CDC5L** | **T424** | 0.03 | 0.00 | 0.03 | 0.02 | -0.10 | 0.01 | 0.368 |
| **CDC5L** | **T385; S/T** | 0.04 | 0.13 | -0.08 | -0.10 | -0.14 | 0.12 | 0.529 |
| **CDC5L** | **T438; T/S** | 0.09 | -0.07 | 0.20 | -0.14 | 0.11 | -0.23 | 0.294 |
| **CDC5L** | **S437; T438; T442** | -0.02 | 0.08 | 0.27 | -0.06 | 0.03 | -0.37 | 0.174 |
| **CDC5L** | **T424** | 0.17 | 0.26 | 0.19 | -0.19 | -0.14 | -0.43 | 0.007 |
| **CDC6** | **S45** | -0.25 | -0.42 | 0.11 | 0.13 | 0.04 | 0.28 | 0.122 |
| **CDC6** | **S74** | 0.00 | -0.16 | -0.09 | 0.06 | -0.09 | 0.24 | 0.238 |

| | | | | | | | | |
|---|---|---|---|---|---|---|---|---|
| **CDC6** | **S74** | 0.08 | -0.12 | -0.07 | -0.10 | -0.04 | 0.22 | 0.619 |
| **CDC7** | **S27** | 0.03 | -0.17 | 0.01 | -0.22 | 0.14 | 0.16 | 0.629 |
| **CDCA2** | **S400** | -0.80 | 0.03 | -0.02 | 0.23 | 0.24 | 0.10 | 0.177 |
| **CDCA2** | **S710** | -0.01 | -0.09 | 0.07 | -0.23 | 0.05 | 0.17 | 0.971 |
| **CDCA2** | **S98** | 0.05 | -0.13 | 0.07 | -0.08 | -0.17 | 0.22 | 0.967 |
| **CDCA2** | **S756** | 0.00 | 0.01 | 0.01 | 0.09 | 0.18 | -0.34 | 0.865 |
| **CDCA3** | **S222** | -0.34 | -0.27 | 0.02 | 0.57 | -0.19 | 0.02 | 0.264 |
| **CDCA3** | **S94** | 0.03 | -0.09 | 0.05 | 0.12 | 0.02 | -0.15 | 0.998 |
| **CDCA3** | **S29** | -0.10 | 0.13 | 0.07 | 0.03 | -0.16 | 0.02 | 0.495 |
| **CDCA3** | **S87** | -0.02 | 0.23 | 0.40 | -0.36 | -0.25 | -0.16 | 0.027 |
| **CDCA3** | **S68** | 0.07 | 0.23 | 0.45 | -0.27 | -0.20 | -0.50 | 0.016 |
| **CDCA5** | **T111** | -0.41 | -0.21 | -0.42 | 0.04 | -0.01 | 0.69 | 0.066 |
| **CDCA5** | **S21** | -0.07 | -0.14 | -0.51 | 0.06 | 0.09 | 0.42 | 0.078 |
| **CDCA5** | **S/T** | -0.25 | 0.06 | -0.34 | -0.01 | 0.08 | 0.36 | 0.126 |
| **CDCA5** | **S/T** | 0.02 | -0.14 | -0.20 | -0.03 | -0.08 | 0.36 | 0.29 |
| **CDCA5** | **S75** | 0.25 | -0.02 | 0.05 | -0.08 | -0.29 | 0.04 | 0.177 |
| **CDCA7L** | **S21** | 0.05 | 0.04 | -0.08 | 0.09 | -0.16 | 0.04 | 0.862 |
| **CDCA8** | **T204** | -0.41 | -0.43 | -0.30 | 0.21 | 0.30 | 0.38 | 4E-04 |
| **CDCA8** | **S219** | 0.06 | 0.04 | 0.05 | -0.18 | 0.07 | -0.05 | 0.233 |
| **CDCA8** | **S219** | 0.10 | 0.12 | 0.02 | -0.23 | 0.18 | -0.25 | 0.276 |
| **CDCP1** | **S797** | 0.01 | 0.08 | 0.21 | -0.07 | 0.11 | -0.43 | 0.245 |
| **CDK1** | **S39** | -0.52 | 0.14 | 0.24 | 0.06 | -0.09 | 0.05 | 0.83 |
| **CDK1** | **T14** | 0.27 | 0.30 | 0.26 | -0.33 | -0.14 | -0.61 | 0.01 |
| **CDK10** | **T350** | 0.27 | -0.19 | 0.12 | 0.19 | -0.14 | -0.35 | 0.466 |
| **CDK10** | **T196** | 0.28 | 0.43 | 0.26 | -0.14 | -0.12 | -1.24 | 0.093 |
| **CDK11A** | **T583** | 0.21 | -0.24 | -0.34 | -0.33 | 0.19 | 0.35 | 0.498 |
| **CDK11B** | **T/S/Y** | -0.38 | -0.36 | -0.25 | 0.84 | -0.47 | 0.14 | 0.26 |
| **CDK11B** | **T/S/Y** | -0.16 | -0.24 | -0.22 | 0.45 | -0.12 | 0.16 | 0.088 |
| **CDK11B** | **S752** | 0.01 | -0.15 | -0.21 | 0.11 | 0.01 | 0.19 | 0.059 |
| **CDK11B** | **S283** | -0.15 | -0.15 | -0.12 | 0.17 | 0.49 | -0.44 | 0.475 |
| **CDK11B** | **T/S/Y** | 0.24 | -0.43 | -0.16 | 0.11 | -0.29 | 0.36 | 0.545 |
| **CDK11B** | **S277; S283** | -0.14 | 0.01 | -0.07 | 0.14 | -0.12 | 0.15 | 0.276 |
| **CDK11B** | **S713** | -0.10 | -0.20 | 0.11 | -0.22 | 0.29 | 0.04 | 0.603 |
| **CDK11B** | **S277** | 0.04 | 0.01 | -0.10 | -0.03 | 0.05 | 0.02 | 0.574 |
| **CDK11B** | **S115** | 0.20 | 0.20 | 0.02 | -0.12 | 0.04 | -0.44 | 0.11 |
| **CDK11B** | **S113; S115** | 0.30 | 0.15 | 0.21 | -0.29 | -0.05 | -0.49 | 0.02 |
| **CDK12** | **T255** | -0.70 | -0.42 | -0.42 | 0.70 | -0.53 | 0.65 | 0.13 |
| **CDK12** | **S236; S241; S/Y** | -0.10 | -0.62 | 0.33 | 0.32 | -0.07 | -0.06 | 0.563 |
| **CDK12** | **S/Y** | -0.14 | -0.17 | 0.04 | -0.03 | -0.20 | 0.41 | 0.486 |
| **CDK12** | **S274; S276** | -0.10 | 0.03 | -0.11 | 0.01 | -0.11 | 0.25 | 0.383 |
| **CDK12** | **T692** | 0.02 | -0.10 | -0.08 | 0.13 | -0.05 | 0.07 | 0.169 |
| **CDK12** | **S1083** | -0.11 | -0.05 | 0.02 | 0.01 | -0.02 | 0.13 | 0.218 |
| **CDK12** | **T893** | 0.03 | 0.02 | -0.18 | -0.08 | 0.22 | -0.03 | 0.542 |
| **CDK12** | **S133** | 0.06 | -0.11 | -0.06 | 0.17 | -0.07 | -0.01 | 0.493 |
| **CDK12** | **S423** | 0.04 | -0.11 | -0.01 | 0.09 | -0.04 | 0.03 | 0.371 |
| **CDK12** | **T893** | 0.06 | -0.02 | -0.11 | 0.01 | -0.01 | 0.05 | 0.491 |
| **CDK12** | **S276** | -0.03 | -0.16 | 0.15 | -0.02 | 0.09 | -0.04 | 0.805 |
| **CDK12** | **S357; S359** | 0.10 | -0.02 | -0.10 | 0.06 | -0.12 | 0.07 | 0.934 |

| | | | | | | | | |
|---|---|---|---|---|---|---|---|---|
| **CDK12** | **S332; S334** | 0.01 | -0.07 | 0.10 | 0.09 | -0.08 | -0.06 | 0.708 |
| **CDK12** | **S434** | 0.06 | -0.07 | 0.05 | 0.04 | 0.09 | -0.19 | 0.731 |
| **CDK12** | **T1244** | 0.06 | -0.09 | 0.08 | 0.10 | 0.05 | -0.22 | 0.746 |
| **CDK12** | **S423** | -0.07 | 0.05 | 0.07 | 0.02 | 0.12 | -0.21 | 0.715 |
| **CDK12** | **S1083** | 0.18 | 0.10 | -0.26 | 0.20 | 0.07 | -0.39 | 0.849 |
| **CDK12** | **S359** | -0.02 | -0.08 | 0.15 | 0.03 | 0.12 | -0.24 | 0.731 |
| **CDK12** | **T1244** | -0.02 | 0.07 | 0.05 | 0.02 | 0.00 | -0.12 | 0.256 |
| **CDK12** | **S332** | 0.04 | 0.00 | 0.08 | -0.10 | 0.14 | -0.18 | 0.452 |
| **CDK12** | **S456** | 0.03 | 0.33 | -0.32 | -0.59 | 0.03 | 0.30 | 0.769 |
| **CDK12** | **T1244** | 0.16 | 0.06 | -0.09 | -0.22 | 0.14 | -0.10 | 0.461 |
| **CDK12** | **S274; S276** | 0.18 | 0.06 | -0.08 | 0.08 | -0.11 | -0.15 | 0.346 |
| **CDK12** | **S18** | -0.37 | 0.18 | 0.31 | -0.04 | 0.16 | -0.38 | 0.657 |
| **CDK12** | **S276** | 0.22 | 0.02 | -0.09 | -0.02 | 0.19 | -0.40 | 0.54 |
| **CDK12** | **S385** | -0.09 | 0.15 | 0.09 | 0.08 | 0.13 | -0.45 | 0.537 |
| **CDK12** | **S1075** | 0.04 | 0.08 | 0.06 | 0.08 | 0.03 | -0.34 | 0.365 |
| **CDK12** | **S276** | 0.24 | -0.07 | -0.03 | -0.17 | 0.37 | -0.49 | 0.621 |
| **CDK12** | **S/T** | -0.07 | 0.02 | 0.25 | 0.17 | 0.03 | -0.49 | 0.504 |
| **CDK12** | **S323; S325** | -0.18 | 0.11 | 0.29 | -0.10 | -0.12 | -0.06 | 0.283 |
| **CDK12** | **T514** | 0.24 | 0.19 | -0.10 | -0.03 | -0.04 | -0.33 | 0.175 |
| **CDK12** | **S341; S345** | 0.05 | 0.24 | 0.04 | 0.08 | -0.27 | -0.20 | 0.129 |
| **CDK12** | **S323; S325; S/Y** | 0.21 | -0.09 | 0.20 | 0.06 | 0.00 | -0.50 | 0.279 |
| **CDK12** | **S/T** | 0.31 | -0.02 | 0.15 | -0.18 | 0.03 | -0.38 | 0.099 |
| **CDK12** | **S236; S241; S/Y** | 0.28 | 0.09 | 0.08 | -0.35 | -0.21 | 0.02 | 0.059 |
| **CDK12** | **S1083** | 0.38 | -0.37 | 0.25 | 0.01 | 0.40 | -1.32 | 0.534 |
| **CDK12** | **S** | 0.20 | 0.13 | 0.25 | -0.12 | -0.07 | -0.52 | 0.042 |
| **CDK12** | **S236; S249** | 0.08 | 0.20 | 0.30 | -0.24 | 0.04 | -0.54 | 0.069 |
| **CDK12** | **T514** | 0.21 | 0.51 | -0.13 | -0.20 | 0.02 | -0.67 | 0.156 |
| **CDK12** | **S** | 0.11 | 0.27 | 0.33 | -0.26 | -0.09 | -0.57 | 0.025 |
| **CDK13** | **S397** | -0.38 | 0.18 | -0.61 | -0.32 | 0.34 | 0.45 | 0.272 |
| **CDK13** | **S413** | -0.16 | -0.09 | -0.24 | 0.20 | 0.31 | -0.09 | 0.075 |
| **CDK13** | **T871** | -0.12 | -0.05 | -0.01 | -0.01 | 0.32 | -0.19 | 0.546 |
| **CDK13** | **S383** | 0.11 | -0.13 | -0.07 | 0.23 | -0.10 | -0.09 | 0.756 |
| **CDK13** | **S367** | -0.01 | 0.12 | 0.01 | 0.02 | 0.04 | -0.20 | 0.37 |
| **CDK13** | **S437; S439** | 0.02 | 0.07 | 0.05 | -0.03 | -0.06 | -0.05 | 0.004 |
| **CDK13** | **T871** | 0.15 | 0.06 | -0.05 | -0.14 | 0.02 | -0.06 | 0.197 |
| **CDK13** | **S317** | 0.08 | 0.05 | 0.06 | 0.14 | -0.04 | -0.35 | 0.358 |
| **CDK13** | **T1246** | 0.07 | 0.15 | 0.13 | -0.14 | 0.03 | -0.29 | 0.062 |
| **CDK13** | **S525** | 0.15 | 0.20 | 0.06 | 0.02 | -0.25 | -0.24 | 0.039 |
| **CDK13** | **T494** | 0.19 | 0.16 | 0.08 | -0.07 | -0.10 | -0.32 | 0.023 |
| **CDK13** | **S317; S325** | 0.24 | 0.18 | 0.05 | -0.18 | 0.03 | -0.42 | 0.071 |
| **CDK13** | **S505** | 0.17 | 0.34 | 0.03 | -0.04 | -0.06 | -0.63 | 0.116 |
| **CDK13** | **T588** | 0.24 | 0.12 | 0.24 | -0.25 | -0.05 | -0.43 | 0.019 |
| **CDK13** | **S525** | 0.39 | 0.08 | 0.19 | -0.13 | -0.35 | -0.33 | 0.013 |
| **CDK13** | **S525** | 0.34 | 0.22 | 0.10 | -0.34 | -0.04 | -0.45 | 0.023 |
| **CDK13** | **S342; S349** | 0.29 | 0.17 | 0.27 | -0.01 | -0.36 | -0.57 | 0.03 |
| **CDK17** | **S137** | -0.47 | -0.10 | 0.14 | -0.18 | 0.16 | 0.31 | 0.357 |
| **CDK18** | **S74** | 0.37 | 0.09 | -0.11 | -0.60 | -0.18 | 0.23 | 0.335 |
| **CDK5RAP2** | **S613** | -0.15 | -0.07 | -0.04 | -0.04 | 0.08 | 0.20 | 0.096 |

| | | | | | | | | |
|---|---|---|---|---|---|---|---|---|
| **CDK5RAP2** | **S1074** | 0.06 | -0.07 | -0.21 | 0.04 | 0.13 | 0.02 | 0.171 |
| **CDK5RAP2** | **S1061** | 0.33 | -0.09 | -0.37 | -0.20 | 0.17 | 0.03 | 0.865 |
| **CDK5RAP2** | **S706** | 0.00 | 0.01 | -0.12 | -0.13 | 0.44 | -0.31 | 0.889 |
| **CDK5RAP2** | **S1887** | -0.12 | 0.03 | 0.08 | -0.01 | -0.07 | 0.08 | 0.92 |
| **CDK5RAP2** | **S255** | 0.14 | -0.21 | 0.14 | -0.33 | 0.17 | 0.01 | 0.713 |
| **CDK5RAP2** | **S1490** | -0.03 | 0.05 | 0.09 | 0.15 | -0.06 | -0.21 | 0.509 |
| **CDK5RAP2** | **S/T** | 0.24 | 0.03 | -0.15 | 0.08 | -0.15 | -0.09 | 0.511 |
| **CDK5RAP2** | **S1350** | 0.04 | -0.11 | 0.21 | -0.04 | -0.32 | 0.16 | 0.519 |
| **CDK5RAP2** | **S/T/Y** | 0.00 | -0.01 | 0.23 | 0.01 | -0.27 | -0.01 | 0.241 |
| **CDK7** | **T170** | 0.13 | 0.07 | -0.05 | 0.15 | -0.13 | -0.21 | 0.396 |
| **CDKAL1** | **S53** | -0.23 | 0.02 | 0.31 | -0.21 | -0.54 | 0.43 | 0.683 |
| **CDKN1B** | **S10** | 0.10 | 0.13 | 0.30 | -0.08 | -0.20 | -0.35 | 0.019 |
| **CDR2L** | **S308** | -0.38 | -0.08 | -0.18 | 0.33 | -0.29 | 0.41 | 0.201 |
| **CDR2L** | **S308** | -0.28 | 0.02 | -0.36 | 0.32 | -0.45 | 0.48 | 0.353 |
| **CDR2L** | **S393** | -0.29 | 0.53 | 0.14 | -0.13 | -0.07 | -0.39 | 0.279 |
| **CDS2** | **S41** | -0.10 | 0.02 | -0.25 | 0.05 | 0.05 | 0.20 | 0.085 |
| **CDS2** | **S21** | 0.22 | 0.23 | 0.12 | -0.17 | -0.25 | -0.23 | 6E-04 |
| **CDT1** | **S93** | 0.34 | 0.31 | 0.00 | 0.11 | -0.60 | -0.39 | 0.097 |
| **CDT1** | **S/T** | 0.34 | 0.31 | 0.00 | 0.11 | -0.60 | -0.39 | 0.097 |
| **CDYL** | **S88** | -0.37 | -0.24 | 0.03 | 0.03 | 0.00 | 0.42 | 0.128 |
| **CDYL** | **S88** | 0.16 | -0.24 | -0.31 | -0.28 | -0.11 | 0.57 | 0.557 |
| **CDYL** | **S88** | -0.08 | 0.07 | 0.05 | 0.06 | -0.03 | -0.09 | 0.652 |
| **CDYL** | **S216** | -0.03 | 0.08 | 0.01 | 0.03 | -0.06 | -0.01 | 0.478 |
| **CDYL** | **S216** | 0.03 | 0.13 | 0.11 | -0.08 | -0.20 | -0.02 | 0.032 |
| **CEACAM20** | **S568** | -0.19 | 0.32 | 0.22 | -0.04 | -0.47 | 0.01 | 0.263 |
| **CEBPA** | **S266** | -0.38 | -0.53 | -0.05 | 0.45 | -0.17 | 0.40 | 0.092 |
| **CEBPB** | **T235** | -0.45 | -0.41 | -0.52 | 0.37 | 0.06 | 0.57 | 0.006 |
| **CEBPZ** | **T/S** | 0.00 | -0.10 | -0.45 | 0.14 | -0.38 | 0.55 | 0.394 |
| **CEBPZ** | **T/S** | 0.06 | -0.17 | -0.18 | 0.13 | -0.24 | 0.31 | 0.404 |
| **CEBPZ** | **S629** | 0.03 | -0.03 | -0.09 | 0.08 | 0.08 | -0.08 | 0.434 |
| **CEBPZ** | **S835** | 0.05 | -0.01 | -0.14 | 0.10 | 0.14 | -0.18 | 0.678 |
| **CEBPZ** | **S629** | 0.05 | -0.02 | -0.06 | 0.08 | 0.10 | -0.16 | 0.856 |
| **CEBPZ** | **S629** | 0.03 | -0.05 | -0.01 | 0.13 | 0.10 | -0.24 | 0.951 |
| **CEBPZ** | **S629** | 0.04 | 0.02 | -0.03 | 0.09 | 0.07 | -0.21 | 0.834 |
| **CEBPZ** | **S835** | 0.08 | 0.03 | -0.06 | 0.04 | 0.06 | -0.16 | 0.667 |
| **CEBPZ** | **S629** | 0.08 | -0.05 | 0.03 | -0.03 | 0.10 | -0.14 | 0.615 |
| **CEL** | **T59** | -0.43 | -0.25 | -0.51 | 0.49 | 0.12 | 0.29 | 0.006 |
| **CELF1** | **S28** | 0.07 | -0.13 | 0.00 | 0.14 | 0.22 | -0.37 | 0.934 |
| **CELSR3** | **S3175** | -0.04 | 0.50 | 0.55 | -1.50 | -0.45 | 0.51 | 0.251 |
| **CENPA** | **S19** | -0.32 | -0.51 | -1.18 | 0.38 | -0.17 | 0.90 | 0.062 |
| **CENPA** | **S17** | -0.50 | -0.18 | -0.07 | 0.38 | -0.05 | 0.26 | 0.067 |
| **CENPA** | **S17; S19** | -0.03 | -0.07 | -0.02 | -0.01 | 0.02 | 0.10 | 0.102 |
| **CENPA** | **S17; S19** | 0.07 | -0.05 | -0.07 | 0.06 | -0.04 | 0.02 | 0.594 |
| **CENPA** | **S19** | 0.01 | -0.08 | 0.05 | -0.01 | 0.03 | 0.00 | 0.73 |
| **CENPA** | **S17; S19** | -0.05 | 0.04 | 0.02 | -0.28 | 0.13 | 0.10 | 0.888 |
| **CENPB** | **S307** | -0.08 | -0.16 | -0.06 | -0.15 | 0.18 | 0.22 | 0.206 |
| **CENPB** | **S156; S/T** | 0.00 | 0.04 | -0.04 | 0.22 | -0.33 | 0.06 | 0.92 |
| **CENPB** | **S156** | -0.03 | 0.30 | 0.09 | -0.06 | -0.07 | -0.30 | 0.101 |

| | | | | | | | | |
|---|---|---|---|---|---|---|---|---|
| **CENPC** | **S/T** | -0.08 | -0.26 | -0.36 | 0.21 | -0.12 | 0.44 | 0.088 |
| **CENPC** | **S501** | -0.06 | -0.19 | -0.31 | 0.13 | 0.20 | 0.16 | 0.009 |
| **CENPC** | **S110** | -0.15 | -0.12 | -0.29 | 0.10 | -0.03 | 0.39 | 0.066 |
| **CENPC** | **S/T/Y** | -0.07 | -0.07 | -0.13 | 0.15 | -0.03 | 0.13 | 0.045 |
| **CENPC** | **S146** | -0.31 | 0.01 | 0.01 | -0.03 | -0.09 | 0.34 | 0.378 |
| **CENPC** | **S96** | -0.12 | -0.14 | 0.05 | -0.06 | -0.04 | 0.26 | 0.35 |
| **CENPC** | **S311** | -0.01 | -0.02 | -0.15 | 0.14 | 0.04 | -0.01 | 0.132 |
| **CENPC** | **S308; S311** | -0.17 | 0.03 | 0.03 | -0.06 | 0.01 | 0.14 | 0.481 |
| **CENPC** | **T/S** | 0.00 | 0.11 | -0.11 | -0.11 | -0.04 | 0.14 | 0.966 |
| **CENPC** | **S/T** | -0.29 | 0.03 | 0.25 | -0.36 | 0.31 | -0.08 | 0.884 |
| **CENPC** | **S538** | -0.12 | 0.12 | 0.11 | -0.05 | -0.11 | 0.02 | 0.413 |
| **CENPC** | **S333** | 0.10 | 0.04 | -0.01 | -0.10 | 0.12 | -0.17 | 0.352 |
| **CENPC** | **S225** | -0.05 | 0.15 | 0.09 | -0.31 | 0.01 | 0.06 | 0.326 |
| **CENPC** | **S225** | 0.11 | 0.10 | 0.01 | -0.15 | 0.01 | -0.09 | 0.064 |
| **CENPC** | **S73** | 0.14 | 0.02 | 0.08 | -0.16 | -0.10 | 0.00 | 0.05 |
| **CENPC** | **S538** | 0.07 | 0.05 | 0.12 | -0.10 | -0.07 | -0.09 | 0.002 |
| **CENPC** | **S333** | 0.02 | -0.02 | 0.23 | -0.24 | 0.16 | -0.21 | 0.306 |
| **CENPC** | **S613** | 0.17 | -0.01 | 0.17 | -0.22 | -0.05 | -0.09 | 0.043 |
| **CENPC** | **S177** | 0.12 | 0.04 | 0.17 | 0.32 | -0.03 | -0.91 | 0.434 |
| **CENPC** | **S73** | 0.12 | 0.22 | 0.27 | -0.26 | -0.23 | -0.22 | 7E-04 |
| **CENPC** | **T130** | 0.29 | 0.29 | 0.03 | -0.48 | -0.08 | -0.19 | 0.037 |
| **CENPC** | **S73** | 0.35 | 0.19 | 0.05 | -0.25 | 0.02 | -0.54 | 0.069 |
| **CENPC** | **S600** | 0.26 | 0.13 | 0.28 | 0.15 | -0.38 | -0.68 | 0.102 |
| **CENPC** | **S73** | -0.05 | 0.51 | 0.39 | -0.57 | -0.18 | -0.44 | 0.03 |
| **CENPC** | **S73** | 0.30 | 0.35 | 0.32 | -0.63 | 0.02 | -0.77 | 0.033 |
| **CENPE** | **S2654** | -0.25 | 0.07 | -0.10 | 0.13 | 0.14 | -0.03 | 0.185 |
| **CENPF** | **S2900; S2911** | -0.39 | -0.25 | -0.34 | 0.31 | 0.14 | 0.35 | 0.002 |
| **CENPF** | **T/S** | -0.12 | -0.15 | -0.47 | -0.05 | 0.51 | 0.08 | 0.106 |
| **CENPF** | **T/S** | -0.40 | 0.04 | -0.32 | 0.26 | 0.29 | -0.01 | 0.071 |
| **CENPF** | **S3054** | -0.33 | -0.13 | -0.19 | 0.44 | -0.39 | 0.37 | 0.266 |
| **CENPF** | **S2922** | -0.11 | -0.09 | -0.30 | 0.17 | -0.01 | 0.27 | 0.042 |
| **CENPF** | **S/T** | -0.13 | -0.15 | -0.20 | 0.18 | -0.05 | 0.29 | 0.043 |
| **CENPF** | **S1654** | -0.12 | 0.03 | -0.28 | 0.16 | 0.07 | 0.09 | 0.068 |
| **CENPF** | **S2900** | -0.30 | 0.04 | 0.26 | 0.09 | -0.15 | 0.00 | 0.912 |
| **CENPF** | **S1324** | 0.15 | -0.31 | 0.15 | 0.13 | 0.19 | -0.43 | 0.895 |
| **CENPF** | **S3023** | 0.20 | -0.03 | -0.21 | 0.48 | -0.10 | -0.56 | 0.891 |
| **CENPF** | **S876** | -0.01 | 0.16 | -0.09 | -0.13 | -0.07 | 0.11 | 0.651 |
| **CENPF** | **S821** | 0.03 | 0.07 | 0.10 | -0.03 | -0.01 | -0.18 | 0.076 |
| **CENPF** | **S1726** | -0.01 | 0.08 | 0.14 | 0.05 | 0.06 | -0.38 | 0.361 |
| **CENPF** | **S3079** | -0.03 | 0.14 | 0.13 | 0.00 | -0.35 | 0.06 | 0.286 |
| **CENPF** | **S3079** | -0.12 | 0.19 | 0.17 | -0.04 | 0.03 | -0.27 | 0.266 |
| **CENPF** | **S1726** | 0.24 | 0.00 | 0.12 | -0.12 | 0.07 | -0.38 | 0.149 |
| **CENPF** | **S2998** | -0.06 | 0.26 | 0.35 | -0.25 | 0.10 | -0.60 | 0.141 |
| **CENPI** | **S284** | 0.02 | -0.14 | -0.23 | 0.10 | -0.02 | 0.22 | 0.099 |
| **CENPJ** | **S260** | -0.12 | 0.16 | -0.07 | 0.07 | 0.05 | -0.12 | 0.945 |
| **CENPL** | **S53** | -0.11 | -0.01 | -0.13 | 0.29 | -0.07 | -0.02 | 0.281 |
| **CENPL** | **S53** | 0.16 | -0.01 | -0.16 | -0.09 | 0.00 | 0.08 | 0.998 |
| **CENPL** | **S53** | 0.09 | -0.07 | 0.06 | -0.02 | -0.13 | 0.05 | 0.436 |

| | | | | | | | | |
|---|---|---|---|---|---|---|---|---|
| **CENPN** | **S218** | 0.17 | 0.04 | -0.22 | -0.39 | -0.16 | 0.42 | 0.895 |
| **CENPN** | **S282** | 0.14 | -0.23 | 0.16 | -0.14 | 0.01 | 0.02 | 0.684 |
| **CENPN** | **S282** | 0.15 | -0.08 | 0.05 | 0.18 | 0.15 | -0.59 | 0.661 |
| **CENPN** | **S226** | 0.29 | 0.11 | 0.02 | -0.18 | -0.38 | 0.05 | 0.102 |
| **CENPN** | **S226** | 0.45 | 0.18 | 0.14 | -0.21 | -1.11 | 0.10 | 0.152 |
| **CENPO** | **S263** | -0.22 | -0.11 | -0.32 | -0.04 | 0.46 | 0.09 | 0.078 |
| **CENPO** | **S263** | -0.07 | -0.14 | 0.01 | -0.05 | 0.04 | 0.19 | 0.196 |
| **CENPO** | **S35** | 0.34 | 0.08 | 0.06 | -0.27 | -0.34 | 0.02 | 0.068 |
| **CENPQ** | **S31** | 0.02 | -0.14 | -0.10 | 0.04 | -0.06 | 0.20 | 0.222 |
| **CENPQ** | **S50** | 0.06 | -0.02 | 0.09 | -0.02 | 0.04 | -0.17 | 0.265 |
| **CENPT** | **S343** | -0.03 | -0.17 | -0.38 | 0.04 | 0.28 | 0.17 | 0.045 |
| **CENPT** | **S400; S403; S407** | -0.28 | -0.29 | 0.06 | 0.25 | 0.03 | 0.15 | 0.073 |
| **CENPT** | **S47** | 0.20 | -0.05 | -0.29 | 0.18 | -0.21 | 0.09 | 0.716 |
| **CENPT** | **S397** | 0.05 | -0.06 | -0.03 | 0.10 | -0.19 | 0.11 | 0.879 |
| **CENPT** | **S343** | 0.06 | -0.07 | 0.12 | -0.04 | -0.14 | 0.05 | 0.347 |
| **CENPT** | **S397; S403** | 0.09 | -0.12 | 0.35 | 0.03 | -0.16 | -0.27 | 0.209 |
| **CENPU** | **S139** | -0.46 | 0.00 | -0.06 | 0.17 | -0.14 | 0.35 | 0.211 |
| **CENPU** | **S136; S139; S141** | -0.11 | 0.01 | -0.21 | -0.11 | 0.03 | 0.33 | 0.264 |
| **CENPU** | **S** | 0.06 | -0.01 | -0.35 | -0.04 | 0.01 | 0.27 | 0.326 |
| **CENPU** | **S111; S116; S/T** | 0.00 | -0.08 | 0.00 | -0.07 | 0.03 | 0.10 | 0.491 |
| **CENPU** | **S194** | 0.12 | 0.02 | -0.21 | -0.04 | -0.03 | 0.12 | 0.734 |
| **CENPU** | **S107; T110** | -0.16 | -0.02 | 0.18 | -0.18 | -0.02 | 0.15 | 0.906 |
| **CENPU** | **S204** | 0.19 | -0.01 | -0.14 | -0.08 | 0.09 | -0.08 | 0.772 |
| **CENPU** | **S136; S139** | -0.01 | 0.03 | 0.16 | -0.10 | -0.19 | 0.08 | 0.226 |
| **CENPU** | **S170** | 0.26 | 0.12 | 0.08 | 0.01 | -0.34 | -0.21 | 0.046 |
| **CEP120** | **S347** | -0.20 | -0.01 | -0.12 | 0.25 | 0.13 | -0.09 | 0.143 |
| **CEP120** | **S935** | -0.16 | -0.07 | 0.03 | 0.26 | -0.12 | 0.02 | 0.371 |
| **CEP128** | **S797** | -0.22 | -0.09 | -0.01 | 0.01 | 0.04 | 0.23 | 0.102 |
| **CEP128** | **S331** | -0.14 | -0.04 | -0.11 | 0.10 | 0.03 | 0.13 | 0.012 |
| **CEP128** | **S1033** | -0.05 | -0.09 | 0.15 | 0.11 | -0.04 | -0.10 | 0.905 |
| **CEP128** | **S298** | -0.34 | 0.41 | 0.08 | -0.09 | -0.05 | -0.12 | 0.55 |
| **CEP128** | **S/T** | -0.09 | 0.37 | 0.10 | -0.11 | -0.34 | -0.03 | 0.152 |
| **CEP128** | **S854** | 0.12 | 0.22 | 0.10 | -0.22 | -0.13 | -0.15 | 0.002 |
| **CEP128** | **S167** | 0.39 | -0.13 | 0.19 | -0.13 | -0.06 | -0.40 | 0.129 |
| **CEP128** | **S1008** | 0.13 | 0.16 | 0.24 | -0.32 | -0.36 | 0.04 | 0.042 |
| **CEP128** | **S1008** | 0.29 | 0.16 | 0.17 | -0.43 | -0.01 | -0.33 | 0.024 |
| **CEP128** | **S1061** | 0.32 | 0.31 | 0.00 | -0.20 | -0.17 | -0.40 | 0.022 |
| **CEP131** | **S78** | -0.16 | -0.11 | -0.10 | 0.14 | 0.18 | 0.01 | 0.013 |
| **CEP131** | **T413; S414** | -0.29 | -0.08 | 0.00 | 0.03 | -0.03 | 0.30 | 0.167 |
| **CEP131** | **S223; S224** | -0.07 | -0.07 | -0.02 | 0.00 | 0.09 | 0.06 | 0.031 |
| **CEP131** | **S78** | 0.16 | -0.64 | 0.29 | -0.17 | -0.33 | 0.42 | 0.916 |
| **CEP131** | **S417** | -0.17 | -0.01 | 0.11 | -0.02 | -0.03 | 0.10 | 0.716 |
| **CEP131** | **S150** | -0.08 | 0.10 | 0.05 | -0.09 | 0.02 | 0.00 | 0.473 |
| **CEP131** | **T/S** | -0.33 | 0.37 | -0.02 | 0.32 | -0.56 | 0.00 | 0.81 |
| **CEP131** | **S208** | -0.07 | 0.10 | 0.06 | 0.31 | -0.13 | -0.36 | 0.682 |
| **CEP131** | **S798** | 0.05 | 0.01 | 0.13 | -0.04 | -0.26 | 0.08 | 0.29 |
| **CEP131** | **S114** | 0.13 | 0.13 | -0.01 | -0.03 | -0.15 | -0.09 | 0.042 |
| **CEP131** | **S221** | 0.12 | 0.08 | 0.08 | 0.01 | -0.12 | -0.19 | 0.03 |

| | | | | | | | | |
|---|---|---|---|---|---|---|---|---|
| **CEP131** | **T/S/Y** | 0.11 | 0.27 | 0.05 | -0.07 | -0.13 | -0.30 | 0.031 |
| **CEP131** | **S381** | 0.08 | 0.21 | 0.17 | -0.14 | -0.13 | -0.25 | 0.003 |
| **CEP131** | **S308** | 0.18 | 0.22 | 0.08 | -0.20 | -0.17 | -0.17 | 0.001 |
| **CEP131** | **S381** | 0.16 | 0.23 | 0.14 | -0.14 | -0.10 | -0.39 | 0.016 |
| **CEP131** | **S/T** | 0.16 | 0.08 | 0.39 | -0.13 | -0.05 | -0.63 | 0.08 |
| **CEP131** | **S105; S/T** | 0.26 | 0.16 | 0.31 | -0.35 | -0.28 | -0.25 | 5E-04 |
| **CEP131** | **S/T** | 0.28 | 0.29 | 0.18 | -0.11 | -0.14 | -0.75 | 0.053 |
| **CEP131** | **S525** | 0.25 | 0.25 | 0.36 | -0.34 | -0.32 | -0.43 | 2E-04 |
| **CEP131** | **S89** | 0.27 | 0.32 | 0.29 | -0.12 | -0.23 | -0.88 | 0.041 |
| **CEP152** | **S1449** | -0.08 | -0.09 | -0.35 | 0.04 | 0.26 | 0.14 | 0.043 |
| **CEP152** | **T1351** | 0.12 | -0.17 | -0.24 | 0.15 | 0.21 | -0.13 | 0.308 |
| **CEP152** | **S1247** | -0.36 | 0.03 | 0.14 | -0.11 | 0.07 | 0.17 | 0.556 |
| **CEP152** | **S1455** | -0.18 | 0.06 | 0.08 | 0.08 | -0.61 | 0.38 | 0.909 |
| **CEP152** | **S133** | 0.22 | 0.06 | -0.17 | -0.01 | 0.02 | -0.17 | 0.513 |
| **CEP162** | **S/T/Y** | 0.43 | 0.09 | -0.08 | -0.72 | -0.32 | 0.30 | 0.306 |
| **CEP164** | **S531** | -0.15 | -0.07 | -0.22 | -0.08 | 0.16 | 0.29 | 0.081 |
| **CEP164** | **S472** | 0.00 | -0.16 | -0.26 | 0.05 | 0.08 | 0.24 | 0.053 |
| **CEP164** | **S472** | -0.21 | 0.11 | -0.24 | 0.25 | -0.10 | 0.12 | 0.251 |
| **CEP164** | **S588** | 0.38 | -0.02 | -0.02 | 0.28 | -0.25 | -0.58 | 0.358 |
| **CEP170** | **S930; S933** | -0.08 | -0.08 | -0.08 | 0.20 | 0.01 | 0.00 | 0.079 |
| **CEP170** | **S933** | -0.19 | 0.07 | 0.10 | -0.11 | -0.11 | 0.20 | 0.993 |
| **CEP170** | **T/S** | -0.11 | 0.03 | 0.23 | -0.11 | 0.13 | -0.22 | 0.47 |
| **CEP170** | **S1522; S1529** | 0.10 | 0.08 | 0.37 | -0.17 | -0.37 | -0.13 | 0.028 |
| **CEP170B** | **S224** | -0.46 | -0.64 | -0.62 | 0.24 | 0.47 | 0.51 | 7E-04 |
| **CEP170B** | **S1179** | -0.29 | -0.66 | -0.38 | 0.43 | 0.24 | 0.33 | 0.004 |
| **CEP170B** | **S988** | -0.30 | -0.28 | -0.27 | 0.30 | -0.48 | 0.67 | 0.257 |
| **CEP170B** | **S360** | -0.14 | -0.16 | -0.25 | 0.14 | 0.17 | 0.17 | 6E-04 |
| **CEP170B** | **S772** | -0.24 | 0.15 | -0.06 | 0.00 | -0.12 | 0.23 | 0.605 |
| **CEP170B** | **S565; T569** | 0.42 | 0.13 | -0.58 | 0.24 | -0.27 | -0.17 | 0.871 |
| **CEP192** | **S2110** | -0.05 | -0.13 | -0.08 | 0.07 | -0.04 | 0.20 | 0.084 |
| **CEP192** | **S** | -0.11 | -0.12 | 0.14 | 0.22 | 0.01 | -0.19 | 0.792 |
| **CEP192** | **S196** | 0.17 | -0.16 | 0.19 | -0.45 | -0.13 | 0.25 | 0.478 |
| **CEP250** | **S2179** | -0.20 | -0.13 | -0.63 | 0.00 | 0.30 | 0.42 | 0.05 |
| **CEP250** | **S2322** | -0.05 | 0.42 | -0.10 | -0.35 | 0.06 | -0.11 | 0.335 |
| **CEP250** | **S2229** | 0.15 | 0.31 | 0.37 | -0.28 | -0.29 | -0.48 | 0.002 |
| **CEP290** | **S1610** | 0.01 | 0.15 | 0.11 | 0.01 | 0.04 | -0.38 | 0.228 |
| **CEP350** | **T878** | -0.09 | -0.12 | -0.22 | 0.19 | 0.04 | 0.15 | 0.011 |
| **CEP350** | **S1818** | -0.10 | -0.04 | -0.05 | 0.25 | -0.21 | 0.09 | 0.469 |
| **CEP350** | **S1551** | -0.03 | 0.06 | 0.05 | -0.33 | 0.16 | 0.04 | 0.668 |
| **CEP350** | **T2114** | 0.04 | 0.17 | 0.09 | -0.18 | 0.10 | -0.27 | 0.144 |
| **CEP350** | **S567** | 0.17 | 0.15 | 0.07 | -0.19 | -0.04 | -0.20 | 0.009 |
| **CEP350** | **S567** | 0.11 | 0.19 | 0.13 | -0.10 | -0.01 | -0.41 | 0.063 |
| **CEP350** | **S1259** | -0.22 | 0.26 | 0.58 | 0.06 | -0.51 | -0.51 | 0.156 |
| **CEP41** | **S121** | -0.03 | -0.07 | 0.20 | -0.37 | -0.06 | 0.25 | 0.655 |
| **CEP43** | **T169** | -0.24 | -0.13 | -0.32 | 0.17 | 0.31 | 0.11 | 0.006 |
| **CEP43** | **S156; S160** | -0.24 | -0.07 | -0.19 | 0.21 | 0.08 | 0.16 | 0.007 |
| **CEP43** | **S160** | -0.25 | 0.07 | -0.28 | 0.05 | 0.07 | 0.26 | 0.098 |
| **CEP55** | **S428** | -0.42 | -0.33 | 0.37 | 0.14 | -0.16 | 0.22 | 0.519 |

| | | | | | | | | |
|---|---|---|---|---|---|---|---|---|
| **CEP55** | **S436** | 0.11 | 0.22 | -0.16 | -0.09 | -0.06 | -0.06 | 0.32 |
| **CEP57** | **T250** | -0.30 | -0.10 | -1.14 | 0.15 | 0.32 | 0.53 | 0.065 |
| **CEP57** | **S53** | 0.09 | -0.09 | -0.39 | 0.11 | -0.07 | 0.26 | 0.244 |
| **CEP57** | **S388** | 0.08 | -0.11 | -0.07 | 0.04 | -0.01 | 0.07 | 0.335 |
| **CEP57** | **S454** | 0.30 | 0.09 | 0.04 | -0.38 | -0.08 | -0.05 | 0.079 |
| **CEP68** | **S447** | -0.06 | -0.04 | -0.17 | 0.08 | -0.16 | 0.29 | 0.305 |
| **CEP68** | **S478** | 0.02 | -0.13 | -0.08 | 0.13 | -0.25 | 0.24 | 0.545 |
| **CEP72** | **S404** | 0.08 | 0.08 | -0.08 | 0.11 | 0.06 | -0.29 | 0.669 |
| **CEP78** | **S644** | 0.16 | -0.35 | -0.13 | 0.13 | 0.14 | -0.01 | 0.285 |
| **CEP78** | **S485** | -0.01 | -0.13 | -0.06 | 0.06 | -0.15 | 0.25 | 0.382 |
| **CEP78** | **S346** | 0.24 | 0.06 | 0.08 | -0.33 | -0.04 | -0.07 | 0.066 |
| **CEP78** | **S325** | 0.48 | 0.04 | -0.01 | -0.23 | 0.21 | -0.79 | 0.254 |
| **CEP83** | **S698** | -0.01 | 0.11 | 0.14 | -0.34 | -0.06 | 0.11 | 0.27 |
| **CEP95** | **S353** | -0.28 | -0.18 | -0.13 | 0.24 | -0.07 | 0.31 | 0.047 |
| **CEP95** | **S419; S422** | 0.00 | 0.26 | 0.26 | -0.25 | 0.04 | -0.45 | 0.08 |
| **CERS6** | **S381** | 0.09 | -0.05 | -0.09 | -0.21 | -0.13 | 0.33 | 0.956 |
| **CERS6** | **S341; S345; S** | 0.22 | 0.34 | 0.26 | -0.16 | -0.12 | -0.86 | 0.054 |
| **CERT1** | **S** | -0.16 | 0.12 | 0.13 | -0.17 | -0.50 | 0.41 | 0.707 |
| **CETN2** | **S20** | 0.04 | 0.00 | -0.11 | 0.03 | 0.09 | -0.07 | 0.586 |
| **CETN2** | **S20** | 0.04 | 0.05 | -0.10 | 0.07 | 0.09 | -0.18 | 0.988 |
| **CETN2** | **S20** | 0.14 | 0.09 | 0.01 | -0.13 | 0.14 | -0.29 | 0.257 |
| **CETN2** | **S20** | 0.10 | 0.09 | 0.07 | -0.15 | 0.16 | -0.33 | 0.243 |
| **CFAP410** | **T/S** | -0.05 | 0.00 | -0.70 | 0.50 | -0.42 | 0.34 | 0.344 |
| **CFL1** | **T25** | 0.21 | 0.11 | -0.17 | -0.21 | 0.31 | -0.36 | 0.588 |
| **CGN** | **S137; S140** | -0.18 | -0.06 | -0.03 | -0.14 | 0.10 | 0.26 | 0.255 |
| **CGN** | **S137; S150** | -0.30 | 0.10 | 0.00 | -0.15 | 0.04 | 0.25 | 0.526 |
| **CGN** | **S155** | 0.09 | -0.17 | 0.07 | 0.13 | -0.04 | -0.11 | 0.977 |
| **CGN** | **S338** | 0.25 | 0.24 | -0.47 | -0.14 | 0.01 | -0.01 | 0.826 |
| **CGN** | **S137** | -0.08 | -0.10 | 0.28 | 0.01 | -0.16 | 0.01 | 0.593 |
| **CGN** | **S153** | -0.01 | -0.02 | 0.14 | 0.08 | -0.14 | -0.06 | 0.382 |
| **CGN** | **T/S/Y** | 0.13 | 0.13 | -0.03 | -0.05 | -0.13 | -0.07 | 0.049 |
| **CGN** | **S/T/Y** | 0.11 | 0.37 | -0.28 | 0.40 | -0.44 | -0.43 | 0.536 |
| **CHAF1A** | **S775; T783** | -0.40 | -0.18 | -0.26 | 0.27 | 0.07 | 0.34 | 0.008 |
| **CHAF1A** | **T865** | -0.35 | 0.04 | -0.49 | 0.02 | 0.21 | 0.38 | 0.068 |
| **CHAF1A** | **T/S** | -0.37 | -0.19 | -0.20 | 0.12 | -0.06 | 0.52 | 0.069 |
| **CHAF1A** | **S206** | -0.12 | -0.30 | -0.19 | 0.16 | 0.27 | 0.09 | 0.007 |
| **CHAF1A** | **S54** | -0.12 | -0.18 | -0.25 | 0.07 | 0.03 | 0.36 | 0.039 |
| **CHAF1A** | **S255; T259** | -0.13 | -0.20 | -0.20 | 0.14 | 0.03 | 0.30 | 0.016 |
| **CHAF1A** | **S71** | -0.03 | -0.04 | -0.25 | 0.33 | -0.24 | 0.15 | 0.366 |
| **CHAF1A** | **S772** | 0.20 | -0.52 | 0.07 | 0.10 | -0.73 | 0.51 | 0.916 |
| **CHAF1A** | **S951** | 0.00 | -0.01 | -0.04 | -0.13 | 0.06 | 0.11 | 0.707 |
| **CHAF1A** | **T259** | -0.01 | -0.11 | 0.13 | -0.11 | 0.04 | 0.04 | 0.854 |
| **CHAF1A** | **S141** | -0.09 | -0.07 | 0.19 | -0.15 | -0.07 | 0.16 | 0.822 |
| **CHAF1A** | **S775** | 0.18 | 0.07 | -0.16 | -0.02 | -0.06 | -0.04 | 0.513 |
| **CHAF1A** | **T/S** | 0.11 | -0.59 | 0.40 | 0.55 | -0.57 | -0.32 | 0.856 |
| **CHAF1A** | **S141** | 0.12 | 0.01 | 0.40 | -0.27 | -0.09 | -0.28 | 0.039 |
| **CHAF1B** | **S/T** | -0.36 | -0.16 | -0.49 | -0.01 | 0.41 | 0.37 | 0.023 |
| **CHAF1B** | **S458** | 0.06 | -0.19 | -0.26 | 0.05 | -0.07 | 0.33 | 0.198 |

| | | | | | | | | |
|---|---|---|---|---|---|---|---|---|
| **CHAF1B** | **S410** | -0.12 | -0.14 | -0.11 | 0.05 | -0.12 | 0.37 | 0.196 |
| **CHAF1B** | **S458** | 0.09 | -0.08 | -0.14 | 0.06 | -0.11 | 0.15 | 0.5 |
| **CHAF1B** | **T432; T433** | -0.05 | -0.06 | 0.15 | -0.09 | 0.01 | 0.03 | 0.718 |
| **CHAF1B** | **S538** | 0.03 | 0.42 | -0.26 | 0.49 | -0.22 | -0.88 | 0.577 |
| **CHAF1B** | **S538** | 0.20 | 0.11 | 0.17 | -0.12 | 0.02 | -0.49 | 0.086 |
| **CHAF1B** | **S/T** | 0.16 | 0.13 | 0.30 | -0.40 | -0.01 | -0.32 | 0.027 |
| **CHAMP1** | **S572** | -0.68 | -0.86 | -0.64 | 0.26 | 0.57 | 0.58 | 7E-04 |
| **CHAMP1** | **S651; S652; S653** | -0.10 | -0.08 | -0.20 | 0.22 | -0.05 | 0.17 | 0.059 |
| **CHAMP1** | **S241** | -0.12 | -0.11 | -0.10 | -0.01 | -0.10 | 0.37 | 0.249 |
| **CHAMP1** | **S476** | -0.06 | -0.15 | -0.11 | 0.07 | -0.06 | 0.27 | 0.121 |
| **CHAMP1** | **S241** | -0.25 | -0.04 | -0.02 | 0.15 | -0.05 | 0.17 | 0.14 |
| **CHAMP1** | **S445** | -0.01 | -0.05 | -0.24 | 0.01 | -0.08 | 0.31 | 0.258 |
| **CHAMP1** | **S196** | -0.46 | -0.05 | 0.18 | 0.13 | 0.27 | -0.20 | 0.489 |
| **CHAMP1** | **S282; S297** | 0.12 | -0.10 | -0.29 | 0.07 | 0.03 | 0.13 | 0.238 |
| **CHAMP1** | **S382; S386** | -0.01 | -0.08 | -0.15 | 0.04 | -0.02 | 0.18 | 0.113 |
| **CHAMP1** | **S653; S** | -0.10 | -0.10 | -0.03 | 0.05 | -0.06 | 0.21 | 0.162 |
| **CHAMP1** | **S108** | 0.06 | 0.03 | -0.36 | 0.41 | 0.00 | -0.28 | 0.607 |
| **CHAMP1** | **S282** | -0.05 | -0.08 | -0.08 | 0.13 | -0.10 | 0.15 | 0.176 |
| **CHAMP1** | **S416** | 0.00 | -0.10 | -0.09 | 0.04 | -0.07 | 0.20 | 0.233 |
| **CHAMP1** | **S651; S652; S653** | 0.03 | -0.07 | -0.12 | 0.05 | -0.08 | 0.18 | 0.298 |
| **CHAMP1** | **S653; S** | -0.09 | -0.06 | -0.02 | 0.19 | -0.21 | 0.15 | 0.492 |
| **CHAMP1** | **S204; S214** | 0.06 | -0.07 | -0.13 | -0.17 | 0.09 | 0.19 | 0.543 |
| **CHAMP1** | **S459** | 0.01 | 0.03 | -0.12 | 0.04 | 0.01 | 0.03 | 0.333 |
| **CHAMP1** | **S432; S436** | 0.03 | -0.04 | -0.06 | 0.09 | -0.12 | 0.08 | 0.616 |
| **CHAMP1** | **S204; S214** | 0.06 | 0.01 | -0.12 | -0.03 | 0.02 | 0.06 | 0.591 |
| **CHAMP1** | **S627; S632; S633** | 0.02 | -0.04 | -0.02 | -0.10 | 0.05 | 0.09 | 0.71 |
| **CHAMP1** | **S507** | 0.13 | -0.05 | -0.10 | -0.04 | -0.15 | 0.17 | 0.992 |
| **CHAMP1** | **S427; S432** | -0.07 | 0.15 | -0.09 | 0.05 | -0.20 | 0.12 | 0.935 |
| **CHAMP1** | **S214** | 0.01 | 0.03 | 0.01 | 0.02 | -0.05 | -0.02 | 0.231 |
| **CHAMP1** | **S427** | 0.10 | 0.01 | -0.05 | -0.09 | -0.13 | 0.15 | 0.697 |
| **CHAMP1** | **S653** | 0.07 | 0.05 | -0.03 | 0.00 | -0.04 | -0.04 | 0.159 |
| **CHAMP1** | **S627** | 0.16 | 0.00 | -0.05 | -0.01 | -0.06 | -0.05 | 0.311 |
| **CHAMP1** | **S108** | 0.13 | 0.06 | -0.03 | -0.12 | 0.07 | -0.13 | 0.237 |
| **CHAMP1** | **S386** | 0.16 | 0.05 | -0.01 | -0.06 | -0.16 | -0.01 | 0.093 |
| **CHAMP1** | **S108** | -0.09 | 0.24 | 0.06 | 0.01 | -0.10 | -0.15 | 0.246 |
| **CHAMP1** | **S436** | 0.14 | 0.05 | 0.04 | -0.19 | -0.13 | 0.06 | 0.106 |
| **CHAMP1** | **S542** | 0.01 | 0.06 | 0.18 | -0.17 | -0.05 | -0.06 | 0.046 |
| **CHAMP1** | **S/Y** | 0.14 | 0.04 | 0.08 | -0.15 | -0.03 | -0.10 | 0.016 |
| **CHAMP1** | **T/S** | 0.02 | 0.09 | 0.15 | 0.02 | 0.00 | -0.32 | 0.189 |
| **CHAMP1** | **S603** | -0.26 | 0.01 | 0.47 | -0.43 | 0.00 | 0.05 | 0.482 |
| **CHAMP1** | **S632** | 0.13 | -0.02 | 0.19 | -0.12 | -0.09 | -0.12 | 0.028 |
| **CHAMP1** | **S405** | 0.15 | 0.06 | 0.15 | -0.32 | 0.07 | -0.18 | 0.084 |
| **CHAMP1** | **S632; S633** | 0.14 | 0.19 | 0.09 | -0.16 | -0.15 | -0.15 | 7E-04 |
| **CHAMP1** | **S87** | 0.20 | 0.21 | 0.10 | -0.26 | -0.09 | -0.24 | 0.005 |
| **CHAMP1** | **S736** | -0.15 | 0.32 | 0.31 | -0.28 | -0.05 | -0.30 | 0.1 |
| **CHAMP1** | **S376** | 0.18 | 0.22 | 0.14 | -0.39 | -0.03 | -0.22 | 0.02 |
| **CHAMP1** | **S405** | 0.27 | 0.24 | 0.01 | -0.23 | 0.14 | -0.65 | 0.158 |
| **CHAMP1** | **S459** | 0.14 | 0.26 | 0.19 | -0.21 | -0.05 | -0.45 | 0.025 |

| | | | | | | | | |
|---|---|---|---|---|---|---|---|---|
| **CHAMP1** | **S603** | 0.25 | 0.14 | 0.26 | -0.28 | -0.12 | -0.40 | 0.006 |
| **CHAMP1** | **S87** | 0.38 | 0.10 | 0.23 | -0.46 | -0.13 | -0.29 | 0.014 |
| **CHAMP1** | **S603** | 0.24 | 0.14 | 0.35 | -0.41 | -0.14 | -0.35 | 0.006 |
| **CHCHD3** | **S29** | -0.15 | -0.66 | -0.02 | -0.16 | 0.07 | 0.62 | 0.21 |
| **CHCHD3** | **S50** | 0.13 | 0.05 | 0.07 | -0.15 | -0.05 | -0.06 | 0.014 |
| **CHCHD3** | **S29** | 0.21 | 0.02 | 0.14 | -0.02 | -0.10 | -0.32 | 0.06 |
| **CHCHD6** | **S13** | -0.17 | 0.10 | -0.29 | 0.07 | 0.04 | 0.19 | 0.159 |
| **CHCHD6** | **S126** | 0.03 | 0.21 | -0.33 | 0.12 | -0.04 | -0.05 | 0.794 |
| **CHCHD6** | **S72** | 0.05 | -0.34 | 0.28 | 0.04 | 0.00 | -0.09 | 0.95 |
| **CHCHD6** | **S43** | 0.26 | 0.05 | -0.43 | -0.39 | -0.45 | 0.61 | 0.927 |
| **CHCHD6** | **S** | 0.02 | 0.10 | 0.06 | -0.04 | 0.07 | -0.22 | 0.225 |
| **CHCHD6** | **T153** | 0.23 | 0.09 | 0.03 | -0.08 | -0.01 | -0.31 | 0.087 |
| **CHCHD6** | **S74** | 0.07 | 0.23 | 0.27 | -0.70 | 0.13 | -0.22 | 0.14 |
| **CHD1** | **S215; S216** | -0.13 | -0.23 | -0.43 | 0.07 | 0.12 | 0.44 | 0.032 |
| **CHD1** | **S1081** | -0.13 | -0.21 | -0.25 | 0.38 | 0.05 | 0.06 | 0.035 |
| **CHD1** | **S1081** | -0.19 | -0.15 | -0.06 | 0.04 | 0.21 | 0.11 | 0.015 |
| **CHD1** | **T/S** | 0.01 | -0.14 | -0.17 | 0.06 | 0.15 | 0.06 | 0.04 |
| **CHD1** | **S1689** | 0.11 | 0.10 | -0.03 | 0.00 | -0.17 | -0.03 | 0.144 |
| **CHD1** | **S1677** | 0.02 | 0.14 | 0.10 | -0.21 | 0.01 | -0.09 | 0.064 |
| **CHD1** | **S/Y** | 0.27 | 0.12 | -0.09 | -0.12 | -0.08 | -0.13 | 0.12 |
| **CHD1** | **T1700** | 0.09 | 0.00 | 0.22 | -0.14 | -0.34 | 0.10 | 0.18 |
| **CHD1** | **S1096; S1100** | 0.23 | 0.27 | 0.09 | -0.12 | -0.02 | -0.63 | 0.082 |
| **CHD1** | **S/T** | 0.34 | 0.33 | 0.11 | -0.35 | -0.47 | -0.17 | 0.007 |
| **CHD1** | **S1100; S1102** | 0.27 | 0.37 | 0.24 | -0.39 | -0.24 | -0.48 | 0.001 |
| **CHD2** | **S207; S208** | -0.09 | -0.19 | -0.14 | 0.04 | 0.08 | 0.25 | 0.02 |
| **CHD2** | **S1728** | -0.04 | -0.03 | -0.31 | -0.17 | 0.45 | -0.03 | 0.367 |
| **CHD2** | **S1789** | 0.01 | 0.09 | -0.33 | 0.17 | 0.08 | -0.07 | 0.397 |
| **CHD2** | **S1085; S1087** | -0.02 | -0.15 | -0.02 | 0.19 | 0.14 | -0.18 | 0.4 |
| **CHD2** | **S1365** | -0.03 | -0.04 | -0.11 | 0.07 | -0.03 | 0.12 | 0.081 |
| **CHD2** | **S156** | -0.22 | 0.12 | -0.01 | 0.16 | 0.26 | -0.42 | 0.877 |
| **CHD2** | **S1085; S1087; S1091** | 0.01 | -0.01 | -0.04 | -0.02 | 0.11 | -0.05 | 0.622 |
| **CHD2** | **T240; S242** | -0.07 | 0.13 | -0.09 | -0.04 | 0.11 | -0.06 | 0.88 |
| **CHD2** | **T/S** | 0.26 | -0.37 | 0.01 | 0.41 | -0.17 | -0.30 | 0.967 |
| **CHD2** | **S1373** | 0.19 | 0.17 | 0.09 | -0.38 | 0.02 | -0.18 | 0.053 |
| **CHD2** | **S156; S165; S168** | 0.25 | 0.21 | 0.02 | -0.27 | 0.13 | -0.49 | 0.131 |
| **CHD2** | **S242** | 0.13 | 0.37 | -0.06 | -0.25 | 0.21 | -0.63 | 0.246 |
| **CHD2** | **S156; S168** | 0.11 | 0.29 | 0.12 | 0.02 | -0.03 | -0.68 | 0.157 |
| **CHD2** | **S156; S165; S168** | 0.27 | 0.24 | 0.06 | -0.28 | 0.11 | -0.58 | 0.108 |
| **CHD2** | **S156; S168** | 0.25 | 0.13 | 0.22 | -0.04 | -0.15 | -0.55 | 0.049 |
| **CHD2** | **S156** | 0.19 | 0.31 | 0.06 | -0.05 | 0.03 | -0.76 | 0.157 |
| **CHD3** | **S/T** | 0.00 | -0.13 | -0.18 | 0.20 | 0.14 | -0.05 | 0.098 |
| **CHD3** | **S712** | 0.06 | -0.12 | -0.18 | 0.09 | 0.13 | -0.02 | 0.152 |
| **CHD3** | **S1601; S1605** | -0.22 | -0.06 | 0.12 | -0.18 | -0.17 | 0.40 | 0.768 |
| **CHD3** | **S597** | 0.07 | -0.14 | 0.04 | -0.03 | 0.03 | 0.03 | 0.77 |
| **CHD3** | **S597** | 0.06 | 0.01 | -0.08 | 0.03 | -0.03 | 0.00 | 0.937 |
| **CHD3** | **S1601** | 0.03 | 0.00 | 0.01 | 0.05 | -0.22 | 0.11 | 0.767 |
| **CHD3** | **S88; S90** | -0.07 | 0.08 | 0.04 | -0.04 | 0.05 | -0.07 | 0.558 |
| **CHD3** | **T1557; T/S** | 0.18 | -0.03 | -0.07 | -0.27 | -0.14 | 0.26 | 0.687 |

| | | | | | | | | |
|---|---|---|---|---|---|---|---|---|
| CHD3 | S1549; T1557 | 0.13 | -0.03 | 0.11 | -0.30 | -0.25 | 0.25 | 0.41 |
| CHD4 | S1570; S1576 | -1.16 | -1.17 | -1.27 | 0.61 | 0.74 | 0.59 | 6E-06 |
| CHD4 | T1653 | -1.22 | -0.73 | -0.96 | 0.53 | 0.24 | 0.87 | 0.003 |
| CHD4 | S1308 | -0.93 | -0.63 | -0.86 | 0.49 | 0.36 | 0.66 | 5E-04 |
| CHD4 | S1570 | -0.57 | -0.61 | -0.64 | 0.45 | 0.53 | 0.29 | 1E-04 |
| CHD4 | T1653 | -0.25 | -0.60 | -0.99 | 0.37 | 0.69 | 0.12 | 0.02 |
| CHD4 | S1570 | -0.48 | -0.61 | -0.55 | 0.62 | 0.40 | 0.13 | 0.003 |
| CHD4 | S1349 | -0.03 | -0.75 | -0.81 | 0.36 | 0.39 | 0.34 | 0.024 |
| CHD4 | S1349 | -0.57 | -0.49 | -0.47 | 0.28 | 0.33 | 0.51 | 3E-04 |
| CHD4 | S325; S/Y/T | -0.28 | -0.28 | -0.68 | 0.69 | -0.30 | 0.39 | 0.106 |
| CHD4 | S1308 | -0.45 | -0.25 | -0.38 | 0.23 | 0.28 | 0.34 | 6E-04 |
| CHD4 | T1549; T1553; S/T | -0.26 | -0.46 | -0.39 | 0.40 | -0.12 | 0.52 | 0.035 |
| CHD4 | S1544; T1545; T1553 | -0.26 | -0.35 | -0.38 | 0.39 | -0.09 | 0.45 | 0.029 |
| CHD4 | S1602 | -0.52 | -0.12 | -0.29 | 0.15 | 0.37 | 0.22 | 0.014 |
| CHD4 | S1602 | -0.40 | -0.29 | -0.18 | 0.21 | 0.21 | 0.30 | 0.002 |
| CHD4 | S1602 | -0.46 | -0.43 | 0.05 | 0.14 | 0.24 | 0.28 | 0.042 |
| CHD4 | T1679 | -0.10 | -0.46 | -0.26 | 0.14 | 0.13 | 0.38 | 0.02 |
| CHD4 | S1602 | -0.45 | -0.22 | -0.12 | 0.23 | -0.03 | 0.42 | 0.045 |
| CHD4 | T703 | -0.21 | -0.25 | -0.30 | 0.16 | 0.48 | -0.05 | 0.047 |
| CHD4 | T1679 | 0.09 | -0.34 | -0.48 | -0.27 | 0.36 | 0.39 | 0.221 |
| CHD4 | Y/T | -0.44 | -0.03 | -0.02 | 0.11 | 0.04 | 0.25 | 0.122 |
| CHD4 | S/Y/T | -0.14 | -0.14 | -0.19 | 0.17 | -0.02 | 0.25 | 0.025 |
| CHD4 | S1531; S1535 | -0.03 | -0.15 | -0.19 | 0.20 | 0.09 | 0.05 | 0.022 |
| CHD4 | T703 | -0.15 | -0.10 | -0.11 | 0.16 | 0.09 | 0.08 | 0.001 |
| CHD4 | S428 | -0.11 | -0.15 | -0.09 | 0.15 | -0.06 | 0.22 | 0.061 |
| CHD4 | T367 | -0.06 | -0.12 | -0.08 | 0.01 | 0.08 | 0.16 | 0.022 |
| CHD4 | T367 | -0.18 | 0.02 | -0.09 | -0.01 | -0.13 | 0.33 | 0.376 |
| CHD4 | S/Y/T | 0.09 | -0.24 | -0.05 | 0.18 | -0.06 | 0.04 | 0.359 |
| CHD4 | T1540; T1553 | -0.07 | -0.02 | -0.10 | 0.04 | -0.02 | 0.15 | 0.095 |
| CHD4 | S428 | -0.09 | 0.00 | -0.08 | 0.12 | -0.12 | 0.15 | 0.28 |
| CHD4 | T1549; T1553 | -0.03 | -0.02 | -0.09 | 0.01 | 0.00 | 0.13 | 0.132 |
| CHD4 | T367 | 0.03 | -0.05 | -0.08 | -0.04 | 0.02 | 0.12 | 0.306 |
| CHD4 | Y/T | 0.05 | -0.07 | -0.05 | -0.03 | 0.05 | 0.04 | 0.368 |
| CHD4 | T1545 | 0.03 | 0.02 | -0.06 | 0.01 | 0.01 | -0.02 | 0.977 |
| CHD4 | S103; S105; S108 | 0.02 | 0.02 | -0.04 | -0.06 | -0.08 | 0.14 | 0.97 |
| CHD4 | S85; S103; S108 | 0.04 | 0.03 | -0.04 | 0.04 | -0.12 | 0.03 | 0.649 |
| CHD4 | S85 | 0.00 | 0.04 | 0.02 | 0.07 | -0.20 | 0.05 | 0.603 |
| CHD4 | S310; S319; S/Y/T | 0.07 | 0.08 | -0.07 | 0.04 | -0.12 | 0.00 | 0.474 |
| CHD4 | S310; S319 | 0.13 | -0.13 | 0.08 | -0.02 | -0.08 | 0.01 | 0.517 |
| CHD4 | S103; S105; S108 | 0.10 | 0.03 | -0.03 | 0.01 | -0.08 | -0.04 | 0.198 |
| CHD4 | S1648 | -0.07 | 0.02 | 0.18 | -0.13 | -0.03 | 0.02 | 0.349 |
| CHD4 | S85 | -0.04 | 0.10 | 0.07 | -0.10 | -0.11 | 0.06 | 0.232 |
| CHD4 | T1553 | 0.01 | 0.14 | 0.02 | -0.07 | 0.01 | -0.12 | 0.101 |
| CHD4 | S/Y/T | 0.02 | 0.09 | 0.03 | -0.39 | -0.19 | 0.33 | 0.58 |
| CHD4 | S310; S319; S/Y/T | 0.30 | 0.20 | -0.37 | -0.02 | -0.31 | 0.07 | 0.602 |
| CHD4 | S1535 | 0.00 | 0.08 | 0.13 | -0.24 | 0.14 | -0.15 | 0.276 |
| CHD4 | S85; S86 | 0.10 | 0.06 | 0.08 | -0.15 | -0.01 | -0.10 | 0.018 |
| CHD4 | S515; S531 | 0.10 | 0.11 | 0.08 | -0.20 | 0.03 | -0.15 | 0.043 |

| | | | | | | | | |
|---|---|---|---|---|---|---|---|---|
| **CHD4** | **S310; S/Y/T** | 0.05 | 0.13 | 0.12 | -0.06 | -0.24 | -0.03 | 0.044 |
| **CHD4** | **S319** | 0.24 | 0.14 | -0.04 | -0.20 | -0.09 | -0.10 | 0.048 |
| **CHD4** | **S103; S105** | 0.18 | 0.19 | 0.04 | -0.26 | -0.10 | -0.10 | 0.015 |
| **CHD4** | **S103; S/T/Y** | 0.19 | 0.20 | 0.13 | -0.30 | -0.07 | -0.23 | 0.006 |
| **CHD4** | **S531** | 0.19 | 0.00 | 0.31 | -0.55 | -0.09 | -0.01 | 0.115 |
| **CHD4** | **S1535** | 0.24 | 0.19 | 0.18 | -0.29 | 0.00 | -0.47 | 0.03 |
| **CHD4** | **S85; S86** | 0.09 | 0.45 | 0.03 | -0.75 | 0.01 | -0.07 | 0.171 |
| **CHD4** | **S1535** | 0.21 | 0.21 | 0.24 | -0.30 | -0.01 | -0.51 | 0.028 |
| **CHD4** | **S105; S108** | 0.31 | 0.29 | 0.26 | -0.26 | -0.17 | -0.68 | 0.014 |
| **CHD4** | **S108** | 0.48 | 0.33 | 0.18 | -0.34 | -0.09 | -1.04 | 0.05 |
| **CHD6** | **S1673** | -0.19 | -0.44 | -0.26 | 0.40 | -0.08 | 0.37 | 0.036 |
| **CHD6** | **S2680** | -0.31 | -0.37 | 0.00 | 0.35 | -0.29 | 0.41 | 0.2 |
| **CHD6** | **S1723** | -0.25 | -0.15 | 0.07 | 0.03 | 0.09 | 0.17 | 0.111 |
| **CHD7** | **S782** | -0.08 | -0.41 | 0.03 | 0.14 | 0.04 | 0.20 | 0.119 |
| **CHD7** | **S2559** | 0.00 | -0.11 | 0.14 | 0.08 | -0.03 | -0.11 | 0.755 |
| **CHD7** | **S2272; S2275** | 0.05 | 0.27 | 0.02 | 0.08 | -0.11 | -0.38 | 0.18 |
| **CHD8** | **S2008** | -0.09 | -0.04 | -0.36 | 0.27 | -0.03 | 0.16 | 0.088 |
| **CHD8** | **S2519** | -0.05 | -0.13 | -0.17 | 0.02 | 0.05 | 0.24 | 0.054 |
| **CHD8** | **S2069; S2071** | 0.02 | 0.01 | -0.25 | -0.01 | -0.12 | 0.29 | 0.448 |
| **CHD8** | **S2557** | 0.11 | -0.27 | -0.04 | -0.06 | 0.04 | 0.17 | 0.439 |
| **CHD8** | **S2046** | 0.02 | -0.16 | 0.01 | -0.15 | 0.14 | 0.12 | 0.529 |
| **CHD8** | **S/T/Y** | -0.03 | 0.05 | -0.04 | -0.12 | -0.10 | 0.22 | 0.971 |
| **CHD8** | **S/Y** | 0.29 | -0.33 | 0.00 | -0.30 | -0.12 | 0.33 | 0.96 |
| **CHD8** | **S1424** | 0.20 | 0.01 | 0.29 | -0.28 | 0.00 | -0.32 | 0.047 |
| **CHD8** | **S1995** | 0.15 | 0.17 | 0.18 | -0.20 | -0.01 | -0.39 | 0.029 |
| **CHD8** | **S549; S550; S562** | 0.28 | 0.21 | 0.23 | -0.24 | -0.02 | -0.69 | 0.049 |
| **CHD9** | **S611** | -0.14 | 0.08 | -0.07 | -0.03 | 0.20 | -0.07 | 0.516 |
| **CHD9** | **S1754** | 0.12 | -0.05 | 0.06 | -0.11 | -0.08 | 0.03 | 0.227 |
| **CHD9** | **S550** | 0.06 | 0.04 | 0.07 | -0.24 | 0.02 | 0.03 | 0.245 |
| **CHD9** | **S2303** | 0.10 | 0.15 | 0.01 | -0.04 | -0.18 | -0.06 | 0.04 |
| **CHD9** | **T2113** | 0.20 | 0.05 | 0.12 | -0.13 | 0.02 | -0.31 | 0.067 |
| **CHD9** | **S1468; S1472** | 0.16 | 0.19 | 0.12 | -0.11 | -0.10 | -0.34 | 0.013 |
| **CHD9** | **S2079** | 0.16 | 0.18 | 0.15 | -0.18 | 0.05 | -0.48 | 0.078 |
| **CHD9** | **S2058; S2059** | 0.56 | -0.08 | 0.12 | -0.49 | -0.16 | -0.19 | 0.089 |
| **CHD9** | **S2058; S2059** | 0.20 | 0.32 | 0.22 | -0.30 | -0.09 | -0.53 | 0.014 |
| **CHD9** | **S1472** | 0.26 | 0.28 | 0.20 | -0.30 | -0.05 | -0.61 | 0.026 |
| **CHD9** | **S2058; S2059** | 0.32 | 0.31 | 0.27 | -0.46 | -0.06 | -0.68 | 0.018 |
| **CHERP** | **S186** | -0.24 | -0.23 | 0.05 | 0.07 | -0.17 | 0.41 | 0.284 |
| **CHERP** | **S815; S817; T819** | -0.02 | -0.10 | 0.05 | -0.08 | -0.02 | 0.16 | 0.61 |
| **CHERP** | **S815; S817; T819** | 0.01 | 0.06 | 0.04 | 0.00 | -0.07 | -0.04 | 0.058 |
| **CHERP** | **S806** | 0.03 | 0.40 | 0.27 | -0.32 | 0.16 | -0.88 | 0.148 |
| **CHMP1A** | **S101** | -0.40 | -0.59 | -0.69 | 0.63 | 0.22 | 0.31 | 0.003 |
| **CHMP2A** | **S203** | -0.33 | -0.19 | -0.27 | 0.21 | 0.08 | 0.36 | 0.007 |
| **CHMP2A** | **S203** | -0.09 | -0.17 | -0.24 | 0.09 | -0.12 | 0.43 | 0.149 |
| **CHMP2B** | **S187** | -0.04 | -0.35 | -0.58 | 0.36 | 0.18 | 0.20 | 0.027 |
| **CHMP3** | **S200** | -0.12 | -0.06 | 0.07 | -0.06 | 0.04 | 0.12 | 0.394 |
| **CHMP3** | **S200; S215** | -0.25 | -0.09 | 0.21 | -0.14 | 0.03 | 0.17 | 0.722 |
| **CHMP3** | **S200** | -0.02 | 0.08 | -0.11 | 0.05 | 0.05 | -0.05 | 0.657 |

| | | | | | | | | |
|---|---|---|---|---|---|---|---|---|
| CHMP3 | S200 | 0.04 | 0.02 | 0.09 | -0.10 | -0.03 | -0.04 | 0.027 |
| CHMP3 | S200 | 0.08 | 0.15 | 0.02 | -0.04 | 0.09 | -0.36 | 0.247 |
| CHP1 | T7 | -0.40 | -0.25 | -0.07 | 0.11 | -0.08 | 0.51 | 0.1 |
| CHP1 | T7 | -0.22 | -0.24 | -0.19 | 0.28 | -0.05 | 0.31 | 0.026 |
| CHTF18 | S63 | -0.47 | -0.39 | 0.10 | 0.16 | 0.15 | 0.29 | 0.07 |
| CHTOP | T242 | 0.12 | -0.14 | -0.26 | 0.13 | 0.11 | 0.00 | 0.227 |
| CHTOP | T246 | 0.11 | -0.04 | -0.35 | 0.26 | 0.05 | -0.10 | 0.399 |
| CHTOP | T242 | -0.10 | -0.03 | -0.06 | 0.16 | -0.14 | 0.14 | 0.285 |
| CHTOP | T33 | 0.20 | 0.20 | -0.18 | -0.22 | 0.13 | -0.21 | 0.37 |
| CIAPIN1 | S | 0.01 | 0.29 | -0.12 | 0.11 | 0.18 | -0.63 | 0.569 |
| CIC | S904 | -0.16 | 0.04 | -0.27 | 0.40 | 0.22 | -0.40 | 0.476 |
| CIC | S1246 | -0.12 | -0.01 | 0.01 | 0.13 | -0.09 | 0.07 | 0.359 |
| CIC | T1291 | 0.02 | -0.01 | -0.08 | 0.04 | -0.01 | 0.03 | 0.236 |
| CIC | S191 | 0.10 | -0.14 | 0.02 | -0.21 | 0.07 | 0.13 | 0.983 |
| CIC | S1409 | -0.22 | 0.02 | 0.33 | -0.18 | 0.13 | -0.16 | 0.58 |
| CIC | S173 | -0.01 | 0.32 | 0.23 | 0.08 | -0.14 | -0.68 | 0.16 |
| CINP | S | 0.16 | 0.31 | -0.06 | -0.23 | -0.10 | -0.16 | 0.062 |
| CIPC | S/T | -0.10 | -0.03 | 0.01 | -0.24 | 0.15 | 0.17 | 0.65 |
| CIR1 | S365 | 0.13 | 0.05 | 0.19 | -0.03 | -0.17 | -0.21 | 0.019 |
| CIR1 | S353 | 0.10 | 0.30 | -0.04 | -0.18 | -0.07 | -0.18 | 0.065 |
| CIR1 | Y/S/T | 0.23 | 0.05 | 0.11 | -0.19 | -0.15 | -0.11 | 0.008 |
| CIR1 | S365 | 0.11 | 0.20 | 0.09 | -0.20 | -0.10 | -0.14 | 0.003 |
| CIR1 | S202 | 0.14 | 0.09 | 0.18 | -0.15 | -0.06 | -0.25 | 0.008 |
| CIR1 | S394 | 0.58 | 0.39 | 0.01 | -0.44 | -0.14 | -0.89 | 0.041 |
| CIRBP | S163 | 0.04 | 0.29 | 0.05 | -0.23 | 0.07 | -0.30 | 0.112 |
| CIRBP | T169 | 0.07 | 0.37 | 0.11 | -0.08 | -0.22 | -0.37 | 0.033 |
| CIT | S480 | -1.02 | -0.03 | -0.01 | -0.29 | 0.56 | 0.32 | 0.262 |
| CIT | S440 | -0.04 | -0.10 | 0.15 | -0.02 | -0.03 | 0.02 | 0.864 |
| CIT | S582 | 0.09 | 0.17 | 0.30 | -0.17 | -0.11 | -0.39 | 0.016 |
| CIZ1 | S/T | -0.05 | -0.26 | -0.14 | 0.23 | -0.08 | 0.23 | 0.083 |
| CIZ1 | T225 | -0.08 | -0.15 | -0.06 | -0.02 | 0.14 | 0.16 | 0.041 |
| CIZ1 | S350 | 0.01 | -0.14 | -0.14 | -0.08 | 0.21 | 0.11 | 0.159 |
| CIZ1 | T244 | 0.09 | -0.13 | -0.13 | -0.09 | 0.05 | 0.17 | 0.378 |
| CIZ1 | S209 | -0.01 | 0.02 | 0.01 | 0.03 | -0.27 | 0.18 | 0.847 |
| CIZ1 | S838 | 0.06 | 0.06 | 0.37 | -0.27 | 0.00 | -0.33 | 0.065 |
| CKAP2 | T579 | -0.22 | -0.33 | -0.28 | 0.16 | 0.11 | 0.40 | 0.007 |
| CKAP2 | T579 | -0.21 | -0.22 | -0.17 | 0.10 | -0.01 | 0.40 | 0.042 |
| CKAP2 | T | -0.14 | -0.46 | 0.06 | 0.10 | 0.22 | 0.12 | 0.1 |
| CKAP2 | S534 | -0.10 | -0.12 | -0.31 | 0.18 | -0.01 | 0.27 | 0.036 |
| CKAP2 | S534 | -0.16 | -0.13 | -0.17 | 0.12 | 0.28 | 0.01 | 0.022 |
| CKAP2 | S534 | -0.12 | -0.15 | -0.05 | 0.07 | 0.29 | -0.09 | 0.154 |
| CKAP4 | S17 | 0.06 | -0.06 | -0.05 | 0.15 | 0.00 | -0.12 | 0.759 |
| CKMT1A | T410 | 0.33 | -0.46 | -0.13 | 0.24 | -0.17 | 0.05 | 0.645 |
| CKMT1A | S174 | 0.01 | 0.12 | 0.08 | -0.15 | -0.03 | -0.04 | 0.052 |
| CLASP1 | S | -0.10 | 0.09 | -0.21 | -0.19 | 0.22 | 0.12 | 0.462 |
| CLASP1 | S572 | 0.14 | -0.04 | -0.06 | 0.04 | -0.19 | 0.09 | 0.763 |
| CLASP1 | S | -0.07 | -0.03 | 0.19 | -0.13 | 0.05 | -0.03 | 0.508 |
| CLASP1 | S1196 | 0.10 | 0.23 | 0.09 | -0.19 | 0.02 | -0.33 | 0.052 |

| | | | | | | | | |
|---|---|---|---|---|---|---|---|---|
| CLASP1 | S572 | 0.35 | 0.09 | 0.16 | -0.32 | -0.11 | -0.30 | 0.013 |
| CLASP2 | S | -0.29 | -0.07 | 0.05 | -0.28 | 0.36 | 0.12 | 0.461 |
| CLASRP | S454; S456 | 0.18 | -0.31 | -0.16 | -0.12 | 0.03 | 0.29 | 0.429 |
| CLASRP | S101 | 0.08 | 0.10 | 0.02 | 0.00 | -0.12 | -0.10 | 0.038 |
| CLASRP | S101 | -0.04 | 0.00 | 0.23 | 0.10 | 0.00 | -0.37 | 0.399 |
| CLASRP | S294 | 0.30 | 0.21 | 0.09 | -0.14 | -0.15 | -0.43 | 0.018 |
| CLASRP | S547 | 0.23 | 0.32 | 0.32 | -0.27 | -0.32 | -0.52 | 0.001 |
| CLASRP | S294 | 0.46 | 0.31 | 0.20 | -0.37 | -0.16 | -0.83 | 0.022 |
| CLCC1 | T482 | 0.29 | -0.31 | 0.31 | -0.34 | 0.33 | -0.54 | 0.448 |
| CLCC1 | S509 | 0.47 | 0.66 | -0.07 | -0.74 | -0.13 | -0.82 | 0.041 |
| CLCN2 | S731 | 0.06 | 0.09 | 0.03 | -0.10 | 0.02 | -0.11 | 0.059 |
| CLCN2 | S712 | 0.23 | 0.16 | 0.19 | -0.26 | -0.08 | -0.34 | 0.006 |
| CLDN23 | S/T | 0.11 | 0.16 | 0.22 | -0.55 | 0.09 | -0.17 | 0.117 |
| CLDN3 | S209 | -0.19 | -0.29 | -0.24 | 0.25 | 0.26 | 0.10 | 0.002 |
| CLGN | S576 | -0.28 | -0.12 | 0.14 | 0.14 | -0.01 | 0.08 | 0.305 |
| CLIC1 | S156 | 0.09 | 0.10 | -0.12 | -0.38 | 0.18 | 0.07 | 0.748 |
| CLIC3 | S49 | 0.09 | -0.33 | -0.18 | 0.23 | 0.16 | -0.04 | 0.159 |
| CLINT1 | T308 | -0.13 | 0.10 | -0.44 | 0.25 | -0.08 | 0.20 | 0.216 |
| CLINT1 | T308 | 0.18 | 0.04 | 0.23 | 0.03 | -0.53 | -0.09 | 0.125 |
| CLIP1 | T182 | -0.14 | 0.08 | -0.06 | 0.22 | 0.13 | -0.30 | 0.769 |
| CLK1 | S37 | 0.06 | -0.25 | -0.79 | 0.29 | -0.12 | 0.49 | 0.15 |
| CLK1 | S140 | -0.19 | -0.27 | -0.25 | 0.27 | -0.07 | 0.37 | 0.036 |
| CLK1 | T138; S140 | -0.02 | 0.04 | -0.06 | 0.02 | 0.00 | 0.01 | 0.483 |
| CLK1 | T/S | 0.16 | 0.00 | -0.06 | -0.05 | -0.10 | 0.04 | 0.389 |
| CLK1 | S37 | 0.26 | 0.32 | 0.13 | -0.07 | 0.03 | -1.02 | 0.157 |
| CLK1 | S/Y | 0.42 | 0.39 | 0.25 | -0.05 | -0.35 | -1.26 | 0.069 |
| CLK2 | S50 | 0.30 | 0.29 | -0.19 | -0.35 | 0.18 | -0.41 | 0.26 |
| CLK3 | S215 | 0.10 | 0.32 | 0.14 | -0.23 | -0.25 | -0.18 | 0.004 |
| CLK3 | S226 | 0.19 | 0.29 | 0.19 | -0.10 | -0.23 | -0.49 | 0.014 |
| CLK3 | S157 | 0.32 | 0.31 | 0.14 | -0.11 | -0.28 | -0.61 | 0.02 |
| CLK4 | S33 | -0.23 | -0.15 | -0.48 | 0.37 | -0.04 | 0.33 | 0.038 |
| CLK4 | S138 | 0.00 | -0.03 | -0.12 | 0.05 | 0.03 | 0.06 | 0.062 |
| CLK4 | S | -0.05 | -0.05 | 0.19 | -0.16 | -0.24 | 0.24 | 0.645 |
| CLMN | S301 | -0.05 | -0.03 | -0.14 | 0.15 | -0.01 | 0.07 | 0.07 |
| CLMN | S402 | -0.01 | -0.10 | -0.05 | -0.08 | 0.07 | 0.16 | 0.227 |
| CLMN | S635 | 0.19 | -0.15 | -0.21 | -0.24 | -0.27 | 0.50 | 0.854 |
| CLMN | S635 | 0.02 | -0.01 | -0.13 | -0.27 | -0.21 | 0.48 | 0.869 |
| CLMN | S635 | -0.01 | -0.03 | 0.21 | -0.14 | -0.07 | 0.01 | 0.238 |
| CLN3 | S12; S14 | 0.04 | 0.16 | 0.14 | -0.24 | -0.14 | 0.00 | 0.037 |
| CLN3 | S12 | 0.09 | 0.10 | 0.23 | -0.25 | -0.17 | -0.05 | 0.016 |
| CLN3 | S14 | -0.12 | 0.14 | 0.66 | -0.42 | -0.25 | -0.32 | 0.077 |
| CLN6 | S31 | -0.02 | -0.02 | 0.06 | 0.08 | 0.09 | -0.20 | 0.856 |
| CLNS1A | S102 | 0.03 | -0.05 | 0.13 | -0.19 | 0.18 | -0.14 | 0.527 |
| CLOCK | S394; T398; S403 | -0.23 | -0.35 | -0.20 | 0.40 | -0.23 | 0.40 | 0.104 |
| CLOCK | S403; S406; S408 | -0.23 | -0.35 | -0.20 | 0.40 | -0.23 | 0.40 | 0.104 |
| CLOCK | S427 | -0.01 | -0.12 | -0.23 | 0.21 | -0.09 | 0.19 | 0.127 |
| CLOCK | S406; S408 | -0.05 | -0.13 | -0.15 | 0.18 | -0.15 | 0.24 | 0.18 |
| CLOCK | S406; S408 | -0.05 | -0.13 | -0.15 | 0.18 | -0.15 | 0.24 | 0.18 |

| | | | | | | | | |
|---|---|---|---|---|---|---|---|---|
| CLOCK | T461 | -0.08 | -0.08 | 0.00 | 0.03 | -0.03 | 0.14 | 0.151 |
| CLSPN | S/T | 0.00 | -0.24 | -0.25 | 0.33 | -0.36 | 0.36 | 0.331 |
| CLSPN | S | -0.11 | -0.15 | -0.04 | -0.13 | 0.11 | 0.27 | 0.202 |
| CLSPN | S762 | 0.06 | 0.14 | 0.00 | -0.08 | -0.09 | -0.04 | 0.039 |
| CLSPN | S/T | 0.24 | 0.06 | 0.06 | -0.09 | 0.04 | -0.38 | 0.13 |
| CLSPN | S1012; S1018; S1020 | -0.15 | 0.29 | 0.13 | -0.33 | -0.76 | 0.48 | 0.49 |
| CLSPN | S1012; S1018; S1020 | -0.15 | 0.29 | 0.13 | -0.33 | -0.76 | 0.48 | 0.49 |
| CLSPN | S846 | 0.18 | 0.16 | 0.13 | -0.23 | -0.16 | -0.14 | 4E-04 |
| CLSPN | S83 | 0.25 | 0.25 | 0.18 | -0.61 | -0.14 | -0.10 | 0.037 |
| CLSTN3 | T951 | -0.06 | 0.26 | -0.64 | -0.90 | 0.34 | 0.48 | 0.822 |
| CLTC | T394 | 0.03 | -0.12 | 0.00 | -0.09 | -0.13 | 0.27 | 0.74 |
| CLTC | S1494 | 0.34 | -0.04 | 0.50 | -0.41 | -0.49 | -0.18 | 0.028 |
| CMTR1 | S51; S53 | 0.07 | 0.07 | 0.07 | -0.09 | -0.24 | 0.09 | 0.186 |
| CMTR1 | S/T | 0.20 | -0.10 | 0.16 | -0.07 | 0.00 | -0.25 | 0.183 |
| CNNM3 | S673 | 0.39 | 0.19 | -0.98 | 0.02 | -0.54 | 0.42 | 0.853 |
| CNNM3 | S453 | 0.01 | 0.14 | -0.08 | 0.04 | 0.01 | -0.13 | 0.59 |
| CNNM4 | S664 | -0.09 | -0.14 | -0.19 | 0.10 | -0.08 | 0.34 | 0.109 |
| CNOT1 | Y/S | 0.07 | -0.26 | 0.00 | -0.10 | -0.02 | 0.26 | 0.506 |
| COBL | S235 | 0.09 | -0.11 | -0.02 | 0.02 | 0.20 | -0.21 | 0.9 |
| COBL | S/T | 0.03 | -0.15 | 0.14 | 0.08 | -0.10 | -0.02 | 0.822 |
| COIL | T122 | -0.01 | -0.01 | -0.11 | 0.01 | -0.04 | 0.15 | 0.273 |
| COIL | S566 | -0.04 | -0.07 | 0.02 | -0.06 | 0.01 | 0.13 | 0.384 |
| COL15A1 | S1052 | 0.07 | -0.42 | -0.38 | 0.45 | -0.12 | 0.19 | 0.143 |
| COPA | S173 | -0.31 | 0.04 | 0.16 | 0.20 | 0.10 | -0.27 | 0.825 |
| COPA | S173 | 0.14 | -0.12 | 0.12 | -0.16 | 0.17 | -0.19 | 0.48 |
| COPB2 | S859 | 0.10 | 0.16 | 0.00 | -0.10 | -0.01 | -0.19 | 0.054 |
| COPB2 | S/T | 0.24 | 0.28 | 0.21 | -0.07 | -0.09 | -0.86 | 0.088 |
| COPB2 | S859 | 0.13 | 0.50 | 0.30 | -0.38 | 0.00 | -1.02 | 0.069 |
| COPE | S99 | 0.06 | -0.06 | -0.17 | -0.20 | 0.13 | 0.20 | 0.52 |
| COPG1 | S/T | -0.11 | -0.22 | 0.24 | -0.12 | 0.09 | 0.07 | 0.785 |
| CPA4 | T90 | 0.14 | 0.02 | 0.21 | -0.26 | 0.02 | -0.18 | 0.061 |
| CPA4 | T90 | 0.14 | 0.02 | 0.21 | -0.26 | 0.02 | -0.18 | 0.061 |
| CPA4 | T90 | 0.14 | 0.02 | 0.21 | -0.26 | 0.02 | -0.18 | 0.061 |
| CPD | T1368 | -0.01 | 0.02 | 0.04 | 0.09 | -0.21 | 0.05 | 0.71 |
| CPD | T1368 | 0.10 | 0.19 | 0.16 | -0.05 | -0.26 | -0.21 | 0.009 |
| CPEB3 | S192 | 0.34 | -0.19 | 0.20 | -0.01 | -0.28 | -0.15 | 0.212 |
| CPEB4 | S97 | -0.49 | -0.07 | -0.25 | 0.23 | -0.08 | 0.46 | 0.073 |
| CPEB4 | S455 | 0.02 | 0.32 | 0.48 | -0.21 | -0.40 | -0.48 | 0.015 |
| CPNE3 | S243 | -0.01 | -0.01 | -0.02 | 0.08 | 0.09 | -0.14 | 0.789 |
| CPSF1 | S | -0.91 | -0.57 | 0.38 | -0.07 | 0.53 | 0.13 | 0.255 |
| CPSF1 | S766 | -0.19 | -0.46 | -0.26 | 0.23 | 0.36 | 0.15 | 0.006 |
| CPSF1 | S766 | -0.15 | -0.33 | -0.22 | 0.12 | 0.30 | 0.18 | 0.004 |
| CPSF1 | S | -0.91 | -0.37 | 0.51 | 0.01 | 0.25 | 0.10 | 0.421 |
| CPSF1 | S756 | 0.03 | 0.10 | 0.07 | -0.01 | 0.03 | -0.25 | 0.19 |
| CPSF1 | S756 | 0.06 | 0.09 | 0.08 | -0.09 | -0.12 | -0.03 | 0.005 |
| CPSF1 | S756 | 0.00 | 0.11 | 0.17 | -0.25 | 0.11 | -0.20 | 0.168 |
| CPSF2 | S419; S420; S423 | -0.05 | -0.17 | -0.06 | 0.08 | -0.05 | 0.22 | 0.114 |
| CPSF2 | S419; S420; S423 | -0.10 | -0.04 | -0.05 | 0.02 | -0.15 | 0.29 | 0.415 |

| | | | | | | | | |
|---|---|---|---|---|---|---|---|---|
| **CPSF2** | **S419; S420** | 0.05 | -0.25 | 0.22 | 0.29 | -0.72 | 0.18 | 0.81 |
| **CPSF3** | **T681** | -0.11 | 0.00 | -0.16 | 0.28 | -0.20 | 0.14 | 0.336 |
| **CPSF4** | **S212; S/T** | 0.00 | -0.24 | -0.29 | 0.27 | 0.24 | -0.07 | 0.083 |
| **CPSF6** | **T407** | 0.03 | 0.06 | -0.16 | 0.20 | -0.02 | -0.14 | 0.765 |
| **CPSF6** | **T407** | 0.17 | 0.00 | -0.17 | 0.03 | -0.09 | 0.04 | 0.955 |
| **CPSF6** | **T407** | 0.11 | -0.04 | 0.01 | -0.03 | 0.13 | -0.22 | 0.579 |
| **CPSF6** | **T407** | -0.05 | 0.13 | 0.14 | -0.01 | -0.14 | -0.09 | 0.101 |
| **CPSF7** | **T203** | -0.02 | -0.23 | -0.02 | 0.12 | -0.19 | 0.27 | 0.364 |
| **CPSF7** | **S409** | -0.28 | -0.18 | 0.25 | 0.06 | 0.03 | 0.06 | 0.499 |
| **CPSF7** | **S60** | -0.01 | -0.16 | 0.03 | 0.21 | -0.09 | -0.02 | 0.511 |
| **CPSF7** | **T203** | -0.18 | -0.15 | 0.20 | -0.06 | -0.01 | 0.15 | 0.662 |
| **CPSF7** | **S60** | 0.10 | 0.06 | 0.09 | -0.01 | -0.03 | -0.23 | 0.077 |
| **CPSF7** | **S48** | 0.13 | -0.05 | 0.19 | -0.03 | -0.06 | -0.22 | 0.108 |
| **CPSF7** | **T203** | -0.21 | -0.05 | 0.51 | -0.37 | 0.02 | -0.07 | 0.418 |
| **CRACR2B** | **S127** | -0.09 | -0.03 | -0.14 | 0.17 | 0.02 | 0.05 | 0.04 |
| **CRACR2B** | **S174** | -0.02 | -0.15 | -0.02 | -0.01 | -0.29 | 0.40 | 0.655 |
| **CRACR2B** | **S392** | 0.12 | 0.09 | -0.10 | -0.10 | 0.01 | -0.03 | 0.354 |
| **CRACR2B** | **S127** | 0.28 | 0.08 | -0.04 | -0.20 | 0.09 | -0.28 | 0.185 |
| **CRAMP1** | **S645** | -0.33 | -0.41 | 0.09 | 0.25 | 0.18 | 0.10 | 0.069 |
| **CRAMP1** | **S645** | -0.09 | -0.27 | -0.25 | 0.06 | -0.17 | 0.55 | 0.191 |
| **CRAMP1** | **S307** | 0.00 | -0.03 | 0.02 | 0.07 | -0.18 | 0.11 | 0.986 |
| **CRAMP1** | **S533** | 0.02 | 0.23 | 0.02 | 0.12 | -0.07 | -0.40 | 0.278 |
| **CREB1** | **S** | 0.04 | -0.36 | -0.59 | 0.09 | -0.24 | 0.70 | 0.217 |
| **CREB1** | **S/T** | 0.11 | -0.56 | -0.09 | 0.30 | 0.06 | 0.04 | 0.219 |
| **CREB1** | **S128** | -0.04 | -0.29 | -0.14 | -0.02 | 0.14 | 0.29 | 0.063 |
| **CREB1** | **S326** | 0.10 | 0.05 | -0.08 | -0.24 | 0.12 | 0.03 | 0.686 |
| **CREB3L1** | **S244** | 0.31 | 0.03 | 0.43 | -0.54 | 0.09 | -0.64 | 0.072 |
| **CREB3L4** | **S/T/Y** | 0.02 | -0.01 | -0.13 | 0.10 | -0.21 | 0.19 | 0.635 |
| **CREB3L4** | **S109** | 0.04 | 0.16 | -0.11 | -0.03 | 0.15 | -0.25 | 0.629 |
| **CREB3L4** | **S63** | 0.08 | 0.04 | 0.07 | -0.11 | 0.02 | -0.11 | 0.04 |
| **CREB3L4** | **S100** | 0.18 | 0.22 | 0.23 | 0.04 | 0.09 | -1.20 | 0.252 |
| **CREM** | **S129** | -1.01 | 0.07 | -0.05 | 0.15 | 0.29 | 0.22 | 0.184 |
| **CRKL** | **S107** | -0.14 | -0.10 | 0.19 | -0.21 | 0.13 | 0.09 | 0.898 |
| **CRKL** | **S107** | 0.21 | -0.02 | 0.19 | 0.17 | -0.46 | -0.22 | 0.208 |
| **CRNKL1** | **T/S** | -0.01 | -0.34 | -0.08 | 0.18 | -0.06 | 0.24 | 0.12 |
| **CRNN** | **T261** | -0.30 | -0.62 | -0.24 | -0.55 | -0.34 | 1.15 | 0.44 |
| **CROCC** | **S1451** | -1.12 | -1.02 | -0.87 | 0.60 | -0.60 | 1.21 | 0.059 |
| **CROCC** | **S2009** | -0.15 | 0.13 | -0.21 | -0.05 | 0.08 | 0.16 | 0.311 |
| **CROCC** | **S462** | -0.07 | -0.11 | -0.05 | -0.19 | 0.06 | 0.31 | 0.407 |
| **CROCC** | **S1490; T1495; S1496** | -0.05 | 0.23 | -0.36 | -0.14 | 0.15 | 0.09 | 0.661 |
| **CROCC** | **S** | 0.08 | -0.15 | -0.04 | -0.20 | -0.03 | 0.28 | 0.753 |
| **CROCC2** | **S440** | -1.01 | -0.89 | -0.86 | 0.31 | -1.29 | 1.46 | 0.249 |
| **CROCC2** | **S440** | -1.01 | -0.89 | -0.86 | 0.31 | -1.29 | 1.46 | 0.249 |
| **CRYBG1** | **S141** | -0.14 | -0.20 | -0.06 | 0.06 | 0.00 | 0.28 | 0.064 |
| **CRYBG1** | **S103** | -0.16 | -0.02 | -0.06 | 0.15 | -0.01 | 0.08 | 0.07 |
| **CRYBG1** | **S484** | -0.07 | -0.06 | -0.09 | -0.04 | 0.01 | 0.22 | 0.16 |
| **CRYBG1** | **S533** | 0.03 | 0.01 | -0.10 | 0.19 | -0.17 | 0.02 | 0.768 |
| **CRYBG1** | **S22** | -0.16 | 0.12 | 0.03 | -0.16 | 0.18 | -0.04 | 0.979 |

| | | | | | | | | |
|---|---|---|---|---|---|---|---|---|
| CRYBG1 | S132 | -0.10 | 0.02 | 0.18 | 0.00 | -0.22 | 0.08 | 0.533 |
| CRYBG1 | S103 | 0.20 | 0.00 | -0.06 | -0.01 | 0.00 | -0.15 | 0.356 |
| CRYBG2 | S474 | 0.14 | -0.15 | -0.13 | 0.08 | 0.11 | -0.08 | 0.481 |
| CRYBG3 | S1280 | -0.02 | -0.12 | 0.07 | -0.09 | -0.03 | 0.17 | 0.671 |
| CSDE1 | S123 | -0.10 | 0.08 | -0.11 | 0.14 | -0.17 | 0.13 | 0.584 |
| CSDE1 | S123 | 0.03 | 0.07 | -0.18 | 0.16 | -0.24 | 0.12 | 0.789 |
| CSNK1A1 | S311 | -0.59 | 0.07 | -0.48 | 0.16 | 0.50 | 0.05 | 0.079 |
| CSNK1A1 | S/T | 0.30 | 0.32 | -0.22 | 0.15 | 0.05 | -0.94 | 0.385 |
| CSNK1D | S382 | -0.24 | -0.01 | -0.16 | 0.22 | 0.02 | 0.13 | 0.042 |
| CSNK1D | T347; S356 | 0.38 | -0.44 | -0.03 | -0.18 | -0.01 | 0.15 | 0.96 |
| CSNK1D | T/S | -0.16 | -0.03 | 0.25 | -0.12 | 0.10 | -0.08 | 0.718 |
| CSNK1D | T347; S/T | 0.05 | 0.10 | -0.25 | 0.50 | 0.21 | -1.04 | 0.885 |
| CSNK1D | T/S | 0.12 | 0.07 | 0.10 | 0.11 | -0.11 | -0.35 | 0.183 |
| CSNK1D | T/S | 0.01 | 0.01 | 0.27 | 0.07 | 0.03 | -0.49 | 0.322 |
| CSNK1D | S356 | 0.14 | 0.15 | 0.12 | 0.20 | -0.40 | -0.33 | 0.17 |
| CSNK1E | S363 | 0.06 | 0.13 | 0.23 | -0.16 | -0.06 | -0.25 | 0.016 |
| CSNK1G2 | S359 | -0.14 | -0.16 | -0.24 | 0.29 | -0.04 | 0.20 | 0.035 |
| CSPP1 | S966 | -0.30 | 0.11 | -0.22 | -0.52 | 0.15 | 0.53 | 0.599 |
| CSPP1 | T242 | -0.02 | 0.09 | -0.20 | -0.19 | -0.02 | 0.28 | 0.694 |
| CSPP1 | S920 | 0.00 | 0.04 | -0.11 | -0.02 | 0.01 | 0.08 | 0.434 |
| CSPP1 | S31 | -0.48 | 0.02 | 0.33 | 0.17 | -0.41 | 0.18 | 0.941 |
| CSPP1 | S459 | 0.06 | 0.17 | 0.16 | -0.12 | -0.10 | -0.23 | 0.007 |
| CSPP1 | S459 | -0.13 | 0.60 | 0.04 | 0.03 | -0.23 | -0.58 | 0.204 |
| CSPP1 | T242 | 0.02 | 0.27 | 0.51 | -0.66 | -0.31 | -0.11 | 0.043 |
| CSTF2T | S/Y/T | -0.18 | -0.01 | 0.18 | -0.07 | -0.07 | 0.12 | 0.983 |
| CSTF2T | S563 | -0.01 | 0.12 | -0.02 | -0.01 | 0.05 | -0.15 | 0.411 |
| CSTF2T | S563 | 0.03 | 0.13 | 0.10 | -0.09 | -0.21 | 0.02 | 0.074 |
| CTCF | T717 | -0.23 | -0.28 | -0.45 | 0.42 | 0.24 | 0.10 | 0.007 |
| CTCF | T177 | -0.47 | -0.14 | -0.27 | 0.34 | 0.13 | 0.24 | 0.009 |
| CTCF | T/S | -0.91 | -0.02 | 0.02 | -0.19 | 0.45 | 0.29 | 0.248 |
| CTCF | T717 | -0.24 | -0.27 | -0.23 | 0.34 | 0.20 | 0.08 | 0.004 |
| CTCF | T289 | -0.28 | -0.20 | -0.24 | 0.15 | 0.12 | 0.34 | 0.004 |
| CTCF | T717 | -0.25 | -0.22 | -0.13 | 0.16 | 0.16 | 0.20 | 6E-04 |
| CTCF | T/S | -0.06 | -0.19 | -0.06 | 0.03 | 0.19 | 0.06 | 0.034 |
| CTCF | T709 | -0.22 | 0.12 | -0.21 | 0.16 | -0.02 | 0.11 | 0.208 |
| CTCF | T709 | -0.12 | 0.12 | -0.22 | 0.10 | 0.01 | 0.09 | 0.25 |
| CTDSP2 | S53 | -0.38 | -0.13 | -0.33 | -0.10 | -0.03 | 0.69 | 0.154 |
| CTDSPL | S33 | 0.33 | 0.33 | -0.11 | -0.72 | -0.04 | -0.02 | 0.176 |
| CTDSPL2 | S/T | -0.14 | -0.44 | -0.29 | 0.19 | 0.20 | 0.32 | 0.006 |
| CTDSPL2 | S104 | -0.04 | -0.15 | -0.25 | 0.29 | 0.00 | 0.08 | 0.063 |
| CTDSPL2 | T/S | -0.18 | -0.15 | -0.11 | 0.07 | -0.07 | 0.36 | 0.111 |
| CTDSPL2 | S165 | -0.17 | -0.20 | -0.01 | 0.02 | -0.06 | 0.35 | 0.182 |
| CTDSPL2 | S165 | -0.11 | -0.03 | -0.09 | -0.22 | 0.01 | 0.36 | 0.483 |
| CTDSPL2 | S104 | 0.08 | -0.26 | 0.05 | 0.14 | -0.02 | -0.03 | 0.559 |
| CTDSPL2 | S134 | -0.03 | -0.06 | 0.10 | 0.00 | 0.11 | -0.13 | 0.89 |
| CTDSPL2 | S165 | -0.12 | -0.03 | 0.15 | 0.05 | -0.31 | 0.19 | 0.887 |
| CTDSPL2 | S104 | 0.18 | 0.07 | -0.18 | -0.01 | 0.07 | -0.16 | 0.666 |
| CTDSPL2 | S104 | 0.14 | 0.13 | 0.00 | 0.04 | 0.02 | -0.38 | 0.241 |

| | | | | | | | | |
|---|---|---|---|---|---|---|---|---|
| CTDSPL2 | S28 | 0.30 | 0.19 | 0.11 | -0.22 | -0.16 | -0.33 | 0.004 |
| CTNNA1 | T654; S655 | -0.17 | -0.06 | -0.02 | 0.16 | -0.31 | 0.32 | 0.506 |
| CTNNA1 | T654; S655 | -0.18 | -0.05 | 0.00 | 0.08 | -0.18 | 0.27 | 0.404 |
| CTNNA1 | S641; T645 | -0.11 | -0.02 | 0.17 | -0.30 | -0.18 | 0.35 | 0.81 |
| CTNNA1 | S641 | -0.03 | 0.11 | 0.07 | -0.02 | -0.20 | 0.05 | 0.277 |
| CTNNA1 | T658; T/S | 0.04 | -0.19 | 0.45 | -0.16 | -0.10 | -0.15 | 0.284 |
| CTNNA1 | T/S | -0.09 | 0.19 | 0.31 | -0.32 | -0.37 | 0.13 | 0.177 |
| CTNNA1 | T654; S655; T658 | 0.32 | 0.18 | 0.10 | -0.27 | -0.19 | -0.25 | 0.003 |
| CTNNB1 | S552 | 0.18 | -1.40 | 0.29 | -0.21 | 0.10 | 0.41 | 0.515 |
| CTNNB1 | T/S | 0.04 | -0.16 | -0.16 | 0.23 | -0.23 | 0.21 | 0.368 |
| CTNNBL1 | S545 | -0.02 | -0.15 | -0.14 | 0.27 | -0.26 | 0.21 | 0.359 |
| CTNND1 | S252 | -0.68 | -0.67 | -0.51 | 0.39 | -0.09 | 0.84 | 0.022 |
| CTNND1 | S230 | -0.05 | -0.38 | -0.49 | -0.20 | 0.17 | 0.65 | 0.138 |
| CTNND1 | S346 | -0.32 | -0.32 | -0.12 | 0.22 | -0.06 | 0.44 | 0.048 |
| CTNND1 | S230 | 0.11 | -0.04 | -0.17 | -0.03 | -0.01 | 0.12 | 0.53 |
| CTNND2 | S267 | -0.21 | -0.15 | -0.20 | 0.20 | -0.09 | 0.35 | 0.058 |
| CTNND2 | S267; S276 | -0.19 | -0.25 | -0.14 | 0.08 | -0.15 | 0.49 | 0.155 |
| CTNND2 | S276 | -0.02 | -0.13 | 0.05 | -0.02 | 0.06 | 0.04 | 0.365 |
| CTNND2 | S537 | -0.03 | 0.03 | -0.04 | 0.05 | -0.09 | 0.07 | 0.717 |
| CTNND2 | S537 | 0.20 | 0.03 | 0.06 | 0.10 | -0.01 | -0.47 | 0.287 |
| CTPS1 | S574; S575 | 0.03 | -0.13 | -0.19 | 0.08 | 0.18 | 0.00 | 0.092 |
| CTPS1 | S574 | 0.10 | 0.13 | 0.24 | -0.22 | 0.05 | -0.40 | 0.065 |
| CTPS2 | S574 | 0.09 | 0.52 | 0.22 | -0.32 | -0.64 | -0.16 | 0.026 |
| CTR9 | S | -0.01 | -0.08 | -0.10 | 0.08 | -0.11 | 0.20 | 0.276 |
| CTR9 | S970 | 0.10 | 0.05 | -0.29 | 0.16 | -0.12 | 0.06 | 0.622 |
| CTR9 | T925 | 0.11 | 0.09 | -0.06 | 0.09 | -0.01 | -0.25 | 0.416 |
| CTR9 | S970 | 0.25 | 0.08 | -0.09 | 0.07 | -0.10 | -0.27 | 0.276 |
| CTR9 | S1097 | -0.01 | 0.04 | 0.20 | 0.15 | 0.04 | -0.52 | 0.429 |
| CTR9 | T925 | 0.15 | -0.04 | 0.14 | 0.30 | -0.16 | -0.55 | 0.439 |
| CTTN | S447 | -0.11 | -0.13 | -0.28 | -0.01 | -0.14 | 0.52 | 0.229 |
| CTTN | S418 | -0.07 | 0.03 | -0.07 | 0.15 | -0.04 | -0.01 | 0.345 |
| CTTN | T401 | -0.06 | 0.07 | 0.30 | -0.16 | -0.07 | -0.14 | 0.101 |
| CTTN | T401; S405 | -0.06 | 0.16 | 0.26 | -0.15 | -0.03 | -0.24 | 0.084 |
| CTTN | S405 | 0.08 | 0.15 | 0.14 | -0.06 | -0.01 | -0.37 | 0.076 |
| CUL3 | S737 | 0.43 | -0.21 | 0.13 | -0.13 | 0.29 | -0.85 | 0.415 |
| CUL4B | T49 | -0.07 | -0.05 | -0.05 | -0.02 | 0.06 | 0.12 | 0.051 |
| CUL4B | T79 | -0.10 | 0.02 | 0.06 | -0.01 | -0.08 | 0.10 | 0.882 |
| CUL4B | T79 | -0.10 | 0.02 | 0.06 | -0.01 | -0.08 | 0.10 | 0.882 |
| CUL4B | S/T | 0.24 | 0.19 | 0.14 | -0.28 | 0.03 | -0.45 | 0.044 |
| CUL4B | S145 | 0.32 | 0.25 | 0.08 | -0.32 | 0.03 | -0.54 | 0.054 |
| CUL9 | S2436 | -0.18 | 0.07 | 0.01 | 0.17 | 0.32 | -0.53 | 0.942 |
| CUL9 | S1457 | -0.49 | 0.15 | 0.38 | 0.18 | 0.12 | -0.60 | 0.765 |
| CUSTOS | S179 | -0.10 | -0.11 | -0.01 | -0.16 | 0.05 | 0.28 | 0.378 |
| CUSTOS | S248 | -0.12 | -0.03 | 0.02 | 0.13 | 0.02 | -0.04 | 0.286 |
| CUSTOS | S179 | 0.07 | 0.25 | 0.07 | -0.13 | -0.21 | -0.10 | 0.014 |
| CUX1 | S1321 | -0.16 | 0.07 | -0.13 | 0.27 | -0.19 | 0.07 | 0.46 |
| CUX1 | S | -0.15 | 0.13 | -0.27 | 0.61 | -0.06 | -0.52 | 0.774 |
| CUX1 | S1312 | -0.15 | 0.13 | -0.27 | 0.61 | -0.06 | -0.52 | 0.774 |

| | | | | | | | | |
|---|---|---|---|---|---|---|---|---|
| CUX1 | S1270 | 0.13 | 0.10 | -0.18 | 0.01 | -0.07 | -0.02 | 0.684 |
| CWC22 | S61 | -0.04 | -0.11 | 0.20 | -0.15 | -0.05 | 0.11 | 0.729 |
| CWC22 | S61 | 0.00 | 0.13 | 0.01 | -0.05 | 0.03 | -0.13 | 0.195 |
| CWC22 | S829; S831 | 0.09 | 0.05 | 0.01 | -0.01 | -0.10 | -0.05 | 0.037 |
| CWC22 | S28; S39 | -0.01 | -0.08 | 0.26 | -0.25 | 0.17 | -0.15 | 0.448 |
| CWC22 | S61 | 0.15 | 0.17 | -0.03 | 0.04 | -0.05 | -0.32 | 0.175 |
| CWC22 | S91; S93 | 0.25 | 0.11 | 0.03 | -0.05 | -0.06 | -0.35 | 0.073 |
| CWC22 | S39 | 0.12 | 0.06 | 0.21 | -0.32 | 0.14 | -0.31 | 0.139 |
| CWC22 | S829 | 0.21 | 0.15 | 0.07 | -0.16 | -0.08 | -0.25 | 0.009 |
| CWC22 | S792 | 0.18 | 0.25 | 0.07 | -0.24 | -0.03 | -0.32 | 0.023 |
| CWC25 | S | 0.27 | 0.36 | 0.32 | -0.36 | -0.06 | -0.91 | 0.038 |
| CWF19L2 | S479 | 0.05 | 0.07 | 0.14 | -0.16 | -0.16 | 0.03 | 0.054 |
| CXXC5 | T53 | -0.04 | -0.17 | -0.08 | 0.01 | 0.11 | 0.15 | 0.032 |
| CYB561 | S236 | -0.25 | 0.12 | 0.11 | -0.24 | 0.11 | 0.11 | 0.997 |
| CYB561 | S247 | 0.05 | 0.09 | 0.19 | -0.14 | -0.05 | -0.17 | 0.012 |
| CYB5B | T/S/Y | -0.20 | 0.00 | -0.11 | 0.08 | -0.12 | 0.30 | 0.229 |
| CYBRD1 | S232 | -0.12 | -0.07 | -0.06 | 0.13 | -0.07 | 0.16 | 0.115 |
| CYBRD1 | S232 | -0.17 | 0.09 | -0.08 | 0.13 | -0.03 | 0.05 | 0.309 |
| CYLD | S399 | -0.13 | -0.04 | -0.26 | 0.21 | -0.29 | 0.39 | 0.312 |
| DAAM1 | S1027 | 0.13 | -0.29 | -0.29 | 0.26 | 0.01 | 0.09 | 0.161 |
| DAAM1 | S1027 | -0.06 | -0.09 | 0.06 | 0.06 | 0.00 | 0.02 | 0.319 |
| DAAM1 | S/T | 0.24 | 0.33 | 0.44 | 0.34 | -0.09 | -1.50 | 0.248 |
| DAB2IP | S747 | -0.37 | 0.10 | 0.02 | 0.40 | 0.18 | -0.55 | 0.787 |
| DAB2IP | S978 | 0.01 | -0.21 | 0.05 | 0.05 | -0.20 | 0.25 | 0.625 |
| DAB2IP | S702 | 0.14 | -0.07 | -0.02 | -0.06 | -0.06 | 0.06 | 0.662 |
| DAB2IP | S995 | 0.01 | -0.06 | 0.28 | -0.20 | -0.17 | 0.09 | 0.287 |
| DACH1 | S491; S497 | -0.41 | -0.57 | -0.53 | 0.35 | 0.11 | 0.61 | 0.005 |
| DACH1 | S491; S493; S497 | -0.16 | -0.11 | -0.17 | 0.13 | -0.06 | 0.31 | 0.069 |
| DACH1 | T727 | -0.05 | -0.05 | -0.17 | 0.06 | 0.07 | 0.12 | 0.016 |
| DACH1 | S493; S497 | -0.08 | 0.04 | -0.03 | 0.05 | -0.14 | 0.15 | 0.651 |
| DACH1 | S | 0.02 | 0.06 | 0.03 | 0.04 | -0.18 | 0.02 | 0.346 |
| DACH1 | S519 | 0.50 | 0.51 | 0.22 | -0.99 | -0.15 | -0.76 | 0.018 |
| DAP | S51 | -0.09 | 0.31 | -0.19 | 0.56 | -0.04 | -1.01 | 0.737 |
| DAP | S/T | 0.24 | 0.42 | -0.03 | 0.14 | -0.38 | -0.67 | 0.133 |
| DAP3 | S280 | -0.19 | 0.22 | -0.23 | 0.00 | -0.20 | 0.31 | 0.65 |
| DAXX | S688; S702 | -0.18 | -0.14 | -0.17 | 0.18 | -0.16 | 0.37 | 0.135 |
| DAXX | S737; S739 | -0.02 | -0.12 | -0.05 | 0.10 | -0.08 | 0.14 | 0.189 |
| DAXX | S688 | -0.20 | 0.04 | 0.07 | 0.13 | -0.68 | 0.41 | 0.965 |
| DAXX | S495 | 0.10 | 0.05 | -0.08 | -0.05 | 0.11 | -0.14 | 0.629 |
| DAXX | S495 | 0.08 | 0.07 | -0.05 | -0.01 | 0.12 | -0.26 | 0.518 |
| DBF4 | T553 | -0.33 | -0.27 | -0.36 | 0.30 | 0.24 | 0.25 | 6E-05 |
| DBF4 | S312 | 0.12 | -0.16 | -0.34 | 0.13 | -0.28 | 0.39 | 0.429 |
| DBF4 | S381 | -0.14 | -0.01 | 0.14 | 0.12 | 0.17 | -0.35 | 0.943 |
| DBF4 | S508 | 0.22 | -0.11 | 0.11 | -0.31 | 0.05 | -0.01 | 0.325 |
| DBF4 | S359 | 0.11 | 0.20 | 0.16 | -0.35 | -0.15 | -0.04 | 0.024 |
| DBF4 | S381 | 0.28 | 0.23 | 0.33 | -1.06 | -0.29 | 0.08 | 0.105 |
| DBN1 | S345; T346 | -0.16 | -0.12 | 0.25 | 0.39 | -0.09 | -0.43 | 0.897 |
| DBN1 | S381 | 0.03 | -0.12 | 0.09 | 0.38 | -0.33 | -0.16 | 0.871 |

| | | | | | | | | |
|---|---|---|---|---|---|---|---|---|
| DBN1 | S601 | -0.04 | 0.10 | 0.06 | 0.16 | -0.11 | -0.20 | 0.507 |
| DBN1 | S337 | -0.11 | 0.01 | 0.23 | -0.04 | -0.14 | 0.01 | 0.424 |
| DBN1 | S601 | -0.12 | 0.13 | 0.15 | 0.27 | -0.21 | -0.33 | 0.512 |
| DBN1 | S/T | 0.02 | 0.13 | 0.15 | 0.21 | -0.44 | -0.17 | 0.29 |
| DBN1 | S/T/Y | -0.02 | 0.08 | 0.28 | 0.05 | -0.14 | -0.33 | 0.136 |
| DBN1 | S142 | -0.01 | 0.28 | 0.32 | 0.09 | -0.22 | -0.68 | 0.13 |
| DBN1 | S337 | 0.04 | 0.39 | 0.45 | -0.11 | -0.14 | -1.13 | 0.104 |
| DBN1 | S337 | 0.16 | 0.32 | 0.46 | -0.32 | -0.08 | -0.95 | 0.05 |
| DBNL | T291 | 0.02 | -0.24 | 0.30 | -0.12 | -0.06 | 0.04 | 0.674 |
| DBNL | S232 | 0.28 | 0.36 | 0.12 | 0.09 | -0.22 | -1.03 | 0.135 |
| DBP | S86 | -0.04 | 0.09 | -0.06 | -0.06 | -0.16 | 0.19 | 0.952 |
| DBP | S164 | 0.03 | 0.01 | 0.13 | -0.03 | -0.56 | 0.29 | 0.561 |
| DBP | S161 | 0.11 | 0.14 | 0.21 | -0.29 | -0.15 | -0.10 | 0.007 |
| DCAF10 | S92 | -0.29 | -0.29 | -0.49 | 0.28 | 0.23 | 0.34 | 1E-03 |
| DCAF10 | S63 | -0.06 | 0.05 | 0.24 | 0.04 | -0.16 | -0.15 | 0.197 |
| DCAF5 | S648 | -0.08 | -0.18 | -0.24 | 0.12 | 0.06 | 0.26 | 0.013 |
| DCAF5 | T/S | -0.06 | -0.08 | -0.32 | 0.33 | -0.06 | 0.11 | 0.117 |
| DCAF5 | S794 | 0.05 | -0.08 | -0.04 | 0.04 | 0.03 | 0.01 | 0.286 |
| DCAF5 | T497 | 0.06 | -0.28 | 0.15 | -0.61 | 0.14 | 0.35 | 0.955 |
| DCAF5 | T497 | 0.21 | -0.12 | 0.10 | -0.50 | 0.12 | 0.07 | 0.488 |
| DCAF5 | S648 | 0.26 | 0.12 | 0.20 | -0.08 | 0.18 | -0.99 | 0.243 |
| DCBLD1 | S513 | -0.31 | 0.02 | -0.14 | 0.20 | 0.02 | 0.15 | 0.073 |
| DCLK1 | S307 | -0.08 | -0.29 | -0.12 | 0.02 | -0.24 | 0.54 | 0.319 |
| DCLK1 | S337 | -0.32 | 0.28 | 0.05 | 0.09 | -0.07 | -0.10 | 0.864 |
| DCLRE1A | S111 | -0.20 | -0.16 | -0.37 | 0.14 | -0.24 | 0.60 | 0.175 |
| DCLRE1A | S238 | 0.03 | 0.12 | -0.09 | -0.06 | -0.12 | 0.09 | 0.618 |
| DCP1A | S525 | -0.17 | 0.08 | 0.07 | -0.07 | 0.02 | 0.06 | 0.904 |
| DCP1B | S283 | 0.05 | -0.09 | -0.18 | 0.16 | 0.13 | -0.11 | 0.291 |
| DCP1B | S147 | 0.08 | -0.22 | 0.10 | -0.01 | 0.00 | 0.02 | 0.861 |
| DCP1B | T/S | 0.51 | -0.43 | -0.02 | -0.71 | 0.19 | 0.14 | 0.736 |
| DCUN1D5 | S9 | 0.09 | -0.33 | -0.21 | 0.34 | 0.00 | 0.02 | 0.178 |
| DDB2 | S26 | 0.08 | 0.02 | 0.06 | 0.11 | 0.02 | -0.33 | 0.412 |
| DDHD1 | S723 | 0.36 | 0.01 | 0.02 | 0.10 | -0.09 | -0.56 | 0.244 |
| DDHD1 | S723 | 0.36 | 0.01 | 0.02 | 0.10 | -0.09 | -0.56 | 0.244 |
| DDX1 | S481 | 0.05 | -0.15 | 0.21 | -0.29 | 0.38 | -0.35 | 0.647 |
| DDX1 | S481 | 0.16 | 0.02 | 0.03 | -0.30 | 0.66 | -1.15 | 0.561 |
| DDX10 | S539 | -0.07 | -0.13 | -0.11 | 0.08 | 0.03 | 0.17 | 0.014 |
| DDX10 | S7 | 0.09 | 0.18 | 0.20 | -0.26 | -0.03 | -0.24 | 0.015 |
| DDX17 | S672 | -0.66 | -0.67 | 0.13 | -0.31 | 0.70 | 0.28 | 0.192 |
| DDX17 | S599 | -0.10 | 0.01 | -0.01 | 0.04 | 0.04 | 0.01 | 0.153 |
| DDX17 | T145 | -0.19 | 0.24 | 0.11 | 0.20 | -0.28 | -0.17 | 0.518 |
| DDX17 | S64 | -0.11 | 0.28 | 0.03 | -0.57 | 0.25 | -0.03 | 0.526 |
| DDX17 | S64 | 0.42 | 0.07 | 0.17 | -0.08 | -0.52 | -0.24 | 0.04 |
| DDX18 | S627 | -0.22 | -0.26 | -0.18 | 0.40 | -0.08 | 0.22 | 0.047 |
| DDX18 | S627 | 0.30 | -0.63 | 0.08 | -0.08 | 0.22 | -0.06 | 0.729 |
| DDX20 | S/Y/T | -0.18 | -0.07 | -0.04 | 0.13 | 0.09 | 0.06 | 0.016 |
| DDX20 | S672 | 0.08 | -0.05 | -0.27 | 0.29 | -0.27 | 0.14 | 0.533 |
| DDX20 | S/Y | 0.04 | -0.12 | -0.06 | -0.10 | 0.14 | 0.08 | 0.362 |

| | | | | | | | | |
|---|---|---|---|---|---|---|---|---|
| **DDX20** | **S703** | 0.02 | -0.10 | 0.00 | 0.01 | -0.03 | 0.09 | 0.361 |
| **DDX20** | **S/T** | 0.02 | -0.08 | 0.03 | -0.16 | 0.09 | 0.09 | 0.845 |
| **DDX21** | **S171** | -0.10 | -0.06 | -0.20 | 0.28 | -0.01 | 0.04 | 0.089 |
| **DDX21** | **S** | 0.17 | -0.17 | -0.38 | 0.20 | 0.08 | 0.01 | 0.256 |
| **DDX21** | **S121** | -0.02 | -0.12 | -0.07 | 0.09 | 0.15 | -0.05 | 0.103 |
| **DDX21** | **S** | -0.09 | -0.08 | -0.04 | 0.11 | 0.29 | -0.25 | 0.489 |
| **DDX21** | **S466** | -0.10 | -0.25 | 0.18 | 0.07 | -0.01 | 0.08 | 0.45 |
| **DDX21** | **S168; S171** | -0.06 | 0.00 | 0.04 | 0.02 | -0.04 | 0.04 | 0.77 |
| **DDX21** | **S** | 0.03 | 0.07 | -0.09 | -0.08 | 0.07 | -0.01 | 0.92 |
| **DDX21** | **S** | 0.06 | 0.05 | -0.07 | 0.06 | -0.09 | -0.01 | 0.693 |
| **DDX21** | **S168; S171; S173** | 0.08 | 0.01 | -0.02 | -0.04 | -0.20 | 0.15 | 0.636 |
| **DDX21** | **S168** | -0.01 | 0.05 | 0.06 | 0.02 | 0.02 | -0.15 | 0.306 |
| **DDX21** | **S168; S171; S173** | 0.07 | 0.07 | -0.01 | -0.04 | -0.10 | 0.00 | 0.086 |
| **DDX21** | **S168; S171** | 0.04 | 0.14 | -0.05 | 0.04 | -0.15 | -0.03 | 0.304 |
| **DDX21** | **S89** | -0.01 | -0.10 | 0.25 | -0.03 | -0.18 | 0.03 | 0.428 |
| **DDX21** | **S71** | 0.05 | -0.03 | 0.14 | -0.15 | 0.06 | -0.08 | 0.222 |
| **DDX21** | **S71** | 0.08 | 0.04 | 0.07 | -0.04 | -0.06 | -0.09 | 0.003 |
| **DDX21** | **S89** | 0.20 | -0.02 | 0.03 | -0.04 | -0.09 | -0.10 | 0.098 |
| **DDX21** | **S89** | 0.16 | 0.09 | 0.02 | -0.13 | -0.05 | -0.11 | 0.016 |
| **DDX21** | **S171; S173** | 0.12 | 0.17 | -0.01 | -0.11 | -0.11 | -0.08 | 0.021 |
| **DDX21** | **S121** | 0.19 | 0.00 | 0.13 | -0.27 | -0.08 | -0.01 | 0.082 |
| **DDX21** | **S** | 0.09 | 0.24 | 0.04 | -0.28 | 0.07 | -0.22 | 0.097 |
| **DDX21** | **S71** | 0.20 | 0.14 | 0.06 | -0.16 | 0.02 | -0.33 | 0.056 |
| **DDX21** | **S89** | 0.16 | 0.21 | 0.06 | -0.25 | -0.02 | -0.22 | 0.021 |
| **DDX21** | **S89** | 0.22 | 0.13 | 0.11 | -0.17 | -0.10 | -0.25 | 0.004 |
| **DDX21** | **S** | 0.20 | 0.14 | 0.15 | -0.19 | 0.00 | -0.40 | 0.037 |
| **DDX21** | **S168; S171; S173** | 0.22 | 0.20 | 0.14 | -0.27 | -0.09 | -0.29 | 0.004 |
| **DDX21** | **S164** | 0.13 | 0.33 | 0.21 | 0.03 | -0.28 | -0.64 | 0.064 |
| **DDX21** | **S121** | 0.22 | 0.24 | 0.25 | -0.40 | -0.06 | -0.42 | 0.01 |
| **DDX23** | **S143** | -3.94 | -3.64 | -4.05 | 1.16 | 0.92 | 0.74 | 1E-05 |
| **DDX23** | **S14** | 0.16 | -0.05 | 0.05 | -0.06 | -0.04 | -0.08 | 0.133 |
| **DDX24** | **S295** | -0.04 | -0.23 | -0.06 | -0.05 | -0.06 | 0.37 | 0.276 |
| **DDX24** | **S287** | -0.01 | -0.03 | -0.10 | 0.04 | 0.08 | 0.03 | 0.038 |
| **DDX24** | **S94** | 0.11 | 0.10 | -0.12 | 0.08 | 0.17 | -0.42 | 0.688 |
| **DDX24** | **S82** | 0.08 | 0.12 | 0.03 | 0.04 | -0.09 | -0.20 | 0.092 |
| **DDX27** | **S/T/Y** | -0.11 | -0.34 | 0.21 | 0.08 | 0.16 | -0.08 | 0.485 |
| **DDX27** | **S79** | -0.11 | 0.40 | -0.58 | -0.26 | 0.24 | 0.10 | 0.728 |
| **DDX27** | **S746** | 0.19 | 0.42 | -0.08 | -0.16 | -0.18 | -0.33 | 0.06 |
| **DDX3X** | **S269** | -1.08 | -1.65 | -1.36 | 0.70 | 0.53 | 0.81 | 4E-04 |
| **DDX3X** | **S508** | -0.02 | -0.14 | 0.09 | -0.17 | -0.06 | 0.27 | 0.818 |
| **DDX3X** | **S102** | -0.21 | 0.09 | 0.11 | -0.10 | -0.19 | 0.23 | 0.925 |
| **DDX3X** | **S61** | 0.07 | -0.13 | 0.09 | -0.01 | 0.05 | -0.08 | 0.751 |
| **DDX3X** | **S83** | -0.07 | 0.47 | 0.38 | -0.20 | -0.16 | -0.74 | 0.066 |
| **DDX41** | **S/Y** | -0.32 | 0.05 | -0.08 | 0.20 | -0.03 | 0.13 | 0.167 |
| **DDX41** | **S23** | 0.03 | 0.05 | 0.01 | 0.03 | 0.15 | -0.31 | 0.634 |
| **DDX41** | **S21; S23** | 0.03 | -0.02 | 0.09 | 0.09 | -0.13 | -0.09 | 0.357 |
| **DDX41** | **S** | 0.18 | 0.02 | 0.12 | -0.02 | -0.22 | -0.11 | 0.038 |
| **DDX41** | **S** | 0.22 | 0.04 | 0.13 | 0.12 | -0.27 | -0.34 | 0.122 |

| | | | | | | | | |
|---|---|---|---|---|---|---|---|---|
| **DDX42** | **S96** | -0.15 | -0.05 | 0.15 | -0.06 | 0.08 | 0.01 | 0.787 |
| **DDX42** | **T169** | 0.01 | 0.36 | -0.37 | -0.17 | 0.25 | -0.23 | 0.856 |
| **DDX42** | **S96** | 0.00 | 0.05 | 0.03 | 0.07 | -0.03 | -0.12 | 0.399 |
| **DDX42** | **Y/S/T** | 0.09 | 0.09 | -0.11 | 0.10 | 0.05 | -0.26 | 0.671 |
| **DDX42** | **S185** | -0.03 | 0.12 | 0.12 | -0.10 | 0.00 | -0.14 | 0.084 |
| **DDX42** | **Y/S/T** | -0.01 | 0.11 | 0.15 | -0.04 | 0.01 | -0.25 | 0.127 |
| **DDX42** | **Y/S/T** | 0.12 | 0.21 | -0.03 | -0.06 | 0.01 | -0.31 | 0.139 |
| **DDX42** | **Y/S/T** | 0.12 | 0.13 | 0.12 | 0.00 | -0.04 | -0.40 | 0.099 |
| **DDX42** | **S185** | 0.16 | 0.14 | 0.11 | -0.05 | -0.05 | -0.38 | 0.056 |
| **DDX42** | **Y/S/T** | 0.22 | 0.08 | 0.13 | -0.04 | 0.04 | -0.54 | 0.155 |
| **DDX42** | **T193** | -0.03 | 0.05 | 0.20 | 0.45 | 0.15 | -1.48 | 0.578 |
| **DDX42** | **S185** | 0.18 | 0.01 | 0.35 | -0.35 | 0.00 | -0.33 | 0.055 |
| **DDX46** | **S199** | -0.98 | -1.06 | -1.05 | 0.59 | 0.51 | 0.68 | 7E-06 |
| **DDX46** | **S804** | 0.33 | -0.57 | -0.27 | 0.46 | -0.42 | 0.16 | 0.554 |
| **DDX46** | **S804** | -0.04 | 0.00 | -0.07 | 0.07 | -0.01 | 0.04 | 0.084 |
| **DDX46** | **S26** | 0.07 | 0.03 | 0.15 | 0.15 | -0.29 | -0.15 | 0.255 |
| **DDX46** | **S804** | 0.18 | 0.11 | 0.02 | 0.02 | -0.04 | -0.34 | 0.13 |
| **DDX46** | **S** | 0.30 | 0.14 | 0.25 | -0.15 | -0.16 | -0.54 | 0.02 |
| **DDX5** | **T224** | -0.21 | -0.31 | -0.32 | 0.21 | 0.19 | 0.31 | 5E-04 |
| **DDX50** | **T266** | -0.20 | -0.15 | -0.30 | 0.11 | 0.15 | 0.29 | 0.005 |
| **DDX50** | **S/T/Y** | 0.03 | -0.35 | -0.05 | -0.01 | 0.06 | 0.26 | 0.188 |
| **DDX50** | **S82** | 0.07 | -0.34 | 0.08 | 0.12 | 0.07 | -0.06 | 0.499 |
| **DDX50** | **T266** | 0.12 | 0.28 | -0.07 | -0.31 | 0.21 | -0.37 | 0.27 |
| **DDX50** | **S/T** | 0.26 | -0.07 | 0.21 | -0.33 | -0.03 | -0.13 | 0.093 |
| **DDX50** | **S41** | 0.21 | 0.21 | 0.01 | -0.16 | -0.15 | -0.18 | 0.01 |
| **DDX50** | **S41** | 0.17 | 0.21 | 0.09 | -0.12 | -0.06 | -0.37 | 0.028 |
| **DDX50** | **S82** | 0.20 | 0.19 | 0.25 | -0.22 | -0.13 | -0.40 | 0.005 |
| **DDX51** | **S83** | -0.05 | -0.11 | -0.02 | -0.07 | -0.04 | 0.26 | 0.362 |
| **DDX51** | **S83** | 0.15 | 0.11 | 0.18 | -0.23 | -0.17 | -0.09 | 0.002 |
| **DDX51** | **S83** | 0.15 | 0.19 | 0.20 | -0.19 | -0.08 | -0.35 | 0.008 |
| **DDX54** | **S39; S41** | -0.31 | -0.15 | -0.23 | 0.25 | -0.14 | 0.43 | 0.078 |
| **DDX54** | **S644** | 0.00 | -0.01 | -0.07 | 0.10 | 0.05 | -0.07 | 0.376 |
| **DDX54** | **T/S** | 0.02 | 0.38 | 0.32 | -0.20 | -0.02 | -0.78 | 0.085 |
| **DDX55** | **S544** | -0.39 | -0.02 | -0.19 | 0.35 | -0.14 | 0.25 | 0.125 |
| **DDX55** | **S544** | -0.05 | -0.08 | -0.07 | 0.11 | 0.02 | 0.06 | 0.011 |
| **DDX55** | **S544** | -0.09 | -0.07 | 0.09 | -0.01 | -0.11 | 0.17 | 0.707 |
| **DDX55** | **S594** | -0.05 | -0.08 | 0.08 | -0.01 | -0.05 | 0.10 | 0.669 |
| **DDX55** | **S544** | 0.02 | -0.03 | 0.05 | -0.11 | 0.00 | 0.07 | 0.669 |
| **DDX55** | **S544** | 0.07 | -0.03 | 0.02 | -0.04 | 0.02 | -0.05 | 0.311 |
| **DDX55** | **T141** | -0.65 | 0.36 | 0.18 | -1.15 | 0.39 | 0.23 | 0.825 |
| **DDX59** | **S160** | -0.33 | -0.49 | -0.79 | 0.41 | 0.32 | 0.41 | 0.003 |
| **DEAF1** | **S176** | -0.20 | 0.00 | -0.01 | 0.18 | -0.09 | 0.09 | 0.273 |
| **DEAF1** | **T312** | 0.05 | 0.04 | 0.07 | -0.05 | -0.10 | -0.01 | 0.024 |
| **DEAF1** | **T312** | 0.17 | 0.07 | 0.09 | -0.14 | -0.09 | -0.13 | 0.003 |
| **DEF6** | **S597** | -0.49 | -0.06 | 0.21 | 0.45 | -0.08 | -0.23 | 0.616 |
| **DEF6** | **S502** | 0.02 | -0.12 | -0.06 | 0.29 | -0.23 | 0.04 | 0.619 |
| **DEK** | **S306; S307** | -0.22 | -0.13 | 0.12 | 0.07 | -0.16 | 0.26 | 0.439 |
| **DEK** | **S210** | -0.15 | -0.14 | 0.17 | -0.06 | 0.29 | -0.18 | 0.749 |

| | | | | | | | | |
|---|---|---|---|---|---|---|---|---|
| **DEK** | **S32; S51** | -0.01 | 0.02 | -0.05 | 0.00 | 0.11 | -0.07 | 0.678 |
| **DEK** | **S227; S231; S232** | 0.09 | 0.11 | 0.07 | -0.09 | -0.07 | -0.12 | 5E-04 |
| **DEK** | **S32** | 0.22 | 0.23 | -0.05 | -0.31 | 0.01 | -0.18 | 0.09 |
| **DEK** | **S** | 0.18 | 0.17 | 0.11 | -0.19 | 0.00 | -0.34 | 0.03 |
| **DEK** | **S303; S306; S307** | 0.32 | 0.33 | 0.14 | -0.34 | 0.01 | -0.77 | 0.054 |
| **DEK** | **S306; S307** | 0.39 | 0.53 | 0.35 | -0.50 | -0.28 | -1.21 | 0.019 |
| **DENND1A** | **S523** | 0.49 | 0.27 | 0.36 | -0.61 | -0.15 | -0.85 | 0.014 |
| **DENND4A** | **S1240** | -0.28 | 0.29 | -0.21 | 0.45 | -0.20 | -0.22 | 0.795 |
| **DENND4B** | **S736** | 0.06 | 0.01 | 0.43 | -0.19 | -0.01 | -0.47 | 0.107 |
| **DENR** | **S73** | 0.26 | 0.11 | 0.13 | -0.16 | -0.09 | -0.34 | 0.015 |
| **DENR** | **S73** | 0.28 | 0.15 | 0.08 | -0.21 | -0.06 | -0.34 | 0.02 |
| **DENR** | **S73** | 0.40 | 0.22 | -0.06 | -0.26 | -0.19 | -0.23 | 0.039 |
| **DFFA** | **S315** | 0.03 | 0.16 | 0.10 | -0.31 | -0.05 | 0.02 | 0.118 |
| **DGCR8** | **S153** | 0.11 | 0.06 | 0.16 | -0.09 | -0.18 | -0.09 | 0.006 |
| **DGCR8** | **S377** | 0.18 | 0.22 | 0.01 | 0.05 | -0.07 | -0.50 | 0.162 |
| **DHCR7** | **S14** | -0.16 | -0.22 | -0.05 | 0.16 | -0.05 | 0.25 | 0.061 |
| **DHCR7** | **S14** | -0.07 | -0.11 | -0.05 | 0.12 | -0.02 | 0.12 | 0.034 |
| **DHCR7** | **S14** | 0.12 | -0.09 | -0.07 | 0.03 | -0.05 | 0.05 | 0.763 |
| **DHCR7** | **S14** | 0.17 | -0.03 | -0.05 | 0.02 | -0.10 | -0.03 | 0.453 |
| **DHCR7** | **S25** | 0.03 | -0.07 | 0.22 | -0.06 | -0.19 | 0.04 | 0.296 |
| **DHX16** | **S103; S106; S112** | -0.30 | 0.05 | -0.09 | 0.00 | 0.05 | 0.24 | 0.169 |
| **DHX16** | **S103; S106** | -0.02 | 0.08 | 0.10 | -0.18 | -0.16 | 0.15 | 0.36 |
| **DHX16** | **S160** | 0.18 | 0.14 | 0.21 | -0.28 | 0.05 | -0.41 | 0.049 |
| **DHX16** | **S106** | 0.28 | 0.26 | 0.24 | -0.28 | -0.15 | -0.56 | 0.009 |
| **DHX29** | **S200** | 0.20 | 0.25 | 0.28 | -0.22 | -0.11 | -0.59 | 0.02 |
| **DHX30** | **S495** | -0.33 | -0.28 | -0.07 | 0.01 | 0.05 | 0.48 | 0.076 |
| **DHX30** | **S226** | -0.03 | -0.29 | -0.15 | 0.05 | -0.01 | 0.35 | 0.1 |
| **DHX38** | **S234** | -0.02 | -0.09 | 0.01 | -0.03 | 0.10 | 0.02 | 0.255 |
| **DHX38** | **T117** | -0.03 | 0.11 | 0.11 | -0.28 | -0.07 | 0.12 | 0.319 |
| **DHX38** | **T269** | 0.31 | -0.27 | 0.13 | 0.03 | -0.07 | -0.22 | 0.491 |
| **DHX8** | **S460** | 0.03 | -0.05 | 0.08 | 0.00 | 0.19 | -0.30 | 0.707 |
| **DHX8** | **S460** | 0.05 | 0.07 | -0.01 | -0.25 | 0.22 | -0.13 | 0.555 |
| **DHX9** | **S87** | 0.05 | -0.18 | -0.37 | -0.49 | 0.19 | 0.55 | 0.487 |
| **DHX9** | **S125** | 0.10 | -0.09 | -0.06 | -0.13 | -0.07 | 0.22 | 0.86 |
| **DHX9** | **T835** | 0.16 | -0.23 | 0.12 | -0.02 | -0.06 | 0.00 | 0.763 |
| **DHX9** | **T160; T177** | 0.00 | 0.08 | 0.00 | -0.06 | 0.11 | -0.13 | 0.512 |
| **DHX9** | **S87** | -0.08 | 0.28 | 0.02 | -0.17 | 0.06 | -0.17 | 0.288 |
| **DHX9** | **S87** | 0.03 | 0.27 | -0.05 | -0.18 | 0.02 | -0.14 | 0.188 |
| **DHX9** | **S87** | 0.17 | 0.12 | 0.19 | -0.21 | 0.00 | -0.36 | 0.032 |
| **DHX9** | **S452** | 0.36 | 0.39 | 0.19 | -0.29 | -0.20 | -0.80 | 0.019 |
| **DIDO1** | **S809** | -0.17 | -0.05 | -0.29 | 0.14 | -0.02 | 0.31 | 0.055 |
| **DIDO1** | **S805; S809** | -0.21 | -0.01 | -0.12 | 0.15 | -0.07 | 0.22 | 0.106 |
| **DIDO1** | **S1522; S** | -0.11 | 0.07 | -0.28 | -0.06 | 0.10 | 0.23 | 0.209 |
| **DIDO1** | **T151; S152; S154** | -0.17 | 0.11 | -0.15 | -0.18 | -0.08 | 0.38 | 0.604 |
| **DIDO1** | **S111; S114** | -0.07 | 0.11 | -0.18 | -0.06 | -0.01 | 0.18 | 0.494 |
| **DIDO1** | **S114** | 0.12 | 0.05 | -0.32 | 0.21 | -0.12 | 0.00 | 0.651 |
| **DIDO1** | **S1040** | 0.04 | 0.00 | -0.15 | 0.16 | -0.02 | -0.06 | 0.49 |
| **DIDO1** | **S805** | -0.07 | 0.14 | -0.20 | 0.05 | -0.29 | 0.29 | 0.774 |

| | | | | | | | | |
|---|---|---|---|---|---|---|---|---|
| DIDO1 | S1456 | -0.06 | -0.02 | 0.00 | 0.18 | 0.00 | -0.13 | 0.645 |
| DIDO1 | T109; S111; S117 | -0.02 | -0.07 | 0.07 | -0.01 | -0.09 | 0.11 | 0.917 |
| DIDO1 | S523 | 0.05 | 0.10 | -0.16 | -0.18 | 0.06 | 0.11 | 0.999 |
| DIDO1 | S805; S809 | -0.06 | -0.06 | 0.12 | 0.03 | -0.03 | -0.01 | 0.969 |
| DIDO1 | T151; S152 | 0.00 | 0.09 | -0.07 | -0.07 | -0.01 | 0.06 | 0.898 |
| DIDO1 | S1456 | 0.02 | 0.17 | -0.10 | 0.09 | -0.12 | -0.09 | 0.529 |
| DIDO1 | T351 | -0.02 | 0.20 | -0.06 | 0.14 | -0.23 | -0.08 | 0.526 |
| DIDO1 | S154 | 0.40 | 0.32 | 0.17 | -0.21 | -0.25 | -0.73 | 0.018 |
| DIP2B | S/T | 0.25 | -0.22 | -0.06 | 0.19 | 0.12 | -0.38 | 0.949 |
| DIP2B | T140 | 0.18 | 0.12 | 0.01 | -0.22 | 0.04 | -0.18 | 0.086 |
| DIP2B | T140 | 0.42 | 0.32 | 0.04 | -0.67 | -0.38 | -0.02 | 0.047 |
| DIS3 | S634 | -0.06 | 0.00 | -0.24 | 0.18 | -0.12 | 0.19 | 0.22 |
| DISP2 | S/Y | -0.65 | 0.35 | -0.35 | 0.43 | -0.14 | 0.07 | 0.377 |
| DISP2 | S1373 | -0.23 | 0.06 | -0.04 | 0.15 | -0.21 | 0.21 | 0.478 |
| DISP2 | S1252 | -0.13 | -0.06 | 0.12 | 0.00 | -0.01 | 0.06 | 0.643 |
| DISP2 | S1175 | -0.15 | -0.02 | 0.15 | -0.08 | -0.09 | 0.17 | 0.961 |
| DKC1 | S21 | 0.00 | -0.17 | -0.25 | 0.15 | 0.01 | 0.21 | 0.049 |
| DKC1 | S21 | -0.07 | 0.00 | -0.26 | 0.09 | 0.00 | 0.20 | 0.101 |
| DKC1 | S21 | -0.01 | -0.11 | -0.06 | 0.01 | -0.06 | 0.21 | 0.241 |
| DKC1 | S453; S455; T458 | -0.01 | -0.10 | 0.07 | 0.12 | -0.06 | -0.04 | 0.779 |
| DKC1 | S494 | 0.03 | -0.04 | -0.02 | 0.04 | -0.01 | -0.01 | 0.645 |
| DKC1 | S494 | 0.09 | 0.00 | -0.08 | 0.03 | -0.01 | -0.03 | 0.959 |
| DKC1 | S513 | -0.05 | 0.05 | 0.03 | -0.01 | -0.02 | -0.01 | 0.503 |
| DKC1 | S451; S453 | -0.08 | 0.07 | 0.07 | -0.11 | -0.04 | 0.08 | 0.582 |
| DKC1 | S451; S453 | 0.12 | 0.11 | 0.04 | -0.07 | -0.09 | -0.13 | 0.003 |
| DKC1 | S451 | 0.21 | 0.18 | 0.04 | -0.12 | -0.09 | -0.28 | 0.018 |
| DKC1 | S451 | 0.38 | 0.34 | 0.20 | -0.38 | -0.10 | -0.79 | 0.025 |
| DKC1 | S453 | 0.32 | 0.41 | 0.27 | -0.40 | -0.15 | -0.84 | 0.018 |
| DKC1 | S/T | 0.40 | 0.55 | 0.25 | -0.56 | -0.16 | -1.16 | 0.027 |
| DLG1 | S138 | 0.04 | -0.12 | -0.17 | -0.13 | 0.36 | -0.06 | 0.45 |
| DLG1 | S138 | 0.04 | -0.12 | -0.17 | -0.13 | 0.36 | -0.06 | 0.45 |
| DLG5 | S773 | -0.02 | 0.06 | -0.32 | 0.18 | -0.22 | 0.24 | 0.434 |
| DLG5 | T984 | 0.10 | -0.11 | -0.16 | 0.16 | -0.12 | 0.10 | 0.404 |
| DLG5 | S1232 | -0.12 | 0.01 | -0.05 | 0.08 | 0.14 | -0.07 | 0.225 |
| DLG5 | S/T/Y | -0.15 | -0.23 | 0.18 | 0.53 | -0.19 | -0.34 | 0.832 |
| DLG5 | T1011 | 0.00 | 0.13 | 0.10 | -0.10 | -0.10 | -0.05 | 0.019 |
| DLG5 | S295 | 0.11 | 0.09 | 0.17 | -0.32 | -0.05 | -0.05 | 0.048 |
| DLG5 | S1209 | 0.03 | 0.22 | 0.32 | -0.23 | -0.14 | -0.30 | 0.014 |
| DLG5 | S295 | 0.14 | 0.34 | 0.31 | -0.35 | -0.22 | -0.41 | 0.002 |
| DLGAP4 | T915 | -0.04 | -0.18 | 0.08 | -0.02 | -0.11 | 0.24 | 0.547 |
| DLGAP4 | S973 | -0.08 | 0.07 | -0.08 | 0.16 | -0.19 | 0.10 | 0.66 |
| DLGAP4 | T915 | -0.03 | -0.14 | 0.09 | -0.02 | 0.24 | -0.18 | 0.781 |
| DLGAP5 | T/S | 0.20 | 0.05 | 0.48 | -0.43 | -0.14 | -0.36 | 0.022 |
| DMAP1 | S454 | 0.00 | -0.07 | -0.22 | 0.18 | 0.01 | 0.07 | 0.092 |
| DMAP1 | T445 | 0.11 | 0.03 | 0.00 | -0.11 | -0.03 | -0.02 | 0.079 |
| DMAP1 | S454 | 0.26 | -0.04 | -0.07 | -0.10 | -0.14 | 0.05 | 0.398 |
| DMAP1 | T409 | -0.24 | 0.36 | 0.10 | 0.14 | -0.28 | -0.20 | 0.44 |
| DMAP1 | T445 | -0.06 | 0.25 | 0.26 | 0.25 | -0.37 | -0.54 | 0.227 |

| | | | | | | | | |
|---|---|---|---|---|---|---|---|---|
| DMWD | T/S | -0.14 | -0.18 | -0.01 | 0.17 | 0.16 | -0.04 | 0.073 |
| DMXL1 | S1908 | -0.03 | 0.21 | 0.15 | -0.16 | -0.06 | -0.15 | 0.037 |
| DNAJA1 | S335 | -0.30 | 0.05 | -0.30 | -0.11 | 0.06 | 0.46 | 0.198 |
| DNAJA2 | S123 | -0.12 | -0.19 | -0.04 | 0.29 | 0.02 | -0.01 | 0.108 |
| DNAJB1 | S171 | 0.02 | 0.01 | 0.05 | -0.25 | -0.05 | 0.19 | 0.63 |
| DNAJB2 | S313 | -0.16 | -0.14 | -0.14 | 0.12 | 0.04 | 0.23 | 0.007 |
| DNAJB6 | S277 | -0.11 | -0.04 | 0.02 | 0.02 | -0.01 | 0.12 | 0.186 |
| DNAJC1 | S479; S480 | -0.15 | -0.13 | -0.04 | -0.06 | -0.09 | 0.38 | 0.299 |
| DNAJC1 | S479; S480 | 0.07 | 0.12 | -0.06 | -0.01 | -0.11 | -0.02 | 0.222 |
| DNAJC1 | S480 | 0.13 | 0.10 | -0.11 | 0.05 | 0.10 | -0.32 | 0.546 |
| DNAJC1 | S479; S480 | 0.19 | 0.13 | 0.05 | -0.22 | -0.16 | -0.05 | 0.014 |
| DNAJC11 | T99 | 0.01 | -0.24 | -0.22 | -0.19 | 0.21 | 0.33 | 0.209 |
| DNAJC2 | S47 | 0.10 | 0.01 | -0.21 | 0.21 | -0.08 | -0.08 | 0.75 |
| DNAJC21 | S370 | -0.13 | 0.06 | 0.16 | -0.23 | 0.17 | -0.07 | 0.632 |
| DNAJC21 | S370 | -0.06 | 0.14 | 0.16 | -0.27 | 0.11 | -0.13 | 0.239 |
| DNAJC30 | S137 | -0.36 | 0.25 | 0.37 | -0.13 | -0.11 | -0.14 | 0.396 |
| DNAJC30 | S39 | 0.13 | 0.22 | 0.13 | -0.15 | -0.02 | -0.40 | 0.04 |
| DNMT1 | S714 | -0.22 | -0.08 | 0.34 | -0.21 | 0.04 | 0.04 | 0.777 |
| DNMT1 | S714 | -0.16 | 0.19 | 0.02 | -0.43 | 0.17 | 0.11 | 0.762 |
| DNMT1 | S714 | 0.11 | 0.10 | 0.05 | -0.12 | -0.06 | -0.11 | 0.002 |
| DNMT3A | S75 | -0.29 | -0.57 | -0.51 | 0.29 | 0.17 | 0.55 | 0.005 |
| DNMT3A | T/S | -0.29 | -0.14 | -0.09 | 0.43 | 0.00 | -0.01 | 0.117 |
| DNMT3A | S | -0.05 | 0.01 | -0.28 | 0.20 | -0.08 | 0.15 | 0.182 |
| DNMT3A | T/S | 0.07 | 0.00 | -0.13 | 0.04 | -0.09 | 0.11 | 0.653 |
| DNMT3A | S243; T260; S267 | 0.13 | 0.05 | -0.06 | -0.36 | -0.11 | 0.27 | 0.615 |
| DNMT3A | S75 | 0.22 | 0.02 | -0.10 | 0.19 | -0.23 | -0.16 | 0.529 |
| DNMT3A | T138 | 0.20 | -0.05 | 0.00 | 0.07 | -0.35 | 0.07 | 0.506 |
| DNMT3A | S243; T257 | 0.09 | 0.18 | -0.08 | -0.13 | -0.01 | -0.07 | 0.182 |
| DNMT3A | S75 | 0.24 | 0.00 | -0.03 | -0.21 | 0.14 | -0.18 | 0.33 |
| DNMT3A | T251 | 0.11 | 0.10 | 0.07 | -0.10 | -0.07 | -0.12 | 5E-04 |
| DNMT3A | S243; T257 | 0.16 | 0.23 | -0.07 | -0.18 | -0.04 | -0.16 | 0.089 |
| DNMT3A | S243; T260; S267 | 0.24 | -0.01 | 0.27 | -0.70 | -0.13 | 0.12 | 0.198 |
| DNMT3B | S | -0.16 | -0.35 | 0.01 | -0.12 | -0.06 | 0.52 | 0.291 |
| DNMT3B | T383 | -0.06 | -0.21 | 0.02 | -0.02 | -0.01 | 0.24 | 0.232 |
| DNMT3B | S34 | -0.07 | -0.07 | 0.12 | 0.02 | -0.04 | 0.03 | 0.932 |
| DNMT3B | S65 | -0.11 | -0.05 | 0.23 | -0.18 | -0.04 | 0.11 | 0.664 |
| DNMT3B | S | -0.09 | -0.09 | 0.31 | 0.06 | -0.06 | -0.18 | 0.544 |
| DNMT3B | S82 | 0.05 | 0.06 | 0.17 | -0.09 | 0.05 | -0.28 | 0.126 |
| DNTTIP2 | S117 | -0.33 | -0.79 | -0.53 | 0.21 | -0.02 | 0.84 | 0.037 |
| DNTTIP2 | S481 | -0.62 | -0.41 | 0.03 | 0.51 | 0.26 | -0.06 | 0.088 |
| DNTTIP2 | T610 | -0.05 | -0.36 | -0.51 | 0.02 | 0.42 | 0.27 | 0.037 |
| DNTTIP2 | S21 | -0.17 | -0.07 | -0.44 | 0.21 | 0.39 | -0.07 | 0.083 |
| DNTTIP2 | S481 | -0.35 | -0.20 | 0.01 | 0.03 | 0.46 | -0.09 | 0.185 |
| DNTTIP2 | T232; S236; T241 | -0.24 | -0.09 | -0.14 | 0.20 | 0.07 | 0.15 | 0.006 |
| DNTTIP2 | S141; S148 | -0.10 | 0.02 | -0.25 | 0.08 | 0.09 | 0.13 | 0.062 |
| DNTTIP2 | S/T | -0.38 | 0.09 | -0.08 | 0.26 | 0.23 | -0.24 | 0.396 |
| DNTTIP2 | S141; S145; S148 | -0.16 | -0.14 | 0.04 | 0.06 | 0.05 | 0.13 | 0.078 |
| DNTTIP2 | T232 | -0.06 | -0.07 | -0.12 | -0.13 | 0.13 | 0.22 | 0.221 |

| | | | | | | | | |
|---|---|---|---|---|---|---|---|---|
| **DNTTIP2** | **S434** | -0.21 | -0.13 | 0.12 | 0.15 | -0.15 | 0.17 | 0.42 |
| **DNTTIP2** | **S184** | -0.18 | -0.15 | 0.17 | 0.33 | -0.06 | -0.18 | 0.691 |
| **DNTTIP2** | **S141** | 0.12 | -0.08 | -0.17 | 0.08 | -0.07 | 0.10 | 0.466 |
| **DNTTIP2** | **S88** | -0.12 | 0.05 | -0.01 | 0.06 | 0.16 | -0.16 | 0.681 |
| **DNTTIP2** | **S397; S/T** | 0.05 | 0.09 | -0.11 | -0.08 | 0.09 | -0.05 | 0.825 |
| **DNTTIP2** | **S474** | 0.06 | 0.12 | -0.11 | -0.02 | -0.03 | -0.03 | 0.508 |
| **DNTTIP2** | **S397; S414** | 0.12 | 0.06 | -0.09 | 0.06 | -0.07 | -0.10 | 0.431 |
| **DNTTIP2** | **S397** | 0.02 | -0.03 | 0.11 | -0.08 | -0.03 | 0.00 | 0.186 |
| **DNTTIP2** | **S184** | -0.07 | 0.05 | 0.12 | -0.05 | 0.09 | -0.17 | 0.442 |
| **DNTTIP2** | **S397** | 0.21 | -0.12 | 0.10 | -0.11 | -0.07 | -0.04 | 0.234 |
| **DNTTIP2** | **S340** | 0.09 | -0.02 | 0.24 | -0.03 | -0.26 | -0.07 | 0.095 |
| **DNTTIP2** | **S397** | 0.18 | 0.28 | 0.23 | -0.38 | -0.24 | -0.21 | 0.001 |
| **DOCK2** | **S1685** | 0.04 | 0.05 | -0.09 | 0.07 | 0.02 | -0.09 | 0.984 |
| **DOCK4** | **S1700** | 0.10 | 0.03 | -0.06 | -0.05 | -0.19 | 0.14 | 0.614 |
| **DOCK4** | **S/T/Y** | 0.10 | 0.36 | -0.28 | 0.09 | -0.33 | -0.06 | 0.513 |
| **DOCK6** | **S** | 0.05 | 0.14 | -0.06 | 0.02 | -0.15 | -0.02 | 0.308 |
| **DOCK7** | **S946** | -0.01 | -0.74 | -0.38 | 0.65 | -0.30 | 0.34 | 0.16 |
| **DOCK7** | **S900** | -0.30 | -0.18 | -0.22 | 0.13 | 0.01 | 0.43 | 0.032 |
| **DOCK7** | **S900** | -0.19 | -0.15 | -0.13 | 0.13 | -0.05 | 0.32 | 0.056 |
| **DOCK7** | **S1432** | -0.16 | -0.16 | 0.17 | 0.01 | -0.03 | 0.14 | 0.486 |
| **DOCK9** | **S927** | -0.03 | 0.06 | -0.21 | 0.15 | -0.02 | 0.03 | 0.299 |
| **DOK2** | **S129; T144** | 0.19 | 0.21 | -0.63 | 0.02 | -0.03 | 0.08 | 0.724 |
| **DOP1B** | **S/T** | -0.04 | 0.25 | -0.69 | -0.42 | -0.33 | 0.75 | 0.751 |
| **DOP1B** | **S/T** | 0.04 | 0.33 | -0.51 | -0.46 | -0.17 | 0.48 | 0.999 |
| **DOP1B** | **S714** | 0.10 | -0.21 | -0.01 | -0.46 | -0.27 | 0.59 | 0.984 |
| **DOT1L** | **S1104** | -0.69 | -0.96 | -0.70 | 0.70 | -0.66 | 1.00 | 0.095 |
| **DOT1L** | **S/T** | 0.06 | -0.38 | -0.08 | -0.06 | 0.02 | 0.34 | 0.264 |
| **DOT1L** | **T900; S902** | -0.42 | -0.14 | 0.12 | 0.46 | -0.66 | 0.32 | 0.642 |
| **DOT1L** | **S826; S834** | -0.01 | 0.04 | -0.02 | 0.03 | -0.04 | 0.00 | 0.852 |
| **DOT1L** | **T/S/Y** | 0.11 | 0.10 | -0.05 | -0.12 | 0.16 | -0.24 | 0.383 |
| **DOT1L** | **S902; S/T** | -0.05 | -0.04 | 0.28 | -0.09 | -0.05 | -0.09 | 0.267 |
| **DOT1L** | **S1065** | 0.12 | 0.05 | 0.04 | -0.11 | -0.14 | 0.03 | 0.065 |
| **DOT1L** | **S1001; S** | 0.07 | 0.22 | 0.04 | -0.04 | -0.12 | -0.20 | 0.034 |
| **DOT1L** | **S374** | 0.07 | 0.09 | 0.16 | -0.14 | 0.01 | -0.24 | 0.043 |
| **DOT1L** | **S1001** | -0.08 | 0.09 | 0.30 | 0.01 | -0.56 | 0.10 | 0.347 |
| **DOT1L** | **S1153** | 0.10 | 0.15 | 0.24 | -0.26 | -0.17 | -0.13 | 0.003 |
| **DOT1L** | **S902** | -0.04 | 0.14 | 0.44 | -0.22 | 0.07 | -0.59 | 0.145 |
| **DOT1L** | **S297** | 0.33 | 0.32 | 0.02 | 0.09 | -0.51 | -0.49 | 0.075 |
| **DOT1L** | **T355** | 0.31 | 0.27 | 0.29 | -0.47 | -0.14 | -0.53 | 0.005 |
| **DOT1L** | **S471** | 0.04 | 0.41 | 0.54 | -0.65 | -0.06 | -0.75 | 0.035 |
| **DOT1L** | **S448** | 0.48 | -0.16 | 0.62 | -0.88 | -1.21 | 0.27 | 0.147 |
| **DPF2** | **S118** | -1.21 | -1.06 | -1.00 | 0.49 | 0.62 | 0.72 | 5E-05 |
| **DPF2** | **T176** | -0.15 | -0.03 | -0.13 | 0.18 | 0.11 | -0.01 | 0.04 |
| **DPF2** | **S142** | -0.14 | -0.10 | -0.03 | 0.01 | 0.07 | 0.16 | 0.039 |
| **DPF2** | **S142** | -0.08 | -0.03 | -0.01 | 0.01 | 0.08 | 0.03 | 0.046 |
| **DPF2** | **T176** | 0.17 | 0.18 | 0.05 | -0.20 | 0.05 | -0.31 | 0.069 |
| **DROSHA** | **S807** | 0.09 | -0.04 | 0.03 | -0.17 | 0.26 | -0.22 | 0.687 |
| **DSC2** | **S864; S868** | -0.26 | -0.15 | -0.02 | 0.06 | -0.07 | 0.36 | 0.146 |

| | | | | | | | | |
|---|---|---|---|---|---|---|---|---|
| DSC2 | S868 | -0.18 | -0.07 | 0.02 | 0.15 | -0.06 | 0.11 | 0.165 |
| DSG2 | S703 | -0.15 | -0.24 | -0.16 | 0.26 | 0.06 | 0.16 | 0.006 |
| DSG2 | T650 | -0.09 | 0.41 | -0.82 | 0.12 | 0.07 | 0.04 | 0.541 |
| DSG2 | S782 | -0.11 | 0.04 | -0.12 | 0.10 | -0.01 | 0.08 | 0.141 |
| DSN1 | S81 | -0.25 | -0.41 | -0.07 | 0.03 | -0.03 | 0.54 | 0.109 |
| DSN1 | S331 | -0.31 | 0.03 | -0.11 | -0.06 | -0.23 | 0.52 | 0.457 |
| DSN1 | S125 | 0.10 | -0.24 | -0.19 | 0.01 | 0.44 | -0.26 | 0.487 |
| DSN1 | S81 | 0.04 | -0.05 | -0.09 | 0.04 | -0.28 | 0.28 | 0.785 |
| DSN1 | S27 | 0.13 | 0.14 | 0.25 | -0.30 | -0.07 | -0.23 | 0.01 |
| DSN1 | S | 0.02 | 0.12 | 0.39 | -0.11 | -0.27 | -0.27 | 0.032 |
| DSP | S2825; S/Y | -0.39 | 0.01 | 0.05 | 0.19 | 0.08 | 0.01 | 0.245 |
| DSP | S/T | -0.11 | -0.07 | -0.05 | 0.11 | 0.10 | 0.01 | 0.014 |
| DSP | S53 | -0.02 | -0.06 | -0.17 | 0.09 | -0.34 | 0.39 | 0.583 |
| DSP | S22 | -0.10 | -0.07 | 0.00 | -0.01 | -0.13 | 0.27 | 0.463 |
| DSP | S53 | 0.01 | 0.05 | -0.19 | 0.05 | -0.46 | 0.41 | 0.875 |
| DSP | S53 | -0.04 | 0.09 | 0.03 | -0.16 | 0.21 | -0.17 | 0.639 |
| DSP | S176 | 0.11 | 0.18 | -0.03 | -0.01 | -0.15 | -0.13 | 0.071 |
| DSP | S2825 | -0.75 | 0.21 | 0.54 | 0.17 | 0.21 | -0.94 | 0.749 |
| DSP | S/T/Y | 0.23 | -0.10 | 0.25 | -0.03 | -0.55 | 0.06 | 0.248 |
| DST | S3970 | -0.71 | 0.12 | -0.39 | 0.17 | 0.22 | 0.32 | 0.083 |
| DTL | S512 | -0.16 | 0.00 | -0.06 | 0.09 | -0.22 | 0.29 | 0.468 |
| DTL | S490 | 0.32 | -0.11 | 0.22 | 0.36 | 0.47 | -1.50 | 0.602 |
| DTNA | S458 | -0.26 | -0.22 | -0.36 | 0.28 | 0.12 | 0.29 | 0.002 |
| DTNA | S/T | -0.15 | -0.06 | -0.25 | 0.09 | 0.01 | 0.29 | 0.048 |
| DTNA | S/T | -0.20 | 0.00 | 0.04 | 0.10 | -0.04 | 0.08 | 0.293 |
| DTNA | S662 | -0.09 | 0.25 | 0.25 | -0.04 | -0.20 | -0.25 | 0.084 |
| DTWD2 | S132 | -0.39 | -0.21 | -0.19 | 0.23 | 0.31 | 0.11 | 0.005 |
| DTX2 | S360 | -0.01 | -0.10 | -0.13 | 0.03 | 0.02 | 0.18 | 0.067 |
| DTX3L | S202 | -0.13 | 0.00 | 0.18 | 0.18 | -0.31 | 0.02 | 0.766 |
| DUS3L | S276; S277 | -0.12 | -0.14 | 0.24 | -0.19 | 0.12 | 0.04 | 0.989 |
| DUT | S99 | 0.12 | 0.01 | -0.45 | -0.10 | -0.28 | 0.50 | 0.645 |
| DYNC1LI1 | S207 | 0.08 | -0.18 | -0.24 | 0.27 | -0.08 | 0.09 | 0.218 |
| DYNC1LI1 | S207 | 0.06 | -0.01 | -0.14 | 0.04 | -0.03 | 0.06 | 0.448 |
| DYNC1LI1 | S510 | -0.05 | 0.06 | 0.03 | -0.32 | -0.12 | 0.33 | 0.806 |
| DYNC1LI1 | S421 | 0.30 | -0.15 | 0.39 | 0.22 | -0.39 | -0.68 | 0.21 |
| DYNC1LI2 | S194 | 0.03 | 0.05 | -0.05 | 0.08 | -0.13 | 0.01 | 0.727 |
| DYNC1LI2 | S407 | 0.09 | 0.07 | -0.11 | 0.00 | 0.00 | -0.05 | 0.647 |
| DYNC2I1 | S247 | 0.10 | 0.22 | 0.14 | -0.02 | 0.02 | -0.58 | 0.151 |
| DYRK1A | Y321 | 0.02 | 0.12 | 0.02 | -0.10 | 0.14 | -0.23 | 0.353 |
| DYRK2 | Y382 | 0.10 | 0.06 | -0.02 | 0.04 | -0.19 | 0.00 | 0.274 |
| E2F1 | S375 | -0.05 | -0.06 | -0.04 | 0.20 | -0.03 | -0.04 | 0.302 |
| EAF1 | S165 | 0.07 | -0.04 | 0.12 | 0.04 | -0.07 | -0.12 | 0.211 |
| EAF1 | S165; T/S | 0.00 | 0.05 | 0.14 | -0.14 | 0.00 | -0.06 | 0.086 |
| EAF1 | T157 | 0.29 | -0.03 | 0.07 | -0.01 | -0.75 | 0.21 | 0.393 |
| EAF1 | S165; T/S | 0.28 | 0.32 | -0.12 | -0.61 | 0.28 | -0.41 | 0.248 |
| EAF1 | T/S | 0.41 | 0.17 | 0.20 | -0.37 | -0.44 | -0.18 | 0.006 |
| EAF2 | S154 | 0.09 | 0.12 | 0.09 | -0.12 | -0.02 | -0.19 | 0.015 |
| EBAG9 | T/S/Y | 0.30 | -0.14 | -0.02 | -0.18 | 0.08 | -0.09 | 0.509 |

| | | | | | | | | |
|---|---|---|---|---|---|---|---|---|
| EBAG9 | T/S/Y | 0.30 | -0.14 | -0.02 | -0.18 | 0.08 | -0.09 | 0.509 |
| EBAG9 | S36 | 0.12 | 0.20 | -0.04 | -0.10 | -0.29 | 0.05 | 0.156 |
| ECSIT | S253 | -0.56 | 0.45 | 0.27 | -0.06 | -0.59 | 0.17 | 0.601 |
| EDC3 | T173 | -0.68 | -0.52 | -0.26 | 0.56 | 0.28 | 0.21 | 0.007 |
| EDC3 | S131 | 0.31 | 0.20 | 0.13 | -0.23 | -0.03 | -0.54 | 0.037 |
| EDC4 | S879 | -0.08 | 0.10 | 0.07 | -0.11 | 0.12 | -0.12 | 0.538 |
| EDC4 | S879 | 0.05 | 0.23 | 0.17 | -0.11 | -0.10 | -0.32 | 0.021 |
| EDC4 | S844 | 0.11 | 0.23 | 0.15 | -0.10 | -0.10 | -0.38 | 0.024 |
| EDC4 | S844 | 0.19 | 0.09 | 0.34 | -0.32 | -0.05 | -0.39 | 0.022 |
| EED | S29; S34 | -0.04 | -0.17 | -0.13 | 0.09 | -0.12 | 0.31 | 0.192 |
| EED | S29; T55 | -0.12 | 0.02 | -0.22 | -0.20 | -0.04 | 0.45 | 0.438 |
| EED | S29; S34; T55 | -0.30 | 0.10 | 0.11 | -0.21 | 0.20 | 0.04 | 0.834 |
| EED | S29; S34; T55 | -0.19 | 0.16 | 0.17 | 0.23 | -0.11 | -0.37 | 0.582 |
| EED | S34 | -0.27 | 0.26 | 0.21 | -0.07 | -0.19 | -0.01 | 0.416 |
| EED | S34; S46 | -0.05 | 0.14 | 0.25 | -0.26 | -0.12 | -0.01 | 0.102 |
| EED | S34; T55 | 0.24 | 0.17 | 0.12 | -0.18 | -0.14 | -0.30 | 0.003 |
| EED | S34 | 0.41 | 0.00 | 0.14 | -0.59 | -0.35 | 0.16 | 0.15 |
| EEF1A1 | S291 | -0.19 | -0.31 | -0.52 | 0.10 | -0.25 | 0.78 | 0.161 |
| EEF1A1 | S300 | -0.10 | -0.27 | -0.07 | 0.00 | -0.24 | 0.53 | 0.361 |
| EEF1B2 | S106 | 0.02 | 0.07 | -0.21 | 0.06 | -0.11 | 0.14 | 0.593 |
| EEF1B2 | S106 | 0.23 | 0.36 | 0.09 | -0.23 | -0.08 | -0.55 | 0.033 |
| EEF1D | T147; S162 | -0.26 | -0.40 | -0.39 | 0.17 | 0.05 | 0.57 | 0.02 |
| EEF1D | T147; S162 | 0.00 | -0.08 | -0.18 | 0.14 | 0.03 | 0.07 | 0.059 |
| EEF1D | S162 | 0.11 | -0.01 | -0.40 | 0.20 | -0.36 | 0.31 | 0.598 |
| EEF1D | S162 | 0.11 | -0.01 | -0.40 | 0.20 | -0.36 | 0.31 | 0.598 |
| EEF1D | S162 | -0.01 | 0.04 | -0.08 | 0.11 | -0.07 | 0.00 | 0.678 |
| EEF1D | S162 | 0.02 | 0.02 | -0.07 | 0.14 | -0.05 | -0.07 | 0.792 |
| EEF1D | S133 | -0.11 | 0.11 | 0.01 | 0.01 | 0.20 | -0.26 | 0.886 |
| EEF1D | S133 | 0.04 | 0.18 | 0.02 | -0.06 | -0.04 | -0.16 | 0.054 |
| EEF1D | S133 | 0.01 | 0.28 | 0.10 | -0.12 | 0.00 | -0.34 | 0.093 |
| EEF2 | S279 | -0.10 | -0.06 | 0.27 | -0.01 | -0.06 | -0.07 | 0.521 |
| EEF2K | S500 | -0.16 | -0.46 | 0.11 | -0.14 | 0.38 | 0.11 | 0.269 |
| EEPD1 | S173 | -0.07 | 0.00 | -0.04 | 0.11 | -0.04 | 0.03 | 0.26 |
| EEPD1 | S106; S | 0.06 | 0.29 | 0.36 | -0.13 | -0.61 | -0.18 | 0.039 |
| EFCC1 | S480 | -0.10 | 0.16 | -0.15 | -0.02 | -0.04 | 0.12 | 0.68 |
| EFR3A | S694 | 0.09 | 0.19 | -0.03 | -0.21 | -0.09 | 0.01 | 0.12 |
| EFR3A | S220 | 0.08 | 0.15 | 0.12 | -0.14 | 0.00 | -0.24 | 0.028 |
| EFR3B | S216 | -0.06 | 0.10 | 0.59 | 0.03 | 0.06 | -1.34 | 0.279 |
| EFS | S287 | -0.05 | -0.30 | -0.09 | 0.20 | -0.27 | 0.37 | 0.295 |
| EFTUD2 | T86 | -0.79 | -0.47 | -0.58 | 0.56 | 0.40 | 0.31 | 9E-04 |
| EGLN1 | S125 | 0.03 | 0.41 | 0.08 | -0.58 | -0.14 | 0.01 | 0.124 |
| EHBP1 | S/T | -0.65 | 0.07 | -0.33 | 0.21 | -0.12 | 0.52 | 0.143 |
| EHBP1 | S/T | -0.28 | -0.14 | -0.42 | -0.03 | 0.66 | -0.04 | 0.124 |
| EHD1 | S456 | 0.11 | 0.12 | -0.28 | 0.23 | 0.07 | -0.35 | 0.988 |
| EHMT1 | S998 | -0.41 | -0.51 | -0.56 | 0.33 | 0.05 | 0.66 | 0.01 |
| EHMT1 | S1048 | 0.02 | -0.61 | -0.04 | 0.19 | 0.36 | -0.09 | 0.205 |
| EHMT1 | S483 | 0.03 | -0.02 | -0.40 | 0.23 | -0.08 | 0.16 | 0.232 |
| EHMT1 | S1048 | -0.23 | 0.22 | -0.05 | -0.01 | 0.12 | -0.09 | 0.885 |

| | | | | | | | | |
|---|---|---|---|---|---|---|---|---|
| EHMT1 | S483 | 0.01 | 0.03 | -0.07 | -0.57 | 0.07 | 0.37 | 0.922 |
| EHMT1 | T652 | -0.02 | -0.07 | 0.20 | -0.01 | -0.14 | 0.02 | 0.446 |
| EHMT2 | S1204 | -1.26 | -1.16 | -0.29 | 0.52 | 0.68 | 0.36 | 0.012 |
| EHMT2 | T555 | -0.22 | -0.18 | -0.28 | 0.07 | 0.09 | 0.41 | 0.021 |
| EHMT2 | S40 | -0.24 | -0.26 | -0.12 | 0.14 | 0.19 | 0.20 | 0.001 |
| EHMT2 | S569 | -0.14 | -0.23 | -0.07 | 0.47 | -0.59 | 0.32 | 0.562 |
| EHMT2 | S246 | -0.06 | 0.04 | -0.07 | -0.20 | 0.11 | 0.15 | 0.679 |
| EHMT2 | S246 | 0.08 | 0.09 | 0.03 | -0.31 | 0.15 | -0.09 | 0.33 |
| EHMT2 | S232 | 0.25 | -0.08 | 0.11 | -0.30 | 0.04 | -0.09 | 0.197 |
| EHMT2 | S119 | 0.30 | 0.26 | 0.33 | -0.34 | -0.23 | -0.59 | 0.003 |
| EHMT2 | S173 | 0.43 | 0.59 | 0.25 | -1.50 | -0.16 | 0.09 | 0.135 |
| EI24 | S326; S330 | 0.28 | 0.24 | 0.22 | -0.28 | -0.44 | -0.20 | 0.002 |
| EIF2A | S506 | 0.10 | 0.18 | 0.14 | -0.17 | -0.04 | -0.27 | 0.014 |
| EIF2AK2 | S456 | -0.03 | -0.04 | -0.22 | 0.38 | 0.11 | -0.31 | 0.501 |
| EIF2AK2 | T/S/Y | -0.14 | 0.08 | -0.05 | 0.15 | -0.14 | 0.07 | 0.599 |
| EIF2AK2 | S542 | 0.09 | 0.14 | 0.01 | -0.06 | 0.12 | -0.36 | 0.278 |
| EIF2AK3 | T982 | -0.34 | -0.02 | 0.11 | 0.00 | -0.23 | 0.37 | 0.586 |
| EIF2AK3 | S715 | 0.00 | 0.17 | 0.11 | -0.26 | -0.05 | 0.00 | 0.1 |
| EIF2AK3 | S1094 | 0.16 | 0.32 | 0.17 | -0.24 | -0.16 | -0.38 | 0.005 |
| EIF2AK3 | S555 | 0.31 | 0.26 | 0.24 | -0.32 | -0.21 | -0.48 | 0.002 |
| EIF2AK4 | T/S | -0.07 | 0.06 | -0.22 | -0.34 | 0.10 | 0.37 | 0.626 |
| EIF2AK4 | T667 | 0.16 | 0.07 | 0.07 | -0.33 | -0.24 | 0.19 | 0.231 |
| EIF2B4 | S130 | -0.27 | -0.18 | 0.02 | 0.04 | 0.01 | 0.31 | 0.105 |
| EIF2B5 | S544 | -0.08 | -0.15 | 0.04 | 0.11 | 0.06 | 0.00 | 0.123 |
| EIF2B5 | S544 | 0.05 | -0.10 | -0.13 | 0.22 | 0.03 | -0.11 | 0.398 |
| EIF2S1 | S158 | 0.09 | -0.03 | -0.18 | 0.04 | 0.12 | -0.05 | 0.469 |
| EIF2S2 | S105 | -0.07 | 0.07 | 0.07 | -0.07 | 0.16 | -0.18 | 0.654 |
| EIF3A | S1188 | -0.13 | -0.27 | 0.20 | 0.08 | -0.30 | 0.30 | 0.698 |
| EIF3A | S882 | -0.02 | 0.02 | 0.33 | -0.32 | -0.13 | 0.04 | 0.177 |
| EIF3A | S584 | 0.40 | -0.01 | 0.45 | -0.28 | -0.11 | -0.80 | 0.055 |
| EIF3B | S/T | -0.22 | -0.13 | -0.06 | 0.23 | -0.15 | 0.25 | 0.153 |
| EIF3B | S152; S154; S164 | 0.00 | -0.06 | -0.13 | 0.11 | -0.04 | 0.10 | 0.127 |
| EIF3B | S/T | -0.23 | -0.04 | 0.12 | 0.10 | -0.15 | 0.15 | 0.574 |
| EIF3B | S164; S | -0.06 | 0.00 | -0.05 | 0.11 | -0.06 | 0.05 | 0.238 |
| EIF3B | S152; S154; S164 | 0.02 | 0.13 | -0.02 | 0.00 | -0.12 | -0.02 | 0.195 |
| EIF3CL | S39 | -0.03 | -0.08 | 0.06 | 0.02 | -0.12 | 0.14 | 0.762 |
| EIF3CL | S/T | -0.08 | 0.09 | 0.01 | -0.10 | -0.20 | 0.23 | 0.849 |
| EIF3CL | S39 | 0.08 | 0.11 | 0.08 | -0.08 | -0.06 | -0.16 | 0.004 |
| EIF3CL | S166; S/T | 0.02 | 0.17 | 0.10 | 0.03 | 0.02 | -0.42 | 0.22 |
| EIF3D | T548 | 0.00 | -0.08 | -0.04 | 0.01 | 0.01 | 0.10 | 0.089 |
| EIF3D | T/S/Y | 0.01 | 0.15 | 0.09 | -0.27 | 0.20 | -0.24 | 0.306 |
| EIF3G | T/S | -0.05 | -0.14 | -0.08 | 0.20 | 0.13 | -0.08 | 0.114 |
| EIF3H | S183 | -0.36 | -0.14 | -0.10 | 0.20 | 0.13 | 0.18 | 0.012 |
| EIF3H | S183 | 0.06 | -0.06 | -0.36 | 0.05 | 0.16 | 0.08 | 0.168 |
| EIF3J | T109 | -0.19 | 0.11 | -0.15 | 0.03 | 0.17 | 0.00 | 0.251 |
| EIF3J | T109 | -0.16 | 0.02 | -0.08 | 0.02 | 0.06 | 0.12 | 0.085 |
| EIF3J | T109 | -0.13 | 0.13 | -0.02 | -0.10 | 0.14 | -0.04 | 0.969 |
| EIF4B | S597 | -0.42 | -0.13 | -0.63 | 0.34 | -0.09 | 0.57 | 0.051 |

| | | | | | | | | |
|---|---|---|---|---|---|---|---|---|
| EIF4B | S406 | -0.02 | 0.05 | -0.09 | 0.02 | -0.05 | 0.09 | 0.576 |
| EIF4B | S459 | 0.28 | -0.07 | 0.07 | 0.02 | -0.04 | -0.32 | 0.232 |
| EIF4B | S504 | 0.14 | 0.17 | 0.33 | -0.31 | -0.04 | -0.43 | 0.021 |
| EIF4B | S504 | -0.03 | 0.23 | 0.46 | -0.44 | -0.19 | -0.23 | 0.036 |
| EIF4B | S445 | 0.28 | 0.27 | 0.14 | -0.29 | -0.16 | -0.39 | 0.003 |
| EIF4B | S459 | 0.37 | 0.31 | 0.21 | -0.38 | -0.11 | -0.71 | 0.018 |
| EIF4B | S459 | 0.42 | 0.36 | 0.22 | -0.63 | -0.20 | -0.52 | 0.005 |
| EIF4EBP1 | T70 | -0.01 | 0.31 | -0.38 | 0.15 | -0.16 | -0.01 | 0.933 |
| EIF4EBP1 | T46; Y/S/T | -0.04 | -0.19 | 0.39 | -0.80 | -0.13 | 0.44 | 0.615 |
| EIF4EBP2 | T50; T/S/Y | 0.25 | -0.17 | 0.31 | -0.20 | -0.26 | -0.05 | 0.141 |
| EIF4ENIF1 | S564 | 0.15 | -0.28 | -0.15 | 0.54 | -0.11 | -0.35 | 0.704 |
| EIF4ENIF1 | S78 | -0.25 | -0.07 | 0.17 | 0.25 | -0.29 | 0.11 | 0.747 |
| EIF4ENIF1 | S752 | 0.12 | 0.02 | 0.06 | -0.05 | -0.20 | 0.04 | 0.144 |
| EIF4ENIF1 | S/T | 0.15 | 0.16 | -0.11 | 0.13 | -0.20 | -0.17 | 0.345 |
| EIF4G1 | S1209 | -0.08 | -0.09 | -0.06 | 0.04 | -0.02 | 0.20 | 0.083 |
| EIF4G1 | S1209 | -0.11 | 0.01 | -0.07 | 0.21 | 0.03 | -0.09 | 0.314 |
| EIF4G1 | S1209 | 0.17 | -0.01 | 0.03 | 0.08 | -0.08 | -0.23 | 0.261 |
| EIF4G1 | S1187 | 0.12 | -0.08 | 0.24 | 0.00 | -0.04 | -0.30 | 0.197 |
| EIF4G1 | S1231 | 0.18 | 0.14 | 0.03 | -0.10 | -0.07 | -0.22 | 0.017 |
| EIF4G1 | T205 | -0.12 | 0.14 | 0.29 | -0.66 | -0.35 | 0.42 | 0.433 |
| EIF4G2 | S | -0.19 | 0.01 | 0.42 | 0.06 | -0.28 | -0.14 | 0.387 |
| EIF4G2 | S395 | 0.10 | -0.04 | 0.26 | -0.03 | -0.05 | -0.29 | 0.121 |
| EIF4G2 | T | 0.50 | 0.24 | 0.43 | -0.01 | -0.39 | -1.71 | 0.104 |
| EIF5 | S/Y | 0.12 | 0.06 | -0.15 | -0.07 | -0.04 | 0.05 | 0.767 |
| EIF5 | S/Y | 0.10 | 0.12 | -0.06 | -0.14 | -0.15 | 0.10 | 0.295 |
| EIF5 | S389; S390 | 0.23 | 0.28 | 0.04 | -0.26 | -0.21 | -0.17 | 0.006 |
| EIF5B | S135; S137 | -0.12 | -0.14 | -0.08 | -0.01 | -0.02 | 0.31 | 0.136 |
| EIF5B | S164 | 0.13 | -0.05 | -0.11 | 0.01 | -0.02 | 0.03 | 0.795 |
| EIF5B | S107; S113 | 0.22 | -0.07 | -0.14 | -0.08 | 0.02 | 0.01 | 0.875 |
| EIF5B | S182; S183; S190 | -0.06 | 0.15 | -0.04 | -0.09 | 0.08 | -0.06 | 0.638 |
| EIF5B | S190 | 0.17 | 0.11 | 0.09 | -0.09 | -0.09 | -0.22 | 0.007 |
| EIF5B | T301 | 0.20 | 0.25 | -0.05 | -0.22 | -0.04 | -0.20 | 0.058 |
| EIF5B | S1168 | 0.14 | 0.12 | 0.16 | -0.40 | 0.05 | -0.15 | 0.076 |
| EIF5B | Y/S | 0.14 | 0.24 | 0.11 | -0.33 | -0.12 | -0.12 | 0.011 |
| EIF5B | T301 | 0.28 | 0.31 | -0.10 | -0.31 | -0.07 | -0.24 | 0.068 |
| EIF5B | S113 | 0.18 | 0.23 | 0.20 | -0.17 | -0.13 | -0.42 | 0.008 |
| EIF5B | S196 | 0.26 | 0.15 | 0.22 | -0.19 | -0.08 | -0.51 | 0.024 |
| EIF5B | S137 | 0.24 | 0.21 | 0.20 | -0.22 | -0.12 | -0.43 | 0.006 |
| EIF5B | S214 | 0.32 | 0.19 | 0.19 | -0.35 | 0.01 | -0.57 | 0.038 |
| EIF5B | S164 | 0.32 | 0.17 | 0.25 | -0.35 | -0.09 | -0.48 | 0.011 |
| EIF5B | S182; S190 | 0.20 | 0.30 | 0.27 | -0.40 | -0.28 | -0.26 | 4E-04 |
| EIF5B | S214 | 0.31 | 0.19 | 0.37 | -0.48 | -0.13 | -0.51 | 0.008 |
| EIF5B | S113 | 0.48 | 0.19 | 0.25 | -0.32 | -0.25 | -0.67 | 0.01 |
| ELAC2 | S199 | -0.67 | -0.51 | -0.34 | 0.69 | -0.24 | 0.51 | 0.052 |
| ELF1 | S163; S167 | -0.03 | -0.01 | 0.01 | -0.01 | -0.12 | 0.15 | 0.831 |
| ELF1 | S168 | 0.08 | 0.12 | 0.04 | -0.02 | -0.12 | -0.12 | 0.016 |
| ELF1 | S168 | 0.07 | 0.08 | 0.17 | -0.29 | -0.03 | -0.04 | 0.067 |
| ELF1 | S187 | 0.16 | 0.22 | 0.06 | -0.32 | 0.15 | -0.38 | 0.128 |

| | | | | | | | | |
|---|---|---|---|---|---|---|---|---|
| **ELF1** | **S334** | 0.24 | 0.12 | 0.11 | -0.14 | -0.09 | -0.31 | 0.012 |
| **ELF2** | **S191** | -0.31 | 0.11 | -0.01 | -0.04 | -0.07 | 0.26 | 0.503 |
| **ELF2** | **S372** | 0.07 | -0.36 | 0.33 | -0.09 | -0.07 | 0.04 | 0.826 |
| **ELF2** | **T/S** | 0.21 | 0.12 | -0.04 | -0.20 | -0.15 | 0.02 | 0.099 |
| **ELF2** | **S191** | 0.06 | 0.15 | 0.09 | -0.10 | 0.08 | -0.34 | 0.155 |
| **ELF4** | **T/S** | 0.02 | -0.05 | 0.18 | 0.02 | -0.06 | -0.13 | 0.264 |
| **ELF4** | **S188** | 0.05 | 0.12 | 0.12 | -0.15 | -0.05 | -0.11 | 0.006 |
| **ELK4** | **S180** | -0.07 | -0.47 | -0.08 | 0.17 | 0.09 | 0.24 | 0.054 |
| **ELL** | **S319** | 0.24 | 0.21 | 0.20 | -0.23 | 0.05 | -0.67 | 0.076 |
| **ELL2** | **S/Y** | -0.27 | -0.02 | 0.25 | 0.36 | 0.03 | -0.52 | 0.918 |
| **ELL2** | **S503** | 0.12 | -0.08 | 0.34 | 0.07 | -0.10 | -0.47 | 0.222 |
| **ELL2** | **S** | 0.31 | 0.13 | -0.03 | -0.01 | 0.00 | -0.54 | 0.191 |
| **ELL3** | **S239** | -0.03 | -0.13 | -0.38 | 0.08 | 0.05 | 0.33 | 0.074 |
| **ELOA** | **S151** | 0.09 | 0.26 | -0.01 | -0.18 | -0.02 | -0.18 | 0.062 |
| **EMC2** | **T121** | 0.10 | 0.18 | 0.15 | -0.29 | -0.01 | -0.20 | 0.023 |
| **EMC4** | **S36** | -0.15 | -0.47 | -0.40 | 0.14 | 0.16 | 0.49 | 0.016 |
| **EMC4** | **S36** | 0.16 | -0.03 | 0.07 | 0.00 | 0.17 | -0.47 | 0.447 |
| **EMD** | **S141** | -0.42 | -0.45 | -0.57 | 0.28 | 0.23 | 0.55 | 0.002 |
| **EMD** | **S143** | -0.34 | -0.62 | -0.38 | 0.20 | 0.12 | 0.64 | 0.014 |
| **EMD** | **S/T** | -0.39 | -0.38 | -0.45 | 0.21 | -0.01 | 0.66 | 0.026 |
| **EMD** | **S/T** | -0.22 | -0.50 | -0.38 | 0.19 | 0.21 | 0.45 | 0.005 |
| **EMD** | **S49; S52** | -0.10 | -0.08 | -0.17 | 0.17 | 0.05 | 0.11 | 0.007 |
| **EMD** | **S87** | -0.08 | -0.25 | 0.03 | 0.02 | 0.06 | 0.18 | 0.12 |
| **EMD** | **S49** | -0.16 | -0.13 | -0.01 | 0.12 | -0.18 | 0.30 | 0.293 |
| **EMD** | **S60** | -0.22 | -0.24 | 0.20 | -0.15 | 0.14 | 0.19 | 0.448 |
| **EMD** | **S163** | 0.18 | -0.16 | -0.04 | 0.09 | -0.21 | 0.10 | 0.996 |
| **EMD** | **S110** | -0.02 | -0.13 | 0.14 | -0.17 | 0.24 | -0.10 | 0.974 |
| **EMD** | **S87** | 0.09 | 0.04 | -0.11 | -0.03 | -0.15 | 0.14 | 0.844 |
| **EMD** | **S49** | 0.14 | 0.04 | -0.10 | -0.03 | -0.10 | 0.03 | 0.491 |
| **EME1** | **S111** | -0.18 | -0.07 | -0.31 | 0.13 | 0.07 | 0.27 | 0.022 |
| **EME1** | **S84** | -0.04 | 0.13 | -0.14 | -0.05 | -0.02 | 0.10 | 0.79 |
| **EME1** | **S117** | 0.02 | 0.01 | -0.04 | -0.04 | -0.12 | 0.15 | 0.968 |
| **EMG1** | **S14** | 0.19 | -0.06 | 0.03 | 0.22 | -0.17 | -0.28 | 0.48 |
| **EML4** | **S/T** | -0.22 | -0.13 | -0.03 | 0.16 | 0.11 | 0.07 | 0.018 |
| **EML4** | **S/T** | -0.22 | -0.13 | -0.03 | 0.16 | 0.11 | 0.07 | 0.018 |
| **EML4** | **S176** | 0.05 | -0.01 | -0.24 | 0.08 | -0.27 | 0.31 | 0.592 |
| **EML4** | **S146** | 0.08 | -0.31 | 0.06 | 0.20 | -0.03 | -0.04 | 0.544 |
| **EMSY** | **S168; S173** | -0.39 | -0.40 | -0.21 | 0.13 | 0.26 | 0.40 | 0.004 |
| **EMSY** | **S209** | -0.13 | 0.00 | -0.33 | 0.44 | -0.63 | 0.36 | 0.589 |
| **EMSY** | **T207; S209** | 0.06 | -0.09 | -0.02 | -0.04 | -0.19 | 0.24 | 0.894 |
| **EMSY** | **S/T** | 0.11 | 0.02 | -0.06 | -0.18 | 0.02 | 0.07 | 0.626 |
| **EMSY** | **S195** | 0.05 | 0.13 | 0.08 | -0.09 | -0.10 | -0.08 | 0.002 |
| **EMSY** | **S407** | 0.15 | 0.09 | 0.06 | -0.21 | 0.10 | -0.23 | 0.121 |
| **EMSY** | **S209** | 0.23 | 0.28 | 0.13 | -0.45 | 0.15 | -0.57 | 0.092 |
| **EMSY** | **S238** | 0.11 | 0.40 | 0.28 | -0.33 | -0.42 | -0.25 | 0.004 |
| **ENTR1** | **S243** | -0.33 | 0.03 | -0.19 | 0.00 | -0.18 | 0.51 | 0.309 |
| **ENTREP3** | **S389** | -0.40 | -0.25 | -0.24 | 0.45 | -0.50 | 0.57 | 0.237 |
| **EP300** | **S1726** | -0.23 | -0.20 | 0.06 | -0.08 | 0.15 | 0.24 | 0.163 |

| | | | | | | | | |
|---|---|---|---|---|---|---|---|---|
| EP300 | T887 | 0.19 | -0.12 | 0.08 | -0.08 | 0.07 | -0.17 | 0.401 |
| EP300 | S1038 | 0.07 | 0.22 | -0.16 | -0.04 | 0.22 | -0.40 | 0.606 |
| EP300 | S1038 | 0.08 | 0.11 | 0.00 | -0.08 | 0.07 | -0.21 | 0.182 |
| EP300 | S1726 | 0.00 | 0.12 | 0.33 | -0.14 | -0.47 | 0.05 | 0.132 |
| EP400 | S2086 | -0.82 | -0.50 | -0.76 | 0.59 | 0.39 | 0.40 | 7E-04 |
| EP400 | S736 | 0.11 | -0.06 | 0.04 | 0.07 | -0.22 | 0.02 | 0.527 |
| EP400 | S722 | 0.27 | 0.32 | 0.12 | -0.44 | -0.17 | -0.28 | 0.006 |
| EP400 | S1732 | 0.57 | 0.17 | 0.13 | -0.36 | -0.57 | -0.25 | 0.015 |
| EPB41 | S/T | -0.45 | 0.02 | 0.32 | 0.07 | 0.02 | -0.09 | 0.879 |
| EPB41 | S555 | -0.04 | 0.17 | 0.30 | -0.05 | -0.23 | -0.23 | 0.053 |
| EPB41 | S188 | 0.08 | 0.17 | 0.25 | -0.59 | -0.17 | 0.10 | 0.136 |
| EPB41L1 | S784 | -0.11 | 0.10 | 0.02 | -0.03 | -0.10 | 0.10 | 0.86 |
| EPB41L1 | S441 | 0.52 | 0.21 | 0.18 | -0.17 | -0.42 | -0.64 | 0.015 |
| EPB41L2 | S | 0.00 | -0.04 | 0.13 | 0.02 | 0.09 | -0.21 | 0.553 |
| EPB41L4A | S426 | -1.71 | -0.36 | 0.03 | 1.02 | -0.14 | -0.06 | 0.213 |
| EPB41L4A | S426 | -0.75 | -0.12 | -0.03 | 0.67 | -0.36 | 0.18 | 0.286 |
| EPB41L4A | S448 | 0.02 | -0.11 | 0.13 | 0.00 | -0.05 | -0.01 | 0.645 |
| EPB41L4A | S541; S543 | 0.04 | -0.02 | 0.05 | 0.15 | -0.11 | -0.12 | 0.632 |
| EPB41L4A | S448 | -0.02 | -0.10 | 0.21 | -0.08 | 0.05 | -0.08 | 0.582 |
| EPB41L4A | S614 | 0.13 | -0.09 | 0.20 | 0.06 | -0.12 | -0.24 | 0.219 |
| EPB41L4A | S543 | -0.04 | 0.14 | 0.21 | -0.15 | -0.12 | -0.08 | 0.053 |
| EPB41L4A | S389 | 0.10 | 0.16 | 0.15 | -0.13 | -0.08 | -0.23 | 0.004 |
| EPB41L4A | S402 | 0.04 | 0.20 | 0.29 | -0.02 | -0.17 | -0.47 | 0.06 |
| EPB41L4A | S366 | 0.40 | 0.10 | 0.31 | -0.41 | -0.05 | -0.64 | 0.031 |
| EPB41L5 | S418; S436 | -0.36 | -0.37 | 0.21 | 0.31 | 0.42 | -0.50 | 0.51 |
| EPB41L5 | S418 | -0.03 | -0.02 | -0.16 | -0.37 | 0.13 | 0.34 | 0.656 |
| EPHA1 | T780 | 0.08 | 0.11 | 0.17 | -0.12 | -0.15 | -0.14 | 7E-04 |
| EPHA2 | S570 | -0.02 | -0.36 | 0.05 | -0.15 | 0.03 | 0.35 | 0.379 |
| EPHA2 | S/T | -0.15 | -0.43 | 0.30 | -1.27 | 0.34 | 0.53 | 0.955 |
| EPHB3 | T865 | -0.16 | 0.01 | 0.10 | 0.18 | -0.46 | 0.21 | 0.971 |
| EPHB4 | T976 | -0.22 | -0.07 | 0.27 | -0.20 | 0.00 | 0.16 | 0.982 |
| EPN2 | S/T/Y | -0.75 | 0.15 | -0.47 | 0.49 | 0.28 | -0.07 | 0.133 |
| EPN2 | S/T/Y | -0.47 | 0.02 | -0.06 | 0.23 | -0.06 | 0.23 | 0.17 |
| EPN3 | S/Y | -0.05 | -0.14 | -0.23 | 0.42 | -0.18 | 0.07 | 0.252 |
| EPPK1 | S4850 | -0.60 | -0.67 | -0.64 | 0.74 | -0.24 | 0.64 | 0.031 |
| EPPK1 | S1299 | -0.08 | -0.02 | -0.29 | 0.05 | 0.22 | 0.08 | 0.07 |
| EPPK1 | S4645 | -0.03 | -0.17 | -0.04 | 0.03 | 0.06 | 0.14 | 0.053 |
| EPPK1 | S1821 | 0.21 | -0.16 | 0.01 | -0.57 | -0.43 | 0.61 | 0.717 |
| EPRS1 | S/T/Y | 0.12 | -0.23 | -0.83 | 0.50 | -0.15 | 0.24 | 0.206 |
| EPRS1 | S/T/Y | -0.14 | -0.14 | -0.25 | 0.22 | 0.15 | 0.10 | 0.003 |
| EPRS1 | S886 | 0.09 | -0.02 | -0.05 | 0.04 | 0.01 | -0.08 | 0.722 |
| EPRS1 | S/T/Y | -0.11 | 0.08 | 0.14 | -0.18 | 0.08 | -0.03 | 0.494 |
| EPRS1 | S/T/Y | 0.12 | 0.07 | 0.13 | -0.15 | -0.05 | -0.15 | 0.004 |
| EPS15 | S814 | -0.03 | -0.19 | -0.11 | 0.17 | 0.29 | -0.20 | 0.268 |
| EPS15 | T777 | -0.09 | -0.10 | -0.05 | 0.00 | 0.07 | 0.15 | 0.027 |
| EPS15 | S563 | -0.05 | 0.03 | -0.18 | 0.19 | -0.14 | 0.11 | 0.357 |
| EPS15 | S563 | 0.01 | -0.02 | -0.17 | 0.11 | 0.18 | -0.15 | 0.41 |
| EPS15 | S796 | -0.12 | 0.11 | 0.14 | -0.06 | 0.11 | -0.21 | 0.471 |

| | | | | | | | | |
|---|---|---|---|---|---|---|---|---|
| **EPS15** | **S324** | -0.06 | 0.02 | 0.17 | -0.19 | 0.14 | -0.12 | 0.441 |
| **EPS15L1** | **S672** | -0.59 | 0.06 | -0.04 | 0.49 | -0.18 | 0.05 | 0.336 |
| **EPS15L1** | **T366** | -0.07 | -0.06 | -0.11 | 0.19 | -0.02 | 0.05 | 0.078 |
| **EPS8** | **S58** | 0.03 | -0.31 | -0.08 | 0.11 | 0.11 | 0.10 | 0.086 |
| **EPS8** | **T** | -0.09 | 0.11 | -0.17 | 0.15 | -0.04 | 0.01 | 0.425 |
| **EPS8** | **S661** | 0.03 | 0.06 | 0.07 | -0.25 | 0.16 | -0.11 | 0.387 |
| **EPS8L1** | **T187** | 0.01 | 0.11 | 0.27 | -0.41 | -0.07 | 0.00 | 0.123 |
| **EPS8L2** | **S570** | 0.11 | 0.12 | 0.20 | 0.18 | -0.18 | -0.58 | 0.206 |
| **ERAL1** | **S173** | 0.14 | -0.31 | -0.15 | 0.03 | 0.28 | -0.05 | 0.299 |
| **ERBB2** | **T733** | -0.02 | -0.57 | -0.41 | 0.64 | -0.01 | 0.05 | 0.1 |
| **ERBB2** | **T701** | -0.29 | -0.10 | 0.03 | 0.12 | -0.02 | 0.21 | 0.125 |
| **ERBB2** | **S1049; S1050; S1051** | -0.16 | -0.06 | -0.09 | 0.01 | 0.08 | 0.19 | 0.033 |
| **ERBB2** | **T/S** | -0.35 | -0.15 | 0.19 | -0.13 | 0.08 | 0.27 | 0.41 |
| **ERBB2** | **T/S** | -0.20 | 0.02 | -0.10 | 0.18 | -0.03 | 0.10 | 0.112 |
| **ERBB2** | **S1054; S1066** | -0.07 | -0.18 | -0.02 | 0.01 | 0.13 | 0.10 | 0.045 |
| **ERBB2** | **S1107** | -0.49 | 0.12 | 0.15 | -0.08 | 0.10 | 0.10 | 0.629 |
| **ERBB2** | **T701** | -0.01 | 0.01 | -0.15 | 0.09 | -0.07 | 0.12 | 0.277 |
| **ERBB2** | **S1054** | -0.09 | 0.00 | -0.06 | 0.02 | 0.03 | 0.09 | 0.06 |
| **ERBB2** | **T1240** | -0.05 | 0.09 | -0.17 | 0.05 | 0.09 | -0.03 | 0.411 |
| **ERBB2** | **S1151** | 0.12 | 0.09 | -0.31 | 0.08 | 0.31 | -0.42 | 0.94 |
| **ERBB2** | **S1151** | -0.17 | 0.08 | 0.09 | 0.05 | 0.17 | -0.28 | 0.902 |
| **ERBB2** | **S1083** | -0.07 | 0.20 | 0.03 | -0.03 | -0.06 | -0.09 | 0.252 |
| **ERBB2** | **T701** | -0.04 | 0.10 | 0.17 | -0.15 | 0.03 | -0.14 | 0.126 |
| **ERBB2** | **S1083** | 0.11 | 0.17 | 0.04 | -0.04 | 0.15 | -0.54 | 0.312 |
| **ERBB3** | **S686** | -0.06 | -0.09 | -0.02 | 0.09 | -0.08 | 0.16 | 0.208 |
| **ERBB3** | **S982** | -0.16 | 0.34 | -0.09 | -0.13 | 0.52 | -0.83 | 0.693 |
| **ERBB3** | **S982** | 0.15 | 0.08 | 0.14 | -0.12 | 0.10 | -0.43 | 0.149 |
| **ERBIN** | **S1179** | -0.42 | -0.28 | -0.34 | 0.20 | -0.04 | 0.61 | 0.036 |
| **ERBIN** | **T485** | -0.50 | -0.31 | -0.20 | 0.52 | 0.00 | 0.25 | 0.027 |
| **ERBIN** | **S1310** | -0.05 | -0.46 | -0.01 | 0.39 | -0.38 | 0.31 | 0.384 |
| **ERBIN** | **S/T** | -0.24 | -0.11 | -0.08 | 0.07 | 0.18 | 0.14 | 0.009 |
| **ERBIN** | **S852** | -0.11 | -0.09 | -0.09 | 0.04 | 0.13 | 0.10 | 0.003 |
| **ERBIN** | **S/T** | 0.21 | -0.12 | -0.24 | -0.11 | 0.06 | 0.14 | 0.63 |
| **ERBIN** | **S715** | 0.24 | -0.20 | -0.19 | -0.18 | 0.13 | 0.13 | 0.69 |
| **ERBIN** | **S602; S603** | 0.08 | 0.11 | -0.04 | -0.01 | 0.01 | -0.16 | 0.228 |
| **ERBIN** | **S1310** | -0.02 | -0.12 | 0.30 | -0.24 | -0.33 | 0.30 | 0.57 |
| **ERBIN** | **S440** | -0.01 | 0.04 | 0.21 | 0.00 | -0.35 | 0.05 | 0.279 |
| **ERCC5** | **S526** | 0.49 | -0.65 | -0.24 | -0.05 | 0.20 | -0.02 | 0.634 |
| **ERCC5** | **T338** | 0.05 | -0.07 | 0.18 | -0.11 | -0.19 | 0.12 | 0.383 |
| **ERCC6** | **S429; S430** | -0.19 | -0.17 | -0.21 | 0.22 | -0.05 | 0.31 | 0.032 |
| **ERCC6** | **T303** | -0.19 | -0.11 | -0.27 | -0.13 | 0.02 | 0.52 | 0.181 |
| **ERCC6** | **S486; S489** | -0.24 | 0.06 | 0.00 | -0.02 | 0.12 | 0.05 | 0.335 |
| **ERCC6** | **S554** | -0.08 | -0.07 | 0.01 | 0.05 | -0.03 | 0.12 | 0.152 |
| **ERCC6** | **S158** | 0.09 | -0.08 | 0.08 | -0.02 | 0.11 | -0.21 | 0.54 |
| **ERCC6** | **T303** | 0.30 | 0.27 | -0.68 | -1.12 | 0.41 | 0.21 | 0.83 |
| **ERCC6** | **S489** | 0.06 | 0.23 | -0.09 | -0.17 | -0.26 | 0.17 | 0.398 |
| **ERCC6** | **S/T** | 0.20 | 0.07 | 0.13 | -0.16 | -0.13 | -0.15 | 0.002 |
| **ERCC6** | **S** | 0.44 | 0.23 | 0.19 | -0.40 | -0.36 | -0.34 | 0.001 |

| | | | | | | | | |
|---|---|---|---|---|---|---|---|---|
| ERCC6L | T/S | 0.18 | -0.16 | 0.14 | -0.33 | -0.69 | 0.55 | 0.62 |
| ERF | S431; S/T | -0.17 | 0.05 | -0.20 | 0.01 | 0.09 | 0.19 | 0.091 |
| ERF | S431; S435; T441 | 0.14 | -0.21 | -0.14 | 0.27 | 0.07 | -0.20 | 0.537 |
| ERF | S21 | 0.21 | 0.17 | 0.27 | -0.34 | -0.07 | -0.38 | 0.008 |
| ERGIC3 | S | 0.06 | 0.05 | 0.13 | 0.01 | -0.14 | -0.14 | 0.036 |
| ERGIC3 | S116 | 0.17 | 0.28 | 0.24 | -0.34 | -0.17 | -0.32 | 0.001 |
| ERH | S24 | 0.18 | 0.11 | 0.07 | -0.26 | -0.03 | -0.11 | 0.026 |
| ERH | S24 | 0.09 | 0.13 | 0.19 | -0.12 | -0.09 | -0.26 | 0.008 |
| ERH | S24 | 0.18 | 0.18 | 0.10 | -0.03 | -0.13 | -0.39 | 0.036 |
| ERMP1 | S53 | -0.14 | -0.05 | -0.03 | 0.02 | -0.11 | 0.28 | 0.314 |
| ERMP1 | S53 | 0.02 | -0.07 | -0.13 | 0.00 | -0.08 | 0.24 | 0.349 |
| ERMP1 | S44 | 0.23 | 0.15 | 0.01 | -0.24 | -0.20 | -0.01 | 0.042 |
| ERMP1 | S53 | 0.24 | 0.07 | 0.10 | 0.15 | 0.02 | -0.79 | 0.319 |
| ESCO1 | S200 | -0.53 | -0.10 | -0.02 | 0.04 | -0.08 | 0.50 | 0.191 |
| ESCO1 | S279 | -0.43 | -0.10 | -0.07 | 0.38 | -0.20 | 0.27 | 0.176 |
| ESCO2 | S512 | -0.07 | -0.24 | 0.21 | -0.06 | -0.02 | 0.13 | 0.765 |
| ESCO2 | S223 | 0.11 | -0.26 | 0.17 | 0.03 | -0.09 | 0.00 | 0.843 |
| ESCO2 | S75 | 0.57 | 0.52 | -0.83 | -0.41 | -0.25 | -0.11 | 0.501 |
| ESF1 | S180 | -0.66 | -0.42 | 0.14 | 0.42 | -0.02 | 0.27 | 0.117 |
| ESF1 | S657; S663 | -0.10 | -0.12 | -0.18 | 0.17 | -0.07 | 0.25 | 0.067 |
| ESF1 | S657; S663 | -0.04 | -0.25 | 0.04 | 0.20 | -0.19 | 0.18 | 0.392 |
| ESF1 | S657 | -0.15 | 0.10 | -0.14 | 0.14 | -0.16 | 0.17 | 0.454 |
| ESF1 | S657 | 0.02 | 0.02 | -0.12 | 0.02 | -0.09 | 0.13 | 0.619 |
| ESF1 | S77; S79; S82 | 0.12 | 0.04 | 0.09 | 0.03 | -0.02 | -0.30 | 0.166 |
| ESF1 | S198 | 0.11 | 0.16 | 0.15 | -0.21 | 0.02 | -0.29 | 0.031 |
| ESF1 | S153 | 0.24 | 0.20 | 0.31 | -0.33 | -0.11 | -0.51 | 0.01 |
| ESF1 | S82 | 0.43 | 0.46 | 0.37 | -0.62 | -0.36 | -0.87 | 0.002 |
| ESF1 | S/Y | 0.38 | 0.75 | 0.47 | -0.86 | -0.34 | -1.77 | 0.024 |
| ESPN | S634 | 0.18 | 0.17 | 0.00 | -0.04 | -0.08 | -0.28 | 0.053 |
| ESRP1 | S237 | -0.41 | -0.29 | -0.02 | 0.10 | 0.01 | 0.44 | 0.072 |
| ESRP1 | S543 | 0.09 | -0.09 | -0.07 | -0.02 | -0.06 | 0.14 | 0.613 |
| ESRP1 | S543 | -0.01 | -0.06 | 0.09 | 0.15 | -0.21 | 0.01 | 0.868 |
| ESRP2 | S573 | -0.13 | 0.13 | 0.32 | -0.03 | 0.00 | -0.37 | 0.253 |
| ESRP2 | S573 | -0.23 | 0.09 | 0.75 | -0.49 | 0.00 | -0.54 | 0.177 |
| ESRRA | S19; S22; T31 | -0.07 | -0.04 | 0.00 | 0.11 | -0.09 | 0.08 | 0.331 |
| ESRRA | S26 | 0.03 | -0.05 | -0.04 | 0.00 | -0.09 | 0.14 | 0.624 |
| ESS2 | T437; S446 | -0.07 | 0.00 | 0.03 | 0.05 | -0.12 | 0.10 | 0.819 |
| ESS2 | S292 | 0.04 | 0.06 | -0.04 | 0.00 | -0.08 | 0.02 | 0.431 |
| ESS2 | S391; S395 | -0.02 | 0.02 | 0.10 | -0.11 | -0.21 | 0.19 | 0.587 |
| ESS2 | S391; S395 | -0.04 | 0.18 | -0.03 | -0.07 | -0.11 | 0.05 | 0.43 |
| ESS2 | S395 | -0.12 | -0.34 | 0.52 | 0.33 | -0.53 | -0.15 | 0.723 |
| ESS2 | S421 | 0.25 | 0.08 | -0.03 | -0.12 | 0.00 | -0.22 | 0.105 |
| ESS2 | T333 | 0.03 | 0.07 | 0.22 | -0.22 | -0.11 | -0.03 | 0.046 |
| ESS2 | S446 | 0.25 | 0.17 | -0.01 | -0.15 | -0.10 | -0.21 | 0.026 |
| ESS2 | T437 | 0.15 | 0.21 | 0.09 | -0.16 | -0.09 | -0.25 | 0.006 |
| ESS2 | T386 | 0.02 | 0.35 | 0.22 | -0.48 | -0.45 | 0.14 | 0.11 |
| ESYT1 | S1034 | -0.02 | -0.02 | 0.10 | 0.01 | 0.05 | -0.13 | 0.591 |
| ESYT2 | S676 | -0.03 | -0.10 | -0.10 | 0.10 | 0.08 | 0.04 | 0.008 |

| | | | | | | | | |
|---|---|---|---|---|---|---|---|---|
| ESYT2 | T/S | -0.02 | -0.16 | 0.14 | 0.16 | -0.12 | -0.02 | 0.85 |
| ESYT2 | S758; S761 | -0.08 | 0.07 | 0.02 | 0.21 | -0.33 | 0.05 | 0.888 |
| ETV6 | S439 | -0.17 | -0.34 | -0.13 | 0.17 | 0.16 | 0.22 | 0.004 |
| ETV6 | S439 | -0.12 | -0.30 | -0.15 | 0.27 | -0.09 | 0.28 | 0.062 |
| ETV6 | S213 | -0.08 | -0.04 | -0.17 | 0.07 | -0.08 | 0.25 | 0.159 |
| ETV6 | T187; S/T | 0.02 | 0.11 | 0.17 | -0.13 | 0.20 | -0.48 | 0.31 |
| ETV6 | S193 | -0.17 | -0.15 | 0.60 | -0.19 | -0.03 | -0.25 | 0.391 |
| EXO1 | S639 | -0.09 | -0.02 | 0.02 | 0.06 | 0.22 | -0.23 | 0.715 |
| EXO1 | S746 | 0.23 | 0.08 | 0.13 | -0.10 | -0.33 | -0.07 | 0.026 |
| EXOSC1 | S21 | 0.35 | 0.01 | -0.03 | 0.25 | -0.21 | -0.55 | 0.353 |
| EXOSC2 | S124 | -0.38 | -0.48 | 0.33 | 0.09 | 0.09 | 0.18 | 0.309 |
| EXOSC7 | S177 | -0.34 | -0.22 | -0.32 | 0.41 | 0.13 | 0.17 | 0.005 |
| EXOSC9 | S306 | 0.07 | -0.01 | 0.00 | 0.12 | -0.23 | 0.03 | 0.704 |
| EXPH5 | S706 | -0.30 | -0.21 | -0.11 | 0.10 | 0.20 | 0.24 | 0.005 |
| EXPH5 | S1821 | -0.46 | 0.13 | 0.02 | 0.06 | -0.09 | 0.24 | 0.448 |
| EXPH5 | S910 | -0.14 | -0.11 | 0.04 | -0.04 | -0.03 | 0.25 | 0.316 |
| EXPH5 | S1238 | -0.04 | -0.13 | -0.02 | 0.07 | 0.02 | 0.09 | 0.039 |
| EXPH5 | S393 | -0.01 | 0.01 | -0.08 | -0.20 | 0.08 | 0.17 | 0.731 |
| EXPH5 | S1753 | 0.07 | 0.00 | -0.14 | 0.05 | -0.16 | 0.15 | 0.764 |
| EXPH5 | S604 | 0.41 | -0.22 | -0.41 | -0.83 | 0.12 | 0.50 | 0.994 |
| EXPH5 | S948 | -0.12 | 0.02 | 0.12 | 0.01 | -0.09 | 0.06 | 0.879 |
| EXPH5 | S910 | -0.16 | 0.15 | 0.03 | -0.09 | 0.05 | 0.01 | 0.872 |
| EXPH5 | S/T | -0.20 | -0.05 | 0.27 | 0.24 | -0.16 | -0.18 | 0.831 |
| EXPH5 | S948 | -0.06 | 0.03 | 0.12 | -0.05 | -0.12 | 0.06 | 0.439 |
| EXPH5 | S1444 | -0.19 | -0.03 | 0.32 | -0.03 | 0.06 | -0.19 | 0.625 |
| EXPH5 | S604 | 0.02 | -0.02 | 0.14 | -0.22 | 0.14 | -0.08 | 0.432 |
| EXPH5 | S393 | 0.06 | 0.03 | 0.08 | -0.10 | -0.04 | -0.03 | 0.009 |
| EXPH5 | T/S | -0.01 | 0.02 | 0.15 | -0.40 | 0.28 | -0.14 | 0.534 |
| EXPH5 | S | -0.01 | 0.07 | 0.19 | -0.29 | 0.07 | -0.07 | 0.21 |
| EXPH5 | T465 | -0.10 | 0.09 | 0.58 | -0.18 | 0.11 | -0.87 | 0.232 |
| EYA3 | S/T | -0.32 | -0.11 | 0.06 | 0.12 | 0.02 | 0.18 | 0.126 |
| EYA3 | S438 | 0.11 | -0.29 | -0.02 | 0.32 | -0.31 | 0.08 | 0.68 |
| EYA3 | T/S | 0.12 | -0.08 | 0.02 | -0.11 | 0.12 | -0.10 | 0.631 |
| EZH2 | S362; S363 | -0.23 | -0.24 | -0.22 | 0.19 | -0.02 | 0.39 | 0.025 |
| EZH2 | T367; S/T | -0.22 | -0.06 | -0.24 | 0.09 | 0.04 | 0.32 | 0.036 |
| EZH2 | T339 | -0.29 | -0.03 | -0.16 | 0.09 | 0.14 | 0.19 | 0.019 |
| EZH2 | S412 | -0.03 | -0.04 | -0.22 | 0.10 | 0.02 | 0.14 | 0.064 |
| EZH2 | S/T | 0.14 | -0.05 | -0.08 | -0.06 | 0.12 | -0.09 | 0.871 |
| EZH2 | T487 | -0.33 | 0.22 | 0.06 | -0.64 | -0.09 | 0.51 | 0.89 |
| EZH2 | T487 | 0.05 | 0.07 | 0.05 | -0.07 | -0.07 | -0.03 | 0.001 |
| EZH2 | T487 | 0.08 | 0.14 | 0.04 | -0.19 | 0.05 | -0.14 | 0.086 |
| EZH2 | S/T | 0.33 | 0.10 | -0.04 | 0.04 | -0.25 | -0.27 | 0.124 |
| FAAP100 | S667 | -0.03 | -0.22 | -0.02 | 0.41 | -0.10 | -0.12 | 0.455 |
| FAF2 | S355 | -0.08 | 0.35 | -0.39 | 0.31 | -0.02 | -0.35 | 0.95 |
| FAM104A | S21 | -0.07 | -0.10 | 0.25 | -0.01 | -0.14 | 0.04 | 0.635 |
| FAM110A | S | -0.19 | 0.13 | -0.11 | -0.03 | -0.13 | 0.28 | 0.58 |
| FAM110A | T140 | 0.41 | -0.28 | -0.24 | 0.39 | -0.17 | -0.31 | 0.976 |
| FAM110A | S189 | 0.13 | 0.13 | -0.10 | -0.05 | -0.28 | 0.12 | 0.439 |

| | | | | | | | | |
|---|---|---|---|---|---|---|---|---|
| FAM111A | S26 | 0.30 | 0.12 | -0.53 | 0.10 | 0.21 | -0.40 | 0.979 |
| FAM117A | S178 | -0.13 | -0.10 | 0.38 | 0.11 | -0.29 | -0.07 | 0.547 |
| FAM117A | S207 | 0.20 | 0.06 | -0.08 | -0.02 | 0.06 | -0.26 | 0.352 |
| FAM117A | S213 | 0.15 | 0.39 | -0.07 | -0.22 | -0.15 | -0.21 | 0.062 |
| FAM117B | S/T | -0.23 | -0.15 | -0.30 | -0.45 | -0.10 | 0.82 | 0.451 |
| FAM117B | S106 | -0.38 | -0.17 | 0.03 | 0.12 | 0.30 | -0.01 | 0.108 |
| FAM117B | S192 | 0.34 | 0.05 | -0.04 | 0.12 | -0.29 | -0.29 | 0.201 |
| FAM120A | S513 | -0.07 | -0.02 | -0.08 | -0.04 | 0.13 | 0.08 | 0.105 |
| FAM120A | T655 | -0.12 | 0.04 | -0.07 | 0.14 | 0.21 | -0.26 | 0.634 |
| FAM120A | S1023 | 0.36 | 0.23 | 0.29 | -0.39 | 0.01 | -0.84 | 0.049 |
| FAM133B | S82 | 0.42 | 0.18 | 0.17 | -0.41 | -0.17 | -0.39 | 0.007 |
| FAM133B | S191; S196 | 0.29 | 0.26 | 0.30 | -0.40 | -0.08 | -0.64 | 0.016 |
| FAM156A | S11 | -0.14 | -0.09 | 0.09 | 0.02 | 0.11 | -0.01 | 0.335 |
| FAM174C | S120 | -0.45 | -0.33 | 0.27 | 0.12 | -0.13 | 0.34 | 0.342 |
| FAM174C | S120 | 0.17 | 0.14 | -0.17 | 0.13 | 0.04 | -0.39 | 0.575 |
| FAM193A | S1144 | -0.35 | 0.33 | -0.22 | -0.88 | 0.51 | 0.19 | 0.969 |
| FAM193A | S393 | -0.18 | -0.01 | 0.21 | 0.05 | -0.03 | -0.06 | 0.876 |
| FAM234A | S21 | 0.08 | 0.31 | 0.42 | -0.40 | -0.16 | -0.49 | 0.011 |
| FAM234A | S21 | 0.19 | 0.05 | 0.56 | -0.23 | -0.13 | -0.78 | 0.064 |
| FAM241A | S56 | -0.05 | 0.09 | -0.02 | -0.10 | 0.11 | -0.04 | 0.792 |
| FAM241A | S56 | 0.04 | 0.06 | 0.00 | -0.16 | 0.22 | -0.21 | 0.579 |
| FAM241A | S56 | 0.04 | 0.06 | 0.00 | -0.16 | 0.22 | -0.21 | 0.579 |
| FAM246B | S35 | -0.09 | -0.11 | 0.38 | -0.27 | 0.01 | -0.01 | 0.463 |
| FAM53C | S162 | -0.38 | 0.11 | 0.06 | -0.08 | 0.15 | 0.08 | 0.52 |
| FAM76B | T215 | 0.01 | 0.01 | 0.06 | -0.14 | 0.01 | 0.04 | 0.384 |
| FAM76B | T215 | 0.04 | 0.06 | 0.06 | 0.01 | -0.03 | -0.16 | 0.098 |
| FAM76B | T215 | 0.04 | 0.00 | 0.21 | 0.00 | -0.09 | -0.19 | 0.109 |
| FAM76B | S193 | 0.05 | 0.12 | 0.16 | -0.01 | -0.04 | -0.35 | 0.096 |
| FAM76B | T215 | 0.28 | -0.10 | 0.22 | 0.15 | -0.31 | -0.38 | 0.197 |
| FAM76B | S193 | -0.10 | 0.31 | 0.16 | 0.01 | 0.20 | -0.86 | 0.381 |
| FAM76B | S338 | 0.28 | -0.06 | 0.29 | -0.13 | -0.14 | -0.35 | 0.052 |
| FAM83B | S466 | 0.00 | 0.10 | -0.01 | 0.10 | -0.07 | -0.14 | 0.459 |
| FAM83B | S869 | 0.15 | 0.04 | -0.07 | 0.06 | -0.25 | 0.04 | 0.488 |
| FAM83B | S/T | 0.02 | 0.13 | 0.14 | 0.04 | -0.29 | -0.08 | 0.115 |
| FAM83B | S729 | -0.05 | -0.01 | 0.35 | -0.17 | -0.13 | -0.06 | 0.181 |
| FAM83B | T/S | 0.12 | 0.22 | 0.00 | -0.14 | -0.02 | -0.23 | 0.051 |
| FAM83D | S493 | -0.64 | -0.55 | -0.63 | 0.81 | -0.23 | 0.51 | 0.035 |
| FAM83D | S/T/Y | -0.05 | 0.00 | -0.11 | 0.00 | -0.05 | 0.19 | 0.279 |
| FAM83D | S462 | -0.05 | -0.10 | 0.07 | -0.05 | 0.05 | 0.07 | 0.459 |
| FAM83G | S634 | -0.07 | 0.10 | -0.39 | -0.06 | 0.17 | 0.17 | 0.256 |
| FAM83G | S616 | 0.13 | -0.16 | -0.22 | 0.41 | -0.28 | 0.01 | 0.605 |
| FAM83G | S760 | -0.18 | -0.15 | 0.13 | -0.05 | 0.15 | 0.06 | 0.363 |
| FAM83G | S356 | 0.00 | -0.07 | -0.09 | 0.18 | -0.10 | 0.06 | 0.307 |
| FAM83H | S1048 | -1.63 | -1.50 | -1.27 | 0.90 | 0.57 | 0.94 | 1E-04 |
| FAM83H | S342 | -0.93 | -1.08 | -1.32 | 0.54 | 0.32 | 0.93 | 0.001 |
| FAM83H | S1009 | -0.66 | -0.05 | -0.37 | 0.38 | 0.33 | 0.10 | 0.034 |
| FAM83H | S1098 | -0.50 | -0.44 | -0.09 | 0.30 | 0.36 | 0.15 | 0.012 |
| FAM83H | S892 | -0.36 | -0.35 | -0.30 | 0.03 | 0.19 | 0.55 | 0.019 |

| | | | | | | | | |
|---|---|---|---|---|---|---|---|---|
| **FAM83H** | **S936** | -0.42 | -0.16 | -0.31 | 0.31 | 0.29 | 0.11 | 0.005 |
| **FAM83H** | **S881** | 0.00 | -0.25 | -0.55 | 0.05 | 0.33 | 0.25 | 0.058 |
| **FAM83H** | **S998** | -0.19 | -0.27 | -0.19 | 0.14 | -0.05 | 0.42 | 0.049 |
| **FAM83H** | **S998** | -0.11 | -0.31 | -0.14 | 0.06 | 0.08 | 0.33 | 0.035 |
| **FAM83H** | **S411** | -0.50 | -0.06 | 0.12 | 0.15 | 0.17 | 0.03 | 0.239 |
| **FAM83H** | **S/T** | -0.22 | -0.13 | -0.09 | -0.23 | 0.44 | 0.11 | 0.274 |
| **FAM83H** | **S881** | -0.08 | -0.21 | -0.01 | 0.16 | 0.00 | 0.11 | 0.065 |
| **FAM83H** | **S925** | -0.12 | -0.11 | -0.08 | 0.09 | -0.05 | 0.22 | 0.073 |
| **FAM83H** | **S903** | -0.12 | -0.10 | -0.08 | 0.07 | 0.04 | 0.16 | 0.006 |
| **FAM83H** | **S892** | -0.12 | -0.15 | 0.00 | 0.25 | -0.01 | 0.00 | 0.15 |
| **FAM83H** | **S647** | -0.09 | -0.12 | -0.04 | 0.09 | -0.05 | 0.18 | 0.095 |
| **FAM83H** | **S925** | -0.12 | 0.03 | -0.05 | 0.12 | 0.00 | 0.00 | 0.212 |
| **FAM83H** | **S914** | -0.05 | -0.04 | -0.04 | -0.05 | -0.03 | 0.20 | 0.361 |
| **FAM83H** | **S1003** | -0.07 | -0.03 | -0.04 | 0.01 | 0.03 | 0.08 | 0.034 |
| **FAM83H** | **S411** | 0.04 | -0.21 | 0.04 | 0.01 | 0.12 | -0.02 | 0.434 |
| **FAM83H** | **S1003** | -0.27 | 0.02 | 0.10 | -0.25 | 0.08 | 0.24 | 0.709 |
| **FAM83H** | **S936; S945** | 0.05 | -0.05 | -0.09 | 0.19 | -0.27 | 0.12 | 0.77 |
| **FAM83H** | **S998** | 0.00 | -0.09 | 0.00 | -0.18 | -0.18 | 0.37 | 0.878 |
| **FAM83H** | **S998** | 0.06 | 0.04 | -0.15 | 0.03 | -0.11 | 0.11 | 0.782 |
| **FAM83H** | **S914** | -0.08 | -0.05 | 0.10 | 0.07 | 0.07 | -0.12 | 0.841 |
| **FAM83H** | **S523** | -0.01 | -0.03 | 0.06 | -0.02 | 0.01 | -0.02 | 0.512 |
| **FAM83H** | **S998** | -0.05 | 0.05 | 0.03 | 0.07 | -0.07 | -0.03 | 0.655 |
| **FAM83H** | **S998** | -0.05 | -0.08 | 0.15 | -0.12 | -0.10 | 0.17 | 0.825 |
| **FAM83H** | **S516** | 0.01 | 0.00 | 0.03 | -0.05 | 0.19 | -0.22 | 0.765 |
| **FAM83H** | **S647** | 0.06 | 0.04 | -0.04 | -0.03 | 0.08 | -0.12 | 0.548 |
| **FAM83H** | **S998** | 0.02 | 0.04 | 0.00 | 0.01 | -0.20 | 0.10 | 0.606 |
| **FAM83H** | **S936** | 0.09 | 0.07 | -0.06 | 0.01 | -0.04 | -0.08 | 0.254 |
| **FAM83H** | **S998** | -0.04 | 0.03 | 0.22 | -0.21 | -0.22 | 0.16 | 0.345 |
| **FAM83H** | **S998** | -0.19 | -0.11 | 0.72 | -0.25 | -0.36 | -0.10 | 0.281 |
| **FAM83H** | **T/S** | 0.27 | 0.19 | 0.21 | -0.02 | -0.16 | -0.70 | 0.068 |
| **FAM8A1** | **S229** | 0.26 | -0.04 | 0.25 | -0.19 | 0.14 | -0.58 | 0.188 |
| **FAM91A1** | **S671** | 0.03 | -0.14 | -0.05 | 0.20 | -0.06 | -0.01 | 0.355 |
| **FAN1** | **S197** | 0.22 | -0.19 | -0.23 | -0.13 | 0.03 | 0.23 | 0.594 |
| **FAN1** | **S197** | -0.09 | 0.10 | -0.15 | 0.09 | 0.08 | -0.05 | 0.389 |
| **FANCB** | **T555** | 0.27 | 0.25 | 0.11 | 0.10 | -0.11 | -0.92 | 0.175 |
| **FANCD2** | **S881** | -0.40 | -0.49 | -0.35 | 0.43 | -0.07 | 0.53 | 0.021 |
| **FANCD2** | **S/T** | -0.21 | -0.48 | 0.03 | 0.12 | 0.18 | 0.24 | 0.056 |
| **FANCE** | **S180** | -0.18 | -0.67 | 0.11 | 0.10 | 0.28 | 0.18 | 0.136 |
| **FANCI** | **S/T** | -0.02 | -0.47 | -0.39 | 0.24 | 0.11 | 0.35 | 0.027 |
| **FANCI** | **S/T** | -0.02 | -0.12 | -0.42 | 0.01 | 0.33 | 0.11 | 0.093 |
| **FANCI** | **T/S** | 0.20 | 0.19 | 0.27 | -0.44 | -0.07 | -0.28 | 0.012 |
| **FANCM** | **S1437** | 0.01 | -0.07 | -0.21 | 0.17 | -0.03 | 0.10 | 0.122 |
| **FANCM** | **S34** | 0.35 | 0.18 | 0.24 | -0.38 | -0.06 | -0.55 | 0.018 |
| **FANCM** | **S34** | 0.27 | 0.44 | 0.32 | -0.53 | -0.15 | -0.76 | 0.011 |
| **FARP1** | **S** | -0.09 | -0.06 | 0.35 | 0.24 | -0.94 | 0.18 | 0.59 |
| **FARP1** | **S/T** | -0.01 | 0.10 | 0.38 | -0.32 | -0.12 | -0.14 | 0.056 |
| **FARP1** | **S427** | -0.03 | 0.09 | 0.60 | -0.38 | -0.28 | -0.24 | 0.056 |
| **FARP1** | **S433** | 0.25 | 0.27 | 0.20 | -0.46 | -0.11 | -0.32 | 0.007 |

| | | | | | | | | |
|---|---|---|---|---|---|---|---|---|
| FARP2 | S428 | -0.03 | 0.03 | 0.06 | -0.01 | 0.06 | -0.12 | 0.511 |
| FARSA | T/S/Y | -0.24 | 0.08 | -0.01 | 0.18 | -0.37 | 0.25 | 0.748 |
| FARSA | S193 | 0.18 | -0.34 | 0.16 | 0.25 | -0.43 | 0.05 | 0.871 |
| FASN | S831 | -0.34 | -0.33 | -0.27 | 0.18 | 0.13 | 0.44 | 0.004 |
| FASN | S1411 | -0.38 | -0.27 | -0.21 | 0.24 | 0.29 | 0.19 | 8E-04 |
| FASTK | T/S | 0.41 | -0.02 | 0.37 | -1.28 | 0.19 | -0.25 | 0.203 |
| FBL | S124 | 0.00 | 0.05 | -0.09 | 0.01 | -0.17 | 0.18 | 0.872 |
| FBL | S124 | 0.18 | 0.04 | 0.05 | 0.01 | -0.13 | -0.18 | 0.06 |
| FBL | S126 | 0.20 | 0.41 | 0.16 | -0.30 | -0.08 | -0.62 | 0.027 |
| FBRSL1 | S844 | -0.26 | -0.15 | -0.44 | 0.22 | 0.27 | 0.21 | 0.004 |
| FBRSL1 | S31; S41; S | -0.02 | -0.04 | 0.31 | -0.66 | 0.05 | 0.18 | 0.468 |
| FBXL20 | T417 | 0.09 | 0.22 | 0.15 | -0.25 | -0.16 | -0.12 | 0.004 |
| FBXO28 | S344 | -0.37 | 0.01 | -0.08 | 0.10 | -0.21 | 0.41 | 0.308 |
| FBXO28 | S344 | -0.07 | -0.20 | 0.17 | -0.02 | -0.21 | 0.27 | 0.805 |
| FBXO28 | S344 | -0.13 | -0.06 | 0.21 | -0.13 | -0.34 | 0.34 | 0.842 |
| FBXO28 | S344 | -0.02 | 0.00 | 0.09 | -0.13 | 0.01 | 0.04 | 0.456 |
| FBXO38 | S654 | -0.07 | 0.09 | 0.00 | 0.13 | -0.07 | -0.10 | 0.849 |
| FBXO38 | S654 | 0.10 | -0.24 | 0.21 | 0.06 | -0.01 | -0.16 | 0.718 |
| FBXO4 | S12 | -0.22 | 0.09 | 0.26 | -0.09 | -0.11 | 0.02 | 0.527 |
| FBXW11 | S65 | -0.05 | 0.41 | -0.03 | -0.08 | -0.08 | -0.27 | 0.195 |
| FBXW11 | S65 | 0.44 | 0.14 | 0.36 | -0.86 | -0.37 | -0.11 | 0.032 |
| FBXW5 | T/S | -0.14 | -0.10 | 0.98 | -0.45 | -0.20 | -0.78 | 0.148 |
| FCHO2 | S403 | 0.23 | 0.06 | 0.35 | -0.28 | -0.04 | -0.49 | 0.037 |
| FDXR | S317 | -0.09 | -0.05 | -0.14 | 0.16 | -0.09 | 0.19 | 0.125 |
| FGD1 | S365 | 0.45 | -0.19 | 0.07 | -0.10 | -0.05 | -0.31 | 0.261 |
| FGF13 | S208 | -0.20 | -0.01 | -0.03 | 0.00 | 0.06 | 0.15 | 0.113 |
| FGF13 | T222 | 0.11 | 0.27 | 0.33 | -0.25 | -0.09 | -0.55 | 0.023 |
| FGFR4 | Y642 | -0.11 | -0.08 | -0.11 | 0.14 | -0.10 | 0.23 | 0.132 |
| FGFR4 | S573 | -0.02 | 0.14 | 0.15 | -0.11 | -0.14 | -0.05 | 0.038 |
| FGFR4 | Y642 | 0.09 | 0.28 | -0.04 | -0.08 | 0.00 | -0.32 | 0.137 |
| FGFR4 | S419 | 0.33 | 0.21 | 0.25 | -0.19 | -0.19 | -0.62 | 0.016 |
| FGFR4 | S419 | 0.30 | 0.36 | 0.28 | -0.24 | -0.28 | -0.71 | 0.009 |
| FHOD1 | S486 | 0.05 | 0.05 | 0.20 | -0.06 | -0.07 | -0.20 | 0.036 |
| FHOD1 | S523 | 0.30 | 0.09 | 0.05 | -0.06 | -0.27 | -0.18 | 0.032 |
| FHOD1 | S367 | 0.24 | 0.22 | -0.01 | -0.24 | -0.07 | -0.22 | 0.027 |
| FILIP1L | T/S | 0.13 | -0.01 | 0.13 | -0.19 | 0.03 | -0.12 | 0.094 |
| FIP1L1 | S/T | -0.11 | -0.04 | -0.22 | 0.28 | -0.04 | 0.08 | 0.093 |
| FIP1L1 | S492 | -0.19 | 0.00 | -0.08 | 0.14 | -0.06 | 0.16 | 0.136 |
| FIP1L1 | S492 | -0.14 | -0.04 | -0.06 | 0.14 | -0.16 | 0.21 | 0.289 |
| FIP1L1 | S492 | -0.09 | -0.08 | -0.05 | 0.17 | -0.14 | 0.16 | 0.248 |
| FIP1L1 | S/T | -0.03 | 0.02 | -0.12 | 0.21 | -0.21 | 0.10 | 0.584 |
| FIP1L1 | S500; S/T | 0.00 | -0.05 | -0.05 | 0.02 | -0.05 | 0.13 | 0.288 |
| FIP1L1 | S85; S87; S89 | -0.05 | -0.07 | 0.01 | 0.10 | -0.06 | 0.06 | 0.287 |
| FIP1L1 | S492; S500 | 0.05 | 0.07 | 0.02 | 0.07 | -0.08 | -0.15 | 0.21 |
| FIP1L1 | S87; S89 | 0.24 | 0.42 | 0.28 | -0.24 | -0.27 | -0.77 | 0.015 |
| FIP1L1 | S89 | 0.27 | 0.64 | 0.45 | -0.59 | -0.36 | -1.23 | 0.014 |
| FKBP15 | S956 | -0.21 | -0.12 | -0.15 | -0.33 | 0.60 | 0.00 | 0.413 |
| FKBP15 | S956 | -0.02 | -0.37 | 0.24 | 0.19 | -0.05 | -0.06 | 0.725 |

| | | | | | | | | |
|---|---|---|---|---|---|---|---|---|
| **FKBP3** | **S36** | 0.33 | -0.34 | 0.31 | 0.28 | -0.13 | -0.77 | 0.463 |
| **FKBP5** | **S445** | -0.51 | -0.39 | -0.32 | 0.51 | 0.13 | 0.28 | 0.004 |
| **FKBP8** | **S296** | 0.14 | 0.07 | 0.19 | -0.15 | 0.00 | -0.31 | 0.042 |
| **FLCN** | **S62** | 0.03 | -0.06 | -0.14 | 0.08 | -0.01 | 0.08 | 0.146 |
| **FLII** | **S436** | 0.18 | 0.12 | -0.06 | -0.11 | -0.07 | -0.09 | 0.088 |
| **FLII** | **S406** | -0.08 | 0.55 | -0.18 | 0.18 | -0.21 | -0.49 | 0.419 |
| **FLII** | **S856** | 0.21 | 0.17 | 0.31 | -0.32 | 0.02 | -0.59 | 0.045 |
| **FLII** | **S436** | 0.26 | 0.24 | 0.21 | -0.32 | -0.05 | -0.52 | 0.018 |
| **FLII** | **S436** | 0.40 | 0.17 | 0.39 | -0.56 | -0.13 | -0.62 | 0.012 |
| **FLNA** | **S1533** | -0.10 | 0.07 | -0.50 | 0.32 | 0.11 | -0.03 | 0.187 |
| **FLNA** | **T2599** | -0.03 | 0.11 | -0.21 | 0.06 | 0.11 | -0.07 | 0.5 |
| **FLNA** | **S2152** | -0.16 | 0.23 | -0.08 | -0.12 | -0.02 | 0.10 | 0.947 |
| **FLNA** | **S2327** | 0.07 | 0.05 | 0.05 | -0.05 | -0.34 | 0.16 | 0.414 |
| **FLNA** | **S1630** | 0.03 | 0.03 | 0.23 | -0.20 | 0.10 | -0.24 | 0.171 |
| **FLNA** | **S2152** | 0.32 | 0.30 | -0.01 | 0.05 | -0.20 | -0.69 | 0.114 |
| **FLNA** | **S1459** | 0.16 | 0.20 | 0.33 | -0.22 | -0.03 | -0.66 | 0.052 |
| **FLNA** | **S1459** | 0.28 | 0.30 | 0.24 | -0.25 | -0.06 | -0.81 | 0.045 |
| **FLNA** | **S1084** | 0.06 | 0.38 | 0.52 | -0.60 | -0.30 | -0.42 | 0.009 |
| **FLNB** | **T/S** | -0.37 | -0.70 | -0.49 | 0.16 | 0.63 | 0.31 | 0.006 |
| **FLNB** | **S2478** | -0.08 | -0.35 | -0.31 | -0.29 | 0.41 | 0.40 | 0.165 |
| **FLNB** | **S1505** | -0.12 | 0.20 | 0.03 | -0.14 | 0.14 | -0.15 | 0.54 |
| **FLOT1** | **Y203** | -0.08 | 0.07 | -0.04 | -0.10 | 0.00 | 0.13 | 0.723 |
| **FLOT1** | **S385** | -0.06 | -0.03 | 0.05 | -0.20 | 0.14 | 0.07 | 0.868 |
| **FLOT1** | **Y203** | 0.02 | -0.06 | 0.10 | 0.01 | 0.18 | -0.30 | 0.726 |
| **FLOT1** | **S/T** | 0.01 | 0.01 | 0.12 | 0.01 | 0.18 | -0.38 | 0.549 |
| **FLOT2** | **S385** | -0.14 | 0.25 | -0.84 | -0.04 | 0.04 | 0.43 | 0.333 |
| **FLVCR1** | **S536** | -0.34 | -0.13 | -0.36 | 0.21 | 0.13 | 0.34 | 0.006 |
| **FLVCR1** | **S536** | -0.16 | -0.12 | -0.28 | 0.19 | 0.04 | 0.26 | 0.011 |
| **FLVCR1** | **S56** | -0.57 | 0.23 | 0.23 | -0.02 | 0.19 | -0.23 | 0.957 |
| **FLYWCH1** | **S503** | -0.01 | -0.18 | -0.15 | 0.08 | 0.04 | 0.19 | 0.031 |
| **FLYWCH1** | **S21** | -0.10 | -0.04 | -0.15 | 0.06 | 0.10 | 0.12 | 0.007 |
| **FLYWCH1** | **S696** | -0.05 | -0.06 | 0.05 | -0.02 | 0.06 | 0.01 | 0.466 |
| **FLYWCH2** | **S21** | 0.25 | 0.32 | -0.08 | 0.03 | -0.46 | -0.19 | 0.118 |
| **FMNL2** | **S171** | 0.08 | -0.28 | 0.05 | 0.18 | -0.09 | 0.02 | 0.551 |
| **FMR1** | **S620** | -0.37 | 0.07 | -0.23 | 0.14 | 0.07 | 0.23 | 0.077 |
| **FMR1** | **S620** | 0.01 | 0.00 | -0.19 | 0.09 | -0.12 | 0.18 | 0.385 |
| **FMR1** | **T492; S/T/Y** | -0.08 | -0.17 | 0.07 | 0.06 | -0.12 | 0.20 | 0.414 |
| **FMR1** | **S514; T518** | -0.01 | -0.21 | 0.16 | 0.22 | -0.12 | -0.09 | 0.884 |
| **FN1** | **S2475** | 0.01 | -0.59 | -0.22 | 0.09 | 0.04 | 0.45 | 0.101 |
| **FNBP1** | **S296** | 0.23 | 0.25 | 0.26 | -0.25 | -0.05 | -0.67 | 0.035 |
| **FNBP1L** | **S295** | 0.20 | 0.23 | 0.08 | -0.05 | -0.04 | -0.55 | 0.091 |
| **FNBP1L** | **S501** | 0.26 | 0.15 | 0.42 | -0.32 | -0.05 | -0.76 | 0.042 |
| **FNBP4** | **T** | 0.10 | -0.14 | 0.09 | 0.05 | -0.11 | -0.02 | 0.691 |
| **FNBP4** | **S432** | 0.05 | 0.06 | -0.01 | 0.02 | -0.10 | -0.02 | 0.159 |
| **FNBP4** | **S499; S508** | 0.09 | 0.18 | -0.03 | 0.02 | -0.10 | -0.19 | 0.118 |
| **FNBP4** | **T** | 0.13 | 0.06 | 0.07 | 0.00 | -0.02 | -0.27 | 0.113 |
| **FNBP4** | **S435; S** | 0.07 | 0.14 | 0.09 | 0.00 | -0.11 | -0.22 | 0.037 |
| **FNBP4** | **S18** | 0.14 | 0.11 | 0.14 | -0.03 | -0.09 | -0.32 | 0.035 |

| | | | | | | | | |
|---|---|---|---|---|---|---|---|---|
| FNBP4 | S464 | 0.05 | 0.44 | -0.12 | -0.26 | -0.67 | 0.28 | 0.352 |
| FNBP4 | S499 | 0.12 | 0.20 | 0.17 | -0.17 | -0.05 | -0.36 | 0.018 |
| FNBP4 | T479 | 0.18 | 0.12 | 0.29 | -0.34 | 0.00 | -0.40 | 0.029 |
| FNBP4 | S499; S508 | 0.31 | 0.20 | 0.18 | -0.29 | -0.03 | -0.57 | 0.031 |
| FNBP4 | T479 | 0.21 | 0.25 | 0.37 | -0.25 | -0.20 | -0.61 | 0.01 |
| FNDC3A | S203 | -0.12 | 0.01 | 0.16 | 0.08 | -0.06 | -0.09 | 0.693 |
| FNDC3A | S213 | 0.05 | 0.14 | 0.05 | 0.05 | -0.17 | -0.16 | 0.09 |
| FNDC3A | S213 | -0.02 | 0.03 | 0.25 | 0.18 | -0.26 | -0.27 | 0.294 |
| FNDC3A | S203 | 0.14 | 0.15 | 0.02 | 0.00 | 0.02 | -0.39 | 0.174 |
| FNDC3B | S208 | -0.01 | 0.10 | -0.49 | 0.39 | 0.36 | -0.68 | 0.712 |
| FOS | S/T | 0.03 | -0.39 | 0.02 | 0.14 | -0.03 | 0.16 | 0.242 |
| FOSL2 | S16; S19 | -0.31 | -0.25 | -0.15 | 0.38 | -0.33 | 0.44 | 0.183 |
| FOSL2 | S233; S234 | -0.18 | -0.03 | -0.01 | 0.01 | 0.04 | 0.15 | 0.114 |
| FOSL2 | S200 | -0.25 | 0.11 | -0.08 | 0.21 | -0.06 | 0.02 | 0.374 |
| FOSL2 | S200 | -0.14 | -0.02 | -0.02 | -0.06 | 0.33 | -0.15 | 0.56 |
| FOSL2 | S200 | 0.03 | -0.07 | -0.12 | 0.19 | 0.20 | -0.29 | 0.617 |
| FOSL2 | S230 | -0.14 | -0.11 | 0.21 | -0.14 | -0.01 | 0.15 | 0.93 |
| FOSL2 | S19 | -0.21 | -0.46 | 0.58 | 0.07 | -0.47 | 0.19 | 0.92 |
| FOSL2 | S308 | -0.01 | -0.04 | 0.25 | -0.25 | -0.08 | 0.08 | 0.312 |
| FOXA1 | S285 | -0.02 | -0.21 | -0.21 | 0.20 | 0.02 | 0.17 | 0.031 |
| FOXA1 | S234 | -0.01 | -0.14 | -0.30 | 0.20 | -0.03 | 0.21 | 0.076 |
| FOXA1 | S331 | 0.03 | 0.06 | -0.02 | -0.11 | -0.01 | 0.05 | 0.42 |
| FOXA1 | S307 | 0.02 | 0.01 | 0.19 | -0.07 | -0.15 | -0.03 | 0.087 |
| FOXA1 | S307 | 0.11 | -0.02 | 0.16 | -0.29 | 0.13 | -0.15 | 0.248 |
| FOXA1 | S307 | 0.13 | 0.05 | 0.10 | -0.07 | -0.12 | -0.11 | 0.002 |
| FOXA1 | T334; S338 | 0.22 | 0.07 | 0.00 | 0.02 | -0.21 | -0.14 | 0.089 |
| FOXA1 | S307 | -0.08 | 0.17 | 0.24 | -0.26 | -0.18 | 0.04 | 0.141 |
| FOXA1 | S331 | 0.13 | 0.14 | 0.14 | -0.25 | 0.02 | -0.24 | 0.032 |
| FOXJ2 | S164; S172 | -0.15 | 0.04 | -0.02 | 0.06 | -0.13 | 0.17 | 0.512 |
| FOXJ2 | S46 | -0.18 | -0.02 | 0.08 | -0.12 | 0.24 | -0.04 | 0.638 |
| FOXJ3 | S223 | -0.05 | -0.10 | -0.01 | 0.22 | -0.02 | -0.07 | 0.35 |
| FOXJ3 | S223 | -0.25 | 0.37 | -0.31 | 0.17 | -0.40 | 0.23 | 0.84 |
| FOXK1 | S420 | -0.43 | -0.48 | -0.42 | 0.50 | -0.46 | 0.72 | 0.127 |
| FOXK1 | S420; S428 | -0.01 | -0.14 | -0.09 | 0.14 | 0.05 | 0.03 | 0.043 |
| FOXK1 | S257; S/T | -0.01 | -0.15 | -0.01 | -0.16 | 0.03 | 0.26 | 0.475 |
| FOXK1 | S420; S428 | -0.08 | -0.02 | -0.04 | -0.07 | 0.09 | 0.12 | 0.203 |
| FOXK1 | S243; S257 | -0.04 | 0.02 | -0.08 | 0.09 | 0.04 | -0.05 | 0.293 |
| FOXK1 | S459 | -0.49 | 0.38 | 0.08 | 0.12 | 0.12 | -0.39 | 0.894 |
| FOXK1 | S213 | 0.12 | 0.08 | -0.12 | 0.01 | -0.03 | -0.07 | 0.524 |
| FOXK1 | S420 | 0.00 | -0.01 | 0.10 | -0.05 | -0.15 | 0.09 | 0.47 |
| FOXK1 | S445 | 0.03 | 0.02 | 0.12 | 0.00 | -0.01 | -0.17 | 0.146 |
| FOXK1 | S213; S223 | 0.11 | 0.02 | 0.05 | 0.06 | -0.18 | -0.09 | 0.159 |
| FOXK1 | S101 | 0.14 | 0.05 | 0.06 | -0.16 | -0.07 | -0.03 | 0.023 |
| FOXK1 | S441 | 0.09 | 0.10 | 0.12 | -0.31 | 0.13 | -0.20 | 0.157 |
| FOXK2 | S239 | -0.07 | -0.43 | -0.23 | 0.23 | 0.28 | 0.10 | 0.02 |
| FOXK2 | S398 | -0.06 | 0.08 | 0.07 | -0.14 | -0.09 | 0.13 | 0.554 |
| FOXK2 | S398 | -0.01 | 0.02 | 0.12 | -0.09 | -0.02 | -0.02 | 0.13 |
| FOXK2 | S170 | 0.18 | 0.02 | -0.04 | -0.02 | 0.01 | -0.17 | 0.255 |

| | | | | | | | | |
|---|---|---|---|---|---|---|---|---|
| FOXK2 | S170 | 0.10 | 0.02 | 0.09 | -0.04 | -0.09 | -0.10 | 0.011 |
| FOXK2 | S428 | -0.02 | 0.31 | 0.67 | -0.05 | -0.80 | -0.63 | 0.055 |
| FOXN3 | T/S/Y | 0.41 | 0.34 | 0.11 | -0.61 | -0.40 | -0.12 | 0.016 |
| FOXO3 | S284 | -0.04 | -0.08 | -0.26 | 0.22 | -0.03 | 0.15 | 0.077 |
| FOXO3 | S/T | -0.04 | -0.17 | -0.01 | 0.13 | -0.26 | 0.28 | 0.493 |
| FOXP1 | S/T/Y | -0.18 | -0.09 | -0.18 | 0.15 | -0.06 | 0.28 | 0.056 |
| FOXP1 | T/S/Y | -0.01 | 0.00 | -0.02 | 0.05 | -0.08 | 0.05 | 0.715 |
| FRA10AC1 | S278; S283 | 0.17 | -0.16 | 0.16 | -0.23 | -0.15 | 0.14 | 0.426 |
| FRA10AC1 | S278 | 0.06 | 0.19 | 0.00 | 0.00 | -0.10 | -0.18 | 0.081 |
| FRA10AC1 | S278 | 0.10 | 0.24 | -0.01 | -0.16 | -0.02 | -0.20 | 0.064 |
| FRA10AC1 | S278 | 0.15 | 0.21 | -0.02 | -0.12 | -0.04 | -0.22 | 0.053 |
| FRMD4A | S711 | 0.01 | 0.08 | 0.08 | -0.08 | -0.01 | -0.08 | 0.027 |
| FRS2 | S428 | -0.10 | -0.05 | 0.11 | 0.10 | 0.02 | -0.10 | 0.795 |
| FRS2 | T227 | 0.18 | 0.23 | 0.39 | -0.45 | 0.04 | -0.68 | 0.048 |
| FRS2 | S/T | 0.38 | 0.27 | 0.12 | -0.53 | 0.20 | -0.79 | 0.107 |
| FTSJ3 | S/T | -0.22 | -0.43 | -0.40 | 0.07 | 0.19 | 0.53 | 0.016 |
| FTSJ3 | S688 | -0.26 | -0.32 | -0.20 | 0.31 | 0.04 | 0.29 | 0.008 |
| FTSJ3 | S333; S335; S336 | -0.12 | -0.17 | -0.13 | 0.24 | -0.09 | 0.21 | 0.069 |
| FTSJ3 | S644 | -0.01 | -0.06 | -0.16 | 0.08 | -0.05 | 0.17 | 0.147 |
| FTSJ3 | S599 | -0.02 | -0.11 | -0.07 | 0.10 | -0.01 | 0.10 | 0.042 |
| FTSJ3 | S356 | -0.10 | -0.10 | 0.01 | 0.15 | 0.00 | 0.02 | 0.101 |
| FTSJ3 | S336; S/T | -0.02 | -0.04 | -0.06 | 0.08 | -0.13 | 0.15 | 0.418 |
| FTSJ3 | T573 | -0.08 | -0.01 | 0.02 | -0.22 | 0.27 | -0.02 | 0.837 |
| FTSJ3 | T467 | -0.04 | 0.05 | -0.04 | 0.00 | -0.12 | 0.15 | 0.82 |
| FTSJ3 | S335 | 0.07 | 0.07 | -0.05 | 0.03 | -0.10 | -0.03 | 0.34 |
| FTSJ3 | S356 | 0.06 | 0.03 | 0.01 | 0.07 | -0.06 | -0.12 | 0.289 |
| FTSJ3 | S356 | 0.06 | 0.03 | 0.01 | 0.07 | -0.06 | -0.12 | 0.289 |
| FTSJ3 | S458; S471; T/S/Y | -0.01 | 0.13 | 0.04 | -0.01 | -0.06 | -0.09 | 0.101 |
| FTSJ3 | S584 | -0.08 | 0.26 | 0.11 | -0.13 | 0.24 | -0.55 | 0.377 |
| FTSJ3 | S584 | 0.07 | 0.19 | 0.22 | -0.21 | 0.05 | -0.42 | 0.068 |
| FTSJ3 | S471; S/T/Y | 0.00 | 0.30 | 0.18 | -0.15 | 0.07 | -0.54 | 0.14 |
| FTSJ3 | S584 | 0.15 | 0.17 | 0.18 | -0.03 | -0.02 | -0.58 | 0.11 |
| FUBP3 | T76 | -0.28 | -0.41 | -0.31 | 0.29 | -0.04 | 0.51 | 0.023 |
| FUBP3 | T76 | -0.02 | -0.28 | -0.48 | 0.36 | -0.16 | 0.37 | 0.113 |
| FUBP3 | S296 | -0.01 | 0.05 | 0.14 | -0.03 | 0.15 | -0.35 | 0.42 |
| FUT8 | S278 | 0.14 | 0.17 | 0.27 | -0.19 | -0.24 | -0.25 | 6E-04 |
| FXR1 | S409 | -0.06 | 0.17 | -0.51 | 0.09 | -0.30 | 0.43 | 0.522 |
| FXR1 | S587 | -0.17 | 0.05 | -0.05 | 0.20 | 0.06 | -0.12 | 0.422 |
| FXR1 | T/S | -0.09 | 0.05 | -0.08 | 0.16 | 0.08 | -0.14 | 0.517 |
| FXR1 | S406; S409 | 0.00 | 0.11 | -0.08 | -0.12 | -0.02 | 0.10 | 0.775 |
| FXR1 | S409 | 0.26 | 0.05 | -0.16 | 0.09 | -0.06 | -0.22 | 0.492 |
| FXR1 | S/T/Y | 0.18 | 0.24 | -0.14 | 0.01 | -0.31 | -0.06 | 0.225 |
| FXR1 | S/T/Y | 0.15 | 0.12 | 0.11 | -0.22 | -0.14 | -0.06 | 0.005 |
| FXR1 | S406; S409 | 0.00 | 0.24 | 0.18 | -0.22 | 0.02 | -0.31 | 0.064 |
| FXR1 | S409 | 0.29 | 0.11 | 0.08 | -0.16 | -0.17 | -0.21 | 0.007 |
| FXR1 | S406; S409 | 0.16 | 0.05 | 0.26 | -0.29 | -0.23 | -0.03 | 0.025 |
| FXR1 | S409 | 0.01 | 0.07 | 0.49 | -0.40 | 0.24 | -0.74 | 0.204 |
| FXR2 | S/T/Y | -0.31 | -0.32 | -0.25 | 0.39 | -0.34 | 0.54 | 0.144 |

| | | | | | | | | |
|---|---|---|---|---|---|---|---|---|
| FXR2 | S566 | -0.18 | -0.16 | -0.19 | 0.09 | -0.09 | 0.42 | 0.099 |
| FXR2 | S/T | -0.22 | 0.00 | -0.20 | 0.14 | -0.06 | 0.27 | 0.098 |
| FXR2 | S453 | 0.01 | 0.05 | 0.11 | 0.05 | 0.24 | -0.61 | 0.56 |
| FXR2 | S453 | 0.07 | 0.11 | 0.08 | -0.07 | 0.15 | -0.42 | 0.285 |
| FXR2 | S603 | 0.05 | 0.31 | 0.35 | -0.82 | 0.22 | -0.47 | 0.138 |
| FXYD3 | T76 | -0.10 | -0.07 | 0.03 | 0.15 | -0.19 | 0.15 | 0.506 |
| FYTTD1 | S23 | -0.06 | -0.16 | -0.12 | 0.11 | 0.04 | 0.17 | 0.011 |
| FYTTD1 | S16 | 0.02 | -0.09 | -0.01 | 0.02 | 0.19 | -0.16 | 0.707 |
| FYTTD1 | S118 | 0.16 | -0.05 | -0.20 | 0.29 | -0.08 | -0.19 | 0.851 |
| FZR1 | S151 | 0.03 | -0.64 | 0.16 | 0.36 | -0.18 | 0.07 | 0.468 |
| FZR1 | S138 | 0.01 | 0.03 | 0.14 | -0.04 | -0.11 | -0.04 | 0.065 |
| FZR1 | S70 | 0.21 | 0.13 | 0.07 | -0.17 | -0.22 | -0.06 | 0.009 |
| G3BP1 | S149 | -0.09 | 0.01 | 0.01 | 0.14 | -0.06 | -0.03 | 0.654 |
| G3BP1 | S149 | -0.02 | -0.01 | 0.04 | 0.10 | -0.06 | -0.06 | 0.906 |
| G3BP1 | S232 | -0.02 | 0.11 | 0.07 | 0.07 | -0.07 | -0.17 | 0.246 |
| G3BP1 | S232 | 0.16 | 0.01 | 0.01 | -0.02 | 0.02 | -0.20 | 0.217 |
| G3BP2 | S141; S149 | -0.32 | -0.08 | 0.13 | 0.07 | 0.24 | -0.11 | 0.403 |
| G3BP2 | S/T | 0.00 | -0.04 | 0.03 | 0.09 | 0.00 | -0.09 | 0.961 |
| GAK | S73 | 0.03 | -0.16 | -0.04 | 0.05 | 0.11 | 0.01 | 0.139 |
| GALNT10 | S481 | -0.31 | 0.19 | 0.38 | -0.30 | -0.21 | 0.11 | 0.413 |
| GALNT2 | S94 | 0.10 | 0.19 | 0.12 | -0.23 | -0.24 | 0.00 | 0.025 |
| GALNT7 | S103 | -0.03 | 0.06 | 0.54 | 0.04 | -0.41 | -0.41 | 0.121 |
| GAREM1 | S/Y | -0.27 | -0.20 | 0.26 | 0.13 | 0.10 | -0.09 | 0.546 |
| GAS2L1 | S620 | -0.24 | -0.36 | -0.26 | 0.01 | -0.02 | 0.63 | 0.085 |
| GAS2L1 | S620 | -0.07 | -0.30 | 0.04 | -0.11 | -0.01 | 0.37 | 0.33 |
| GAS2L1 | S479 | -0.16 | -0.11 | 0.05 | 0.21 | -0.23 | 0.19 | 0.458 |
| GAS2L1 | T334 | -0.15 | 0.00 | 0.11 | -0.09 | -0.19 | 0.27 | 0.943 |
| GAS2L1 | S352 | -0.01 | -0.09 | 0.29 | 0.02 | -0.25 | -0.02 | 0.377 |
| GATA2 | T188; S195; S196 | 0.01 | -0.31 | -0.01 | 0.32 | -0.06 | -0.02 | 0.316 |
| GATA2 | S73 | 0.07 | -0.14 | -0.09 | -0.17 | 0.13 | 0.16 | 0.489 |
| GATA2 | S192 | -0.21 | -0.03 | 0.11 | 0.13 | -0.27 | 0.21 | 0.728 |
| GATA3 | S381 | -0.40 | -0.61 | -0.46 | 0.17 | 0.15 | 0.71 | 0.013 |
| GATA3 | S381 | -0.52 | -0.60 | -0.27 | 0.26 | 0.45 | 0.32 | 0.002 |
| GATA3 | S162 | 0.05 | -0.03 | -0.04 | 0.00 | -0.13 | 0.13 | 0.943 |
| GATA6 | T/S | -0.39 | 0.16 | 0.62 | -0.62 | -0.01 | -0.09 | 0.35 |
| GATAD2A | S548 | 0.25 | -0.95 | -0.16 | 0.06 | 0.05 | 0.40 | 0.285 |
| GATAD2A | T327; S340 | 0.17 | 0.10 | -0.63 | -0.11 | 0.19 | 0.13 | 0.53 |
| GATAD2A | S100 | -0.20 | -0.02 | 0.01 | 0.00 | -0.04 | 0.22 | 0.268 |
| GATAD2A | S100; S107; S114 | -0.02 | -0.17 | 0.05 | 0.02 | -0.15 | 0.23 | 0.549 |
| GATAD2A | S100; S107 | -0.05 | 0.00 | 0.03 | -0.01 | -0.10 | 0.11 | 0.917 |
| GATAD2A | T49 | -0.05 | 0.09 | 0.07 | 0.00 | -0.21 | 0.08 | 0.431 |
| GATAD2A | S | 0.05 | 0.24 | 0.06 | -0.03 | -0.12 | -0.25 | 0.05 |
| GATAD2A | T188 | 0.36 | 0.06 | 0.19 | -0.16 | -0.18 | -0.41 | 0.018 |
| GATAD2B | T120; S122; S | -0.12 | -0.11 | -0.02 | 0.10 | -0.14 | 0.26 | 0.259 |
| GATAD2B | T120; S | -0.06 | -0.02 | -0.05 | -0.08 | -0.07 | 0.26 | 0.493 |
| GATAD2B | S122; S135 | -0.01 | -0.01 | -0.02 | -0.04 | -0.05 | 0.13 | 0.643 |
| GATAD2B | S135 | 0.08 | -0.17 | 0.07 | -0.06 | 0.16 | -0.10 | 0.951 |
| GATAD2B | S135 | 0.15 | -0.03 | -0.15 | -0.12 | -0.21 | 0.30 | 0.998 |

| | | | | | | | | |
|---|---|---|---|---|---|---|---|---|
| GATAD2B | T120 | 0.07 | 0.06 | 0.13 | -0.31 | -0.04 | 0.05 | 0.159 |
| GATAD2B | S486 | 0.32 | 0.37 | 0.07 | -0.32 | -0.12 | -0.52 | 0.019 |
| GBF1 | S1784 | -0.16 | -0.12 | 0.03 | 0.10 | 0.18 | -0.07 | 0.177 |
| GBF1 | S314 | -0.17 | 0.19 | -0.23 | 0.06 | 0.00 | 0.11 | 0.416 |
| GBF1 | S314 | -0.01 | 0.04 | -0.16 | 0.01 | -0.03 | 0.14 | 0.363 |
| GBF1 | S1298 | 0.02 | -0.02 | -0.04 | 0.15 | -0.05 | -0.08 | 0.802 |
| GBF1 | S/Y | 0.12 | -0.02 | -0.12 | -0.07 | -0.11 | 0.16 | 0.987 |
| GBF1 | S1773 | 0.11 | -0.16 | 0.05 | 0.15 | 0.13 | -0.33 | 0.921 |
| GBF1 | S1318 | 0.11 | -0.04 | -0.05 | 0.10 | -0.01 | -0.13 | 0.807 |
| GCC2 | S1483 | 0.03 | -0.15 | 0.22 | 0.15 | -0.01 | -0.29 | 0.628 |
| GEMIN4 | T84 | -0.29 | -0.09 | 0.02 | 0.07 | 0.01 | 0.24 | 0.121 |
| GFPT1 | S261 | 0.24 | 0.19 | -0.07 | -0.22 | -0.05 | -0.15 | 0.076 |
| GGA3 | S159 | 0.08 | 0.00 | 0.02 | -0.06 | 0.29 | -0.41 | 0.682 |
| GHDC | S87 | 0.02 | -0.16 | -0.15 | -0.09 | 0.21 | 0.12 | 0.178 |
| GIGYF2 | S236 | -0.59 | 0.11 | 0.23 | 0.41 | -0.20 | -0.18 | 0.791 |
| GIGYF2 | S593 | -0.23 | 0.30 | -0.01 | -0.01 | -0.15 | 0.04 | 0.734 |
| GIGYF2 | S26 | 0.15 | 0.10 | 0.10 | -0.08 | -0.06 | -0.25 | 0.017 |
| GINS2 | S182 | -0.20 | -0.13 | 0.01 | 0.13 | 0.12 | 0.04 | 0.041 |
| GIPC1 | S232 | 0.22 | 0.44 | 0.26 | -0.19 | -0.16 | -0.98 | 0.053 |
| GIT1 | S388 | -0.16 | -0.13 | -0.04 | 0.23 | -0.04 | 0.10 | 0.076 |
| GIT1 | S592 | 0.08 | -0.24 | -0.04 | -0.06 | 0.03 | 0.19 | 0.364 |
| GIT1 | S592 | 0.03 | 0.01 | 0.02 | -0.12 | -0.10 | 0.15 | 0.659 |
| GIT1 | S592 | 0.24 | -0.01 | -0.01 | -0.07 | -0.15 | -0.03 | 0.156 |
| GLE1 | S88; S96 | -0.02 | -0.12 | -0.08 | 0.01 | -0.18 | 0.33 | 0.457 |
| GLE1 | S88 | 0.05 | 0.06 | 0.06 | -0.15 | 0.01 | -0.04 | 0.068 |
| GLE1 | S88; T102 | 0.17 | 0.21 | 0.01 | -0.10 | -0.13 | -0.21 | 0.017 |
| GLE1 | S88 | 0.26 | 0.19 | -0.02 | -0.10 | -0.30 | -0.12 | 0.041 |
| GLI3 | S445 | 0.23 | 0.01 | 0.05 | -0.16 | -0.01 | -0.16 | 0.077 |
| GLI3 | S664 | 0.13 | 0.27 | 0.01 | -0.44 | -0.29 | 0.19 | 0.201 |
| GLI3 | S664 | 0.35 | 0.13 | 0.07 | -0.16 | -0.84 | 0.18 | 0.217 |
| GLYR1 | S130 | 0.22 | -0.17 | -0.14 | 0.08 | -0.15 | 0.11 | 0.779 |
| GLYR1 | S122 | 0.10 | 0.08 | 0.03 | 0.02 | -0.06 | -0.19 | 0.082 |
| GLYR1 | S122 | 0.15 | 0.20 | 0.10 | -0.28 | 0.33 | -0.78 | 0.285 |
| GMIP | S919 | -0.22 | -0.06 | 0.17 | 0.18 | -0.28 | 0.14 | 0.804 |
| GMIP | S234 | -0.12 | -0.03 | 0.21 | -0.16 | -0.16 | 0.21 | 0.725 |
| GMIP | S437 | 0.02 | 0.24 | -0.01 | 0.03 | -0.22 | -0.10 | 0.18 |
| GMIP | S437; T439 | 0.09 | 0.16 | 0.05 | -0.05 | -0.13 | -0.14 | 0.009 |
| GMPS | T/S | -0.28 | -0.06 | 0.44 | -0.49 | 0.61 | -0.67 | 0.655 |
| GNAS | S/T/Y | 0.08 | -0.39 | -0.06 | 0.09 | -0.02 | 0.22 | 0.229 |
| GNAS | S995 | -0.09 | 0.06 | 0.27 | -0.10 | 0.04 | -0.22 | 0.25 |
| GNAS | S581 | 0.08 | 0.28 | 0.21 | 0.12 | -0.53 | -0.33 | 0.095 |
| GNB1 | T/S | -0.54 | 0.10 | -0.03 | -0.17 | 0.39 | 0.09 | 0.371 |
| GNB2 | T31 | 0.06 | -0.11 | -0.15 | 0.02 | -0.12 | 0.26 | 0.4 |
| GNG12 | S7 | -0.02 | 0.07 | 0.24 | -0.26 | 0.05 | -0.13 | 0.151 |
| GNL1 | S324 | -0.36 | 0.05 | 0.13 | 0.15 | -0.18 | 0.13 | 0.634 |
| GNL2 | S580; S581; S582 | 0.16 | 0.21 | 0.08 | -0.32 | -0.18 | -0.01 | 0.031 |
| GNL3L | S465 | 0.21 | 0.19 | -0.15 | -0.16 | 0.15 | -0.34 | 0.345 |
| GOLGA1 | S30 | 0.21 | -0.14 | 0.06 | 0.11 | 0.08 | -0.41 | 0.592 |

| | | | | | | | | |
|---|---|---|---|---|---|---|---|---|
| GOLGA2 | S66 | -0.04 | -0.09 | -0.45 | 0.06 | 0.15 | 0.27 | 0.07 |
| GOLGA2 | S273 | -0.59 | -0.06 | 0.05 | 0.40 | -0.16 | 0.16 | 0.263 |
| GOLGA2 | S66 | -0.01 | 0.04 | -0.12 | 0.15 | -0.26 | 0.17 | 0.756 |
| GOLGA2 | S438 | -0.12 | 0.01 | 0.03 | 0.17 | 0.25 | -0.43 | 0.912 |
| GOLGA2 | S66 | 0.51 | 0.10 | -0.23 | -1.09 | 0.64 | -0.67 | 0.425 |
| GOLGA2 | S37 | 0.19 | 0.22 | 0.28 | -0.38 | 0.16 | -0.75 | 0.108 |
| GOLGA3 | S1392 | -0.45 | 0.05 | 0.04 | 0.36 | -0.12 | 0.00 | 0.417 |
| GOLGA3 | S1392 | -0.45 | 0.05 | 0.04 | 0.36 | -0.12 | 0.00 | 0.417 |
| GOLGA4 | S71 | -0.51 | -0.47 | -0.42 | 0.43 | 0.26 | 0.36 | 1E-04 |
| GOLGB1 | S3010 | -0.20 | -0.74 | -0.09 | 0.50 | 0.27 | -0.05 | 0.086 |
| GOLGB1 | S1568 | -0.04 | -0.02 | 0.10 | -0.04 | -0.16 | 0.14 | 0.759 |
| GOLGB1 | S649 | -0.26 | 0.12 | 0.10 | 0.38 | 0.07 | -0.62 | 0.892 |
| GOLGB1 | T/S | 0.05 | -0.07 | 0.07 | 0.23 | -0.21 | -0.12 | 0.74 |
| GOLGB1 | S1564 | 0.02 | -0.01 | 0.08 | -0.16 | -0.26 | 0.27 | 0.666 |
| GOLGB1 | S3124 | 0.27 | -0.04 | -0.09 | 0.19 | 0.00 | -0.43 | 0.582 |
| GOLGB1 | S/T | 0.10 | 0.09 | 0.12 | -0.16 | 0.20 | -0.44 | 0.278 |
| GOLGB1 | S | 0.20 | 0.45 | -0.09 | -0.28 | 0.11 | -0.62 | 0.164 |
| GOLGB1 | S671 | 0.20 | 0.45 | -0.09 | -0.28 | 0.11 | -0.62 | 0.164 |
| GOLGB1 | S/T | 0.15 | 0.27 | 0.27 | 0.04 | -0.04 | -1.05 | 0.175 |
| GOLIM4 | Y506 | -0.15 | -0.10 | -0.28 | 0.29 | -0.27 | 0.37 | 0.215 |
| GOLIM4 | Y506 | -0.16 | 0.04 | -0.36 | 0.39 | -0.23 | 0.18 | 0.271 |
| GOLIM4 | S364 | -0.16 | -0.01 | 0.15 | -0.02 | 0.00 | 0.03 | 0.896 |
| GOLIM4 | S364 | 0.34 | 0.06 | -0.27 | -0.19 | -0.01 | -0.01 | 0.562 |
| GOLIM4 | S364 | -0.03 | 0.17 | 0.14 | -0.02 | -0.03 | -0.26 | 0.118 |
| GOLIM4 | S583 | 0.23 | 0.36 | 0.12 | -0.15 | -0.35 | -0.37 | 0.006 |
| GOLM2 | Y389 | 0.13 | -0.36 | -0.03 | 0.08 | -0.18 | 0.27 | 0.506 |
| GOLM2 | Y389 | 0.00 | 0.17 | 0.04 | 0.06 | -0.20 | -0.08 | 0.191 |
| GOLM2 | Y379; Y389 | 0.03 | 0.03 | 0.19 | -0.07 | 0.13 | -0.38 | 0.295 |
| GOLM2 | S332 | 0.10 | 0.13 | 0.11 | 0.03 | -0.17 | -0.23 | 0.04 |
| GOLM2 | S | 0.31 | -0.06 | 0.07 | 0.07 | -0.14 | -0.34 | 0.212 |
| GOLM2 | S366 | 0.14 | 0.23 | 0.20 | -0.14 | -0.14 | -0.40 | 0.009 |
| GOLM2 | S233 | 0.46 | 0.32 | 0.31 | -0.30 | -0.55 | -0.64 | 0.002 |
| GOLPH3L | S112 | 0.03 | -0.39 | 0.12 | -0.21 | 0.08 | 0.27 | 0.57 |
| GON4L | S/T | 0.32 | -0.59 | -0.22 | -0.05 | 0.06 | 0.29 | 0.407 |
| GON4L | S/T | 0.19 | -0.20 | -0.31 | -0.19 | 0.12 | 0.28 | 0.433 |
| GON4L | S206 | -0.28 | 0.35 | 0.10 | -0.03 | 0.00 | -0.25 | 0.496 |
| GORASP1 | T216; T237 | -0.02 | -0.03 | -0.11 | 0.16 | -0.10 | 0.08 | 0.282 |
| GORASP2 | T415 | -0.01 | 0.00 | -0.21 | -0.21 | 0.15 | 0.23 | 0.436 |
| GORASP2 | T433; S451 | -0.06 | -0.03 | -0.03 | -0.30 | 0.08 | 0.27 | 0.754 |
| GORASP2 | S451 | -0.02 | 0.10 | 0.08 | -0.27 | 0.17 | -0.11 | 0.427 |
| GORASP2 | S451 | 0.02 | 0.20 | 0.04 | -0.27 | 0.00 | -0.03 | 0.138 |
| GPALPP1 | S28 | 0.36 | 0.01 | 0.04 | -0.12 | -0.14 | -0.23 | 0.062 |
| GPALPP1 | S105 | 0.43 | 0.35 | 0.21 | -0.57 | 0.03 | -0.94 | 0.046 |
| GPATCH1 | S/Y/T | 0.12 | 0.24 | 0.14 | -0.15 | -0.06 | -0.38 | 0.024 |
| GPATCH1 | S/T | 0.25 | -0.16 | 0.36 | -0.18 | 0.12 | -0.60 | 0.225 |
| GPATCH1 | S8 | 0.18 | 0.36 | 0.05 | -0.34 | -0.15 | -0.21 | 0.016 |
| GPATCH1 | S715 | 0.15 | 0.33 | 0.24 | -0.34 | -0.18 | -0.36 | 0.002 |
| GPATCH1 | S203 | 0.22 | 0.30 | 0.25 | -0.34 | -0.05 | -0.61 | 0.023 |

| | | | | | | | | |
|---|---|---|---|---|---|---|---|---|
| GPATCH2 | S115; S117 | -0.05 | -0.05 | -0.01 | 0.09 | -0.15 | 0.16 | 0.486 |
| GPATCH2L | S/T | 0.02 | -0.24 | -0.52 | -0.18 | 0.35 | 0.36 | 0.147 |
| GPATCH2L | S447 | -0.31 | -0.05 | -0.29 | 0.17 | 0.09 | 0.29 | 0.017 |
| GPATCH4 | S137; S | 0.23 | 0.22 | 0.06 | -0.18 | -0.01 | -0.44 | 0.05 |
| GPATCH8 | S1065 | -1.24 | -0.91 | -0.50 | 0.40 | 0.70 | 0.47 | 0.004 |
| GPATCH8 | S1065; S1068 | -0.64 | -0.55 | -0.62 | 0.70 | -0.19 | 0.61 | 0.026 |
| GPATCH8 | S494 | -0.50 | -0.53 | -0.48 | 0.37 | 0.38 | 0.37 | 5E-07 |
| GPATCH8 | S911; S914 | 0.18 | -0.09 | -0.46 | 0.26 | 0.23 | -0.26 | 0.474 |
| GPATCH8 | S908; S911; S914 | -0.24 | -0.08 | 0.18 | -0.02 | 0.15 | -0.02 | 0.573 |
| GPATCH8 | S758 | 0.08 | 0.01 | -0.15 | 0.19 | -0.10 | -0.06 | 0.812 |
| GPATCH8 | S981 | 0.04 | 0.10 | -0.18 | 0.08 | 0.04 | -0.10 | 0.825 |
| GPATCH8 | S740 | 0.15 | 0.01 | -0.09 | 0.13 | -0.11 | -0.11 | 0.635 |
| GPATCH8 | S1014 | 0.05 | 0.01 | 0.02 | -0.24 | -0.10 | 0.23 | 0.665 |
| GPATCH8 | S1107 | 0.21 | 0.13 | -0.07 | 0.06 | -0.20 | -0.17 | 0.171 |
| GPATCH8 | S1068; S1076 | 0.22 | 0.09 | 0.03 | -0.15 | -0.08 | -0.14 | 0.016 |
| GPATCH8 | S1035 | 0.20 | 0.19 | -0.03 | -0.36 | -0.13 | 0.05 | 0.129 |
| GPATCH8 | S1081 | 0.16 | 0.16 | 0.08 | -0.01 | -0.25 | -0.20 | 0.022 |
| GPATCH8 | S1081 | 0.17 | 0.18 | 0.06 | 0.05 | -0.29 | -0.25 | 0.057 |
| GPI | T109 | -0.26 | -0.02 | -0.14 | -0.08 | 0.30 | 0.14 | 0.121 |
| GPKOW | S42 | 0.12 | 0.12 | -0.05 | -0.06 | 0.17 | -0.37 | 0.415 |
| GPRC5A | S301 | -0.06 | -0.01 | -0.14 | 0.18 | -0.13 | 0.12 | 0.289 |
| GPRC5C | S383 | -0.91 | -0.86 | -2.22 | 0.99 | 0.23 | 0.62 | 0.017 |
| GPRC5C | T423 | -0.25 | 0.47 | -0.59 | 0.06 | -0.10 | 0.18 | 0.63 |
| GPRC5C | T423 | -0.01 | -0.06 | -0.01 | 0.06 | 0.19 | -0.20 | 0.725 |
| GPRC5C | T423 | -0.14 | 0.15 | -0.02 | 0.52 | -0.74 | -0.06 | 0.827 |
| GPRC5C | T423 | -0.05 | 0.21 | 0.01 | -0.07 | -0.06 | -0.05 | 0.204 |
| GPRC5C | S/Y/T | -0.40 | 0.38 | 0.25 | 0.24 | -0.48 | -0.22 | 0.503 |
| GPRC5C | T423 | 0.19 | 0.32 | -0.01 | -0.22 | -0.34 | -0.05 | 0.043 |
| GPRC5C | S/Y/T | -0.31 | 0.31 | 0.46 | 0.05 | -0.33 | -0.42 | 0.239 |
| GPRIN1 | S615 | -0.10 | 0.06 | 0.22 | 0.01 | -0.34 | 0.08 | 0.41 |
| GPRIN1 | T/S | -0.10 | 0.03 | 0.31 | 0.04 | -0.22 | -0.12 | 0.277 |
| GPRIN1 | S799 | 0.10 | 0.29 | 0.10 | -0.31 | -0.12 | -0.13 | 0.015 |
| GPRIN1 | S615 | 0.29 | 0.24 | 0.37 | -0.45 | -0.03 | -0.76 | 0.03 |
| GPS1 | S474 | 0.02 | -0.35 | 0.15 | -0.13 | 0.12 | 0.12 | 0.621 |
| GPSM1 | S492 | -0.19 | -0.17 | -0.18 | 0.03 | -0.02 | 0.42 | 0.081 |
| GPSM2 | T526 | -0.16 | -0.46 | -0.15 | 0.04 | 0.21 | 0.37 | 0.028 |
| GRAMD1A | T277 | 0.14 | 0.08 | 0.13 | 0.04 | -0.35 | -0.10 | 0.096 |
| GRAMD1A | S16 | 0.20 | 0.22 | 0.03 | -0.38 | 0.07 | -0.25 | 0.082 |
| GRAMD1A | S267 | 0.25 | -0.01 | 0.29 | -0.02 | -0.20 | -0.44 | 0.061 |
| GRAMD2B | S72 | 0.16 | 0.05 | 0.14 | -0.06 | -0.05 | -0.27 | 0.042 |
| GRAMD4 | S159 | -0.19 | -0.14 | 0.09 | 0.00 | 0.10 | 0.10 | 0.198 |
| GRAMD4 | S24 | 0.05 | 0.31 | 0.05 | -0.19 | -0.13 | -0.16 | 0.028 |
| GRAMD4 | S28 | 0.10 | 0.24 | 0.21 | -0.12 | -0.09 | -0.44 | 0.03 |
| GRAMD4 | S/T | 0.11 | 0.27 | 0.24 | -0.01 | -0.19 | -0.60 | 0.061 |
| GRB10 | S104 | -0.24 | -0.14 | -0.17 | 0.17 | 0.17 | 0.14 | 5E-04 |
| GRB7 | S163 | -0.25 | -0.01 | 0.06 | 0.23 | -0.35 | 0.23 | 0.661 |
| GRB7 | S163 | -0.25 | -0.01 | 0.06 | 0.23 | -0.35 | 0.23 | 0.661 |
| GRHL1 | T208 | 0.03 | -0.08 | -0.42 | 0.28 | 0.03 | 0.08 | 0.135 |

| | | | | | | | | |
|---|---|---|---|---|---|---|---|---|
| GRHL1 | S/T | -0.04 | -0.12 | -0.17 | 0.19 | -0.01 | 0.12 | 0.041 |
| GRHL1 | T208 | 0.10 | -0.14 | -0.03 | 0.19 | -0.08 | -0.07 | 0.761 |
| GRHL1 | S95 | 0.05 | 0.14 | 0.04 | -0.01 | -0.20 | -0.04 | 0.075 |
| GRHL2 | S210 | 0.04 | -0.02 | 0.11 | -0.19 | 0.15 | -0.12 | 0.414 |
| GRHL2 | S210 | 0.18 | 0.03 | -0.01 | -0.07 | -0.03 | -0.13 | 0.094 |
| GRHL2 | S210; T211; S214 | 0.22 | 0.14 | 0.08 | -0.25 | -0.06 | -0.19 | 0.01 |
| GRHL2 | S217 | 0.17 | 0.26 | -0.02 | -0.01 | 0.09 | -0.67 | 0.254 |
| GRHL2 | S210; T211 | 0.25 | 0.12 | 0.10 | -0.26 | -0.06 | -0.23 | 0.013 |
| GRHL2 | T211; S214 | 0.26 | 0.15 | 0.15 | -0.39 | -0.16 | -0.10 | 0.014 |
| GRHL3 | T/S/Y | 0.03 | 0.38 | 0.45 | -0.23 | -0.31 | -0.62 | 0.019 |
| GRHL3 | S493 | 0.43 | 0.18 | 0.32 | -0.56 | -0.04 | -0.67 | 0.025 |
| GRIN2C | T1196 | 0.27 | 0.13 | 0.44 | -0.68 | -0.34 | -0.10 | 0.026 |
| GRIP1 | S43 | -0.56 | 0.11 | -0.28 | -0.40 | 0.17 | 0.62 | 0.357 |
| GRIPAP1 | S | 0.07 | -0.34 | 0.25 | 0.11 | -0.18 | 0.01 | 0.969 |
| GRK2 | S670 | 0.20 | 0.10 | -0.31 | 0.08 | -0.04 | -0.08 | 0.934 |
| GRK5 | S484 | -0.05 | 0.08 | -0.05 | -0.19 | 0.12 | 0.06 | 0.987 |
| GRSF1 | S244 | -0.02 | 0.10 | 0.08 | 0.03 | -0.19 | -0.02 | 0.23 |
| GRWD1 | S/T/Y | -0.30 | -0.26 | 0.05 | 0.05 | 0.12 | 0.27 | 0.069 |
| GSE1 | S1101 | -0.03 | -0.06 | -0.15 | -0.25 | 0.07 | 0.34 | 0.477 |
| GSE1 | T433 | -0.08 | 0.06 | 0.24 | -0.15 | -0.05 | -0.05 | 0.183 |
| GSK3B | S/T | -0.23 | 0.02 | -0.19 | 0.14 | 0.20 | 0.00 | 0.067 |
| GSK3B | S215 | 0.01 | -0.08 | -0.16 | 0.08 | -0.08 | 0.19 | 0.207 |
| GSS | S415 | 0.06 | 0.08 | 0.03 | -0.19 | -0.09 | 0.08 | 0.192 |
| GTF2F1 | S224 | -0.25 | -0.18 | -0.18 | 0.02 | 0.01 | 0.45 | 0.067 |
| GTF2F1 | S221; S224 | -0.14 | 0.00 | -0.08 | 0.04 | -0.04 | 0.19 | 0.173 |
| GTF2F1 | S221; S224 | -0.13 | 0.03 | 0.01 | 0.05 | -0.06 | 0.09 | 0.434 |
| GTF2F1 | S385; T389 | 0.03 | -0.12 | 0.11 | -0.13 | -0.12 | 0.20 | 0.874 |
| GTF2F1 | S385 | 0.12 | -0.03 | 0.01 | -0.07 | -0.04 | 0.00 | 0.204 |
| GTF2F1 | S433 | 0.01 | 0.03 | 0.14 | -0.06 | 0.10 | -0.25 | 0.29 |
| GTF2F1 | S217; S221; S224 | 0.13 | -0.07 | 0.11 | -0.36 | -0.18 | 0.27 | 0.496 |
| GTF2F2 | S248 | 0.13 | 0.07 | 0.50 | -0.19 | -0.35 | -0.36 | 0.021 |
| GTF2I | S784 | -0.14 | -0.29 | -0.35 | 0.39 | 0.22 | 0.01 | 0.02 |
| GTF2I | S674 | -0.01 | -0.22 | -0.13 | -0.06 | 0.23 | 0.13 | 0.1 |
| GTF2I | T703 | -0.06 | -0.14 | 0.02 | -0.07 | 0.22 | 0.00 | 0.331 |
| GTF2I | S914 | -0.30 | -0.10 | 0.21 | 0.03 | -0.14 | 0.23 | 0.6 |
| GTF2I | S674 | 0.04 | -0.04 | -0.15 | 0.10 | -0.11 | 0.14 | 0.381 |
| GTF2I | S412 | -0.11 | 0.11 | -0.12 | -0.20 | 0.40 | -0.17 | 0.815 |
| GTF2I | S823 | 0.19 | 0.09 | 0.02 | -0.17 | 0.15 | -0.37 | 0.225 |
| GTF2I | T558 | 0.09 | 0.21 | 0.17 | -0.31 | 0.13 | -0.41 | 0.105 |
| GTF2IRD1 | S448 | 0.06 | -0.14 | 0.08 | -0.06 | -0.06 | 0.10 | 0.975 |
| GTF3A | S347 | -0.58 | -1.09 | -0.29 | 0.68 | 0.44 | 0.12 | 0.02 |
| GTF3A | S347 | -0.54 | -0.67 | -0.06 | 0.28 | 0.23 | 0.43 | 0.019 |
| GTF3C1 | S1653 | -0.28 | -0.01 | -0.88 | 0.13 | 0.27 | 0.43 | 0.071 |
| GTF3C1 | S754 | -0.11 | -0.37 | -0.05 | 0.19 | 0.17 | 0.09 | 0.033 |
| GTF3C1 | T/S | -0.49 | -0.03 | -0.02 | 0.27 | 0.39 | -0.31 | 0.329 |
| GTF3C1 | S1868 | -0.04 | -0.12 | -0.32 | 0.23 | -0.10 | 0.27 | 0.112 |
| GTF3C1 | S1068 | -0.14 | -0.05 | -0.26 | 0.03 | 0.15 | 0.22 | 0.025 |
| GTF3C1 | S1865 | -0.09 | -0.13 | -0.12 | 0.12 | -0.01 | 0.20 | 0.024 |

| | | | | | | | | |
|---|---|---|---|---|---|---|---|---|
| **GTF3C1** | **S1865** | 0.00 | -0.21 | -0.02 | 0.22 | 0.16 | -0.20 | 0.404 |
| **GTF3C1** | **S1865; S1868** | -0.16 | -0.13 | 0.11 | -0.03 | 0.12 | 0.06 | 0.299 |
| **GTF3C1** | **T514** | 0.09 | 0.00 | -0.25 | -0.06 | 0.10 | 0.09 | 0.452 |
| **GTF3C1** | **S1868** | -0.04 | 0.00 | -0.10 | 0.12 | -0.05 | 0.06 | 0.187 |
| **GTF3C1** | **S1865** | -0.15 | -0.19 | 0.18 | 0.00 | 0.15 | -0.04 | 0.547 |
| **GTF3C1** | **S1865** | 0.02 | -0.05 | -0.09 | 0.12 | -0.12 | 0.10 | 0.448 |
| **GTF3C1** | **S1068** | -0.02 | -0.01 | -0.07 | -0.09 | 0.17 | 0.01 | 0.447 |
| **GTF3C1** | **S1865; S1868** | -0.09 | -0.12 | 0.11 | 0.05 | 0.14 | -0.12 | 0.613 |
| **GTF3C1** | **S** | -0.04 | -0.03 | -0.01 | 0.00 | -0.04 | 0.11 | 0.324 |
| **GTF3C1** | **S1868** | 0.00 | -0.04 | 0.01 | 0.03 | 0.02 | -0.02 | 0.498 |
| **GTF3C1** | **S667** | 0.01 | -0.17 | 0.12 | 0.20 | 0.07 | -0.27 | 0.944 |
| **GTF3C1** | **S1868** | 0.04 | -0.06 | 0.13 | -0.08 | 0.01 | -0.04 | 0.315 |
| **GTF3C1** | **S1062** | 0.12 | 0.08 | -0.04 | -0.01 | 0.12 | -0.32 | 0.423 |
| **GTF3C1** | **S1865** | 0.12 | 0.07 | 0.00 | 0.13 | -0.12 | -0.24 | 0.286 |
| **GTF3C1** | **S1865** | -0.01 | 0.19 | 0.02 | -0.04 | -0.10 | -0.08 | 0.093 |
| **GTF3C1** | **S1068** | 0.01 | 0.10 | 0.08 | -0.11 | 0.42 | -0.75 | 0.566 |
| **GTF3C1** | **S1062; S1068** | 0.10 | 0.09 | 0.15 | -0.25 | -0.26 | 0.11 | 0.118 |
| **GTF3C1** | **S1062; S1068** | 0.18 | 0.14 | 0.15 | -0.23 | -0.06 | -0.25 | 0.005 |
| **GTF3C1** | **S1062; S1068** | 0.25 | 0.17 | 0.12 | -0.18 | -0.10 | -0.34 | 0.008 |
| **GTF3C2** | **S901** | -0.17 | -0.16 | -0.27 | 0.19 | -0.02 | 0.33 | 0.028 |
| **GTF3C2** | **S167** | 0.14 | 0.09 | -0.03 | -0.02 | -0.06 | -0.15 | 0.077 |
| **GTF3C3** | **S43** | 0.00 | -0.12 | -0.26 | 0.18 | -0.05 | 0.19 | 0.098 |
| **GTF3C3** | **S43** | 0.26 | -0.24 | -0.33 | 0.15 | 0.09 | -0.03 | 0.417 |
| **GTF3C4** | **S611** | -0.05 | -0.05 | -0.12 | 0.09 | 0.09 | 0.03 | 0.01 |
| **GTF3C4** | **S611** | 0.04 | 0.06 | -0.15 | 0.10 | -0.01 | -0.05 | 0.711 |
| **GTF3C4** | **S611** | -0.04 | -0.04 | 0.06 | -0.04 | -0.04 | 0.09 | 0.844 |
| **GTF3C4** | **S611** | 0.10 | -0.04 | 0.10 | 0.08 | -0.20 | -0.07 | 0.277 |
| **GTF3C4** | **S611** | 0.10 | -0.02 | 0.26 | -0.18 | 0.04 | -0.26 | 0.111 |
| **GTPBP1** | **S47** | 0.23 | 0.28 | -0.50 | -0.30 | -0.12 | 0.23 | 0.827 |
| **GTPBP1** | **S47** | 0.12 | 0.34 | -0.21 | -0.27 | 0.08 | -0.16 | 0.349 |
| **GTPBP4** | **S468; S470; S472** | 0.04 | 0.00 | 0.10 | -0.09 | 0.05 | -0.11 | 0.166 |
| **GTPBP4** | **S468; S470; S472** | 0.03 | 0.04 | 0.09 | -0.15 | 0.02 | -0.05 | 0.104 |
| **GTPBP4** | **S470; S472** | 0.05 | 0.11 | 0.10 | -0.08 | -0.10 | -0.09 | 9E-04 |
| **GTPBP4** | **S470; S472** | 0.10 | 0.01 | 0.17 | -0.13 | -0.01 | -0.17 | 0.04 |
| **GTPBP4** | **S470** | 0.19 | 0.23 | 0.16 | -0.38 | -0.07 | -0.23 | 0.01 |
| **GTPBP4** | **S470** | 0.18 | 0.33 | 0.27 | -0.32 | -0.21 | -0.44 | 0.002 |
| **GTSE1** | **S** | -0.43 | -0.40 | -0.37 | 0.38 | 0.28 | 0.28 | 5E-05 |
| **GTSE1** | **S580** | -0.21 | -0.32 | -0.10 | 0.40 | 0.07 | 0.05 | 0.044 |
| **GTSE1** | **S461** | -0.12 | 0.01 | -0.18 | 0.21 | 0.24 | -0.24 | 0.381 |
| **GTSE1** | **S575** | -0.06 | 0.02 | 0.03 | 0.07 | 0.07 | -0.14 | 0.961 |
| **GUCY1A1** | **S/T/Y** | 0.21 | -0.12 | 0.19 | -0.07 | 0.08 | -0.37 | 0.277 |
| **GYS1** | **S698** | -0.01 | -0.08 | 0.54 | -0.19 | 0.10 | -0.60 | 0.252 |
| **GZF1** | **T/S/Y** | -0.27 | -0.26 | -0.13 | 0.40 | -0.33 | 0.39 | 0.203 |
| **GZF1** | **S613** | 0.25 | 0.11 | 0.05 | -0.01 | -0.04 | -0.45 | 0.118 |
| **GZF1** | **S265** | 0.25 | 0.33 | 0.17 | -0.22 | 0.02 | -0.82 | 0.08 |
| **H1-0** | **T153** | -0.41 | -0.75 | -0.64 | 0.42 | 0.28 | 0.54 | 0.001 |
| **H1-0** | **T119** | -0.26 | -0.52 | -0.48 | 0.28 | 0.13 | 0.52 | 0.006 |
| **H1-0** | **S92** | -0.18 | -0.23 | -0.12 | 0.19 | 0.05 | 0.22 | 0.006 |

| | | | | | | | | |
|---|---|---|---|---|---|---|---|---|
| **H1-0** | **S66** | -0.20 | -0.11 | -0.18 | 0.19 | 0.27 | -0.04 | 0.037 |
| **H1-0** | **S98** | 0.07 | 0.05 | -0.03 | 0.09 | -0.08 | -0.11 | 0.393 |
| **H1-10** | **S31** | -0.34 | -0.34 | -0.38 | 0.36 | 0.26 | 0.23 | 1E-04 |
| **H1-10** | **S27; S31** | -0.20 | -0.19 | -0.19 | 0.13 | 0.14 | 0.24 | 5E-04 |
| **H1-10** | **S31** | -0.16 | -0.12 | -0.32 | 0.26 | 0.44 | -0.28 | 0.204 |
| **H1-2** | **S36** | -0.31 | -0.50 | -0.57 | 0.24 | 0.32 | 0.46 | 0.001 |
| **H1-2** | **S36** | -0.38 | -0.51 | -0.39 | 0.37 | 0.28 | 0.33 | 1E-04 |
| **H1-2** | **T146** | -0.23 | -0.56 | -0.35 | 0.41 | 0.05 | 0.40 | 0.012 |
| **H1-2** | **T146** | -0.20 | -0.45 | -0.39 | 0.31 | 0.07 | 0.42 | 0.009 |
| **H1-2** | **S113** | -0.09 | -0.28 | -0.30 | 0.21 | 0.01 | 0.32 | 0.023 |
| **H1-2** | **T146** | 0.02 | -0.15 | -0.08 | 0.22 | 0.24 | -0.33 | 0.592 |
| **H1-2** | **S104** | 0.04 | -0.10 | -0.09 | 0.07 | -0.08 | 0.13 | 0.322 |
| **H1-2** | **T146** | 0.14 | -0.06 | -0.08 | 0.24 | 0.30 | -0.77 | 0.841 |
| **H1-2** | **T165** | -0.07 | 0.36 | 0.19 | -0.36 | -0.02 | -0.23 | 0.08 |
| **H1-2** | **S58** | 0.33 | 0.45 | 0.60 | -0.57 | -0.35 | -1.35 | 0.018 |
| **H1-3** | **T147** | -0.01 | -0.33 | -0.24 | 0.25 | 0.11 | 0.13 | 0.029 |
| **H1-3** | **S114** | -0.05 | -0.22 | -0.30 | 0.16 | 0.04 | 0.29 | 0.027 |
| **H1-3** | **T147** | -0.09 | -0.31 | 0.00 | 0.20 | 0.04 | 0.12 | 0.073 |
| **H1-4** | **T146** | -0.13 | -0.34 | -0.25 | 0.20 | 0.08 | 0.32 | 0.008 |
| **H1-4** | **T146** | 0.00 | -0.27 | -0.08 | 0.21 | 0.15 | -0.05 | 0.117 |
| **H1-4** | **S187** | 0.22 | 0.01 | -0.07 | -0.06 | -0.08 | -0.04 | 0.262 |
| **H1-5** | **T138** | -0.12 | -0.36 | -0.40 | 0.26 | 0.04 | 0.40 | 0.018 |
| **H1-5** | **T138** | -0.10 | -0.33 | -0.33 | 0.32 | 0.13 | 0.19 | 0.008 |
| **H1-5** | **S189** | 0.11 | -0.12 | -0.34 | 0.12 | -0.03 | 0.20 | 0.215 |
| **H1-5** | **S189** | 0.18 | -0.13 | -0.28 | 0.05 | -0.02 | 0.14 | 0.407 |
| **H1-5** | **S61** | 0.05 | 0.30 | 0.33 | -0.56 | -0.02 | -0.31 | 0.044 |
| **H2AC4** | **S123** | -0.09 | -0.25 | -0.39 | 0.28 | 0.24 | 0.08 | 0.014 |
| **H2AX** | **S122** | 0.18 | -0.14 | -0.12 | -0.08 | 0.07 | 0.06 | 0.735 |
| **H2AX** | **S122** | 0.30 | 0.20 | 0.18 | -0.31 | 0.00 | -0.53 | 0.034 |
| **H2AZ1** | **S99** | 0.22 | 0.20 | 0.18 | -0.12 | -0.14 | -0.47 | 0.018 |
| **H2BC13** | **S39** | -0.03 | -0.39 | -0.39 | 0.14 | 0.00 | 0.48 | 0.062 |
| **H2BC13** | **S39** | 0.10 | 0.00 | -0.41 | 0.04 | -0.20 | 0.35 | 0.505 |
| **H2BC26** | **T89** | -0.06 | 0.06 | -0.14 | -0.05 | -0.03 | 0.20 | 0.431 |
| **H2BC26** | **S39** | 0.19 | -0.09 | -0.07 | -0.07 | -0.14 | 0.15 | 0.798 |
| **H2BC26** | **S39** | 0.23 | -0.06 | -0.05 | 0.14 | -0.05 | -0.25 | 0.547 |
| **H2BC26** | **S57** | 0.18 | 0.08 | 0.21 | -0.10 | -0.06 | -0.39 | 0.037 |
| **H2BC26** | **S113** | 0.20 | 0.22 | 0.18 | -0.29 | 0.00 | -0.44 | 0.026 |
| **H3-3A** | **S32** | -0.78 | -1.03 | -0.79 | 0.87 | 0.22 | 0.44 | 0.003 |
| **H3-3A** | **S29** | -0.75 | -0.64 | -0.61 | 0.66 | 0.22 | 0.44 | 0.001 |
| **H3-3A** | **T81** | -0.08 | -0.18 | -0.30 | 0.21 | 0.13 | 0.15 | 0.007 |
| **H3-7** | **S29** | -0.57 | -1.11 | -1.06 | 0.83 | 0.10 | 0.62 | 0.007 |
| **H3-7** | **S29** | -0.59 | -0.68 | -0.85 | 0.61 | 0.09 | 0.65 | 0.004 |
| **H3-7** | **T81** | -0.16 | 0.02 | -0.26 | -0.30 | 0.53 | 0.00 | 0.454 |
| **H3-7** | **T59** | 0.05 | -0.04 | 0.00 | -0.07 | 0.13 | -0.08 | 0.924 |
| **H3Y2** | **S58** | 0.00 | -0.06 | -0.06 | 0.02 | 0.19 | -0.10 | 0.422 |
| **H4C1** | **S48** | -0.13 | -0.09 | -0.10 | 0.10 | 0.01 | 0.19 | 0.016 |
| **H4C1** | **T74** | -0.03 | -0.08 | 0.07 | 0.18 | -0.01 | -0.15 | 0.873 |
| **H4C1** | **S48** | -0.09 | 0.04 | 0.09 | -0.17 | -0.02 | 0.13 | 0.757 |

| | | | | | | | | |
|---|---|---|---|---|---|---|---|---|
| HABP4 | S108 | -0.18 | 0.20 | 0.42 | -0.21 | -0.14 | -0.22 | 0.132 |
| HABP4 | S108 | 0.18 | 0.23 | 0.25 | 0.02 | -0.30 | -0.54 | 0.039 |
| HACD3 | S114 | 0.08 | 0.09 | 0.14 | -0.05 | -0.01 | -0.29 | 0.071 |
| HACD3 | S114 | 0.09 | 0.11 | 0.18 | -0.06 | -0.01 | -0.38 | 0.084 |
| HACD3 | S135 | 0.18 | -0.07 | 0.29 | -0.24 | -0.03 | -0.20 | 0.08 |
| HACD3 | S135 | 0.04 | 0.10 | 0.31 | -0.13 | -0.13 | -0.27 | 0.025 |
| HACD3 | S135 | 0.19 | 0.05 | 0.23 | -0.15 | -0.12 | -0.26 | 0.008 |
| HACD3 | T/S | 0.37 | 0.14 | 0.14 | -0.35 | -0.09 | -0.34 | 0.014 |
| HARS2 | S67 | -0.02 | -0.04 | -0.16 | 0.13 | 0.08 | -0.01 | 0.071 |
| HASPIN | S220 | -0.19 | -0.69 | -1.19 | 0.54 | 0.07 | 0.65 | 0.03 |
| HASPIN | S147 | -0.38 | 0.01 | -0.34 | 0.21 | 0.18 | 0.20 | 0.024 |
| HASPIN | T97 | 0.16 | -0.01 | -0.17 | -0.02 | 0.05 | -0.03 | 0.921 |
| HASPIN | S147 | -0.08 | 0.09 | 0.19 | -0.13 | 0.03 | -0.12 | 0.204 |
| HAUS6 | S552 | 0.11 | 0.01 | -0.49 | 0.19 | -0.13 | 0.20 | 0.389 |
| HAUS6 | T584 | -0.01 | -0.17 | -0.10 | 0.30 | -0.26 | 0.16 | 0.413 |
| HAX1 | S/T | 0.15 | -0.48 | -0.28 | -0.17 | 0.55 | -0.01 | 0.319 |
| HCFC1 | S1507 | 0.03 | -0.14 | -0.06 | 0.06 | -0.18 | 0.24 | 0.513 |
| HCFC1 | S598 | -0.16 | -0.06 | 0.24 | -0.24 | 0.00 | 0.17 | 0.872 |
| HCFC1 | S666 | 0.40 | 0.38 | -0.25 | -1.19 | 0.07 | 0.07 | 0.325 |
| HDAC1 | S393 | -0.02 | -0.10 | -0.12 | 0.06 | 0.10 | 0.07 | 0.008 |
| HDAC1 | S393 | 0.06 | -0.22 | 0.00 | -0.04 | 0.03 | 0.15 | 0.39 |
| HDAC1 | S393 | -0.19 | 0.02 | 0.05 | 0.16 | -0.02 | -0.06 | 0.566 |
| HDAC1 | S421; S423 | 0.05 | -0.10 | 0.19 | -0.01 | -0.44 | 0.22 | 0.588 |
| HDAC1 | S406 | 0.16 | 0.32 | -0.26 | -0.38 | 0.32 | -0.35 | 0.508 |
| HDAC1 | S421; S423 | 0.17 | 0.12 | 0.17 | -0.23 | 0.04 | -0.35 | 0.042 |
| HDAC1 | S421 | 0.30 | 0.28 | 0.27 | -0.41 | -0.11 | -0.59 | 0.01 |
| HDAC2 | S394 | -0.27 | 0.16 | -0.44 | -0.38 | 0.36 | 0.34 | 0.386 |
| HDAC2 | S422; S424 | -0.05 | -0.05 | -0.05 | 0.04 | 0.07 | 0.03 | 0.002 |
| HDAC2 | S422; S424 | -0.27 | -0.06 | 0.21 | 0.02 | -0.10 | 0.15 | 0.71 |
| HDAC2 | S394 | 0.03 | 0.02 | -0.08 | -0.05 | 0.15 | -0.08 | 0.84 |
| HDAC2 | S394 | 0.02 | 0.04 | -0.09 | 0.09 | 0.09 | -0.17 | 0.872 |
| HDAC2 | S394 | 0.22 | -0.29 | 0.05 | 0.27 | -0.06 | -0.31 | 0.912 |
| HDAC2 | S394 | 0.06 | 0.14 | 0.05 | -0.01 | -0.04 | -0.23 | 0.081 |
| HDAC2 | S424 | 0.10 | 0.20 | 0.17 | -0.38 | 0.02 | -0.19 | 0.047 |
| HDAC3 | S424 | -0.22 | -0.23 | -0.32 | 0.37 | -0.08 | 0.32 | 0.033 |
| HDAC3 | S424 | -0.18 | -0.29 | -0.29 | 0.28 | -0.04 | 0.37 | 0.026 |
| HDAC6 | S22 | 0.05 | -0.19 | 0.15 | 0.11 | -0.27 | 0.10 | 0.9 |
| HDGF | S102 | 0.16 | -0.11 | -0.54 | 0.27 | -0.26 | 0.30 | 0.381 |
| HDGF | S132; S133 | -0.04 | 0.01 | -0.14 | 0.08 | -0.13 | 0.19 | 0.364 |
| HDGF | S132; S133 | 0.18 | 0.08 | -0.14 | -0.03 | -0.05 | -0.06 | 0.424 |
| HDGF | S133 | 0.17 | 0.16 | 0.01 | -0.09 | -0.05 | -0.24 | 0.036 |
| HDGF | S165 | 0.20 | 0.18 | -0.03 | -0.14 | -0.10 | -0.16 | 0.033 |
| HDGF | S165 | 0.17 | 0.12 | 0.16 | -0.14 | -0.05 | -0.33 | 0.018 |
| HDGF | S165 | 0.33 | 0.31 | 0.02 | -0.16 | -0.04 | -0.71 | 0.086 |
| HDGFL2 | S599 | -0.25 | -0.26 | -0.50 | 0.35 | 0.12 | 0.33 | 0.006 |
| HDGFL2 | S399 | -0.34 | -0.04 | -0.25 | 0.22 | -0.09 | 0.37 | 0.083 |
| HDGFL2 | T/S | -0.16 | -0.11 | -0.09 | -0.08 | 0.28 | 0.12 | 0.101 |
| HDGFL2 | T/S | 0.00 | -0.29 | 0.00 | -0.05 | 0.60 | -0.51 | 0.759 |

| | | | | | | | | |
|---|---|---|---|---|---|---|---|---|
| HDGFL2 | S299 | 0.09 | 0.07 | -0.19 | 0.04 | 0.00 | -0.03 | 0.881 |
| HDGFL2 | T421 | 0.03 | -0.02 | 0.00 | -0.02 | -0.14 | 0.14 | 0.89 |
| HDGFL2 | S | 0.00 | 0.00 | 0.04 | 0.03 | -0.12 | 0.04 | 0.598 |
| HDGFL2 | S369; S370 | 0.01 | 0.13 | -0.08 | 0.07 | -0.11 | -0.02 | 0.642 |
| HDGFL2 | S137 | 0.32 | 0.20 | -0.59 | -0.63 | 0.09 | 0.30 | 0.892 |
| HDGFL2 | S366; S369; S370 | 0.17 | -0.13 | 0.04 | -0.14 | -0.13 | 0.15 | 0.616 |
| HDGFL2 | S652 | 0.06 | -0.04 | 0.16 | -0.11 | -0.02 | -0.06 | 0.112 |
| HDGFL2 | S240 | 0.09 | 0.05 | 0.18 | -0.08 | -0.11 | -0.16 | 0.006 |
| HDGFL2 | S240 | 0.17 | -0.03 | 0.21 | -0.03 | -0.15 | -0.22 | 0.054 |
| HDGFL2 | S153 | 0.14 | 0.10 | 0.22 | -0.19 | -0.04 | -0.31 | 0.016 |
| HDGFL2 | S144; S153 | 0.13 | 0.27 | 0.08 | -0.18 | 0.01 | -0.40 | 0.056 |
| HDGFL2 | S454 | 0.40 | 0.18 | 0.03 | -0.30 | -0.06 | -0.41 | 0.038 |
| HDGFL3 | S/T/Y | 0.24 | 0.01 | 0.06 | -0.17 | 0.11 | -0.32 | 0.188 |
| HDGFL3 | S/T/Y | 0.10 | 0.61 | 0.16 | -0.29 | -0.22 | -0.74 | 0.036 |
| HDLBP | S31 | -0.08 | -0.47 | 0.17 | 0.12 | 0.18 | -0.02 | 0.329 |
| HDLBP | S31 | -0.06 | -0.10 | 0.04 | 0.04 | 0.01 | 0.07 | 0.155 |
| HEATR1 | S1190 | -0.05 | -0.39 | -0.34 | 0.32 | 0.16 | 0.15 | 0.017 |
| HEATR1 | S1190 | -0.03 | -0.38 | -0.30 | 0.13 | 0.02 | 0.41 | 0.053 |
| HEATR1 | S1190 | -0.16 | -0.14 | -0.16 | 0.24 | -0.29 | 0.39 | 0.265 |
| HEATR3 | S15 | -0.34 | -0.14 | -0.01 | -0.06 | 0.09 | 0.37 | 0.134 |
| HEATR3 | S15 | -0.34 | -0.14 | -0.01 | -0.06 | 0.09 | 0.37 | 0.134 |
| HEATR3 | S15 | -0.15 | -0.08 | -0.02 | 0.17 | -0.21 | 0.23 | 0.363 |
| HEATR3 | S15 | 0.02 | -0.02 | -0.24 | 0.37 | -0.17 | -0.04 | 0.519 |
| HEATR5B | S1737 | 0.15 | -0.07 | 0.04 | 0.15 | -0.26 | -0.06 | 0.506 |
| HELLS | S832 | -0.07 | -0.19 | 0.10 | -0.09 | 0.00 | 0.21 | 0.49 |
| HELLS | S503 | -0.09 | 0.01 | 0.15 | 0.07 | -0.17 | 0.00 | 0.613 |
| HELZ | S/T/Y | -0.55 | -0.30 | 0.25 | 0.28 | 0.30 | -0.20 | 0.323 |
| HELZ | S1763 | 0.00 | 0.22 | -0.41 | 0.08 | -0.07 | 0.10 | 0.627 |
| HELZ | T1163 | -0.04 | 0.12 | -0.14 | 0.09 | 0.23 | -0.33 | 0.935 |
| HELZ | S1126 | 0.05 | 0.01 | 0.08 | -0.07 | 0.09 | -0.18 | 0.272 |
| HELZ2 | S338 | 0.26 | 0.12 | 0.26 | -0.01 | -0.35 | -0.45 | 0.026 |
| HERC1 | T2701 | -0.86 | -0.17 | 0.02 | 0.34 | 0.40 | -0.05 | 0.131 |
| HERC1 | S1510; S1521 | -0.15 | -0.11 | 0.19 | 0.19 | -0.28 | 0.08 | 0.91 |
| HERC2 | S2928 | -0.02 | 0.07 | -0.08 | 0.00 | 0.02 | 0.00 | 0.745 |
| HERC2 | S2928 | 0.02 | 0.14 | 0.02 | 0.06 | -0.08 | -0.19 | 0.193 |
| HERC2 | T272 | 0.03 | 0.31 | 0.24 | -0.50 | -0.03 | -0.20 | 0.051 |
| HERC2 | S1588 | 0.39 | 0.35 | 0.36 | -0.03 | -1.79 | -0.22 | 0.134 |
| HERPUD2 | T90 | 0.01 | -0.02 | -0.31 | 0.29 | -0.15 | 0.11 | 0.313 |
| HES1 | S/T | 0.16 | 0.18 | 0.06 | -0.12 | 0.44 | -1.23 | 0.426 |
| HEXIM2 | S76 | -0.15 | 0.13 | -0.26 | -0.06 | -0.04 | 0.30 | 0.389 |
| HEXIM2 | S29; T32 | 0.00 | -0.01 | 0.23 | -0.42 | -0.02 | 0.14 | 0.393 |
| HEXIM2 | S29 | 0.21 | 0.16 | 0.46 | -0.54 | -0.13 | -0.41 | 0.013 |
| HGH1 | T390 | -0.55 | -0.54 | 0.03 | 0.40 | 0.18 | 0.21 | 0.039 |
| HID1 | T584 | -0.08 | 0.10 | 0.01 | -0.01 | -0.02 | 0.01 | 0.739 |
| HID1 | S653 | 0.01 | 0.05 | 0.07 | 0.00 | -0.13 | -0.01 | 0.117 |
| HID1 | S653 | 0.26 | 0.00 | -0.02 | -0.05 | 0.19 | -0.50 | 0.412 |
| HID1 | S653 | 0.28 | 0.17 | 0.03 | -0.13 | -0.08 | -0.35 | 0.034 |
| HIPK1 | Y352 | -0.21 | -0.45 | -0.07 | -0.06 | 0.19 | 0.44 | 0.076 |

| | | | | | | | | |
|---|---|---|---|---|---|---|---|---|
| HIPK3 | Y/S/T | 0.10 | 0.43 | 0.22 | -0.40 | -0.18 | -0.37 | 0.008 |
| HIRA | T586 | -0.28 | -0.28 | -0.08 | -0.01 | 0.22 | 0.32 | 0.03 |
| HIRA | S/T | 0.17 | -0.15 | -0.32 | 0.18 | 0.06 | 0.00 | 0.321 |
| HIRA | S612 | 0.40 | 0.47 | 0.12 | -0.33 | -0.31 | -0.70 | 0.009 |
| HIRIP3 | S339; S/T | -0.26 | 0.06 | -0.07 | -0.09 | 0.08 | 0.23 | 0.274 |
| HIRIP3 | S550; S555 | 0.02 | -0.02 | 0.03 | -0.04 | -0.16 | 0.16 | 0.832 |
| HIRIP3 | S125 | -0.07 | -0.07 | 0.17 | -0.10 | -0.12 | 0.16 | 0.845 |
| HIRIP3 | S159; S160 | 0.03 | -0.16 | 0.17 | -0.08 | 0.11 | -0.10 | 0.773 |
| HIRIP3 | T84 | 0.00 | 0.09 | 0.08 | -0.08 | -0.26 | 0.14 | 0.364 |
| HIRIP3 | S223 | -0.07 | 0.03 | 0.20 | 0.13 | -0.13 | -0.21 | 0.39 |
| HIRIP3 | S125 | 0.17 | 0.03 | 0.03 | -0.21 | -0.14 | 0.09 | 0.192 |
| HIRIP3 | T84; S87 | 0.10 | 0.00 | 0.13 | -0.23 | -0.11 | 0.07 | 0.167 |
| HIRIP3 | S555 | 0.04 | 0.09 | 0.12 | -0.10 | -0.14 | -0.02 | 0.018 |
| HIRIP3 | S357 | 0.08 | 0.19 | 0.19 | -0.20 | -0.08 | -0.24 | 0.006 |
| HIRIP3 | S330; S332; S333 | 0.27 | 0.10 | 0.10 | -0.35 | -0.02 | -0.18 | 0.036 |
| HIRIP3 | S125 | 0.19 | 0.15 | 0.16 | 0.03 | -0.13 | -0.52 | 0.087 |
| HIRIP3 | S359; S363; S370 | 0.20 | 0.25 | 0.13 | -0.38 | 0.05 | -0.38 | 0.044 |
| HIRIP3 | S339; S/T | 0.21 | 0.27 | 0.15 | -0.27 | -0.19 | -0.28 | 5E-04 |
| HIRIP3 | S289; S291 | 0.29 | -0.06 | 0.40 | -0.28 | -0.25 | -0.24 | 0.029 |
| HIRIP3 | S227 | 0.31 | 0.22 | 0.18 | -0.41 | -0.09 | -0.38 | 0.008 |
| HIRIP3 | S339 | 0.23 | 0.23 | 0.28 | -0.31 | -0.11 | -0.49 | 0.008 |
| HIRIP3 | S227 | 0.39 | 0.20 | 0.19 | -0.47 | -0.16 | -0.35 | 0.006 |
| HIRIP3 | S289; S291 | 0.22 | 0.27 | 0.35 | -0.48 | -0.05 | -0.56 | 0.017 |
| HIRIP3 | S102 | 0.43 | 0.22 | 0.25 | -0.34 | -0.32 | -0.49 | 0.001 |
| HIRIP3 | S291 | 0.59 | -0.02 | 0.28 | -0.38 | -0.07 | -0.79 | 0.063 |
| HIRIP3 | S363; S370 | 0.28 | 0.30 | 0.29 | -0.41 | 0.06 | -0.89 | 0.061 |
| HIRIP3 | S291 | 0.61 | 0.03 | 0.23 | -0.39 | -0.03 | -0.92 | 0.076 |
| HIRIP3 | S/Y | 0.55 | 0.32 | 0.19 | -0.28 | -0.39 | -0.80 | 0.012 |
| HIRIP3 | S125 | 0.18 | 0.45 | 0.63 | -0.77 | -0.28 | -0.88 | 0.01 |
| HIVEP1 | S148 | -0.14 | -0.13 | -0.34 | 0.26 | 0.11 | 0.15 | 0.011 |
| HIVEP1 | S1180 | -0.17 | -0.50 | 0.17 | -0.09 | 0.14 | 0.30 | 0.283 |
| HIVEP1 | S779 | -0.12 | 0.06 | 0.10 | -0.04 | 0.04 | -0.07 | 0.673 |
| HIVEP1 | S1735 | 0.19 | -0.05 | -0.07 | -0.15 | 0.05 | 0.01 | 0.626 |
| HIVEP1 | S1874; S1884 | 0.09 | -0.03 | 0.08 | 0.13 | -0.14 | -0.15 | 0.362 |
| HIVEP1 | S1051 | -0.08 | 0.25 | 0.16 | -0.27 | -0.26 | 0.11 | 0.183 |
| HIVEP1 | S1884 | 0.31 | 0.20 | -0.05 | -0.38 | -0.14 | -0.05 | 0.079 |
| HIVEP1 | S2599 | 0.28 | 0.10 | 0.23 | -0.29 | 0.04 | -0.50 | 0.051 |
| HIVEP1 | S65 | 0.11 | 0.36 | 0.27 | -0.13 | -0.02 | -0.91 | 0.109 |
| HIVEP2 | S1443; S1446 | -0.12 | 0.02 | -0.05 | 0.05 | 0.10 | -0.01 | 0.125 |
| HIVEP2 | S1031 | 0.05 | 0.43 | 0.06 | -0.23 | -0.14 | -0.29 | 0.041 |
| HJURP | S | -0.19 | -0.42 | -0.17 | 0.08 | -0.25 | 0.67 | 0.204 |
| HJURP | S473 | -0.44 | -0.18 | 0.19 | 0.11 | -0.10 | 0.29 | 0.321 |
| HJURP | S | -0.17 | -0.19 | 0.07 | 0.11 | -0.01 | 0.16 | 0.138 |
| HLA-A | S343 | -0.29 | 0.30 | 0.41 | -0.70 | -0.17 | 0.15 | 0.311 |
| HLA-A | S359 | -0.01 | 0.27 | 0.25 | -0.48 | -0.23 | 0.06 | 0.098 |
| HLA-A | S359 | 0.37 | 0.14 | 0.27 | -0.51 | -0.19 | -0.28 | 0.007 |
| HLA-C | S351; S357; S360 | -0.27 | -0.16 | -0.13 | -0.13 | -0.07 | 0.58 | 0.256 |
| HLA-C | S351; S357; S360 | -0.27 | -0.16 | -0.13 | -0.13 | -0.07 | 0.58 | 0.256 |

| | | | | | | | | |
|---|---|---|---|---|---|---|---|---|
| **HLA-C** | **S360** | 0.15 | 0.05 | 0.44 | -0.19 | -0.41 | -0.21 | 0.024 |
| **HLA-C** | **S353** | -0.08 | 0.29 | 0.52 | -0.28 | -0.58 | -0.14 | 0.057 |
| **HLTF** | **S/T/Y** | 0.11 | 0.00 | 0.13 | -0.06 | -0.07 | -0.13 | 0.02 |
| **HMCES** | **S50** | -0.15 | -0.08 | 0.21 | 0.26 | 0.25 | -0.70 | 0.874 |
| **HMG20A** | **S105** | -0.07 | -0.29 | -0.41 | 0.20 | 0.02 | 0.39 | 0.035 |
| **HMG20A** | **S105** | 0.05 | -0.38 | -0.18 | -0.03 | 0.22 | 0.23 | 0.108 |
| **HMGA1** | **S49** | -1.42 | -0.90 | -1.77 | 0.72 | 0.62 | 0.69 | 0.001 |
| **HMGA1** | **S44** | -1.12 | -1.33 | -1.36 | 0.71 | 0.64 | 0.64 | 2E-05 |
| **HMGA1** | **S9** | -0.88 | -1.17 | -1.01 | 0.58 | 0.68 | 0.51 | 8E-05 |
| **HMGA1** | **S9** | -0.63 | -0.84 | -0.85 | 0.23 | 0.65 | 0.58 | 0.001 |
| **HMGA1** | **S36** | -0.35 | -0.45 | -0.53 | 0.29 | 0.24 | 0.47 | 9E-04 |
| **HMGA1** | **S36** | -0.38 | -0.44 | -0.46 | 0.34 | 0.19 | 0.44 | 6E-04 |
| **HMGA1** | **S99; S102; S103** | -0.29 | -0.37 | -0.12 | 0.37 | 0.01 | 0.25 | 0.02 |
| **HMGA1** | **S99; S102; S103** | -0.23 | -0.29 | -0.25 | 0.25 | 0.05 | 0.33 | 0.005 |
| **HMGA1** | **T53** | -0.20 | -0.31 | -0.22 | 0.21 | 0.03 | 0.37 | 0.013 |
| **HMGA1** | **S36** | -0.02 | -0.19 | -0.50 | 0.39 | 0.17 | -0.01 | 0.083 |
| **HMGA1** | **S99; S102; S103** | -0.12 | -0.26 | -0.25 | 0.13 | 0.09 | 0.32 | 0.01 |
| **HMGA1** | **S99; S102; S103** | -0.20 | -0.06 | 0.09 | 0.07 | -0.12 | 0.19 | 0.453 |
| **HMGA1** | **S102; S103** | 0.04 | 0.03 | -0.09 | 0.03 | -0.03 | 0.01 | 0.833 |
| **HMGA1** | **S102** | 0.05 | 0.00 | -0.04 | 0.03 | -0.02 | -0.03 | 0.71 |
| **HMGA1** | **S102; S103** | 0.06 | 0.02 | -0.02 | 0.02 | -0.05 | -0.04 | 0.275 |
| **HMGA1** | **S** | 0.11 | 0.10 | -0.11 | -0.01 | -0.03 | -0.07 | 0.423 |
| **HMGA1** | **S102; S103** | 0.03 | 0.02 | 0.24 | -0.22 | 0.04 | -0.15 | 0.118 |
| **HMGA1** | **S102; S103** | -0.25 | 0.17 | 0.36 | -0.22 | -0.43 | 0.19 | 0.396 |
| **HMGA1** | **S** | 0.50 | -0.14 | 0.38 | -0.42 | -0.15 | -0.46 | 0.055 |
| **HMGCR** | **S872** | 0.26 | 0.31 | -0.14 | -0.40 | 0.03 | -0.18 | 0.157 |
| **HMGCS1** | **S516** | 0.02 | 0.00 | -0.37 | -0.13 | 0.14 | 0.27 | 0.289 |
| **HMGCS1** | **S495** | 0.12 | 0.44 | -0.46 | -0.05 | 0.02 | -0.23 | 0.682 |
| **HMGN1** | **S7** | -0.12 | -0.34 | -0.28 | 0.01 | 0.23 | 0.36 | 0.022 |
| **HMGN1** | **S86** | -0.04 | -0.11 | 0.09 | -0.05 | 0.21 | -0.12 | 0.781 |
| **HMGN1** | **S86; S89** | -0.03 | 0.06 | -0.01 | -0.10 | 0.11 | -0.04 | 0.793 |
| **HMGN1** | **S8** | 0.00 | -0.06 | 0.10 | -0.07 | 0.12 | -0.10 | 0.748 |
| **HMGN3** | **S6** | -0.09 | -0.20 | -0.19 | 0.12 | 0.04 | 0.26 | 0.016 |
| **HMGN3** | **S6** | -0.20 | -0.12 | -0.12 | 0.05 | 0.05 | 0.28 | 0.03 |
| **HMGN3** | **S6** | 0.16 | 0.04 | -0.07 | -0.04 | -0.03 | -0.09 | 0.226 |
| **HMGN3** | **S78** | 0.11 | 0.12 | -0.09 | 0.04 | 0.15 | -0.42 | 0.541 |
| **HMGN3** | **S78** | 0.24 | 0.15 | -0.14 | -0.29 | 0.04 | -0.06 | 0.293 |
| **HMGN3** | **S78** | 0.13 | 0.15 | 0.03 | -0.23 | 0.19 | -0.36 | 0.239 |
| **HMGN3** | **S78** | 0.05 | 0.15 | 0.26 | -0.06 | -0.07 | -0.42 | 0.061 |
| **HMGN4** | **S80** | -1.14 | -1.31 | -1.34 | 0.56 | 0.71 | 0.71 | 2E-05 |
| **HMGXB4** | **S502; S512** | -0.14 | -0.18 | -0.48 | 0.18 | 0.18 | 0.28 | 0.013 |
| **HMGXB4** | **S/Y/T** | 0.07 | -0.16 | 0.03 | 0.07 | -0.27 | 0.21 | 0.877 |
| **HMGXB4** | **S502; S512** | 0.09 | -0.04 | -0.03 | -0.14 | 0.01 | 0.09 | 0.825 |
| **HMGXB4** | **S/T** | 0.06 | 0.19 | -0.23 | -0.01 | -0.03 | -0.02 | 0.845 |
| **HMGXB4** | **S101** | 0.16 | -0.16 | 0.10 | 0.05 | -0.18 | 0.00 | 0.557 |
| **HMGXB4** | **S497** | 0.10 | 0.13 | -0.03 | -0.30 | -0.13 | 0.17 | 0.358 |
| **HMGXB4** | **S497** | 0.26 | 0.08 | -0.03 | -0.11 | 0.06 | -0.31 | 0.18 |
| **HNRNPA0** | **S280** | 0.26 | 0.36 | 0.32 | -0.21 | -0.15 | -0.97 | 0.047 |

| | | | | | | | | |
|---|---|---|---|---|---|---|---|---|
| HNRNPA1 | T103 | -0.16 | 0.04 | 0.20 | 0.32 | -0.30 | -0.20 | 0.718 |
| HNRNPA1 | S4; S6 | 0.17 | 0.07 | -0.04 | -0.07 | -0.08 | -0.07 | 0.095 |
| HNRNPA1 | S6 | 0.07 | 0.03 | 0.13 | -0.10 | -0.01 | -0.14 | 0.029 |
| HNRNPA1 | S199 | 0.32 | 0.17 | 0.32 | -0.43 | -0.18 | -0.40 | 0.003 |
| HNRNPA1 | S362 | 0.30 | 0.34 | 0.33 | -0.48 | -0.17 | -0.63 | 0.005 |
| HNRNPA1 | S364 | 0.33 | 0.41 | 0.36 | -0.48 | -0.17 | -0.93 | 0.016 |
| HNRNPA1 | S6 | 0.35 | 0.54 | 0.35 | -0.47 | -0.22 | -1.24 | 0.028 |
| HNRNPA2B1 | S259 | -0.49 | -1.07 | -1.17 | 0.05 | -0.21 | 1.30 | 0.067 |
| HNRNPA2B1 | S259 | -0.05 | -0.24 | -0.29 | 0.01 | -0.16 | 0.55 | 0.227 |
| HNRNPA2B1 | S259 | -0.14 | -0.19 | -0.20 | 0.11 | -0.12 | 0.43 | 0.122 |
| HNRNPA2B1 | S225 | -0.05 | -0.16 | -0.17 | 0.09 | 0.04 | 0.21 | 0.021 |
| HNRNPA2B1 | S102 | -0.07 | -0.09 | -0.17 | 0.16 | 0.20 | -0.07 | 0.075 |
| HNRNPA2B1 | S198 | -0.04 | -0.19 | -0.10 | 0.05 | -0.01 | 0.26 | 0.085 |
| HNRNPA2B1 | S231 | -0.08 | -0.02 | 0.04 | -0.04 | 0.09 | 0.00 | 0.476 |
| HNRNPA2B1 | S212 | -0.18 | -0.07 | 0.21 | 0.09 | -0.14 | 0.04 | 0.946 |
| HNRNPA2B1 | S149 | 0.06 | 0.03 | 0.06 | -0.17 | 0.24 | -0.26 | 0.492 |
| HNRNPA2B1 | S225 | 0.23 | 0.21 | -0.21 | -0.11 | -0.33 | 0.11 | 0.393 |
| HNRNPA2B1 | S344 | 0.21 | 0.11 | 0.06 | -0.27 | 0.17 | -0.39 | 0.175 |
| HNRNPA2B1 | S344 | 0.14 | 0.30 | 0.02 | -0.16 | -0.06 | -0.33 | 0.04 |
| HNRNPA2B1 | S344 | 0.31 | 0.24 | -0.09 | -0.30 | 0.12 | -0.43 | 0.16 |
| HNRNPA2B1 | T159 | 0.42 | 0.53 | 0.24 | -0.33 | -0.31 | -1.21 | 0.031 |
| HNRNPA3 | S356; S358 | 0.16 | -0.13 | 0.07 | 0.04 | 0.05 | -0.22 | 0.57 |
| HNRNPA3 | S358 | 0.10 | 0.01 | 0.05 | -0.11 | -0.07 | 0.01 | 0.054 |
| HNRNPA3 | S116 | 0.07 | 0.07 | 0.08 | -0.06 | -0.13 | -0.06 | 0.002 |
| HNRNPAB | T33; T43 | -0.42 | -0.26 | -0.76 | 0.32 | -0.02 | 0.67 | 0.031 |
| HNRNPAB | T33 | -0.24 | -0.26 | -0.69 | 0.00 | -0.22 | 0.88 | 0.169 |
| HNRNPAB | T33 | -0.09 | -0.28 | -0.64 | 0.04 | -0.01 | 0.66 | 0.102 |
| HNRNPAB | T33 | -0.36 | -0.05 | -0.78 | 0.19 | -0.65 | 0.93 | 0.334 |
| HNRNPAB | T33; T43 | -0.02 | -0.28 | -0.64 | -0.01 | 0.08 | 0.58 | 0.112 |
| HNRNPAB | S242 | 0.05 | -0.20 | -0.15 | 0.04 | 0.13 | 0.11 | 0.075 |
| HNRNPAB | T33; T43 | -0.02 | -0.29 | 0.17 | 0.04 | 0.13 | -0.07 | 0.61 |
| HNRNPC | S253; S260 | 0.07 | -0.25 | -0.32 | 0.03 | -0.02 | 0.38 | 0.165 |
| HNRNPC | S233 | 0.11 | -0.27 | -0.21 | 0.00 | 0.23 | 0.08 | 0.172 |
| HNRNPC | S233 | -0.02 | -0.17 | -0.16 | 0.10 | 0.20 | 0.00 | 0.051 |
| HNRNPC | S233 | 0.03 | -0.15 | -0.21 | 0.09 | 0.25 | -0.07 | 0.166 |
| HNRNPC | S260 | -0.01 | -0.28 | -0.03 | -0.10 | -0.11 | 0.43 | 0.423 |
| HNRNPC | S253; S260 | -0.04 | -0.03 | -0.24 | 0.34 | 0.25 | -0.43 | 0.567 |
| HNRNPC | S233 | -0.03 | -0.09 | -0.09 | 0.00 | 0.02 | 0.18 | 0.094 |
| HNRNPC | S260 | -0.15 | -0.19 | 0.13 | -0.19 | 0.06 | 0.26 | 0.526 |
| HNRNPC | S299 | -0.02 | -0.03 | -0.07 | -0.05 | -0.02 | 0.18 | 0.358 |
| HNRNPC | S233 | 0.03 | -0.05 | -0.02 | -0.03 | 0.03 | 0.03 | 0.471 |
| HNRNPC | S233 | 0.02 | -0.01 | -0.01 | 0.05 | -0.18 | 0.12 | 0.991 |
| HNRNPC | S260 | 0.04 | -0.32 | 0.25 | 0.14 | -0.18 | -0.01 | 0.957 |
| HNRNPC | S299 | 0.07 | 0.02 | -0.05 | -0.10 | -0.02 | 0.08 | 0.682 |
| HNRNPC | S306 | 0.10 | -0.10 | 0.13 | -0.06 | -0.15 | 0.06 | 0.379 |
| HNRNPC | S241 | 0.08 | 0.02 | 0.09 | -0.13 | 0.00 | -0.07 | 0.037 |
| HNRNPC | S253; S260 | 0.04 | 0.11 | 0.04 | -0.13 | 0.01 | -0.09 | 0.054 |
| HNRNPC | S240; S241 | 0.19 | -0.06 | 0.09 | -0.13 | -0.17 | 0.05 | 0.182 |

| | | | | | | | | |
|---|---|---|---|---|---|---|---|---|
| **HNRNPC** | **S260** | 0.14 | 0.06 | 0.09 | -0.14 | -0.01 | -0.16 | 0.02 |
| **HNRNPC** | **S260** | 0.14 | 0.07 | 0.10 | -0.14 | -0.04 | -0.17 | 0.008 |
| **HNRNPC** | **S253; S260** | 0.18 | 0.01 | 0.15 | -0.25 | -0.07 | -0.05 | 0.046 |
| **HNRNPC** | **S** | 0.12 | 0.03 | 0.19 | -0.11 | -0.06 | -0.22 | 0.02 |
| **HNRNPC** | **S260** | 0.19 | 0.08 | 0.08 | -0.24 | -0.01 | -0.15 | 0.033 |
| **HNRNPC** | **S260** | 0.06 | 0.12 | 0.20 | -0.17 | 0.04 | -0.32 | 0.072 |
| **HNRNPC** | **S162** | 0.01 | -0.02 | 0.35 | -0.11 | 0.10 | -0.47 | 0.247 |
| **HNRNPC** | **S260** | 0.15 | 0.09 | 0.15 | -0.20 | 0.05 | -0.30 | 0.054 |
| **HNRNPC** | **S238; S241** | 0.21 | 0.11 | 0.10 | -0.20 | -0.08 | -0.18 | 0.005 |
| **HNRNPC** | **S260** | 0.16 | 0.16 | 0.11 | -0.20 | -0.01 | -0.28 | 0.021 |
| **HNRNPC** | **S260** | 0.20 | 0.12 | 0.10 | -0.18 | 0.01 | -0.33 | 0.041 |
| **HNRNPC** | **S260** | 0.09 | 0.13 | 0.21 | -0.27 | -0.04 | -0.19 | 0.014 |
| **HNRNPC** | **S233** | 0.12 | 0.09 | 0.24 | -0.22 | 0.01 | -0.31 | 0.035 |
| **HNRNPC** | **S260** | 0.13 | 0.05 | 0.29 | -0.21 | -0.08 | -0.24 | 0.018 |
| **HNRNPC** | **S260** | 0.12 | 0.15 | 0.19 | -0.22 | -0.03 | -0.28 | 0.012 |
| **HNRNPC** | **S238; S241** | 0.27 | 0.12 | 0.27 | -0.31 | -0.08 | -0.40 | 0.011 |
| **HNRNPC** | **S241** | 0.35 | 0.20 | 0.12 | -0.20 | -0.01 | -0.65 | 0.067 |
| **HNRNPC** | **S241** | 0.37 | 0.27 | 0.06 | -0.25 | 0.02 | -0.74 | 0.082 |
| **HNRNPC** | **S241** | 0.28 | 0.20 | 0.32 | -0.47 | 0.01 | -0.58 | 0.029 |
| **HNRNPCL1** | **S** | -0.02 | 0.19 | 0.07 | -0.28 | 0.14 | -0.16 | 0.267 |
| **HNRNPCL1** | **S240** | 0.33 | 0.01 | 0.12 | -0.15 | -0.05 | -0.37 | 0.061 |
| **HNRNPCL3** | **S196** | -0.30 | -0.27 | -0.36 | 0.23 | 0.23 | 0.31 | 1E-04 |
| **HNRNPD** | **S71; S** | -0.08 | -0.20 | -0.15 | -0.02 | -0.01 | 0.38 | 0.129 |
| **HNRNPD** | **T193** | -0.06 | -0.10 | 0.03 | 0.08 | 0.22 | -0.20 | 0.594 |
| **HNRNPD** | **S** | 0.02 | -0.02 | 0.11 | 0.00 | -0.12 | 0.00 | 0.246 |
| **HNRNPD** | **S** | 0.19 | 0.05 | 0.17 | -0.12 | -0.08 | -0.27 | 0.016 |
| **HNRNPD** | **S82; S83** | 0.30 | 0.05 | 0.13 | -0.13 | -0.11 | -0.32 | 0.027 |
| **HNRNPD** | **S** | 0.27 | 0.01 | 0.45 | -0.88 | 0.09 | -0.30 | 0.123 |
| **HNRNPD** | **S** | -0.03 | 0.30 | 0.54 | -0.19 | -0.29 | -0.63 | 0.039 |
| **HNRNPDL** | **S241** | -0.24 | -0.17 | -0.27 | 0.21 | 0.23 | 0.16 | 3E-04 |
| **HNRNPDL** | **S/T/Y** | -0.22 | -0.14 | -0.16 | 0.12 | 0.20 | 0.13 | 8E-04 |
| **HNRNPDL** | **T178** | -0.12 | 0.06 | 0.01 | 0.04 | 0.08 | -0.08 | 0.671 |
| **HNRNPDL** | **S/T/Y** | -0.03 | 0.30 | -0.31 | 0.18 | -0.07 | -0.16 | 0.991 |
| **HNRNPF** | **S/Y** | -0.99 | -1.07 | -1.09 | 0.71 | 0.63 | 0.46 | 3E-05 |
| **HNRNPH1** | **S63** | 0.12 | 0.05 | 0.12 | -0.09 | 0.04 | -0.27 | 0.086 |
| **HNRNPH1** | **S23** | 0.10 | 0.12 | 0.19 | 0.09 | -0.16 | -0.43 | 0.115 |
| **HNRNPH2** | **S90** | -0.25 | -0.21 | 0.05 | 0.02 | 0.10 | 0.23 | 0.081 |
| **HNRNPH2** | **S272** | -0.19 | -0.22 | 0.14 | -0.03 | 0.05 | 0.20 | 0.304 |
| **HNRNPH2** | **S23** | -0.09 | 0.01 | -0.16 | 0.14 | 0.11 | -0.04 | 0.118 |
| **HNRNPH2** | **S104** | -0.06 | -0.07 | 0.14 | -0.05 | -0.05 | 0.07 | 0.899 |
| **HNRNPH2** | **S104** | 0.04 | 0.09 | 0.20 | -0.18 | 0.03 | -0.23 | 0.064 |
| **HNRNPH3** | **S298** | -0.05 | 0.02 | -0.02 | 0.26 | -0.14 | -0.10 | 0.85 |
| **HNRNPK** | **S379** | -0.45 | -0.86 | -0.90 | 0.44 | 0.28 | 0.69 | 0.003 |
| **HNRNPK** | **S379** | -0.04 | -0.70 | -1.05 | 0.04 | 0.26 | 0.77 | 0.06 |
| **HNRNPK** | **S284** | -0.07 | -0.05 | -0.03 | 0.00 | -0.01 | 0.15 | 0.144 |
| **HNRNPK** | **S284** | -0.10 | -0.07 | 0.18 | -0.11 | 0.08 | -0.01 | 0.889 |
| **HNRNPK** | **S284** | 0.14 | 0.09 | 0.18 | -0.10 | -0.14 | -0.21 | 0.002 |
| **HNRNPK** | **S75** | -0.41 | 0.13 | 0.62 | 0.10 | -0.67 | -0.12 | 0.409 |

| | | | | | | | | |
|---|---|---|---|---|---|---|---|---|
| **HNRNPL** | **S101** | -0.13 | 0.06 | -0.10 | 0.11 | 0.06 | -0.01 | 0.171 |
| **HNRNPL** | **S543** | -0.22 | 0.18 | -0.09 | -0.10 | 0.19 | -0.01 | 0.662 |
| **HNRNPL** | **T487** | 0.03 | -0.14 | 0.18 | -0.09 | 0.20 | -0.23 | 0.719 |
| **HNRNPL** | **S185** | -0.05 | 0.00 | 0.17 | -0.14 | -0.08 | 0.08 | 0.384 |
| **HNRNPL** | **S291** | 0.37 | 0.14 | -0.29 | -0.26 | -0.16 | 0.08 | 0.442 |
| **HNRNPL** | **T/S/Y** | 0.37 | 0.14 | -0.29 | -0.26 | -0.16 | 0.08 | 0.442 |
| **HNRNPL** | **S589** | 0.52 | -0.03 | -0.25 | -0.40 | 0.03 | -0.04 | 0.458 |
| **HNRNPL** | **S52** | 0.17 | 0.07 | 0.10 | -0.09 | 0.00 | -0.31 | 0.061 |
| **HNRNPL** | **S298** | 0.57 | 0.34 | 0.14 | -0.81 | -0.14 | -0.59 | 0.021 |
| **HNRNPLL** | **S68** | 0.07 | -0.52 | 0.15 | 0.01 | -0.02 | 0.21 | 0.493 |
| **HNRNPM** | **S29** | -0.37 | -0.23 | 0.04 | 0.15 | 0.39 | -0.12 | 0.161 |
| **HNRNPM** | **S29** | -0.27 | -0.27 | 0.18 | 0.10 | 0.11 | 0.08 | 0.225 |
| **HNRNPR** | **S/T/Y** | -0.01 | 0.03 | 0.09 | 0.06 | -0.56 | 0.26 | 0.663 |
| **HNRNPU** | **S59** | -0.85 | -0.81 | -0.76 | 0.53 | 0.79 | 0.15 | 0.002 |
| **HNRNPU** | **S59** | -0.71 | -0.69 | -0.66 | 0.48 | 0.71 | 0.14 | 0.002 |
| **HNRNPU** | **S59** | -0.70 | -0.68 | -0.56 | 0.45 | 0.64 | 0.21 | 0.001 |
| **HNRNPU** | **S59** | -0.64 | -0.66 | -0.62 | 0.43 | 0.63 | 0.24 | 7E-04 |
| **HNRNPU** | **S59; S66** | -0.71 | -0.63 | -0.31 | 0.39 | 0.38 | 0.41 | 0.002 |
| **HNRNPU** | **S59; S66** | -0.49 | -0.02 | -0.40 | 0.12 | -0.03 | 0.56 | 0.086 |
| **HNRNPU** | **S271** | -0.13 | -0.30 | -0.32 | 0.16 | 0.06 | 0.40 | 0.017 |
| **HNRNPU** | **S271** | -0.10 | -0.20 | -0.33 | 0.11 | -0.02 | 0.41 | 0.057 |
| **HNRNPU** | **S271** | -0.07 | -0.10 | -0.33 | 0.07 | -0.07 | 0.40 | 0.136 |
| **HNRNPU** | **S267** | 0.04 | -0.14 | -0.21 | 0.08 | -0.06 | 0.24 | 0.175 |
| **HNRNPU** | **S271** | 0.03 | 0.07 | -0.21 | 0.03 | 0.13 | -0.07 | 0.568 |
| **HNRNPU** | **S271** | -0.06 | 0.04 | -0.07 | -0.15 | -0.01 | 0.21 | 0.683 |
| **HNRNPUL1** | **S194; T209** | -0.47 | -0.25 | -0.57 | 0.07 | 0.01 | 0.78 | 0.053 |
| **HNRNPUL1** | **S194** | -0.34 | -0.39 | -0.35 | 0.18 | 0.21 | 0.46 | 0.002 |
| **HNRNPUL1** | **S/Y** | -0.04 | -0.19 | -0.68 | 0.31 | 0.32 | 0.05 | 0.066 |
| **HNRNPUL1** | **S194; T209** | -0.54 | -0.14 | -0.10 | -0.11 | 0.00 | 0.63 | 0.178 |
| **HNRNPUL1** | **S/Y** | -0.29 | -0.16 | -0.10 | 0.26 | -0.08 | 0.28 | 0.062 |
| **HNRNPUL1** | **S194** | -0.29 | 0.09 | -0.12 | -0.05 | 0.06 | 0.25 | 0.234 |
| **HNRNPUL1** | **T/Y/S** | 0.01 | 0.13 | -0.39 | 0.31 | -0.03 | -0.12 | 0.539 |
| **HNRNPUL2** | **S597** | -0.90 | -1.05 | -1.04 | 0.72 | 0.47 | 0.54 | 6E-05 |
| **HNRNPUL2** | **S636** | -0.46 | -0.44 | -0.30 | 0.06 | 0.28 | 0.56 | 0.01 |
| **HNRNPUL2** | **S224** | -0.16 | -0.23 | -0.03 | 0.25 | 0.03 | 0.08 | 0.039 |
| **HNRNPUL2** | **S185** | 0.06 | -0.67 | 0.49 | -0.05 | -0.35 | 0.23 | 0.965 |
| **HNRNPUL2** | **S193** | 0.04 | -0.07 | 0.07 | 0.02 | -0.02 | -0.05 | 0.611 |
| **HNRNPUL2** | **S161** | 0.06 | -0.04 | 0.02 | -0.08 | 0.06 | -0.03 | 0.575 |
| **HNRNPUL2** | **T165; S168** | -0.20 | 0.05 | 0.24 | -0.15 | 0.04 | -0.02 | 0.62 |
| **HNRNPUL2** | **S228** | 0.09 | -0.08 | 0.14 | -0.14 | -0.13 | 0.09 | 0.311 |
| **HNRNPUL2** | **S161** | 0.08 | 0.01 | 0.09 | -0.01 | 0.00 | -0.19 | 0.139 |
| **HNRNPUL2** | **S655** | 0.11 | -0.02 | 0.19 | -0.16 | -0.22 | 0.06 | 0.131 |
| **HNRNPUL2** | **S161** | 0.16 | 0.06 | 0.14 | -0.13 | 0.09 | -0.40 | 0.141 |
| **HNRNPUL2** | **S193** | 0.34 | 0.22 | 0.34 | -0.36 | -0.05 | -0.87 | 0.041 |
| **HOMER3** | **S256** | -0.07 | -0.06 | -0.01 | 0.01 | -0.08 | 0.18 | 0.334 |
| **HOMEZ** | **S351** | -0.27 | -0.09 | -0.08 | 0.22 | 0.11 | 0.07 | 0.021 |
| **HOMEZ** | **T451** | -0.12 | -0.09 | -0.14 | 0.10 | 0.02 | 0.20 | 0.015 |
| **HOMEZ** | **T451** | -0.22 | 0.19 | -0.30 | 0.08 | -0.01 | 0.19 | 0.288 |

| | | | | | | | | |
|---|---|---|---|---|---|---|---|---|
| HOOK2 | T/S | -0.44 | -0.50 | 0.16 | 0.23 | 0.15 | 0.21 | 0.096 |
| HOOK2 | S163 | -0.19 | 0.55 | -0.41 | 0.06 | -0.32 | 0.08 | 0.9 |
| HOOK3 | S/T | -0.07 | 0.41 | 0.18 | -0.21 | 0.01 | -0.49 | 0.116 |
| HOXC10 | S210 | -0.11 | -0.11 | -0.09 | -0.02 | 0.29 | 0.00 | 0.123 |
| HOXC10 | S206; S210 | -0.12 | -0.13 | -0.01 | 0.12 | 0.05 | 0.08 | 0.022 |
| HOXC10 | S189 | 0.25 | 0.28 | -0.93 | -0.33 | 0.22 | 0.15 | 0.751 |
| HOXC10 | S189 | -0.06 | -0.05 | -0.05 | 0.04 | 0.00 | 0.11 | 0.034 |
| HOXC10 | T8 | -0.06 | -0.11 | 0.07 | 0.02 | -0.05 | 0.12 | 0.447 |
| HOXC10 | T/S | 0.03 | 0.03 | 0.02 | 0.01 | -0.07 | -0.02 | 0.082 |
| HOXC10 | T8 | 0.11 | 0.03 | 0.09 | -0.37 | 0.11 | -0.02 | 0.298 |
| HOXC10 | T244; S245 | 0.24 | 0.17 | 0.15 | -0.09 | -0.12 | -0.48 | 0.033 |
| HOXC9 | S159 | -0.30 | -0.37 | -0.12 | 0.16 | 0.09 | 0.39 | 0.015 |
| HOXC9 | S159 | -0.07 | -0.07 | -0.31 | -0.13 | 0.28 | 0.22 | 0.142 |
| HP1BP3 | S156 | -0.05 | -0.15 | -0.10 | 0.10 | 0.26 | -0.12 | 0.187 |
| HP1BP3 | S | -0.10 | -0.13 | -0.05 | -0.03 | 0.22 | 0.06 | 0.08 |
| HP1BP3 | S441; S442; S446 | -0.02 | -0.07 | -0.13 | 0.02 | 0.09 | 0.10 | 0.023 |
| HP1BP3 | S249 | -0.05 | -0.20 | 0.02 | -0.05 | 0.22 | 0.02 | 0.246 |
| HP1BP3 | S156 | -0.15 | -0.29 | 0.22 | -0.20 | 0.39 | -0.11 | 0.703 |
| HP1BP3 | T51 | 0.05 | 0.04 | 0.09 | -0.12 | 0.08 | -0.17 | 0.174 |
| HP1BP3 | S446; S | 0.29 | 0.05 | 0.04 | -0.08 | 0.13 | -0.57 | 0.242 |
| HP1BP3 | T51 | 0.15 | 0.16 | 0.18 | -0.24 | 0.08 | -0.45 | 0.076 |
| HS1BP3 | S194 | 0.00 | 0.16 | -0.03 | 0.00 | 0.03 | -0.19 | 0.329 |
| HSD17B8 | S60 | 0.14 | 0.12 | 0.18 | -0.14 | -0.12 | -0.22 | 9E-04 |
| HSF1 | S303 | -0.15 | -0.24 | -0.22 | 0.20 | 0.28 | 0.05 | 0.007 |
| HSF1 | S363 | -0.11 | -0.12 | -0.21 | 0.25 | 0.13 | 0.01 | 0.021 |
| HSF1 | S303; S307 | 0.10 | 0.26 | -0.04 | -0.33 | 0.13 | -0.21 | 0.21 |
| HSP90AA1 | S263 | 0.11 | 0.06 | -0.03 | -0.02 | -0.17 | 0.04 | 0.27 |
| HSP90AA1 | S263 | 0.22 | 0.15 | 0.04 | -0.14 | -0.09 | -0.23 | 0.013 |
| HSP90AA1 | S263 | 0.19 | 0.13 | 0.12 | 0.12 | -0.48 | -0.18 | 0.132 |
| HSP90AA1 | S263 | 0.25 | 0.24 | -0.03 | -0.16 | -0.03 | -0.35 | 0.064 |
| HSP90AB1 | S226 | 0.07 | -0.04 | -0.20 | 0.07 | -0.03 | 0.10 | 0.311 |
| HSP90AB2P | S177 | 0.17 | 0.15 | 0.06 | -0.10 | -0.01 | -0.32 | 0.052 |
| HSP90AB2P | S177 | 0.20 | 0.21 | 0.01 | -0.23 | -0.09 | -0.14 | 0.019 |
| HSP90AB2P | S177 | 0.32 | 0.25 | 0.23 | -0.32 | -0.01 | -0.75 | 0.043 |
| HSP90B1 | S64 | -0.28 | 0.22 | -0.01 | 0.09 | -0.28 | 0.17 | 0.943 |
| HSP90B1 | S306 | 0.25 | 0.20 | 0.31 | -0.42 | 0.11 | -0.76 | 0.075 |
| HSPA14 | S76 | -0.15 | -0.24 | 0.06 | 0.05 | 0.07 | 0.18 | 0.098 |
| HSPA1B | S254 | 0.13 | -0.05 | -0.08 | 0.06 | 0.09 | -0.17 | 0.953 |
| HSPA8 | S/T/Y | -1.50 | -1.00 | -1.67 | 1.00 | 1.03 | 0.18 | 0.003 |
| HSPB1 | S199 | -0.71 | -0.82 | -0.96 | 0.41 | 0.59 | 0.56 | 1E-04 |
| HSPB1 | S82 | -0.28 | -0.35 | -0.41 | 0.32 | 0.28 | 0.23 | 2E-04 |
| HSPB1 | S15 | -0.44 | -0.33 | -0.19 | 0.37 | 0.18 | 0.22 | 0.003 |
| HSPD1 | S/T | 0.14 | 0.28 | -0.23 | 0.07 | 0.07 | -0.44 | 0.509 |
| HSPD1 | S70 | -0.07 | 0.13 | 0.18 | 0.23 | -0.54 | -0.05 | 0.447 |
| HSPH1 | S809 | 0.23 | 0.25 | 0.21 | -0.25 | -0.07 | -0.52 | 0.017 |
| HTATSF1 | S481 | -0.40 | -0.43 | -0.51 | 0.53 | 0.18 | 0.30 | 0.002 |
| HTATSF1 | S485 | -0.04 | -0.05 | -0.75 | 0.15 | 0.31 | 0.16 | 0.116 |
| HTATSF1 | S481 | -0.22 | -0.13 | -0.32 | 0.38 | 0.13 | 0.06 | 0.02 |

| | | | | | | | | |
|---|---|---|---|---|---|---|---|---|
| HTATSF1 | S481 | -0.09 | -0.22 | -0.35 | 0.13 | 0.24 | 0.19 | 0.007 |
| HTATSF1 | S193 | -0.44 | 0.07 | -0.32 | 0.25 | 0.46 | -0.24 | 0.212 |
| HTATSF1 | S481 | -0.07 | -0.01 | -0.22 | 0.19 | 0.07 | 0.01 | 0.081 |
| HTATSF1 | S607 | 0.04 | -0.22 | -0.11 | 0.01 | 0.10 | 0.14 | 0.098 |
| HTATSF1 | S597; S607 | 0.03 | -0.05 | -0.25 | 0.15 | -0.09 | 0.16 | 0.244 |
| HTATSF1 | S597; S600 | 0.03 | -0.06 | -0.23 | 0.15 | -0.10 | 0.16 | 0.244 |
| HTATSF1 | S579 | 0.05 | 0.15 | -0.26 | 0.09 | -0.09 | 0.02 | 0.842 |
| HTATSF1 | S579 | 0.03 | 0.05 | -0.13 | 0.04 | -0.18 | 0.15 | 0.891 |
| HTATSF1 | S498 | 0.14 | 0.00 | -0.18 | 0.18 | -0.07 | -0.11 | 0.909 |
| HTATSF1 | S453 | 0.16 | -0.13 | 0.02 | 0.03 | -0.03 | -0.07 | 0.705 |
| HTATSF1 | S597; S600; S607 | 0.10 | 0.06 | -0.09 | 0.09 | -0.24 | 0.05 | 0.644 |
| HTATSF1 | S597; S600; S607 | 0.04 | 0.03 | 0.02 | 0.02 | -0.25 | 0.12 | 0.554 |
| HTATSF1 | S561 | 0.06 | 0.06 | 0.00 | -0.08 | 0.26 | -0.38 | 0.606 |
| HTATSF1 | S597; S600; S607 | 0.13 | -0.07 | 0.08 | -0.07 | -0.26 | 0.14 | 0.44 |
| HTATSF1 | S702 | 0.22 | 0.14 | -0.26 | 0.30 | 0.11 | -0.76 | 0.701 |
| HTATSF1 | S616; S624 | 0.14 | 0.15 | -0.08 | 0.06 | -0.03 | -0.28 | 0.294 |
| HTATSF1 | S713 | 0.16 | 0.02 | 0.11 | -0.25 | -0.08 | 0.00 | 0.066 |
| HTATSF1 | S624 | 0.14 | 0.08 | 0.10 | -0.30 | 0.18 | -0.27 | 0.204 |
| HTATSF1 | S | 0.16 | 0.12 | 0.08 | -0.03 | -0.04 | -0.36 | 0.076 |
| HTATSF1 | S713; S714; S721 | 0.10 | 0.19 | 0.10 | -0.17 | 0.02 | -0.30 | 0.047 |
| HTATSF1 | S676 | 0.22 | 0.19 | -0.01 | -0.04 | -0.01 | -0.44 | 0.124 |
| HTATSF1 | S452; S453 | 0.23 | 0.39 | -0.18 | -0.51 | 0.06 | -0.17 | 0.207 |
| HTATSF1 | S702; S714 | 0.20 | 0.19 | 0.08 | 0.04 | -0.01 | -0.66 | 0.188 |
| HTATSF1 | S453 | 0.28 | 0.18 | -0.02 | 0.02 | 0.09 | -0.77 | 0.272 |
| HTATSF1 | S642 | 0.19 | 0.22 | 0.10 | -0.10 | 0.02 | -0.57 | 0.104 |
| HTATSF1 | S452; S453 | 0.27 | 0.14 | 0.16 | -0.20 | 0.10 | -0.66 | 0.116 |
| HTATSF1 | S702 | 0.34 | 0.25 | 0.23 | -0.26 | -0.24 | -0.55 | 0.004 |
| HUWE1 | S1907 | -0.16 | -0.18 | -0.12 | 0.16 | -0.14 | 0.35 | 0.125 |
| HUWE1 | S3662 | -0.02 | -0.21 | -0.08 | -0.15 | -0.14 | 0.48 | 0.479 |
| HYCC2 | T | 0.05 | 0.04 | 0.10 | -0.12 | 0.05 | -0.15 | 0.109 |
| HYCC2 | S450 | 0.18 | 0.19 | -0.10 | -0.23 | 0.19 | -0.32 | 0.317 |
| IARS1 | S1047 | -0.10 | -0.15 | -0.29 | 0.17 | 0.09 | 0.21 | 0.007 |
| IBTK | S1045 | -0.04 | -0.03 | -0.06 | 0.17 | 0.01 | -0.06 | 0.293 |
| ICA1 | S/T | -0.04 | -0.02 | -0.10 | 0.06 | 0.30 | -0.26 | 0.613 |
| ICE1 | S892 | -0.29 | -0.01 | -0.30 | 0.09 | -0.07 | 0.44 | 0.117 |
| ICE1 | T/S | -0.25 | -0.10 | -0.21 | 0.17 | -0.12 | 0.39 | 0.097 |
| ICE1 | S1891; S1903 | -0.11 | -0.05 | -0.02 | 0.00 | 0.12 | 0.06 | 0.05 |
| ICE1 | S925 | -0.03 | -0.09 | -0.02 | 0.07 | -0.03 | 0.08 | 0.116 |
| ICE1 | S255 | -0.07 | -0.03 | -0.01 | 0.06 | -0.16 | 0.19 | 0.548 |
| ICE1 | S1692 | -0.12 | 0.06 | 0.00 | -0.05 | 0.12 | -0.02 | 0.688 |
| ICE1 | S516 | 0.08 | -0.11 | 0.15 | 0.04 | -0.07 | -0.12 | 0.384 |
| ICE1 | S1854 | -0.08 | 0.16 | 0.10 | 0.01 | -0.06 | -0.15 | 0.212 |
| ICE1 | S533 | -0.02 | 0.16 | 0.09 | -0.27 | 0.00 | 0.01 | 0.209 |
| ICE1 | S939 | 0.13 | 0.16 | 0.10 | -0.02 | -0.36 | -0.07 | 0.062 |
| ICE2 | S519 | -0.05 | -0.06 | 0.05 | 0.07 | -0.01 | 0.01 | 0.375 |
| ICE2 | S519 | -0.05 | -0.06 | 0.05 | 0.07 | -0.01 | 0.01 | 0.375 |
| ICE2 | S418 | -0.05 | 0.22 | -0.13 | 0.05 | -0.30 | 0.14 | 0.801 |
| IFIH1 | S301 | 0.20 | -0.22 | -0.40 | 0.27 | 0.04 | 0.00 | 0.287 |

| | | | | | | | | |
|---|---|---|---|---|---|---|---|---|
| IFIH1 | S301 | 0.10 | -0.03 | -0.28 | 0.17 | 0.02 | -0.02 | 0.358 |
| IFNAR1 | T494 | 0.30 | 0.33 | 0.39 | -0.40 | -0.46 | -0.48 | 2E-05 |
| IFT70B | S125; S132; S134 | -0.30 | 0.23 | 0.31 | -0.59 | 0.01 | 0.15 | 0.494 |
| IGF1R | T1366 | -0.18 | 0.12 | -0.40 | -0.06 | 0.05 | 0.36 | 0.233 |
| IGF2R | S2484 | -0.13 | -0.20 | -0.19 | 0.15 | -0.30 | 0.50 | 0.282 |
| IGF2R | S2347 | 0.02 | -0.12 | 0.14 | 0.21 | 0.02 | -0.35 | 0.779 |
| IGF2R | S2409 | 0.33 | -0.06 | 0.19 | -0.44 | -0.20 | 0.05 | 0.122 |
| IGF2R | S2409 | -0.06 | 0.32 | 0.39 | -0.58 | -0.31 | 0.02 | 0.084 |
| IGHMBP2 | S716 | -0.30 | 0.11 | -0.08 | 0.03 | -0.07 | 0.25 | 0.355 |
| IGHMBP2 | S716 | -0.38 | 0.59 | -0.57 | -0.21 | -0.54 | 0.59 | 0.897 |
| IGSF9 | S809 | 0.23 | -0.32 | 0.06 | -0.19 | 0.20 | -0.06 | 0.972 |
| IGSF9 | S797 | 0.00 | 0.19 | -0.02 | -0.37 | 0.15 | -0.02 | 0.467 |
| IK | S417 | -0.11 | -0.03 | -0.04 | -0.22 | 0.18 | 0.18 | 0.49 |
| IKBKB | S672 | 0.17 | -0.39 | -0.48 | 0.46 | 0.17 | -0.15 | 0.222 |
| IKBKB | S672 | 0.08 | 0.09 | -0.23 | 0.12 | 0.00 | -0.08 | 0.806 |
| IKBKG | S387 | -0.29 | -0.19 | -0.76 | 0.37 | 0.01 | 0.50 | 0.036 |
| IKBKG | S387 | -0.22 | 0.05 | -0.17 | 0.06 | 0.14 | 0.10 | 0.067 |
| IKZF2 | S78; S79 | -0.22 | -0.29 | -1.05 | 0.45 | -0.16 | 0.66 | 0.083 |
| IKZF4 | S491 | 0.21 | -0.03 | -0.02 | -0.27 | -0.09 | 0.15 | 0.44 |
| IL17RA | S629 | -0.20 | 0.26 | -0.05 | 0.41 | -0.17 | -0.41 | 0.835 |
| IL22RA1 | S420 | -0.81 | -0.43 | 0.40 | 0.94 | -0.26 | -0.69 | 0.673 |
| ILF2 | S/T | -0.75 | -0.75 | -0.86 | 0.23 | 0.33 | 0.87 | 0.003 |
| ILF2 | T/S | -0.09 | 0.03 | -0.41 | 0.23 | 0.22 | -0.07 | 0.157 |
| ILF3 | S860 | -0.04 | -0.36 | -0.67 | 0.09 | -0.11 | 0.71 | 0.13 |
| ILF3 | S73 | -0.62 | -0.18 | -0.18 | 0.63 | -0.13 | 0.16 | 0.108 |
| ILF3 | S860 | -0.05 | -0.27 | -0.61 | 0.11 | -0.07 | 0.60 | 0.116 |
| ILF3 | T592 | -0.40 | -0.28 | -0.06 | 0.03 | 0.14 | 0.42 | 0.044 |
| ILF3 | S56 | -0.26 | -0.19 | -0.21 | 0.23 | 0.14 | 0.20 | 3E-04 |
| ILF3 | S | 0.08 | -0.28 | -0.48 | 0.20 | 0.21 | 0.13 | 0.07 |
| ILF3 | S62 | -0.12 | -0.05 | -0.28 | 0.19 | -0.12 | 0.29 | 0.132 |
| ILF3 | T592 | -0.08 | -0.18 | -0.16 | 0.12 | 0.24 | 0.01 | 0.023 |
| ILF3 | S190 | 0.07 | -0.09 | -0.35 | 0.30 | 0.25 | -0.31 | 0.423 |
| ILF3 | S62 | -0.14 | -0.05 | -0.14 | 0.13 | -0.04 | 0.20 | 0.057 |
| ILF3 | S62; S | 0.04 | -0.20 | -0.14 | 0.20 | -0.10 | 0.16 | 0.187 |
| ILF3 | S62; S | 0.12 | -0.26 | -0.11 | -0.03 | 0.12 | 0.11 | 0.291 |
| ILF3 | S56; S62 | 0.02 | -0.11 | -0.08 | -0.01 | 0.13 | 0.04 | 0.126 |
| ILF3 | S62 | -0.18 | 0.02 | 0.05 | -0.11 | -0.09 | 0.27 | 0.68 |
| ILF3 | S382 | 0.07 | 0.02 | -0.14 | 0.07 | 0.02 | -0.06 | 0.725 |
| ILF3 | S382 | 0.08 | 0.01 | -0.13 | 0.06 | 0.05 | -0.07 | 0.755 |
| ILF3 | S476 | 0.03 | -0.18 | 0.21 | -0.14 | 0.17 | -0.14 | 0.739 |
| ILF3 | S810 | -0.08 | -0.09 | 0.23 | 0.04 | 0.10 | -0.25 | 0.72 |
| ILF3 | T/S | -0.34 | 0.31 | 0.14 | 0.13 | 0.10 | -0.51 | 0.674 |
| ILF3 | T368 | -0.17 | 0.45 | -0.17 | 0.13 | 0.06 | -0.48 | 0.655 |
| ILF3 | S382 | 0.13 | 0.12 | -0.07 | 0.01 | 0.09 | -0.34 | 0.392 |
| ILF3 | S382 | 0.21 | 0.09 | -0.09 | 0.02 | 0.08 | -0.37 | 0.396 |
| ILF3 | S/Y | 0.32 | -0.26 | -0.06 | -0.35 | -0.84 | 0.69 | 0.754 |
| ILF3 | Y/S | 0.22 | 0.21 | -0.10 | -0.32 | -0.68 | 0.39 | 0.396 |
| ILF3 | S783 | 0.02 | 0.01 | 0.30 | 0.09 | -1.07 | 0.28 | 0.477 |

| | | | | | | | | |
|---|---|---|---|---|---|---|---|---|
| ILF3 | S863 | 0.11 | 0.18 | 0.14 | 0.88 | -1.50 | -0.22 | 0.571 |
| ILF3 | S783 | 0.10 | 0.09 | 0.37 | -0.05 | -0.67 | -0.03 | 0.131 |
| ILF3 | T592 | 0.27 | 0.13 | 0.11 | -0.04 | 0.45 | -1.81 | 0.411 |
| IMMT | S103 | -0.04 | 0.10 | -0.35 | 0.09 | 0.19 | -0.05 | 0.313 |
| IMMT | S186 | -0.07 | -0.10 | 0.05 | 0.13 | 0.03 | -0.05 | 0.302 |
| IMMT | S388 | 0.06 | 0.11 | -0.28 | -0.21 | 0.33 | -0.10 | 0.857 |
| IMMT | S447 | 0.20 | 0.04 | -0.05 | 0.03 | -0.18 | -0.07 | 0.238 |
| IMMT | S388 | 0.00 | 0.19 | 0.05 | -0.27 | 0.21 | -0.26 | 0.319 |
| IMMT | S/T/Y | 0.43 | -0.01 | -0.01 | -0.35 | -0.19 | 0.01 | 0.154 |
| IMP3 | S153 | -0.31 | -0.30 | -0.20 | 0.23 | 0.24 | 0.20 | 2E-04 |
| IMP4 | S56 | -0.12 | 0.12 | -0.32 | 0.22 | 0.01 | 0.04 | 0.241 |
| IMP4 | S56 | -0.05 | 0.02 | -0.19 | 0.27 | -0.23 | 0.12 | 0.473 |
| INCENP | S119 | -0.48 | 0.02 | -0.76 | 0.31 | 0.64 | -0.18 | 0.115 |
| INCENP | S119 | -0.42 | -0.08 | -0.64 | -0.27 | 0.58 | 0.42 | 0.112 |
| INCENP | T292 | -0.13 | -0.21 | -0.78 | -0.10 | 0.50 | 0.36 | 0.084 |
| INCENP | T478; S481 | -0.29 | -0.45 | -0.14 | 0.02 | 0.29 | 0.39 | 0.02 |
| INCENP | S197 | -0.13 | -0.33 | 0.09 | -0.07 | 0.12 | 0.25 | 0.222 |
| INCENP | S263; S269 | -0.01 | -0.21 | -0.17 | 0.02 | -0.29 | 0.52 | 0.433 |
| INCENP | S197; T219 | -0.11 | -0.17 | -0.03 | 0.24 | -0.05 | 0.08 | 0.112 |
| INCENP | S828; S831; T832 | -0.15 | -0.06 | -0.04 | 0.01 | 0.10 | 0.12 | 0.03 |
| INCENP | S828; S831; T832 | -0.11 | -0.01 | -0.11 | -0.02 | 0.18 | 0.06 | 0.086 |
| INCENP | S72 | -0.21 | 0.01 | -0.02 | 0.03 | 0.06 | 0.12 | 0.123 |
| INCENP | S306 | 0.12 | -0.37 | 0.03 | -0.11 | 0.09 | 0.17 | 0.504 |
| INCENP | S72 | 0.03 | -0.07 | -0.15 | -0.11 | 0.08 | 0.18 | 0.314 |
| INCENP | S93 | 0.18 | -0.12 | -0.20 | -0.06 | 0.23 | -0.07 | 0.634 |
| INCENP | S275 | 0.21 | -0.22 | -0.09 | 0.00 | 0.02 | 0.04 | 0.712 |
| INCENP | S197 | -0.10 | 0.02 | -0.02 | -0.13 | 0.27 | -0.09 | 0.741 |
| INCENP | S197; T204 | -0.07 | 0.02 | 0.00 | -0.04 | 0.04 | 0.05 | 0.477 |
| INCENP | S72 | 0.08 | 0.00 | -0.06 | 0.07 | 0.04 | -0.15 | 0.793 |
| INCENP | S481 | 0.07 | -0.01 | 0.00 | -0.05 | 0.03 | -0.04 | 0.311 |
| INCENP | S263 | -0.08 | 0.12 | 0.19 | -0.51 | -0.05 | 0.21 | 0.443 |
| INCENP | S263 | 0.14 | 0.09 | 0.12 | -0.24 | 0.05 | -0.21 | 0.058 |
| INCENP | S311; S312 | 0.16 | 0.28 | -0.02 | -0.01 | -0.04 | -0.47 | 0.141 |
| INCENP | T219 | 0.22 | 0.20 | 0.20 | -0.11 | -0.10 | -0.56 | 0.038 |
| INCENP | S306; S311 | 0.32 | 0.27 | 0.27 | -0.59 | -0.18 | -0.34 | 0.006 |
| INF2 | S1204 | 0.06 | -0.09 | 0.07 | -0.03 | -0.02 | 0.00 | 0.585 |
| INF2 | T/S | -0.20 | 0.12 | 0.21 | -0.10 | -0.07 | 0.00 | 0.467 |
| INF2 | S1188 | 0.10 | 0.15 | 0.05 | -0.11 | -0.10 | -0.12 | 0.002 |
| INF2 | S/T | -0.02 | 0.50 | 0.12 | -0.40 | -0.01 | -0.39 | 0.083 |
| ING4 | S150 | 0.12 | 0.12 | -0.17 | -0.07 | -0.02 | 0.00 | 0.602 |
| ING5 | S148; T152 | 0.05 | -0.05 | -0.11 | -0.01 | -0.01 | 0.12 | 0.33 |
| ING5 | S118 | 0.20 | 0.06 | 0.03 | -0.07 | -0.01 | -0.24 | 0.081 |
| ING5 | T152 | 0.32 | 0.16 | 0.18 | -0.19 | -0.22 | -0.39 | 0.003 |
| ING5 | S118 | 0.21 | 0.19 | 0.25 | -0.27 | 0.06 | -0.63 | 0.067 |
| ING5 | S118 | 0.37 | 0.17 | 0.17 | -0.62 | 0.14 | -0.49 | 0.083 |
| INO80B | S99 | -0.19 | 0.01 | -0.22 | -0.07 | 0.15 | 0.26 | 0.107 |
| INO80B | S130 | -0.02 | 0.00 | -0.27 | 0.06 | -0.05 | 0.24 | 0.222 |
| INO80B | S99 | 0.06 | -0.06 | -0.13 | -0.06 | -0.06 | 0.23 | 0.494 |

| | | | | | | | | |
|---|---|---|---|---|---|---|---|---|
| **INO80B** | **S99** | 0.09 | -0.08 | 0.07 | -0.10 | -0.07 | 0.08 | 0.49 |
| **INO80B** | **S351** | 0.14 | 0.14 | -0.03 | -0.10 | -0.06 | -0.11 | 0.037 |
| **INO80B** | **S63** | 0.21 | 0.02 | 0.09 | -0.22 | 0.20 | -0.40 | 0.263 |
| **INO80B** | **S63** | 0.17 | 0.27 | 0.14 | -0.19 | -0.04 | -0.49 | 0.036 |
| **INO80C** | **S26** | 0.07 | 0.11 | -0.12 | 0.14 | -0.83 | 0.37 | 0.754 |
| **INO80C** | **S26** | 0.13 | 0.17 | -0.06 | 0.01 | -0.17 | -0.12 | 0.134 |
| **INO80E** | **S227** | -0.22 | -0.21 | -0.04 | 0.07 | -0.04 | 0.36 | 0.1 |
| **INO80E** | **S223; S227** | -0.09 | -0.04 | -0.16 | -0.13 | -0.18 | 0.48 | 0.519 |
| **INPP4B** | **S487** | -0.12 | -0.07 | -0.05 | 0.02 | 0.14 | 0.06 | 0.02 |
| **INPP5D** | **S1111** | -0.23 | 0.32 | 0.31 | -0.16 | -0.04 | -0.35 | 0.193 |
| **INTS1** | **S283; S288; T290** | -0.43 | -0.39 | -0.17 | 0.39 | -0.09 | 0.44 | 0.038 |
| **INTS1** | **S1142** | -0.36 | -0.30 | -0.15 | 0.22 | 0.22 | 0.23 | 0.001 |
| **INTS1** | **S1318** | -0.05 | -0.21 | -0.34 | 0.13 | 0.32 | 0.06 | 0.031 |
| **INTS1** | **S1395** | 0.06 | -0.71 | 0.03 | 0.04 | 0.08 | 0.31 | 0.259 |
| **INTS1** | **S307** | 0.09 | -0.26 | 0.05 | -0.03 | 0.03 | 0.09 | 0.58 |
| **INTS1** | **S307** | -0.13 | 0.11 | -0.04 | 0.21 | 0.15 | -0.38 | 0.952 |
| **INTS1** | **S1327** | 0.22 | 0.03 | 0.11 | -0.18 | 0.02 | -0.24 | 0.062 |
| **INTS1** | **T83** | 0.24 | 0.02 | 0.29 | -0.22 | -0.03 | -0.43 | 0.044 |
| **INTS10** | **S381; S382** | -0.20 | -0.09 | -0.14 | 0.37 | -0.30 | 0.24 | 0.308 |
| **INTS10** | **S231** | 0.08 | -0.34 | 0.12 | 0.15 | -0.04 | -0.02 | 0.669 |
| **INTS10** | **S/T** | -0.09 | 0.05 | 0.02 | 0.27 | -0.22 | -0.07 | 0.997 |
| **INTS10** | **S231** | -0.16 | 0.21 | -0.07 | 0.04 | 0.07 | -0.13 | 0.985 |
| **INTS10** | **S231** | 0.12 | -0.03 | 0.09 | -0.07 | 0.09 | -0.23 | 0.267 |
| **INTS10** | **S231** | 0.16 | 0.20 | 0.13 | -0.31 | 0.04 | -0.30 | 0.041 |
| **INTS10** | **S231** | 0.26 | 0.12 | 0.33 | -0.10 | -0.10 | -0.74 | 0.069 |
| **INTS10** | **S231** | 0.23 | 0.26 | 0.30 | -0.34 | -0.05 | -0.64 | 0.023 |
| **INTS10** | **S231** | 0.37 | 0.33 | 0.10 | -0.18 | -0.12 | -0.81 | 0.055 |
| **INTS12** | **S378** | -0.01 | -0.16 | -0.18 | 0.09 | 0.10 | 0.12 | 0.015 |
| **INTS12** | **S/T** | 0.10 | -0.25 | 0.06 | -0.12 | 0.31 | -0.17 | 0.85 |
| **INTS12** | **S/T** | -0.22 | 0.24 | 0.14 | -0.47 | 0.20 | -0.02 | 0.57 |
| **INTS13** | **S626** | 0.11 | 0.01 | -0.24 | -0.03 | -0.09 | 0.19 | 0.655 |
| **INTS13** | **S626** | 0.26 | -0.19 | -0.21 | 0.00 | -0.18 | 0.22 | 0.772 |
| **INTS13** | **S626** | 0.06 | 0.16 | 0.09 | -0.09 | -0.09 | -0.17 | 0.006 |
| **INTS13** | **S626** | 0.25 | 0.09 | 0.05 | -0.22 | -0.30 | 0.06 | 0.09 |
| **INTS3** | **S502** | -0.18 | -0.20 | -0.27 | 0.26 | 0.25 | 0.04 | 0.007 |
| **INTS3** | **S502** | -0.05 | 0.04 | -0.32 | -0.05 | 0.05 | 0.27 | 0.231 |
| **INTS3** | **S1040; S1042** | 0.08 | -0.15 | 0.16 | -0.12 | 0.04 | -0.03 | 0.573 |
| **INTS3** | **S537** | 0.13 | 0.13 | -0.01 | -0.26 | -0.20 | 0.15 | 0.24 |
| **INTS3** | **S537** | 0.09 | 0.20 | -0.01 | -0.22 | -0.12 | 0.02 | 0.095 |
| **INTS3** | **S1042** | 0.34 | 0.23 | 0.35 | -0.52 | 0.03 | -0.81 | 0.04 |
| **INTS5** | **S279** | 0.03 | -0.31 | 0.25 | -0.13 | -0.21 | 0.27 | 0.942 |
| **INTS5** | **S1010** | 0.10 | 0.12 | 0.11 | -0.21 | 0.01 | -0.16 | 0.026 |
| **INTS7** | **S809** | -0.33 | -0.06 | -0.04 | -0.03 | 0.22 | 0.17 | 0.097 |
| **IPO5** | **S/Y/T** | -0.31 | -0.22 | 0.20 | 0.22 | 0.09 | -0.06 | 0.337 |
| **IPO5** | **S827** | -0.12 | -0.24 | 0.22 | 0.06 | 0.02 | 0.03 | 0.577 |
| **IPO5** | **S827** | -0.10 | 0.01 | -0.01 | -0.14 | -0.08 | 0.28 | 0.698 |
| **IPO5** | **S827** | 0.05 | -0.19 | 0.24 | -0.07 | -0.11 | 0.04 | 0.573 |
| **IQCE** | **S/T/Y** | -0.57 | 0.31 | 0.07 | -0.02 | -0.37 | 0.36 | 0.885 |

| | | | | | | | | |
|---|---|---|---|---|---|---|---|---|
| IQSEC1 | S105 | -0.05 | -0.19 | -0.14 | 0.38 | -0.27 | 0.16 | 0.33 |
| IRF2BP1 | S384 | -0.08 | -0.18 | -0.32 | 0.22 | 0.15 | 0.13 | 0.008 |
| IRF2BP1 | S453 | -0.24 | 0.15 | 0.02 | 0.02 | 0.13 | -0.11 | 0.783 |
| IRF2BP2 | S360 | 0.17 | 0.24 | -0.18 | -0.12 | -0.15 | -0.01 | 0.281 |
| IRF2BP2 | S360 | 0.26 | 0.25 | -0.18 | -0.24 | -0.19 | 0.01 | 0.201 |
| IRF2BP2 | S175 | 0.25 | 0.26 | 0.04 | -0.19 | -0.23 | -0.23 | 0.006 |
| IRF2BP2 | S457; S460 | 0.18 | 0.10 | 0.31 | -0.26 | 0.02 | -0.51 | 0.051 |
| IRF2BPL | S215 | -0.06 | -0.22 | -0.16 | 0.08 | -0.09 | 0.37 | 0.129 |
| IRF2BPL | S69 | 0.23 | -0.08 | -0.33 | 0.22 | -0.55 | 0.30 | 0.883 |
| IRF2BPL | S547 | 0.10 | 0.17 | -0.12 | -0.05 | -0.11 | -0.01 | 0.316 |
| IRF2BPL | S662 | 0.42 | 0.28 | 0.28 | -0.73 | -0.34 | -0.25 | 0.008 |
| IRF3 | S175 | 0.24 | -0.17 | -0.32 | 0.26 | 0.10 | -0.21 | 0.583 |
| IRF6 | S47 | 0.22 | 0.10 | -0.05 | -0.22 | -0.11 | 0.02 | 0.128 |
| IRX5 | S248 | 0.01 | 0.11 | 0.03 | -0.08 | 0.03 | -0.10 | 0.12 |
| IRX5 | T237 | 0.14 | 0.29 | 0.08 | -0.74 | 0.21 | -0.20 | 0.215 |
| IRX5 | S274 | 0.07 | 0.23 | 0.36 | -0.35 | 0.00 | -0.50 | 0.041 |
| ISL2 | S157 | -0.28 | -0.12 | 0.00 | 0.18 | -0.05 | 0.21 | 0.103 |
| ISY1 | S222; S227 | -0.45 | -0.37 | -0.39 | 0.32 | 0.14 | 0.46 | 0.002 |
| ISY1 | Y215; S227 | -0.37 | -0.25 | -0.38 | 0.33 | 0.07 | 0.39 | 0.005 |
| ISY1 | S222 | 0.25 | 0.11 | 0.06 | -0.14 | -0.18 | -0.15 | 0.008 |
| ISY1 | S222 | 0.27 | 0.13 | 0.06 | -0.16 | -0.23 | -0.14 | 0.009 |
| ITGB1BP1 | S41 | 0.27 | 0.03 | -0.04 | -0.18 | -0.10 | -0.03 | 0.135 |
| ITGB3BP | S71 | -0.06 | -0.10 | -0.14 | 0.05 | 0.09 | 0.15 | 0.005 |
| ITGB4 | T1530 | -0.22 | 0.01 | 0.17 | -0.08 | 0.22 | -0.14 | 0.943 |
| ITPKA | S95 | 0.01 | 0.06 | -0.33 | 0.03 | 0.04 | 0.14 | 0.287 |
| ITPKA | T/S | -0.06 | 0.19 | 0.03 | -0.08 | -0.40 | 0.23 | 0.531 |
| ITPKB | S120 | -0.45 | 0.13 | -0.20 | 0.05 | -0.01 | 0.35 | 0.204 |
| ITPKB | S65 | 0.02 | 0.03 | -0.05 | 0.07 | -0.05 | -0.02 | 0.966 |
| ITPKB | T33 | -0.05 | 0.25 | 0.30 | -0.41 | 0.10 | -0.34 | 0.117 |
| ITPR1 | S1598 | 0.11 | 0.02 | -0.05 | -0.09 | -0.03 | 0.02 | 0.308 |
| ITPR1 | S1577 | 0.09 | -0.05 | 0.14 | 0.26 | -0.16 | -0.37 | 0.479 |
| ITPR2 | S1872 | -0.17 | 0.01 | -0.05 | 0.00 | -0.12 | 0.28 | 0.381 |
| ITPR2 | S1160 | -0.06 | -0.09 | -0.01 | 0.09 | 0.00 | 0.07 | 0.051 |
| ITPR3 | S934; S942 | -0.10 | -0.15 | -0.27 | 0.09 | 0.17 | 0.20 | 0.006 |
| ITPR3 | S934 | 0.03 | -0.16 | -0.21 | 0.30 | -0.23 | 0.19 | 0.318 |
| ITPR3 | S940 | -0.14 | -0.05 | -0.14 | -0.02 | -0.17 | 0.43 | 0.363 |
| ITPR3 | S1832; S1838 | -0.01 | -0.08 | -0.01 | 0.16 | 0.18 | -0.28 | 0.743 |
| ITPR3 | S1832 | 0.13 | -0.13 | -0.05 | -0.03 | 0.26 | -0.24 | 0.951 |
| ITPRID2 | S668 | 0.06 | -0.06 | -0.07 | 0.15 | -0.12 | 0.02 | 0.682 |
| ITPRID2 | S1165; T1168; S/T | -0.02 | 0.07 | -0.06 | -0.15 | -0.05 | 0.18 | 0.988 |
| ITPRID2 | T1156 | 0.05 | 0.02 | -0.04 | -0.05 | -0.08 | 0.10 | 0.73 |
| ITPRID2 | S318 | 0.07 | -0.08 | 0.05 | 0.09 | -0.04 | -0.09 | 0.743 |
| ITPRID2 | S380 | 0.26 | -0.09 | -0.10 | 0.18 | -0.12 | -0.18 | 0.708 |
| ITPRID2 | S380 | -0.02 | 0.18 | -0.07 | 0.14 | -0.02 | -0.25 | 0.629 |
| ITPRID2 | S593 | 0.06 | 0.00 | 0.05 | 0.01 | -0.04 | -0.09 | 0.092 |
| ITPRID2 | T87 | -0.01 | -0.07 | 0.18 | 0.09 | -0.07 | -0.15 | 0.493 |
| ITPRID2 | S/T | 0.02 | 0.08 | 0.01 | 0.00 | -0.04 | -0.09 | 0.076 |
| ITPRID2 | S92 | 0.07 | 0.02 | 0.03 | 0.00 | -0.02 | -0.12 | 0.1 |

| | | | | | | | | |
|---|---|---|---|---|---|---|---|---|
| **ITPRID2** | **S92** | 0.19 | -0.10 | 0.09 | -0.09 | -0.22 | 0.09 | 0.339 |
| **ITPRID2** | **S739; S746** | 0.03 | 0.05 | 0.12 | -0.09 | -0.06 | -0.07 | 0.009 |
| **ITPRID2** | **S737** | 0.20 | 0.05 | -0.02 | 0.09 | -0.05 | -0.32 | 0.291 |
| **ITPRID2** | **S380** | 0.01 | 0.18 | 0.05 | -0.05 | -0.20 | -0.03 | 0.086 |
| **ITPRID2** | **S1165; T1168** | -0.07 | 0.17 | 0.21 | -0.04 | -0.13 | -0.19 | 0.081 |
| **ITPRID2** | **S593** | 0.08 | 0.19 | 0.07 | -0.04 | -0.02 | -0.33 | 0.082 |
| **ITPRID2** | **T776** | 0.06 | 0.06 | 0.23 | -0.11 | -0.03 | -0.26 | 0.046 |
| **ITPRID2** | **S466** | 0.08 | 0.29 | -0.04 | 0.00 | 0.11 | -0.56 | 0.32 |
| **ITPRID2** | **S466** | 0.08 | 0.29 | -0.04 | 0.00 | 0.11 | -0.56 | 0.32 |
| **ITPRID2** | **S803** | 0.07 | 0.26 | 0.14 | -0.26 | 0.16 | -0.53 | 0.153 |
| **ITPRID2** | **T87** | 0.20 | -0.15 | 0.45 | -0.07 | -0.17 | -0.43 | 0.126 |
| **ITPRID2** | **S1134** | 0.25 | 0.01 | 0.31 | -0.21 | -0.10 | -0.37 | 0.025 |
| **ITPRID2** | **S593** | 0.29 | 0.00 | 0.39 | -0.35 | -0.15 | -0.34 | 0.02 |
| **ITPRID2** | **S156** | 0.38 | 0.26 | 0.15 | -0.24 | 0.15 | -1.18 | 0.163 |
| **ITPRID2** | **S156; S159** | 0.18 | 0.50 | 0.28 | -0.55 | -0.16 | -0.61 | 0.011 |
| **ITSN2** | **S884; S889** | -0.08 | 0.16 | 0.09 | -0.02 | 0.20 | -0.42 | 0.523 |
| **ITSN2** | **S889** | 0.05 | 0.03 | 0.30 | -0.10 | 0.18 | -0.61 | 0.288 |
| **IWS1** | **S313; S315** | 0.02 | -0.29 | -0.50 | 0.13 | -0.17 | 0.56 | 0.171 |
| **IWS1** | **S172; S185** | -0.17 | 0.01 | -0.50 | -0.07 | 0.18 | 0.39 | 0.124 |
| **IWS1** | **S183; S185** | -0.24 | -0.01 | -0.37 | 0.04 | 0.16 | 0.31 | 0.044 |
| **IWS1** | **T67; S69; S84** | -0.22 | -0.15 | -0.02 | 0.07 | -0.09 | 0.34 | 0.169 |
| **IWS1** | **T67; S69; S82** | -0.18 | -0.10 | -0.02 | 0.05 | -0.06 | 0.27 | 0.156 |
| **IWS1** | **S** | 0.06 | -0.12 | -0.22 | 0.12 | 0.31 | -0.22 | 0.402 |
| **IWS1** | **T67; S84** | 0.01 | -0.27 | -0.02 | -0.10 | -0.15 | 0.42 | 0.508 |
| **IWS1** | **S131; S133; S146** | -0.03 | -0.18 | -0.02 | -0.12 | -0.10 | 0.38 | 0.493 |
| **IWS1** | **S398; S400** | 0.07 | -0.16 | -0.11 | 0.03 | -0.12 | 0.26 | 0.39 |
| **IWS1** | **S157** | 0.01 | -0.16 | 0.00 | 0.10 | -0.20 | 0.21 | 0.564 |
| **IWS1** | **S222** | -0.07 | 0.01 | -0.06 | 0.04 | -0.02 | 0.10 | 0.136 |
| **IWS1** | **S222; S224** | -0.13 | 0.00 | 0.05 | -0.02 | 0.00 | 0.10 | 0.441 |
| **IWS1** | **S222** | 0.00 | -0.02 | -0.01 | 0.03 | 0.05 | -0.05 | 0.559 |
| **IWS1** | **S82; T/S** | -0.04 | -0.07 | 0.12 | -0.09 | -0.12 | 0.18 | 0.912 |
| **IWS1** | **S438; S440** | 0.06 | 0.12 | -0.07 | -0.18 | 0.08 | -0.03 | 0.449 |
| **IWS1** | **S** | -0.03 | 0.01 | 0.14 | -0.11 | 0.00 | -0.02 | 0.246 |
| **IWS1** | **S222; S224** | 0.16 | 0.05 | -0.07 | -0.21 | 0.11 | -0.06 | 0.437 |
| **IWS1** | **S198** | 0.03 | 0.02 | 0.10 | -0.08 | -0.06 | -0.01 | 0.039 |
| **IWS1** | **S196; S198** | 0.09 | 0.03 | 0.06 | -0.02 | -0.27 | 0.07 | 0.264 |
| **IWS1** | **S261** | 0.10 | 0.04 | 0.10 | -0.16 | -0.05 | -0.06 | 0.012 |
| **IWS1** | **S196; S198** | 0.08 | 0.09 | 0.14 | -0.11 | -0.16 | -0.07 | 0.002 |
| **IWS1** | **S398; S400** | 0.16 | 0.17 | -0.03 | -0.03 | -0.05 | -0.27 | 0.103 |
| **IWS1** | **S287; S289** | 0.07 | 0.13 | 0.12 | -0.09 | -0.18 | -0.07 | 0.004 |
| **IWS1** | **S211; S213; S224** | 0.07 | 0.14 | 0.09 | -0.41 | -0.12 | 0.15 | 0.24 |
| **IWS1** | **S261; S263** | 0.00 | 0.06 | 0.27 | -0.18 | -0.09 | -0.11 | 0.051 |
| **IWS1** | **S363; S365** | 0.14 | 0.04 | 0.16 | -0.31 | -0.13 | 0.04 | 0.084 |
| **IWS1** | **S398** | 0.22 | 0.13 | 0.05 | -0.13 | -0.13 | -0.18 | 0.005 |
| **IWS1** | **S54** | 0.17 | 0.18 | 0.05 | -0.29 | 0.06 | -0.23 | 0.073 |
| **IWS1** | **S261; S263** | 0.16 | 0.10 | 0.21 | -0.24 | -0.13 | -0.16 | 0.002 |
| **IWS1** | **S440** | 0.22 | 0.10 | 0.16 | -0.22 | -0.06 | -0.26 | 0.009 |
| **IWS1** | **S300** | 0.23 | 0.13 | 0.17 | -0.18 | -0.14 | -0.30 | 0.003 |

| | | | | | | | | |
|---|---|---|---|---|---|---|---|---|
| **IWS1** | **S287; S289** | 0.16 | 0.15 | 0.23 | -0.30 | -0.08 | -0.24 | 0.005 |
| **IWS1** | **S289** | 0.11 | 0.23 | 0.17 | -0.35 | 0.13 | -0.43 | 0.095 |
| **IWS1** | **S720** | 0.38 | 0.04 | 0.29 | -0.31 | 0.01 | -0.66 | 0.062 |
| **IWS1** | **S438; S440** | 0.31 | 0.26 | 0.16 | -0.21 | -0.06 | -0.69 | 0.046 |
| **IWS1** | **S420; S422; S426** | 0.39 | 0.28 | 0.17 | -0.25 | -0.14 | -0.73 | 0.027 |
| **IWS1** | **S720** | 0.48 | 0.00 | 0.38 | -0.37 | -0.07 | -0.79 | 0.051 |
| **IWS1** | **S422; S426; S429** | 0.36 | 0.37 | 0.23 | -0.47 | -0.22 | -0.59 | 0.003 |
| **JADE1** | **S603** | -0.08 | -0.08 | -0.07 | -0.05 | -0.03 | 0.28 | 0.262 |
| **JADE1** | **S603** | 0.05 | 0.09 | -0.05 | -0.01 | -0.07 | -0.01 | 0.263 |
| **JADE1** | **S743** | -0.08 | 0.29 | -0.12 | 0.25 | -0.19 | -0.23 | 0.699 |
| **JADE1** | **S810** | 0.07 | 0.15 | 0.19 | -0.25 | -0.07 | -0.15 | 0.01 |
| **JADE1** | **S703** | 0.34 | 0.29 | 0.03 | -0.12 | -0.40 | -0.28 | 0.019 |
| **JADE3** | **S85** | -0.02 | 0.04 | -0.31 | 0.01 | -0.02 | 0.24 | 0.281 |
| **JADE3** | **S774; S776** | -0.11 | 0.11 | -0.08 | -0.02 | 0.06 | 0.04 | 0.493 |
| **JADE3** | **S774** | 0.01 | 0.26 | 0.00 | -0.04 | -0.22 | -0.05 | 0.143 |
| **JARID2** | **T209** | -0.27 | -0.37 | 0.74 | -0.38 | -0.37 | 0.25 | 0.647 |
| **JARID2** | **S455** | 0.08 | 0.09 | 0.16 | -0.54 | 0.01 | 0.09 | 0.274 |
| **JDP2** | **T148** | -0.05 | -0.25 | 0.05 | 0.12 | 0.06 | 0.04 | 0.171 |
| **JDP2** | **T148** | 0.00 | 0.30 | 0.17 | -0.25 | 0.06 | -0.39 | 0.09 |
| **JDP2** | **T148** | 0.07 | 0.45 | 0.01 | 0.14 | -0.45 | -0.43 | 0.147 |
| **JMJD1C** | **S1058** | 0.12 | 0.09 | -0.03 | -0.43 | -0.11 | 0.27 | 0.519 |
| **JMJD1C** | **S2053** | -0.08 | 0.06 | 0.25 | 0.01 | -0.22 | -0.05 | 0.242 |
| **JMJD1C** | **S652** | 0.05 | 0.18 | -0.02 | 0.19 | -0.32 | -0.15 | 0.356 |
| **JMY** | **S713** | 0.10 | 0.07 | 0.02 | 0.31 | -0.32 | -0.27 | 0.494 |
| **JMY** | **S115** | -0.09 | 0.05 | 0.36 | 0.01 | -0.05 | -0.37 | 0.249 |
| **JOSD1** | **S15** | 0.27 | -0.35 | 0.48 | 0.11 | -0.28 | -0.49 | 0.314 |
| **JPH1** | **S413** | -0.11 | -0.30 | -0.19 | 0.35 | 0.11 | 0.05 | 0.025 |
| **JPH1** | **S475; S478** | 0.04 | -0.06 | -0.01 | 0.15 | -0.14 | 0.00 | 0.912 |
| **JPH1** | **S452** | -0.07 | 0.19 | 0.02 | 0.03 | -0.15 | -0.04 | 0.359 |
| **JPH1** | **S162** | 0.03 | 0.08 | 0.14 | -0.07 | 0.06 | -0.27 | 0.15 |
| **JPH1** | **S625** | 0.11 | 0.19 | 0.09 | -0.03 | -0.16 | -0.25 | 0.016 |
| **JPH1** | **T460; S465** | 0.19 | -0.05 | 0.48 | 0.10 | -0.66 | -0.34 | 0.136 |
| **JPT1** | **S131** | -0.36 | 0.18 | -0.27 | 0.28 | -0.23 | 0.25 | 0.335 |
| **JPT1** | **S87** | 0.12 | 0.07 | -0.23 | -0.02 | 0.02 | 0.02 | 0.87 |
| **JPT1** | **S87** | 0.36 | 0.41 | 0.06 | -0.37 | -0.24 | -0.46 | 0.008 |
| **JPT1** | **S87** | 0.47 | 0.41 | 0.14 | -0.40 | -0.45 | -0.53 | 0.002 |
| **JPT2** | **S97** | -0.07 | -0.03 | -0.08 | 0.13 | -0.26 | 0.26 | 0.55 |
| **JRK** | **S457** | -0.06 | 0.22 | 0.07 | 0.00 | -0.11 | -0.15 | 0.148 |
| **JUN** | **S63** | -0.72 | -0.45 | -0.35 | 0.24 | 0.70 | 0.10 | 0.016 |
| **JUN** | **S243** | -0.05 | -0.12 | -0.14 | 0.13 | 0.10 | 0.06 | 0.004 |
| **JUN** | **S243** | 0.09 | -0.18 | -0.07 | 0.09 | -0.11 | 0.15 | 0.417 |
| **JUN** | **S243** | 0.09 | -0.02 | -0.13 | 0.06 | 0.05 | -0.06 | 0.616 |
| **JUNB** | **S117** | -0.14 | -0.14 | -0.05 | -0.19 | 0.20 | 0.26 | 0.244 |
| **JUNB** | **S117** | -0.11 | -0.15 | -0.01 | -0.17 | 0.18 | 0.22 | 0.267 |
| **JUNB** | **T255** | -0.01 | -0.15 | 0.01 | -0.07 | -0.24 | 0.37 | 0.733 |
| **JUNB** | **S251; T255; S259** | -0.04 | 0.05 | -0.12 | 0.01 | -0.09 | 0.17 | 0.547 |
| **JUNB** | **T255; S259** | -0.17 | -0.08 | 0.16 | -0.04 | -0.05 | 0.15 | 0.711 |
| **JUNB** | **T255; S259** | -0.11 | 0.02 | 0.03 | -0.04 | 0.01 | 0.08 | 0.504 |

| | | | | | | | | |
|---|---|---|---|---|---|---|---|---|
| **JUNB** | **T255** | 0.07 | -0.02 | 0.18 | -0.11 | -0.18 | 0.03 | 0.133 |
| **JUND** | **S100** | -0.29 | -0.33 | -0.32 | 0.25 | 0.30 | 0.21 | 4E-05 |
| **JUND** | **S255; S259** | -0.03 | -0.08 | 0.04 | 0.15 | -0.03 | -0.06 | 0.583 |
| **JUND** | **S259** | -0.04 | 0.03 | -0.04 | -0.02 | 0.05 | 0.02 | 0.312 |
| **JUND** | **S/T** | 0.11 | -0.12 | 0.00 | -0.08 | 0.01 | 0.08 | 0.95 |
| **JUND** | **S255** | -0.24 | -0.10 | 0.32 | -0.09 | 0.11 | -0.07 | 0.924 |
| **KANK1** | **S881** | -0.21 | 0.02 | 0.02 | 0.02 | -0.02 | 0.15 | 0.297 |
| **KANK1** | **S/Y** | 0.14 | 0.25 | 0.36 | -1.11 | -0.24 | 0.16 | 0.168 |
| **KANK2** | **S375** | -0.16 | -0.25 | -0.04 | 0.16 | 0.15 | 0.10 | 0.01 |
| **KANK2** | **S323** | -0.12 | -0.04 | -0.14 | 0.11 | -0.23 | 0.34 | 0.349 |
| **KANSL1** | **S991** | -0.03 | 0.03 | -0.14 | 0.00 | -0.19 | 0.29 | 0.623 |
| **KANSL1** | **S1045** | 0.08 | -0.26 | 0.16 | -0.01 | 0.06 | -0.06 | 0.961 |
| **KANSL1** | **S994; T1003** | 0.08 | -0.34 | 0.28 | 0.03 | -0.01 | -0.12 | 0.846 |
| **KANSL1** | **S1045** | -0.10 | 0.08 | 0.19 | -0.11 | -0.06 | -0.01 | 0.277 |
| **KANSL1** | **S249** | 0.08 | 0.05 | 0.08 | -0.14 | 0.04 | -0.14 | 0.071 |
| **KANSL1** | **T1022** | -0.02 | -0.28 | 0.43 | -0.45 | -0.22 | 0.33 | 0.649 |
| **KANSL1** | **S1082** | 0.07 | -0.02 | 0.14 | -0.47 | 0.17 | 0.02 | 0.482 |
| **KANSL2** | **S149** | -0.06 | -0.13 | 0.08 | 0.04 | -0.01 | 0.07 | 0.354 |
| **KANSL3** | **S536** | 0.03 | -0.01 | -0.16 | -0.07 | -0.04 | 0.22 | 0.479 |
| **KANSL3** | **S633** | -0.02 | 0.02 | 0.17 | 0.24 | -0.58 | 0.03 | 0.558 |
| **KANSL3** | **S515** | 0.08 | 0.14 | 0.20 | -0.05 | -0.09 | -0.35 | 0.037 |
| **KANSL3** | **S523; S536** | 0.24 | 0.10 | 0.13 | -0.27 | -0.12 | -0.14 | 0.006 |
| **KAT5** | **S90** | -0.07 | 0.00 | 0.25 | -0.11 | -0.05 | -0.06 | 0.244 |
| **KAT5** | **S199** | 0.12 | 0.23 | 0.41 | -0.92 | 0.00 | -0.17 | 0.105 |
| **KAT6A** | **S473** | -0.47 | -0.26 | -0.20 | 0.04 | 0.55 | 0.11 | 0.039 |
| **KAT6A** | **S941** | -0.19 | -0.28 | -0.12 | 0.26 | 0.15 | 0.09 | 0.006 |
| **KAT6A** | **S941** | -0.42 | -0.02 | -0.12 | 0.20 | 0.11 | 0.17 | 0.049 |
| **KAT6A** | **T/S/Y** | -0.21 | -0.34 | -0.06 | 0.56 | -0.26 | 0.11 | 0.244 |
| **KAT6A** | **T/S/Y** | -0.10 | -0.25 | -0.17 | 0.46 | -0.35 | 0.24 | 0.307 |
| **KAT6A** | **S1089** | -0.14 | -0.09 | -0.15 | 0.17 | 0.04 | 0.13 | 0.004 |
| **KAT6A** | **S1113** | -0.07 | -0.07 | -0.08 | 0.06 | -0.05 | 0.19 | 0.112 |
| **KAT6A** | **S1136** | 0.06 | -0.05 | -0.19 | 0.05 | -0.14 | 0.24 | 0.457 |
| **KAT6A** | **S828** | 0.09 | -0.04 | -0.10 | 0.03 | 0.10 | -0.09 | 0.698 |
| **KAT6A** | **S678** | 0.07 | 0.04 | -0.07 | -0.07 | 0.19 | -0.19 | 0.773 |
| **KAT6A** | **S448** | -0.07 | 0.09 | 0.09 | -0.11 | -0.16 | 0.13 | 0.485 |
| **KAT6A** | **S784** | -0.30 | 0.55 | -0.31 | 0.20 | -0.82 | 0.27 | 0.839 |
| **KAT6A** | **S/T** | 0.19 | -0.01 | 0.25 | -0.23 | -0.49 | 0.15 | 0.178 |
| **KAT6A** | **S448** | 0.26 | 0.08 | 0.14 | -0.19 | -0.12 | -0.24 | 0.006 |
| **KAT6A** | **S/T** | 0.16 | 0.29 | 0.26 | -0.29 | -0.06 | -0.54 | 0.021 |
| **KAT6B** | **T1275** | -0.05 | -0.08 | 0.10 | 0.10 | 0.12 | -0.22 | 0.942 |
| **KAT6B** | **T1275** | 0.11 | 0.00 | 0.04 | 0.19 | 0.09 | -0.55 | 0.584 |
| **KAT6B** | **S1502** | 0.03 | 0.06 | 0.15 | -0.08 | -0.06 | -0.12 | 0.016 |
| **KAT6B** | **S998** | -0.02 | 0.12 | 0.19 | -0.43 | -0.16 | 0.20 | 0.306 |
| **KAT6B** | **S1303** | -0.04 | 0.29 | 0.25 | -0.30 | -0.01 | -0.31 | 0.061 |
| **KAT6B** | **S1502** | 0.10 | 0.17 | 0.29 | 0.00 | -0.08 | -0.66 | 0.114 |
| **KAT7** | **S64** | -0.46 | -0.22 | -0.56 | 0.29 | 0.47 | 0.18 | 0.005 |
| **KAT7** | **S124; T128** | 0.04 | -0.18 | -0.04 | 0.15 | -0.06 | 0.07 | 0.279 |
| **KAT7** | **S99; S100; S102** | -0.07 | 0.05 | 0.06 | -0.16 | 0.20 | -0.12 | 0.77 |

| | | | | | | | | |
|---|---|---|---|---|---|---|---|---|
| KAT7 | S100; S102 | 0.00 | 0.00 | 0.08 | -0.10 | 0.05 | -0.04 | 0.344 |
| KAT7 | S102 | 0.04 | 0.05 | 0.18 | -0.24 | -0.01 | -0.05 | 0.087 |
| KAT7 | S138 | 0.03 | 0.09 | 0.16 | -0.08 | -0.09 | -0.14 | 0.01 |
| KAT7 | S | 0.10 | 0.19 | 0.10 | -0.10 | -0.05 | -0.28 | 0.023 |
| KAT7 | S138 | -0.06 | 0.12 | 0.30 | 0.07 | -0.11 | -0.44 | 0.196 |
| KAT7 | S124; T128 | 0.12 | 0.11 | 0.17 | -0.19 | -0.02 | -0.24 | 0.015 |
| KAT7 | S124; T130; S138 | 0.28 | 0.14 | 0.01 | -0.03 | -0.21 | -0.25 | 0.043 |
| KAT7 | T128; T130; S143 | 0.14 | 0.17 | 0.14 | -0.08 | -0.21 | -0.21 | 0.002 |
| KAT7 | T88 | 0.17 | 0.08 | 0.21 | -0.20 | -0.03 | -0.31 | 0.021 |
| KAT7 | S56; S57 | 0.13 | 0.08 | 0.26 | -0.20 | 0.00 | -0.37 | 0.045 |
| KAT7 | S57 | 0.26 | 0.12 | 0.28 | -0.17 | -0.09 | -0.56 | 0.035 |
| KAT7 | S57 | 0.21 | 0.21 | 0.31 | -0.40 | -0.03 | -0.49 | 0.018 |
| KAT7 | T88 | 0.22 | 0.17 | 0.38 | -0.37 | -0.19 | -0.41 | 0.003 |
| KAT7 | T128 | 0.35 | 0.22 | 0.21 | -0.29 | -0.34 | -0.34 | 2E-04 |
| KAT7 | T128 | 0.44 | 0.35 | 0.38 | -0.51 | -0.41 | -0.71 | 5E-04 |
| KAT8 | S42 | 0.00 | -0.11 | -0.09 | 0.09 | -0.03 | 0.13 | 0.092 |
| KAT8 | S348 | 0.09 | 0.07 | 0.13 | -0.18 | -0.17 | 0.02 | 0.038 |
| KATNA1 | S170 | -0.05 | 0.34 | 0.07 | -0.33 | -0.03 | -0.08 | 0.151 |
| KATNIP | S/T | -0.22 | -0.11 | -0.36 | -0.02 | 0.39 | 0.19 | 0.039 |
| KATNIP | S/T | -0.10 | -0.13 | 0.07 | 0.14 | 0.00 | 0.00 | 0.295 |
| KATNIP | S660 | 0.32 | -0.13 | 0.26 | -0.09 | -0.42 | -0.07 | 0.131 |
| KCNK2 | S348 | 0.14 | 0.08 | -0.10 | -0.12 | 0.04 | -0.05 | 0.38 |
| KCNK2 | S366 | 0.09 | 0.06 | 0.23 | -0.02 | 0.09 | -0.59 | 0.241 |
| KCNMA1 | Y1086 | 0.00 | -0.03 | -0.13 | -0.02 | -0.02 | 0.18 | 0.282 |
| KCNMA1 | S | -0.06 | -0.04 | 0.04 | -0.08 | 0.01 | 0.12 | 0.609 |
| KCNMA1 | S978 | -0.03 | 0.00 | 0.01 | -0.05 | 0.19 | -0.12 | 0.876 |
| KCNMA1 | S | -0.18 | 0.30 | -0.01 | 0.04 | -0.07 | -0.13 | 0.577 |
| KCNMA1 | S778 | 0.16 | 0.12 | 0.27 | -0.36 | 0.13 | -0.49 | 0.094 |
| KCTD18 | S393 | -0.56 | 0.07 | 0.09 | -0.05 | 0.65 | -0.58 | 0.752 |
| KCTD18 | T259 | 0.38 | 0.45 | -0.35 | -0.55 | -0.15 | -0.05 | 0.243 |
| KCTD3 | T/S | 0.27 | 0.56 | 0.30 | -0.54 | -0.19 | -0.91 | 0.016 |
| KDF1 | S137 | -0.29 | -0.05 | 0.01 | 0.01 | -0.06 | 0.32 | 0.243 |
| KDF1 | T/S | 0.03 | 0.29 | -0.05 | -0.05 | -0.08 | -0.19 | 0.147 |
| KDF1 | S/T | -0.02 | 0.35 | 0.01 | -0.05 | -0.19 | -0.17 | 0.12 |
| KDM1A | T/S | -0.46 | -0.44 | -1.11 | -0.36 | -0.44 | 1.35 | 0.241 |
| KDM1A | S69 | -0.67 | -0.31 | 0.32 | -0.17 | 0.43 | 0.11 | 0.365 |
| KDM1A | S69 | -0.16 | -0.15 | -0.22 | 0.32 | 0.12 | 0.01 | 0.025 |
| KDM1A | S131; S137 | -0.15 | -0.13 | -0.18 | 0.08 | -0.01 | 0.33 | 0.05 |
| KDM1A | S131; S137 | -0.31 | 0.01 | -0.13 | 0.11 | -0.10 | 0.33 | 0.172 |
| KDM1A | S69 | -0.30 | -0.07 | -0.08 | -0.13 | -0.06 | 0.50 | 0.306 |
| KDM1A | S131; S137 | -0.04 | -0.06 | -0.26 | 0.09 | 0.04 | 0.19 | 0.051 |
| KDM1A | S131; S137 | 0.05 | -0.09 | -0.37 | -0.02 | 0.53 | -0.28 | 0.477 |
| KDM1A | S166 | -0.14 | -0.09 | -0.01 | -0.01 | -0.04 | 0.25 | 0.225 |
| KDM1A | S166 | 0.08 | -0.03 | -0.26 | -0.11 | -0.03 | 0.29 | 0.488 |
| KDM1A | S166 | -0.08 | 0.05 | -0.14 | 0.05 | -0.05 | 0.15 | 0.249 |
| KDM1A | S69 | 0.04 | 0.06 | -0.35 | 0.49 | -0.39 | -0.04 | 0.734 |
| KDM1A | S166 | 0.09 | -0.07 | -0.17 | 0.06 | -0.03 | 0.10 | 0.345 |
| KDM1A | S/Y | -0.15 | 0.08 | -0.06 | 0.11 | -0.17 | 0.15 | 0.585 |

| | | | | | | | | |
|---|---|---|---|---|---|---|---|---|
| **KDM1A** | **S166** | -0.15 | 0.13 | -0.02 | -0.19 | 0.25 | -0.07 | 0.928 |
| **KDM1A** | **S849** | 0.00 | -0.09 | 0.06 | -0.14 | 0.33 | -0.23 | 0.982 |
| **KDM1A** | **S137** | -0.07 | 0.10 | 0.04 | 0.13 | -0.09 | -0.13 | 0.604 |
| **KDM1A** | **S137** | -0.01 | 0.05 | 0.14 | 0.00 | -0.13 | -0.07 | 0.095 |
| **KDM1A** | **S137** | -0.12 | 0.33 | -0.04 | -0.27 | 0.00 | 0.04 | 0.473 |
| **KDM1A** | **S849** | 0.07 | 0.13 | 0.10 | 0.04 | -0.19 | -0.19 | 0.05 |
| **KDM1A** | **S69** | 0.18 | 0.40 | -0.33 | 0.32 | -0.43 | -0.38 | 0.488 |
| **KDM1A** | **S137** | 0.33 | 0.28 | 0.07 | -0.11 | -0.11 | -0.66 | 0.061 |
| **KDM1B** | **S17** | -0.03 | -0.02 | 0.03 | 0.13 | -0.21 | 0.08 | 0.955 |
| **KDM1B** | **S247** | 0.13 | 0.07 | -0.14 | -0.13 | -0.12 | 0.15 | 0.69 |
| **KDM1B** | **S247** | 0.27 | 0.27 | -0.14 | -0.20 | -0.07 | -0.23 | 0.107 |
| **KDM2A** | **S433** | -0.57 | 0.20 | 0.00 | 0.28 | 0.04 | -0.10 | 0.474 |
| **KDM2A** | **S558** | -0.06 | -0.09 | -0.05 | -0.08 | 0.22 | 0.04 | 0.233 |
| **KDM2A** | **T713; S718** | -0.03 | 0.00 | 0.05 | -0.13 | 0.04 | 0.06 | 0.769 |
| **KDM2A** | **S28** | 0.11 | 0.03 | -0.10 | 0.02 | -0.11 | 0.04 | 0.698 |
| **KDM2A** | **S692** | 0.03 | 0.02 | 0.00 | -0.05 | -0.05 | 0.05 | 0.339 |
| **KDM2A** | **S869** | -0.20 | 0.32 | -0.10 | -0.29 | -0.03 | 0.19 | 0.828 |
| **KDM2A** | **T713; S718** | -0.03 | 0.00 | 0.10 | -0.14 | 0.04 | 0.02 | 0.505 |
| **KDM2A** | **S692** | 0.08 | 0.03 | -0.02 | 0.01 | -0.03 | -0.07 | 0.216 |
| **KDM2A** | **T550; S558** | -0.01 | -0.05 | 0.13 | -0.03 | 0.11 | -0.18 | 0.581 |
| **KDM2A** | **S28** | 0.04 | 0.08 | 0.10 | -0.18 | -0.01 | -0.05 | 0.044 |
| **KDM2A** | **T713; S718** | -0.05 | 0.19 | 0.08 | -0.10 | -0.05 | -0.10 | 0.086 |
| **KDM2A** | **S692** | 0.10 | 0.06 | 0.16 | -0.06 | -0.08 | -0.22 | 0.017 |
| **KDM2A** | **S692** | 0.19 | 0.15 | 0.22 | -0.16 | -0.06 | -0.44 | 0.025 |
| **KDM2A** | **S28** | 0.32 | 0.23 | 0.21 | -0.22 | -0.08 | -0.71 | 0.04 |
| **KDM2B** | **S795** | -0.38 | -0.42 | -0.55 | 0.33 | 0.20 | 0.48 | 0.001 |
| **KDM2B** | **S951** | -0.12 | -0.20 | -0.20 | 0.21 | 0.20 | 0.04 | 0.007 |
| **KDM2B** | **S445** | -0.15 | -0.32 | 0.14 | 0.20 | -0.27 | 0.27 | 0.461 |
| **KDM2B** | **T444** | 0.09 | -0.09 | -0.03 | 0.10 | -0.16 | 0.08 | 0.86 |
| **KDM2B** | **S57** | 0.11 | 0.03 | -0.03 | 0.07 | -0.04 | -0.16 | 0.364 |
| **KDM2B** | **S1031** | 0.11 | 0.13 | 0.13 | -0.16 | -0.09 | -0.15 | 4E-04 |
| **KDM2B** | **S497** | 0.20 | 0.14 | 0.08 | -0.06 | -0.16 | -0.26 | 0.01 |
| **KDM2B** | **S914** | 0.23 | 0.01 | 0.20 | -0.27 | -0.07 | -0.18 | 0.023 |
| **KDM2B** | **S474** | 0.24 | 0.06 | 0.24 | -0.27 | 0.00 | -0.39 | 0.038 |
| **KDM2B** | **S975; S979** | 0.22 | 0.18 | 0.26 | -0.37 | 0.05 | -0.52 | 0.043 |
| **KDM3A** | **S325** | 0.09 | -0.05 | -0.21 | 0.11 | 0.06 | -0.02 | 0.302 |
| **KDM3A** | **S325** | 0.09 | -0.05 | -0.21 | 0.11 | 0.06 | -0.02 | 0.302 |
| **KDM3A** | **S445** | 0.10 | 0.17 | 0.15 | 0.07 | -0.41 | -0.17 | 0.092 |
| **KDM3B** | **S798** | -0.32 | 0.07 | -0.75 | 0.21 | 0.21 | 0.31 | 0.074 |
| **KDM3B** | **S727; T729** | 0.13 | 0.03 | -0.16 | 0.34 | -0.40 | -0.06 | 0.866 |
| **KDM4A** | **S/T** | 0.18 | 0.44 | -0.23 | -0.30 | 0.28 | -0.67 | 0.35 |
| **KDM4B** | **S1039** | -0.07 | 0.08 | -0.04 | -0.37 | 0.36 | -0.06 | 0.956 |
| **KDM5A** | **S1666** | 0.17 | -0.17 | -0.46 | -0.25 | 0.02 | 0.49 | 0.441 |
| **KDM5A** | **S1365** | 0.18 | -0.04 | -0.28 | 0.09 | 0.06 | -0.05 | 0.585 |
| **KDM5A** | **S204** | -0.10 | 0.11 | -0.10 | 0.05 | -0.06 | 0.08 | 0.511 |
| **KDM5A** | **T/S** | -0.01 | 0.04 | -0.10 | 0.06 | -0.07 | 0.08 | 0.519 |
| **KDM5A** | **S1666** | 0.06 | 0.01 | -0.07 | -0.09 | 0.14 | -0.07 | 0.945 |
| **KDM5A** | **S1111** | 0.09 | -0.03 | -0.05 | -0.05 | 0.02 | 0.01 | 0.864 |

| | | | | | | | | |
|---|---|---|---|---|---|---|---|---|
| KDM5A | S204 | -0.06 | 0.05 | 0.08 | -0.09 | -0.03 | 0.04 | 0.476 |
| KDM5A | S1111 | 0.10 | -0.07 | 0.04 | -0.04 | 0.00 | -0.03 | 0.394 |
| KDM5A | S1331 | 0.16 | 0.01 | 0.09 | -0.18 | -0.07 | -0.04 | 0.038 |
| KDM5A | S1603 | 0.35 | 0.53 | 0.12 | -0.10 | -0.30 | -1.20 | 0.073 |
| KDM5A | S1603 | 0.25 | 0.46 | 0.11 | 0.31 | -0.21 | -1.95 | 0.266 |
| KDM5B | S1456 | -0.23 | -0.21 | -0.13 | 0.17 | 0.01 | 0.31 | 0.019 |
| KDM5B | S1384 | 0.35 | 0.46 | -0.08 | -0.43 | -0.09 | -0.46 | 0.05 |
| KDM5C | S1359 | -0.05 | -0.05 | 0.15 | -0.06 | 0.16 | -0.18 | 0.735 |
| KDM5C | S1359 | 0.17 | 0.07 | 0.26 | -0.22 | -0.22 | -0.15 | 0.004 |
| KDM6A | S818; S829 | -0.18 | -0.37 | -0.33 | 0.43 | 0.01 | 0.25 | 0.018 |
| KDM6B | S224 | -0.39 | 0.20 | -0.31 | 0.13 | 0.03 | 0.22 | 0.197 |
| KDM6B | S627 | -0.07 | -0.15 | -0.24 | 0.21 | -0.07 | 0.24 | 0.064 |
| KDM7A | S503 | 0.03 | -0.38 | -0.39 | -0.02 | -0.36 | 0.75 | 0.355 |
| KDM7A | S604 | -0.59 | -0.05 | -0.09 | 0.76 | 0.18 | -0.75 | 0.553 |
| KDM7A | S604 | 0.20 | -0.23 | -0.09 | 0.03 | -0.01 | 0.07 | 0.616 |
| KDM7A | T601 | 0.13 | -0.15 | 0.11 | -0.25 | -0.19 | 0.28 | 0.703 |
| KDM7A | S/T/Y | 0.02 | 0.21 | 0.00 | -0.01 | -0.20 | -0.06 | 0.13 |
| KHDRBS1 | S58 | -0.51 | -0.01 | 0.08 | 0.02 | -0.08 | 0.36 | 0.339 |
| KHDRBS1 | S20 | -0.52 | -0.57 | 0.59 | -0.24 | 0.24 | 0.13 | 0.631 |
| KHDRBS1 | S | -0.33 | -0.11 | 0.14 | -0.19 | 0.08 | 0.32 | 0.456 |
| KHSRP | S480 | 0.04 | -0.14 | -0.03 | 0.15 | -0.18 | 0.12 | 0.57 |
| KHSRP | S181 | 0.07 | -0.11 | 0.04 | 0.03 | -0.06 | 0.03 | 0.981 |
| KIAA0319L | T/S | -0.03 | 0.13 | 0.00 | -0.31 | -0.03 | 0.19 | 0.609 |
| KIAA0319L | S978 | 0.05 | 0.18 | 0.06 | -0.18 | 0.03 | -0.18 | 0.06 |
| KIAA0586 | S406 | 0.18 | -0.27 | -0.37 | 0.16 | 0.13 | 0.08 | 0.175 |
| KIAA0753 | S287 | 0.43 | 0.17 | -0.52 | -0.26 | 0.20 | -0.24 | 0.717 |
| KIAA0753 | S826 | 0.37 | 0.11 | 0.17 | -0.56 | 0.01 | -0.29 | 0.052 |
| KIAA1143 | S50 | 0.39 | 0.11 | -0.03 | -0.34 | 0.04 | -0.31 | 0.108 |
| KIAA1143 | S50 | 0.41 | 0.19 | -0.06 | -0.59 | 0.14 | -0.31 | 0.16 |
| KIAA1191 | S183 | 0.27 | 0.18 | 0.10 | -0.05 | -0.67 | -0.01 | 0.127 |
| KIAA1191 | S183 | 0.33 | 0.10 | 0.10 | 0.10 | -1.02 | 0.04 | 0.274 |
| KIAA1217 | T/S/Y | -0.14 | 0.12 | -0.17 | 0.09 | -0.03 | 0.09 | 0.327 |
| KIAA1217 | Y | -0.06 | -0.18 | 0.25 | -0.03 | -0.08 | 0.06 | 0.885 |
| KIAA1217 | S1648 | 0.01 | 0.03 | 0.13 | -0.30 | 0.12 | -0.03 | 0.368 |
| KIAA1217 | S1044 | 0.43 | -0.19 | -0.04 | 0.07 | -0.07 | -0.32 | 0.484 |
| KIAA1328 | S499 | 0.06 | 0.12 | -0.20 | -0.33 | 0.04 | 0.24 | 0.963 |
| KIAA1328 | S | -0.17 | -0.07 | 0.19 | 0.10 | -0.60 | 0.36 | 0.923 |
| KIAA1328 | S273 | 0.14 | 0.09 | 0.27 | 0.00 | -0.06 | -0.58 | 0.121 |
| KIAA1328 | T/S | 0.08 | 0.30 | 0.13 | -0.23 | 0.02 | -0.42 | 0.056 |
| KIAA1522 | S404; S406; S/T | -0.31 | -0.31 | -0.24 | 0.35 | -0.03 | 0.36 | 0.017 |
| KIAA1522 | S404; S/T | -0.51 | -0.29 | 0.01 | 0.13 | -0.19 | 0.59 | 0.184 |
| KIAA1522 | T593 | -0.13 | -0.33 | -0.04 | -0.45 | 0.40 | 0.35 | 0.411 |
| KIAA1522 | S669; S673 | -0.09 | -0.17 | -0.11 | 0.21 | -0.38 | 0.41 | 0.444 |
| KIAA1522 | S570 | -0.06 | 0.10 | -0.28 | 0.06 | -0.19 | 0.30 | 0.492 |
| KIAA1522 | S409 | 0.01 | -0.15 | -0.05 | 0.26 | -0.17 | 0.06 | 0.43 |
| KIAA1522 | S379 | 0.08 | -0.37 | 0.12 | 0.14 | -0.12 | 0.09 | 0.627 |
| KIAA1522 | S545 | -0.09 | 0.03 | 0.03 | 0.00 | 0.04 | -0.01 | 0.613 |
| KIAA1522 | S979 | 0.14 | -0.07 | -0.01 | -0.06 | -0.10 | 0.09 | 0.628 |

| | | | | | | | | |
|---|---|---|---|---|---|---|---|---|
| KIAA1522 | S421 | -0.13 | -0.08 | 0.29 | 0.06 | -0.10 | -0.09 | 0.64 |
| KIAA1522 | S929 | 0.04 | 0.07 | 0.13 | -0.11 | 0.04 | -0.20 | 0.084 |
| KIAA1522 | S669; S673 | 0.33 | -0.18 | 0.17 | -0.18 | -0.10 | -0.11 | 0.203 |
| KIAA1522 | S929 | 0.00 | 0.13 | 0.18 | -0.02 | 0.08 | -0.46 | 0.245 |
| KIAA1522 | S327 | 0.24 | -0.08 | 0.29 | -0.66 | 0.03 | -0.01 | 0.224 |
| KIAA1671 | S1488 | -0.47 | 0.17 | -0.61 | 0.24 | 0.36 | 0.06 | 0.112 |
| KIAA1671 | T600 | -0.50 | -0.11 | -0.10 | 0.14 | 0.28 | 0.15 | 0.038 |
| KIAA1671 | S558 | -0.18 | -0.32 | -0.21 | 0.26 | -0.21 | 0.47 | 0.115 |
| KIAA1671 | S1786 | -0.04 | -0.23 | -0.07 | 0.10 | 0.03 | 0.17 | 0.045 |
| KIAA1671 | S774 | -0.14 | -0.13 | -0.03 | 0.14 | -0.08 | 0.21 | 0.102 |
| KIAA1671 | S/T | -0.17 | -0.05 | 0.00 | 0.11 | -0.22 | 0.27 | 0.45 |
| KIAA1671 | S1224 | -0.13 | -0.04 | 0.00 | 0.08 | -0.13 | 0.19 | 0.378 |
| KIAA1671 | S797 | -0.44 | 0.37 | -0.11 | 0.21 | 0.12 | -0.33 | 0.86 |
| KIAA1671 | S465 | -0.03 | -0.04 | 0.11 | -0.12 | 0.12 | -0.05 | 0.731 |
| KIAA1671 | S1312 | 0.07 | 0.04 | 0.01 | 0.00 | -0.24 | 0.10 | 0.446 |
| KIAA1671 | S1019 | 0.02 | 0.06 | 0.08 | -0.05 | -0.09 | -0.04 | 0.008 |
| KIAA1671 | T634 | -0.31 | 0.04 | 0.49 | -0.45 | -0.14 | 0.16 | 0.501 |
| KIAA1671 | S1224 | 0.17 | 0.11 | 0.09 | -0.04 | -0.09 | -0.29 | 0.027 |
| KIAA1671 | S/T/Y | 0.20 | -0.15 | 0.33 | -0.39 | -0.14 | 0.03 | 0.195 |
| KIAA1958 | T/S | 0.16 | 0.08 | 0.14 | -0.30 | 0.09 | -0.24 | 0.087 |
| KIDINS220 | S1521; S1526 | 0.03 | -0.19 | -0.05 | 0.31 | -0.16 | 0.01 | 0.471 |
| KIDINS220 | S1296 | -0.02 | -0.23 | 0.05 | -0.05 | -0.07 | 0.26 | 0.452 |
| KIDINS220 | S1521 | 0.00 | 0.06 | -0.02 | 0.04 | -0.09 | 0.01 | 0.624 |
| KIDINS220 | S1555 | 0.00 | 0.07 | 0.01 | 0.07 | -0.06 | -0.09 | 0.369 |
| KIDINS220 | S1463 | 0.01 | -0.09 | 0.23 | 0.00 | -0.44 | 0.19 | 0.558 |
| KIDINS220 | S1652 | -0.01 | 0.41 | -0.14 | -0.13 | 0.15 | -0.42 | 0.395 |
| KIDINS220 | S918 | 0.16 | 0.06 | 0.11 | -0.10 | 0.00 | -0.26 | 0.048 |
| KIDINS220 | S885 | 0.03 | 0.03 | 0.25 | 0.00 | -0.04 | -0.33 | 0.145 |
| KIDINS220 | S1593; S1594; S1598 | 0.13 | 0.16 | 0.11 | -0.07 | -0.15 | -0.22 | 0.004 |
| KIDINS220 | S/T | -0.09 | 0.30 | 0.07 | 0.02 | 0.33 | -0.96 | 0.503 |
| KIDINS220 | S918 | 0.21 | 0.18 | 0.03 | -0.18 | -0.05 | -0.26 | 0.019 |
| KIDINS220 | S1362 | 0.20 | 0.28 | 0.12 | -0.11 | -0.24 | -0.36 | 0.008 |
| KIDINS220 | S885 | 0.35 | 0.06 | 0.15 | -0.04 | -0.05 | -0.65 | 0.115 |
| KIDINS220 | S1593; S1594; S1598 | 0.27 | 0.16 | 0.25 | -0.11 | -0.40 | -0.33 | 0.006 |
| KIF11 | T926 | -0.15 | -0.15 | 0.04 | -0.01 | 0.05 | 0.20 | 0.129 |
| KIF13B | S | -0.35 | -0.52 | -0.09 | 0.16 | 0.44 | 0.14 | 0.023 |
| KIF14 | S346 | 0.00 | -0.21 | -0.12 | 0.28 | 0.08 | -0.08 | 0.168 |
| KIF14 | S346 | 0.07 | 0.18 | 0.27 | -0.16 | -0.02 | -0.45 | 0.051 |
| KIF16B | S1017 | -0.19 | -0.04 | -0.18 | 0.07 | 0.15 | 0.15 | 0.009 |
| KIF16B | S1052 | 0.04 | 0.08 | 0.09 | -0.25 | -0.10 | 0.11 | 0.217 |
| KIF18B | S447; S480 | -0.12 | -0.29 | 0.05 | 0.21 | -0.11 | 0.19 | 0.21 |
| KIF18B | S404 | -0.06 | -0.06 | -0.16 | 0.12 | 0.05 | 0.10 | 0.011 |
| KIF18B | S480; S/T | -0.30 | 0.07 | -0.10 | -0.27 | 0.00 | 0.46 | 0.502 |
| KIF18B | T674 | 0.02 | -0.27 | 0.05 | -0.27 | 0.26 | 0.13 | 0.599 |
| KIF18B | S447; S480 | -0.38 | 0.03 | 0.19 | 0.23 | -0.16 | 0.01 | 0.723 |
| KIF18B | S682 | -0.05 | 0.02 | 0.07 | -0.34 | 0.02 | 0.23 | 0.798 |
| KIF18B | S447 | -0.10 | 0.08 | 0.11 | -0.19 | 0.11 | -0.03 | 0.542 |
| KIF18B | S447; S480 | -0.51 | 0.26 | 0.28 | 0.18 | -0.05 | -0.36 | 0.791 |

| | | | | | | | | |
|---|---|---|---|---|---|---|---|---|
| **KIF18B** | **S662** | 0.29 | 0.20 | -0.20 | 0.10 | 0.14 | -0.77 | 0.463 |
| **KIF1B** | **S1487** | 0.22 | -0.57 | -0.22 | -0.48 | 0.40 | 0.34 | 0.489 |
| **KIF1C** | **S494** | -0.08 | -0.14 | -0.08 | 0.23 | 0.20 | -0.18 | 0.237 |
| **KIF1C** | **S674; S676** | 0.00 | -0.15 | -0.04 | -0.13 | 0.00 | 0.28 | 0.414 |
| **KIF1C** | **S817** | -0.01 | -0.03 | -0.03 | -0.01 | -0.14 | 0.19 | 0.719 |
| **KIF1C** | **T1083** | -0.04 | 0.04 | 0.25 | 0.00 | -0.27 | -0.02 | 0.213 |
| **KIF1C** | **S915** | -0.10 | 0.20 | 0.23 | -0.07 | 0.01 | -0.35 | 0.175 |
| **KIF1C** | **S676** | 0.12 | 0.32 | 0.30 | -0.23 | -0.10 | -0.62 | 0.029 |
| **KIF20A** | **S433** | -0.24 | -0.17 | 0.03 | 0.22 | 0.22 | -0.12 | 0.167 |
| **KIF20A** | **S556** | 0.13 | -0.09 | -0.01 | -0.01 | 0.03 | -0.05 | 0.772 |
| **KIF20A** | **S532** | -0.08 | 0.13 | 0.13 | -0.09 | 0.08 | -0.20 | 0.303 |
| **KIF20A** | **S825** | 0.01 | 0.07 | 0.17 | -0.26 | 0.06 | -0.09 | 0.151 |
| **KIF20B** | **S/T** | 0.33 | 0.04 | 0.05 | -0.50 | -0.08 | 0.03 | 0.159 |
| **KIF21A** | **S1212** | 0.05 | 0.10 | -0.14 | -0.20 | 0.21 | -0.06 | 0.898 |
| **KIF21A** | **S853** | -0.04 | 0.28 | -0.19 | 0.24 | 0.02 | -0.44 | 0.759 |
| **KIF21A** | **S1239** | 0.15 | 0.26 | -0.07 | -0.16 | 0.03 | -0.27 | 0.13 |
| **KIF21A** | **S853** | 0.43 | 0.36 | -0.36 | -0.47 | -0.14 | -0.06 | 0.257 |
| **KIF22** | **S562** | -1.69 | -1.59 | -1.73 | 0.63 | 0.73 | 0.89 | 1E-05 |
| **KIF22** | **S581** | -0.99 | 0.46 | -0.10 | -0.12 | -0.38 | 0.59 | 0.661 |
| **KIF23** | **S924; T927** | -0.18 | -0.15 | -0.09 | 0.19 | 0.17 | 0.03 | 0.01 |
| **KIF23** | **S911** | -0.05 | -0.16 | -0.09 | 0.13 | -0.05 | 0.20 | 0.075 |
| **KIF23** | **T927** | -0.10 | -0.11 | 0.10 | 0.02 | -0.13 | 0.19 | 0.604 |
| **KIF23** | **T927** | -0.07 | -0.14 | 0.11 | -0.06 | -0.07 | 0.20 | 0.655 |
| **KIF23** | **S716** | 0.20 | 0.01 | 0.06 | 0.06 | -0.06 | -0.32 | 0.189 |
| **KIF23** | **S902** | 0.12 | 0.13 | 0.09 | -0.09 | -0.04 | -0.24 | 0.02 |
| **KIF23** | **S867** | 0.14 | 0.16 | 0.13 | -0.23 | -0.09 | -0.16 | 0.002 |
| **KIF23** | **S867** | 0.11 | 0.12 | 0.22 | -0.11 | -0.21 | -0.19 | 0.002 |
| **KIF23** | **S902** | 0.18 | 0.17 | 0.48 | -0.26 | -0.29 | -0.51 | 0.008 |
| **KIF24** | **S584** | -0.14 | -0.14 | 0.18 | 0.04 | -0.22 | 0.22 | 0.787 |
| **KIF24** | **S636; S646** | 0.21 | 0.17 | -0.39 | 0.61 | -0.16 | -0.87 | 0.782 |
| **KIF26A** | **S879** | -0.19 | -0.23 | -0.22 | 0.16 | -0.11 | 0.46 | 0.08 |
| **KIF26A** | **S/T** | -0.09 | -0.06 | -0.13 | 0.15 | 0.11 | 0.01 | 0.017 |
| **KIF2A** | **S586** | 0.03 | -0.20 | 0.03 | 0.13 | -0.11 | 0.10 | 0.458 |
| **KIF2A** | **S140** | -0.34 | 0.19 | -0.01 | 0.44 | -0.12 | -0.33 | 0.866 |
| **KIF2A** | **S/T** | 0.17 | -0.38 | 0.12 | 0.19 | -0.09 | -0.09 | 0.883 |
| **KIF2A** | **S140** | 0.01 | 0.43 | -0.12 | 0.02 | -0.27 | -0.18 | 0.256 |
| **KIF3B** | **T672** | 0.18 | 0.41 | 0.15 | -0.18 | -0.07 | -0.74 | 0.061 |
| **KIF4A** | **T1181** | -0.45 | -0.04 | -0.16 | 0.01 | 0.18 | 0.34 | 0.066 |
| **KIF4A** | **S1126** | -0.03 | -0.29 | 0.08 | -0.07 | 0.28 | -0.04 | 0.438 |
| **KIF4A** | **S1225** | 0.07 | -0.18 | -0.05 | 0.19 | -0.14 | 0.07 | 0.476 |
| **KIF4A** | **S961** | -0.40 | -0.22 | 0.45 | -0.21 | 0.01 | 0.20 | 0.851 |
| **KIF4A** | **T1181** | 0.07 | -0.12 | 0.14 | -0.48 | 0.18 | 0.11 | 0.688 |
| **KLC1** | **S460** | 0.06 | 0.13 | -0.02 | -0.04 | -0.12 | -0.03 | 0.081 |
| **KLC2** | **S521** | -0.16 | -0.03 | -0.07 | 0.04 | -0.04 | 0.23 | 0.132 |
| **KLC2** | **S582** | 0.03 | -0.17 | -0.19 | -0.19 | -0.17 | 0.54 | 0.542 |
| **KLC2** | **S521** | -0.16 | -0.01 | 0.01 | 0.08 | -0.12 | 0.16 | 0.406 |
| **KLC2** | **S445** | 0.10 | 0.33 | 0.22 | -0.50 | 0.00 | -0.33 | 0.039 |
| **KLF10** | **S249** | 0.39 | -0.26 | 0.04 | -0.56 | 0.04 | 0.16 | 0.576 |

| | | | | | | | | |
|---|---|---|---|---|---|---|---|---|
| KLF16 | S238; T/S | 0.10 | 0.21 | -0.14 | -0.05 | 0.02 | -0.17 | 0.338 |
| KLF16 | S109 | 0.11 | 0.02 | 0.04 | -0.35 | -0.15 | 0.26 | 0.493 |
| KLF16 | S/T | 0.06 | 0.13 | 0.16 | -0.20 | -0.13 | -0.06 | 0.007 |
| KLF2 | S177 | 0.01 | -0.13 | -0.04 | 0.10 | -0.17 | 0.20 | 0.446 |
| KLF3 | S250 | -0.04 | -0.08 | 0.05 | -0.05 | 0.02 | 0.09 | 0.523 |
| KLF3 | S92; S | 0.04 | 0.05 | -0.14 | 0.22 | 0.07 | -0.29 | 0.934 |
| KLF3 | S71 | 0.08 | 0.10 | 0.03 | -0.10 | 0.04 | -0.17 | 0.091 |
| KLF4 | S254 | -0.51 | -0.36 | -0.12 | 0.34 | -0.14 | 0.51 | 0.065 |
| KLF4 | S254 | -0.26 | -0.63 | 0.54 | -0.50 | 0.07 | 0.39 | 0.822 |
| KLHDC7A | S72 | -0.12 | -0.12 | 0.15 | 0.14 | -0.10 | 0.03 | 0.662 |
| KLHL22 | T605 | -0.05 | -0.25 | 0.03 | 0.06 | 0.27 | -0.11 | 0.298 |
| KLKB1 | S | 0.09 | -0.02 | -0.08 | 0.08 | 0.03 | -0.12 | 0.98 |
| KLKB1 | S | 0.13 | 0.22 | -0.13 | -0.09 | -0.10 | -0.07 | 0.208 |
| KLRG2 | S158 | -0.49 | -0.25 | -0.03 | 0.04 | 0.14 | 0.43 | 0.06 |
| KLRG2 | S204 | 0.16 | -0.25 | -0.31 | -0.13 | 0.13 | 0.30 | 0.294 |
| KMT2A | S1837; T1845 | -0.18 | -0.20 | -0.14 | 0.14 | -0.04 | 0.33 | 0.047 |
| KMT2A | S/T | -0.18 | -0.29 | -0.01 | -0.20 | 0.24 | 0.33 | 0.194 |
| KMT2A | T2147; S/T/Y | -0.49 | 0.04 | -0.03 | 0.33 | 0.06 | -0.03 | 0.226 |
| KMT2A | S2201 | -0.12 | -0.18 | -0.13 | 0.01 | 0.08 | 0.29 | 0.036 |
| KMT2A | S/T | -0.10 | -0.26 | -0.04 | -0.11 | 0.28 | 0.16 | 0.141 |
| KMT2A | S1685 | -0.19 | -0.09 | -0.03 | 0.03 | 0.13 | 0.12 | 0.027 |
| KMT2A | S153 | 0.12 | -0.15 | -0.25 | 0.08 | -0.02 | 0.17 | 0.246 |
| KMT2A | S/T | -0.23 | 0.03 | 0.02 | 0.08 | 0.03 | 0.05 | 0.26 |
| KMT2A | S2938 | 0.29 | -0.24 | -0.23 | 0.22 | -0.30 | 0.14 | 0.749 |
| KMT2A | S153 | -0.03 | -0.04 | -0.06 | 0.02 | -0.07 | 0.17 | 0.329 |
| KMT2A | S2156; S/T | 0.10 | -0.01 | -0.20 | -0.07 | 0.08 | 0.08 | 0.533 |
| KMT2A | S1837 | -0.05 | -0.03 | -0.02 | -0.01 | -0.03 | 0.13 | 0.289 |
| KMT2A | S/T | -0.05 | -0.02 | -0.04 | -0.12 | -0.23 | 0.38 | 0.812 |
| KMT2A | S2098 | 0.05 | 0.01 | -0.10 | -0.03 | -0.18 | 0.23 | 0.903 |
| KMT2A | S/T | 0.05 | 0.01 | -0.09 | 0.07 | -0.15 | 0.09 | 0.887 |
| KMT2A | S3644 | 0.04 | -0.02 | -0.01 | 0.01 | 0.21 | -0.27 | 0.873 |
| KMT2A | T337 | 0.14 | 0.07 | -0.19 | -0.06 | 0.05 | -0.05 | 0.803 |
| KMT2A | T506 | 0.10 | -0.03 | 0.03 | -0.10 | -0.23 | 0.19 | 0.586 |
| KMT2A | S/T | 0.05 | -0.10 | 0.22 | 0.04 | -0.06 | -0.19 | 0.309 |
| KMT2A | S2151 | 0.19 | -0.01 | 0.04 | -0.05 | -0.08 | -0.11 | 0.072 |
| KMT2A | S3511; S3518 | -0.03 | 0.01 | 0.33 | -0.65 | 0.01 | 0.16 | 0.384 |
| KMT2A | S161; S165 | 0.20 | 0.11 | 0.10 | -0.20 | -0.03 | -0.25 | 0.016 |
| KMT2A | S459 | -0.03 | -0.03 | 0.41 | -0.63 | -0.06 | 0.14 | 0.334 |
| KMT2A | S1119 | 0.15 | 0.22 | 0.06 | -0.27 | 0.31 | -0.70 | 0.291 |
| KMT2B | T/S | -0.11 | -0.05 | -0.43 | -0.17 | 0.12 | 0.48 | 0.199 |
| KMT2B | S2288 | -0.07 | -0.04 | -0.30 | 0.01 | 0.00 | 0.33 | 0.14 |
| KMT2B | T667; S687 | -0.41 | 0.02 | -0.07 | -0.28 | 0.03 | 0.52 | 0.423 |
| KMT2B | T588 | -0.22 | -0.05 | -0.21 | 0.30 | -0.51 | 0.46 | 0.479 |
| KMT2B | S1807 | -0.07 | -0.36 | 0.10 | 0.18 | 0.04 | 0.05 | 0.238 |
| KMT2B | S844 | -0.08 | -0.11 | -0.05 | 0.09 | -0.10 | 0.23 | 0.194 |
| KMT2B | S2348 | -0.04 | -0.11 | -0.03 | 0.08 | -0.14 | 0.21 | 0.339 |
| KMT2B | S1836 | -0.19 | -0.06 | 0.03 | -0.23 | -0.27 | 0.54 | 0.764 |
| KMT2B | S861 | -0.03 | -0.11 | 0.02 | 0.06 | -0.10 | 0.13 | 0.434 |

| | | | | | | | | |
|---|---|---|---|---|---|---|---|---|
| KMT2B | T172 | -0.01 | -0.06 | -0.02 | -0.12 | 0.05 | 0.15 | 0.496 |
| KMT2B | S493 | -0.02 | 0.03 | -0.08 | 0.07 | -0.25 | 0.20 | 0.828 |
| KMT2B | S632 | -0.02 | 0.02 | -0.03 | 0.11 | -0.06 | -0.03 | 0.781 |
| KMT2B | S118; S135; S136 | -0.09 | 0.00 | -0.02 | 0.16 | -0.64 | 0.40 | 0.979 |
| KMT2B | S1836 | 0.06 | -0.07 | 0.00 | -0.02 | 0.00 | 0.02 | 0.921 |
| KMT2B | S1035 | 0.03 | 0.10 | -0.13 | 0.00 | -0.04 | 0.03 | 0.988 |
| KMT2B | S1917 | 0.33 | -0.12 | -0.26 | -0.21 | -0.09 | 0.24 | 0.987 |
| KMT2B | S1095 | 0.08 | -0.09 | 0.05 | -0.02 | 0.04 | -0.06 | 0.692 |
| KMT2B | T2083 | 0.08 | 0.04 | -0.03 | -0.16 | -0.08 | 0.14 | 0.531 |
| KMT2B | S351 | -0.06 | 0.15 | 0.43 | -0.26 | -0.09 | -0.30 | 0.064 |
| KMT2C | S4267 | -0.13 | 0.02 | -0.18 | 0.17 | -0.01 | 0.10 | 0.084 |
| KMT2C | S | 0.02 | -0.16 | -0.15 | -0.09 | 0.13 | 0.21 | 0.169 |
| KMT2C | S4034 | 0.02 | -0.14 | -0.06 | 0.11 | -0.10 | 0.15 | 0.267 |
| KMT2C | S46 | -0.02 | -0.07 | 0.00 | 0.15 | -0.01 | -0.07 | 0.482 |
| KMT2C | S1987 | -0.04 | -0.08 | 0.05 | 0.23 | -0.11 | -0.08 | 0.758 |
| KMT2C | S200 | 0.14 | -0.36 | 0.17 | -0.21 | -0.04 | 0.21 | 0.99 |
| KMT2C | S4267 | 0.20 | -0.15 | -0.05 | -0.34 | 0.14 | 0.12 | 0.895 |
| KMT2C | T3977 | 0.00 | 0.08 | -0.01 | 0.07 | 0.04 | -0.20 | 0.554 |
| KMT2C | S3758 | -0.03 | 0.08 | 0.10 | -0.14 | -0.08 | 0.07 | 0.25 |
| KMT2C | S3758 | -0.01 | 0.01 | 0.24 | -0.05 | -0.07 | -0.17 | 0.114 |
| KMT2C | S/T | 0.18 | 0.30 | -0.10 | -0.75 | 0.43 | -0.38 | 0.383 |
| KMT2D | S4738 | -0.32 | -0.60 | -0.48 | 0.32 | 0.01 | 0.64 | 0.016 |
| KMT2D | T1865 | -0.07 | -0.15 | -0.23 | 0.05 | 0.07 | 0.27 | 0.03 |
| KMT2D | S2274 | -0.17 | -0.14 | 0.05 | 0.09 | 0.04 | 0.10 | 0.08 |
| KMT2D | T1843 | -0.14 | -0.01 | -0.10 | 0.13 | -0.15 | 0.23 | 0.266 |
| KMT2D | S2251 | -0.44 | 0.15 | 0.02 | 0.21 | -0.15 | 0.12 | 0.501 |
| KMT2D | S1671 | -0.04 | -0.01 | 0.00 | 0.04 | -0.05 | 0.07 | 0.448 |
| KMT2D | T/S | -0.08 | -0.03 | 0.07 | 0.03 | -0.06 | 0.06 | 0.68 |
| KMT2D | S/T | 0.30 | -0.10 | 0.11 | -0.06 | -0.15 | -0.17 | 0.126 |
| KMT2D | S373 | 0.12 | 0.36 | -0.12 | -0.27 | -0.09 | -0.08 | 0.153 |
| KMT2D | S/T | 0.50 | -0.38 | 0.17 | 0.07 | -0.87 | 0.14 | 0.484 |
| KMT2D | S3130 | 0.12 | 0.31 | 0.15 | 0.30 | -0.94 | -0.30 | 0.232 |
| KMT2E | S845 | -0.01 | 0.08 | -0.04 | 0.29 | -0.60 | 0.13 | 0.809 |
| KMT5A | T181 | -0.10 | 0.03 | -0.35 | 0.11 | 0.27 | -0.04 | 0.153 |
| KMT5A | T181 | -0.07 | -0.14 | -0.18 | -0.02 | 0.28 | 0.08 | 0.058 |
| KMT5B | S/T/Y | -0.20 | 0.28 | 0.01 | -0.09 | -0.40 | 0.27 | 0.679 |
| KMT5B | S532 | 0.07 | 0.09 | 0.08 | -0.04 | -0.21 | -0.02 | 0.045 |
| KMT5B | S726 | 0.22 | 0.13 | 0.10 | -0.15 | -0.15 | -0.19 | 0.001 |
| KMT5C | S290 | -0.04 | -0.04 | -0.12 | 0.16 | -0.10 | 0.12 | 0.209 |
| KMT5C | S290 | -0.16 | 0.22 | 0.15 | 0.16 | -0.02 | -0.47 | 0.464 |
| KMT5C | T416 | 0.39 | -0.57 | 0.33 | -0.93 | -0.12 | 0.39 | 0.614 |
| KNL1 | S60 | -0.47 | -0.18 | -0.20 | 0.23 | 0.29 | 0.18 | 0.007 |
| KNL1 | S1076 | -0.04 | -0.23 | 0.00 | 0.08 | 0.17 | -0.02 | 0.141 |
| KNL1 | S32 | -0.02 | 0.08 | -0.09 | 0.02 | -0.06 | 0.06 | 0.759 |
| KNL1 | T1017 | 0.25 | -0.98 | 0.63 | -0.63 | -0.53 | 0.51 | 0.782 |
| KNOP1 | S185 | 0.07 | -0.29 | -0.15 | 0.08 | -0.16 | 0.36 | 0.305 |
| KNOP1 | T/S | 0.14 | -0.02 | -0.09 | 0.00 | 0.03 | -0.07 | 0.767 |
| KNOP1 | S42 | 0.25 | 0.37 | 0.12 | -0.23 | -0.16 | -0.56 | 0.017 |

| | | | | | | | | |
|---|---|---|---|---|---|---|---|---|
| KPNA1 | S95 | -0.31 | -0.02 | -0.01 | 0.22 | -0.17 | 0.22 | 0.287 |
| KPNA2 | T/S | -0.07 | 0.02 | -0.01 | -0.06 | -0.07 | 0.17 | 0.68 |
| KPNA2 | S/T | -0.17 | -0.23 | 0.33 | -0.09 | -0.38 | 0.38 | 0.982 |
| KPNA2 | S62 | -0.02 | 0.04 | 0.14 | -0.10 | 0.06 | -0.13 | 0.204 |
| KPNA3 | S60 | 0.07 | 0.10 | 0.16 | -0.12 | 0.07 | -0.35 | 0.123 |
| KPNA6 | S113 | 0.26 | -0.04 | -0.12 | -0.14 | 0.05 | -0.04 | 0.582 |
| KRBA1 | S355 | 0.03 | 0.04 | 0.01 | -0.08 | -0.28 | 0.23 | 0.67 |
| KRI1 | S639; S | -0.32 | -0.15 | -0.08 | 0.22 | 0.13 | 0.13 | 0.009 |
| KRI1 | S171 | -0.22 | -0.15 | -0.18 | 0.23 | 0.13 | 0.12 | 0.001 |
| KRI1 | S628 | -0.05 | -0.06 | -0.18 | 0.13 | 0.10 | 0.04 | 0.02 |
| KRI1 | S628; S639 | -0.18 | -0.08 | -0.01 | 0.11 | 0.15 | -0.01 | 0.061 |
| KRI1 | S171 | -0.03 | -0.07 | -0.18 | 0.14 | -0.05 | 0.16 | 0.093 |
| KRI1 | S639 | -0.01 | 0.02 | -0.26 | 0.06 | 0.04 | 0.12 | 0.173 |
| KRI1 | S628 | -0.13 | 0.10 | -0.13 | 0.05 | 0.00 | 0.09 | 0.282 |
| KRT18 | S319 | -0.20 | -0.38 | -0.10 | 0.19 | 0.05 | 0.33 | 0.023 |
| KRT18 | S305 | -0.05 | 0.05 | -0.06 | 0.14 | 0.24 | -0.42 | 0.977 |
| KRT18 | S7 | 0.05 | -0.09 | 0.05 | 0.12 | 0.18 | -0.39 | 0.865 |
| KRT18 | S305 | -0.01 | -0.05 | 0.09 | -0.03 | 0.35 | -0.46 | 0.83 |
| KRT18 | S100 | 0.17 | 0.18 | -0.09 | -0.18 | 0.05 | -0.18 | 0.185 |
| KRT19 | S397 | -0.33 | -0.52 | 0.15 | 0.16 | 0.05 | 0.32 | 0.127 |
| KRT19 | S234 | -0.21 | -0.09 | -0.23 | 0.47 | 0.00 | -0.06 | 0.147 |
| KRT19 | S35 | -0.14 | -0.17 | -0.16 | 0.18 | -0.04 | 0.27 | 0.033 |
| KRT19 | S35; S40 | -0.03 | -0.44 | 0.08 | -0.09 | 0.06 | 0.32 | 0.312 |
| KRT19 | S397 | -0.05 | -0.08 | -0.21 | 0.20 | 0.07 | 0.04 | 0.04 |
| KRT19 | T323 | -0.14 | -0.06 | -0.08 | 0.10 | -0.18 | 0.30 | 0.296 |
| KRT19 | S14 | -0.11 | -0.09 | 0.05 | -0.10 | 0.18 | 0.05 | 0.398 |
| KRT19 | S93 | -0.01 | 0.07 | -0.08 | 0.07 | 0.07 | -0.13 | 0.898 |
| KRT19 | S397 | -0.13 | -0.14 | 0.27 | 0.00 | 0.16 | -0.23 | 0.914 |
| KRT23 | S65 | -0.10 | 0.14 | 0.00 | 0.02 | 0.21 | -0.35 | 0.779 |
| KRT5 | S75 | -0.23 | -0.31 | -0.04 | -0.17 | 0.17 | 0.44 | 0.15 |
| KRT5 | S181 | 0.02 | -0.15 | 0.15 | -0.17 | 0.31 | -0.25 | 0.842 |
| KRT7 | S254 | -0.13 | -0.30 | -0.43 | 0.34 | 0.09 | 0.27 | 0.01 |
| KRT8 | S24 | -1.81 | -1.26 | -1.11 | 1.42 | 0.25 | -0.05 | 0.018 |
| KRT8 | S280 | -0.18 | -0.37 | -0.32 | 0.23 | 0.00 | 0.46 | 0.022 |
| KRT8 | S258 | -0.10 | -0.26 | -0.17 | 0.15 | 0.29 | 0.01 | 0.024 |
| KRT8 | S280 | -0.19 | -0.16 | -0.16 | 0.20 | 0.24 | 0.00 | 0.014 |
| KRT8 | S253 | -0.07 | -0.15 | -0.20 | 0.19 | 0.16 | 0.03 | 0.013 |
| KRT8 | S475 | 0.02 | -0.19 | -0.20 | 0.14 | 0.26 | -0.10 | 0.157 |
| KRT8 | S31 | 0.03 | -0.12 | -0.09 | 0.08 | 0.19 | -0.12 | 0.339 |
| KRT8 | S291 | -0.03 | -0.05 | -0.08 | 0.18 | 0.22 | -0.31 | 0.658 |
| KRT8 | S330 | 0.06 | -0.04 | -0.10 | 0.04 | 0.08 | -0.06 | 0.508 |
| KRT8 | S34 | 0.11 | -0.03 | 0.03 | -0.14 | 0.16 | -0.15 | 0.51 |
| KRT82 | S133 | 0.33 | -0.17 | -0.31 | -0.54 | 0.74 | -0.53 | 0.91 |
| KRTCAP3 | S214 | 0.15 | -0.05 | -0.10 | -0.19 | 0.13 | 0.02 | 0.908 |
| KRTCAP3 | S214 | 0.15 | 0.06 | -0.05 | -0.18 | 0.08 | -0.08 | 0.301 |
| KTN1 | T153 | -0.13 | 0.03 | -0.06 | 0.21 | -0.05 | -0.01 | 0.316 |
| KTN1 | S75 | 0.10 | 0.20 | -0.33 | 0.00 | -0.11 | 0.09 | 0.979 |
| KTN1 | S75; S77 | -0.05 | 0.02 | 0.04 | 0.07 | -0.04 | -0.04 | 0.906 |

| | | | | | | | | |
|---|---|---|---|---|---|---|---|---|
| **KTN1** | **S75** | -0.02 | 0.03 | 0.01 | 0.03 | 0.00 | -0.05 | 0.653 |
| **KTN1** | **S174** | -0.14 | 0.17 | 0.01 | 0.05 | -0.15 | 0.03 | 0.759 |
| **KTN1** | **S75** | 0.00 | 0.10 | 0.09 | -0.03 | -0.07 | -0.10 | 0.027 |
| **KTN1** | **S75** | -0.05 | 0.13 | 0.13 | -0.17 | -0.21 | 0.13 | 0.295 |
| **KTN1** | **S75** | 0.14 | 0.02 | 0.11 | -0.25 | -0.02 | -0.03 | 0.078 |
| **KTN1** | **T153** | 0.09 | 0.09 | 0.10 | -0.09 | 0.03 | -0.25 | 0.069 |
| **KTN1** | **S75; S79** | 0.05 | 0.14 | 0.12 | -0.09 | -0.03 | -0.22 | 0.026 |
| **KTN1** | **S/T** | -0.13 | 0.25 | 0.07 | -0.80 | 0.49 | -0.20 | 0.571 |
| **KTN1** | **S108** | -0.13 | 0.25 | 0.07 | -0.80 | 0.49 | -0.20 | 0.571 |
| **KTN1** | **S75** | 0.05 | 0.21 | 0.27 | -0.19 | -0.14 | -0.31 | 0.009 |
| **L1RE1** | **S/T** | -0.06 | 0.12 | -0.14 | 0.05 | -0.15 | 0.14 | 0.753 |
| **L1RE1** | **S/T** | -0.04 | -0.10 | 0.23 | -0.34 | -0.10 | 0.26 | 0.677 |
| **L1RE1** | **S18** | 0.04 | 0.09 | 0.32 | -0.24 | 0.02 | -0.34 | 0.073 |
| **L3MBTL2** | **S689** | -0.35 | 0.06 | -0.34 | 0.23 | -0.35 | 0.51 | 0.304 |
| **LAD1** | **S177** | -0.01 | 0.01 | -0.25 | 0.05 | -0.04 | 0.21 | 0.223 |
| **LAD1** | **S301** | -0.22 | 0.03 | -0.02 | -0.15 | 0.03 | 0.29 | 0.437 |
| **LAD1** | **S301** | -0.10 | 0.06 | -0.13 | 0.05 | -0.05 | 0.15 | 0.278 |
| **LAD1** | **S272** | 0.09 | 0.21 | -0.11 | 0.04 | -0.01 | -0.25 | 0.355 |
| **LAD1** | **S123** | 0.34 | 0.13 | -0.27 | 0.04 | 0.00 | -0.36 | 0.467 |
| **LAD1** | **S356** | 0.10 | 0.24 | 0.03 | -0.14 | -0.04 | -0.24 | 0.035 |
| **LAD1** | **S356** | 0.13 | 0.23 | 0.04 | -0.13 | -0.05 | -0.28 | 0.03 |
| **LAMTOR1** | **S27** | -0.08 | 0.33 | -0.93 | 0.51 | -0.26 | 0.02 | 0.506 |
| **LAMTOR1** | **S27** | -0.21 | -0.15 | -0.09 | 0.06 | 0.15 | 0.19 | 0.005 |
| **LAMTOR1** | **S42** | -0.29 | 0.25 | 0.05 | -0.14 | -0.11 | 0.16 | 0.853 |
| **LAMTOR1** | **S56** | -0.13 | 0.34 | 0.09 | -0.21 | 0.07 | -0.25 | 0.243 |
| **LAMTOR1** | **S42** | 0.06 | 0.44 | 0.08 | -0.10 | -0.26 | -0.36 | 0.039 |
| **LARP1** | **S1040** | -0.42 | -0.32 | -0.25 | 0.44 | -0.16 | 0.44 | 0.053 |
| **LARP1** | **T724** | -0.18 | -0.15 | -0.10 | 0.03 | 0.13 | 0.22 | 0.01 |
| **LARP1** | **T68** | -0.21 | 0.12 | -0.11 | 0.27 | -0.19 | 0.06 | 0.545 |
| **LARP1** | **S548** | 0.00 | -0.15 | -0.01 | 0.12 | 0.00 | 0.03 | 0.148 |
| **LARP1** | **S766; S774** | -0.04 | -0.14 | 0.04 | 0.12 | -0.05 | 0.05 | 0.269 |
| **LARP1** | **S75; S90** | -0.03 | -0.03 | -0.05 | -0.04 | -0.04 | 0.18 | 0.368 |
| **LARP1** | **T724** | -0.27 | -0.04 | 0.17 | 0.07 | -0.17 | 0.19 | 0.68 |
| **LARP1** | **S** | -0.41 | -0.07 | 0.30 | 0.07 | -0.38 | 0.32 | 0.841 |
| **LARP1** | **S631** | -0.06 | -0.13 | 0.13 | -0.01 | 0.10 | -0.04 | 0.677 |
| **LARP1** | **S774** | 0.08 | 0.01 | 0.03 | 0.08 | 0.02 | -0.24 | 0.449 |
| **LARP1** | **T526** | 0.08 | 0.07 | 0.01 | 0.03 | -0.12 | -0.08 | 0.104 |
| **LARP1** | **T788; T** | -0.02 | -0.03 | 0.24 | -0.27 | 0.06 | -0.02 | 0.339 |
| **LARP1** | **S75** | 0.02 | 0.13 | 0.11 | -0.12 | -0.06 | -0.10 | 0.009 |
| **LARP1** | **T/S** | 0.04 | 0.13 | 0.13 | -0.09 | -0.02 | -0.23 | 0.037 |
| **LARP1** | **S90** | -0.02 | 0.16 | 0.14 | -0.11 | -0.52 | 0.22 | 0.351 |
| **LARP4** | **S597** | -0.38 | -0.28 | -0.23 | 0.26 | -0.21 | 0.58 | 0.097 |
| **LARP4** | **S597** | -0.09 | -0.25 | -0.07 | 0.67 | -0.47 | -0.07 | 0.633 |
| **LARP4** | **S722** | 0.09 | 0.06 | 0.06 | 0.00 | -0.06 | -0.16 | 0.043 |
| **LARP4** | **S647** | 0.09 | -0.11 | 0.22 | -0.19 | 0.10 | -0.16 | 0.318 |
| **LARP4** | **S647** | 0.14 | -0.07 | 0.19 | 0.06 | 0.22 | -0.76 | 0.474 |
| **LARP4** | **S647** | 0.10 | 0.46 | -0.04 | -0.31 | 0.35 | -1.00 | 0.303 |
| **LARP4B** | **S601** | -0.07 | 0.08 | 0.02 | 0.03 | -0.07 | 0.00 | 0.745 |

| | | | | | | | | |
|---|---|---|---|---|---|---|---|---|
| LARP4B | S736 | 0.06 | 0.00 | 0.09 | -0.02 | 0.04 | -0.20 | 0.219 |
| LARP4B | S664 | 0.05 | 0.09 | 0.13 | -0.13 | -0.09 | -0.07 | 0.003 |
| LARP4B | S736; S/T | 0.16 | 0.14 | 0.07 | -0.09 | -0.02 | -0.32 | 0.05 |
| LARP7 | S/T | 0.04 | -0.08 | -0.07 | 0.11 | 0.05 | -0.05 | 0.291 |
| LARP7 | S337; T338 | 0.05 | 0.06 | -0.10 | 0.12 | -0.11 | -0.02 | 0.954 |
| LARP7 | S261 | 0.02 | 0.03 | 0.02 | -0.03 | -0.07 | 0.02 | 0.16 |
| LARP7 | S261 | 0.07 | 0.10 | 0.09 | -0.14 | -0.08 | -0.06 | 0.002 |
| LARP7 | S258; S261 | 0.12 | 0.08 | 0.22 | -0.13 | -0.12 | -0.22 | 0.005 |
| LARP7 | S | 0.14 | 0.07 | 0.23 | -0.03 | -0.11 | -0.39 | 0.048 |
| LAS1L | S560 | -0.43 | -0.49 | -0.32 | -0.22 | 0.14 | 0.83 | 0.102 |
| LAS1L | S560 | -0.27 | -0.25 | -0.18 | -0.08 | 0.07 | 0.54 | 0.094 |
| LAS1L | S560 | -0.13 | -0.13 | -0.16 | -0.03 | 0.11 | 0.28 | 0.043 |
| LAS1L | S523 | 0.26 | -0.59 | -0.31 | -0.04 | -0.66 | 0.79 | 0.64 |
| LAS1L | S617 | 0.03 | -0.56 | 0.05 | -0.02 | -0.28 | 0.54 | 0.49 |
| LAS1L | S560 | 0.07 | -0.21 | -0.06 | -0.15 | 0.34 | -0.05 | 0.541 |
| LAS1L | S560 | 0.02 | -0.05 | -0.11 | 0.00 | -0.03 | 0.15 | 0.258 |
| LAS1L | S617 | -0.16 | 0.00 | 0.08 | -0.46 | 0.20 | 0.23 | 0.946 |
| LAS1L | S560 | 0.09 | 0.06 | -0.16 | 0.05 | -0.03 | -0.02 | 0.965 |
| LAS1L | S617 | -0.03 | 0.06 | 0.02 | 0.08 | 0.13 | -0.31 | 0.74 |
| LAS1L | S617 | -0.01 | 0.01 | 0.08 | -0.14 | 0.00 | 0.04 | 0.372 |
| LASP1 | S146 | 0.04 | -0.09 | -0.25 | 0.05 | 0.33 | -0.14 | 0.321 |
| LATS1 | S464 | -0.58 | -0.09 | -0.42 | -0.03 | 0.45 | 0.37 | 0.04 |
| LATS1 | T246 | -0.03 | 0.00 | -0.20 | 0.00 | 0.26 | -0.07 | 0.311 |
| LATS2 | T/S | 0.07 | -0.32 | -0.15 | -0.01 | -0.30 | 0.53 | 0.482 |
| LBR | S86 | 0.01 | 0.03 | 0.00 | 0.02 | -0.10 | 0.02 | 0.462 |
| LBR | S84; S86 | 0.08 | 0.05 | -0.04 | 0.00 | -0.03 | -0.06 | 0.212 |
| LBR | S99 | 0.14 | 0.11 | -0.02 | 0.04 | -0.17 | -0.12 | 0.121 |
| LBR | S99 | 0.13 | 0.03 | 0.10 | 0.02 | -0.14 | -0.17 | 0.046 |
| LBR | S99 | 0.09 | 0.06 | 0.16 | 0.00 | -0.29 | -0.07 | 0.074 |
| LBR | T200 | 0.28 | -0.06 | 0.09 | -0.13 | 0.12 | -0.39 | 0.251 |
| LBR | S86 | 0.16 | 0.08 | 0.17 | -0.24 | 0.06 | -0.30 | 0.059 |
| LBR | S67; S71 | 0.17 | 0.21 | 0.17 | -0.18 | -0.03 | -0.46 | 0.034 |
| LBR | S71 | 0.16 | 0.17 | 0.54 | -0.89 | -0.09 | -0.29 | 0.058 |
| LBR | S67 | 0.35 | 0.29 | 0.31 | -0.44 | -0.08 | -0.79 | 0.022 |
| LBR | S | 0.67 | 0.09 | 0.46 | -0.76 | -0.34 | -0.76 | 0.01 |
| LCOR | S426 | -0.26 | 0.02 | 0.22 | -0.12 | 0.05 | 0.05 | 0.997 |
| LCOR | S249 | 0.11 | 0.16 | -0.24 | 0.25 | 0.00 | -0.39 | 0.813 |
| LCOR | S311 | -0.30 | 0.27 | 0.21 | -0.02 | 0.10 | -0.39 | 0.52 |
| LDB1 | T61 | -0.11 | -0.03 | 0.25 | 0.02 | -0.24 | 0.06 | 0.546 |
| LDLRAP1 | S186 | -0.29 | -0.48 | -0.38 | 0.28 | 0.32 | 0.30 | 3E-04 |
| LEMD2 | S453 | -0.26 | -0.17 | 0.01 | 0.03 | 0.14 | 0.19 | 0.047 |
| LEMD2 | S134; S138; S139 | -0.10 | -0.09 | -0.20 | 0.27 | -0.15 | 0.20 | 0.158 |
| LEMD2 | S166 | 0.10 | -0.17 | 0.07 | -0.10 | 0.25 | -0.20 | 0.91 |
| LEMD2 | S138; S139 | -0.14 | 0.31 | 0.07 | -0.17 | 0.05 | -0.20 | 0.286 |
| LEMD2 | S499 | 0.21 | 0.14 | 0.18 | -0.26 | 0.03 | -0.40 | 0.04 |
| LEMD2 | S82 | 0.21 | 0.22 | 0.27 | -0.37 | -0.10 | -0.39 | 0.005 |
| LEMD3 | S187 | -0.07 | -0.26 | -0.25 | 0.15 | 0.03 | 0.31 | 0.027 |
| LEMD3 | S | -0.10 | -0.10 | -0.27 | -0.11 | 0.01 | 0.45 | 0.203 |

| | | | | | | | | |
|---|---|---|---|---|---|---|---|---|
| **LEMD3** | **S/T** | -0.12 | -0.36 | 0.07 | 0.11 | 0.10 | 0.13 | 0.11 |
| **LEMD3** | **S352** | -0.09 | -0.14 | -0.19 | 0.02 | -0.12 | 0.42 | 0.203 |
| **LEMD3** | **S352** | 0.04 | -0.06 | -0.08 | -0.01 | -0.19 | 0.27 | 0.68 |
| **LEMD3** | **T209** | -0.47 | -0.39 | 0.69 | -0.25 | -0.02 | 0.11 | 0.992 |
| **LEMD3** | **S144** | -0.02 | -0.01 | 0.10 | -0.06 | -0.09 | 0.07 | 0.514 |
| **LENG8** | **S351** | 0.09 | 0.00 | 0.05 | -0.02 | -0.12 | -0.01 | 0.092 |
| **LEO1** | **S658** | 0.04 | -0.24 | -0.33 | 0.20 | -0.05 | 0.28 | 0.097 |
| **LEO1** | **S630** | 0.18 | 0.02 | -0.10 | -0.01 | 0.05 | -0.17 | 0.486 |
| **LEO1** | **S179** | 0.24 | 0.06 | -0.05 | -0.09 | 0.00 | -0.20 | 0.162 |
| **LEO1** | **S614** | 0.04 | 0.12 | 0.21 | -0.09 | -0.10 | -0.22 | 0.017 |
| **LEO1** | **S614** | 0.04 | 0.12 | 0.21 | -0.09 | -0.10 | -0.22 | 0.017 |
| **LEO1** | **S630** | 0.26 | 0.16 | -0.02 | -0.16 | 0.09 | -0.43 | 0.153 |
| **LEO1** | **S610** | 0.12 | 0.27 | 0.11 | -0.36 | 0.09 | -0.35 | 0.079 |
| **LEO1** | **S610; S614** | 0.44 | 0.24 | 0.28 | -0.24 | -0.16 | -0.96 | 0.041 |
| **LETM1** | **T469** | -0.02 | -0.20 | -0.08 | 0.18 | -0.35 | 0.35 | 0.512 |
| **LIG1** | **S141** | 0.07 | -0.05 | -0.05 | -0.12 | -0.09 | 0.22 | 0.907 |
| **LIG1** | **S51** | 0.15 | -0.01 | -0.09 | -0.16 | -0.13 | 0.20 | 0.736 |
| **LIG1** | **S51** | 0.17 | 0.04 | 0.04 | -0.08 | -0.30 | 0.09 | 0.207 |
| **LIG1** | **S66** | 0.11 | 0.14 | 0.05 | -0.15 | -0.11 | -0.07 | 0.004 |
| **LIG1** | **T183** | 0.33 | 0.24 | 0.04 | -0.21 | -0.11 | -0.42 | 0.022 |
| **LIG1** | **T183** | 0.28 | 0.19 | 0.16 | -0.20 | -0.33 | -0.22 | 0.001 |
| **LIG1** | **S66; S76** | 0.27 | 0.23 | 0.13 | -0.32 | -0.19 | -0.24 | 0.001 |
| **LIG1** | **S141** | 0.25 | 0.23 | 0.20 | -0.24 | -0.14 | -0.43 | 0.004 |
| **LIG3** | **S853** | -0.19 | -0.26 | -0.27 | 0.24 | 0.01 | 0.35 | 0.013 |
| **LIG3** | **S853** | -0.09 | -0.05 | -0.26 | 0.13 | 0.09 | 0.14 | 0.019 |
| **LIG3** | **S853** | -0.06 | 0.07 | -0.18 | 0.03 | 0.03 | 0.09 | 0.23 |
| **LIG3** | **S248** | -0.05 | -0.10 | 0.12 | 0.05 | 0.07 | -0.09 | 0.829 |
| **LIG3** | **S242** | 0.19 | 0.20 | 0.41 | -0.74 | 0.07 | -0.45 | 0.059 |
| **LIG3** | **S210** | 0.23 | 0.25 | 0.38 | -0.31 | -0.13 | -0.69 | 0.018 |
| **LIMA1** | **S686; S692** | -0.21 | -0.42 | -0.41 | 0.27 | 0.06 | 0.47 | 0.011 |
| **LIMA1** | **S686; S692** | -0.12 | -0.28 | -0.34 | 0.25 | 0.11 | 0.26 | 0.005 |
| **LIMA1** | **S686** | -0.32 | -0.07 | -0.21 | 0.11 | -0.13 | 0.47 | 0.14 |
| **LIMA1** | **S671; S692** | -0.03 | -0.23 | -0.32 | 0.11 | -0.22 | 0.51 | 0.224 |
| **LIMA1** | **S692; S698** | -0.21 | -0.18 | -0.06 | 0.18 | 0.15 | 0.07 | 0.008 |
| **LIMA1** | **S617** | -0.09 | -0.33 | -0.04 | 0.26 | -0.03 | 0.15 | 0.088 |
| **LIMA1** | **S617** | 0.06 | -0.01 | -0.48 | 0.19 | 0.17 | -0.04 | 0.249 |
| **LIMA1** | **S230** | -0.27 | -0.15 | 0.13 | -0.07 | 0.11 | 0.19 | 0.282 |
| **LIMA1** | **S230** | -0.27 | -0.15 | 0.13 | -0.07 | 0.11 | 0.19 | 0.282 |
| **LIMA1** | **S686** | -0.09 | -0.06 | -0.12 | 0.13 | 0.00 | 0.13 | 0.018 |
| **LIMA1** | **S132** | 0.07 | -0.11 | -0.20 | -0.11 | 0.02 | 0.27 | 0.379 |
| **LIMA1** | **S686** | 0.02 | -0.03 | -0.14 | 0.11 | -0.08 | 0.11 | 0.278 |
| **LIMA1** | **S686** | -0.09 | -0.11 | 0.05 | 0.17 | -0.15 | 0.11 | 0.461 |
| **LIMA1** | **S686** | -0.08 | 0.03 | -0.09 | 0.18 | -0.04 | -0.02 | 0.341 |
| **LIMA1** | **S726** | 0.09 | -0.15 | -0.08 | 0.14 | 0.12 | -0.15 | 0.522 |
| **LIMA1** | **S225** | 0.11 | 0.01 | -0.25 | 0.16 | -0.14 | 0.08 | 0.615 |
| **LIMA1** | **S132** | -0.15 | 0.01 | 0.06 | 0.03 | 0.04 | 0.00 | 0.46 |
| **LIMA1** | **S686** | -0.04 | 0.03 | -0.07 | 0.13 | -0.10 | 0.03 | 0.546 |
| **LIMA1** | **S371; S** | 0.01 | -0.08 | -0.01 | 0.06 | -0.08 | 0.08 | 0.492 |

| | | | | | | | | |
|---|---|---|---|---|---|---|---|---|
| **LIMA1** | **S** | 0.03 | -0.04 | -0.01 | 0.10 | -0.04 | -0.04 | 0.837 |
| **LIMA1** | **S490** | 0.01 | 0.03 | -0.04 | 0.06 | -0.11 | 0.03 | 0.868 |
| **LIMA1** | **S698** | 0.03 | 0.11 | 0.11 | -0.09 | 0.00 | -0.19 | 0.042 |
| **LIMA1** | **S225** | 0.29 | 0.08 | -0.05 | -0.02 | -0.29 | -0.08 | 0.137 |
| **LIMA1** | **S609** | 0.10 | 0.17 | 0.26 | -0.10 | -0.17 | -0.36 | 0.013 |
| **LIMA1** | **S362** | 0.18 | 0.37 | 0.28 | -0.24 | -0.13 | -0.71 | 0.027 |
| **LIMCH1** | **S973** | -0.22 | -0.01 | -0.30 | 0.24 | -0.12 | 0.30 | 0.112 |
| **LIMCH1** | **S875** | -0.09 | -0.29 | -0.14 | 0.36 | -0.03 | 0.11 | 0.071 |
| **LIMCH1** | **S493** | -0.22 | -0.06 | -0.20 | -0.07 | 0.09 | 0.38 | 0.105 |
| **LIMCH1** | **S973** | -0.07 | -0.08 | -0.03 | 0.19 | -0.22 | 0.16 | 0.479 |
| **LIMCH1** | **S718** | -0.05 | 0.09 | -0.16 | 0.19 | -0.09 | 0.00 | 0.538 |
| **LIMCH1** | **S377** | 0.10 | -0.04 | -0.17 | 0.42 | -0.15 | -0.26 | 0.854 |
| **LIMCH1** | **S585** | -0.18 | 0.29 | -0.25 | 0.23 | 0.29 | -0.60 | 0.96 |
| **LIMCH1** | **S201** | -0.21 | 0.14 | 0.09 | -0.12 | 0.02 | 0.05 | 0.866 |
| **LIMCH1** | **S523** | -0.03 | 0.07 | 0.07 | 0.05 | -0.10 | -0.07 | 0.255 |
| **LIMCH1** | **T215; S217** | -0.02 | 0.04 | 0.18 | -0.08 | -0.03 | -0.10 | 0.091 |
| **LIMCH1** | **S523** | 0.25 | -0.21 | 0.18 | 0.00 | -0.12 | -0.17 | 0.334 |
| **LIMCH1** | **S201** | 0.18 | -0.04 | 0.21 | -0.04 | 0.04 | -0.44 | 0.194 |
| **LIMCH1** | **S169** | 0.09 | 0.16 | 0.16 | -0.05 | -0.19 | -0.22 | 0.008 |
| **LIMCH1** | **S231** | 0.13 | 0.16 | 0.17 | -0.15 | -0.19 | -0.18 | 3E-05 |
| **LIMCH1** | **S231** | 0.12 | 0.17 | 0.19 | -0.14 | -0.18 | -0.22 | 4E-04 |
| **LIMCH1** | **S601** | 0.02 | 0.47 | 0.03 | -0.35 | 0.02 | -0.37 | 0.105 |
| **LIMCH1** | **S169** | 0.19 | 0.18 | 0.19 | -0.18 | -0.14 | -0.34 | 0.003 |
| **LIMCH1** | **S217** | 0.21 | 0.21 | 0.30 | -0.26 | -0.10 | -0.55 | 0.016 |
| **LIMCH1** | **T/S** | 0.28 | -0.03 | 0.82 | -1.50 | 0.03 | 0.03 | 0.216 |
| **LIMK2** | **S** | -0.13 | -0.45 | -0.76 | 0.02 | 0.64 | 0.25 | 0.043 |
| **LIN37** | **S135** | -0.39 | -0.36 | -0.10 | 0.27 | -0.01 | 0.41 | 0.03 |
| **LIN37** | **S135; S137** | -0.17 | -0.06 | -0.07 | 0.00 | 0.18 | 0.09 | 0.039 |
| **LIN37** | **S182** | -0.52 | 0.11 | 0.06 | -0.14 | 0.54 | -0.29 | 0.66 |
| **LIN37** | **S135; S137** | -0.11 | 0.08 | -0.16 | 0.04 | 0.04 | 0.09 | 0.196 |
| **LIN37** | **T167** | -0.57 | -0.01 | 0.31 | -0.10 | -0.25 | 0.40 | 0.748 |
| **LIN37** | **S138** | 0.04 | -0.10 | -0.08 | -0.04 | 0.04 | 0.13 | 0.243 |
| **LIN37** | **S182** | 0.36 | -0.36 | -0.23 | 0.52 | -0.79 | 0.10 | 0.967 |
| **LIN37** | **S182; S202** | -0.18 | 0.21 | -0.06 | -0.17 | -0.01 | 0.16 | 0.991 |
| **LIN37** | **S182** | 0.04 | -0.07 | 0.13 | 0.11 | -0.15 | -0.08 | 0.507 |
| **LIN52** | **S53** | -0.22 | -0.12 | -0.10 | 0.20 | -0.01 | 0.19 | 0.025 |
| **LIN54** | **S264** | -0.33 | -0.23 | 0.31 | 0.23 | -0.25 | 0.14 | 0.642 |
| **LIPE** | **S/T** | -0.01 | -0.04 | -0.07 | -0.01 | -0.07 | 0.19 | 0.408 |
| **LLGL2** | **S680** | -0.14 | -0.03 | 0.18 | 0.08 | 0.04 | -0.15 | 0.914 |
| **LMBRD1** | **S528; S531** | 0.02 | -0.03 | 0.16 | -0.07 | -0.24 | 0.12 | 0.402 |
| **LMF2** | **S670** | 0.09 | -0.08 | -0.10 | 0.69 | -0.49 | -0.44 | 0.904 |
| **LMNA** | **S414** | -0.32 | -0.48 | -0.43 | 0.40 | -0.48 | 0.76 | 0.165 |
| **LMNA** | **S406; S/T** | -0.17 | -0.24 | -0.35 | 0.13 | -0.34 | 0.68 | 0.245 |
| **LMNA** | **S51** | -0.17 | -0.21 | -0.18 | 0.19 | 0.34 | -0.06 | 0.042 |
| **LMNA** | **S636** | -0.09 | -0.21 | -0.18 | 0.17 | 0.24 | 0.00 | 0.021 |
| **LMNA** | **S/T** | 0.19 | -0.49 | -0.17 | 0.12 | 0.05 | 0.18 | 0.242 |
| **LMNA** | **S403** | -0.07 | -0.30 | -0.14 | 0.36 | -0.48 | 0.40 | 0.427 |
| **LMNA** | **S429** | -0.10 | -0.18 | -0.01 | 0.17 | 0.06 | 0.04 | 0.039 |

| | | | | | | | | |
|---|---|---|---|---|---|---|---|---|
| LMNA | S/T | 0.09 | -0.21 | -0.15 | 0.14 | -0.34 | 0.35 | 0.568 |
| LMNA | S599 | 0.09 | -0.15 | -0.09 | -0.05 | 0.15 | 0.02 | 0.381 |
| LMNA | S533 | -0.15 | -0.16 | 0.15 | 0.00 | 0.32 | -0.23 | 0.681 |
| LMNA | S/T | -0.14 | -0.18 | 0.14 | 0.18 | -0.44 | 0.31 | 0.769 |
| LMNA | S618; S619 | 0.16 | -0.25 | -0.03 | 0.30 | -0.23 | -0.03 | 0.807 |
| LMNA | S601 | 0.02 | -0.08 | -0.01 | -0.03 | -0.04 | 0.13 | 0.464 |
| LMNA | S149 | -0.03 | -0.04 | -0.03 | 0.11 | -0.32 | 0.24 | 0.812 |
| LMNA | S406; S407 | 0.06 | -0.09 | -0.02 | 0.01 | -0.35 | 0.31 | 0.964 |
| LMNA | S107 | 0.27 | -0.09 | -0.22 | -0.13 | 0.12 | 0.00 | 0.978 |
| LMNA | S403; S/T | 0.10 | 0.01 | -0.12 | 0.16 | -0.07 | -0.11 | 0.994 |
| LMNA | S18; T19 | -0.18 | 0.01 | 0.19 | 0.04 | 0.03 | -0.13 | 0.845 |
| LMNA | S392 | 0.00 | -0.08 | 0.12 | -0.07 | 0.03 | 0.00 | 0.668 |
| LMNA | S525 | 0.08 | 0.05 | -0.08 | -0.07 | 0.11 | -0.11 | 0.656 |
| LMNA | S533 | 0.01 | 0.00 | 0.10 | -0.12 | 0.03 | -0.02 | 0.246 |
| LMNA | S392 | -0.03 | -0.01 | 0.15 | -0.04 | -0.04 | -0.04 | 0.234 |
| LMNA | S423 | 0.11 | -0.03 | 0.04 | 0.07 | -0.14 | -0.06 | 0.3 |
| LMNA | S390; S392 | 0.11 | 0.01 | 0.04 | -0.06 | -0.03 | -0.08 | 0.031 |
| LMNA | S390 | 0.09 | 0.13 | -0.05 | -0.07 | 0.03 | -0.14 | 0.179 |
| LMNA | S71 | 0.14 | 0.04 | 0.00 | -0.13 | 0.08 | -0.15 | 0.207 |
| LMNA | S390 | 0.14 | 0.04 | 0.02 | -0.07 | 0.02 | -0.16 | 0.094 |
| LMNA | S458 | -0.11 | 0.19 | 0.10 | -0.05 | 0.20 | -0.41 | 0.514 |
| LMNA | S463 | 0.08 | 0.06 | 0.09 | -0.22 | -0.01 | -0.03 | 0.078 |
| LMNA | S22 | -0.05 | 0.18 | 0.13 | 0.00 | -0.12 | -0.18 | 0.096 |
| LMNA | S403; S404; S406 | 0.22 | 0.05 | 0.01 | -0.09 | -0.17 | -0.04 | 0.06 |
| LMNA | S301 | 0.15 | -0.16 | 0.30 | 0.00 | -0.09 | -0.26 | 0.244 |
| LMNA | S390; S392 | 0.22 | 0.03 | 0.07 | -0.18 | -0.16 | -0.02 | 0.045 |
| LMNA | S12 | 0.11 | 0.07 | 0.18 | -0.17 | 0.01 | -0.24 | 0.037 |
| LMNA | S403; S406 | 0.10 | 0.39 | -0.14 | 0.00 | -0.27 | -0.17 | 0.208 |
| LMNA | S403; S404; T409 | 0.24 | -0.03 | 0.12 | -0.07 | -0.52 | 0.13 | 0.279 |
| LMNA | S463 | 0.16 | 0.32 | 0.01 | -0.44 | 0.03 | -0.20 | 0.088 |
| LMNA | S66 | -0.01 | 0.06 | 0.65 | -0.45 | -0.31 | -0.23 | 0.061 |
| LMNB1 | S375 | -0.50 | -0.02 | -0.04 | 0.09 | 0.06 | 0.29 | 0.123 |
| LMNB1 | T575 | -0.26 | -0.11 | 0.01 | 0.24 | -0.11 | 0.16 | 0.177 |
| LMNB1 | S210 | 0.09 | -0.11 | -0.21 | 0.14 | -0.10 | 0.15 | 0.326 |
| LMNB1 | T575 | -0.10 | -0.04 | 0.09 | 0.04 | -0.06 | 0.06 | 0.639 |
| LMNB1 | S23; T | -0.05 | -0.06 | 0.05 | 0.16 | 0.12 | -0.26 | 0.867 |
| LMNB1 | S58 | -0.17 | 0.07 | 0.07 | 0.12 | -0.24 | 0.11 | 0.99 |
| LMNB1 | S284 | 0.21 | 0.10 | -0.24 | -0.36 | 0.04 | 0.15 | 0.73 |
| LMNB1 | S210 | 0.20 | 0.06 | -0.13 | 0.03 | 0.01 | -0.21 | 0.474 |
| LMNB1 | S232 | -0.02 | 0.05 | 0.12 | -0.19 | 0.06 | -0.05 | 0.25 |
| LMNB1 | S23 | 0.09 | 0.00 | 0.13 | -0.09 | 0.00 | -0.15 | 0.05 |
| LMNB1 | S278 | 0.16 | -0.02 | 0.14 | -0.28 | 0.15 | -0.23 | 0.222 |
| LMNB1 | S479 | 0.25 | 0.12 | 0.05 | -0.10 | -0.06 | -0.32 | 0.04 |
| LMNB1 | S288 | 0.10 | 0.05 | 0.33 | -0.31 | -0.04 | -0.23 | 0.04 |
| LMNB1 | S200 | 0.19 | 0.05 | 0.27 | -0.31 | 0.04 | -0.35 | 0.054 |
| LMNB1 | S278 | 0.02 | 0.25 | 0.35 | -0.36 | 0.01 | -0.44 | 0.05 |
| LMNB1 | S284 | 0.32 | 0.30 | 0.20 | -0.42 | -0.04 | -0.62 | 0.022 |
| LMNB2 | T518 | 0.01 | -0.06 | -0.26 | 0.07 | -0.17 | 0.33 | 0.351 |

| | | | | | | | | |
|---|---|---|---|---|---|---|---|---|
| LMNB2 | S134 | 0.00 | -0.13 | -0.08 | 0.13 | 0.14 | -0.08 | 0.187 |
| LMNB2 | S316 | 0.02 | -0.01 | -0.22 | 0.21 | -0.22 | 0.16 | 0.488 |
| LMNB2 | S559 | -0.04 | -0.02 | -0.10 | 0.04 | -0.01 | 0.12 | 0.088 |
| LMNB2 | S168 | -0.36 | 0.34 | -0.12 | -0.04 | -0.16 | 0.23 | 0.829 |
| LMNB2 | S224 | -0.15 | -0.09 | 0.18 | 0.00 | -0.13 | 0.15 | 0.852 |
| LMNB2 | S424 | 0.10 | -0.11 | 0.01 | -0.04 | -0.04 | 0.07 | 0.967 |
| LMNB2 | T34 | 0.10 | 0.01 | 0.06 | -0.03 | -0.01 | -0.15 | 0.077 |
| LMNB2 | S405; S | 0.13 | 0.05 | 0.01 | -0.10 | -0.03 | -0.07 | 0.037 |
| LMNB2 | S424; S426 | 0.12 | 0.06 | 0.14 | -0.15 | 0.00 | -0.21 | 0.028 |
| LMNB2 | S407 | 0.10 | 0.09 | 0.19 | -0.14 | -0.07 | -0.21 | 0.007 |
| LMNB2 | S405 | 0.21 | 0.15 | 0.06 | -0.15 | -0.13 | -0.20 | 0.003 |
| LMO7 | S1026 | -0.75 | -0.19 | 0.02 | -0.27 | 0.24 | 0.59 | 0.225 |
| LMO7 | S873 | 0.00 | 0.02 | -0.20 | 0.01 | 0.12 | 0.04 | 0.209 |
| LMO7 | S867 | -0.07 | -0.03 | -0.07 | 0.03 | 0.18 | -0.05 | 0.166 |
| LMO7 | S246 | 0.06 | -0.10 | -0.14 | 0.04 | -0.05 | 0.16 | 0.282 |
| LMO7 | S1593 | -0.03 | -0.09 | -0.03 | -0.22 | -0.04 | 0.34 | 0.654 |
| LMO7 | S1423 | -0.02 | -0.02 | 0.04 | -0.04 | 0.00 | 0.03 | 0.998 |
| LMO7 | S704 | 0.03 | 0.02 | 0.01 | 0.08 | -0.14 | 0.00 | 0.56 |
| LMO7 | S805 | -0.10 | 0.14 | 0.04 | 0.09 | -0.03 | -0.15 | 0.585 |
| LMO7 | S805 | 0.16 | 0.02 | -0.01 | -0.16 | -0.01 | -0.02 | 0.185 |
| LMO7 | S276 | 0.04 | 0.09 | 0.05 | -0.21 | -0.10 | 0.11 | 0.236 |
| LMO7 | S988 | 0.10 | 0.08 | 0.21 | -0.22 | -0.36 | 0.12 | 0.129 |
| LMO7 | T1048 | 0.10 | 0.25 | 0.51 | -0.41 | -0.12 | -0.63 | 0.024 |
| LMTK2 | S601 | -0.10 | -0.18 | 0.06 | 0.07 | -0.08 | 0.18 | 0.277 |
| LMTK2 | S600; S601 | -0.02 | -0.15 | 0.02 | 0.02 | 0.06 | 0.06 | 0.148 |
| LMTK2 | S601 | -0.08 | -0.11 | 0.06 | -0.01 | -0.07 | 0.19 | 0.47 |
| LNX1 | S441 | 0.46 | -0.35 | 0.32 | -0.21 | -0.39 | -0.04 | 0.255 |
| LPAR2 | T347 | -0.27 | 0.08 | -0.08 | 0.07 | -0.20 | 0.32 | 0.438 |
| LPAR2 | T347 | -0.06 | -0.15 | 0.16 | -0.05 | -0.12 | 0.19 | 0.864 |
| LRATD2 | T289 | 0.03 | 0.22 | 0.47 | -0.35 | -0.13 | -0.47 | 0.027 |
| LRBA | S/T | -0.04 | -0.24 | -0.11 | -0.28 | 0.18 | 0.37 | 0.347 |
| LRBA | S/T | -0.04 | -0.24 | -0.11 | -0.28 | 0.18 | 0.37 | 0.347 |
| LRBA | S10 | -0.21 | 0.04 | -0.17 | 0.06 | -0.07 | 0.29 | 0.179 |
| LRBA | S1118 | -0.04 | 0.04 | 0.11 | -0.15 | 0.11 | -0.08 | 0.449 |
| LRCH1 | T568 | -0.19 | -1.34 | -0.65 | 0.07 | -0.05 | 1.05 | 0.087 |
| LRCH1 | S532; S536 | -0.12 | -0.04 | -0.04 | 0.12 | 0.11 | -0.05 | 0.107 |
| LRCH1 | S536 | 0.03 | -0.23 | 0.01 | 0.08 | 0.61 | -0.91 | 0.992 |
| LRCH1 | S370 | 0.10 | -0.08 | 0.12 | 0.11 | -0.42 | 0.09 | 0.545 |
| LRCH1 | S370 | 0.02 | 0.13 | 0.06 | -0.11 | 0.00 | -0.11 | 0.046 |
| LRCH1 | S370 | 0.04 | 0.11 | 0.10 | -0.08 | 0.02 | -0.21 | 0.068 |
| LRCH3 | S324 | -0.29 | -0.35 | 0.04 | 0.19 | 0.17 | 0.14 | 0.038 |
| LRCH3 | S419 | -0.07 | 0.23 | 0.08 | 0.31 | -0.30 | -0.39 | 0.433 |
| LRCH3 | S324 | 0.05 | 0.20 | 0.11 | -0.25 | 0.16 | -0.36 | 0.169 |
| LRCH4 | S346 | -0.13 | -0.28 | 0.03 | 0.33 | -0.13 | 0.11 | 0.23 |
| LRCH4 | S521 | 0.21 | 0.07 | -0.11 | 0.00 | -0.27 | 0.05 | 0.397 |
| LRIF1 | S386 | 0.02 | -0.11 | -0.10 | -0.08 | 0.23 | 0.01 | 0.304 |
| LRIF1 | T732 | 0.28 | -0.17 | -0.21 | -0.28 | -0.15 | 0.37 | 0.96 |
| LRIF1 | S17 | 0.28 | 0.11 | -0.35 | 0.07 | -0.52 | 0.24 | 0.784 |

| | | | | | | | | |
|---|---|---|---|---|---|---|---|---|
| LRIF1 | S/T | 0.08 | 0.22 | -0.18 | -0.34 | 0.20 | -0.07 | 0.594 |
| LRPPRC | T1136 | 0.01 | 0.12 | -0.07 | 0.06 | -0.11 | -0.03 | 0.563 |
| LRRC31 | S10 | 0.44 | 0.04 | 0.35 | -0.24 | -0.29 | -0.56 | 0.015 |
| LRRC41 | S357 | 0.20 | -0.35 | -0.16 | 0.08 | -0.17 | 0.29 | 0.467 |
| LRRC45 | S337 | 0.01 | -0.80 | -0.03 | -0.11 | 0.31 | 0.35 | 0.205 |
| LRRC47 | S520 | -0.15 | 0.11 | 0.00 | -0.03 | -0.05 | 0.10 | 0.819 |
| LRRC47 | S520 | 0.00 | 0.28 | -0.11 | 0.22 | -0.29 | -0.20 | 0.483 |
| LRRC47 | S/T/Y | 0.20 | -0.53 | 0.48 | -0.49 | -0.11 | 0.17 | 0.614 |
| LRRC47 | S520 | 0.09 | 0.13 | 0.06 | -0.12 | 0.01 | -0.20 | 0.035 |
| LRRC47 | T/S/Y | 0.37 | 0.20 | -0.24 | -0.54 | 0.18 | -0.16 | 0.363 |
| LRRC47 | S520 | 0.36 | 0.18 | 0.10 | 0.12 | -0.16 | -0.90 | 0.168 |
| LRRFIP1 | S16 | 0.09 | 0.08 | 0.01 | 0.08 | -0.02 | -0.27 | 0.286 |
| LRRFIP1 | S521 | 0.04 | 0.19 | 0.25 | -0.15 | 0.05 | -0.50 | 0.107 |
| LRRFIP1 | S714 | 0.39 | 0.18 | 0.07 | -0.32 | 0.10 | -0.65 | 0.102 |
| LRRFIP1 | S521 | 0.12 | 0.26 | 0.34 | -0.32 | -0.15 | -0.43 | 0.007 |
| LRRFIP2 | S328 | 0.07 | 0.05 | -0.14 | 0.02 | -0.10 | 0.08 | 0.909 |
| LRRFIP2 | S373 | 0.06 | -0.08 | 0.05 | 0.00 | -0.17 | 0.13 | 0.817 |
| LRRFIP2 | S328 | 0.00 | 0.06 | -0.01 | -0.10 | 0.10 | -0.06 | 0.656 |
| LRRFIP2 | S/T/Y | 0.35 | 0.14 | 0.13 | -0.11 | -0.02 | -0.71 | 0.102 |
| LRRFIP2 | S324 | 0.20 | 0.05 | 0.44 | -0.37 | 0.00 | -0.56 | 0.053 |
| LRRFIP2 | S320; S323 | 0.13 | 0.33 | 0.47 | -0.30 | -0.18 | -0.82 | 0.027 |
| LRRK1 | T/Y | -0.20 | -0.42 | -0.57 | 0.42 | -0.06 | 0.50 | 0.03 |
| LRRK1 | T964; T967 | -0.13 | -0.31 | -0.75 | 0.46 | -0.12 | 0.47 | 0.067 |
| LRWD1 | S243 | -0.14 | -0.19 | -0.03 | 0.19 | -0.27 | 0.34 | 0.348 |
| LRWD1 | S212 | -0.06 | -0.08 | -0.09 | 0.14 | -0.04 | 0.11 | 0.066 |
| LRWD1 | S212 | 0.12 | -0.06 | -0.26 | -0.08 | 0.06 | 0.17 | 0.434 |
| LRWD1 | S251 | 0.01 | -0.06 | -0.09 | 0.11 | -0.10 | 0.11 | 0.299 |
| LRWD1 | S251; S259 | -0.13 | -0.16 | 0.15 | 0.05 | -0.15 | 0.19 | 0.622 |
| LRWD1 | S251 | 0.01 | -0.04 | -0.08 | 0.16 | -0.05 | -0.02 | 0.405 |
| LRWD1 | S251; S259 | -0.05 | -0.03 | 0.00 | 0.06 | -0.08 | 0.09 | 0.446 |
| LRWD1 | S243 | -0.02 | 0.09 | 0.14 | -0.06 | 0.04 | -0.23 | 0.16 |
| LSG1 | T124 | -0.26 | -0.19 | -0.22 | 0.16 | 0.06 | 0.34 | 0.009 |
| LSM11 | S21 | -0.44 | -0.08 | 0.31 | 0.16 | -0.10 | 0.03 | 0.689 |
| LSM12 | T75 | 0.17 | -0.02 | 0.15 | 0.25 | -0.32 | -0.33 | 0.314 |
| LSM14A | S183 | 0.10 | 0.07 | 0.15 | -0.16 | 0.05 | -0.25 | 0.068 |
| LSM14A | S216 | 0.19 | 0.24 | -0.10 | -0.06 | -0.07 | -0.27 | 0.127 |
| LSM14A | S183 | 0.33 | 0.37 | 0.44 | -0.50 | -0.08 | -1.17 | 0.038 |
| LSM14B | S296 | -0.46 | 0.28 | 0.17 | -0.28 | 0.36 | -0.28 | 0.848 |
| LSR | S364 | -0.55 | 0.08 | -0.18 | 0.13 | 0.44 | -0.10 | 0.193 |
| LSR | S424 | -0.05 | -0.05 | -0.26 | 0.02 | 0.01 | 0.27 | 0.117 |
| LSR | S493 | -0.14 | 0.02 | -0.03 | 0.02 | -0.05 | 0.16 | 0.288 |
| LSR | S464 | -0.08 | 0.16 | 0.13 | -0.18 | -0.21 | 0.12 | 0.292 |
| LSR | S493 | 0.17 | 0.19 | -0.01 | -0.10 | -0.09 | -0.21 | 0.029 |
| LTBP1 | S1414 | 0.05 | 0.02 | -0.06 | 0.18 | -0.55 | 0.23 | 0.86 |
| LUC7L3 | S431 | -0.37 | -0.07 | 0.29 | -0.09 | -0.19 | 0.30 | 0.825 |
| LUC7L3 | S425 | 0.18 | 0.21 | 0.48 | -0.57 | -0.12 | -0.47 | 0.015 |
| LUZP1 | S639 | -0.04 | 0.02 | -0.32 | 0.32 | 0.03 | -0.09 | 0.276 |
| LUZP1 | S574 | 0.02 | 0.11 | -0.15 | 0.01 | 0.12 | -0.13 | 0.945 |

| | | | | | | | | |
|---|---|---|---|---|---|---|---|---|
| LYAR | S244 | -0.32 | -0.09 | -0.20 | 0.40 | 0.16 | -0.07 | 0.07 |
| LYAR | S244 | -0.42 | -0.07 | -0.13 | 0.53 | -0.10 | 0.00 | 0.191 |
| LZTS2 | S311 | -0.04 | -0.07 | -0.24 | 0.07 | 0.10 | 0.14 | 0.026 |
| LZTS2 | S301; S | -0.19 | 0.19 | -0.16 | 0.10 | 0.03 | -0.01 | 0.521 |
| LZTS2 | S346 | -0.12 | -0.11 | 0.16 | 0.01 | 0.02 | 0.02 | 0.663 |
| LZTS2 | S346 | 0.12 | -0.19 | 0.08 | -0.30 | 0.11 | 0.13 | 0.882 |
| LZTS2 | S346 | 0.12 | -0.19 | 0.08 | -0.30 | 0.11 | 0.13 | 0.882 |
| LZTS2 | S296 | 0.19 | -0.09 | 0.07 | 0.29 | -0.10 | -0.47 | 0.566 |
| LZTS2 | S570 | 0.18 | 0.11 | 0.23 | -0.04 | -0.19 | -0.39 | 0.022 |
| MACF1 | S/T | -0.07 | -0.17 | -0.14 | -0.07 | -0.32 | 0.58 | 0.515 |
| MACF1 | S280 | 0.11 | -0.14 | -0.06 | 0.11 | -0.07 | 0.04 | 0.58 |
| MACF1 | T409 | 0.08 | -0.21 | 0.18 | -0.67 | 0.30 | 0.13 | 0.784 |
| MACF1 | S3927 | -0.13 | 0.17 | 0.19 | -0.06 | 0.23 | -0.53 | 0.456 |
| MACO1 | S332 | -0.20 | 0.07 | -0.26 | -0.05 | -0.11 | 0.43 | 0.341 |
| MACO1 | S360 | 0.02 | -0.09 | 0.29 | 0.58 | -0.65 | -0.53 | 0.538 |
| MACO1 | S305 | 0.10 | 0.03 | 0.21 | 0.05 | 0.19 | -0.79 | 0.395 |
| MACO1 | T/S/Y | 0.19 | 0.13 | 0.16 | -0.32 | -0.28 | 0.04 | 0.039 |
| MACO1 | S332 | 0.16 | 0.17 | 0.29 | -0.35 | -0.06 | -0.33 | 0.012 |
| MACROH2A1 | S170; S173 | -0.42 | -0.45 | -0.38 | 0.32 | 0.02 | 0.58 | 0.012 |
| MACROH2A1 | S190 | -0.09 | -0.20 | -0.64 | 0.28 | 0.25 | 0.20 | 0.032 |
| MACROH2A1 | T129 | -0.13 | -0.22 | -0.36 | 0.20 | 0.15 | 0.25 | 0.004 |
| MACROH2A1 | T178 | -0.12 | -0.16 | -0.22 | 0.16 | 0.14 | 0.15 | 5E-04 |
| MACROH2A1 | T129 | -0.08 | -0.16 | -0.22 | 0.10 | 0.15 | 0.15 | 0.003 |
| MACROH2A1 | S313 | 0.00 | -0.09 | 0.21 | 0.00 | -0.06 | -0.07 | 0.402 |
| MACROH2A2 | S129 | -0.08 | -0.04 | -0.23 | 0.15 | 0.05 | 0.12 | 0.026 |
| MACROH2A2 | S/T | 0.35 | -0.06 | 0.09 | -0.56 | -0.20 | 0.20 | 0.282 |
| MACROH2A2 | S173 | 0.45 | 0.15 | -0.09 | -0.29 | -0.36 | -0.01 | 0.109 |
| MAD1L1 | S428 | -0.02 | -0.07 | -0.20 | -0.01 | -0.07 | 0.31 | 0.251 |
| MAD1L1 | S428 | -0.11 | 0.03 | -0.20 | -0.15 | -0.03 | 0.37 | 0.423 |
| MAD2L1BP | S102 | -0.05 | 0.01 | -0.14 | 0.08 | 0.08 | 0.02 | 0.079 |
| MADD | S1059 | -0.19 | -0.27 | -0.50 | 0.40 | 0.15 | 0.21 | 0.009 |
| MADD | S813 | 0.16 | -0.25 | -0.49 | -0.13 | -0.05 | 0.54 | 0.327 |
| MADD | S1239 | 0.00 | -0.14 | -0.21 | 0.07 | 0.04 | 0.21 | 0.054 |
| MADD | S1059 | 0.05 | -0.02 | -0.09 | 0.12 | 0.00 | -0.06 | 0.574 |
| MADD | S930 | 0.07 | -0.11 | 0.02 | 0.04 | -0.20 | 0.14 | 0.993 |
| MAFA | S336 | -0.14 | -0.55 | 0.44 | 0.41 | -0.48 | 0.02 | 0.87 |
| MAFB | S313 | -0.11 | 0.06 | -0.05 | 0.04 | -0.12 | 0.15 | 0.565 |
| MAFB | S315; S317; S318 | 0.01 | -0.06 | 0.09 | -0.08 | -0.16 | 0.17 | 0.749 |
| MAFB | S317; S318 | 0.21 | 0.14 | -0.06 | -0.14 | -0.06 | -0.13 | 0.073 |
| MAFG | S124 | -0.14 | -0.04 | 0.09 | -0.01 | 0.15 | -0.07 | 0.609 |
| MAFK | S25 | 0.02 | -0.05 | -0.12 | 0.05 | 0.05 | 0.04 | 0.076 |
| MAFK | S25 | -0.03 | 0.04 | 0.05 | -0.07 | -0.04 | 0.04 | 0.361 |
| MAGED2 | S162 | -1.21 | -1.53 | -1.63 | 0.50 | 0.57 | 1.00 | 4E-04 |
| MAGED2 | S162 | -1.04 | -0.67 | -1.07 | 0.78 | 0.57 | 0.27 | 0.002 |
| MAK16 | S232 | -0.04 | -0.17 | -0.18 | 0.08 | 0.04 | 0.23 | 0.025 |
| MAK16 | S229; S232 | 0.10 | -0.06 | -0.33 | 0.05 | 0.05 | 0.14 | 0.237 |
| MAK16 | S197; S/T | 0.00 | -0.15 | -0.12 | 0.14 | 0.02 | 0.09 | 0.046 |
| MAK16 | S | 0.05 | -0.12 | -0.11 | 0.09 | -0.04 | 0.10 | 0.179 |

| | | | | | | | | |
|---|---|---|---|---|---|---|---|---|
| **MAK16** | **S229; S232** | 0.12 | -0.07 | -0.06 | 0.00 | -0.11 | 0.10 | 0.977 |
| **MALT1** | **S42** | -0.14 | 0.01 | -0.15 | -0.03 | 0.33 | -0.08 | 0.295 |
| **MAP1S** | **S759** | 0.05 | -0.28 | -0.12 | -0.06 | 0.20 | 0.16 | 0.158 |
| **MAP1S** | **S759** | 0.16 | -0.19 | -0.19 | -0.15 | 0.27 | 0.04 | 0.485 |
| **MAP1S** | **S759** | -0.67 | 0.08 | 0.35 | -0.24 | 0.15 | 0.12 | 0.796 |
| **MAP1S** | **S475** | 0.02 | 0.14 | -0.13 | 0.12 | -0.27 | 0.09 | 0.86 |
| **MAP2K2** | **T394** | 0.04 | 0.02 | 0.00 | -0.43 | 0.13 | 0.16 | 0.749 |
| **MAP3K1** | **S1018** | -0.01 | -0.32 | -0.01 | -0.06 | 0.09 | 0.25 | 0.201 |
| **MAP3K1** | **S292** | 0.00 | 0.06 | -0.16 | -0.09 | 0.01 | 0.16 | 0.561 |
| **MAP3K1** | **S21** | 0.20 | 0.27 | 0.04 | -0.30 | -0.18 | -0.12 | 0.014 |
| **MAP3K1** | **S252** | 0.06 | 0.34 | 0.14 | 0.01 | -0.26 | -0.43 | 0.058 |
| **MAP3K11** | **S524** | -0.16 | 0.10 | -0.14 | 0.05 | 0.03 | 0.10 | 0.211 |
| **MAP3K11** | **S740** | -0.05 | 0.01 | 0.06 | 0.10 | -0.10 | -0.04 | 0.809 |
| **MAP3K11** | **S507** | -0.05 | 0.08 | 0.00 | 0.02 | 0.11 | -0.17 | 0.792 |
| **MAP3K2** | **S/Y** | -0.03 | -0.17 | -0.11 | 0.10 | -0.02 | 0.21 | 0.063 |
| **MAP3K20** | **T628** | -0.14 | -0.90 | 0.42 | 0.39 | -0.19 | 0.05 | 0.53 |
| **MAP3K20** | **S637** | 0.19 | 0.09 | -0.04 | 0.00 | 0.11 | -0.43 | 0.35 |
| **MAP3K20** | **S691** | 0.10 | 0.01 | 0.21 | -0.02 | -0.18 | -0.15 | 0.04 |
| **MAP3K20** | **S727** | 0.61 | 0.26 | -0.18 | -0.23 | -0.63 | -0.15 | 0.105 |
| **MAP3K3** | **S176** | -0.17 | -0.11 | -0.18 | 0.16 | -0.30 | 0.46 | 0.309 |
| **MAP4** | **S696** | 0.01 | -0.09 | -0.11 | 0.19 | -0.20 | 0.16 | 0.431 |
| **MAP4** | **S280** | 0.02 | 0.24 | -0.42 | 0.07 | 0.02 | 0.00 | 0.7 |
| **MAP4** | **S696** | 0.07 | -0.13 | -0.07 | 0.23 | -0.15 | 0.02 | 0.589 |
| **MAP4** | **S825** | 0.26 | 0.03 | -0.41 | 0.05 | -0.22 | 0.19 | 0.848 |
| **MAP4** | **T521** | 0.24 | 0.02 | -0.14 | 0.08 | -0.19 | -0.06 | 0.495 |
| **MAP4** | **S787** | 0.27 | 0.12 | -0.13 | -0.05 | -0.14 | -0.12 | 0.19 |
| **MAP4** | **S941** | 0.18 | 0.15 | 0.05 | -0.23 | 0.17 | -0.42 | 0.186 |
| **MAP4** | **S636** | 0.23 | 0.23 | 0.02 | -0.09 | 0.01 | -0.51 | 0.109 |
| **MAP4** | **S1073** | 0.64 | -0.12 | 0.26 | -0.22 | -0.10 | -0.91 | 0.114 |
| **MAP4K5** | **S335** | -0.42 | 0.00 | -0.14 | -0.22 | 0.52 | 0.06 | 0.285 |
| **MAP7** | **S365** | -0.27 | -0.10 | 0.03 | 0.09 | -0.06 | 0.27 | 0.173 |
| **MAP7** | **S315** | -0.11 | -0.14 | 0.12 | 0.12 | -0.27 | 0.22 | 0.714 |
| **MAP7** | **T277** | -0.10 | -0.16 | 0.21 | 0.06 | 0.01 | -0.06 | 0.859 |
| **MAP7D1** | **S366** | -0.18 | -0.25 | -0.23 | 0.09 | 0.29 | 0.19 | 0.003 |
| **MAP7D1** | **S116; S125; S/T** | -0.14 | -0.18 | -0.19 | 0.04 | -0.17 | 0.50 | 0.216 |
| **MAP7D1** | **S410** | -0.24 | 0.06 | -0.12 | 0.22 | -0.08 | 0.11 | 0.203 |
| **MAP7D1** | **S/T** | -0.05 | 0.05 | -0.26 | 0.16 | 0.15 | -0.09 | 0.265 |
| **MAP7D1** | **S116; S125** | -0.11 | -0.04 | -0.06 | 0.10 | -0.09 | 0.17 | 0.173 |
| **MAP7D1** | **S/T** | -0.15 | -0.09 | 0.07 | 0.21 | -0.12 | 0.04 | 0.433 |
| **MAP7D1** | **S410** | 0.00 | 0.11 | -0.24 | 0.02 | 0.00 | 0.08 | 0.513 |
| **MAP7D1** | **S112; S116** | -0.17 | -0.07 | 0.14 | 0.17 | 0.01 | -0.11 | 0.654 |
| **MAP7D1** | **S116; S/T** | -0.15 | 0.10 | 0.10 | -0.20 | -0.01 | 0.13 | 0.745 |
| **MAP7D1** | **S125** | -0.06 | 0.06 | 0.12 | 0.02 | -0.04 | -0.11 | 0.261 |
| **MAP7D1** | **S399** | 0.08 | 0.20 | -0.21 | 0.27 | 0.03 | -0.49 | 0.748 |
| **MAP7D1** | **S446** | 0.06 | 0.00 | 0.08 | -0.28 | 0.18 | -0.09 | 0.451 |
| **MAP7D1** | **S742** | 0.16 | 0.02 | 0.14 | -0.23 | -0.08 | -0.05 | 0.034 |
| **MAP7D1** | **S460** | 0.24 | 0.19 | 0.17 | -0.23 | -0.14 | -0.32 | 0.002 |
| **MAP7D1** | **S544** | 0.44 | 0.10 | 0.13 | -0.32 | -0.20 | -0.31 | 0.013 |

| | | | | | | | | |
|---|---|---|---|---|---|---|---|---|
| MAPK6 | S189 | -0.23 | -0.01 | -0.26 | 0.21 | 0.27 | -0.07 | 0.082 |
| MAPKAP1 | T/S | -0.35 | -0.01 | -0.11 | -0.11 | 0.15 | 0.33 | 0.161 |
| MAPT | S713; S717 | 0.05 | 0.00 | -0.11 | 0.10 | -0.14 | 0.09 | 0.703 |
| MAPT | T498 | 0.09 | 0.16 | 0.22 | -0.28 | -0.15 | -0.10 | 0.007 |
| MAPT | S519 | 0.40 | 0.17 | 0.07 | -0.29 | -0.30 | -0.18 | 0.011 |
| MAPT | S579 | 0.57 | 0.19 | -0.13 | -0.60 | 0.01 | -0.34 | 0.127 |
| MAPT | S721 | 0.39 | 0.30 | 0.00 | -0.32 | -0.16 | -0.40 | 0.02 |
| MAPT | S516; S519 | 0.21 | 0.58 | 0.11 | -0.21 | -0.29 | -0.77 | 0.033 |
| MARCHF7 | S544 | 0.28 | -0.20 | 0.24 | -0.37 | -0.02 | -0.04 | 0.264 |
| MARCKS | S81 | -0.24 | -0.16 | -0.18 | 0.13 | 0.01 | 0.34 | 0.023 |
| MARCKS | S27 | 0.02 | 0.05 | 0.01 | 0.06 | -0.09 | -0.05 | 0.319 |
| MARCKS | S145 | 0.01 | 0.05 | 0.16 | 0.02 | -0.21 | -0.06 | 0.119 |
| MARCKS | T150 | 0.26 | 0.00 | -0.04 | 0.13 | -0.36 | -0.06 | 0.368 |
| MARCKS | S46 | -0.01 | 0.04 | 0.22 | -0.24 | 0.13 | -0.19 | 0.246 |
| MARCKS | S46 | -0.01 | 0.04 | 0.22 | -0.24 | 0.13 | -0.19 | 0.246 |
| MARCKS | T150 | -0.06 | -0.05 | 0.42 | -0.25 | -0.09 | -0.05 | 0.245 |
| MARCKSL1 | S165 | -0.32 | 0.00 | -0.16 | 0.14 | -0.17 | 0.39 | 0.209 |
| MARCKSL1 | T148 | 0.00 | -0.38 | 0.17 | 0.09 | -0.12 | 0.16 | 0.582 |
| MARCKSL1 | T85 | 0.08 | 0.11 | 0.02 | -0.09 | -0.06 | -0.07 | 0.01 |
| MARCKSL1 | S22 | 0.04 | 0.12 | 0.09 | -0.03 | -0.24 | 0.01 | 0.102 |
| MARCKSL1 | S36 | 0.09 | 0.26 | 0.21 | -0.23 | -0.10 | -0.32 | 0.007 |
| MARF1 | S716 | 0.11 | 0.05 | 0.06 | 0.14 | 0.13 | -0.66 | 0.49 |
| MARK2 | S456 | -0.15 | -0.23 | 0.02 | 0.22 | -0.10 | 0.19 | 0.148 |
| MARK2 | T/S/Y | -0.07 | 0.02 | -0.20 | 0.05 | -0.16 | 0.30 | 0.362 |
| MARK2 | S486 | -0.01 | 0.02 | 0.15 | -0.07 | 0.03 | -0.13 | 0.177 |
| MARK2 | S40 | 0.26 | 0.07 | -0.07 | -0.06 | -0.51 | 0.19 | 0.412 |
| MARK4 | T214 | 0.11 | 0.20 | -0.14 | -0.17 | 0.01 | -0.04 | 0.343 |
| MARK4 | S438 | 0.07 | -0.05 | 0.18 | -0.01 | -0.24 | 0.02 | 0.241 |
| MARVELD2 | S387 | -0.17 | -0.03 | 0.15 | 0.15 | 0.05 | -0.20 | 0.932 |
| MARVELD2 | S389 | 0.00 | 0.04 | 0.16 | -0.24 | -0.13 | 0.13 | 0.284 |
| MARVELD2 | S389 | 0.00 | 0.04 | 0.16 | -0.24 | -0.13 | 0.13 | 0.284 |
| MARVELD2 | S116 | 0.16 | 0.38 | 0.04 | -0.35 | -0.30 | -0.07 | 0.029 |
| MAST2 | S74 | 0.32 | -0.10 | -0.19 | -0.18 | 0.15 | -0.08 | 0.82 |
| MAST2 | S1381 | 0.19 | 0.51 | -0.13 | -0.32 | -0.25 | -0.19 | 0.077 |
| MAST4 | T2516 | -2.13 | -1.71 | -1.68 | 1.34 | -1.45 | 1.18 | 0.075 |
| MASTL | S598 | -0.20 | -0.21 | -0.20 | 0.31 | 0.16 | 0.05 | 0.007 |
| MASTL | S598 | -0.12 | -0.26 | -0.08 | 0.11 | 0.08 | 0.21 | 0.012 |
| MASTL | S453 | 0.16 | -0.04 | 0.33 | -0.11 | 0.03 | -0.49 | 0.146 |
| MATR3 | S208 | -0.60 | -0.50 | -0.65 | 0.40 | 0.66 | 0.13 | 0.004 |
| MATR3 | T150 | -0.05 | -0.29 | -0.31 | 0.13 | 0.13 | 0.29 | 0.016 |
| MATR3 | S596; S598 | 0.02 | -0.16 | -0.22 | 0.13 | 0.01 | 0.18 | 0.063 |
| MATR3 | S766 | -0.02 | 0.08 | -0.46 | -0.17 | 0.03 | 0.40 | 0.404 |
| MATR3 | S206 | -0.02 | -0.08 | -0.15 | 0.01 | -0.11 | 0.30 | 0.308 |
| MATR3 | S188 | -0.22 | 0.01 | 0.04 | 0.01 | -0.09 | 0.22 | 0.443 |
| MATR3 | S598 | -0.02 | -0.07 | 0.02 | 0.07 | -0.09 | 0.09 | 0.509 |
| MATR3 | S766 | 0.12 | -0.04 | -0.15 | -0.24 | 0.04 | 0.20 | 0.882 |
| MATR3 | S766 | 0.08 | 0.00 | -0.01 | -0.03 | -0.10 | 0.05 | 0.373 |
| MATR3 | S157 | -0.10 | -0.08 | 0.24 | -0.08 | -0.10 | 0.08 | 0.677 |

| | | | | | | | | |
|---|---|---|---|---|---|---|---|---|
| MATR3 | S598; S604 | 0.19 | 0.05 | -0.10 | -0.09 | -0.20 | 0.10 | 0.414 |
| MATR3 | S604 | 0.18 | 0.09 | -0.03 | -0.09 | 0.01 | -0.18 | 0.116 |
| MATR3 | S596; S598; S604 | 0.28 | 0.10 | -0.16 | 0.02 | -0.13 | -0.17 | 0.298 |
| MATR3 | S11 | 0.04 | 0.16 | 0.09 | -0.13 | 0.07 | -0.28 | 0.127 |
| MATR3 | S598; S604 | 0.20 | 0.10 | 0.01 | -0.11 | -0.15 | -0.08 | 0.025 |
| MATR3 | S598 | 0.23 | -0.01 | 0.07 | -0.32 | 0.17 | -0.22 | 0.254 |
| MATR3 | S705 | 0.20 | 0.00 | 0.11 | -0.23 | 0.13 | -0.28 | 0.183 |
| MATR3 | S604 | 0.18 | 0.13 | 0.15 | 0.00 | -0.14 | -0.40 | 0.046 |
| MAVS | T/S/Y | -0.07 | -0.41 | -0.51 | 0.30 | 0.24 | 0.24 | 0.012 |
| MAVS | T/S/Y | -0.53 | -0.12 | 0.21 | -0.09 | 0.09 | 0.30 | 0.364 |
| MAVS | T/S/Y | -0.53 | -0.12 | 0.21 | -0.09 | 0.09 | 0.30 | 0.364 |
| MAVS | S/T | -0.02 | -0.13 | -0.13 | 0.14 | -0.03 | 0.14 | 0.063 |
| MAVS | S165 | 0.04 | 0.27 | -0.97 | 0.51 | -1.13 | 0.45 | 0.816 |
| MAVS | S165 | -0.17 | -0.16 | 0.17 | -0.08 | 0.14 | 0.05 | 0.516 |
| MAVS | S/T | 0.18 | -0.11 | -0.15 | 0.06 | -0.51 | 0.37 | 0.998 |
| MAX | Y/S/T | -0.02 | -0.03 | -0.15 | 0.04 | 0.07 | 0.07 | 0.044 |
| MAX | S/T/Y | -0.03 | -0.03 | -0.13 | 0.07 | -0.10 | 0.19 | 0.273 |
| MAX | S/T | -0.07 | 0.05 | -0.02 | 0.03 | -0.10 | 0.10 | 0.763 |
| MAX | S144 | -0.13 | 0.07 | 0.18 | -0.10 | 0.01 | -0.06 | 0.397 |
| MAX | S140; S142; S144 | 0.12 | 0.10 | 0.05 | -0.05 | -0.03 | -0.21 | 0.039 |
| MAZ | S302 | -0.57 | -0.41 | -0.51 | 0.38 | 0.23 | 0.48 | 6E-04 |
| MAZ | S302 | -0.33 | -0.23 | -0.53 | 0.41 | 0.10 | 0.32 | 0.007 |
| MBD1 | S311 | -0.50 | -0.16 | -0.37 | 0.24 | 0.00 | 0.52 | 0.03 |
| MBD1 | T505 | -0.40 | -0.48 | -0.13 | 0.27 | 0.07 | 0.44 | 0.017 |
| MBD1 | S311 | -0.26 | -0.17 | 0.02 | 0.11 | -0.07 | 0.30 | 0.141 |
| MBD1 | S297 | 0.07 | -0.02 | -0.44 | 0.25 | -0.09 | 0.15 | 0.286 |
| MBD1 | S391; S399 | -0.06 | -0.07 | -0.17 | 0.11 | -0.07 | 0.23 | 0.109 |
| MBD1 | S37 | -0.01 | -0.39 | 0.24 | 0.05 | -0.04 | 0.09 | 0.673 |
| MBD1 | S297 | 0.02 | -0.15 | 0.07 | -0.08 | 0.10 | 0.03 | 0.673 |
| MBD1 | S297 | 0.03 | 0.16 | 0.03 | -0.14 | -0.07 | -0.03 | 0.04 |
| MBD1 | S/Y | 0.19 | 0.09 | -0.05 | -0.17 | -0.16 | 0.07 | 0.204 |
| MBD1 | S297 | 0.12 | 0.12 | 0.13 | -0.16 | -0.06 | -0.19 | 0.003 |
| MBD2 | S407 | 0.00 | -0.20 | -0.08 | 0.21 | -0.02 | 0.06 | 0.112 |
| MBD2 | S181 | 0.04 | 0.00 | -0.11 | -0.05 | 0.16 | -0.06 | 0.666 |
| MBD2 | S44 | -0.07 | 0.00 | 0.19 | -0.18 | 0.05 | -0.02 | 0.441 |
| MBD2 | S407 | 0.33 | -0.05 | 0.05 | -0.24 | 0.05 | -0.22 | 0.168 |
| MBD2 | S44 | 0.17 | 0.17 | 0.03 | -0.10 | -0.02 | -0.31 | 0.051 |
| MBD2 | S44 | 0.33 | 0.07 | 0.00 | -0.10 | -0.07 | -0.31 | 0.076 |
| MBD3 | S37 | -0.03 | -0.14 | -0.18 | -0.52 | 0.41 | 0.26 | 0.6 |
| MBD3 | S85 | 0.33 | 0.20 | 0.27 | -0.49 | -0.22 | -0.31 | 0.002 |
| MBIP | S91 | 0.13 | 0.00 | -0.09 | 0.19 | -0.15 | -0.11 | 0.785 |
| MBOAT2 | T474 | -0.03 | -0.12 | -0.34 | 0.13 | 0.05 | 0.24 | 0.05 |
| MBOAT2 | T474 | -0.12 | -0.26 | 0.12 | 0.19 | 0.09 | -0.06 | 0.289 |
| MBOAT7 | S285 | -0.04 | -0.19 | -0.21 | 0.02 | -0.05 | 0.38 | 0.134 |
| MBOAT7 | S285 | 0.08 | 0.03 | 0.06 | -0.05 | -0.10 | -0.04 | 0.008 |
| MBOAT7 | T/S | 0.09 | 0.09 | 0.36 | -0.18 | -0.49 | -0.01 | 0.07 |
| MBOAT7 | S291 | 0.32 | 0.29 | 0.19 | -0.26 | -0.23 | -0.52 | 0.004 |
| MCC | S828 | -0.07 | 0.27 | -0.34 | -0.44 | -0.16 | 0.51 | 0.965 |

| | | | | | | | | |
|---|---|---|---|---|---|---|---|---|
| **MCM10** | **S/Y/T** | -0.26 | -0.26 | 0.23 | -0.18 | 0.13 | 0.24 | 0.491 |
| **MCM10** | **S644** | 0.19 | -0.21 | 0.03 | 0.08 | -0.21 | 0.07 | 0.899 |
| **MCM10** | **S54** | 0.06 | 0.00 | 0.04 | -0.06 | 0.14 | -0.19 | 0.516 |
| **MCM2** | **S170** | -0.24 | -0.21 | -0.37 | 0.21 | 0.12 | 0.35 | 0.004 |
| **MCM2** | **T158** | -0.22 | -0.18 | -0.24 | 0.21 | 0.02 | 0.32 | 0.011 |
| **MCM2** | **S229** | -0.07 | -0.36 | 0.15 | 0.05 | -0.10 | 0.25 | 0.417 |
| **MCM2** | **S229** | -0.07 | -0.36 | 0.15 | 0.05 | -0.10 | 0.25 | 0.417 |
| **MCM2** | **S53** | 0.07 | -0.11 | 0.03 | 0.05 | -0.20 | 0.13 | 0.993 |
| **MCM2** | **S27; T/S** | 0.12 | -0.08 | 0.15 | 0.09 | -0.14 | -0.17 | 0.286 |
| **MCM2** | **T/S** | 0.07 | -0.02 | 0.19 | -0.08 | -0.20 | 0.00 | 0.107 |
| **MCM2** | **S27** | 0.16 | 0.00 | 0.11 | -0.02 | -0.28 | -0.01 | 0.124 |
| **MCM2** | **S139** | 0.19 | 0.00 | 0.08 | -0.01 | -0.25 | -0.05 | 0.111 |
| **MCM2** | **S41** | 0.14 | 0.07 | 0.13 | -0.21 | -0.04 | -0.12 | 0.012 |
| **MCM2** | **S/T** | 0.24 | 0.25 | 0.36 | -0.37 | -0.11 | -0.64 | 0.014 |
| **MCM3** | **S711; T713; T722** | -0.03 | -0.16 | 0.04 | -0.01 | 0.19 | -0.04 | 0.345 |
| **MCM3** | **S275** | 0.03 | 0.02 | -0.13 | 0.16 | 0.21 | -0.36 | 0.89 |
| **MCM3** | **S711; T713; T722** | 0.02 | -0.09 | 0.16 | 0.04 | 0.20 | -0.41 | 0.697 |
| **MCM3** | **T722; S/T/Y** | 0.14 | 0.04 | -0.02 | -0.02 | -0.24 | 0.07 | 0.303 |
| **MCM3** | **S/T/Y** | 0.38 | 0.20 | -0.15 | -0.40 | -0.10 | -0.05 | 0.165 |
| **MCM3** | **S711** | 0.21 | 0.15 | 0.16 | -0.22 | -0.16 | -0.21 | 1E-04 |
| **MCM3** | **S711; T722** | 0.26 | 0.14 | 0.11 | -0.05 | -0.19 | -0.36 | 0.02 |
| **MCM3** | **S/T/Y** | 0.29 | 0.15 | 0.14 | -0.17 | -0.22 | -0.29 | 0.002 |
| **MCM3** | **S711** | 0.31 | 0.22 | 0.19 | -0.22 | -0.21 | -0.45 | 0.003 |
| **MCM3AP** | **S300** | -0.10 | -0.21 | -0.06 | 0.61 | -0.12 | -0.34 | 0.583 |
| **MCM3AP** | **S430** | 0.32 | 0.10 | 0.02 | -0.19 | 0.00 | -0.34 | 0.07 |
| **MCM3AP** | **S538** | 0.36 | 0.19 | -0.08 | -0.27 | -0.15 | -0.15 | 0.058 |
| **MCM4** | **S120** | -0.26 | -0.40 | -0.12 | 0.06 | 0.32 | 0.25 | 0.013 |
| **MCM4** | **S131; S142; S145** | 0.05 | -0.25 | -0.02 | 0.19 | -0.08 | 0.06 | 0.344 |
| **MCM4** | **S131** | -0.06 | 0.07 | 0.04 | 0.31 | -0.23 | -0.22 | 0.752 |
| **MCM4** | **S326** | 0.13 | 0.04 | 0.12 | -0.13 | -0.13 | -0.06 | 0.005 |
| **MCM5** | **S315** | -0.20 | 0.12 | -0.15 | 0.36 | -0.06 | -0.14 | 0.518 |
| **MCM5** | **S315** | -0.04 | -0.23 | 0.08 | -0.27 | 0.07 | 0.31 | 0.622 |
| **MCM6** | **S762** | -0.45 | -0.61 | -0.76 | 0.45 | 0.39 | 0.43 | 4E-04 |
| **MCM7** | **S121** | 0.31 | -0.06 | -0.30 | 0.08 | -0.50 | 0.30 | 0.944 |
| **MCM7** | **S678** | -0.10 | 0.24 | 0.21 | -0.78 | -0.38 | 0.47 | 0.417 |
| **MCM8** | **S630** | -0.26 | -0.46 | -0.13 | 0.23 | 0.30 | 0.16 | 0.008 |
| **MCM9** | **S883** | 0.06 | -0.19 | 0.00 | -0.13 | 0.26 | -0.05 | 0.641 |
| **MCM9** | **S1073** | -0.05 | 0.02 | -0.02 | -0.07 | 0.05 | 0.06 | 0.544 |
| **MCM9** | **S1088** | -0.36 | -0.21 | 0.70 | -0.59 | 0.15 | -0.05 | 0.623 |
| **MCMBP** | **S154** | 0.06 | 0.05 | 0.02 | -0.02 | -0.01 | -0.12 | 0.055 |
| **MCOLN1** | **T34** | -0.20 | 0.07 | -0.04 | 0.37 | -0.25 | -0.04 | 0.697 |
| **MCOLN1** | **T34** | -0.07 | 0.08 | 0.05 | 0.10 | -0.05 | -0.12 | 0.634 |
| **MCOLN1** | **S547** | 0.21 | 0.14 | 0.06 | -0.03 | -0.12 | -0.31 | 0.034 |
| **MCPH1** | **S333** | 0.31 | -0.08 | 0.01 | -0.14 | -0.03 | -0.12 | 0.219 |
| **MCRIP1** | **S21** | 0.02 | 0.01 | -0.03 | -0.17 | 0.07 | 0.08 | 0.951 |
| **MCRIP2** | **S/T/Y** | -0.17 | -0.22 | 0.05 | 0.04 | 0.08 | 0.18 | 0.085 |
| **MCRIP2** | **S/T/Y** | -0.02 | -0.14 | 0.00 | 0.20 | -0.14 | 0.07 | 0.405 |
| **MCRIP2** | **S82** | -0.11 | 0.06 | -0.03 | 0.39 | -0.01 | -0.42 | 0.953 |

| | | | | | | | | |
|---|---|---|---|---|---|---|---|---|
| **MCRS1** | **S282** | -0.04 | -0.03 | -0.06 | -0.01 | -0.06 | 0.18 | 0.324 |
| **MCRS1** | **S282** | 0.05 | 0.01 | -0.04 | -0.05 | -0.03 | 0.07 | 0.827 |
| **MCRS1** | **S22** | 0.05 | 0.03 | 0.03 | -0.09 | 0.02 | -0.04 | 0.072 |
| **MCRS1** | **S85** | 0.26 | 0.00 | -0.18 | 0.32 | -0.29 | -0.22 | 0.708 |
| **MDC1** | **S/T** | -1.09 | -1.13 | -1.34 | 0.70 | 0.09 | 0.99 | 0.003 |
| **MDC1** | **S485; S495; S498** | -0.47 | -0.87 | -1.02 | 0.55 | 0.19 | 0.71 | 0.005 |
| **MDC1** | **T1800** | -0.51 | -0.38 | -0.63 | -0.05 | 0.55 | 0.54 | 0.016 |
| **MDC1** | **T1485** | -0.50 | -0.67 | -0.47 | -0.39 | 0.08 | 1.08 | 0.14 |
| **MDC1** | **S1086** | -0.51 | -0.24 | -0.65 | 0.07 | 0.69 | 0.19 | 0.025 |
| **MDC1** | **S1086** | -0.51 | -0.24 | -0.65 | 0.07 | 0.69 | 0.19 | 0.025 |
| **MDC1** | **S/T** | -0.99 | -0.64 | 0.12 | -0.02 | -0.32 | 0.99 | 0.233 |
| **MDC1** | **T1526** | -0.02 | -0.35 | -0.74 | 0.13 | -0.04 | 0.64 | 0.103 |
| **MDC1** | **T1526** | -0.02 | -0.35 | -0.74 | 0.13 | -0.04 | 0.64 | 0.103 |
| **MDC1** | **T1198** | -0.35 | -0.34 | -0.33 | 0.10 | 0.48 | 0.22 | 0.006 |
| **MDC1** | **T1157** | -0.21 | -0.28 | -0.52 | 0.15 | 0.39 | 0.27 | 0.006 |
| **MDC1** | **S1033** | -0.26 | -0.31 | -0.43 | 0.24 | 0.08 | 0.47 | 0.008 |
| **MDC1** | **S/T/Y** | 0.09 | -0.61 | -0.44 | 0.19 | -0.14 | 0.58 | 0.147 |
| **MDC1** | **T966** | -0.18 | -0.22 | -0.61 | 0.24 | -0.41 | 0.74 | 0.216 |
| **MDC1** | **T1198** | -0.35 | -0.31 | -0.21 | 0.07 | 0.41 | 0.23 | 0.008 |
| **MDC1** | **S1033** | -0.19 | -0.27 | -0.41 | 0.20 | 0.01 | 0.47 | 0.025 |
| **MDC1** | **S376** | -0.27 | -0.23 | -0.35 | 0.16 | 0.09 | 0.43 | 0.009 |
| **MDC1** | **S793** | -0.29 | -0.13 | -0.34 | 0.25 | 0.07 | 0.32 | 0.009 |
| **MDC1** | **S/T** | -0.18 | -0.21 | -0.35 | 0.20 | -0.01 | 0.41 | 0.025 |
| **MDC1** | **S793** | -0.08 | -0.22 | -0.35 | 0.15 | -0.01 | 0.39 | 0.05 |
| **MDC1** | **S495; S505; S** | -0.29 | -0.12 | -0.23 | 0.06 | 0.13 | 0.34 | 0.016 |
| **MDC1** | **S376** | -0.18 | -0.19 | -0.25 | 0.12 | 0.02 | 0.37 | 0.023 |
| **MDC1** | **T1526; T1548** | -0.23 | -0.11 | -0.27 | 0.13 | 0.20 | 0.19 | 0.002 |
| **MDC1** | **S495; S498** | -0.08 | -0.16 | -0.39 | 0.14 | 0.41 | -0.05 | 0.083 |
| **MDC1** | **T1157** | -0.34 | 0.04 | -0.31 | 0.33 | -0.18 | 0.30 | 0.159 |
| **MDC1** | **T378** | -0.15 | -0.13 | -0.31 | 0.26 | -0.22 | 0.41 | 0.152 |
| **MDC1** | **T/S** | -0.07 | -0.19 | -0.28 | 0.15 | 0.18 | 0.15 | 0.005 |
| **MDC1** | **T1671** | -0.24 | 0.01 | -0.32 | -0.01 | 0.39 | 0.06 | 0.104 |
| **MDC1** | **T1157** | -0.09 | -0.65 | 0.15 | -0.09 | 0.32 | 0.17 | 0.286 |
| **MDC1** | **S108** | -0.14 | -0.08 | -0.29 | 0.10 | 0.03 | 0.31 | 0.039 |
| **MDC1** | **T1302** | -0.12 | -0.39 | 0.07 | 0.11 | 0.18 | 0.08 | 0.117 |
| **MDC1** | **T1403** | -0.59 | 0.03 | 0.08 | -0.22 | 0.23 | 0.29 | 0.386 |
| **MDC1** | **T/S** | -0.16 | -0.03 | -0.27 | 0.05 | -0.27 | 0.52 | 0.347 |
| **MDC1** | **S329** | -0.12 | -0.11 | -0.17 | 0.21 | 0.02 | 0.13 | 0.012 |
| **MDC1** | **S/T** | -0.15 | -0.20 | -0.07 | 0.17 | -0.25 | 0.38 | 0.273 |
| **MDC1** | **T1239** | -0.20 | -0.17 | 0.00 | 0.20 | 0.10 | 0.03 | 0.041 |
| **MDC1** | **S/T** | -0.55 | 0.11 | 0.07 | 0.23 | 0.16 | -0.16 | 0.456 |
| **MDC1** | **T1567** | -0.06 | 0.09 | -0.36 | -0.07 | 0.33 | -0.02 | 0.363 |
| **MDC1** | **S299; T301** | -0.17 | -0.10 | -0.02 | -0.11 | -0.01 | 0.35 | 0.302 |
| **MDC1** | **S329** | -0.08 | -0.08 | -0.09 | 0.13 | 0.03 | 0.09 | 0.004 |
| **MDC1** | **S513** | -0.17 | 0.01 | -0.11 | 0.23 | 0.03 | -0.03 | 0.151 |
| **MDC1** | **T1485** | -0.06 | 0.05 | -0.25 | -0.17 | 0.17 | 0.20 | 0.354 |
| **MDC1** | **T1444** | 0.11 | -0.64 | 0.22 | -0.36 | 0.08 | 0.36 | 0.71 |
| **MDC1** | **S1820** | -0.03 | -0.11 | -0.06 | 0.11 | 0.10 | -0.02 | 0.054 |

| | | | | | | | | |
|---|---|---|---|---|---|---|---|---|
| **MDC1** | **S1820** | -0.04 | -0.02 | -0.14 | 0.14 | 0.10 | -0.06 | 0.159 |
| **MDC1** | **T1485** | -0.10 | 0.15 | -0.27 | -0.23 | 0.11 | 0.25 | 0.577 |
| **MDC1** | **S793** | -0.19 | -0.07 | 0.09 | 0.02 | 0.03 | 0.10 | 0.285 |
| **MDC1** | **S/T** | -0.04 | 0.02 | -0.14 | -0.08 | -0.05 | 0.26 | 0.479 |
| **MDC1** | **T1444; T1466** | -0.23 | 0.03 | 0.06 | 0.23 | -0.15 | 0.02 | 0.601 |
| **MDC1** | **S376** | -0.36 | 0.20 | 0.05 | -0.52 | 0.18 | 0.27 | 0.96 |
| **MDC1** | **S498** | -0.06 | 0.17 | -0.13 | -0.06 | 0.18 | -0.14 | 0.984 |
| **MDC1** | **S168** | -0.20 | 0.09 | 0.06 | -0.52 | 0.03 | 0.38 | 0.95 |
| **MDC1** | **T1630** | 0.07 | -0.40 | 0.30 | -0.03 | -0.34 | 0.25 | 0.911 |
| **MDC1** | **S505** | -0.09 | 0.10 | 0.07 | 0.06 | -0.17 | 0.02 | 0.595 |
| **MDC1** | **S641** | -0.01 | -0.06 | 0.11 | 0.00 | -0.36 | 0.24 | 0.78 |
| **MDC1** | **T1526** | 0.03 | 0.13 | -0.11 | 0.10 | -0.48 | 0.23 | 0.775 |
| **MDC1** | **S793** | 0.18 | -0.04 | -0.07 | -0.40 | -0.02 | 0.26 | 0.742 |
| **MDC1** | **T404** | 0.10 | -0.04 | 0.04 | -0.15 | 0.19 | -0.18 | 0.568 |
| **MDC1** | **T1384** | -0.02 | 0.06 | 0.07 | -0.01 | 0.11 | -0.24 | 0.492 |
| **MDC1** | **T301** | 0.01 | 0.00 | 0.11 | -0.06 | 0.07 | -0.15 | 0.286 |
| **MDC1** | **S780** | 0.19 | -0.24 | 0.22 | 0.01 | -0.20 | -0.06 | 0.424 |
| **MDC1** | **T1630** | 0.09 | 0.02 | 0.11 | -0.08 | -0.25 | 0.08 | 0.19 |
| **MDC1** | **T1384** | 0.09 | 0.16 | 0.15 | -0.20 | 0.11 | -0.42 | 0.122 |
| **MDC1** | **T1548** | 0.08 | 0.29 | 0.09 | -0.54 | 0.20 | -0.29 | 0.183 |
| **MDC1** | **S780** | 0.21 | 0.11 | 0.19 | -0.25 | -0.07 | -0.28 | 0.007 |
| **MDC1** | **T1425** | 0.07 | 0.53 | -0.02 | -0.23 | -0.38 | -0.17 | 0.065 |
| **MDC1** | **S** | 0.33 | 0.32 | 0.25 | -0.60 | -0.33 | -0.24 | 0.003 |
| **MDM1** | **S356** | -0.38 | -0.21 | -0.16 | 0.33 | 0.26 | 0.02 | 0.016 |
| **MDM1** | **T/S** | -0.12 | -0.20 | 0.11 | -0.26 | 0.10 | 0.29 | 0.579 |
| **MDM1** | **S263** | 0.16 | 0.23 | 0.34 | -0.39 | -0.08 | -0.43 | 0.012 |
| **MDN1** | **T4898** | 0.13 | -0.39 | -0.22 | 0.07 | 0.06 | 0.25 | 0.159 |
| **MDN1** | **S4538** | -0.09 | 0.01 | -0.02 | 0.17 | 0.08 | -0.18 | 0.661 |
| **MDN1** | **T4898** | 0.14 | 0.19 | 0.27 | 0.02 | -0.28 | -0.49 | 0.042 |
| **MECP2** | **S149** | -2.06 | -1.38 | -1.50 | 0.73 | 0.96 | 0.83 | 3E-04 |
| **MECP2** | **S423; S426** | -1.20 | -1.05 | -1.23 | 0.51 | 0.44 | 0.91 | 3E-04 |
| **MECP2** | **S423** | -1.05 | -1.24 | -1.12 | 0.65 | 0.47 | 0.75 | 6E-05 |
| **MECP2** | **S401** | -0.23 | -0.57 | -0.46 | 0.31 | 0.54 | 0.08 | 0.012 |
| **MECP2** | **S68; S80** | -0.57 | -0.39 | -0.19 | 0.36 | 0.26 | 0.28 | 0.004 |
| **MECP2** | **S80** | -0.07 | -0.10 | -0.11 | 0.07 | 0.00 | 0.19 | 0.038 |
| **MECP2** | **S229** | 0.03 | -0.01 | -0.16 | -0.01 | 0.12 | 0.02 | 0.262 |
| **MECP2** | **S360** | 0.11 | 0.01 | 0.11 | -0.12 | 0.02 | -0.14 | 0.065 |
| **MECP2** | **S360** | 0.17 | 0.10 | 0.15 | -0.17 | 0.10 | -0.45 | 0.126 |
| **MED1** | **S1049; T1051; T1057** | -0.07 | -0.54 | -0.03 | 0.07 | -0.23 | 0.56 | 0.289 |
| **MED1** | **S1453** | -0.02 | -0.02 | -0.01 | -0.06 | -0.18 | 0.26 | 0.869 |
| **MED1** | **T1159** | 0.03 | 0.07 | -0.04 | 0.14 | -0.24 | 0.01 | 0.671 |
| **MED1** | **S238** | -0.28 | 0.21 | 0.11 | -0.35 | -0.05 | 0.25 | 0.793 |
| **MED1** | **T805** | -0.10 | 0.16 | 0.07 | 0.05 | -0.24 | 0.03 | 0.448 |
| **MED1** | **S771** | -0.06 | -0.01 | 0.21 | -0.05 | 0.12 | -0.25 | 0.475 |
| **MED1** | **S1447; S1453** | 0.23 | -0.02 | -0.06 | 0.13 | -0.08 | -0.25 | 0.45 |
| **MED1** | **S1447** | 0.26 | 0.14 | -0.11 | 0.01 | -0.24 | -0.11 | 0.183 |
| **MED1** | **S/T** | 0.09 | 0.17 | 0.02 | 0.11 | -0.14 | -0.31 | 0.189 |
| **MED1** | **S1401** | 0.24 | 0.31 | 0.34 | -0.29 | -0.15 | -0.73 | 0.018 |

| | | | | | | | | |
|---|---|---|---|---|---|---|---|---|
| MED1 | S/Y/T | 0.54 | 0.43 | 0.14 | -0.47 | -0.20 | -0.97 | 0.023 |
| MED1 | S/Y/T | 0.47 | 0.42 | 0.31 | -0.48 | -0.36 | -0.88 | 0.004 |
| MED12 | S1258 | -0.12 | -0.18 | 0.03 | 0.11 | -0.13 | 0.25 | 0.26 |
| MED12 | S698 | 0.15 | -0.06 | -0.17 | 0.23 | -0.18 | -0.01 | 0.819 |
| MED12 | S635 | 0.08 | -0.02 | -0.06 | 0.28 | -0.15 | -0.17 | 0.924 |
| MED13 | S395 | -0.03 | 0.14 | 0.25 | 0.11 | -0.15 | -0.41 | 0.191 |
| MED13 | S890 | 0.13 | 0.16 | 0.05 | 0.25 | -0.22 | -0.50 | 0.284 |
| MED17 | S55 | -0.13 | -0.03 | -0.20 | -0.01 | 0.02 | 0.30 | 0.11 |
| MED24 | S/T | 0.08 | -0.19 | -0.49 | 0.23 | -0.12 | 0.34 | 0.175 |
| MED24 | S873 | -0.15 | 0.25 | 0.17 | 0.10 | -0.15 | -0.30 | 0.281 |
| MED26 | S447 | -0.09 | 0.10 | -0.21 | 0.07 | 0.24 | -0.17 | 0.495 |
| MED26 | S373 | 0.22 | 0.06 | 0.09 | 0.05 | -0.28 | -0.19 | 0.075 |
| MEF2A | S98 | -0.05 | -0.08 | -0.07 | 0.05 | -0.09 | 0.21 | 0.223 |
| MEF2D | S98; S110 | -0.29 | -0.18 | -0.07 | 0.19 | -0.06 | 0.31 | 0.06 |
| MEF2D | S98 | 0.07 | -0.02 | -0.08 | -0.08 | -0.01 | 0.11 | 0.842 |
| MEIS1 | S196 | 0.15 | 0.13 | 0.13 | -0.11 | -0.05 | -0.30 | 0.019 |
| MELK | S505 | 0.00 | 0.03 | -0.09 | 0.12 | 0.12 | -0.20 | 0.777 |
| MELK | S356 | -0.14 | 0.10 | 0.03 | -0.31 | 0.23 | 0.03 | 0.94 |
| MEN1 | S492 | 0.10 | -0.24 | -0.28 | -0.13 | 0.27 | 0.19 | 0.212 |
| MEPCE | S330 | -0.42 | -0.15 | 0.09 | 0.48 | 0.24 | -0.51 | 0.53 |
| MEPCE | S330 | -0.12 | 0.32 | -0.43 | 0.42 | -0.63 | 0.15 | 0.893 |
| MEPCE | S254 | 0.01 | 0.01 | -0.07 | 0.18 | 0.02 | -0.17 | 0.8 |
| MEPCE | S69 | -0.18 | 0.15 | 0.20 | 0.10 | -0.14 | -0.18 | 0.432 |
| MEPCE | T213; S/T | -0.03 | 0.01 | 0.19 | 0.17 | -0.07 | -0.33 | 0.441 |
| MEPCE | S60 | -0.01 | 0.22 | 0.00 | -0.01 | 0.01 | -0.26 | 0.255 |
| METTL16 | S498 | -0.08 | 0.36 | 0.28 | -0.53 | -0.03 | -0.18 | 0.096 |
| METTL16 | S498 | 0.12 | 0.37 | 0.17 | -0.54 | -0.03 | -0.27 | 0.04 |
| METTL16 | S498 | 0.27 | 0.37 | 0.07 | -0.55 | -0.02 | -0.35 | 0.038 |
| METTL22 | S132 | -0.24 | -0.41 | -0.46 | 0.20 | 0.00 | 0.60 | 0.028 |
| METTL3 | S/T | -1.28 | -1.18 | -1.07 | 1.19 | -1.18 | 0.97 | 0.119 |
| METTL3 | S43 | 0.03 | -0.02 | 0.14 | 0.26 | -0.11 | -0.39 | 0.546 |
| MEX3A | S338 | 0.14 | -0.01 | -0.01 | 0.31 | -0.18 | -0.35 | 0.607 |
| MEX3C | S537 | -0.09 | 0.10 | 0.22 | -0.24 | 0.21 | -0.29 | 0.373 |
| MEX3D | S514 | -0.30 | -0.07 | 0.22 | -0.18 | 0.30 | -0.06 | 0.755 |
| MFAP1 | S132; S133 | -0.18 | -0.25 | 0.15 | -0.10 | 0.17 | 0.15 | 0.323 |
| MFAP1 | S118 | -0.21 | -0.13 | 0.08 | -0.04 | -0.13 | 0.36 | 0.449 |
| MFAP1 | T364 | 0.05 | -0.14 | -0.05 | 0.02 | -0.07 | 0.18 | 0.395 |
| MFAP1 | S116; S118 | -0.02 | -0.09 | -0.02 | 0.06 | -0.02 | 0.08 | 0.082 |
| MFAP1 | T267 | -0.01 | -0.23 | 0.18 | 0.00 | -0.27 | 0.24 | 0.96 |
| MFAP1 | T267 | 0.08 | 0.02 | 0.00 | -0.20 | 0.14 | -0.06 | 0.486 |
| MFAP1 | S116 | 0.13 | 0.13 | -0.09 | -0.05 | 0.02 | -0.18 | 0.255 |
| MFAP1 | T267 | 0.26 | -0.09 | 0.07 | -0.24 | -0.19 | 0.13 | 0.314 |
| MFAP1 | S/Y | 0.17 | 0.17 | 0.02 | 0.07 | -0.17 | -0.32 | 0.099 |
| MFAP1 | S132; S133 | 0.24 | 0.06 | 0.07 | -0.54 | 0.00 | 0.06 | 0.223 |
| MFAP1 | S52 | 0.23 | 0.24 | 0.10 | -0.06 | -0.09 | -0.57 | 0.066 |
| MFAP1 | S52 | 0.28 | 0.21 | 0.15 | -0.07 | -0.14 | -0.60 | 0.049 |
| MFF | S157 | 0.19 | 0.28 | 0.21 | -0.24 | -0.19 | -0.39 | 0.002 |
| MFSD13A | S85 | 0.06 | -0.02 | 0.04 | -0.07 | 0.15 | -0.19 | 0.58 |

| | | | | | | | | |
|---|---|---|---|---|---|---|---|---|
| **MFSD2A** | **S/T** | -0.26 | 0.18 | -0.10 | 0.13 | -0.24 | 0.20 | 0.662 |
| **MFSD5** | **S440** | -0.05 | 0.02 | 0.03 | 0.09 | -0.06 | -0.04 | 0.901 |
| **MFSD6** | **T10** | -0.13 | 0.11 | 0.13 | 0.00 | -0.29 | 0.13 | 0.572 |
| **MFSD6** | **T10** | -0.08 | 0.10 | 0.19 | -0.11 | -0.06 | -0.06 | 0.151 |
| **MGA** | **S989** | -0.15 | 0.13 | 0.11 | -0.14 | 0.28 | -0.32 | 0.682 |
| **MGAT4A** | **S474** | 0.11 | 0.07 | 0.55 | -0.31 | -0.25 | -0.39 | 0.024 |
| **MGMT** | **S201** | 0.25 | 0.24 | 0.03 | -0.13 | -0.22 | -0.27 | 0.01 |
| **MIA2** | **S1243** | -0.01 | -0.33 | -0.19 | 0.14 | 0.32 | -0.02 | 0.074 |
| **MIA2** | **S1202; S1204** | -0.13 | -0.11 | -0.20 | -0.02 | -0.09 | 0.44 | 0.195 |
| **MIA2** | **S1198; S1202; S1204** | 0.06 | -0.19 | -0.04 | 0.05 | -0.10 | 0.19 | 0.394 |
| **MIA2** | **S1204** | 0.10 | 0.11 | -0.15 | 0.04 | -0.18 | 0.05 | 0.68 |
| **MIA2** | **S745** | 0.26 | 0.26 | -0.07 | -0.11 | -0.07 | -0.38 | 0.086 |
| **MIA3** | **S/T** | -0.03 | -0.11 | -0.29 | 0.13 | 0.19 | 0.06 | 0.035 |
| **MIA3** | **S1906** | -0.11 | 0.02 | -0.10 | 0.09 | 0.00 | 0.08 | 0.08 |
| **MIA3** | **S727** | -0.06 | -0.04 | -0.06 | 0.04 | -0.06 | 0.16 | 0.193 |
| **MIA3** | **S727** | 0.00 | -0.07 | -0.06 | 0.06 | -0.09 | 0.14 | 0.309 |
| **MIA3** | **S/T** | 0.23 | 0.08 | -0.28 | -0.12 | 0.22 | -0.22 | 0.808 |
| **MIA3** | **S1906** | 0.02 | 0.16 | -0.11 | 0.05 | 0.00 | -0.14 | 0.602 |
| **MIB1** | **S805** | 0.04 | 0.06 | -0.21 | 0.23 | -0.28 | 0.08 | 0.804 |
| **MIB1** | **S805** | -0.14 | 0.25 | -0.12 | 0.13 | -0.22 | 0.05 | 0.947 |
| **MIB1** | **S805** | -0.07 | 0.13 | 0.03 | -0.10 | 0.16 | -0.17 | 0.587 |
| **MIB1** | **S805** | -0.04 | 0.24 | -0.08 | 0.13 | 0.00 | -0.31 | 0.588 |
| **MIB2** | **S251** | -0.58 | -0.55 | 0.01 | 0.30 | 0.14 | 0.39 | 0.034 |
| **MICAL3** | **S1649** | -0.03 | 0.13 | -0.17 | 0.11 | -0.02 | -0.04 | 0.675 |
| **MICAL3** | **S1156** | 0.01 | -0.28 | 0.21 | 0.17 | -0.14 | -0.04 | 0.913 |
| **MICAL3** | **S1310** | -0.11 | -0.05 | 0.12 | 0.03 | -0.11 | 0.10 | 0.891 |
| **MICAL3** | **S1134** | 0.08 | -0.18 | 0.09 | -0.02 | -0.02 | 0.02 | 0.979 |
| **MICAL3** | **S1794; S1800; S1801** | -0.05 | 0.03 | 0.05 | 0.00 | -0.03 | -0.01 | 0.502 |
| **MICAL3** | **S1337** | 0.19 | 0.19 | 0.05 | -0.82 | 0.36 | -0.26 | 0.329 |
| **MICALL1** | **S640; S644** | 0.31 | 0.00 | 0.08 | -0.24 | -0.18 | -0.04 | 0.066 |
| **MICALL1** | **S578** | 0.40 | -0.03 | 0.22 | -0.11 | -0.37 | -0.26 | 0.039 |
| **MICALL2** | **S249** | -0.22 | -0.19 | 0.04 | 0.07 | -0.14 | 0.36 | 0.258 |
| **MIDEAS** | **S700** | -0.15 | -0.17 | -0.11 | 0.18 | -0.01 | 0.22 | 0.022 |
| **MIDEAS** | **T704; T715** | -0.09 | -0.07 | -0.05 | 0.15 | -0.42 | 0.36 | 0.688 |
| **MIDEAS** | **T704** | 0.03 | 0.17 | 0.05 | -0.12 | -0.12 | -0.03 | 0.035 |
| **MIDEAS** | **S461** | -0.03 | 0.21 | 0.12 | -0.45 | -0.21 | 0.25 | 0.342 |
| **MIEF1** | **S59** | -0.04 | -0.14 | -0.07 | 0.08 | -0.05 | 0.18 | 0.101 |
| **MIER1** | **S369; S372** | -0.22 | -0.34 | 0.26 | 0.15 | 0.04 | 0.03 | 0.407 |
| **MIER1** | **S488** | -0.13 | -0.04 | -0.03 | 0.05 | 0.06 | 0.08 | 0.016 |
| **MIER1** | **S52** | -0.10 | -0.05 | 0.03 | 0.07 | 0.07 | -0.04 | 0.238 |
| **MIER1** | **S483** | -0.08 | -0.06 | 0.04 | -0.09 | -0.11 | 0.26 | 0.68 |
| **MIER1** | **S377** | 0.04 | 0.06 | -0.09 | -0.03 | -0.24 | 0.21 | 0.881 |
| **MIER1** | **S377** | 0.06 | 0.04 | -0.03 | -0.06 | -0.22 | 0.18 | 0.667 |
| **MIER1** | **S488** | 0.23 | -0.05 | -0.03 | -0.17 | -0.07 | 0.06 | 0.39 |
| **MIER1** | **S166** | 0.11 | 0.11 | 0.07 | -0.23 | 0.01 | -0.11 | 0.044 |
| **MIER1** | **S377** | 0.13 | 0.16 | 0.03 | -0.22 | -0.09 | -0.04 | 0.032 |
| **MIER1** | **S166** | 0.12 | 0.08 | 0.19 | -0.20 | -0.14 | -0.11 | 0.002 |
| **MIER1** | **S377** | 0.26 | 0.29 | -0.15 | -0.12 | -0.18 | -0.18 | 0.109 |

| | | | | | | | | |
|---|---|---|---|---|---|---|---|---|
| **MIER1** | **S377** | 0.40 | 0.23 | -0.10 | -0.30 | -0.16 | -0.22 | 0.06 |
| **MIER1** | **S377** | 0.35 | 0.10 | 0.10 | -0.25 | -0.24 | -0.17 | 0.01 |
| **MIER1** | **S488** | 0.16 | 0.22 | 0.12 | -1.50 | 0.54 | 0.23 | 0.552 |
| **MIER1** | **S483; S488** | 0.47 | 0.50 | -0.11 | -0.18 | -0.24 | -0.87 | 0.072 |
| **MIER3** | **S52** | -0.02 | 0.24 | 0.10 | 0.10 | 0.00 | -0.53 | 0.294 |
| **MIER3** | **S156** | -0.11 | 0.43 | 0.15 | -0.40 | 0.16 | -0.43 | 0.201 |
| **MIGA1** | **S122** | 0.20 | 0.20 | -0.15 | -0.21 | 0.16 | -0.29 | 0.343 |
| **MIGA2** | **S111** | -0.07 | -0.23 | 0.24 | 0.06 | -0.26 | 0.19 | 0.944 |
| **MIIP** | **S303** | 0.05 | -0.09 | -0.30 | -0.58 | -0.44 | 0.85 | 0.915 |
| **MINK1** | **S778** | 0.24 | 0.33 | 0.13 | -0.35 | -0.10 | -0.41 | 0.01 |
| **MIOS** | **S766** | 0.30 | -0.07 | 0.09 | -0.39 | -0.04 | 0.02 | 0.22 |
| **MIS18A** | **S227** | 0.25 | 0.11 | -0.14 | -0.10 | -0.03 | -0.14 | 0.247 |
| **MIS18BP1** | **T161** | -0.46 | -1.01 | -0.23 | 0.08 | 0.13 | 0.83 | 0.052 |
| **MIS18BP1** | **S860** | -0.07 | -0.55 | -0.22 | 0.35 | -0.05 | 0.35 | 0.064 |
| **MIS18BP1** | **S/T** | -0.36 | -0.18 | -0.05 | 0.10 | 0.23 | 0.18 | 0.02 |
| **MIS18BP1** | **S/T** | -0.11 | -0.44 | 0.01 | -0.03 | 0.13 | 0.32 | 0.132 |
| **MIS18BP1** | **S1104** | -0.13 | -0.03 | -0.32 | 0.12 | 0.16 | 0.14 | 0.025 |
| **MIS18BP1** | **T702** | -0.50 | 0.25 | -0.30 | 0.50 | -0.20 | 0.00 | 0.405 |
| **MIS18BP1** | **T653** | -0.06 | -0.15 | -0.24 | 0.13 | -0.25 | 0.44 | 0.282 |
| **MIS18BP1** | **S991** | -0.18 | -0.13 | -0.10 | 0.21 | 0.09 | 0.07 | 0.006 |
| **MIS18BP1** | **S110** | -0.14 | -0.11 | 0.03 | 0.07 | 0.02 | 0.11 | 0.07 |
| **MIS18BP1** | **S824** | 0.14 | -0.04 | -0.10 | -0.10 | 0.04 | 0.05 | 0.979 |
| **MIS18BP1** | **S772; S773** | -0.03 | 0.04 | -0.04 | 0.17 | -0.48 | 0.23 | 0.949 |
| **MIS18BP1** | **S634** | -0.17 | 0.07 | 0.13 | -0.11 | -0.05 | 0.10 | 0.799 |
| **MIS18BP1** | **S172** | -0.01 | -0.07 | 0.17 | -0.05 | -0.09 | 0.04 | 0.506 |
| **MIS18BP1** | **S** | -0.27 | 0.06 | 0.27 | -0.07 | -0.26 | 0.18 | 0.753 |
| **MIS18BP1** | **S135** | 0.09 | -0.13 | 0.27 | 0.01 | -0.12 | -0.17 | 0.255 |
| **MIS18BP1** | **S541** | 0.13 | 0.18 | 0.23 | -0.38 | 0.21 | -0.56 | 0.144 |
| **MIS18BP1** | **S541** | 0.13 | 0.23 | 0.20 | -0.58 | 0.17 | -0.33 | 0.123 |
| **MIS18BP1** | **S541** | 0.24 | 0.39 | -0.05 | -0.21 | 0.06 | -0.66 | 0.131 |
| **MIS18BP1** | **S365** | 0.11 | -0.02 | 0.56 | -0.20 | -0.36 | -0.30 | 0.05 |
| **MISP** | **S394** | -1.03 | 0.22 | -0.13 | 0.12 | 0.23 | 0.22 | 0.246 |
| **MISP** | **S394** | -0.20 | -0.30 | -0.18 | 0.19 | 0.02 | 0.36 | 0.015 |
| **MISP** | **T377** | -0.01 | -0.01 | -0.41 | 0.12 | 0.31 | -0.08 | 0.217 |
| **MISP** | **S** | -0.10 | 0.12 | -0.03 | -0.03 | -0.02 | 0.05 | 0.964 |
| **MISP3** | **S91** | -0.16 | 0.02 | -0.15 | -0.20 | 0.02 | 0.39 | 0.416 |
| **MISP3** | **S91** | -0.01 | 0.21 | -0.06 | -0.01 | -0.11 | -0.04 | 0.318 |
| **MKI67** | **S1740** | -0.76 | -0.93 | -0.93 | 0.52 | 0.36 | 0.72 | 3E-04 |
| **MKI67** | **T2212** | -0.36 | -0.83 | -0.89 | 0.34 | 0.49 | 0.53 | 0.003 |
| **MKI67** | **S1740** | -0.57 | -0.67 | -0.59 | 0.34 | 0.35 | 0.57 | 2E-04 |
| **MKI67** | **S2344; S/T** | -0.10 | -0.61 | -0.72 | 0.58 | 0.22 | 0.19 | 0.024 |
| **MKI67** | **T2259** | -0.43 | -0.40 | -0.53 | 0.33 | 0.32 | 0.38 | 5E-05 |
| **MKI67** | **S2135** | -0.27 | -0.43 | -0.69 | -0.03 | 0.57 | 0.43 | 0.023 |
| **MKI67** | **T2389** | -0.50 | -0.22 | -0.58 | 0.32 | 0.22 | 0.44 | 0.004 |
| **MKI67** | **S2072** | -0.42 | -0.41 | -0.43 | 0.27 | 0.32 | 0.38 | 3E-05 |
| **MKI67** | **S2793** | -0.38 | -0.33 | -0.58 | 0.38 | -0.03 | 0.57 | 0.019 |
| **MKI67** | **T2406** | -1.53 | -0.07 | 0.00 | -0.56 | 0.40 | 0.77 | 0.311 |
| **MKI67** | **T1897** | -0.31 | -0.34 | -0.55 | 0.23 | 0.32 | 0.38 | 0.001 |

| | | | | | | | | |
|---|---|---|---|---|---|---|---|---|
| MKI67 | T2325 | -0.36 | -0.30 | -0.53 | 0.15 | 0.39 | 0.38 | 0.003 |
| MKI67 | T2426 | -0.43 | -0.43 | -0.26 | 0.24 | 0.18 | 0.45 | 0.003 |
| MKI67 | T1335 | -0.17 | -0.34 | -0.65 | -0.33 | 0.69 | 0.35 | 0.131 |
| MKI67 | T2389 | -0.28 | -0.35 | -0.42 | 0.17 | 0.15 | 0.50 | 0.006 |
| MKI67 | S1861; S1864 | -0.28 | -0.33 | -0.48 | 0.38 | -0.19 | 0.57 | 0.058 |
| MKI67 | S2223; T2231 | -0.23 | -0.29 | -0.43 | 0.21 | 0.12 | 0.42 | 0.006 |
| MKI67 | T2380; S2382 | -0.31 | -0.24 | -0.34 | 0.18 | 0.22 | 0.33 | 5E-04 |
| MKI67 | T1906 | -0.22 | -0.33 | -0.34 | 0.36 | 0.11 | 0.26 | 0.003 |
| MKI67 | T2268 | -0.30 | -0.28 | -0.28 | 0.14 | 0.35 | 0.22 | 0.001 |
| MKI67 | S2695 | -0.27 | -0.43 | -0.17 | 0.01 | 0.32 | 0.36 | 0.018 |
| MKI67 | T1923 | -0.23 | -0.28 | -0.30 | 0.08 | 0.25 | 0.35 | 0.004 |
| MKI67 | T2065 | -0.28 | -0.34 | -0.23 | 0.13 | -0.08 | 0.57 | 0.064 |
| MKI67 | T1355 | -0.60 | -0.02 | -0.23 | 0.48 | -0.04 | 0.18 | 0.098 |
| MKI67 | S2588 | -0.15 | -0.24 | -0.41 | 0.01 | 0.23 | 0.39 | 0.023 |
| MKI67 | S2588 | -0.16 | -0.28 | -0.23 | 0.14 | 0.19 | 0.25 | 1E-03 |
| MKI67 | S2223 | -0.42 | -0.13 | -0.10 | -0.05 | 0.33 | 0.25 | 0.064 |
| MKI67 | T1849 | -0.23 | -0.25 | -0.15 | 0.37 | -0.02 | 0.18 | 0.03 |
| MKI67 | S2453 | -0.12 | 0.01 | -0.59 | 0.24 | 0.48 | -0.27 | 0.252 |
| MKI67 | T/S | -0.22 | -0.05 | -0.35 | 0.17 | 0.16 | 0.21 | 0.012 |
| MKI67 | T2446 | -0.31 | -0.15 | -0.15 | 0.04 | 0.21 | 0.27 | 0.013 |
| MKI67 | S2708 | -0.10 | -0.17 | -0.29 | 0.07 | 0.15 | 0.27 | 0.011 |
| MKI67 | S2941 | -0.10 | -0.23 | -0.20 | 0.00 | 0.25 | 0.21 | 0.018 |
| MKI67 | T2380 | 0.07 | -0.15 | -0.49 | -0.06 | 0.10 | 0.38 | 0.184 |
| MKI67 | S1983; T/S | -0.21 | -0.07 | -0.24 | 0.21 | 0.24 | -0.01 | 0.028 |
| MKI67 | S2261 | -0.03 | -0.30 | -0.19 | 0.03 | 0.09 | 0.32 | 0.054 |
| MKI67 | T2380 | -0.23 | -0.11 | -0.17 | 0.07 | 0.27 | 0.10 | 0.011 |
| MKI67 | T2085 | -0.13 | -0.18 | -0.19 | 0.07 | 0.16 | 0.22 | 0.003 |
| MKI67 | S648 | -0.13 | -0.56 | 0.15 | 0.19 | -0.03 | 0.23 | 0.238 |
| MKI67 | T1849 | -0.19 | 0.22 | -0.62 | 0.21 | -0.36 | 0.46 | 0.429 |
| MKI67 | T/S | -0.08 | 0.07 | -0.51 | 0.02 | 0.40 | -0.05 | 0.257 |
| MKI67 | S2466 | -0.03 | -0.08 | -0.29 | 0.15 | 0.01 | 0.18 | 0.061 |
| MKI67 | S1815 | 0.13 | -0.24 | -0.31 | 0.25 | -0.18 | 0.24 | 0.29 |
| MKI67 | T2446 | -0.11 | -0.17 | -0.03 | 0.15 | 0.12 | 0.01 | 0.029 |
| MKI67 | T2446 | -0.10 | -0.07 | -0.12 | 0.07 | 0.07 | 0.14 | 0.002 |
| MKI67 | S2223 | 0.06 | 0.02 | -0.38 | 0.16 | 0.02 | 0.06 | 0.281 |
| MKI67 | S1861 | -0.03 | -0.10 | -0.08 | 0.11 | 0.01 | 0.08 | 0.021 |
| MKI67 | T2949 | 0.08 | -0.18 | -0.14 | -0.17 | 0.35 | -0.02 | 0.487 |
| MKI67 | S1815 | -0.51 | 0.27 | 0.00 | 0.18 | 0.24 | -0.36 | 0.751 |
| MKI67 | S1118 | -0.11 | 0.01 | -0.06 | 0.17 | 0.16 | -0.21 | 0.5 |
| MKI67 | S2344 | 0.01 | 0.06 | -0.22 | 0.02 | -0.04 | 0.15 | 0.433 |
| MKI67 | S1131 | 0.00 | 0.04 | -0.17 | 0.07 | -0.14 | 0.18 | 0.513 |
| MKI67 | S2828 | -0.01 | -0.01 | -0.08 | 0.00 | -0.05 | 0.15 | 0.344 |
| MKI67 | S2105 | -0.01 | -0.04 | -0.03 | -0.03 | 0.09 | 0.02 | 0.253 |
| MKI67 | S1983 | -0.04 | 0.06 | -0.05 | -0.02 | -0.06 | 0.11 | 0.741 |
| MKI67 | S357 | -0.49 | 0.15 | 0.26 | -0.10 | -0.13 | 0.19 | 0.964 |
| MKI67 | S1376 | -0.08 | 0.11 | -0.04 | 0.05 | -0.03 | -0.02 | 0.989 |
| MKI67 | S2466 | 0.14 | 0.05 | -0.20 | 0.00 | 0.12 | -0.14 | 0.967 |
| MKI67 | T/S | -0.04 | -0.06 | 0.11 | -0.03 | 0.13 | -0.14 | 0.879 |

| | | | | | | | | |
|---|---|---|---|---|---|---|---|---|
| **MKI67** | **T/S** | -0.29 | 0.21 | 0.09 | -0.13 | 0.05 | 0.01 | 0.874 |
| **MKI67** | **S2344** | 0.00 | 0.03 | 0.01 | -0.16 | 0.15 | -0.04 | 0.756 |
| **MKI67** | **S584** | 0.04 | -0.05 | 0.06 | -0.07 | 0.05 | -0.03 | 0.556 |
| **MKI67** | **S1131** | -0.09 | 0.11 | 0.04 | -0.01 | 0.03 | -0.07 | 0.603 |
| **MKI67** | **S2638** | 0.21 | -0.15 | -0.01 | -0.28 | -0.23 | 0.35 | 0.775 |
| **MKI67** | **S1376** | 0.03 | 0.05 | 0.03 | -0.01 | -0.01 | -0.10 | 0.063 |
| **MKI67** | **S1861** | 0.05 | 0.14 | -0.09 | -0.07 | 0.18 | -0.25 | 0.594 |
| **MKI67** | **S2223** | -0.10 | 0.09 | 0.11 | -0.26 | 0.17 | -0.05 | 0.589 |
| **MKI67** | **S2105** | 0.05 | 0.02 | 0.04 | 0.01 | 0.12 | -0.28 | 0.52 |
| **MKI67** | **S2105** | 0.04 | 0.02 | 0.06 | -0.05 | 0.07 | -0.16 | 0.278 |
| **MKI67** | **S1253** | 0.05 | 0.05 | 0.02 | -0.08 | -0.25 | 0.18 | 0.497 |
| **MKI67** | **S2828** | 0.09 | 0.03 | 0.03 | -0.07 | 0.02 | -0.11 | 0.074 |
| **MKI67** | **S308** | 0.15 | 0.01 | 0.02 | -0.14 | 0.01 | -0.06 | 0.129 |
| **MKI67** | **T2450** | 0.28 | 0.03 | -0.07 | -0.07 | 0.14 | -0.40 | 0.384 |
| **MKI67** | **S2505** | 0.06 | 0.01 | 0.20 | -0.18 | 0.03 | -0.16 | 0.095 |
| **MKI67** | **S357** | 0.10 | 0.18 | 0.02 | -0.07 | -0.10 | -0.17 | 0.018 |
| **MKI67** | **S1983** | 0.04 | 0.09 | 0.19 | -0.08 | 0.04 | -0.34 | 0.128 |
| **MKI67** | **T/S** | 0.02 | 0.27 | 0.02 | -0.44 | 0.29 | -0.30 | 0.343 |
| **MKI67** | **S2505** | 0.12 | 0.09 | 0.17 | -0.21 | 0.13 | -0.40 | 0.147 |
| **MKI67** | **S308** | 0.32 | 0.14 | 0.06 | -0.25 | -0.10 | -0.27 | 0.016 |
| **MLEC** | **Y/S** | 0.12 | -0.26 | -0.04 | 0.05 | 0.10 | 0.00 | 0.4 |
| **MLF2** | **S** | -0.16 | 0.14 | -0.04 | 0.21 | -0.12 | -0.06 | 0.824 |
| **MLF2** | **S238** | 0.05 | -0.01 | 0.09 | 0.03 | -0.04 | -0.12 | 0.196 |
| **MLLT1** | **S359; S361** | -0.01 | 0.02 | 0.02 | -0.05 | -0.08 | 0.10 | 0.729 |
| **MLLT1** | **S361** | 0.05 | 0.01 | -0.01 | -0.05 | -0.14 | 0.13 | 0.691 |
| **MLLT1** | **S438; S447; S454** | 0.00 | 0.10 | -0.04 | -0.08 | -0.08 | 0.08 | 0.519 |
| **MLLT1** | **S292** | 0.08 | 0.07 | -0.07 | -0.03 | -0.09 | 0.04 | 0.426 |
| **MLLT1** | **S440** | -0.06 | 0.20 | 0.04 | -0.11 | -0.23 | 0.11 | 0.343 |
| **MLLT1** | **S475** | 0.33 | 0.15 | -0.13 | 0.06 | -0.16 | -0.36 | 0.211 |
| **MLLT1** | **S440; S447** | 0.25 | -0.03 | 0.17 | 0.11 | -0.34 | -0.27 | 0.14 |
| **MLLT1** | **S411; S414** | 0.02 | 0.34 | 0.03 | -0.36 | -0.36 | 0.20 | 0.231 |
| **MLLT1** | **S475** | 0.29 | 0.18 | 0.13 | -0.13 | -0.10 | -0.50 | 0.031 |
| **MLLT1** | **S** | 0.63 | 0.72 | 0.00 | -0.99 | -0.95 | -0.36 | 0.016 |
| **MLLT10** | **S/Y** | -0.15 | -0.04 | -0.29 | 0.28 | -0.08 | 0.20 | 0.09 |
| **MLLT10** | **S370** | -0.51 | -0.17 | 0.21 | -0.72 | 0.15 | 0.62 | 0.717 |
| **MLLT10** | **S686; S689** | 0.04 | 0.02 | -0.18 | -0.13 | -0.04 | 0.25 | 0.633 |
| **MLLT10** | **S689** | 0.12 | 0.16 | -0.02 | 0.07 | -0.19 | -0.18 | 0.135 |
| **MLLT6** | **S423** | 0.12 | 0.02 | -0.10 | 0.06 | -0.10 | -0.01 | 0.693 |
| **MLLT6** | **S258** | -0.03 | 0.21 | -0.02 | 0.00 | -0.16 | -0.02 | 0.283 |
| **MLLT6** | **S/T** | 0.18 | 0.05 | -0.07 | 0.08 | -0.10 | -0.18 | 0.328 |
| **MLLT6** | **S222** | 0.16 | 0.14 | 0.05 | 0.00 | -0.01 | -0.39 | 0.134 |
| **MLLT6** | **S382** | 0.42 | 0.09 | -0.12 | -0.01 | -0.17 | -0.33 | 0.174 |
| **MLLT6** | **S382** | 0.05 | 0.43 | 0.07 | -0.08 | -0.05 | -0.62 | 0.128 |
| **MLPH** | **S/T** | 0.10 | -0.90 | -0.50 | 0.16 | 0.49 | 0.21 | 0.08 |
| **MLPH** | **S337** | -0.06 | -0.14 | -0.42 | 0.24 | -0.03 | 0.30 | 0.062 |
| **MLPH** | **S402** | -0.06 | -0.01 | -0.15 | 0.19 | -0.01 | 0.01 | 0.134 |
| **MLPH** | **S444** | -0.33 | -0.08 | 0.17 | 0.03 | 0.09 | 0.06 | 0.392 |
| **MLPH** | **S314** | 0.03 | -0.13 | -0.04 | -0.02 | 0.09 | 0.06 | 0.181 |

| | | | | | | | | |
|---|---|---|---|---|---|---|---|---|
| MLPH | S231 | -0.08 | 0.03 | -0.01 | -0.06 | 0.00 | 0.11 | 0.547 |
| MLPH | S231; S233 | -0.12 | -0.09 | 0.17 | -0.08 | -0.18 | 0.24 | 0.99 |
| MLPH | S159 | -0.02 | 0.12 | -0.02 | -0.03 | 0.14 | -0.22 | 0.634 |
| MLPH | S397 | -0.05 | 0.05 | 0.09 | 0.07 | 0.03 | -0.21 | 0.524 |
| MLPH | S481 | -0.06 | 0.08 | 0.09 | 0.03 | -0.22 | 0.05 | 0.461 |
| MLPH | S/T | 0.07 | 0.28 | -0.30 | -0.24 | -0.28 | 0.32 | 0.757 |
| MLPH | T457; T458; S463 | 0.03 | 0.01 | 0.15 | 0.22 | -0.01 | -0.49 | 0.501 |
| MLPH | S314 | 0.07 | 0.13 | 0.03 | -0.01 | -0.19 | -0.06 | 0.055 |
| MLPH | S155 | -0.16 | 0.05 | 0.33 | 0.08 | -0.01 | -0.39 | 0.426 |
| MLPH | S397; T/S | -0.01 | 0.05 | 0.17 | 0.34 | -0.20 | -0.49 | 0.501 |
| MLPH | S444 | 0.07 | 0.12 | 0.08 | -0.16 | 0.06 | -0.20 | 0.086 |
| MLPH | S/T | -0.11 | 0.06 | 0.33 | -0.29 | -0.13 | 0.06 | 0.266 |
| MLPH | T/S | -0.02 | 0.04 | 0.31 | -0.05 | -0.01 | -0.35 | 0.171 |
| MLPH | S266 | 0.02 | 0.09 | 0.24 | -0.23 | -0.12 | -0.05 | 0.037 |
| MLPH | T458 | -0.27 | 0.20 | 0.38 | -0.65 | -0.28 | 0.33 | 0.427 |
| MLPH | S159; S162; T165 | 0.03 | 0.00 | 0.39 | -0.16 | -0.15 | -0.20 | 0.067 |
| MLPH | S162; T165 | 0.16 | 0.01 | 0.29 | -0.14 | -0.03 | -0.37 | 0.064 |
| MLPH | S162 | 0.08 | 0.14 | 0.26 | -0.22 | 0.06 | -0.42 | 0.076 |
| MLPH | S162; T165 | 0.07 | 0.13 | 0.29 | -0.20 | -0.06 | -0.31 | 0.022 |
| MLPH | S159; S162; T165 | -0.06 | 0.22 | 0.33 | -0.39 | -0.07 | -0.15 | 0.072 |
| MLPH | T401; S402 | 0.13 | 0.14 | 0.15 | 0.36 | -0.29 | -0.75 | 0.315 |
| MLPH | T165 | 0.03 | 0.30 | 0.30 | -0.33 | -0.09 | -0.35 | 0.02 |
| MME | S6 | 0.03 | -0.03 | -0.17 | -0.08 | 0.20 | 0.02 | 0.358 |
| MMTAG2 | S217; S220 | -0.03 | -0.09 | 0.03 | 0.06 | -0.05 | 0.08 | 0.323 |
| MMTAG2 | S217 | -0.02 | -0.01 | -0.04 | 0.14 | 0.03 | -0.11 | 0.594 |
| MMTAG2 | S216; S217; S220 | -0.11 | -0.07 | 0.17 | -0.13 | -0.15 | 0.25 | 0.981 |
| MMTAG2 | S155 | 0.14 | -0.07 | 0.01 | 0.02 | -0.03 | -0.09 | 0.447 |
| MMTAG2 | S123 | 0.03 | 0.09 | 0.26 | -0.45 | 0.38 | -0.56 | 0.336 |
| MON1B | S61 | -0.14 | 0.03 | -0.03 | 0.03 | -0.23 | 0.28 | 0.662 |
| MON1B | S59 | -0.17 | 0.10 | 0.04 | 0.02 | -0.17 | 0.15 | 0.944 |
| MORC2 | S615 | -0.20 | -0.20 | -0.16 | 0.29 | 0.11 | 0.10 | 0.005 |
| MORC2 | S777; S779 | 0.02 | 0.00 | 0.02 | 0.00 | 0.08 | -0.12 | 0.635 |
| MORC2 | S725 | -0.09 | -0.21 | 0.31 | 0.34 | -0.38 | -0.12 | 0.836 |
| MORC2 | S725; S730 | 0.05 | 0.19 | -0.01 | -0.03 | 0.05 | -0.28 | 0.23 |
| MORC2 | S779 | 0.16 | 0.16 | 0.05 | -0.12 | -0.15 | -0.14 | 0.002 |
| MORC2 | S777 | 0.15 | 0.02 | 0.29 | -0.24 | 0.06 | -0.40 | 0.091 |
| MORC2 | S779 | 0.30 | 0.45 | 0.05 | -0.21 | -0.35 | -0.48 | 0.012 |
| MORC4 | S465 | -0.02 | -0.17 | -0.06 | 0.03 | -0.09 | 0.26 | 0.263 |
| MORC4 | S465 | -0.21 | 0.01 | -0.01 | 0.14 | -0.01 | 0.06 | 0.172 |
| MORC4 | T621 | 0.00 | -0.18 | 0.15 | 0.00 | -0.03 | 0.04 | 0.886 |
| MOV10 | S969 | -0.12 | -0.11 | 0.41 | 0.17 | -0.13 | -0.34 | 0.519 |
| MPDZ | S483 | -0.16 | -0.17 | -0.27 | -0.18 | 0.12 | 0.50 | 0.157 |
| MPHOSPH10 | S163; S167; S171 | -0.26 | -0.37 | -0.34 | 0.25 | -0.08 | 0.55 | 0.039 |
| MPHOSPH10 | S163 | -0.10 | -0.11 | -0.06 | 0.18 | -0.07 | 0.13 | 0.094 |
| MPHOSPH10 | S163; S171 | -0.06 | 0.04 | -0.17 | 0.15 | -0.16 | 0.16 | 0.417 |
| MPHOSPH10 | S242 | -0.05 | -0.01 | 0.08 | 0.19 | -0.14 | -0.10 | 0.855 |
| MPHOSPH10 | S242 | -0.05 | -0.01 | 0.08 | 0.19 | -0.14 | -0.10 | 0.855 |
| MPHOSPH10 | S163 | 0.05 | 0.07 | 0.15 | -0.18 | 0.23 | -0.42 | 0.328 |

| | | | | | | | | |
|---|---|---|---|---|---|---|---|---|
| **MPHOSPH6** | **S110** | -0.11 | -0.16 | -0.15 | 0.11 | -0.12 | 0.36 | 0.141 |
| **MPHOSPH6** | **T147** | -0.06 | -0.12 | -0.01 | 0.07 | -0.10 | 0.19 | 0.257 |
| **MPHOSPH6** | **S110** | -0.02 | -0.05 | -0.01 | 0.01 | -0.02 | 0.09 | 0.176 |
| **MPHOSPH6** | **S110** | -0.08 | 0.03 | 0.00 | -0.06 | -0.14 | 0.22 | 0.82 |
| **MPHOSPH6** | **T147** | 0.10 | 0.03 | -0.05 | 0.05 | -0.15 | 0.01 | 0.514 |
| **MPHOSPH8** | **S264; S272** | -0.37 | -0.15 | -0.14 | 0.16 | 0.26 | 0.15 | 0.007 |
| **MPHOSPH8** | **S264; S279** | -0.16 | -0.21 | -0.14 | 0.12 | 0.16 | 0.18 | 3E-04 |
| **MPHOSPH8** | **S51** | -0.09 | -0.20 | -0.13 | 0.13 | -0.07 | 0.30 | 0.081 |
| **MPHOSPH8** | **S/T** | -0.16 | -0.18 | 0.05 | 0.09 | -0.12 | 0.28 | 0.26 |
| **MPHOSPH8** | **S164** | 0.00 | -0.11 | -0.17 | -0.06 | -0.03 | 0.31 | 0.269 |
| **MPHOSPH8** | **S20** | -0.10 | -0.25 | 0.09 | 0.16 | -0.19 | 0.22 | 0.39 |
| **MPHOSPH8** | **S403** | 0.04 | -0.09 | -0.19 | 0.05 | -0.09 | 0.24 | 0.275 |
| **MPHOSPH8** | **S** | -0.12 | -0.03 | -0.07 | 0.10 | 0.00 | 0.12 | 0.03 |
| **MPHOSPH8** | **T/S/Y** | 0.00 | -0.06 | -0.13 | 0.08 | -0.06 | 0.15 | 0.174 |
| **MPHOSPH8** | **S164** | 0.05 | -0.07 | -0.14 | 0.03 | 0.00 | 0.12 | 0.191 |
| **MPHOSPH8** | **S/T** | -0.05 | -0.02 | -0.08 | 0.11 | -0.12 | 0.14 | 0.324 |
| **MPHOSPH8** | **S319** | -0.06 | 0.03 | -0.13 | 0.20 | 0.10 | -0.18 | 0.492 |
| **MPHOSPH8** | **S284** | -0.06 | 0.04 | -0.06 | -0.03 | -0.04 | 0.14 | 0.483 |
| **MPHOSPH8** | **S425** | 0.09 | 0.06 | -0.24 | 0.07 | -0.04 | 0.04 | 0.672 |
| **MPHOSPH8** | **S403** | 0.00 | -0.09 | 0.02 | -0.19 | -0.03 | 0.26 | 0.802 |
| **MPHOSPH8** | **S319** | -0.08 | 0.08 | 0.02 | 0.20 | -0.11 | -0.13 | 0.846 |
| **MPHOSPH8** | **S164** | 0.07 | -0.01 | 0.01 | -0.05 | 0.04 | -0.05 | 0.361 |
| **MPHOSPH8** | **S138** | 0.16 | 0.12 | 0.22 | -0.14 | -0.02 | -0.44 | 0.045 |
| **MPHOSPH9** | **S516** | -0.14 | -0.16 | -0.04 | 0.08 | 0.08 | 0.15 | 0.006 |
| **MPND** | **S214** | -0.05 | -0.09 | -0.04 | 0.02 | 0.05 | 0.10 | 0.011 |
| **MPRIP** | **S224; S/T** | -0.29 | -0.04 | -0.45 | -0.02 | 0.00 | 0.58 | 0.127 |
| **MPRIP** | **S220; S224** | -0.32 | -0.06 | -0.20 | 0.06 | 0.05 | 0.37 | 0.056 |
| **MPRIP** | **S289; S292** | -0.09 | -0.25 | -0.20 | 0.26 | -0.10 | 0.29 | 0.067 |
| **MPRIP** | **S365** | -0.24 | 0.05 | -0.39 | 0.23 | -0.28 | 0.44 | 0.27 |
| **MPRIP** | **T383** | -0.29 | -0.24 | 0.17 | 0.02 | 0.05 | 0.22 | 0.237 |
| **MPRIP** | **S977** | -0.02 | 0.02 | -0.17 | 0.08 | 0.05 | 0.02 | 0.149 |
| **MPRIP** | **S292** | 0.08 | 0.03 | -0.20 | -0.18 | -0.10 | 0.30 | 0.82 |
| **MPRIP** | **S977** | -0.01 | 0.09 | -0.14 | 0.13 | -0.08 | -0.01 | 0.734 |
| **MPRIP** | **S619** | -0.01 | 0.05 | 0.02 | -0.01 | 0.03 | -0.09 | 0.302 |
| **MPRIP** | **S292** | -0.03 | 0.04 | 0.10 | -0.19 | -0.08 | 0.13 | 0.447 |
| **MPRIP** | **S365** | 0.23 | -0.09 | 0.07 | -0.13 | -0.03 | -0.08 | 0.188 |
| **MPRIP** | **S365** | 0.01 | 0.06 | 0.14 | -0.08 | 0.17 | -0.36 | 0.353 |
| **MPRIP** | **S220** | 0.18 | 0.04 | 0.03 | -0.21 | -0.22 | 0.13 | 0.21 |
| **MPRIP** | **S504** | 0.16 | -0.08 | 0.19 | 0.08 | -0.03 | -0.40 | 0.287 |
| **MPRIP** | **S218; S224** | -0.05 | 0.24 | 0.12 | -0.06 | -0.30 | -0.02 | 0.129 |
| **MPRIP** | **S993** | 0.21 | 0.20 | 0.00 | -0.05 | -0.06 | -0.37 | 0.074 |
| **MPRIP** | **S891** | 0.10 | 0.23 | 0.15 | -0.15 | 0.00 | -0.41 | 0.049 |
| **MPRIP** | **S993** | 0.33 | 0.16 | 0.01 | -0.20 | 0.18 | -0.69 | 0.206 |
| **MPRIP** | **T646** | -0.02 | 0.50 | -0.09 | -0.22 | 0.42 | -1.13 | 0.417 |
| **MPRIP** | **S619** | 0.18 | 0.27 | 0.21 | -0.37 | 0.08 | -0.57 | 0.059 |
| **MPRIP** | **S224** | 0.49 | 0.06 | 0.06 | -0.12 | -1.02 | 0.14 | 0.228 |
| **MPRIP** | **S220** | 0.44 | 0.21 | 0.47 | -0.65 | -0.60 | -0.32 | 0.002 |
| **MPZL1** | **S260** | -0.20 | -0.16 | 0.03 | 0.08 | 0.20 | 0.01 | 0.083 |

| | | | | | | | | |
|---|---|---|---|---|---|---|---|---|
| MPZL1 | S/T/Y | 0.10 | 0.31 | 0.10 | -0.30 | -0.50 | 0.13 | 0.114 |
| MRE11 | S688; S689 | -0.32 | -0.23 | -0.08 | 0.04 | 0.21 | 0.28 | 0.018 |
| MRE11 | S688; S689 | -0.01 | -0.18 | -0.15 | -0.04 | 0.03 | 0.29 | 0.142 |
| MRE11 | S688 | 0.12 | -0.06 | -0.13 | 0.17 | -0.07 | -0.06 | 0.723 |
| MRE11 | S/T | 0.05 | 0.07 | -0.07 | 0.11 | -0.12 | -0.06 | 0.667 |
| MREG | S213 | 0.13 | -0.17 | -0.11 | 0.15 | -0.09 | 0.05 | 0.506 |
| MRGBP | S195 | 0.04 | 0.88 | 0.36 | -1.50 | -0.05 | -0.19 | 0.127 |
| MRPL23 | S76 | 0.17 | 0.10 | 0.14 | -0.05 | -0.14 | -0.29 | 0.015 |
| MRPL4 | S40 | -0.24 | 0.13 | -0.05 | 0.21 | -0.23 | 0.12 | 0.63 |
| MRPS15 | S244 | 0.24 | 0.29 | -0.03 | -0.12 | -0.21 | -0.25 | 0.028 |
| MRPS17 | S128 | 0.37 | 0.17 | 0.11 | -0.42 | -0.18 | -0.20 | 0.012 |
| MRPS18B | S38 | 0.23 | 0.14 | -0.22 | -0.07 | -0.17 | 0.03 | 0.475 |
| MRPS18B | S38 | 0.26 | 0.17 | -0.28 | 0.08 | -0.07 | -0.26 | 0.527 |
| MRPS36 | S61 | 0.01 | 0.07 | 0.00 | -0.02 | -0.12 | 0.05 | 0.346 |
| MRPS5 | S386 | 0.04 | -0.01 | -0.10 | -0.13 | 0.05 | 0.14 | 0.639 |
| MRTFA | S385 | 0.28 | 0.11 | 0.15 | -0.08 | -0.72 | 0.07 | 0.159 |
| MRTO4 | S80 | -0.16 | -0.21 | -0.02 | 0.24 | 0.08 | 0.03 | 0.049 |
| MRTO4 | S/T | -0.20 | 0.05 | -0.05 | 0.02 | -0.01 | 0.17 | 0.244 |
| MRTO4 | S233; S235 | 0.02 | 0.10 | 0.19 | -0.32 | -0.11 | 0.06 | 0.13 |
| MSANTD7 | S | -0.20 | -0.05 | -0.02 | -0.09 | -0.27 | 0.49 | 0.609 |
| MSANTD7 | S225 | -0.09 | 0.35 | 0.03 | -0.18 | -0.16 | -0.02 | 0.207 |
| MSH6 | S14 | -0.06 | -0.24 | -0.28 | 0.19 | 0.05 | 0.26 | 0.018 |
| MSH6 | S137 | -0.10 | -0.01 | -0.28 | -0.11 | 0.13 | 0.30 | 0.175 |
| MSH6 | S830 | -0.09 | -0.08 | 0.18 | 0.20 | 0.26 | -0.66 | 0.825 |
| MSH6 | S261 | 0.37 | -0.02 | -0.25 | -0.08 | 0.20 | -0.34 | 0.684 |
| MSL1 | S205 | -0.14 | -0.04 | -0.13 | 0.15 | -0.09 | 0.21 | 0.115 |
| MSL1 | S161 | 0.00 | -0.06 | 0.20 | -0.66 | 0.12 | 0.22 | 0.621 |
| MSL1 | S442 | -0.31 | 0.39 | 0.09 | -0.19 | -0.24 | 0.13 | 0.539 |
| MSL1 | S205 | 0.44 | 0.04 | -0.42 | -1.09 | 0.58 | -0.14 | 0.679 |
| MSL1 | T396; S/T | 0.17 | 0.20 | 0.12 | -0.12 | -0.27 | -0.16 | 0.003 |
| MSL1 | S126 | 0.08 | 0.15 | 0.30 | -0.42 | 0.04 | -0.27 | 0.057 |
| MSL1 | S362 | 0.42 | 0.19 | 0.23 | -0.40 | -0.02 | -0.71 | 0.036 |
| MSL3 | T334; S/T | -0.08 | 0.16 | -0.28 | 0.19 | 0.00 | -0.04 | 0.47 |
| MSL3 | S400 | 0.15 | -0.13 | -0.13 | 0.12 | -0.07 | 0.03 | 0.582 |
| MSL3 | T334 | 0.05 | 0.15 | -0.31 | 0.27 | -0.46 | 0.17 | 0.919 |
| MSL3 | S309 | -0.06 | 0.06 | -0.05 | -0.08 | 0.18 | -0.07 | 0.779 |
| MSL3 | S311; T334 | 0.13 | 0.04 | -0.17 | -0.13 | -0.13 | 0.22 | 0.926 |
| MSL3 | S400; T405 | 0.14 | 0.27 | 0.04 | -0.19 | -0.39 | 0.03 | 0.073 |
| MSL3 | S367 | 0.42 | 0.29 | 0.28 | -0.50 | -0.28 | -0.53 | 0.001 |
| MSX2 | S91 | -0.15 | 0.08 | -0.17 | 0.35 | -0.38 | 0.15 | 0.638 |
| MSX2 | S123 | 0.00 | 0.25 | 0.06 | 0.02 | -0.27 | -0.10 | 0.123 |
| MSX2 | S123 | -0.14 | 0.33 | 0.18 | -0.03 | -0.27 | -0.16 | 0.147 |
| MSX2 | S123 | 0.03 | 0.19 | 0.25 | -0.04 | -0.14 | -0.36 | 0.043 |
| MTA1 | S386 | -0.84 | -0.40 | -0.01 | -0.21 | 0.18 | 0.76 | 0.148 |
| MTA1 | S522 | -0.20 | -0.27 | -0.41 | 0.03 | 0.29 | 0.38 | 0.012 |
| MTA1 | S386 | -0.17 | -0.22 | 0.09 | -0.03 | 0.08 | 0.21 | 0.193 |
| MTA1 | S386 | -0.06 | 0.11 | 0.00 | -0.09 | 0.22 | -0.21 | 0.771 |
| MTA1 | S/T | 0.01 | 0.06 | 0.00 | 0.02 | 0.03 | -0.13 | 0.452 |

| | | | | | | | | |
|---|---|---|---|---|---|---|---|---|
| **MTA1** | **S449** | 0.07 | 0.13 | -0.05 | 0.07 | 0.01 | -0.27 | 0.371 |
| **MTA1** | **S576** | 0.06 | 0.13 | 0.00 | -0.32 | 0.02 | 0.08 | 0.35 |
| **MTA1** | **S576** | -0.09 | 0.17 | 0.10 | 0.27 | -0.20 | -0.34 | 0.491 |
| **MTA1** | **T564** | 0.20 | -0.15 | 0.14 | -0.18 | -0.32 | 0.22 | 0.459 |
| **MTA1** | **S449** | 0.09 | 0.09 | 0.07 | -0.13 | 0.06 | -0.22 | 0.092 |
| **MTA1** | **S576** | 0.20 | 0.18 | 0.09 | -0.30 | -0.11 | -0.13 | 0.008 |
| **MTA2** | **S54** | -0.22 | 0.10 | -0.26 | 0.08 | -0.10 | 0.32 | 0.24 |
| **MTA2** | **S587** | -0.07 | -0.06 | -0.18 | 0.09 | 0.07 | 0.12 | 0.008 |
| **MTA2** | **S435** | 0.21 | -0.49 | -0.22 | -0.12 | -1.14 | 0.92 | 0.938 |
| **MTA3** | **S430** | -0.11 | -0.31 | -0.15 | 0.62 | -0.06 | -0.22 | 0.311 |
| **MTA3** | **S428** | 0.03 | 0.08 | -0.09 | 0.03 | 0.06 | -0.13 | 0.789 |
| **MTA3** | **S428** | 0.03 | 0.00 | 0.01 | -0.01 | 0.02 | -0.05 | 0.242 |
| **MTA3** | **S428; S430** | -0.06 | 0.00 | 0.15 | -0.09 | -0.11 | 0.09 | 0.517 |
| **MTA3** | **S519** | 0.20 | 0.26 | 0.12 | -0.48 | 0.05 | -0.29 | 0.055 |
| **MTBP** | **S639** | 0.16 | 0.43 | 0.39 | 0.16 | -0.67 | -1.05 | 0.082 |
| **MTCH2** | **S98** | -0.12 | 0.11 | 0.21 | -0.24 | 0.05 | -0.06 | 0.316 |
| **MTCL1** | **S306** | 0.07 | -0.03 | 0.14 | 0.09 | -0.32 | 0.01 | 0.382 |
| **MTDH** | **S/T** | 0.28 | -0.18 | -0.80 | -0.08 | 0.53 | -0.10 | 0.403 |
| **MTDH** | **S298** | -0.11 | 0.03 | 0.14 | 0.02 | 0.21 | -0.38 | 0.723 |
| **MTDH** | **S426** | 0.01 | 0.06 | 0.08 | 0.06 | -0.18 | -0.04 | 0.238 |
| **MTDH** | **S308** | 0.25 | -0.06 | 0.01 | -0.05 | 0.14 | -0.36 | 0.425 |
| **MTDH** | **S426** | 0.07 | 0.09 | 0.09 | -0.09 | -0.05 | -0.13 | 0.002 |
| **MTDH** | **T143** | 0.07 | 0.09 | 0.16 | -0.01 | 0.04 | -0.42 | 0.18 |
| **MTDH** | **S568** | 0.25 | 0.26 | 0.41 | -0.36 | -0.21 | -0.65 | 0.007 |
| **MTDH** | **S494** | -0.14 | 0.28 | 0.76 | -0.53 | -0.03 | -0.94 | 0.098 |
| **MTFR1** | **S119** | -0.17 | -0.08 | -0.13 | 0.09 | 0.16 | 0.10 | 0.002 |
| **MTFR1** | **S119** | -0.36 | 0.02 | -0.06 | 0.06 | -0.13 | 0.37 | 0.283 |
| **MTFR1** | **S290** | 0.06 | 0.01 | -0.06 | 0.10 | 0.04 | -0.16 | 0.896 |
| **MTFR1L** | **S103** | 0.18 | 0.17 | 0.19 | -0.32 | 0.18 | -0.58 | 0.133 |
| **MTMR1** | **S43** | 0.16 | 0.28 | -0.17 | -0.25 | -0.17 | 0.07 | 0.289 |
| **MTMR10** | **S769** | 0.33 | 0.14 | 0.13 | -0.09 | -0.34 | -0.28 | 0.011 |
| **MTMR3** | **S633** | -0.07 | -0.15 | 0.13 | 0.08 | -0.04 | 0.04 | 0.562 |
| **MTMR3** | **S613** | -0.37 | -0.05 | 0.50 | -0.39 | -0.11 | 0.21 | 0.7 |
| **MTMR3** | **T731** | 0.04 | 0.15 | -0.01 | -0.10 | -0.19 | 0.08 | 0.226 |
| **MTMR3** | **S/Y/T** | -0.13 | 0.16 | 0.21 | -0.27 | -0.09 | 0.06 | 0.277 |
| **MTOR** | **S2448** | 0.05 | -0.18 | -0.09 | -0.05 | 0.05 | 0.19 | 0.221 |
| **MTOR** | **S2448** | 0.05 | -0.10 | -0.14 | -0.07 | 0.06 | 0.18 | 0.26 |
| **MTSS2** | **S612** | -0.02 | -0.04 | 0.10 | -0.17 | 0.03 | 0.09 | 0.776 |
| **MTUS1** | **S1245** | -0.15 | -0.11 | 0.01 | -0.05 | 0.08 | 0.19 | 0.132 |
| **MTUS1** | **S1203** | -0.13 | 0.14 | -0.02 | 0.23 | -0.06 | -0.21 | 0.968 |
| **MTUS1** | **S958** | -0.09 | 0.27 | 0.29 | -0.14 | -0.02 | -0.45 | 0.114 |
| **MTX3** | **S311** | -0.04 | -0.21 | 0.04 | 0.05 | 0.08 | 0.07 | 0.137 |
| **MTX3** | **S311** | -0.08 | 0.01 | 0.10 | 0.03 | -0.17 | 0.09 | 0.78 |
| **MTX3** | **S284** | 0.01 | 0.13 | -0.06 | -0.17 | -0.19 | 0.23 | 0.676 |
| **MTX3** | **S284** | -0.11 | 0.00 | 0.21 | 0.03 | -0.17 | 0.02 | 0.559 |
| **MTX3** | **S284** | -0.02 | 0.16 | 0.26 | -0.20 | -0.09 | -0.17 | 0.031 |
| **MTX3** | **S249** | 0.19 | 0.36 | 0.08 | -0.26 | -0.17 | -0.32 | 0.008 |
| **MUS81** | **S95** | 0.03 | -0.14 | 0.20 | 0.05 | 0.01 | -0.18 | 0.584 |

| | | | | | | | | |
|---|---|---|---|---|---|---|---|---|
| **MUS81** | **S95** | 0.03 | -0.14 | 0.20 | 0.05 | 0.01 | -0.18 | 0.584 |
| **MVB12A** | **S170** | -0.27 | -0.01 | -0.10 | -0.01 | 0.27 | 0.07 | 0.103 |
| **MVB12A** | **S170** | 0.00 | -0.43 | 0.03 | 0.37 | -0.08 | 0.00 | 0.319 |
| **MYBBP1A** | **S1159; S1163** | -0.20 | -0.12 | -0.22 | 0.39 | -0.26 | 0.26 | 0.202 |
| **MYBBP1A** | **T1227** | -0.09 | -0.23 | -0.13 | -0.02 | 0.15 | 0.26 | 0.036 |
| **MYBBP1A** | **T1227** | -0.05 | -0.06 | -0.32 | 0.18 | -0.02 | 0.20 | 0.079 |
| **MYBBP1A** | **S1308** | 0.09 | -0.08 | -0.22 | 0.10 | 0.15 | -0.07 | 0.297 |
| **MYBBP1A** | **S1241** | -0.05 | 0.27 | -0.41 | -0.25 | 0.06 | 0.26 | 0.744 |
| **MYBBP1A** | **S1163** | 0.05 | 0.04 | -0.17 | -0.16 | 0.22 | -0.01 | 0.748 |
| **MYBBP1A** | **S775** | 0.01 | 0.03 | -0.09 | 0.03 | 0.22 | -0.25 | 0.904 |
| **MYBBP1A** | **S1186** | -0.18 | 0.15 | 0.00 | 0.27 | 0.05 | -0.37 | 0.984 |
| **MYBBP1A** | **S1163** | 0.15 | -0.02 | -0.12 | 0.08 | -0.01 | -0.09 | 0.922 |
| **MYBBP1A** | **S1186** | -0.10 | 0.03 | 0.09 | -0.14 | 0.15 | -0.05 | 0.83 |
| **MYBBP1A** | **S1267** | 0.14 | -0.01 | -0.06 | -0.13 | -0.01 | 0.06 | 0.546 |
| **MYBBP1A** | **S775** | -0.07 | 0.15 | 0.10 | -0.21 | 0.05 | -0.06 | 0.269 |
| **MYBBP1A** | **S1267** | 0.20 | 0.24 | 0.12 | -0.20 | -0.08 | -0.40 | 0.015 |
| **MYBBP1A** | **S1186** | 0.25 | 0.05 | 0.24 | -0.20 | 0.05 | -0.55 | 0.089 |
| **MYBBP1A** | **S1267** | 0.34 | 0.25 | 0.03 | -0.26 | -0.10 | -0.41 | 0.022 |
| **MYBBP1A** | **S1267** | 0.27 | 0.27 | 0.19 | -0.33 | 0.01 | -0.63 | 0.04 |
| **MYBBP1A** | **S1267** | 0.23 | 0.23 | 0.40 | -0.21 | -0.20 | -0.75 | 0.025 |
| **MYBL2** | **S282** | -0.47 | -0.43 | -0.19 | 0.17 | 0.41 | 0.27 | 0.004 |
| **MYBL2** | **S282** | -0.53 | -0.41 | -0.15 | 0.09 | 0.24 | 0.49 | 0.017 |
| **MYBL2** | **T266** | 0.02 | -0.62 | -0.30 | 0.16 | -0.02 | 0.51 | 0.099 |
| **MYBL2** | **T266** | -0.15 | -0.16 | -0.38 | 0.31 | 0.19 | 0.08 | 0.013 |
| **MYBL2** | **T505** | -0.08 | -0.34 | -0.14 | 0.08 | 0.36 | 0.01 | 0.063 |
| **MYC** | **S** | 0.25 | 0.02 | 0.24 | -0.13 | -0.17 | -0.28 | 0.013 |
| **MYC** | **S77** | 0.31 | 0.12 | 0.36 | -0.25 | -0.05 | -0.77 | 0.053 |
| **MYCBP2** | **S3932** | -0.03 | -0.20 | -0.28 | 0.28 | 0.08 | 0.08 | 0.033 |
| **MYCBP2** | **S2920** | -0.04 | -0.02 | -0.02 | 0.04 | -0.13 | 0.15 | 0.584 |
| **MYEF2** | **S571** | -0.30 | -0.33 | -0.12 | 0.08 | 0.14 | 0.39 | 0.017 |
| **MYEF2** | **S17** | -0.01 | -0.02 | -0.09 | 0.16 | -0.14 | 0.07 | 0.469 |
| **MYEF2** | **T13; S17** | 0.04 | 0.09 | -0.07 | -0.15 | -0.14 | 0.19 | 0.673 |
| **MYEF2** | **S17** | 0.00 | 0.09 | 0.00 | 0.05 | -0.09 | -0.06 | 0.322 |
| **MYH10** | **S1956** | -0.02 | 0.18 | -0.03 | -0.42 | -0.12 | 0.30 | 0.616 |
| **MYH10** | **S1952** | -0.01 | 0.07 | 0.11 | -0.02 | -0.13 | -0.04 | 0.068 |
| **MYH10** | **S559** | -0.13 | 0.17 | 0.36 | -0.19 | 0.03 | -0.35 | 0.173 |
| **MYH14** | **S1989** | 0.02 | -0.12 | 0.12 | 0.01 | 0.01 | -0.06 | 0.772 |
| **MYH14** | **S1969; S1989** | -0.03 | -0.03 | 0.10 | -0.21 | 0.06 | 0.08 | 0.765 |
| **MYH14** | **S45** | 0.17 | -0.10 | -0.01 | -0.13 | -0.11 | 0.16 | 0.714 |
| **MYH14** | **S1461** | 0.18 | -0.12 | 0.11 | -0.09 | 0.01 | -0.12 | 0.269 |
| **MYH14** | **S1969** | -0.08 | 0.12 | 0.25 | -0.03 | -0.04 | -0.26 | 0.161 |
| **MYH14** | **S1980; S1983; S1989** | 0.05 | 0.07 | 0.21 | -0.77 | 0.10 | 0.14 | 0.389 |
| **MYH14** | **S1980; S1983; S1989** | 0.05 | 0.07 | 0.21 | -0.77 | 0.10 | 0.14 | 0.389 |
| **MYH14** | **S1969; S1989** | -0.03 | 0.32 | 0.18 | -0.35 | -0.05 | -0.17 | 0.063 |
| **MYH9** | **T1151** | -0.18 | -0.24 | -0.22 | 0.10 | 0.06 | 0.38 | 0.02 |
| **MYH9** | **S1808** | -0.21 | -0.29 | 0.16 | 0.22 | -0.21 | 0.22 | 0.398 |
| **MYH9** | **S1943** | 0.10 | -0.05 | 0.06 | 0.03 | -0.18 | 0.02 | 0.389 |
| **MYH9** | **S1714** | 0.06 | 0.04 | 0.09 | -0.13 | -0.02 | -0.07 | 0.016 |

| | | | | | | | | |
|---|---|---|---|---|---|---|---|---|
| MYH9 | S1943 | 0.09 | 0.04 | 0.11 | -0.05 | -0.10 | -0.11 | 0.004 |
| MYH9 | S1714 | 0.11 | 0.03 | 0.17 | -0.08 | -0.03 | -0.22 | 0.042 |
| MYH9 | T725 | 0.10 | 0.06 | 0.13 | -0.14 | 0.07 | -0.27 | 0.107 |
| MYH9 | T/S | 0.17 | 0.19 | 0.06 | -0.10 | -0.11 | -0.26 | 0.011 |
| MYH9 | S1943 | 0.23 | 0.25 | 0.17 | -0.16 | 0.02 | -0.72 | 0.088 |
| MYH9 | S1943 | 0.04 | 0.46 | 0.36 | -0.16 | -0.20 | -0.87 | 0.056 |
| MYO18A | T/S | -0.40 | -0.41 | -0.32 | 0.61 | -0.35 | 0.45 | 0.109 |
| MYO18A | S332 | 0.20 | -0.20 | -0.16 | 0.05 | -0.11 | 0.17 | 0.579 |
| MYO18A | S140 | -0.02 | 0.04 | -0.14 | 0.11 | -0.11 | 0.11 | 0.455 |
| MYO18A | S154; T155; S157 | -0.20 | -0.06 | 0.12 | -0.31 | -0.06 | 0.39 | 0.822 |
| MYO18A | S83 | -0.07 | 0.16 | -0.13 | -0.05 | -0.14 | 0.19 | 0.935 |
| MYO18A | S1998; S2002 | 0.03 | -0.05 | 0.04 | 0.04 | -0.09 | 0.03 | 0.77 |
| MYO18A | S2007 | 0.20 | -0.08 | -0.08 | 0.09 | -0.04 | -0.11 | 0.755 |
| MYO18A | S2020 | 0.02 | -0.07 | 0.10 | 0.05 | -0.02 | -0.10 | 0.545 |
| MYO18A | S2020 | 0.08 | 0.07 | 0.14 | -0.09 | -0.09 | -0.13 | 0.002 |
| MYO18A | S1998 | 0.14 | 0.09 | 0.15 | -0.14 | -0.15 | -0.12 | 2E-04 |
| MYO18A | S1640 | 0.06 | 0.08 | 0.24 | -0.13 | -0.09 | -0.22 | 0.016 |
| MYO18A | S2041 | 0.15 | 0.20 | 0.14 | -0.06 | -0.07 | -0.45 | 0.05 |
| MYO19 | S685 | 0.00 | -0.05 | 0.16 | 0.05 | 0.00 | -0.19 | 0.44 |
| MYO1C | S408 | -0.15 | -0.19 | 0.04 | 0.02 | -0.16 | 0.37 | 0.359 |
| MYO5B | S1446 | -0.48 | -0.23 | -0.40 | -0.01 | 0.72 | 0.05 | 0.066 |
| MYO5B | S978 | -0.18 | -0.22 | -0.17 | 0.17 | -0.07 | 0.38 | 0.055 |
| MYO5B | S1290 | 0.11 | 0.01 | 0.07 | -0.19 | 0.21 | -0.27 | 0.386 |
| MYO5B | S1199 | 0.28 | 0.68 | -0.24 | -0.24 | -0.14 | -0.76 | 0.132 |
| MYO5C | S966 | -0.09 | -0.28 | 0.06 | -0.01 | 0.09 | 0.18 | 0.168 |
| MYO9A | S1299 | 0.00 | 0.10 | 0.17 | 0.09 | -0.33 | -0.08 | 0.218 |
| MYO9A | S/Y | -0.30 | 0.05 | 0.58 | -0.57 | -0.17 | 0.13 | 0.395 |
| MYO9B | S2141 | -0.32 | -0.14 | -0.20 | 0.54 | -0.31 | 0.20 | 0.227 |
| MYO9B | S | -0.09 | -0.09 | -0.33 | 0.50 | -0.18 | 0.03 | 0.259 |
| MYO9B | S1405 | -0.11 | -0.02 | -0.30 | 0.34 | -0.20 | 0.19 | 0.238 |
| MYO9B | S | -0.03 | -0.42 | 0.15 | -0.04 | -0.11 | 0.33 | 0.496 |
| MYO9B | S1115; S1122 | 0.12 | 0.00 | -0.07 | -0.22 | -0.02 | 0.16 | 0.756 |
| MYO9B | S1992 | -0.12 | 0.14 | 0.02 | 0.15 | -0.33 | 0.08 | 0.787 |
| MYO9B | S1992; S/T | -0.08 | 0.20 | 0.07 | -0.11 | -0.09 | -0.01 | 0.183 |
| MYO9B | T1271 | 0.02 | 0.29 | -0.11 | -0.06 | -0.24 | 0.03 | 0.332 |
| MYO9B | S1115 | 0.04 | 0.03 | 0.18 | 0.04 | -0.38 | 0.03 | 0.276 |
| MYO9B | T1346 | 0.07 | -0.11 | 0.30 | -0.12 | -0.10 | -0.09 | 0.197 |
| MYO9B | S1354 | 0.15 | 0.12 | 0.02 | 0.17 | -0.08 | -0.47 | 0.3 |
| MYO9B | S1290 | 0.01 | 0.12 | 0.21 | 0.00 | -0.16 | -0.23 | 0.052 |
| MYOF | S174 | -0.17 | -0.47 | -0.24 | -0.01 | -0.17 | 0.73 | 0.181 |
| MYOF | S | -0.27 | 0.18 | -0.38 | -0.05 | -0.20 | 0.52 | 0.433 |
| MYPOP | S202 | 0.03 | -0.12 | -0.01 | 0.12 | -0.05 | 0.03 | 0.365 |
| MYPOP | S378 | 0.07 | 0.32 | -0.04 | 0.28 | -0.58 | -0.24 | 0.339 |
| MYRIP | S622; S626 | 0.16 | -0.88 | -0.21 | -0.08 | 0.36 | 0.32 | 0.203 |
| MYRIP | T307 | -0.19 | 0.01 | -0.23 | 0.32 | -0.22 | 0.20 | 0.256 |
| MYRIP | S557; S561 | -0.11 | -0.12 | -0.10 | 0.02 | 0.06 | 0.22 | 0.029 |
| MYRIP | S280 | -0.01 | 0.01 | 0.14 | -0.25 | -0.01 | 0.10 | 0.415 |
| MYRIP | S | 0.20 | 0.23 | 0.33 | -0.27 | -0.15 | -0.52 | 0.008 |

| | | | | | | | | |
|---|---|---|---|---|---|---|---|---|
| MYSM1 | S340 | 0.06 | -0.16 | 0.12 | 0.08 | -0.19 | 0.06 | 0.84 |
| MYT1 | S91; S94 | -0.12 | -0.30 | -0.15 | 0.27 | -0.09 | 0.28 | 0.062 |
| MYT1 | S333 | 0.08 | -0.18 | -0.12 | 0.13 | -0.05 | 0.12 | 0.229 |
| MYT1 | S/Y/T | -0.01 | -0.10 | 0.00 | -0.18 | 0.22 | 0.04 | 0.62 |
| MYT1 | S692 | 0.09 | -0.04 | 0.22 | -0.02 | -0.25 | -0.03 | 0.15 |
| MYT1 | S610 | 0.16 | 0.18 | 0.09 | -0.24 | -0.06 | -0.19 | 0.007 |
| MYT1 | S697 | 0.30 | 0.13 | 0.21 | -0.29 | -0.03 | -0.46 | 0.026 |
| MYT1 | S697 | 0.41 | -0.03 | 0.33 | 0.07 | -0.04 | -1.30 | 0.227 |
| MZT2A | S152 | -0.10 | -0.18 | -0.06 | -0.26 | 0.35 | 0.16 | 0.345 |
| N4BP1 | T242 | -0.16 | -0.58 | -0.15 | 0.02 | 0.14 | 0.50 | 0.062 |
| N4BP1 | S300 | -0.20 | -0.12 | -0.13 | 0.26 | -0.01 | 0.14 | 0.028 |
| N4BP3 | S176 | -0.21 | -0.12 | -0.29 | 0.01 | 0.27 | 0.24 | 0.017 |
| NAA30 | S152 | 0.02 | -0.06 | -0.11 | 0.10 | 0.08 | -0.06 | 0.219 |
| NAA30 | S55 | 0.05 | 0.11 | -0.28 | -0.21 | -0.18 | 0.38 | 0.882 |
| NAALAD2 | T735; S/T | 0.18 | -0.08 | 0.14 | -0.08 | -0.39 | 0.14 | 0.325 |
| NAB2 | S162 | -0.26 | -0.29 | -0.75 | 0.18 | 0.05 | 0.66 | 0.041 |
| NACA | S2029 | 0.13 | -0.09 | 0.04 | 0.02 | -0.16 | 0.04 | 0.57 |
| NACA | T/S | -0.42 | 0.16 | 0.28 | 0.19 | 0.13 | -0.54 | 0.814 |
| NACA | T/S | 0.04 | 0.15 | 0.12 | -0.08 | -0.09 | -0.17 | 0.007 |
| NACA | T/S | 0.15 | 0.09 | 0.12 | -0.08 | -0.06 | -0.28 | 0.024 |
| NADK2 | S/Y/T | -0.17 | 0.13 | 0.06 | -0.15 | -0.15 | 0.23 | 0.864 |
| NAF1 | S315 | 0.08 | 0.14 | 0.39 | -0.13 | -0.22 | -0.40 | 0.022 |
| NAF1 | S315 | 0.25 | 0.28 | 0.25 | -0.46 | -0.01 | -0.54 | 0.022 |
| NAGK | S76 | -0.22 | 0.14 | 0.34 | -0.30 | -0.04 | -0.02 | 0.341 |
| NAP1L1 | T62 | -0.29 | -0.55 | -0.16 | 0.40 | 0.10 | 0.28 | 0.014 |
| NAP1L1 | S10 | -0.01 | -0.23 | 0.05 | -0.12 | 0.26 | 0.00 | 0.457 |
| NAP1L4 | S125 | -0.12 | -0.06 | 0.04 | 0.04 | 0.00 | 0.09 | 0.179 |
| NAP1L4 | S125 | 0.02 | 0.01 | 0.02 | 0.08 | -0.09 | -0.06 | 0.506 |
| NAPG | T287 | -0.21 | 0.11 | 0.30 | -0.08 | 0.22 | -0.48 | 0.512 |
| NASP | S421 | -0.24 | -0.32 | -0.15 | 0.09 | 0.13 | 0.36 | 0.011 |
| NASP | T477; S480 | -0.19 | 0.11 | -0.08 | 0.20 | -0.08 | 0.00 | 0.459 |
| NASP | S451 | 0.14 | -0.13 | -0.15 | -0.13 | 0.06 | 0.17 | 0.567 |
| NASP | S726 | 0.08 | 0.02 | -0.21 | 0.10 | -0.08 | 0.07 | 0.533 |
| NAV1 | S1253 | 0.12 | -0.05 | 0.02 | -0.21 | -0.13 | 0.20 | 0.601 |
| NBEAL2 | T1867 | -0.06 | -0.21 | -0.01 | 0.05 | 0.44 | -0.34 | 0.56 |
| NBEAL2 | S2739 | -0.41 | 0.03 | 0.12 | 0.22 | -0.42 | 0.30 | 0.685 |
| NBEAL2 | S2739 | -0.11 | 0.35 | 0.25 | -0.12 | -0.08 | -0.44 | 0.105 |
| NBN | S343 | -0.48 | -0.51 | -0.23 | 0.23 | 0.57 | 0.10 | 0.014 |
| NBN | S432 | -0.37 | 0.10 | -0.42 | 0.27 | -0.01 | 0.28 | 0.098 |
| NBN | T599 | -0.04 | -0.07 | -0.11 | 0.08 | 0.02 | 0.09 | 0.009 |
| NBR1 | S656 | -0.17 | 0.14 | -0.14 | -0.08 | 0.06 | 0.15 | 0.441 |
| NCAM2 | S786 | -0.74 | 0.11 | 0.28 | -0.29 | 0.37 | -0.01 | 0.721 |
| NCAPD2 | S585 | -0.05 | -0.53 | -0.05 | 0.19 | 0.25 | 0.06 | 0.093 |
| NCAPD2 | S585 | -0.04 | -0.11 | -0.31 | 0.00 | 0.22 | 0.18 | 0.05 |
| NCAPD2 | S585 | 0.10 | 0.01 | -0.20 | 0.06 | 0.20 | -0.20 | 0.741 |
| NCAPD2 | T1339 | 0.23 | -0.06 | 0.13 | -0.11 | -0.25 | 0.01 | 0.133 |
| NCAPD3 | S1321 | -0.21 | -0.13 | -0.20 | 0.04 | -0.19 | 0.53 | 0.223 |
| NCAPD3 | S530; S531 | -0.20 | 0.00 | 0.29 | 0.19 | -0.09 | -0.28 | 0.654 |

| | | | | | | | | |
|---|---|---|---|---|---|---|---|---|
| NCAPG | T931 | -0.26 | -0.54 | -0.33 | 0.18 | 0.28 | 0.43 | 0.004 |
| NCAPG | S674 | -0.41 | -0.38 | -0.34 | 0.23 | 0.20 | 0.45 | 0.001 |
| NCAPG | S674 | -0.38 | -0.43 | -0.32 | 0.19 | 0.18 | 0.50 | 0.004 |
| NCAPG2 | T605 | -0.14 | -0.28 | -0.38 | 0.15 | 0.24 | 0.27 | 0.004 |
| NCAPH | T49 | -0.07 | -0.59 | -0.14 | 0.21 | -0.14 | 0.50 | 0.139 |
| NCAPH | S408 | -0.16 | -0.17 | -0.13 | 0.12 | 0.02 | 0.27 | 0.017 |
| NCAPH | S201 | 0.15 | 0.18 | -0.18 | -0.02 | -0.02 | -0.15 | 0.4 |
| NCAPH2 | S284 | -0.41 | -0.37 | -0.23 | 0.40 | -0.22 | 0.53 | 0.075 |
| NCAPH2 | S492 | -0.21 | -0.19 | -0.46 | 0.02 | 0.22 | 0.44 | 0.026 |
| NCAPH2 | S466 | -0.31 | -0.15 | -0.39 | -0.09 | 0.43 | 0.32 | 0.045 |
| NCAPH2 | S308 | -0.25 | -0.23 | -0.16 | 0.16 | 0.18 | 0.21 | 2E-04 |
| NCAPH2 | S466 | -0.20 | -0.44 | 0.02 | 0.15 | 0.25 | 0.12 | 0.051 |
| NCAPH2 | S284 | -0.15 | -0.33 | 0.05 | 0.18 | -0.06 | 0.24 | 0.138 |
| NCAPH2 | S | 0.14 | 0.02 | -0.56 | 0.07 | 0.11 | 0.11 | 0.345 |
| NCBP1 | S7 | -0.19 | -0.24 | -0.14 | 0.23 | 0.04 | 0.23 | 0.007 |
| NCBP1 | T/S | -0.20 | -0.27 | 0.17 | -0.12 | 0.21 | 0.13 | 0.364 |
| NCBP1 | T/S | 0.19 | 0.05 | 0.09 | -0.10 | 0.02 | -0.31 | 0.09 |
| NCBP3 | S25 | -0.03 | -0.01 | -0.01 | 0.04 | -0.15 | 0.15 | 0.74 |
| NCBP3 | S399; S/Y | 0.17 | -0.12 | -0.05 | 0.06 | -0.25 | 0.14 | 0.899 |
| NCBP3 | S30 | 0.08 | 0.06 | -0.02 | -0.04 | 0.15 | -0.27 | 0.491 |
| NCBP3 | S500 | 0.16 | 0.07 | -0.06 | -0.02 | -0.05 | -0.12 | 0.151 |
| NCBP3 | S444 | 0.06 | 0.07 | 0.14 | 0.04 | -0.16 | -0.18 | 0.061 |
| NCBP3 | S415 | 0.11 | 0.06 | 0.24 | -0.12 | 0.11 | -0.52 | 0.177 |
| NCBP3 | S25 | 0.07 | 0.06 | 0.25 | 0.17 | 0.02 | -0.78 | 0.343 |
| NCBP3 | S415 | 0.42 | -0.12 | 0.45 | -0.21 | -0.12 | -0.76 | 0.088 |
| NCKAP5 | S459 | 0.04 | 0.40 | 0.04 | -0.36 | 0.16 | -0.45 | 0.172 |
| NCKIPSD | T | -0.04 | -0.05 | -0.08 | 0.13 | -0.04 | 0.06 | 0.105 |
| NCL | S591 | -0.39 | -0.58 | -0.43 | 0.33 | 0.55 | 0.16 | 0.003 |
| NCL | S619 | -0.41 | -0.47 | -0.32 | 0.32 | 0.24 | 0.36 | 2E-04 |
| NCL | T76 | 0.00 | -0.39 | -0.12 | 0.50 | -0.08 | -0.06 | 0.262 |
| NCL | S | -0.04 | -0.30 | -0.19 | 0.61 | -0.14 | -0.15 | 0.341 |
| NCL | S | -0.03 | -0.09 | -0.13 | 0.01 | -0.25 | 0.39 | 0.515 |
| NCL | S458 | -0.02 | 0.00 | -0.17 | 0.17 | 0.15 | -0.18 | 0.426 |
| NCL | S34; S | -0.14 | 0.07 | 0.00 | -0.01 | 0.11 | -0.05 | 0.626 |
| NCL | S145; S153 | 0.10 | 0.03 | -0.11 | 0.00 | -0.01 | -0.02 | 0.814 |
| NCL | S153 | 0.09 | 0.09 | 0.02 | 0.08 | 0.02 | -0.33 | 0.347 |
| NCL | S28; S34; S | 0.16 | 0.04 | 0.01 | -0.19 | -0.14 | 0.09 | 0.191 |
| NCL | S206 | 0.09 | 0.04 | 0.09 | -0.11 | 0.07 | -0.21 | 0.129 |
| NCL | S153 | 0.16 | 0.18 | -0.10 | -0.32 | 0.09 | -0.08 | 0.295 |
| NCL | S145; S153 | 0.12 | 0.10 | 0.09 | -0.08 | 0.01 | -0.28 | 0.063 |
| NCL | S28; S34; S | 0.27 | -0.08 | 0.15 | 0.16 | -0.41 | -0.19 | 0.252 |
| NCL | S67 | 0.26 | 0.20 | -0.07 | -0.18 | -0.04 | -0.23 | 0.069 |
| NCL | S34; S | 0.35 | 0.27 | -0.17 | -0.12 | 0.07 | -0.58 | 0.223 |
| NCL | S67 | 0.19 | 0.10 | 0.22 | 0.00 | -0.23 | -0.38 | 0.033 |
| NCL | S41; S42 | 0.20 | 0.26 | 0.06 | -0.04 | -0.08 | -0.50 | 0.074 |
| NCL | S28; S41; S42 | 0.24 | 0.26 | 0.15 | -0.16 | 0.07 | -0.83 | 0.125 |
| NCL | S28; S | 0.24 | 0.49 | 0.06 | -0.17 | 0.04 | -1.15 | 0.152 |
| NCL | T/S | 0.12 | 0.59 | 0.71 | -0.75 | -0.50 | -1.07 | 0.007 |

| | | | | | | | | |
|---|---|---|---|---|---|---|---|---|
| NCOA1 | S22 | 0.05 | 0.02 | 0.03 | -0.08 | -0.24 | 0.18 | 0.549 |
| NCOA2 | S736 | -0.19 | -0.36 | -0.20 | 0.33 | 0.25 | 0.04 | 0.012 |
| NCOA2 | S29 | 0.00 | -0.23 | -0.02 | -0.33 | 0.29 | 0.20 | 0.534 |
| NCOA2 | S699 | 0.02 | -0.02 | -0.07 | 0.08 | 0.02 | -0.03 | 0.313 |
| NCOA3 | S214 | -0.60 | 0.08 | -0.26 | 0.06 | -0.08 | 0.55 | 0.186 |
| NCOA3 | S214 | -0.45 | 0.08 | -0.18 | -0.68 | 0.13 | 0.69 | 0.619 |
| NCOA5 | S21 | -0.08 | -0.11 | -0.16 | 0.19 | -0.11 | 0.22 | 0.12 |
| NCOA5 | S162 | -0.24 | -0.14 | 0.08 | 0.06 | -0.03 | 0.22 | 0.198 |
| NCOA5 | S96 | -0.16 | -0.24 | 0.11 | 0.09 | 0.20 | -0.06 | 0.253 |
| NCOA5 | S151 | -0.26 | -0.04 | 0.13 | 0.23 | -0.34 | 0.19 | 0.712 |
| NCOA5 | S378; S381 | -0.11 | -0.15 | 0.13 | -0.04 | -0.13 | 0.26 | 0.636 |
| NCOA5 | S126 | -0.06 | 0.04 | 0.08 | -0.13 | 0.10 | -0.04 | 0.623 |
| NCOA5 | S29 | 0.03 | -0.02 | 0.18 | -0.11 | 0.05 | -0.14 | 0.197 |
| NCOA5 | S29; S34 | 0.07 | 0.05 | 0.13 | -0.11 | 0.00 | -0.16 | 0.031 |
| NCOA5 | S381 | 0.03 | 0.06 | 0.17 | -0.15 | -0.13 | 0.00 | 0.048 |
| NCOR1 | S821 | -0.36 | -0.60 | -0.16 | 0.42 | 0.18 | 0.26 | 0.01 |
| NCOR1 | S2120 | -0.21 | -0.34 | -0.27 | 0.31 | 0.26 | 0.10 | 0.003 |
| NCOR1 | S1281 | -0.06 | -0.21 | -0.15 | 0.07 | 0.00 | 0.29 | 0.054 |
| NCOR1 | S224 | -0.08 | -0.13 | -0.17 | -0.15 | 0.39 | 0.05 | 0.23 |
| NCOR1 | S1472 | -0.15 | -0.01 | -0.20 | 0.21 | -0.07 | 0.17 | 0.103 |
| NCOR1 | S1196 | -0.04 | -0.31 | 0.14 | 0.02 | 0.29 | -0.20 | 0.604 |
| NCOR1 | S1980; S1981 | -0.05 | 0.04 | -0.17 | 0.18 | -0.16 | 0.11 | 0.445 |
| NCOR1 | S1196 | -0.13 | -0.09 | 0.06 | -0.05 | 0.09 | 0.10 | 0.267 |
| NCOR1 | S1977 | 0.08 | -0.12 | -0.11 | -0.01 | 0.13 | 0.01 | 0.282 |
| NCOR1 | S992 | 0.19 | -0.07 | -0.02 | 0.01 | 0.09 | -0.25 | 0.538 |
| NCOR1 | S | 0.06 | 0.06 | 0.05 | -0.03 | 0.06 | -0.21 | 0.231 |
| NCOR1 | S | 0.14 | 0.10 | -0.05 | -0.05 | 0.10 | -0.29 | 0.32 |
| NCOR1 | S158; S172 | 0.03 | 0.26 | 0.13 | -0.13 | 0.19 | -0.67 | 0.257 |
| NCOR2 | S2203 | -0.12 | -0.31 | -0.21 | 0.20 | 0.10 | 0.25 | 0.005 |
| NCOR2 | S2194 | -0.14 | -0.13 | -0.24 | 0.24 | -0.10 | 0.29 | 0.071 |
| NCOR2 | S956 | -0.03 | 0.06 | -0.42 | 0.23 | -0.04 | 0.11 | 0.248 |
| NCOR2 | S215 | -0.32 | -0.04 | 0.00 | 0.09 | 0.42 | -0.28 | 0.444 |
| NCOR2 | S1971 | -0.14 | -0.06 | -0.06 | 0.09 | 0.17 | -0.02 | 0.053 |
| NCOR2 | S149; S152 | 0.06 | -0.11 | -0.19 | 0.11 | 0.09 | 0.01 | 0.13 |
| NCOR2 | S215 | -0.27 | -0.05 | -0.05 | 0.14 | 0.63 | -0.74 | 0.763 |
| NCOR2 | S2223 | -0.17 | 0.09 | -0.09 | -0.14 | 0.12 | 0.14 | 0.471 |
| NCOR2 | S/T | 0.28 | -0.83 | 0.31 | -0.41 | 0.13 | 0.19 | 0.918 |
| NCOR2 | S1539 | -0.24 | 0.15 | -0.01 | -0.21 | 0.06 | 0.20 | 0.802 |
| NCOR2 | S2223 | -0.11 | 0.47 | -0.37 | 0.06 | 0.08 | -0.30 | 0.859 |
| NCOR2 | S2393 | 0.28 | -0.09 | -0.12 | -0.03 | -0.11 | 0.02 | 0.658 |
| NCOR2 | S1025 | 0.01 | -0.07 | 0.22 | -0.26 | 0.06 | 0.00 | 0.421 |
| NCOR2 | T553; S554 | 0.11 | -0.22 | 0.42 | -0.38 | 0.02 | -0.08 | 0.319 |
| NDC1 | S406 | -0.11 | -0.23 | -0.20 | 0.18 | 0.04 | 0.25 | 0.009 |
| NDC1 | S619 | -0.06 | 0.11 | -0.37 | -0.21 | 0.38 | 0.04 | 0.466 |
| NDC1 | S394 | 0.07 | 0.14 | 0.03 | -0.20 | 0.22 | -0.35 | 0.326 |
| NDC1 | S406 | 0.20 | 0.12 | 0.05 | -0.08 | 0.01 | -0.38 | 0.09 |
| NDC1 | S406 | 0.32 | 0.06 | 0.30 | -0.29 | 0.00 | -0.60 | 0.053 |
| NDE1 | S282 | 0.10 | 0.13 | 0.04 | -0.55 | 0.13 | 0.03 | 0.364 |

| | | | | | | | | |
|---|---|---|---|---|---|---|---|---|
| NDFIP1 | S/Y/T | 0.08 | 0.02 | -0.08 | -0.04 | 0.16 | -0.17 | 0.858 |
| NDFIP1 | S66 | 0.06 | 0.11 | 0.06 | -0.10 | 0.05 | -0.21 | 0.101 |
| NDUFB4 | S26 | 0.02 | 0.24 | 0.31 | -0.19 | 0.01 | -0.57 | 0.083 |
| NDUFB5 | S182 | -0.25 | -0.15 | -0.07 | -0.08 | 0.19 | 0.29 | 0.073 |
| NDUFV3 | S105 | -0.03 | 0.11 | -0.02 | 0.09 | -0.04 | -0.13 | 0.567 |
| NDUFV3 | S101 | 0.05 | 0.05 | 0.11 | 0.01 | -0.01 | -0.24 | 0.136 |
| NECTIN1 | S/Y | -0.14 | -0.24 | -0.03 | 0.12 | -0.18 | 0.37 | 0.232 |
| NECTIN1 | S508 | -0.15 | -0.02 | 0.09 | 0.00 | -0.09 | 0.15 | 0.696 |
| NECTIN1 | S508 | -0.15 | -0.02 | 0.09 | 0.00 | -0.09 | 0.15 | 0.696 |
| NECTIN1 | S422 | 0.22 | -0.33 | 0.26 | -0.14 | 0.00 | -0.10 | 0.546 |
| NECTIN1 | S422 | 0.15 | 0.15 | 0.15 | -0.12 | -0.35 | -0.05 | 0.024 |
| NECTIN1 | S422 | 0.05 | 0.31 | 0.10 | -0.14 | -0.43 | -0.01 | 0.078 |
| NECTIN1 | S422 | 0.23 | 0.22 | 0.06 | -0.29 | 0.05 | -0.39 | 0.056 |
| NECTIN4 | S426 | -0.03 | 0.04 | 0.00 | -0.02 | -0.03 | 0.04 | 0.93 |
| NECTIN4 | S426 | -0.20 | 0.08 | 0.14 | 0.02 | -0.37 | 0.25 | 0.857 |
| NECTIN4 | S444 | -0.03 | 0.15 | 0.15 | -0.09 | -0.15 | -0.06 | 0.046 |
| NEDD4L | S/T | 0.09 | 0.03 | 0.11 | -0.45 | -0.37 | 0.41 | 0.48 |
| NEIL1 | T/S | -0.22 | -0.08 | -0.28 | 0.17 | 0.11 | 0.22 | 0.006 |
| NEIL2 | S/T | -0.20 | -0.18 | 0.06 | 0.12 | 0.26 | -0.12 | 0.239 |
| NEIL3 | S450 | 0.04 | -0.10 | -0.18 | 0.12 | 0.15 | -0.06 | 0.189 |
| NEK1 | S1126 | -0.29 | 0.03 | -0.01 | 0.18 | 0.07 | -0.02 | 0.215 |
| NEK1 | S664 | -0.01 | -0.01 | 0.17 | -0.01 | 0.05 | -0.22 | 0.349 |
| NEK1 | S418 | 0.12 | -0.17 | 0.20 | -0.31 | 0.11 | -0.02 | 0.498 |
| NEK1 | S664 | 0.20 | 0.15 | -0.04 | 0.02 | -0.20 | -0.18 | 0.099 |
| NEK2 | S | -0.07 | -0.13 | -0.03 | -0.08 | 0.05 | 0.23 | 0.189 |
| NEK4 | S661 | -0.39 | -0.10 | 0.02 | 0.03 | 0.21 | 0.15 | 0.098 |
| NEK4 | S563 | 0.03 | 0.00 | 0.08 | 0.00 | -0.06 | -0.05 | 0.072 |
| NEK4 | S461 | 0.16 | 0.10 | -0.04 | -0.31 | 0.07 | -0.02 | 0.273 |
| NEK9 | S | 0.35 | 0.13 | -0.42 | -0.11 | 0.10 | -0.17 | 0.762 |
| NELFA | S363 | 0.01 | 0.04 | 0.11 | -0.22 | -0.16 | 0.17 | 0.377 |
| NELFE | S | -0.35 | -0.35 | -0.37 | 0.53 | -0.34 | 0.52 | 0.105 |
| NELFE | S181 | -0.24 | -0.09 | -0.16 | 0.20 | -0.14 | 0.33 | 0.118 |
| NEMF | S417 | -0.29 | -0.08 | 0.13 | -0.02 | -0.07 | 0.27 | 0.44 |
| NEMF | S747; S748 | 0.09 | 0.00 | -0.30 | -0.01 | -0.09 | 0.26 | 0.499 |
| NEMF | S798 | 0.04 | 0.07 | -0.22 | 0.09 | 0.16 | -0.18 | 0.699 |
| NEMF | S747; S748 | 0.25 | 0.13 | -0.24 | -0.13 | 0.00 | -0.06 | 0.522 |
| NEMF | S831 | 0.17 | 0.18 | -0.10 | -0.13 | -0.01 | -0.14 | 0.148 |
| NEMF | S417 | 0.03 | 0.14 | 0.19 | -0.12 | -0.10 | -0.18 | 0.008 |
| NES | S768 | -0.19 | -0.18 | 0.01 | 0.39 | -0.25 | 0.11 | 0.353 |
| NES | S680 | -0.13 | 0.22 | 0.39 | -0.37 | -0.10 | -0.13 | 0.106 |
| NES | S680 | -0.13 | 0.22 | 0.39 | -0.37 | -0.10 | -0.13 | 0.106 |
| NET1 | S131 | -0.01 | -0.98 | -0.16 | 0.56 | 0.00 | 0.19 | 0.139 |
| NEU3 | S309 | -0.26 | 0.02 | -0.03 | -0.01 | -0.03 | 0.26 | 0.265 |
| NEURL4 | S502 | -0.03 | -0.10 | -0.09 | -0.06 | 0.15 | 0.11 | 0.102 |
| NF1 | S2515 | -0.74 | -0.44 | -0.45 | 0.14 | -0.02 | 0.89 | 0.042 |
| NF1 | S2597 | -0.09 | -0.14 | -0.36 | 0.68 | 0.05 | -0.44 | 0.428 |
| NF1 | T2565 | 0.05 | 0.01 | 0.03 | 0.06 | -0.20 | 0.03 | 0.464 |
| NF1 | S2543 | 0.20 | 0.27 | -0.32 | 0.25 | -0.22 | -0.32 | 0.594 |

| | | | | | | | | |
|---|---|---|---|---|---|---|---|---|
| **NF1** | **S2188** | -0.04 | -0.03 | 0.27 | 0.21 | -0.31 | -0.18 | 0.425 |
| **NF2** | **S518** | 0.09 | 0.05 | -0.15 | 0.02 | 0.14 | -0.17 | 0.991 |
| **NFAT5** | **S145** | -0.37 | 0.01 | 0.06 | -0.09 | 0.16 | 0.18 | 0.324 |
| **NFAT5** | **S229** | 0.10 | 0.09 | 0.12 | 0.05 | -0.18 | -0.21 | 0.057 |
| **NFAT5** | **S561** | 0.26 | 0.33 | -0.05 | -0.44 | -0.04 | -0.20 | 0.069 |
| **NFATC2IP** | **S204** | -0.03 | -0.11 | -0.20 | 0.17 | -0.03 | 0.17 | 0.059 |
| **NFATC2IP** | **S/T** | 0.22 | -0.15 | -0.36 | -0.13 | 0.15 | 0.18 | 0.447 |
| **NFE2L2** | **S215** | -0.04 | -0.04 | -0.25 | 0.14 | -0.02 | 0.16 | 0.091 |
| **NFE2L2** | **S433** | -0.10 | 0.00 | -0.09 | 0.22 | -0.01 | -0.05 | 0.268 |
| **NFE2L2** | **S215** | -0.20 | 0.11 | -0.08 | 0.13 | -0.07 | 0.08 | 0.402 |
| **NFIA** | **S280** | -0.32 | -0.18 | -0.33 | 0.24 | 0.28 | 0.17 | 9E-04 |
| **NFIA** | **S265** | -0.11 | -0.19 | -0.39 | 0.18 | -0.03 | 0.39 | 0.049 |
| **NFIA** | **S280** | -0.13 | -0.06 | -0.17 | 0.12 | 0.17 | 0.03 | 0.012 |
| **NFIA** | **S300** | 0.01 | -0.03 | -0.26 | 0.16 | -0.21 | 0.26 | 0.385 |
| **NFIA** | **S287** | -0.09 | -0.09 | 0.04 | -0.01 | 0.00 | 0.14 | 0.24 |
| **NFIA** | **S287** | -0.03 | -0.05 | -0.04 | 0.02 | 0.08 | 0.01 | 0.028 |
| **NFIA** | **S300** | -0.04 | -0.04 | -0.01 | 0.20 | -0.46 | 0.24 | 0.923 |
| **NFIA** | **S300** | 0.07 | -0.05 | -0.09 | 0.28 | -0.52 | 0.18 | 0.988 |
| **NFIA** | **S300; S319** | -0.02 | 0.23 | 0.03 | 0.02 | -0.40 | 0.06 | 0.329 |
| **NFIB** | **S328** | 0.05 | -0.23 | -0.30 | 0.08 | 0.20 | 0.13 | 0.057 |
| **NFIB** | **S328** | 0.09 | -0.27 | -0.31 | -0.04 | -0.04 | 0.44 | 0.238 |
| **NFIB** | **S** | 0.05 | -0.08 | -0.12 | 0.01 | 0.07 | 0.06 | 0.162 |
| **NFIB** | **S294; S295** | -0.01 | -0.12 | 0.18 | 0.09 | 0.06 | -0.23 | 0.762 |
| **NFIB** | **S284; S295** | -0.09 | 0.19 | -0.08 | -0.17 | -0.28 | 0.34 | 0.832 |
| **NFIB** | **S312** | 0.17 | 0.06 | 0.07 | -0.12 | 0.07 | -0.30 | 0.125 |
| **NFIB** | **S312** | 0.18 | 0.08 | 0.14 | -0.10 | 0.06 | -0.45 | 0.12 |
| **NFIB** | **S295** | 0.22 | -0.06 | 0.28 | -0.33 | 0.33 | -0.73 | 0.301 |
| **NFIB** | **S295** | 0.18 | 0.18 | 0.09 | -0.23 | 0.39 | -0.99 | 0.347 |
| **NFIC** | **S323** | -0.14 | -0.27 | 0.00 | 0.00 | 0.20 | 0.16 | 0.06 |
| **NFIC** | **S470** | -0.10 | 0.05 | -0.06 | 0.08 | -0.02 | 0.04 | 0.228 |
| **NFIC** | **S194** | 0.05 | -0.08 | -0.02 | -0.03 | 0.08 | 0.01 | 0.514 |
| **NFIC** | **S339** | 0.41 | -0.24 | -0.29 | -0.42 | 0.06 | 0.28 | 0.977 |
| **NFIC** | **S277** | 0.03 | -0.04 | 0.16 | -0.15 | -0.02 | -0.01 | 0.208 |
| **NFIC** | **S343** | 0.12 | 0.02 | 0.05 | -0.20 | 0.05 | -0.06 | 0.178 |
| **NFIC** | **S286; S305** | 0.01 | 0.14 | 0.04 | -0.10 | 0.06 | -0.16 | 0.159 |
| **NFIC** | **S343** | 0.10 | -0.01 | 0.10 | -0.07 | -0.02 | -0.12 | 0.046 |
| **NFIC** | **S323** | 0.10 | 0.02 | 0.08 | 0.02 | -0.12 | -0.11 | 0.057 |
| **NFIC** | **S294; S305** | 0.04 | 0.11 | 0.19 | -0.14 | -0.05 | -0.19 | 0.018 |
| **NFIC** | **S305** | 0.13 | 0.13 | 0.09 | -0.17 | -0.01 | -0.21 | 0.016 |
| **NFIC** | **S294; S302; S305** | 0.13 | 0.09 | 0.16 | -0.28 | -0.03 | -0.12 | 0.023 |
| **NFIC** | **T307** | 0.13 | 0.20 | 0.06 | -0.22 | -0.07 | -0.15 | 0.01 |
| **NFIC** | **S284; T293; S304** | 0.24 | -0.02 | 0.24 | -0.28 | -0.23 | -0.03 | 0.046 |
| **NFIC** | **S323** | 0.11 | 0.19 | 0.20 | -0.20 | -0.05 | -0.33 | 0.013 |
| **NFIC** | **S333** | 0.18 | 0.14 | 0.22 | -0.44 | 0.21 | -0.50 | 0.141 |
| **NFIC** | **S323** | 0.20 | 0.23 | 0.24 | -0.29 | -0.02 | -0.54 | 0.029 |
| **NFIC** | **S333** | 0.21 | 0.25 | 0.34 | -0.30 | -0.01 | -0.77 | 0.051 |
| **NFIC** | **S333** | 0.22 | 0.22 | 0.37 | -0.34 | 0.01 | -0.77 | 0.052 |
| **NFIC** | **S323** | 0.31 | 0.26 | 0.25 | -0.37 | 0.00 | -0.73 | 0.038 |

| | | | | | | | | |
|---|---|---|---|---|---|---|---|---|
| **NFIL3** | **S301** | -0.05 | -0.05 | -0.06 | 0.09 | -0.09 | 0.14 | 0.221 |
| **NFIL3** | **S461** | -0.03 | -0.21 | 0.08 | -0.26 | -0.35 | 0.57 | 0.904 |
| **NFIL3** | **T274; S285; S286** | 0.35 | -0.04 | 0.15 | -0.26 | -0.03 | -0.26 | 0.068 |
| **NFIX** | **S280; S288** | -0.11 | -0.05 | -0.06 | 0.20 | 0.01 | -0.02 | 0.124 |
| **NFIX** | **S280** | -0.08 | 0.02 | -0.12 | 0.09 | -0.06 | 0.13 | 0.198 |
| **NFIX** | **S301** | 0.01 | 0.08 | -0.16 | 0.05 | -0.09 | 0.08 | 0.718 |
| **NFIX** | **S301** | 0.12 | -0.07 | -0.11 | -0.02 | -0.11 | 0.16 | 0.801 |
| **NFIX** | **S265; T/S** | 0.08 | -0.09 | -0.02 | -0.06 | 0.20 | -0.13 | 0.883 |
| **NFIX** | **S280** | -0.01 | 0.00 | 0.02 | -0.07 | 0.00 | 0.05 | 0.757 |
| **NFIX** | **S268** | 0.16 | -0.03 | -0.03 | -0.02 | -0.06 | -0.03 | 0.334 |
| **NFIX** | **S301** | 0.16 | 0.08 | -0.11 | -0.08 | -0.07 | 0.00 | 0.337 |
| **NFKB2** | **S222** | -0.18 | -0.17 | -0.45 | 0.21 | 0.03 | 0.40 | 0.027 |
| **NFKBIL1** | **S150** | -0.01 | 0.03 | -0.02 | -0.04 | 0.06 | -0.02 | 0.969 |
| **NFKBIL1** | **S150** | 0.24 | -0.26 | 0.01 | -0.13 | -0.84 | 0.59 | 0.796 |
| **NFRKB** | **S1175; T/S** | -0.35 | -0.58 | -0.45 | 0.34 | -0.24 | 0.77 | 0.068 |
| **NFRKB** | **T1169; T1170** | 0.07 | -0.61 | -0.81 | 0.37 | -0.49 | 0.78 | 0.219 |
| **NFRKB** | **S228** | -0.12 | -0.17 | -0.10 | 0.23 | -0.07 | 0.17 | 0.066 |
| **NFRKB** | **S209; S210; S222** | -0.14 | -0.21 | 0.11 | 0.15 | -0.11 | 0.16 | 0.321 |
| **NFRKB** | **S351** | 0.20 | -0.10 | -0.17 | -0.14 | 0.20 | -0.05 | 0.854 |
| **NFRKB** | **S351** | 0.18 | 0.07 | -0.08 | -0.07 | -0.03 | -0.08 | 0.18 |
| **NFRKB** | **S1291** | 0.10 | 0.11 | 0.02 | -0.87 | 0.15 | 0.24 | 0.545 |
| **NFX1** | **S326** | 0.16 | 0.06 | 0.47 | -0.18 | 0.05 | -0.90 | 0.14 |
| **NFXL1** | **S331** | -0.59 | -0.15 | 0.19 | 0.25 | -0.31 | 0.38 | 0.396 |
| **NFXL1** | **S** | -0.08 | -0.32 | -0.05 | -0.21 | -0.09 | 0.57 | 0.402 |
| **NFXL1** | **S50** | -0.15 | -0.17 | 0.18 | -0.05 | -0.08 | 0.22 | 0.643 |
| **NFXL1** | **S331** | 0.02 | -0.14 | 0.13 | 0.13 | -0.44 | 0.21 | 0.881 |
| **NFYA** | **S326** | -0.21 | 0.01 | -0.06 | -0.19 | 0.15 | 0.24 | 0.374 |
| **NFYA** | **S326** | 0.02 | -0.01 | -0.13 | 0.09 | 0.00 | 0.02 | 0.217 |
| **NFYA** | **S326** | 0.09 | -0.04 | 0.01 | 0.02 | 0.06 | -0.15 | 0.589 |
| **NFYA** | **S326** | 0.03 | 0.00 | 0.06 | -0.16 | -0.06 | 0.11 | 0.438 |
| **NGDN** | **S** | -0.14 | -0.22 | -0.21 | 0.01 | -0.39 | 0.68 | 0.409 |
| **NGDN** | **S142; S143** | -0.33 | -0.18 | 0.24 | -0.08 | 0.25 | 0.02 | 0.473 |
| **NGDN** | **S214** | 0.01 | 0.18 | -0.05 | -0.03 | 0.01 | -0.14 | 0.297 |
| **NHERF1** | **S280** | 0.25 | 0.37 | -0.04 | -0.54 | -0.15 | -0.05 | 0.084 |
| **NHERF1** | **S302** | 0.18 | 0.31 | 0.14 | -0.04 | -0.44 | -0.31 | 0.02 |
| **NHERF1** | **S290** | 0.31 | 0.30 | 0.11 | -0.14 | -0.28 | -0.48 | 0.01 |
| **NHERF2** | **S254** | -0.12 | 0.01 | 0.09 | 0.08 | -0.12 | 0.05 | 0.908 |
| **NHERF2** | **S254** | -0.06 | 0.06 | 0.11 | 0.06 | -0.16 | -0.02 | 0.402 |
| **NHERF2** | **S/T** | 0.13 | 0.29 | -0.28 | -0.08 | 0.10 | -0.24 | 0.584 |
| **NHS** | **T997** | -0.17 | -0.11 | -0.13 | 0.08 | -0.03 | 0.29 | 0.06 |
| **NHS** | **S/T** | 0.23 | -0.11 | -0.20 | -0.12 | 0.24 | -0.11 | 0.862 |
| **NHS** | **S418** | -0.04 | 0.11 | 0.09 | -0.15 | 0.05 | -0.07 | 0.219 |
| **NHS** | **T332** | -0.03 | 0.38 | -0.07 | -0.02 | -0.13 | -0.21 | 0.246 |
| **NHS** | **S/T** | -0.16 | 0.32 | 0.12 | -0.40 | -0.13 | 0.14 | 0.349 |
| **NHSL1** | **S328** | 0.07 | -0.30 | 0.45 | -0.48 | 0.29 | -0.28 | 0.513 |
| **NIBAN2** | **S696; S** | 0.04 | 0.15 | 0.11 | -0.02 | -0.32 | 0.00 | 0.114 |
| **NIBAN2** | **S** | 0.13 | 0.24 | 0.14 | -0.10 | -0.20 | -0.29 | 0.004 |
| **NIBAN2** | **S** | 0.31 | 0.24 | 0.06 | -0.31 | -0.28 | -0.14 | 0.008 |

| | | | | | | | | |
|---|---|---|---|---|---|---|---|---|
| NIFK | T234; T238 | -0.17 | 0.02 | 0.07 | -0.06 | -0.11 | 0.21 | 0.772 |
| NIFK | T238 | 0.13 | 0.13 | 0.01 | -0.37 | 0.23 | -0.23 | 0.311 |
| NIFK | S218 | 0.14 | 0.04 | 0.18 | -0.15 | -0.22 | -0.05 | 0.016 |
| NIFK | T223 | 0.31 | 0.24 | 0.32 | -0.37 | -0.20 | -0.54 | 0.003 |
| NIFK | T223 | 0.33 | 0.42 | 0.32 | -0.56 | -0.01 | -1.03 | 0.04 |
| NIFK | T223 | 0.40 | 0.49 | 0.49 | -0.73 | -0.13 | -1.49 | 0.035 |
| NIPAL2 | S360 | 0.28 | 0.29 | 0.10 | -0.96 | -0.55 | 0.37 | 0.205 |
| NIPBL | S2509; S2513; S | -0.24 | -0.38 | -0.31 | -0.06 | 0.06 | 0.66 | 0.082 |
| NIPBL | S1089; S1090; S1096 | -0.52 | -0.21 | -0.07 | -0.05 | 0.18 | 0.47 | 0.08 |
| NIPBL | S1089; S1090; S1096 | -0.24 | -0.03 | -0.07 | 0.00 | 0.06 | 0.24 | 0.089 |
| NIPBL | S2513 | -0.20 | -0.08 | -0.05 | 0.13 | -0.30 | 0.40 | 0.418 |
| NIPBL | S2511; S2513 | -0.10 | -0.08 | -0.10 | 0.08 | -0.14 | 0.29 | 0.238 |
| NIPBL | S350 | -0.21 | 0.05 | -0.11 | 0.10 | -0.06 | 0.19 | 0.192 |
| NIPBL | S1150; S1152; S1160 | -0.12 | -0.07 | -0.03 | 0.05 | 0.00 | 0.16 | 0.062 |
| NIPBL | S1152; S1154; S1160 | 0.08 | -0.18 | -0.10 | 0.04 | -0.04 | 0.17 | 0.27 |
| NIPBL | T746 | -0.05 | -0.06 | -0.06 | 0.07 | -0.06 | 0.15 | 0.153 |
| NIPBL | T599 | -0.20 | -0.10 | 0.17 | -0.07 | -0.13 | 0.28 | 0.695 |
| NIPBL | S318 | 0.05 | -0.04 | -0.10 | 0.01 | -0.02 | 0.10 | 0.327 |
| NIPBL | T735 | -0.01 | -0.04 | 0.08 | -0.03 | -0.06 | 0.05 | 0.647 |
| NIPBL | S2513 | -0.26 | 0.22 | 0.04 | -0.20 | -0.23 | 0.32 | 0.894 |
| NIPBL | S280 | -0.13 | 0.28 | -0.14 | 0.15 | 0.10 | -0.34 | 0.878 |
| NIPBL | S1154; S1160 | 0.01 | 0.09 | 0.01 | 0.02 | 0.09 | -0.25 | 0.457 |
| NIPBL | S318 | 0.08 | 0.10 | -0.05 | -0.01 | -0.09 | -0.04 | 0.176 |
| NIPBL | S280 | 0.04 | 0.04 | 0.05 | 0.04 | -0.28 | 0.08 | 0.447 |
| NIPBL | S2509; S2511; S2513 | 0.21 | 0.11 | -0.14 | 0.06 | -0.41 | 0.09 | 0.479 |
| NIPBL | S2513 | 0.04 | 0.06 | 0.13 | -0.13 | 0.01 | -0.12 | 0.04 |
| NIPBL | S2509; S2513 | 0.10 | 0.10 | 0.07 | -0.05 | -0.06 | -0.19 | 0.013 |
| NIPBL | S2509; S2511; S2521 | 0.27 | 0.15 | -0.15 | 0.03 | -0.36 | -0.02 | 0.311 |
| NIPBL | S | -0.02 | 0.22 | 0.09 | -0.06 | -0.08 | -0.19 | 0.063 |
| NIPBL | S2513; S2515; S | 0.29 | -0.03 | 0.08 | -0.20 | -0.06 | -0.13 | 0.07 |
| NIPBL | S2513; S2515 | -0.04 | 0.18 | 0.26 | -0.20 | -0.13 | -0.12 | 0.038 |
| NIPBL | S1150; S1160 | 0.14 | 0.08 | 0.22 | -0.18 | -0.07 | -0.26 | 0.01 |
| NIPBL | S274; S280 | 0.24 | 0.07 | 0.26 | -0.53 | 0.01 | -0.19 | 0.064 |
| NIPBL | S2513 | 0.20 | 0.24 | 0.20 | -0.42 | -0.30 | -0.06 | 0.011 |
| NIPBL | S2511; S2513 | 0.33 | 0.12 | 0.18 | -0.20 | -0.03 | -0.57 | 0.052 |
| NIPBL | S2482; S2485; S2493 | 0.28 | 0.24 | 0.18 | -0.19 | -0.17 | -0.52 | 0.011 |
| NIPBL | S1096; S/Y | 0.11 | 0.34 | 0.33 | -0.39 | 0.07 | -0.78 | 0.072 |
| NIPBL | S1096; S/Y | 0.11 | 0.34 | 0.33 | -0.39 | 0.07 | -0.78 | 0.072 |
| NIPBL | S1096 | 0.24 | 0.59 | 0.41 | -0.22 | -0.54 | -1.18 | 0.025 |
| NIPBL | S/Y | 0.58 | 0.57 | 0.09 | -0.21 | -0.76 | -1.01 | 0.02 |
| NKAP | S149 | 0.27 | 0.16 | 0.16 | -0.12 | -0.02 | -0.63 | 0.078 |
| NKAP | S157 | 0.26 | 0.25 | 0.15 | -0.20 | -0.25 | -0.34 | 7E-04 |
| NKAPD1 | S80 | -0.05 | 0.10 | 0.03 | 0.04 | 0.15 | -0.30 | 0.663 |
| NKAPD1 | S125; T131 | -0.06 | 0.15 | 0.05 | -0.25 | 0.08 | 0.00 | 0.423 |
| NKAPD1 | S188 | 0.02 | 0.04 | 0.37 | -0.43 | -0.02 | -0.10 | 0.125 |
| NKRF | S/T | 0.13 | -0.12 | 0.04 | -0.02 | -0.46 | 0.32 | 0.791 |
| NKTR | S/T | -0.32 | -0.06 | -0.07 | 0.02 | -0.02 | 0.37 | 0.14 |
| NKTR | S390 | -0.07 | -0.05 | -0.13 | 0.35 | -0.35 | 0.15 | 0.556 |

| | | | | | | | | |
|---|---|---|---|---|---|---|---|---|
| **NKTR** | **S866** | -0.01 | -0.05 | -0.08 | 0.20 | -0.03 | -0.04 | 0.326 |
| **NKTR** | **S821** | 0.11 | 0.21 | -0.31 | 0.35 | -0.06 | -0.48 | 0.833 |
| **NKTR** | **S/T** | -0.11 | -0.12 | 0.27 | -0.34 | 0.31 | -0.13 | 0.791 |
| **NKTR** | **S379** | 0.10 | 0.01 | 0.01 | 0.13 | -0.10 | -0.17 | 0.415 |
| **NKTR** | **T361** | 0.14 | 0.02 | 0.05 | -0.28 | 0.06 | -0.04 | 0.219 |
| **NKTR** | **S613** | 0.12 | 0.04 | 0.13 | 0.08 | -0.27 | -0.13 | 0.131 |
| **NKTR** | **S358; S359; T361** | -0.01 | 0.23 | 0.08 | 0.10 | 0.08 | -0.62 | 0.365 |
| **NKTR** | **S463** | 0.19 | 0.13 | 0.06 | -0.14 | 0.00 | -0.32 | 0.047 |
| **NKTR** | **T1155** | 0.23 | 0.29 | -0.06 | -0.20 | 0.01 | -0.38 | 0.093 |
| **NKTR** | **S671** | 0.20 | 0.19 | 0.11 | -0.24 | -0.15 | -0.18 | 8E-04 |
| **NKTR** | **S1335** | 0.22 | -0.11 | 0.62 | -0.94 | -0.02 | -0.21 | 0.149 |
| **NKX2-2** | **S199** | -0.18 | 0.30 | 0.11 | -0.10 | -0.17 | -0.02 | 0.291 |
| **NKX2-2** | **S199** | 0.15 | 0.24 | -0.09 | -0.17 | 0.06 | -0.25 | 0.182 |
| **NME3** | **T111** | -0.04 | 0.01 | -0.48 | -0.01 | -0.05 | 0.43 | 0.247 |
| **NMNAT1** | **S117** | 0.07 | 0.32 | -0.27 | 0.08 | -0.14 | -0.13 | 0.614 |
| **NMNAT1** | **S117** | 0.14 | 0.22 | 0.08 | 0.08 | -0.17 | -0.46 | 0.107 |
| **NMNAT1** | **S/T** | 0.08 | 0.30 | 0.10 | -0.10 | -0.04 | -0.47 | 0.076 |
| **NMT1** | **S47** | 0.08 | -0.24 | 0.01 | 0.00 | -0.18 | 0.27 | 0.647 |
| **NMT1** | **Y327; T332** | 0.08 | 0.08 | 0.08 | -0.08 | 0.04 | -0.23 | 0.095 |
| **NMT1** | **S47** | 0.25 | 0.15 | -0.09 | -0.14 | 0.09 | -0.33 | 0.226 |
| **NMT2** | **S38** | 0.16 | 0.16 | -0.01 | 0.07 | -0.02 | -0.43 | 0.236 |
| **NOA1** | **S419** | 0.01 | 0.11 | -0.16 | -0.08 | -0.29 | 0.33 | 0.995 |
| **NOB1** | **S201** | -0.55 | -0.03 | -0.17 | 0.20 | 0.34 | 0.04 | 0.068 |
| **NOC2L** | **S672; S673** | -0.04 | -0.02 | -0.08 | 0.05 | -0.02 | 0.10 | 0.093 |
| **NOC2L** | **S49; S56** | 0.15 | -0.09 | -0.18 | 0.15 | -0.09 | 0.03 | 0.595 |
| **NOC2L** | **S672; S673** | -0.01 | 0.10 | -0.19 | 0.37 | -0.21 | -0.14 | 0.845 |
| **NOC2L** | **S49** | 0.12 | -0.04 | -0.15 | 0.16 | -0.07 | -0.04 | 0.741 |
| **NOC2L** | **S49** | 0.14 | -0.06 | -0.13 | 0.03 | 0.05 | -0.04 | 0.749 |
| **NOC2L** | **S592** | 0.09 | -0.01 | -0.04 | 0.10 | -0.03 | -0.11 | 0.693 |
| **NOC2L** | **S49; S** | 0.08 | 0.02 | -0.03 | 0.01 | 0.00 | -0.08 | 0.347 |
| **NOC2L** | **S672** | -0.02 | 0.15 | 0.05 | -0.17 | 0.07 | -0.11 | 0.215 |
| **NOC2L** | **S672** | 0.28 | 0.29 | 0.11 | -0.17 | -0.09 | -0.62 | 0.042 |
| **NOC3L** | **S787** | 0.10 | 0.26 | 0.21 | -0.07 | -0.10 | -0.52 | 0.052 |
| **NOL12** | **T124** | -0.14 | -0.12 | -0.03 | 0.09 | 0.20 | -0.03 | 0.073 |
| **NOL12** | **S139; S140** | 0.23 | 0.26 | 0.20 | -0.23 | -0.10 | -0.54 | 0.017 |
| **NOL12** | **S134** | 0.40 | 0.33 | 0.36 | -0.53 | -0.09 | -0.97 | 0.025 |
| **NOL3** | **S/T** | -0.32 | -0.15 | -0.02 | -0.13 | 0.23 | 0.30 | 0.136 |
| **NOL3** | **S147** | -0.32 | -0.15 | -0.02 | -0.13 | 0.23 | 0.30 | 0.136 |
| **NOL6** | **S811** | -0.48 | 0.46 | -0.26 | -0.25 | 0.13 | 0.19 | 0.74 |
| **NOL6** | **S283** | -0.03 | -0.10 | -0.02 | -0.03 | -0.15 | 0.28 | 0.55 |
| **NOL8** | **S365** | -0.19 | -0.23 | -0.15 | 0.15 | -0.07 | 0.39 | 0.06 |
| **NOL8** | **S378; T381** | 0.18 | -0.15 | -0.62 | 0.04 | 0.03 | 0.34 | 0.265 |
| **NOL8** | **S378; T381** | -0.12 | -0.07 | -0.10 | 0.04 | -0.05 | 0.27 | 0.125 |
| **NOL8** | **S660** | 0.02 | -0.11 | -0.07 | 0.13 | -0.21 | 0.20 | 0.521 |
| **NOL8** | **S378; T381** | -0.04 | -0.03 | -0.04 | 0.05 | -0.01 | 0.06 | 0.028 |
| **NOL8** | **S612** | -0.07 | 0.00 | -0.04 | -0.33 | -0.09 | 0.43 | 0.868 |
| **NOL8** | **S612** | -0.07 | 0.00 | -0.04 | -0.33 | -0.09 | 0.43 | 0.868 |
| **NOL8** | **S329** | -0.10 | -0.01 | 0.14 | -0.16 | 0.09 | 0.02 | 0.8 |

| | | | | | | | | |
|---|---|---|---|---|---|---|---|---|
| **NOL8** | **T/S** | -0.19 | 0.38 | -0.21 | 0.18 | -0.34 | 0.05 | 0.915 |
| **NOL8** | **S378; T381** | 0.04 | 0.02 | 0.08 | -0.08 | 0.08 | -0.16 | 0.256 |
| **NOL8** | **S1099** | 0.17 | -0.09 | 0.06 | 0.00 | -0.31 | 0.12 | 0.506 |
| **NOL8** | **S890** | 0.07 | 0.16 | -0.04 | -0.09 | -0.13 | 0.00 | 0.116 |
| **NOL8** | **S378** | 0.09 | 0.05 | 0.06 | 0.03 | -0.09 | -0.15 | 0.06 |
| **NOL8** | **S378** | 0.18 | 0.03 | 0.02 | -0.02 | -0.10 | -0.13 | 0.067 |
| **NOL8** | **T888; S890** | 0.04 | 0.05 | 0.15 | -0.15 | -0.01 | -0.10 | 0.031 |
| **NOL8** | **S/T** | 0.16 | 0.09 | 0.03 | -0.08 | -0.07 | -0.16 | 0.016 |
| **NOL8** | **S660** | 0.29 | 0.02 | 0.00 | -0.10 | -0.09 | -0.18 | 0.089 |
| **NOL8** | **S** | 0.09 | 0.04 | 0.20 | -0.17 | 0.06 | -0.28 | 0.094 |
| **NOL8** | **S268** | 0.22 | 0.12 | 0.10 | -0.21 | 0.01 | -0.30 | 0.036 |
| **NOL8** | **T/S** | 0.27 | -0.06 | 0.21 | -0.24 | -0.32 | 0.04 | 0.102 |
| **NOL8** | **T888; S890** | 0.13 | 0.07 | 0.26 | -0.42 | -0.01 | -0.12 | 0.067 |
| **NOL8** | **S** | 0.23 | 0.38 | 0.20 | -0.30 | 0.06 | -0.96 | 0.093 |
| **NOL8** | **S890** | 0.33 | 0.34 | 0.38 | -0.31 | -0.10 | -1.20 | 0.058 |
| **NOLC1** | **T349; T350** | -0.22 | -0.19 | -0.32 | 0.27 | 0.07 | 0.28 | 0.005 |
| **NOLC1** | **S563** | 0.04 | 0.00 | -0.18 | 0.01 | -0.05 | 0.16 | 0.385 |
| **NOLC1** | **S698** | 0.00 | -0.04 | -0.02 | -0.08 | -0.08 | 0.19 | 0.749 |
| **NOLC1** | **S643** | 0.00 | -0.04 | 0.04 | -0.03 | 0.14 | -0.13 | 0.934 |
| **NOLC1** | **S508** | 0.21 | -0.33 | 0.16 | -0.21 | 0.02 | 0.08 | 0.814 |
| **NOLC1** | **S362; S363; S365** | -0.05 | 0.14 | -0.05 | 0.22 | 0.00 | -0.32 | 0.784 |
| **NOLC1** | **S538** | 0.10 | 0.04 | 0.04 | -0.03 | -0.10 | -0.06 | 0.015 |
| **NOLC1** | **S83** | 0.21 | 0.03 | 0.01 | -0.11 | 0.02 | -0.20 | 0.114 |
| **NOLC1** | **S/T** | 0.24 | 0.12 | -0.08 | -0.08 | -0.03 | -0.22 | 0.132 |
| **NOLC1** | **S83** | 0.28 | -0.03 | 0.06 | -0.18 | 0.02 | -0.21 | 0.127 |
| **NOLC1** | **S397** | 0.36 | 0.07 | -0.14 | -0.45 | -0.13 | 0.16 | 0.362 |
| **NOLC1** | **S538** | 0.31 | 0.17 | -0.01 | -0.13 | 0.01 | -0.47 | 0.109 |
| **NOLC1** | **S623** | 0.24 | 0.15 | 0.22 | -0.02 | -0.15 | -0.62 | 0.065 |
| **NOLC1** | **S264** | 0.38 | 0.31 | 0.22 | -0.67 | 0.01 | -0.59 | 0.031 |
| **NOLC1** | **S87; S90; S91** | 0.36 | 0.41 | 0.32 | -0.46 | -0.27 | -0.78 | 0.005 |
| **NOM1** | **S317; S320; S321** | -0.09 | -0.24 | -0.20 | 0.07 | 0.14 | 0.25 | 0.009 |
| **NOM1** | **S317; S320; S321** | 0.01 | -0.30 | -0.14 | 0.00 | 0.07 | 0.29 | 0.101 |
| **NOM1** | **S334** | 0.05 | 0.02 | -0.10 | 0.23 | -0.09 | -0.15 | 0.957 |
| **NOM1** | **S320; S321** | 0.31 | 0.32 | 0.24 | -0.31 | 0.09 | -1.10 | 0.106 |
| **NOM1** | **S320; S321** | 0.41 | 0.36 | 0.26 | -0.34 | -0.04 | -1.21 | 0.07 |
| **NONO** | **S262** | -0.10 | -0.28 | -0.23 | 0.24 | 0.24 | 0.04 | 0.012 |
| **NONO** | **T428** | -0.78 | 0.61 | 0.25 | 0.40 | -0.65 | -0.44 | 0.647 |
| **NONO** | **T450** | 0.16 | 0.00 | 0.40 | -1.16 | 0.27 | -0.14 | 0.293 |
| **NOP14** | **S96** | 0.08 | 0.08 | 0.24 | -0.11 | -0.07 | -0.29 | 0.028 |
| **NOP14** | **S96** | 0.21 | 0.18 | 0.20 | -0.31 | 0.08 | -0.53 | 0.066 |
| **NOP2** | **S812** | 0.02 | -0.40 | -0.54 | 0.10 | 0.41 | 0.19 | 0.049 |
| **NOP2** | **S58** | -0.16 | -0.31 | 0.03 | 0.06 | 0.33 | -0.04 | 0.152 |
| **NOP2** | **S786** | -0.02 | -0.13 | -0.04 | 0.16 | -0.15 | 0.15 | 0.354 |
| **NOP2** | **S181; T185; T195** | -0.03 | -0.09 | -0.03 | 0.03 | -0.04 | 0.15 | 0.19 |
| **NOP2** | **S181; T185; T195** | 0.02 | -0.03 | -0.02 | 0.04 | -0.08 | 0.05 | 0.752 |
| **NOP2** | **S181; T185** | 0.13 | -0.14 | 0.00 | 0.13 | -0.12 | -0.02 | 0.977 |
| **NOP2** | **S786** | 0.05 | 0.02 | -0.04 | -0.03 | -0.13 | 0.11 | 0.778 |
| **NOP2** | **T185** | 0.01 | 0.04 | -0.02 | 0.05 | -0.05 | -0.04 | 0.529 |

| | | | | | | | | |
|---|---|---|---|---|---|---|---|---|
| NOP2 | S181; T185 | -0.02 | -0.15 | 0.24 | -0.08 | -0.08 | 0.07 | 0.694 |
| NOP2 | S666 | -0.08 | 0.04 | 0.01 | -0.56 | -0.23 | 0.58 | 0.879 |
| NOP2 | T185; T195 | 0.06 | 0.02 | 0.02 | -0.03 | -0.03 | -0.04 | 0.011 |
| NOP2 | S181 | 0.11 | 0.07 | 0.00 | -0.03 | -0.05 | -0.11 | 0.042 |
| NOP2 | S732 | 0.07 | 0.17 | 0.07 | -0.10 | -0.14 | -0.10 | 0.004 |
| NOP2 | S732 | 0.15 | 0.04 | 0.15 | -0.03 | -0.21 | -0.13 | 0.022 |
| NOP2 | S181 | 0.16 | -0.01 | 0.28 | -0.08 | -0.22 | -0.20 | 0.028 |
| NOP2 | S67 | 0.42 | 0.38 | 0.45 | -0.66 | 0.07 | -1.70 | 0.082 |
| NOP56 | S591 | -0.77 | -0.80 | -0.73 | 0.59 | 0.75 | 0.06 | 0.004 |
| NOP56 | T538 | -0.30 | -0.21 | -0.36 | 0.21 | 0.23 | 0.28 | 3E-04 |
| NOP56 | S563 | -0.18 | -0.26 | -0.43 | 0.20 | 0.43 | 0.05 | 0.017 |
| NOP56 | S/T | 0.06 | 0.05 | -0.18 | -0.13 | 0.08 | 0.10 | 0.748 |
| NOP56 | S519; S520 | 0.04 | 0.02 | -0.06 | 0.12 | -0.06 | -0.07 | 0.973 |
| NOP56 | S520 | 0.07 | 0.05 | -0.02 | 0.01 | 0.06 | -0.19 | 0.421 |
| NOP56 | S520 | 0.05 | 0.09 | 0.01 | -0.01 | 0.01 | -0.17 | 0.17 |
| NOP56 | S570 | 0.12 | 0.03 | 0.02 | -0.02 | -0.01 | -0.15 | 0.107 |
| NOP56 | S406 | 0.15 | 0.06 | -0.02 | 0.11 | -0.08 | -0.26 | 0.316 |
| NOP56 | S/T | 0.05 | -0.16 | 0.26 | 0.18 | 0.10 | -0.59 | 0.608 |
| NOP56 | S569; S570 | 0.15 | 0.05 | 0.02 | -0.09 | -0.02 | -0.12 | 0.031 |
| NOP56 | S/T | 0.08 | 0.09 | 0.03 | -0.12 | 0.27 | -0.44 | 0.482 |
| NOP56 | T468 | 0.18 | 0.23 | -0.03 | -0.30 | 0.05 | -0.21 | 0.099 |
| NOP56 | S/T | 0.20 | 0.41 | -0.09 | -0.70 | -0.02 | -0.02 | 0.194 |
| NOP56 | S569 | 0.33 | 0.28 | 0.31 | -0.40 | 0.08 | -1.09 | 0.083 |
| NOP58 | S502 | -0.06 | -0.14 | -0.18 | 0.10 | 0.05 | 0.20 | 0.013 |
| NOP58 | S502; S514 | 0.05 | -0.09 | -0.07 | 0.10 | -0.12 | 0.12 | 0.477 |
| NOP58 | S514 | 0.07 | 0.04 | -0.13 | 0.02 | 0.01 | -0.01 | 0.815 |
| NOP58 | S502; S514 | 0.09 | -0.03 | -0.06 | -0.02 | -0.07 | 0.08 | 0.959 |
| NOP58 | S502 | -0.01 | 0.12 | -0.10 | -0.04 | 0.16 | -0.14 | 0.923 |
| NOP58 | S502 | 0.08 | 0.05 | -0.06 | -0.02 | 0.15 | -0.23 | 0.667 |
| NOP58 | S502; S514 | 0.12 | 0.17 | 0.09 | -0.10 | -0.06 | -0.25 | 0.013 |
| NOP58 | S502; S514 | 0.23 | 0.18 | 0.11 | -0.15 | -0.03 | -0.46 | 0.042 |
| NOP58 | S514 | 0.20 | 0.16 | 0.17 | -0.07 | 0.06 | -0.70 | 0.152 |
| NOPCHAP1 | S178 | 0.33 | -0.09 | -0.01 | -0.25 | 0.01 | -0.06 | 0.31 |
| NOPCHAP1 | S15 | 0.32 | 0.03 | -0.05 | -0.25 | -0.07 | -0.05 | 0.162 |
| NOS1AP | T/S | -0.41 | 0.03 | 0.05 | -0.51 | 0.23 | 0.39 | 0.667 |
| NOS1AP | S266 | 0.15 | -0.11 | 0.08 | 0.00 | -0.14 | 0.01 | 0.404 |
| NOSIP | S138 | -0.42 | -0.50 | 0.09 | -0.03 | 0.27 | 0.37 | 0.097 |
| NOVA1 | S/T | 0.21 | -0.07 | 0.19 | -0.17 | -0.03 | -0.19 | 0.081 |
| NPAT | T/S | 0.34 | 0.11 | 0.19 | -0.45 | 0.07 | -0.44 | 0.057 |
| NPDC1 | S275; S276 | 0.02 | -0.15 | -0.16 | 0.01 | -0.17 | 0.37 | 0.383 |
| NPDC1 | S229 | -0.05 | -0.03 | 0.00 | 0.05 | -0.15 | 0.16 | 0.644 |
| NPDC1 | S275; S276 | 0.15 | 0.07 | 0.00 | -0.34 | 0.03 | 0.05 | 0.298 |
| NPDC1 | T/S/Y | 0.05 | 0.11 | 0.21 | -0.11 | -0.16 | -0.13 | 0.007 |
| NPDC1 | S276 | -0.06 | 0.00 | 0.38 | -0.45 | -0.09 | 0.09 | 0.279 |
| NPHP4 | T/S | 0.14 | 0.08 | -0.18 | 0.16 | -0.32 | 0.06 | 0.792 |
| NPM1 | S125 | -0.11 | -0.09 | -0.26 | 0.22 | -0.02 | 0.19 | 0.038 |
| NPM1 | S125 | -0.08 | -0.05 | -0.21 | 0.15 | 0.03 | 0.12 | 0.029 |
| NPM1 | S70 | -0.10 | -0.10 | -0.10 | 0.22 | -0.11 | 0.15 | 0.131 |

| | | | | | | | | |
|---|---|---|---|---|---|---|---|---|
| NPM1 | S70 | 0.01 | 0.05 | -0.14 | 0.03 | 0.05 | 0.00 | 0.453 |
| NPM1 | S70 | 0.03 | 0.00 | -0.08 | -0.02 | 0.20 | -0.16 | 0.838 |
| NPM1 | S70 | 0.08 | 0.22 | -0.22 | 0.05 | -0.09 | -0.09 | 0.62 |
| NPM1 | S242 | 0.22 | 0.10 | -0.05 | -0.20 | -0.08 | -0.02 | 0.112 |
| NQO1 | S82 | -0.39 | -0.45 | -0.54 | 0.40 | 0.26 | 0.37 | 2E-04 |
| NQO1 | S13 | -0.14 | -0.02 | -0.36 | 0.35 | -0.26 | 0.29 | 0.24 |
| NR2C1 | S215 | -1.39 | -0.20 | 0.00 | 0.07 | 0.32 | 0.54 | 0.137 |
| NR2C1 | S215 | -0.08 | -0.39 | 0.03 | 0.25 | 0.05 | 0.07 | 0.128 |
| NR2C2 | S68 | -0.34 | -0.46 | -0.25 | 0.19 | 0.39 | 0.26 | 0.002 |
| NR2F1 | T51 | -0.25 | 0.04 | -0.26 | 0.25 | -0.14 | 0.26 | 0.162 |
| NR2F1 | S77 | -0.18 | 0.32 | -0.24 | 0.10 | -0.56 | 0.35 | 0.992 |
| NR2F1 | T51; T63; S/T | 0.28 | 0.07 | -0.22 | -0.49 | 0.02 | 0.21 | 0.643 |
| NR2F1 | T63 | 0.12 | 0.19 | 0.03 | -0.09 | -0.42 | 0.10 | 0.194 |
| NR2F1 | T51; T63 | -0.15 | 0.26 | 0.27 | -0.62 | -0.04 | 0.11 | 0.297 |
| NR2F1 | T63 | 0.04 | 0.38 | 0.11 | 0.22 | -0.54 | -0.43 | 0.177 |
| NR2F1 | T51; T54 | 0.23 | 0.43 | 0.13 | -0.44 | -0.43 | -0.14 | 0.01 |
| NR2F1 | T51; T54 | 0.41 | 0.14 | 0.52 | -0.50 | -0.30 | -0.70 | 0.006 |
| NR2F6 | S34; S40 | -0.13 | -0.04 | -0.14 | 0.20 | -0.12 | 0.19 | 0.156 |
| NR2F6 | S34 | -0.10 | 0.14 | -0.25 | 0.06 | -0.12 | 0.21 | 0.458 |
| NR2F6 | S34 | 0.02 | -0.04 | -0.09 | 0.19 | -0.04 | -0.06 | 0.464 |
| NR3C1 | T505 | -0.80 | 0.06 | -0.68 | 0.51 | 0.48 | -0.08 | 0.078 |
| NRBP1 | T433 | -0.23 | -0.11 | -0.15 | 0.03 | -0.14 | 0.47 | 0.202 |
| NRDC | S94 | -0.03 | 0.27 | 0.25 | -0.03 | -0.32 | -0.23 | 0.053 |
| NRIP1 | S807 | -0.02 | -0.45 | 0.11 | -0.25 | 0.19 | 0.28 | 0.463 |
| NRIP1 | T/S | 0.14 | -0.32 | 0.26 | -0.43 | 0.45 | -0.35 | 0.701 |
| NRK | S1084 | -0.20 | -0.28 | -0.10 | 0.25 | 0.13 | 0.13 | 0.006 |
| NRK | S1084 | -0.20 | -0.28 | -0.10 | 0.25 | 0.13 | 0.13 | 0.006 |
| NRK | S365 | -0.30 | -0.10 | -0.05 | 0.11 | 0.18 | 0.11 | 0.025 |
| NRK | S718 | -0.09 | 0.16 | -0.22 | 0.01 | 0.07 | 0.03 | 0.488 |
| NRK | S1084 | -0.08 | 0.06 | 0.16 | -0.15 | 0.21 | -0.25 | 0.518 |
| NRK | S788 | 0.04 | 0.17 | -0.07 | 0.22 | -0.13 | -0.30 | 0.516 |
| NRK | S1031; S1034 | 0.25 | 0.02 | 0.09 | -0.15 | -0.21 | -0.06 | 0.033 |
| NRK | S1031; S1034 | 0.12 | 0.16 | 0.18 | -0.36 | 0.15 | -0.38 | 0.114 |
| NRK | S/Y/T | 0.17 | 0.28 | 0.16 | -0.28 | -0.28 | -0.16 | 0.001 |
| NRK | S1149 | 0.49 | 0.22 | 0.16 | -0.22 | -0.58 | -0.33 | 0.011 |
| NSD1 | S1210 | -0.40 | -0.03 | -0.22 | 0.10 | 0.06 | 0.36 | 0.05 |
| NSD1 | T1208 | -0.60 | 0.20 | -0.14 | 0.17 | -0.21 | 0.37 | 0.368 |
| NSD1 | S/T | -0.16 | -0.18 | -0.09 | 0.10 | -0.03 | 0.30 | 0.058 |
| NSD1 | S1334 | -0.18 | -0.06 | -0.07 | 0.01 | 0.22 | 0.04 | 0.065 |
| NSD1 | S948 | 0.00 | -0.28 | 0.05 | 0.12 | -0.08 | 0.15 | 0.321 |
| NSD1 | S1498 | 0.07 | 0.00 | -0.12 | 0.01 | -0.09 | 0.12 | 0.727 |
| NSD1 | S2471 | 0.13 | 0.08 | -0.08 | -0.03 | -0.15 | 0.03 | 0.321 |
| NSD1 | S486 | 0.13 | 0.12 | -0.04 | -0.08 | -0.24 | 0.08 | 0.24 |
| NSD1 | T/S | 0.25 | 0.22 | -0.04 | -0.28 | 0.10 | -0.37 | 0.129 |
| NSD1 | S2369 | 0.21 | 0.22 | 0.26 | -0.55 | -0.29 | -0.03 | 0.027 |
| NSD2 | S437 | -0.01 | -0.17 | -0.14 | 0.13 | -0.18 | 0.31 | 0.27 |
| NSD2 | T114; T115 | 0.05 | -0.08 | -0.05 | -0.07 | -0.06 | 0.19 | 0.671 |
| NSD2 | T114; T115 | -0.08 | 0.05 | 0.02 | -0.04 | 0.17 | -0.14 | 0.996 |

| | | | | | | | | |
|---|---|---|---|---|---|---|---|---|
| NSD2 | S380 | 0.11 | -0.10 | 0.01 | -0.13 | 0.06 | 0.03 | 0.824 |
| NSD2 | T544 | -0.26 | 0.01 | 0.31 | -0.28 | -0.10 | 0.22 | 0.762 |
| NSD2 | S121; T | 0.22 | 0.09 | -0.06 | -0.13 | -0.14 | -0.01 | 0.115 |
| NSD2 | S/Y/T | 0.22 | -0.03 | 0.06 | -0.13 | -0.18 | 0.02 | 0.121 |
| NSD2 | S1214 | -0.03 | 0.08 | 0.18 | -0.07 | 0.24 | -0.52 | 0.45 |
| NSD2 | S194 | 0.17 | 0.13 | 0.20 | -0.21 | -0.10 | -0.26 | 0.002 |
| NSD2 | T115 | 0.30 | 0.07 | 0.16 | -0.20 | -0.25 | -0.16 | 0.006 |
| NSD2 | S643 | 0.25 | 0.16 | 0.21 | -0.23 | -0.20 | -0.30 | 4E-04 |
| NSD2 | S637 | 0.34 | 0.17 | 0.20 | -0.29 | -0.24 | -0.33 | 9E-04 |
| NSD2 | T/S/Y | 0.30 | 0.26 | 0.14 | -0.23 | -0.60 | -0.07 | 0.032 |
| NSD3 | S489 | -0.53 | -0.46 | -0.39 | 0.25 | 0.27 | 0.51 | 0.001 |
| NSD3 | S671 | 0.05 | -0.06 | -0.08 | -0.17 | -0.06 | 0.28 | 0.76 |
| NSD3 | S561 | -0.03 | 0.00 | -0.08 | 0.19 | -0.44 | 0.27 | 0.861 |
| NSD3 | T675 | 0.08 | 0.22 | -0.01 | -0.19 | 0.04 | -0.18 | 0.106 |
| NSD3 | S671 | 0.08 | 0.22 | -0.01 | -0.19 | 0.04 | -0.18 | 0.106 |
| NSD3 | S150 | 0.23 | 0.07 | 0.08 | -0.33 | 0.07 | -0.19 | 0.1 |
| NSD3 | S/T | 0.13 | 0.08 | 0.36 | -0.19 | -0.16 | -0.34 | 0.015 |
| NSF | S739 | -0.24 | 0.07 | -0.04 | 0.14 | -0.26 | 0.25 | 0.568 |
| NSFL1C | S/T | 0.12 | -0.10 | 0.07 | 0.14 | -0.06 | -0.21 | 0.592 |
| NSMCE3 | S60; S64 | -0.44 | 0.18 | -0.14 | 0.37 | -0.31 | 0.18 | 0.476 |
| NSMCE3 | S303 | -0.03 | -0.20 | -0.01 | -0.13 | 0.14 | 0.19 | 0.268 |
| NSMCE3 | S64 | -0.04 | -0.10 | -0.10 | 0.02 | 0.01 | 0.18 | 0.071 |
| NSMCE3 | S64 | -0.19 | 0.18 | -0.01 | -0.11 | -0.25 | 0.30 | 0.937 |
| NSMCE4A | S32 | -0.46 | -0.27 | -0.22 | 0.20 | -0.32 | 0.71 | 0.165 |
| NSMCE4A | S61; S63 | -0.20 | -0.04 | -0.16 | 0.03 | -0.08 | 0.38 | 0.167 |
| NSMCE4A | T345 | 0.02 | -0.06 | -0.04 | -0.19 | -0.02 | 0.25 | 0.763 |
| NSMCE4A | T345 | 0.04 | -0.07 | 0.03 | -0.31 | -0.31 | 0.47 | 0.859 |
| NSMCE4A | T58; S61; S63 | 0.07 | 0.15 | 0.01 | 0.04 | -0.24 | -0.06 | 0.15 |
| NSMCE4A | S63 | 0.11 | 0.15 | -0.01 | -0.22 | 0.05 | -0.10 | 0.137 |
| NSMCE4A | T375 | 0.41 | -0.39 | 0.16 | 0.28 | -0.38 | -0.29 | 0.58 |
| NSRP1 | T275 | -0.03 | 0.01 | -0.10 | 0.15 | 0.01 | -0.05 | 0.336 |
| NSRP1 | S | -0.48 | -0.21 | 0.42 | -0.55 | -0.09 | 0.55 | 0.888 |
| NSRP1 | S248; S/T | -0.03 | 0.24 | -0.13 | -0.60 | 0.14 | 0.22 | 0.732 |
| NSRP1 | S248 | 0.23 | 0.02 | 0.05 | -0.03 | -0.13 | -0.17 | 0.053 |
| NSRP1 | S254 | 0.19 | 0.25 | 0.36 | -0.05 | -0.15 | -0.96 | 0.088 |
| NSUN2 | S139 | -0.24 | -0.21 | -0.06 | 0.35 | 0.08 | 0.01 | 0.054 |
| NSUN2 | S743; S751 | -0.25 | -0.21 | -0.03 | 0.26 | 0.11 | 0.05 | 0.033 |
| NSUN2 | S743; S751 | -0.26 | -0.09 | -0.12 | 0.31 | -0.13 | 0.20 | 0.123 |
| NSUN2 | S456 | -0.05 | -0.13 | -0.02 | 0.08 | 0.24 | -0.17 | 0.401 |
| NSUN2 | S456 | -0.09 | -0.01 | -0.05 | -0.33 | 0.53 | -0.20 | 0.858 |
| NSUN2 | S751 | -0.10 | -0.07 | 0.14 | 0.01 | -0.09 | 0.09 | 0.898 |
| NSUN2 | S743 | -0.06 | -0.04 | 0.15 | 0.08 | -0.03 | -0.11 | 0.704 |
| NSUN4 | S206 | 0.14 | 0.08 | 0.29 | -0.09 | -0.06 | -0.48 | 0.066 |
| NSUN5P2 | S143 | -0.12 | -0.07 | 0.10 | 0.02 | 0.19 | -0.15 | 0.695 |
| NSUN5P2 | T258; S260 | 0.04 | -0.04 | 0.00 | 0.03 | -0.08 | 0.04 | 0.965 |
| NTHL1 | S71 | -0.01 | -0.03 | -0.22 | 0.02 | -0.03 | 0.23 | 0.197 |
| NTHL1 | S53 | 0.10 | -0.03 | -0.17 | 0.02 | -0.08 | 0.13 | 0.607 |
| NTHL1 | S71 | 0.10 | 0.11 | -0.07 | -0.02 | -0.04 | -0.09 | 0.18 |

| | | | | | | | | |
|---|---|---|---|---|---|---|---|---|
| **NUCKS1** | **S54** | -0.28 | -0.32 | -0.48 | 0.02 | 0.55 | 0.23 | 0.019 |
| **NUCKS1** | **S54; S58; S61** | -0.18 | -0.36 | -0.53 | -0.14 | 0.61 | 0.28 | 0.065 |
| **NUCKS1** | **S58; S61** | -0.11 | -0.21 | -0.28 | -0.16 | 0.50 | 0.10 | 0.155 |
| **NUCKS1** | **S19** | -0.14 | -0.21 | -0.24 | -0.23 | 0.21 | 0.45 | 0.164 |
| **NUCKS1** | **S204; S214** | -0.01 | -0.17 | -0.33 | 0.04 | 0.17 | 0.22 | 0.042 |
| **NUCKS1** | **T34** | -0.01 | -0.04 | -0.39 | -0.10 | 0.16 | 0.29 | 0.188 |
| **NUCKS1** | **S19** | -0.13 | -0.18 | -0.09 | -0.31 | 0.16 | 0.42 | 0.356 |
| **NUCKS1** | **S54; S61** | -0.07 | -0.08 | -0.22 | -0.19 | 0.44 | 0.00 | 0.343 |
| **NUCKS1** | **T202; S214** | -0.01 | -0.08 | -0.21 | -0.12 | 0.16 | 0.22 | 0.195 |
| **NUCKS1** | **S132; S144** | -0.10 | -0.02 | -0.13 | 0.00 | 0.01 | 0.20 | 0.105 |
| **NUCKS1** | **S19** | -0.03 | -0.09 | -0.12 | -0.15 | 0.05 | 0.29 | 0.337 |
| **NUCKS1** | **S30** | 0.11 | -0.09 | -0.23 | 0.11 | 0.21 | -0.17 | 0.468 |
| **NUCKS1** | **T179; S181** | 0.03 | -0.12 | -0.10 | 0.03 | -0.10 | 0.23 | 0.325 |
| **NUCKS1** | **S75; S79** | -0.04 | -0.10 | 0.06 | 0.02 | -0.07 | 0.12 | 0.551 |
| **NUCKS1** | **S214** | -0.06 | 0.12 | -0.09 | -0.07 | -0.10 | 0.18 | 0.898 |
| **NUCKS1** | **S19** | 0.02 | 0.03 | -0.03 | -0.15 | 0.00 | 0.13 | 0.866 |
| **NUCKS1** | **S75; S79** | 0.05 | 0.12 | -0.19 | -0.68 | 0.08 | 0.40 | 0.862 |
| **NUCKS1** | **S223; S234** | -0.03 | -0.06 | 0.15 | -0.31 | 0.23 | -0.04 | 0.721 |
| **NUCKS1** | **S113** | 0.17 | 0.04 | -0.13 | -0.14 | 0.10 | -0.07 | 0.593 |
| **NUCKS1** | **S144** | 0.21 | -0.01 | -0.09 | -0.17 | -0.03 | 0.05 | 0.485 |
| **NUCKS1** | **S113** | 0.17 | 0.13 | -0.13 | -0.21 | 0.11 | -0.11 | 0.395 |
| **NUCKS1** | **S113** | 0.18 | 0.18 | -0.19 | -0.09 | 0.05 | -0.18 | 0.402 |
| **NUCKS1** | **T202** | 0.30 | 0.04 | -0.16 | -0.25 | 0.04 | -0.04 | 0.412 |
| **NUCKS1** | **S113** | 0.12 | 0.14 | -0.07 | -0.34 | 0.07 | 0.03 | 0.363 |
| **NUCKS1** | **S113** | 0.23 | 0.07 | -0.10 | -0.24 | 0.12 | -0.14 | 0.341 |
| **NUCKS1** | **S113** | 0.24 | 0.13 | -0.10 | -0.29 | 0.13 | -0.18 | 0.273 |
| **NUCKS1** | **S214** | 0.15 | 0.15 | -0.01 | -0.33 | -0.08 | 0.06 | 0.172 |
| **NUCKS1** | **S144** | 0.37 | 0.15 | -0.19 | -0.27 | -0.06 | -0.11 | 0.209 |
| **NUCKS1** | **S79** | 0.30 | 0.20 | -0.12 | -0.31 | -0.01 | -0.14 | 0.142 |
| **NUCKS1** | **S214** | 0.21 | 0.09 | 0.10 | -0.34 | 0.02 | -0.14 | 0.059 |
| **NUCKS1** | **S240** | 0.20 | -0.15 | 0.39 | 0.14 | -0.60 | -0.19 | 0.243 |
| **NUCKS1** | **S223** | 0.38 | 0.10 | 0.12 | -0.33 | -0.15 | -0.25 | 0.012 |
| **NUCKS1** | **S223** | 0.32 | 0.16 | 0.16 | -0.35 | -0.04 | -0.39 | 0.017 |
| **NUCKS1** | **S181** | 0.36 | 0.18 | 0.11 | -0.42 | -0.25 | -0.13 | 0.014 |
| **NUCKS1** | **S214** | 0.37 | 0.27 | 0.00 | -0.42 | 0.04 | -0.46 | 0.063 |
| **NUCKS1** | **S61** | 0.33 | 0.27 | 0.18 | -0.50 | 0.03 | -0.55 | 0.035 |
| **NUCKS1** | **S61** | 0.29 | 0.40 | 0.27 | -0.61 | -0.17 | -0.50 | 0.006 |
| **NUDT16** | **S183; S185; S187** | -0.17 | -0.24 | -0.22 | 0.25 | 0.11 | 0.18 | 0.001 |
| **NUF2** | **S247** | 0.07 | -0.04 | -0.02 | -0.26 | 0.21 | 0.00 | 0.875 |
| **NUFIP1** | **S403** | 0.05 | 0.14 | 0.06 | -0.35 | 0.06 | -0.01 | 0.233 |
| **NUFIP1** | **S403** | -0.09 | 0.56 | 0.17 | -0.68 | 0.01 | -0.27 | 0.127 |
| **NUFIP2** | **S304** | 0.01 | 0.07 | -0.21 | 0.14 | 0.03 | -0.07 | 0.503 |
| **NUFIP2** | **S212** | 0.01 | 0.08 | -0.07 | 0.16 | -0.15 | -0.05 | 0.857 |
| **NUFIP2** | **S212; S214** | -0.05 | -0.03 | 0.10 | 0.23 | -0.19 | -0.10 | 0.839 |
| **NUFIP2** | **S629** | 0.12 | 0.16 | 0.01 | -0.03 | -0.13 | -0.16 | 0.029 |
| **NUFIP2** | **S629** | 0.12 | 0.14 | 0.04 | -0.02 | -0.13 | -0.18 | 0.019 |
| **NUFIP2** | **S652** | 0.08 | 0.08 | 0.21 | -0.12 | -0.10 | -0.21 | 0.008 |
| **NUFIP2** | **S112** | 0.19 | -0.01 | 0.23 | 0.09 | -0.13 | -0.47 | 0.156 |

| | | | | | | | | |
|---|---|---|---|---|---|---|---|---|
| **NUMA1** | **S395** | -1.68 | -0.88 | -0.83 | 0.55 | 0.04 | 1.07 | 0.014 |
| **NUMA1** | **S1769** | -0.99 | -0.82 | -1.05 | 0.55 | 0.67 | 0.48 | 7E-05 |
| **NUMA1** | **S820** | -0.99 | -0.78 | -0.80 | 0.54 | 0.44 | 0.61 | 8E-05 |
| **NUMA1** | **S/T/Y** | -0.23 | -0.23 | -0.59 | 0.25 | 0.34 | 0.23 | 0.008 |
| **NUMA1** | **S1181** | -0.89 | -0.28 | 0.01 | -0.43 | 0.23 | 0.78 | 0.261 |
| **NUMA1** | **S1792** | -0.28 | -0.21 | -0.40 | 0.10 | 0.38 | 0.24 | 0.005 |
| **NUMA1** | **S1682** | -0.44 | -0.34 | -0.10 | 0.50 | -0.03 | 0.21 | 0.047 |
| **NUMA1** | **S801** | -0.56 | -0.08 | -0.20 | 0.40 | 0.06 | 0.19 | 0.044 |
| **NUMA1** | **S1830** | -0.12 | -0.35 | -0.12 | 0.16 | 0.16 | 0.20 | 0.01 |
| **NUMA1** | **S169** | -0.05 | -0.10 | -0.26 | 0.09 | -0.05 | 0.31 | 0.108 |
| **NUMA1** | **S1991** | -0.06 | -0.13 | -0.11 | 0.02 | 0.00 | 0.25 | 0.088 |
| **NUMA1** | **S1991** | 0.11 | 0.14 | -0.61 | -0.17 | 0.40 | -0.08 | 0.603 |
| **NUMA1** | **S271** | 0.01 | -0.03 | -0.06 | 0.01 | -0.01 | 0.09 | 0.184 |
| **NUMA1** | **T2000** | 0.12 | -0.02 | -0.19 | -0.06 | -0.27 | 0.35 | 0.853 |
| **NUMA1** | **S1757** | 0.20 | -0.09 | 0.14 | -0.18 | 0.02 | -0.12 | 0.168 |
| **NUMA1** | **S1757** | 0.04 | 0.09 | 0.13 | -0.17 | -0.03 | -0.08 | 0.02 |
| **NUMA1** | **T2000** | 0.00 | 0.13 | 0.36 | -0.44 | -0.19 | 0.02 | 0.096 |
| **NUMB** | **S425** | 0.01 | -0.08 | -0.02 | 0.15 | -0.08 | 0.01 | 0.481 |
| **NUMB** | **S425** | 0.02 | 0.09 | 0.02 | 0.05 | -0.10 | -0.08 | 0.191 |
| **NUMB** | **S634** | -0.16 | 0.13 | 0.37 | -0.23 | 0.05 | -0.27 | 0.224 |
| **NUP107** | **S37** | -0.12 | -0.27 | -0.44 | 0.33 | 0.16 | 0.18 | 0.01 |
| **NUP107** | **S11** | 0.14 | -0.06 | 0.14 | -0.10 | 0.11 | -0.28 | 0.274 |
| **NUP133** | **S480** | -0.55 | -0.06 | -0.27 | -0.23 | 0.17 | 0.63 | 0.171 |
| **NUP133** | **S131** | 0.15 | -0.07 | -0.08 | 0.29 | -0.20 | -0.17 | 0.893 |
| **NUP153** | **T97** | -1.27 | -0.34 | 0.41 | 0.26 | -0.12 | 0.44 | 0.31 |
| **NUP153** | **S217** | -0.25 | -0.09 | -0.50 | 0.22 | 0.04 | 0.40 | 0.033 |
| **NUP153** | **S334** | 0.00 | -0.05 | -0.46 | 0.14 | -0.02 | 0.28 | 0.152 |
| **NUP153** | **T588** | 0.06 | -0.06 | -0.40 | 0.22 | -0.02 | 0.12 | 0.197 |
| **NUP153** | **T588** | -0.10 | 0.07 | 0.10 | 0.03 | -0.05 | -0.06 | 0.514 |
| **NUP153** | **S209** | -0.10 | 0.05 | 0.16 | -0.21 | 0.21 | -0.17 | 0.588 |
| **NUP153** | **S192** | 0.14 | 0.11 | -0.10 | 0.03 | -0.05 | -0.15 | 0.293 |
| **NUP153** | **S614** | 0.24 | 0.09 | -0.10 | -0.07 | -0.01 | -0.19 | 0.207 |
| **NUP153** | **S320** | 0.19 | 0.03 | 0.03 | 0.07 | -0.01 | -0.36 | 0.258 |
| **NUP155** | **S992** | 0.21 | 0.11 | 0.06 | -0.07 | -0.23 | -0.13 | 0.013 |
| **NUP155** | **S992** | 0.38 | 0.20 | -0.15 | -0.37 | -0.18 | -0.01 | 0.15 |
| **NUP188** | **S1709** | -0.30 | -0.15 | -0.30 | 0.46 | 0.22 | -0.10 | 0.057 |
| **NUP188** | **S/T** | -0.13 | -0.02 | -0.34 | 0.28 | 0.11 | 0.03 | 0.067 |
| **NUP188** | **S1717; S/T** | -0.25 | 0.15 | -0.38 | 0.11 | 0.05 | 0.22 | 0.163 |
| **NUP205** | **S/T** | 0.69 | 0.07 | -0.63 | 0.19 | -0.02 | -0.83 | 0.62 |
| **NUP210** | **T1844** | 0.08 | 0.00 | -0.06 | 0.18 | -0.11 | -0.12 | 0.804 |
| **NUP214** | **S430** | -0.03 | -0.08 | -0.40 | 0.22 | 0.02 | 0.19 | 0.074 |
| **NUP214** | **S430; S433; T434** | -0.04 | -0.29 | -0.18 | 0.28 | -0.08 | 0.22 | 0.08 |
| **NUP214** | **T987; S989; S991** | 0.00 | -0.21 | -0.13 | 0.24 | 0.00 | 0.06 | 0.091 |
| **NUP214** | **S986** | -0.05 | 0.01 | -0.18 | 0.07 | -0.17 | 0.27 | 0.394 |
| **NUP214** | **S430** | -0.19 | -0.14 | 0.16 | 0.15 | 0.05 | -0.07 | 0.461 |
| **NUP214** | **S433; T437** | 0.09 | -0.04 | -0.07 | 0.04 | -0.16 | 0.12 | 0.929 |
| **NUP214** | **S657** | 0.17 | 0.13 | -0.10 | 0.03 | -0.04 | -0.21 | 0.27 |
| **NUP214** | **S** | 0.06 | 0.01 | 0.23 | -0.02 | -0.21 | -0.12 | 0.063 |

| | | | | | | | | |
|---|---|---|---|---|---|---|---|---|
| NUP214 | S678 | 0.25 | 0.41 | 0.28 | -0.12 | -0.17 | -1.10 | 0.072 |
| NUP35 | S/T | -0.19 | -0.02 | 0.03 | 0.08 | 0.39 | -0.41 | 0.755 |
| NUP35 | T308 | 0.08 | 0.00 | -0.13 | 0.19 | 0.11 | -0.30 | 0.93 |
| NUP35 | T106 | -0.01 | 0.14 | -0.05 | -0.16 | 0.00 | 0.06 | 0.522 |
| NUP35 | T273 | 0.10 | 0.09 | 0.24 | -0.13 | 0.15 | -0.59 | 0.209 |
| NUP35 | S73 | 0.21 | 0.29 | 0.20 | -0.33 | -0.14 | -0.39 | 0.003 |
| NUP35 | S73 | 0.25 | 0.61 | 0.10 | -1.50 | 0.31 | -0.04 | 0.275 |
| NUP42 | S106 | 0.02 | -0.07 | -0.12 | 0.10 | 0.02 | 0.04 | 0.086 |
| NUP50 | S221 | 0.11 | -0.03 | -0.02 | -0.04 | -0.01 | -0.01 | 0.474 |
| NUP54 | S289 | -0.12 | 0.34 | 0.20 | -0.19 | 0.06 | -0.40 | 0.171 |
| NUP58 | T331 | -0.05 | 0.10 | -0.31 | 0.32 | -0.15 | 0.01 | 0.467 |
| NUP62 | S408 | -0.10 | 0.03 | -0.27 | 0.09 | 0.33 | -0.16 | 0.294 |
| NUP62 | S418 | -0.01 | 0.19 | -0.05 | 0.01 | -0.12 | -0.05 | 0.315 |
| NUP85 | T90 | -0.01 | -0.14 | -0.08 | -0.12 | 0.05 | 0.26 | 0.287 |
| NUP88 | T525 | -0.02 | -0.07 | 0.02 | 0.02 | 0.25 | -0.24 | 0.812 |
| NUP88 | S517 | 0.15 | 0.12 | 0.02 | -0.02 | -0.04 | -0.25 | 0.07 |
| NUP88 | S/T | 0.13 | 0.18 | 0.27 | -0.14 | 0.16 | -0.89 | 0.2 |
| NUP93 | S614 | 0.01 | -0.09 | 0.24 | 0.20 | -0.08 | -0.37 | 0.522 |
| NUP93 | T/S | 0.13 | 0.16 | 0.05 | -0.07 | -0.01 | -0.30 | 0.064 |
| NUP93 | S52 | 0.43 | 0.38 | 0.48 | -0.38 | -0.30 | -1.43 | 0.036 |
| NUP98 | T/Y | -0.34 | -0.24 | -0.25 | 0.56 | -0.03 | 0.09 | 0.057 |
| NUP98 | S1465 | -0.31 | 0.22 | -0.40 | 0.05 | 0.11 | 0.21 | 0.221 |
| NUP98 | T1000 | -0.11 | -0.32 | -0.02 | 0.15 | -0.08 | 0.30 | 0.127 |
| NUP98 | S612 | -0.11 | -0.30 | 0.02 | 0.21 | 0.08 | 0.05 | 0.078 |
| NUP98 | S1803 | -0.23 | -0.18 | 0.07 | 0.22 | -0.16 | 0.21 | 0.267 |
| NUP98 | S683 | -0.46 | -0.03 | 0.19 | 0.09 | 0.01 | 0.11 | 0.432 |
| NUP98 | S1465 | -0.02 | -0.03 | -0.09 | 0.00 | -0.11 | 0.23 | 0.428 |
| NUP98 | S623 | -0.02 | 0.04 | 0.02 | -0.14 | 0.02 | 0.07 | 0.649 |
| NUP98 | S623 | 0.01 | 0.03 | 0.03 | -0.01 | -0.02 | -0.05 | 0.029 |
| NUP98 | S683 | 0.00 | 0.30 | -0.34 | 0.10 | 0.45 | -0.88 | 0.83 |
| NUP98 | S888 | 0.07 | 0.02 | 0.28 | -0.13 | -0.18 | -0.13 | 0.03 |
| NUP98 | S1023 | 0.22 | 0.04 | 0.17 | -0.19 | -0.11 | -0.18 | 0.007 |
| NUP98 | S612 | 0.39 | 0.24 | 0.27 | -0.19 | -0.17 | -0.91 | 0.043 |
| NUSAP1 | S135 | -0.12 | -0.07 | -0.01 | 0.09 | -0.03 | 0.11 | 0.087 |
| NUSAP1 | S135 | -0.10 | -0.05 | 0.00 | 0.12 | -0.06 | 0.09 | 0.196 |
| NUSAP1 | S60 | 0.15 | 0.37 | 0.27 | -0.28 | -0.13 | -0.62 | 0.018 |
| NUSAP1 | T/S | 0.21 | 0.34 | 0.25 | -0.62 | 0.27 | -0.90 | 0.125 |
| NVL | T138 | -0.01 | -0.52 | -0.63 | -0.03 | 0.47 | 0.38 | 0.054 |
| NVL | T138 | -0.29 | -0.07 | -0.35 | 0.25 | 0.28 | 0.04 | 0.02 |
| NVL | T138 | -0.08 | -0.17 | -0.15 | 0.16 | 0.32 | -0.15 | 0.16 |
| NVL | S185 | 0.02 | -0.17 | -0.27 | -0.20 | 0.48 | 0.01 | 0.343 |
| NVL | T138 | -0.29 | -0.12 | 0.02 | 0.04 | 0.45 | -0.23 | 0.378 |
| NVL | T138 | -0.10 | -0.12 | -0.08 | 0.21 | -0.15 | 0.20 | 0.186 |
| NVL | S191 | 0.01 | -0.02 | -0.05 | -0.16 | 0.09 | 0.11 | 0.748 |
| NVL | S439 | 0.04 | -0.15 | 0.17 | 0.01 | 0.08 | -0.19 | 0.676 |
| NVL | S | 0.02 | 0.06 | 0.02 | -0.04 | -0.13 | 0.05 | 0.204 |
| NVL | S211 | 0.09 | 0.12 | 0.08 | 0.05 | -0.04 | -0.36 | 0.164 |
| NXF1 | S9 | -0.28 | -0.08 | -0.57 | 0.63 | -0.03 | 0.05 | 0.104 |

| | | | | | | | | |
|---|---|---|---|---|---|---|---|---|
| **NXF1** | **S111** | -0.29 | -0.24 | -0.29 | 0.31 | -0.03 | 0.38 | 0.018 |
| **NXF1** | **S223** | -0.07 | -0.76 | 0.12 | 0.48 | -0.51 | 0.35 | 0.448 |
| **OASL** | **S396** | 0.22 | 0.05 | 0.05 | 0.09 | -0.10 | -0.37 | 0.182 |
| **OBI1** | **S210** | -0.16 | -0.29 | -0.05 | 0.19 | -0.11 | 0.33 | 0.107 |
| **OBSL1** | **S191** | -0.58 | -0.49 | -0.46 | 0.72 | -0.43 | 0.59 | 0.093 |
| **OBSL1** | **S10** | 0.20 | 0.15 | 0.11 | -0.22 | -0.03 | -0.27 | 0.014 |
| **OCIAD1** | **S213** | 0.09 | -0.09 | -0.18 | 0.07 | 0.13 | -0.04 | 0.312 |
| **OCIAD1** | **S108** | 0.23 | -0.02 | 0.07 | -0.17 | -0.09 | -0.05 | 0.073 |
| **OCIAD1** | **S209** | 0.18 | 0.05 | 0.07 | -0.31 | 0.10 | -0.15 | 0.156 |
| **OCIAD1** | **S108** | 0.22 | 0.12 | 0.23 | -0.15 | -0.14 | -0.37 | 0.008 |
| **OCIAD1** | **S123** | 0.21 | 0.29 | 0.16 | -0.25 | -0.30 | -0.23 | 3E-04 |
| **OCIAD1** | **S123** | 0.31 | 0.26 | 0.10 | -0.16 | -0.26 | -0.39 | 0.006 |
| **OCLN** | **S358** | -0.57 | -0.17 | 0.13 | -0.06 | 0.03 | 0.45 | 0.252 |
| **OCLN** | **S321** | -0.19 | -0.03 | -0.02 | 0.07 | -0.01 | 0.15 | 0.102 |
| **OCLN** | **S310** | -0.06 | -0.04 | -0.02 | 0.03 | -0.06 | 0.14 | 0.244 |
| **OCLN** | **S408** | 0.04 | -0.02 | -0.14 | 0.03 | -0.23 | 0.28 | 0.695 |
| **OCLN** | **S321** | -0.28 | -0.01 | 0.20 | 0.02 | 0.01 | 0.02 | 0.758 |
| **OCLN** | **S321** | -0.40 | 0.26 | 0.04 | 0.26 | -0.02 | -0.27 | 0.941 |
| **OCLN** | **T/S/Y** | 0.06 | -0.02 | -0.06 | -0.24 | -0.05 | 0.27 | 0.989 |
| **OCLN** | **S358** | -0.07 | 0.04 | 0.05 | -0.15 | 0.04 | 0.07 | 0.783 |
| **OCLN** | **S369** | -0.34 | 0.35 | -0.07 | 0.37 | 0.02 | -0.56 | 0.918 |
| **ODF2** | **S106; S109; T110** | -0.12 | 0.05 | -0.02 | 0.12 | -0.08 | 0.04 | 0.464 |
| **ODF2** | **S261** | -0.05 | 0.00 | 0.06 | 0.13 | -0.07 | -0.09 | 0.882 |
| **ODF2** | **T92** | -0.03 | 0.14 | 0.06 | -0.05 | 0.03 | -0.18 | 0.193 |
| **ODF2** | **T92** | 0.15 | 0.14 | 0.04 | -0.26 | 0.10 | -0.23 | 0.119 |
| **ODF2** | **S109; T110** | 0.12 | 0.18 | 0.21 | -0.25 | -0.18 | -0.16 | 6E-04 |
| **ODF2L** | **S9** | 0.00 | -0.09 | 0.29 | -0.14 | -0.02 | -0.08 | 0.285 |
| **ODF2L** | **S9** | 0.24 | -0.10 | 0.27 | -0.14 | 0.03 | -0.43 | 0.152 |
| **OFD1** | **S686** | -0.06 | -0.30 | -0.02 | -0.06 | 0.24 | 0.14 | 0.129 |
| **OFD1** | **S735** | -0.33 | 0.07 | -0.04 | 0.37 | -0.11 | -0.05 | 0.425 |
| **OFD1** | **S774** | 0.02 | 0.15 | 0.05 | -0.29 | -0.01 | 0.05 | 0.233 |
| **OFD1** | **S762** | -0.09 | 0.39 | 0.03 | 0.10 | 0.03 | -0.68 | 0.365 |
| **OGFR** | **S361; S378** | -0.87 | -0.90 | -1.05 | 0.68 | 0.51 | 0.49 | 5E-05 |
| **OGFR** | **S315** | -0.51 | -0.36 | -0.43 | 0.25 | 0.22 | 0.52 | 0.002 |
| **OIP5** | **S** | -0.25 | 0.33 | -0.20 | -0.32 | 0.20 | 0.11 | 0.881 |
| **OIP5** | **S225** | 0.18 | 0.18 | 0.25 | -0.16 | -0.45 | -0.13 | 0.014 |
| **ONECUT2** | **S/T** | -0.19 | 0.01 | 0.10 | -0.20 | 0.24 | -0.02 | 0.84 |
| **ORAI1** | **S13** | -0.08 | -0.04 | 0.21 | -0.09 | -0.04 | 0.02 | 0.524 |
| **ORC1** | **T337** | -0.26 | -0.50 | 0.10 | 0.21 | 0.17 | 0.14 | 0.09 |
| **ORC1** | **S311** | -0.07 | -0.20 | -0.16 | -0.04 | 0.13 | 0.28 | 0.057 |
| **ORC1** | **S273** | 0.20 | -0.37 | -0.22 | -0.01 | 0.08 | 0.22 | 0.285 |
| **ORC1** | **S199** | 0.01 | -0.03 | 0.00 | -0.02 | 0.03 | 0.02 | 0.457 |
| **ORC1** | **T230** | 0.09 | -0.11 | 0.02 | 0.11 | -0.13 | 0.01 | 0.965 |
| **ORC1** | **S311** | 0.09 | 0.03 | -0.10 | -0.07 | 0.22 | -0.21 | 0.849 |
| **ORC1** | **S287** | 0.16 | 0.00 | 0.13 | -0.18 | 0.14 | -0.32 | 0.212 |
| **ORC1** | **T/S** | 0.15 | 0.24 | 0.07 | -0.55 | 0.02 | -0.05 | 0.144 |
| **ORC2** | **S280** | 0.11 | 0.06 | -0.29 | -0.10 | 0.05 | 0.13 | 0.668 |
| **ORC2** | **T226** | 0.08 | 0.16 | -0.29 | -0.21 | 0.18 | 0.02 | 0.928 |

| | | | | | | | | |
|---|---|---|---|---|---|---|---|---|
| ORC6 | T195 | 0.06 | -0.07 | 0.17 | 0.13 | -0.15 | -0.19 | 0.379 |
| ORMDL2 | T74 | -0.07 | -0.03 | -0.03 | -1.08 | -0.19 | 0.81 | 0.852 |
| OSBP | S240 | 0.00 | 0.01 | 0.01 | 0.11 | -0.23 | 0.07 | 0.832 |
| OSBP | S190; S193 | 0.12 | 0.12 | 0.17 | -0.15 | -0.17 | -0.15 | 7E-05 |
| OSBP | S190 | 0.08 | 0.19 | 0.25 | -0.05 | -0.25 | -0.31 | 0.015 |
| OSBP | S190; S193 | 0.13 | 0.28 | 0.18 | -0.21 | -0.10 | -0.39 | 0.011 |
| OSBP | S193 | 0.06 | 0.17 | 0.44 | -0.25 | 0.05 | -0.74 | 0.105 |
| OSBPL10 | S57 | 0.11 | 0.11 | 0.16 | -0.35 | 0.22 | -0.36 | 0.208 |
| OSBPL2 | S341 | -0.33 | -0.08 | -0.16 | -0.07 | 0.15 | 0.38 | 0.082 |
| OSBPL3 | S437 | -0.03 | -0.13 | -0.02 | 0.24 | -0.08 | -0.01 | 0.362 |
| OSBPL3 | S/T | -0.04 | 0.19 | -0.03 | 0.08 | 0.04 | -0.28 | 0.533 |
| OSBPL3 | S | -0.14 | 0.01 | 0.27 | -0.26 | 0.13 | -0.07 | 0.531 |
| OSBPL3 | S34 | 0.17 | 0.11 | -0.06 | -0.12 | -0.10 | -0.03 | 0.099 |
| OSBPL3 | S304 | 0.13 | 0.09 | 0.21 | -0.06 | 0.00 | -0.48 | 0.107 |
| OSBPL3 | S | 0.05 | -0.11 | 0.47 | -0.88 | 0.33 | -0.23 | 0.365 |
| OSBPL3 | S410 | 0.77 | -0.08 | 0.36 | -0.44 | 0.41 | -1.50 | 0.231 |
| OSBPL5 | S325; S327; S/T | -0.02 | 0.07 | -0.02 | -0.04 | -0.02 | 0.02 | 0.581 |
| OSBPL5 | S325 | -0.02 | 0.13 | 0.10 | -0.01 | -0.09 | -0.13 | 0.059 |
| OSBPL5 | T64; S67 | 0.14 | -0.02 | 0.15 | -0.23 | 0.06 | -0.15 | 0.128 |
| OSBPL5 | T64 | 0.08 | 0.04 | 0.27 | -0.10 | -0.08 | -0.26 | 0.038 |
| OSBPL5 | S747 | 0.07 | 0.15 | 0.36 | -0.20 | -0.15 | -0.34 | 0.014 |
| OSBPL5 | S88 | 0.37 | 0.36 | 0.38 | -0.40 | -0.28 | -0.91 | 0.01 |
| OSBPL6 | S9 | 0.01 | 0.00 | -0.09 | 0.04 | -0.10 | 0.12 | 0.546 |
| OSBPL6 | S44 | 0.04 | 0.06 | 0.11 | -0.04 | 0.10 | -0.32 | 0.26 |
| OSBPL7 | S236 | -0.67 | 0.34 | -0.35 | 0.15 | -0.03 | 0.30 | 0.307 |
| OSBPL8 | S314 | -0.08 | -0.08 | -0.21 | 0.37 | 0.05 | -0.13 | 0.233 |
| OSBPL8 | T/S/Y | -0.19 | -0.16 | 0.17 | 0.05 | 0.22 | -0.14 | 0.541 |
| OSBPL8 | S807; S808; S810 | -0.17 | -0.11 | 0.21 | -0.02 | -0.26 | 0.27 | 0.933 |
| OSBPL8 | S/Y/T | -0.28 | 0.07 | 0.16 | 0.35 | -0.53 | 0.06 | 0.947 |
| OSBPL8 | S358; T/S/Y | -0.08 | 0.04 | 0.30 | -0.29 | -0.02 | -0.02 | 0.238 |
| OSBPL8 | S810 | 0.20 | 0.13 | 0.04 | 0.10 | -0.23 | -0.31 | 0.11 |
| OSBPL8 | S353; T357 | 0.00 | 0.13 | 0.38 | -0.47 | 0.03 | -0.21 | 0.104 |
| OSBPL8 | S65 | 0.11 | 0.21 | 0.47 | -0.55 | 0.12 | -0.71 | 0.082 |
| OSTF1 | S213 | 0.13 | 0.25 | 0.00 | -0.10 | -0.03 | -0.31 | 0.072 |
| OTUD3 | S224 | -0.12 | -0.10 | -0.14 | -0.12 | 0.12 | 0.30 | 0.146 |
| OTUD4 | S1023; S1024 | -0.10 | -0.07 | -0.09 | 0.01 | 0.00 | 0.22 | 0.091 |
| OTUD4 | S1006 | 0.00 | 0.08 | -0.22 | 0.07 | 0.16 | -0.14 | 0.581 |
| OTUD4 | S546 | -0.03 | -0.04 | 0.14 | 0.13 | 0.05 | -0.28 | 0.691 |
| OTUD4 | S443 | 0.12 | 0.14 | -0.10 | -0.12 | 0.01 | -0.08 | 0.241 |
| OTUD5 | S64 | -0.16 | 0.07 | -0.07 | 0.04 | 0.10 | 0.01 | 0.217 |
| OTUD5 | S64 | -0.14 | 0.24 | -0.22 | -0.24 | 0.19 | 0.10 | 0.804 |
| OTUD7B | S467 | 0.21 | 0.30 | 0.13 | -0.25 | -0.01 | -0.55 | 0.043 |
| OTUD7B | S464 | 0.25 | 0.42 | 0.00 | -0.33 | -0.13 | -0.39 | 0.026 |
| OVOL1 | T158 | -0.21 | -0.17 | -0.29 | 0.27 | 0.18 | 0.12 | 0.002 |
| OVOL1 | S239 | -0.13 | -0.30 | -0.18 | 0.19 | -0.17 | 0.44 | 0.125 |
| OVOL1 | S112 | -0.40 | -0.20 | 0.03 | 0.10 | 0.27 | 0.11 | 0.063 |
| OVOL2 | S269 | 0.12 | 0.05 | 0.04 | -0.06 | -0.12 | -0.05 | 0.012 |
| OVOL2 | S269 | 0.06 | 0.10 | 0.17 | -0.12 | -0.06 | -0.17 | 0.007 |

| | | | | | | | | |
|---|---|---|---|---|---|---|---|---|
| OVOL2 | S269 | 0.23 | -0.34 | 0.46 | -0.42 | 0.03 | -0.18 | 0.318 |
| OXSR1 | S339 | 0.22 | 0.05 | -0.16 | -0.01 | 0.01 | -0.14 | 0.527 |
| P2RX4 | S384 | -0.22 | -0.03 | -0.18 | 0.22 | -0.05 | 0.20 | 0.06 |
| PABPC1 | S629 | 0.11 | -0.49 | -0.29 | 0.20 | 0.31 | 0.00 | 0.12 |
| PABPC1 | S96 | 0.01 | -0.01 | 0.12 | 0.06 | -0.16 | -0.04 | 0.308 |
| PABPC4 | T/S | -0.53 | 0.29 | 0.19 | 0.10 | -0.22 | 0.02 | 0.943 |
| PABPN1 | S95 | -0.90 | -0.88 | -0.93 | 0.53 | 0.41 | 0.70 | 6E-05 |
| PABPN1 | S90; S95 | -0.65 | -0.85 | -0.64 | 0.41 | 0.33 | 0.66 | 7E-04 |
| PABPN1 | S150 | -0.55 | -0.37 | -0.27 | 0.19 | 0.31 | 0.42 | 0.003 |
| PABPN1 | S150 | -0.47 | -0.34 | -0.29 | 0.10 | 0.07 | 0.64 | 0.029 |
| PABPN1 | S95 | -0.25 | 0.08 | -0.77 | 0.08 | 0.29 | 0.31 | 0.105 |
| PABPN1 | S150 | -0.12 | -0.24 | 0.14 | -0.19 | 0.20 | 0.15 | 0.488 |
| PACS1 | S495 | -0.32 | -0.21 | 0.68 | -0.34 | 0.05 | -0.13 | 0.611 |
| PACSIN3 | T324 | -0.30 | -0.11 | -0.40 | 0.21 | 0.08 | 0.37 | 0.015 |
| PACSIN3 | S319 | -0.17 | -0.11 | 0.07 | -0.03 | -0.05 | 0.25 | 0.353 |
| PACSIN3 | S354 | -0.48 | -0.30 | 0.52 | -0.30 | 0.00 | 0.30 | 0.815 |
| PACSIN3 | S383 | 0.02 | -0.16 | 0.07 | 0.11 | -0.17 | 0.09 | 0.765 |
| PACSIN3 | S354 | 0.26 | 0.30 | 0.39 | -0.49 | -0.33 | -0.40 | 3E-04 |
| PAG1 | S/Y/T | 0.28 | -0.02 | -0.05 | -0.26 | 0.30 | -0.39 | 0.482 |
| PAGR1 | S237 | 0.08 | 0.07 | 0.04 | -0.16 | 0.01 | -0.06 | 0.055 |
| PAICS | S27 | 0.55 | 0.31 | -0.05 | -0.22 | -0.39 | -0.49 | 0.029 |
| PAK1 | T/S | -0.20 | 0.03 | -0.11 | -0.17 | -0.56 | 0.69 | 0.85 |
| PAK1IP1 | S320 | 0.04 | 0.04 | -0.02 | 0.02 | 0.05 | -0.13 | 0.502 |
| PAK2 | T169 | -0.29 | 0.03 | 0.07 | 0.04 | 0.14 | -0.04 | 0.434 |
| PAK2 | S141 | 0.45 | 0.22 | -0.03 | -0.11 | -0.10 | -0.66 | 0.095 |
| PAK4 | S104 | -0.04 | 0.11 | 0.33 | -0.18 | -0.06 | -0.23 | 0.068 |
| PAK6 | S616 | -0.15 | -0.16 | 0.01 | 0.20 | 0.10 | -0.04 | 0.094 |
| PALB2 | S357 | -0.03 | -0.16 | -0.32 | -0.01 | 0.13 | 0.31 | 0.065 |
| PALB2 | S660 | 0.14 | 0.00 | -0.02 | -0.04 | -0.01 | -0.08 | 0.2 |
| PALB2 | S209 | 0.11 | -0.01 | 0.19 | 0.03 | -0.45 | 0.06 | 0.29 |
| PALB2 | S209 | 0.14 | -0.07 | 0.19 | -0.04 | -0.57 | 0.21 | 0.423 |
| PALB2 | S376 | 0.11 | 0.23 | 0.05 | -0.30 | -0.17 | 0.01 | 0.053 |
| PALB2 | S387 | 0.18 | 0.14 | 0.23 | -0.15 | -0.15 | -0.34 | 0.005 |
| PALS1 | S245 | -0.20 | -0.38 | -0.58 | 0.29 | -0.19 | 0.67 | 0.077 |
| PALS1 | S25 | -0.23 | 0.08 | 0.24 | -0.31 | -0.25 | 0.34 | 0.693 |
| PALS2 | S106 | 0.16 | 0.00 | 0.25 | 0.05 | -0.23 | -0.31 | 0.085 |
| PALS2 | S106 | 0.09 | 0.25 | 0.12 | -0.32 | 0.02 | -0.25 | 0.044 |
| PAN2 | S1189 | -0.04 | -0.07 | -0.08 | -0.02 | -0.07 | 0.25 | 0.306 |
| PAN3 | S354 | -0.29 | -0.42 | -0.65 | 0.40 | -0.34 | 0.76 | 0.099 |
| PAN3 | S/T | -0.63 | -0.97 | 0.23 | 0.10 | 0.06 | 0.64 | 0.15 |
| PAPOLA | S545 | 0.46 | 0.31 | 0.47 | -0.75 | -0.43 | -0.62 | 7E-04 |
| PAPOLG | S648 | -0.15 | 0.10 | -0.24 | 0.07 | -0.12 | 0.28 | 0.322 |
| PARD3 | S852 | -0.06 | 0.04 | -0.38 | 0.07 | -0.03 | 0.28 | 0.198 |
| PARD3 | S/T | -0.03 | -0.05 | -0.06 | 0.16 | -0.16 | 0.11 | 0.449 |
| PARD3 | S383 | -0.10 | -0.05 | 0.13 | 0.00 | 0.08 | -0.08 | 0.956 |
| PARD3 | S/T | -0.03 | 0.18 | 0.09 | -0.14 | -0.39 | 0.19 | 0.335 |
| PARD3B | S338 | 0.46 | -0.33 | 0.00 | -0.52 | 0.17 | 0.01 | 0.643 |
| PARD3B | S/T | 0.11 | 0.19 | 0.22 | -0.18 | -0.04 | -0.39 | 0.025 |

| | | | | | | | | |
|---|---|---|---|---|---|---|---|---|
| PARD6B | S11 | -0.37 | 0.26 | 0.14 | 0.06 | -0.22 | 0.04 | 0.821 |
| PARG | S291 | -0.09 | 0.16 | -0.08 | 0.00 | -0.10 | 0.09 | 0.962 |
| PARN | T166 | 0.05 | -0.44 | -0.53 | 0.12 | 0.29 | 0.28 | 0.047 |
| PARN | S619 | -0.06 | -0.26 | -0.43 | 0.63 | 0.19 | -0.38 | 0.272 |
| PARN | S619 | 0.02 | -0.07 | -0.19 | 0.19 | -0.01 | 0.04 | 0.152 |
| PARN | S557 | -0.01 | -0.34 | 0.15 | 0.20 | 0.12 | -0.19 | 0.601 |
| PARN | S163 | -0.12 | -0.01 | 0.02 | -0.11 | 0.14 | 0.06 | 0.441 |
| PARN | S163 | -0.04 | 0.17 | 0.05 | -0.07 | 0.04 | -0.18 | 0.212 |
| PARN | S/T | 0.09 | -0.11 | 0.24 | -0.19 | -0.06 | -0.01 | 0.238 |
| PARN | S570; S/T | 0.06 | 0.09 | 0.25 | 0.03 | 0.01 | -0.55 | 0.206 |
| PARN | S583 | 0.11 | 0.04 | 0.36 | -0.19 | -0.07 | -0.37 | 0.043 |
| PARP1 | S179 | -0.38 | -0.27 | -0.55 | 0.31 | 0.21 | 0.41 | 0.002 |
| PARP1 | S782 | -0.29 | -0.24 | -0.06 | 0.34 | -0.29 | 0.37 | 0.212 |
| PARP1 | S75 | 0.10 | 0.00 | 0.03 | -0.04 | -0.06 | -0.04 | 0.037 |
| PARP1 | S362 | 0.18 | 0.16 | -0.21 | -0.24 | 0.26 | -0.26 | 0.603 |
| PARP1 | S362 | 0.16 | 0.05 | -0.04 | -0.03 | 0.17 | -0.38 | 0.485 |
| PARP1 | S/T | 0.02 | 0.30 | 0.00 | 0.08 | 0.21 | -0.87 | 0.451 |
| PARP1 | T368 | 0.21 | 0.21 | 0.10 | -0.26 | 0.01 | -0.38 | 0.036 |
| PARP10 | S925 | -0.25 | -0.23 | 0.06 | 0.18 | 0.07 | 0.11 | 0.066 |
| PARP10 | S431 | -0.14 | -0.14 | 0.29 | -0.02 | -0.01 | -0.03 | 0.899 |
| PARP12 | S263 | -0.39 | 0.02 | -0.20 | 0.09 | -0.14 | 0.46 | 0.198 |
| PARP14 | S823 | -0.21 | -0.12 | -0.11 | 0.18 | -0.17 | 0.35 | 0.164 |
| PARP14 | S786 | 0.14 | 0.00 | -0.11 | 0.10 | 0.16 | -0.36 | 0.799 |
| PARP2 | S226 | -0.13 | -0.09 | -0.04 | 0.04 | 0.19 | 0.02 | 0.048 |
| PARP2 | S/T | 0.08 | 0.08 | 0.18 | -0.03 | -0.38 | 0.01 | 0.129 |
| PARP2 | S34 | 0.07 | 0.04 | 0.25 | -0.16 | -0.20 | -0.04 | 0.036 |
| PARP2 | S34 | 0.16 | 0.14 | 0.28 | -0.16 | -0.04 | -0.53 | 0.045 |
| PARP2 | S34 | 0.03 | 0.25 | 0.33 | -0.28 | -0.17 | -0.30 | 0.01 |
| PARP2 | S34 | 0.13 | 0.26 | 0.31 | -0.28 | -0.19 | -0.39 | 0.003 |
| PARP2 | S34 | 0.32 | 0.40 | 0.48 | -0.54 | -0.20 | -1.06 | 0.017 |
| PATJ | S1212 | -0.21 | -0.13 | -0.09 | 0.17 | 0.10 | 0.12 | 0.003 |
| PATJ | S1212 | 0.09 | -0.32 | -0.15 | 0.23 | -0.27 | 0.29 | 0.383 |
| PATJ | S645 | -0.15 | -0.13 | -0.05 | 0.06 | -0.06 | 0.28 | 0.125 |
| PATJ | S522 | -0.16 | -0.09 | -0.06 | 0.06 | 0.02 | 0.20 | 0.034 |
| PATJ | S645 | 0.00 | -0.09 | 0.07 | 0.14 | -0.16 | 0.02 | 0.96 |
| PATJ | S645 | -0.09 | 0.00 | 0.15 | -0.05 | 0.08 | -0.11 | 0.626 |
| PATJ | S455 | 0.20 | 0.15 | 0.12 | -0.14 | -0.07 | -0.33 | 0.016 |
| PATL1 | S179 | -0.29 | 0.07 | -0.04 | 0.14 | 0.08 | 0.00 | 0.231 |
| PATL1 | T194 | 0.18 | 0.02 | 0.10 | -0.36 | 0.12 | -0.12 | 0.2 |
| PATZ1 | T576 | -0.03 | 0.15 | 0.00 | -0.27 | -0.28 | 0.32 | 0.607 |
| PAXBP1 | S295 | 0.01 | -0.05 | -0.08 | -0.03 | 0.00 | 0.15 | 0.28 |
| PAXBP1 | T242 | 0.06 | -0.20 | 0.02 | -0.09 | -0.10 | 0.26 | 0.677 |
| PAXBP1 | S295 | -0.04 | -0.07 | 0.03 | -0.06 | -0.05 | 0.18 | 0.577 |
| PAXBP1 | S155 | 0.03 | -0.08 | 0.05 | -0.05 | -0.04 | 0.09 | 0.951 |
| PAXBP1 | S16 | 0.11 | 0.05 | 0.14 | -0.12 | -0.07 | -0.14 | 0.004 |
| PAXBP1 | S154; S155 | -0.08 | 0.12 | 0.24 | -0.04 | -0.45 | 0.11 | 0.319 |
| PAXBP1 | S16 | 0.16 | -0.01 | 0.22 | -0.16 | -0.13 | -0.13 | 0.018 |
| PAXIP1 | S235 | -0.01 | 0.28 | 0.43 | 0.12 | -0.43 | -0.67 | 0.105 |

| | | | | | | | | |
|---|---|---|---|---|---|---|---|---|
| PAXX | S148 | -0.15 | -0.30 | -0.21 | 0.16 | 0.35 | 0.05 | 0.015 |
| PAXX | S148 | 0.03 | 0.12 | 0.27 | -0.22 | 0.07 | -0.36 | 0.1 |
| PBRM1 | S948 | -0.43 | -0.59 | -0.72 | 0.46 | 0.60 | 0.13 | 0.004 |
| PBRM1 | S178 | -0.06 | -0.10 | -0.26 | -0.07 | -0.10 | 0.47 | 0.288 |
| PBRM1 | S1453 | -0.62 | -0.18 | 0.35 | 0.09 | -0.48 | 0.50 | 0.665 |
| PBRM1 | S1405 | 0.00 | -0.29 | 0.02 | -0.23 | 0.01 | 0.40 | 0.508 |
| PBRM1 | S1453 | -0.02 | -0.10 | -0.10 | 0.07 | 0.03 | 0.10 | 0.012 |
| PBRM1 | S501; T/S | -0.07 | -0.13 | -0.01 | 0.07 | -0.06 | 0.18 | 0.155 |
| PBRM1 | S505 | 0.04 | -0.02 | -0.17 | 0.13 | 0.05 | -0.05 | 0.306 |
| PBRM1 | S636 | 0.08 | -0.13 | -0.07 | 0.10 | 0.03 | -0.02 | 0.345 |
| PBRM1 | S178 | 0.07 | -0.06 | -0.12 | -0.15 | -0.13 | 0.33 | 0.769 |
| PBRM1 | S1278 | 0.06 | 0.03 | -0.16 | 0.14 | -0.05 | -0.03 | 0.655 |
| PBRM1 | S636 | -0.03 | 0.01 | -0.03 | 0.03 | -0.03 | 0.04 | 0.315 |
| PBRM1 | T/S | 0.02 | -0.16 | 0.25 | 0.18 | -0.36 | -0.02 | 0.638 |
| PBRM1 | S131 | 0.26 | 0.20 | -0.33 | 0.11 | -0.01 | -0.34 | 0.623 |
| PBRM1 | S/Y | 0.02 | 0.21 | -0.04 | -0.25 | -0.16 | 0.15 | 0.354 |
| PBRM1 | S987 | 0.29 | 0.11 | -0.03 | -0.09 | -0.07 | -0.27 | 0.081 |
| PBXIP1 | S469 | 0.01 | -0.56 | -0.13 | 0.22 | 0.04 | 0.28 | 0.093 |
| PBXIP1 | S | 0.12 | 0.18 | 0.18 | 0.05 | -0.35 | -0.28 | 0.046 |
| PCBP1 | S27 | -0.10 | -0.29 | -0.28 | 0.26 | 0.14 | 0.18 | 0.004 |
| PCBP1 | S190 | 0.37 | 0.14 | 0.16 | -0.27 | 0.06 | -0.71 | 0.086 |
| PCBP1 | S190 | 0.23 | 0.35 | 0.28 | -0.30 | 0.03 | -1.01 | 0.082 |
| PCBP2 | S/T | -0.06 | -0.10 | -0.05 | -0.10 | 0.00 | 0.27 | 0.31 |
| PCDH8 | S | 0.39 | 0.47 | -0.07 | -0.14 | -0.35 | -0.60 | 0.042 |
| PCF11 | S/T | 0.01 | -0.12 | -0.02 | -0.10 | -0.02 | 0.22 | 0.487 |
| PCF11 | S511 | 0.08 | 0.02 | -0.04 | -0.02 | 0.21 | -0.29 | 0.749 |
| PCF11 | S705 | -0.12 | 0.02 | 0.16 | 0.05 | -0.22 | 0.08 | 0.7 |
| PCF11 | S511 | 0.12 | 0.04 | -0.11 | 0.20 | 0.08 | -0.42 | 0.756 |
| PCF11 | S705 | 0.19 | -0.12 | 0.06 | 0.05 | -0.07 | -0.13 | 0.438 |
| PCF11 | S705 | 0.24 | -0.25 | 0.12 | 0.18 | -0.18 | -0.19 | 0.631 |
| PCF11 | S509; S511 | -0.26 | -0.01 | 0.44 | -0.24 | -0.03 | -0.02 | 0.513 |
| PCF11 | S487 | 0.07 | 0.12 | 0.14 | -0.01 | -0.11 | -0.24 | 0.033 |
| PCF11 | S489; S494 | 0.19 | 0.31 | 0.04 | -0.34 | 0.08 | -0.41 | 0.079 |
| PCIF1 | S116 | 0.00 | 0.09 | -0.02 | 0.05 | -0.18 | 0.05 | 0.597 |
| PCM1 | S1891 | -0.90 | -0.65 | -0.56 | 0.57 | 0.36 | 0.46 | 6E-04 |
| PCM1 | S | -0.27 | -0.35 | -0.11 | 0.33 | 0.30 | -0.05 | 0.036 |
| PCM1 | S68; S69 | -0.24 | -0.19 | -0.04 | 0.21 | 0.06 | 0.15 | 0.016 |
| PCM1 | S65; S68; S69 | -0.07 | -0.35 | -0.07 | 0.11 | -0.06 | 0.34 | 0.122 |
| PCM1 | S643 | -0.13 | -0.03 | -0.15 | 0.13 | -0.16 | 0.28 | 0.232 |
| PCM1 | S1287 | -0.18 | -0.15 | 0.08 | 0.18 | -0.23 | 0.23 | 0.446 |
| PCM1 | S1776 | 0.04 | -0.05 | -0.02 | -0.07 | -0.02 | 0.11 | 0.717 |
| PCM1 | T/S/Y | -0.01 | 0.04 | -0.05 | -0.05 | 0.14 | -0.08 | 0.928 |
| PCM1 | S1304 | -0.02 | -0.04 | 0.06 | 0.11 | -0.12 | 0.01 | 0.957 |
| PCM1 | S2023 | 0.00 | -0.13 | 0.10 | 0.01 | -0.26 | 0.22 | 0.996 |
| PCM1 | T/S/Y | -0.11 | 0.06 | 0.06 | 0.00 | -0.02 | 0.01 | 0.954 |
| PCM1 | T530; S/Y/T | -0.21 | 0.00 | 0.24 | -0.25 | -0.08 | 0.22 | 0.82 |
| PCM1 | S960 | 0.21 | -0.17 | 0.04 | 0.03 | -0.19 | 0.04 | 0.651 |
| PCM1 | S815 | -0.23 | 0.21 | 0.08 | 0.30 | -0.39 | -0.10 | 0.732 |

| | | | | | | | | |
|---|---|---|---|---|---|---|---|---|
| **PCM1** | **S1923** | -0.13 | 0.16 | 0.09 | -0.02 | -0.13 | 0.01 | 0.416 |
| **PCM1** | **S977** | -0.58 | 0.14 | 0.46 | -0.01 | -0.02 | -0.21 | 0.791 |
| **PCM1** | **S372** | 0.01 | 0.05 | 0.08 | 0.01 | -0.21 | 0.04 | 0.289 |
| **PCM1** | **S861** | 0.05 | 0.10 | 0.09 | -0.21 | 0.04 | -0.09 | 0.084 |
| **PCM1** | **S180** | -0.17 | 0.32 | -0.02 | 0.05 | 0.33 | -0.78 | 0.654 |
| **PCM1** | **S65** | -0.23 | 0.18 | 0.27 | 0.13 | -0.29 | -0.16 | 0.41 |
| **PCM1** | **S1730** | 0.11 | 0.10 | 0.06 | 0.03 | 0.00 | -0.34 | 0.175 |
| **PCM1** | **S1335** | -0.05 | 0.28 | 0.06 | -0.15 | 0.08 | -0.28 | 0.217 |
| **PCM1** | **S991** | 0.15 | 0.10 | 0.06 | -0.12 | 0.01 | -0.23 | 0.042 |
| **PCM1** | **S537** | 0.42 | 0.14 | -0.30 | 0.02 | -0.18 | -0.24 | 0.388 |
| **PCM1** | **S1776** | 0.11 | 0.13 | 0.12 | -0.11 | -0.07 | -0.23 | 0.005 |
| **PCM1** | **S960** | -0.01 | 0.10 | 0.27 | 0.04 | -0.14 | -0.34 | 0.126 |
| **PCM1** | **S580** | 0.03 | 0.26 | 0.10 | -0.03 | -0.25 | -0.17 | 0.039 |
| **PCM1** | **S537** | 0.13 | 0.08 | 0.22 | -0.10 | -0.08 | -0.31 | 0.022 |
| **PCM1** | **S65; S69** | 0.06 | 0.10 | 0.28 | -0.11 | -0.18 | -0.22 | 0.013 |
| **PCM1** | **S1322** | 0.17 | 0.21 | 0.10 | -0.20 | -0.28 | -0.05 | 0.011 |
| **PCM1** | **S168** | 0.23 | 0.21 | 0.14 | -0.12 | -0.24 | -0.31 | 0.003 |
| **PCM1** | **S1322; S1323** | 0.19 | 0.22 | 0.20 | -0.12 | -0.25 | -0.35 | 0.003 |
| **PCM1** | **S65** | 0.10 | 0.24 | 0.30 | -0.27 | -0.13 | -0.36 | 0.007 |
| **PCM1** | **S110** | 0.18 | 0.13 | 0.31 | -0.06 | -0.19 | -0.53 | 0.035 |
| **PCM1** | **S872** | 0.15 | 0.16 | 0.34 | -0.26 | -0.14 | -0.39 | 0.007 |
| **PCM1** | **S119** | 0.29 | 0.24 | 0.27 | -0.15 | -0.29 | -0.55 | 0.007 |
| **PCM1** | **S1435** | 0.43 | 0.21 | 0.36 | -0.46 | -0.46 | -0.40 | 3E-04 |
| **PCM1** | **S1765** | 0.27 | 0.23 | 0.48 | -0.57 | -0.06 | -0.75 | 0.024 |
| **PCM1** | **S1260** | 0.26 | 0.45 | 0.35 | -0.37 | -0.30 | -0.79 | 0.007 |
| **PCM1** | **S1257** | 0.32 | 0.42 | 0.25 | -0.26 | -0.06 | -1.24 | 0.081 |
| **PCNP** | **S119** | -0.17 | -0.21 | -0.08 | 0.03 | 0.29 | 0.08 | 0.029 |
| **PCNP** | **S119** | 0.00 | -0.23 | 0.11 | -0.04 | 0.18 | -0.05 | 0.599 |
| **PCNP** | **S53** | 0.08 | 0.07 | -0.05 | -0.05 | 0.24 | -0.35 | 0.652 |
| **PCNP** | **S119** | 0.30 | -0.31 | 0.10 | 0.32 | -1.03 | 0.21 | 0.698 |
| **PCNP** | **S119** | 0.09 | 0.18 | 0.17 | 0.26 | -0.70 | -0.22 | 0.26 |
| **PCNT** | **S1653** | -0.19 | -0.38 | -0.33 | 0.14 | 0.26 | 0.33 | 0.002 |
| **PCNT** | **T** | -0.41 | -0.31 | -0.07 | -0.20 | 0.00 | 0.69 | 0.212 |
| **PCNT** | **T/S** | -0.14 | -1.02 | 0.33 | 0.03 | 0.08 | 0.33 | 0.357 |
| **PCNT** | **T/S** | -0.14 | -1.02 | 0.33 | 0.03 | 0.08 | 0.33 | 0.357 |
| **PCNT** | **S2433** | 0.02 | -0.19 | -0.50 | -0.11 | 0.50 | 0.07 | 0.181 |
| **PCNT** | **S1703** | -0.09 | -0.28 | -0.20 | 0.12 | 0.12 | 0.25 | 0.007 |
| **PCNT** | **S2352** | 0.05 | -0.59 | 0.09 | 0.19 | 0.24 | -0.14 | 0.389 |
| **PCNT** | **S2352; T2367** | -0.12 | 0.21 | -0.37 | -0.02 | -0.13 | 0.32 | 0.53 |
| **PCNT** | **S3274** | 0.11 | -0.07 | -0.04 | -0.02 | 0.07 | -0.05 | 0.961 |
| **PCNT** | **S1632** | 0.00 | 0.01 | 0.02 | -0.10 | 0.00 | 0.06 | 0.61 |
| **PCNT** | **S1630** | 0.24 | -0.21 | 0.01 | -0.46 | 0.21 | 0.09 | 0.798 |
| **PCNT** | **S/T** | -0.46 | 0.42 | 0.08 | 0.18 | -0.52 | 0.08 | 0.782 |
| **PCNT** | **S2177** | 0.03 | 0.13 | 0.02 | -0.19 | 0.23 | -0.28 | 0.447 |
| **PCNT** | **S2177** | -0.05 | 0.03 | 0.19 | -0.54 | 0.18 | 0.07 | 0.548 |
| **PCNT** | **S/T** | 0.07 | 0.27 | 0.04 | -0.44 | -0.23 | 0.17 | 0.203 |
| **PCNX1** | **S117** | 0.15 | 0.33 | 0.06 | -0.01 | -0.04 | -0.67 | 0.143 |
| **PCNX3** | **S246** | -0.10 | -0.31 | -0.16 | 0.13 | 0.15 | 0.22 | 0.006 |

| | | | | | | | | |
|---|---|---|---|---|---|---|---|---|
| **PCNX3** | **S1909** | -0.15 | -0.15 | -0.16 | -0.02 | 0.06 | 0.34 | 0.064 |
| **PCNX3** | **S287** | 0.03 | 0.00 | -0.22 | 0.15 | 0.00 | 0.02 | 0.268 |
| **PCNX3** | **S/T** | -0.11 | 0.15 | -0.12 | -0.16 | 0.03 | 0.18 | 0.736 |
| **PCNX3** | **S287** | 0.13 | 0.00 | -0.12 | -0.05 | 0.08 | -0.06 | 0.864 |
| **PCNX3** | **S1955** | 0.19 | 0.15 | 0.05 | -0.17 | -0.20 | -0.06 | 0.01 |
| **PCNX3** | **S/T** | 0.01 | 0.27 | 0.13 | -0.18 | -0.07 | -0.21 | 0.027 |
| **PCNX3** | **S1955** | 0.16 | 0.39 | 0.06 | -0.17 | 0.07 | -0.74 | 0.134 |
| **PCYT1A** | **S315** | -0.14 | -0.03 | -0.36 | 0.15 | 0.35 | -0.08 | 0.118 |
| **PCYT1A** | **S329; S331** | -0.12 | -0.13 | -0.12 | 0.03 | 0.08 | 0.22 | 0.014 |
| **PCYT1A** | **S343; S347** | 0.03 | 0.08 | -0.07 | -0.16 | 0.04 | 0.05 | 0.677 |
| **PCYT1A** | **S343** | 0.06 | -0.07 | 0.11 | -0.04 | 0.26 | -0.39 | 0.667 |
| **PDAP1** | **S60; S63** | -0.21 | -0.20 | -0.05 | -0.07 | -0.33 | 0.63 | 0.475 |
| **PDAP1** | **S60; S63** | -0.02 | -0.11 | -0.07 | 0.06 | 0.04 | 0.09 | 0.014 |
| **PDAP1** | **S19** | 0.03 | 0.02 | -0.18 | -0.06 | 0.19 | -0.03 | 0.485 |
| **PDAP1** | **S60** | -0.04 | 0.36 | 0.19 | -0.37 | -0.11 | -0.14 | 0.055 |
| **PDAP1** | **S/Y** | 0.22 | 0.23 | 0.16 | -0.28 | -0.26 | -0.17 | 5E-04 |
| **PDCD11** | **S/T** | -0.12 | -0.13 | -0.06 | -0.01 | 0.07 | 0.21 | 0.046 |
| **PDCD11** | **S1498** | -0.04 | -0.05 | 0.01 | -0.02 | -0.01 | 0.11 | 0.297 |
| **PDCD11** | **S1360** | 0.28 | 0.12 | 0.10 | 0.09 | -0.34 | -0.38 | 0.08 |
| **PDCD11** | **S/T** | 0.20 | 0.18 | 0.36 | -0.04 | -0.34 | -0.57 | 0.026 |
| **PDCD11** | **S438** | 0.11 | 0.09 | 0.54 | -0.37 | -0.05 | -0.58 | 0.051 |
| **PDCD2L** | **S20** | -0.14 | -0.13 | -0.09 | -0.08 | -0.24 | 0.53 | 0.472 |
| **PDE4DIP** | **S1205; S/T** | -0.14 | -0.01 | 0.05 | 0.12 | -0.02 | -0.01 | 0.433 |
| **PDHA1** | **S232** | -0.23 | -0.04 | -0.24 | 0.31 | 0.28 | -0.20 | 0.17 |
| **PDHA1** | **S293; S300** | -0.15 | -0.19 | 0.11 | -0.32 | 0.07 | 0.37 | 0.634 |
| **PDHA1** | **S293** | -0.08 | 0.03 | 0.35 | 0.07 | 0.10 | -0.66 | 0.404 |
| **PDHA1** | **S293** | 0.00 | 0.24 | 0.36 | -0.05 | 0.14 | -1.08 | 0.247 |
| **PDIA6** | **S428** | 0.22 | 0.15 | 0.02 | -0.12 | 0.01 | -0.33 | 0.074 |
| **PDPK1** | **S241** | -0.08 | 0.13 | -0.19 | 0.08 | -0.01 | 0.05 | 0.432 |
| **PDS5A** | **S1305** | -0.05 | -0.12 | 0.06 | 0.18 | -0.41 | 0.25 | 0.852 |
| **PDS5A** | **S1305** | -0.04 | 0.00 | -0.03 | 0.06 | -0.23 | 0.20 | 0.801 |
| **PDS5A** | **T1208** | 0.19 | 0.38 | -0.02 | -0.10 | -0.07 | -0.53 | 0.091 |
| **PDS5B** | **S/Y/T** | -0.14 | -0.33 | -0.53 | 0.33 | 0.36 | 0.10 | 0.013 |
| **PDS5B** | **S1176** | -0.12 | -0.48 | -0.36 | 0.30 | 0.15 | 0.31 | 0.008 |
| **PDS5B** | **S1407; S1416; S1417** | -0.21 | -0.30 | -0.28 | -0.09 | 0.03 | 0.62 | 0.11 |
| **PDS5B** | **S1176** | -0.12 | -0.25 | -0.42 | 0.31 | 0.50 | -0.25 | 0.137 |
| **PDS5B** | **S1165; S1166** | -0.74 | -0.05 | 0.04 | 0.03 | 0.07 | 0.42 | 0.199 |
| **PDS5B** | **S1166; S/T** | -0.97 | 0.05 | 0.13 | 0.36 | -0.12 | 0.21 | 0.341 |
| **PDS5B** | **S1406** | -0.12 | -0.21 | -0.16 | 0.13 | -0.06 | 0.34 | 0.062 |
| **PDS5B** | **S1257; S1259** | -0.12 | -0.18 | -0.14 | 0.14 | 0.03 | 0.22 | 0.009 |
| **PDS5B** | **S1416; S1417; S/T** | -0.21 | -0.05 | -0.15 | 0.29 | 0.14 | -0.08 | 0.098 |
| **PDS5B** | **S1162; S1166** | -0.31 | -0.11 | 0.02 | -0.06 | -0.04 | 0.40 | 0.266 |
| **PDS5B** | **S1358** | -0.11 | -0.03 | -0.16 | 0.12 | -0.18 | 0.30 | 0.279 |
| **PDS5B** | **S1358; T1370** | 0.04 | -0.02 | -0.26 | 0.20 | -0.04 | 0.04 | 0.266 |
| **PDS5B** | **S1358** | -0.02 | -0.04 | -0.12 | 0.02 | -0.09 | 0.23 | 0.304 |
| **PDS5B** | **S1283** | -0.02 | -0.04 | -0.09 | 0.13 | -0.13 | 0.13 | 0.344 |
| **PDS5B** | **S1358; S1366** | 0.06 | -0.10 | -0.07 | 0.00 | -0.04 | 0.14 | 0.365 |
| **PDS5B** | **S1283** | 0.06 | -0.34 | 0.16 | 0.14 | 0.02 | -0.10 | 0.73 |

| | | | | | | | | |
|---|---|---|---|---|---|---|---|---|
| **PDS5B** | **T/S** | -0.04 | -0.08 | 0.02 | -0.02 | -0.12 | 0.22 | 0.588 |
| **PDS5B** | **S1417; S/T** | 0.05 | -0.08 | -0.04 | -0.13 | -0.02 | 0.20 | 0.709 |
| **PDS5B** | **S1379; S1383** | 0.01 | -0.02 | -0.02 | 0.02 | -0.06 | 0.07 | 0.642 |
| **PDS5B** | **S1191** | -0.02 | -0.05 | 0.03 | -0.19 | 0.01 | 0.19 | 0.911 |
| **PDS5B** | **S1087** | -0.23 | 0.15 | 0.04 | -0.05 | -0.09 | 0.15 | 0.93 |
| **PDS5B** | **S1283** | 0.10 | 0.03 | -0.13 | 0.00 | 0.09 | -0.11 | 0.952 |
| **PDS5B** | **S1406** | -0.01 | 0.11 | -0.09 | -0.06 | -0.06 | 0.10 | 0.897 |
| **PDS5B** | **S1166** | -0.18 | 0.23 | -0.08 | -0.28 | -0.04 | 0.25 | 0.943 |
| **PDS5B** | **S1283** | 0.02 | 0.06 | -0.05 | -0.01 | -0.03 | 0.01 | 0.594 |
| **PDS5B** | **T1220** | 0.02 | 0.06 | -0.02 | -0.07 | 0.02 | -0.01 | 0.373 |
| **PDS5B** | **S1176** | -0.03 | 0.04 | 0.03 | 0.00 | 0.20 | -0.28 | 0.791 |
| **PDS5B** | **S1283** | 0.07 | 0.07 | -0.08 | -0.09 | 0.04 | -0.02 | 0.514 |
| **PDS5B** | **S1257** | 0.05 | -0.05 | 0.11 | -0.01 | -0.17 | 0.06 | 0.402 |
| **PDS5B** | **S1406; S/T** | 0.05 | 0.02 | 0.07 | -0.05 | -0.15 | 0.05 | 0.183 |
| **PDS5B** | **S1283** | 0.09 | 0.08 | -0.03 | -0.15 | -0.02 | 0.02 | 0.213 |
| **PDS5B** | **S1257** | 0.04 | 0.04 | 0.15 | -0.02 | -0.18 | -0.05 | 0.065 |
| **PDS5B** | **S1383** | 0.12 | 0.15 | -0.05 | -0.11 | 0.03 | -0.17 | 0.147 |
| **PDS5B** | **S1182** | 0.12 | 0.21 | -0.11 | -0.07 | 0.08 | -0.28 | 0.311 |
| **PDS5B** | **S1166** | 0.08 | -0.02 | 0.18 | 0.04 | -0.07 | -0.23 | 0.158 |
| **PDS5B** | **S1283** | 0.08 | 0.07 | 0.10 | -0.11 | -0.04 | -0.11 | 0.003 |
| **PDS5B** | **S/T** | -0.10 | 0.03 | 0.18 | 0.19 | 0.29 | -0.87 | 0.681 |
| **PDS5B** | **S1166** | 0.07 | 0.13 | 0.18 | -0.19 | -0.03 | -0.20 | 0.014 |
| **PDS5B** | **S1166** | -0.04 | 0.21 | 0.21 | -0.27 | 0.02 | -0.19 | 0.085 |
| **PDS5B** | **S1182** | 0.13 | 0.17 | 0.18 | -0.15 | 0.02 | -0.43 | 0.057 |
| **PDS5B** | **S1182** | 0.16 | 0.14 | 0.23 | -0.20 | -0.07 | -0.36 | 0.012 |
| **PDS5B** | **S1182** | 0.14 | 0.18 | 0.20 | -0.11 | -0.01 | -0.52 | 0.069 |
| **PDS5B** | **S/Y/T** | 0.25 | 0.17 | 0.14 | -0.34 | -0.04 | -0.29 | 0.014 |
| **PDS5B** | **S1166** | 0.15 | 0.19 | 0.22 | -0.33 | -0.01 | -0.34 | 0.019 |
| **PDXDC1** | **S387** | 0.00 | -0.32 | -0.11 | 0.09 | 0.15 | 0.13 | 0.045 |
| **PDZD2** | **S/T** | -0.06 | -0.06 | -0.47 | -0.14 | -0.37 | 0.75 | 0.492 |
| **PDZD2** | **S/T** | -0.15 | -0.05 | -0.28 | -0.06 | -0.29 | 0.61 | 0.427 |
| **PDZD2** | **T/S** | -0.19 | -0.19 | 0.05 | 0.15 | -0.10 | 0.22 | 0.187 |
| **PDZD2** | **S2037** | -0.09 | -0.06 | 0.11 | 0.23 | -0.19 | -0.04 | 0.924 |
| **PDZD2** | **S1270** | 0.01 | 0.09 | 0.16 | -0.34 | 0.33 | -0.39 | 0.407 |
| **PDZD2** | **S1270** | -0.14 | 0.11 | 0.33 | -0.14 | 0.25 | -0.59 | 0.409 |
| **PEA15** | **S116** | 0.19 | 0.21 | -0.04 | -0.10 | -0.13 | -0.18 | 0.035 |
| **PELP1** | **T1014** | -0.18 | -0.32 | -0.46 | 0.47 | -0.15 | 0.38 | 0.058 |
| **PELP1** | **S1033** | -0.40 | -0.13 | -0.31 | 0.24 | 0.08 | 0.36 | 0.01 |
| **PELP1** | **S1043; T1069** | -0.19 | -0.21 | -0.24 | 0.34 | -0.02 | 0.21 | 0.023 |
| **PELP1** | **S1043; T1069** | -0.60 | -0.11 | 0.06 | 0.35 | -0.16 | 0.27 | 0.217 |
| **PELP1** | **T1090** | -0.19 | -0.04 | -0.27 | 0.15 | 0.05 | 0.24 | 0.024 |
| **PELP1** | **T** | -0.32 | -0.01 | -0.15 | 0.18 | 0.04 | 0.20 | 0.044 |
| **PELP1** | **S/T** | -0.11 | -0.08 | -0.10 | 0.14 | -0.12 | 0.23 | 0.169 |
| **PELP1** | **S1043** | -0.28 | -0.09 | 0.19 | -0.01 | 0.35 | -0.27 | 0.737 |
| **PELP1** | **S481** | 0.12 | -0.05 | -0.16 | 0.17 | -0.11 | 0.01 | 0.695 |
| **PELP1** | **S734; S/T** | 0.01 | -0.04 | 0.12 | -0.25 | -0.05 | 0.18 | 0.63 |
| **PELP1** | **T745** | 0.03 | 0.02 | 0.07 | -0.04 | -0.02 | -0.06 | 0.014 |
| **PELP1** | **S1043** | 0.20 | 0.16 | -0.01 | -0.09 | -0.02 | -0.29 | 0.066 |

| | | | | | | | | |
|---|---|---|---|---|---|---|---|---|
| PER1 | S592 | -0.48 | 0.13 | -0.06 | 0.40 | 0.10 | -0.26 | 0.456 |
| PER2 | S977 | 0.17 | -0.08 | -0.09 | 0.07 | -0.13 | 0.04 | 0.925 |
| PES1 | S307 | -0.41 | -0.34 | -0.29 | 0.54 | 0.09 | 0.16 | 0.014 |
| PES1 | T302 | -0.38 | -0.18 | -0.28 | 0.33 | 0.03 | 0.32 | 0.011 |
| PEX1 | S1211 | 0.02 | -0.14 | 0.03 | 0.04 | 0.04 | 0.01 | 0.358 |
| PEX11G | T/S | -0.02 | 0.15 | 0.13 | -0.50 | 0.45 | -0.42 | 0.471 |
| PEX14 | S232 | 0.11 | -0.05 | 0.23 | -0.17 | -0.09 | -0.07 | 0.08 |
| PEX16 | S158 | -0.06 | 0.36 | 0.45 | -0.37 | -0.19 | -0.44 | 0.028 |
| PEX19 | S66 | -0.57 | 0.16 | 0.16 | 0.13 | -0.44 | 0.33 | 0.797 |
| PEX19 | S147 | -0.19 | 0.15 | -0.03 | 0.40 | -0.40 | -0.07 | 0.995 |
| PGAM5 | S188 | 0.56 | 0.01 | -0.31 | -0.08 | -0.10 | -0.26 | 0.417 |
| PGAM5 | S80 | 0.24 | 0.17 | 0.17 | -0.19 | -0.14 | -0.35 | 0.003 |
| PGM3 | T/S | -0.28 | 0.03 | 0.07 | 0.06 | 0.32 | -0.29 | 0.679 |
| PGM3 | T/S | 0.01 | 0.11 | -0.32 | 0.18 | 0.33 | -0.46 | 0.774 |
| PGM3 | S64 | -0.07 | -0.21 | 0.16 | -0.03 | 0.15 | -0.03 | 0.588 |
| PGM3 | S64 | 0.26 | -0.22 | -0.20 | -0.51 | 0.24 | 0.26 | 0.879 |
| PGRMC1 | S57 | -0.64 | 0.19 | -0.13 | -0.04 | -0.15 | 0.51 | 0.395 |
| PGRMC1 | S57 | -0.13 | -0.11 | -0.09 | 0.09 | -0.07 | 0.27 | 0.107 |
| PGRMC1 | S181 | -0.36 | 0.12 | 0.37 | -0.16 | -0.37 | 0.23 | 0.639 |
| PGRMC1 | S181 | -0.33 | 0.11 | 0.36 | -0.17 | -0.31 | 0.20 | 0.603 |
| PGRMC1 | S181 | -0.04 | 0.06 | 0.30 | -0.26 | -0.15 | 0.02 | 0.147 |
| PGRMC1 | S181 | -0.09 | 0.12 | 0.31 | -0.28 | -0.02 | -0.11 | 0.139 |
| PGRMC2 | S104 | -0.83 | -0.71 | -0.87 | 0.51 | 0.57 | 0.45 | 3E-05 |
| PGRMC2 | S90 | -0.10 | -0.15 | 0.04 | 0.11 | 0.07 | 0.01 | 0.099 |
| PGRMC2 | T211 | 0.13 | 0.11 | 0.14 | -0.22 | -0.17 | -0.03 | 0.01 |
| PGRMC2 | T211 | 0.23 | 0.19 | 0.04 | -0.25 | -0.18 | -0.10 | 0.01 |
| PGRMC2 | S/T/Y | 0.20 | 0.30 | 0.11 | -0.24 | -0.24 | -0.25 | 0.001 |
| PGRMC2 | T211 | 0.37 | 0.28 | 0.22 | -0.38 | -0.21 | -0.52 | 0.003 |
| PGRMC2 | T211 | 0.34 | 0.35 | 0.19 | -0.36 | -0.22 | -0.56 | 0.004 |
| PHACTR4 | S118 | -0.25 | 0.26 | 0.18 | 0.06 | -0.28 | -0.05 | 0.468 |
| PHACTR4 | S118 | -0.25 | 0.26 | 0.18 | 0.06 | -0.28 | -0.05 | 0.468 |
| PHB1 | S213 | -0.03 | 0.03 | 0.11 | -0.03 | -0.17 | 0.08 | 0.398 |
| PHB2 | S291 | 0.01 | -0.05 | -0.10 | 0.13 | -0.13 | 0.13 | 0.38 |
| PHB2 | S91 | 0.23 | 0.04 | 0.07 | -0.25 | 0.05 | -0.21 | 0.085 |
| PHC1 | S786 | -0.02 | -0.22 | -0.14 | -0.15 | 0.24 | 0.22 | 0.168 |
| PHC1 | S664 | -0.11 | -0.12 | -0.12 | 0.12 | -0.13 | 0.29 | 0.158 |
| PHC1 | S895 | 0.02 | 0.04 | -0.02 | -0.08 | -0.07 | 0.10 | 0.612 |
| PHC1 | S786 | 0.31 | 0.31 | -0.05 | -0.25 | -0.13 | -0.34 | 0.035 |
| PHC2 | T/S/Y | 0.02 | 0.01 | -0.02 | 0.07 | -0.18 | 0.09 | 0.901 |
| PHC2 | T617 | 0.04 | 0.07 | -0.03 | -0.10 | -0.05 | 0.06 | 0.353 |
| PHC2 | T617 | 0.10 | 0.18 | 0.01 | -0.22 | 0.05 | -0.15 | 0.096 |
| PHC3 | S842 | -0.19 | -0.24 | -0.32 | 0.08 | 0.10 | 0.43 | 0.019 |
| PHC3 | S272 | 0.03 | -0.26 | 0.00 | 0.07 | -0.16 | 0.26 | 0.421 |
| PHC3 | S263 | -0.01 | -0.01 | -0.19 | 0.14 | -0.03 | 0.07 | 0.159 |
| PHC3 | S761; S762 | -0.05 | 0.09 | 0.05 | -0.06 | 0.00 | -0.04 | 0.251 |
| PHC3 | S761; S762; T764 | 0.15 | -0.18 | 0.05 | -0.10 | 0.38 | -0.44 | 0.821 |
| PHC3 | S272 | 0.06 | 0.02 | 0.06 | -0.12 | -0.04 | 0.02 | 0.089 |
| PHC3 | S761; S762 | 0.11 | 0.10 | 0.08 | -0.05 | -0.01 | -0.27 | 0.064 |

| | | | | | | | | |
|---|---|---|---|---|---|---|---|---|
| **PHC3** | **S/T** | 0.08 | -0.04 | 0.35 | -0.48 | 0.01 | -0.04 | 0.195 |
| **PHC3** | **S/T** | 0.08 | 0.16 | 0.19 | -0.12 | -0.04 | -0.33 | 0.029 |
| **PHC3** | **T609; S616** | 0.01 | 0.12 | 0.30 | -0.38 | 0.11 | -0.27 | 0.132 |
| **PHC3** | **T609; S616** | 0.15 | 0.22 | 0.32 | -0.34 | 0.05 | -0.63 | 0.058 |
| **PHC3** | **T609** | 0.00 | 0.52 | 0.22 | -0.75 | -0.08 | -0.21 | 0.081 |
| **PHF1** | **S522** | -0.02 | -0.01 | -0.20 | 0.20 | -0.05 | 0.06 | 0.187 |
| **PHF1** | **S515** | 0.19 | 0.06 | 0.10 | -0.12 | -0.09 | -0.16 | 0.005 |
| **PHF10** | **S270** | -0.01 | -0.07 | -0.24 | 0.19 | -0.05 | 0.13 | 0.127 |
| **PHF10** | **S56** | 0.41 | 0.35 | -0.01 | -0.24 | -0.25 | -0.48 | 0.02 |
| **PHF12** | **T671** | -0.60 | -0.10 | 0.10 | 0.00 | -0.07 | 0.47 | 0.281 |
| **PHF12** | **S/Y/T** | -0.27 | -0.04 | 0.00 | 0.16 | -0.13 | 0.22 | 0.242 |
| **PHF12** | **T880** | -0.04 | -0.08 | 0.01 | 0.02 | -0.08 | 0.15 | 0.428 |
| **PHF12** | **S554** | -0.11 | 0.17 | -0.12 | 0.17 | 0.03 | -0.18 | 0.868 |
| **PHF12** | **T671** | 0.07 | -0.31 | 0.27 | -0.15 | 0.03 | 0.03 | 0.826 |
| **PHF12** | **T130; S134** | 0.11 | 0.03 | 0.13 | -0.16 | 0.00 | -0.13 | 0.037 |
| **PHF12** | **S131** | 0.10 | 0.20 | 0.13 | -0.40 | 0.10 | -0.22 | 0.1 |
| **PHF12** | **S691** | 0.29 | 0.16 | 0.16 | -0.32 | -0.09 | -0.33 | 0.008 |
| **PHF12** | **S662** | 0.28 | 0.63 | 0.38 | -0.92 | -0.08 | -1.14 | 0.027 |
| **PHF14** | **S84** | -1.07 | -1.07 | -0.82 | 0.55 | 0.73 | 0.45 | 2E-04 |
| **PHF14** | **S936** | -0.71 | -1.10 | -0.99 | 0.25 | 0.72 | 0.67 | 0.001 |
| **PHF14** | **S936** | -0.90 | -0.63 | -0.82 | 0.25 | 0.68 | 0.54 | 0.001 |
| **PHF14** | **T/S** | -0.03 | -0.45 | -0.36 | 0.19 | 0.35 | 0.13 | 0.023 |
| **PHF14** | **S835** | -0.09 | -0.09 | -0.33 | 0.14 | 0.12 | 0.18 | 0.017 |
| **PHF14** | **T155; S/T** | -0.33 | -0.16 | 0.15 | 0.05 | 0.07 | 0.16 | 0.228 |
| **PHF14** | **S602** | -0.14 | -0.22 | 0.06 | 0.21 | -0.12 | 0.16 | 0.234 |
| **PHF14** | **T152; T/S** | -0.11 | -0.24 | 0.12 | -0.02 | 0.01 | 0.20 | 0.316 |
| **PHF14** | **S57; S59** | -0.18 | -0.13 | 0.17 | -0.09 | -0.14 | 0.29 | 0.733 |
| **PHF14** | **S290** | -0.10 | -0.06 | 0.05 | -0.02 | -0.05 | 0.16 | 0.493 |
| **PHF14** | **T287; S290; S294** | -0.06 | -0.10 | 0.14 | -0.03 | -0.04 | 0.07 | 0.966 |
| **PHF14** | **S530** | 0.03 | -0.03 | 0.00 | -0.07 | 0.09 | -0.02 | 0.949 |
| **PHF14** | **S/T/Y** | -0.39 | -0.02 | 0.32 | 0.45 | -0.34 | -0.24 | 0.971 |
| **PHF14** | **S/T/Y** | -0.02 | -0.10 | 0.13 | -0.03 | -0.11 | 0.10 | 0.847 |
| **PHF14** | **S308** | -0.08 | -0.01 | 0.11 | 0.11 | 0.04 | -0.20 | 0.852 |
| **PHF14** | **S298** | 0.05 | 0.04 | 0.02 | -0.07 | 0.06 | -0.11 | 0.225 |
| **PHF14** | **T171** | 0.10 | 0.08 | -0.05 | -0.21 | -0.01 | 0.07 | 0.388 |
| **PHF14** | **T287** | 0.05 | 0.01 | 0.09 | -0.02 | -0.14 | 0.00 | 0.103 |
| **PHF14** | **T287; S290** | 0.04 | -0.01 | 0.11 | -0.10 | 0.14 | -0.20 | 0.398 |
| **PHF14** | **S91** | 0.06 | 0.10 | -0.02 | -0.37 | -0.01 | 0.18 | 0.539 |
| **PHF14** | **S530** | 0.18 | 0.12 | -0.03 | -0.02 | 0.19 | -0.56 | 0.394 |
| **PHF14** | **S/T** | 0.10 | -0.04 | 0.31 | -0.19 | -0.23 | -0.03 | 0.08 |
| **PHF14** | **S/T** | 0.10 | -0.04 | 0.31 | -0.19 | -0.23 | -0.03 | 0.08 |
| **PHF14** | **S/T/Y** | 0.18 | 0.10 | 0.13 | -0.15 | -0.10 | -0.21 | 0.002 |
| **PHF2** | **S899** | -0.22 | -0.41 | -0.11 | 0.44 | 0.22 | -0.08 | 0.067 |
| **PHF2** | **S705** | -0.17 | -0.37 | -0.13 | 0.02 | 0.18 | 0.35 | 0.026 |
| **PHF2** | **S458; S459** | -0.24 | -0.17 | -0.08 | 0.17 | -0.12 | 0.35 | 0.11 |
| **PHF2** | **S539; S/T** | -0.06 | -0.39 | 0.01 | 0.16 | -0.03 | 0.24 | 0.138 |
| **PHF2** | **S853** | -0.10 | -0.21 | 0.00 | 0.08 | 0.02 | 0.17 | 0.065 |
| **PHF2** | **S474** | 0.02 | -0.09 | -0.23 | 0.16 | -0.02 | 0.13 | 0.1 |

| | | | | | | | | |
|---|---|---|---|---|---|---|---|---|
| PHF2 | S/T | -0.01 | -0.03 | -0.14 | -0.14 | 0.00 | 0.28 | 0.448 |
| PHF2 | T479 | 0.02 | 0.02 | -0.20 | 0.06 | -0.03 | 0.12 | 0.3 |
| PHF2 | S/T | -0.02 | 0.06 | -0.19 | 0.16 | -0.02 | -0.01 | 0.387 |
| PHF2 | S539 | -0.07 | 0.00 | -0.06 | 0.01 | 0.16 | -0.05 | 0.266 |
| PHF2 | S1059 | 0.00 | -0.05 | -0.02 | 0.01 | -0.02 | 0.08 | 0.199 |
| PHF2 | S681 | 0.09 | 0.00 | -0.09 | 0.01 | -0.04 | 0.01 | 0.943 |
| PHF2 | S458 | -0.10 | 0.01 | 0.12 | -0.14 | -0.06 | 0.15 | 0.827 |
| PHF2 | S853 | 0.02 | -0.06 | 0.11 | -0.05 | -0.01 | -0.02 | 0.38 |
| PHF2 | S474 | 0.30 | 0.03 | -0.19 | -0.18 | -0.03 | 0.02 | 0.517 |
| PHF2 | S882 | 0.10 | 0.13 | -0.06 | -0.10 | 0.00 | -0.09 | 0.143 |
| PHF2 | S905 | 0.11 | 0.02 | 0.06 | 0.05 | -0.17 | -0.08 | 0.143 |
| PHF2 | S/Y | -0.33 | 0.47 | 0.04 | -0.26 | 0.07 | -0.15 | 0.531 |
| PHF2 | S681 | 0.08 | 0.10 | 0.10 | -0.02 | 0.00 | -0.30 | 0.114 |
| PHF2 | S853 | 0.21 | -0.10 | 0.15 | 0.34 | -0.34 | -0.41 | 0.433 |
| PHF2 | S905 | 0.14 | 0.15 | 0.11 | -0.16 | -0.06 | -0.22 | 0.004 |
| PHF2 | S625 | 0.30 | 0.15 | -0.06 | -0.28 | 0.04 | -0.23 | 0.114 |
| PHF2 | S733; S734; S738 | -0.05 | 0.33 | 0.22 | -0.19 | -0.23 | -0.17 | 0.032 |
| PHF20 | S248 | -0.19 | -0.20 | -0.23 | 0.06 | 0.07 | 0.39 | 0.026 |
| PHF20 | S880 | -0.15 | -0.05 | 0.06 | -0.05 | 0.00 | 0.17 | 0.381 |
| PHF20 | S902 | 0.05 | -0.01 | -0.07 | -0.05 | -0.18 | 0.23 | 0.934 |
| PHF20 | S159 | 0.20 | 0.08 | 0.08 | -0.16 | -0.03 | -0.21 | 0.018 |
| PHF20 | S488 | 0.13 | 0.04 | 0.20 | -0.14 | -0.10 | -0.16 | 0.006 |
| PHF20 | S/T | 0.07 | -0.03 | 0.35 | -0.03 | -0.25 | -0.19 | 0.095 |
| PHF20 | S902 | 0.04 | 0.11 | 0.43 | -0.13 | -0.19 | -0.40 | 0.04 |
| PHF20L1 | S881; S888; S889 | -0.60 | -0.21 | -0.28 | -0.01 | -0.65 | 1.02 | 0.386 |
| PHF20L1 | S433 | 0.25 | 0.13 | 0.01 | -0.22 | -0.03 | -0.19 | 0.041 |
| PHF20L1 | S395 | 0.20 | 0.28 | 0.34 | -0.46 | 0.08 | -0.77 | 0.06 |
| PHF23 | S74 | 0.13 | 0.03 | -0.10 | 0.05 | -0.21 | 0.08 | 0.683 |
| PHF23 | S | 0.01 | 0.15 | 0.17 | -0.14 | -0.28 | 0.03 | 0.081 |
| PHF3 | S1614 | -0.45 | -0.56 | 0.26 | -0.18 | -0.01 | 0.61 | 0.33 |
| PHF3 | S125 | 0.01 | -0.17 | 0.03 | 0.01 | -0.10 | 0.19 | 0.527 |
| PHF3 | S283 | 0.12 | 0.00 | -0.14 | 0.08 | -0.18 | 0.08 | 0.99 |
| PHF3 | S1642 | 0.15 | 0.03 | -0.04 | 0.02 | -0.15 | -0.04 | 0.229 |
| PHF3 | S1133 | 0.12 | 0.19 | 0.05 | -0.15 | 0.05 | -0.33 | 0.089 |
| PHF3 | S1133 | 0.17 | 0.09 | 0.10 | 0.08 | -0.20 | -0.30 | 0.085 |
| PHF3 | S1614 | -0.03 | -0.08 | 0.51 | -0.17 | -0.01 | -0.37 | 0.213 |
| PHF3 | S1614 | 0.06 | 0.23 | 0.23 | -0.27 | -0.01 | -0.33 | 0.029 |
| PHF6 | S155 | -0.01 | -0.11 | -0.13 | 0.21 | -0.22 | 0.21 | 0.366 |
| PHF6 | S155 | -0.07 | -0.03 | -0.07 | 0.13 | 0.00 | 0.04 | 0.044 |
| PHF6 | S204 | 0.03 | 0.02 | -0.01 | -0.18 | 0.05 | 0.08 | 0.76 |
| PHF8 | S854; S857 | -0.17 | -0.22 | -0.30 | 0.10 | 0.07 | 0.40 | 0.021 |
| PHF8 | S1021 | -0.29 | -0.06 | -0.27 | 0.18 | 0.23 | 0.12 | 0.009 |
| PHF8 | T886 | -0.05 | -0.33 | -0.13 | 0.00 | -0.01 | 0.41 | 0.133 |
| PHF8 | T705; T706 | -0.09 | -0.10 | -0.26 | 0.11 | 0.05 | 0.24 | 0.02 |
| PHF8 | S826; S840 | -0.33 | -0.13 | 0.06 | 0.10 | 0.32 | -0.12 | 0.242 |
| PHF8 | S857 | 0.06 | 0.00 | -0.06 | 0.02 | -0.04 | 0.03 | 0.901 |
| PHF8 | S880 | 0.03 | 0.09 | 0.23 | -0.18 | 0.09 | -0.32 | 0.131 |
| PHF8 | S804 | -0.30 | 0.04 | 0.60 | -0.05 | -0.06 | -0.47 | 0.362 |

| | | | | | | | | |
|---|---|---|---|---|---|---|---|---|
| PHF8 | S120 | -0.17 | 0.28 | 0.32 | -0.22 | -0.05 | -0.29 | 0.126 |
| PHF8 | S880 | 0.15 | 0.23 | 0.15 | -0.19 | 0.05 | -0.51 | 0.076 |
| PHIP | S1315 | -0.34 | -0.44 | -0.69 | 0.27 | 0.29 | 0.51 | 0.003 |
| PHIP | S1457 | -0.48 | -0.76 | -0.25 | 0.48 | 0.71 | -0.29 | 0.077 |
| PHIP | S1651 | -0.37 | -0.25 | -0.48 | 0.14 | 0.36 | 0.36 | 0.003 |
| PHIP | S1457 | -0.47 | -0.47 | -0.14 | 0.22 | 0.55 | 0.02 | 0.03 |
| PHIP | S1525 | -0.32 | -0.31 | -0.39 | 0.19 | 0.22 | 0.41 | 0.001 |
| PHIP | S1548 | -0.27 | -0.37 | -0.35 | 0.20 | 0.42 | 0.17 | 0.002 |
| PHIP | S1762 | -0.21 | -0.34 | -0.02 | 0.23 | 0.29 | -0.06 | 0.072 |
| PHIP | S1377 | -0.13 | -0.14 | -0.24 | 0.32 | -0.03 | 0.14 | 0.045 |
| PHIP | S692 | -0.07 | -0.30 | -0.11 | 0.12 | 0.32 | -0.04 | 0.08 |
| PHIP | S674; S683 | -0.27 | -0.15 | -0.02 | 0.09 | -0.03 | 0.30 | 0.09 |
| PHIP | S1762 | -0.03 | -0.15 | -0.18 | 0.08 | 0.08 | 0.17 | 0.013 |
| PHIP | S1377 | -0.15 | -0.11 | -0.11 | 0.26 | -0.11 | 0.16 | 0.111 |
| PHIP | S674 | -0.06 | -0.21 | -0.07 | -0.17 | 0.01 | 0.40 | 0.333 |
| PHIP | S879; S880; S881 | -0.11 | 0.04 | -0.20 | 0.25 | -0.04 | 0.01 | 0.226 |
| PHIP | S1281; S1283 | -0.07 | -0.09 | -0.07 | 0.04 | -0.02 | 0.19 | 0.074 |
| PHIP | S1377 | -0.03 | 0.11 | -0.14 | 0.04 | 0.09 | -0.09 | 0.689 |
| PHIP | S1783 | 0.02 | -0.08 | 0.06 | -0.05 | -0.08 | 0.13 | 0.975 |
| PHIP | S1651 | -0.19 | 0.21 | 0.05 | -0.34 | 0.00 | 0.20 | 0.725 |
| PHIP | S1243 | 0.29 | -0.02 | -0.02 | -0.10 | -0.07 | -0.11 | 0.157 |
| PHIP | T1480 | 0.08 | 0.00 | 0.22 | -0.25 | 0.02 | -0.11 | 0.104 |
| PHIP | S1783 | 0.08 | 0.07 | 0.24 | -0.21 | 0.05 | -0.30 | 0.076 |
| PHIP | S911 | 0.17 | 0.12 | 0.19 | -0.21 | 0.09 | -0.48 | 0.099 |
| PHIP | S/T | -0.07 | 0.20 | 0.36 | -0.10 | -0.47 | -0.06 | 0.104 |
| PHIP | S911 | 0.32 | 0.20 | -0.01 | -0.19 | -0.05 | -0.39 | 0.049 |
| PHIP | S1281 | 0.36 | 0.42 | 0.27 | -0.39 | -0.16 | -0.95 | 0.024 |
| PHIP | S1283 | 0.46 | 0.48 | 0.56 | -0.66 | -0.40 | -1.48 | 0.014 |
| PHKB | S701 | 0.02 | 0.08 | 0.01 | -0.21 | 0.20 | -0.15 | 0.514 |
| PHLDB2 | S204 | -0.20 | -0.02 | 0.50 | -0.09 | -0.38 | 0.03 | 0.387 |
| PHLDB2 | S334 | 0.22 | 0.00 | 0.14 | -0.07 | 0.00 | -0.35 | 0.107 |
| PHLDB3 | T499 | -0.30 | -0.41 | -0.48 | 0.36 | 0.17 | 0.39 | 0.001 |
| PHRF1 | S915; T917 | -0.42 | -0.37 | -0.07 | 0.10 | -0.07 | 0.59 | 0.095 |
| PHRF1 | S1165 | -0.04 | -0.23 | -0.40 | 0.16 | 0.05 | 0.33 | 0.039 |
| PHRF1 | S/T | -0.16 | 0.04 | -0.20 | -0.01 | -0.03 | 0.30 | 0.207 |
| PHRF1 | S936; S948; S955 | -0.19 | 0.04 | -0.18 | -0.21 | 0.22 | 0.24 | 0.305 |
| PHRF1 | S1413 | 0.18 | -0.01 | -0.49 | -0.32 | 0.61 | -0.27 | 0.778 |
| PHRF1 | S73; S76 | -0.08 | -0.10 | 0.00 | 0.10 | -0.27 | 0.28 | 0.58 |
| PHRF1 | S814 | 0.12 | -0.17 | -0.09 | 0.03 | 0.14 | -0.06 | 0.461 |
| PHRF1 | S1202 | -0.01 | -0.05 | -0.05 | -0.05 | -0.05 | 0.19 | 0.456 |
| PHRF1 | S936; S948 | -0.13 | 0.01 | 0.02 | 0.18 | -0.05 | -0.04 | 0.513 |
| PHRF1 | S1095 | -0.03 | -0.06 | 0.07 | 0.04 | -0.07 | 0.04 | 0.808 |
| PHRF1 | S1229 | 0.04 | 0.02 | -0.08 | 0.07 | -0.02 | -0.04 | 0.882 |
| PHRF1 | S101 | 0.12 | -0.11 | -0.03 | -0.04 | 0.00 | 0.04 | 0.934 |
| PHRF1 | S1359; S1360 | 0.05 | -0.01 | -0.05 | -0.07 | -0.08 | 0.14 | 0.976 |
| PHRF1 | S1124 | 0.07 | -0.05 | -0.01 | -0.05 | 0.06 | -0.03 | 0.871 |
| PHRF1 | S915 | 0.01 | 0.03 | -0.01 | 0.03 | -0.12 | 0.06 | 0.728 |
| PHRF1 | S936 | 0.02 | 0.07 | 0.00 | -0.01 | -0.06 | -0.02 | 0.053 |

| | | | | | | | | |
|---|---|---|---|---|---|---|---|---|
| PHRF1 | S991 | 0.00 | 0.14 | -0.02 | 0.03 | -0.04 | -0.13 | 0.271 |
| PHRF1 | S1360 | 0.11 | 0.01 | 0.02 | -0.02 | -0.14 | 0.01 | 0.167 |
| PHRF1 | S1371 | 0.08 | 0.11 | -0.05 | 0.03 | 0.04 | -0.24 | 0.371 |
| PHRF1 | S973 | 0.05 | 0.07 | 0.07 | -0.07 | 0.02 | -0.16 | 0.07 |
| PHRF1 | S1202 | 0.07 | 0.08 | 0.08 | -0.16 | 0.01 | -0.10 | 0.029 |
| PHRF1 | S973 | -0.07 | 0.06 | 0.25 | -0.13 | -0.08 | -0.06 | 0.144 |
| PHRF1 | S867 | 0.12 | 0.00 | 0.19 | -0.07 | -0.05 | -0.23 | 0.053 |
| PHRF1 | S1202 | 0.13 | 0.10 | 0.15 | -0.13 | -0.08 | -0.20 | 0.003 |
| PHRF1 | S845; S846 | 0.15 | 0.23 | -0.02 | -0.14 | 0.08 | -0.38 | 0.154 |
| PHRF1 | S1034 | 0.06 | 0.15 | 0.17 | -0.12 | -0.01 | -0.30 | 0.043 |
| PHRF1 | S1359; S1360; S1371 | 0.14 | 0.19 | 0.10 | -0.13 | -0.13 | -0.22 | 0.002 |
| PHRF1 | S814 | 0.44 | 0.23 | -0.34 | -0.67 | 0.29 | -0.27 | 0.42 |
| PHRF1 | S1360; S1371 | 0.19 | 0.14 | 0.16 | -0.09 | -0.14 | -0.33 | 0.011 |
| PHRF1 | T/S | 0.34 | 0.21 | -0.12 | -1.15 | -0.21 | 0.43 | 0.4 |
| PHRF1 | S/Y/T | 0.25 | 0.08 | 0.35 | -0.42 | -0.09 | -0.34 | 0.015 |
| PHRF1 | S867 | 0.36 | 0.28 | 0.34 | -0.41 | -0.14 | -0.80 | 0.016 |
| PI4K2A | S47; S51 | -0.05 | 0.02 | 0.26 | -0.25 | -0.11 | 0.08 | 0.272 |
| PI4K2B | S17 | -0.20 | -0.14 | -0.02 | 0.18 | -0.07 | 0.21 | 0.093 |
| PI4KA | S/T | -0.11 | 0.53 | 0.28 | -0.51 | -0.26 | -0.18 | 0.06 |
| PI4KB | S266 | -0.10 | -0.77 | -0.27 | -0.02 | 0.17 | 0.62 | 0.084 |
| PI4KB | S428 | 0.00 | -0.23 | -0.09 | 0.19 | 0.08 | 0.01 | 0.082 |
| PI4KB | S511 | 0.38 | -0.07 | -0.53 | -0.15 | 0.46 | -0.37 | 0.879 |
| PIAS1 | S488 | 0.19 | -0.29 | -0.17 | -0.06 | 0.08 | 0.18 | 0.38 |
| PIAS1 | S503 | 0.01 | -0.17 | -0.05 | 0.22 | -0.04 | -0.01 | 0.267 |
| PIAS1 | S488 | 0.40 | 0.02 | 0.07 | -0.43 | -0.29 | 0.09 | 0.129 |
| PIEZO1 | S165 | -0.06 | 0.17 | -0.08 | -0.20 | 0.06 | 0.08 | 0.801 |
| PIEZO1 | S1646 | 0.07 | 0.15 | 0.07 | 0.01 | -0.08 | -0.26 | 0.069 |
| PIEZO1 | T734 | 0.00 | 0.15 | 0.23 | -0.29 | 0.04 | -0.20 | 0.082 |
| PIF1 | S151 | -0.16 | -0.27 | -0.17 | 0.27 | 0.16 | 0.09 | 0.004 |
| PIF1 | S27 | -0.01 | -0.11 | -0.12 | 0.06 | -0.08 | 0.24 | 0.208 |
| PIF1 | S27 | 0.09 | -0.22 | -0.08 | -0.06 | 0.09 | 0.15 | 0.293 |
| PIGW | S416 | -0.03 | 0.11 | -0.01 | 0.18 | -0.01 | -0.28 | 0.715 |
| PIK3C2B | S155 | 0.12 | 0.20 | -0.06 | 0.02 | -0.10 | -0.23 | 0.15 |
| PIK3C3 | S244 | 0.49 | -0.09 | -0.30 | 0.01 | 0.07 | -0.34 | 0.668 |
| PITPNM1 | S300 | -0.07 | 0.17 | 0.07 | -0.12 | -0.15 | 0.06 | 0.25 |
| PITPNM1 | S896 | 0.25 | -0.01 | 0.03 | 0.34 | -0.26 | -0.51 | 0.422 |
| PITPNM1 | S300 | -0.10 | 0.20 | 0.21 | -0.26 | 0.19 | -0.35 | 0.284 |
| PITPNM1 | S373 | -0.30 | 0.57 | 0.16 | -0.53 | 0.25 | -0.51 | 0.32 |
| PITPNM3 | S109 | -0.05 | -0.31 | 0.10 | -0.26 | 0.09 | 0.33 | 0.547 |
| PITX1 | S46; S/T | 0.18 | 0.13 | 0.04 | -0.17 | -0.19 | -0.03 | 0.018 |
| PKMYT1 | S94 | -0.03 | -0.37 | -0.33 | 0.28 | 0.18 | 0.14 | 0.019 |
| PKMYT1 | S160 | -0.02 | 0.02 | -0.28 | 0.00 | 0.25 | -0.02 | 0.256 |
| PKMYT1 | S469 | 0.01 | 0.34 | -0.09 | -0.18 | 0.20 | -0.40 | 0.381 |
| PKN1 | S/T/Y | 0.42 | -1.06 | 0.21 | -0.15 | -0.30 | 0.39 | 0.817 |
| PKN1 | S/T | 0.09 | 0.04 | 0.07 | -0.07 | 0.01 | -0.16 | 0.053 |
| PKN1 | S916 | 0.06 | 0.13 | 0.20 | -0.27 | -0.01 | -0.17 | 0.03 |
| PKN2 | S583 | -0.11 | -0.15 | -0.11 | 0.09 | 0.05 | 0.18 | 0.004 |
| PKN2 | T958 | -0.18 | -0.06 | -0.04 | 0.08 | 0.01 | 0.17 | 0.052 |

| | | | | | | | | |
|---|---|---|---|---|---|---|---|---|
| **PKN2** | **T958** | 0.11 | 0.13 | -0.26 | -0.19 | 0.17 | -0.03 | 0.969 |
| **PKN2** | **T958** | 0.02 | 0.02 | -0.01 | 0.11 | -0.02 | -0.13 | 0.736 |
| **PKNOX1** | **S41** | 0.13 | 0.00 | 0.07 | -0.04 | -0.11 | -0.06 | 0.039 |
| **PKNOX1** | **S41** | 0.03 | 0.13 | 0.18 | -0.13 | -0.05 | -0.19 | 0.016 |
| **PKP2** | **S155** | 0.21 | 0.03 | -0.14 | -0.10 | -0.03 | 0.00 | 0.498 |
| **PKP3** | **S707** | -0.64 | -0.58 | -0.55 | 0.49 | 0.22 | 0.52 | 5E-04 |
| **PKP3** | **S359** | -0.53 | -0.43 | -0.60 | 0.41 | 0.32 | 0.40 | 1E-04 |
| **PKP3** | **S180** | -0.32 | -0.34 | -0.29 | 0.27 | 0.15 | 0.35 | 6E-04 |
| **PKP3** | **S123** | -0.25 | -0.37 | -0.22 | 0.28 | 0.15 | 0.27 | 0.001 |
| **PKP3** | **T178; S180** | -0.31 | -0.34 | -0.14 | 0.23 | 0.05 | 0.37 | 0.012 |
| **PKP3** | **S196** | -0.15 | -0.14 | -0.18 | 0.22 | 0.18 | 0.02 | 0.009 |
| **PKP3** | **S196** | 0.00 | -0.02 | -0.11 | 0.13 | 0.00 | -0.01 | 0.211 |
| **PKP3** | **S238** | -0.04 | -0.07 | -0.01 | 0.10 | -0.03 | 0.04 | 0.127 |
| **PKP3** | **S199** | 0.26 | -0.43 | 0.00 | -0.28 | -0.05 | 0.34 | 0.838 |
| **PKP3** | **S** | -0.05 | 0.17 | 0.10 | 0.05 | 0.09 | -0.44 | 0.385 |
| **PKP4** | **S337** | -0.69 | 0.03 | -0.15 | 0.28 | -0.17 | 0.44 | 0.185 |
| **PKP4** | **S221** | 0.00 | -0.34 | -0.19 | 0.02 | -0.04 | 0.43 | 0.151 |
| **PKP4** | **S447** | -0.05 | -0.12 | -0.12 | 0.01 | 0.25 | 0.00 | 0.093 |
| **PKP4** | **S406** | 0.15 | -0.31 | -0.08 | -0.01 | 0.10 | 0.11 | 0.359 |
| **PKP4** | **S327; S** | -0.08 | -0.17 | 0.11 | -0.10 | 0.11 | 0.12 | 0.46 |
| **PKP4** | **S273; T283** | 0.03 | -0.17 | 0.11 | 0.00 | -0.05 | 0.07 | 0.882 |
| **PKP4** | **S337** | -0.03 | 0.02 | 0.01 | -0.06 | 0.09 | -0.04 | 0.922 |
| **PKP4** | **S/T** | -0.12 | -0.01 | 0.17 | -0.10 | 0.12 | -0.08 | 0.758 |
| **PKP4** | **S314** | 0.08 | 0.02 | 0.08 | -0.15 | 0.04 | -0.09 | 0.096 |
| **PKP4** | **S281** | 0.23 | 0.39 | -0.25 | -0.30 | 0.01 | -0.24 | 0.236 |
| **PKP4** | **S** | 0.06 | 0.01 | 0.42 | -0.18 | 0.11 | -0.62 | 0.186 |
| **PLCB1** | **S1199** | -0.02 | -0.14 | -0.09 | -0.09 | 0.05 | 0.27 | 0.216 |
| **PLCB3** | **S1105** | -0.01 | -0.11 | 0.05 | 0.08 | -0.12 | 0.10 | 0.609 |
| **PLCB3** | **S1105** | 0.28 | 0.19 | 0.16 | -0.24 | -0.06 | -0.47 | 0.019 |
| **PLCH1** | **S/T** | 0.08 | 0.16 | -0.17 | -0.14 | -0.24 | 0.24 | 0.701 |
| **PLD1** | **S629** | 0.10 | 0.13 | -0.36 | -0.18 | 0.35 | -0.15 | 0.844 |
| **PLD2** | **S134** | 0.09 | -0.34 | 0.33 | -0.67 | 0.01 | 0.33 | 0.724 |
| **PLEC** | **S1047** | -0.06 | -0.68 | -0.39 | 0.38 | 0.33 | 0.13 | 0.027 |
| **PLEC** | **S2802** | -0.10 | -0.51 | 0.04 | 0.21 | -0.19 | 0.37 | 0.242 |
| **PLEC** | **S4613** | 0.08 | -0.09 | -0.24 | 0.19 | -0.17 | 0.17 | 0.388 |
| **PLEC** | **S4622; T/S/Y** | -0.42 | 0.12 | 0.02 | 0.42 | -0.36 | 0.05 | 0.668 |
| **PLEC** | **S1732** | -0.11 | 0.07 | -0.11 | 0.26 | 0.05 | -0.21 | 0.604 |
| **PLEC** | **T4030** | -0.17 | 0.11 | -0.07 | -0.11 | 0.30 | -0.11 | 0.699 |
| **PLEC** | **S1435** | -0.04 | 0.03 | -0.03 | 0.17 | -0.02 | -0.11 | 0.792 |
| **PLEC** | **S4386; S4396** | -0.02 | 0.04 | -0.01 | -0.12 | 0.03 | 0.07 | 0.889 |
| **PLEC** | **S4386** | 0.12 | 0.16 | -0.02 | -0.07 | 0.01 | -0.23 | 0.115 |
| **PLEC** | **S4675** | 0.07 | 0.03 | 0.12 | -0.67 | 0.00 | 0.29 | 0.522 |
| **PLEC** | **S720** | -0.04 | 0.15 | 0.25 | -0.16 | 0.07 | -0.35 | 0.148 |
| **PLEC** | **S1721** | 0.11 | 0.27 | -0.01 | -0.12 | 0.01 | -0.33 | 0.102 |
| **PLEKHA5** | **S855** | 0.09 | 0.02 | -0.50 | 0.21 | 0.05 | 0.02 | 0.319 |
| **PLEKHA6** | **S867** | -0.02 | -0.27 | -0.38 | 0.33 | -0.08 | 0.29 | 0.076 |
| **PLEKHA6** | **S940** | -0.14 | 0.09 | -0.09 | 0.13 | 0.17 | -0.19 | 0.577 |
| **PLEKHA6** | **S336** | 0.02 | 0.00 | -0.05 | -0.15 | -0.02 | 0.18 | 0.903 |

| | | | | | | | | |
|---|---|---|---|---|---|---|---|---|
| PLEKHA6 | S/T | 0.02 | -0.30 | 0.25 | -0.16 | 0.16 | -0.05 | 0.972 |
| PLEKHA6 | S476 | 0.11 | -0.12 | 0.08 | -0.50 | 0.41 | -0.13 | 0.738 |
| PLEKHA6 | S251 | -0.17 | 0.08 | 0.33 | -0.29 | -0.09 | 0.05 | 0.34 |
| PLEKHA6 | S459 | 0.26 | 0.26 | 0.03 | -0.10 | -0.16 | -0.41 | 0.029 |
| PLEKHA6 | S777 | 0.12 | 0.22 | 0.25 | -0.19 | -0.10 | -0.40 | 0.011 |
| PLEKHA7 | S907; T/S | -0.14 | -0.05 | -0.38 | 0.28 | 0.05 | 0.15 | 0.043 |
| PLEKHA7 | S903 | 0.06 | 0.09 | -0.34 | -0.26 | 0.06 | 0.29 | 0.701 |
| PLEKHA7 | S612 | -0.07 | -0.04 | 0.04 | 0.01 | 0.06 | -0.01 | 0.349 |
| PLEKHA7 | S986 | 0.02 | -0.07 | 0.17 | 0.10 | -0.07 | -0.18 | 0.419 |
| PLEKHA7 | S612 | -0.20 | 0.45 | -0.05 | 0.04 | -0.32 | -0.04 | 0.472 |
| PLEKHF2 | S226 | 0.00 | 0.17 | -0.05 | -0.03 | 0.02 | -0.13 | 0.345 |
| PLEKHG3 | S741 | -0.05 | -0.18 | -0.07 | -0.45 | 0.32 | 0.28 | 0.59 |
| PLEKHG3 | S433 | 0.03 | -0.14 | -0.05 | -0.07 | 0.07 | 0.13 | 0.272 |
| PLEKHG3 | S962 | -0.03 | -0.12 | 0.00 | 0.08 | -0.32 | 0.31 | 0.714 |
| PLEKHG3 | S76 | -0.06 | 0.02 | -0.05 | 0.17 | -0.11 | 0.01 | 0.574 |
| PLEKHG3 | S1154; S1169 | -0.11 | 0.06 | -0.02 | 0.19 | -0.17 | 0.03 | 0.749 |
| PLEKHG3 | S1081 | -0.01 | -0.16 | 0.12 | 0.02 | 0.11 | -0.10 | 0.786 |
| PLEKHG3 | S1081 | -0.04 | -0.15 | 0.21 | 0.03 | 0.00 | -0.08 | 0.816 |
| PLEKHG3 | S | 0.06 | -0.02 | 0.00 | -0.24 | -0.18 | 0.31 | 0.777 |
| PLEKHG3 | S1081 | 0.20 | 0.21 | -0.32 | -0.17 | 0.13 | -0.14 | 0.669 |
| PLEKHG3 | S962 | -0.06 | -0.09 | 0.25 | -0.45 | -0.22 | 0.40 | 0.681 |
| PLEKHG3 | S741 | 0.04 | 0.23 | -0.06 | -0.18 | 0.13 | -0.21 | 0.311 |
| PLEKHG3 | S1037 | 0.15 | 0.17 | -0.03 | -0.12 | -0.18 | -0.01 | 0.066 |
| PLEKHG3 | S1154; S/T | 0.21 | -0.05 | 0.14 | -0.06 | -0.30 | 0.01 | 0.158 |
| PLEKHG3 | S647 | 0.23 | 0.31 | 0.39 | -0.43 | -0.20 | -0.57 | 0.004 |
| PLEKHG3 | S1028 | 0.19 | 0.33 | 0.46 | -0.45 | -0.17 | -0.72 | 0.012 |
| PLEKHG3 | S962 | 0.39 | 0.29 | 0.33 | -0.56 | -0.25 | -0.55 | 0.002 |
| PLEKHG4 | S64 | -0.32 | -0.11 | -0.32 | 0.45 | -0.04 | 0.19 | 0.046 |
| PLEKHG4 | S64 | -0.36 | -0.11 | -0.24 | 0.30 | 0.01 | 0.28 | 0.023 |
| PLEKHG4 | S64 | -0.23 | -0.02 | -0.22 | 0.17 | -0.05 | 0.29 | 0.071 |
| PLEKHG4 | S414 | 0.12 | -0.11 | 0.41 | -0.13 | -0.10 | -0.28 | 0.124 |
| PLEKHG6 | S/T | 0.09 | -0.21 | 0.27 | -0.47 | 0.29 | -0.12 | 0.595 |
| PLEKHN1 | S559 | -0.16 | 0.08 | 0.02 | -0.02 | 0.12 | -0.06 | 0.714 |
| PLK2 | S/T | 0.38 | -0.19 | -0.04 | -0.01 | -0.03 | -0.20 | 0.511 |
| PLK4 | S665 | -0.04 | -0.25 | -0.34 | 0.18 | 0.02 | 0.33 | 0.037 |
| PLPP3 | S19 | -0.11 | 0.00 | -0.12 | -0.05 | 0.01 | 0.24 | 0.203 |
| PLPP3 | S19 | 0.20 | -0.09 | 0.05 | -0.44 | 0.48 | -0.44 | 0.585 |
| PLPP4 | S257 | -0.35 | -0.49 | -0.10 | 0.53 | -0.19 | 0.33 | 0.09 |
| PLPP4 | S257 | 0.10 | -0.50 | -0.38 | 0.30 | -0.18 | 0.42 | 0.17 |
| PLPP6 | S63 | 0.35 | 0.22 | 0.37 | -0.24 | -0.34 | -0.67 | 0.006 |
| PML | S403 | -0.11 | -0.20 | -0.34 | 0.32 | 0.02 | 0.20 | 0.022 |
| PML | S518; S527 | -0.26 | -0.31 | -0.08 | 0.21 | -0.04 | 0.36 | 0.043 |
| PML | S36 | -0.06 | -0.18 | 0.09 | 0.09 | -0.01 | 0.06 | 0.303 |
| PML | S505 | 0.05 | 0.07 | 0.17 | 0.01 | 0.08 | -0.46 | 0.28 |
| PNISR | S465 | -0.69 | -0.49 | -0.72 | 0.45 | 0.66 | 0.16 | 0.003 |
| PNISR | S211 | 0.09 | -0.46 | -0.32 | 0.11 | -0.14 | 0.51 | 0.195 |
| PNISR | T485 | 0.03 | -0.12 | -0.18 | 0.26 | -0.17 | 0.13 | 0.316 |
| PNISR | S290 | 0.29 | 0.00 | 0.02 | -0.19 | -0.01 | -0.16 | 0.11 |

| | | | | | | | | |
|---|---|---|---|---|---|---|---|---|
| PNKP | S114; T118; T122 | -0.37 | -0.56 | -0.70 | 0.40 | 0.43 | 0.34 | 7E-04 |
| PNKP | S114; T118 | -0.54 | -0.33 | -0.63 | 0.45 | 0.28 | 0.38 | 1E-03 |
| PNKP | S114; T118 | -0.26 | -0.27 | -0.42 | 0.27 | 0.25 | 0.26 | 4E-04 |
| PNKP | T118 | 0.01 | -0.01 | 0.05 | -0.05 | -0.02 | 0.02 | 0.267 |
| PNKP | T118 | 0.07 | 0.10 | 0.09 | -0.09 | -0.10 | -0.09 | 1E-04 |
| PNN | S381 | -0.44 | -0.59 | -0.49 | 0.31 | 0.61 | 0.16 | 0.004 |
| PNN | S381 | -0.56 | -0.28 | -0.54 | 0.54 | 0.49 | -0.05 | 0.02 |
| PNN | S66 | -0.35 | -0.34 | -0.65 | 0.42 | 0.29 | 0.30 | 0.002 |
| PNN | S381 | -0.39 | -0.30 | -0.50 | 0.41 | 0.39 | 0.11 | 0.003 |
| PNN | S66 | -0.21 | -0.27 | -0.32 | 0.21 | 0.22 | 0.24 | 1E-04 |
| PNN | S534 | 0.02 | -0.41 | -0.40 | 0.17 | 0.23 | 0.24 | 0.028 |
| PNN | S381 | -0.39 | -0.26 | -0.06 | 0.20 | 0.03 | 0.35 | 0.032 |
| PNN | S417; T/S | -0.25 | -0.07 | -0.14 | 0.25 | -0.05 | 0.20 | 0.053 |
| PNN | S347 | -0.02 | -0.19 | -0.21 | 0.04 | -0.11 | 0.39 | 0.196 |
| PNN | S347 | -0.05 | -0.15 | -0.17 | 0.01 | -0.13 | 0.40 | 0.249 |
| PNN | S413 | -0.13 | -0.05 | -0.17 | 0.16 | 0.25 | -0.12 | 0.142 |
| PNN | S413 | -0.12 | -0.09 | -0.14 | 0.12 | 0.25 | -0.08 | 0.1 |
| PNN | S413 | -0.17 | -0.13 | -0.02 | 0.11 | 0.13 | 0.06 | 0.013 |
| PNN | S413 | -0.04 | -0.16 | -0.12 | 0.03 | 0.13 | 0.13 | 0.014 |
| PNN | S347 | 0.01 | -0.10 | -0.16 | 0.09 | -0.04 | 0.18 | 0.111 |
| PNN | S347 | -0.25 | 0.07 | 0.00 | -0.07 | -0.05 | 0.25 | 0.501 |
| PNN | S422 | 0.10 | -0.22 | 0.01 | -0.02 | 0.12 | -0.02 | 0.581 |
| PNN | S347 | 0.07 | 0.00 | -0.16 | 0.12 | -0.05 | 0.00 | 0.584 |
| PNN | S347 | 0.08 | 0.00 | -0.10 | 0.06 | -0.03 | -0.03 | 0.927 |
| PNN | S347 | 0.11 | 0.02 | -0.10 | 0.00 | 0.03 | -0.07 | 0.757 |
| PNN | S347 | 0.13 | 0.01 | -0.11 | 0.02 | -0.01 | -0.05 | 0.759 |
| PNN | S347 | 0.05 | 0.07 | -0.07 | 0.04 | 0.02 | -0.10 | 0.652 |
| PNN | S552 | -0.14 | 0.14 | 0.04 | -0.01 | -0.33 | 0.23 | 0.795 |
| PNN | S347 | 0.03 | 0.05 | 0.01 | 0.01 | -0.05 | -0.05 | 0.056 |
| PNN | S417 | 0.13 | 0.02 | -0.02 | -0.09 | 0.01 | -0.06 | 0.158 |
| PNN | S417 | 0.19 | 0.00 | -0.03 | -0.04 | 0.15 | -0.33 | 0.456 |
| PNN | S422 | -0.02 | 0.22 | 0.08 | -0.07 | 0.05 | -0.32 | 0.184 |
| PNN | T/S | 0.09 | 0.11 | 0.17 | -0.15 | -0.08 | -0.19 | 0.003 |
| PNN | S | 0.05 | 0.15 | 0.19 | -0.05 | 0.06 | -0.50 | 0.175 |
| PNN | S422 | 0.18 | 0.10 | 0.13 | -0.03 | 0.09 | -0.61 | 0.216 |
| PNN | S100 | 0.20 | 0.15 | 0.09 | -0.06 | 0.01 | -0.50 | 0.111 |
| PNN | S100 | 0.24 | 0.18 | 0.07 | -0.11 | 0.02 | -0.54 | 0.102 |
| PNPLA6 | T464 | -0.09 | 0.03 | -0.02 | 0.06 | 0.14 | -0.14 | 0.629 |
| PNPLA6 | S1318; T1322 | 0.02 | 0.13 | -0.04 | -0.03 | 0.02 | -0.12 | 0.268 |
| PNPLA6 | S1318 | 0.10 | -0.02 | 0.10 | 0.00 | -0.16 | -0.04 | 0.106 |
| PNPLA6 | S354 | 0.10 | 0.15 | 0.38 | -0.43 | -0.30 | -0.06 | 0.026 |
| PNPLA7 | S1288 | -0.36 | -0.41 | 0.18 | 0.08 | 0.11 | 0.26 | 0.15 |
| PNPLA7 | S355 | -0.04 | 0.11 | 0.39 | 0.01 | -0.81 | 0.07 | 0.27 |
| POF1B | S56 | 0.02 | 0.01 | 0.09 | 0.00 | 0.12 | -0.28 | 0.493 |
| POGK | S199 | -0.08 | -0.28 | 0.16 | 0.08 | 0.01 | 0.07 | 0.393 |
| POGZ | S1364; T1368 | -0.27 | -0.13 | 0.15 | 0.11 | -0.14 | 0.22 | 0.421 |
| POGZ | S314 | -0.22 | 0.14 | -0.05 | -0.03 | 0.16 | -0.03 | 0.541 |
| POGZ | S425 | 0.11 | 0.05 | -0.04 | -0.06 | 0.02 | -0.09 | 0.199 |

| | | | | | | | | |
|---|---|---|---|---|---|---|---|---|
| POGZ | S445 | -0.09 | 0.26 | 0.13 | -0.08 | 0.05 | -0.35 | 0.221 |
| POGZ | S445 | 0.41 | 0.14 | -0.11 | -0.23 | -0.25 | -0.08 | 0.108 |
| POGZ | T1368 | 0.25 | 0.17 | 0.08 | -0.03 | -0.38 | -0.17 | 0.033 |
| POGZ | T448 | 0.35 | 0.23 | -0.01 | -0.26 | -0.14 | -0.30 | 0.021 |
| POLA2 | T127 | -0.10 | -0.42 | -0.08 | -0.07 | 0.06 | 0.46 | 0.145 |
| POLD3 | S307 | -0.31 | -0.19 | -0.13 | 0.18 | -0.08 | 0.40 | 0.064 |
| POLD3 | S356 | -0.11 | -0.13 | -0.19 | -0.14 | 0.01 | 0.45 | 0.229 |
| POLD3 | S307 | 0.20 | 0.06 | -0.12 | -0.10 | 0.08 | -0.15 | 0.433 |
| POLD3 | S348; S356 | 0.22 | -0.10 | 0.07 | -0.11 | -0.17 | 0.05 | 0.291 |
| POLD3 | S407; T411; S/Y | -0.02 | 0.26 | 0.15 | 0.13 | -0.37 | -0.26 | 0.156 |
| POLDIP3 | S/T | 0.06 | -0.29 | -0.01 | 0.44 | -0.41 | 0.05 | 0.709 |
| POLDIP3 | S383 | 0.09 | -0.25 | 0.01 | 0.24 | -0.07 | -0.07 | 0.622 |
| POLDIP3 | S275 | -0.08 | 0.43 | -0.24 | -0.36 | 0.01 | 0.10 | 0.659 |
| POLDIP3 | S/T | -0.12 | -0.03 | 0.51 | 0.16 | -0.42 | -0.30 | 0.309 |
| POLDIP3 | S383 | 0.22 | 0.25 | 0.28 | -0.49 | 0.02 | -0.50 | 0.029 |
| POLE | S1940 | -0.25 | -0.39 | -0.39 | 0.50 | 0.25 | 0.04 | 0.013 |
| POLI | T/S | -0.41 | -0.31 | 0.13 | -0.07 | 0.25 | 0.27 | 0.158 |
| POLM | S12 | -0.11 | -0.03 | 0.36 | -0.35 | 0.16 | -0.13 | 0.431 |
| POLR1A | S1489 | -0.48 | -0.16 | -0.50 | 0.34 | 0.23 | 0.31 | 0.004 |
| POLR1A | T357 | -0.26 | -0.42 | -0.34 | 0.31 | -0.12 | 0.55 | 0.045 |
| POLR1D | S104 | 0.04 | 0.00 | -0.12 | -0.01 | 0.00 | 0.08 | 0.43 |
| POLR1E | S230 | -0.30 | -0.43 | 0.24 | 0.02 | 0.22 | 0.12 | 0.258 |
| POLR1F | T322 | -0.15 | 0.07 | -0.15 | 0.33 | 0.03 | -0.20 | 0.483 |
| POLR1F | S316 | 0.14 | 0.00 | 0.08 | 0.02 | -0.14 | -0.10 | 0.078 |
| POLR1F | S328 | 0.18 | 0.25 | 0.18 | -0.35 | -0.02 | -0.37 | 0.018 |
| POLR1G | S/T | -0.27 | -0.09 | 0.19 | -0.50 | 0.15 | 0.35 | 0.847 |
| POLR1G | S490 | 0.05 | -0.12 | 0.04 | 0.01 | -0.06 | 0.08 | 0.761 |
| POLR1G | S285 | -0.06 | 0.18 | 0.00 | -0.09 | 0.24 | -0.34 | 0.615 |
| POLR1G | S285 | -0.06 | 0.18 | 0.00 | -0.09 | 0.24 | -0.34 | 0.615 |
| POLR1G | S27 | -0.12 | 0.19 | 0.11 | 0.04 | 0.23 | -0.60 | 0.568 |
| POLR1G | S285 | 0.12 | 0.11 | 0.19 | -0.22 | -0.02 | -0.23 | 0.015 |
| POLR2A | S1941; S/T/Y | 0.00 | -0.16 | 0.01 | -0.04 | 0.01 | 0.17 | 0.3 |
| POLR2A | S1889; T/S/Y | 0.16 | 0.14 | -0.04 | -0.10 | -0.06 | -0.14 | 0.05 |
| POLR2A | S1966 | 0.22 | -0.02 | 0.03 | -0.47 | -0.24 | 0.34 | 0.464 |
| POLR2A | T1863; S1868 | 0.19 | 0.09 | 0.05 | -0.07 | -0.08 | -0.21 | 0.02 |
| POLR2A | S1847; S1857 | 0.18 | 0.11 | 0.07 | -0.05 | -0.13 | -0.22 | 0.012 |
| POLR2A | S1857 | 0.28 | 0.15 | -0.08 | -0.08 | -0.15 | -0.18 | 0.081 |
| POLR2A | S1924; S1931 | 0.24 | 0.10 | 0.09 | -0.10 | -0.13 | -0.26 | 0.011 |
| POLR2A | S1934 | 0.29 | 0.10 | 0.03 | -0.13 | -0.07 | -0.30 | 0.04 |
| POLR2A | S1875; S1882 | 0.19 | 0.20 | 0.08 | -0.12 | -0.13 | -0.28 | 0.007 |
| POLR2A | S1882 | 0.23 | 0.24 | 0.04 | -0.18 | -0.12 | -0.31 | 0.012 |
| POLR2A | T1894 | 0.57 | 0.14 | 0.05 | -0.65 | 0.02 | -0.47 | 0.072 |
| POLR2B | S861 | 0.28 | -0.64 | -0.15 | -0.09 | 0.13 | 0.27 | 0.392 |
| POLR2B | S75 | -0.21 | 0.01 | 0.09 | 0.19 | -0.06 | -0.05 | 0.628 |
| POLR2B | S75 | 0.13 | -0.07 | 0.10 | -0.12 | -0.03 | -0.04 | 0.167 |
| POLR2M | S/T | 0.13 | 0.03 | -0.35 | 0.03 | 0.28 | -0.20 | 0.64 |
| POLR3E | S503 | 0.14 | -0.01 | 0.06 | 0.28 | -0.18 | -0.39 | 0.477 |
| POLR3E | S503 | 0.14 | -0.01 | 0.06 | 0.28 | -0.18 | -0.39 | 0.477 |

| | | | | | | | | |
|---|---|---|---|---|---|---|---|---|
| POM121 | S380 | -0.58 | -0.52 | 0.00 | 0.57 | -0.57 | 0.57 | 0.258 |
| POM121 | S434; S435 | -0.04 | 0.09 | -0.32 | -0.05 | 0.11 | 0.16 | 0.295 |
| POM121 | S435 | 0.05 | -0.19 | -0.08 | 0.10 | -0.10 | 0.19 | 0.291 |
| POM121 | S179 | -0.03 | -0.13 | -0.04 | 0.07 | 0.00 | 0.11 | 0.044 |
| POM121 | S184 | -0.08 | 0.04 | -0.17 | -0.03 | 0.01 | 0.19 | 0.239 |
| POM121 | S269 | -0.01 | -0.13 | -0.06 | -0.09 | 0.24 | 0.02 | 0.303 |
| POM121 | S184; S188 | -0.08 | -0.01 | 0.05 | 0.00 | -0.10 | 0.13 | 0.744 |
| POM121 | S434; S439 | -0.21 | -0.16 | 0.32 | 0.01 | 0.08 | -0.11 | 0.969 |
| POM121 | S174; S179 | 0.01 | -0.09 | 0.09 | -0.07 | -0.16 | 0.20 | 0.913 |
| POM121 | S435 | 0.10 | -0.20 | 0.11 | -0.15 | 0.09 | 0.02 | 0.896 |
| POM121 | S188 | 0.13 | 0.00 | -0.04 | -0.02 | -0.17 | 0.08 | 0.506 |
| POM121 | S179; S184; S/T | -0.09 | 0.17 | 0.22 | -0.15 | -0.15 | -0.05 | 0.107 |
| POM121 | S108 | 0.21 | 0.07 | 0.01 | -0.20 | 0.23 | -0.42 | 0.317 |
| POM121 | T447 | 0.30 | 0.15 | 0.06 | -0.25 | -0.04 | -0.31 | 0.026 |
| POM121 | S445; T447 | 0.19 | 0.15 | 0.22 | -0.29 | -0.12 | -0.26 | 0.002 |
| POM121 | S435 | 0.22 | 0.20 | 0.39 | -0.62 | -0.03 | -0.42 | 0.026 |
| POM121C | S95 | -0.16 | 0.00 | 0.04 | -0.37 | 0.29 | 0.10 | 0.838 |
| POMGNT1 | S66 | -0.31 | 0.07 | -0.02 | 0.11 | -0.13 | 0.22 | 0.39 |
| POMT1 | S727 | 0.11 | 0.02 | 0.24 | -0.47 | -0.03 | 0.02 | 0.168 |
| POP1 | S367 | -0.51 | -0.54 | -0.26 | 0.73 | 0.20 | -0.04 | 0.04 |
| POP1 | S/Y | 0.02 | 0.03 | 0.04 | 0.04 | -0.05 | -0.09 | 0.182 |
| POP1 | S730 | 0.07 | 0.09 | 0.06 | -0.04 | -0.07 | -0.12 | 0.003 |
| POP1 | S730 | 0.13 | 0.24 | 0.11 | -0.06 | -0.06 | -0.46 | 0.062 |
| POP5 | S154 | -0.10 | -0.12 | -0.06 | -0.01 | 0.01 | 0.25 | 0.112 |
| POP5 | S154 | -0.11 | -0.08 | -0.02 | 0.06 | -0.07 | 0.20 | 0.184 |
| POU2F1 | S366 | -0.16 | -0.22 | 0.14 | 0.26 | 0.06 | -0.14 | 0.436 |
| POU2F1 | S385 | 0.12 | -0.08 | 0.22 | 0.09 | -0.17 | -0.24 | 0.224 |
| POU2F1 | S448 | 0.24 | 0.08 | 0.23 | -0.50 | 0.07 | -0.25 | 0.076 |
| POU3F2 | S341 | 0.15 | 0.57 | -0.01 | -0.69 | -0.41 | 0.07 | 0.108 |
| PPA2 | S317 | -0.08 | -0.01 | 0.08 | 0.18 | -0.19 | 0.00 | 0.981 |
| PPAN | S359 | 0.12 | 0.08 | 0.25 | -0.16 | -0.01 | -0.35 | 0.045 |
| PPFIA1 | S668 | -0.33 | -0.52 | -0.26 | 0.19 | 0.32 | 0.35 | 0.002 |
| PPFIA1 | T/S | -0.27 | -0.57 | -0.16 | 0.15 | 0.36 | 0.27 | 0.012 |
| PPFIA1 | T/S | 0.01 | 0.12 | 0.08 | -0.18 | 0.01 | -0.06 | 0.086 |
| PPFIBP1 | S466 | -1.20 | -0.11 | -0.82 | 0.62 | 0.60 | 0.03 | 0.039 |
| PPFIBP1 | S794 | -0.21 | -0.07 | -0.10 | 0.29 | 0.03 | 0.01 | 0.076 |
| PPFIBP1 | S794 | -0.08 | 0.06 | -0.19 | 0.16 | -0.04 | 0.06 | 0.235 |
| PPFIBP2 | S387 | -0.01 | -0.07 | 0.00 | 0.07 | 0.08 | -0.09 | 0.502 |
| PPFIBP2 | S329 | 0.03 | 0.02 | -0.06 | 0.12 | -0.14 | 0.01 | 0.974 |
| PPFIBP2 | S414 | -0.13 | 0.19 | 0.02 | 0.05 | -0.06 | -0.09 | 0.607 |
| PPFIBP2 | S387 | 0.06 | 0.23 | -0.18 | -0.02 | -0.08 | -0.05 | 0.509 |
| PPHLN1 | S133 | 0.07 | -0.05 | 0.03 | -0.05 | 0.03 | -0.04 | 0.441 |
| PPHLN1 | S | 0.15 | 0.22 | -0.08 | 0.08 | -0.23 | -0.20 | 0.186 |
| PPHLN1 | T204 | 0.19 | 0.08 | 0.08 | -0.15 | -0.03 | -0.22 | 0.021 |
| PPHLN1 | S201; S205 | 0.04 | 0.13 | 0.18 | -0.19 | 0.12 | -0.36 | 0.155 |
| PPHLN1 | S205; S210 | 0.17 | 0.12 | 0.18 | -0.06 | -0.17 | -0.30 | 0.009 |
| PPHLN1 | S140 | 0.23 | 0.18 | 0.11 | -0.16 | -0.04 | -0.42 | 0.032 |
| PPHLN1 | T/S | 0.02 | 0.15 | 0.34 | -0.49 | 0.16 | -0.37 | 0.138 |

| | | | | | | | | |
|---|---|---|---|---|---|---|---|---|
| PPHLN1 | S121 | 0.30 | 0.22 | 0.03 | -0.09 | -0.03 | -0.60 | 0.096 |
| PPID | S201 | -0.18 | 0.05 | -0.03 | -0.12 | 0.11 | 0.14 | 0.42 |
| PPIG | S696 | -0.62 | -0.62 | -0.74 | 0.55 | 0.70 | 0.01 | 0.007 |
| PPIG | S259 | -0.07 | 0.14 | -0.04 | -0.11 | -0.13 | 0.18 | 0.804 |
| PPIG | T343; S347 | -0.01 | 0.12 | -0.03 | 0.08 | -0.04 | -0.15 | 0.479 |
| PPIG | S356; T358 | -0.05 | 0.07 | 0.09 | -0.01 | -0.08 | -0.02 | 0.204 |
| PPIG | S259 | 0.07 | 0.05 | 0.05 | -0.11 | -0.04 | -0.03 | 0.01 |
| PPIG | S356; T358 | 0.10 | 0.04 | 0.04 | 0.01 | -0.25 | 0.04 | 0.234 |
| PPIG | T358 | 0.12 | 0.02 | 0.04 | 0.03 | 0.01 | -0.25 | 0.234 |
| PPIG | S256; S257; S259 | -0.01 | 0.00 | 0.20 | -0.12 | -0.07 | -0.01 | 0.163 |
| PPIG | S577; T579 | 0.04 | 0.20 | -0.04 | -0.01 | -0.10 | -0.11 | 0.147 |
| PPIG | S256; S257; S259 | 0.00 | -0.05 | 0.27 | 0.18 | -0.21 | -0.26 | 0.371 |
| PPIG | S/T | -0.10 | 0.19 | 0.15 | -0.11 | 0.14 | -0.34 | 0.321 |
| PPIG | S744; S745; T748 | 0.12 | 0.04 | 0.12 | -0.05 | -0.10 | -0.14 | 0.008 |
| PPIG | S397 | -0.08 | 0.17 | 0.17 | 0.09 | 0.00 | -0.42 | 0.336 |
| PPIG | T611 | 0.09 | 0.08 | 0.15 | -0.06 | 0.02 | -0.34 | 0.107 |
| PPIG | S415 | 0.09 | 0.09 | 0.17 | -0.06 | -0.05 | -0.28 | 0.034 |
| PPIG | S/T | -0.01 | 0.19 | 0.18 | -0.07 | -0.08 | -0.25 | 0.043 |
| PPIG | S/T | -0.08 | 0.22 | 0.25 | -0.27 | 0.00 | -0.19 | 0.099 |
| PPIG | S356; T358 | 0.19 | 0.15 | 0.07 | -0.16 | -0.18 | -0.12 | 0.002 |
| PPIG | S533 | 0.18 | 0.14 | 0.02 | 0.17 | 0.09 | -0.83 | 0.397 |
| PPIG | S695 | -0.10 | 0.35 | 0.16 | -0.06 | 0.12 | -0.67 | 0.285 |
| PPIG | S290 | 0.28 | 0.28 | -0.11 | -0.01 | -0.09 | -0.49 | 0.153 |
| PPIG | S753 | -0.10 | 0.43 | 0.15 | -0.12 | -0.04 | -0.48 | 0.144 |
| PPIG | S259; S/T | 0.09 | 0.29 | 0.18 | -0.19 | -0.17 | -0.29 | 0.004 |
| PPIG | S257; S259 | 0.49 | -0.09 | 0.15 | -0.50 | 0.04 | -0.31 | 0.131 |
| PPIG | S256; S257; S259 | 0.19 | 0.23 | 0.19 | -0.15 | -0.21 | -0.35 | 0.002 |
| PPIG | T358 | 0.19 | 0.19 | 0.20 | -0.20 | 0.03 | -0.57 | 0.064 |
| PPIG | S290 | 0.29 | 0.25 | 0.08 | -0.12 | -0.07 | -0.61 | 0.06 |
| PPIG | S535 | 0.27 | 0.20 | 0.19 | -0.07 | -0.31 | -0.42 | 0.01 |
| PPIG | S687 | 0.26 | 0.26 | 0.18 | -0.14 | -0.10 | -0.67 | 0.044 |
| PPIG | S717 | 0.23 | 0.26 | 0.24 | -0.07 | -0.35 | -0.49 | 0.012 |
| PPIG | S744; S745; S753 | 0.28 | 0.29 | 0.15 | -0.18 | -0.09 | -0.66 | 0.04 |
| PPIG | S744 | 0.27 | 0.42 | 0.08 | -0.32 | -0.16 | -0.51 | 0.015 |
| PPIG | S658 | 0.38 | 0.38 | 0.13 | -0.36 | -0.15 | -0.68 | 0.016 |
| PPIG | S546 | 0.31 | 0.34 | 0.28 | -0.34 | -0.19 | -0.69 | 0.009 |
| PPIG | S | 0.16 | 0.36 | 0.42 | -0.56 | 0.07 | -0.90 | 0.058 |
| PPIG | S658 | 0.42 | 0.32 | 0.26 | -0.31 | -0.12 | -1.02 | 0.042 |
| PPIG | S413; S415 | 0.35 | 0.39 | 0.31 | -0.42 | -0.06 | -1.09 | 0.044 |
| PPIL1 | T/S | -0.94 | -0.76 | -0.50 | 0.43 | 0.56 | 0.44 | 8E-04 |
| PPIL1 | S149 | -0.75 | -0.64 | -0.80 | 0.55 | 0.60 | 0.28 | 4E-04 |
| PPIL4 | S178 | -0.20 | -0.20 | 0.07 | 0.12 | -0.04 | 0.21 | 0.152 |
| PPIL4 | S178 | -0.08 | 0.01 | -0.02 | 0.13 | -0.05 | 0.00 | 0.422 |
| PPM1D | T529 | -0.28 | -0.15 | -0.40 | 0.17 | 0.09 | 0.41 | 0.014 |
| PPM1D | S40 | -0.16 | 0.00 | -0.31 | 0.04 | 0.21 | 0.15 | 0.05 |
| PPM1D | S40 | 0.01 | -0.04 | -0.41 | 0.41 | -0.02 | -0.07 | 0.279 |
| PPM1D | S40 | 0.17 | -0.09 | -0.20 | 0.09 | 0.16 | -0.18 | 0.697 |
| PPM1D | S40 | -0.06 | -0.01 | -0.03 | 0.06 | -0.16 | 0.18 | 0.595 |

| | | | | | | | | |
|---|---|---|---|---|---|---|---|---|
| **PPM1H** | **S124** | 0.11 | 0.03 | 0.04 | -0.02 | -0.20 | 0.02 | 0.158 |
| **PPP1CA** | **T320** | 0.05 | 0.07 | 0.32 | -0.05 | -0.04 | -0.45 | 0.114 |
| **PPP1CB** | **T316** | 0.03 | -0.02 | -0.09 | 0.01 | 0.12 | -0.06 | 0.475 |
| **PPP1CB** | **T316** | 0.02 | -0.04 | 0.09 | -0.09 | 0.03 | -0.01 | 0.459 |
| **PPP1CC** | **T311** | -0.21 | 0.01 | -0.15 | 0.25 | -0.22 | 0.23 | 0.294 |
| **PPP1R10** | **S545** | 0.07 | -0.12 | -0.06 | 0.35 | -0.13 | -0.18 | 0.785 |
| **PPP1R10** | **T256** | 0.01 | -0.13 | 0.04 | 0.00 | 0.09 | -0.02 | 0.466 |
| **PPP1R10** | **S545** | -0.03 | 0.07 | 0.03 | 0.01 | 0.03 | -0.12 | 0.431 |
| **PPP1R10** | **S542; S545** | 0.43 | 0.02 | -0.17 | -0.25 | -0.35 | 0.17 | 0.381 |
| **PPP1R12A** | **S871** | -0.36 | -0.37 | 0.13 | 0.34 | 0.17 | -0.05 | 0.152 |
| **PPP1R12A** | **S299** | 0.05 | -0.05 | -0.43 | 0.41 | -0.02 | -0.08 | 0.314 |
| **PPP1R12A** | **S/T** | 0.15 | 0.14 | -0.24 | -0.05 | -0.06 | 0.03 | 0.771 |
| **PPP1R12A** | **S/T** | -0.05 | 0.30 | -0.08 | 0.29 | -0.19 | -0.40 | 0.555 |
| **PPP1R12A** | **S871** | 0.06 | 0.05 | 0.35 | -0.19 | 0.03 | -0.41 | 0.103 |
| **PPP1R12A** | **S479** | 0.10 | 0.20 | 0.38 | -0.04 | -0.18 | -0.68 | 0.065 |
| **PPP1R12B** | **S687** | -0.06 | -0.06 | 0.29 | -0.21 | 0.00 | -0.01 | 0.39 |
| **PPP1R12B** | **S687** | -0.06 | -0.06 | 0.29 | -0.21 | 0.00 | -0.01 | 0.39 |
| **PPP1R12C** | **S509** | -0.32 | -0.10 | -0.23 | 0.22 | 0.21 | 0.13 | 0.005 |
| **PPP1R13B** | **S566** | -0.32 | -0.09 | 0.34 | -0.25 | -0.03 | 0.23 | 0.998 |
| **PPP1R15B** | **S** | -0.38 | -0.16 | -0.01 | 0.27 | 0.33 | -0.19 | 0.177 |
| **PPP1R15B** | **S508** | 0.03 | 0.04 | -0.13 | -0.22 | 0.37 | -0.18 | 0.961 |
| **PPP1R16A** | **S478** | -0.33 | -0.14 | 0.03 | 0.23 | -0.30 | 0.37 | 0.345 |
| **PPP1R16A** | **S/T** | -0.33 | -0.14 | 0.03 | 0.23 | -0.30 | 0.37 | 0.345 |
| **PPP1R18** | **S/T** | -0.13 | -0.61 | 0.11 | -0.22 | 0.36 | 0.27 | 0.279 |
| **PPP1R18** | **S/T** | -0.15 | -0.36 | -0.03 | -0.27 | 0.37 | 0.28 | 0.234 |
| **PPP1R18** | **S125** | -0.19 | 0.09 | -0.07 | 0.23 | -0.12 | 0.02 | 0.498 |
| **PPP1R18** | **S224** | 0.03 | 0.16 | 0.11 | -0.05 | -0.09 | -0.19 | 0.019 |
| **PPP1R18** | **S432** | 0.04 | 0.45 | -0.07 | -1.50 | 0.21 | 0.58 | 0.599 |
| **PPP1R1C** | **S65** | 0.25 | -0.24 | -0.09 | -0.09 | -0.19 | 0.26 | 0.919 |
| **PPP1R35** | **S52** | 0.16 | 0.26 | -0.53 | -0.27 | 0.10 | 0.13 | 0.948 |
| **PPP1R3D** | **S46** | -0.83 | -0.90 | -0.95 | 0.60 | 0.55 | 0.49 | 7E-06 |
| **PPP1R3D** | **S74; S77** | -0.33 | -0.36 | -0.31 | 0.32 | 0.08 | 0.40 | 0.004 |
| **PPP1R3D** | **S** | 0.01 | -0.24 | -0.05 | 0.10 | 0.33 | -0.23 | 0.423 |
| **PPP1R9A** | **S187; S190** | -0.75 | -0.59 | -0.26 | 0.66 | -0.59 | 0.73 | 0.152 |
| **PPP1R9A** | **S840** | -0.19 | -0.09 | 0.22 | 0.20 | -0.20 | 0.00 | 0.909 |
| **PPP1R9B** | **S100** | -0.21 | -0.45 | -0.51 | 0.26 | 0.07 | 0.54 | 0.014 |
| **PPP1R9B** | **S** | -0.21 | -0.44 | -0.19 | 0.18 | 0.19 | 0.32 | 0.005 |
| **PPP1R9B** | **S192** | -0.08 | -0.04 | 0.07 | -0.01 | 0.07 | -0.02 | 0.636 |
| **PPP1R9B** | **S192** | -0.05 | 0.20 | 0.43 | -0.55 | 0.02 | -0.26 | 0.1 |
| **PPP6R3** | **S202** | -1.14 | -0.13 | -0.37 | 0.17 | 0.52 | 0.37 | 0.049 |
| **PPP6R3** | **S617** | -0.08 | -0.17 | -0.43 | 0.25 | 0.20 | 0.12 | 0.02 |
| **PPP6R3** | **S617** | -0.05 | -0.13 | -0.02 | 0.09 | 0.00 | 0.10 | 0.041 |
| **PPP6R3** | **S** | -0.05 | -0.09 | -0.01 | 0.19 | -0.09 | 0.03 | 0.325 |
| **PPP6R3** | **T631** | 0.13 | 0.22 | 0.11 | -0.11 | -0.13 | -0.29 | 0.007 |
| **PPRC1** | **T1391** | -0.09 | -0.03 | -0.05 | -0.01 | -0.09 | 0.24 | 0.363 |
| **PPRC1** | **S1411; S1413** | 0.05 | -0.07 | 0.04 | 0.04 | -0.08 | 0.01 | 0.751 |
| **PPRC1** | **S237** | 0.05 | 0.00 | 0.13 | -0.10 | 0.05 | -0.16 | 0.148 |
| **PPRC1** | **S1413** | 0.27 | 0.17 | 0.12 | -0.01 | -0.18 | -0.51 | 0.051 |

| | | | | | | | | |
|---|---|---|---|---|---|---|---|---|
| PPRC1 | S548 | 0.30 | 0.06 | 0.29 | -0.24 | -0.05 | -0.54 | 0.037 |
| PQBP1 | S | 0.12 | -0.04 | 0.52 | -0.33 | -0.08 | -0.39 | 0.072 |
| PRAG1 | S | 0.27 | -0.20 | -0.01 | -0.03 | -0.12 | 0.05 | 0.733 |
| PRAG1 | S706; S712 | -0.13 | 0.21 | 0.11 | -0.04 | 0.16 | -0.38 | 0.469 |
| PRCC | S212 | 0.05 | -0.15 | -0.08 | 0.03 | -0.05 | 0.17 | 0.267 |
| PRCC | S267 | -0.02 | -0.16 | 0.40 | -0.22 | -0.12 | 0.02 | 0.379 |
| PRCC | S159 | 0.28 | 0.20 | 0.04 | -0.14 | -0.12 | -0.35 | 0.021 |
| PRDM10 | S1090 | -0.20 | -0.15 | -0.46 | 0.35 | 0.05 | 0.26 | 0.02 |
| PRDM10 | S1090 | -0.20 | -0.15 | -0.46 | 0.35 | 0.05 | 0.26 | 0.02 |
| PRDM10 | S424 | 0.15 | -0.14 | -0.29 | 0.24 | -0.01 | -0.01 | 0.345 |
| PRDM11 | S63 | 0.12 | 0.13 | 0.42 | -0.17 | -0.09 | -0.63 | 0.057 |
| PRDM2 | S/T | -0.22 | -0.11 | -0.27 | 0.16 | -0.12 | 0.43 | 0.097 |
| PRDM2 | S/T | -0.10 | -0.35 | 0.06 | 0.21 | 0.03 | 0.08 | 0.148 |
| PRDM2 | S1484 | 0.07 | -0.02 | -0.28 | -0.02 | 0.06 | 0.16 | 0.289 |
| PRDM2 | S | -0.12 | -0.19 | 0.03 | -0.41 | 0.43 | 0.11 | 0.622 |
| PRDM2 | S914; T/S/Y | 0.00 | -0.29 | 0.09 | -0.10 | -0.07 | 0.31 | 0.547 |
| PRDM2 | S1118 | 0.01 | -0.02 | -0.12 | -0.12 | -0.05 | 0.27 | 0.561 |
| PRDM2 | S739; S743 | -0.01 | -0.11 | 0.09 | 0.06 | -0.18 | 0.13 | 0.898 |
| PRDM2 | S796 | 0.06 | -0.02 | -0.05 | 0.09 | -0.05 | -0.04 | 0.933 |
| PRDM2 | S781 | -0.20 | 0.19 | 0.01 | 0.34 | -0.46 | -0.01 | 0.884 |
| PRDM2 | S1501 | -0.01 | -0.17 | 0.31 | 0.08 | -0.24 | -0.04 | 0.542 |
| PRDM2 | S447 | 0.23 | 0.25 | -0.33 | 0.06 | -0.15 | -0.16 | 0.543 |
| PRDM2 | S643 | 0.21 | 0.02 | 0.21 | -0.30 | 0.11 | -0.37 | 0.107 |
| PRDM4 | T641 | -0.13 | -0.16 | -0.22 | 0.14 | 0.11 | 0.20 | 0.001 |
| PRDX2 | S112 | 0.07 | 0.05 | -0.11 | 0.04 | -0.02 | -0.03 | 0.926 |
| PRICKLE2 | S753 | -0.32 | -0.45 | -0.04 | 0.29 | -0.02 | 0.36 | 0.044 |
| PRICKLE2 | T/S/Y | 0.19 | -0.05 | 0.09 | -0.20 | -0.01 | -0.05 | 0.139 |
| PRICKLE3 | S453 | -0.32 | -0.13 | -0.49 | 0.26 | -0.27 | 0.62 | 0.135 |
| PRIM1 | S/T | -0.41 | -0.43 | -0.25 | 0.03 | 0.61 | 0.16 | 0.027 |
| PRKAB1 | S24 | 0.13 | 0.22 | 0.05 | -0.28 | 0.09 | -0.30 | 0.092 |
| PRKAB1 | S/Y | 0.02 | 0.27 | 0.34 | -0.45 | -0.27 | -0.06 | 0.035 |
| PRKAB1 | S/Y | 0.07 | 0.22 | 0.47 | -0.47 | -0.17 | -0.34 | 0.015 |
| PRKAB2 | S/Y | 0.03 | -0.12 | -0.03 | -0.02 | 0.13 | 0.00 | 0.296 |
| PRKACA | S339 | -0.03 | -0.21 | -0.25 | 0.30 | -0.11 | 0.22 | 0.103 |
| PRKACB | S339 | -0.23 | 0.03 | -0.11 | 0.11 | 0.14 | 0.03 | 0.078 |
| PRKAG2 | S/Y/T | 0.05 | 0.07 | 0.40 | -0.60 | 0.44 | -0.79 | 0.287 |
| PRKAR1A | S83 | -0.04 | -0.01 | 0.03 | 0.01 | -0.09 | 0.10 | 0.878 |
| PRKAR1A | S83 | 0.15 | 0.02 | -0.03 | -0.38 | -0.19 | 0.32 | 0.582 |
| PRKAR1A | S83 | 0.12 | 0.18 | 0.14 | -0.51 | 0.15 | -0.20 | 0.153 |
| PRKAR1A | S83 | 0.19 | 0.24 | 0.33 | -0.39 | -0.02 | -0.58 | 0.027 |
| PRKAR2A | S99 | -0.48 | -0.47 | -0.30 | 0.24 | -0.08 | 0.70 | 0.039 |
| PRKAR2A | S99 | -0.40 | -0.28 | -0.46 | 0.19 | 0.15 | 0.53 | 0.007 |
| PRKAR2A | S78; S80 | -0.17 | -0.34 | -0.33 | 0.17 | 0.09 | 0.42 | 0.011 |
| PRKAR2A | S78; S80 | -0.09 | -0.01 | -0.09 | 0.07 | 0.13 | -0.02 | 0.067 |
| PRKAR2A | S78 | -0.04 | 0.12 | -0.01 | -0.02 | 0.03 | -0.09 | 0.465 |
| PRKAR2A | S80 | 0.08 | 0.14 | 0.08 | -0.06 | 0.04 | -0.33 | 0.127 |
| PRKCD | S304 | -0.29 | -0.30 | -0.01 | -0.09 | 0.09 | 0.45 | 0.13 |
| PRKCD | S645 | 0.10 | -0.01 | -0.18 | 0.20 | -0.04 | -0.09 | 0.674 |

| | | | | | | | | |
|---|---|---|---|---|---|---|---|---|
| PRKCE | S | 0.05 | -0.66 | 0.04 | 0.37 | -0.13 | 0.13 | 0.321 |
| PRKDC | T2638 | -0.44 | -0.45 | -0.45 | 0.27 | 0.38 | 0.37 | 3E-05 |
| PRKDC | S2624 | -0.25 | -0.58 | -0.28 | 0.14 | 0.47 | 0.25 | 0.01 |
| PRKDC | S3205 | -0.26 | -0.22 | -0.27 | 0.24 | 0.25 | 0.14 | 3E-04 |
| PRKDC | T3299 | -0.01 | -0.41 | -0.37 | -0.11 | 0.23 | 0.46 | 0.093 |
| PRKDC | S/T/Y | -0.24 | -0.16 | -0.22 | 0.18 | 0.22 | 0.15 | 3E-04 |
| PRKDC | T/S/Y | -0.22 | -0.16 | -0.18 | 0.12 | 0.13 | 0.24 | 0.001 |
| PRKDC | S3821 | -0.32 | -0.45 | 0.14 | -0.04 | -0.03 | 0.49 | 0.233 |
| PRKDC | S3205 | -0.23 | -0.17 | -0.10 | 0.15 | 0.14 | 0.16 | 0.001 |
| PRKDC | S3205 | -0.07 | -0.27 | -0.10 | 0.17 | 0.16 | 0.06 | 0.019 |
| PRKDC | S3205 | -0.35 | -0.26 | 0.13 | 0.07 | -0.18 | 0.43 | 0.309 |
| PRKDC | S3432 | 0.28 | -0.12 | -0.24 | -0.03 | 0.03 | 0.02 | 0.843 |
| PRKDC | S4026 | -0.08 | 0.25 | -0.15 | 0.26 | -0.09 | -0.28 | 0.846 |
| PRKDC | S3366 | 0.20 | 0.14 | 0.08 | -0.42 | 0.13 | -0.23 | 0.128 |
| PRKRA | S18 | 0.02 | -0.16 | -0.06 | 0.11 | 0.25 | -0.21 | 0.473 |
| PRKRA | S18 | -0.03 | -0.07 | -0.09 | 0.27 | 0.18 | -0.33 | 0.613 |
| PRKRIP1 | T171; S179; T181 | -0.28 | -0.02 | 0.00 | 0.03 | 0.00 | 0.22 | 0.19 |
| PRLR | S356 | -0.25 | 0.22 | -0.26 | 0.05 | -0.22 | 0.35 | 0.522 |
| PROSER1 | S613 | 0.24 | 0.06 | 0.24 | -0.16 | -0.10 | -0.38 | 0.021 |
| PRPF3 | S619 | -0.06 | -0.16 | -0.13 | 0.16 | -0.19 | 0.31 | 0.237 |
| PRPF3 | S619 | 0.10 | 0.12 | -0.22 | 0.08 | -0.11 | 0.00 | 0.952 |
| PRPF3 | S619 | 0.19 | -0.01 | -0.03 | 0.04 | 0.00 | -0.22 | 0.358 |
| PRPF3 | S619 | 0.18 | 0.12 | 0.07 | 0.03 | -0.07 | -0.41 | 0.122 |
| PRPF31 | S450 | 0.19 | -0.06 | 0.23 | -0.34 | 0.20 | -0.34 | 0.242 |
| PRPF31 | S450; T455 | 0.12 | 0.09 | 0.27 | -0.22 | -0.10 | -0.22 | 0.007 |
| PRPF31 | T455 | 0.38 | 0.22 | 0.21 | 0.30 | -1.05 | -0.59 | 0.147 |
| PRPF38A | S194 | 0.01 | -0.01 | -0.14 | -0.11 | 0.05 | 0.17 | 0.434 |
| PRPF38A | S279; S281 | 0.18 | 0.08 | -0.11 | 0.07 | -0.05 | -0.21 | 0.384 |
| PRPF38A | S193; S194 | 0.09 | 0.06 | 0.13 | -0.12 | 0.05 | -0.23 | 0.082 |
| PRPF38A | S193; S194 | 0.11 | 0.07 | 0.13 | -0.12 | 0.02 | -0.24 | 0.047 |
| PRPF38A | S193 | 0.12 | 0.11 | 0.15 | -0.14 | 0.01 | -0.31 | 0.041 |
| PRPF38B | S491 | -0.05 | 0.02 | -0.07 | 0.14 | 0.00 | -0.06 | 0.446 |
| PRPF38B | S | 0.28 | -0.08 | -0.18 | 0.00 | 0.10 | -0.16 | 0.862 |
| PRPF38B | S527; S529 | 0.13 | 0.09 | 0.15 | 0.09 | -0.30 | -0.23 | 0.094 |
| PRPF38B | S491 | 0.18 | 0.13 | 0.12 | 0.00 | -0.18 | -0.33 | 0.033 |
| PRPF38B | S529 | 0.13 | 0.18 | 0.21 | -0.13 | -0.20 | -0.27 | 0.001 |
| PRPF38B | S527 | 0.25 | 0.24 | 0.26 | -0.20 | -0.16 | -0.57 | 0.013 |
| PRPF38B | S491 | 0.36 | 0.35 | 0.14 | -0.23 | -0.14 | -0.79 | 0.036 |
| PRPF40A | S787 | -0.09 | -0.14 | -0.32 | -0.05 | -0.10 | 0.54 | 0.223 |
| PRPF40A | S885; S888 | 0.28 | 0.11 | 0.08 | -0.53 | 0.29 | -0.45 | 0.222 |
| PRPF40A | S933; S938 | 0.23 | 0.24 | 0.10 | -0.16 | -0.18 | -0.33 | 0.004 |
| PRPF40A | T373 | 0.19 | 0.14 | 0.25 | -0.18 | -0.15 | -0.35 | 0.004 |
| PRPF40A | T932; S933; S938 | 0.19 | 0.24 | 0.19 | -0.22 | -0.14 | -0.38 | 0.003 |
| PRPF40A | T932; S935; S938 | 0.28 | 0.34 | 0.18 | -0.25 | -0.11 | -0.68 | 0.027 |
| PRPF40A | S938 | 0.29 | 0.32 | 0.18 | -0.32 | -0.06 | -0.66 | 0.028 |
| PRPF40A | S723 | 0.35 | 0.34 | 0.12 | -0.16 | -0.47 | -0.42 | 0.007 |
| PRPF40A | T373 | 0.31 | 0.31 | 0.21 | -0.28 | -0.06 | -0.78 | 0.038 |
| PRPF40A | T373 | 0.27 | 0.20 | 0.41 | -0.23 | -0.19 | -0.74 | 0.023 |

| | | | | | | | | |
|---|---|---|---|---|---|---|---|---|
| **PRPF40A** | **S938** | 0.34 | 0.41 | 0.21 | -0.30 | -0.07 | -1.02 | 0.055 |
| **PRPF40A** | **S935; S938** | 0.35 | 0.48 | 0.31 | -0.42 | -0.18 | -1.10 | 0.028 |
| **PRPF40A** | **S834** | 0.15 | 0.34 | 0.67 | -0.46 | -0.41 | -0.84 | 0.01 |
| **PRPF40A** | **S883; S888** | 0.50 | 0.57 | 0.33 | -0.69 | -0.24 | -1.42 | 0.024 |
| **PRPF40B** | **S764** | -0.11 | -0.25 | 0.13 | 0.00 | -0.09 | 0.26 | 0.438 |
| **PRPF40B** | **S832** | 0.40 | 0.27 | -0.27 | -0.10 | -0.34 | -0.12 | 0.212 |
| **PRPF40B** | **S852** | 0.19 | 0.09 | 0.21 | -0.24 | -0.09 | -0.24 | 0.005 |
| **PRPF40B** | **S832** | 0.29 | 0.10 | 0.12 | -0.03 | -0.14 | -0.45 | 0.052 |
| **PRPF4B** | **S839; Y849** | -0.77 | -0.72 | -0.26 | 0.63 | -0.12 | 0.59 | 0.032 |
| **PRPF4B** | **S569** | -0.09 | -0.18 | -0.17 | 0.19 | 0.10 | 0.11 | 0.002 |
| **PRPF4B** | **S518; S519; S520** | -0.04 | -0.07 | -0.29 | -0.03 | -0.11 | 0.43 | 0.279 |
| **PRPF4B** | **S257** | -0.06 | -0.08 | -0.11 | 0.13 | -0.08 | 0.18 | 0.131 |
| **PRPF4B** | **S20; S23; S32** | -0.08 | -0.14 | 0.00 | 0.20 | 0.12 | -0.14 | 0.293 |
| **PRPF4B** | **Y/S** | -0.16 | -0.01 | -0.06 | -0.21 | -0.24 | 0.53 | 0.699 |
| **PRPF4B** | **S277** | 0.11 | 0.07 | -0.36 | 0.05 | -0.01 | 0.08 | 0.56 |
| **PRPF4B** | **T576; S578; S580** | -0.07 | 0.11 | -0.18 | 0.01 | -0.14 | 0.22 | 0.604 |
| **PRPF4B** | **S578; S580** | -0.01 | -0.07 | -0.03 | 0.00 | -0.07 | 0.17 | 0.373 |
| **PRPF4B** | **S87; S93** | 0.03 | -0.02 | -0.11 | 0.13 | -0.05 | 0.01 | 0.415 |
| **PRPF4B** | **Y/S** | -0.18 | -0.03 | 0.04 | -0.31 | -0.16 | 0.49 | 0.821 |
| **PRPF4B** | **S20; S23; S32** | 0.02 | -0.07 | -0.04 | -0.10 | 0.03 | 0.14 | 0.519 |
| **PRPF4B** | **S565; S/Y/T** | -0.25 | 0.11 | 0.07 | -0.04 | 0.03 | 0.05 | 0.767 |
| **PRPF4B** | **S294** | 0.03 | -0.03 | -0.05 | 0.05 | 0.15 | -0.17 | 0.829 |
| **PRPF4B** | **S** | 0.06 | -0.07 | -0.01 | 0.08 | -0.04 | -0.03 | 0.875 |
| **PRPF4B** | **T847** | -0.01 | 0.02 | -0.02 | -0.05 | 0.06 | 0.00 | 0.827 |
| **PRPF4B** | **T847** | 0.00 | 0.11 | -0.07 | 0.16 | -0.22 | -0.01 | 0.78 |
| **PRPF4B** | **S368** | 0.03 | 0.00 | 0.03 | 0.01 | 0.10 | -0.19 | 0.639 |
| **PRPF4B** | **S277** | 0.10 | 0.04 | -0.06 | 0.08 | -0.09 | -0.09 | 0.475 |
| **PRPF4B** | **S93** | 0.14 | 0.07 | -0.14 | 0.10 | 0.08 | -0.29 | 0.719 |
| **PRPF4B** | **S569** | -0.02 | 0.21 | -0.11 | -0.05 | -0.05 | 0.00 | 0.565 |
| **PRPF4B** | **S427; S431** | 0.13 | 0.01 | -0.06 | 0.04 | -0.02 | -0.12 | 0.438 |
| **PRPF4B** | **S292; S294** | 0.08 | 0.00 | 0.01 | 0.01 | -0.11 | 0.01 | 0.266 |
| **PRPF4B** | **S93** | 0.17 | 0.08 | -0.11 | 0.11 | -0.05 | -0.26 | 0.455 |
| **PRPF4B** | **S20; S23** | 0.27 | 0.17 | -0.30 | -0.04 | -0.10 | -0.07 | 0.545 |
| **PRPF4B** | **S427; S431** | 0.12 | 0.04 | 0.02 | -0.06 | -0.03 | -0.11 | 0.027 |
| **PRPF4B** | **S20; S23** | 0.32 | 0.02 | -0.18 | 0.16 | -0.03 | -0.40 | 0.545 |
| **PRPF4B** | **S578; S580** | -0.01 | 0.02 | 0.23 | -0.12 | -0.16 | 0.01 | 0.138 |
| **PRPF4B** | **S277** | 0.20 | 0.09 | -0.04 | -0.03 | -0.03 | -0.23 | 0.139 |
| **PRPF4B** | **S520; S** | 0.07 | 0.10 | 0.09 | -0.32 | -0.09 | 0.10 | 0.202 |
| **PRPF4B** | **S20; S23** | -0.03 | 0.26 | 0.05 | -0.14 | 0.08 | -0.27 | 0.218 |
| **PRPF4B** | **S257** | 0.22 | 0.12 | -0.02 | 0.07 | -0.22 | -0.22 | 0.127 |
| **PRPF4B** | **S580** | 0.07 | 0.21 | 0.06 | -0.09 | -0.03 | -0.28 | 0.052 |
| **PRPF4B** | **S144** | 0.00 | 0.21 | 0.15 | -0.28 | -0.18 | 0.04 | 0.08 |
| **PRPF4B** | **S277** | 0.11 | 0.16 | 0.10 | -0.02 | -0.05 | -0.35 | 0.067 |
| **PRPF4B** | **S257** | 0.21 | 0.14 | 0.05 | 0.03 | -0.01 | -0.54 | 0.175 |
| **PRPF4B** | **S431** | 0.13 | 0.14 | 0.14 | -0.09 | 0.11 | -0.54 | 0.182 |
| **PRPF4B** | **S431; S437** | 0.13 | 0.15 | 0.15 | -0.09 | 0.05 | -0.47 | 0.112 |
| **PRPF4B** | **S20; S23** | 0.13 | 0.22 | 0.09 | -0.09 | -0.16 | -0.26 | 0.008 |
| **PRPF4B** | **S20; S23** | -0.07 | 0.58 | -0.18 | 0.07 | -0.65 | -0.03 | 0.393 |

| | | | | | | | | |
|---|---|---|---|---|---|---|---|---|
| **PRPF4B** | **S330** | 0.25 | 0.28 | -0.10 | -0.04 | -0.11 | -0.39 | 0.115 |
| **PRPF4B** | **S23** | 0.21 | 0.16 | 0.12 | -0.12 | -0.02 | -0.44 | 0.051 |
| **PRPF4B** | **S20; S23** | 0.60 | 0.06 | -0.09 | -0.08 | -0.23 | -0.51 | 0.133 |
| **PRPF4B** | **S23** | 0.17 | 0.39 | 0.11 | -0.21 | -0.11 | -0.52 | 0.028 |
| **PRPF4B** | **S93** | 0.12 | 0.32 | 0.22 | -0.27 | 0.05 | -0.65 | 0.072 |
| **PRPF4B** | **S431; S437** | 0.24 | 0.22 | 0.23 | -0.27 | 0.03 | -0.68 | 0.06 |
| **PRPF4B** | **S93** | 0.28 | 0.24 | 0.13 | 0.03 | -0.05 | -0.96 | 0.166 |
| **PRPF4B** | **S144** | 0.20 | 0.30 | 0.25 | -0.27 | -0.07 | -0.64 | 0.027 |
| **PRPF4B** | **S144** | 0.25 | 0.37 | 0.30 | -0.30 | -0.22 | -0.70 | 0.01 |
| **PRPF4B** | **S144** | 0.31 | 0.45 | 0.39 | -0.54 | -0.11 | -1.12 | 0.031 |
| **PRPF6** | **T266** | 0.23 | -0.63 | -0.33 | 0.29 | 0.15 | 0.07 | 0.188 |
| **PRPF6** | **T266** | 0.05 | -0.15 | 0.02 | -0.24 | 0.01 | 0.25 | 0.848 |
| **PRPF8** | **T2064** | -0.34 | 0.41 | -0.05 | 0.04 | -0.35 | 0.14 | 0.824 |
| **PRPSAP2** | **S227** | -0.30 | -0.34 | 0.32 | 0.04 | -0.20 | 0.33 | 0.564 |
| **PRR11** | **S307** | 0.23 | 0.22 | 0.15 | -0.22 | 0.15 | -0.76 | 0.144 |
| **PRR12** | **S332** | -0.17 | -0.33 | -0.10 | 0.14 | 0.05 | 0.32 | 0.023 |
| **PRR12** | **T1561; S1568** | -0.19 | -0.17 | -0.04 | 0.20 | -0.06 | 0.20 | 0.069 |
| **PRR12** | **S917** | -0.08 | -0.15 | -0.14 | 0.25 | -0.13 | 0.19 | 0.128 |
| **PRR12** | **T1561; S1568** | -0.23 | 0.08 | -0.10 | -0.06 | 0.04 | 0.23 | 0.278 |
| **PRR12** | **S1308** | -0.10 | 0.00 | -0.05 | 0.13 | -0.17 | 0.16 | 0.457 |
| **PRR12** | **T1705** | 0.04 | 0.08 | -0.15 | -0.09 | -0.05 | 0.15 | 0.901 |
| **PRR12** | **S865** | -0.12 | 0.01 | 0.26 | -0.06 | -0.03 | -0.09 | 0.377 |
| **PRR12** | **S/T** | -0.20 | 0.51 | -0.20 | 0.11 | -0.20 | -0.17 | 0.654 |
| **PRR12** | **S651** | -0.05 | 0.17 | 0.15 | -0.27 | 0.12 | -0.17 | 0.228 |
| **PRR12** | **S1568** | 0.22 | 0.25 | -0.04 | -0.10 | -0.07 | -0.35 | 0.066 |
| **PRR12** | **S1135** | 0.27 | 0.11 | 0.08 | -0.66 | 0.22 | -0.22 | 0.231 |
| **PRR12** | **S1568** | 0.19 | 0.27 | 0.12 | -0.18 | -0.13 | -0.39 | 0.01 |
| **PRR14** | **T363** | -0.30 | -0.08 | -0.24 | 0.23 | 0.22 | 0.08 | 0.01 |
| **PRR14** | **S277** | -0.14 | -0.15 | 0.11 | -0.07 | -0.03 | 0.24 | 0.456 |
| **PRR14** | **S277** | 0.04 | -0.03 | 0.10 | -0.04 | -0.02 | -0.06 | 0.106 |
| **PRR14** | **S93** | 0.10 | 0.06 | -0.02 | -0.08 | -0.13 | 0.05 | 0.197 |
| **PRR14** | **S277** | 0.06 | 0.06 | 0.06 | -0.12 | 0.10 | -0.18 | 0.218 |
| **PRR14** | **S304** | 0.22 | -0.06 | 0.16 | -0.17 | 0.08 | -0.30 | 0.172 |
| **PRR14L** | **S1994** | -0.12 | -0.07 | -0.26 | -0.27 | 0.27 | 0.34 | 0.251 |
| **PRR15** | **S114** | 0.15 | 0.17 | -0.15 | -0.14 | -0.38 | 0.24 | 0.511 |
| **PRR15** | **S49** | 0.20 | -0.01 | 0.01 | -0.10 | -0.40 | 0.21 | 0.446 |
| **PRR15** | **S49** | 0.17 | 0.15 | 0.03 | -0.02 | -0.30 | -0.08 | 0.053 |
| **PRR36** | **S110** | 0.08 | -0.04 | 0.26 | -0.25 | -0.18 | 0.06 | 0.159 |
| **PRRC2A** | **S1306** | 0.04 | -0.66 | -0.45 | 0.39 | 0.05 | 0.34 | 0.057 |
| **PRRC2A** | **S1306** | -0.29 | -0.18 | -0.54 | 0.18 | 0.40 | 0.20 | 0.009 |
| **PRRC2A** | **S456** | -0.13 | -0.21 | -0.44 | 0.15 | 0.28 | 0.21 | 0.01 |
| **PRRC2A** | **S1306** | -0.19 | -0.16 | -0.26 | 0.24 | 0.01 | 0.26 | 0.013 |
| **PRRC2A** | **S1089; S1092** | -0.39 | 0.06 | -0.26 | 0.26 | -0.07 | 0.29 | 0.112 |
| **PRRC2A** | **S1325** | -0.33 | -0.37 | 0.21 | 0.04 | 0.01 | 0.31 | 0.254 |
| **PRRC2A** | **S380** | -0.05 | -0.13 | -0.17 | 0.21 | -0.05 | 0.15 | 0.063 |
| **PRRC2A** | **S1306; S/T** | -0.09 | -0.19 | 0.02 | 0.18 | -0.05 | 0.10 | 0.148 |
| **PRRC2A** | **S1306** | -0.08 | -0.20 | 0.05 | 0.02 | -0.05 | 0.23 | 0.263 |
| **PRRC2A** | **T825** | -0.20 | -0.04 | 0.00 | -0.32 | -0.24 | 0.60 | 0.775 |

| | | | | | | | | |
|---|---|---|---|---|---|---|---|---|
| **PRRC2A** | **T1347** | -0.04 | -0.07 | 0.03 | 0.07 | 0.14 | -0.16 | 0.655 |
| **PRRC2A** | **S1314; S1327; S1328** | -0.14 | 0.22 | -0.13 | 0.01 | -0.03 | 0.04 | 0.82 |
| **PRRC2A** | **S** | -0.19 | 0.06 | 0.10 | -0.05 | 0.05 | 0.00 | 0.913 |
| **PRRC2A** | **T878** | -0.09 | -0.06 | 0.13 | -0.03 | -0.14 | 0.17 | 0.984 |
| **PRRC2A** | **S1306; S/T** | -0.30 | 0.31 | -0.04 | 0.02 | 0.02 | -0.08 | 0.994 |
| **PRRC2A** | **S342; S350** | 0.00 | 0.04 | -0.05 | 0.00 | -0.05 | 0.04 | 0.969 |
| **PRRC2A** | **S1004** | -0.14 | 0.03 | 0.14 | -0.18 | -0.03 | 0.15 | 0.821 |
| **PRRC2A** | **S1306; S1327; S1328** | -0.10 | 0.17 | 0.03 | -0.01 | -0.07 | -0.04 | 0.414 |
| **PRRC2A** | **S1089** | 0.06 | 0.07 | 0.07 | -0.03 | -0.09 | -0.11 | 0.005 |
| **PRRC2A** | **T610** | 0.07 | 0.18 | 0.06 | -0.11 | -0.01 | -0.23 | 0.039 |
| **PRRC2A** | **S1219** | 0.05 | 0.17 | 0.20 | -0.24 | -0.10 | -0.14 | 0.008 |
| **PRRC2A** | **S1085; S1089** | 0.06 | 0.12 | 0.24 | -0.20 | 0.04 | -0.35 | 0.068 |
| **PRRC2A** | **T782** | 0.14 | 0.29 | 0.02 | -0.22 | 0.03 | -0.37 | 0.075 |
| **PRRC2A** | **S380** | 0.16 | 0.09 | 0.21 | -0.04 | -0.01 | -0.52 | 0.117 |
| **PRRC2A** | **S** | -0.06 | 0.27 | 0.11 | 0.26 | 0.23 | -1.37 | 0.505 |
| **PRRC2B** | **S556** | -0.53 | 0.11 | -0.20 | 0.07 | 0.08 | 0.32 | 0.145 |
| **PRRC2B** | **S1136** | -0.37 | 0.04 | -0.06 | 0.03 | -0.35 | 0.52 | 0.518 |
| **PRRC2B** | **S388** | -0.02 | -0.06 | -0.12 | 0.03 | 0.07 | 0.09 | 0.023 |
| **PRRC2B** | **S907** | -0.25 | 0.04 | 0.05 | 0.00 | 0.33 | -0.25 | 0.689 |
| **PRRC2B** | **S1395** | 0.04 | 0.09 | -0.09 | -0.16 | 0.02 | 0.08 | 0.722 |
| **PRRC2B** | **S1395** | -0.15 | 0.26 | 0.02 | -0.09 | 0.08 | -0.17 | 0.507 |
| **PRRC2B** | **S480** | -0.02 | 0.08 | 0.08 | -0.05 | 0.20 | -0.35 | 0.523 |
| **PRRC2C** | **T1965** | -0.16 | -0.50 | -0.01 | 0.29 | 0.42 | -0.25 | 0.206 |
| **PRRC2C** | **S1503; S/T** | -0.31 | -0.08 | -0.21 | 0.16 | -0.22 | 0.50 | 0.183 |
| **PRRC2C** | **S1500; S1503** | 0.00 | -0.50 | -0.15 | -0.03 | -0.39 | 0.73 | 0.424 |
| **PRRC2C** | **T1965** | -0.13 | -0.15 | -0.08 | 0.19 | 0.14 | 0.01 | 0.015 |
| **PRRC2C** | **S1489; S1502; S1503** | -0.18 | 0.20 | -0.40 | 0.13 | 0.08 | 0.09 | 0.269 |
| **PRRC2C** | **T/S** | -0.12 | -0.08 | -0.08 | 0.14 | 0.01 | 0.12 | 0.013 |
| **PRRC2C** | **S1503; S/T** | -0.10 | -0.06 | -0.05 | -0.15 | -0.02 | 0.33 | 0.448 |
| **PRRC2C** | **S2105** | 0.02 | 0.01 | -0.17 | 0.07 | -0.10 | 0.15 | 0.441 |
| **PRRC2C** | **T1498; S1500; S1503** | -0.26 | 0.02 | 0.08 | -0.84 | 0.11 | 0.55 | 0.991 |
| **PRRC2C** | **S1242; S/T** | -0.03 | 0.01 | 0.03 | 0.03 | -0.19 | 0.13 | 0.889 |
| **PRRC2C** | **S1246** | 0.04 | 0.03 | 0.06 | -0.05 | -0.10 | 0.01 | 0.057 |
| **PRRC2C** | **T1498; S1500; S/T** | -0.17 | 0.40 | -0.13 | -0.20 | 0.15 | -0.15 | 0.655 |
| **PRRC2C** | **S2694** | 0.15 | 0.04 | -0.05 | -0.14 | -0.20 | 0.16 | 0.449 |
| **PRRC2C** | **S1503** | -0.01 | 0.14 | 0.17 | -0.18 | -0.01 | -0.14 | 0.052 |
| **PRRC2C** | **S878** | -0.04 | 0.30 | 0.05 | -0.04 | -0.34 | 0.00 | 0.198 |
| **PRRC2C** | **S1248; S1249** | 0.10 | 0.12 | 0.19 | -0.15 | -0.09 | -0.22 | 0.003 |
| **PRRC2C** | **S2694** | 0.21 | 0.07 | 0.15 | -0.18 | 0.01 | -0.34 | 0.047 |
| **PRRC2C** | **S1503** | 0.06 | 0.31 | 0.12 | -0.04 | -0.19 | -0.35 | 0.04 |
| **PRRC2C** | **S1248** | 0.18 | 0.25 | 0.21 | -0.23 | -0.15 | -0.38 | 0.003 |
| **PRSS12** | **S/T** | -0.42 | -0.43 | -0.67 | 0.80 | -0.56 | 0.54 | 0.148 |
| **PRUNE1** | **S321** | -0.20 | 0.18 | 0.04 | 0.04 | 0.06 | -0.15 | 0.875 |
| **PSEN1** | **S366; S367** | -0.10 | -0.15 | -0.08 | -0.03 | -0.17 | 0.44 | 0.374 |
| **PSEN1** | **S** | -0.22 | 0.00 | 0.06 | 0.00 | -0.07 | 0.20 | 0.443 |
| **PSEN1** | **S** | -0.18 | 0.02 | 0.02 | -0.07 | -0.32 | 0.43 | 0.799 |
| **PSEN1** | **S319; T327** | -0.10 | 0.14 | 0.05 | -0.04 | -0.19 | 0.11 | 0.547 |
| **PSEN1** | **S324** | -0.42 | 0.06 | 0.45 | 0.28 | -0.57 | -0.07 | 0.688 |

| | | | | | | | | |
|---|---|---|---|---|---|---|---|---|
| **PSEN1** | **S324** | -0.22 | 0.20 | 0.35 | 0.03 | -0.28 | -0.20 | 0.253 |
| **PSEN1** | **S43** | 0.05 | 0.17 | 0.21 | -0.08 | 0.02 | -0.46 | 0.108 |
| **PSEN2** | **S/T** | 0.26 | 0.31 | 0.27 | -0.67 | -0.01 | -0.43 | 0.029 |
| **PSEN2** | **S/T** | 0.09 | 0.54 | 0.21 | -0.55 | -0.14 | -0.45 | 0.022 |
| **PSIP1** | **T141** | -0.19 | -0.40 | -0.36 | 0.19 | 0.10 | 0.46 | 0.01 |
| **PSIP1** | **S206** | -0.03 | -0.13 | -0.10 | 0.01 | 0.11 | 0.12 | 0.027 |
| **PSIP1** | **S118; T122; S129** | 0.08 | -0.09 | -0.12 | 0.02 | -0.06 | 0.15 | 0.386 |
| **PSIP1** | **S/T** | 0.01 | 0.00 | -0.13 | 0.02 | 0.05 | 0.04 | 0.193 |
| **PSIP1** | **T122** | 0.12 | 0.03 | 0.00 | -0.10 | 0.00 | -0.05 | 0.09 |
| **PSIP1** | **T122; S129** | 0.13 | 0.05 | 0.01 | -0.11 | 0.07 | -0.17 | 0.174 |
| **PSIP1** | **S106** | 0.08 | 0.09 | 0.03 | -0.15 | -0.05 | 0.00 | 0.047 |
| **PSIP1** | **S273; S275** | 0.02 | 0.14 | 0.15 | 0.12 | -0.07 | -0.44 | 0.234 |
| **PSIP1** | **S106; S118; T122** | 0.19 | 0.09 | 0.09 | -0.24 | 0.11 | -0.31 | 0.113 |
| **PSMA5** | **S56** | -0.38 | -0.43 | -0.45 | 0.29 | 0.48 | 0.18 | 0.001 |
| **PSMA5** | **S16** | -0.21 | -0.41 | -0.19 | 0.39 | 0.19 | 0.08 | 0.012 |
| **PSMA5** | **S56** | -0.10 | -0.27 | -0.25 | 0.16 | 0.24 | 0.13 | 0.004 |
| **PSMB10** | **S/T** | -0.16 | -0.49 | 0.28 | 0.08 | 0.04 | 0.14 | 0.403 |
| **PSMC6** | **S244** | -0.11 | -0.28 | -0.26 | 0.16 | -0.29 | 0.57 | 0.222 |
| **PSMD1** | **T273** | -0.16 | -0.31 | -0.09 | 0.06 | -0.02 | 0.40 | 0.081 |
| **PSMD1** | **T273** | -0.01 | -0.13 | -0.34 | -0.12 | 0.03 | 0.44 | 0.222 |
| **PSMD2** | **S16** | -0.19 | -0.37 | -0.20 | 0.17 | 0.21 | 0.26 | 0.002 |
| **PSMD2** | **S16** | 0.17 | -0.17 | -0.54 | -0.04 | 0.27 | 0.15 | 0.241 |
| **PSMD2** | **S16** | -0.07 | -0.13 | -0.28 | 0.15 | 0.21 | 0.06 | 0.018 |
| **PSMD2** | **S16** | -0.27 | 0.26 | -0.13 | 0.44 | -0.05 | -0.45 | 0.929 |
| **PSPC1** | **S** | 0.21 | -0.09 | -0.19 | -0.17 | -0.13 | 0.29 | 0.915 |
| **PSPC1** | **S477** | 0.30 | -0.14 | 0.03 | 0.12 | -0.11 | -0.28 | 0.412 |
| **PTBP1** | **S459** | -0.14 | -0.15 | -0.18 | 0.25 | -0.11 | 0.26 | 0.077 |
| **PTBP3** | **S454** | -0.23 | -0.54 | -0.04 | 0.43 | 0.16 | 0.03 | 0.064 |
| **PTBP3** | **S56** | 0.05 | -0.05 | 0.00 | 0.00 | -0.14 | 0.12 | 0.944 |
| **PTCD3** | **T670; S671; S681** | -0.02 | 0.10 | -0.08 | -0.13 | 0.03 | 0.09 | 0.97 |
| **PTDSS1** | **S442** | 0.19 | 0.19 | -0.39 | -0.10 | -0.12 | 0.14 | 0.927 |
| **PTDSS1** | **S11** | 0.13 | 0.11 | -0.07 | -0.02 | 0.08 | -0.25 | 0.374 |
| **PTDSS1** | **S417** | 0.05 | 0.06 | 0.23 | -0.36 | 0.11 | -0.16 | 0.16 |
| **PTDSS2** | **S16** | -0.18 | -0.18 | -0.19 | 0.26 | -0.26 | 0.40 | 0.187 |
| **PTDSS2** | **S16** | 0.21 | 0.02 | 0.35 | -0.78 | -0.34 | 0.24 | 0.191 |
| **PTGES3** | **S113** | -0.19 | -0.17 | -0.17 | 0.17 | 0.13 | 0.17 | 2E-05 |
| **PTGES3** | **S113** | -0.10 | -0.08 | -0.22 | 0.14 | 0.17 | 0.04 | 0.012 |
| **PTOV1** | **S36** | -0.08 | -0.22 | 0.46 | -0.09 | -0.31 | 0.09 | 0.538 |
| **PTPN1** | **S352** | -0.39 | -0.25 | -0.07 | 0.65 | -0.01 | -0.17 | 0.215 |
| **PTPN1** | **S352** | -0.08 | -0.11 | -0.19 | 0.05 | -0.05 | 0.33 | 0.115 |
| **PTPN1** | **S50** | -0.01 | -0.09 | 0.00 | 0.00 | -0.06 | 0.15 | 0.414 |
| **PTPN1** | **S386** | 0.10 | -0.02 | -0.03 | -0.03 | -0.18 | 0.14 | 0.711 |
| **PTPN1** | **S352** | 0.13 | 0.00 | -0.03 | -0.20 | 0.15 | -0.08 | 0.532 |
| **PTPN12** | **S449** | -0.04 | -0.05 | -0.26 | 0.28 | 0.07 | -0.06 | 0.156 |
| **PTPN12** | **T519** | 0.17 | 0.25 | 0.15 | -0.05 | -0.39 | -0.23 | 0.016 |
| **PTPN12** | **S435** | 0.53 | 0.19 | 0.15 | -0.33 | -0.25 | -0.60 | 0.013 |
| **PTPN14** | **S803** | -0.49 | 0.06 | -0.12 | -0.03 | 0.23 | 0.23 | 0.153 |
| **PTPN14** | **S803** | -0.49 | 0.06 | -0.12 | -0.03 | 0.23 | 0.23 | 0.153 |

| | | | | | | | | |
|---|---|---|---|---|---|---|---|---|
| **PTPN14** | **S593** | 0.11 | 0.08 | -0.17 | 0.01 | -0.01 | -0.04 | 0.838 |
| **PTPN14** | **S314** | 0.14 | 0.10 | 0.44 | -0.22 | -0.04 | -0.64 | 0.065 |
| **PTPN14** | **S314** | 0.25 | 0.32 | 0.31 | 0.02 | 0.39 | -1.50 | 0.319 |
| **PTPN2** | **S304** | 0.10 | 0.13 | 0.12 | 0.04 | -0.10 | -0.35 | 0.091 |
| **PTPN2** | **S304** | 0.17 | 0.28 | 0.10 | -0.22 | -0.24 | -0.18 | 0.002 |
| **PTPN3** | **S359** | -0.04 | 0.04 | -0.14 | 0.11 | 0.04 | -0.02 | 0.25 |
| **PTPN3** | **S372** | -0.17 | 0.17 | -0.08 | 0.03 | -0.06 | 0.09 | 0.694 |
| **PTPRA** | **S189** | 0.10 | 0.11 | 0.03 | -0.12 | -0.15 | 0.01 | 0.037 |
| **PTPRK** | **S835** | -0.32 | 0.20 | -0.25 | -0.17 | 0.20 | 0.22 | 0.379 |
| **PTPRK** | **S856** | -0.03 | 0.07 | 0.05 | -0.04 | 0.03 | -0.09 | 0.244 |
| **PUM1** | **S709** | -0.07 | 0.17 | -0.22 | -0.19 | 0.09 | 0.17 | 0.715 |
| **PUM1** | **S709** | 0.11 | 0.08 | -0.06 | -0.09 | 0.05 | -0.10 | 0.272 |
| **PUM2** | **S136** | -0.17 | -0.09 | -0.05 | 0.41 | -0.36 | 0.14 | 0.514 |
| **PUM2** | **S136** | -0.07 | -0.06 | -0.07 | 0.19 | -0.01 | 0.00 | 0.131 |
| **PUM2** | **S587** | 0.05 | 0.06 | -0.25 | 0.18 | 0.13 | -0.24 | 0.697 |
| **PUM2** | **S82** | -0.08 | 0.02 | -0.06 | -0.22 | 0.06 | 0.25 | 0.657 |
| **PURB** | **S304** | -0.34 | -0.25 | 0.02 | 0.15 | 0.17 | 0.17 | 0.03 |
| **PURB** | **S/Y** | -0.08 | -0.35 | 0.09 | 0.14 | -0.10 | 0.22 | 0.276 |
| **PWP1** | **S50** | -0.64 | -0.02 | -0.04 | 0.04 | -0.41 | 0.70 | 0.421 |
| **PWP1** | **S50** | -0.38 | -0.06 | -0.11 | -0.01 | -0.29 | 0.62 | 0.367 |
| **PWP1** | **S485** | 0.05 | -0.07 | 0.06 | -0.29 | 0.01 | 0.20 | 0.797 |
| **PWP1** | **S50** | 0.47 | 0.28 | 0.01 | -0.52 | 0.06 | -0.60 | 0.07 |
| **PWP2** | **S891; S898; S902** | 0.03 | -0.03 | 0.09 | -0.08 | -0.18 | 0.15 | 0.574 |
| **PWP2** | **S898; S913** | 0.07 | 0.18 | 0.12 | -0.17 | -0.07 | -0.16 | 0.004 |
| **PWP2** | **S891; S898; S902** | 0.11 | 0.27 | 0.19 | -0.27 | -0.07 | -0.34 | 0.011 |
| **PWP2** | **S891; S898; S902** | 0.49 | 0.17 | 0.25 | -0.35 | -0.19 | -0.69 | 0.015 |
| **PWWP2A** | **S81** | -0.02 | -0.12 | -0.26 | 0.08 | -0.01 | 0.28 | 0.091 |
| **PWWP2A** | **S116; S119** | -0.11 | -0.09 | -0.07 | -0.06 | -0.07 | 0.35 | 0.306 |
| **PWWP2A** | **S584** | 0.17 | -0.13 | -0.02 | -0.42 | -0.06 | 0.34 | 0.83 |
| **PWWP2A** | **S119** | -0.07 | 0.03 | 0.28 | -0.10 | -0.05 | -0.13 | 0.18 |
| **PWWP2A** | **S119** | 0.02 | 0.12 | 0.34 | -0.26 | -0.02 | -0.32 | 0.053 |
| **PWWP2B** | **S186** | -0.16 | 0.31 | -0.13 | -0.20 | -0.03 | 0.14 | 0.845 |
| **PWWP3A** | **S540** | 0.03 | -0.06 | -0.24 | -0.08 | 0.05 | 0.25 | 0.256 |
| **PWWP3A** | **S235** | -0.12 | -0.02 | -0.12 | -0.17 | 0.00 | 0.36 | 0.402 |
| **PWWP3A** | **S/T** | 0.34 | -0.22 | -0.14 | -0.14 | 0.15 | -0.08 | 0.944 |
| **PYCR1** | **S301** | 0.03 | 0.24 | 0.14 | -0.12 | -0.19 | -0.16 | 0.01 |
| **PYCR2** | **S304** | 0.46 | -0.33 | 0.19 | -0.64 | 0.03 | 0.04 | 0.415 |
| **PYGO2** | **T302** | -0.22 | -0.14 | -0.50 | 0.30 | 0.15 | 0.25 | 0.012 |
| **QSER1** | **T1341; S1348** | -0.10 | -0.15 | -0.05 | -0.13 | -0.01 | 0.36 | 0.318 |
| **QSOX2** | **S578** | 0.08 | 0.02 | 0.19 | -0.10 | -0.11 | -0.10 | 0.016 |
| **R3HCC1** | **Y138** | 0.19 | 0.03 | -0.35 | -0.10 | -0.18 | 0.31 | 0.824 |
| **RAB10** | **S/T** | -0.38 | 0.18 | -0.37 | 0.77 | -0.38 | -0.23 | 0.585 |
| **RAB11FIP1** | **S342; S343; S345** | -0.13 | -0.02 | -0.13 | 0.05 | -0.09 | 0.27 | 0.201 |
| **RAB11FIP1** | **S202** | 0.02 | 0.05 | -0.03 | -0.02 | 0.02 | -0.06 | 0.364 |
| **RAB11FIP1** | **S345** | 0.00 | 0.03 | 0.06 | -0.02 | -0.01 | -0.06 | 0.081 |
| **RAB11FIP1** | **S500** | -0.07 | -0.05 | 0.26 | -0.17 | 0.15 | -0.19 | 0.486 |
| **RAB11FIP1** | **S528** | -0.05 | 0.08 | 0.15 | -0.13 | 0.07 | -0.14 | 0.246 |
| **RAB11FIP1** | **S339** | 0.14 | 0.26 | 0.10 | -0.23 | -0.20 | -0.15 | 0.003 |

| | | | | | | | | |
|---|---|---|---|---|---|---|---|---|
| RAB11FIP1 | S357 | 0.22 | 0.35 | -0.01 | -0.25 | 0.13 | -0.66 | 0.151 |
| RAB11FIP5 | S482 | -0.03 | -0.14 | 0.10 | 0.23 | -0.09 | -0.10 | 0.781 |
| RAB11FIP5 | S307 | -0.14 | 0.03 | 0.10 | 0.01 | 0.02 | -0.02 | 0.905 |
| RAB11FIP5 | S393 | -0.09 | -0.01 | 0.18 | -0.25 | 0.00 | 0.13 | 0.655 |
| RAB11FIP5 | S538 | 0.16 | 0.19 | 0.56 | -0.64 | 0.29 | -1.30 | 0.148 |
| RAB12 | S21 | 0.14 | -0.24 | 0.43 | 0.08 | -0.48 | -0.11 | 0.335 |
| RAB13 | S178 | 0.15 | 0.08 | 0.19 | -0.27 | -0.08 | -0.11 | 0.011 |
| RAB1B | S17 | 0.20 | -0.68 | 0.21 | -0.22 | -0.45 | 0.57 | 0.897 |
| RAB1B | S111 | 0.10 | 0.28 | 0.22 | -0.06 | -0.19 | -0.49 | 0.032 |
| RAB27A | S206 | -0.21 | -0.13 | -0.15 | 0.16 | -0.18 | 0.41 | 0.17 |
| RAB27A | S206 | -0.03 | -0.10 | 0.06 | -0.01 | -0.17 | 0.22 | 0.759 |
| RAB27A | S/T | -0.07 | 0.23 | 0.36 | 0.07 | -0.30 | -0.46 | 0.117 |
| RAB27B | S203 | -0.24 | -0.54 | 0.19 | 0.07 | 0.12 | 0.25 | 0.189 |
| RAB29 | S183 | 0.08 | -0.24 | -0.06 | -0.16 | 0.21 | 0.12 | 0.421 |
| RAB8A | S181 | 0.35 | 0.27 | -0.07 | -0.13 | 0.16 | -0.90 | 0.237 |
| RAB9A | S179 | 0.18 | 0.12 | -0.07 | -0.13 | -0.23 | 0.09 | 0.236 |
| RABAC1 | S/T | -0.30 | -0.32 | 0.06 | -0.12 | -0.02 | 0.53 | 0.25 |
| RABEP1 | S407 | -0.11 | -0.01 | -0.47 | 0.10 | 0.05 | 0.32 | 0.093 |
| RABEP2 | S/T | -0.44 | -0.23 | 0.15 | 0.32 | -0.15 | 0.20 | 0.247 |
| RABGAP1 | T996 | -0.13 | -0.05 | -0.26 | 0.10 | 0.21 | 0.08 | 0.018 |
| RABL6 | S596; T599 | 0.01 | -0.12 | -0.30 | -0.20 | 0.16 | 0.35 | 0.263 |
| RABL6 | S425; S427 | -0.22 | 0.01 | 0.05 | -0.18 | -0.07 | 0.34 | 0.66 |
| RABL6 | S596; T599 | -0.05 | -0.11 | 0.06 | -0.01 | -0.10 | 0.19 | 0.58 |
| RABL6 | S454 | 0.78 | -0.22 | -1.02 | -1.50 | 0.34 | 0.73 | 0.992 |
| RABL6 | S425; S427 | 0.09 | 0.06 | -0.06 | -0.12 | -0.01 | 0.04 | 0.413 |
| RABL6 | S/T | 0.04 | 0.01 | 0.04 | -0.17 | 0.04 | 0.02 | 0.357 |
| RABL6 | S492 | 0.05 | -0.04 | 0.11 | 0.06 | -0.42 | 0.17 | 0.616 |
| RABL6 | T599 | 0.09 | 0.13 | 0.03 | -0.21 | -0.08 | 0.02 | 0.079 |
| RABL6 | S427 | 0.13 | 0.07 | 0.11 | -0.06 | -0.15 | -0.13 | 0.003 |
| RABL6 | S402 | 0.36 | -0.10 | 0.11 | 0.01 | 0.05 | -0.62 | 0.29 |
| RACGAP1 | S | -0.04 | -0.13 | -0.02 | 0.39 | -0.71 | 0.26 | 0.903 |
| RACGAP1 | S206 | -0.25 | 0.38 | 0.06 | -0.28 | 0.07 | -0.08 | 0.495 |
| RACGAP1 | S203 | 0.31 | 0.06 | 0.05 | -0.24 | -0.18 | -0.07 | 0.038 |
| RACK1 | S276 | 0.04 | 0.06 | 0.20 | -0.11 | -0.05 | -0.17 | 0.029 |
| RAD18 | S99 | 0.10 | 0.47 | -0.48 | -0.06 | 0.08 | -0.30 | 0.692 |
| RAD18 | S471 | 0.06 | 0.04 | 0.36 | -0.12 | 0.00 | -0.44 | 0.111 |
| RAD21 | T623 | -0.14 | -0.34 | -0.36 | 0.29 | 0.01 | 0.37 | 0.018 |
| RAD21 | S459 | -0.12 | -0.23 | 0.11 | 0.23 | -0.28 | 0.20 | 0.542 |
| RAD21 | T394 | -0.10 | -0.03 | 0.13 | 0.05 | 0.05 | -0.11 | 0.981 |
| RAD23A | S123 | 0.07 | 0.02 | -0.10 | -0.15 | 0.08 | 0.05 | 0.971 |
| RAD23B | S160 | -0.14 | 0.07 | -0.06 | -0.02 | 0.19 | -0.06 | 0.453 |
| RAD23B | T/S | -0.06 | 0.28 | -0.15 | -0.42 | -0.18 | 0.37 | 0.741 |
| RAD50 | S635 | -0.40 | -0.22 | 0.06 | 0.26 | 0.28 | -0.10 | 0.14 |
| RAD51AP1 | S120 | 0.47 | -0.11 | -0.17 | 0.18 | -0.10 | -0.45 | 0.532 |
| RAD54L2 | T1260 | -0.29 | -0.70 | -0.43 | 0.68 | 0.03 | 0.27 | 0.024 |
| RAD54L2 | S1155 | -0.07 | 0.03 | -0.18 | 0.00 | 0.06 | 0.13 | 0.14 |
| RAD54L2 | T1260 | -0.15 | 0.26 | -0.11 | -0.08 | 0.22 | -0.22 | 0.89 |
| RAD9A | S/T | 0.15 | -0.08 | 0.09 | -0.77 | 0.01 | 0.37 | 0.624 |

| | | | | | | | | |
|---|---|---|---|---|---|---|---|---|
| **RAF1** | **S259** | -0.51 | 0.01 | 0.06 | -0.10 | -0.21 | 0.53 | 0.504 |
| **RAF1** | **T/S** | -0.12 | 0.01 | -0.08 | -0.05 | -0.03 | 0.24 | 0.307 |
| **RAF1** | **S259** | -0.10 | -0.10 | 0.28 | -0.01 | -0.10 | -0.01 | 0.647 |
| **RAI1** | **S1492** | -0.69 | -0.72 | -0.61 | 0.67 | -0.42 | 0.83 | 0.059 |
| **RAI1** | **S1374** | -0.18 | -0.70 | -0.38 | 0.27 | 0.04 | 0.58 | 0.031 |
| **RAI1** | **S805** | -0.47 | -0.32 | -0.26 | 0.23 | 0.47 | 0.11 | 0.007 |
| **RAI1** | **S1358** | -0.04 | -0.22 | -0.13 | -0.03 | 0.00 | 0.35 | 0.148 |
| **RAI1** | **S1374** | -0.03 | -0.21 | -0.15 | 0.09 | -0.09 | 0.32 | 0.146 |
| **RAI1** | **S805** | -0.19 | 0.01 | -0.19 | 0.22 | -0.07 | 0.16 | 0.116 |
| **RAI1** | **S683** | -0.03 | -0.13 | -0.16 | 0.12 | 0.09 | 0.07 | 0.008 |
| **RAI1** | **S1492** | -0.07 | -0.20 | -0.02 | 0.14 | 0.13 | 0.00 | 0.056 |
| **RAI1** | **S1358** | -0.06 | -0.07 | -0.10 | 0.02 | -0.01 | 0.20 | 0.081 |
| **RAI1** | **S** | 0.09 | -0.06 | -0.13 | 0.04 | 0.08 | -0.04 | 0.447 |
| **RAI1** | **S** | 0.00 | 0.03 | -0.12 | -0.07 | 0.02 | 0.13 | 0.456 |
| **RAI1** | **S1013** | -0.05 | 0.02 | -0.07 | -0.06 | -0.08 | 0.22 | 0.621 |
| **RAI1** | **S1064** | 0.05 | -0.15 | 0.01 | -0.32 | 0.20 | 0.15 | 0.841 |
| **RAI1** | **S1122; S** | 0.00 | -0.04 | 0.02 | -0.07 | 0.04 | 0.05 | 0.76 |
| **RAI1** | **Y** | 0.04 | 0.09 | -0.09 | 0.01 | -0.01 | -0.05 | 0.597 |
| **RAI1** | **S1266** | 0.10 | 0.02 | -0.07 | -0.14 | 0.03 | 0.04 | 0.613 |
| **RAI1** | **S1108; S** | 0.07 | -0.01 | 0.04 | -0.02 | -0.05 | -0.02 | 0.049 |
| **RAI1** | **T1136** | 0.16 | -0.06 | 0.08 | 0.00 | -0.20 | -0.01 | 0.228 |
| **RAI1** | **T/S** | 0.11 | 0.36 | -0.27 | 0.20 | -0.08 | -0.50 | 0.513 |
| **RAI1** | **S1133** | 0.19 | 0.06 | 0.09 | -0.25 | -0.04 | -0.09 | 0.034 |
| **RAI1** | **S339** | -0.15 | 0.49 | -0.01 | -0.02 | -0.57 | 0.06 | 0.36 |
| **RAI1** | **S1013** | 0.14 | 0.25 | 0.14 | -0.19 | -0.01 | -0.45 | 0.04 |
| **RAI1** | **S1266** | 0.25 | 0.25 | 0.25 | -0.35 | -0.10 | -0.48 | 0.007 |
| **RAI1** | **S/T** | 0.31 | 0.08 | 0.40 | -0.28 | -0.28 | -0.45 | 0.006 |
| **RAI1** | **S/T** | 0.30 | 0.24 | 0.25 | -0.26 | -0.09 | -0.68 | 0.027 |
| **RAI1** | **S1133** | 0.34 | 0.33 | 0.17 | -0.33 | -0.18 | -0.55 | 0.006 |
| **RAI14** | **S281** | 0.10 | -0.32 | -0.25 | -0.22 | -0.10 | 0.58 | 0.436 |
| **RAI14** | **S414; S419** | -0.23 | -0.03 | -0.01 | 0.11 | 0.16 | -0.04 | 0.14 |
| **RAI14** | **S293; T297** | 0.05 | 0.10 | -0.05 | 0.02 | -0.36 | 0.17 | 0.615 |
| **RAI14** | **S358** | -0.31 | 0.36 | -0.01 | -0.62 | 0.33 | 0.02 | 0.779 |
| **RALBP1** | **S92; S93** | -0.09 | -0.04 | 0.07 | -0.21 | 0.12 | 0.12 | 0.802 |
| **RALBP1** | **S92; S93; S99** | -0.33 | 0.19 | 0.12 | -0.02 | 0.03 | -0.05 | 0.973 |
| **RALBP1** | **S/T** | -0.03 | 0.09 | -0.02 | -0.01 | 0.03 | -0.06 | 0.587 |
| **RALBP1** | **S92; S93** | 0.02 | 0.04 | 0.11 | -0.12 | -0.09 | 0.03 | 0.093 |
| **RALBP1** | **S** | -0.03 | 0.36 | 0.03 | -0.09 | -0.39 | 0.02 | 0.182 |
| **RALGAPA1** | **S773** | -0.21 | 0.15 | 0.09 | 0.17 | -0.21 | -0.05 | 0.813 |
| **RALGAPA1** | **S773** | -0.08 | 0.10 | 0.07 | 0.09 | -0.04 | -0.14 | 0.548 |
| **RALGAPA1** | **S797** | 0.05 | 0.17 | 0.00 | -0.17 | 0.23 | -0.36 | 0.391 |
| **RALGAPA1** | **S740** | 0.01 | 0.19 | 0.30 | -0.41 | -0.02 | -0.16 | 0.064 |
| **RALGAPA1** | **S860** | 0.10 | 0.37 | 0.21 | -0.37 | 0.05 | -0.59 | 0.061 |
| **RALGAPA2** | **S820** | 0.10 | -0.05 | -0.56 | -0.13 | -0.06 | 0.50 | 0.39 |
| **RALGAPA2** | **S379** | -0.17 | 0.09 | 0.21 | -0.19 | -0.17 | 0.16 | 0.53 |
| **RALGAPA2** | **S819** | 0.08 | 0.25 | 0.12 | -0.09 | -0.16 | -0.26 | 0.011 |
| **RALGAPA2** | **S821** | 0.09 | 0.19 | 0.20 | -0.48 | -0.10 | -0.01 | 0.073 |
| **RALGAPA2** | **S821** | 0.00 | 0.22 | 0.28 | 0.02 | -0.20 | -0.43 | 0.078 |

| | | | | | | | | |
|---|---|---|---|---|---|---|---|---|
| RALGAPB | T/S | -0.26 | -0.23 | 0.13 | 0.12 | 0.18 | -0.01 | 0.197 |
| RALGAPB | S1453 | 0.43 | 0.03 | 0.04 | -0.36 | -0.41 | 0.10 | 0.139 |
| RALGPS2 | S329 | -0.18 | -0.08 | -0.38 | 0.42 | 0.03 | 0.06 | 0.068 |
| RALY | T286; S288 | -0.32 | -0.39 | -0.09 | 0.63 | -0.47 | 0.29 | 0.28 |
| RALY | S264 | -0.13 | -0.18 | -0.06 | -0.01 | 0.12 | 0.21 | 0.031 |
| RALY | S288 | -0.15 | -0.07 | 0.14 | 0.07 | -0.19 | 0.17 | 0.778 |
| RALY | S135 | 0.03 | -0.01 | -0.04 | -0.03 | -0.06 | 0.10 | 0.865 |
| RALY | S135 | 0.08 | 0.00 | -0.03 | -0.05 | 0.02 | -0.03 | 0.346 |
| RALY | S239 | 0.12 | 0.04 | -0.10 | -0.49 | -0.10 | 0.39 | 0.756 |
| RALY | T286; S288 | 0.04 | -0.08 | 0.23 | -0.19 | -0.14 | 0.10 | 0.34 |
| RALY | T286; S295; T298 | 0.02 | 0.13 | 0.07 | -0.11 | 0.09 | -0.23 | 0.196 |
| RALY | S/T | 0.04 | 0.08 | 0.13 | -0.06 | -0.15 | -0.05 | 0.017 |
| RALY | T298 | 0.11 | 0.13 | 0.15 | -0.26 | -0.23 | 0.06 | 0.057 |
| RALY | S239 | 0.04 | 0.30 | 0.23 | -0.10 | 0.09 | -0.81 | 0.178 |
| RALY | S177 | 0.15 | 0.27 | 0.21 | -0.44 | -0.14 | -0.19 | 0.009 |
| RALY | S264 | 0.15 | 0.53 | 0.08 | -0.34 | -0.46 | -0.18 | 0.023 |
| RALY | T286; S288 | -0.14 | 0.34 | 0.67 | -1.29 | -0.03 | -0.24 | 0.149 |
| RAMACL | S36 | -0.01 | -0.07 | 0.10 | -0.03 | -0.19 | 0.17 | 0.831 |
| RANBP1 | S60 | 0.00 | -0.53 | -0.01 | 0.41 | 0.14 | -0.16 | 0.268 |
| RANBP1 | S188 | -0.17 | -0.21 | -0.11 | -0.09 | 0.29 | 0.21 | 0.065 |
| RANBP10 | S365; S369 | 0.03 | -0.09 | 0.19 | -0.07 | -0.07 | 0.00 | 0.339 |
| RANBP10 | S369 | 0.22 | -0.16 | 0.06 | 0.15 | -0.09 | -0.24 | 0.548 |
| RANBP2 | T1396; S1400 | 0.09 | -0.43 | -0.23 | 0.35 | 0.16 | -0.07 | 0.158 |
| RANBP2 | S2741 | 0.00 | 0.00 | -0.19 | 0.15 | -0.06 | 0.07 | 0.236 |
| RANBP2 | S3207 | -0.07 | 0.02 | -0.07 | 0.25 | 0.03 | -0.20 | 0.653 |
| RANBP2 | S2741 | 0.03 | -0.03 | -0.06 | 0.21 | -0.14 | -0.04 | 0.799 |
| RANBP2 | S1993 | -0.06 | -0.02 | 0.07 | 0.02 | -0.12 | 0.10 | 0.95 |
| RANBP2 | T1255 | -0.08 | 0.11 | 0.07 | 0.07 | -0.22 | 0.02 | 0.508 |
| RANBP2 | S1993 | 0.15 | 0.09 | -0.15 | 0.12 | 0.01 | -0.27 | 0.618 |
| RANBP2 | S1160 | 0.01 | 0.08 | 0.04 | 0.06 | -0.09 | -0.12 | 0.201 |
| RANBP2 | T2504 | -0.02 | 0.03 | 0.13 | -0.01 | -0.18 | 0.03 | 0.262 |
| RANBP2 | T1396 | 0.30 | -0.27 | 0.07 | 0.23 | -0.08 | -0.38 | 0.669 |
| RANBP2 | S2462; S/T | 0.08 | -0.04 | 0.13 | 0.11 | -0.07 | -0.25 | 0.332 |
| RANBP2 | S2276; S2280 | 0.20 | 0.05 | -0.08 | 0.07 | 0.03 | -0.32 | 0.434 |
| RANBP2 | S837 | 0.08 | 0.11 | 0.02 | 0.05 | -0.13 | -0.14 | 0.107 |
| RANBP2 | T/S | 0.15 | 0.04 | 0.01 | -0.04 | 0.02 | -0.22 | 0.156 |
| RANBP2 | S1573 | 0.04 | 0.14 | 0.01 | 0.11 | 0.06 | -0.44 | 0.452 |
| RANBP2 | S | 0.51 | -0.49 | 0.01 | 0.28 | -1.00 | 0.18 | 0.723 |
| RANBP2 | S2668 | -0.01 | 0.18 | 0.13 | -0.07 | 0.02 | -0.30 | 0.116 |
| RANBP2 | S2454 | 0.05 | 0.01 | 0.27 | -0.12 | -0.15 | -0.10 | 0.051 |
| RANBP2 | T2293 | 0.07 | -0.11 | 0.38 | -0.51 | 0.21 | -0.21 | 0.322 |
| RANBP2 | S730 | 0.02 | 0.16 | 0.24 | 0.04 | -0.18 | -0.37 | 0.085 |
| RANBP2 | S21 | -0.08 | 0.15 | 0.36 | -0.08 | -0.51 | 0.02 | 0.18 |
| RANBP2 | S2805 | 0.30 | 0.16 | 0.01 | -0.01 | 0.00 | -0.61 | 0.173 |
| RANBP2 | S21 | -0.01 | 0.48 | 0.14 | -0.15 | -0.13 | -0.54 | 0.072 |
| RANBP2 | S833; S | 0.28 | 0.17 | 0.28 | -0.24 | -0.20 | -0.46 | 0.003 |
| RANBP2 | S781 | 0.39 | 0.16 | 0.29 | -0.45 | -0.23 | -0.38 | 0.002 |
| RANBP2 | S955 | 0.34 | 0.18 | 0.43 | -0.48 | -0.12 | -0.68 | 0.014 |

| | | | | | | | | |
|---|---|---|---|---|---|---|---|---|
| **RAP1GAP** | **S515** | 0.22 | -0.14 | 0.09 | -0.03 | -0.05 | -0.13 | 0.318 |
| **RAPGEF3** | **S864** | -0.10 | -0.36 | 0.12 | -0.11 | -0.08 | 0.41 | 0.448 |
| **RASAL2** | **S1119** | -0.04 | -0.27 | -0.14 | 0.14 | 0.03 | 0.22 | 0.03 |
| **RASAL2** | **S928** | 0.20 | -0.10 | 0.11 | -0.27 | 0.11 | -0.09 | 0.336 |
| **RASEF** | **S93** | 0.26 | -0.20 | -0.20 | 0.04 | -0.11 | 0.14 | 0.681 |
| **RB1** | **T821** | -0.24 | -0.22 | -0.29 | 0.13 | 0.20 | 0.30 | 9E-04 |
| **RB1** | **S37** | -0.15 | -0.14 | -0.12 | 0.03 | 0.16 | 0.19 | 0.007 |
| **RB1** | **S37** | -0.01 | -0.09 | -0.02 | 0.00 | 0.04 | 0.07 | 0.069 |
| **RB1** | **T373** | 0.00 | 0.05 | -0.15 | 0.02 | -0.02 | 0.09 | 0.414 |
| **RB1** | **T373** | -0.02 | 0.02 | 0.09 | 0.12 | -0.03 | -0.20 | 0.534 |
| **RB1** | **T/S** | 0.00 | 0.13 | 0.16 | -0.15 | -0.06 | -0.11 | 0.024 |
| **RB1** | **S788; S/Y** | 0.26 | -0.11 | 0.08 | 0.37 | -0.47 | -0.32 | 0.492 |
| **RB1** | **S249** | 0.07 | 0.16 | 0.10 | -0.29 | -0.04 | -0.05 | 0.051 |
| **RB1** | **T373** | 0.27 | 0.22 | 0.07 | -0.19 | -0.27 | -0.20 | 0.004 |
| **RB1CC1** | **S647** | -0.17 | -0.15 | -0.11 | 0.18 | -0.06 | 0.24 | 0.047 |
| **RB1CC1** | **S647** | -0.38 | -0.10 | 0.02 | 0.39 | -0.18 | 0.11 | 0.269 |
| **RB1CC1** | **S222** | -0.07 | 0.02 | 0.11 | -0.13 | -0.08 | 0.12 | 0.606 |
| **RBBP4** | **S146** | -0.02 | 0.13 | 0.01 | -0.28 | -0.04 | 0.16 | 0.517 |
| **RBBP5** | **S525** | -0.80 | -0.28 | -0.99 | 0.33 | 0.51 | 0.50 | 0.007 |
| **RBBP5** | **S433** | -0.09 | -0.59 | -0.13 | 0.17 | -0.01 | 0.45 | 0.088 |
| **RBBP6** | **S772** | -0.07 | -0.74 | -0.36 | 0.12 | 0.26 | 0.46 | 0.037 |
| **RBBP6** | **S854** | -0.17 | -0.28 | -0.27 | 0.04 | -0.01 | 0.52 | 0.073 |
| **RBBP6** | **S748; S750** | -0.27 | -0.07 | -0.03 | 0.58 | -0.15 | -0.24 | 0.52 |
| **RBBP6** | **S780** | 0.03 | -0.09 | -0.03 | 0.24 | -0.20 | 0.01 | 0.75 |
| **RBBP6** | **S1277** | -0.09 | 0.10 | 0.04 | -0.05 | -0.13 | 0.10 | 0.65 |
| **RBBP6** | **S1277** | 0.35 | 0.01 | -0.35 | -0.45 | -0.41 | 0.53 | 0.777 |
| **RBBP6** | **S1277** | -0.18 | 0.25 | 0.15 | -0.14 | -0.33 | 0.15 | 0.393 |
| **RBBP6** | **S1705** | 0.10 | 0.25 | -0.05 | 0.05 | -0.22 | -0.18 | 0.149 |
| **RBBP6** | **S1179** | 0.16 | 0.16 | 0.03 | -0.10 | -0.09 | -0.20 | 0.01 |
| **RBBP6** | **S1644; S1648; S1651** | 0.25 | 0.06 | 0.04 | -0.11 | -0.08 | -0.21 | 0.031 |
| **RBBP6** | **T984** | 0.11 | 0.16 | 0.08 | -0.07 | 0.00 | -0.34 | 0.077 |
| **RBBP6** | **T984** | 0.14 | 0.17 | 0.10 | -0.08 | 0.00 | -0.41 | 0.082 |
| **RBBP6** | **S1644; S1648; S1651** | 0.28 | 0.17 | 0.02 | -0.07 | -0.13 | -0.37 | 0.044 |
| **RBBP6** | **T984** | 0.24 | -0.05 | 0.28 | -0.06 | 0.00 | -0.55 | 0.153 |
| **RBBP6** | **S1221** | 0.19 | 0.14 | 0.26 | -0.15 | -0.08 | -0.49 | 0.028 |
| **RBBP6** | **S1282; S/T/Y** | 0.74 | -0.19 | -0.01 | -0.37 | -0.01 | -0.54 | 0.207 |
| **RBBP6** | **S703** | 0.39 | 0.26 | 0.07 | -0.37 | -0.17 | -0.35 | 0.009 |
| **RBBP6** | **S1277** | 0.36 | 0.39 | 0.34 | -0.42 | 0.13 | -1.72 | 0.133 |
| **RBBP8NL** | **T242** | -0.09 | -0.16 | -0.06 | 0.04 | -0.03 | 0.25 | 0.099 |
| **RBBP8NL** | **S487** | 0.13 | -0.09 | -0.18 | 0.12 | -0.16 | 0.15 | 0.563 |
| **RBBP8NL** | **S196** | 0.09 | 0.01 | -0.21 | 0.19 | 0.03 | -0.16 | 0.709 |
| **RBBP8NL** | **S251** | 0.15 | -0.05 | -0.04 | 0.00 | 0.03 | -0.11 | 0.56 |
| **RBBP8NL** | **S145** | 0.26 | 0.20 | -0.40 | -0.04 | 0.03 | -0.15 | 0.756 |
| **RBBP8NL** | **S487** | -0.16 | 0.14 | 0.19 | 0.11 | 0.19 | -0.65 | 0.588 |
| **RBBP8NL** | **S645** | 0.28 | 0.08 | 0.17 | -0.26 | 0.11 | -0.55 | 0.107 |
| **RBL1** | **S1041** | 0.26 | -0.25 | -0.17 | 0.24 | -0.16 | -0.01 | 0.708 |
| **RBL1** | **S650** | 0.52 | 0.45 | 0.45 | -0.93 | -0.56 | -0.65 | 5E-04 |
| **RBL2** | **S1112** | -0.40 | -0.10 | -0.35 | 0.17 | -0.06 | 0.53 | 0.063 |

| | | | | | | | | |
|---|---|---|---|---|---|---|---|---|
| RBL2 | S1112 | 0.18 | -0.10 | -0.25 | -0.38 | 0.19 | 0.26 | 0.759 |
| RBL2 | S662 | 0.56 | -0.19 | -0.48 | 0.02 | -0.11 | -0.01 | 0.987 |
| RBL2 | S639 | -0.17 | 0.21 | 0.05 | -0.14 | 0.01 | 0.01 | 0.602 |
| RBL2 | S952 | -0.02 | 0.05 | 0.33 | -0.15 | 0.05 | -0.36 | 0.163 |
| RBL2 | S/T/Y | 0.20 | 0.21 | 0.12 | -0.20 | -0.27 | -0.14 | 0.001 |
| RBL2 | T694 | -0.15 | 0.51 | 0.10 | -0.03 | 0.00 | -0.69 | 0.254 |
| RBM10 | S733; S736; S738 | -0.21 | -0.16 | -0.29 | 0.23 | -0.02 | 0.34 | 0.024 |
| RBM10 | T/S/Y | -0.06 | -0.26 | -0.05 | -0.33 | 0.31 | 0.27 | 0.39 |
| RBM10 | S797 | 0.04 | -0.05 | -0.07 | 0.03 | 0.09 | -0.05 | 0.394 |
| RBM10 | S733 | -0.03 | 0.04 | -0.03 | -0.01 | -0.07 | 0.09 | 0.843 |
| RBM10 | S89 | 0.09 | 0.09 | -0.13 | 0.11 | -0.06 | -0.12 | 0.737 |
| RBM10 | S736; S738 | 0.11 | 0.07 | -0.05 | 0.06 | -0.15 | -0.05 | 0.319 |
| RBM10 | S738 | 0.02 | 0.11 | 0.02 | 0.01 | -0.12 | -0.05 | 0.095 |
| RBM10 | S60 | -0.25 | 0.22 | 0.46 | -0.14 | -0.89 | 0.23 | 0.352 |
| RBM10 | S723 | 0.18 | 0.21 | 0.22 | -0.21 | -0.03 | -0.51 | 0.031 |
| RBM10 | S89 | 0.20 | 0.12 | 0.30 | -0.24 | -0.02 | -0.52 | 0.038 |
| RBM12B | S254 | -0.18 | -0.26 | -0.26 | 0.20 | 0.24 | 0.16 | 2E-04 |
| RBM12B | S874 | -0.27 | -0.05 | -0.19 | 0.15 | -0.12 | 0.38 | 0.125 |
| RBM12B | S710 | -0.07 | -0.12 | -0.10 | 0.07 | -0.14 | 0.31 | 0.24 |
| RBM12B | S638 | -0.04 | -0.26 | 0.04 | 0.01 | -0.10 | 0.28 | 0.356 |
| RBM12B | S591 | -0.15 | -0.13 | 0.10 | -0.07 | 0.04 | 0.18 | 0.367 |
| RBM12B | S638 | -0.07 | 0.03 | 0.02 | -0.05 | 0.01 | 0.05 | 0.876 |
| RBM12B | S575 | 0.02 | 0.05 | 0.05 | 0.02 | 0.06 | -0.22 | 0.371 |
| RBM12B | S250; S254 | 0.03 | 0.05 | 0.06 | -0.05 | -0.04 | -0.05 | 4E-04 |
| RBM12B | S839 | -0.05 | 0.12 | 0.19 | -0.08 | -0.04 | -0.17 | 0.09 |
| RBM12B | S839 | 0.12 | 0.12 | 0.26 | -0.24 | -0.01 | -0.33 | 0.026 |
| RBM14 | T206 | -0.25 | -0.34 | -0.27 | 0.23 | 0.14 | 0.34 | 0.001 |
| RBM14 | T572 | -0.04 | -0.14 | -0.28 | 0.27 | 0.10 | 0.02 | 0.052 |
| RBM14 | S/Y/T | 0.02 | 0.05 | -0.29 | 0.21 | -0.05 | 0.01 | 0.385 |
| RBM14 | T572 | -0.03 | -0.09 | -0.07 | 0.06 | -0.05 | 0.17 | 0.132 |
| RBM14 | T572 | -0.04 | 0.04 | -0.17 | 0.14 | 0.03 | -0.01 | 0.212 |
| RBM14 | S/Y/T | -0.22 | 0.07 | -0.03 | -0.09 | -0.05 | 0.27 | 0.502 |
| RBM14 | S649 | 0.19 | -0.19 | -0.05 | 0.28 | -0.03 | -0.26 | 0.952 |
| RBM14 | S582 | -0.01 | 0.07 | 0.10 | -0.07 | 0.10 | -0.23 | 0.304 |
| RBM14 | S/T | 0.30 | 0.12 | -0.08 | 0.11 | 0.02 | -0.64 | 0.343 |
| RBM15 | S294 | -0.15 | -0.41 | -0.34 | 0.10 | 0.03 | 0.55 | 0.045 |
| RBM15 | T568 | -0.11 | -0.29 | -0.21 | 0.07 | -0.06 | 0.47 | 0.094 |
| RBM15 | S259 | -0.09 | -0.08 | -0.17 | 0.10 | 0.08 | 0.14 | 0.002 |
| RBM15 | S670 | -0.09 | -0.06 | -0.02 | 0.01 | 0.21 | -0.06 | 0.257 |
| RBM15 | S97 | 0.09 | 0.15 | -0.33 | 0.32 | -0.15 | -0.19 | 0.918 |
| RBM15 | S208 | -0.10 | -0.02 | 0.10 | -0.06 | 0.16 | -0.10 | 0.968 |
| RBM15 | S622 | -0.01 | -0.04 | 0.07 | -0.04 | 0.03 | -0.01 | 0.724 |
| RBM15 | S109 | -0.04 | 0.11 | -0.08 | 0.21 | -0.32 | 0.06 | 0.934 |
| RBM15 | S622 | 0.27 | -0.02 | -0.16 | 0.17 | -0.34 | -0.01 | 0.673 |
| RBM15 | S700 | 0.04 | 0.04 | 0.09 | -0.03 | -0.02 | -0.12 | 0.028 |
| RBM15 | S686 | -0.19 | 0.04 | 0.33 | -0.09 | 0.21 | -0.43 | 0.528 |
| RBM15 | S700 | 0.00 | 0.03 | 0.25 | -0.03 | 0.04 | -0.36 | 0.224 |
| RBM15 | S752 | 0.12 | 0.30 | 0.05 | -0.08 | 0.01 | -0.54 | 0.128 |

| | | | | | | | | |
|---|---|---|---|---|---|---|---|---|
| **RBM15** | **S670; S674** | 0.10 | 0.13 | 0.26 | -0.15 | -0.05 | -0.39 | 0.031 |
| **RBM15** | **S83** | 0.17 | 0.11 | 0.20 | -0.08 | -0.01 | -0.51 | 0.083 |
| **RBM15** | **S724** | 0.24 | 0.06 | 0.26 | -0.20 | -0.15 | -0.31 | 0.007 |
| **RBM15** | **S741** | 0.25 | 0.29 | 0.11 | -0.37 | -0.07 | -0.35 | 0.012 |
| **RBM15** | **S656** | 0.18 | 0.19 | 0.29 | -0.26 | -0.03 | -0.54 | 0.031 |
| **RBM15** | **S/Y/T** | 0.22 | 0.28 | 0.19 | -0.28 | -0.03 | -0.54 | 0.027 |
| **RBM15** | **S741** | 0.28 | 0.22 | 0.20 | -0.29 | -0.05 | -0.55 | 0.023 |
| **RBM15** | **S604** | 0.22 | 0.06 | 0.39 | -0.26 | 0.03 | -0.69 | 0.08 |
| **RBM15** | **S128** | 0.35 | 0.11 | 0.27 | -0.15 | -0.04 | -0.81 | 0.084 |
| **RBM15** | **S83** | 0.32 | 0.39 | 0.53 | -0.61 | -0.21 | -1.09 | 0.016 |
| **RBM15B** | **T631** | -0.23 | -0.25 | 0.00 | 0.09 | 0.47 | -0.21 | 0.257 |
| **RBM15B** | **T532** | 0.04 | -0.21 | -0.18 | -0.01 | 0.08 | 0.23 | 0.102 |
| **RBM15B** | **S562** | -0.05 | -0.05 | -0.15 | 0.17 | 0.00 | 0.06 | 0.055 |
| **RBM15B** | **S267** | 0.00 | -0.09 | -0.09 | 0.04 | 0.07 | 0.07 | 0.018 |
| **RBM15B** | **S265; S267** | -0.23 | -0.05 | 0.07 | -0.17 | -0.08 | 0.37 | 0.6 |
| **RBM15B** | **S113** | 0.04 | -0.04 | -0.10 | 0.18 | -0.14 | 0.03 | 0.594 |
| **RBM15B** | **S/T** | 0.13 | -0.15 | 0.02 | 0.21 | -0.22 | -0.03 | 0.951 |
| **RBM15B** | **S109** | 0.02 | 0.09 | 0.00 | 0.05 | 0.08 | -0.27 | 0.519 |
| **RBM15B** | **S109** | 0.00 | 0.04 | 0.10 | 0.05 | -0.09 | -0.12 | 0.164 |
| **RBM15B** | **S609** | 0.08 | 0.03 | 0.10 | -0.04 | -0.05 | -0.14 | 0.019 |
| **RBM15B** | **S265; S267** | 0.15 | 0.05 | 0.09 | -0.37 | 0.13 | -0.12 | 0.217 |
| **RBM15B** | **T238** | 0.20 | 0.05 | 0.22 | -0.35 | -0.05 | -0.14 | 0.031 |
| **RBM15B** | **S267** | 0.06 | 0.10 | 0.47 | -0.40 | 0.00 | -0.45 | 0.064 |
| **RBM15B** | **S552** | 0.25 | 0.20 | 0.22 | -0.26 | -0.02 | -0.58 | 0.036 |
| **RBM17** | **S155** | -0.08 | -0.14 | 0.25 | 0.07 | -0.18 | 0.04 | 0.828 |
| **RBM17** | **T71** | 0.13 | 0.09 | 0.10 | -0.08 | -0.13 | -0.14 | 6E-04 |
| **RBM17** | **S222** | 0.22 | 0.24 | -0.08 | 0.11 | -0.19 | -0.41 | 0.184 |
| **RBM17** | **S222** | 0.22 | 0.21 | -0.01 | -0.03 | -0.07 | -0.41 | 0.093 |
| **RBM17** | **S222** | 0.20 | 0.14 | 0.14 | -0.13 | -0.14 | -0.28 | 0.003 |
| **RBM17** | **S222** | 0.23 | 0.03 | 0.43 | 0.02 | -0.33 | -0.62 | 0.066 |
| **RBM19** | **T108** | -0.16 | 0.00 | -0.27 | 0.11 | 0.21 | 0.05 | 0.044 |
| **RBM23** | **S149** | 0.12 | 0.13 | 0.11 | -0.06 | -0.10 | -0.24 | 0.009 |
| **RBM23** | **S128** | 0.42 | 0.16 | -0.19 | 0.01 | -0.17 | -0.36 | 0.212 |
| **RBM25** | **S677; S683** | 0.10 | 0.27 | -0.03 | -0.04 | 0.15 | -0.60 | 0.316 |
| **RBM25** | **S583** | 0.33 | 0.15 | 0.16 | -0.16 | -0.16 | -0.46 | 0.015 |
| **RBM25** | **S** | 0.21 | 0.19 | 0.34 | -0.15 | -0.15 | -0.66 | 0.032 |
| **RBM25** | **S583** | 0.33 | 0.29 | 0.16 | -0.46 | 0.08 | -0.68 | 0.059 |
| **RBM25** | **S677** | 0.31 | 0.25 | 0.24 | -0.26 | -0.08 | -0.71 | 0.03 |
| **RBM25** | **S677** | 0.35 | 0.33 | 0.36 | -0.31 | -0.20 | -0.96 | 0.025 |
| **RBM26** | **S127** | 0.14 | -0.09 | -0.24 | 0.33 | -0.10 | -0.10 | 0.585 |
| **RBM26** | **S127** | 0.02 | -0.01 | -0.08 | 0.15 | -0.08 | 0.00 | 0.54 |
| **RBM26** | **S616** | 0.11 | -0.11 | 0.15 | -0.22 | -0.05 | 0.08 | 0.401 |
| **RBM26** | **S127** | 0.20 | -0.17 | 0.23 | -0.80 | 0.28 | 0.00 | 0.502 |
| **RBM27** | **S120** | 0.21 | 0.14 | -0.05 | 0.03 | -0.10 | -0.29 | 0.139 |
| **RBM28** | **S** | -0.15 | -0.13 | 0.28 | -0.25 | -0.07 | 0.23 | 0.887 |
| **RBM33** | **S765** | 0.03 | -0.01 | -0.01 | 0.04 | 0.06 | -0.11 | 0.902 |
| **RBM34** | **S28** | -0.34 | -0.07 | -0.01 | 0.06 | 0.09 | 0.21 | 0.081 |
| **RBM34** | **S28** | -0.04 | 0.03 | 0.02 | 0.00 | 0.04 | -0.05 | 0.937 |

| | | | | | | | | |
|---|---|---|---|---|---|---|---|---|
| **RBM34** | **S14** | 0.38 | -0.06 | 0.05 | -0.18 | 0.12 | -0.46 | 0.243 |
| **RBM39** | **S136** | -0.15 | -0.22 | -0.28 | 0.25 | -0.30 | 0.50 | 0.202 |
| **RBM39** | **S337** | 0.14 | -0.23 | -0.18 | 0.02 | 0.08 | 0.13 | 0.247 |
| **RBM39** | **S136** | 0.03 | -0.04 | -0.12 | 0.03 | -0.04 | 0.14 | 0.285 |
| **RBM39** | **S136** | 0.05 | 0.06 | -0.04 | -0.02 | -0.01 | -0.04 | 0.218 |
| **RBM39** | **S97** | 0.25 | -0.06 | -0.10 | 0.09 | -0.25 | 0.02 | 0.634 |
| **RBM39** | **S97** | 0.20 | 0.10 | 0.24 | -0.26 | -0.07 | -0.30 | 0.009 |
| **RBM4** | **S86** | 0.13 | -0.05 | -0.01 | -0.17 | 0.26 | -0.21 | 0.72 |
| **RBM4** | **S309** | 0.03 | 0.08 | 0.05 | -0.02 | 0.05 | -0.21 | 0.245 |
| **RBM4B** | **S86** | -0.24 | 0.05 | 0.16 | -1.21 | -0.03 | 0.66 | 0.763 |
| **RBM5** | **S78** | -0.12 | -0.19 | -0.22 | 0.24 | 0.07 | 0.14 | 0.005 |
| **RBM5** | **S621; S624** | -0.20 | -0.10 | -0.02 | 0.12 | -0.13 | 0.27 | 0.206 |
| **RBM5** | **S624** | -0.08 | -0.09 | -0.07 | 0.07 | -0.06 | 0.20 | 0.125 |
| **RBM5** | **S624** | -0.09 | -0.04 | -0.10 | 0.03 | -0.12 | 0.28 | 0.291 |
| **RBM5** | **S59** | -0.15 | -0.11 | 0.10 | 0.02 | -0.05 | 0.17 | 0.392 |
| **RBM6** | **T344; S362** | -0.59 | -0.83 | -0.59 | 0.51 | -0.02 | 0.76 | 0.011 |
| **RBM6** | **S353; S360; S362** | -0.43 | -0.27 | -0.48 | 0.46 | 0.32 | 0.11 | 0.005 |
| **RBM6** | **S353; S362** | -0.10 | -0.23 | -0.23 | 0.25 | -0.11 | 0.32 | 0.071 |
| **RBM6** | **S362** | 0.19 | 0.04 | -0.42 | 0.20 | -0.35 | 0.20 | 0.778 |
| **RBM6** | **S891** | 0.15 | 0.07 | 0.08 | 0.00 | -0.12 | -0.21 | 0.033 |
| **RBM6** | **S1025** | 0.17 | 0.08 | 0.19 | -0.17 | -0.04 | -0.29 | 0.016 |
| **RBM6** | **S362** | 0.34 | 0.06 | 0.09 | 0.02 | -0.29 | -0.33 | 0.062 |
| **RBM6** | **S1025** | 0.18 | 0.24 | 0.20 | -0.13 | -0.22 | -0.38 | 0.004 |
| **RBM6** | **S461** | 0.19 | 0.09 | 0.33 | -0.32 | -0.15 | -0.26 | 0.007 |
| **RBM7** | **S136** | -0.23 | -0.02 | -0.46 | 0.13 | 0.33 | 0.12 | 0.041 |
| **RBM8A** | **S42** | -0.01 | -0.05 | -0.01 | 0.03 | -0.12 | 0.14 | 0.644 |
| **RBMS3** | **S111** | -0.01 | -0.31 | -0.22 | 0.17 | 0.08 | 0.21 | 0.026 |
| **RBMX** | **S157** | -0.09 | -0.10 | -0.07 | 0.17 | 0.01 | 0.07 | 0.021 |
| **RBMX** | **S261** | -0.01 | -0.09 | -0.16 | 0.03 | 0.06 | 0.15 | 0.042 |
| **RBMX** | **S88** | 0.00 | -0.15 | -0.01 | 0.05 | -0.12 | 0.20 | 0.403 |
| **RBMX** | **S352** | 0.02 | 0.05 | 0.01 | 0.02 | -0.04 | -0.06 | 0.112 |
| **RBMX** | **S168** | 0.02 | 0.01 | 0.09 | -0.12 | -0.08 | 0.07 | 0.278 |
| **RBMX** | **S88** | 0.18 | 0.19 | 0.17 | -0.23 | -0.05 | -0.36 | 0.012 |
| **RBMX2** | **S186** | -0.06 | -0.01 | 0.10 | -0.06 | -0.10 | 0.12 | 0.783 |
| **RBMX2** | **S187** | 0.08 | 0.02 | 0.03 | -0.07 | -0.02 | -0.04 | 0.02 |
| **RBMXL1** | **S58** | -0.42 | -0.59 | -0.72 | 0.25 | 0.17 | 0.74 | 0.008 |
| **RBMXL1** | **S326** | -0.18 | -0.24 | 0.00 | -0.01 | 0.19 | 0.19 | 0.057 |
| **RBMXL1** | **S208** | -0.02 | -0.14 | -0.08 | 0.06 | -0.04 | 0.20 | 0.124 |
| **RBMXL1** | **S303** | 0.05 | -0.05 | -0.10 | 0.03 | -0.16 | 0.20 | 0.63 |
| **RBMXL1** | **S189** | 0.12 | -0.10 | -0.10 | 0.16 | -0.21 | 0.09 | 0.769 |
| **RBMXL1** | **S208** | 0.06 | 0.12 | -0.17 | -0.03 | 0.01 | -0.02 | 0.872 |
| **RBMXL1** | **S249** | 0.03 | -0.06 | 0.00 | 0.27 | -0.47 | 0.13 | 0.948 |
| **RBMXL1** | **S326** | -0.06 | -0.02 | 0.16 | -0.16 | -0.07 | 0.12 | 0.595 |
| **RBMXL1** | **S351** | 0.08 | 0.01 | 0.01 | -0.08 | -0.10 | 0.07 | 0.295 |
| **RBMXL1** | **S287** | 0.04 | -0.02 | 0.13 | -0.05 | 0.03 | -0.14 | 0.211 |
| **RBMXL1** | **S277** | 0.34 | 0.06 | -0.15 | 0.06 | -0.10 | -0.28 | 0.324 |
| **RBMXL1** | **S219** | 0.18 | 0.14 | -0.01 | -0.20 | 0.02 | -0.16 | 0.068 |
| **RBMXL1** | **S314** | 0.07 | 0.13 | 0.12 | -0.23 | 0.02 | -0.16 | 0.038 |

| | | | | | | | | |
|---|---|---|---|---|---|---|---|---|
| **RBMXL1** | **S208** | 0.14 | 0.10 | 0.09 | -0.17 | -0.01 | -0.18 | 0.014 |
| **RBMXL1** | **S189** | -0.25 | 0.06 | 0.57 | -0.47 | 0.00 | -0.14 | 0.301 |
| **RBMXL1** | **S250** | 0.26 | 0.16 | 0.05 | -0.17 | -0.18 | -0.19 | 0.005 |
| **RBMXL1** | **S88** | 0.22 | 0.27 | 0.17 | -0.37 | 0.01 | -0.47 | 0.029 |
| **RCC1** | **S11** | -0.16 | -0.27 | -0.51 | 0.26 | 0.03 | 0.44 | 0.024 |
| **RCC1** | **S11** | -0.08 | -0.16 | -0.19 | 0.12 | 0.13 | 0.15 | 1E-03 |
| **RCC1** | **S11** | 0.19 | 0.13 | 0.02 | -0.02 | 0.09 | -0.51 | 0.238 |
| **RCC2** | **S43; S45; S46** | -0.07 | 0.00 | -0.05 | 0.23 | 0.06 | -0.21 | 0.63 |
| **RCC2** | **S50; S51** | -0.01 | 0.02 | 0.07 | -0.14 | 0.01 | 0.04 | 0.376 |
| **RCC2** | **S50** | 0.22 | 0.12 | 0.25 | -0.28 | -0.06 | -0.36 | 0.012 |
| **RCOR1** | **S460** | 0.06 | -0.06 | -0.25 | 0.14 | -0.14 | 0.20 | 0.348 |
| **RCOR1** | **S260** | 0.02 | -0.15 | -0.10 | 0.13 | -0.16 | 0.22 | 0.341 |
| **RCOR1** | **S260** | 0.18 | -0.10 | -0.23 | 0.16 | -0.19 | 0.12 | 0.65 |
| **RCOR1** | **S260** | -0.04 | 0.01 | -0.04 | 0.05 | -0.06 | 0.08 | 0.35 |
| **RCOR1** | **S260** | -0.02 | 0.09 | -0.07 | -0.03 | -0.05 | 0.07 | 0.976 |
| **RCOR1** | **S260** | 0.23 | -0.06 | -0.12 | -0.01 | -0.19 | 0.10 | 0.714 |
| **RCOR1** | **S460** | 0.33 | -0.04 | -0.02 | -0.13 | -0.10 | -0.09 | 0.179 |
| **RCOR1** | **S460** | 0.48 | 0.21 | -0.09 | -0.30 | -0.03 | -0.46 | 0.089 |
| **RCOR3** | **S375; T376** | 0.17 | 0.02 | 0.10 | -0.19 | 0.07 | -0.20 | 0.104 |
| **RCOR3** | **S156; S171** | 0.14 | 0.27 | 0.13 | -0.35 | -0.12 | -0.17 | 0.009 |
| **RCOR3** | **S156; S171** | 0.24 | 0.12 | 0.25 | -0.23 | -0.29 | -0.20 | 8E-04 |
| **RCOR3** | **S/T** | 0.43 | 0.09 | 0.13 | -0.28 | -0.16 | -0.38 | 0.017 |
| **RECQL** | **S/Y** | -0.22 | 0.16 | 0.10 | 0.15 | -0.11 | -0.12 | 0.797 |
| **RECQL4** | **S251** | 0.07 | -0.01 | 0.31 | -0.15 | -0.09 | -0.20 | 0.057 |
| **RECQL5** | **T839** | -0.06 | -0.14 | 0.16 | -0.05 | 0.14 | -0.09 | 0.902 |
| **RECQL5** | **S727** | 0.14 | 0.03 | -0.20 | -0.38 | -0.08 | 0.37 | 0.947 |
| **RECQL5** | **S815** | -0.04 | 0.09 | 0.05 | 0.00 | -0.03 | -0.07 | 0.183 |
| **RECQL5** | **S815** | -0.08 | 0.20 | 0.10 | -0.06 | 0.04 | -0.25 | 0.236 |
| **RECQL5** | **S727** | 0.21 | -0.05 | 0.10 | 0.09 | -0.32 | -0.10 | 0.236 |
| **REEP4** | **S194; T196; S202** | -0.36 | -0.14 | -0.29 | 0.13 | -0.07 | 0.54 | 0.071 |
| **REEP4** | **S257** | 0.01 | -0.12 | -0.08 | -0.09 | -0.16 | 0.37 | 0.572 |
| **REEP4** | **S194; S202** | 0.13 | -0.01 | 0.05 | 0.03 | 0.00 | -0.23 | 0.263 |
| **REEP4** | **S202** | 0.04 | 0.15 | 0.03 | -0.10 | -0.08 | -0.06 | 0.018 |
| **REEP4** | **S152** | 0.09 | 0.16 | -0.04 | 0.04 | -0.23 | -0.07 | 0.187 |
| **REEP4** | **T196** | 0.22 | 0.26 | 0.33 | -0.33 | -0.09 | -0.62 | 0.017 |
| **RELCH** | **S170** | -0.23 | -0.16 | -0.30 | 0.19 | 0.29 | 0.11 | 0.003 |
| **RELCH** | **S/T** | -0.05 | -0.38 | 0.03 | 0.24 | 0.11 | -0.03 | 0.175 |
| **REPS1** | **S/T** | -0.47 | 0.21 | -0.13 | 0.38 | -0.06 | -0.08 | 0.44 |
| **REPS1** | **S/T** | -0.24 | 0.03 | -0.11 | 0.22 | -0.10 | 0.15 | 0.195 |
| **REPS1** | **S562** | -0.16 | 0.01 | 0.00 | 0.13 | 0.16 | -0.18 | 0.521 |
| **REPS1** | **S537** | -0.67 | 0.17 | 0.30 | 0.22 | -0.32 | 0.07 | 0.885 |
| **REPS1** | **T173; S174** | -0.16 | 0.13 | -0.09 | 0.02 | -0.39 | 0.37 | 0.888 |
| **REPS1** | **S162; S170; S174** | -0.07 | -0.07 | 0.12 | 0.13 | -0.03 | -0.10 | 0.926 |
| **REPS1** | **S162; S170; S174** | -0.11 | -0.04 | 0.14 | 0.11 | 0.05 | -0.19 | 0.961 |
| **REPS1** | **Y/S** | 0.08 | 0.10 | -0.18 | -0.19 | 0.07 | 0.09 | 0.935 |
| **REPS1** | **S154; T173; S174** | 0.10 | -0.11 | 0.13 | -0.38 | 0.09 | 0.10 | 0.573 |
| **REPS1** | **S709** | 0.10 | 0.10 | 0.00 | -0.07 | 0.07 | -0.23 | 0.2 |
| **REPS1** | **S174; S/T** | 0.08 | 0.04 | 0.10 | -0.02 | -0.43 | 0.16 | 0.392 |

| | | | | | | | | |
|---|---|---|---|---|---|---|---|---|
| REPS1 | S/T | 0.32 | -0.20 | 0.14 | -0.38 | 0.06 | -0.04 | 0.373 |
| REPS2 | S493 | -0.18 | -0.14 | -0.23 | 0.16 | -0.15 | 0.41 | 0.12 |
| REPS2 | S254 | -0.27 | -0.09 | -0.05 | 0.01 | -0.02 | 0.34 | 0.138 |
| REPS2 | S | 0.16 | -0.03 | -0.34 | 0.09 | 0.30 | -0.30 | 0.668 |
| RER1 | S95 | -0.09 | -0.02 | -0.06 | 0.08 | -0.03 | 0.10 | 0.078 |
| RER1 | S95 | -0.01 | 0.05 | -0.08 | 0.10 | -0.03 | -0.04 | 0.69 |
| RERE | S594 | 0.04 | 0.01 | 0.18 | -0.01 | -0.13 | -0.11 | 0.067 |
| RERE | S613 | 0.58 | 0.23 | 0.20 | -0.47 | -0.14 | -0.85 | 0.027 |
| RESF1 | S1358 | -0.14 | 0.24 | -0.06 | 0.10 | -0.18 | 0.01 | 0.803 |
| REST | S864 | -0.07 | 0.25 | 0.04 | 0.02 | -0.13 | -0.15 | 0.224 |
| RET | S696 | -0.06 | -0.21 | -0.18 | 0.10 | -0.19 | 0.42 | 0.224 |
| RETREG3 | S435; S436 | 0.00 | -0.02 | -0.14 | 0.13 | -0.02 | 0.04 | 0.181 |
| RETSAT | S484 | 0.00 | -0.29 | -0.14 | 0.20 | -0.09 | 0.25 | 0.122 |
| REV1 | S301 | -0.02 | -0.15 | 0.16 | 0.15 | 0.36 | -0.71 | 0.86 |
| REV1 | S1108 | 0.38 | 0.03 | 0.08 | 0.32 | -0.09 | -1.20 | 0.357 |
| REV3L | S1406 | 0.10 | -0.44 | -0.29 | 0.17 | 0.40 | -0.10 | 0.159 |
| REV3L | S1724 | -0.23 | 0.02 | 0.09 | -0.02 | 0.10 | 0.02 | 0.526 |
| REV3L | S1030 | 0.04 | -0.07 | 0.06 | 0.03 | -0.09 | 0.02 | 0.688 |
| REV3L | T1041 | 0.24 | 0.02 | -0.12 | 0.14 | -0.39 | 0.02 | 0.561 |
| REV3L | S2171 | -0.06 | 0.23 | -0.01 | 0.27 | -0.38 | -0.16 | 0.534 |
| REV3L | S1130 | -0.07 | 0.19 | 0.30 | -0.54 | 0.10 | -0.13 | 0.205 |
| REV3L | S1119 | 0.11 | 0.04 | 0.30 | -0.24 | -0.43 | 0.10 | 0.117 |
| REV3L | S1123; S1124 | 0.20 | 0.11 | 0.26 | -0.23 | -0.04 | -0.42 | 0.024 |
| REV3L | S1086 | 0.37 | 0.11 | 0.25 | -0.26 | -0.47 | -0.19 | 0.008 |
| REXO1 | S287; S289 | -0.18 | -0.30 | -0.19 | 0.17 | 0.12 | 0.28 | 0.003 |
| REXO1 | S358 | -0.23 | -0.09 | -0.35 | -0.06 | 0.15 | 0.44 | 0.07 |
| REXO1 | S610 | 0.03 | 0.04 | -0.07 | 0.04 | -0.07 | 0.02 | 0.944 |
| REXO1 | S289 | 0.00 | 0.10 | 0.06 | -0.24 | -0.17 | 0.21 | 0.456 |
| REXO4 | S419 | -0.06 | 0.00 | -0.24 | -0.08 | -0.02 | 0.34 | 0.282 |
| REXO4 | S111 | 0.05 | -0.03 | 0.03 | -0.01 | 0.10 | -0.16 | 0.662 |
| RFC1 | S190 | -0.38 | -0.70 | -0.81 | 0.42 | 0.43 | 0.44 | 0.001 |
| RFC1 | S190 | -0.31 | -0.57 | -0.63 | 0.33 | 0.25 | 0.51 | 0.002 |
| RFC1 | S190 | -0.34 | -0.54 | -0.49 | 0.32 | 0.25 | 0.46 | 8E-04 |
| RFC1 | S108 | -0.16 | -0.07 | -0.10 | 0.08 | 0.10 | 0.14 | 0.002 |
| RFC1 | S/T/Y | -0.08 | 0.08 | -0.18 | 0.05 | 0.00 | 0.11 | 0.235 |
| RFC1 | S368 | 0.03 | 0.05 | -0.04 | -0.02 | -0.09 | 0.06 | 0.579 |
| RFC1 | S71 | 0.06 | 0.07 | 0.12 | -0.06 | -0.12 | -0.10 | 0.002 |
| RFC1 | S312 | 0.08 | 0.06 | 0.16 | -0.18 | 0.14 | -0.32 | 0.19 |
| RFC1 | S156 | 0.15 | 0.18 | 0.12 | -0.26 | 0.08 | -0.36 | 0.068 |
| RFFL | S240 | -0.28 | 0.13 | -0.02 | 0.08 | 0.27 | -0.28 | 0.716 |
| RFFL | S229 | -0.05 | -0.08 | 0.15 | 0.00 | -0.12 | 0.09 | 0.868 |
| RFTN1 | S199 | 0.01 | -0.06 | 0.07 | 0.08 | -0.13 | 0.03 | 0.875 |
| RFX1 | S204 | -0.20 | -0.38 | 0.28 | 0.43 | -0.40 | 0.06 | 0.685 |
| RFX1 | S204 | 0.40 | 0.22 | 0.32 | -0.55 | -0.68 | -0.06 | 0.019 |
| RFX5 | S185 | -0.27 | -0.06 | -0.12 | 0.07 | 0.02 | 0.30 | 0.057 |
| RFX7 | T891 | -0.25 | -0.13 | 0.14 | -0.09 | -0.13 | 0.37 | 0.549 |
| RFX7 | S225 | -0.23 | 0.15 | -0.11 | -0.27 | 0.00 | 0.36 | 0.684 |
| RFX7 | S282 | 0.05 | -0.04 | -0.02 | -0.03 | -0.10 | 0.13 | 0.999 |

| | | | | | | | | |
|---|---|---|---|---|---|---|---|---|
| RFX7 | S282 | 0.05 | 0.01 | 0.08 | 0.02 | -0.11 | -0.06 | 0.091 |
| RFX7 | S321 | 0.11 | 0.04 | -0.01 | -0.14 | 0.16 | -0.18 | 0.416 |
| RFX7 | S225 | 0.00 | 0.03 | 0.20 | -0.07 | -0.22 | 0.03 | 0.169 |
| RFX7 | S202 | 0.24 | 0.27 | 0.32 | -0.30 | -0.25 | -0.48 | 0.001 |
| RGL3 | S569; S573 | -0.27 | -0.15 | -0.21 | 0.18 | 0.01 | 0.35 | 0.02 |
| RGL3 | S555 | -0.11 | -0.22 | -0.12 | 0.09 | 0.09 | 0.23 | 0.008 |
| RGL3 | S569 | -0.19 | -0.07 | 0.10 | -0.12 | 0.13 | 0.12 | 0.452 |
| RGP1 | S179 | -0.11 | -0.22 | 0.09 | 0.10 | -0.02 | 0.13 | 0.21 |
| RGS12 | S850 | 0.16 | 0.24 | -0.13 | 0.11 | -0.41 | -0.07 | 0.319 |
| RGS3 | S806 | -0.29 | -0.21 | -0.24 | 0.23 | 0.00 | 0.38 | 0.016 |
| RGS3 | S1044 | 0.33 | -0.15 | 0.25 | -0.27 | -0.13 | -0.15 | 0.102 |
| RHBDF1 | S76 | -0.15 | -0.02 | 0.26 | -0.13 | -0.03 | 0.03 | 0.586 |
| RHBDF2 | S385 | -0.04 | 0.24 | -0.22 | -0.02 | 0.12 | -0.13 | 0.971 |
| RHOBTB2 | S344 | 0.02 | -0.09 | 0.08 | -0.19 | 0.10 | 0.06 | 0.932 |
| RHOG | T180 | -0.14 | 0.13 | 0.33 | -0.29 | -0.22 | 0.10 | 0.248 |
| RHOV | T18 | 0.26 | 0.14 | 0.02 | -0.28 | -0.22 | 0.00 | 0.046 |
| RIC1 | T/S | 0.01 | -0.03 | 0.04 | 0.26 | -0.09 | -0.23 | 0.875 |
| RIC8A | S436; T441 | -0.30 | -0.17 | -0.35 | 0.24 | 0.19 | 0.26 | 9E-04 |
| RIC8A | S436; T441 | -0.13 | -0.26 | -0.24 | 0.22 | 0.00 | 0.31 | 0.018 |
| RICTOR | S21 | -0.12 | -0.02 | 0.07 | -0.02 | -0.14 | 0.20 | 0.747 |
| RICTOR | T1172 | 0.02 | 0.02 | 0.05 | -0.03 | -0.03 | -0.02 | 0.003 |
| RICTOR | S/T | 0.07 | 0.02 | 0.01 | -0.04 | 0.00 | -0.07 | 0.061 |
| RICTOR | S1591 | 0.23 | -0.21 | 0.28 | 0.23 | -0.24 | -0.45 | 0.385 |
| RIF1 | S1876 | -0.76 | -0.82 | -0.88 | 0.36 | 0.64 | 0.54 | 1E-04 |
| RIF1 | T1980 | -0.24 | -0.26 | -0.18 | -0.07 | 0.09 | 0.51 | 0.085 |
| RIF1 | S1851; S1854 | -0.11 | -0.30 | -0.17 | 0.33 | 0.04 | 0.13 | 0.025 |
| RIF1 | S1851; S1854 | -0.11 | -0.30 | -0.17 | 0.33 | 0.04 | 0.13 | 0.025 |
| RIF1 | S/T | -0.08 | -0.01 | -0.28 | -0.11 | 0.26 | 0.15 | 0.173 |
| RIF1 | S2161 | -0.19 | -0.04 | -0.05 | 0.01 | -0.03 | 0.26 | 0.171 |
| RIF1 | S/T/Y | -0.31 | -0.08 | 0.14 | 0.06 | 0.16 | -0.02 | 0.355 |
| RIF1 | S2161 | -0.02 | -0.03 | -0.12 | -0.08 | -0.03 | 0.26 | 0.398 |
| RIF1 | S1616 | -0.07 | 0.02 | -0.11 | -0.12 | -0.07 | 0.30 | 0.562 |
| RIF1 | S1554 | 0.11 | -0.05 | -0.10 | -0.17 | 0.01 | 0.17 | 0.92 |
| RIF1 | S2161 | 0.02 | -0.06 | 0.04 | 0.08 | -0.07 | -0.01 | 0.976 |
| RIF1 | S1688 | 0.12 | -0.13 | 0.07 | -0.39 | -0.08 | 0.31 | 0.755 |
| RIF1 | S | 0.06 | -0.01 | 0.12 | -0.31 | -0.13 | 0.20 | 0.431 |
| RIF1 | S | 0.06 | -0.01 | 0.12 | -0.31 | -0.13 | 0.20 | 0.431 |
| RIF1 | S2144 | -0.16 | 0.29 | 0.16 | -0.03 | -0.14 | -0.19 | 0.191 |
| RIF1 | S1854 | 0.20 | 0.02 | 0.10 | -0.16 | -0.10 | -0.11 | 0.014 |
| RIF1 | S782 | 0.24 | 0.12 | 0.04 | -0.15 | 0.09 | -0.45 | 0.146 |
| RIF1 | S1454 | 0.17 | 0.47 | -0.21 | -0.37 | 0.12 | -0.39 | 0.241 |
| RIF1 | S2465 | 0.12 | 0.07 | 0.41 | 0.08 | -0.21 | -0.70 | 0.127 |
| RIF1 | T1161 | 0.32 | 0.36 | 0.53 | -0.55 | -0.23 | -1.05 | 0.015 |
| RIN1 | S333; T/S | -0.27 | -0.27 | -0.12 | 0.15 | -0.09 | 0.46 | 0.077 |
| RIN1 | S333 | -0.03 | 0.06 | 0.18 | -0.34 | 0.04 | 0.04 | 0.325 |
| RING1 | S170; S190 | -0.30 | 0.07 | -0.32 | 0.15 | 0.18 | 0.13 | 0.056 |
| RING1 | S187; S188; S190 | 0.08 | -0.20 | -0.30 | 0.05 | 0.03 | 0.27 | 0.132 |
| RING1 | S187; S188; S190 | -0.17 | -0.07 | -0.15 | 0.11 | 0.03 | 0.21 | 0.014 |

| | | | | | | | | |
|---|---|---|---|---|---|---|---|---|
| RING1 | T166; T167; T/S | -0.23 | -0.39 | 0.28 | 0.17 | -0.04 | 0.10 | 0.418 |
| RING1 | S187; S188; S190 | -0.28 | -0.07 | 0.07 | -0.05 | -0.09 | 0.34 | 0.412 |
| RING1 | S38 | 0.01 | -0.07 | -0.16 | 0.01 | 0.10 | 0.09 | 0.065 |
| RING1 | S140 | -0.11 | 0.02 | -0.10 | 0.16 | -0.19 | 0.19 | 0.402 |
| RING1 | S229; S232 | -0.03 | -0.05 | -0.05 | 0.02 | -0.03 | 0.13 | 0.153 |
| RING1 | S170; S/T | 0.04 | -0.03 | 0.01 | 0.03 | -0.03 | -0.01 | 0.713 |
| RING1 | S170 | 0.04 | 0.12 | -0.04 | -0.04 | -0.09 | 0.00 | 0.177 |
| RING1 | S/T | -0.01 | 0.08 | 0.09 | -0.06 | -0.06 | -0.05 | 0.024 |
| RING1 | S170 | 0.07 | 0.12 | -0.02 | -0.02 | -0.10 | -0.06 | 0.072 |
| RING1 | S38 | 0.11 | 0.12 | 0.03 | -0.18 | 0.03 | -0.14 | 0.056 |
| RING1 | S/T | 0.02 | -0.06 | 0.40 | -0.35 | -0.10 | -0.01 | 0.19 |
| RIOK2 | S/Y | -0.37 | 0.15 | 0.15 | 0.02 | 0.00 | -0.01 | 0.881 |
| RIOX1 | S109 | 0.03 | -0.57 | -0.48 | 0.54 | 0.19 | -0.01 | 0.077 |
| RIOX1 | S109 | 0.01 | 0.23 | -0.17 | 0.13 | -0.33 | 0.05 | 0.708 |
| RIPK1 | S610 | 0.05 | -0.46 | 0.10 | 0.20 | 0.15 | -0.14 | 0.445 |
| RIPK1 | S610 | -0.02 | -0.15 | -0.05 | 0.07 | 0.07 | 0.06 | 0.026 |
| RLF | S632 | 0.18 | -0.11 | -0.15 | 0.07 | -0.09 | 0.08 | 0.712 |
| RLF | S1273 | 0.13 | -0.18 | 0.01 | -0.08 | -0.01 | 0.11 | 0.848 |
| RLF | S632 | -0.01 | 0.00 | -0.03 | -0.02 | -0.10 | 0.14 | 0.797 |
| RLF | S1043 | 0.12 | 0.08 | 0.13 | 0.11 | -0.06 | -0.47 | 0.218 |
| RLIM | S230 | 0.21 | -0.05 | 0.12 | -0.03 | -0.12 | -0.16 | 0.082 |
| RMDN3 | S46 | -0.09 | -0.11 | -0.13 | 0.06 | 0.06 | 0.18 | 0.008 |
| RMDN3 | S46 | -0.08 | -0.11 | -0.10 | 0.11 | 0.02 | 0.14 | 0.008 |
| RMDN3 | S50; S/Y/T | -0.09 | -0.07 | 0.22 | -0.23 | -0.09 | 0.21 | 0.742 |
| RMDN3 | S212 | 0.11 | 0.31 | 0.00 | -0.72 | 0.12 | -0.03 | 0.271 |
| RMI2 | S20 | -0.30 | -0.11 | -0.43 | 0.35 | -0.37 | 0.56 | 0.192 |
| RNASEH2A | S259 | -0.05 | 0.33 | -0.23 | -0.15 | -0.14 | 0.16 | 0.781 |
| RNF11 | S14 | -0.12 | -0.14 | 0.01 | 0.08 | -0.03 | 0.18 | 0.116 |
| RNF11 | S25 | -0.28 | 0.27 | -0.20 | -0.58 | 0.14 | 0.40 | 0.867 |
| RNF113A | S253 | -0.05 | 0.08 | -0.15 | 0.19 | 0.02 | -0.12 | 0.557 |
| RNF113A | S/Y | 0.16 | -0.05 | -0.16 | 0.14 | -0.09 | -0.03 | 0.845 |
| RNF113A | S253 | 0.11 | -0.02 | -0.08 | 0.04 | 0.04 | -0.10 | 0.889 |
| RNF113A | S329 | -0.40 | 0.32 | 0.29 | 0.17 | 0.10 | -0.79 | 0.564 |
| RNF14 | S30 | 0.10 | -0.30 | -0.34 | 0.10 | 0.28 | 0.06 | 0.1 |
| RNF14 | S451 | -0.28 | 0.02 | 0.20 | -0.35 | -0.08 | 0.37 | 0.998 |
| RNF168 | T208 | 0.05 | -0.03 | 0.09 | 0.11 | -0.03 | -0.22 | 0.469 |
| RNF168 | S481 | 0.05 | 0.07 | 0.09 | -0.02 | -0.12 | -0.07 | 0.009 |
| RNF168 | S/T | 0.05 | 0.07 | 0.09 | -0.07 | 0.03 | -0.19 | 0.091 |
| RNF169 | S403 | -0.03 | -0.12 | -0.01 | 0.08 | 0.02 | 0.05 | 0.049 |
| RNF169 | S/T | 0.03 | 0.12 | -0.10 | -0.07 | -0.10 | 0.11 | 0.722 |
| RNF169 | S520 | -0.06 | 0.03 | 0.09 | -0.27 | 0.01 | 0.17 | 0.722 |
| RNF169 | T554 | 0.16 | 0.10 | 0.15 | -0.17 | -0.08 | -0.21 | 0.002 |
| RNF169 | S644 | -0.01 | 0.19 | 0.27 | -0.15 | -0.39 | 0.00 | 0.078 |
| RNF185 | T106 | 0.04 | -0.02 | -0.12 | 0.13 | 0.00 | -0.03 | 0.398 |
| RNF185 | S11 | -0.13 | -0.05 | -0.01 | -0.66 | 0.51 | 0.09 | 0.906 |
| RNF185 | S11 | -0.13 | -0.05 | -0.01 | -0.66 | 0.51 | 0.09 | 0.906 |
| RNF185 | S11 | -0.13 | -0.05 | -0.01 | -0.66 | 0.51 | 0.09 | 0.906 |
| RNF19A | S284 | 0.15 | 0.00 | 0.12 | 0.04 | -0.06 | -0.30 | 0.148 |

| | | | | | | | | |
|---|---|---|---|---|---|---|---|---|
| RNF19A | S574 | -0.47 | 0.50 | 0.19 | 0.35 | -0.81 | -0.19 | 0.542 |
| RNF20 | S138 | -0.18 | -0.07 | -0.03 | -0.02 | -0.07 | 0.31 | 0.254 |
| RNF213 | S5110 | -0.42 | -0.48 | -0.33 | 0.48 | -0.48 | 0.71 | 0.153 |
| RNF213 | S1258 | -0.06 | -0.15 | -0.14 | -0.09 | 0.32 | 0.06 | 0.161 |
| RNF213 | S1258 | 0.04 | 0.03 | 0.04 | 0.17 | -0.10 | -0.21 | 0.497 |
| RNF213 | S217 | -0.78 | -0.13 | 0.84 | -0.54 | 0.09 | -0.06 | 0.789 |
| RNF216 | S419 | -0.07 | 0.19 | -0.21 | 0.40 | 0.35 | -1.17 | 0.84 |
| RNF31 | S466 | -0.17 | -0.24 | -0.16 | 0.15 | 0.11 | 0.23 | 0.001 |
| RNF6 | S468 | 0.29 | -0.06 | 0.24 | -0.47 | 0.00 | -0.13 | 0.119 |
| RNMT | T77 | 0.25 | 0.11 | 0.45 | -0.41 | -0.09 | -0.57 | 0.021 |
| RNPC3 | S21 | 0.06 | -0.08 | -0.13 | 0.13 | -0.04 | 0.04 | 0.283 |
| RNPC3 | S21 | 0.09 | 0.14 | 0.09 | 0.00 | 0.00 | -0.37 | 0.131 |
| RNPC3 | S108 | 0.15 | 0.21 | 0.04 | -0.13 | -0.05 | -0.27 | 0.028 |
| RNPS1 | S52 | 0.15 | 0.08 | -0.09 | 0.02 | -0.02 | -0.15 | 0.323 |
| RNPS1 | S155; S157 | 0.20 | 0.10 | -0.16 | 0.06 | -0.07 | -0.15 | 0.457 |
| RNPS1 | S155; S157 | 0.18 | 0.08 | -0.03 | -0.03 | -0.03 | -0.21 | 0.123 |
| RNPS1 | S155; S157 | 0.18 | 0.12 | 0.08 | -0.06 | -0.03 | -0.35 | 0.058 |
| RNPS1 | S251 | 0.31 | 0.31 | 0.27 | -0.21 | -0.06 | -1.04 | 0.075 |
| RPA2 | S/T/Y | -0.28 | -0.19 | 0.01 | -0.33 | 0.19 | 0.43 | 0.363 |
| RPA2 | S/T/Y | -0.32 | -0.26 | 0.27 | -0.05 | -0.09 | 0.31 | 0.521 |
| RPL12 | S165 | -0.01 | -0.30 | -0.32 | 0.21 | 0.24 | 0.08 | 0.026 |
| RPL12 | S38 | 0.07 | 0.05 | 0.15 | 0.01 | -0.03 | -0.26 | 0.11 |
| RPL13 | S109 | -0.06 | -0.04 | 0.02 | 0.13 | 0.10 | -0.16 | 0.643 |
| RPL13 | S140 | 0.12 | 0.06 | 0.07 | 0.01 | 0.09 | -0.41 | 0.291 |
| RPL13 | S106 | -0.05 | 0.12 | 0.22 | -0.02 | 0.15 | -0.54 | 0.353 |
| RPL15 | S34 | 0.17 | 0.30 | 0.31 | -0.80 | 0.01 | -0.27 | 0.065 |
| RPL17 | S5 | 0.31 | 0.32 | 0.51 | -0.53 | -0.22 | -0.91 | 0.011 |
| RPL18A | S123 | -0.35 | 0.07 | -0.06 | 0.28 | 0.16 | -0.20 | 0.359 |
| RPL18A | S71 | 0.06 | 0.12 | -0.07 | -0.13 | 0.07 | -0.08 | 0.375 |
| RPL27 | S86 | 0.16 | 0.06 | 0.36 | -0.03 | -0.19 | -0.51 | 0.058 |
| RPL29 | S158 | 0.25 | 0.35 | 0.05 | -0.28 | 0.03 | -0.61 | 0.071 |
| RPL30 | S10 | 0.05 | 0.06 | 0.03 | 0.04 | -0.01 | -0.19 | 0.214 |
| RPL34 | S12 | 0.51 | -0.17 | -0.11 | 0.07 | -0.24 | -0.21 | 0.451 |
| RPL4 | S66 | -0.10 | -0.03 | 0.12 | 0.18 | -0.03 | -0.17 | 0.99 |
| RPL4 | S66 | -0.10 | -0.03 | 0.12 | 0.18 | -0.03 | -0.17 | 0.99 |
| RPL4 | S295 | 0.01 | 0.08 | 0.11 | 0.26 | -0.14 | -0.40 | 0.457 |
| RPL5 | S185 | -0.13 | -0.16 | -0.21 | -0.02 | 0.09 | 0.35 | 0.051 |
| RPLP0 | S304; S307 | -0.17 | -0.08 | -0.01 | 0.18 | -0.09 | 0.14 | 0.156 |
| RPLP0 | S304 | -0.06 | 0.06 | 0.02 | 0.11 | -0.01 | -0.14 | 0.784 |
| RPLP0 | S307 | 0.00 | -0.04 | 0.20 | -0.01 | 0.05 | -0.22 | 0.352 |
| RPLP0 | S304 | 0.03 | 0.10 | 0.02 | 0.13 | 0.14 | -0.53 | 0.575 |
| RPLP0 | S304; S307 | 0.23 | 0.13 | 0.07 | 0.00 | -0.16 | -0.35 | 0.051 |
| RPLP0 | S307 | 0.10 | 0.28 | 0.20 | -0.07 | -0.09 | -0.56 | 0.063 |
| RPLP2 | S102; S105 | -0.04 | -0.11 | -0.02 | 0.16 | 0.03 | -0.03 | 0.13 |
| RPLP2 | S102; S105 | -0.11 | -0.13 | 0.07 | 0.47 | -0.15 | -0.27 | 0.775 |
| RPLP2 | S102; S105 | 0.01 | -0.07 | 0.05 | 0.07 | -0.04 | -0.03 | 0.978 |
| RPLP2 | S105 | -0.08 | 0.18 | -0.06 | 0.08 | -0.15 | 0.00 | 0.767 |
| RPLP2 | S102 | 0.13 | 0.19 | 0.15 | 0.01 | 0.04 | -0.69 | 0.195 |

| | | | | | | | | |
|---|---|---|---|---|---|---|---|---|
| RPLP2 | S102 | 0.14 | 0.22 | 0.14 | -0.01 | -0.02 | -0.62 | 0.133 |
| RPLP2 | S105 | 0.22 | 0.14 | 0.22 | -0.11 | -0.01 | -0.63 | 0.082 |
| RPP25 | S162 | 0.27 | 0.12 | 0.07 | 0.17 | -0.39 | -0.38 | 0.141 |
| RPP25 | S/Y | 0.19 | 0.25 | 0.04 | 0.02 | -0.21 | -0.40 | 0.055 |
| RPP30 | S251 | 0.38 | 0.23 | 0.06 | -0.23 | -0.27 | -0.34 | 0.007 |
| RPP38 | T/S | 0.16 | 0.04 | -0.10 | 0.12 | 0.04 | -0.33 | 0.601 |
| RPRD1A | S156 | -0.08 | -0.15 | -0.09 | 0.05 | -0.01 | 0.25 | 0.062 |
| RPRD1A | S156 | -0.02 | 0.04 | 0.07 | -0.04 | -0.06 | 0.00 | 0.11 |
| RPRD1B | S128; S | -0.10 | -0.27 | -0.08 | 0.00 | 0.20 | 0.19 | 0.033 |
| RPRD1B | S134 | 0.21 | 0.11 | 0.03 | -0.14 | 0.02 | -0.26 | 0.067 |
| RPRD1B | S134 | 0.16 | 0.12 | 0.07 | -0.10 | -0.02 | -0.27 | 0.032 |
| RPRD1B | S171 | 0.20 | 0.16 | 0.45 | -0.52 | -0.12 | -0.43 | 0.014 |
| RPRD2 | T723 | -0.15 | -0.02 | -0.65 | 0.19 | -0.24 | 0.58 | 0.215 |
| RPRD2 | T723 | -0.04 | -0.28 | -0.27 | 0.42 | -0.44 | 0.39 | 0.331 |
| RPRD2 | S614 | 0.08 | -0.22 | -0.27 | 0.07 | -0.05 | 0.31 | 0.182 |
| RPRD2 | T723 | -0.04 | -0.17 | -0.15 | 0.08 | -0.08 | 0.30 | 0.132 |
| RPRD2 | S374 | 0.03 | -0.06 | -0.08 | 0.03 | 0.02 | 0.05 | 0.124 |
| RPRD2 | S374 | 0.03 | -0.02 | -0.10 | 0.02 | -0.01 | 0.08 | 0.276 |
| RPRD2 | S | -0.07 | -0.05 | 0.08 | 0.10 | -0.18 | 0.10 | 0.858 |
| RPRD2 | T358 | 0.01 | 0.01 | 0.17 | -0.10 | -0.01 | -0.09 | 0.096 |
| RPRD2 | S593 | 0.11 | 0.03 | 0.06 | -0.06 | -0.12 | -0.05 | 0.011 |
| RPRD2 | S593 | 0.12 | 0.08 | 0.01 | -0.08 | -0.07 | -0.07 | 0.01 |
| RPRD2 | T664 | 0.03 | 0.12 | 0.10 | -0.09 | -0.01 | -0.18 | 0.034 |
| RPRD2 | S758 | 0.05 | 0.15 | 0.12 | -0.12 | -0.09 | -0.14 | 0.003 |
| RPRD2 | S665 | 0.13 | 0.09 | 0.15 | -0.08 | 0.01 | -0.35 | 0.074 |
| RPRD2 | S479 | 0.43 | 0.19 | 0.35 | -0.58 | -0.14 | -0.59 | 0.01 |
| RPS10 | S146 | -0.23 | -0.12 | 0.15 | -0.04 | 0.04 | 0.15 | 0.396 |
| RPS10 | S146 | -0.14 | -0.08 | 0.14 | 0.07 | -0.10 | 0.08 | 0.719 |
| RPS14 | S/T | -0.39 | -0.10 | 0.07 | -0.46 | 0.07 | 0.56 | 0.577 |
| RPS2 | S264 | -0.05 | -0.09 | -0.44 | 0.44 | -0.06 | 0.07 | 0.156 |
| RPS26 | T96 | 0.11 | 0.08 | 0.25 | -0.17 | 0.00 | -0.35 | 0.05 |
| RPS27 | S11 | 0.03 | -0.05 | -0.05 | 0.07 | 0.05 | -0.04 | 0.285 |
| RPS27 | S11 | -0.06 | 0.12 | 0.00 | 0.17 | 0.00 | -0.26 | 0.712 |
| RPS27L | S11 | 0.14 | -0.19 | -0.21 | 0.43 | -0.28 | -0.02 | 0.613 |
| RPS3 | T221 | -0.03 | -0.09 | -0.15 | -0.05 | 0.11 | 0.18 | 0.092 |
| RPS3 | T221 | 0.02 | -0.06 | -0.10 | 0.12 | 0.11 | -0.11 | 0.36 |
| RPS3A | S236 | 0.07 | -0.12 | 0.07 | 0.10 | 0.02 | -0.16 | 0.823 |
| RPS3A | S236 | 0.06 | -0.04 | 0.19 | -0.18 | -0.06 | 0.02 | 0.188 |
| RPS3A | S263 | 0.08 | -0.03 | 0.23 | -0.17 | -0.01 | -0.13 | 0.093 |
| RPS5 | S142 | -0.26 | -0.17 | 0.14 | 0.12 | -0.21 | 0.29 | 0.434 |
| RPS6 | S240; S/T | -0.16 | -0.26 | -0.15 | 0.25 | 0.13 | 0.12 | 0.003 |
| RPS6KA4 | S347 | -0.13 | -0.46 | 0.02 | 0.16 | 0.10 | 0.20 | 0.077 |
| RPS6KA4 | S745 | 0.06 | 0.01 | 0.06 | -0.08 | 0.09 | -0.15 | 0.313 |
| RPS6KC1 | S282 | 0.07 | 0.27 | -0.03 | -0.17 | -0.26 | 0.05 | 0.151 |
| RPS8 | T130 | -0.19 | -0.31 | -0.48 | 0.35 | 0.27 | 0.16 | 0.004 |
| RPS8 | T130 | -0.30 | -0.49 | -0.19 | 0.58 | 0.02 | 0.14 | 0.041 |
| RPS9 | S163 | 0.01 | -0.17 | -0.23 | 0.10 | -0.02 | 0.26 | 0.087 |
| RPTOR | S722 | 0.03 | -0.32 | -0.34 | 0.25 | 0.13 | 0.14 | 0.036 |

| | | | | | | | | |
|---|---|---|---|---|---|---|---|---|
| **RPTOR** | **S863; T/S** | 0.05 | -0.04 | -0.16 | -0.24 | 0.27 | 0.06 | 0.632 |
| **RPTOR** | **S877** | 0.11 | 0.16 | 0.08 | 0.07 | -0.16 | -0.31 | 0.089 |
| **RPTOR** | **S877** | 0.28 | 0.13 | 0.18 | -0.18 | -0.29 | -0.22 | 0.001 |
| **RPTOR** | **S863** | 0.09 | -0.05 | 0.53 | -0.10 | -0.07 | -0.65 | 0.147 |
| **RPUSD2** | **S68** | 0.08 | 0.05 | 0.31 | -0.06 | -0.12 | -0.34 | 0.057 |
| **RRAGC** | **S95** | -0.13 | 0.02 | 0.05 | -0.04 | 0.04 | 0.04 | 0.648 |
| **RRAGC** | **S95** | -0.03 | 0.14 | -0.16 | -0.02 | -0.01 | 0.07 | 0.755 |
| **RRAGC** | **T394** | -0.05 | -0.03 | 0.07 | 0.09 | -0.28 | 0.15 | 0.963 |
| **RRAGC** | **T394** | 0.15 | 0.13 | 0.13 | 0.08 | -0.20 | -0.36 | 0.083 |
| **RRAS** | **S201** | 0.17 | -0.04 | -0.10 | 0.11 | -0.07 | -0.08 | 0.821 |
| **RRBP1** | **T191** | -0.12 | -0.35 | 0.10 | 0.01 | 0.11 | 0.19 | 0.185 |
| **RRBP1** | **T255** | -0.02 | -0.04 | -0.02 | -0.02 | -0.25 | 0.29 | 0.838 |
| **RRBP1** | **S573** | -0.08 | -0.02 | 0.16 | 0.10 | -0.23 | 0.03 | 0.695 |
| **RRBP1** | **T** | 0.08 | -0.14 | 0.18 | -0.43 | -0.05 | 0.26 | 0.639 |
| **RRBP1** | **T255** | 0.08 | 0.08 | 0.08 | -0.09 | -0.16 | -0.01 | 0.02 |
| **RRBP1** | **S** | 0.12 | 0.18 | -0.05 | -0.32 | -0.03 | 0.05 | 0.244 |
| **RRBP1** | **S533** | 0.16 | 0.21 | 0.02 | -0.03 | -0.15 | -0.26 | 0.033 |
| **RRBP1** | **S** | 0.11 | 0.17 | 0.15 | -0.19 | -0.13 | -0.17 | 4E-04 |
| **RRBP1** | **T235** | 0.15 | 0.10 | 0.17 | 0.00 | 0.01 | -0.54 | 0.165 |
| **RRBP1** | **T255** | 0.15 | 0.15 | 0.16 | -0.07 | -0.21 | -0.24 | 0.004 |
| **RRBP1** | **S583** | 0.12 | 0.20 | 0.20 | -0.16 | -0.29 | -0.15 | 0.002 |
| **RRBP1** | **S583** | 0.25 | 0.29 | 0.04 | 0.00 | -0.25 | -0.48 | 0.053 |
| **RRBP1** | **T275** | 0.20 | 0.35 | 0.13 | -0.14 | -0.26 | -0.41 | 0.009 |
| **RRBP1** | **T275** | 0.29 | 0.23 | 0.39 | -0.40 | -0.20 | -0.58 | 0.004 |
| **RRBP1** | **T235** | 0.10 | 0.33 | 0.53 | -0.39 | -0.27 | -0.67 | 0.011 |
| **RREB1** | **S1429** | -0.09 | -0.40 | 0.02 | -0.01 | 0.00 | 0.37 | 0.195 |
| **RREB1** | **S/T** | -0.09 | -0.40 | 0.02 | -0.01 | 0.00 | 0.37 | 0.195 |
| **RREB1** | **S/T** | -0.14 | -0.29 | 0.08 | 0.03 | -0.11 | 0.34 | 0.295 |
| **RREB1** | **S1167** | 0.04 | -0.19 | -0.13 | 0.17 | 0.06 | 0.01 | 0.111 |
| **RREB1** | **S1167; S1174; S1175** | -0.17 | -0.10 | 0.01 | 0.12 | -0.05 | 0.17 | 0.118 |
| **RREB1** | **S1320** | -0.23 | -0.05 | 0.03 | 0.05 | -0.07 | 0.23 | 0.257 |
| **RREB1** | **S1356; S1357; S1360** | 0.09 | 0.11 | -0.44 | 0.01 | -0.04 | 0.19 | 0.532 |
| **RREB1** | **S1388** | -0.26 | -0.13 | 0.18 | -0.08 | 0.14 | 0.10 | 0.454 |
| **RREB1** | **S231** | -0.01 | -0.08 | -0.01 | 0.04 | -0.03 | 0.09 | 0.178 |
| **RREB1** | **S161** | 0.02 | -0.02 | -0.07 | 0.01 | -0.08 | 0.13 | 0.529 |
| **RREB1** | **S175** | 0.11 | -0.03 | -0.12 | -0.04 | 0.07 | -0.01 | 0.829 |
| **RREB1** | **S/T** | -0.11 | 0.20 | -0.14 | -0.13 | -0.16 | 0.28 | 0.953 |
| **RREB1** | **S175** | 0.08 | -0.18 | 0.08 | 0.03 | -0.09 | 0.06 | 0.943 |
| **RREB1** | **S161** | 0.17 | -0.68 | 0.30 | 0.25 | 0.32 | -0.78 | 0.995 |
| **RREB1** | **S1475** | -0.06 | 0.02 | 0.04 | -0.13 | -0.01 | 0.12 | 0.949 |
| **RREB1** | **S42** | 0.09 | 0.00 | -0.09 | -0.03 | -0.14 | 0.15 | 0.952 |
| **RREB1** | **S180** | -0.13 | 0.13 | 0.01 | 0.08 | -0.09 | -0.01 | 0.902 |
| **RREB1** | **S1140** | 0.05 | 0.02 | -0.05 | 0.02 | -0.10 | 0.05 | 0.827 |
| **RREB1** | **S231** | -0.18 | 0.10 | 0.09 | -0.21 | 0.10 | 0.07 | 0.913 |
| **RREB1** | **S1360** | 0.12 | -0.05 | -0.06 | -0.06 | -0.21 | 0.21 | 0.855 |
| **RREB1** | **S1140** | 0.12 | -0.02 | -0.05 | -0.05 | -0.05 | 0.05 | 0.632 |
| **RREB1** | **S1122** | 0.04 | -0.02 | 0.03 | -0.03 | -0.02 | 0.00 | 0.156 |
| **RREB1** | **S1167** | 0.12 | 0.05 | -0.10 | 0.04 | -0.14 | 0.02 | 0.608 |

| | | | | | | | | |
|---|---|---|---|---|---|---|---|---|
| RREB1 | S1356; S1357; S1360 | 0.29 | -0.25 | -0.04 | 0.43 | -0.10 | -0.54 | 0.843 |
| RREB1 | S36 | 0.03 | 0.02 | 0.14 | -0.12 | -0.03 | -0.05 | 0.058 |
| RREB1 | S1134; S1135; S1140 | 0.03 | 0.02 | 0.15 | -0.10 | -0.05 | -0.06 | 0.039 |
| RREB1 | S1134; S1135; S/T | -0.10 | 0.15 | 0.19 | -0.28 | -0.08 | 0.06 | 0.252 |
| RREB1 | S161 | 0.11 | 0.15 | 0.01 | -0.08 | -0.08 | -0.14 | 0.016 |
| RREB1 | S42 | 0.15 | 0.15 | 0.10 | -0.31 | 0.03 | -0.19 | 0.045 |
| RREB1 | S1356; S1357; S1360 | 0.08 | 0.33 | -0.02 | -0.25 | -0.32 | 0.07 | 0.134 |
| RREB1 | S1134; S1135 | 0.15 | 0.12 | 0.15 | -0.12 | -0.02 | -0.33 | 0.032 |
| RREB1 | S1219 | 0.26 | 0.19 | 0.10 | -0.29 | -0.19 | -0.15 | 0.003 |
| RREB1 | S36; S42 | 0.29 | 0.15 | 0.14 | -0.14 | -0.09 | -0.48 | 0.032 |
| RREB1 | S36 | 0.24 | 0.17 | 0.31 | -0.33 | -0.03 | -0.57 | 0.027 |
| RREB1 | S36; S42 | 0.25 | 0.21 | 0.33 | -0.32 | 0.01 | -0.78 | 0.055 |
| RREB1 | S161 | 0.20 | 0.28 | 0.35 | -0.32 | 0.10 | -1.04 | 0.107 |
| RREB1 | S42 | 0.47 | 0.48 | 0.05 | -0.68 | -0.22 | -0.51 | 0.014 |
| RRM2 | S/T | -0.07 | -0.11 | -0.29 | 0.00 | -0.11 | 0.47 | 0.221 |
| RRN3 | S44 | -0.43 | -0.89 | -0.29 | 0.28 | 0.08 | 0.70 | 0.026 |
| RRN3 | S170; S172 | -0.09 | 0.00 | 0.18 | -0.02 | 0.08 | -0.17 | 0.549 |
| RRN3 | S170; S172 | 0.03 | 0.20 | 0.09 | -0.24 | 0.01 | -0.14 | 0.062 |
| RRP1 | S383 | -0.19 | -0.27 | -0.21 | 0.26 | -0.02 | 0.32 | 0.018 |
| RRP1 | S291 | 0.01 | -0.36 | 0.00 | 0.19 | 0.18 | -0.09 | 0.242 |
| RRP1 | S291 | -0.08 | -0.12 | 0.03 | 0.07 | 0.02 | 0.06 | 0.095 |
| RRP1 | S383 | -0.10 | -0.07 | 0.15 | 0.01 | 0.09 | -0.10 | 0.941 |
| RRP12 | T77 | -0.18 | -0.18 | -0.25 | 0.23 | -0.17 | 0.42 | 0.106 |
| RRP12 | S1080 | -0.03 | -0.22 | -0.18 | 0.07 | -0.14 | 0.41 | 0.201 |
| RRP12 | S180 | -0.35 | 0.02 | -0.19 | 0.23 | -0.65 | 0.59 | 0.579 |
| RRP12 | S1080 | 0.33 | -0.26 | -0.29 | 0.40 | -0.25 | -0.10 | 0.77 |
| RRP15 | T104 | -0.28 | -0.60 | -0.79 | 0.66 | 0.39 | 0.06 | 0.016 |
| RRP15 | S11 | 0.00 | 0.04 | 0.09 | -0.12 | 0.04 | -0.05 | 0.184 |
| RRP15 | S266; S/T | 0.18 | 0.30 | 0.41 | -0.24 | -0.14 | -0.87 | 0.04 |
| RRP1B | S513 | -0.03 | -0.45 | -0.33 | 0.15 | 0.04 | 0.45 | 0.051 |
| RRP1B | S706 | -0.45 | -0.11 | -0.05 | 0.00 | 0.60 | -0.21 | 0.29 |
| RRP1B | S732; S735 | 0.01 | -0.11 | 0.15 | 0.06 | -0.15 | 0.03 | 0.714 |
| RRP1B | S392 | 0.06 | 0.01 | 0.00 | 0.04 | 0.01 | -0.12 | 0.395 |
| RRP1B | S/T | -0.11 | 0.15 | 0.20 | 0.07 | -0.05 | -0.31 | 0.299 |
| RRP1B | S245 | 0.05 | 0.06 | 0.17 | -0.08 | -0.01 | -0.23 | 0.06 |
| RRP1B | S350 | 0.15 | 0.06 | 0.20 | -0.16 | 0.08 | -0.41 | 0.11 |
| RRP1B | S/T | 0.17 | 0.18 | 0.32 | -0.26 | 0.00 | -0.62 | 0.051 |
| RRP36 | S73 | 0.19 | 0.06 | 0.10 | -0.21 | 0.00 | -0.20 | 0.033 |
| RRP36 | S73 | 0.38 | 0.17 | 0.19 | -0.27 | -0.03 | -0.67 | 0.048 |
| RRP7A | S19 | 0.05 | -0.29 | -0.13 | 0.09 | 0.10 | 0.14 | 0.076 |
| RRP8 | S126 | -0.09 | -0.21 | 0.03 | 0.05 | -0.02 | 0.21 | 0.154 |
| RRP8 | S171; T/S | 0.28 | -0.07 | -0.32 | 0.32 | 0.04 | -0.41 | 0.953 |
| RRP8 | S176 | 0.14 | 0.11 | 0.24 | -0.21 | -0.17 | -0.18 | 0.001 |
| RRP9 | S50; S51; S53 | -0.16 | -0.06 | 0.10 | -0.10 | -0.12 | 0.28 | 0.687 |
| RRP9 | S51; S53 | -0.09 | 0.06 | 0.08 | 0.08 | -0.21 | 0.06 | 0.75 |
| RRP9 | S53 | 0.01 | 0.13 | 0.19 | -0.18 | -0.09 | -0.10 | 0.016 |
| RRP9 | S53 | 0.24 | 0.44 | 0.32 | -0.32 | -0.28 | -0.76 | 0.008 |
| RRP9 | S51; S53 | 0.34 | 0.48 | 0.26 | -0.44 | -0.04 | -1.23 | 0.059 |

| | | | | | | | | |
|---|---|---|---|---|---|---|---|---|
| **RRS1** | **T56** | -0.10 | 0.11 | -0.11 | 0.25 | -0.20 | -0.01 | 0.773 |
| **RSBN1** | **S81** | -0.02 | 0.02 | -0.12 | 0.10 | 0.05 | -0.05 | 0.317 |
| **RSF1** | **S524** | -1.04 | -1.03 | -0.71 | 0.71 | 0.57 | 0.37 | 5E-04 |
| **RSF1** | **S1277; T1278; S1282** | -0.34 | -0.16 | -0.58 | 0.03 | -0.07 | 0.74 | 0.104 |
| **RSF1** | **S492** | -0.44 | -0.32 | -0.06 | 0.25 | 0.31 | 0.11 | 0.017 |
| **RSF1** | **S492** | -0.44 | -0.32 | -0.06 | 0.25 | 0.31 | 0.11 | 0.017 |
| **RSF1** | **S1403** | -0.07 | 0.12 | -0.93 | 0.31 | 0.41 | -0.21 | 0.284 |
| **RSF1** | **T/S** | -0.09 | -0.39 | -0.35 | 0.04 | -0.19 | 0.69 | 0.176 |
| **RSF1** | **S1345** | 0.03 | -0.41 | -0.13 | 0.05 | 0.12 | 0.25 | 0.09 |
| **RSF1** | **S748** | -0.05 | -0.17 | -0.22 | 0.08 | -0.02 | 0.31 | 0.065 |
| **RSF1** | **S540** | -0.09 | -0.09 | -0.21 | 0.10 | -0.11 | 0.34 | 0.153 |
| **RSF1** | **S266** | 0.12 | -0.24 | -0.25 | 0.28 | -0.27 | 0.23 | 0.387 |
| **RSF1** | **S227** | 0.03 | -0.09 | -0.24 | 0.05 | 0.01 | 0.20 | 0.117 |
| **RSF1** | **S748** | 0.06 | -0.08 | -0.26 | 0.03 | 0.08 | 0.13 | 0.141 |
| **RSF1** | **S1105** | 0.15 | -0.32 | -0.02 | 0.07 | 0.30 | -0.27 | 0.687 |
| **RSF1** | **S473** | 0.03 | -0.11 | -0.06 | 0.02 | -0.05 | 0.15 | 0.299 |
| **RSF1** | **S1247** | -0.31 | 0.03 | 0.13 | 0.01 | -0.12 | 0.20 | 0.653 |
| **RSF1** | **S1282** | -0.17 | -0.13 | 0.15 | -0.20 | 0.25 | 0.04 | 0.655 |
| **RSF1** | **S1359** | -0.01 | -0.11 | 0.02 | -0.06 | 0.14 | 0.02 | 0.399 |
| **RSF1** | **T1305** | 0.23 | -0.20 | -0.08 | -0.20 | 0.22 | -0.03 | 0.956 |
| **RSF1** | **S1359** | 0.24 | -0.18 | -0.11 | -0.11 | 0.26 | -0.19 | 0.99 |
| **RSF1** | **S1345** | 0.01 | -0.01 | 0.00 | -0.06 | 0.06 | -0.01 | 0.977 |
| **RSF1** | **S604** | 0.03 | -0.02 | -0.01 | -0.13 | 0.11 | 0.01 | 0.929 |
| **RSF1** | **S473** | 0.05 | -0.04 | 0.03 | -0.03 | 0.05 | -0.06 | 0.566 |
| **RSF1** | **S1359; S1375** | -0.08 | -0.01 | 0.12 | -0.23 | 0.09 | 0.09 | 0.835 |
| **RSF1** | **S604** | 0.07 | 0.02 | -0.04 | -0.11 | 0.07 | -0.01 | 0.616 |
| **RSF1** | **S1245** | 0.03 | 0.07 | -0.04 | -0.21 | 0.07 | 0.06 | 0.672 |
| **RSF1** | **S622; S629** | 0.04 | 0.04 | 0.00 | -0.04 | 0.04 | -0.07 | 0.242 |
| **RSF1** | **S604** | 0.06 | 0.14 | -0.12 | -0.14 | -0.02 | 0.07 | 0.584 |
| **RSF1** | **T1305** | 0.11 | -0.17 | 0.12 | -0.37 | 0.13 | 0.12 | 0.757 |
| **RSF1** | **T1305** | -0.09 | -0.04 | 0.21 | -0.06 | -0.12 | 0.07 | 0.582 |
| **RSF1** | **S622** | 0.16 | -0.03 | -0.04 | -0.20 | 0.23 | -0.17 | 0.63 |
| **RSF1** | **T1305; S1310** | 0.24 | -0.42 | 0.28 | -0.21 | 0.05 | -0.07 | 0.672 |
| **RSF1** | **S397** | 0.14 | 0.01 | 0.02 | -0.13 | -0.07 | 0.02 | 0.124 |
| **RSF1** | **T408** | 0.07 | 0.07 | 0.02 | -0.11 | 0.11 | -0.19 | 0.264 |
| **RSF1** | **S622** | 0.13 | 0.07 | -0.03 | -0.15 | 0.02 | -0.06 | 0.148 |
| **RSF1** | **S1359; S1375** | 0.08 | 0.08 | 0.02 | -0.10 | 0.07 | -0.18 | 0.17 |
| **RSF1** | **S1096; S1098; S1105** | 0.05 | 0.03 | 0.15 | -0.18 | -0.07 | 0.00 | 0.065 |
| **RSF1** | **S874** | 0.10 | 0.12 | 0.03 | -0.21 | -0.05 | 0.00 | 0.073 |
| **RSF1** | **S1105** | 0.13 | 0.18 | 0.01 | -0.03 | 0.02 | -0.37 | 0.151 |
| **RSF1** | **T1305; S1310** | 0.43 | -0.60 | 0.36 | -0.61 | 0.02 | 0.06 | 0.574 |
| **RSF1** | **S1098; S1105** | 0.20 | 0.11 | 0.13 | -0.11 | -0.05 | -0.33 | 0.026 |
| **RSF1** | **S874** | 0.23 | 0.19 | 0.16 | -0.14 | -0.19 | -0.36 | 0.004 |
| **RSF1** | **S397** | 0.25 | 0.12 | 0.21 | -0.12 | -0.03 | -0.59 | 0.069 |
| **RSF1** | **S604** | 0.18 | 0.29 | 0.18 | -0.42 | 0.31 | -0.94 | 0.193 |
| **RSF1** | **S1226** | 0.41 | 0.51 | 0.83 | -0.75 | -0.79 | -1.73 | 0.008 |
| **RSF1** | **S1226** | 0.21 | 0.71 | 0.81 | -0.82 | -0.70 | -1.85 | 0.014 |
| **RSL1D1** | **T423; S427** | -0.45 | -0.58 | -0.29 | 0.57 | 0.18 | 0.21 | 0.007 |

| | | | | | | | | |
|---|---|---|---|---|---|---|---|---|
| **RSL1D1** | **S392; S396** | -0.25 | -0.47 | -0.36 | 0.50 | -0.10 | 0.39 | 0.033 |
| **RSL1D1** | **T415** | -0.37 | -0.41 | -0.19 | 0.32 | 0.21 | 0.25 | 0.001 |
| **RSL1D1** | **S392; S396** | -0.14 | -0.37 | -0.02 | 0.11 | 0.25 | 0.09 | 0.047 |
| **RSL1D1** | **T340** | -0.05 | -0.16 | -0.12 | 0.16 | 0.15 | -0.02 | 0.034 |
| **RSL1D1** | **S325** | -0.09 | 0.02 | -0.13 | 0.02 | 0.11 | 0.05 | 0.07 |
| **RSL1D1** | **S361** | 0.04 | -0.09 | -0.15 | -0.04 | -0.03 | 0.25 | 0.318 |
| **RSL1D1** | **T106** | 0.16 | -0.15 | -0.11 | 0.00 | -0.02 | 0.11 | 0.577 |
| **RSL1D1** | **S443** | 0.06 | -0.02 | -0.09 | 0.07 | 0.03 | -0.05 | 0.645 |
| **RSL1D1** | **S361** | 0.05 | 0.01 | -0.08 | 0.05 | -0.05 | 0.01 | 0.829 |
| **RSL1D1** | **S427** | 0.12 | 0.05 | -0.02 | 0.03 | -0.12 | -0.08 | 0.148 |
| **RSL1D1** | **S92** | 0.27 | 0.01 | 0.06 | -0.07 | -0.12 | -0.20 | 0.057 |
| **RSRC1** | **S254** | 0.30 | -0.02 | -0.08 | 0.06 | -0.14 | -0.17 | 0.336 |
| **RSRC1** | **S61** | 0.34 | 0.17 | 0.16 | -0.31 | -0.04 | -0.48 | 0.025 |
| **RSRC2** | **T/S** | 0.25 | 0.23 | 0.12 | -0.18 | -0.17 | -0.37 | 0.005 |
| **RSRC2** | **S32** | 0.23 | 0.16 | 0.26 | -0.19 | -0.07 | -0.56 | 0.031 |
| **RSRC2** | **S32** | 0.12 | 0.30 | 0.35 | -0.24 | -0.09 | -0.67 | 0.034 |
| **RSRC2** | **T/S** | 0.17 | 0.29 | 0.36 | -0.34 | -0.19 | -0.52 | 0.005 |
| **RTCB** | **S300** | -0.15 | -0.06 | 0.10 | -0.21 | 0.36 | -0.12 | 0.833 |
| **RTEL1** | **S847** | -0.81 | 0.24 | 0.01 | -0.13 | 0.37 | 0.05 | 0.457 |
| **RTKN** | **S115** | -0.18 | -0.23 | -0.46 | 0.43 | -0.09 | 0.32 | 0.048 |
| **RTKN** | **S529** | -0.20 | -0.21 | -0.17 | 0.08 | 0.15 | 0.28 | 0.003 |
| **RTKN** | **S106** | -0.35 | -0.04 | 0.04 | 0.11 | -0.01 | 0.19 | 0.176 |
| **RTN1** | **S563** | -0.15 | 0.11 | -0.82 | 0.10 | 0.00 | 0.46 | 0.203 |
| **RTN4** | **S181** | -0.10 | 0.03 | 0.16 | -0.22 | -0.03 | 0.13 | 0.61 |
| **RUBCN** | **S388** | -0.07 | -0.04 | 0.15 | 0.07 | 0.02 | -0.16 | 0.703 |
| **RUNX1** | **S212** | -0.08 | -0.18 | -0.04 | 0.33 | -0.08 | -0.01 | 0.246 |
| **RUNX1** | **T14; S21** | -0.04 | 0.01 | 0.09 | 0.00 | -0.15 | 0.07 | 0.587 |
| **RUNX1** | **T14** | 0.12 | 0.00 | 0.03 | -0.13 | 0.00 | -0.03 | 0.137 |
| **RUNX1** | **S21** | 0.14 | 0.01 | 0.08 | -0.48 | 0.44 | -0.41 | 0.488 |
| **RUSC1** | **S831** | 0.01 | -0.08 | -0.35 | 0.33 | -0.53 | 0.38 | 0.558 |
| **RWDD2B** | **S173** | -0.33 | 0.05 | -0.27 | 0.13 | 0.02 | 0.30 | 0.079 |
| **RXYLT1** | **S63** | 0.15 | 0.13 | 0.12 | -0.15 | -0.14 | -0.16 | 2E-05 |
| **RYBP** | **S227** | -0.07 | 0.01 | -0.10 | -0.02 | 0.13 | 0.03 | 0.143 |
| **RYBP** | **S/T** | -0.11 | 0.17 | -0.21 | 0.13 | -0.04 | 0.02 | 0.538 |
| **RYBP** | **S/T** | 0.20 | 0.00 | -0.05 | -0.11 | -0.11 | 0.04 | 0.292 |
| **RYBP** | **S99** | 0.20 | 0.19 | -0.09 | -0.27 | -0.01 | -0.08 | 0.138 |
| **RYBP** | **S201** | 0.09 | 0.25 | 0.15 | -0.61 | -0.03 | 0.00 | 0.144 |
| **RYBP** | **S201** | 0.32 | 0.39 | 0.03 | -0.39 | -0.21 | -0.35 | 0.01 |
| **S100A11** | **S6** | 0.11 | -0.03 | 0.04 | -0.08 | -0.02 | -0.03 | 0.146 |
| **S100A14** | **S77** | 0.16 | 0.18 | -0.12 | 0.00 | -0.03 | -0.25 | 0.251 |
| **S100A14** | **S73** | -0.04 | 0.13 | 0.24 | -0.13 | 0.10 | -0.38 | 0.192 |
| **S100A16** | **S101; S** | -0.16 | -0.07 | 0.02 | -0.03 | -0.09 | 0.28 | 0.396 |
| **S100A16** | **S** | -0.05 | -0.04 | 0.23 | -0.14 | -0.06 | 0.03 | 0.36 |
| **S100A16** | **S** | -0.15 | 0.02 | 0.34 | -0.25 | -0.20 | 0.16 | 0.437 |
| **S1PR3** | **S326** | -0.12 | -0.16 | -0.42 | 0.08 | 0.00 | 0.47 | 0.077 |
| **SAAL1** | **S6** | 0.05 | 0.10 | 0.11 | -0.16 | 0.10 | -0.24 | 0.157 |
| **SAAL1** | **S56** | 0.28 | -0.06 | 0.06 | -0.45 | -0.32 | 0.32 | 0.405 |
| **SAC3D1** | **S402** | 0.00 | -0.01 | -0.03 | 0.12 | 0.01 | -0.10 | 0.743 |

| | | | | | | | | |
|---|---|---|---|---|---|---|---|---|
| SAC3D1 | S402 | 0.00 | 0.08 | 0.04 | -0.07 | -0.16 | 0.09 | 0.318 |
| SAFB | S383; S384 | -0.12 | -0.58 | 0.22 | 0.02 | 0.15 | 0.17 | 0.319 |
| SAFB | S79 | -0.02 | -0.15 | -0.16 | 0.15 | -0.07 | 0.21 | 0.1 |
| SAFB | S604 | -0.08 | -0.04 | -0.14 | 0.14 | -0.09 | 0.18 | 0.14 |
| SAFB | S195 | -0.09 | -0.04 | -0.06 | -0.05 | 0.02 | 0.20 | 0.18 |
| SAFB | S196 | 0.02 | -0.07 | -0.07 | -0.08 | 0.03 | 0.15 | 0.352 |
| SAFB | S601; S604 | 0.01 | 0.02 | -0.08 | 0.04 | -0.13 | 0.13 | 0.744 |
| SAFB | S383 | -0.12 | -0.04 | 0.11 | 0.10 | 0.15 | -0.24 | 0.9 |
| SAFB | S383 | 0.16 | 0.13 | 0.15 | -0.09 | 0.18 | -0.71 | 0.249 |
| SAFB | S604 | 0.26 | 0.34 | 0.08 | -0.11 | -0.11 | -0.69 | 0.063 |
| SAFB2 | S287 | -1.82 | -1.73 | -1.63 | 0.74 | 0.67 | 0.87 | 7E-06 |
| SAFB2 | S262 | -0.73 | -0.74 | -0.93 | 0.65 | 0.39 | 0.48 | 2E-04 |
| SAFB2 | S233 | -0.40 | -0.43 | -0.47 | 0.28 | 0.38 | 0.34 | 2E-05 |
| SAFB2 | S233; S246 | -0.30 | -0.33 | -0.36 | 0.19 | 0.30 | 0.31 | 1E-04 |
| SAFB2 | T244; S245 | -0.28 | -0.24 | -0.35 | 0.22 | 0.20 | 0.30 | 3E-04 |
| SAFB2 | T201 | -0.28 | -0.23 | -0.16 | 0.24 | 0.17 | 0.17 | 6E-04 |
| SAFB2 | S330; S343 | -0.21 | -0.05 | -0.20 | 0.08 | 0.06 | 0.27 | 0.026 |
| SAFB2 | S233 | -0.17 | -0.13 | -0.13 | 0.11 | 0.22 | 0.05 | 0.005 |
| SAFB2 | S/T | -0.09 | -0.11 | -0.18 | 0.13 | 0.09 | 0.12 | 0.001 |
| SAFB2 | S207 | -0.18 | -0.15 | -0.04 | 0.13 | 0.05 | 0.15 | 0.01 |
| SAFB2 | S513 | -0.09 | -0.12 | -0.14 | 0.11 | 0.30 | -0.12 | 0.155 |
| SAFB2 | S343; S/T | -0.12 | -0.08 | -0.10 | 0.04 | 0.09 | 0.15 | 0.005 |
| SAFB2 | S321; S324; S343 | -0.18 | -0.18 | 0.08 | 0.04 | 0.11 | 0.10 | 0.118 |
| SAFB2 | S613 | 0.00 | 0.07 | -0.28 | -0.09 | 0.12 | 0.14 | 0.374 |
| SAFB2 | S/T | -0.14 | -0.06 | 0.00 | 0.11 | -0.08 | 0.15 | 0.192 |
| SAFB2 | S343 | -0.05 | -0.02 | 0.05 | 0.00 | -0.06 | 0.08 | 0.813 |
| SAFB2 | S321; S324; S343 | -0.01 | -0.06 | 0.06 | 0.03 | 0.01 | -0.03 | 0.871 |
| SAFB2 | S827 | -0.15 | 0.03 | 0.11 | 0.08 | -0.05 | -0.04 | 0.978 |
| SAFB2 | S787 | -0.09 | 0.07 | 0.10 | -0.04 | 0.11 | -0.18 | 0.556 |
| SAFB2 | S/T | -0.06 | 0.14 | 0.09 | -0.10 | 0.30 | -0.49 | 0.56 |
| SAFB2 | S246 | 0.24 | 0.32 | 0.27 | -0.34 | -0.04 | -0.71 | 0.031 |
| SAFB2 | S246 | 0.40 | 0.27 | 0.29 | -0.40 | 0.01 | -1.03 | 0.06 |
| SAMD1 | S161 | -0.16 | -0.12 | -0.10 | 0.02 | 0.08 | 0.24 | 0.022 |
| SAMD1 | T107 | -0.07 | -0.05 | -0.09 | 0.02 | 0.08 | 0.10 | 0.006 |
| SAMD1 | S261 | 0.00 | -0.18 | -0.01 | -0.10 | 0.24 | 0.01 | 0.392 |
| SAMD1 | T107 | -0.09 | -0.04 | 0.04 | -0.01 | -0.04 | 0.13 | 0.431 |
| SAMD1 | S161 | -0.26 | -0.01 | 0.45 | -0.40 | -0.09 | 0.14 | 0.523 |
| SAMD1 | S425 | 0.39 | 0.04 | 0.33 | -0.13 | -0.24 | -0.64 | 0.035 |
| SAMD15 | T348 | 0.14 | 0.16 | 0.09 | -0.17 | -0.06 | -0.20 | 0.005 |
| SAMD4B | S271 | 0.43 | -0.68 | -0.43 | 0.48 | -0.62 | 0.31 | 0.59 |
| SAMD4B | S271 | 0.07 | -0.23 | 0.01 | 0.39 | -0.37 | -0.01 | 0.839 |
| SAMHD1 | S33 | -0.28 | -0.32 | -0.42 | 0.69 | -0.54 | 0.41 | 0.229 |
| SAMHD1 | S33 | 0.12 | -0.11 | -0.29 | 0.04 | 0.18 | 0.02 | 0.256 |
| SAMHD1 | S278 | 0.10 | -0.10 | -0.02 | -0.11 | 0.05 | 0.05 | 0.968 |
| SAP130 | S855; T856 | -0.22 | -0.29 | -0.53 | 0.23 | -0.13 | 0.63 | 0.07 |
| SAP130 | S855; T/S | -0.24 | -0.21 | -0.46 | 0.15 | -0.07 | 0.58 | 0.064 |
| SAP130 | S875 | -0.01 | -0.01 | -0.18 | 0.14 | -0.10 | 0.13 | 0.272 |
| SAP130 | S875 | 0.13 | -0.11 | -0.12 | -0.09 | -0.05 | 0.21 | 0.677 |

| | | | | | | | | |
|---|---|---|---|---|---|---|---|---|
| SAP130 | T/S | 0.55 | -0.41 | 0.07 | -0.01 | -0.67 | 0.15 | 0.553 |
| SAP18 | S119 | 0.14 | -0.25 | 0.07 | 0.02 | 0.06 | -0.07 | 0.927 |
| SAP30L | S/T | -0.30 | -0.24 | -0.32 | 0.37 | -0.27 | 0.50 | 0.109 |
| SART1 | S486 | -0.17 | -0.14 | -0.25 | 0.11 | -0.12 | 0.44 | 0.119 |
| SART1 | T392 | -0.18 | -0.26 | -0.11 | -0.13 | 0.49 | 0.07 | 0.156 |
| SART1 | T392 | -0.17 | -0.08 | -0.27 | -0.01 | 0.32 | 0.13 | 0.044 |
| SART1 | S591; S596; S598 | -0.19 | 0.10 | -0.09 | 0.17 | -0.08 | 0.06 | 0.391 |
| SART1 | S448 | -0.06 | 0.01 | -0.08 | 0.02 | -0.05 | 0.15 | 0.261 |
| SART1 | S448 | 0.05 | -0.05 | -0.05 | 0.05 | -0.13 | 0.12 | 0.716 |
| SART1 | S265 | 0.09 | 0.15 | -0.26 | -0.13 | -0.09 | 0.19 | 0.974 |
| SART1 | S591; S596; S598 | 0.10 | 0.00 | -0.04 | -0.11 | 0.05 | -0.02 | 0.505 |
| SART1 | S/T | 0.11 | 0.01 | -0.03 | -0.01 | -0.10 | 0.02 | 0.358 |
| SART1 | S591; S596; S598 | -0.01 | 0.09 | 0.01 | 0.07 | -0.08 | -0.09 | 0.374 |
| SART1 | S591; S596; S598 | 0.09 | 0.05 | -0.04 | -0.04 | -0.05 | -0.01 | 0.161 |
| SART1 | S591; S596; S598 | 0.12 | 0.05 | -0.03 | 0.01 | 0.03 | -0.20 | 0.304 |
| SART1 | S596; S598 | 0.06 | 0.10 | 0.04 | -0.11 | -0.07 | -0.02 | 0.014 |
| SART1 | S474; S486 | 0.07 | 0.07 | 0.07 | -0.16 | -0.02 | -0.04 | 0.031 |
| SART1 | S474; S486 | 0.05 | 0.12 | 0.05 | -0.01 | -0.02 | -0.20 | 0.084 |
| SART1 | S621 | 0.12 | 0.03 | 0.09 | -0.09 | -0.07 | -0.10 | 0.003 |
| SART1 | S591; S/T | 0.08 | 0.14 | 0.05 | -0.22 | -0.13 | 0.06 | 0.098 |
| SART1 | S598 | 0.07 | 0.11 | 0.11 | -0.13 | -0.04 | -0.15 | 0.006 |
| SART1 | S448 | 0.13 | 0.08 | 0.13 | -0.10 | -0.10 | -0.16 | 1E-03 |
| SART1 | S474; S486 | -0.01 | 0.20 | 0.20 | -0.11 | -0.12 | -0.20 | 0.024 |
| SART1 | S591; S598 | 0.20 | 0.16 | 0.12 | -0.16 | 0.02 | -0.42 | 0.059 |
| SART1 | S598 | 0.16 | 0.19 | 0.16 | -0.19 | -0.09 | -0.31 | 0.005 |
| SART1 | S591; S/T | 0.12 | 0.22 | 0.20 | -0.31 | -0.01 | -0.33 | 0.021 |
| SASH1 | S90 | -0.16 | -0.21 | -0.24 | 0.13 | 0.11 | 0.28 | 0.003 |
| SASH1 | S821 | -0.53 | 0.08 | 0.12 | -0.31 | 0.16 | 0.31 | 0.586 |
| SASH1 | S614 | 0.09 | -0.03 | -0.10 | -0.10 | 0.01 | 0.11 | 0.822 |
| SASH1 | S/T | -0.25 | 0.10 | 0.17 | -0.25 | 0.31 | -0.19 | 0.828 |
| SASH1 | T/S/Y | 0.51 | -0.08 | -0.36 | -0.05 | -0.01 | -0.17 | 0.716 |
| SASH1 | S101 | 0.05 | -0.03 | 0.20 | -0.21 | 0.14 | -0.20 | 0.283 |
| SASH1 | T/S/Y | 0.12 | 0.11 | 0.06 | -0.03 | -0.03 | -0.26 | 0.066 |
| SASH1 | S837 | 0.34 | 0.20 | -0.21 | 0.19 | -0.61 | -0.13 | 0.366 |
| SBF1 | T/S | -0.34 | 0.26 | -0.16 | -0.04 | 0.20 | 0.00 | 0.516 |
| SCAF1 | S614 | -0.59 | -0.30 | -0.44 | 0.24 | 0.20 | 0.54 | 0.005 |
| SCAF1 | S/T | -0.30 | -0.50 | -0.14 | -0.18 | 0.24 | 0.60 | 0.097 |
| SCAF1 | S949 | -0.22 | -0.24 | -0.18 | 0.16 | 0.16 | 0.24 | 3E-04 |
| SCAF1 | S/T | 0.04 | -0.08 | -0.47 | 0.56 | -0.40 | 0.11 | 0.458 |
| SCAF1 | T989; S992 | -0.11 | -0.25 | -0.05 | 0.45 | 0.19 | -0.38 | 0.428 |
| SCAF1 | S498 | -0.13 | -0.12 | -0.04 | -0.04 | 0.14 | 0.16 | 0.061 |
| SCAF1 | S448; S453 | 0.21 | -0.50 | -0.02 | 0.08 | -0.17 | 0.26 | 0.551 |
| SCAF1 | S725 | -0.01 | -0.15 | -0.09 | 0.04 | -0.08 | 0.25 | 0.218 |
| SCAF1 | S498; S500 | -0.23 | -0.11 | 0.09 | 0.09 | -0.16 | 0.26 | 0.4 |
| SCAF1 | S965 | 0.04 | -0.25 | 0.00 | 0.20 | -0.28 | 0.21 | 0.558 |
| SCAF1 | S448; S453 | -0.22 | 0.04 | -0.01 | -0.13 | 0.10 | 0.18 | 0.386 |
| SCAF1 | S500 | -0.01 | 0.06 | -0.16 | 0.12 | -0.09 | 0.07 | 0.489 |
| SCAF1 | S498; S500 | -0.06 | -0.03 | -0.01 | 0.04 | -0.08 | 0.14 | 0.335 |

| | | | | | | | | |
|---|---|---|---|---|---|---|---|---|
| **SCAF1** | **S719; S** | 0.06 | -0.10 | -0.07 | -0.02 | -0.13 | 0.22 | 0.641 |
| **SCAF1** | **S548** | -0.15 | -0.05 | 0.11 | -0.04 | 0.06 | 0.06 | 0.543 |
| **SCAF1** | **T1001** | -0.02 | -0.03 | -0.02 | 0.09 | -0.05 | 0.02 | 0.326 |
| **SCAF1** | **T849** | 0.02 | -0.11 | 0.10 | -0.10 | -0.11 | 0.17 | 0.885 |
| **SCAF1** | **S874** | 0.20 | 0.00 | -0.17 | 0.05 | -0.12 | 0.01 | 0.8 |
| **SCAF1** | **S498; S500** | 0.09 | 0.06 | -0.11 | -0.02 | -0.13 | 0.09 | 0.723 |
| **SCAF1** | **S389** | -0.27 | 0.12 | 0.16 | -0.38 | 0.04 | 0.22 | 0.869 |
| **SCAF1** | **S389** | -0.27 | 0.12 | 0.16 | -0.38 | 0.04 | 0.22 | 0.869 |
| **SCAF1** | **T976** | 0.06 | 0.03 | -0.02 | -0.02 | -0.08 | 0.02 | 0.203 |
| **SCAF1** | **S734; S738** | 0.16 | 0.01 | -0.04 | -0.02 | -0.11 | -0.01 | 0.254 |
| **SCAF1** | **S874** | 0.12 | 0.03 | -0.01 | -0.06 | -0.14 | 0.05 | 0.237 |
| **SCAF1** | **T989; T/S** | 0.02 | 0.03 | 0.09 | -0.01 | -0.12 | -0.02 | 0.087 |
| **SCAF1** | **S725** | 0.06 | 0.06 | 0.01 | 0.02 | 0.06 | -0.24 | 0.388 |
| **SCAF1** | **S239** | 0.07 | 0.04 | 0.04 | -0.04 | -0.19 | 0.07 | 0.251 |
| **SCAF1** | **S680** | 0.17 | 0.14 | -0.15 | -0.19 | 0.03 | -0.03 | 0.402 |
| **SCAF1** | **S734; S738** | 0.07 | -0.02 | 0.12 | -0.11 | -0.10 | 0.03 | 0.129 |
| **SCAF1** | **T/Y/S** | 0.05 | 0.16 | -0.04 | -0.19 | -0.04 | 0.03 | 0.233 |
| **SCAF1** | **T/S** | 0.02 | -0.03 | 0.18 | -0.33 | -0.02 | 0.12 | 0.411 |
| **SCAF1** | **T989** | 0.10 | 0.08 | 0.02 | -0.10 | -0.15 | 0.04 | 0.085 |
| **SCAF1** | **S612; S614** | 0.21 | 0.04 | -0.02 | -0.20 | 0.05 | -0.11 | 0.172 |
| **SCAF1** | **S548** | 0.17 | -0.02 | 0.11 | -0.10 | -0.16 | -0.03 | 0.049 |
| **SCAF1** | **S612; S614** | 0.19 | 0.16 | -0.02 | -0.21 | 0.01 | -0.19 | 0.068 |
| **SCAF1** | **T989; S992** | 0.24 | 0.08 | 0.02 | -0.11 | 0.04 | -0.33 | 0.126 |
| **SCAF1** | **S564; S569** | 0.26 | 0.08 | 0.02 | -0.09 | -0.44 | 0.09 | 0.196 |
| **SCAF1** | **S734** | 0.18 | 0.15 | 0.08 | -0.14 | -0.07 | -0.26 | 0.009 |
| **SCAF1** | **S719; S** | 0.23 | 0.11 | 0.12 | -0.03 | -0.14 | -0.35 | 0.031 |
| **SCAF1** | **S936** | 0.19 | 0.17 | 0.13 | -0.40 | 0.06 | -0.25 | 0.058 |
| **SCAF1** | **S548** | 0.44 | 0.38 | 0.30 | -0.66 | -0.02 | -1.01 | 0.032 |
| **SCAF1** | **S548** | 0.50 | 0.32 | 0.26 | -0.57 | 0.15 | -1.48 | 0.105 |
| **SCAF11** | **S533** | -0.44 | -0.49 | -0.55 | 0.35 | 0.55 | 0.18 | 0.002 |
| **SCAF11** | **S587** | -0.21 | -0.19 | -0.35 | 0.26 | 0.29 | 0.09 | 0.004 |
| **SCAF11** | **S338; S341; S344** | -0.12 | -0.18 | -0.22 | 0.11 | -0.30 | 0.53 | 0.302 |
| **SCAF11** | **S338; S341; S344** | -0.08 | -0.15 | -0.24 | 0.08 | -0.25 | 0.50 | 0.295 |
| **SCAF11** | **S338** | 0.04 | -0.10 | -0.25 | -0.24 | 0.21 | 0.25 | 0.376 |
| **SCAF11** | **S338** | 0.18 | 0.02 | -0.21 | -0.10 | 0.01 | 0.06 | 0.996 |
| **SCAF11** | **S608; S/T** | -0.11 | 0.05 | 0.10 | -0.06 | 0.10 | -0.10 | 0.752 |
| **SCAF11** | **S/T** | 0.08 | 0.01 | -0.05 | -0.13 | -0.06 | 0.14 | 0.752 |
| **SCAF11** | **S401; S402; T410** | -0.06 | -0.04 | 0.14 | -0.11 | -0.13 | 0.18 | 0.802 |
| **SCAF11** | **S816; S818** | 0.01 | 0.01 | 0.03 | -0.06 | -0.18 | 0.17 | 0.713 |
| **SCAF11** | **S608** | 0.12 | -0.09 | 0.03 | -0.01 | 0.09 | -0.16 | 0.624 |
| **SCAF11** | **S400; S401; T410** | 0.02 | 0.00 | 0.07 | -0.07 | -0.11 | 0.08 | 0.347 |
| **SCAF11** | **S338** | 0.14 | 0.09 | -0.11 | 0.01 | -0.19 | 0.02 | 0.413 |
| **SCAF11** | **S816** | 0.04 | 0.02 | 0.10 | -0.02 | -0.11 | -0.04 | 0.037 |
| **SCAF11** | **S882** | 0.06 | 0.03 | 0.08 | -0.09 | -0.01 | -0.08 | 0.016 |
| **SCAF11** | **S608** | 0.11 | -0.02 | 0.08 | -0.07 | -0.22 | 0.09 | 0.259 |
| **SCAF11** | **S796; S802** | 0.19 | 0.10 | -0.07 | -0.18 | -0.12 | 0.06 | 0.216 |
| **SCAF11** | **S/T** | 0.04 | -0.15 | 0.30 | -0.22 | -0.32 | 0.24 | 0.499 |
| **SCAF11** | **S796; S798** | -0.06 | 0.11 | 0.19 | -0.24 | -0.13 | 0.08 | 0.216 |

| | | | | | | | | |
|---|---|---|---|---|---|---|---|---|
| **SCAF11** | **S** | 0.13 | 0.15 | -0.03 | -0.12 | 0.09 | -0.27 | 0.204 |
| **SCAF11** | **S963** | 0.09 | 0.03 | 0.14 | -0.09 | -0.23 | 0.04 | 0.099 |
| **SCAF11** | **S796; S798** | 0.03 | 0.08 | 0.20 | -0.12 | -0.05 | -0.18 | 0.027 |
| **SCAF11** | **S830; S832** | 0.15 | 0.10 | 0.10 | -0.15 | -0.15 | -0.08 | 7E-04 |
| **SCAF11** | **S771** | 0.04 | 0.15 | 0.17 | -0.30 | -0.07 | -0.03 | 0.056 |
| **SCAF11** | **S796** | 0.24 | 0.03 | 0.07 | -0.34 | 0.17 | -0.26 | 0.207 |
| **SCAF11** | **S816** | 0.09 | 0.24 | 0.07 | -0.11 | -0.23 | -0.11 | 0.013 |
| **SCAF11** | **S771; S776** | 0.15 | 0.10 | 0.19 | -0.19 | -0.25 | -0.06 | 0.007 |
| **SCAF11** | **S776** | 0.23 | 0.14 | 0.16 | -0.19 | -0.13 | -0.29 | 0.002 |
| **SCAF11** | **S405** | 0.18 | 0.12 | 0.23 | -0.28 | -0.20 | -0.14 | 0.002 |
| **SCAF11** | **S902** | 0.32 | 0.19 | 0.13 | -0.35 | -0.06 | -0.39 | 0.016 |
| **SCAF11** | **S608** | 0.20 | 0.23 | 0.29 | -0.34 | -0.04 | -0.52 | 0.019 |
| **SCAF4** | **S/T** | -0.34 | 0.03 | -0.05 | -0.05 | 0.18 | 0.16 | 0.185 |
| **SCAF4** | **T237** | 0.14 | 0.16 | 0.36 | -0.33 | 0.01 | -0.54 | 0.044 |
| **SCAF8** | **S273** | -0.30 | -0.11 | 0.04 | 0.31 | -0.06 | 0.06 | 0.198 |
| **SCAF8** | **S617** | 0.01 | -0.10 | 0.07 | -0.05 | 0.03 | 0.03 | 0.858 |
| **SCAF8** | **S617** | 0.08 | 0.10 | -0.10 | -0.14 | 0.03 | 0.01 | 0.501 |
| **SCAF8** | **S/T** | 0.17 | 0.24 | -0.23 | -0.07 | 0.16 | -0.37 | 0.503 |
| **SCAF8** | **T614** | 0.22 | 0.28 | -0.09 | -0.16 | 0.04 | -0.39 | 0.144 |
| **SCAMP3** | **S76** | 0.04 | -0.02 | 0.53 | 0.08 | -0.59 | -0.28 | 0.162 |
| **SCAP** | **T/S/Y** | 0.05 | -0.36 | 0.25 | 0.27 | 0.07 | -0.44 | 0.956 |
| **SCAP** | **S851** | 0.01 | -0.06 | 0.08 | -0.04 | -0.15 | 0.13 | 0.778 |
| **SCAP** | **S937** | 0.17 | 0.18 | 0.15 | -0.14 | -0.03 | -0.42 | 0.033 |
| **SCD** | **T199** | 0.15 | -0.11 | 0.09 | -0.11 | -0.07 | 0.03 | 0.349 |
| **SCD** | **S198** | 0.24 | 0.00 | 0.12 | -0.16 | -0.12 | -0.13 | 0.023 |
| **SCFD1** | **S303** | 0.25 | 0.29 | 0.25 | -0.30 | -0.05 | -0.67 | 0.03 |
| **SCLT1** | **S681** | 0.15 | 0.07 | 0.64 | 0.34 | -0.85 | -1.13 | 0.159 |
| **SCMH1** | **S570** | 0.21 | 0.37 | 0.09 | -0.23 | -0.12 | -0.47 | 0.02 |
| **SCML2** | **S590** | -0.05 | -0.23 | -0.13 | 0.22 | 0.05 | 0.08 | 0.026 |
| **SCML2** | **S511** | -0.09 | -0.12 | 0.14 | -0.22 | -0.28 | 0.44 | 0.98 |
| **SCML2** | **S511; S/T/Y** | -0.31 | 0.20 | 0.09 | -0.02 | -0.07 | 0.05 | 0.964 |
| **SCML2** | **S499; S511** | 0.12 | 0.03 | -0.06 | -0.01 | -0.16 | 0.06 | 0.469 |
| **SCML2** | **S511** | 0.25 | 0.06 | -0.26 | -0.01 | -0.31 | 0.17 | 0.756 |
| **SCNM1** | **S183** | -0.20 | -0.27 | 0.02 | 0.06 | -0.06 | 0.36 | 0.15 |
| **SCNM1** | **S219** | -0.27 | 0.05 | -0.05 | 0.14 | -0.02 | 0.11 | 0.205 |
| **SCRIB** | **S/T/Y** | -0.57 | -0.36 | -0.53 | 0.33 | 0.07 | 0.63 | 0.009 |
| **SCRIB** | **S1378** | -0.27 | -0.55 | -0.59 | 0.23 | 0.02 | 0.71 | 0.026 |
| **SCRIB** | **S708** | -0.47 | -0.44 | -0.35 | 0.43 | 0.07 | 0.45 | 0.005 |
| **SCRIB** | **S1486** | -0.43 | -0.46 | -0.18 | 0.23 | 0.21 | 0.40 | 0.004 |
| **SCRIB** | **S875** | -0.32 | 0.05 | -0.52 | 0.32 | -0.08 | 0.35 | 0.1 |
| **SCRIB** | **S835** | -0.23 | -0.07 | -0.10 | -0.06 | 0.14 | 0.26 | 0.078 |
| **SCRIB** | **S23** | -0.30 | -0.45 | 0.29 | -0.15 | -0.02 | 0.44 | 0.452 |
| **SCRIB** | **S1348** | -0.02 | -0.17 | -0.13 | 0.21 | -0.04 | 0.11 | 0.079 |
| **SCRIB** | **S504** | -0.33 | -0.03 | 0.04 | -0.02 | 0.12 | 0.16 | 0.207 |
| **SCRIB** | **S1475** | -0.23 | -0.08 | 0.05 | 0.01 | 0.05 | 0.17 | 0.145 |
| **SCRIB** | **S853** | -0.18 | 0.09 | -0.10 | 0.02 | 0.02 | 0.13 | 0.242 |
| **SCRIB** | **S1348** | -0.06 | -0.06 | -0.06 | 0.17 | -0.07 | 0.07 | 0.177 |
| **SCRIB** | **S1140** | -0.13 | 0.00 | 0.00 | 0.03 | 0.11 | -0.02 | 0.224 |

| | | | | | | | | |
|---|---|---|---|---|---|---|---|---|
| SCRIB | S504 | -0.15 | 0.01 | 0.06 | -0.20 | 0.01 | 0.23 | 0.78 |
| SCRIB | S1220 | -0.16 | 0.07 | 0.03 | 0.01 | 0.11 | -0.07 | 0.681 |
| SCRIB | T1545; S1547 | 0.03 | -0.16 | 0.10 | -0.03 | 0.05 | 0.00 | 0.805 |
| SCRIB | S1541; T1545; T1549 | -0.17 | 0.11 | 0.02 | -0.19 | -0.03 | 0.22 | 0.932 |
| SCRIB | T1549; S1566 | 0.02 | -0.23 | 0.18 | -0.03 | -0.18 | 0.18 | 0.991 |
| SCRIB | S1439 | 0.03 | 0.10 | -0.13 | -0.04 | -0.05 | 0.08 | 0.968 |
| SCRIB | S/T | -0.16 | -0.04 | 0.19 | -0.06 | 0.08 | -0.04 | 0.961 |
| SCRIB | S1445 | 0.04 | 0.16 | -0.25 | -0.33 | -0.09 | 0.35 | 0.968 |
| SCRIB | S1437 | 0.12 | 0.07 | -0.16 | -0.20 | 0.05 | 0.09 | 0.843 |
| SCRIB | S1439 | 0.05 | 0.07 | -0.07 | -0.04 | -0.05 | 0.03 | 0.505 |
| SCRIB | S835 | -0.01 | 0.05 | 0.02 | 0.06 | 0.02 | -0.14 | 0.563 |
| SCRIB | S1309 | 0.01 | 0.12 | -0.07 | -0.08 | 0.28 | -0.33 | 0.755 |
| SCRIB | S1566 | -0.07 | 0.07 | 0.12 | 0.02 | 0.06 | -0.23 | 0.43 |
| SCRIB | S835 | -0.03 | 0.11 | 0.13 | -0.07 | 0.01 | -0.18 | 0.114 |
| SCRIB | S/T | -0.13 | 0.34 | 0.07 | -0.20 | 0.15 | -0.33 | 0.323 |
| SCRIB | S | 0.37 | 0.07 | 0.35 | -0.32 | -0.12 | -0.59 | 0.022 |
| SCRN2 | T52 | 0.09 | -0.19 | -0.20 | 0.05 | 0.09 | 0.13 | 0.131 |
| SDAD1 | S459 | -1.47 | -1.29 | -0.71 | 0.54 | 0.66 | 0.66 | 0.002 |
| SDAD1 | S585 | -0.04 | -0.03 | -0.05 | -0.14 | -0.16 | 0.36 | 0.733 |
| SDAD1 | S585 | 0.16 | 0.08 | 0.08 | -0.04 | -0.10 | -0.21 | 0.018 |
| SDC1 | S285 | -0.18 | -0.13 | 0.03 | 0.00 | -0.06 | 0.29 | 0.249 |
| SDC1 | S287 | -0.12 | -0.05 | -0.04 | -0.06 | -0.07 | 0.30 | 0.369 |
| SDC1 | S285; S287 | -0.24 | -0.11 | 0.09 | -0.40 | -0.12 | 0.57 | 0.758 |
| SDC1 | T115; T122 | -0.13 | 0.06 | -0.01 | -0.28 | 0.18 | 0.13 | 0.806 |
| SDC1 | S287 | -0.11 | 0.03 | 0.19 | -0.11 | -0.08 | 0.06 | 0.477 |
| SDC2 | S187 | -0.06 | -0.17 | -0.17 | 0.08 | 0.08 | 0.19 | 0.009 |
| SDC4 | S97 | -0.05 | -0.10 | -0.21 | 0.05 | 0.22 | 0.05 | 0.041 |
| SDE2 | S278 | -0.31 | 0.18 | 0.15 | -0.12 | -0.21 | 0.22 | 0.832 |
| SDE2 | S278 | 0.01 | 0.29 | 0.10 | -0.09 | -0.08 | -0.31 | 0.059 |
| SEC13 | S313 | -0.47 | -0.45 | -0.68 | 0.19 | -0.32 | 0.99 | 0.103 |
| SEC13 | S313 | -0.40 | -0.45 | -0.56 | 0.28 | -0.38 | 0.89 | 0.119 |
| SEC14L1 | S/T/Y | -0.46 | -0.45 | 0.40 | -0.23 | 0.01 | 0.44 | 0.525 |
| SEC16A | S314 | -0.24 | -0.27 | 0.02 | 0.11 | 0.36 | -0.07 | 0.124 |
| SEC16A | S1327 | -0.31 | -0.27 | 0.10 | 0.15 | 0.07 | 0.18 | 0.094 |
| SEC16A | S1069 | -0.07 | 0.08 | -0.35 | 0.16 | -0.07 | 0.19 | 0.239 |
| SEC16A | S1327 | -0.11 | -0.14 | -0.07 | 0.03 | -0.02 | 0.28 | 0.1 |
| SEC16A | T/S | 0.02 | 0.25 | -0.74 | 0.39 | 0.33 | -0.67 | 0.72 |
| SEC16A | T/S/Y | 0.03 | -0.14 | -0.11 | -0.03 | 0.04 | 0.19 | 0.155 |
| SEC16A | S2037 | -0.30 | -0.03 | 0.10 | -0.09 | 0.19 | 0.08 | 0.38 |
| SEC16A | S1207 | -0.04 | -0.17 | 0.00 | -0.02 | -0.09 | 0.28 | 0.381 |
| SEC16A | S2042 | -0.08 | -0.01 | -0.06 | 0.15 | 0.01 | -0.01 | 0.139 |
| SEC16A | S314 | -0.15 | -0.06 | -0.03 | -0.53 | 0.03 | 0.53 | 0.796 |
| SEC16A | S1226 | 0.02 | -0.03 | -0.13 | 0.10 | 0.07 | -0.05 | 0.248 |
| SEC16A | S1350; S1353 | 0.04 | 0.03 | -0.21 | 0.26 | -0.18 | 0.02 | 0.628 |
| SEC16A | S1222 | 0.04 | -0.04 | -0.12 | 0.13 | 0.06 | -0.09 | 0.406 |
| SEC16A | S/T/Y | -0.13 | -0.01 | 0.03 | 0.07 | -0.05 | 0.09 | 0.332 |
| SEC16A | S569 | -0.05 | 0.00 | -0.06 | 0.11 | -0.05 | 0.04 | 0.257 |
| SEC16A | T1440; S1441 | -0.05 | 0.17 | -0.21 | 0.04 | -0.05 | 0.08 | 0.7 |

| | | | | | | | | |
|---|---|---|---|---|---|---|---|---|
| **SEC16A** | **T/S** | 0.00 | 0.17 | -0.27 | 0.21 | 0.07 | -0.25 | 0.827 |
| **SEC16A** | **S589** | -0.04 | 0.39 | -0.48 | 0.25 | 0.19 | -0.57 | 0.996 |
| **SEC16A** | **S1350** | 0.05 | 0.09 | -0.16 | 0.17 | 0.04 | -0.23 | 0.984 |
| **SEC16A** | **S1305** | -0.03 | 0.13 | 0.00 | 0.03 | 0.04 | -0.19 | 0.468 |
| **SEC16A** | **S1069** | 0.04 | 0.17 | -0.04 | 0.01 | -0.12 | -0.10 | 0.156 |
| **SEC16A** | **S1356** | 0.35 | 0.04 | -0.24 | -0.11 | -0.04 | -0.08 | 0.495 |
| **SEC16A** | **S1964** | 0.29 | -0.01 | -0.09 | 0.04 | 0.09 | -0.41 | 0.464 |
| **SEC16A** | **S2083** | 0.11 | 0.09 | 0.15 | -0.04 | 0.00 | -0.38 | 0.099 |
| **SEC16A** | **S2271** | -0.10 | 0.20 | 0.27 | -0.19 | 0.07 | -0.33 | 0.17 |
| **SEC22B** | **S137** | -0.20 | -0.10 | 0.48 | -0.33 | -0.26 | 0.23 | 0.554 |
| **SEC22B** | **S137** | 0.24 | 0.15 | 0.22 | -0.53 | -0.08 | -0.14 | 0.033 |
| **SEC31A** | **S527** | -0.10 | 0.02 | -0.10 | 0.13 | 0.00 | 0.04 | 0.094 |
| **SEC31A** | **S799** | 0.00 | -0.05 | -0.07 | 0.01 | 0.01 | 0.11 | 0.101 |
| **SEC31A** | **S799** | 0.14 | 0.11 | -0.24 | -0.16 | 0.05 | 0.06 | 0.92 |
| **SEC62** | **S/T** | -0.22 | 0.19 | -0.34 | 0.37 | -0.23 | 0.09 | 0.446 |
| **SEC62** | **T155; T158** | -0.05 | -0.07 | 0.11 | 0.12 | -0.23 | 0.09 | 0.961 |
| **SEC62** | **S356** | 0.23 | -0.17 | 0.04 | 0.21 | -0.09 | -0.30 | 0.662 |
| **SEC62** | **S/T** | 0.04 | 0.14 | 0.08 | 0.08 | -0.65 | 0.16 | 0.438 |
| **SEC62** | **S356** | 0.16 | -0.01 | 0.11 | 0.34 | -0.61 | -0.17 | 0.458 |
| **SEC62** | **S335** | 0.07 | 0.13 | 0.29 | -0.16 | -0.20 | -0.20 | 0.006 |
| **SEC62** | **T375** | 0.20 | 0.19 | 0.12 | -0.22 | -0.07 | -0.30 | 0.007 |
| **SEC62** | **T375** | 0.18 | 0.23 | 0.09 | -0.02 | -0.53 | -0.07 | 0.091 |
| **SEC62** | **S307; S313** | 0.24 | 0.18 | 0.11 | -0.20 | -0.22 | -0.19 | 6E-04 |
| **SEC62** | **T158** | 0.07 | 0.03 | 0.42 | -0.09 | -0.39 | -0.16 | 0.067 |
| **SEC62** | **S335** | 0.19 | 0.16 | 0.29 | -0.29 | -0.10 | -0.37 | 0.007 |
| **SEC62** | **S117** | 0.26 | 0.27 | 0.13 | -0.22 | -0.22 | -0.35 | 0.001 |
| **SEC62** | **T158** | 0.24 | 0.21 | 0.23 | -0.23 | -0.19 | -0.41 | 0.002 |
| **SEC62** | **T158** | 0.29 | 0.41 | 0.32 | -0.29 | -0.18 | -0.98 | 0.031 |
| **SEC62** | **T332; S335** | 0.41 | 0.13 | 0.29 | 0.45 | -0.58 | -1.62 | 0.225 |
| **SEH1L** | **S/T** | 0.02 | -0.05 | 0.02 | 0.08 | -0.04 | -0.04 | 0.978 |
| **SEL1L** | **T/S** | -0.11 | -0.15 | 0.01 | 0.05 | 0.35 | -0.23 | 0.467 |
| **SELENON** | **S13** | -0.11 | -0.11 | 0.00 | 0.05 | 0.19 | -0.04 | 0.143 |
| **SELENOS** | **S140** | 0.28 | 0.06 | 0.12 | -0.10 | -0.45 | -0.03 | 0.074 |
| **SELENOS** | **S140** | 0.27 | 0.28 | 0.02 | -0.05 | -0.47 | -0.18 | 0.048 |
| **SENP2** | **S32** | -0.21 | -0.68 | -0.50 | 0.54 | 0.20 | 0.26 | 0.01 |
| **SENP2** | **S/T** | 0.07 | -0.14 | 0.05 | -0.03 | -0.27 | 0.26 | 0.978 |
| **SENP3** | **T/S** | -0.13 | -0.15 | -0.13 | 0.16 | 0.03 | 0.17 | 0.006 |
| **SENP3** | **S212** | 0.01 | -0.02 | -0.06 | 0.11 | -0.03 | -0.02 | 0.422 |
| **SENP3** | **S212** | -0.02 | 0.01 | 0.01 | 0.01 | -0.04 | 0.04 | 0.789 |
| **SENP3** | **S/T/Y** | 0.04 | -0.01 | -0.05 | -0.01 | -0.07 | 0.08 | 0.921 |
| **SENP3** | **S232** | 0.09 | 0.00 | -0.09 | 0.07 | 0.01 | -0.10 | 0.93 |
| **SENP3** | **S232** | 0.01 | 0.15 | -0.08 | 0.15 | -0.19 | -0.08 | 0.596 |
| **SENP3** | **S169; S181** | -0.09 | 0.07 | 0.12 | -0.08 | -0.02 | 0.00 | 0.36 |
| **SENP3** | **S25** | -0.03 | -0.18 | 0.27 | -0.34 | -0.10 | 0.27 | 0.745 |
| **SENP3** | **S169; S181** | 0.07 | 0.09 | 0.16 | -0.05 | -0.12 | -0.18 | 0.01 |
| **SENP3** | **S169** | -0.36 | 0.42 | 0.36 | -0.25 | -0.28 | -0.11 | 0.234 |
| **SENP3** | **S169** | 0.11 | 0.18 | 0.27 | -0.23 | -0.16 | -0.27 | 0.002 |
| **SENP5** | **S363** | -0.03 | 0.03 | -0.15 | 0.20 | -0.24 | 0.13 | 0.614 |

| | | | | | | | | |
|---|---|---|---|---|---|---|---|---|
| SENP5 | S293 | 0.22 | -0.13 | 0.32 | 0.15 | -0.07 | -0.69 | 0.304 |
| SENP6 | S/T | 0.11 | -1.42 | 0.08 | 0.12 | 0.37 | 0.14 | 0.293 |
| SENP6 | S919 | -0.19 | -0.27 | -0.04 | 0.25 | 0.14 | 0.04 | 0.024 |
| SENP6 | S350 | -0.13 | -0.15 | -0.01 | -0.75 | 0.01 | 0.67 | 0.867 |
| SENP6 | S352 | -0.16 | 0.13 | 0.09 | -0.29 | -0.12 | 0.26 | 0.733 |
| SENP7 | S11 | -0.26 | -0.10 | -0.02 | 0.12 | 0.17 | 0.05 | 0.037 |
| SEPTIN2 | S218 | 0.09 | 0.06 | -0.02 | 0.02 | -0.14 | -0.01 | 0.239 |
| SEPTIN2 | S218 | 0.10 | 0.09 | -0.04 | -0.01 | -0.19 | 0.04 | 0.279 |
| SEPTIN2 | S218 | 0.50 | -0.02 | -0.07 | -0.55 | -0.59 | 0.38 | 0.352 |
| SEPTIN9 | S30 | 0.06 | -0.13 | -0.15 | 0.17 | -0.06 | 0.08 | 0.223 |
| SEPTIN9 | S30 | 0.15 | 0.00 | -0.11 | -0.03 | -0.06 | 0.02 | 0.655 |
| SERBP1 | S252 | -0.26 | -0.43 | -0.41 | 0.32 | 0.14 | 0.40 | 0.002 |
| SERBP1 | S/T | 0.20 | -0.60 | -0.37 | 0.04 | -0.24 | 0.63 | 0.313 |
| SERBP1 | S/T | 0.01 | -0.08 | -0.06 | -0.08 | -0.02 | 0.21 | 0.416 |
| SERBP1 | S247 | -0.03 | -0.25 | 0.45 | 0.22 | -0.16 | -0.41 | 0.563 |
| SERBP1 | S | 0.23 | 0.28 | -0.20 | -0.10 | 0.01 | -0.31 | 0.257 |
| SERBP1 | S234 | 0.09 | 0.07 | 0.18 | -0.01 | -0.10 | -0.28 | 0.05 |
| SERINC1 | T352 | -0.06 | 0.01 | 0.12 | -0.02 | 0.08 | -0.15 | 0.552 |
| SET | S7 | -0.02 | 0.04 | 0.04 | -0.10 | 0.04 | -0.01 | 0.401 |
| SETBP1 | S893 | 0.09 | 0.10 | -0.06 | 0.29 | -0.70 | 0.10 | 0.661 |
| SETBP1 | S1567 | 0.23 | -0.05 | 0.08 | -0.43 | 0.00 | 0.08 | 0.32 |
| SETBP1 | S1567 | 0.27 | 0.28 | -0.13 | -0.11 | 0.03 | -0.47 | 0.183 |
| SETD1A | S470 | -0.16 | 0.18 | 0.21 | -0.21 | -0.11 | 0.04 | 0.281 |
| SETD1B | S1283 | -0.31 | 0.03 | 0.11 | -0.06 | 0.04 | 0.14 | 0.53 |
| SETD1B | S1298 | 0.00 | 0.02 | -0.12 | 0.00 | -0.01 | 0.10 | 0.356 |
| SETD2 | S249 | -2.41 | -1.11 | -2.40 | 1.03 | 0.32 | 0.91 | 0.005 |
| SETD2 | S321; S323 | -0.12 | -0.26 | -0.22 | 0.13 | -0.24 | 0.53 | 0.203 |
| SETD2 | S458; S459; S461 | -0.05 | -0.10 | -0.11 | 0.03 | -0.19 | 0.35 | 0.396 |
| SETD2 | T1853 | 0.00 | -0.08 | -0.15 | 0.21 | -0.08 | 0.06 | 0.209 |
| SETD2 | S744; S754 | 0.07 | -0.01 | -0.06 | -0.07 | 0.16 | -0.10 | 0.955 |
| SETD2 | S131 | 0.05 | 0.04 | -0.03 | -0.15 | -0.03 | 0.10 | 0.564 |
| SETD2 | S459; S461 | 0.10 | 0.07 | -0.10 | -0.02 | 0.00 | -0.05 | 0.483 |
| SETD2 | T1872 | -0.01 | 0.01 | 0.10 | -0.07 | -0.04 | 0.01 | 0.206 |
| SETD2 | S2080; S2082 | 0.05 | -0.04 | 0.07 | 0.03 | -0.27 | 0.12 | 0.616 |
| SETD2 | S614 | 0.14 | -0.08 | 0.08 | 0.03 | -0.09 | -0.11 | 0.251 |
| SETD2 | S754 | 0.13 | 0.13 | -0.07 | -0.04 | 0.03 | -0.20 | 0.226 |
| SETD2 | S131 | 0.08 | 0.12 | 0.05 | -0.12 | 0.03 | -0.19 | 0.062 |
| SETD2 | S323 | 0.02 | 0.19 | 0.20 | -0.19 | -0.06 | -0.22 | 0.019 |
| SETD2 | S754; S/T | 0.09 | 0.27 | 0.10 | 0.07 | -0.22 | -0.43 | 0.086 |
| SETD2 | S532 | 0.18 | 0.02 | 0.27 | -0.31 | 0.16 | -0.46 | 0.146 |
| SETD2 | T1872 | 0.21 | 0.25 | 0.08 | -0.01 | -0.15 | -0.51 | 0.066 |
| SETD2 | S744 | 0.09 | 0.22 | 0.25 | -0.04 | -0.27 | -0.35 | 0.018 |
| SETD2 | S624 | 0.25 | 0.15 | 0.21 | -0.24 | -0.08 | -0.41 | 0.011 |
| SETD2 | S744 | 0.30 | 0.21 | 0.24 | -0.10 | -0.23 | -0.65 | 0.027 |
| SETD2 | S/T/Y | 0.54 | 0.14 | 0.28 | -0.38 | -0.48 | -0.44 | 0.003 |
| SETD2 | S461 | 0.36 | 0.34 | 0.26 | -0.38 | -0.16 | -0.77 | 0.014 |
| SETD5 | S865 | -0.10 | 0.06 | -0.63 | 0.13 | 0.27 | 0.11 | 0.142 |
| SETD5 | S1198 | -0.04 | 0.00 | -0.22 | 0.21 | -0.06 | 0.07 | 0.196 |

| | | | | | | | | |
|---|---|---|---|---|---|---|---|---|
| **SETD5** | **S541** | 0.05 | -0.16 | 0.14 | -0.06 | 0.00 | 0.01 | 0.829 |
| **SETD5** | **S541** | 0.11 | -0.10 | 0.03 | -0.11 | 0.02 | 0.03 | 0.675 |
| **SETD5** | **S191** | 0.04 | 0.01 | 0.21 | -0.38 | -0.11 | 0.15 | 0.287 |
| **SETD5** | **S191** | 0.15 | -0.17 | 0.26 | 0.27 | 0.01 | -0.77 | 0.512 |
| **SETDB1** | **S1066** | -0.06 | -0.35 | -0.22 | 0.14 | 0.04 | 0.34 | 0.035 |
| **SETDB1** | **T361** | 0.09 | 0.02 | -0.02 | 0.05 | 0.26 | -0.50 | 0.695 |
| **SETMAR** | **S508** | 0.07 | -0.05 | -0.14 | 0.12 | -0.05 | 0.02 | 0.432 |
| **SETX** | **S1017; S1019** | -0.24 | -0.17 | -0.13 | 0.22 | -0.10 | 0.32 | 0.068 |
| **SETX** | **S1366** | 0.01 | 0.00 | -0.02 | 0.13 | -0.12 | -0.01 | 0.92 |
| **SETX** | **S947** | 0.04 | 0.00 | -0.01 | 0.10 | -0.07 | -0.08 | 0.68 |
| **SF1** | **S82** | -0.31 | -0.35 | -0.10 | 0.23 | 0.37 | 0.01 | 0.025 |
| **SF1** | **S80; S82** | -0.14 | -0.11 | 0.00 | 0.28 | -0.24 | 0.13 | 0.429 |
| **SF1** | **S80; S82** | 0.02 | 0.14 | 0.15 | -0.11 | -0.02 | -0.21 | 0.033 |
| **SF1** | **S80; S82** | 0.06 | 0.21 | 0.21 | -0.17 | -0.04 | -0.37 | 0.031 |
| **SF3A1** | **S320; S329** | -1.19 | -0.98 | -0.84 | 0.47 | 0.96 | 0.21 | 0.003 |
| **SF3A1** | **S359** | -0.07 | -0.19 | -0.37 | 0.16 | -0.12 | 0.45 | 0.115 |
| **SF3A1** | **S359** | -0.53 | -0.20 | 0.10 | 0.17 | -0.17 | 0.43 | 0.237 |
| **SF3A1** | **S359** | -0.04 | -0.12 | -0.29 | 0.13 | -0.01 | 0.27 | 0.061 |
| **SF3A1** | **S344** | -0.02 | -0.28 | -0.05 | -0.04 | 0.21 | 0.12 | 0.123 |
| **SF3A1** | **S359** | 0.07 | -0.11 | -0.30 | 0.05 | 0.01 | 0.22 | 0.172 |
| **SF3A1** | **S451** | 0.00 | 0.00 | -0.03 | 0.06 | -0.20 | 0.15 | 0.919 |
| **SF3A1** | **S451** | -0.15 | -0.22 | 0.28 | -0.16 | -0.45 | 0.48 | 0.973 |
| **SF3A1** | **S329** | 0.00 | 0.01 | 0.12 | -0.04 | 0.03 | -0.13 | 0.198 |
| **SF3A1** | **S329** | 0.11 | -0.01 | 0.28 | -0.10 | 0.02 | -0.38 | 0.127 |
| **SF3A1** | **S329** | 0.18 | 0.14 | 0.08 | -0.21 | -0.06 | -0.17 | 0.006 |
| **SF3A1** | **S329** | 0.16 | 0.21 | 0.12 | -0.14 | -0.17 | -0.24 | 1E-03 |
| **SF3A1** | **S329** | 0.28 | 0.14 | 0.24 | -0.30 | -0.02 | -0.51 | 0.03 |
| **SF3A3** | **S295** | -1.68 | -1.15 | -1.34 | 0.28 | 0.45 | 1.17 | 0.003 |
| **SF3A3** | **S365; S367; S369** | 0.22 | -0.02 | 0.07 | -0.11 | -0.13 | -0.06 | 0.063 |
| **SF3A3** | **S367** | 0.28 | 0.05 | 0.01 | -0.33 | -0.36 | 0.23 | 0.272 |
| **SF3B1** | **T434; T436** | -0.28 | 0.13 | -0.41 | 0.00 | 0.16 | 0.28 | 0.139 |
| **SF3B1** | **T326; T328; S332** | -0.14 | -0.18 | -0.17 | -0.02 | -0.01 | 0.43 | 0.115 |
| **SF3B1** | **S488** | 0.18 | 0.04 | -0.77 | 0.19 | -0.08 | 0.21 | 0.404 |
| **SF3B1** | **S488** | -0.05 | -0.12 | -0.23 | 0.17 | -0.03 | 0.22 | 0.056 |
| **SF3B1** | **T244; T248** | 0.04 | -0.24 | -0.20 | 0.15 | -0.11 | 0.29 | 0.172 |
| **SF3B1** | **S194** | -0.12 | -0.12 | 0.00 | -0.04 | -0.07 | 0.31 | 0.317 |
| **SF3B1** | **T248** | 0.01 | -0.18 | -0.01 | 0.06 | -0.11 | 0.20 | 0.383 |
| **SF3B1** | **T211** | 0.01 | -0.04 | -0.10 | -0.02 | 0.02 | 0.13 | 0.193 |
| **SF3B1** | **T434; T436** | 0.02 | -0.04 | -0.11 | 0.06 | -0.10 | 0.15 | 0.364 |
| **SF3B1** | **T257; T261** | 0.04 | -0.10 | -0.05 | -0.05 | 0.09 | 0.07 | 0.259 |
| **SF3B1** | **T313** | 0.07 | -0.04 | -0.16 | 0.14 | 0.14 | -0.20 | 0.631 |
| **SF3B1** | **T211** | 0.00 | -0.17 | 0.10 | -0.03 | 0.19 | -0.12 | 0.794 |
| **SF3B1** | **S/T** | -0.33 | 0.05 | 0.26 | -0.04 | -0.05 | 0.05 | 0.97 |
| **SF3B1** | **T426** | 0.10 | 0.03 | -0.12 | -0.08 | -0.08 | 0.13 | 0.905 |
| **SF3B1** | **T326; T328** | 0.13 | -0.05 | -0.06 | -0.08 | -0.04 | 0.09 | 0.843 |
| **SF3B1** | **T142** | 0.12 | -0.01 | -0.08 | -0.02 | -0.06 | 0.04 | 0.686 |
| **SF3B1** | **T326** | 0.11 | -0.03 | -0.02 | 0.02 | -0.08 | 0.00 | 0.548 |
| **SF3B1** | **S488** | 0.05 | 0.07 | -0.09 | -0.28 | 0.20 | -0.01 | 0.8 |

| | | | | | | | | |
|---|---|---|---|---|---|---|---|---|
| **SF3B1** | **S194** | -0.33 | 0.12 | 0.28 | -0.14 | -0.10 | 0.09 | 0.721 |
| **SF3B1** | **S/T** | -0.08 | 0.04 | 0.15 | 0.17 | -0.06 | -0.27 | 0.566 |
| **SF3B1** | **T211** | 0.35 | 0.09 | -0.34 | -0.15 | 0.14 | -0.19 | 0.678 |
| **SF3B1** | **T211** | 0.06 | -0.07 | 0.20 | -0.13 | 0.06 | -0.15 | 0.255 |
| **SF3B1** | **T223; T227** | 0.04 | -0.05 | 0.21 | -0.03 | 0.01 | -0.21 | 0.224 |
| **SF3B1** | **T223** | 0.12 | 0.12 | 0.04 | -0.13 | -0.07 | -0.11 | 0.003 |
| **SF3B1** | **T261** | -0.02 | 0.02 | 0.29 | -0.17 | 0.04 | -0.22 | 0.167 |
| **SF3B1** | **T211** | 0.14 | 0.02 | 0.15 | -0.18 | -0.02 | -0.14 | 0.027 |
| **SF3B1** | **S129** | 0.23 | 0.09 | 0.02 | -0.20 | 0.15 | -0.38 | 0.204 |
| **SF3B1** | **T350** | 0.21 | 0.12 | 0.02 | -0.05 | 0.03 | -0.41 | 0.148 |
| **SF3B1** | **S129** | 0.15 | 0.01 | 0.21 | -0.12 | 0.03 | -0.35 | 0.1 |
| **SF3B1** | **T207; T211** | -0.02 | 0.08 | 0.31 | -0.12 | -0.32 | -0.01 | 0.116 |
| **SF3B1** | **T296** | 0.22 | 0.03 | 0.14 | -0.12 | -0.01 | -0.33 | 0.062 |
| **SF3B1** | **T227** | 0.30 | 0.16 | -0.06 | -0.28 | -0.50 | 0.21 | 0.234 |
| **SF3B1** | **T303** | 0.44 | -0.18 | 0.23 | -0.28 | -0.06 | -0.30 | 0.129 |
| **SF3B1** | **T278; T/S** | -0.04 | 0.69 | 0.14 | -0.21 | -0.39 | -0.55 | 0.056 |
| **SF3B2** | **S/T** | -0.16 | -0.48 | -0.41 | 0.35 | 0.57 | -0.19 | 0.074 |
| **SF3B2** | **S431; S435; S436** | -0.36 | -0.26 | -0.36 | 0.35 | -0.36 | 0.63 | 0.144 |
| **SF3B2** | **S431; S435; S436** | -0.18 | -0.29 | -0.25 | 0.12 | -0.27 | 0.63 | 0.205 |
| **SF3B2** | **T780** | -0.06 | -0.20 | -0.32 | 0.17 | 0.21 | 0.12 | 0.011 |
| **SF3B2** | **T780** | 0.02 | -0.11 | -0.16 | 0.21 | -0.06 | 0.06 | 0.17 |
| **SF3B2** | **S347** | -0.01 | -0.04 | -0.10 | 0.08 | -0.02 | 0.07 | 0.078 |
| **SF3B2** | **S317** | -0.03 | 0.10 | -0.18 | -0.01 | 0.16 | -0.07 | 0.58 |
| **SF3B2** | **S302; S307; S309** | -0.03 | 0.02 | -0.03 | 0.03 | 0.08 | -0.08 | 0.652 |
| **SF3B2** | **S362** | 0.25 | 0.07 | -0.23 | -0.15 | -0.05 | 0.06 | 0.621 |
| **SF3B2** | **S307; S309** | 0.04 | 0.02 | 0.06 | -0.05 | -0.07 | -0.01 | 0.014 |
| **SF3B2** | **S317** | 0.03 | 0.18 | 0.05 | -0.12 | 0.17 | -0.38 | 0.308 |
| **SF3B2** | **S302** | 0.09 | 0.09 | 0.15 | -0.10 | -0.13 | -0.14 | 5E-04 |
| **SF3B2** | **T780** | 0.17 | 0.10 | 0.05 | -0.15 | 0.15 | -0.40 | 0.212 |
| **SF3B2** | **T780** | 0.18 | 0.04 | 0.16 | -0.15 | -0.01 | -0.26 | 0.035 |
| **SF3B2** | **S861** | 0.29 | 0.30 | -0.22 | -0.14 | 0.05 | -0.41 | 0.258 |
| **SF3B2** | **S435; S436** | 0.30 | 0.08 | 0.17 | -0.03 | -0.11 | -0.54 | 0.075 |
| **SF3B2** | **S309** | 0.32 | 0.19 | 0.29 | -0.19 | -0.12 | -0.78 | 0.041 |
| **SF3B3** | **T1200** | 0.23 | 0.16 | 0.24 | -0.18 | -0.01 | -0.59 | 0.054 |
| **SFPQ** | **S33** | -0.71 | 0.13 | -0.38 | 0.20 | 0.17 | 0.33 | 0.09 |
| **SFPQ** | **S33** | -0.40 | -0.11 | -0.38 | 0.38 | 0.23 | 0.11 | 0.012 |
| **SFPQ** | **S273** | 0.21 | 0.05 | 0.06 | -0.16 | -0.07 | -0.13 | 0.018 |
| **SFPQ** | **S273** | 0.11 | 0.27 | -0.01 | -0.13 | 0.02 | -0.34 | 0.109 |
| **SFSWAP** | **T642** | -1.34 | -1.21 | -1.56 | 0.53 | 0.77 | 0.75 | 9E-05 |
| **SFSWAP** | **S/T** | 0.12 | -0.07 | -0.24 | 0.10 | -0.17 | 0.21 | 0.527 |
| **SFSWAP** | **S/Y** | 0.22 | -0.06 | -0.19 | 0.02 | -0.22 | 0.18 | 0.984 |
| **SFSWAP** | **S/Y** | 0.07 | -0.03 | -0.04 | 0.07 | -0.10 | 0.03 | 0.995 |
| **SFSWAP** | **S604** | 0.04 | 0.16 | -0.12 | -0.33 | -0.05 | 0.23 | 0.69 |
| **SFSWAP** | **S604** | 0.11 | 0.04 | 0.02 | -0.10 | -0.08 | 0.00 | 0.043 |
| **SFSWAP** | **S870** | 0.19 | 0.12 | 0.26 | -0.21 | 0.07 | -0.61 | 0.094 |
| **SFSWAP** | **S866; S870** | 0.25 | 0.19 | 0.19 | -0.20 | -0.24 | -0.31 | 2E-04 |
| **SGMS1** | **S8** | -0.10 | -0.25 | -0.23 | 0.04 | 0.16 | 0.29 | 0.014 |
| **SGO1** | **S256** | -0.09 | -0.22 | 0.12 | -0.09 | -0.04 | 0.27 | 0.516 |

| | | | | | | | | |
|---|---|---|---|---|---|---|---|---|
| SGPP1 | S112 | 0.20 | 0.23 | -0.12 | -0.39 | -0.13 | 0.12 | 0.273 |
| SGTA | S77 | 0.04 | -0.14 | -0.18 | 0.17 | -0.03 | 0.11 | 0.121 |
| SH2D4A | S315 | 0.14 | -0.03 | -0.35 | 0.17 | 0.16 | -0.17 | 0.512 |
| SH3BP1 | S/T | -0.26 | -0.05 | -0.17 | -0.05 | -0.36 | 0.65 | 0.472 |
| SH3BP1 | S262 | -0.05 | 0.18 | 0.12 | 0.05 | -0.14 | -0.19 | 0.154 |
| SH3BP5L | S362 | 0.20 | -0.20 | 0.20 | -0.47 | -0.29 | 0.37 | 0.526 |
| SH3BP5L | S343 | -0.07 | 0.17 | 0.46 | -0.43 | 0.00 | -0.33 | 0.092 |
| SH3D21 | S/T | -0.18 | 0.28 | -0.23 | -0.20 | -0.10 | 0.32 | 0.841 |
| SH3KBP1 | S230 | 0.19 | -0.21 | -0.11 | -0.04 | 0.33 | -0.26 | 0.813 |
| SHANK2 | S1104 | -0.23 | 0.10 | -0.26 | -0.02 | 0.20 | 0.15 | 0.148 |
| SHANK2 | S960 | -0.10 | -0.07 | -0.12 | -0.01 | -0.12 | 0.36 | 0.302 |
| SHANK2 | S475 | -0.12 | -0.05 | 0.02 | 0.19 | 0.10 | -0.18 | 0.511 |
| SHANK2 | S1448 | 0.26 | 0.36 | 0.08 | -0.08 | -0.59 | -0.22 | 0.036 |
| SHB | S310 | 0.17 | -0.07 | -0.01 | -0.29 | -0.24 | 0.34 | 0.692 |
| SHB | S310 | 0.07 | 0.07 | 0.07 | -0.37 | -0.17 | 0.25 | 0.423 |
| SHC1 | S426 | 0.00 | -0.29 | -0.51 | 0.58 | 0.01 | -0.02 | 0.137 |
| SHROOM2 | S413 | -0.34 | -0.39 | -0.60 | 0.49 | 0.33 | 0.17 | 0.003 |
| SHROOM2 | S1173 | -0.25 | -0.29 | -0.12 | 0.14 | 0.18 | 0.25 | 0.003 |
| SHROOM2 | S | -0.16 | 0.00 | 0.14 | 0.16 | -0.16 | 0.00 | 0.956 |
| SHROOM2 | S456 | -0.41 | 0.09 | 0.25 | 0.19 | 0.06 | -0.30 | 0.978 |
| SHROOM2 | S313 | -0.09 | 0.15 | 0.00 | 0.18 | -0.17 | -0.12 | 0.674 |
| SHROOM2 | S974 | -0.11 | 0.18 | 0.06 | 0.00 | -0.16 | 0.01 | 0.421 |
| SHROOM2 | S779 | 0.09 | 0.08 | -0.01 | 0.01 | 0.02 | -0.21 | 0.226 |
| SHROOM2 | S1039 | 0.11 | 0.10 | -0.03 | 0.03 | -0.09 | -0.14 | 0.119 |
| SHROOM3 | S1497 | 0.01 | -0.08 | -0.16 | -0.13 | 0.16 | 0.16 | 0.259 |
| SHROOM3 | S1441 | -0.10 | 0.00 | -0.08 | 0.01 | 0.02 | 0.15 | 0.088 |
| SHROOM3 | S910 | -0.15 | 0.03 | 0.04 | 0.06 | -0.02 | 0.02 | 0.494 |
| SHROOM3 | S443 | -0.08 | 0.02 | 0.06 | 0.16 | -0.10 | -0.09 | 0.939 |
| SHTN1 | S/T | 0.14 | 0.07 | -0.10 | -0.33 | 0.08 | 0.09 | 0.599 |
| SHTN1 | S494 | 0.18 | 0.13 | -0.06 | 0.00 | -0.03 | -0.25 | 0.184 |
| SHTN1 | S534 | 0.20 | 0.07 | 0.12 | -0.24 | -0.36 | 0.12 | 0.119 |
| SHTN1 | S/T | 0.23 | 0.38 | 0.37 | -0.41 | -0.33 | -0.54 | 7E-04 |
| SIDT2 | S404 | -0.03 | 0.06 | 0.15 | 0.05 | -0.07 | -0.19 | 0.196 |
| SIN3A | S832 | -0.03 | -0.09 | -0.15 | 0.16 | -0.01 | 0.10 | 0.044 |
| SIN3A | S832 | 0.03 | -0.02 | -0.15 | 0.12 | -0.03 | 0.04 | 0.259 |
| SIN3A | T434 | 0.05 | -0.03 | -0.14 | 0.15 | 0.06 | -0.10 | 0.46 |
| SIN3A | S860 | 0.03 | -0.03 | 0.07 | 0.07 | -0.09 | -0.07 | 0.42 |
| SIN3A | S860 | 0.06 | 0.04 | 0.14 | -0.14 | -0.06 | -0.07 | 0.013 |
| SIN3A | S1112 | 0.14 | 0.12 | 0.12 | -0.13 | 0.03 | -0.34 | 0.058 |
| SIN3A | S1112 | 0.07 | 0.09 | 0.31 | -0.14 | 0.05 | -0.52 | 0.124 |
| SIN3A | T435 | 0.09 | 0.26 | 0.36 | -0.41 | -0.06 | -0.44 | 0.021 |
| SIN3A | S860 | 0.35 | 0.27 | 0.26 | -0.32 | -0.24 | -0.56 | 0.003 |
| SIN3B | S122 | 0.26 | -0.36 | -0.43 | 0.35 | -0.27 | 0.24 | 0.382 |
| SIN3B | S740 | 0.16 | 0.14 | -0.39 | 0.13 | 0.00 | -0.12 | 0.86 |
| SIPA1L1 | S208 | -0.05 | -0.09 | -0.15 | 0.10 | 0.02 | 0.15 | 0.017 |
| SIPA1L1 | S114 | -0.20 | 0.01 | -0.07 | 0.07 | 0.08 | 0.09 | 0.046 |
| SIPA1L1 | S255 | 0.06 | -0.08 | -0.24 | 0.06 | -0.10 | 0.24 | 0.302 |
| SIPA1L1 | S1647; S/T | -0.22 | -0.03 | 0.01 | 0.02 | -0.06 | 0.24 | 0.272 |

| | | | | | | | | |
|---|---|---|---|---|---|---|---|---|
| SIPA1L1 | S | -0.02 | -0.11 | -0.08 | 0.11 | -0.11 | 0.19 | 0.229 |
| SIPA1L1 | S1149 | 0.02 | -0.09 | -0.11 | -0.03 | 0.01 | 0.17 | 0.218 |
| SIPA1L1 | S1528 | 0.06 | 0.08 | -0.24 | 0.24 | -0.38 | 0.15 | 0.871 |
| SIPA1L1 | S1734 | 0.01 | -0.03 | 0.02 | 0.05 | 0.00 | -0.05 | 0.989 |
| SIPA1L1 | S97 | 0.07 | 0.05 | -0.05 | 0.02 | 0.18 | -0.31 | 0.687 |
| SIPA1L1 | S114 | 0.05 | 0.04 | 0.02 | 0.00 | 0.02 | -0.15 | 0.238 |
| SIPA1L1 | S1585 | 0.07 | 0.11 | 0.08 | -0.02 | -0.02 | -0.24 | 0.07 |
| SIPA1L1 | S1270 | -0.03 | 0.23 | 0.14 | -0.13 | -0.07 | -0.19 | 0.044 |
| SIPA1L1 | S1549 | 0.05 | 0.24 | 0.15 | -0.10 | -0.07 | -0.34 | 0.035 |
| SIPA1L1 | S1433 | 0.03 | 0.22 | 0.42 | -0.19 | -0.18 | -0.47 | 0.027 |
| SIPA1L1 | S1255 | 0.25 | 0.03 | 0.29 | 0.32 | -0.15 | -1.27 | 0.309 |
| SIPA1L1 | S | 0.18 | 0.29 | 0.28 | -0.22 | 0.00 | -0.81 | 0.072 |
| SIPA1L2 | S1461 | -0.20 | -0.16 | 0.07 | 0.24 | 0.09 | -0.09 | 0.232 |
| SIPA1L2 | S1029 | -0.08 | 0.07 | 0.15 | 0.05 | -0.08 | -0.13 | 0.308 |
| SIPA1L3 | S100 | -0.14 | -0.23 | -0.18 | -0.01 | -0.05 | 0.48 | 0.132 |
| SIPA1L3 | T1703 | -0.19 | -0.11 | -0.16 | 0.05 | 0.11 | 0.24 | 0.008 |
| SIPA1L3 | S172 | -0.24 | -0.04 | -0.01 | 0.10 | -0.10 | 0.24 | 0.218 |
| SIPA1L3 | S146 | -0.01 | -0.03 | -0.15 | 0.16 | -0.03 | 0.05 | 0.159 |
| SIPA1L3 | S1544; S1547 | 0.05 | -0.09 | 0.05 | -0.07 | -0.15 | 0.19 | 0.889 |
| SIPA1L3 | S1364 | -0.04 | 0.10 | -0.01 | 0.05 | -0.11 | 0.01 | 0.605 |
| SIPA1L3 | S401 | 0.07 | 0.05 | -0.05 | -0.02 | 0.15 | -0.23 | 0.647 |
| SIPA1L3 | S1364 | 0.06 | 0.15 | -0.10 | -0.13 | 0.09 | -0.10 | 0.46 |
| SIPA1L3 | S1544 | 0.07 | 0.17 | 0.16 | -0.17 | -0.06 | -0.21 | 0.007 |
| SIPA1L3 | T1703 | -0.11 | 0.24 | 0.31 | 0.03 | -0.21 | -0.39 | 0.138 |
| SIPA1L3 | S1254 | 0.34 | 0.30 | 0.23 | -0.62 | -0.03 | -0.50 | 0.021 |
| SIRT6 | T294 | -0.02 | -0.14 | -0.14 | 0.13 | -0.08 | 0.22 | 0.123 |
| SIRT6 | T337 | 0.03 | 0.01 | -0.09 | -0.06 | 0.11 | -0.01 | 0.647 |
| SIRT6 | S330 | 0.08 | 0.11 | 0.01 | 0.13 | -0.03 | -0.35 | 0.359 |
| SIRT6 | S330 | -0.03 | 0.03 | 0.27 | -0.08 | -0.05 | -0.18 | 0.119 |
| SIRT6 | S303 | 0.16 | 0.13 | 0.09 | -0.02 | 0.01 | -0.45 | 0.136 |
| SIRT6 | S330 | 0.08 | 0.01 | 0.35 | -0.20 | -0.03 | -0.31 | 0.069 |
| SIX2 | S212 | 0.12 | 0.08 | 0.28 | -0.15 | -0.19 | -0.21 | 0.006 |
| SIX4 | S44 | -0.01 | -0.13 | -0.03 | -0.05 | 0.00 | 0.19 | 0.278 |
| SKA2 | S101 | 0.00 | 0.04 | -0.38 | 0.23 | 0.10 | -0.07 | 0.267 |
| SKA3 | S155 | 0.02 | 0.03 | -0.37 | 0.27 | 0.11 | -0.14 | 0.346 |
| SKIC2 | S256 | 0.11 | 0.12 | -0.33 | 0.06 | 0.17 | -0.20 | 0.809 |
| SKIC2 | S245 | -0.11 | 0.15 | 0.12 | -0.11 | -0.08 | 0.00 | 0.254 |
| SKP1 | T131 | 0.14 | 0.07 | 0.07 | 0.15 | -0.09 | -0.40 | 0.268 |
| SLBP | S182 | -0.23 | 0.04 | 0.00 | -0.07 | 0.15 | 0.08 | 0.319 |
| SLC12A2 | S940 | 0.00 | 0.02 | -0.01 | -0.03 | 0.05 | -0.03 | 0.767 |
| SLC12A2 | T266 | 0.02 | 0.07 | 0.28 | -0.20 | -0.02 | -0.22 | 0.06 |
| SLC12A2 | S | 0.14 | -0.03 | 0.42 | 0.19 | -0.61 | -0.36 | 0.184 |
| SLC12A2 | S994 | 0.13 | 0.44 | 0.36 | 0.19 | -0.18 | -1.99 | 0.228 |
| SLC12A4 | S967 | 0.05 | 0.04 | 0.11 | -0.11 | -0.01 | -0.09 | 0.025 |
| SLC12A4 | S47 | 0.06 | 0.04 | 0.27 | -0.11 | -0.13 | -0.18 | 0.027 |
| SLC12A6 | S29 | 0.07 | 0.30 | 0.09 | 0.33 | -0.14 | -1.03 | 0.348 |
| SLC12A7 | T30; S40 | -0.23 | -0.15 | 0.45 | -0.30 | -0.05 | 0.14 | 0.734 |
| SLC12A7 | T30 | -0.11 | 0.07 | 0.21 | 0.18 | -0.26 | -0.15 | 0.451 |

| | | | | | | | | |
|---|---|---|---|---|---|---|---|---|
| **SLC12A7** | **S997** | 0.18 | 0.23 | 0.15 | -0.40 | -0.10 | -0.17 | 0.013 |
| **SLC12A7** | **T973** | 0.09 | 0.18 | 0.42 | -0.32 | -0.25 | -0.28 | 0.006 |
| **SLC16A1** | **S461** | -0.21 | -0.08 | 0.41 | 0.29 | -0.46 | -0.12 | 0.66 |
| **SLC16A1** | **S467** | 0.05 | 0.14 | 0.06 | -0.18 | -0.11 | 0.01 | 0.051 |
| **SLC16A1** | **S467** | 0.01 | 0.27 | 0.15 | -0.32 | -0.01 | -0.18 | 0.054 |
| **SLC16A2** | **S512** | 0.10 | 0.39 | 0.25 | -0.17 | -0.26 | -0.51 | 0.012 |
| **SLC16A6** | **S237** | -0.20 | 0.01 | -0.03 | 0.20 | 0.08 | -0.10 | 0.291 |
| **SLC16A6** | **S273** | 0.12 | 0.09 | 0.02 | -0.37 | 0.15 | -0.08 | 0.31 |
| **SLC16A6** | **T/S** | -0.01 | 0.24 | 0.17 | -0.51 | 0.24 | -0.29 | 0.239 |
| **SLC1A4** | **S507** | -0.33 | 0.41 | 0.13 | -0.09 | -0.09 | -0.14 | 0.454 |
| **SLC1A4** | **S507** | 0.04 | 0.12 | 0.29 | -0.39 | -0.10 | -0.06 | 0.06 |
| **SLC1A5** | **S503** | -0.19 | 0.05 | 0.03 | 0.14 | -0.30 | 0.21 | 0.766 |
| **SLC1A5** | **S535** | 0.02 | 0.21 | 0.32 | -0.13 | -0.16 | -0.37 | 0.024 |
| **SLC1A5** | **S493** | 0.05 | 0.33 | 0.43 | -0.55 | -0.23 | -0.28 | 0.015 |
| **SLC20A1** | **S265** | -0.23 | 0.23 | 0.23 | -0.25 | 0.03 | -0.10 | 0.348 |
| **SLC20A2** | **S256** | -0.11 | 0.04 | 0.15 | -0.01 | 0.00 | -0.08 | 0.53 |
| **SLC20A2** | **S256; S259** | -0.05 | 0.12 | 0.45 | -0.23 | -0.13 | -0.30 | 0.063 |
| **SLC20A2** | **S422** | 0.52 | 0.23 | -0.12 | -0.28 | -0.50 | -0.08 | 0.088 |
| **SLC23A2** | **S81** | -0.30 | -0.01 | -0.35 | 0.02 | 0.25 | 0.27 | 0.038 |
| **SLC25A1** | **S17** | -0.70 | -0.64 | -0.52 | 0.00 | -0.29 | 1.15 | 0.111 |
| **SLC25A1** | **S156** | 0.24 | 0.24 | 0.20 | -0.13 | -0.30 | -0.38 | 0.003 |
| **SLC26A6** | **T/S** | -0.09 | 0.12 | -0.16 | 0.02 | -0.10 | 0.17 | 0.569 |
| **SLC27A3** | **S92** | 0.04 | -0.10 | -0.14 | 0.08 | 0.10 | 0.01 | 0.1 |
| **SLC29A1** | **S269** | -0.11 | 0.12 | -0.04 | 0.08 | 0.01 | -0.07 | 0.843 |
| **SLC29A2** | **S252** | -0.21 | -0.22 | 0.38 | 0.31 | -0.44 | 0.00 | 0.94 |
| **SLC29A2** | **S252** | 0.03 | -0.25 | 0.38 | 0.13 | -0.33 | -0.08 | 0.556 |
| **SLC2A1** | **T478** | -0.07 | 0.29 | 0.18 | 0.08 | -0.55 | -0.07 | 0.227 |
| **SLC2A1** | **T478** | 0.04 | 0.17 | 0.33 | -0.19 | -0.05 | -0.43 | 0.046 |
| **SLC2A11** | **S487** | -0.23 | 0.05 | 0.07 | -0.04 | -0.22 | 0.30 | 0.798 |
| **SLC30A6** | **S381** | -0.07 | 0.12 | -0.39 | 0.10 | 0.14 | 0.02 | 0.26 |
| **SLC35B2** | **S427** | -0.21 | 0.02 | -0.38 | 0.24 | -0.09 | 0.31 | 0.112 |
| **SLC35B2** | **S427** | -0.17 | -0.18 | -0.15 | 0.30 | -0.04 | 0.17 | 0.036 |
| **SLC35B2** | **S137** | -0.06 | -0.06 | 0.36 | -0.19 | -0.02 | -0.10 | 0.282 |
| **SLC35B2** | **S427** | 0.13 | 0.16 | 0.06 | 0.17 | 0.06 | -0.81 | 0.373 |
| **SLC35C2** | **S** | -0.21 | -0.08 | -0.06 | 0.25 | -0.02 | 0.07 | 0.075 |
| **SLC35F2** | **S371** | -0.03 | 0.02 | -0.08 | -0.12 | -0.13 | 0.30 | 0.769 |
| **SLC35F6** | **T365** | -0.05 | 0.24 | 0.30 | -0.10 | 0.10 | -0.70 | 0.209 |
| **SLC38A1** | **S52** | -0.15 | -0.03 | 0.15 | 0.02 | -0.05 | 0.04 | 0.898 |
| **SLC38A1** | **S52** | -0.17 | 0.07 | 0.21 | 0.08 | -0.27 | 0.03 | 0.577 |
| **SLC38A1** | **S52; S56** | 0.08 | 0.14 | 0.07 | -0.05 | -0.16 | -0.11 | 0.007 |
| **SLC38A1** | **S49** | 0.13 | -0.17 | 0.34 | -0.18 | -0.17 | -0.03 | 0.223 |
| **SLC38A1** | **S52** | 0.13 | 0.07 | 0.12 | -0.15 | -0.10 | -0.10 | 7E-04 |
| **SLC38A1** | **S52; S56** | 0.03 | 0.14 | 0.30 | -0.16 | -0.17 | -0.23 | 0.012 |
| **SLC38A1** | **S52** | 0.14 | 0.24 | 0.22 | -0.11 | -0.20 | -0.39 | 0.008 |
| **SLC38A1** | **S49** | -0.03 | 0.19 | 0.43 | -0.32 | -0.11 | -0.32 | 0.042 |
| **SLC38A2** | **S/Y** | 0.14 | 0.29 | 0.22 | -0.91 | -0.21 | 0.16 | 0.165 |
| **SLC38A2** | **S/Y** | 0.14 | 0.29 | 0.22 | -0.91 | -0.21 | 0.16 | 0.165 |
| **SLC38A2** | **S/Y/T** | 0.29 | 0.42 | 0.23 | -0.40 | -0.18 | -0.69 | 0.009 |

| | | | | | | | | |
|---|---|---|---|---|---|---|---|---|
| SLC39A7 | S293 | 0.00 | -0.06 | 0.14 | 0.04 | 0.12 | -0.28 | 0.653 |
| SLC41A2 | T148; S/T | 0.07 | -0.04 | -0.08 | -0.09 | -0.12 | 0.23 | 0.868 |
| SLC41A3 | T49 | -0.24 | 0.25 | -0.02 | -0.18 | -0.61 | 0.53 | 0.831 |
| SLC43A2 | S297 | -0.02 | -0.21 | 0.18 | -0.16 | 0.04 | 0.12 | 0.917 |
| SLC44A2 | T14 | 0.22 | 0.30 | 0.35 | -0.54 | -0.19 | -0.38 | 0.004 |
| SLC45A4 | S348 | 0.05 | 0.26 | 0.29 | -0.25 | -0.32 | -0.15 | 0.009 |
| SLC46A1 | S458 | 0.00 | 0.06 | -0.01 | -0.20 | -0.13 | 0.23 | 0.729 |
| SLC4A11 | S308 | -0.35 | 0.18 | -0.39 | 0.39 | -0.51 | 0.38 | 0.475 |
| SLC4A11 | S61; S64 | -0.09 | 0.09 | -0.08 | -0.08 | -0.09 | 0.22 | 0.729 |
| SLC4A1AP | S82 | -0.18 | -0.09 | 0.01 | 0.30 | -0.49 | 0.30 | 0.672 |
| SLC4A2 | S170 | -0.17 | 0.04 | 0.26 | -0.04 | -0.14 | 0.01 | 0.489 |
| SLC4A2 | S132 | -0.36 | 0.04 | 0.58 | -0.41 | 0.00 | -0.09 | 0.447 |
| SLC4A7 | S233 | -0.45 | -0.51 | -0.55 | 0.39 | 0.26 | 0.45 | 2E-04 |
| SLC4A7 | S242 | 0.01 | 0.18 | -0.08 | -0.39 | 0.30 | -0.13 | 0.63 |
| SLC4A7 | S403 | 0.02 | 0.07 | 0.11 | -0.09 | -0.22 | 0.08 | 0.202 |
| SLC4A7 | S84 | 0.10 | 0.24 | 0.15 | -0.23 | -0.14 | -0.19 | 0.002 |
| SLC4A7 | S84 | 0.07 | 0.23 | 0.19 | -0.21 | -0.12 | -0.23 | 0.004 |
| SLC4A7 | S403 | 0.18 | 0.25 | 0.25 | -0.26 | -0.16 | -0.40 | 0.003 |
| SLC7A1 | T620 | -0.25 | -0.18 | -0.54 | 0.31 | 0.31 | 0.15 | 0.009 |
| SLC7A11 | S26 | -0.01 | -0.34 | -0.14 | 0.36 | 0.00 | 0.05 | 0.112 |
| SLC7A11 | S356 | -0.37 | -0.06 | -0.02 | -0.02 | -0.32 | 0.59 | 0.466 |
| SLC9A1 | S785; S788 | -0.10 | -0.16 | -0.05 | 0.05 | -0.17 | 0.36 | 0.316 |
| SLC9A1 | S/T | -0.23 | 0.00 | 0.13 | 0.04 | -0.11 | 0.12 | 0.716 |
| SLC9A1 | S796 | -0.02 | 0.02 | -0.07 | -0.01 | 0.24 | -0.20 | 0.811 |
| SLC9A1 | S785 | -0.09 | -0.01 | 0.07 | 0.02 | 0.04 | -0.04 | 0.698 |
| SLC9A1 | S693 | -0.05 | 0.18 | 0.12 | 0.03 | -0.12 | -0.20 | 0.133 |
| SLC9A1 | S703 | 0.15 | 0.13 | 0.13 | -0.36 | 0.11 | -0.24 | 0.098 |
| SLC9A1 | S/T | 0.09 | 0.12 | 0.28 | -0.38 | 0.03 | -0.24 | 0.058 |
| SLC9A1 | S/T | 0.21 | 0.22 | 0.24 | 0.02 | -0.12 | -0.83 | 0.112 |
| SLC9A1 | S/T | 0.04 | 0.64 | -0.01 | -0.20 | -0.15 | -0.62 | 0.1 |
| SLC9A2 | T746; T749 | -0.02 | -0.11 | 0.19 | -0.14 | -0.17 | 0.20 | 0.705 |
| SLC9A7 | S545 | -0.18 | 0.01 | -0.10 | 0.52 | -0.21 | -0.19 | 0.621 |
| SLCO2A1 | S16; S/T | -0.20 | 0.28 | 0.30 | -0.12 | -0.24 | -0.12 | 0.166 |
| SLF1 | T809 | -0.18 | 0.01 | -0.23 | 0.10 | -0.10 | 0.32 | 0.16 |
| SLF1 | S/T | 0.05 | -0.03 | -0.35 | -1.50 | 0.33 | 0.96 | 0.963 |
| SLF2 | S603; S607; S614 | 0.07 | -0.23 | -0.24 | 0.02 | -0.19 | 0.44 | 0.358 |
| SLF2 | S710; T711 | -0.05 | 0.16 | -0.02 | 0.01 | 0.09 | -0.21 | 0.597 |
| SLF2 | S603; S614 | -0.02 | 0.19 | 0.05 | -0.06 | 0.01 | -0.20 | 0.143 |
| SLF2 | S317 | 0.26 | 0.21 | -0.06 | -0.46 | -0.07 | 0.02 | 0.159 |
| SLF2 | S603; S607 | 0.21 | 0.20 | 0.15 | -0.08 | -0.29 | -0.29 | 0.005 |
| SLFN13 | S170 | 0.16 | 0.22 | -0.14 | 0.27 | -0.13 | -0.53 | 0.456 |
| SLIRP | T/S | -0.05 | -0.13 | -0.11 | 0.12 | -0.08 | 0.21 | 0.111 |
| SLIRP | S102 | -0.01 | -0.11 | -0.17 | 0.12 | -0.33 | 0.38 | 0.503 |
| SLIRP | S102 | 0.22 | 0.21 | -0.28 | -0.17 | 0.07 | -0.13 | 0.521 |
| SLMAP | S148 | -0.02 | 0.13 | -0.08 | -0.01 | 0.03 | -0.05 | 0.731 |
| SLMAP | S452 | 0.03 | -0.02 | 0.04 | 0.06 | -0.03 | -0.08 | 0.476 |
| SLMAP | S280 | 0.00 | 0.13 | 0.10 | -0.14 | -0.23 | 0.11 | 0.211 |
| SLMAP | S452 | 0.39 | 0.04 | 0.07 | -0.31 | 0.12 | -0.48 | 0.14 |

| | | | | | | | | |
|---|---|---|---|---|---|---|---|---|
| **SLTM** | **S126** | -0.62 | -0.95 | -0.79 | 0.52 | 0.45 | 0.54 | 2E-04 |
| **SLTM** | **S126** | -0.73 | -0.47 | -0.41 | 0.46 | 0.45 | 0.24 | 0.002 |
| **SLTM** | **S** | -0.11 | -0.35 | -0.38 | 0.26 | 0.07 | 0.35 | 0.013 |
| **SLTM** | **S144** | -0.19 | -0.16 | -0.31 | 0.28 | 0.02 | 0.25 | 0.012 |
| **SLTM** | **S93** | -0.20 | -0.04 | -0.13 | 0.06 | 0.14 | 0.15 | 0.012 |
| **SLTM** | **S553** | 0.04 | -0.04 | -0.19 | 0.04 | 0.01 | 0.12 | 0.189 |
| **SLTM** | **S1002** | 0.03 | -0.03 | -0.17 | -0.03 | 0.23 | -0.05 | 0.375 |
| **SLTM** | **T74** | -0.15 | 0.09 | -0.06 | 0.10 | 0.07 | -0.07 | 0.453 |
| **SLTM** | **S553** | 0.09 | -0.17 | -0.03 | 0.14 | 0.00 | -0.05 | 0.549 |
| **SLTM** | **S** | 0.12 | -0.06 | -0.14 | -0.06 | 0.42 | -0.42 | 0.995 |
| **SLTM** | **S344** | 0.13 | 0.01 | -0.11 | 0.07 | -0.20 | 0.08 | 0.808 |
| **SLTM** | **S376** | 0.17 | 0.11 | -0.15 | -0.15 | -0.03 | 0.01 | 0.419 |
| **SLTM** | **S888** | -0.06 | 0.07 | 0.18 | -0.34 | 0.11 | -0.02 | 0.393 |
| **SLTM** | **S289** | 0.09 | -0.02 | 0.15 | -0.07 | -0.06 | -0.10 | 0.046 |
| **SLTM** | **T/S** | 0.18 | 0.12 | 0.03 | -0.12 | -0.12 | -0.11 | 0.007 |
| **SLTM** | **S289** | 0.14 | 0.16 | 0.24 | -0.17 | -0.06 | -0.41 | 0.022 |
| **SLTM** | **S371; S/T** | 0.19 | 0.25 | 0.13 | -0.31 | -0.02 | -0.35 | 0.019 |
| **SLTM** | **S289** | 0.26 | 0.28 | 0.20 | -0.32 | 0.09 | -0.79 | 0.082 |
| **SLTM** | **S/T** | 0.36 | 0.37 | 0.33 | -0.24 | -0.23 | -1.07 | 0.036 |
| **SLTM** | **S543** | 0.30 | 0.39 | 0.42 | -0.51 | -0.20 | -0.87 | 0.01 |
| **SLU7** | **S466** | -0.45 | -0.51 | -0.62 | 0.58 | 0.17 | 0.37 | 0.002 |
| **SLU7** | **S235** | 0.09 | -0.02 | 0.06 | -0.22 | -0.05 | 0.11 | 0.394 |
| **SLU7** | **S235** | 0.12 | 0.10 | -0.03 | -0.11 | -0.10 | 0.01 | 0.108 |
| **SLU7** | **S215** | 0.06 | 0.09 | 0.10 | -0.12 | -0.06 | -0.08 | 9E-04 |
| **SLU7** | **S215** | 0.16 | 0.14 | 0.00 | -0.18 | 0.05 | -0.21 | 0.089 |
| **SLX4** | **S1469** | -0.24 | -0.30 | -0.34 | 0.54 | -0.10 | 0.21 | 0.05 |
| **SLX4** | **S1469** | -0.25 | -0.42 | -0.19 | 0.28 | 0.03 | 0.37 | 0.014 |
| **SLX4** | **S1244** | -0.13 | -0.42 | -0.15 | 0.30 | -0.05 | 0.31 | 0.05 |
| **SLX4** | **S467** | -0.11 | -0.03 | -0.46 | 0.36 | -0.12 | 0.22 | 0.147 |
| **SLX4** | **S1028** | -0.11 | -0.27 | -0.08 | 0.16 | 0.10 | 0.15 | 0.009 |
| **SLX4** | **S** | -0.11 | -0.16 | -0.21 | 0.26 | -0.19 | 0.32 | 0.153 |
| **SLX4** | **S584** | -0.40 | -0.14 | 0.20 | 0.10 | -0.18 | 0.30 | 0.456 |
| **SLX4** | **S199** | -0.15 | 0.13 | -0.04 | 0.11 | -0.06 | -0.02 | 0.748 |
| **SLX4** | **S960** | -0.16 | -0.11 | 0.18 | -0.08 | -0.37 | 0.40 | 0.964 |
| **SLX4** | **S1206** | 0.32 | -0.10 | 0.04 | -0.02 | -0.29 | -0.01 | 0.278 |
| **SLX4** | **T/S** | 0.00 | 0.22 | 0.24 | -0.10 | -0.04 | -0.42 | 0.071 |
| **SLX4IP** | **S271** | -0.13 | -0.14 | 0.21 | 0.31 | -0.12 | -0.21 | 0.951 |
| **SLX4IP** | **S** | 0.35 | 0.25 | -0.73 | 0.01 | 0.40 | -0.71 | 0.914 |
| **SLX4IP** | **S212** | -0.26 | 0.21 | 0.08 | -0.46 | 0.14 | 0.17 | 0.814 |
| **SLX4IP** | **S** | -0.02 | 0.16 | 0.22 | 0.02 | 0.01 | -0.50 | 0.209 |
| **SLX4IP** | **S396** | 0.17 | 0.33 | 0.12 | 0.11 | -0.67 | -0.29 | 0.104 |
| **SLX9** | **T34** | -0.22 | -0.11 | -0.07 | 0.18 | 0.03 | 0.15 | 0.016 |
| **SMAP** | **S17** | 0.07 | 0.02 | 0.09 | -0.06 | -0.19 | 0.05 | 0.164 |
| **SMAP** | **S17** | 0.45 | 0.01 | 0.05 | -0.12 | -0.16 | -0.37 | 0.076 |
| **SMAP2** | **S219** | -0.20 | -0.09 | -0.11 | 0.30 | -0.01 | 0.05 | 0.069 |
| **SMARCA2** | **S1410** | -0.78 | -1.29 | -1.22 | 0.71 | 0.54 | 0.57 | 6E-04 |
| **SMARCA2** | **S666** | -0.64 | -0.81 | -0.92 | 0.33 | 0.40 | 0.75 | 0.001 |
| **SMARCA2** | **S591; S** | -0.53 | -0.36 | -0.32 | 0.27 | 0.24 | 0.42 | 0.001 |

| | | | | | | | | |
|---|---|---|---|---|---|---|---|---|
| **SMARCA2** | **S** | -0.19 | 0.07 | -0.23 | 0.07 | 0.06 | 0.17 | 0.094 |
| **SMARCA2** | **S** | -0.14 | -0.16 | 0.08 | 0.12 | -0.30 | 0.31 | 0.58 |
| **SMARCA2** | **S591** | -0.31 | -0.05 | 0.10 | 0.05 | -0.44 | 0.47 | 0.717 |
| **SMARCA2** | **S** | -0.08 | 0.06 | -0.11 | 0.00 | 0.08 | 0.05 | 0.206 |
| **SMARCA2** | **S1377** | 0.15 | 0.07 | -0.17 | -0.05 | -0.03 | 0.01 | 0.72 |
| **SMARCA2** | **S1572; S1585; T1587** | -0.04 | 0.07 | 0.17 | 0.01 | -0.25 | 0.01 | 0.249 |
| **SMARCA2** | **S1572; S1585; T1587** | 0.26 | 0.27 | 0.14 | -0.21 | -0.22 | -0.39 | 0.002 |
| **SMARCA4** | **S1382** | -0.32 | -0.46 | -0.17 | 0.35 | 0.01 | 0.38 | 0.017 |
| **SMARCA4** | **T/S** | -0.21 | -0.20 | -0.12 | 0.06 | 0.05 | 0.35 | 0.031 |
| **SMARCA4** | **S695; S699** | -0.12 | -0.17 | -0.14 | 0.02 | 0.05 | 0.30 | 0.043 |
| **SMARCA4** | **T609** | -0.04 | -0.17 | -0.14 | 0.15 | 0.09 | 0.07 | 0.009 |
| **SMARCA4** | **T596; S613** | -0.12 | -0.07 | -0.06 | 0.19 | -0.10 | 0.13 | 0.152 |
| **SMARCA4** | **S613** | -0.03 | -0.11 | -0.09 | -0.08 | 0.23 | 0.05 | 0.195 |
| **SMARCA4** | **S610** | -0.10 | 0.06 | -0.18 | -0.02 | 0.17 | 0.04 | 0.213 |
| **SMARCA4** | **T/S** | 0.00 | 0.14 | -0.24 | 0.09 | 0.10 | -0.13 | 0.689 |
| **SMARCA4** | **S610** | -0.11 | 0.06 | -0.02 | 0.01 | -0.05 | 0.10 | 0.534 |
| **SMARCA4** | **S/T** | 0.03 | -0.13 | 0.00 | 0.21 | -0.40 | 0.21 | 0.847 |
| **SMARCA4** | **T/S** | 0.07 | 0.02 | -0.08 | -0.09 | -0.02 | 0.10 | 0.93 |
| **SMARCA4** | **S695** | 0.16 | 0.02 | -0.06 | -0.05 | -0.06 | -0.02 | 0.266 |
| **SMARCA4** | **S695** | 0.04 | 0.05 | 0.04 | -0.04 | -0.08 | -0.02 | 0.007 |
| **SMARCA4** | **T/S** | 0.23 | -0.02 | -0.08 | -0.34 | -0.11 | 0.24 | 0.598 |
| **SMARCA4** | **S1452** | 0.12 | 0.07 | -0.02 | -0.20 | 0.04 | -0.02 | 0.24 |
| **SMARCA4** | **S1452** | 0.21 | 0.08 | -0.10 | -0.20 | -0.08 | 0.05 | 0.289 |
| **SMARCA4** | **S1417** | 0.21 | -0.09 | 0.09 | -0.04 | -0.06 | -0.15 | 0.174 |
| **SMARCA4** | **S657; S660; S662** | 0.07 | 0.13 | 0.25 | -0.17 | -0.06 | -0.30 | 0.019 |
| **SMARCA4** | **S660; S662; S/T** | 0.08 | 0.07 | 0.33 | -0.21 | -0.22 | -0.13 | 0.019 |
| **SMARCA4** | **S1570; S1575; S1586** | 0.29 | 0.13 | 0.19 | -0.16 | -0.13 | -0.44 | 0.015 |
| **SMARCA5** | **S137** | -0.38 | -0.74 | -0.45 | 0.44 | 0.16 | 0.52 | 0.004 |
| **SMARCA5** | **S755** | -0.48 | -0.23 | -0.29 | 0.31 | 0.22 | 0.28 | 0.002 |
| **SMARCA5** | **S47; S50; S66** | -0.31 | -0.35 | -0.20 | 0.21 | 0.08 | 0.41 | 0.008 |
| **SMARCA5** | **S66** | 0.03 | -0.01 | -0.24 | 0.16 | -0.10 | 0.11 | 0.332 |
| **SMARCA5** | **S66** | -0.10 | 0.00 | -0.04 | 0.07 | -0.06 | 0.11 | 0.229 |
| **SMARCA5** | **S825** | 0.00 | -0.06 | 0.02 | 0.00 | 0.10 | -0.07 | 0.738 |
| **SMARCA5** | **S541** | -0.14 | -0.04 | 0.17 | -0.17 | -0.15 | 0.27 | 0.949 |
| **SMARCA5** | **S825** | 0.00 | 0.01 | 0.00 | -0.18 | 0.19 | -0.04 | 0.895 |
| **SMARCA5** | **S66; S** | 0.12 | 0.43 | -0.43 | 0.04 | 0.16 | -0.56 | 0.662 |
| **SMARCA5** | **S171** | 0.20 | 0.00 | 0.14 | -0.31 | 0.11 | -0.22 | 0.145 |
| **SMARCAD1** | **S152** | -0.07 | -0.58 | -0.41 | 0.29 | 0.30 | 0.22 | 0.015 |
| **SMARCAD1** | **S124; S127; S132** | -0.19 | -0.15 | -0.21 | 0.10 | -0.01 | 0.37 | 0.043 |
| **SMARCAD1** | **S124; S127; S132** | -0.31 | -0.04 | -0.15 | 0.10 | 0.10 | 0.22 | 0.024 |
| **SMARCAD1** | **S39; S/T** | 0.30 | -0.63 | -0.15 | 0.09 | -0.26 | 0.40 | 0.518 |
| **SMARCAD1** | **S211** | 0.18 | -0.16 | -0.09 | -0.09 | 0.08 | 0.05 | 0.771 |
| **SMARCAD1** | **S/T** | -0.03 | 0.01 | 0.01 | -0.03 | -0.16 | 0.18 | 0.992 |
| **SMARCAD1** | **S132** | -0.03 | 0.01 | 0.01 | -0.03 | -0.16 | 0.18 | 0.992 |
| **SMARCAD1** | **S211** | 0.29 | -0.20 | -0.09 | -0.10 | 0.04 | 0.01 | 0.918 |
| **SMARCAD1** | **S39; S/T** | -0.03 | -0.22 | 0.29 | -0.21 | -0.08 | 0.19 | 0.823 |
| **SMARCAD1** | **S28; S37** | -0.69 | 0.20 | 0.72 | -0.72 | 0.13 | -0.18 | 0.52 |
| **SMARCC1** | **T765** | -0.71 | -0.58 | -0.33 | 0.41 | 0.41 | 0.34 | 0.001 |

| | | | | | | | | |
|---|---|---|---|---|---|---|---|---|
| SMARCC1 | T | -0.05 | 0.02 | -0.48 | -0.01 | -0.16 | 0.51 | 0.332 |
| SMARCC1 | T417 | -0.16 | 0.04 | 0.03 | -0.10 | -0.01 | 0.18 | 0.652 |
| SMARCC1 | S328; S330; T337 | -0.08 | -0.05 | 0.12 | -0.08 | -0.04 | 0.12 | 0.984 |
| SMARCC1 | S330 | 0.02 | 0.03 | -0.05 | -0.08 | -0.02 | 0.09 | 0.989 |
| SMARCC1 | T398 | 0.10 | 0.04 | 0.00 | -0.10 | -0.06 | 0.01 | 0.09 |
| SMARCC1 | S328 | -0.10 | -0.07 | 0.35 | -0.23 | 0.03 | -0.05 | 0.423 |
| SMARCC1 | S328; S330 | 0.10 | 0.10 | 0.01 | -0.11 | 0.03 | -0.15 | 0.07 |
| SMARCC1 | T398 | 0.11 | 0.08 | 0.08 | -0.13 | -0.09 | -0.07 | 0.001 |
| SMARCC1 | T398 | 0.27 | 0.08 | 0.14 | -0.28 | -0.08 | -0.20 | 0.013 |
| SMARCC1 | S395 | 0.26 | 0.12 | 0.16 | -0.79 | -0.12 | 0.13 | 0.188 |
| SMARCC1 | T398 | 0.29 | 0.36 | 0.11 | -0.52 | -0.06 | -0.38 | 0.021 |
| SMARCC1 | T398 | 0.08 | 0.35 | 0.46 | -0.62 | -0.10 | -0.49 | 0.022 |
| SMARCC2 | T376; T378 | -0.23 | -0.31 | -0.55 | 0.48 | -0.02 | 0.35 | 0.023 |
| SMARCC2 | S387 | -0.14 | -0.31 | -0.30 | 0.29 | 0.10 | 0.23 | 0.004 |
| SMARCC2 | T/S | -0.22 | -0.32 | -0.15 | 0.10 | 0.19 | 0.30 | 0.005 |
| SMARCC2 | T378; S387 | -0.40 | -0.16 | -0.06 | 0.24 | 0.25 | 0.02 | 0.04 |
| SMARCC2 | S399 | -0.16 | -0.08 | -0.29 | 0.20 | 0.12 | 0.15 | 0.007 |
| SMARCC2 | S/T | -0.13 | -0.11 | -0.28 | 0.18 | 0.10 | 0.17 | 0.006 |
| SMARCC2 | S/T | -0.26 | 0.13 | -0.40 | 0.02 | 0.28 | 0.11 | 0.152 |
| SMARCC2 | S754 | -0.18 | -0.17 | -0.09 | 0.21 | 0.20 | -0.02 | 0.026 |
| SMARCC2 | T378 | -0.08 | -0.09 | -0.25 | 0.07 | 0.07 | 0.23 | 0.025 |
| SMARCC2 | S347 | 0.02 | 0.01 | -0.16 | 0.03 | 0.07 | 0.02 | 0.239 |
| SMARCC2 | S283; S286 | -0.03 | -0.11 | 0.00 | -0.10 | -0.02 | 0.23 | 0.467 |
| SMARCC2 | S283 | -0.26 | -0.03 | 0.12 | -0.27 | -0.06 | 0.40 | 0.734 |
| SMARCC2 | S806 | 0.05 | -0.10 | -0.03 | -0.17 | 0.10 | 0.12 | 0.716 |
| SMARCC2 | S347 | 0.10 | 0.00 | -0.16 | -0.08 | 0.08 | 0.04 | 0.736 |
| SMARCC2 | S302; S304 | 0.11 | 0.00 | -0.13 | -0.05 | 0.01 | 0.05 | 0.901 |
| SMARCC2 | S347 | 0.11 | -0.04 | -0.10 | -0.25 | 0.18 | 0.07 | 0.95 |
| SMARCC2 | S302; S304 | 0.00 | 0.00 | -0.01 | -0.02 | -0.14 | 0.15 | 0.993 |
| SMARCC2 | S302; S304 | 0.19 | 0.01 | -0.18 | -0.02 | -0.13 | 0.10 | 0.865 |
| SMARCC2 | S347 | 0.11 | 0.07 | -0.14 | -0.01 | 0.09 | -0.15 | 0.713 |
| SMARCC2 | T378 | 0.20 | -0.08 | -0.07 | -0.13 | 0.30 | -0.31 | 0.77 |
| SMARCC2 | S/T | -0.21 | 0.03 | 0.27 | -0.03 | -0.15 | 0.04 | 0.659 |
| SMARCC2 | S302 | 0.14 | 0.09 | -0.07 | -0.11 | 0.03 | -0.10 | 0.216 |
| SMARCC2 | S399 | 0.29 | 0.09 | -0.09 | -0.06 | 0.16 | -0.53 | 0.361 |
| SMARCC2 | S283 | 0.16 | 0.11 | 0.07 | -0.27 | -0.03 | -0.09 | 0.03 |
| SMARCC2 | S399 | 0.24 | 0.17 | -0.05 | -0.11 | 0.13 | -0.49 | 0.245 |
| SMARCC2 | S283 | 0.16 | 0.18 | 0.24 | -0.31 | -0.07 | -0.31 | 0.007 |
| SMARCD2 | T217 | -0.12 | -0.17 | -0.15 | 0.05 | 0.07 | 0.26 | 0.017 |
| SMARCD2 | S203 | 0.00 | 0.05 | -0.16 | -0.04 | 0.09 | 0.03 | 0.427 |
| SMARCD2 | S203 | 0.77 | 0.00 | -0.28 | -0.36 | 0.06 | -0.65 | 0.273 |
| SMARCE1 | S314; S | -0.58 | -0.81 | -0.70 | 0.40 | 0.50 | 0.49 | 9E-05 |
| SMARCE1 | S314 | -0.49 | -0.42 | -0.35 | 0.32 | 0.08 | 0.54 | 0.006 |
| SMARCE1 | T/S | -0.05 | -0.26 | -0.18 | 0.21 | 0.25 | -0.05 | 0.055 |
| SMC1A | S957 | -0.71 | -0.75 | -0.61 | 0.46 | 0.42 | 0.52 | 2E-05 |
| SMC1A | S/T | -0.33 | -0.43 | -0.12 | 0.27 | 0.38 | 0.05 | 0.017 |
| SMC1A | S957 | -0.34 | -0.25 | -0.05 | 0.21 | 0.37 | -0.05 | 0.061 |
| SMC3 | S1083 | -0.60 | -0.71 | -0.12 | 0.44 | 0.17 | 0.41 | 0.015 |

| | | | | | | | | |
|---|---|---|---|---|---|---|---|---|
| SMC4 | S41 | 0.00 | -0.01 | -0.05 | -0.07 | -0.06 | 0.16 | 0.713 |
| SMC4 | S28 | -0.14 | 0.05 | 0.33 | 0.08 | -0.24 | -0.15 | 0.336 |
| SMC5 | S12 | -0.09 | -0.26 | 0.07 | 0.25 | -0.11 | 0.09 | 0.299 |
| SMC5 | T10; S12 | 0.04 | -0.01 | 0.35 | -0.19 | 0.11 | -0.41 | 0.196 |
| SMC5 | S12 | -0.02 | 0.30 | 0.27 | -0.28 | -0.03 | -0.37 | 0.045 |
| SMC6 | S669 | 0.02 | -0.02 | -0.12 | 0.18 | -0.07 | -0.01 | 0.46 |
| SMG6 | S203 | 0.19 | 0.10 | -0.08 | -0.07 | -0.20 | 0.03 | 0.225 |
| SMIM1 | S22; S27 | -0.08 | -0.07 | 0.09 | 0.02 | -0.04 | 0.07 | 0.638 |
| SMIM1 | S17 | -0.10 | -0.08 | 0.16 | 0.03 | -0.21 | 0.16 | 0.974 |
| SMIM1 | S22 | 0.00 | 0.09 | 0.12 | -0.12 | -0.05 | -0.05 | 0.025 |
| SMIM14 | T/S | -0.44 | -0.56 | -0.25 | 0.48 | 0.27 | 0.19 | 0.005 |
| SMN1 | S31 | -0.13 | 0.17 | 0.06 | -0.04 | -0.32 | 0.19 | 0.622 |
| SMN1 | S31 | 0.04 | 0.08 | 0.12 | 0.01 | -0.06 | -0.21 | 0.072 |
| SMPD4 | S169 | 0.26 | -0.02 | -0.27 | -0.34 | -0.07 | 0.31 | 0.924 |
| SMPD4 | S169 | 0.20 | 0.03 | -0.17 | -0.36 | -0.21 | 0.39 | 0.764 |
| SMPD4 | S792 | 0.32 | 0.13 | 0.14 | -0.33 | 0.13 | -0.58 | 0.103 |
| SMTN | S/T | -0.01 | -0.07 | 0.05 | 0.04 | -0.11 | 0.09 | 0.824 |
| SMTN | S514 | 0.03 | -0.14 | 0.21 | 0.10 | -0.06 | -0.19 | 0.547 |
| SMTN | T/S | 0.07 | -0.02 | 0.10 | -0.01 | -0.12 | -0.03 | 0.106 |
| SMTN | S277 | 0.14 | 0.10 | 0.34 | -0.35 | -0.27 | -0.07 | 0.019 |
| SNAP23 | T157 | 0.00 | -0.10 | -0.17 | 0.18 | -0.08 | 0.14 | 0.15 |
| SNAP23 | S136 | -0.03 | 0.08 | -0.05 | -0.02 | 0.02 | 0.00 | 0.977 |
| SNAP23 | S136 | -0.03 | 0.08 | -0.05 | -0.02 | 0.02 | 0.00 | 0.977 |
| SNAP23 | S110 | 0.16 | 0.14 | -0.03 | 0.04 | -0.26 | -0.09 | 0.145 |
| SNAP23 | S110 | 0.04 | 0.18 | 0.29 | -0.30 | 0.05 | -0.37 | 0.064 |
| SNAPC4 | S599 | 0.14 | -0.17 | -0.28 | -0.18 | 0.12 | 0.28 | 0.387 |
| SNAPC4 | S1224 | 0.10 | 0.18 | 0.08 | -0.13 | 0.08 | -0.37 | 0.12 |
| SND1 | S426 | 0.02 | 0.11 | -0.28 | 0.18 | -0.13 | 0.06 | 0.588 |
| SND1 | T240 | -0.07 | -0.02 | -0.04 | -0.07 | 0.01 | 0.17 | 0.322 |
| SND1 | S645 | 0.06 | -0.20 | 0.09 | 0.22 | -0.05 | -0.17 | 0.926 |
| SNIP1 | S | -0.39 | -0.38 | -0.30 | 0.56 | 0.42 | -0.25 | 0.075 |
| SNIP1 | S76 | 0.02 | 0.00 | -0.12 | 0.12 | 0.03 | -0.06 | 0.404 |
| SNIP1 | S74; S76 | 0.12 | -0.01 | -0.03 | 0.00 | -0.09 | 0.00 | 0.342 |
| SNIP1 | S49; S52; T/S | 0.08 | -0.09 | 0.15 | -0.10 | -0.08 | 0.02 | 0.282 |
| SNIP1 | S52; S54 | 0.05 | 0.03 | 0.10 | -0.04 | -0.11 | -0.04 | 0.016 |
| SNIP1 | S52 | -0.01 | 0.06 | 0.14 | 0.05 | -0.17 | -0.09 | 0.163 |
| SNIP1 | S35 | 0.06 | 0.15 | 0.06 | -0.11 | -0.05 | -0.14 | 0.008 |
| SNIP1 | S35 | 0.08 | 0.02 | 0.20 | -0.10 | -0.08 | -0.16 | 0.019 |
| SNIP1 | S153 | 0.14 | 0.26 | 0.17 | -0.28 | 0.14 | -0.61 | 0.115 |
| SNIP1 | S153 | 0.22 | 0.24 | 0.28 | -0.29 | -0.02 | -0.66 | 0.037 |
| SNIP1 | S99 | 0.39 | 0.46 | 0.49 | -0.74 | -0.31 | -0.99 | 0.005 |
| SNRNP200 | S8 | -0.75 | -0.92 | -1.08 | 0.71 | 0.56 | 0.38 | 4E-04 |
| SNRNP200 | T/Y/S | 0.10 | -0.05 | -0.63 | 0.54 | -0.13 | -0.09 | 0.393 |
| SNRNP200 | T/Y/S | 0.10 | -0.05 | -0.63 | 0.54 | -0.13 | -0.09 | 0.393 |
| SNRNP200 | T/Y/S | -0.05 | 0.16 | -0.59 | 0.19 | 0.03 | 0.11 | 0.301 |
| SNRNP200 | S17 | 0.05 | -0.33 | 0.04 | -0.18 | 0.14 | 0.21 | 0.465 |
| SNRNP200 | S207 | 0.01 | -0.06 | -0.08 | 0.11 | 0.06 | -0.05 | 0.212 |
| SNRNP200 | T/Y/S | -0.21 | 0.11 | 0.12 | -0.04 | -0.06 | 0.06 | 0.878 |

| | | | | | | | | |
|---|---|---|---|---|---|---|---|---|
| **SNRNP200** | **S225** | -0.19 | -0.07 | 0.18 | 0.42 | 0.09 | -0.66 | 0.935 |
| **SNRNP200** | **S2002** | -0.19 | -0.13 | 0.26 | -0.56 | -0.29 | 0.60 | 0.886 |
| **SNRNP200** | **S225** | 0.03 | 0.14 | 0.12 | -0.07 | 0.02 | -0.28 | 0.093 |
| **SNRNP48** | **S283; S286** | 0.08 | 0.07 | 0.02 | -0.05 | -0.12 | -0.01 | 0.032 |
| **SNRNP70** | **S410** | -0.29 | 0.09 | -0.06 | 0.14 | 0.11 | -0.03 | 0.262 |
| **SNRNP70** | **S417** | -0.15 | -0.03 | -0.05 | 0.02 | -0.15 | 0.30 | 0.383 |
| **SNRNP70** | **S417** | -0.13 | 0.02 | -0.12 | -0.07 | -0.18 | 0.41 | 0.532 |
| **SNRNP70** | **S410** | -0.21 | 0.06 | 0.07 | 0.04 | 0.12 | -0.10 | 0.702 |
| **SNRNP70** | **S410** | -0.13 | 0.14 | 0.08 | 0.16 | -0.12 | -0.16 | 0.628 |
| **SNRNP70** | **S320** | 0.16 | 0.27 | 0.19 | -0.25 | -0.13 | -0.35 | 0.004 |
| **SNRNP70** | **S320** | 0.16 | 0.37 | 0.19 | -0.18 | -0.24 | -0.46 | 0.008 |
| **SNRNP70** | **S226** | 0.16 | 0.37 | 0.29 | -0.24 | -0.28 | -0.53 | 0.005 |
| **SNRPA1** | **S197** | -0.09 | -0.25 | -0.09 | 0.23 | -0.06 | 0.20 | 0.068 |
| **SNRPA1** | **S178** | 0.05 | 0.11 | -0.19 | -0.15 | 0.30 | -0.20 | 0.963 |
| **SNRPD2** | **T12** | -0.02 | -0.18 | -0.26 | 0.17 | 0.32 | -0.12 | 0.131 |
| **SNRPD2** | **T12** | -0.11 | -0.13 | -0.09 | 0.21 | 0.26 | -0.20 | 0.243 |
| **SNTB2** | **S110** | -0.18 | -0.08 | -0.07 | 0.11 | -0.03 | 0.21 | 0.058 |
| **SNTB2** | **S393** | -0.03 | -0.03 | -0.11 | 0.03 | 0.04 | 0.08 | 0.024 |
| **SNTB2** | **S95** | -0.12 | 0.00 | 0.03 | 0.07 | -0.06 | 0.06 | 0.431 |
| **SNTB2** | **S110** | 0.09 | 0.09 | 0.07 | 0.06 | -0.24 | -0.10 | 0.113 |
| **SNU13** | **S122** | -0.14 | -0.48 | -0.18 | 0.34 | 0.00 | 0.29 | 0.034 |
| **SNW1** | **S224** | -0.03 | -0.52 | -0.19 | 0.10 | 0.20 | 0.30 | 0.045 |
| **SNW1** | **S224** | -0.06 | -0.16 | 0.05 | 0.16 | -0.13 | 0.11 | 0.383 |
| **SNW1** | **S224; S232** | -0.03 | -0.03 | 0.08 | 0.06 | -0.09 | -0.01 | 0.755 |
| **SNW1** | **S224; S232** | 0.03 | 0.01 | 0.05 | 0.00 | -0.08 | -0.02 | 0.071 |
| **SNX1** | **S368** | 0.31 | -0.69 | -0.06 | 0.07 | -0.02 | 0.19 | 0.483 |
| **SNX1** | **S25; S32** | -0.08 | 0.29 | -0.49 | -0.19 | 0.24 | 0.08 | 0.626 |
| **SNX11** | **S168** | -0.44 | 0.05 | 0.32 | -0.05 | -0.26 | 0.24 | 0.999 |
| **SNX15** | **S227** | 0.10 | -0.06 | 0.02 | -0.24 | 0.21 | -0.07 | 0.703 |
| **SNX16** | **S/T/Y** | -0.07 | 0.05 | -0.07 | 0.13 | 0.04 | -0.10 | 0.526 |
| **SNX16** | **S/T/Y** | 0.11 | 0.02 | -0.02 | -0.01 | -0.05 | -0.05 | 0.146 |
| **SNX16** | **S286** | 0.01 | 0.05 | 0.07 | 0.01 | -0.06 | -0.09 | 0.067 |
| **SNX16** | **S/T** | 0.13 | 0.20 | -0.15 | 0.06 | 0.00 | -0.29 | 0.431 |
| **SNX16** | **T100** | 0.06 | 0.05 | 0.16 | -0.10 | -0.23 | 0.02 | 0.071 |
| **SNX17** | **S421** | -0.11 | 0.27 | 0.06 | -0.11 | -0.38 | 0.17 | 0.395 |
| **SNX27** | **S51** | -0.07 | -0.63 | -0.25 | 0.32 | 0.12 | 0.29 | 0.033 |
| **SNX29** | **T/S** | 0.02 | 0.22 | 0.12 | -0.41 | -0.28 | 0.22 | 0.237 |
| **SNX4** | **S22** | -0.02 | -0.07 | 0.27 | -0.22 | -0.48 | 0.36 | 0.552 |
| **SOGA1** | **S1303** | -0.04 | 0.04 | 0.01 | 0.07 | -0.16 | 0.08 | 0.959 |
| **SOGA1** | **S1017** | 0.05 | 0.18 | 0.13 | 0.06 | -0.17 | -0.31 | 0.088 |
| **SON** | **T/S** | -0.36 | -0.43 | -0.69 | 0.43 | -0.33 | 0.78 | 0.083 |
| **SON** | **S1556** | -0.15 | -0.25 | -0.45 | 0.36 | -0.30 | 0.52 | 0.151 |
| **SON** | **S160** | -0.26 | -0.19 | -0.14 | 0.20 | 0.15 | 0.16 | 6E-04 |
| **SON** | **S1556** | -0.09 | -0.14 | -0.27 | 0.23 | -0.21 | 0.36 | 0.173 |
| **SON** | **S1509** | -0.01 | -0.50 | 0.10 | 0.29 | 0.08 | -0.08 | 0.336 |
| **SON** | **S1509** | -0.01 | -0.50 | 0.10 | 0.29 | 0.08 | -0.08 | 0.336 |
| **SON** | **S1509** | -0.19 | 0.03 | -0.12 | -0.03 | 0.10 | 0.18 | 0.124 |
| **SON** | **S152** | 0.06 | -0.07 | -0.22 | 0.04 | -0.06 | 0.21 | 0.283 |

| | | | | | | | | |
|---|---|---|---|---|---|---|---|---|
| **SON** | **S1697** | 0.18 | -0.17 | -0.24 | 0.02 | 0.06 | 0.12 | 0.345 |
| **SON** | **T/S** | 0.08 | -0.06 | -0.22 | 0.09 | 0.06 | 0.02 | 0.242 |
| **SON** | **S154** | 0.00 | -0.05 | -0.06 | 0.01 | 0.05 | 0.04 | 0.036 |
| **SON** | **T959** | -0.09 | 0.00 | 0.03 | 0.04 | 0.02 | -0.01 | 0.391 |
| **SON** | **S1697** | 0.04 | -0.02 | -0.02 | -0.03 | 0.00 | 0.01 | 0.839 |
| **SON** | **S/T** | -0.02 | 0.00 | 0.10 | 0.12 | -0.07 | -0.14 | 0.554 |
| **SON** | **S2029; S2031** | 0.11 | 0.04 | 0.04 | -0.11 | -0.02 | -0.07 | 0.021 |
| **SON** | **S94** | 0.13 | 0.04 | 0.02 | -0.13 | -0.17 | 0.08 | 0.194 |
| **SON** | **S283** | 0.10 | 0.03 | 0.07 | -0.09 | 0.02 | -0.15 | 0.062 |
| **SON** | **S2129** | 0.18 | 0.12 | -0.02 | -0.04 | 0.00 | -0.29 | 0.126 |
| **SON** | **S94** | 0.14 | 0.06 | 0.11 | -0.12 | -0.04 | -0.18 | 0.011 |
| **SON** | **S2129** | 0.14 | 0.13 | 0.06 | -0.17 | 0.04 | -0.24 | 0.061 |
| **SON** | **S1782** | 0.25 | 0.16 | -0.01 | -0.20 | -0.11 | -0.14 | 0.026 |
| **SON** | **S1769** | 0.18 | 0.22 | 0.20 | -0.19 | -0.03 | -0.52 | 0.037 |
| **SON** | **S2013** | -0.23 | 0.31 | 0.61 | -0.65 | 0.10 | -0.56 | 0.153 |
| **SON** | **S1769** | 0.29 | 0.28 | 0.24 | -0.28 | -0.13 | -0.64 | 0.015 |
| **SON** | **S2009; S2013** | 0.33 | 0.30 | 0.22 | -0.27 | -0.13 | -0.75 | 0.026 |
| **SON** | **S1948; S1952** | 0.32 | 0.33 | 0.23 | -0.27 | -0.16 | -0.74 | 0.019 |
| **SON** | **S94** | 0.19 | 0.30 | 0.40 | -0.08 | -0.30 | -0.85 | 0.042 |
| **SON** | **S** | 0.35 | 0.25 | 0.30 | -0.22 | -0.14 | -0.87 | 0.038 |
| **SON** | **T/S** | 0.47 | 0.25 | 0.19 | -1.03 | 0.12 | -0.51 | 0.087 |
| **SORT1** | **S825** | 0.04 | 0.00 | -0.44 | 0.21 | -0.22 | 0.29 | 0.367 |
| **SORT1** | **T/S/Y** | 0.14 | 0.03 | -0.02 | -0.21 | 0.04 | 0.01 | 0.329 |
| **SOWAHB** | **S166** | -0.12 | 0.13 | -0.17 | 0.23 | -0.30 | 0.15 | 0.713 |
| **SOX13** | **S83** | 0.13 | -0.15 | -0.12 | 0.03 | -0.09 | 0.16 | 0.515 |
| **SOX13** | **S386** | 0.11 | 0.02 | -0.20 | 0.11 | -0.02 | -0.03 | 0.701 |
| **SOX13** | **S83** | 0.08 | -0.02 | -0.04 | 0.03 | -0.12 | 0.06 | 0.788 |
| **SOX13** | **S310** | 0.11 | 0.02 | 0.07 | 0.01 | -0.17 | -0.06 | 0.078 |
| **SOX13** | **S/T** | 0.29 | 0.04 | -0.09 | -0.32 | 0.13 | -0.12 | 0.353 |
| **SOX13** | **S** | 0.29 | 0.04 | 0.19 | 0.03 | -0.02 | -0.73 | 0.182 |
| **SP100** | **S157** | -0.26 | -0.12 | -0.15 | 0.23 | -0.06 | 0.28 | 0.049 |
| **SP100** | **S** | -0.04 | -0.06 | -0.16 | 0.22 | -0.09 | 0.09 | 0.176 |
| **SP100** | **S18** | 0.07 | -0.11 | -0.01 | -0.02 | 0.12 | -0.06 | 0.746 |
| **SP110** | **S296** | -0.07 | -0.13 | -0.48 | 0.01 | 0.21 | 0.32 | 0.061 |
| **SP110** | **S380** | 0.06 | -0.27 | -0.35 | 0.00 | 0.11 | 0.35 | 0.103 |
| **SP110** | **S194** | -0.09 | -0.09 | -0.15 | 0.03 | 0.28 | -0.03 | 0.103 |
| **SP110** | **S256** | -0.04 | 0.04 | -0.22 | -0.06 | -0.04 | 0.28 | 0.386 |
| **SP110** | **S244** | 0.05 | 0.21 | -0.08 | 0.01 | -0.08 | -0.13 | 0.248 |
| **SP110** | **S244** | 0.07 | 0.16 | 0.02 | -0.02 | 0.03 | -0.30 | 0.178 |
| **SP110** | **S244** | 0.02 | 0.22 | 0.06 | -0.10 | -0.07 | -0.16 | 0.034 |
| **SP110** | **S380** | 0.09 | 0.28 | 0.21 | -0.33 | -0.02 | -0.34 | 0.023 |
| **SP110** | **S/T** | 0.14 | -0.18 | 0.62 | -0.12 | -0.45 | -0.26 | 0.134 |
| **SP110** | **S380** | 0.39 | 0.25 | 0.18 | -0.54 | -0.16 | -0.35 | 0.007 |
| **SP3** | **S73** | -0.05 | 0.42 | 0.04 | -0.19 | -0.12 | -0.19 | 0.106 |
| **SP3** | **S73** | -0.17 | 0.39 | 0.18 | -0.15 | -0.50 | 0.08 | 0.243 |
| **SP4** | **S770** | -0.22 | -0.03 | 0.00 | 0.10 | 0.07 | 0.06 | 0.085 |
| **SP4** | **S136** | 0.29 | 0.46 | -0.32 | -0.14 | 0.00 | -0.53 | 0.269 |
| **SP4** | **S136** | 0.30 | 0.35 | -0.22 | 0.04 | 0.06 | -0.79 | 0.328 |

| | | | | | | | | |
|---|---|---|---|---|---|---|---|---|
| SPAG1 | S423 | -0.40 | -0.15 | -0.23 | 0.11 | 0.28 | 0.26 | 0.006 |
| SPAG9 | S109 | 0.10 | -0.01 | -0.39 | 0.38 | 0.02 | -0.24 | 0.55 |
| SPECC1 | S315 | -0.12 | 0.20 | 0.18 | -0.05 | -0.01 | -0.26 | 0.215 |
| SPECC1 | T/S | 0.20 | 0.21 | 0.20 | -0.40 | -0.12 | -0.21 | 0.006 |
| SPECC1L | S881 | -0.57 | 0.00 | 0.27 | 0.29 | -0.21 | 0.05 | 0.645 |
| SPECC1L | S220 | -0.03 | 0.05 | 0.12 | 0.02 | -0.12 | -0.06 | 0.173 |
| SPECC1L | S220 | 0.06 | 0.05 | 0.10 | -0.01 | -0.18 | -0.04 | 0.051 |
| SPECC1L | S384 | 0.15 | 0.11 | 0.07 | -0.13 | -0.03 | -0.21 | 0.015 |
| SPECC1L | S/T | 0.10 | 0.04 | 0.20 | 0.10 | -0.19 | -0.31 | 0.13 |
| SPECC1L | S384 | 0.04 | 0.16 | 0.24 | -0.29 | -0.23 | 0.01 | 0.044 |
| SPECC1L | S881 | 0.24 | 0.20 | 0.20 | 0.07 | -0.14 | -0.80 | 0.126 |
| SPECC1L | S/T | 0.40 | 0.21 | 0.24 | -0.16 | -0.17 | -0.82 | 0.042 |
| SPEN | S1333 | -0.74 | -0.11 | 0.11 | 0.18 | 0.09 | 0.27 | 0.177 |
| SPEN | S736 | -0.15 | -0.08 | -0.27 | 0.23 | -0.12 | 0.30 | 0.092 |
| SPEN | S623 | -0.14 | -0.15 | -0.19 | 0.07 | 0.15 | 0.21 | 0.002 |
| SPEN | S736 | -0.19 | -0.11 | -0.20 | 0.15 | -0.25 | 0.46 | 0.237 |
| SPEN | S2493 | -0.10 | 0.00 | -0.39 | 0.14 | -0.14 | 0.36 | 0.204 |
| SPEN | S/T | -0.10 | -0.22 | -0.07 | 0.28 | -0.26 | 0.26 | 0.283 |
| SPEN | S1983 | -0.07 | -0.04 | -0.14 | -0.02 | 0.13 | 0.13 | 0.049 |
| SPEN | S1268; S1278 | -0.07 | -0.03 | -0.11 | 0.02 | -0.12 | 0.27 | 0.319 |
| SPEN | S736; S740 | -0.04 | -0.06 | -0.06 | 0.10 | -0.07 | 0.11 | 0.16 |
| SPEN | S727 | 0.04 | -0.04 | -0.09 | 0.03 | 0.04 | 0.02 | 0.21 |
| SPEN | S725; S727 | -0.02 | -0.10 | 0.06 | 0.16 | -0.12 | 0.01 | 0.716 |
| SPEN | S1380; S1382 | 0.13 | -0.01 | -0.10 | -0.05 | 0.20 | -0.22 | 0.837 |
| SPEN | S1918 | 0.09 | 0.02 | -0.06 | -0.01 | -0.20 | 0.14 | 0.753 |
| SPEN | S2366 | -0.08 | 0.16 | -0.05 | 0.21 | -0.39 | 0.06 | 0.823 |
| SPEN | S2120 | 0.04 | 0.06 | -0.04 | 0.02 | -0.02 | -0.07 | 0.3 |
| SPEN | S190 | 0.08 | 0.08 | -0.05 | 0.00 | -0.04 | -0.08 | 0.209 |
| SPEN | S1222 | 0.11 | 0.02 | -0.02 | -0.01 | -0.29 | 0.16 | 0.57 |
| SPEN | T/S | 0.19 | -0.07 | -0.03 | -0.09 | -0.30 | 0.23 | 0.652 |
| SPEN | S2159 | 0.01 | 0.01 | 0.11 | 0.02 | 0.04 | -0.20 | 0.359 |
| SPEN | S244; S | 0.03 | 0.12 | -0.02 | 0.00 | 0.05 | -0.20 | 0.34 |
| SPEN | S1278 | -0.02 | 0.04 | 0.14 | -0.16 | -0.13 | 0.10 | 0.292 |
| SPEN | S751 | 0.03 | 0.07 | 0.08 | -0.03 | -0.14 | -0.01 | 0.043 |
| SPEN | S847 | 0.07 | -0.06 | 0.16 | -0.13 | -0.15 | 0.08 | 0.264 |
| SPEN | S/T | 0.09 | -0.09 | 0.16 | 0.04 | 0.11 | -0.36 | 0.476 |
| SPEN | S/T | -0.02 | 0.08 | 0.14 | 0.04 | -0.08 | -0.18 | 0.148 |
| SPEN | S770 | 0.11 | -0.04 | 0.13 | -0.05 | 0.02 | -0.21 | 0.162 |
| SPEN | S2120; S2126 | 0.10 | 0.08 | 0.04 | -0.08 | 0.05 | -0.21 | 0.114 |
| SPEN | S2120; S2124 | -0.15 | 0.39 | -0.07 | 0.09 | 0.08 | -0.48 | 0.555 |
| SPEN | S2481; S2493 | 0.09 | 0.11 | 0.05 | -0.10 | -0.08 | -0.08 | 0.001 |
| SPEN | S2120 | 0.28 | 0.01 | -0.03 | 0.01 | -0.09 | -0.23 | 0.194 |
| SPEN | S1006 | 0.12 | 0.08 | 0.09 | -0.01 | -0.01 | -0.31 | 0.112 |
| SPEN | S244 | 0.03 | -0.44 | 0.61 | 0.02 | -0.58 | 0.04 | 0.551 |
| SPEN | T2421 | 0.21 | 0.13 | 0.05 | -0.07 | -0.23 | -0.14 | 0.012 |
| SPEN | S2101 | 0.06 | -0.01 | 0.34 | -0.10 | -0.23 | -0.15 | 0.063 |
| SPEN | S2120 | 0.15 | 0.10 | 0.16 | 0.07 | 0.05 | -0.71 | 0.267 |
| SPEN | S1918 | 0.21 | 0.21 | 0.06 | -0.29 | 0.00 | -0.27 | 0.033 |

| | | | | | | | | |
|---|---|---|---|---|---|---|---|---|
| SPEN | T1619 | 0.16 | 0.17 | 0.15 | -0.20 | -0.12 | -0.23 | 4E-04 |
| SPEN | T2393 | 0.40 | -0.13 | 0.20 | -0.19 | -0.23 | -0.17 | 0.087 |
| SPEN | S768; S770 | 0.11 | 0.17 | 0.26 | -0.28 | -0.15 | -0.20 | 0.002 |
| SPEN | S2126 | 0.05 | 0.20 | 0.33 | -0.13 | -0.17 | -0.39 | 0.021 |
| SPEN | T2393 | 0.45 | -0.19 | 0.30 | -0.42 | -0.40 | 0.02 | 0.133 |
| SPEN | S2493 | 0.26 | -0.03 | 0.38 | -0.45 | 0.03 | -0.37 | 0.075 |
| SPEN | S1062 | 0.29 | 0.07 | 0.27 | -0.18 | -0.12 | -0.47 | 0.023 |
| SPEN | S1918 | 0.32 | 0.32 | 0.12 | -0.41 | -0.04 | -0.53 | 0.023 |
| SPEN | S1918 | 0.35 | 0.37 | 0.01 | -0.65 | 0.07 | -0.43 | 0.075 |
| SPHK2 | S414 | 0.18 | 0.04 | 0.07 | 0.02 | -0.28 | -0.07 | 0.096 |
| SPICE1 | S640; T642 | -0.08 | -0.10 | -0.01 | 0.16 | 0.13 | -0.12 | 0.273 |
| SPICE1 | S640 | -0.12 | 0.26 | -0.24 | 0.05 | 0.17 | -0.18 | 0.823 |
| SPICE1 | T235 | 0.06 | 0.10 | 0.09 | -0.22 | -0.16 | 0.09 | 0.139 |
| SPIN1 | S199 | -0.10 | -0.31 | -0.32 | 0.17 | 0.24 | 0.21 | 0.003 |
| SPIN2A | S195 | -0.06 | -0.27 | -0.12 | 0.43 | -0.11 | 0.03 | 0.197 |
| SPINDOC | S82 | -0.34 | -0.27 | -0.37 | 0.01 | 0.17 | 0.56 | 0.027 |
| SPINDOC | S | -0.33 | -0.23 | -0.19 | 0.10 | 0.06 | 0.45 | 0.025 |
| SPINDOC | S248; S251 | -0.17 | 0.08 | 0.00 | -0.08 | -0.08 | 0.21 | 0.729 |
| SPINDOC | S251 | 0.02 | -0.13 | 0.09 | 0.05 | -0.11 | 0.07 | 0.895 |
| SPINDOC | S308 | 0.01 | 0.04 | -0.04 | -0.02 | -0.08 | 0.08 | 0.909 |
| SPINDOC | T49 | 0.05 | 0.34 | 0.22 | -0.42 | 0.09 | -0.47 | 0.073 |
| SPIRE1 | S387 | -0.14 | 0.07 | 0.04 | 0.02 | -0.04 | 0.04 | 0.839 |
| SPIRE1 | S465 | 0.37 | 0.23 | 0.01 | -0.09 | -0.50 | -0.18 | 0.049 |
| SPOP | S6 | 0.28 | 0.21 | -0.12 | -0.06 | 0.11 | -0.58 | 0.281 |
| SPPL2B | S583 | -0.34 | -0.14 | 0.09 | -0.23 | -0.06 | 0.51 | 0.472 |
| SPPL2B | S583 | -0.34 | -0.14 | 0.09 | -0.23 | -0.06 | 0.51 | 0.472 |
| SPRED2 | S206 | -0.06 | -0.15 | -0.09 | 0.04 | 0.04 | 0.19 | 0.028 |
| SPTAN1 | S | -0.57 | -0.57 | -0.15 | 0.54 | -0.08 | 0.43 | 0.037 |
| SPTAN1 | S1217 | -0.01 | -0.02 | -0.11 | 0.14 | -0.03 | 0.02 | 0.199 |
| SPTAN1 | S1031 | -0.16 | -0.02 | 0.12 | 0.11 | 0.11 | -0.19 | 0.847 |
| SPTAN1 | S1031 | -0.17 | -0.21 | 0.25 | 0.44 | 0.01 | -0.53 | 0.952 |
| SPTAN1 | S1363 | 0.05 | -0.12 | 0.07 | 0.10 | -0.09 | -0.03 | 0.945 |
| SPTBN1 | S2161 | -0.24 | -0.25 | -0.27 | 0.33 | 0.30 | 0.00 | 0.012 |
| SPTBN1 | S1447 | 0.07 | -0.44 | -0.22 | 0.42 | 0.22 | -0.22 | 0.229 |
| SPTBN1 | S2138 | -0.30 | -0.06 | -0.05 | 0.17 | -0.09 | 0.27 | 0.133 |
| SPTBN1 | S2341 | 0.08 | -0.17 | -0.23 | 0.10 | -0.22 | 0.35 | 0.393 |
| SPTBN1 | S2138 | -0.13 | -0.10 | -0.06 | 0.14 | -0.11 | 0.23 | 0.159 |
| SPTBN1 | S257 | 0.08 | -0.13 | -0.14 | -0.02 | -0.20 | 0.35 | 0.577 |
| SPTBN1 | T2320 | -0.18 | -0.01 | -0.01 | 0.33 | -0.20 | -0.01 | 0.557 |
| SPTBN1 | S/T | -0.62 | -0.04 | 0.41 | -0.15 | -0.05 | 0.25 | 0.768 |
| SPTBN1 | S2102 | -0.07 | -0.03 | -0.03 | 0.18 | -0.06 | 0.00 | 0.317 |
| SPTBN1 | S2102 | -0.17 | -0.19 | 0.23 | -0.04 | 0.11 | 0.02 | 0.653 |
| SPTBN1 | S2138 | -0.07 | -0.05 | 0.06 | 0.05 | -0.11 | 0.11 | 0.632 |
| SPTBN1 | S/T | -0.02 | 0.08 | -0.12 | 0.20 | -0.11 | -0.05 | 0.77 |
| SPTBN1 | S/T | -0.21 | -0.08 | 0.23 | 0.07 | -0.04 | 0.00 | 0.822 |
| SPTBN1 | T2155; S2161; S/T | -0.22 | 0.14 | 0.08 | 0.14 | 0.08 | -0.28 | 0.912 |
| SPTBN1 | S817 | -0.21 | 0.16 | 0.37 | -0.08 | -0.07 | -0.27 | 0.247 |
| SPTBN1 | S2161; S2164; S2165 | 0.11 | 0.16 | 0.11 | -0.24 | 0.00 | -0.19 | 0.022 |

| | | | | | | | | |
|---|---|---|---|---|---|---|---|---|
| **SPTBN1** | **S2161; S2164** | 0.02 | 0.15 | 0.23 | -0.09 | 0.03 | -0.43 | 0.124 |
| **SPTBN1** | **S2160; S2161; S2165** | 0.10 | 0.21 | 0.27 | -0.14 | -0.04 | -0.54 | 0.054 |
| **SPTBN2** | **S2171** | -0.43 | -0.02 | -0.09 | 0.37 | -0.02 | 0.08 | 0.137 |
| **SPTBN2** | **S772** | -0.44 | 0.21 | 0.14 | -0.03 | -0.19 | 0.19 | 0.937 |
| **SPTBN2** | **S2171** | -0.55 | 0.31 | 0.27 | -0.30 | 0.14 | -0.06 | 0.797 |
| **SPTY2D1** | **S599** | -0.10 | 0.08 | -0.18 | 0.24 | -0.02 | -0.07 | 0.398 |
| **SPTY2D1** | **Y/S** | 0.01 | 0.06 | -0.28 | -0.01 | -0.16 | 0.31 | 0.535 |
| **SPTY2D1** | **Y/S** | 0.04 | 0.01 | -0.02 | 0.05 | 0.04 | -0.13 | 0.735 |
| **SPTY2D1** | **S471** | 0.13 | 0.01 | -0.11 | 0.11 | -0.12 | -0.05 | 0.759 |
| **SQSTM1** | **S284** | -0.32 | -0.27 | -0.77 | 0.58 | 0.34 | 0.03 | 0.026 |
| **SQSTM1** | **S283** | -0.48 | -0.34 | -0.40 | 0.43 | 0.53 | -0.09 | 0.023 |
| **SQSTM1** | **S28** | -0.17 | -0.30 | -0.27 | 0.28 | 0.20 | 0.16 | 1E-03 |
| **SQSTM1** | **S355** | -0.25 | -0.20 | -0.10 | 0.42 | 0.17 | -0.16 | 0.133 |
| **SQSTM1** | **T269; S272** | -0.17 | -0.12 | -0.21 | 0.10 | 0.09 | 0.26 | 0.007 |
| **SQSTM1** | **S332** | -0.40 | -0.16 | 0.01 | 0.37 | -0.16 | 0.19 | 0.183 |
| **SQSTM1** | **S355** | -0.23 | -0.18 | -0.10 | 0.40 | 0.15 | -0.13 | 0.121 |
| **SQSTM1** | **S/T** | -0.14 | -0.10 | -0.24 | 0.27 | 0.10 | 0.05 | 0.019 |
| **SQSTM1** | **S355** | -0.08 | -0.18 | -0.28 | 0.30 | 0.41 | -0.36 | 0.3 |
| **SQSTM1** | **S/T** | -0.16 | -0.02 | -0.20 | 0.25 | 0.18 | -0.10 | 0.111 |
| **SQSTM1** | **S/T** | 0.01 | -0.30 | 0.05 | 0.34 | -0.46 | 0.21 | 0.697 |
| **SQSTM1** | **S24** | -0.08 | -0.08 | 0.02 | -0.02 | -0.01 | 0.15 | 0.261 |
| **SQSTM1** | **S272** | -0.01 | -0.05 | 0.10 | 0.00 | 0.07 | -0.11 | 0.714 |
| **SQSTM1** | **T269** | -0.01 | 0.03 | 0.14 | 0.01 | 0.01 | -0.19 | 0.239 |
| **SRBD1** | **S151** | -0.42 | -0.43 | -0.38 | 0.44 | 0.12 | 0.38 | 0.002 |
| **SRBD1** | **S964** | -0.19 | -0.35 | -0.09 | 0.13 | 0.17 | 0.23 | 0.01 |
| **SRCAP** | **S2323** | -0.77 | -0.61 | -0.63 | 0.26 | 0.35 | 0.71 | 0.002 |
| **SRCAP** | **S3177** | -0.23 | -0.08 | -0.15 | 0.36 | 0.02 | 0.00 | 0.089 |
| **SRCAP** | **T572** | -0.20 | -0.03 | -0.04 | -0.07 | 0.10 | 0.21 | 0.162 |
| **SRCAP** | **S3161; S3177; T3186** | -0.08 | -0.10 | -0.08 | 0.06 | -0.03 | 0.21 | 0.087 |
| **SRCAP** | **S2787** | -0.14 | -0.01 | -0.03 | -0.02 | 0.02 | 0.17 | 0.172 |
| **SRCAP** | **S2370** | -0.22 | -0.15 | 0.21 | -0.09 | 0.10 | 0.10 | 0.584 |
| **SRCAP** | **S274; S** | -0.14 | -0.07 | 0.07 | 0.09 | -0.04 | 0.06 | 0.33 |
| **SRCAP** | **T572** | -0.09 | 0.00 | -0.04 | -0.04 | 0.03 | 0.12 | 0.217 |
| **SRCAP** | **S2370** | -0.47 | 0.01 | 0.32 | -0.11 | 0.36 | -0.30 | 0.932 |
| **SRCAP** | **S3161** | -0.09 | -0.03 | 0.08 | 0.10 | -0.05 | -0.02 | 0.741 |
| **SRCAP** | **S273; S274; S** | -0.08 | -0.19 | 0.22 | -0.17 | -0.12 | 0.27 | 0.949 |
| **SRCAP** | **S3161; S3177** | -0.01 | -0.07 | 0.06 | 0.01 | -0.05 | 0.06 | 0.846 |
| **SRCAP** | **S568; T572** | -0.03 | -0.04 | 0.06 | -0.05 | -0.03 | 0.09 | 0.928 |
| **SRCAP** | **S1882** | 0.06 | 0.14 | -0.14 | 0.03 | 0.01 | -0.10 | 0.695 |
| **SRCAP** | **S3148** | 0.11 | -0.05 | 0.06 | -0.16 | -0.16 | 0.17 | 0.486 |
| **SRCAP** | **S3016** | 0.08 | 0.15 | 0.21 | -0.37 | 0.01 | -0.16 | 0.052 |
| **SRCAP** | **S274** | 0.24 | 0.16 | 0.08 | -0.30 | -0.14 | -0.11 | 0.01 |
| **SRCIN1** | **T997; S1001; S1003** | -0.24 | -0.54 | -0.44 | 0.22 | 0.22 | 0.48 | 0.005 |
| **SRCIN1** | **S165** | -0.21 | -0.41 | -0.13 | 0.34 | 0.12 | 0.17 | 0.012 |
| **SRCIN1** | **S844** | -0.32 | -0.10 | -0.19 | -0.03 | -0.06 | 0.55 | 0.163 |
| **SRCIN1** | **S844** | -0.30 | -0.07 | -0.15 | -0.07 | -0.06 | 0.50 | 0.215 |
| **SRCIN1** | **S** | -0.35 | 0.20 | -0.29 | -0.03 | -0.09 | 0.41 | 0.362 |
| **SRCIN1** | **S1118** | -0.11 | -0.09 | -0.14 | 0.15 | 0.03 | 0.13 | 0.006 |

| | | | | | | | | |
|---|---|---|---|---|---|---|---|---|
| **SRCIN1** | **S1003** | -0.14 | -0.01 | -0.14 | 0.09 | -0.21 | 0.34 | 0.361 |
| **SRCIN1** | **S/T** | -0.11 | -0.37 | 0.21 | 0.14 | -0.30 | 0.30 | 0.603 |
| **SRCIN1** | **T/S** | -0.22 | 0.17 | -0.11 | 0.14 | 0.22 | -0.28 | 0.682 |
| **SRCIN1** | **S1154** | 0.04 | 0.02 | -0.19 | 0.08 | 0.24 | -0.24 | 0.691 |
| **SRCIN1** | **S1118** | -0.37 | 0.30 | -0.08 | 0.24 | 0.05 | -0.27 | 0.826 |
| **SRCIN1** | **S13** | 0.09 | -0.05 | -0.03 | 0.06 | -0.15 | 0.07 | 0.88 |
| **SRCIN1** | **S987** | -0.02 | 0.06 | 0.02 | -0.10 | 0.00 | 0.03 | 0.377 |
| **SRCIN1** | **S/T** | -0.09 | 0.28 | 0.19 | -0.49 | -0.03 | 0.03 | 0.217 |
| **SREBF1** | **S457** | 0.10 | -0.17 | -0.04 | -0.27 | -0.09 | 0.38 | 0.839 |
| **SREBF1** | **S455; S457** | 0.00 | 0.04 | -0.07 | 0.11 | -0.13 | 0.03 | 0.886 |
| **SREBF2** | **S117** | 0.11 | 0.06 | -0.24 | 0.08 | 0.09 | -0.15 | 0.818 |
| **SREBF2** | **S469** | 0.12 | 0.18 | -0.08 | -0.40 | 0.03 | 0.08 | 0.377 |
| **SREK1** | **S/T** | -0.13 | -0.15 | -0.01 | 0.25 | -0.01 | 0.02 | 0.112 |
| **SREK1** | **S/T** | -0.02 | 0.03 | -0.26 | -0.04 | -0.06 | 0.29 | 0.372 |
| **SREK1** | **S502** | -0.18 | 0.19 | -0.07 | 0.27 | -0.54 | 0.17 | 0.965 |
| **SREK1** | **S** | 0.13 | -0.08 | 0.08 | 0.01 | 0.06 | -0.23 | 0.415 |
| **SREK1** | **T503** | -0.06 | 0.20 | 0.18 | 0.41 | -1.50 | 0.52 | 0.673 |
| **SREK1** | **S203; S204** | 0.30 | 0.21 | 0.07 | -0.25 | -0.12 | -0.31 | 0.009 |
| **SREK1IP1** | **S149** | -0.09 | -0.02 | 0.00 | -0.06 | -0.05 | 0.20 | 0.51 |
| **SRFBP1** | **S367** | 0.07 | -0.82 | -0.21 | 0.17 | 0.01 | 0.47 | 0.144 |
| **SRFBP1** | **S203** | -0.08 | -0.25 | -0.32 | 0.03 | 0.10 | 0.41 | 0.044 |
| **SRFBP1** | **S247; S264** | 0.08 | -0.03 | -0.01 | 0.02 | -0.10 | 0.03 | 0.572 |
| **SRFBP1** | **S273; S279** | 0.05 | 0.05 | 0.06 | -0.08 | -0.11 | 0.01 | 0.035 |
| **SRFBP1** | **S203** | 0.16 | 0.14 | -0.10 | -0.11 | -0.18 | 0.05 | 0.233 |
| **SRFBP1** | **S247; S264** | 0.07 | 0.11 | 0.40 | -0.20 | -0.53 | -0.02 | 0.07 |
| **SRGAP2** | **S206** | 0.12 | 0.01 | 0.05 | -0.20 | -0.10 | 0.10 | 0.243 |
| **SRP72** | **S/T** | 0.15 | 0.15 | 0.18 | -0.18 | -0.10 | -0.27 | 0.002 |
| **SRPK1** | **S51** | -0.02 | 0.07 | -0.05 | 0.01 | 0.02 | -0.02 | 0.987 |
| **SRPK1** | **S309; S311** | -0.22 | -0.04 | 0.24 | -0.10 | -0.10 | 0.16 | 0.963 |
| **SRPK1** | **S311** | -0.03 | 0.10 | 0.13 | 0.02 | -0.15 | -0.10 | 0.106 |
| **SRPK1** | **S51** | 0.11 | -0.11 | 0.21 | -0.25 | -0.03 | 0.03 | 0.289 |
| **SRPK1** | **S33; S37; S51** | 0.22 | 0.14 | -0.08 | 0.07 | -0.19 | -0.20 | 0.193 |
| **SRPK1** | **S51** | 0.08 | 0.08 | 0.13 | -0.51 | 0.12 | 0.00 | 0.311 |
| **SRPK1** | **S37; S51** | 0.28 | 0.14 | -0.02 | -0.38 | -0.08 | -0.03 | 0.102 |
| **SRPK1** | **S311** | 0.13 | 0.12 | 0.18 | -0.14 | -0.02 | -0.33 | 0.032 |
| **SRPK2** | **S** | 0.25 | 0.25 | 0.14 | -0.26 | -0.07 | -0.45 | 0.015 |
| **SRPK2** | **S497** | 0.30 | 0.21 | 0.35 | -0.31 | -0.08 | -0.78 | 0.032 |
| **SRRM1** | **S738** | -0.60 | -0.45 | -0.32 | 0.20 | -0.25 | 0.86 | 0.093 |
| **SRRM1** | **S713** | -0.14 | -0.15 | -0.15 | 0.15 | -0.03 | 0.26 | 0.033 |
| **SRRM1** | **S414; S420** | -0.02 | -0.14 | -0.26 | 0.12 | 0.15 | 0.12 | 0.018 |
| **SRRM1** | **S636; S638** | -0.01 | -0.16 | -0.18 | 0.11 | -0.21 | 0.37 | 0.309 |
| **SRRM1** | **T872; S874** | 0.30 | -0.46 | -0.21 | 0.21 | 0.06 | -0.03 | 0.43 |
| **SRRM1** | **T574** | -0.17 | -0.14 | 0.00 | 0.24 | 0.00 | 0.04 | 0.095 |
| **SRRM1** | **S292** | -0.01 | -0.02 | -0.28 | 0.32 | 0.22 | -0.33 | 0.464 |
| **SRRM1** | **S260** | 0.01 | -0.10 | -0.16 | 0.09 | 0.01 | 0.13 | 0.058 |
| **SRRM1** | **S389** | -0.12 | -0.24 | 0.12 | 0.28 | -0.01 | -0.09 | 0.416 |
| **SRRM1** | **S478** | 0.02 | -0.11 | -0.12 | 0.10 | 0.15 | -0.07 | 0.183 |
| **SRRM1** | **S696** | -0.04 | -0.05 | -0.15 | 0.29 | -0.36 | 0.22 | 0.565 |

| | | | | | | | | |
|---|---|---|---|---|---|---|---|---|
| **SRRM1** | **S628** | 0.03 | -0.06 | -0.14 | 0.01 | 0.07 | 0.08 | 0.118 |
| **SRRM1** | **S463; S465** | -0.06 | -0.07 | -0.03 | 0.04 | -0.10 | 0.20 | 0.313 |
| **SRRM1** | **T572; T574** | -0.15 | -0.10 | 0.11 | 0.14 | -0.02 | -0.01 | 0.423 |
| **SRRM1** | **S675** | 0.13 | -0.05 | -0.21 | 0.13 | 0.05 | -0.07 | 0.522 |
| **SRRM1** | **S740** | 0.03 | -0.08 | -0.06 | 0.12 | -0.06 | 0.05 | 0.308 |
| **SRRM1** | **S** | 0.12 | -0.04 | -0.19 | 0.11 | 0.15 | -0.20 | 0.737 |
| **SRRM1** | **S294** | 0.07 | 0.01 | -0.15 | 0.09 | -0.02 | -0.01 | 0.63 |
| **SRRM1** | **S769; T/S** | 0.20 | -0.24 | -0.03 | 0.12 | -0.12 | 0.03 | 0.821 |
| **SRRM1** | **S769** | 0.16 | 0.03 | -0.26 | 0.09 | -0.01 | -0.04 | 0.805 |
| **SRRM1** | **S797** | 0.03 | -0.02 | -0.06 | -0.01 | 0.14 | -0.08 | 0.667 |
| **SRRM1** | **S795; S797** | -0.09 | -0.03 | 0.07 | 0.09 | -0.18 | 0.12 | 0.82 |
| **SRRM1** | **S234** | -0.03 | -0.05 | 0.02 | -0.10 | -0.08 | 0.21 | 0.821 |
| **SRRM1** | **S605** | 0.13 | -0.01 | -0.16 | 0.00 | -0.01 | 0.03 | 0.822 |
| **SRRM1** | **T872; S874** | 0.05 | -0.03 | -0.05 | 0.10 | 0.08 | -0.17 | 0.861 |
| **SRRM1** | **S551** | -0.03 | 0.09 | -0.08 | -0.01 | 0.13 | -0.11 | 0.862 |
| **SRRM1** | **S696; T/S** | 0.03 | -0.03 | -0.03 | 0.08 | -0.18 | 0.11 | 0.901 |
| **SRRM1** | **T572; T574** | -0.07 | -0.01 | 0.06 | 0.00 | -0.06 | 0.08 | 0.838 |
| **SRRM1** | **S549** | -0.26 | 0.03 | 0.20 | 0.06 | 0.02 | -0.09 | 0.975 |
| **SRRM1** | **S769; S773** | 0.44 | -0.27 | -0.24 | 0.03 | 0.00 | -0.09 | 0.989 |
| **SRRM1** | **S549; S551** | 0.07 | 0.04 | -0.11 | 0.08 | 0.00 | -0.08 | 0.967 |
| **SRRM1** | **S209** | 0.14 | 0.00 | -0.15 | -0.02 | 0.02 | -0.01 | 0.975 |
| **SRRM1** | **S626** | 0.11 | 0.00 | -0.11 | 0.13 | -0.02 | -0.13 | 0.951 |
| **SRRM1** | **S234** | -0.02 | -0.04 | 0.06 | 0.05 | -0.15 | 0.09 | 0.921 |
| **SRRM1** | **T614; S616** | 0.15 | 0.06 | -0.20 | 0.11 | -0.09 | -0.06 | 0.894 |
| **SRRM1** | **S636** | 0.12 | 0.11 | -0.22 | -0.06 | 0.11 | -0.10 | 0.875 |
| **SRRM1** | **S234** | 0.07 | 0.12 | -0.17 | -0.02 | 0.14 | -0.18 | 0.82 |
| **SRRM1** | **S463; S465** | 0.09 | 0.01 | -0.06 | 0.11 | -0.02 | -0.14 | 0.728 |
| **SRRM1** | **S583** | 0.00 | -0.04 | 0.10 | -0.07 | -0.02 | 0.03 | 0.503 |
| **SRRM1** | **S389; S393** | 0.05 | 0.01 | 0.01 | 0.06 | -0.02 | -0.12 | 0.436 |
| **SRRM1** | **S797** | 0.03 | -0.36 | 0.34 | -0.18 | 0.24 | -0.21 | 0.848 |
| **SRRM1** | **S738; S740** | 0.08 | 0.04 | -0.04 | -0.02 | -0.08 | 0.01 | 0.309 |
| **SRRM1** | **S626; S628** | 0.08 | 0.03 | -0.03 | 0.09 | -0.06 | -0.13 | 0.462 |
| **SRRM1** | **S696** | 0.07 | 0.02 | 0.00 | 0.08 | -0.11 | -0.07 | 0.366 |
| **SRRM1** | **S638** | 0.11 | 0.05 | -0.07 | 0.05 | 0.02 | -0.18 | 0.529 |
| **SRRM1** | **S683; S685** | 0.03 | 0.01 | 0.07 | 0.02 | -0.08 | -0.05 | 0.113 |
| **SRRM1** | **S478** | 0.08 | 0.00 | 0.01 | 0.09 | 0.07 | -0.28 | 0.579 |
| **SRRM1** | **S607** | 0.03 | 0.06 | 0.03 | 0.02 | 0.06 | -0.22 | 0.396 |
| **SRRM1** | **S636; S638** | 0.08 | 0.03 | 0.01 | 0.07 | -0.11 | -0.10 | 0.243 |
| **SRRM1** | **S549; S551** | 0.03 | 0.00 | 0.10 | -0.03 | -0.05 | -0.06 | 0.041 |
| **SRRM1** | **T614; S616** | 0.05 | 0.03 | 0.06 | -0.03 | 0.02 | -0.13 | 0.128 |
| **SRRM1** | **S597** | 0.06 | 0.05 | 0.02 | 0.03 | 0.01 | -0.19 | 0.26 |
| **SRRM1** | **S874** | 0.11 | 0.01 | 0.01 | 0.04 | -0.17 | -0.01 | 0.255 |
| **SRRM1** | **S769; S773** | 0.18 | 0.08 | -0.14 | 0.08 | -0.23 | -0.02 | 0.509 |
| **SRRM1** | **S626; S628** | 0.12 | 0.05 | -0.02 | 0.07 | -0.10 | -0.14 | 0.26 |
| **SRRM1** | **S560; S562** | 0.08 | 0.06 | 0.01 | 0.02 | -0.01 | -0.17 | 0.181 |
| **SRRM1** | **S605; S607** | 0.09 | 0.07 | 0.00 | 0.03 | -0.08 | -0.12 | 0.112 |
| **SRRM1** | **S209; S211** | 0.10 | 0.05 | 0.00 | -0.06 | 0.03 | -0.14 | 0.13 |
| **SRRM1** | **S605; S607** | 0.14 | 0.08 | -0.06 | 0.05 | -0.09 | -0.14 | 0.224 |

| | | | | | | | | |
|---|---|---|---|---|---|---|---|---|
| SRRM1 | S562 | 0.05 | 0.10 | 0.02 | -0.03 | -0.01 | -0.14 | 0.069 |
| SRRM1 | S354; S | -0.02 | 0.28 | -0.16 | 0.15 | 0.15 | -0.54 | 0.681 |
| SRRM1 | S675 | 0.16 | 0.00 | 0.00 | -0.02 | 0.04 | -0.21 | 0.271 |
| SRRM1 | S628 | 0.03 | 0.10 | 0.04 | 0.05 | 0.05 | -0.30 | 0.37 |
| SRRM1 | S743; S | 0.08 | 0.07 | 0.03 | 0.03 | -0.11 | -0.11 | 0.061 |
| SRRM1 | T220 | 0.23 | 0.09 | -0.18 | 0.19 | -0.06 | -0.36 | 0.572 |
| SRRM1 | T581; S583 | 0.03 | 0.07 | 0.08 | 0.00 | -0.09 | -0.11 | 0.03 |
| SRRM1 | S560; S562 | 0.06 | 0.05 | 0.08 | 0.01 | -0.10 | -0.12 | 0.029 |
| SRRM1 | S463 | 0.15 | 0.08 | -0.03 | -0.01 | -0.14 | -0.07 | 0.085 |
| SRRM1 | S414 | 0.17 | 0.06 | -0.03 | -0.03 | -0.11 | -0.09 | 0.087 |
| SRRM1 | S549; S551 | 0.06 | 0.06 | 0.08 | 0.04 | -0.03 | -0.24 | 0.169 |
| SRRM1 | S560; S562 | 0.09 | 0.10 | 0.02 | 0.02 | -0.05 | -0.19 | 0.096 |
| SRRM1 | S713; S715 | 0.04 | 0.06 | 0.10 | -0.01 | -0.05 | -0.16 | 0.039 |
| SRRM1 | T220 | 0.11 | 0.18 | -0.07 | -0.03 | -0.06 | -0.17 | 0.139 |
| SRRM1 | S626; S628 | 0.11 | 0.10 | 0.03 | 0.00 | -0.06 | -0.20 | 0.068 |
| SRRM1 | T220 | 0.14 | 0.11 | -0.01 | -0.01 | -0.04 | -0.22 | 0.112 |
| SRRM1 | S653; S655 | 0.11 | 0.08 | 0.05 | 0.01 | -0.04 | -0.23 | 0.092 |
| SRRM1 | S675 | 0.16 | -0.26 | 0.31 | 0.11 | -0.02 | -0.43 | 0.479 |
| SRRM1 | S562 | 0.00 | 0.11 | 0.16 | -0.13 | 0.00 | -0.16 | 0.055 |
| SRRM1 | T572; T574 | 0.10 | 0.10 | 0.06 | -0.01 | -0.05 | -0.23 | 0.053 |
| SRRM1 | S738 | 0.21 | 0.13 | -0.07 | -0.21 | -0.03 | -0.06 | 0.136 |
| SRRM1 | S/T | -0.12 | 0.36 | -0.02 | -0.23 | 0.30 | -0.45 | 0.502 |
| SRRM1 | S791; S797 | 0.09 | 0.06 | 0.13 | 0.02 | -0.15 | -0.18 | 0.039 |
| SRRM1 | S450; S452 | 0.08 | 0.08 | 0.12 | -0.02 | -0.05 | -0.25 | 0.048 |
| SRRM1 | S715 | 0.07 | 0.12 | 0.10 | -0.16 | 0.08 | -0.26 | 0.114 |
| SRRM1 | S725; T727 | 0.17 | 0.12 | 0.01 | -0.05 | -0.05 | -0.22 | 0.047 |
| SRRM1 | T581; S583 | -0.01 | 0.12 | 0.19 | -0.17 | -0.10 | -0.06 | 0.039 |
| SRRM1 | T614; S616 | 0.11 | 0.14 | 0.06 | -0.07 | -0.07 | -0.20 | 0.014 |
| SRRM1 | S211 | 0.23 | 0.12 | -0.04 | -0.13 | 0.07 | -0.30 | 0.174 |
| SRRM1 | S616 | 0.09 | -0.04 | 0.26 | -0.04 | 0.01 | -0.34 | 0.181 |
| SRRM1 | S874 | 0.14 | 0.11 | 0.07 | -0.04 | -0.03 | -0.29 | 0.062 |
| SRRM1 | S463; S465 | 0.05 | 0.18 | 0.09 | -0.11 | 0.01 | -0.27 | 0.058 |
| SRRM1 | S389; S391 | -0.01 | 0.04 | 0.30 | -0.05 | -0.04 | -0.31 | 0.131 |
| SRRM1 | T614; S616 | -0.16 | 0.30 | 0.17 | -0.09 | -0.48 | 0.13 | 0.336 |
| SRRM1 | S743; S | 0.11 | 0.13 | 0.11 | 0.06 | -0.28 | -0.19 | 0.069 |
| SRRM1 | S802; S/T | -0.16 | 0.33 | 0.20 | -0.28 | 0.15 | -0.37 | 0.259 |
| SRRM1 | S560 | 0.16 | 0.08 | 0.17 | -0.07 | -0.01 | -0.41 | 0.081 |
| SRRM1 | S414 | 0.14 | 0.17 | 0.13 | -0.06 | -0.07 | -0.38 | 0.037 |
| SRRM1 | S429; S431 | 0.12 | 0.12 | 0.21 | -0.14 | 0.00 | -0.40 | 0.052 |
| SRRM1 | S393 | 0.15 | 0.13 | 0.16 | -0.13 | 0.09 | -0.52 | 0.135 |
| SRRM1 | S616 | 0.14 | 0.14 | 0.18 | -0.17 | 0.03 | -0.41 | 0.059 |
| SRRM1 | S463 | 0.24 | 0.14 | 0.10 | -0.05 | -0.11 | -0.41 | 0.042 |
| SRRM1 | T614; S616 | -0.05 | 0.12 | 0.41 | -0.28 | -0.12 | -0.19 | 0.067 |
| SRRM1 | S260 | 0.39 | 0.38 | -0.44 | -0.13 | 0.26 | -0.90 | 0.45 |
| SRRM1 | S402; T406 | 0.19 | 0.16 | 0.20 | -0.15 | -0.06 | -0.45 | 0.028 |
| SRRM1 | S797; S802 | 0.18 | 0.29 | 0.07 | -0.18 | 0.02 | -0.51 | 0.075 |
| SRRM1 | S791 | 0.15 | 0.17 | 0.23 | 0.02 | -0.29 | -0.40 | 0.033 |
| SRRM1 | S756 | 0.23 | 0.22 | 0.11 | -0.22 | 0.00 | -0.47 | 0.042 |

| | | | | | | | | |
|---|---|---|---|---|---|---|---|---|
| **SRRM1** | **T727** | 0.19 | 0.22 | 0.15 | -0.19 | 0.09 | -0.64 | 0.116 |
| **SRRM1** | **T406** | 0.18 | 0.22 | 0.19 | -0.27 | 0.05 | -0.51 | 0.052 |
| **SRRM1** | **S452** | 0.18 | 0.26 | 0.18 | -0.18 | 0.03 | -0.64 | 0.078 |
| **SRRM1** | **S685** | 0.21 | 0.23 | 0.19 | -0.26 | 0.04 | -0.59 | 0.057 |
| **SRRM1** | **S414** | 0.20 | 0.17 | 0.29 | -0.28 | -0.10 | -0.42 | 0.008 |
| **SRRM1** | **S754; S756** | 0.29 | 0.21 | 0.14 | -0.14 | -0.06 | -0.63 | 0.055 |
| **SRRM1** | **S795; S797; S802** | 0.20 | 0.43 | 0.03 | -0.19 | 0.05 | -0.81 | 0.13 |
| **SRRM1** | **S463** | 0.28 | 0.22 | 0.22 | -0.23 | -0.06 | -0.61 | 0.031 |
| **SRRM1** | **S713; S715** | 0.21 | 0.21 | 0.29 | -0.23 | 0.00 | -0.70 | 0.057 |
| **SRRM1** | **S429** | 0.12 | 0.23 | 0.38 | -0.24 | -0.09 | -0.62 | 0.033 |
| **SRRM1** | **S211** | 0.25 | 0.44 | 0.05 | -0.39 | 0.01 | -0.62 | 0.056 |
| **SRRM1** | **S** | 0.30 | 0.28 | 0.18 | -0.24 | -0.06 | -0.71 | 0.041 |
| **SRRM1** | **S** | 0.30 | 0.34 | 0.32 | -0.39 | -0.17 | -0.72 | 0.01 |
| **SRRM1** | **S187** | 0.30 | 0.29 | 0.37 | -0.24 | 0.00 | -1.30 | 0.105 |
| **SRRM1** | **S450** | 0.34 | 0.35 | 0.31 | -0.32 | -0.06 | -1.12 | 0.059 |
| **SRRM1** | **S616** | 0.24 | 0.32 | 0.51 | -0.50 | 0.05 | -1.31 | 0.08 |
| **SRRM2** | **S1368** | -0.58 | -0.80 | -0.73 | 0.25 | 0.53 | 0.60 | 7E-04 |
| **SRRM2** | **S1336** | -0.01 | -0.69 | -0.84 | -0.42 | 0.72 | 0.51 | 0.146 |
| **SRRM2** | **T1413; S** | -0.10 | 0.12 | -1.27 | 0.19 | 0.64 | -0.21 | 0.278 |
| **SRRM2** | **S1257** | -0.82 | -0.08 | -0.24 | 0.63 | 0.30 | -0.22 | 0.139 |
| **SRRM2** | **S1233** | -0.34 | -0.29 | -0.34 | 0.35 | 0.39 | 0.03 | 0.007 |
| **SRRM2** | **S1232; S1233** | -0.25 | -0.43 | -0.23 | 0.21 | 0.11 | 0.42 | 0.008 |
| **SRRM2** | **S1232; S1233** | -0.25 | -0.37 | -0.29 | 0.33 | -0.01 | 0.40 | 0.013 |
| **SRRM2** | **S778; S780; S** | -0.13 | -0.28 | -0.18 | 0.17 | 0.02 | 0.31 | 0.019 |
| **SRRM2** | **S1233** | -0.22 | -0.14 | -0.21 | 0.16 | 0.22 | 0.12 | 6E-04 |
| **SRRM2** | **S970; S974; S/T/Y** | -0.40 | -0.32 | 0.10 | 0.19 | -0.11 | 0.38 | 0.16 |
| **SRRM2** | **S2046** | -0.08 | -0.07 | -0.39 | 0.17 | 0.22 | 0.07 | 0.042 |
| **SRRM2** | **S1336** | -0.26 | -0.22 | -0.06 | -0.02 | 0.38 | 0.08 | 0.07 |
| **SRRM2** | **S1179** | -0.18 | -0.20 | -0.16 | 0.31 | -0.27 | 0.36 | 0.198 |
| **SRRM2** | **T/S** | -0.09 | -0.29 | -0.06 | 0.10 | 0.18 | 0.11 | 0.024 |
| **SRRM2** | **S1179** | -0.43 | 0.27 | -0.37 | 0.08 | -0.25 | 0.47 | 0.425 |
| **SRRM2** | **T2104** | -0.08 | -0.10 | -0.24 | 0.32 | -0.09 | 0.12 | 0.114 |
| **SRRM2** | **T2289** | -0.12 | -0.42 | 0.08 | 0.46 | -0.18 | 0.02 | 0.339 |
| **SRRM2** | **S1326** | -0.33 | 0.06 | -0.10 | 0.22 | -0.10 | 0.18 | 0.222 |
| **SRRM2** | **S295; S297** | -0.08 | -0.09 | -0.12 | 0.11 | -0.02 | 0.17 | 0.035 |
| **SRRM2** | **S486** | -0.04 | -0.04 | -0.25 | 0.07 | 0.37 | -0.20 | 0.365 |
| **SRRM2** | **S424; S/T** | -0.04 | -0.03 | -0.17 | 0.01 | -0.12 | 0.30 | 0.333 |
| **SRRM2** | **S1083** | -0.11 | -0.10 | -0.04 | 0.04 | -0.17 | 0.31 | 0.369 |
| **SRRM2** | **S1081; S1083** | -0.06 | -0.07 | -0.09 | 0.09 | 0.01 | 0.10 | 0.012 |
| **SRRM2** | **S2132** | -0.03 | -0.01 | -0.17 | 0.22 | 0.00 | -0.04 | 0.258 |
| **SRRM2** | **S1398; S** | -0.21 | -0.12 | 0.13 | 0.13 | 0.10 | -0.07 | 0.354 |
| **SRRM2** | **T367** | -0.08 | -0.07 | -0.05 | -0.03 | 0.03 | 0.17 | 0.105 |
| **SRRM2** | **S1387** | -0.08 | -0.06 | -0.06 | 0.25 | -0.34 | 0.21 | 0.607 |
| **SRRM2** | **T866** | -0.11 | -0.04 | -0.04 | -0.12 | -0.13 | 0.37 | 0.573 |
| **SRRM2** | **S440; S449** | -0.14 | 0.01 | -0.09 | 0.07 | -0.44 | 0.45 | 0.733 |
| **SRRM2** | **S322; S323** | 0.00 | 0.00 | -0.15 | 0.11 | -0.06 | 0.08 | 0.277 |
| **SRRM2** | **S2132** | 0.08 | -0.11 | -0.11 | 0.10 | 0.02 | 0.00 | 0.29 |
| **SRRM2** | **S1404** | -0.06 | 0.01 | -0.07 | 0.07 | -0.05 | 0.10 | 0.197 |

| | | | | | | | | |
|---|---|---|---|---|---|---|---|---|
| **SRRM2** | **S/Y/T** | -0.08 | -0.06 | 0.00 | -0.13 | 0.28 | -0.05 | 0.557 |
| **SRRM2** | **S2044; S2046** | -0.02 | -0.16 | 0.06 | 0.04 | 0.03 | 0.04 | 0.275 |
| **SRRM2** | **S1501** | -0.12 | 0.02 | -0.02 | 0.08 | 0.02 | 0.01 | 0.196 |
| **SRRM2** | **S1403** | -0.02 | 0.03 | -0.12 | 0.14 | -0.01 | -0.04 | 0.408 |
| **SRRM2** | **S1083** | 0.01 | 0.00 | -0.11 | 0.12 | -0.05 | 0.01 | 0.378 |
| **SRRM2** | **T1003** | -0.07 | 0.00 | -0.02 | 0.13 | -0.04 | -0.02 | 0.404 |
| **SRRM2** | **T2104** | 0.06 | 0.05 | -0.20 | 0.23 | -0.10 | -0.07 | 0.726 |
| **SRRM2** | **S1818** | 0.05 | 0.05 | -0.19 | 0.14 | -0.07 | -0.01 | 0.644 |
| **SRRM2** | **S871; S875; S876** | -0.05 | -0.11 | 0.08 | 0.05 | -0.10 | 0.11 | 0.612 |
| **SRRM2** | **T1413; S1415** | -0.15 | 0.08 | 0.01 | -0.15 | -0.03 | 0.21 | 0.793 |
| **SRRM2** | **T1492** | -0.08 | -0.07 | 0.08 | 0.22 | -0.28 | 0.08 | 0.85 |
| **SRRM2** | **T315; S323** | 0.05 | 0.00 | -0.11 | 0.06 | -0.05 | 0.04 | 0.623 |
| **SRRM2** | **T2022** | -0.01 | 0.05 | -0.10 | 0.11 | 0.06 | -0.13 | 0.73 |
| **SRRM2** | **S1101; S1103** | 0.01 | -0.01 | -0.05 | 0.03 | -0.05 | 0.07 | 0.491 |
| **SRRM2** | **S1497; S1499; S1501** | -0.21 | 0.04 | 0.11 | -0.02 | -0.13 | 0.17 | 0.855 |
| **SRRM2** | **S323** | 0.08 | 0.07 | -0.19 | -0.01 | 0.04 | -0.02 | 0.857 |
| **SRRM2** | **S875; S876** | 0.01 | 0.00 | -0.02 | 0.07 | 0.00 | -0.05 | 0.806 |
| **SRRM2** | **S357; S358; T359** | 0.10 | 0.06 | -0.21 | 0.02 | -0.34 | 0.28 | 0.986 |
| **SRRM2** | **S954** | -0.10 | 0.10 | -0.01 | -0.05 | 0.01 | 0.04 | 0.963 |
| **SRRM2** | **T1472** | 0.03 | 0.08 | -0.12 | 0.11 | -0.03 | -0.09 | 0.982 |
| **SRRM2** | **T/S** | -0.13 | 0.04 | 0.03 | 0.28 | 0.13 | -0.45 | 0.999 |
| **SRRM2** | **S510** | -0.11 | 0.11 | 0.00 | -0.04 | -0.02 | 0.05 | 0.994 |
| **SRRM2** | **S1124** | -0.09 | -0.14 | 0.18 | 0.39 | -0.20 | -0.25 | 0.997 |
| **SRRM2** | **S377; S395; T/S** | -0.08 | -0.05 | 0.13 | 0.04 | -0.04 | -0.02 | 0.949 |
| **SRRM2** | **S/T** | 0.09 | -0.09 | -0.02 | -0.17 | -0.13 | 0.27 | 0.974 |
| **SRRM2** | **S1101; S1103** | -0.06 | -0.11 | 0.16 | 0.01 | -0.15 | 0.12 | 0.963 |
| **SRRM2** | **S1762; S1764** | 0.07 | 0.05 | -0.11 | 0.12 | -0.06 | -0.08 | 0.938 |
| **SRRM2** | **S1443** | -0.04 | 0.06 | 0.00 | 0.02 | -0.11 | 0.07 | 0.83 |
| **SRRM2** | **S1869** | 0.02 | 0.03 | -0.03 | 0.15 | -0.16 | -0.04 | 0.827 |
| **SRRM2** | **S857; T866** | -0.03 | 0.04 | 0.03 | 0.01 | 0.00 | -0.05 | 0.452 |
| **SRRM2** | **S2684; S2688** | 0.05 | 0.09 | -0.11 | -0.09 | 0.05 | -0.01 | 0.736 |
| **SRRM2** | **T1003; S1014** | 0.10 | 0.05 | -0.11 | 0.13 | -0.08 | -0.10 | 0.777 |
| **SRRM2** | **S1691; S/T** | 0.17 | -0.05 | -0.10 | 0.26 | -0.23 | -0.10 | 0.856 |
| **SRRM2** | **T2599** | 0.22 | -0.24 | 0.06 | -0.02 | 0.01 | -0.07 | 0.796 |
| **SRRM2** | **S994** | 0.01 | 0.00 | 0.04 | 0.21 | -0.12 | -0.17 | 0.735 |
| **SRRM2** | **S1329** | 0.00 | 0.01 | 0.05 | 0.06 | -0.18 | 0.05 | 0.584 |
| **SRRM2** | **S2102; T2104** | -0.17 | 0.03 | 0.19 | 0.11 | -0.14 | -0.06 | 0.731 |
| **SRRM2** | **T866; T/S** | -0.20 | 0.07 | 0.16 | 0.17 | -0.37 | 0.09 | 0.822 |
| **SRRM2** | **T1003** | 0.09 | 0.05 | -0.06 | 0.08 | -0.10 | -0.06 | 0.524 |
| **SRRM2** | **S333** | 0.02 | -0.02 | 0.07 | -0.01 | -0.09 | 0.02 | 0.27 |
| **SRRM2** | **T383; S/T** | -0.21 | 0.00 | 0.26 | -0.16 | -0.10 | 0.15 | 0.769 |
| **SRRM2** | **S1124** | 0.02 | -0.02 | 0.06 | 0.24 | -0.16 | -0.18 | 0.721 |
| **SRRM2** | **S/T** | 0.11 | 0.11 | -0.15 | 0.05 | 0.01 | -0.16 | 0.634 |
| **SRRM2** | **T1962** | -0.13 | 0.17 | 0.04 | -0.04 | 0.05 | -0.10 | 0.588 |
| **SRRM2** | **S952; S954** | -0.04 | -0.04 | 0.17 | -0.01 | 0.00 | -0.09 | 0.455 |
| **SRRM2** | **S456; S/T** | 0.07 | 0.07 | -0.05 | 0.07 | 0.00 | -0.18 | 0.474 |
| **SRRM2** | **S992; S/Y** | -0.15 | 0.04 | 0.20 | -0.04 | -0.07 | -0.01 | 0.56 |
| **SRRM2** | **T1880** | 0.02 | 0.06 | 0.03 | 0.02 | -0.09 | -0.04 | 0.1 |

| | | | | | | | | |
|---|---|---|---|---|---|---|---|---|
| SRRM2 | S404 | 0.10 | 0.14 | -0.18 | 0.08 | 0.16 | -0.39 | 0.745 |
| SRRM2 | S2132 | -0.07 | -0.06 | 0.23 | 0.14 | -0.15 | -0.13 | 0.587 |
| SRRM2 | T866 | -0.20 | 0.11 | 0.18 | -0.10 | -0.25 | 0.20 | 0.664 |
| SRRM2 | S440 | 0.01 | 0.10 | 0.01 | -0.07 | -0.12 | 0.06 | 0.226 |
| SRRM2 | S1401; S | -0.13 | 0.08 | 0.15 | 0.11 | -0.33 | 0.06 | 0.628 |
| SRRM2 | S2581 | 0.16 | 0.03 | -0.07 | 0.03 | -0.12 | -0.05 | 0.327 |
| SRRM2 | S1657; S1658 | 0.05 | 0.11 | -0.02 | -0.01 | -0.02 | -0.11 | 0.149 |
| SRRM2 | S2407; T2409 | 0.00 | -0.07 | 0.20 | -0.05 | -0.11 | 0.01 | 0.337 |
| SRRM2 | T1434 | 0.03 | 0.05 | 0.06 | -0.01 | 0.00 | -0.13 | 0.083 |
| SRRM2 | T400 | 0.37 | -0.05 | -0.21 | -0.23 | 0.03 | 0.01 | 0.634 |
| SRRM2 | S1061 | 0.01 | 0.18 | -0.06 | 0.05 | 0.00 | -0.22 | 0.407 |
| SRRM2 | S1112 | -0.07 | 0.13 | 0.07 | -0.34 | 0.14 | 0.02 | 0.534 |
| SRRM2 | S1014 | 0.17 | 0.05 | -0.07 | 0.01 | -0.12 | -0.06 | 0.242 |
| SRRM2 | S536 | 0.08 | 0.14 | -0.08 | 0.19 | -0.10 | -0.29 | 0.513 |
| SRRM2 | S322; S323 | 0.01 | 0.15 | 0.00 | 0.00 | -0.04 | -0.13 | 0.147 |
| SRRM2 | S333 | 0.19 | 0.11 | -0.15 | 0.06 | -0.07 | -0.18 | 0.41 |
| SRRM2 | S | -0.12 | 0.01 | 0.27 | 0.02 | -0.14 | -0.09 | 0.394 |
| SRRM2 | T1003; S1010 | 0.18 | -0.15 | 0.12 | -0.28 | 0.06 | 0.02 | 0.464 |
| SRRM2 | S333 | 0.17 | 0.12 | -0.13 | 0.04 | 0.01 | -0.26 | 0.426 |
| SRRM2 | S1179 | 0.05 | 0.07 | 0.05 | 0.02 | 0.00 | -0.21 | 0.173 |
| SRRM2 | S1179 | 0.08 | 0.08 | 0.01 | -0.04 | 0.05 | -0.21 | 0.199 |
| SRRM2 | S1320 | 0.07 | 0.09 | 0.03 | -0.06 | 0.05 | -0.20 | 0.161 |
| SRRM2 | S838; S839 | 0.13 | 0.05 | 0.01 | 0.03 | -0.19 | -0.04 | 0.146 |
| SRRM2 | S2702; S2706 | 0.09 | 0.03 | 0.07 | -0.05 | -0.19 | 0.03 | 0.108 |
| SRRM2 | T1208 | 0.05 | 0.10 | 0.04 | -0.07 | 0.05 | -0.19 | 0.133 |
| SRRM2 | S456 | 0.15 | 0.11 | -0.06 | -0.01 | 0.07 | -0.29 | 0.334 |
| SRRM2 | S1501 | 0.09 | 0.16 | -0.05 | -0.04 | 0.02 | -0.21 | 0.191 |
| SRRM2 | S424 | 0.17 | 0.15 | -0.12 | -0.07 | -0.24 | 0.07 | 0.325 |
| SRRM2 | S2100; S2102; T2104 | 0.05 | -0.03 | 0.17 | -0.30 | 0.05 | 0.01 | 0.312 |
| SRRM2 | T383; T400 | -0.18 | -0.16 | 0.49 | 0.11 | -0.28 | -0.13 | 0.585 |
| SRRM2 | S424 | 0.17 | 0.09 | -0.05 | -0.11 | -0.15 | 0.03 | 0.156 |
| SRRM2 | T2409 | 0.12 | 0.04 | 0.06 | -0.05 | -0.06 | -0.12 | 0.008 |
| SRRM2 | T2409 | 0.24 | 0.19 | -0.23 | -0.03 | -0.16 | -0.08 | 0.366 |
| SRRM2 | S1497 | -0.04 | 0.09 | 0.17 | -0.03 | 0.03 | -0.25 | 0.2 |
| SRRM2 | S1866; S1869 | 0.22 | 0.08 | -0.10 | 0.16 | -0.12 | -0.31 | 0.388 |
| SRRM2 | T1413; S | 0.16 | 0.08 | -0.01 | 0.09 | -0.22 | -0.13 | 0.184 |
| SRRM2 | S435; S436; S440 | -0.09 | 0.22 | 0.11 | -0.13 | -0.11 | -0.04 | 0.149 |
| SRRM2 | S/T/Y | 0.17 | -0.02 | 0.08 | -0.01 | 0.17 | -0.48 | 0.419 |
| SRRM2 | S1099; S1101; S1102 | 0.16 | 0.06 | 0.05 | -0.01 | -0.03 | -0.27 | 0.103 |
| SRRM2 | S1188 | 0.25 | 0.21 | -0.28 | 0.15 | 0.11 | -0.65 | 0.57 |
| SRRM2 | T848; S854; S857 | 0.00 | 0.09 | 0.16 | -0.07 | 0.15 | -0.41 | 0.316 |
| SRRM2 | S1727; S1729 | 0.11 | 0.13 | 0.04 | -0.04 | -0.02 | -0.25 | 0.066 |
| SRRM2 | S1462; S1463 | -0.34 | 0.16 | 0.39 | -0.21 | -0.38 | 0.20 | 0.514 |
| SRRM2 | S818 | 0.09 | 0.09 | 0.09 | -0.02 | 0.06 | -0.37 | 0.202 |
| SRRM2 | S2702; S2706 | 0.17 | 0.06 | 0.06 | -0.05 | -0.11 | -0.15 | 0.012 |
| SRRM2 | S1618; S1620; S1621 | 0.08 | 0.13 | 0.05 | 0.01 | 0.14 | -0.50 | 0.366 |
| SRRM2 | T2069; S2071 | 0.09 | 0.14 | 0.05 | -0.08 | 0.02 | -0.27 | 0.079 |
| SRRM2 | S | -0.12 | 0.32 | 0.06 | -0.12 | 0.09 | -0.32 | 0.303 |

| | | | | | | | | |
|---|---|---|---|---|---|---|---|---|
| **SRRM2** | **S1854; S1857** | 0.14 | 0.14 | 0.01 | 0.00 | -0.04 | -0.29 | 0.11 |
| **SRRM2** | **S1818; S1822** | 0.03 | 0.16 | 0.10 | 0.03 | -0.01 | -0.36 | 0.184 |
| **SRRM2** | **S1441; S1443** | 0.16 | -0.15 | 0.26 | -0.30 | -0.11 | 0.06 | 0.259 |
| **SRRM2** | **S895** | 0.06 | 0.11 | 0.13 | -0.02 | -0.10 | -0.21 | 0.021 |
| **SRRM2** | **S231; T233** | 0.12 | 0.08 | 0.10 | -0.11 | -0.05 | -0.17 | 0.005 |
| **SRRM2** | **S974; S/Y/T** | 0.11 | 0.11 | 0.08 | 0.00 | -0.15 | -0.19 | 0.021 |
| **SRRM2** | **T856; S857** | 0.09 | 0.09 | 0.13 | -0.10 | 0.03 | -0.28 | 0.073 |
| **SRRM2** | **T111** | 0.14 | 0.07 | 0.11 | -0.04 | -0.03 | -0.30 | 0.067 |
| **SRRM2** | **S854; T856; T866** | 0.06 | 0.06 | 0.20 | 0.02 | -0.08 | -0.30 | 0.1 |
| **SRRM2** | **S472** | 0.23 | 0.18 | -0.09 | 0.03 | -0.08 | -0.35 | 0.194 |
| **SRRM2** | **S424; T/S** | -0.07 | 0.13 | 0.23 | 0.21 | -0.19 | -0.43 | 0.321 |
| **SRRM2** | **S1320** | 0.17 | 0.13 | 0.03 | -0.04 | -0.11 | -0.22 | 0.025 |
| **SRRM2** | **S/T** | 0.09 | 0.17 | 0.07 | -0.06 | -0.05 | -0.27 | 0.038 |
| **SRRM2** | **T983** | 0.16 | 0.17 | 0.00 | -0.14 | -0.01 | -0.24 | 0.05 |
| **SRRM2** | **T1531** | 0.09 | 0.11 | 0.13 | -0.10 | 0.05 | -0.34 | 0.103 |
| **SRRM2** | **S2100; S2102; T2104** | 0.05 | 0.09 | 0.20 | -0.07 | -0.05 | -0.27 | 0.043 |
| **SRRM2** | **S2020; T2022** | 0.12 | 0.15 | 0.08 | -0.12 | 0.03 | -0.31 | 0.07 |
| **SRRM2** | **S931** | 0.11 | 0.36 | -0.14 | -0.19 | -0.40 | 0.13 | 0.291 |
| **SRRM2** | **S950; S952; S954** | 0.02 | 0.07 | 0.24 | -0.51 | 0.15 | -0.08 | 0.271 |
| **SRRM2** | **S2343** | -0.03 | 0.08 | 0.12 | 0.29 | -1.16 | 0.26 | 0.617 |
| **SRRM2** | **S778; S780** | 0.02 | 0.11 | 0.23 | -0.03 | -0.30 | -0.09 | 0.063 |
| **SRRM2** | **S1103; S/T** | 0.08 | 0.16 | 0.13 | -0.09 | -0.03 | -0.29 | 0.033 |
| **SRRM2** | **S404; S408** | 0.19 | 0.11 | 0.07 | -0.10 | -0.26 | -0.05 | 0.023 |
| **SRRM2** | **S1478** | 0.09 | 0.19 | 0.10 | -0.12 | -0.09 | -0.21 | 0.005 |
| **SRRM2** | **T1844** | 0.04 | 0.22 | 0.12 | -0.18 | -0.05 | -0.21 | 0.018 |
| **SRRM2** | **S2132** | 0.11 | 0.17 | 0.10 | 0.03 | -0.08 | -0.39 | 0.104 |
| **SRRM2** | **S2449** | 0.23 | 0.15 | 0.00 | -0.03 | -0.19 | -0.22 | 0.036 |
| **SRRM2** | **S2382** | 0.32 | 0.18 | -0.16 | -0.05 | 0.07 | -0.50 | 0.291 |
| **SRRM2** | **S248; S250** | 0.10 | 0.12 | 0.19 | -0.15 | -0.07 | -0.22 | 0.005 |
| **SRRM2** | **S1731** | 0.10 | 0.17 | 0.12 | -0.11 | 0.01 | -0.36 | 0.065 |
| **SRRM2** | **S2702** | 0.25 | 0.07 | 0.09 | -0.15 | -0.10 | -0.19 | 0.01 |
| **SRRM2** | **S952; S954** | 0.26 | 0.16 | -0.03 | 0.02 | -0.14 | -0.35 | 0.106 |
| **SRRM2** | **S1497; S1499** | 0.14 | 0.18 | 0.08 | -0.13 | -0.09 | -0.24 | 0.006 |
| **SRRM2** | **S1320** | 0.11 | 0.25 | -0.05 | -0.37 | 0.43 | -0.62 | 0.425 |
| **SRRM2** | **S790; T792** | 0.17 | 0.11 | 0.13 | -0.03 | -0.07 | -0.37 | 0.055 |
| **SRRM2** | **S1691; S/T** | 0.10 | 0.12 | 0.20 | -0.09 | -0.07 | -0.31 | 0.026 |
| **SRRM2** | **S351; S353** | 0.01 | 0.20 | 0.21 | -0.16 | -0.10 | -0.21 | 0.014 |
| **SRRM2** | **S1102** | 0.08 | 0.16 | 0.17 | -0.14 | 0.03 | -0.37 | 0.067 |
| **SRRM2** | **T792** | 0.23 | 0.19 | -0.02 | -0.25 | 0.14 | -0.39 | 0.168 |
| **SRRM2** | **S353; S/T** | 0.07 | 0.12 | 0.23 | -0.17 | 0.00 | -0.32 | 0.046 |
| **SRRM2** | **S1672; S** | 0.14 | 0.21 | 0.06 | -0.16 | 0.05 | -0.38 | 0.079 |
| **SRRM2** | **S778; S780** | 0.19 | 0.16 | 0.08 | -0.07 | -0.14 | -0.28 | 0.011 |
| **SRRM2** | **S1762; S1764** | 0.16 | 0.16 | 0.12 | -0.03 | -0.21 | -0.25 | 0.012 |
| **SRRM2** | **S377; S398** | 0.04 | 0.18 | 0.22 | -0.19 | -0.10 | -0.21 | 0.009 |
| **SRRM2** | **S871** | 0.15 | 0.15 | 0.14 | -0.08 | -0.04 | -0.42 | 0.054 |
| **SRRM2** | **S745** | 0.14 | 0.17 | 0.13 | -0.11 | 0.10 | -0.55 | 0.155 |
| **SRRM2** | **S/T** | 0.00 | 0.28 | 0.20 | -0.28 | 0.00 | -0.29 | 0.052 |
| **SRRM2** | **T848; S854** | 0.19 | -0.04 | 0.31 | -0.01 | -0.11 | -0.47 | 0.116 |

| | | | | | | | | |
|---|---|---|---|---|---|---|---|---|
| **SRRM2** | **S377** | 0.15 | 0.17 | 0.16 | -0.40 | -0.17 | 0.01 | 0.042 |
| **SRRM2** | **S837** | 0.08 | 0.21 | 0.19 | -0.15 | -0.43 | 0.00 | 0.056 |
| **SRRM2** | **S1727; S1729; S1731** | 0.13 | 0.15 | 0.22 | -0.24 | -0.04 | -0.29 | 0.011 |
| **SRRM2** | **S1960; T1962** | 0.09 | 0.20 | 0.20 | -0.18 | -0.01 | -0.40 | 0.041 |
| **SRRM2** | **S2280** | 0.01 | 0.04 | 0.41 | -0.51 | -0.13 | 0.03 | 0.152 |
| **SRRM2** | **S1443; S1444** | 0.14 | 0.18 | 0.18 | -0.13 | -0.05 | -0.41 | 0.032 |
| **SRRM2** | **S172** | 0.13 | 0.20 | 0.10 | 0.19 | -0.04 | -0.80 | 0.297 |
| **SRRM2** | **S2692; S2694** | 0.23 | 0.06 | 0.21 | -0.09 | -0.14 | -0.36 | 0.02 |
| **SRRM2** | **S2335** | 0.10 | 0.25 | 0.14 | -0.39 | -0.18 | -0.02 | 0.034 |
| **SRRM2** | **S2581** | 0.21 | 0.16 | 0.14 | 0.03 | -0.21 | -0.43 | 0.049 |
| **SRRM2** | **S820; S** | 0.16 | 0.14 | 0.21 | -0.09 | -0.08 | -0.43 | 0.033 |
| **SRRM2** | **S1565** | 0.18 | 0.26 | 0.08 | -0.21 | 0.00 | -0.43 | 0.046 |
| **SRRM2** | **S914; S** | 0.19 | 0.14 | 0.20 | -0.12 | -0.07 | -0.45 | 0.033 |
| **SRRM2** | **S954** | 0.15 | 0.21 | 0.15 | -0.16 | 0.05 | -0.53 | 0.086 |
| **SRRM2** | **T903; S** | 0.18 | 0.20 | 0.14 | -0.12 | -0.06 | -0.46 | 0.036 |
| **SRRM2** | **T1208** | 0.18 | 0.10 | 0.24 | -0.08 | 0.01 | -0.60 | 0.114 |
| **SRRM2** | **S890; S892** | 0.20 | 0.14 | 0.19 | -0.07 | 0.01 | -0.61 | 0.108 |
| **SRRM2** | **S1620** | 0.19 | 0.23 | 0.13 | -0.20 | -0.02 | -0.46 | 0.037 |
| **SRRM2** | **S323** | 0.15 | 0.20 | 0.21 | -0.13 | -0.06 | -0.48 | 0.035 |
| **SRRM2** | **S2702** | 0.20 | 0.23 | 0.13 | -0.15 | -0.04 | -0.49 | 0.041 |
| **SRRM2** | **S534; S536** | 0.14 | 0.27 | 0.15 | -0.16 | -0.03 | -0.49 | 0.047 |
| **SRRM2** | **S1497** | 0.04 | 0.29 | 0.21 | -0.19 | 0.09 | -0.62 | 0.132 |
| **SRRM2** | **S220** | 0.23 | 0.11 | 0.23 | -0.25 | -0.04 | -0.40 | 0.02 |
| **SRRM2** | **S353** | 0.11 | 0.19 | 0.27 | -0.13 | -0.02 | -0.56 | 0.067 |
| **SRRM2** | **S2407; T2409** | 0.16 | 0.20 | 0.23 | -0.30 | -0.13 | -0.27 | 0.002 |
| **SRRM2** | **S1541; S1552** | 0.19 | 0.24 | 0.16 | -0.17 | -0.11 | -0.43 | 0.014 |
| **SRRM2** | **T1453; S1458; S1460** | 0.19 | 0.14 | 0.28 | 0.00 | -0.44 | -0.32 | 0.029 |
| **SRRM2** | **S1975** | 0.15 | 0.35 | 0.09 | -0.49 | 0.14 | -0.44 | 0.103 |
| **SRRM2** | **S1329** | 0.17 | 0.23 | 0.23 | -0.15 | -0.05 | -0.60 | 0.048 |
| **SRRM2** | **S2688** | 0.41 | 0.22 | -0.01 | -0.12 | -0.12 | -0.57 | 0.067 |
| **SRRM2** | **S972** | 0.29 | 0.26 | 0.10 | -0.11 | -0.36 | -0.32 | 0.007 |
| **SRRM2** | **T2092** | 0.19 | 0.26 | 0.16 | -0.16 | 0.07 | -0.74 | 0.118 |
| **SRRM2** | **S780** | 0.21 | 0.30 | 0.12 | -0.13 | -0.02 | -0.67 | 0.079 |
| **SRRM2** | **S1600; S1601** | 0.20 | 0.23 | 0.22 | -0.25 | 0.00 | -0.58 | 0.041 |
| **SRRM2** | **S1822; S1824** | 0.30 | 0.29 | 0.08 | -0.09 | -0.32 | -0.42 | 0.015 |
| **SRRM2** | **S1731** | 0.22 | 0.32 | 0.11 | -0.10 | 0.01 | -0.81 | 0.123 |
| **SRRM2** | **S1482; S1483** | 0.21 | 0.18 | 0.31 | -0.09 | -0.12 | -0.70 | 0.058 |
| **SRRM2** | **S1132** | 0.14 | 0.21 | 0.33 | -0.19 | 0.02 | -0.75 | 0.086 |
| **SRRM2** | **S1072; S1073** | 0.10 | 0.27 | 0.32 | -0.09 | -0.07 | -0.78 | 0.088 |
| **SRRM2** | **S1581** | 0.19 | 0.31 | 0.20 | -0.26 | 0.02 | -0.69 | 0.06 |
| **SRRM2** | **S1519** | 0.17 | 0.31 | 0.25 | -0.43 | 0.07 | -0.62 | 0.052 |
| **SRRM2** | **S1541** | 0.21 | 0.28 | 0.25 | -0.31 | -0.03 | -0.65 | 0.033 |
| **SRRM2** | **S/T** | 0.24 | 0.28 | 0.23 | -0.28 | -0.01 | -0.70 | 0.047 |
| **SRRM2** | **S1423** | 0.24 | 0.32 | 0.19 | -0.22 | 0.01 | -0.82 | 0.077 |
| **SRRM2** | **S1893** | 0.14 | 0.31 | 0.29 | -0.22 | 0.03 | -0.85 | 0.09 |
| **SRRM2** | **S1916** | 0.21 | 0.28 | 0.25 | -0.20 | 0.06 | -0.91 | 0.107 |
| **SRRM2** | **S745** | 0.24 | 0.29 | 0.21 | -0.16 | 0.08 | -1.03 | 0.141 |
| **SRRM2** | **S1601** | 0.18 | 0.32 | 0.29 | -0.23 | -0.02 | -0.83 | 0.066 |

| | | | | | | | | |
|---|---|---|---|---|---|---|---|---|
| **SRRM2** | **S1423; S1424** | 0.24 | 0.30 | 0.25 | -0.18 | -0.03 | -0.87 | 0.073 |
| **SRRM2** | **S1421** | 0.25 | 0.32 | 0.24 | -0.20 | -0.03 | -0.93 | 0.076 |
| **SRRM2** | **S2398** | 0.29 | 0.22 | 0.36 | -0.40 | -0.06 | -0.69 | 0.022 |
| **SRRM2** | **S2694** | 0.33 | 0.34 | 0.21 | -0.35 | -0.12 | -0.68 | 0.016 |
| **SRRM2** | **S1869** | 0.25 | 0.36 | 0.25 | -0.26 | -0.01 | -0.95 | 0.07 |
| **SRRM2** | **S2449** | 1.11 | -0.40 | 0.27 | -0.48 | 0.22 | -1.50 | 0.239 |
| **SRRM2** | **S745; S746** | 0.36 | 0.49 | 0.32 | -0.39 | -0.12 | -1.38 | 0.059 |
| **SRRM2** | **T1003** | 0.42 | 0.48 | 0.52 | -0.95 | -0.11 | -1.36 | 0.026 |
| **SRRM2** | **S/T** | 0.92 | 0.56 | 0.27 | -0.44 | -0.52 | -1.50 | 0.022 |
| **SRRT** | **S67** | 0.13 | 0.09 | 0.09 | -0.08 | -0.10 | -0.16 | 0.002 |
| **SRRT** | **S319** | 0.20 | 0.13 | 0.07 | -0.13 | -0.07 | -0.26 | 0.014 |
| **SRRT** | **S493** | 0.21 | 0.18 | 0.19 | -0.59 | 0.17 | -0.37 | 0.116 |
| **SRRT** | **S/T** | 0.26 | 0.29 | 0.19 | -0.30 | -0.13 | -0.49 | 0.007 |
| **SRRT** | **S74** | 0.33 | 0.33 | 0.23 | -0.35 | -0.14 | -0.69 | 0.013 |
| **SRRT** | **S67; S74** | 0.33 | 0.40 | 0.18 | -0.19 | -0.30 | -0.75 | 0.018 |
| **SRRT** | **S74** | 0.39 | 0.46 | 0.39 | -0.53 | -0.18 | -1.22 | 0.026 |
| **SRSF1** | **S199** | 0.02 | 0.05 | 0.14 | 0.08 | -0.08 | -0.24 | 0.206 |
| **SRSF1** | **S199; S201** | 0.32 | 0.29 | 0.40 | -0.39 | -0.15 | -0.87 | 0.02 |
| **SRSF10** | **S158** | -0.16 | -0.07 | -0.07 | 0.02 | 0.48 | -0.35 | 0.562 |
| **SRSF10** | **S209** | 0.22 | 0.02 | -0.27 | 0.17 | 0.00 | -0.20 | 0.988 |
| **SRSF10** | **S131; S133** | 0.06 | 0.06 | 0.05 | -0.03 | -0.04 | -0.10 | 0.006 |
| **SRSF10** | **S133** | 0.25 | 0.14 | -0.14 | 0.03 | -0.03 | -0.33 | 0.299 |
| **SRSF10** | **S133** | 0.21 | 0.20 | 0.07 | -0.08 | -0.01 | -0.50 | 0.093 |
| **SRSF10** | **S106** | 0.27 | 0.29 | 0.22 | -0.25 | -0.16 | -0.58 | 0.01 |
| **SRSF10** | **S160** | 0.25 | 0.26 | 0.26 | -0.22 | -0.06 | -0.74 | 0.044 |
| **SRSF10** | **S160** | 0.37 | 0.39 | 0.33 | -0.32 | -0.19 | -1.09 | 0.034 |
| **SRSF11** | **S434** | 0.13 | 0.00 | 0.12 | -0.22 | 0.00 | -0.06 | 0.082 |
| **SRSF11** | **S434** | 0.09 | 0.20 | 0.00 | -0.19 | 0.01 | -0.14 | 0.069 |
| **SRSF11** | **S449** | 0.16 | 0.18 | 0.02 | -0.09 | -0.15 | -0.16 | 0.01 |
| **SRSF11** | **S414** | 0.21 | 0.18 | -0.04 | -0.17 | -0.02 | -0.22 | 0.063 |
| **SRSF11** | **S434** | 0.08 | 0.16 | 0.13 | -0.24 | 0.00 | -0.18 | 0.026 |
| **SRSF11** | **S323; T325** | 0.31 | 0.26 | 0.15 | -0.20 | -0.22 | -0.47 | 0.006 |
| **SRSF11** | **S434** | 0.33 | 0.30 | 0.14 | -0.28 | -0.07 | -0.66 | 0.031 |
| **SRSF2** | **S208; S212** | 0.06 | -0.01 | -0.08 | 0.05 | 0.11 | -0.14 | 0.88 |
| **SRSF2** | **S191** | -0.07 | 0.00 | 0.20 | -0.11 | 0.13 | -0.20 | 0.457 |
| **SRSF2** | **S208** | 0.11 | 0.16 | -0.08 | -0.23 | 0.30 | -0.38 | 0.476 |
| **SRSF2** | **S189; S191** | 0.06 | 0.12 | 0.26 | -0.21 | -0.06 | -0.24 | 0.017 |
| **SRSF3** | **S5** | -0.52 | -0.11 | 0.12 | 0.08 | 0.46 | -0.23 | 0.372 |
| **SRSF3** | **S95** | 0.04 | -0.12 | -0.11 | 0.13 | 0.05 | -0.01 | 0.132 |
| **SRSF3** | **S108** | 0.13 | 0.11 | 0.17 | -0.15 | -0.02 | -0.30 | 0.025 |
| **SRSF4** | **S259; S261** | 0.13 | 0.02 | -0.08 | 0.12 | -0.21 | -0.01 | 0.672 |
| **SRSF4** | **S78** | 0.40 | -0.13 | -0.36 | 0.55 | -0.08 | -0.79 | 0.871 |
| **SRSF4** | **S** | 0.21 | 0.05 | 0.16 | 0.08 | -0.33 | -0.26 | 0.085 |
| **SRSF4** | **S267; S269** | 0.23 | 0.22 | 0.03 | 0.01 | -0.19 | -0.39 | 0.057 |
| **SRSF4** | **S456** | 0.29 | 0.25 | 0.24 | -0.01 | -0.30 | -0.72 | 0.044 |
| **SRSF5** | **S250** | -0.14 | -0.47 | -0.42 | 0.31 | 0.12 | 0.36 | 0.009 |
| **SRSF5** | **S116** | -0.09 | -0.50 | 0.01 | 0.45 | -0.11 | 0.07 | 0.214 |
| **SRSF5** | **S250; S253** | 0.09 | -0.02 | 0.10 | 0.03 | -0.02 | -0.19 | 0.198 |

| | | | | | | | | |
|---|---|---|---|---|---|---|---|---|
| **SRSF5** | **S229; S233** | 0.13 | 0.14 | 0.03 | -0.18 | 0.01 | -0.15 | 0.042 |
| **SRSF5** | **S233** | 0.27 | 0.20 | 0.09 | -0.43 | 0.01 | -0.26 | 0.042 |
| **SRSF6** | **S303** | 0.11 | -0.63 | -0.03 | 0.17 | 0.06 | 0.17 | 0.236 |
| **SRSF6** | **S314; S316** | -0.05 | -0.10 | -0.10 | 0.12 | -0.02 | 0.13 | 0.039 |
| **SRSF6** | **S314; S316** | -0.02 | -0.06 | 0.08 | 0.15 | -0.07 | -0.09 | 0.939 |
| **SRSF6** | **S314; S316** | 0.11 | 0.01 | -0.09 | 0.03 | -0.09 | 0.01 | 0.68 |
| **SRSF6** | **S316** | 0.19 | 0.04 | 0.00 | 0.01 | 0.16 | -0.51 | 0.415 |
| **SRSF6** | **S314; S316** | 0.24 | 0.17 | -0.07 | -0.01 | -0.08 | -0.30 | 0.132 |
| **SRSF6** | **S272** | 0.29 | -0.03 | 0.12 | 0.15 | -0.13 | -0.54 | 0.243 |
| **SRSF6** | **S303** | 0.28 | 0.23 | 0.23 | -0.26 | -0.10 | -0.57 | 0.017 |
| **SRSF7** | **S181; S183** | 0.12 | 0.15 | 0.12 | -0.14 | -0.06 | -0.24 | 0.006 |
| **SRSF8** | **S273; S** | 0.00 | 0.03 | -0.07 | 0.09 | 0.03 | -0.09 | 0.692 |
| **SRSF8** | **S273** | -0.06 | -0.04 | 0.21 | 0.09 | -0.02 | -0.21 | 0.547 |
| **SRSF8** | **S273** | 0.10 | 0.15 | 0.20 | -0.02 | 0.04 | -0.61 | 0.17 |
| **SRSF9** | **S204** | 0.18 | -0.24 | -0.03 | -0.26 | 0.34 | -0.08 | 0.885 |
| **SRSF9** | **S211** | 0.14 | -0.14 | -0.01 | 0.00 | -0.23 | 0.19 | 0.933 |
| **SRSF9** | **S204** | 0.09 | -0.01 | 0.02 | -0.01 | -0.10 | 0.00 | 0.172 |
| **SRSF9** | **S211; S216** | 0.12 | 0.00 | 0.08 | 0.06 | -0.24 | -0.04 | 0.221 |
| **SRSF9** | **S204** | 0.09 | 0.02 | 0.12 | -0.05 | -0.13 | -0.06 | 0.018 |
| **SRSF9** | **S189** | 0.18 | 0.21 | 0.22 | -0.23 | 0.00 | -0.54 | 0.043 |
| **SSB** | **S92** | -0.36 | -0.53 | -0.31 | 0.43 | 0.27 | 0.23 | 0.001 |
| **SSB** | **S92** | -0.15 | -0.40 | -0.64 | 0.46 | 0.28 | 0.15 | 0.015 |
| **SSB** | **S92** | 0.11 | -0.24 | -0.48 | 0.21 | 0.17 | 0.10 | 0.105 |
| **SSB** | **S366** | 0.06 | 0.12 | 0.08 | -0.10 | 0.04 | -0.22 | 0.084 |
| **SSB** | **S366** | 0.03 | 0.09 | 0.13 | -0.13 | 0.07 | -0.23 | 0.12 |
| **SSH1** | **S938** | -0.45 | -0.05 | -0.10 | 0.22 | 0.18 | 0.10 | 0.051 |
| **SSH1** | **S978** | -0.07 | -0.07 | 0.01 | 0.16 | 0.03 | -0.07 | 0.322 |
| **SSH1** | **S515** | -0.07 | -0.04 | 0.25 | 0.08 | -0.19 | -0.07 | 0.469 |
| **SSH2** | **S** | -0.13 | 0.19 | 0.04 | 0.28 | -0.56 | 0.03 | 0.679 |
| **SSH2** | **S487** | 0.12 | 0.12 | 0.09 | -0.03 | -0.11 | -0.21 | 0.011 |
| **SSH2** | **T795** | 0.06 | 0.31 | 0.20 | -0.31 | -0.06 | -0.30 | 0.02 |
| **SSH2** | **S461** | 0.31 | 0.26 | 0.29 | -0.43 | -0.22 | -0.44 | 9E-04 |
| **SSH2** | **S461** | 0.31 | 0.26 | 0.29 | -0.43 | -0.22 | -0.44 | 9E-04 |
| **SSH3** | **S484** | -0.34 | -0.30 | -0.15 | 0.28 | 0.27 | 0.10 | 0.004 |
| **SSH3** | **T/S** | -0.36 | -0.09 | -0.27 | 0.29 | -0.13 | 0.40 | 0.077 |
| **SSH3** | **S/T** | -0.39 | -0.05 | -0.27 | -0.04 | -0.06 | 0.60 | 0.163 |
| **SSH3** | **S9** | -0.26 | -0.20 | -0.18 | 0.19 | 0.27 | 0.09 | 0.002 |
| **SSR3** | **S11** | -0.04 | -0.03 | -0.04 | 0.03 | 0.04 | 0.03 | 2E-04 |
| **SSR3** | **S** | 0.30 | 0.44 | -0.10 | -0.51 | -0.03 | -0.33 | 0.079 |
| **SSRP1** | **S437; S444** | -0.58 | -0.70 | -0.60 | 0.54 | 0.41 | 0.35 | 1E-04 |
| **SSRP1** | **S437; S444** | -0.51 | -0.67 | -0.57 | 0.41 | 0.50 | 0.32 | 1E-04 |
| **SSRP1** | **T111** | 0.03 | -0.55 | -0.17 | 0.05 | -0.11 | 0.52 | 0.206 |
| **SSRP1** | **S444** | 0.01 | -0.02 | -0.12 | -0.06 | 0.04 | 0.13 | 0.307 |
| **SSRP1** | **S444** | -0.01 | -0.02 | 0.00 | 0.04 | 0.02 | -0.04 | 0.468 |
| **SSRP1** | **S444** | 0.02 | 0.04 | -0.03 | -0.06 | 0.06 | -0.02 | 0.666 |
| **SSRP1** | **S667; S671; S672** | 0.05 | 0.03 | 0.06 | 0.02 | -0.12 | -0.04 | 0.089 |
| **SSRP1** | **S444** | -0.04 | 0.11 | 0.10 | -0.05 | -0.02 | -0.12 | 0.116 |
| **SSRP1** | **S667** | 0.06 | 0.02 | 0.10 | 0.06 | -0.05 | -0.22 | 0.196 |

| | | | | | | | | |
|---|---|---|---|---|---|---|---|---|
| SSRP1 | S671; S672; S673 | 0.09 | 0.07 | 0.14 | -0.28 | 0.03 | -0.09 | 0.081 |
| SSRP1 | S667 | 0.17 | 0.31 | 0.12 | -0.27 | -0.18 | -0.24 | 0.002 |
| SSRP1 | S671; S672 | 0.29 | 0.22 | 0.11 | -0.13 | -0.01 | -0.69 | 0.089 |
| SSRP1 | S667; S671; S673 | 0.33 | 0.36 | 0.24 | -0.38 | -0.05 | -0.87 | 0.038 |
| SSX2 | S108 | -0.14 | -0.18 | -0.23 | 0.12 | -0.15 | 0.45 | 0.145 |
| SSX2IP | S540 | -0.28 | 0.11 | -0.08 | -0.13 | -0.53 | 0.63 | 0.85 |
| SSX2IP | S312 | 0.00 | 0.12 | 0.13 | -0.12 | -0.09 | -0.07 | 0.021 |
| ST7 | S386 | -0.09 | 0.10 | -0.32 | 0.16 | 0.09 | 0.02 | 0.204 |
| STAG1 | S1093 | 0.00 | -0.26 | -0.38 | -0.23 | 0.11 | 0.55 | 0.232 |
| STAG1 | S1093 | -0.11 | 0.23 | 0.04 | -0.02 | -0.21 | 0.03 | 0.375 |
| STAG1 | S/T | 0.23 | -0.02 | 0.35 | -0.18 | -0.06 | -0.48 | 0.061 |
| STAG1 | S1065 | 0.27 | 0.31 | 0.12 | -0.28 | -0.05 | -0.55 | 0.027 |
| STAG1 | S1065 | 0.21 | 0.21 | 0.32 | -0.44 | -0.04 | -0.47 | 0.016 |
| STAG1 | S1062 | 0.28 | 0.21 | 0.26 | 0.03 | -0.32 | -0.72 | 0.054 |
| STAG2 | S1161 | -0.04 | -0.35 | -0.26 | 0.05 | 0.08 | 0.39 | 0.052 |
| STAG2 | S1061 | -0.08 | -0.19 | -0.36 | -0.28 | 0.29 | 0.44 | 0.2 |
| STAG2 | S/T | -0.30 | -0.11 | -0.15 | 0.31 | -0.09 | 0.24 | 0.067 |
| STAG2 | T/S | -0.11 | -0.04 | 0.09 | -0.07 | 0.04 | 0.08 | 0.652 |
| STAG3 | S1190 | -0.18 | 0.16 | -0.21 | 0.21 | -0.33 | 0.24 | 0.619 |
| STAG3 | S1174; S1190 | -0.62 | -0.58 | 0.76 | -0.54 | 0.36 | 0.03 | 0.866 |
| STAG3 | S/T | 0.45 | -0.31 | -0.08 | -0.21 | 0.12 | -0.11 | 0.729 |
| STAG3 | S93 | -0.19 | 0.59 | 0.39 | -0.93 | -0.57 | 0.14 | 0.14 |
| STAMBPL1 | S242 | -0.17 | -0.17 | -0.04 | 0.35 | -0.49 | 0.34 | 0.537 |
| STARD10 | S284 | -0.34 | -0.25 | 0.32 | -0.24 | 0.04 | 0.32 | 0.651 |
| STARD3NL | S218 | 0.00 | -0.18 | -0.05 | 0.02 | -0.08 | 0.26 | 0.269 |
| STARD3NL | S193 | -0.02 | 0.14 | 0.12 | -0.29 | 0.01 | 0.00 | 0.2 |
| STAT1 | S740 | -0.29 | -0.39 | -0.60 | 0.50 | 0.21 | 0.26 | 0.004 |
| STAT1 | S727 | 0.18 | -0.60 | -0.58 | 0.44 | -0.11 | 0.33 | 0.147 |
| STAT1 | S727 | -0.41 | -0.60 | 0.03 | 0.57 | 0.31 | -0.25 | 0.152 |
| STAT3 | S691 | -0.19 | 0.11 | -0.14 | -0.62 | 0.56 | 0.01 | 0.882 |
| STAU1 | S176 | -0.01 | -0.04 | 0.01 | 0.01 | 0.05 | -0.02 | 0.39 |
| STAU1 | S390 | 0.13 | -0.04 | 0.01 | -0.21 | -0.02 | 0.11 | 0.527 |
| STAU1 | S278 | 0.21 | 0.05 | -0.19 | -0.20 | 0.42 | -0.47 | 0.734 |
| STAU1 | S390 | 0.12 | 0.07 | 0.12 | -0.07 | 0.05 | -0.36 | 0.134 |
| STAU1 | S390 | 0.05 | 0.22 | 0.08 | -0.08 | 0.13 | -0.51 | 0.242 |
| STAU1 | S390 | 0.19 | 0.17 | 0.09 | -0.06 | -0.08 | -0.39 | 0.041 |
| STAU2 | S416 | -0.12 | -0.10 | 0.00 | 0.10 | -0.13 | 0.22 | 0.277 |
| STAU2 | S188 | -0.04 | 0.00 | -0.14 | 0.03 | 0.04 | 0.10 | 0.067 |
| STAU2 | S188 | 0.04 | 0.08 | -0.04 | -0.07 | 0.06 | -0.08 | 0.375 |
| STAU2 | S416 | 0.09 | 0.06 | -0.07 | 0.10 | -0.12 | -0.09 | 0.506 |
| STAU2 | S188 | -0.30 | -0.16 | 0.45 | 0.35 | -0.62 | 0.00 | 0.822 |
| STAU2 | S485; S492 | -0.01 | 0.05 | 0.14 | -0.10 | -0.14 | 0.03 | 0.125 |
| STAU2 | S/T | 0.13 | 0.02 | 0.14 | 0.04 | -0.22 | -0.14 | 0.081 |
| STAU2 | S485; T488; S492 | 0.09 | -0.01 | 0.21 | -0.06 | -0.21 | -0.05 | 0.065 |
| STIM1 | S616 | 0.03 | -0.11 | -0.08 | 0.11 | -0.01 | 0.05 | 0.131 |
| STIM1 | S668 | 0.10 | 0.20 | 0.03 | -0.06 | -0.05 | -0.26 | 0.047 |
| STIM1 | S575 | 0.17 | 0.21 | 0.21 | -0.50 | 0.00 | -0.24 | 0.038 |
| STIM2 | S | 0.09 | 0.18 | 0.09 | 0.05 | -0.21 | -0.24 | 0.057 |

| | | | | | | | | |
|---|---|---|---|---|---|---|---|---|
| STK10 | T952 | 0.12 | 0.10 | 0.03 | -0.17 | 0.00 | -0.11 | 0.039 |
| STK10 | T952 | 0.33 | 0.07 | 0.14 | -0.47 | -0.08 | -0.12 | 0.052 |
| STK11IP | S482 | -0.13 | -0.14 | -0.22 | 0.17 | -0.08 | 0.31 | 0.068 |
| STK11IP | S761 | -0.03 | 0.07 | -0.07 | 0.03 | -0.01 | 0.01 | 0.682 |
| STK11IP | S392 | 0.16 | 0.00 | 0.06 | -0.08 | -0.24 | 0.06 | 0.187 |
| STK3 | S316 | -0.46 | 0.03 | 0.15 | 0.11 | 0.22 | -0.16 | 0.525 |
| STK3 | S316 | -0.10 | -0.05 | -0.04 | 0.08 | 0.10 | 0.00 | 0.028 |
| STK39 | S385 | -0.19 | 0.10 | 0.05 | 0.15 | 0.09 | -0.26 | 0.975 |
| STMN1 | S38 | 0.34 | 0.36 | -0.09 | -0.33 | -0.16 | -0.28 | 0.04 |
| STOML2 | T327 | 0.02 | -0.01 | -0.16 | -0.11 | -0.03 | 0.25 | 0.518 |
| STOML2 | T/S | 0.08 | -0.02 | 0.15 | 0.17 | 0.07 | -0.58 | 0.491 |
| STOML2 | S345 | 0.30 | 0.43 | 0.37 | -0.44 | -0.18 | -0.95 | 0.018 |
| STON2 | S302 | -0.23 | 0.34 | 0.21 | -0.12 | 0.00 | -0.31 | 0.266 |
| STRIP1 | S335 | -0.12 | -0.15 | -0.01 | 0.26 | -0.21 | 0.17 | 0.318 |
| STRIP1 | S335 | -0.03 | 0.02 | -0.08 | 0.06 | -0.10 | 0.12 | 0.441 |
| STRIP1 | S59 | 0.08 | 0.17 | 0.15 | -0.26 | 0.04 | -0.24 | 0.045 |
| STRN | S245 | -0.01 | -0.09 | -0.03 | -0.01 | 0.03 | 0.10 | 0.112 |
| STRN | S | 0.13 | -0.46 | 0.25 | 0.16 | 0.32 | -0.70 | 0.912 |
| STRN | S/Y/T | -0.25 | 0.61 | -0.01 | 0.14 | -0.22 | -0.55 | 0.364 |
| STRN3 | S229 | 0.00 | 0.24 | 0.23 | -0.43 | 0.04 | -0.20 | 0.087 |
| STRN3 | S229 | 0.21 | 0.10 | 0.55 | -0.47 | -0.11 | -0.60 | 0.027 |
| STRN4 | S167 | -0.07 | -0.04 | -0.08 | 0.02 | -0.08 | 0.23 | 0.25 |
| STT3B | Y511; S | 0.14 | 0.08 | -0.16 | 0.06 | -0.26 | 0.10 | 0.715 |
| STT3B | S498; S499 | 0.07 | 0.06 | 0.14 | -0.01 | -0.11 | -0.19 | 0.03 |
| STT3B | S499 | 0.16 | 0.27 | 0.18 | -0.29 | -0.16 | -0.27 | 0.001 |
| STT3B | S499 | 0.16 | 0.27 | 0.18 | -0.29 | -0.16 | -0.27 | 0.001 |
| STUB1 | S23 | 0.00 | -0.11 | -0.14 | 0.07 | 0.18 | -0.03 | 0.1 |
| STUB1 | S19; S23 | 0.06 | -0.36 | 0.18 | 0.06 | 0.05 | -0.06 | 0.756 |
| STUB1 | S19 | -0.01 | -0.05 | 0.07 | 0.10 | 0.11 | -0.24 | 0.883 |
| STUB1 | S19 | 0.06 | -0.01 | 0.02 | -0.03 | 0.09 | -0.14 | 0.48 |
| STX17 | S/T | 0.20 | 0.24 | 0.22 | -0.18 | 0.06 | -0.79 | 0.107 |
| STX18 | S189 | 0.18 | 0.26 | 0.12 | -0.36 | -0.01 | -0.30 | 0.024 |
| STX4 | S117 | -0.14 | 0.07 | -0.05 | 0.02 | -0.22 | 0.26 | 0.735 |
| STX4 | S36 | -0.19 | 0.11 | 0.09 | 0.21 | 0.06 | -0.36 | 0.872 |
| STX4 | S117 | 0.06 | 0.03 | 0.13 | 0.02 | -0.33 | 0.05 | 0.277 |
| STX7 | S45 | 0.17 | -0.15 | -0.22 | -0.14 | -0.34 | 0.50 | 0.811 |
| STX7 | T79 | 0.04 | -0.11 | 0.17 | 0.03 | -0.31 | 0.13 | 0.618 |
| STX7 | S131 | 0.24 | -0.27 | 0.09 | -0.18 | -0.36 | 0.33 | 0.746 |
| STXBP5 | S/Y | -0.01 | -0.41 | -0.08 | 0.20 | -0.18 | 0.36 | 0.217 |
| STXBP5 | S692 | -0.24 | -0.11 | -0.02 | 0.26 | -0.13 | 0.17 | 0.176 |
| STXBP5 | S902 | 0.06 | -0.08 | -0.09 | 0.23 | -0.17 | 0.02 | 0.639 |
| STXBP5 | S782 | 0.05 | -0.03 | -0.02 | 0.04 | 0.03 | -0.08 | 0.977 |
| STXBP5 | S782; T784 | 0.13 | -0.06 | 0.09 | -0.10 | -0.07 | -0.01 | 0.157 |
| STXBP5 | S785 | 0.05 | 0.25 | 0.29 | -0.32 | 0.03 | -0.46 | 0.05 |
| SUB1 | S19; S | 0.22 | 0.04 | -0.05 | -0.19 | -0.10 | 0.04 | 0.22 |
| SUB1 | S4 | 0.18 | 0.02 | 0.12 | -0.24 | 0.01 | -0.12 | 0.059 |
| SUB1 | S10; S11; S19 | 0.33 | 0.18 | 0.00 | -0.13 | -0.17 | -0.29 | 0.026 |
| SUB1 | S19 | 0.37 | 0.04 | 0.10 | -0.30 | -0.19 | -0.11 | 0.031 |

| | | | | | | | | |
|---|---|---|---|---|---|---|---|---|
| SUB1 | S19 | 0.43 | 0.27 | 0.04 | -0.35 | -0.14 | -0.45 | 0.019 |
| SUB1 | S4 | 0.61 | 0.33 | -0.26 | -0.37 | 0.00 | -0.71 | 0.148 |
| SUB1 | S10; S19 | 0.11 | 0.13 | 0.68 | -0.44 | -0.37 | -0.51 | 0.017 |
| SUCO | S1081 | 0.06 | 0.09 | 0.07 | -0.05 | -0.24 | 0.03 | 0.119 |
| SUDS3 | S234; S236 | -0.21 | -0.09 | -0.12 | 0.26 | -0.28 | 0.32 | 0.288 |
| SUDS3 | S234 | -0.02 | -0.02 | -0.04 | 0.11 | -0.11 | 0.08 | 0.501 |
| SUDS3 | S45; T49; S53 | 0.17 | 0.19 | -0.09 | 0.14 | -0.04 | -0.48 | 0.359 |
| SUDS3 | T49; S/T/Y | 0.40 | 0.28 | -0.17 | -0.43 | -0.04 | -0.22 | 0.125 |
| SUFU | S346 | -0.02 | 0.21 | 0.09 | -0.16 | -0.13 | -0.03 | 0.061 |
| SUGP1 | S411 | -0.01 | 0.01 | -0.13 | -0.03 | -0.11 | 0.24 | 0.525 |
| SUGP1 | S485 | -0.10 | 0.06 | 0.10 | 0.02 | -0.07 | -0.02 | 0.573 |
| SUGP1 | S | -0.24 | -0.03 | 0.43 | 0.10 | -0.29 | -0.10 | 0.553 |
| SUGP1 | S485 | 0.00 | 0.06 | 0.26 | -0.09 | -0.18 | -0.09 | 0.052 |
| SUGP2 | S887 | 0.13 | -0.40 | -0.02 | 0.00 | 0.19 | 0.03 | 0.374 |
| SUGP2 | S572 | -0.08 | -0.11 | 0.18 | 0.10 | -0.25 | 0.12 | 0.957 |
| SUGP2 | S603 | 0.11 | 0.05 | -0.03 | -0.05 | -0.01 | -0.07 | 0.11 |
| SUGP2 | S93 | 0.11 | 0.17 | -0.07 | -0.19 | -0.08 | 0.03 | 0.202 |
| SUGP2 | S277 | 0.16 | 0.11 | 0.21 | -0.26 | -0.01 | -0.28 | 0.019 |
| SUMO3 | S27 | -0.75 | -0.74 | -0.73 | 0.51 | 0.48 | 0.47 | 4E-08 |
| SUN1 | S48 | 0.03 | -0.06 | 0.11 | -0.06 | 0.03 | -0.06 | 0.38 |
| SUN1 | S79 | 0.06 | 0.04 | 0.04 | -0.26 | -0.12 | 0.20 | 0.478 |
| SUN2 | Y140; S/Y/T | -0.19 | -0.13 | -0.34 | 0.20 | 0.13 | 0.24 | 0.004 |
| SUN2 | S/Y/T | 0.01 | 0.01 | -0.08 | 0.01 | -0.04 | 0.08 | 0.488 |
| SUN2 | S/Y/T | 0.09 | -0.04 | 0.02 | 0.03 | -0.11 | 0.00 | 0.423 |
| SUN2 | S110; S116; S/T | 0.03 | 0.00 | 0.07 | 0.00 | -0.13 | 0.03 | 0.279 |
| SUN2 | S12 | -0.17 | 0.11 | 0.16 | -0.03 | -0.35 | 0.20 | 0.642 |
| SUN2 | S38 | 0.18 | 0.14 | 0.06 | -0.18 | -0.04 | -0.20 | 0.013 |
| SUN2 | S116 | 0.14 | 0.12 | 0.12 | -0.16 | -0.08 | -0.19 | 0.001 |
| SUN2 | S12 | 0.20 | 0.09 | 0.14 | -0.16 | -0.15 | -0.17 | 6E-04 |
| SUN2 | S38 | 0.16 | 0.36 | 0.08 | -0.30 | -0.06 | -0.36 | 0.024 |
| SUPT16H | S650 | -0.32 | -0.32 | -0.19 | 0.01 | 0.43 | 0.23 | 0.018 |
| SUPT16H | S982 | -0.02 | -0.12 | -0.15 | 0.12 | 0.17 | -0.03 | 0.067 |
| SUPT16H | S982; S986 | -0.04 | -0.19 | 0.05 | -0.14 | 0.17 | 0.12 | 0.423 |
| SUPT16H | S188 | 0.37 | -0.42 | -0.09 | 0.24 | -0.25 | 0.00 | 0.882 |
| SUPT16H | S1013 | -0.04 | 0.28 | -0.12 | -0.28 | 0.38 | -0.37 | 0.647 |
| SUPT20H | S296 | -0.04 | -0.50 | 0.27 | 0.18 | 0.20 | -0.26 | 0.651 |
| SUPT20H | Y159 | -0.01 | -0.13 | 0.13 | -0.22 | 0.17 | 0.01 | 0.95 |
| SUPT20H | S296 | 0.08 | -0.07 | 0.00 | 0.38 | -0.57 | 0.02 | 0.833 |
| SUPT5H | S | -0.36 | -0.15 | -0.18 | 0.28 | 0.08 | 0.22 | 0.009 |
| SUPT5H | T/S | -0.23 | 0.11 | -0.51 | 0.12 | 0.03 | 0.32 | 0.139 |
| SUPT5H | S | 0.30 | 0.25 | -0.29 | 0.03 | 0.16 | -0.69 | 0.481 |
| SUPT6H | T1523; S1528 | -0.10 | 0.04 | -0.18 | 0.03 | 0.08 | 0.10 | 0.096 |
| SUPT6H | S125 | 0.15 | -0.07 | -0.29 | 0.07 | -0.13 | 0.21 | 0.498 |
| SUPT6H | S73; S75; S78 | -0.10 | -0.03 | 0.06 | -0.23 | -0.08 | 0.32 | 0.878 |
| SUPT6H | S73; S75; S78 | 0.09 | 0.01 | -0.07 | 0.07 | -0.06 | -0.04 | 0.73 |
| SUPT6H | S75; S78 | 0.01 | 0.25 | -0.21 | 0.11 | -0.12 | -0.08 | 0.779 |
| SUPT6H | S125 | 0.26 | 0.08 | -0.12 | 0.01 | -0.10 | -0.17 | 0.248 |
| SUPT6H | T1523 | 0.15 | -0.03 | 0.12 | -0.08 | -0.10 | -0.08 | 0.042 |

| | | | | | | | | |
|---|---|---|---|---|---|---|---|---|
| SUPT6H | S125 | 0.21 | 0.12 | 0.19 | -0.53 | -0.17 | 0.05 | 0.086 |
| SUPT7L | S108 | 0.10 | 0.25 | 0.25 | -0.16 | 0.02 | -0.63 | 0.085 |
| SUPV3L1 | S725 | -0.19 | 0.01 | -0.01 | 0.04 | -0.04 | 0.16 | 0.246 |
| SURF2 | T190 | 0.14 | 0.11 | 0.04 | -0.13 | 0.03 | -0.23 | 0.059 |
| SURF6 | T229 | -0.24 | -0.53 | -0.18 | 0.28 | 0.17 | 0.31 | 0.008 |
| SURF6 | T184 | -0.31 | -0.20 | -0.38 | 0.27 | 0.09 | 0.35 | 0.004 |
| SURF6 | T184 | -0.31 | -0.20 | -0.38 | 0.27 | 0.09 | 0.35 | 0.004 |
| SURF6 | S74 | -0.27 | -0.43 | -0.09 | 0.28 | 0.32 | 0.03 | 0.023 |
| SURF6 | S138 | -0.01 | -0.21 | -0.28 | 0.12 | -0.01 | 0.31 | 0.065 |
| SUV39H1 | S391 | -0.13 | 0.27 | -0.38 | 0.30 | -0.01 | -0.17 | 0.634 |
| SUV39H2 | S382; S384; S388 | -0.13 | -0.38 | -0.25 | 0.23 | 0.11 | 0.29 | 0.006 |
| SUV39H2 | S388 | -0.15 | 0.05 | 0.15 | -0.17 | 0.21 | -0.14 | 0.77 |
| SUV39H2 | S388 | 0.12 | 0.01 | 0.15 | -0.09 | -0.19 | -0.03 | 0.034 |
| SUZ12 | S405 | -0.19 | -0.27 | -0.20 | 0.18 | 0.10 | 0.28 | 0.002 |
| SUZ12 | S695 | -0.29 | -0.11 | -0.16 | 0.04 | -0.07 | 0.47 | 0.125 |
| SUZ12 | S695 | -0.08 | -0.11 | -0.26 | 0.17 | -0.08 | 0.29 | 0.084 |
| SUZ12 | S695 | -0.09 | -0.10 | -0.20 | 0.14 | 0.03 | 0.19 | 0.013 |
| SUZ12 | S541; S546 | -0.13 | -0.08 | -0.17 | -0.08 | 0.23 | 0.18 | 0.078 |
| SUZ12 | S695 | -0.20 | -0.19 | 0.01 | 0.29 | 0.06 | -0.03 | 0.119 |
| SUZ12 | S546 | -0.05 | -0.11 | -0.09 | 0.10 | -0.12 | 0.23 | 0.206 |
| SUZ12 | S546 | -0.01 | 0.10 | -0.01 | -0.02 | 0.05 | -0.11 | 0.391 |
| SVIL | S1322 | -0.18 | -0.05 | -0.11 | 0.16 | -0.01 | 0.16 | 0.036 |
| SVIL | S1322 | -0.33 | 0.10 | -0.03 | -0.04 | -0.06 | 0.29 | 0.42 |
| SVIL | S707 | -0.03 | -0.11 | -0.04 | 0.01 | -0.03 | 0.18 | 0.176 |
| SVIL | S221 | -0.69 | 0.25 | 0.24 | -0.06 | -0.42 | 0.37 | 0.938 |
| SVIL | S707 | 0.09 | -0.01 | -0.13 | 0.04 | -0.14 | 0.14 | 0.802 |
| SVIL | S707 | 0.03 | 0.01 | 0.02 | -0.17 | 0.07 | 0.03 | 0.612 |
| SVIL | S914 | 0.08 | -0.11 | 0.13 | 0.05 | -0.11 | -0.07 | 0.43 |
| SVIL | S968 | 0.07 | 0.20 | 0.02 | -0.11 | -0.03 | -0.18 | 0.037 |
| SVIL | S1120 | 0.10 | 0.06 | 0.14 | -0.18 | -0.05 | -0.09 | 0.009 |
| SVIL | S221 | 0.14 | 0.16 | 0.10 | -0.06 | -0.21 | -0.17 | 0.004 |
| SVIL | S245 | 0.12 | 0.18 | 0.15 | -0.28 | 0.06 | -0.30 | 0.051 |
| SVIL | T/S | 0.09 | 0.05 | 0.45 | -0.37 | -0.19 | -0.18 | 0.035 |
| SYAP1 | T248 | 0.33 | -0.34 | -0.28 | -0.34 | 0.45 | -0.05 | 0.728 |
| SYBU | S396 | 0.26 | 0.09 | 0.01 | -0.25 | -0.33 | 0.12 | 0.159 |
| SYCP2 | S1108 | -0.12 | -0.09 | -0.07 | -0.39 | 0.21 | 0.34 | 0.557 |
| SYMPK | T1257 | -0.46 | -0.27 | -0.08 | 0.53 | 0.32 | -0.31 | 0.178 |
| SYMPK | S870 | 0.07 | -0.22 | -0.19 | -0.06 | 0.18 | 0.18 | 0.159 |
| SYMPK | S1243 | -0.01 | -0.03 | -0.26 | 0.18 | -0.15 | 0.22 | 0.26 |
| SYMPK | S1243; T1257; S1259 | -0.02 | 0.08 | -0.04 | 0.04 | 0.13 | -0.21 | 0.854 |
| SYMPK | S1243 | 0.06 | 0.00 | -0.01 | 0.06 | 0.00 | -0.11 | 0.547 |
| SYMPK | S1243 | 0.10 | 0.10 | -0.11 | 0.22 | -0.16 | -0.19 | 0.657 |
| SYMPK | S1243 | 0.10 | 0.08 | 0.06 | 0.10 | -0.11 | -0.27 | 0.189 |
| SYMPK | S1259 | 0.08 | 0.09 | 0.14 | -0.05 | -0.10 | -0.18 | 0.006 |
| SYNCRIP | S575 | -0.10 | 0.09 | -0.60 | 0.40 | -0.09 | 0.11 | 0.24 |
| SYNCRIP | S575 | -0.35 | 0.05 | 0.16 | 0.39 | -0.13 | -0.26 | 0.869 |
| SYNE1 | T8360 | 0.14 | 0.03 | -0.28 | -0.11 | 0.16 | 0.01 | 0.713 |
| SYNE1 | S8305 | 0.16 | 0.06 | -0.23 | -0.05 | -0.23 | 0.23 | 0.949 |

| | | | | | | | | |
|---|---|---|---|---|---|---|---|---|
| SYNE2 | S4126 | -0.14 | -0.26 | -0.22 | 0.18 | -0.15 | 0.44 | 0.105 |
| SYNE2 | S6392; S | -0.11 | 0.03 | -0.17 | 0.07 | -0.09 | 0.23 | 0.247 |
| SYNE2 | S841 | -0.18 | 0.05 | 0.02 | 0.18 | -0.20 | 0.09 | 0.671 |
| SYNE2 | S6625 | 0.02 | 0.19 | -0.36 | -0.21 | 0.39 | -0.16 | 0.842 |
| SYNE2 | S2781 | 0.02 | 0.00 | -0.04 | -0.07 | 0.04 | 0.05 | 0.792 |
| SYNE2 | S6389 | -0.13 | 0.17 | -0.05 | -0.10 | 0.03 | 0.05 | 0.978 |
| SYNE2 | S6361 | 0.04 | -0.02 | -0.01 | -0.08 | 0.01 | 0.07 | 0.935 |
| SYNE2 | S4108 | 0.02 | -0.07 | 0.13 | 0.03 | -0.08 | -0.03 | 0.472 |
| SYNE2 | S6389 | -0.03 | 0.19 | -0.07 | -0.08 | -0.09 | 0.07 | 0.541 |
| SYNE2 | S4108; S4126 | 0.14 | 0.09 | 0.09 | -0.20 | 0.08 | -0.24 | 0.084 |
| SYNE2 | S | 0.13 | 0.06 | 0.17 | 0.15 | -0.49 | -0.13 | 0.22 |
| SYNE2 | S/T/Y | 0.27 | 0.06 | 0.04 | -0.47 | -0.16 | 0.15 | 0.215 |
| SYNE2 | S4156 | 0.19 | 0.15 | 0.05 | -0.24 | -0.27 | 0.05 | 0.059 |
| SYNE2 | S4108; S4126 | 0.07 | 0.16 | 0.23 | -0.32 | -0.04 | -0.18 | 0.022 |
| SYNE2 | S6361 | 0.07 | 0.27 | 0.27 | -0.35 | -0.01 | -0.40 | 0.032 |
| SYNE2 | S2781 | 0.17 | 0.08 | 0.39 | -0.40 | 0.16 | -0.66 | 0.116 |
| SYNE2 | S6361 | 0.21 | 0.18 | 0.31 | -0.29 | -0.05 | -0.54 | 0.023 |
| SYNE2 | S6361 | 0.23 | 0.23 | 0.27 | -0.40 | 0.04 | -0.58 | 0.042 |
| SYNE4 | S331 | -0.03 | -0.12 | 0.00 | 0.02 | 0.02 | 0.12 | 0.102 |
| SYNJ1 | S1565 | 0.04 | 0.05 | -0.01 | 0.04 | -0.04 | -0.08 | 0.24 |
| SYNJ1 | T1220 | 0.16 | 0.17 | 0.26 | -0.25 | -0.01 | -0.46 | 0.03 |
| SYNRG | S752 | -0.32 | -0.05 | -0.14 | 0.25 | 0.14 | 0.05 | 0.03 |
| SYNRG | S/T | 0.00 | -0.04 | -0.38 | 0.10 | 0.21 | 0.05 | 0.115 |
| SYNRG | S720 | -0.24 | 0.00 | -0.12 | 0.25 | -0.08 | 0.13 | 0.137 |
| SYNRG | S935 | -0.02 | -0.14 | -0.10 | 0.14 | 0.17 | -0.09 | 0.148 |
| SYNRG | S/T | -0.13 | -0.16 | 0.13 | -0.04 | 0.01 | 0.16 | 0.427 |
| SYT7 | S52 | 0.02 | 0.15 | 0.10 | -0.02 | -0.03 | -0.25 | 0.087 |
| SYTL1 | S159 | 0.02 | -0.16 | -0.23 | 0.08 | 0.37 | -0.17 | 0.291 |
| SYTL1 | S216 | -0.16 | 0.11 | -0.23 | 0.35 | -0.05 | -0.09 | 0.406 |
| SYTL1 | S392 | -0.09 | -0.07 | 0.04 | 0.07 | -0.06 | 0.09 | 0.324 |
| SYTL2 | S321 | -0.10 | 0.07 | -0.18 | 0.22 | -0.22 | 0.15 | 0.48 |
| SYTL2 | S535 | -0.06 | -0.04 | -0.06 | 0.02 | 0.01 | 0.12 | 0.046 |
| SYTL2 | S535 | -0.07 | 0.05 | 0.12 | -0.19 | 0.04 | 0.03 | 0.487 |
| SYTL2 | S316 | 0.23 | -0.19 | 0.10 | -0.55 | 0.10 | 0.17 | 0.624 |
| SYTL2 | S571 | 0.16 | 0.13 | 0.05 | -0.17 | -0.05 | -0.14 | 0.009 |
| SYTL2 | S535 | -0.17 | 0.24 | 0.29 | -0.38 | -0.07 | -0.02 | 0.208 |
| SYTL2 | T578 | 0.26 | -0.02 | 0.18 | -0.32 | -0.16 | -0.03 | 0.058 |
| SYTL3 | S231 | -0.48 | 0.30 | -0.17 | 0.01 | 0.34 | -0.17 | 0.56 |
| SYTL4 | S199; T206 | 0.01 | -0.21 | -0.10 | -0.01 | -0.10 | 0.34 | 0.299 |
| SYTL4 | S488 | -0.08 | -0.13 | 0.03 | 0.11 | 0.09 | -0.04 | 0.163 |
| SYTL4 | S204 | -0.13 | 0.01 | 0.20 | -0.08 | -0.02 | 0.00 | 0.592 |
| SYVN1 | S613 | -0.01 | 0.02 | 0.08 | 0.06 | -0.17 | 0.00 | 0.392 |
| TACC2 | S2226 | -0.03 | 0.42 | -0.06 | 0.07 | -0.23 | -0.28 | 0.251 |
| TACC2 | S2317; S2321 | 0.28 | 0.40 | 0.08 | -0.26 | -0.29 | -0.40 | 0.005 |
| TAF1 | S/T | -0.52 | -0.13 | -0.22 | 0.28 | -0.34 | 0.62 | 0.196 |
| TAF1 | S307 | 0.09 | 0.05 | -0.14 | 0.12 | 0.02 | -0.15 | 0.958 |
| TAF1 | S1155; S1158 | 0.08 | -0.05 | -0.01 | -0.07 | -0.20 | 0.22 | 0.851 |
| TAF1 | S324 | 0.16 | -0.01 | 0.30 | -0.18 | 0.11 | -0.52 | 0.159 |

| | | | | | | | | |
|---|---|---|---|---|---|---|---|---|
| **TAF1** | **T/S** | 0.32 | -0.07 | 0.16 | -1.03 | -0.22 | 0.41 | 0.393 |
| **TAF1** | **S1152** | 0.17 | 0.20 | 0.28 | -0.38 | -0.02 | -0.39 | 0.018 |
| **TAF10** | **S44** | -0.01 | 0.00 | 0.45 | -0.12 | -0.15 | -0.29 | 0.105 |
| **TAF11** | **T/S** | 0.08 | -0.08 | -0.11 | -0.07 | 0.23 | -0.08 | 0.622 |
| **TAF12** | **T43; S51** | -0.11 | -0.06 | -0.05 | 0.20 | -0.13 | 0.13 | 0.246 |
| **TAF12** | **T43** | 0.02 | -0.08 | -0.02 | -0.06 | -0.25 | 0.33 | 0.864 |
| **TAF12** | **S51** | 0.03 | 0.00 | 0.03 | -0.03 | 0.03 | -0.07 | 0.229 |
| **TAF12** | **S51** | 0.05 | 0.01 | 0.05 | -0.02 | 0.13 | -0.25 | 0.5 |
| **TAF12** | **S51** | 0.33 | 0.31 | 0.11 | 0.33 | -0.20 | -1.72 | 0.28 |
| **TAF15** | **S97** | -0.45 | -0.14 | 0.08 | -0.30 | -0.28 | 0.74 | 0.583 |
| **TAF15** | **S180** | 0.00 | 0.08 | -0.31 | 0.11 | -0.06 | 0.14 | 0.354 |
| **TAF15** | **S438** | 0.08 | 0.26 | 0.31 | -0.41 | 0.21 | -0.74 | 0.138 |
| **TAF15** | **S226** | 0.30 | 0.26 | 0.31 | -0.46 | -0.10 | -0.57 | 0.009 |
| **TAF1D** | **S204** | -0.07 | 0.10 | 0.15 | -0.06 | -0.08 | -0.06 | 0.146 |
| **TAF1D** | **S206** | -0.04 | 0.24 | 0.13 | -0.20 | 0.01 | -0.21 | 0.088 |
| **TAF2** | **S/T** | -0.45 | 0.10 | -0.38 | 0.57 | -0.16 | 0.08 | 0.217 |
| **TAF2** | **S617; S623** | -0.42 | 0.00 | -0.11 | -0.12 | -0.16 | 0.59 | 0.36 |
| **TAF3** | **S667** | -0.24 | 0.11 | -1.23 | 0.23 | -0.12 | 0.64 | 0.201 |
| **TAF3** | **S345** | -0.18 | 0.05 | -0.43 | 0.34 | -0.06 | 0.15 | 0.142 |
| **TAF3** | **S518** | -0.07 | -0.34 | -0.09 | 0.16 | 0.06 | 0.21 | 0.034 |
| **TAF3** | **S183** | -0.13 | -0.10 | -0.17 | 0.32 | -0.11 | 0.12 | 0.118 |
| **TAF3** | **S183** | 0.00 | -0.05 | -0.11 | 0.12 | -0.02 | 0.05 | 0.115 |
| **TAF3** | **T421; S422** | 0.02 | -0.07 | -0.02 | 0.06 | -0.25 | 0.22 | 0.836 |
| **TAF3** | **S415** | 0.01 | 0.05 | -0.04 | 0.09 | -0.03 | -0.09 | 0.798 |
| **TAF3** | **T364** | 0.06 | 0.00 | 0.11 | 0.04 | -0.13 | -0.09 | 0.13 |
| **TAF3** | **T433** | 0.09 | 0.10 | 0.09 | -0.08 | 0.11 | -0.37 | 0.217 |
| **TAF3** | **S297** | 0.18 | 0.06 | 0.12 | 0.10 | 0.03 | -0.64 | 0.286 |
| **TAF3** | **S297** | 0.19 | 0.13 | 0.15 | -0.18 | 0.06 | -0.46 | 0.079 |
| **TAF4** | **S/T** | -0.40 | -0.06 | 0.46 | -0.23 | 0.34 | -0.34 | 0.828 |
| **TAF6** | **S672** | 0.14 | -0.14 | -0.24 | -0.52 | -0.01 | 0.54 | 0.819 |
| **TAF6** | **S626** | -0.15 | 0.00 | 0.08 | 0.11 | 0.05 | -0.11 | 0.689 |
| **TAF6** | **S626; S634** | -0.08 | 0.04 | 0.00 | 0.04 | 0.04 | -0.03 | 0.49 |
| **TAF6** | **S532** | 0.16 | -0.07 | -0.11 | 0.09 | -0.25 | 0.13 | 0.975 |
| **TAF6** | **S672** | -0.03 | -0.08 | 0.14 | 0.05 | -0.20 | 0.09 | 0.806 |
| **TAF6** | **S653; T660** | 0.04 | -0.03 | 0.05 | 0.16 | -0.06 | -0.19 | 0.653 |
| **TAF6** | **S653** | 0.03 | -0.03 | 0.11 | -0.12 | 0.03 | -0.04 | 0.262 |
| **TAF6** | **S672** | -0.25 | 0.11 | 0.31 | 0.16 | 0.10 | -0.62 | 0.591 |
| **TAF7** | **S213** | -0.42 | -0.13 | -0.12 | 0.16 | 0.11 | 0.29 | 0.021 |
| **TAF7** | **S201** | -0.08 | -0.25 | -0.02 | 0.23 | -0.21 | 0.24 | 0.275 |
| **TAF7** | **S159** | -0.02 | 0.02 | -0.09 | -0.07 | -0.14 | 0.25 | 0.747 |
| **TAF7** | **S201** | 0.04 | -0.08 | 0.04 | 0.16 | -0.27 | 0.07 | 0.941 |
| **TAF7** | **S264** | -0.20 | 0.50 | -0.11 | -0.05 | 0.08 | -0.40 | 0.513 |
| **TAF7** | **S264** | 0.40 | 0.23 | -0.06 | -0.37 | 0.22 | -0.72 | 0.193 |
| **TANC1** | **S214** | -0.22 | 0.28 | -0.11 | -0.23 | -0.27 | 0.40 | 0.958 |
| **TANC1** | **T687** | 0.03 | 0.07 | 0.10 | -0.21 | -0.28 | 0.23 | 0.395 |
| **TANC2** | **S1530** | -0.01 | -0.22 | -0.22 | 0.15 | 0.09 | 0.16 | 0.016 |
| **TANK** | **S208** | -0.27 | -0.36 | 0.03 | 0.17 | -0.25 | 0.49 | 0.233 |
| **TANK** | **S129** | -0.05 | -0.07 | 0.13 | 0.11 | -0.14 | 0.00 | 0.882 |

| | | | | | | | | |
|---|---|---|---|---|---|---|---|---|
| **TAOK1** | **S/T** | 0.03 | -0.28 | 0.07 | -0.18 | 0.05 | 0.25 | 0.572 |
| **TAOK1** | **S421** | -0.07 | 0.07 | -0.12 | 0.06 | 0.00 | 0.04 | 0.29 |
| **TAOK1** | **S965** | -0.04 | 0.01 | 0.38 | -0.23 | -0.06 | -0.15 | 0.14 |
| **TAOK2** | **S827** | 0.11 | 0.13 | 0.13 | -0.12 | 0.08 | -0.41 | 0.125 |
| **TAOK2** | **S827** | 0.21 | 0.08 | 0.14 | -0.27 | 0.17 | -0.46 | 0.159 |
| **TARDBP** | **S** | -0.40 | 0.26 | 0.09 | -0.36 | 0.18 | 0.10 | 0.987 |
| **TASOR** | **S673** | -0.46 | -0.15 | -0.65 | 0.28 | -0.07 | 0.65 | 0.05 |
| **TASOR** | **S971; S979** | -0.62 | 0.00 | -0.29 | 0.32 | 0.33 | 0.03 | 0.062 |
| **TASOR** | **S1651** | 0.05 | -0.19 | -0.38 | 0.05 | 0.10 | 0.28 | 0.087 |
| **TASOR** | **T/S** | -0.10 | -0.13 | -0.19 | -0.21 | 0.25 | 0.29 | 0.194 |
| **TASOR** | **S927** | -0.09 | -0.33 | 0.07 | 0.11 | 0.05 | 0.14 | 0.142 |
| **TASOR** | **S1103** | -0.08 | -0.09 | -0.15 | 0.17 | -0.04 | 0.15 | 0.047 |
| **TASOR** | **S945** | -0.01 | -0.08 | -0.16 | 0.18 | -0.45 | 0.38 | 0.66 |
| **TASOR** | **S927** | 0.05 | -0.06 | -0.15 | 0.07 | -0.03 | 0.11 | 0.21 |
| **TASOR** | **S945** | 0.17 | 0.09 | -0.42 | 0.09 | -0.24 | 0.19 | 0.78 |
| **TASOR** | **S/T/Y** | -0.02 | -0.08 | 0.03 | 0.02 | -0.15 | 0.18 | 0.685 |
| **TASOR** | **S1103** | 0.04 | -0.03 | -0.02 | -0.03 | 0.00 | 0.02 | 0.94 |
| **TASOR** | **S945** | 0.13 | -0.08 | 0.06 | 0.15 | -0.25 | -0.05 | 0.551 |
| **TASOR** | **S1552** | -0.14 | 0.19 | 0.07 | 0.01 | -0.07 | -0.08 | 0.416 |
| **TASOR** | **S/T** | 0.13 | -0.04 | 0.09 | -0.19 | 0.14 | -0.17 | 0.332 |
| **TASOR** | **S633** | 0.09 | 0.05 | 0.06 | 0.03 | -0.03 | -0.23 | 0.151 |
| **TASOR** | **S633** | 0.10 | 0.14 | 0.04 | -0.01 | -0.14 | -0.15 | 0.021 |
| **TASOR** | **S633** | 0.08 | 0.14 | 0.14 | -0.07 | -0.11 | -0.21 | 0.006 |
| **TASOR** | **S636** | 0.19 | 0.14 | 0.03 | 0.01 | -0.08 | -0.35 | 0.089 |
| **TASOR** | **S979** | 0.22 | -0.01 | 0.15 | -0.31 | 0.19 | -0.36 | 0.212 |
| **TASOR** | **S633** | 0.33 | 0.12 | -0.07 | 0.17 | -0.21 | -0.46 | 0.248 |
| **TASOR** | **S1552** | 0.07 | 0.22 | 0.22 | -0.45 | 0.10 | -0.29 | 0.089 |
| **TASOR** | **S633** | 0.24 | 0.20 | 0.15 | -0.15 | -0.02 | -0.57 | 0.059 |
| **TASOR** | **T819** | 0.17 | 0.35 | 0.25 | -0.25 | -0.22 | -0.49 | 0.004 |
| **TASOR2** | **S675** | -0.21 | -0.38 | -0.28 | -0.03 | 0.18 | 0.52 | 0.037 |
| **TASOR2** | **S/Y/T** | -0.32 | -0.30 | -0.04 | 0.18 | 0.26 | 0.12 | 0.015 |
| **TASOR2** | **S219** | -0.22 | -0.22 | -0.14 | 0.12 | -0.03 | 0.39 | 0.049 |
| **TASOR2** | **S1172** | -0.19 | -0.16 | -0.09 | 0.13 | -0.03 | 0.28 | 0.042 |
| **TASOR2** | **S1087** | -0.09 | -0.03 | -0.20 | -0.16 | 0.06 | 0.35 | 0.291 |
| **TASOR2** | **S/Y** | 0.06 | -0.08 | -0.24 | 0.15 | 0.09 | -0.02 | 0.195 |
| **TASOR2** | **T/S** | -0.04 | -0.03 | -0.13 | 0.05 | 0.00 | 0.13 | 0.072 |
| **TASOR2** | **S2037** | -0.03 | -0.07 | -0.02 | 0.04 | 0.02 | 0.06 | 0.013 |
| **TASOR2** | **S685** | 0.00 | 0.03 | -0.10 | 0.03 | -0.12 | 0.15 | 0.643 |
| **TASOR2** | **S1848** | -0.24 | -0.17 | 0.29 | 0.37 | -0.05 | -0.35 | 0.929 |
| **TASOR2** | **S384** | -0.03 | -0.03 | 0.03 | -0.10 | -0.04 | 0.15 | 0.86 |
| **TASOR2** | **S340** | 0.15 | -0.06 | 0.00 | 0.02 | -0.04 | -0.08 | 0.445 |
| **TASOR2** | **S2009** | 0.14 | 0.15 | -0.01 | 0.11 | -0.10 | -0.35 | 0.217 |
| **TASOR2** | **S685** | 0.14 | 0.15 | 0.17 | -0.18 | -0.01 | -0.34 | 0.024 |
| **TASOR2** | **S1541** | 0.07 | 0.19 | 0.43 | -0.25 | -0.30 | -0.31 | 0.009 |
| **TASOR2** | **S2037** | 0.54 | 0.28 | 0.30 | -1.50 | 0.35 | -0.27 | 0.198 |
| **TBC1D10A** | **S43** | -0.38 | 0.04 | 0.04 | -0.15 | 0.07 | 0.28 | 0.423 |
| **TBC1D10A** | **S18** | -0.23 | -0.08 | 0.07 | 0.10 | 0.04 | 0.08 | 0.155 |
| **TBC1D10A** | **S40** | -0.25 | -0.02 | 0.13 | -0.23 | 0.04 | 0.25 | 0.731 |

| | | | | | | | | |
|---|---|---|---|---|---|---|---|---|
| TBC1D10A | S497 | -0.11 | 0.05 | -0.05 | 0.10 | -0.06 | 0.05 | 0.389 |
| TBC1D10A | S40; S43 | -0.16 | 0.09 | 0.11 | -0.24 | -0.10 | 0.24 | 0.803 |
| TBC1D10A | S40; S43 | -0.04 | 0.01 | 0.11 | 0.02 | -0.17 | 0.05 | 0.499 |
| TBC1D10A | S317 | 0.18 | 0.12 | 0.02 | -0.44 | 0.13 | -0.11 | 0.226 |
| TBC1D10B | S22 | -0.05 | -0.07 | 0.15 | -0.30 | -0.26 | 0.40 | 0.809 |
| TBC1D10B | S132 | 0.08 | 0.06 | 0.10 | -0.06 | -0.22 | 0.02 | 0.088 |
| TBC1D10B | S678 | 0.09 | 0.12 | 0.16 | -0.13 | -0.13 | -0.14 | 2E-04 |
| TBC1D10B | T148 | 0.32 | -0.06 | 0.37 | -0.52 | -0.12 | -0.18 | 0.057 |
| TBC1D10B | S22 | 0.01 | 0.97 | -0.32 | -0.28 | -0.70 | -0.32 | 0.189 |
| TBC1D16 | S260 | -0.41 | -0.36 | -0.25 | 0.42 | 0.17 | 0.23 | 0.002 |
| TBC1D16 | S103 | -0.19 | 0.07 | -0.31 | 0.27 | 0.14 | -0.06 | 0.147 |
| TBC1D17 | T620 | -0.12 | 0.11 | -0.05 | -0.02 | -0.09 | 0.15 | 0.752 |
| TBC1D22A | S/T | 0.27 | 0.23 | -0.63 | 0.10 | -0.11 | -0.03 | 0.929 |
| TBC1D30 | S114 | -0.05 | 0.03 | -0.02 | 0.07 | 0.05 | -0.08 | 0.681 |
| TBC1D30 | S800 | 0.08 | -0.03 | 0.05 | -0.22 | -0.26 | 0.30 | 0.644 |
| TBC1D30 | S800 | -0.04 | 0.11 | 0.18 | -0.05 | -0.13 | -0.10 | 0.067 |
| TBC1D30 | S713 | 0.11 | 0.21 | 0.10 | -0.09 | 0.07 | -0.51 | 0.147 |
| TBC1D4 | S591 | 0.07 | 0.05 | 0.24 | -0.17 | 0.03 | -0.26 | 0.073 |
| TBC1D5 | S554 | 0.31 | 0.19 | 0.17 | -0.22 | -0.11 | -0.49 | 0.015 |
| TBC1D9B | S435 | 0.03 | -0.07 | -0.18 | -0.28 | 0.26 | 0.16 | 0.532 |
| TBC1D9B | S435 | 0.02 | -0.13 | 0.08 | -0.01 | -0.10 | 0.12 | 0.881 |
| TBC1D9B | S435 | -0.21 | 0.45 | -0.08 | -0.04 | -0.01 | -0.23 | 0.516 |
| TBKBP1 | S365 | -0.63 | 0.10 | 0.36 | 0.30 | -0.27 | -0.09 | 0.918 |
| TBKBP1 | S534 | 0.01 | 0.22 | -0.13 | 0.01 | 0.06 | -0.20 | 0.578 |
| TBKBP1 | S388 | -0.54 | 0.20 | 0.25 | -0.06 | 0.63 | -1.10 | 0.81 |
| TBKBP1 | S365 | -0.14 | 0.25 | 0.21 | -0.04 | -0.10 | -0.24 | 0.167 |
| TBRG1 | S405 | -0.21 | -0.25 | -0.03 | 0.55 | -0.22 | -0.02 | 0.326 |
| TBX3 | S371; S375 | 0.22 | 0.03 | 0.03 | 0.05 | -0.01 | -0.37 | 0.23 |
| TBX3 | S409 | 0.14 | 0.13 | 0.04 | -0.10 | -0.02 | -0.23 | 0.033 |
| TBX3 | S432 | 0.14 | 0.14 | 0.13 | -0.12 | -0.07 | -0.28 | 0.01 |
| TBX3 | S438 | 0.01 | 0.14 | 0.35 | -0.57 | -0.01 | -0.08 | 0.129 |
| TBX3 | S/T | -0.09 | 0.45 | 0.34 | -0.09 | 0.10 | -1.27 | 0.226 |
| TBX3 | S/T | 0.44 | 0.38 | 0.47 | -0.64 | -0.45 | -0.78 | 4E-04 |
| TC2N | S168 | -0.05 | -0.05 | -0.11 | 0.03 | -0.06 | 0.22 | 0.199 |
| TC2N | S218 | 0.01 | -0.09 | -0.09 | 0.08 | 0.07 | 0.01 | 0.056 |
| TC2N | S327 | 0.09 | -0.21 | -0.08 | -0.26 | 0.38 | -0.01 | 0.64 |
| TC2N | S211; S218 | 0.07 | 0.17 | -0.03 | 0.05 | -0.05 | -0.26 | 0.233 |
| TC2N | S218 | 0.24 | 0.09 | 0.14 | 0.08 | -0.18 | -0.49 | 0.103 |
| TCEA1 | S100 | -0.09 | 0.03 | -0.23 | 0.07 | 0.02 | 0.16 | 0.096 |
| TCEA1 | S100 | 0.03 | -0.11 | -0.07 | 0.02 | -0.06 | 0.18 | 0.307 |
| TCEA1 | S100 | 0.21 | -0.02 | -0.18 | -0.01 | 0.02 | -0.04 | 0.895 |
| TCEA3 | S81 | 0.01 | -0.61 | -0.25 | -0.08 | -0.52 | 0.89 | 0.448 |
| TCEA3 | S115 | 0.15 | 0.11 | 0.07 | -0.14 | -0.18 | -0.03 | 0.013 |
| TCEA3 | S115 | 0.23 | 0.25 | 0.17 | -0.05 | -0.27 | -0.49 | 0.019 |
| TCEANC2 | S14 | 0.06 | -0.03 | 0.12 | -0.08 | -0.05 | -0.04 | 0.087 |
| TCF12 | S333 | -0.61 | -0.28 | -0.09 | 0.25 | 0.17 | 0.34 | 0.023 |
| TCF12 | S67 | -0.37 | 0.35 | -0.28 | 0.07 | 0.23 | -0.14 | 0.564 |
| TCF20 | S559 | -0.18 | -0.11 | -0.10 | -0.11 | -0.10 | 0.48 | 0.329 |

| | | | | | | | | |
|---|---|---|---|---|---|---|---|---|
| TCF20 | S1005 | -0.02 | -0.31 | 0.05 | 0.07 | -0.07 | 0.23 | 0.291 |
| TCF20 | T1671 | -0.06 | -0.14 | -0.04 | 0.09 | -0.05 | 0.18 | 0.098 |
| TCF20 | S871 | 0.03 | -0.08 | -0.07 | 0.05 | 0.02 | 0.05 | 0.081 |
| TCF20 | S1005 | 0.04 | -0.01 | -0.10 | 0.11 | -0.20 | 0.14 | 0.765 |
| TCF20 | S640 | 0.00 | 0.05 | -0.08 | 0.02 | -0.13 | 0.13 | 0.821 |
| TCF20 | S538 | -0.09 | 0.12 | -0.02 | 0.04 | -0.07 | 0.01 | 0.857 |
| TCF20 | T/S | 0.09 | 0.03 | -0.05 | -0.02 | -0.02 | -0.04 | 0.282 |
| TCF20 | T/S | 0.07 | 0.10 | -0.03 | 0.02 | -0.05 | -0.11 | 0.177 |
| TCF20 | S1522 | 0.12 | 0.07 | 0.02 | -0.10 | -0.01 | -0.12 | 0.034 |
| TCF20 | S1335 | 0.35 | -0.41 | 0.36 | -0.39 | -0.18 | 0.07 | 0.412 |
| TCF20 | S640 | 0.21 | 0.15 | 0.05 | -0.10 | -0.14 | -0.22 | 0.008 |
| TCF20 | S1335 | 0.13 | 0.18 | 0.30 | -0.19 | -0.15 | -0.40 | 0.009 |
| TCF7L2 | S60 | -0.15 | -0.11 | -0.31 | 0.20 | -0.01 | 0.29 | 0.033 |
| TCF7L2 | S179 | -0.22 | 0.46 | -0.35 | -0.30 | -0.13 | 0.32 | 0.999 |
| TCIRG1 | S691 | -0.23 | -0.27 | -0.32 | 0.36 | -0.21 | 0.45 | 0.086 |
| TCIRG1 | S685 | -0.06 | -0.07 | 0.12 | -0.01 | -0.33 | 0.28 | 0.953 |
| TCOF1 | S1410 | -0.58 | -0.83 | -0.63 | 0.50 | 0.46 | 0.42 | 1E-04 |
| TCOF1 | S1410 | -0.49 | -0.75 | -0.64 | 0.48 | 0.36 | 0.46 | 2E-04 |
| TCOF1 | S1376; S1378 | -0.18 | -0.33 | -0.28 | 0.29 | -0.23 | 0.51 | 0.116 |
| TCOF1 | S328 | -0.25 | -0.25 | -0.09 | -0.24 | 0.14 | 0.51 | 0.215 |
| TCOF1 | S906 | 0.13 | -0.36 | -0.14 | 0.03 | 0.05 | 0.21 | 0.219 |
| TCOF1 | T533 | 0.01 | -0.22 | 0.16 | -0.05 | -0.01 | 0.09 | 0.852 |
| TCOF1 | S/T | 0.14 | -0.02 | 0.10 | -0.04 | -0.10 | -0.09 | 0.039 |
| TCOF1 | T249 | -0.03 | 0.15 | 0.09 | 0.10 | -0.22 | -0.13 | 0.226 |
| TCOF1 | S87; S88 | 0.10 | 0.13 | 0.03 | -0.11 | -0.07 | -0.10 | 0.005 |
| TCOF1 | T914 | 0.26 | -0.06 | 0.12 | -0.24 | 0.02 | -0.16 | 0.121 |
| TCOF1 | S1378 | 0.23 | 0.13 | 0.02 | -0.24 | 0.00 | -0.21 | 0.046 |
| TCOF1 | S233 | 0.29 | 0.04 | 0.12 | -0.35 | -0.02 | -0.17 | 0.051 |
| TCOF1 | S1350 | 0.29 | 0.13 | 0.08 | -0.26 | -0.07 | -0.24 | 0.015 |
| TCOF1 | S446 | 0.31 | 0.09 | 0.12 | -0.22 | -0.22 | -0.16 | 0.007 |
| TCOF1 | S/T | 0.23 | 0.19 | 0.11 | -0.26 | -0.16 | -0.18 | 0.001 |
| TCOF1 | S967 | 0.18 | -0.16 | 0.48 | 0.21 | -0.51 | -0.47 | 0.224 |
| TCOF1 | S1378 | 0.25 | 0.14 | 0.23 | -0.26 | -0.04 | -0.45 | 0.02 |
| TCOF1 | S446 | 0.39 | 0.06 | 0.24 | -0.38 | -0.03 | -0.46 | 0.033 |
| TCOF1 | S/T | 0.40 | 0.13 | 0.26 | -0.40 | -0.10 | -0.52 | 0.014 |
| TCOF1 | S1111 | 0.24 | 0.35 | 0.23 | -0.35 | -0.29 | -0.38 | 2E-04 |
| TCOF1 | S381 | 0.34 | 0.25 | 0.33 | -0.48 | -0.16 | -0.56 | 0.005 |
| TCOF1 | S381 | 0.23 | 0.26 | 0.48 | -0.57 | -0.40 | -0.31 | 0.002 |
| TCOF1 | T1106 | 0.72 | 0.31 | 0.02 | -0.74 | -0.65 | -0.20 | 0.029 |
| TCP1 | S544 | -0.17 | -0.37 | 0.17 | -0.02 | 0.20 | 0.11 | 0.281 |
| TDP1 | S61 | 0.07 | -0.03 | 0.37 | 0.22 | -0.59 | -0.22 | 0.274 |
| TEF | S140 | 0.32 | 0.09 | 0.17 | -0.22 | -0.18 | -0.30 | 0.005 |
| TEF | S140 | 0.32 | 0.27 | 0.17 | -0.21 | -0.31 | -0.42 | 0.002 |
| TELO2 | S637 | -0.33 | 0.05 | 0.05 | -0.07 | 0.13 | 0.11 | 0.406 |
| TENT4B | S484 | -0.75 | -0.27 | -0.45 | 0.21 | 0.18 | 0.64 | 0.016 |
| TENT4B | S48; S71 | 0.14 | -0.91 | -0.43 | 0.32 | -0.03 | 0.48 | 0.127 |
| TENT4B | S49 | 0.19 | -0.04 | -0.17 | 0.26 | -0.37 | 0.04 | 0.947 |
| TENT4B | S49 | 0.37 | 0.06 | 0.00 | -0.14 | -0.31 | -0.07 | 0.08 |

| | | | | | | | | |
|---|---|---|---|---|---|---|---|---|
| TERF1 | S279 | -0.60 | -0.99 | -0.79 | 0.42 | 0.75 | 0.31 | 0.002 |
| TERF1 | S434; S437 | 0.00 | -0.08 | 0.05 | -0.07 | 0.13 | -0.04 | 0.849 |
| TERF1 | S434 | -0.09 | -0.02 | 0.20 | -0.17 | -0.10 | 0.15 | 0.62 |
| TERF1 | T371 | 0.22 | 0.10 | 0.12 | -0.16 | -0.06 | -0.28 | 0.012 |
| TERF2 | S365 | 0.06 | -0.12 | -0.32 | 0.07 | 0.05 | 0.20 | 0.132 |
| TERF2IP | T230 | -0.83 | -0.88 | -1.11 | 0.71 | 0.33 | 0.62 | 5E-04 |
| TERF2IP | T230 | -0.80 | -0.73 | -0.94 | 0.60 | 0.35 | 0.59 | 2E-04 |
| TERF2IP | S203 | 0.24 | 0.17 | -0.04 | 0.05 | -0.14 | -0.37 | 0.133 |
| TESK1 | S437; S | -0.06 | -0.25 | -0.15 | 0.30 | 0.05 | 0.04 | 0.045 |
| TESK1 | S441 | -0.11 | 0.19 | 0.07 | 0.13 | -0.06 | -0.28 | 0.461 |
| TESK2 | S346 | -0.14 | -0.23 | -0.05 | 0.18 | -0.04 | 0.23 | 0.056 |
| TEX2 | S538 | -0.36 | -0.13 | -0.05 | 0.23 | 0.03 | 0.21 | 0.041 |
| TEX2 | S802 | 0.10 | -0.15 | -0.46 | -0.26 | 0.39 | 0.21 | 0.331 |
| TEX2 | S196 | 0.03 | -0.03 | -0.16 | 0.05 | 0.00 | 0.09 | 0.195 |
| TEX2 | S/T | 0.03 | 0.46 | -0.50 | -0.79 | 0.21 | 0.22 | 0.814 |
| TEX264 | S244 | 0.18 | 0.12 | 0.18 | -0.21 | -0.05 | -0.29 | 0.009 |
| TFAM | S124 | 0.01 | -0.07 | 0.00 | -0.04 | -0.09 | 0.18 | 0.712 |
| TFAM | S193 | 0.18 | 0.13 | 0.00 | -0.17 | 0.09 | -0.29 | 0.135 |
| TFAM | S195 | 0.13 | 0.11 | 0.09 | -0.12 | 0.10 | -0.38 | 0.163 |
| TFAP2A | S223 | 0.11 | -0.15 | 0.09 | 0.10 | -0.14 | -0.03 | 0.733 |
| TFAP2B | T/S | 0.13 | -0.04 | 0.14 | -0.05 | -0.11 | -0.10 | 0.054 |
| TFAP2C | S434 | -0.19 | -0.03 | -0.14 | 0.11 | 0.01 | 0.20 | 0.035 |
| TFAP2C | S434 | 0.14 | 0.37 | -0.60 | 0.13 | -0.05 | -0.17 | 0.996 |
| TFAP2C | S434 | 0.06 | -0.01 | 0.08 | 0.05 | -0.04 | -0.15 | 0.214 |
| TFAP2C | S434 | 0.11 | 0.13 | -0.02 | -0.13 | 0.07 | -0.19 | 0.162 |
| TFAP4 | S139 | -0.05 | -0.15 | -0.05 | -0.01 | -0.12 | 0.32 | 0.333 |
| TFAP4 | S139 | 0.07 | -0.12 | -0.08 | -0.07 | -0.07 | 0.23 | 0.556 |
| TFAP4 | S139 | 0.14 | -0.18 | -0.07 | 0.11 | -0.12 | 0.10 | 0.607 |
| TFAP4 | S121 | 0.18 | 0.58 | 0.05 | -0.31 | -0.19 | -0.61 | 0.035 |
| TFAP4 | S124 | 0.27 | 0.22 | 0.36 | -0.76 | -0.11 | -0.27 | 0.029 |
| TFB1M | S296 | 0.00 | -0.06 | -0.09 | 0.13 | 0.31 | -0.39 | 0.765 |
| TFDP2 | S42 | -0.21 | -0.23 | -0.17 | 0.20 | 0.06 | 0.26 | 0.004 |
| TFDP2 | S24 | -0.45 | -0.19 | 0.04 | 0.00 | 0.20 | 0.29 | 0.095 |
| TFE3 | S556 | -0.14 | -0.15 | -0.15 | 0.22 | 0.05 | 0.12 | 0.005 |
| TFE3 | S548; S556 | -0.30 | -0.19 | 0.13 | 0.14 | -0.06 | 0.21 | 0.225 |
| TFE3 | S568 | -0.12 | -0.01 | -0.01 | 0.10 | -0.02 | 0.04 | 0.177 |
| TFEB | S423 | -0.05 | -0.11 | 0.14 | 0.34 | 0.20 | -0.75 | 0.864 |
| TFIP11 | S392 | -0.11 | -0.02 | -0.29 | 0.35 | -0.03 | 0.01 | 0.155 |
| TFIP11 | S392 | -0.07 | -0.02 | -0.26 | 0.25 | 0.07 | -0.03 | 0.118 |
| TFIP11 | S98 | 0.02 | -0.07 | -0.13 | 0.09 | -0.11 | 0.18 | 0.286 |
| TFIP11 | S98 | -0.08 | -0.10 | 0.04 | 0.03 | 0.02 | 0.08 | 0.132 |
| TFIP11 | S392 | 0.09 | -0.27 | 0.04 | 0.18 | -0.20 | 0.10 | 0.658 |
| TFIP11 | S568 | 0.19 | 0.01 | -0.27 | -0.18 | 0.22 | -0.02 | 0.887 |
| TFIP11 | S98 | 0.11 | 0.05 | -0.19 | 0.08 | -0.20 | 0.11 | 0.976 |
| TFIP11 | S98 | 0.13 | 0.01 | -0.04 | 0.06 | -0.10 | -0.08 | 0.384 |
| TFIP11 | S59 | 0.15 | 0.12 | 0.02 | -0.01 | -0.04 | -0.28 | 0.094 |
| TFIP11 | S210 | 0.11 | 0.14 | 0.12 | -0.10 | 0.01 | -0.36 | 0.07 |
| TFIP11 | S210 | 0.17 | 0.20 | 0.29 | -0.19 | 0.02 | -0.70 | 0.077 |

| | | | | | | | | |
|---|---|---|---|---|---|---|---|---|
| TFPT | S/T | -0.26 | -0.70 | 0.21 | -0.55 | -0.21 | 0.88 | 0.595 |
| TFPT | S180 | -0.10 | -0.09 | -0.14 | 0.02 | -0.08 | 0.33 | 0.187 |
| TFPT | T207 | 0.03 | 0.01 | -0.27 | 0.01 | 0.19 | 0.00 | 0.269 |
| TFPT | S/T | -0.36 | 0.09 | 0.08 | 0.16 | 0.10 | -0.12 | 0.548 |
| TFPT | S180 | -0.10 | -0.01 | -0.06 | 0.04 | -0.02 | 0.13 | 0.097 |
| TFPT | S167 | 0.06 | -0.02 | -0.08 | -0.05 | -0.01 | 0.09 | 0.722 |
| TFPT | S252 | 0.09 | 0.37 | -0.37 | -0.07 | 0.00 | -0.14 | 0.677 |
| TFPT | T/S | 0.17 | 0.07 | 0.01 | 0.15 | -0.22 | -0.22 | 0.242 |
| TFRC | S34 | -0.46 | 0.07 | 0.28 | 0.15 | -0.07 | -0.07 | 0.872 |
| TGFBR1 | S165 | -0.30 | 0.31 | 0.06 | 0.12 | -0.09 | -0.18 | 0.726 |
| TGFBR1 | S165 | -0.30 | 0.31 | 0.06 | 0.12 | -0.09 | -0.18 | 0.726 |
| TGFBRAP1 | S/T | 0.00 | 0.52 | 0.34 | -0.39 | -0.32 | -0.45 | 0.013 |
| TGM1 | S24 | -0.51 | -0.37 | -0.26 | 0.24 | 0.04 | 0.57 | 0.017 |
| TGM1 | S24 | -0.49 | -0.32 | -0.29 | 0.34 | 0.11 | 0.41 | 0.004 |
| TGOLN2 | S71 | 0.51 | 0.25 | 0.16 | -0.50 | -0.33 | -0.39 | 0.004 |
| TGOLN2 | S71 | 0.47 | 0.47 | 0.15 | -0.67 | -0.26 | -0.60 | 0.006 |
| TGOLN2 | S351 | 0.37 | 0.30 | 0.48 | -0.62 | -0.27 | -0.72 | 0.003 |
| TGS1 | S154 | -0.53 | -0.08 | -0.10 | 0.20 | -0.06 | 0.40 | 0.103 |
| TH | S/T/Y | 0.44 | 0.27 | 0.28 | -0.31 | -0.16 | -0.95 | 0.032 |
| THAP12 | S141 | -0.16 | -0.11 | -0.04 | 0.26 | 0.17 | -0.18 | 0.249 |
| THAP9 | S346 | -0.26 | -0.05 | 0.03 | 0.04 | -0.10 | 0.29 | 0.307 |
| THOC1 | T542 | 0.16 | 0.19 | -0.27 | 0.05 | -0.20 | 0.02 | 0.712 |
| THOC1 | S560 | 0.23 | 0.01 | -0.03 | -0.06 | 0.02 | -0.21 | 0.222 |
| THOC2 | T1385; S1393 | -0.33 | -0.08 | -0.12 | 0.37 | 0.03 | 0.04 | 0.078 |
| THOC2 | T1285; T | -0.22 | -0.07 | -0.01 | 0.15 | 0.05 | 0.06 | 0.054 |
| THOC2 | T1285 | -0.04 | 0.04 | 0.05 | 0.18 | -0.14 | -0.11 | 0.724 |
| THOC2 | S1516 | 0.15 | -0.03 | 0.03 | 0.34 | -0.24 | -0.37 | 0.566 |
| THOC2 | S1516 | 0.13 | 0.08 | 0.00 | 0.05 | -0.12 | -0.17 | 0.127 |
| THOC2 | S1516 | 0.10 | 0.15 | -0.03 | 0.08 | -0.15 | -0.18 | 0.177 |
| THOC2 | T1443 | 0.03 | 0.22 | 0.03 | 0.05 | 0.08 | -0.51 | 0.339 |
| THOC2 | S1393 | 0.11 | 0.16 | 0.02 | -0.35 | 0.27 | -0.32 | 0.325 |
| THOC2 | S1417 | -0.01 | 0.29 | 0.10 | -0.53 | -0.10 | 0.12 | 0.227 |
| THOC2 | S/Y | 0.17 | 0.51 | 0.40 | -0.86 | -0.19 | -0.50 | 0.016 |
| THOC3 | S273 | -0.25 | -0.02 | -0.01 | 0.25 | 0.00 | -0.02 | 0.218 |
| THOC5 | T328 | -0.58 | -0.27 | -0.70 | 0.45 | 0.11 | 0.52 | 0.008 |
| THOC5 | T328 | -0.48 | 0.25 | -0.85 | 0.71 | -0.67 | 0.34 | 0.407 |
| THOC5 | S314 | 0.47 | 0.38 | 0.31 | -0.34 | -0.19 | -1.30 | 0.047 |
| THOC5 | S314 | 0.60 | 0.45 | 0.32 | -0.81 | -0.05 | -1.59 | 0.048 |
| THOC7 | S197 | -0.18 | -0.26 | -0.07 | 0.19 | 0.14 | 0.12 | 0.006 |
| THRAP3 | S331 | -0.70 | -0.75 | -0.75 | 0.57 | 0.55 | 0.32 | 1E-04 |
| THRAP3 | S377 | -0.21 | -0.27 | -0.47 | 0.29 | 0.46 | -0.02 | 0.026 |
| THRAP3 | S928; S939 | -0.18 | -0.27 | 0.04 | 0.23 | -0.28 | 0.34 | 0.326 |
| THRAP3 | S939 | -0.19 | -0.14 | -0.08 | 0.07 | -0.25 | 0.47 | 0.33 |
| THRAP3 | S682 | -0.11 | -0.07 | -0.11 | 0.08 | 0.07 | 0.12 | 8E-04 |
| THRAP3 | T874 | -0.18 | -0.05 | -0.04 | -0.01 | 0.12 | 0.14 | 0.06 |
| THRAP3 | S494 | -0.08 | -0.04 | -0.15 | 0.04 | 0.01 | 0.19 | 0.069 |
| THRAP3 | S320; S323 | -0.09 | -0.12 | 0.00 | 0.15 | -0.17 | 0.20 | 0.36 |
| THRAP3 | S379 | -0.06 | -0.12 | -0.02 | 0.22 | -0.11 | 0.05 | 0.292 |

| | | | | | | | | |
|---|---|---|---|---|---|---|---|---|
| THRAP3 | S928 | 0.01 | -0.04 | -0.16 | 0.09 | 0.01 | 0.07 | 0.096 |
| THRAP3 | S232 | 0.03 | 0.02 | -0.23 | 0.14 | 0.05 | -0.04 | 0.307 |
| THRAP3 | S928 | -0.02 | 0.00 | -0.13 | 0.03 | 0.07 | 0.04 | 0.079 |
| THRAP3 | T954 | 0.03 | 0.00 | -0.15 | 0.10 | -0.10 | 0.09 | 0.442 |
| THRAP3 | S928; S939 | 0.01 | -0.12 | 0.02 | 0.16 | -0.22 | 0.11 | 0.722 |
| THRAP3 | S257 | 0.05 | -0.05 | -0.06 | 0.11 | -0.02 | -0.04 | 0.563 |
| THRAP3 | S320 | 0.12 | 0.15 | -0.35 | 0.14 | -0.05 | -0.06 | 0.855 |
| THRAP3 | S240 | 0.03 | -0.04 | 0.00 | 0.02 | -0.03 | 0.03 | 0.754 |
| THRAP3 | S320 | 0.12 | 0.04 | -0.18 | 0.11 | -0.05 | -0.06 | 0.986 |
| THRAP3 | S237; S243 | 0.01 | 0.05 | -0.06 | -0.04 | -0.09 | 0.11 | 0.927 |
| THRAP3 | S805 | 0.02 | 0.09 | -0.05 | 0.00 | 0.04 | -0.10 | 0.539 |
| THRAP3 | S320 | 0.31 | 0.02 | -0.31 | 0.03 | -0.04 | -0.09 | 0.836 |
| THRAP3 | S315; S320 | 0.25 | 0.08 | -0.28 | 0.05 | -0.11 | -0.05 | 0.753 |
| THRAP3 | S315; S320 | 0.32 | -0.09 | -0.11 | 0.04 | -0.20 | -0.02 | 0.559 |
| THRAP3 | S253 | 0.15 | -0.10 | 0.05 | -0.41 | 0.37 | -0.18 | 0.687 |
| THRAP3 | S320 | 0.24 | 0.12 | -0.23 | -0.05 | 0.05 | -0.20 | 0.513 |
| THRAP3 | S320 | 0.35 | 0.14 | -0.37 | 0.01 | 0.10 | -0.38 | 0.649 |
| THRAP3 | S698 | 0.07 | 0.11 | 0.01 | -0.04 | 0.04 | -0.21 | 0.167 |
| THRAP3 | S562 | 0.02 | 0.03 | 0.15 | -0.15 | 0.13 | -0.22 | 0.26 |
| THRAP3 | S248; S253 | 0.14 | 0.12 | 0.01 | -0.07 | 0.01 | -0.24 | 0.087 |
| THRAP3 | S339 | 0.17 | 0.16 | -0.07 | -0.15 | 0.16 | -0.34 | 0.3 |
| THRAP3 | S/T | 0.18 | 0.06 | -0.01 | -0.61 | 0.25 | -0.02 | 0.487 |
| THRAP3 | S786; S/Y | -0.12 | 0.37 | 0.06 | 0.04 | -0.21 | -0.23 | 0.222 |
| THRAP3 | S682 | 0.19 | 0.07 | 0.11 | -0.13 | 0.06 | -0.36 | 0.1 |
| THRAP3 | S698 | 0.06 | 0.18 | 0.16 | -0.09 | 0.06 | -0.45 | 0.131 |
| THRAP3 | S55 | 0.17 | 0.15 | 0.10 | 0.07 | -0.23 | -0.33 | 0.068 |
| THRAP3 | S/Y | 0.46 | 0.00 | -0.07 | 0.09 | -0.53 | -0.13 | 0.263 |
| THRAP3 | S672 | 0.18 | 0.17 | 0.10 | -0.28 | 0.15 | -0.43 | 0.128 |
| THRAP3 | S928 | 0.19 | 0.20 | 0.06 | -0.07 | -0.01 | -0.48 | 0.096 |
| THRAP3 | S928 | 0.24 | 0.19 | 0.12 | -0.11 | -0.01 | -0.57 | 0.08 |
| THRAP3 | S53; S55 | 0.14 | 0.24 | 0.22 | -0.25 | -0.07 | -0.40 | 0.012 |
| THRAP3 | S698 | 0.22 | 0.44 | -0.11 | -0.12 | 0.04 | -0.74 | 0.185 |
| THRAP3 | S53; S55 | 0.23 | 0.34 | 0.15 | -0.24 | -0.15 | -0.50 | 0.01 |
| THRAP3 | S939 | 0.24 | 0.15 | 0.39 | -0.23 | -0.07 | -0.74 | 0.048 |
| THRAP3 | T/S | 0.34 | 0.23 | 0.29 | -0.20 | -0.10 | -0.88 | 0.05 |
| THRAP3 | S/T | 0.27 | 0.24 | 0.35 | -0.49 | 0.05 | -0.75 | 0.045 |
| THRAP3 | S535 | 0.27 | 0.28 | 0.32 | -0.26 | -0.02 | -0.98 | 0.071 |
| THUMPD1 | S/T | 0.10 | 0.16 | 0.19 | -0.20 | -0.02 | -0.30 | 0.02 |
| THUMPD1 | S/T | 0.21 | 0.29 | 0.17 | -0.31 | -0.03 | -0.50 | 0.023 |
| TIAM1 | S231 | -0.32 | -0.05 | -0.27 | 0.25 | 0.10 | 0.21 | 0.013 |
| TIAM1 | S358 | 0.44 | -0.14 | 0.24 | -0.67 | 0.29 | -0.52 | 0.235 |
| TICRR | T1301 | -0.80 | 0.00 | 0.12 | 0.42 | 0.32 | -0.41 | 0.442 |
| TICRR | S441 | -0.12 | 0.12 | -0.25 | -0.23 | 0.08 | 0.32 | 0.505 |
| TICRR | S1413 | -0.25 | 0.18 | -0.07 | -0.04 | -0.33 | 0.38 | 0.839 |
| TICRR | T/S | -0.07 | 0.01 | 0.09 | -0.08 | 0.05 | -0.02 | 0.707 |
| TICRR | S599 | -0.01 | 0.19 | -0.06 | -0.09 | 0.01 | -0.05 | 0.354 |
| TICRR | T1307; S/T | 0.09 | 0.10 | -0.03 | 0.03 | -0.09 | -0.11 | 0.145 |
| TICRR | S1057 | -0.02 | 0.10 | 0.12 | -0.10 | -0.09 | -0.03 | 0.045 |

| | | | | | | | | |
|---|---|---|---|---|---|---|---|---|
| TICRR | S1433 | 0.12 | 0.21 | -0.13 | -0.08 | -0.13 | -0.02 | 0.239 |
| TICRR | S1334 | 0.13 | 0.05 | 0.04 | -0.30 | 0.01 | 0.03 | 0.232 |
| TICRR | S1413 | -0.34 | 0.39 | 0.06 | -0.20 | -0.62 | 0.42 | 0.669 |
| TICRR | T1134 | 0.03 | 0.08 | 0.19 | -0.29 | -0.10 | 0.04 | 0.117 |
| TICRR | S1057; S1064 | 0.16 | 0.11 | 0.05 | -0.20 | -0.10 | -0.05 | 0.015 |
| TICRR | S838; S841 | 0.09 | 0.14 | 0.13 | -0.21 | -0.03 | -0.16 | 0.011 |
| TICRR | S838 | 0.17 | 0.08 | 0.11 | -0.28 | -0.17 | 0.04 | 0.058 |
| TICRR | S1001 | 0.15 | 0.17 | 0.18 | -0.27 | -0.07 | -0.25 | 0.005 |
| TICRR | S292 | 0.22 | 0.21 | 0.05 | -0.61 | 0.05 | -0.07 | 0.15 |
| TIMELESS | S1149 | -0.27 | -0.23 | -0.19 | 0.25 | 0.13 | 0.21 | 5E-04 |
| TIMELESS | S1149 | -0.21 | -0.24 | -0.06 | -0.32 | 0.26 | 0.41 | 0.285 |
| TIMELESS | S1173 | -0.03 | 0.22 | -0.38 | -0.33 | 0.09 | 0.29 | 0.768 |
| TIMELESS | S823 | -0.20 | 0.20 | 0.04 | -0.26 | 0.22 | -0.07 | 0.788 |
| TINF2 | S295 | -0.12 | -0.04 | -0.16 | 0.13 | 0.21 | -0.06 | 0.08 |
| TJAP1 | S300 | -0.08 | -0.07 | -0.14 | 0.18 | 0.06 | 0.03 | 0.022 |
| TJAP1 | S545 | -0.22 | 0.72 | -0.55 | 0.25 | -0.73 | 0.04 | 0.799 |
| TJP1 | S1111 | -0.08 | -0.56 | -0.06 | 0.14 | 0.27 | 0.15 | 0.068 |
| TJP1 | T868 | -0.17 | -0.05 | -0.19 | 0.19 | -0.36 | 0.43 | 0.397 |
| TJP1 | T868 | -0.09 | -0.09 | -0.12 | 0.24 | -0.15 | 0.17 | 0.189 |
| TJP1 | S294; S297; S300 | -0.02 | -0.10 | -0.13 | 0.05 | -0.03 | 0.21 | 0.113 |
| TJP1 | S1617 | -0.12 | -0.04 | -0.09 | 0.16 | -0.09 | 0.15 | 0.146 |
| TJP1 | S297 | 0.02 | -0.12 | -0.14 | 0.10 | -0.11 | 0.22 | 0.241 |
| TJP1 | S1051 | -0.18 | -0.22 | 0.17 | -0.19 | 0.19 | 0.16 | 0.505 |
| TJP1 | T868 | -0.25 | 0.37 | -0.34 | 0.29 | -0.38 | 0.12 | 0.792 |
| TJP1 | S1111 | 0.01 | -0.06 | -0.01 | 0.21 | -0.01 | -0.16 | 0.76 |
| TJP1 | S297; S300 | -0.19 | 0.08 | 0.08 | 0.05 | -0.09 | 0.05 | 0.904 |
| TJP1 | S277 | 0.05 | 0.09 | -0.15 | 0.12 | -0.19 | 0.04 | 0.944 |
| TJP1 | S617 | -0.18 | 0.27 | -0.06 | 0.13 | 0.00 | -0.23 | 0.82 |
| TJP1 | S/Y | -0.09 | 0.14 | 0.00 | -0.17 | 0.21 | -0.14 | 0.74 |
| TJP1 | S/Y | 0.00 | 0.06 | 0.00 | -0.26 | 0.16 | 0.00 | 0.69 |
| TJP1 | S617 | 0.08 | 0.12 | -0.10 | 0.11 | 0.07 | -0.34 | 0.615 |
| TJP1 | S617 | 0.07 | 0.10 | -0.03 | -0.08 | 0.12 | -0.20 | 0.369 |
| TJP1 | S927 | 0.15 | -0.05 | 0.08 | 0.01 | -0.19 | -0.03 | 0.204 |
| TJP1 | S329 | -0.14 | 0.08 | 0.25 | -0.13 | -0.22 | 0.10 | 0.389 |
| TJP1 | S125 | 0.04 | -0.05 | 0.27 | -0.30 | 0.01 | -0.02 | 0.243 |
| TJP1 | S968 | 0.00 | 0.07 | 0.31 | -0.19 | -0.11 | -0.15 | 0.047 |
| TJP1 | S125 | -0.03 | 0.20 | 0.22 | -0.20 | 0.06 | -0.34 | 0.111 |
| TJP1 | S329 | -0.16 | 0.27 | 0.29 | -0.13 | -0.22 | -0.15 | 0.113 |
| TJP1 | S179 | 0.08 | 0.16 | 0.15 | 0.13 | -0.06 | -0.59 | 0.237 |
| TJP1 | T354 | 0.18 | 0.23 | 0.06 | -0.31 | 0.23 | -0.57 | 0.204 |
| TJP1 | S912 | 0.17 | 0.43 | 0.32 | -0.41 | -0.15 | -0.68 | 0.013 |
| TJP2 | S244 | -0.17 | -0.38 | -0.03 | 0.22 | -0.23 | 0.43 | 0.207 |
| TJP2 | S170 | -0.37 | -0.08 | -0.06 | 0.21 | 0.05 | 0.18 | 0.046 |
| TJP2 | S1159 | -0.34 | -0.12 | -0.01 | 0.05 | 0.09 | 0.26 | 0.068 |
| TJP2 | S702 | -0.03 | -0.19 | -0.17 | 0.19 | 0.17 | -0.02 | 0.045 |
| TJP2 | S978; S986 | 0.12 | -0.38 | -0.18 | 0.49 | 0.08 | -0.32 | 0.452 |
| TJP2 | S130 | -0.13 | -0.04 | 0.00 | 0.20 | 0.00 | -0.05 | 0.275 |
| TJP2 | S266 | -0.13 | 0.06 | -0.05 | 0.17 | -0.11 | 0.03 | 0.52 |

| | | | | | | | | |
|---|---|---|---|---|---|---|---|---|
| TJP2 | S/T | -0.16 | 0.15 | -0.04 | -0.03 | 0.11 | -0.05 | 0.793 |
| TJP2 | S/T | -0.08 | 0.08 | 0.03 | 0.11 | 0.00 | -0.15 | 0.809 |
| TJP2 | S986 | 0.03 | 0.14 | -0.01 | 0.02 | -0.11 | -0.09 | 0.137 |
| TJP2 | S174 | 0.08 | 0.12 | 0.13 | -0.06 | -0.06 | -0.25 | 0.023 |
| TJP2 | S986 | 0.03 | 0.21 | 0.11 | -0.10 | 0.01 | -0.32 | 0.081 |
| TJP2 | S431 | 0.14 | 0.17 | 0.24 | -0.03 | -0.13 | -0.52 | 0.055 |
| TJP3 | S368; S371 | -0.44 | -0.37 | -0.18 | 0.33 | 0.04 | 0.40 | 0.013 |
| TJP3 | S319 | -0.27 | -0.07 | -0.14 | 0.19 | -0.12 | 0.32 | 0.116 |
| TJP3 | S319 | -0.15 | -0.05 | -0.17 | 0.24 | -0.13 | 0.20 | 0.137 |
| TJP3 | S164 | -0.11 | -0.10 | -0.14 | 0.13 | -0.09 | 0.26 | 0.096 |
| TJP3 | S203 | -0.07 | -0.05 | -0.11 | 0.14 | -0.09 | 0.15 | 0.15 |
| TJP3 | S | 0.11 | 0.32 | -0.74 | -0.22 | 0.17 | 0.14 | 0.721 |
| TJP3 | S371 | -0.16 | 0.04 | -0.06 | 0.18 | 0.02 | -0.04 | 0.267 |
| TJP3 | S164 | 0.11 | -0.08 | -0.24 | 0.37 | -0.41 | 0.12 | 0.728 |
| TJP3 | S164 | 0.01 | -0.03 | -0.11 | 0.35 | -0.09 | -0.19 | 0.723 |
| TJP3 | S591 | -0.07 | 0.08 | -0.09 | -0.18 | 0.10 | 0.14 | 0.698 |
| TJP3 | S905; S906 | 0.03 | 0.06 | -0.10 | 0.06 | -0.23 | 0.16 | 0.988 |
| TJP3 | S489 | 0.05 | 0.32 | -0.32 | -0.61 | 0.35 | -0.01 | 0.765 |
| TJP3 | S203 | 0.05 | 0.10 | 0.06 | -0.17 | -0.01 | -0.04 | 0.049 |
| TLE1 | S/T | 0.31 | 0.35 | 0.15 | -0.05 | -0.30 | -0.73 | 0.038 |
| TLE3 | S286 | -0.17 | -0.28 | -0.36 | 0.31 | 0.04 | 0.31 | 0.01 |
| TLE3 | S240; S245 | 0.00 | -0.09 | -0.03 | -0.01 | 0.05 | 0.08 | 0.082 |
| TLE4 | S208 | 0.02 | -0.02 | 0.13 | -0.50 | 0.28 | -0.03 | 0.612 |
| TLK1 | S134 | 0.06 | 0.12 | -0.10 | 0.13 | -0.21 | -0.03 | 0.652 |
| TLK2 | S771 | -0.26 | -0.48 | 0.42 | -0.61 | 0.18 | 0.41 | 0.813 |
| TMC4 | S109 | 0.13 | 0.32 | 0.25 | -0.31 | -0.20 | -0.34 | 0.002 |
| TMC5 | S968 | -0.13 | -0.09 | -0.06 | 0.18 | -0.19 | 0.24 | 0.274 |
| TMC6 | S63 | 0.10 | 0.21 | 0.08 | -0.34 | 0.04 | -0.17 | 0.074 |
| TMCO1 | S239 | -0.15 | -0.04 | -0.12 | -0.09 | 0.30 | 0.04 | 0.192 |
| TMEM104 | S/T | -0.09 | 0.16 | 0.05 | 0.02 | -0.17 | 0.02 | 0.434 |
| TMEM104 | S90 | -0.08 | 0.21 | 0.06 | -0.03 | -0.08 | -0.11 | 0.186 |
| TMEM11 | S13 | -0.21 | -0.18 | 0.10 | 0.24 | -0.12 | 0.10 | 0.313 |
| TMEM115 | S320 | 0.13 | 0.22 | 0.39 | -0.22 | -0.24 | -0.46 | 0.007 |
| TMEM115 | T329 | 0.32 | 0.50 | 0.35 | -1.28 | -0.30 | -0.26 | 0.042 |
| TMEM131 | S1261 | -0.37 | -0.02 | 0.17 | -0.05 | 0.05 | 0.16 | 0.501 |
| TMEM132A | S913 | -0.09 | -0.10 | 0.22 | -0.33 | 0.09 | 0.14 | 0.828 |
| TMEM132A | S529 | -0.10 | 0.16 | 0.24 | -0.11 | -0.15 | -0.10 | 0.104 |
| TMEM134 | S70 | -0.37 | 0.14 | 0.13 | 0.18 | -0.50 | 0.26 | 0.965 |
| TMEM184B | S388 | -0.02 | -0.18 | 0.05 | 0.11 | 0.02 | 0.00 | 0.291 |
| TMEM184B | S402 | -0.11 | 0.24 | 0.22 | 0.05 | -0.28 | -0.21 | 0.163 |
| TMEM184C | S417 | -0.20 | 0.20 | 0.17 | 0.13 | -0.40 | 0.00 | 0.513 |
| TMEM185A | S333 | 0.23 | 0.03 | -0.11 | -0.23 | -0.24 | 0.23 | 0.517 |
| TMEM192 | S230 | -0.09 | -0.09 | 0.16 | 0.08 | 0.00 | -0.08 | 0.923 |
| TMEM201 | S611; S612; S615 | -0.05 | 0.01 | -0.02 | 0.16 | -0.22 | 0.10 | 0.768 |
| TMEM201 | S441 | 0.08 | 0.03 | -0.03 | -0.06 | 0.02 | -0.05 | 0.204 |
| TMEM201 | S612; S615 | 0.08 | 0.06 | 0.04 | -0.11 | -0.13 | 0.04 | 0.079 |
| TMEM201 | S614 | 0.15 | 0.27 | 0.05 | -0.14 | 0.01 | -0.43 | 0.073 |
| TMEM201 | S441 | 0.10 | 0.17 | 0.21 | -0.27 | 0.06 | -0.37 | 0.057 |

| | | | | | | | | |
|---|---|---|---|---|---|---|---|---|
| **TMEM209** | **S222** | 0.22 | 0.02 | 0.10 | -0.02 | -0.17 | -0.20 | 0.036 |
| **TMEM209** | **S222** | 0.49 | 0.17 | 0.24 | -0.54 | -0.04 | -0.66 | 0.029 |
| **TMEM209** | **S222** | 0.38 | 0.31 | 0.27 | -0.33 | -0.22 | -0.75 | 0.01 |
| **TMEM209** | **S222** | 0.34 | 0.32 | 0.36 | -0.42 | -0.32 | -0.62 | 1E-03 |
| **TMEM214** | **T97** | 0.33 | 0.24 | -0.08 | -0.25 | -0.16 | -0.17 | 0.047 |
| **TMEM214** | **T97** | 0.30 | 0.17 | 0.19 | -0.19 | -0.15 | -0.46 | 0.011 |
| **TMEM230** | **S24** | -0.07 | -0.05 | -0.21 | 0.16 | 0.13 | 0.01 | 0.033 |
| **TMEM237** | **S62** | -0.03 | 0.00 | -0.04 | 0.04 | -0.09 | 0.11 | 0.512 |
| **TMEM237** | **S49** | -0.16 | 0.23 | 0.48 | 0.20 | -0.21 | -0.93 | 0.261 |
| **TMEM245** | **S327; S332** | -0.04 | -0.05 | 0.01 | -0.13 | 0.17 | 0.03 | 0.591 |
| **TMEM245** | **T36** | 0.02 | 0.03 | 0.15 | -0.08 | -0.14 | 0.00 | 0.067 |
| **TMEM245** | **S332** | 0.05 | 0.06 | 0.16 | -0.07 | -0.10 | -0.12 | 0.008 |
| **TMEM40** | **S137** | 0.01 | -0.03 | 0.04 | -0.09 | -0.02 | 0.09 | 0.808 |
| **TMEM41B** | **T18** | 0.07 | 0.37 | -0.43 | 0.00 | -0.48 | 0.26 | 0.811 |
| **TMEM45B** | **S270; S272** | -0.17 | -0.13 | -0.05 | 0.07 | -0.03 | 0.26 | 0.079 |
| **TMEM45B** | **S270; S272** | -0.23 | 0.00 | 0.04 | -0.18 | 0.19 | 0.13 | 0.483 |
| **TMEM45B** | **S272** | -0.04 | -0.02 | 0.08 | -0.09 | -0.09 | 0.16 | 0.88 |
| **TMEM45B** | **S270** | 0.04 | 0.06 | 0.12 | -0.08 | 0.04 | -0.20 | 0.108 |
| **TMEM51** | **T234** | -0.43 | 0.11 | -0.28 | 0.53 | -0.16 | 0.03 | 0.267 |
| **TMEM51** | **S155; S157** | -0.07 | 0.06 | -0.08 | -0.10 | -0.24 | 0.35 | 0.873 |
| **TMEM54** | **T221** | 0.21 | -0.14 | 0.12 | -0.16 | -0.05 | -0.02 | 0.273 |
| **TMEM63A** | **S739** | 0.07 | 0.16 | 0.17 | -0.14 | -0.09 | -0.21 | 0.004 |
| **TMEM63A** | **S739** | 0.17 | 0.06 | 0.36 | -0.48 | 0.04 | -0.32 | 0.065 |
| **TMEM63B** | **S115** | -0.03 | -0.03 | 0.13 | -0.19 | -0.01 | 0.11 | 0.639 |
| **TMEM63C** | **S96** | 0.06 | 0.05 | 0.34 | -0.18 | -0.01 | -0.35 | 0.078 |
| **TMEM63C** | **S790** | 0.20 | 0.00 | 0.31 | 0.05 | -0.54 | -0.18 | 0.114 |
| **TMEM87A** | **S540** | -0.18 | -0.09 | -0.10 | -0.09 | 0.30 | 0.10 | 0.122 |
| **TMEM87A** | **S540** | -0.04 | 0.16 | 0.11 | 0.01 | -0.17 | -0.09 | 0.114 |
| **TMEM94** | **S** | -0.37 | -0.20 | -0.04 | 0.07 | 0.22 | 0.22 | 0.025 |
| **TMEM94** | **S445** | -0.01 | 0.17 | -0.02 | -0.16 | 0.02 | -0.01 | 0.301 |
| **TMEM94** | **S798** | 0.05 | 0.04 | 0.15 | -0.39 | 0.24 | -0.17 | 0.371 |
| **TMF1** | **S344** | 0.05 | 0.06 | 0.05 | -0.02 | -0.09 | -0.06 | 0.004 |
| **TMF1** | **T401** | 0.00 | 0.50 | -0.02 | -0.37 | -0.07 | -0.20 | 0.119 |
| **TMF1** | **T401** | 0.03 | 0.21 | 0.53 | -0.51 | -0.05 | -0.49 | 0.045 |
| **TMOD3** | **S25** | -0.08 | -0.04 | -0.05 | 0.05 | 0.01 | 0.10 | 0.016 |
| **TMOD3** | **S/T** | -0.02 | 0.00 | 0.06 | -0.09 | -0.01 | 0.05 | 0.52 |
| **TMOD3** | **S25** | -0.05 | -0.01 | 0.21 | 0.21 | -0.10 | -0.32 | 0.526 |
| **TMPO** | **S351** | -0.43 | -0.34 | -0.59 | 0.49 | -0.36 | 0.70 | 0.093 |
| **TMPO** | **S222** | -0.05 | -0.19 | -0.51 | 0.45 | -0.17 | 0.27 | 0.131 |
| **TMPO** | **T160; S/T** | -0.16 | -0.19 | -0.12 | 0.12 | -0.07 | 0.35 | 0.079 |
| **TMPO** | **S66; S67; T74** | -0.08 | -0.04 | -0.05 | 0.05 | -0.11 | 0.21 | 0.324 |
| **TMPO** | **S66** | -0.04 | -0.07 | -0.07 | -0.01 | -0.14 | 0.28 | 0.447 |
| **TMPO** | **S/T** | -0.09 | -0.09 | 0.05 | -0.06 | -0.02 | 0.18 | 0.447 |
| **TMPO** | **T74** | -0.04 | 0.01 | -0.04 | 0.00 | -0.17 | 0.20 | 0.775 |
| **TMPO** | **S424** | -0.07 | 0.09 | -0.06 | 0.07 | 0.14 | -0.20 | 0.871 |
| **TMPO** | **S66; S67** | 0.02 | 0.00 | -0.03 | 0.00 | -0.10 | 0.11 | 0.948 |
| **TMPO** | **S306** | -0.13 | 0.03 | 0.13 | -0.03 | 0.16 | -0.19 | 0.826 |
| **TMPO** | **S156; S159; T160** | -0.13 | -0.04 | 0.20 | -0.08 | 0.11 | -0.09 | 0.808 |

| | | | | | | | | |
|---|---|---|---|---|---|---|---|---|
| TMPO | S/T/Y | 0.11 | -0.26 | 0.23 | 0.01 | -0.31 | 0.14 | 0.711 |
| TMPO | S385 | 0.04 | -0.04 | 0.20 | -0.03 | 0.01 | -0.21 | 0.217 |
| TMPO | S158; S159 | 0.12 | 0.07 | 0.02 | -0.01 | -0.18 | -0.03 | 0.079 |
| TMPO | T160 | 0.06 | 0.03 | 0.15 | -0.15 | -0.06 | -0.06 | 0.021 |
| TMPO | S184 | -0.07 | 0.10 | 0.25 | -0.11 | 0.09 | -0.33 | 0.242 |
| TMPO | S184; S/Y | 0.11 | 0.11 | 0.07 | -0.12 | 0.12 | -0.36 | 0.187 |
| TMPO | T74 | 0.06 | 0.02 | 0.24 | -0.24 | -0.25 | 0.10 | 0.155 |
| TMPO | S184 | 0.13 | 0.09 | 0.14 | -0.36 | 0.14 | -0.22 | 0.151 |
| TMPO | S177; S180; S184 | 0.21 | 0.04 | 0.19 | -0.09 | 0.02 | -0.46 | 0.104 |
| TMPO | S177; S184 | 0.16 | 0.09 | 0.21 | -0.27 | -0.03 | -0.23 | 0.014 |
| TMPO | S184; S/Y | -0.03 | 0.26 | 0.32 | -0.33 | -0.09 | -0.25 | 0.037 |
| TMPO | S184 | 0.07 | 0.15 | 0.29 | -0.47 | 0.22 | -0.47 | 0.16 |
| TMPO | S180 | 0.19 | 0.23 | 0.14 | -0.26 | 0.12 | -0.60 | 0.106 |
| TMPO | S184; S/Y | 0.23 | 0.18 | 0.26 | -0.26 | 0.00 | -0.58 | 0.04 |
| TMPO | S190 | 0.32 | 0.27 | 0.10 | -0.18 | -0.12 | -0.59 | 0.031 |
| TMPO | S177; S180; S184 | 0.26 | 0.13 | 0.26 | 0.13 | -0.05 | -1.12 | 0.225 |
| TMPO | S184 | 0.27 | 0.24 | 0.23 | -0.28 | 0.05 | -0.78 | 0.071 |
| TMPO | S190 | 0.21 | 0.36 | 0.40 | -0.37 | -0.07 | -0.99 | 0.045 |
| TMPO | S180; S184 | 0.39 | 0.32 | 0.37 | -0.35 | -0.12 | -1.18 | 0.047 |
| TMPO | S184 | 0.38 | 0.34 | 0.42 | -0.53 | -0.12 | -1.02 | 0.023 |
| TMPRSS13 | T29 | 0.27 | -0.04 | -0.16 | -0.51 | 0.12 | 0.18 | 0.739 |
| TMPRSS13 | T64 | -0.28 | 0.13 | 0.48 | -0.54 | -0.07 | 0.08 | 0.372 |
| TMPRSS13 | S115 | 0.37 | 0.12 | 0.15 | -0.24 | -0.10 | -0.45 | 0.021 |
| TMTC1 | S290 | -0.09 | -0.07 | 0.08 | 0.06 | 0.01 | 0.00 | 0.388 |
| TMUB1 | T71 | -0.11 | -0.02 | 0.05 | -0.03 | 0.05 | 0.05 | 0.436 |
| TMX1 | S270 | 0.26 | 0.02 | -0.38 | -0.17 | -0.02 | 0.19 | 0.887 |
| TMX1 | S247 | 0.20 | 0.13 | -0.10 | -0.10 | -0.13 | -0.04 | 0.149 |
| TMX1 | S247 | 0.03 | 0.29 | -0.04 | -0.25 | -0.24 | 0.13 | 0.25 |
| TMX2 | S288 | 0.15 | 0.08 | 0.11 | -0.07 | -0.12 | -0.17 | 0.003 |
| TMX2 | S288 | 0.20 | 0.21 | 0.10 | -0.08 | -0.19 | -0.33 | 0.01 |
| TMX2 | S288 | 0.14 | 0.18 | 0.23 | -0.21 | -0.16 | -0.27 | 6E-04 |
| TNC | S72 | 0.01 | -0.16 | -0.14 | 0.01 | -0.14 | 0.34 | 0.336 |
| TNC | S/T | -0.10 | -0.05 | -0.11 | 0.20 | -0.26 | 0.25 | 0.42 |
| TNC | S/T | -0.13 | -0.04 | -0.05 | 0.28 | -0.39 | 0.23 | 0.639 |
| TNC | S72 | 0.13 | -0.12 | -0.09 | 0.03 | -0.13 | 0.16 | 0.72 |
| TNIK | S/Y/T | 0.03 | -0.16 | 0.03 | -0.26 | 0.06 | 0.25 | 0.753 |
| TNIK | S720 | 0.01 | -0.05 | 0.02 | -0.47 | -0.25 | 0.55 | 0.883 |
| TNIK | S548 | 0.07 | -0.13 | 0.33 | -0.31 | -0.12 | 0.08 | 0.309 |
| TNIK | S/T | -0.02 | -0.04 | 0.37 | 0.05 | -0.17 | -0.26 | 0.223 |
| TNIK | S | 0.09 | 0.19 | 0.08 | 0.08 | -0.26 | -0.24 | 0.085 |
| TNIK | S769 | 0.25 | 0.15 | 0.18 | -0.25 | 0.00 | -0.45 | 0.033 |
| TNIP1 | S403 | -0.20 | -0.44 | 0.25 | -0.14 | 0.30 | 0.08 | 0.428 |
| TNKS1BP1 | S221 | -0.18 | -0.33 | -0.02 | 0.40 | -0.46 | 0.36 | 0.404 |
| TNKS1BP1 | S498 | -0.04 | -0.12 | -0.24 | -0.25 | 0.25 | 0.30 | 0.279 |
| TNKS1BP1 | S1103 | -0.20 | 0.16 | -0.25 | 0.23 | 0.16 | -0.18 | 0.408 |
| TNKS1BP1 | S936 | 0.05 | 0.03 | -0.35 | -0.11 | 0.10 | 0.21 | 0.379 |
| TNKS1BP1 | S712 | 0.05 | -0.09 | -0.18 | 0.14 | -0.02 | 0.07 | 0.16 |
| TNKS1BP1 | S893 | -0.15 | 0.12 | -0.08 | -0.03 | -0.03 | 0.14 | 0.561 |

| | | | | | | | | |
|---|---|---|---|---|---|---|---|---|
| **TNKS1BP1** | **S672** | 0.05 | -0.11 | 0.00 | 0.06 | -0.08 | 0.08 | 0.587 |
| **TNKS1BP1** | **S429** | 0.12 | 0.08 | -0.26 | 0.07 | 0.00 | -0.04 | 0.804 |
| **TNKS1BP1** | **S1620** | -0.03 | -0.01 | -0.01 | 0.02 | 0.00 | 0.02 | 0.024 |
| **TNKS1BP1** | **S1473** | -0.05 | 0.20 | -0.23 | 0.29 | -0.05 | -0.24 | 0.899 |
| **TNKS1BP1** | **S601** | 0.04 | -0.01 | -0.04 | -0.12 | 0.03 | 0.09 | 0.967 |
| **TNKS1BP1** | **S435** | -0.08 | 0.12 | -0.02 | -0.11 | 0.04 | 0.05 | 0.898 |
| **TNKS1BP1** | **S893** | 0.07 | -0.07 | 0.11 | 0.06 | 0.00 | -0.18 | 0.471 |
| **TNKS1BP1** | **S893** | -0.03 | 0.05 | 0.09 | 0.03 | -0.22 | 0.06 | 0.482 |
| **TNKS1BP1** | **S494; S498** | -0.04 | 0.04 | 0.13 | -0.09 | -0.04 | -0.01 | 0.186 |
| **TNKS1BP1** | **S1297** | 0.15 | 0.05 | -0.20 | -0.66 | -0.39 | 0.67 | 0.775 |
| **TNKS1BP1** | **S872** | 0.10 | 0.05 | 0.07 | 0.08 | -0.02 | -0.33 | 0.265 |
| **TNKS1BP1** | **S1385** | 0.15 | 0.08 | -0.01 | -0.01 | 0.04 | -0.29 | 0.218 |
| **TNKS1BP1** | **S1545** | 0.08 | 0.16 | 0.01 | -0.07 | 0.00 | -0.19 | 0.078 |
| **TNKS1BP1** | **S228** | 0.20 | 0.12 | -0.09 | 0.02 | -0.06 | -0.24 | 0.207 |
| **TNKS1BP1** | **S1385** | 0.03 | 0.25 | 0.01 | -0.18 | -0.13 | -0.03 | 0.078 |
| **TNKS1BP1** | **S872** | 0.28 | 0.01 | 0.09 | 0.11 | -0.15 | -0.43 | 0.182 |
| **TNKS1BP1** | **S1439** | -0.02 | 0.23 | 0.27 | 0.02 | -0.09 | -0.55 | 0.13 |
| **TNKS1BP1** | **S1138** | -0.01 | 0.10 | 0.43 | -0.23 | -0.01 | -0.43 | 0.094 |
| **TNRC18** | **S1127** | -0.21 | 0.28 | -0.63 | 0.43 | 0.12 | -0.26 | 0.443 |
| **TNRC18** | **S1989** | -0.06 | -0.12 | -0.09 | 0.09 | -0.15 | 0.28 | 0.264 |
| **TNRC18** | **S991** | 0.01 | -0.08 | -0.17 | 0.11 | -0.10 | 0.20 | 0.216 |
| **TNRC18** | **S1857** | -0.01 | -0.01 | -0.17 | 0.04 | -0.11 | 0.22 | 0.353 |
| **TNRC18** | **T1250** | -0.05 | 0.08 | -0.17 | -0.04 | 0.17 | -0.02 | 0.459 |
| **TNRC18** | **T/S** | 0.13 | 0.10 | -0.45 | -0.53 | -0.28 | 0.66 | 0.968 |
| **TNRC18** | **S2375** | 0.14 | 0.01 | -0.20 | -0.04 | -0.21 | 0.24 | 0.923 |
| **TNRC18** | **S2543** | 0.40 | -0.87 | 0.29 | 0.02 | 0.00 | -0.16 | 0.979 |
| **TNRC18** | **S2543** | 0.11 | 0.03 | -0.11 | 0.08 | -0.15 | 0.02 | 0.766 |
| **TNRC18** | **T2146** | 0.03 | 0.05 | 0.05 | -0.09 | -0.01 | -0.03 | 0.026 |
| **TNRC18** | **S1957** | 0.10 | 0.03 | 0.06 | -0.09 | -0.22 | 0.09 | 0.201 |
| **TNRC18** | **S999** | 0.13 | 0.11 | -0.01 | -0.17 | -0.28 | 0.15 | 0.262 |
| **TNRC18** | **S1957** | 0.13 | 0.16 | -0.03 | 0.07 | -0.19 | -0.17 | 0.145 |
| **TNRC18** | **S2471** | 0.16 | 0.12 | 0.03 | -0.03 | -0.13 | -0.18 | 0.022 |
| **TNRC18** | **S1540** | 0.05 | 0.18 | 0.31 | -0.19 | -0.20 | -0.25 | 0.007 |
| **TNRC18** | **S2716** | 0.29 | 0.02 | 0.30 | -0.19 | -0.42 | -0.15 | 0.021 |
| **TNRC18** | **S1127; S1136** | 0.22 | 0.50 | -0.01 | 0.05 | -0.12 | -1.05 | 0.178 |
| **TNRC6A** | **S739** | -0.28 | -0.02 | 0.12 | 0.19 | -0.05 | 0.00 | 0.479 |
| **TNRC6A** | **S991** | -0.03 | 0.03 | -0.13 | 0.01 | 0.08 | 0.03 | 0.165 |
| **TNRC6B** | **S879** | 0.02 | -0.18 | 0.12 | 0.03 | -0.16 | 0.14 | 0.892 |
| **TNRC6C** | **S714** | -0.23 | -0.10 | -0.15 | 0.10 | -0.14 | 0.42 | 0.159 |
| **TNRC6C** | **S/T** | 0.12 | 0.26 | -0.20 | 0.01 | 0.08 | -0.34 | 0.491 |
| **TNS1** | **S1373** | -0.24 | 0.27 | -0.10 | -0.21 | 0.22 | -0.02 | 0.924 |
| **TNS2** | **S835** | 0.18 | -0.24 | 0.20 | 0.12 | -0.16 | -0.17 | 0.529 |
| **TNS3** | **S660** | -0.58 | -0.46 | -0.17 | 0.30 | 0.22 | 0.40 | 0.006 |
| **TNS3** | **S660** | -0.55 | -0.52 | -0.12 | 0.13 | 0.19 | 0.55 | 0.023 |
| **TNS3** | **S332** | -0.09 | -0.18 | -0.23 | 0.37 | 0.26 | -0.27 | 0.224 |
| **TNS3** | **S332** | -0.11 | -0.18 | -0.08 | 0.13 | 0.33 | -0.15 | 0.189 |
| **TNS3** | **S/T** | -0.35 | 0.18 | -0.03 | -0.15 | -0.01 | 0.27 | 0.637 |
| **TOE1** | **S428** | -0.11 | -0.42 | 0.11 | 0.08 | -0.02 | 0.27 | 0.227 |

| | | | | | | | | |
|---|---|---|---|---|---|---|---|---|
| **TOE1** | **S428** | -0.01 | -0.13 | -0.25 | 0.28 | -0.20 | 0.23 | 0.234 |
| **TOM1** | **S355** | -0.27 | -0.07 | -0.20 | 0.10 | 0.04 | 0.33 | 0.034 |
| **TOM1L1** | **S323** | 0.02 | 0.07 | -0.07 | -0.05 | 0.07 | -0.05 | 0.823 |
| **TOM1L1** | **S323** | -0.05 | 0.11 | 0.05 | -0.21 | 0.15 | -0.08 | 0.51 |
| **TOM1L1** | **S/T** | 0.34 | 0.20 | 0.38 | -0.38 | -0.21 | -0.59 | 0.005 |
| **TOMM20** | **S135; S138** | -0.13 | -0.69 | -0.16 | 0.01 | 0.44 | 0.28 | 0.062 |
| **TOMM20** | **S135** | -0.16 | -0.07 | -0.15 | 0.12 | 0.19 | 0.04 | 0.01 |
| **TOMM22** | **S45** | -0.09 | -0.23 | -0.08 | 0.24 | -0.05 | 0.15 | 0.062 |
| **TOMM34** | **S93** | 0.15 | -0.07 | -0.44 | 0.18 | 0.07 | 0.03 | 0.291 |
| **TOMM34** | **S186** | -0.16 | -0.05 | 0.01 | 0.00 | -0.08 | 0.25 | 0.321 |
| **TOMM34** | **S186** | 0.28 | 0.10 | 0.15 | -0.38 | 0.24 | -0.59 | 0.175 |
| **TOMM70** | **S91** | -0.16 | -0.16 | 0.09 | 0.04 | 0.04 | 0.13 | 0.172 |
| **TOMM70** | **S91** | -0.02 | 0.09 | 0.22 | -0.07 | -0.10 | -0.15 | 0.046 |
| **TOMM70** | **S91** | 0.11 | 0.15 | 0.19 | -0.11 | -0.12 | -0.27 | 0.005 |
| **TOMM70** | **S91** | 0.12 | 0.21 | 0.19 | -0.12 | -0.17 | -0.31 | 0.004 |
| **TOP1** | **S97** | -0.38 | -0.41 | -0.29 | 0.50 | 0.15 | 0.19 | 0.005 |
| **TOP1** | **S112** | -0.07 | -0.12 | -0.29 | 0.09 | -0.01 | 0.33 | 0.069 |
| **TOP1** | **S112** | -0.08 | -0.16 | -0.16 | 0.07 | 0.11 | 0.19 | 0.005 |
| **TOP1** | **S112** | 0.02 | -0.05 | -0.25 | 0.05 | 0.05 | 0.15 | 0.111 |
| **TOP1** | **S/Y** | 0.02 | 0.00 | -0.23 | 0.13 | 0.02 | 0.04 | 0.202 |
| **TOP1** | **S73** | 0.01 | 0.00 | -0.08 | 0.04 | 0.15 | -0.13 | 0.649 |
| **TOP1** | **S112** | 0.12 | 0.10 | -0.21 | -0.02 | -0.02 | 0.02 | 0.937 |
| **TOP1** | **S394** | -0.08 | 0.10 | -0.01 | 0.03 | 0.00 | -0.05 | 0.824 |
| **TOP1** | **S112** | 0.02 | 0.06 | 0.04 | 0.02 | -0.05 | -0.10 | 0.076 |
| **TOP1** | **S112** | 0.23 | -0.13 | 0.14 | -0.20 | 0.02 | -0.12 | 0.229 |
| **TOP1** | **S/Y** | 0.28 | 0.16 | 0.09 | -0.10 | -0.04 | -0.52 | 0.068 |
| **TOP2A** | **S1449** | -0.68 | -0.49 | -0.58 | 0.24 | 0.53 | 0.46 | 6E-04 |
| **TOP2A** | **S1449** | -0.41 | -0.38 | -0.49 | 0.14 | 0.20 | 0.61 | 0.008 |
| **TOP2A** | **S1374; S1377** | -0.17 | -0.10 | -0.27 | 0.17 | 0.15 | 0.15 | 0.002 |
| **TOP2A** | **T/S** | -0.17 | -0.34 | -0.04 | -0.03 | 0.17 | 0.32 | 0.067 |
| **TOP2A** | **S1525** | -0.13 | -0.18 | -0.10 | -0.01 | 0.08 | 0.29 | 0.053 |
| **TOP2A** | **S1247** | -0.01 | -0.18 | -0.07 | 0.03 | 0.22 | -0.02 | 0.132 |
| **TOP2A** | **S1354** | -0.07 | -0.14 | 0.01 | 0.14 | -0.17 | 0.19 | 0.372 |
| **TOP2A** | **S1106** | -0.04 | -0.06 | -0.09 | -0.09 | -0.02 | 0.27 | 0.364 |
| **TOP2A** | **S1469** | 0.02 | -0.11 | -0.01 | -0.16 | -0.10 | 0.32 | 0.741 |
| **TOP2A** | **S1247** | 0.02 | -0.14 | 0.03 | -0.14 | 0.28 | -0.10 | 0.773 |
| **TOP2A** | **S1425** | 0.19 | -0.34 | 0.07 | -0.23 | 0.04 | 0.19 | 0.89 |
| **TOP2A** | **S1106** | 0.06 | 0.05 | -0.12 | -0.05 | -0.01 | 0.06 | 0.978 |
| **TOP2A** | **S1247** | 0.04 | 0.08 | -0.11 | -0.18 | 0.30 | -0.19 | 0.883 |
| **TOP2A** | **T602** | 0.08 | -0.01 | 0.00 | -0.16 | 0.10 | -0.03 | 0.51 |
| **TOP2A** | **S1377** | 0.12 | -0.01 | -0.02 | -0.14 | 0.03 | 0.01 | 0.411 |
| **TOP2A** | **S1213** | -0.06 | 0.22 | -0.08 | 0.01 | 0.24 | -0.43 | 0.706 |
| **TOP2A** | **S1393** | 0.08 | 0.09 | -0.03 | -0.17 | -0.01 | 0.04 | 0.28 |
| **TOP2A** | **S1469** | 0.07 | 0.11 | 0.07 | -0.15 | -0.16 | 0.03 | 0.053 |
| **TOP2A** | **S/T** | 0.09 | 0.14 | 0.02 | -0.35 | -0.20 | 0.22 | 0.331 |
| **TOP2A** | **S1351; S1354** | 0.01 | 0.03 | 0.23 | -0.03 | 0.00 | -0.29 | 0.161 |
| **TOP2A** | **S1469; S1471; S1474** | 0.22 | 0.15 | -0.04 | -0.18 | -0.11 | -0.09 | 0.046 |
| **TOP2A** | **S1213** | 0.04 | 0.13 | 0.22 | -0.33 | 0.20 | -0.40 | 0.197 |

| | | | | | | | | |
|---|---|---|---|---|---|---|---|---|
| **TOP2A** | **S1213** | 0.12 | 0.10 | 0.30 | -0.35 | 0.13 | -0.45 | 0.107 |
| **TOP2A** | **S1295; S1299; S1302** | 0.24 | 0.27 | 0.09 | -0.30 | -0.12 | -0.29 | 0.006 |
| **TOP2B** | **T1292** | -0.16 | -0.21 | -0.10 | -0.08 | 0.29 | 0.19 | 0.065 |
| **TOP2B** | **S1466** | -0.13 | -0.18 | -0.03 | 0.03 | 0.07 | 0.20 | 0.034 |
| **TOP2B** | **S1400** | -0.09 | -0.03 | -0.20 | 0.07 | -0.06 | 0.26 | 0.138 |
| **TOP2B** | **S1236** | -0.09 | -0.04 | -0.17 | 0.06 | 0.05 | 0.17 | 0.023 |
| **TOP2B** | **S1613** | -0.10 | -0.10 | -0.08 | 0.05 | 0.04 | 0.17 | 0.014 |
| **TOP2B** | **S1466; S1471** | -0.49 | 0.06 | 0.13 | -0.04 | 0.20 | 0.04 | 0.457 |
| **TOP2B** | **S1400** | -0.16 | 0.21 | -0.32 | -0.06 | 0.23 | 0.03 | 0.434 |
| **TOP2B** | **S1552** | -0.15 | -0.05 | -0.04 | -0.08 | 0.18 | 0.12 | 0.156 |
| **TOP2B** | **S/T** | -0.40 | -0.05 | 0.19 | 0.11 | 0.14 | -0.08 | 0.478 |
| **TOP2B** | **S675** | -0.02 | -0.17 | -0.02 | -0.15 | 0.39 | -0.10 | 0.552 |
| **TOP2B** | **S1375** | -0.02 | -0.21 | 0.05 | -0.01 | -0.12 | 0.26 | 0.495 |
| **TOP2B** | **S1613** | 0.07 | -0.24 | 0.06 | -0.02 | -0.01 | 0.11 | 0.577 |
| **TOP2B** | **S1400** | 0.14 | -0.02 | -0.18 | -0.10 | -0.04 | 0.17 | 0.812 |
| **TOP2B** | **S1375** | 0.04 | -0.07 | -0.01 | -0.11 | 0.03 | 0.11 | 0.786 |
| **TOP2B** | **S1522; S1524** | 0.04 | 0.06 | -0.10 | 0.09 | 0.01 | -0.12 | 0.946 |
| **TOP2B** | **S1524** | 0.26 | -0.02 | -0.24 | -0.18 | 0.12 | 0.00 | 0.916 |
| **TOP2B** | **S1236** | 0.05 | 0.01 | -0.03 | -0.23 | 0.13 | 0.05 | 0.822 |
| **TOP2B** | **S1336** | -0.01 | 0.05 | -0.02 | 0.08 | 0.12 | -0.25 | 0.826 |
| **TOP2B** | **S1581** | 0.09 | -0.08 | 0.08 | -0.09 | 0.06 | -0.07 | 0.402 |
| **TOP2B** | **S1400** | 0.14 | 0.11 | -0.16 | -0.18 | 0.07 | -0.01 | 0.608 |
| **TOP2B** | **S1344** | 0.08 | 0.09 | -0.03 | -0.09 | -0.25 | 0.16 | 0.447 |
| **TOP2B** | **S1550; S1552** | 0.03 | -0.02 | 0.15 | -0.10 | -0.04 | -0.03 | 0.107 |
| **TOP2B** | **S1424** | 0.20 | 0.05 | -0.10 | -0.26 | -0.10 | 0.16 | 0.492 |
| **TOP2B** | **S** | -0.15 | 0.23 | 0.08 | -0.19 | 0.06 | -0.07 | 0.408 |
| **TOP2B** | **S1552** | 0.01 | 0.10 | 0.05 | -0.34 | 0.11 | 0.02 | 0.427 |
| **TOP2B** | **S1550** | 0.21 | -0.01 | 0.01 | -0.27 | 0.04 | -0.01 | 0.271 |
| **TOP2B** | **S1550** | 0.10 | 0.00 | 0.17 | -0.19 | -0.06 | -0.06 | 0.038 |
| **TOP2B** | **S1400; S1424** | 0.07 | 0.25 | -0.01 | -0.12 | -0.14 | -0.09 | 0.048 |
| **TOP2B** | **S1550; S1552** | 0.18 | 0.09 | 0.05 | -0.20 | -0.08 | -0.06 | 0.02 |
| **TOP2B** | **S1336** | 0.16 | 0.15 | 0.02 | -0.05 | -0.16 | -0.15 | 0.017 |
| **TOP2B** | **S1424** | 0.25 | -0.19 | 0.24 | 0.14 | -0.15 | -0.41 | 0.319 |
| **TOP2B** | **S1466** | 0.14 | 0.10 | 0.14 | -0.20 | -0.02 | -0.20 | 0.012 |
| **TOP2B** | **S1400; T/S/Y** | 0.20 | 0.09 | 0.09 | -0.28 | 0.01 | -0.16 | 0.042 |
| **TOP2B** | **S1466** | 0.17 | 0.10 | 0.15 | -0.28 | -0.04 | -0.16 | 0.015 |
| **TOP2B** | **S1336; S1340; S1344** | 0.09 | 0.27 | 0.09 | -0.20 | -0.07 | -0.24 | 0.015 |
| **TOP2B** | **S1524** | 0.25 | 0.21 | 0.10 | -0.41 | -0.10 | -0.16 | 0.017 |
| **TOP2B** | **S1336; S1340; S/T** | 0.34 | 0.33 | 0.24 | -0.43 | -0.01 | -0.84 | 0.039 |
| **TOP2B** | **S1524** | 0.34 | 0.40 | 0.24 | -0.40 | -0.08 | -0.90 | 0.031 |
| **TOP2B** | **S1336; S1344** | 0.32 | 0.47 | 0.39 | -0.53 | -0.17 | -1.07 | 0.021 |
| **TOPORS** | **S743** | -0.18 | -0.25 | -0.04 | -0.15 | 0.13 | 0.38 | 0.172 |
| **TOPORS** | **T491; S/Y** | -0.10 | -0.11 | 0.00 | 0.28 | -0.19 | 0.07 | 0.41 |
| **TOPORS** | **S866** | 0.05 | -0.03 | -0.11 | 0.06 | 0.03 | 0.00 | 0.322 |
| **TOPORS** | **S718** | 0.15 | -0.14 | 0.02 | -0.47 | 0.09 | 0.25 | 0.836 |
| **TOPORS** | **S501** | -0.14 | -0.01 | 0.27 | -0.16 | -0.16 | 0.14 | 0.537 |
| **TOPORS** | **S98** | 0.10 | -0.04 | 0.43 | -0.12 | -0.03 | -0.49 | 0.133 |
| **TOPORS** | **S585** | 0.10 | 0.13 | 0.30 | -0.20 | -0.17 | -0.25 | 0.005 |

| | | | | | | | | |
|---|---|---|---|---|---|---|---|---|
| **TOPORS** | **S194** | 0.18 | 0.05 | 0.38 | -0.34 | -0.16 | -0.23 | 0.015 |
| **TOPORS** | **S98** | 0.25 | 0.14 | 0.22 | -0.18 | -0.04 | -0.55 | 0.042 |
| **TOR1AIP1** | **S156; S157; S164** | -0.10 | -0.20 | -0.14 | 0.06 | -0.10 | 0.39 | 0.146 |
| **TOR1AIP1** | **T/S/Y** | -0.07 | 0.04 | -0.16 | 0.04 | 0.05 | 0.08 | 0.118 |
| **TOR1AIP1** | **S/Y** | -0.23 | 0.04 | 0.11 | 0.03 | 0.07 | -0.04 | 0.678 |
| **TOR1AIP1** | **S156; S157** | 0.00 | -0.02 | -0.03 | 0.05 | -0.15 | 0.14 | 0.778 |
| **TOR1AIP1** | **S179** | -0.15 | 0.27 | -0.18 | -0.24 | 0.12 | 0.11 | 0.935 |
| **TOR1AIP1** | **T47** | 0.10 | 0.24 | -0.25 | 0.03 | -0.10 | -0.08 | 0.638 |
| **TOR1AIP1** | **S/T** | 0.06 | 0.19 | 0.01 | -0.12 | 0.00 | -0.17 | 0.068 |
| **TOR1AIP1** | **S230; S231; S/T** | -0.30 | 0.08 | 0.41 | 0.07 | 0.09 | -0.55 | 0.558 |
| **TOR1AIP1** | **S227; S230; S231** | -0.03 | -0.01 | 0.32 | 0.02 | 0.02 | -0.41 | 0.31 |
| **TOR1AIP1** | **S156** | 0.16 | 0.10 | 0.08 | -0.07 | -0.16 | -0.14 | 0.003 |
| **TOR1AIP1** | **S156; S157** | 0.24 | 0.23 | -0.11 | -0.33 | -0.26 | 0.13 | 0.22 |
| **TOR1AIP1** | **S215** | 0.24 | 0.10 | 0.10 | -0.17 | -0.13 | -0.19 | 0.004 |
| **TOR1AIP1** | **S157** | 0.21 | 0.13 | 0.09 | -0.07 | -0.22 | -0.20 | 0.006 |
| **TOR1AIP1** | **S/T** | 0.25 | 0.17 | -0.03 | -0.64 | -0.13 | 0.21 | 0.29 |
| **TOR1AIP1** | **S231; S/T** | 0.21 | 0.22 | 0.04 | -0.24 | -0.32 | 0.00 | 0.037 |
| **TOR1AIP1** | **S215** | 0.11 | 0.37 | 0.31 | -0.18 | -0.33 | -0.51 | 0.008 |
| **TOR1AIP1** | **S215** | 0.35 | 0.36 | 0.62 | -0.49 | -0.29 | -1.41 | 0.03 |
| **TOX4** | **T/S** | 0.05 | 0.04 | -0.04 | 0.00 | -0.06 | 0.01 | 0.467 |
| **TOX4** | **S182; T/S** | -0.13 | 0.14 | 0.12 | -0.20 | -0.09 | 0.12 | 0.474 |
| **TOX4** | **T176; S181; S182** | -0.05 | 0.08 | 0.12 | -0.11 | -0.11 | 0.06 | 0.252 |
| **TP53BP1** | **S/T** | -0.99 | -1.39 | -1.39 | 0.71 | 0.17 | 0.98 | 0.002 |
| **TP53BP1** | **T593** | -1.06 | -0.62 | -1.06 | 0.54 | 0.34 | 0.74 | 0.001 |
| **TP53BP1** | **T855** | -0.55 | -0.85 | -0.54 | 0.29 | 0.56 | 0.47 | 0.001 |
| **TP53BP1** | **S294; S301; T302** | -0.71 | -0.51 | -0.70 | 0.45 | 0.26 | 0.60 | 8E-04 |
| **TP53BP1** | **S1068** | -0.55 | -1.02 | -0.29 | 0.45 | 0.33 | 0.47 | 0.009 |
| **TP53BP1** | **S1068** | -0.50 | -0.61 | -0.65 | 0.34 | 0.17 | 0.69 | 0.003 |
| **TP53BP1** | **S1068** | -0.45 | -0.53 | -0.75 | 0.35 | 0.37 | 0.50 | 6E-04 |
| **TP53BP1** | **S222** | -0.37 | -0.48 | -0.78 | 0.50 | 0.64 | -0.07 | 0.022 |
| **TP53BP1** | **S280** | -0.51 | -0.41 | -0.64 | 0.32 | 0.38 | 0.44 | 3E-04 |
| **TP53BP1** | **S999** | -0.46 | -0.25 | -0.52 | 0.19 | 0.54 | 0.19 | 0.007 |
| **TP53BP1** | **S/T** | -0.24 | -0.25 | -0.73 | 0.05 | 0.42 | 0.42 | 0.025 |
| **TP53BP1** | **S1342** | -0.42 | -0.25 | -0.43 | 0.25 | 0.30 | 0.32 | 5E-04 |
| **TP53BP1** | **S580** | -0.25 | -0.36 | -0.42 | 0.30 | 0.28 | 0.24 | 3E-04 |
| **TP53BP1** | **S294; S301** | -0.30 | -0.31 | -0.41 | 0.24 | 0.05 | 0.49 | 0.011 |
| **TP53BP1** | **S834; S/T** | -0.13 | -0.72 | -0.29 | -0.52 | 0.71 | 0.41 | 0.232 |
| **TP53BP1** | **S294; S310** | -0.19 | -0.41 | -0.31 | 0.28 | 0.15 | 0.31 | 0.002 |
| **TP53BP1** | **S831; S834** | -0.36 | -0.36 | -0.18 | 0.28 | 0.36 | 0.08 | 0.006 |
| **TP53BP1** | **S1368** | -0.41 | -0.25 | -0.26 | 0.11 | 0.14 | 0.46 | 0.012 |
| **TP53BP1** | **S999** | -0.50 | -0.02 | -0.37 | 0.25 | 0.25 | 0.20 | 0.021 |
| **TP53BP1** | **S1426** | -0.15 | -0.27 | -0.43 | 0.10 | 0.36 | 0.24 | 0.009 |
| **TP53BP1** | **S580** | -0.13 | -0.33 | -0.38 | 0.18 | 0.24 | 0.26 | 0.003 |
| **TP53BP1** | **S809** | -0.21 | -0.30 | -0.32 | 0.23 | 0.13 | 0.33 | 0.002 |
| **TP53BP1** | **S580** | -0.03 | -0.56 | -0.26 | 0.02 | 0.35 | 0.28 | 0.053 |
| **TP53BP1** | **S398** | -0.18 | -0.30 | -0.38 | 0.20 | 0.56 | -0.13 | 0.072 |
| **TP53BP1** | **S1101** | -0.14 | -0.26 | -0.42 | 0.16 | 0.11 | 0.39 | 0.014 |
| **TP53BP1** | **T/S** | -0.36 | -0.37 | -0.10 | -0.01 | 0.49 | 0.15 | 0.048 |

| | | | | | | | | |
|---|---|---|---|---|---|---|---|---|
| **TP53BP1** | **S78** | -0.20 | -0.41 | -0.18 | 0.17 | 0.09 | 0.38 | 0.014 |
| **TP53BP1** | **S398** | -0.20 | -0.28 | -0.25 | 0.13 | 0.40 | 0.07 | 0.012 |
| **TP53BP1** | **T591** | -0.17 | -0.19 | -0.36 | -0.03 | 0.25 | 0.36 | 0.032 |
| **TP53BP1** | **S1678** | -0.03 | -0.50 | -0.16 | -0.09 | 0.14 | 0.47 | 0.135 |
| **TP53BP1** | **S** | -0.02 | -0.38 | -0.20 | -0.05 | 0.39 | 0.13 | 0.095 |
| **TP53BP1** | **S580** | -0.28 | -0.22 | -0.14 | 0.22 | 0.50 | -0.28 | 0.198 |
| **TP53BP1** | **T1609** | -0.17 | -0.05 | -0.36 | 0.10 | 0.21 | 0.18 | 0.023 |
| **TP53BP1** | **S1705** | -0.37 | -0.17 | 0.02 | -0.04 | 0.32 | 0.14 | 0.111 |
| **TP53BP1** | **S/T** | -0.15 | -0.21 | -0.12 | 0.03 | 0.29 | 0.09 | 0.022 |
| **TP53BP1** | **S580** | -0.21 | -0.23 | -0.09 | -0.35 | 0.15 | 0.54 | 0.324 |
| **TP53BP1** | **T/S** | -0.14 | 0.12 | -0.47 | 0.42 | -0.16 | 0.09 | 0.309 |
| **TP53BP1** | **T1285; S1288** | -0.13 | -0.11 | -0.17 | 0.23 | 0.02 | 0.12 | 0.014 |
| **TP53BP1** | **S1295; S1296** | -0.13 | -0.11 | -0.17 | 0.23 | 0.02 | 0.12 | 0.014 |
| **TP53BP1** | **S525; Y/S/T** | 0.03 | -0.27 | -0.15 | -0.15 | 0.04 | 0.40 | 0.287 |
| **TP53BP1** | **S1114** | -0.06 | -0.11 | -0.18 | -0.01 | 0.23 | 0.10 | 0.049 |
| **TP53BP1** | **S1462** | -0.21 | -0.08 | -0.06 | 0.14 | -0.03 | 0.19 | 0.056 |
| **TP53BP1** | **S1678; T/S** | -0.10 | -0.06 | -0.18 | 0.09 | 0.09 | 0.13 | 0.005 |
| **TP53BP1** | **S580** | 0.16 | -0.31 | -0.19 | 0.04 | 0.27 | -0.05 | 0.295 |
| **TP53BP1** | **T633; S/T** | -0.31 | -0.04 | 0.03 | 0.12 | -0.18 | 0.30 | 0.352 |
| **TP53BP1** | **S639** | 0.07 | -0.19 | -0.20 | 0.16 | 0.37 | -0.33 | 0.485 |
| **TP53BP1** | **S831** | -0.05 | -0.17 | -0.05 | 0.03 | 0.16 | 0.06 | 0.034 |
| **TP53BP1** | **S1114** | -0.06 | -0.01 | -0.19 | 0.01 | 0.20 | 0.03 | 0.11 |
| **TP53BP1** | **S1028** | -0.08 | -0.10 | -0.09 | -0.06 | -0.03 | 0.31 | 0.242 |
| **TP53BP1** | **S105** | -0.19 | -0.08 | 0.00 | -0.15 | 0.10 | 0.27 | 0.296 |
| **TP53BP1** | **S727** | -0.42 | 0.33 | -0.29 | -0.52 | 0.05 | 0.53 | 0.719 |
| **TP53BP1** | **S630** | 0.01 | -0.30 | 0.04 | -0.11 | -0.09 | 0.37 | 0.489 |
| **TP53BP1** | **S552** | 0.00 | -0.07 | -0.15 | 0.01 | 0.10 | 0.10 | 0.054 |
| **TP53BP1** | **S1114** | -0.04 | -0.06 | -0.11 | 0.05 | 0.21 | -0.08 | 0.204 |
| **TP53BP1** | **S1028** | 0.04 | -0.06 | -0.18 | -0.07 | 0.16 | 0.09 | 0.243 |
| **TP53BP1** | **S754; S755** | 0.01 | -0.12 | -0.08 | 0.14 | -0.03 | 0.07 | 0.114 |
| **TP53BP1** | **S625; S630** | -0.19 | 0.02 | -0.02 | 0.01 | -0.08 | 0.22 | 0.355 |
| **TP53BP1** | **S523; S525** | -0.61 | 0.14 | 0.12 | -0.63 | 0.03 | 0.58 | 0.808 |
| **TP53BP1** | **S635; S639; S640** | -0.08 | -0.09 | 0.00 | 0.14 | -0.02 | 0.03 | 0.124 |
| **TP53BP1** | **S500** | -0.02 | -0.02 | -0.08 | -0.08 | 0.27 | -0.09 | 0.567 |
| **TP53BP1** | **S380** | 0.01 | -0.06 | -0.05 | -0.05 | 0.19 | -0.05 | 0.478 |
| **TP53BP1** | **S294** | 0.07 | 0.02 | -0.18 | 0.01 | -0.09 | 0.15 | 0.62 |
| **TP53BP1** | **S366** | -0.12 | 0.06 | 0.00 | -0.03 | 0.12 | -0.04 | 0.632 |
| **TP53BP1** | **S1462** | 0.01 | -0.01 | -0.06 | -0.17 | 0.18 | 0.04 | 0.739 |
| **TP53BP1** | **S265** | 0.00 | -0.06 | 0.00 | 0.02 | 0.17 | -0.14 | 0.739 |
| **TP53BP1** | **S/T** | -0.07 | 0.04 | -0.02 | 0.08 | 0.10 | -0.13 | 0.724 |
| **TP53BP1** | **S639; S640** | -0.06 | 0.02 | -0.01 | 0.03 | 0.04 | -0.04 | 0.495 |
| **TP53BP1** | **S552** | 0.03 | -0.03 | -0.03 | -0.03 | 0.01 | 0.06 | 0.559 |
| **TP53BP1** | **S639** | -0.01 | 0.08 | -0.10 | 0.10 | 0.19 | -0.30 | 0.97 |
| **TP53BP1** | **S265** | 0.08 | -0.07 | -0.04 | 0.03 | 0.24 | -0.28 | 0.973 |
| **TP53BP1** | **S1462** | -0.16 | 0.01 | 0.13 | -0.19 | 0.10 | 0.09 | 0.973 |
| **TP53BP1** | **S552** | 0.01 | 0.00 | -0.01 | -0.06 | 0.08 | -0.03 | 0.946 |
| **TP53BP1** | **S727** | 0.05 | 0.03 | -0.08 | -0.02 | -0.02 | 0.04 | 0.948 |
| **TP53BP1** | **S1678; T/S** | 0.15 | -0.02 | -0.13 | 0.08 | -0.06 | -0.05 | 0.928 |

| | | | | | | | | |
|---|---|---|---|---|---|---|---|---|
| **TP53BP1** | **S1295; S1296** | -0.25 | 0.27 | -0.03 | -0.21 | -0.02 | 0.17 | 0.935 |
| **TP53BP1** | **T/S** | 0.08 | 0.04 | -0.08 | 0.04 | -0.02 | -0.06 | 0.674 |
| **TP53BP1** | **S294** | 0.15 | 0.03 | -0.16 | 0.07 | -0.25 | 0.10 | 0.842 |
| **TP53BP1** | **S/T** | 0.16 | 0.12 | -0.25 | 0.07 | -0.06 | -0.08 | 0.82 |
| **TP53BP1** | **S/T** | 0.01 | 0.07 | 0.01 | -0.09 | 0.01 | -0.01 | 0.186 |
| **TP53BP1** | **S784** | 0.04 | -0.07 | 0.11 | -0.21 | -0.04 | 0.14 | 0.6 |
| **TP53BP1** | **S1678** | 0.11 | -0.01 | 0.00 | -0.13 | -0.02 | 0.04 | 0.37 |
| **TP53BP1** | **S1462; S1481** | -0.28 | 0.04 | 0.29 | -0.32 | -0.05 | 0.21 | 0.771 |
| **TP53BP1** | **S552** | 0.08 | 0.02 | 0.01 | -0.08 | 0.10 | -0.15 | 0.361 |
| **TP53BP1** | **T/S** | 0.02 | -0.04 | 0.13 | -0.16 | 0.08 | -0.05 | 0.385 |
| **TP53BP1** | **S727** | 0.12 | 0.16 | -0.16 | -0.21 | -0.03 | 0.08 | 0.517 |
| **TP53BP1** | **S/T** | -0.06 | 0.14 | 0.04 | -0.11 | -0.24 | 0.19 | 0.537 |
| **TP53BP1** | **S1028** | 0.07 | -0.01 | 0.13 | -0.17 | 0.00 | -0.04 | 0.111 |
| **TP53BP1** | **S316** | 0.05 | 0.03 | 0.10 | -0.14 | 0.17 | -0.27 | 0.356 |
| **TP53BP1** | **S1635** | 0.11 | 0.08 | 0.06 | -0.20 | 0.03 | -0.10 | 0.058 |
| **TP53BP1** | **T/S** | 0.08 | 0.12 | 0.05 | -0.21 | 0.08 | -0.15 | 0.127 |
| **TP53BP1** | **S630** | 0.20 | 0.06 | 0.01 | -0.03 | -0.07 | -0.20 | 0.064 |
| **TP53BP1** | **S/T** | 0.15 | 0.19 | 0.04 | -0.16 | 0.12 | -0.45 | 0.16 |
| **TP53BP1** | **S525** | 0.28 | 0.16 | -0.04 | -0.07 | -0.10 | -0.31 | 0.072 |
| **TP53BP1** | **S1462** | 0.13 | 0.28 | 0.06 | 0.10 | -0.25 | -0.44 | 0.106 |
| **TP53BP1** | **S1678** | 0.19 | 0.21 | 0.12 | -0.39 | 0.11 | -0.36 | 0.082 |
| **TP53BP1** | **S525** | 0.38 | 0.05 | 0.12 | -0.33 | -0.27 | -0.06 | 0.036 |
| **TP53BP1** | **S1635; T1638** | 0.07 | 0.06 | 0.76 | -0.60 | -0.29 | -0.42 | 0.042 |
| **TP53BP1** | **T1648** | 0.33 | 0.32 | 0.50 | -0.54 | -0.26 | -0.86 | 0.007 |
| **TP53BP2** | **S698** | 0.07 | 0.09 | -0.02 | 0.27 | -0.23 | -0.25 | 0.549 |
| **TP53I11** | **S14** | 0.14 | 0.04 | 0.14 | 0.02 | -0.09 | -0.30 | 0.077 |
| **TP53I11** | **S14** | 0.03 | 0.21 | 0.19 | -0.16 | -0.07 | -0.27 | 0.022 |
| **TPD52L1** | **S149** | 0.04 | 0.08 | 0.27 | -0.16 | 0.11 | -0.44 | 0.165 |
| **TPD52L2** | **S166** | 0.08 | -0.03 | 0.49 | -0.35 | 0.03 | -0.42 | 0.111 |
| **TPR** | **S2034** | -0.51 | -0.22 | -0.05 | 0.30 | 0.16 | 0.17 | 0.03 |
| **TPR** | **S2037** | -0.31 | -0.25 | -0.01 | 0.06 | 0.11 | 0.32 | 0.041 |
| **TPR** | **S/T** | 0.02 | -0.11 | -0.31 | 0.11 | -0.08 | 0.30 | 0.179 |
| **TPR** | **S379** | 0.02 | 0.04 | -0.21 | 0.03 | 0.04 | 0.06 | 0.295 |
| **TPR** | **S1185** | 0.11 | -0.01 | -0.23 | -0.08 | 0.05 | 0.13 | 0.556 |
| **TPR** | **S379** | 0.01 | 0.02 | -0.13 | 0.00 | 0.02 | 0.08 | 0.292 |
| **TPR** | **S2034; S2037** | -0.42 | 0.12 | 0.28 | -0.03 | -0.14 | 0.09 | 0.911 |
| **TPRN** | **T102** | -0.38 | -0.39 | -0.54 | 0.41 | -0.40 | 0.76 | 0.115 |
| **TPRN** | **S264; S274** | -0.37 | -0.41 | -0.30 | 0.35 | -0.21 | 0.60 | 0.065 |
| **TPRN** | **S362** | -0.41 | 0.30 | -0.59 | 0.20 | 0.24 | 0.02 | 0.238 |
| **TPRN** | **S362** | -0.19 | 0.15 | -0.41 | 0.00 | 0.22 | 0.14 | 0.201 |
| **TPRN** | **S362** | -0.05 | -0.15 | -0.15 | 0.13 | 0.12 | 0.07 | 0.005 |
| **TPRN** | **S** | -0.11 | -0.14 | -0.04 | -0.12 | 0.39 | -0.05 | 0.35 |
| **TPRN** | **S274** | -0.16 | 0.00 | 0.05 | 0.02 | -0.18 | 0.23 | 0.695 |
| **TPRN** | **S274** | -0.09 | 0.10 | -0.14 | -0.22 | -0.17 | 0.41 | 0.824 |
| **TPRN** | **S418** | -0.03 | 0.05 | 0.18 | -0.15 | 0.11 | -0.20 | 0.254 |
| **TPRN** | **S241** | 0.38 | 0.22 | 0.11 | -0.16 | -0.25 | -0.50 | 0.013 |
| **TPRN** | **S241** | 0.35 | 0.25 | 0.15 | -0.21 | -0.19 | -0.55 | 0.012 |
| **TPX2** | **S** | 0.13 | -0.04 | -0.01 | 0.01 | -0.21 | 0.11 | 0.609 |

| | | | | | | | | |
|---|---|---|---|---|---|---|---|---|
| **TPX2** | **S738** | 0.09 | 0.17 | 0.18 | -0.14 | -0.02 | -0.34 | 0.03 |
| **TRA2A** | **S236** | -0.11 | -0.02 | -0.05 | -0.01 | 0.08 | 0.11 | 0.058 |
| **TRA2A** | **S86; T88** | 0.19 | 0.08 | 0.17 | -0.16 | -0.13 | -0.21 | 0.002 |
| **TRA2A** | **S98; S100** | 0.25 | 0.18 | -0.01 | 0.15 | -0.03 | -0.75 | 0.283 |
| **TRA2A** | **S86** | 0.10 | 0.27 | 0.17 | -0.06 | -0.43 | -0.17 | 0.029 |
| **TRA2A** | **S16; S20** | 0.15 | 0.29 | 0.27 | -0.40 | 0.15 | -0.75 | 0.1 |
| **TRA2A** | **S100** | 0.26 | 0.35 | 0.30 | -0.26 | -0.05 | -1.01 | 0.065 |
| **TRA2A** | **S100** | 0.22 | 0.34 | 0.36 | -0.35 | -0.02 | -0.95 | 0.054 |
| **TRA2B** | **S22; S26** | 0.01 | 0.00 | -0.14 | 0.10 | -0.04 | 0.07 | 0.231 |
| **TRA2B** | **S95; S97; S99** | 0.08 | -0.21 | 0.06 | -0.11 | -0.11 | 0.24 | 0.843 |
| **TRA2B** | **S264; S266** | 0.02 | 0.03 | -0.07 | 0.05 | -0.07 | 0.04 | 0.737 |
| **TRA2B** | **S95; S97; S99** | 0.07 | 0.04 | 0.08 | 0.03 | -0.09 | -0.14 | 0.058 |
| **TRA2B** | **S95; S99** | -0.09 | 0.07 | 0.21 | -0.04 | -0.07 | -0.11 | 0.196 |
| **TRA2B** | **S95; S99** | 0.07 | 0.09 | 0.06 | 0.02 | -0.10 | -0.16 | 0.045 |
| **TRA2B** | **S264; S266** | 0.10 | 0.08 | 0.05 | -0.02 | -0.01 | -0.22 | 0.077 |
| **TRA2B** | **S264; S266** | 0.15 | 0.07 | 0.02 | 0.01 | -0.08 | -0.20 | 0.082 |
| **TRA2B** | **S99** | 0.03 | 0.22 | -0.05 | 0.07 | 0.17 | -0.57 | 0.516 |
| **TRA2B** | **S266** | 0.15 | 0.18 | 0.08 | -0.11 | -0.02 | -0.34 | 0.044 |
| **TRA2B** | **S97; S99** | 0.10 | 0.17 | 0.18 | -0.07 | -0.15 | -0.29 | 0.009 |
| **TRA2B** | **S26; S29** | -0.09 | 0.41 | 0.22 | -0.13 | 0.01 | -0.63 | 0.153 |
| **TRA2B** | **S282; S284** | 0.25 | 0.27 | 0.15 | -0.27 | 0.00 | -0.58 | 0.041 |
| **TRA2B** | **S266** | 0.22 | 0.28 | 0.24 | -0.19 | -0.08 | -0.69 | 0.04 |
| **TRA2B** | **S26** | 0.35 | 0.23 | 0.19 | -0.01 | -0.53 | -0.46 | 0.026 |
| **TRA2B** | **S97; S99** | 0.27 | 0.30 | 0.18 | -0.13 | -0.08 | -0.80 | 0.067 |
| **TRA2B** | **S97** | 0.32 | 0.22 | 0.36 | -0.08 | -0.12 | -1.21 | 0.107 |
| **TRAF4** | **S426** | -0.06 | -0.07 | -0.23 | 0.17 | -0.03 | 0.19 | 0.063 |
| **TRAF7** | **S108** | 0.18 | 0.03 | 0.11 | 0.06 | 0.00 | -0.46 | 0.233 |
| **TRAFD1** | **S/T** | -0.41 | -0.66 | -0.59 | 0.39 | 0.22 | 0.56 | 0.002 |
| **TRAFD1** | **S415** | -0.17 | -0.06 | -0.05 | 0.11 | 0.09 | 0.06 | 0.013 |
| **TRAFD1** | **T/S** | -0.51 | -0.05 | 0.24 | 0.04 | 0.09 | 0.09 | 0.463 |
| **TRAK1** | **S393** | -0.02 | 0.04 | 0.01 | -0.03 | -0.12 | 0.11 | 0.754 |
| **TRAM1** | **S365** | -0.07 | -0.04 | 0.06 | 0.04 | -0.19 | 0.18 | 0.866 |
| **TRAM1** | **S365** | -0.03 | 0.01 | 0.08 | 0.02 | -0.19 | 0.09 | 0.624 |
| **TRAM1** | **S365** | 0.08 | 0.08 | 0.14 | -0.09 | -0.21 | -0.04 | 0.018 |
| **TRAM1** | **S365** | -0.01 | 0.15 | 0.21 | -0.15 | -0.25 | -0.01 | 0.058 |
| **TRAM1** | **S365** | 0.28 | 0.34 | 0.23 | -0.34 | -0.15 | -0.62 | 0.009 |
| **TRAM1** | **S365** | 0.26 | 0.24 | 0.42 | -0.38 | -0.26 | -0.54 | 0.002 |
| **TRAM1** | **S365** | 0.39 | 0.38 | 0.47 | -0.57 | -0.31 | -0.96 | 0.006 |
| **TRAM1** | **S365** | 0.35 | 0.49 | 0.62 | -0.66 | -0.36 | -1.44 | 0.017 |
| **TRAPPC10** | **T184** | -0.15 | -0.11 | -0.71 | -0.12 | 0.19 | 0.58 | 0.129 |
| **TRAPPC9** | **S953** | -0.23 | 0.00 | -0.08 | 0.06 | -0.26 | 0.41 | 0.458 |
| **TRIB3** | **S327** | -0.08 | 0.03 | 0.08 | -0.25 | 0.39 | -0.28 | 0.803 |
| **TRIM13** | **S171** | -0.16 | 0.04 | -0.45 | 0.06 | 0.05 | 0.34 | 0.121 |
| **TRIM16** | **S/T** | -0.15 | 0.01 | -0.45 | 0.34 | 0.12 | 0.02 | 0.098 |
| **TRIM24** | **S1028** | -0.16 | -0.13 | -0.12 | 0.19 | -0.07 | 0.23 | 0.053 |
| **TRIM24** | **S811** | 0.01 | 0.09 | -0.23 | 0.07 | -0.11 | 0.13 | 0.58 |
| **TRIM24** | **S1025; S1028** | 0.10 | 0.02 | 0.12 | -0.11 | -0.02 | -0.12 | 0.02 |
| **TRIM24** | **S744** | 0.07 | 0.19 | 0.29 | -0.18 | -0.09 | -0.39 | 0.02 |

| | | | | | | | | |
|---|---|---|---|---|---|---|---|---|
| **TRIM24** | **S/T** | -0.19 | 0.33 | 0.45 | -0.05 | -0.51 | -0.26 | 0.118 |
| **TRIM24** | **S811** | 0.24 | 0.24 | 0.30 | 0.05 | -0.40 | -0.69 | 0.05 |
| **TRIM24** | **S1028** | 0.19 | 0.31 | 0.41 | -0.24 | -0.19 | -0.81 | 0.027 |
| **TRIM25** | **S100** | 0.08 | -0.12 | -0.01 | 0.40 | -0.25 | -0.20 | 0.988 |
| **TRIM25** | **S100** | 0.06 | 0.07 | -0.05 | -0.08 | 0.14 | -0.16 | 0.564 |
| **TRIM28** | **S/T** | -0.51 | -0.39 | -0.55 | 0.32 | 0.49 | 0.26 | 6E-04 |
| **TRIM28** | **S473** | -0.18 | -0.23 | -0.39 | 0.26 | 0.21 | 0.20 | 0.002 |
| **TRIM28** | **S594; S/T** | -0.41 | 0.11 | -0.14 | 0.14 | -0.10 | 0.30 | 0.237 |
| **TRIM28** | **S752** | -0.06 | -0.08 | -0.10 | 0.08 | 0.09 | 0.06 | 5E-04 |
| **TRIM28** | **S/T** | -0.13 | -0.09 | 0.16 | 0.04 | 0.22 | -0.24 | 0.883 |
| **TRIM28** | **S594** | -0.06 | 0.17 | 0.04 | 0.02 | -0.18 | -0.01 | 0.306 |
| **TRIM28** | **T/S** | -0.40 | 0.44 | 0.05 | 0.28 | -0.35 | -0.24 | 0.691 |
| **TRIM28** | **S752** | 0.21 | 0.04 | 0.25 | -0.63 | 0.13 | -0.17 | 0.162 |
| **TRIM3** | **S427** | -0.02 | 0.03 | -0.10 | 0.03 | -0.01 | 0.06 | 0.269 |
| **TRIM3** | **S7** | -0.07 | 0.02 | 0.04 | 0.04 | -0.08 | 0.03 | 0.989 |
| **TRIM3** | **S437** | 0.05 | 0.08 | -0.03 | -0.04 | -0.06 | -0.01 | 0.126 |
| **TRIM3** | **S7** | 0.03 | 0.03 | 0.08 | 0.03 | 0.02 | -0.21 | 0.273 |
| **TRIM3** | **S427** | 0.01 | 0.09 | 0.07 | -0.08 | -0.01 | -0.08 | 0.023 |
| **TRIM3** | **S437** | 0.04 | 0.21 | 0.07 | -0.13 | 0.11 | -0.38 | 0.195 |
| **TRIM41** | **S571** | 0.39 | -0.02 | -0.13 | 0.32 | -0.96 | 0.04 | 0.544 |
| **TRIM8** | **S321** | -0.05 | 0.12 | -0.03 | -0.10 | 0.10 | -0.05 | 0.734 |
| **TRIP10** | **S296** | 0.06 | 0.09 | -0.17 | 0.04 | -0.32 | 0.24 | 0.97 |
| **TRIP12** | **S312** | -0.05 | -0.15 | -0.06 | 0.12 | -0.12 | 0.23 | 0.204 |
| **TRIP12** | **S1078** | 0.01 | -0.10 | -0.14 | -0.02 | 0.10 | 0.12 | 0.084 |
| **TRIP12** | **S1030; T/S** | -0.05 | -0.10 | -0.02 | 0.02 | 0.01 | 0.12 | 0.065 |
| **TRIP12** | **S1317; S1322** | -0.13 | 0.00 | 0.09 | -0.18 | -0.01 | 0.20 | 0.882 |
| **TRIP12** | **S85; S88** | -0.17 | 0.08 | 0.06 | -0.06 | 0.11 | -0.04 | 0.916 |
| **TRIP12** | **S991** | 0.01 | -0.13 | 0.17 | -0.13 | 0.01 | 0.04 | 0.685 |
| **TRIP12** | **S1030** | -0.01 | 0.05 | 0.05 | -0.16 | 0.18 | -0.15 | 0.57 |
| **TRIP12** | **S/T** | -0.05 | 0.15 | 0.05 | -0.01 | -0.29 | 0.12 | 0.47 |
| **TRIP12** | **S1376** | -0.04 | 0.09 | 0.14 | -0.19 | -0.02 | 0.00 | 0.184 |
| **TRIP12** | **S1016** | 0.13 | 0.05 | 0.01 | -0.19 | -0.04 | 0.02 | 0.137 |
| **TRIP12** | **S77** | 0.10 | 0.04 | 0.07 | -0.20 | 0.06 | -0.11 | 0.126 |
| **TRIP12** | **S312** | 0.07 | 0.13 | 0.07 | -0.27 | 0.03 | -0.07 | 0.105 |
| **TRIP12** | **T/S** | 0.12 | 0.19 | -0.03 | -0.39 | -0.03 | 0.07 | 0.244 |
| **TRIP12** | **S100** | 0.17 | 0.20 | -0.03 | -0.27 | -0.11 | -0.02 | 0.076 |
| **TRIP12** | **S1036** | 0.22 | 0.12 | 0.02 | -0.21 | -0.11 | -0.08 | 0.025 |
| **TRIP12** | **S/T** | 0.21 | 0.15 | 0.02 | -0.27 | -0.17 | 0.01 | 0.052 |
| **TRIP12** | **S1016** | 0.20 | 0.20 | 0.04 | -0.17 | 0.03 | -0.38 | 0.073 |
| **TRIP12** | **S942** | 0.03 | 0.25 | 0.33 | -0.19 | -0.24 | -0.32 | 0.01 |
| **TRIP12** | **S85** | 0.26 | 0.20 | 0.36 | -0.55 | 0.04 | -0.62 | 0.04 |
| **TRMT1** | **S625** | 0.02 | 0.02 | -0.11 | 0.12 | 0.03 | -0.09 | 0.597 |
| **TRMT10A** | **S318** | -0.01 | 0.06 | 0.27 | -0.13 | -0.23 | 0.00 | 0.105 |
| **TRMT112** | **S125** | -0.80 | -0.04 | -0.17 | 0.52 | -0.10 | 0.26 | 0.132 |
| **TRMT112** | **S119** | 0.14 | 0.03 | -0.67 | 0.37 | -0.07 | 0.01 | 0.402 |
| **TRMT1L** | **T26** | 0.11 | 0.05 | -0.10 | -0.12 | 0.10 | -0.06 | 0.65 |
| **TRMT2A** | **T/S** | 0.01 | -0.05 | -0.16 | 0.21 | -0.05 | 0.02 | 0.247 |
| **TRMT2A** | **S602** | 0.08 | 0.37 | -0.21 | -0.40 | 0.02 | 0.03 | 0.419 |

| | | | | | | | | |
|---|---|---|---|---|---|---|---|---|
| TRMT2A | S602 | 0.05 | 0.45 | -0.16 | -0.06 | -0.01 | -0.40 | 0.283 |
| TRMT6 | S/T | -0.68 | -0.60 | -0.66 | 0.50 | 0.46 | 0.36 | 2E-05 |
| TRMT6 | T291 | -0.49 | -0.58 | -0.59 | 0.48 | 0.24 | 0.46 | 4E-04 |
| TRMT61A | T/S | -0.11 | 0.06 | -0.07 | 0.07 | 0.01 | 0.04 | 0.236 |
| TRPM4 | S255 | 0.02 | -0.33 | -0.31 | 0.02 | 0.16 | 0.32 | 0.06 |
| TRPM4 | S1103 | -0.09 | -0.14 | -0.06 | 0.17 | -0.03 | 0.13 | 0.043 |
| TRPM4 | S728 | -0.04 | -0.07 | -0.02 | 0.10 | -0.05 | 0.06 | 0.171 |
| TRPM7 | S1404 | -0.61 | -0.35 | -0.41 | 0.50 | -0.26 | 0.64 | 0.061 |
| TRPM7 | S101 | -0.82 | 0.12 | 0.22 | 0.49 | -0.55 | 0.12 | 0.712 |
| TRPM7 | S/T | 0.07 | -0.09 | -0.31 | -0.17 | -0.27 | 0.57 | 0.622 |
| TRPM7 | S/T | -0.05 | 0.03 | -0.03 | 0.03 | -0.08 | 0.10 | 0.601 |
| TRPM7 | S1569 | -0.29 | 0.25 | 0.26 | -0.24 | 0.03 | -0.11 | 0.411 |
| TRPM7 | S101 | -0.02 | -0.14 | 0.56 | -0.11 | 0.02 | -0.54 | 0.276 |
| TRPM7 | S/T | 0.09 | 0.25 | 0.16 | -0.24 | 0.07 | -0.46 | 0.077 |
| TRPS1 | S99 | -1.25 | -1.51 | -1.27 | 0.89 | 0.52 | 0.60 | 1E-04 |
| TRPS1 | T/S | -0.46 | -0.52 | -0.29 | 0.33 | 0.24 | 0.41 | 8E-04 |
| TRPS1 | S803; S805 | -0.21 | -0.24 | -0.37 | 0.28 | 0.13 | 0.27 | 0.002 |
| TRPS1 | T142 | -0.18 | -0.17 | -0.24 | 0.19 | 0.22 | 0.10 | 9E-04 |
| TRPS1 | T142; S147 | -0.03 | -0.07 | -0.47 | 0.27 | -0.10 | 0.28 | 0.146 |
| TRPS1 | S1062; S1066 | -0.02 | -0.08 | -0.24 | 0.12 | -0.12 | 0.27 | 0.199 |
| TRPS1 | S90 | 0.02 | -0.16 | -0.18 | 0.10 | -0.21 | 0.35 | 0.347 |
| TRPS1 | T323 | 0.16 | -0.13 | -0.28 | 0.11 | -0.12 | 0.20 | 0.404 |
| TRPS1 | S391 | 0.02 | -0.02 | -0.20 | -0.02 | 0.25 | -0.06 | 0.352 |
| TRPS1 | S178 | 0.02 | 0.27 | -0.53 | 0.20 | -0.21 | 0.10 | 0.705 |
| TRPS1 | S1041 | 0.15 | 0.04 | -0.23 | -0.10 | 0.27 | -0.20 | 0.995 |
| TRPS1 | S1081 | 0.02 | 0.05 | -0.05 | -0.01 | 0.04 | -0.07 | 0.631 |
| TRPS1 | S1041 | 0.11 | 0.06 | -0.13 | 0.04 | -0.12 | 0.03 | 0.721 |
| TRPS1 | S216 | 0.09 | -0.01 | -0.03 | -0.16 | 0.22 | -0.15 | 0.742 |
| TRPS1 | S178 | 0.04 | 0.06 | -0.01 | -0.03 | -0.11 | 0.05 | 0.307 |
| TRPS1 | S803 | 0.09 | 0.11 | -0.10 | -0.08 | -0.15 | 0.11 | 0.528 |
| TRPS1 | S/T | 0.04 | 0.03 | 0.09 | -0.08 | 0.00 | -0.09 | 0.025 |
| TRPS1 | S1085 | 0.10 | 0.07 | -0.01 | -0.08 | 0.08 | -0.19 | 0.247 |
| TRPS1 | S14 | 0.17 | 0.00 | -0.02 | -0.22 | -0.12 | 0.14 | 0.388 |
| TRPS1 | S978 | 0.20 | -0.05 | 0.00 | 0.15 | -0.17 | -0.17 | 0.416 |
| TRPS1 | S178 | -0.02 | 0.01 | 0.19 | -0.05 | -0.19 | 0.04 | 0.253 |
| TRPS1 | S1081; S1085 | 0.11 | 0.15 | -0.04 | -0.06 | -0.14 | -0.03 | 0.085 |
| TRPS1 | S178 | 0.05 | 0.07 | 0.14 | -0.05 | -0.07 | -0.16 | 0.015 |
| TRPS1 | S978 | 0.08 | 0.15 | 0.13 | -0.21 | -0.21 | 0.01 | 0.029 |
| TRPS1 | T751 | 0.15 | 0.08 | 0.24 | 0.02 | 0.05 | -0.72 | 0.212 |
| TRPS1 | S216 | 0.22 | 0.27 | 0.13 | -0.35 | 0.03 | -0.47 | 0.041 |
| TRPS1 | S1066 | 0.33 | 0.26 | 0.31 | -0.21 | -0.06 | -1.04 | 0.072 |
| TRPT1 | T/S | 0.19 | -0.02 | -0.15 | -0.02 | -0.09 | 0.06 | 0.864 |
| TRPT1 | S240 | 0.14 | 0.11 | -0.07 | -0.09 | -0.15 | 0.03 | 0.198 |
| TRRAP | S2051 | -0.24 | -0.14 | -0.12 | 0.28 | -0.06 | 0.21 | 0.045 |
| TSC2 | S | 0.01 | 0.01 | -0.07 | 0.07 | 0.09 | -0.12 | 0.669 |
| TSC2 | S1411 | 0.29 | -0.02 | 0.00 | -0.04 | -0.15 | -0.13 | 0.134 |
| TSC22D3 | S102 | -0.07 | -0.30 | 0.12 | -0.11 | 0.06 | 0.25 | 0.396 |
| TSC22D3 | S122 | 0.01 | -0.19 | 0.03 | 0.05 | 0.07 | 0.02 | 0.25 |

| | | | | | | | | |
|---|---|---|---|---|---|---|---|---|
| TSC22D4 | S62 | -0.32 | -0.21 | 0.29 | -0.07 | 0.21 | 0.00 | 0.574 |
| TSEN34 | S | 0.01 | -0.39 | -0.46 | -0.11 | 0.29 | 0.44 | 0.09 |
| TSEN34 | S | 0.12 | -0.10 | -0.12 | -0.30 | 0.16 | 0.18 | 0.809 |
| TSG101 | T220 | -0.08 | -0.12 | 0.04 | -0.05 | -0.03 | 0.21 | 0.368 |
| TSG101 | S309 | 0.42 | -0.18 | -0.11 | 0.14 | 0.02 | -0.44 | 0.627 |
| TSG101 | S309 | 0.25 | -0.02 | -0.03 | 0.51 | -0.08 | -1.07 | 0.583 |
| TSHZ1 | S489 | 0.04 | -0.06 | 0.17 | 0.24 | -0.20 | -0.26 | 0.507 |
| TSHZ1 | S489 | 0.04 | 0.21 | 0.06 | 0.03 | -0.23 | -0.16 | 0.073 |
| TSNARE1 | S378 | -0.01 | -0.04 | -0.10 | 0.17 | 0.16 | -0.21 | 0.535 |
| TSNARE1 | S47 | 0.04 | -0.01 | 0.06 | 0.13 | -0.10 | -0.14 | 0.505 |
| TSPYL1 | S/T | -0.36 | -0.35 | -0.24 | 0.24 | 0.07 | 0.45 | 0.008 |
| TSPYL2 | S16 | -0.04 | -0.40 | -0.44 | 0.18 | -0.07 | 0.54 | 0.082 |
| TTC14 | S525 | 0.10 | 0.11 | 0.32 | 0.06 | -0.15 | -0.61 | 0.124 |
| TTC14 | S671 | 0.38 | 0.57 | 0.56 | -0.69 | -0.60 | -1.12 | 0.002 |
| TTC3 | S1819 | -0.09 | -0.04 | 0.24 | 0.05 | -0.13 | -0.07 | 0.499 |
| TTC33 | S197 | 0.04 | 0.07 | 0.40 | -0.72 | 0.09 | -0.10 | 0.202 |
| TTC7A | S647 | 0.06 | -0.04 | 0.07 | 0.00 | 0.07 | -0.17 | 0.459 |
| TTC7A | S51 | 0.43 | -0.47 | 0.28 | -0.23 | -0.12 | -0.08 | 0.468 |
| TTC9 | S15 | 0.13 | 0.10 | -0.21 | -0.08 | -0.08 | 0.10 | 0.843 |
| TTC9 | S15 | 0.17 | 0.10 | -0.17 | -0.05 | -0.02 | -0.06 | 0.501 |
| TTF1 | S403 | -0.21 | -0.11 | -0.21 | -0.06 | 0.23 | 0.27 | 0.041 |
| TTF1 | S478; S481; S487 | -0.10 | -0.10 | -0.11 | 0.16 | -0.12 | 0.22 | 0.144 |
| TTF1 | S487 | -0.02 | 0.20 | -0.12 | 0.09 | 0.03 | -0.21 | 0.726 |
| TTF1 | S472 | -0.12 | 1.22 | -0.94 | -0.61 | -0.30 | -0.40 | 0.484 |
| TTLL12 | S16 | -0.28 | -0.16 | -0.56 | 0.30 | 0.41 | 0.06 | 0.021 |
| TTYH3 | S496 | 0.11 | -0.11 | 0.09 | 0.01 | -0.30 | 0.15 | 0.61 |
| TUBA1B | S439 | -0.07 | -0.09 | 0.13 | -0.02 | 0.15 | -0.11 | 0.89 |
| TUBA1B | S439 | -0.01 | 0.08 | 0.06 | -0.07 | 0.09 | -0.17 | 0.317 |
| TUBA1C | T73 | -0.30 | -0.70 | -0.54 | 0.03 | 0.41 | 0.62 | 0.014 |
| TUBA1C | T334 | -0.13 | -0.18 | -0.19 | 0.28 | 0.05 | 0.11 | 0.012 |
| TUBA1C | S439 | -0.19 | -0.53 | 0.41 | 0.06 | 0.13 | -0.04 | 0.61 |
| TUBA1C | S48 | 0.06 | -0.07 | 0.03 | -0.11 | 0.17 | -0.11 | 0.831 |
| TUBA4A | T109 | -0.05 | -0.04 | -0.46 | 0.28 | 0.00 | 0.17 | 0.104 |
| TUBA4A | S158 | 0.07 | -0.15 | 0.01 | 0.10 | 0.07 | -0.12 | 0.701 |
| TUBA4A | T223 | -0.14 | 0.25 | 0.06 | 0.03 | -0.09 | -0.15 | 0.377 |
| TUBB | S48 | 0.03 | -0.04 | -0.12 | -0.05 | 0.34 | -0.23 | 0.751 |
| TUBB | S48 | 0.13 | 0.03 | -0.19 | -0.07 | 0.39 | -0.42 | 0.928 |
| TUBB | S75 | 0.00 | 0.19 | -0.10 | -0.14 | -0.02 | 0.04 | 0.526 |
| TUBGCP3 | S896 | -0.05 | -0.41 | -0.23 | 0.12 | 0.39 | 0.05 | 0.045 |
| TUBGCP3 | S515 | -0.28 | -0.02 | -0.13 | -0.07 | 0.15 | 0.28 | 0.105 |
| TXLNA | S | 0.13 | 0.25 | 0.01 | -0.04 | -0.18 | -0.22 | 0.033 |
| TXLNA | S515 | 0.04 | 0.43 | -0.08 | -0.08 | -0.10 | -0.31 | 0.157 |
| TXLNG | S97 | 0.20 | -0.12 | 0.34 | -0.72 | 0.37 | -0.39 | 0.33 |
| TYK2 | Y292 | -0.06 | -0.03 | -0.13 | 0.18 | 0.14 | -0.13 | 0.252 |
| U2AF1 | S | -0.92 | -0.64 | -0.68 | 0.59 | 0.67 | 0.15 | 0.003 |
| U2AF2 | S79 | 0.12 | 0.08 | -0.01 | -0.11 | -0.06 | -0.02 | 0.053 |
| U2SURP | S767 | 0.31 | -0.19 | -0.29 | -0.16 | -0.11 | 0.31 | 0.787 |
| U2SURP | S238 | -0.18 | 0.00 | 0.14 | -0.07 | -0.33 | 0.34 | 0.976 |

| | | | | | | | | |
|---|---|---|---|---|---|---|---|---|
| U2SURP | T919 | 0.14 | 0.04 | 0.02 | -0.02 | -0.10 | -0.10 | 0.041 |
| U2SURP | S946; S948 | 0.25 | 0.16 | 0.01 | -0.16 | 0.00 | -0.34 | 0.063 |
| U2SURP | S930; S934 | 0.02 | 0.21 | 0.28 | -0.28 | 0.18 | -0.61 | 0.17 |
| U2SURP | T919 | 0.28 | 0.21 | 0.09 | -0.04 | -0.10 | -0.60 | 0.074 |
| UBA52 | T12 | 0.12 | -0.20 | -0.06 | -0.32 | -0.17 | 0.47 | 0.887 |
| UBAC1 | S98 | -0.03 | 0.00 | -0.38 | 0.42 | 0.30 | -0.55 | 0.58 |
| UBAP2 | S254 | 0.04 | -0.03 | 0.06 | 0.01 | 0.04 | -0.12 | 0.456 |
| UBAP2L | S609 | 0.15 | -0.24 | -0.57 | -0.19 | 0.29 | 0.35 | 0.242 |
| UBAP2L | S607; S/Y | -0.24 | -0.07 | 0.17 | 0.03 | -0.11 | 0.18 | 0.596 |
| UBAP2L | S609 | -0.15 | -0.12 | 0.17 | 0.01 | -0.12 | 0.17 | 0.704 |
| UBAP2L | S467 | -0.05 | 0.02 | 0.19 | -0.01 | 0.16 | -0.38 | 0.496 |
| UBAP2L | S/T | 0.01 | -0.03 | 0.29 | -0.12 | 0.07 | -0.30 | 0.227 |
| UBAP2L | S/T | -0.01 | 0.19 | 0.15 | -0.18 | 0.22 | -0.51 | 0.294 |
| UBAP2L | S467 | 0.11 | 0.24 | 0.12 | -0.04 | 0.00 | -0.54 | 0.119 |
| UBAP2L | S467; S477 | 0.09 | 0.16 | 0.25 | -0.12 | -0.06 | -0.42 | 0.041 |
| UBAP2L | S476; S477; S/T | 0.20 | 0.16 | 0.21 | -0.06 | -0.16 | -0.47 | 0.027 |
| UBAP2L | S476; S477; S/T | 0.20 | 0.16 | 0.21 | -0.06 | -0.16 | -0.47 | 0.027 |
| UBAP2L | S476; S477 | 0.61 | 0.01 | -0.12 | -0.05 | -0.15 | -0.57 | 0.2 |
| UBAP2L | S467 | 0.28 | 0.09 | 0.18 | -0.76 | 0.48 | -0.71 | 0.279 |
| UBAP2L | S/T | 0.10 | 0.49 | 0.08 | 0.01 | -0.45 | -0.47 | 0.062 |
| UBE2J1 | S266 | -0.28 | 0.05 | 0.39 | -0.34 | 0.01 | 0.06 | 0.559 |
| UBE2J1 | S266 | 0.08 | 0.16 | 0.08 | -0.14 | -0.27 | 0.05 | 0.077 |
| UBE2J1 | S266 | 0.06 | 0.13 | 0.18 | -0.09 | -0.13 | -0.19 | 0.004 |
| UBE2O | S896 | 0.05 | 0.03 | -0.02 | -0.07 | -0.15 | 0.14 | 0.622 |
| UBE2O | S896 | 0.54 | -0.18 | -0.32 | -0.28 | 0.01 | 0.05 | 0.774 |
| UBE2O | S896 | 0.12 | 0.07 | 0.03 | -0.15 | 0.19 | -0.32 | 0.335 |
| UBE2O | T/S | 0.32 | 0.02 | 0.03 | -0.25 | 0.07 | -0.28 | 0.138 |
| UBE2O | T/S | 0.37 | 0.19 | 0.06 | -0.04 | -0.02 | -0.81 | 0.147 |
| UBE2O | T/S | 0.47 | 0.21 | -0.05 | 0.09 | -0.12 | -0.99 | 0.206 |
| UBE2O | S401 | 0.55 | 0.10 | 0.14 | -0.41 | 0.11 | -0.93 | 0.115 |
| UBE2O | S401 | 0.64 | 0.14 | 0.52 | -1.11 | 0.14 | -1.58 | 0.075 |
| UBN1 | S336 | -0.06 | -0.36 | -0.08 | 0.02 | 0.06 | 0.33 | 0.093 |
| UBN1 | S336; S338 | -0.11 | -0.06 | -0.07 | 0.07 | -0.04 | 0.19 | 0.096 |
| UBN1 | S336 | -0.09 | -0.02 | -0.02 | -0.06 | 0.07 | 0.11 | 0.231 |
| UBN1 | S173; S175 | -0.03 | 0.03 | 0.00 | 0.03 | 0.04 | -0.08 | 0.975 |
| UBN1 | S660 | 0.05 | -0.16 | 0.09 | -0.10 | -0.22 | 0.28 | 0.97 |
| UBN1 | S173; S175 | 0.05 | -0.04 | 0.01 | -0.16 | 0.04 | 0.08 | 0.862 |
| UBN1 | S175 | 0.33 | 0.24 | 0.16 | -0.29 | 0.00 | -0.68 | 0.047 |
| UBN2 | S13 | -0.41 | -1.05 | -0.30 | 0.16 | 0.04 | 0.86 | 0.053 |
| UBQLN4 | S122 | 0.09 | -0.12 | -0.04 | 0.10 | -0.40 | 0.27 | 0.951 |
| UBQLN4 | S114 | -0.03 | -0.08 | 0.14 | 0.19 | -0.40 | 0.10 | 0.83 |
| UBQLN4 | T95; S/T | -0.02 | 0.19 | 0.12 | -0.20 | -0.21 | 0.08 | 0.151 |
| UBR1 | T21 | 0.08 | -0.09 | 0.18 | 0.22 | -0.01 | -0.48 | 0.53 |
| UBR4 | S | 0.05 | -0.24 | -0.26 | 0.02 | 0.13 | 0.23 | 0.079 |
| UBR4 | S181 | -0.20 | 0.03 | -0.15 | 0.19 | 0.04 | 0.05 | 0.077 |
| UBR4 | T/S | -0.10 | -0.03 | -0.02 | 0.29 | -0.37 | 0.15 | 0.728 |
| UBR4 | S/T | -0.14 | 0.07 | 0.11 | -0.15 | 0.16 | -0.08 | 0.775 |
| UBR4 | S178 | -0.14 | 0.53 | 0.04 | -0.11 | -0.05 | -0.44 | 0.218 |

| | | | | | | | | |
|---|---|---|---|---|---|---|---|---|
| UBR5 | S/T | -0.68 | -0.48 | -1.20 | 0.40 | 0.52 | 0.56 | 0.004 |
| UBR5 | S327 | -0.03 | -0.07 | -0.15 | 0.00 | 0.04 | 0.19 | 0.082 |
| UBR5 | S1549 | -0.18 | -0.23 | 0.16 | 0.07 | 0.00 | 0.13 | 0.311 |
| UBTF | S484 | -0.05 | -0.06 | -0.32 | -0.09 | 0.15 | 0.28 | 0.143 |
| UBTF | S484 | -0.10 | -0.22 | -0.03 | 0.20 | -0.08 | 0.18 | 0.114 |
| UBTF | S389 | 0.22 | 0.21 | 0.11 | -0.40 | 0.14 | -0.43 | 0.095 |
| UBXN1 | S/Y/T | -0.33 | 0.29 | -0.44 | 0.44 | -0.01 | -0.17 | 0.442 |
| UBXN6 | S96 | -0.17 | -0.21 | -0.17 | 0.22 | -0.11 | 0.35 | 0.07 |
| UBXN7 | S297 | -1.29 | -0.95 | -0.73 | 0.72 | 0.55 | 0.44 | 0.001 |
| UBXN7 | S288 | 0.06 | 0.03 | 0.08 | -0.07 | 0.00 | -0.11 | 0.032 |
| UCKL1 | T547 | -0.05 | -0.16 | -0.01 | 0.01 | 0.07 | 0.12 | 0.058 |
| UFD1 | S247 | 0.08 | 0.24 | 0.05 | -0.16 | -0.22 | -0.04 | 0.033 |
| UFL1 | S458 | 0.14 | 0.00 | 0.11 | 0.09 | -0.44 | 0.03 | 0.327 |
| UFL1 | S458 | 0.09 | 0.10 | 0.10 | -0.01 | 0.00 | -0.32 | 0.123 |
| UFL1 | S458 | 0.37 | 0.13 | 0.17 | -0.26 | -0.36 | -0.20 | 0.004 |
| UGDH | T/S | 0.27 | -0.58 | 0.19 | -0.07 | -0.12 | 0.15 | 0.938 |
| UHRF1 | S108 | 0.02 | -0.07 | -0.09 | 0.16 | -0.24 | 0.18 | 0.577 |
| UHRF1 | S/T/Y | -0.04 | -0.08 | 0.04 | 0.02 | -0.08 | 0.14 | 0.524 |
| UHRF1 | S91 | 0.02 | -0.05 | 0.02 | 0.04 | -0.20 | 0.14 | 0.981 |
| UHRF1 | S95 | 0.24 | 0.06 | -0.22 | -0.11 | -0.30 | 0.24 | 0.695 |
| UHRF1 | S287 | 0.00 | 0.14 | 0.24 | 0.00 | -0.08 | -0.39 | 0.108 |
| UHRF1 | S287 | 0.05 | 0.16 | 0.20 | -0.11 | -0.03 | -0.34 | 0.043 |
| UHRF1 | S91 | 0.18 | 0.11 | 0.16 | -0.19 | -0.09 | -0.22 | 0.002 |
| UHRF1 | S287 | 0.03 | 0.21 | 0.27 | -0.16 | 0.01 | -0.49 | 0.078 |
| UHRF1 | S91 | 0.28 | 0.18 | 0.07 | -0.22 | -0.17 | -0.23 | 0.004 |
| UHRF2 | S667 | -0.06 | -0.11 | 0.00 | -0.01 | -0.04 | 0.21 | 0.253 |
| UIMC1 | S677 | -0.02 | -0.04 | -0.16 | 0.08 | 0.03 | 0.09 | 0.037 |
| UIMC1 | S653 | 0.09 | -0.05 | 0.03 | -0.07 | 0.02 | -0.02 | 0.394 |
| UIMC1 | S653 | -0.24 | -0.05 | 0.35 | -0.33 | 0.01 | 0.15 | 0.749 |
| ULK1 | S | -0.58 | 0.17 | 0.22 | 0.29 | -0.19 | -0.09 | 0.831 |
| ULK1 | S450 | 0.31 | -0.05 | 0.13 | -0.03 | -0.08 | -0.38 | 0.121 |
| UNC13B | S176 | -0.17 | -0.44 | -0.58 | 0.74 | 0.24 | -0.24 | 0.106 |
| UNC13B | S367 | -0.26 | 0.21 | -0.07 | 0.27 | 0.07 | -0.33 | 0.85 |
| Uncharacterizı | S294 | 0.36 | 1.00 | -0.66 | -0.09 | -0.71 | -0.92 | 0.211 |
| UNG | T60 | -0.02 | -0.43 | 0.11 | 0.18 | -0.38 | 0.37 | 0.566 |
| UNG | S63; S64 | -0.39 | 0.04 | 0.10 | 0.01 | -0.10 | 0.25 | 0.506 |
| UNG | S23 | 0.21 | -0.09 | -0.09 | 0.32 | -0.07 | -0.38 | 0.812 |
| UNG | T60 | 0.02 | 0.24 | -0.12 | -0.43 | -0.07 | 0.26 | 0.616 |
| UPF1 | S1107; S1110 | -0.10 | 0.00 | -0.06 | 0.03 | -0.12 | 0.23 | 0.389 |
| UPF1 | S1110; S/T/Y | -0.15 | -0.04 | 0.19 | -0.09 | -0.02 | 0.07 | 0.9 |
| UPF1 | S1107 | 0.00 | 0.15 | 0.24 | -0.27 | -0.04 | -0.13 | 0.046 |
| UPF3B | S409 | -0.04 | 0.02 | -0.08 | -0.05 | 0.11 | 0.03 | 0.268 |
| UQCRFS1 | S99 | 0.07 | 0.04 | -0.07 | 0.17 | -0.04 | -0.22 | 0.734 |
| UQCRH | S13 | 0.00 | -0.13 | -0.19 | 0.23 | -0.20 | 0.23 | 0.267 |
| URB1 | S1097 | -0.16 | -0.15 | -0.40 | 0.06 | 0.04 | 0.46 | 0.056 |
| URB1 | S9 | -0.08 | -0.11 | -0.54 | 0.49 | 0.32 | -0.34 | 0.246 |
| USF2 | S222 | 0.06 | 0.18 | 0.20 | -0.09 | -0.06 | -0.36 | 0.041 |
| USO1 | S952 | -0.20 | 0.17 | 0.03 | 0.30 | -0.17 | -0.22 | 0.899 |

| | | | | | | | | |
|---|---|---|---|---|---|---|---|---|
| USO1 | S942 | -0.08 | 0.24 | -0.13 | 0.17 | -0.02 | -0.24 | 0.806 |
| USP1 | S67 | -0.24 | -0.31 | 0.01 | 0.03 | 0.12 | 0.31 | 0.062 |
| USP1 | S67 | -0.06 | -0.17 | -0.19 | -0.06 | 0.08 | 0.32 | 0.104 |
| USP1 | S313 | -0.09 | 0.21 | 0.12 | -0.12 | -0.24 | 0.06 | 0.222 |
| USP10 | S718 | -0.06 | -0.04 | -0.22 | 0.10 | 0.04 | 0.15 | 0.035 |
| USP10 | S365 | -0.04 | 0.16 | -0.29 | 0.05 | 0.28 | -0.25 | 0.697 |
| USP10 | T100 | -0.41 | -0.05 | 0.67 | -0.23 | 0.19 | -0.53 | 0.534 |
| USP10 | S576 | 0.02 | 0.23 | 0.12 | -0.26 | -0.01 | -0.16 | 0.048 |
| USP13 | S630 | -0.14 | -0.64 | 0.27 | 0.40 | 0.07 | -0.20 | 0.449 |
| USP16 | S552 | -0.45 | 0.18 | -0.44 | 0.21 | 0.20 | 0.13 | 0.116 |
| USP16 | S552 | -0.28 | -0.27 | -0.10 | 0.11 | 0.19 | 0.25 | 0.005 |
| USP16 | S415 | 0.28 | 0.11 | 0.09 | -0.17 | -0.12 | -0.27 | 0.009 |
| USP16 | S415 | 0.25 | 0.14 | 0.10 | -0.18 | -0.08 | -0.31 | 0.012 |
| USP20 | S854 | 0.07 | -0.11 | -0.14 | 0.09 | -0.08 | 0.15 | 0.286 |
| USP20 | S112 | -0.04 | 0.09 | 0.06 | -0.28 | -0.03 | 0.15 | 0.546 |
| USP20 | S132 | 0.07 | 0.09 | -0.03 | -0.07 | -0.18 | 0.10 | 0.366 |
| USP20 | S112 | 0.04 | 0.25 | 0.34 | -0.40 | -0.07 | -0.31 | 0.023 |
| USP20 | T377; S | -0.38 | 0.21 | 0.77 | -0.73 | -0.14 | -0.23 | 0.209 |
| USP24 | S1943 | 0.01 | -0.23 | -0.02 | 0.13 | 0.11 | -0.03 | 0.175 |
| USP24 | S2047 | 0.01 | -0.07 | 0.07 | -0.12 | 0.21 | -0.13 | 0.893 |
| USP28 | S1053 | -0.08 | -0.40 | -0.26 | -0.02 | -0.04 | 0.59 | 0.134 |
| USP30 | S477; T480 | -0.30 | -0.21 | 0.07 | -0.06 | 0.15 | 0.27 | 0.142 |
| USP30 | S477; T480 | -0.30 | -0.21 | 0.07 | -0.06 | 0.15 | 0.27 | 0.142 |
| USP34 | S3406 | -0.13 | -0.47 | -0.07 | 0.10 | 0.22 | 0.24 | 0.035 |
| USP34 | S2488 | -0.38 | -0.09 | -0.15 | 0.07 | 0.14 | 0.30 | 0.027 |
| USP34 | T/S | -0.21 | -0.06 | 0.18 | -0.46 | -0.01 | 0.40 | 0.982 |
| USP36 | T653 | -0.02 | -0.11 | -0.34 | 0.08 | 0.07 | 0.25 | 0.062 |
| USP36 | S713 | -0.02 | -0.48 | 0.02 | 0.22 | 0.22 | -0.06 | 0.195 |
| USP36 | S18 | -0.15 | -0.11 | 0.01 | 0.11 | -0.04 | 0.15 | 0.099 |
| USP36 | S742 | -0.02 | -0.10 | -0.06 | 0.01 | -0.12 | 0.25 | 0.382 |
| USP36 | S682 | -0.08 | -0.07 | 0.02 | 0.11 | -0.12 | 0.12 | 0.394 |
| USP36 | S682 | -0.13 | -0.13 | 0.15 | 0.14 | -0.16 | 0.10 | 0.66 |
| USP36 | S546 | 0.05 | -0.04 | -0.11 | 0.23 | -0.02 | -0.14 | 0.641 |
| USP36 | S952 | -0.02 | 0.01 | -0.06 | 0.09 | -0.16 | 0.13 | 0.664 |
| USP36 | S682 | -0.09 | -0.01 | 0.04 | 0.00 | -0.19 | 0.21 | 0.829 |
| USP36 | T581 | 0.05 | 0.08 | -0.11 | -0.04 | -0.02 | 0.03 | 0.816 |
| USP36 | S614 | 0.19 | 0.03 | -0.06 | 0.13 | -0.30 | -0.05 | 0.429 |
| USP36 | S871 | -0.02 | 0.32 | 0.06 | 0.08 | -0.39 | -0.15 | 0.183 |
| USP39 | S82 | 0.22 | 0.31 | 0.18 | -0.13 | -0.12 | -0.68 | 0.045 |
| USP42 | S754; S759 | -0.42 | -0.54 | -0.20 | 0.19 | 0.16 | 0.52 | 0.012 |
| USP42 | S754 | -0.19 | -0.05 | -0.14 | 0.09 | -0.06 | 0.29 | 0.094 |
| USP42 | S/T | 0.01 | -0.13 | -0.20 | 0.46 | -0.10 | -0.15 | 0.428 |
| USP42 | S919 | -0.69 | 0.61 | -0.27 | 0.13 | 0.04 | -0.15 | 0.774 |
| USP42 | S754 | -0.25 | -0.09 | 0.19 | 0.35 | -0.17 | -0.12 | 0.753 |
| USP42 | S919 | 0.32 | -0.10 | -0.36 | -0.42 | 0.32 | 0.06 | 0.922 |
| USP42 | S936 | 0.10 | -0.03 | -0.05 | 0.04 | -0.13 | 0.05 | 0.811 |
| USP42 | S754 | 0.13 | -0.09 | 0.05 | -0.21 | -0.07 | 0.15 | 0.587 |
| USP42 | T/S | 0.16 | -0.07 | 0.02 | 0.01 | 0.01 | -0.13 | 0.419 |

| | | | | | | | | |
|---|---|---|---|---|---|---|---|---|
| USP42 | S856 | -0.01 | 0.08 | 0.06 | -0.02 | -0.05 | -0.06 | 0.037 |
| USP42 | S1166 | 0.13 | 0.02 | 0.07 | -0.11 | -0.13 | 0.00 | 0.036 |
| USP42 | S75 | 0.50 | 0.16 | -0.61 | 0.32 | -0.10 | -0.68 | 0.719 |
| USP42 | S1222; S1226 | -0.01 | 0.18 | 0.16 | -0.03 | -0.16 | -0.19 | 0.041 |
| USP42 | S1181 | 0.14 | 0.20 | 0.04 | 0.03 | -0.13 | -0.35 | 0.084 |
| USP42 | S1181 | 0.22 | 0.11 | 0.24 | -0.29 | -0.12 | -0.26 | 0.003 |
| USP42 | S432 | 0.24 | 0.66 | 0.10 | -0.47 | -0.13 | -0.89 | 0.04 |
| USP42 | S615 | 0.42 | 0.47 | 0.52 | -0.44 | -0.91 | -0.82 | 0.001 |
| USP5 | T623 | 0.13 | -0.05 | -0.17 | 0.23 | -0.34 | 0.13 | 0.845 |
| USP6NL | S435 | -0.21 | -0.26 | -0.20 | 0.22 | 0.16 | 0.21 | 9E-05 |
| USP6NL | S546 | 0.06 | -0.24 | -0.11 | -0.05 | 0.11 | 0.19 | 0.179 |
| USP6NL | S396 | -0.28 | -0.18 | 0.21 | -0.10 | 0.21 | 0.06 | 0.458 |
| USP6NL | S716 | 0.03 | 0.19 | -0.27 | 0.10 | 0.03 | -0.13 | 0.92 |
| USP8 | S718 | 0.05 | -0.24 | -0.43 | 0.12 | 0.00 | 0.37 | 0.102 |
| USP8 | S718 | 0.07 | 0.02 | -0.13 | -0.09 | -0.12 | 0.22 | 0.899 |
| USP8 | S/Y/T | 0.09 | 0.01 | 0.25 | 0.40 | -0.53 | -0.45 | 0.367 |
| USP8 | T577 | 0.02 | -0.03 | 0.53 | -0.57 | 0.05 | -0.23 | 0.168 |
| USPL1 | S200 | -0.35 | 0.02 | 0.01 | -0.19 | 0.10 | 0.32 | 0.387 |
| UTP11 | T90 | -0.12 | -0.01 | -0.12 | 0.11 | -0.01 | 0.13 | 0.042 |
| UTP11 | S241 | 0.05 | -0.14 | -0.07 | 0.06 | -0.02 | 0.10 | 0.192 |
| UTP14A | S445 | -0.21 | -0.29 | -0.36 | 0.17 | 0.36 | 0.18 | 0.002 |
| UTP14A | S77 | -0.12 | -0.21 | -0.01 | 0.21 | 0.24 | -0.18 | 0.234 |
| UTP14A | S445 | -0.06 | -0.14 | -0.01 | 0.06 | 0.12 | 0.01 | 0.057 |
| UTP14A | S29; S31 | 0.13 | -0.07 | -0.19 | -0.11 | 0.06 | 0.14 | 0.586 |
| UTP14A | S405; S407 | 0.01 | -0.08 | -0.04 | -0.05 | 0.05 | 0.10 | 0.26 |
| UTP14A | S405; S407 | -0.15 | -0.14 | 0.21 | -0.10 | -0.08 | 0.20 | 0.834 |
| UTP14A | S407 | 0.10 | 0.09 | -0.11 | 0.00 | -0.17 | 0.06 | 0.56 |
| UTP14A | S405 | 0.02 | 0.16 | 0.06 | 0.03 | -0.11 | -0.20 | 0.084 |
| UTP18 | T204; S210; S | -0.29 | -0.10 | -0.12 | 0.25 | -0.04 | 0.22 | 0.046 |
| UTP18 | S124 | -0.08 | -0.11 | -0.12 | 0.33 | -0.19 | 0.11 | 0.277 |
| UTP18 | S206; S210 | -0.13 | -0.13 | -0.03 | 0.17 | -0.14 | 0.21 | 0.194 |
| UTP18 | S121; S124 | 0.08 | -0.01 | -0.07 | -0.08 | 0.03 | 0.04 | 0.96 |
| UTP18 | S/T | 0.18 | 0.36 | 0.22 | -0.37 | -0.18 | -0.40 | 0.003 |
| UTP18 | S210 | 0.32 | 0.42 | 0.40 | -0.50 | -0.18 | -0.96 | 0.015 |
| UTP20 | T1741 | -0.24 | -0.30 | -0.34 | 0.25 | 0.09 | 0.38 | 0.004 |
| UTP20 | T1741 | 0.17 | 0.14 | 0.09 | -0.06 | -0.10 | -0.29 | 0.018 |
| UTP20 | S2637 | 0.19 | 0.08 | 0.34 | -0.24 | -0.22 | -0.27 | 0.004 |
| UTP25 | T149 | -0.06 | -0.13 | 0.14 | -0.21 | 0.13 | 0.09 | 0.917 |
| UTP25 | S153 | -0.01 | -0.13 | 0.11 | -0.01 | -0.10 | 0.13 | 0.884 |
| UTP3 | S365; S368 | -0.15 | -0.29 | -0.02 | -0.07 | 0.17 | 0.28 | 0.101 |
| UTP3 | S37 | -0.05 | 0.02 | -0.05 | 0.09 | -0.06 | 0.05 | 0.365 |
| UTP3 | S37 | -0.09 | 0.01 | 0.00 | 0.16 | -0.13 | 0.03 | 0.635 |
| UTP3 | S37 | -0.06 | 0.02 | -0.02 | 0.05 | -0.04 | 0.06 | 0.357 |
| UTP3 | T/S/Y | -0.09 | 0.19 | -0.04 | 0.09 | -0.09 | -0.08 | 0.664 |
| UTP3 | S368 | 0.15 | 0.18 | 0.21 | -0.19 | -0.08 | -0.36 | 0.01 |
| UVSSA | S281; S287 | 0.01 | -0.31 | 0.04 | 0.12 | -0.14 | 0.21 | 0.385 |
| VAMP2 | S80 | -0.11 | 0.18 | 0.09 | -0.14 | -0.25 | 0.17 | 0.452 |
| VAMP2 | S75 | 0.27 | 0.23 | 0.19 | -0.33 | -0.34 | -0.16 | 0.001 |

| | | | | | | | | |
|---|---|---|---|---|---|---|---|---|
| **VAMP4** | **S30** | -0.03 | -0.03 | 0.00 | 0.13 | 0.04 | -0.12 | 0.655 |
| **VAMP4** | **S30** | 0.09 | 0.06 | 0.02 | 0.18 | -0.06 | -0.35 | 0.423 |
| **VANGL1** | **T80; S86; S88** | -0.03 | -0.09 | 0.17 | -0.15 | -0.16 | 0.22 | 0.769 |
| **VANGL1** | **S/T** | 0.16 | 0.09 | -0.13 | 0.24 | 0.01 | -0.48 | 0.632 |
| **VANGL1** | **S/T** | 0.16 | 0.09 | -0.13 | 0.24 | 0.01 | -0.48 | 0.632 |
| **VAPA** | **S/T** | -0.27 | 0.03 | -0.51 | 0.56 | -0.14 | 0.08 | 0.186 |
| **VAPB** | **S156; S160** | -0.07 | -0.04 | -0.05 | 0.07 | -0.09 | 0.17 | 0.26 |
| **VAPB** | **S156** | 0.00 | 0.04 | 0.02 | -0.04 | -0.04 | 0.01 | 0.124 |
| **VAPB** | **T150** | 0.02 | 0.04 | 0.16 | -0.31 | 0.15 | -0.11 | 0.29 |
| **VAPB** | **S158** | 0.17 | 0.13 | -0.01 | -0.10 | -0.08 | -0.15 | 0.028 |
| **VCP** | **S197** | -0.29 | -0.29 | -0.25 | 0.08 | -0.24 | 0.70 | 0.171 |
| **VCPIP1** | **S768** | -0.09 | 0.01 | -0.08 | 0.20 | -0.21 | 0.14 | 0.5 |
| **VCPIP1** | **S747** | 0.17 | 0.09 | -0.04 | -0.06 | -0.01 | -0.17 | 0.116 |
| **VDAC1** | **T70** | -0.27 | -0.90 | 0.04 | 0.05 | 0.13 | 0.56 | 0.119 |
| **VDAC1** | **S104** | -0.15 | 0.19 | -0.28 | 0.10 | 0.19 | -0.12 | 0.466 |
| **VDAC1** | **S104** | 0.08 | -0.01 | 0.05 | -0.07 | 0.10 | -0.17 | 0.333 |
| **VDAC1** | **T/S** | -0.06 | 0.06 | 0.17 | 0.10 | -0.10 | -0.19 | 0.35 |
| **VDAC1** | **T/S** | 0.21 | 0.01 | 0.11 | -0.01 | -0.08 | -0.29 | 0.082 |
| **VDAC1** | **S57** | 0.19 | 0.17 | 0.19 | -0.12 | -0.05 | -0.50 | 0.044 |
| **VDAC1** | **T/S** | 0.21 | 0.15 | 0.18 | -0.07 | 0.00 | -0.63 | 0.111 |
| **VDAC2** | **S115** | 0.00 | -0.13 | -0.10 | 0.05 | 0.08 | 0.09 | 0.027 |
| **VDAC2** | **S48** | 0.10 | -0.14 | -0.15 | 0.01 | -0.10 | 0.24 | 0.443 |
| **VDAC2** | **S115** | 0.01 | 0.04 | -0.02 | 0.05 | 0.14 | -0.24 | 0.856 |
| **VDAC2** | **T68** | 0.13 | 0.21 | 0.18 | -0.28 | 0.00 | -0.34 | 0.022 |
| **VDAC2** | **S46** | 0.19 | 0.26 | 0.09 | -0.20 | -0.05 | -0.38 | 0.022 |
| **VDAC2** | **S46** | 0.14 | 0.27 | 0.19 | -0.13 | -0.18 | -0.39 | 0.008 |
| **VDAC3** | **S55** | 0.04 | -0.08 | 0.03 | 0.24 | -0.23 | -0.04 | 0.955 |
| **VEZF1** | **S197** | -0.16 | -0.34 | -0.22 | 0.20 | 0.19 | 0.23 | 0.001 |
| **VEZT** | **S696** | 0.08 | -0.37 | -0.05 | 0.13 | 0.07 | 0.09 | 0.199 |
| **VEZT** | **S696** | 0.23 | -0.18 | 0.04 | 0.11 | 0.27 | -0.68 | 0.699 |
| **VIRMA** | **T184** | 0.07 | -0.01 | -0.03 | 0.11 | -0.27 | 0.10 | 0.807 |
| **VIRMA** | **S222** | 0.05 | 0.02 | 0.10 | 0.02 | -0.13 | -0.07 | 0.071 |
| **VIRMA** | **S** | 0.00 | 0.29 | -0.09 | 0.08 | 0.00 | -0.37 | 0.411 |
| **VIRMA** | **S173** | 0.19 | 0.13 | 0.21 | -0.10 | -0.22 | -0.29 | 0.003 |
| **VIRMA** | **S173** | -0.05 | 0.27 | 0.35 | -0.18 | -0.04 | -0.54 | 0.083 |
| **VIRMA** | **S1579** | 0.21 | 0.19 | 0.22 | -0.19 | -0.08 | -0.47 | 0.018 |
| **VIRMA** | **T1708** | 0.23 | 0.12 | 0.32 | -0.17 | -0.39 | -0.24 | 0.005 |
| **VIRMA** | **S173** | 0.18 | 0.21 | 0.27 | -0.18 | -0.09 | -0.55 | 0.026 |
| **VIRMA** | **S173** | 0.25 | 0.39 | 0.34 | -0.30 | -0.14 | -0.96 | 0.036 |
| **VPS13D** | **S1138** | 0.02 | -0.30 | -0.50 | -0.03 | 0.54 | 0.04 | 0.137 |
| **VPS13D** | **S2692** | 0.05 | -0.02 | -0.32 | -0.07 | 0.01 | 0.29 | 0.327 |
| **VPS13D** | **S1765** | -0.19 | -0.17 | 0.09 | 0.33 | -0.22 | 0.08 | 0.453 |
| **VPS13D** | **S/T/Y** | 0.06 | 0.23 | -0.58 | 0.26 | -0.03 | -0.09 | 0.628 |
| **VPS13D** | **S1138** | -0.12 | 0.14 | -0.04 | 0.20 | 0.16 | -0.43 | 0.934 |
| **VPS13D** | **S2079** | 0.43 | -0.50 | 0.28 | -0.15 | 0.32 | -0.78 | 0.557 |
| **VPS13D** | **S1709** | 0.40 | 0.28 | 0.23 | -0.42 | -0.75 | -0.07 | 0.024 |
| **VPS50** | **S559** | -0.22 | -0.23 | -0.24 | 0.31 | -0.15 | 0.38 | 0.071 |
| **VPS50** | **S494** | 0.18 | -0.09 | -0.11 | -0.02 | -0.02 | 0.05 | 0.926 |

| | | | | | | | | |
|---|---|---|---|---|---|---|---|---|
| **VPS50** | **S561** | -0.05 | 0.07 | 0.13 | -0.03 | -0.26 | 0.09 | 0.372 |
| **VPS53** | **S377** | -0.28 | -0.05 | -0.30 | 0.18 | 0.20 | 0.16 | 0.009 |
| **VPS53** | **S377** | -0.21 | -0.03 | -0.27 | 0.10 | -0.08 | 0.38 | 0.117 |
| **VPS72** | **S127** | 0.03 | -0.24 | 0.04 | -0.10 | -0.10 | 0.31 | 0.608 |
| **VPS72** | **S127** | -0.06 | -0.04 | 0.09 | -0.02 | -0.36 | 0.31 | 0.936 |
| **VRK1** | **S376** | -0.37 | -0.04 | 0.00 | -0.10 | 0.17 | 0.26 | 0.201 |
| **VRK1** | **S376** | -0.28 | 0.00 | 0.20 | 0.16 | -0.26 | 0.11 | 0.889 |
| **VRK1** | **S342** | -0.11 | 0.33 | 0.02 | 0.27 | -0.26 | -0.40 | 0.43 |
| **VRK2** | **T336** | -0.23 | -0.19 | 0.08 | 0.04 | -0.08 | 0.32 | 0.255 |
| **VRK3** | **S83** | 0.07 | 0.10 | 0.01 | -0.08 | -0.13 | 0.01 | 0.064 |
| **VRK3** | **S/T** | 0.17 | 0.07 | 0.00 | -0.10 | -0.18 | 0.02 | 0.092 |
| **VRK3** | **T/S** | 0.22 | 0.18 | 0.10 | -0.05 | -0.37 | -0.17 | 0.023 |
| **VRK3** | **S71** | 0.31 | 0.16 | 0.14 | -0.39 | -0.24 | -0.10 | 0.011 |
| **VRK3** | **S59** | 0.30 | 0.35 | 0.08 | -0.56 | 0.04 | -0.45 | 0.047 |
| **VTI1B** | **T103** | -0.10 | 0.05 | 0.06 | -0.05 | 0.18 | -0.15 | 0.908 |
| **WAC** | **S53** | 0.10 | -0.02 | -0.10 | 0.05 | 0.00 | -0.03 | 0.824 |
| **WAPL** | **S221** | -0.04 | -0.08 | -0.28 | -0.01 | 0.04 | 0.31 | 0.123 |
| **WAPL** | **S221; S226** | -0.14 | 0.03 | 0.03 | 0.04 | -0.07 | 0.10 | 0.535 |
| **WAPL** | **S459** | -0.01 | 0.04 | 0.01 | 0.05 | -0.21 | 0.10 | 0.727 |
| **WAPL** | **S/Y** | -0.01 | 0.03 | 0.04 | 0.06 | -0.13 | 0.01 | 0.537 |
| **WAPL** | **S77** | 0.06 | -0.06 | 0.06 | -0.21 | 0.12 | 0.01 | 0.66 |
| **WASF2** | **S497; S/Y** | -0.06 | 0.15 | -0.14 | -0.13 | -0.01 | 0.17 | 0.822 |
| **WASHC2A** | **T/S** | 0.20 | -0.12 | -0.04 | 0.04 | 0.15 | -0.30 | 0.782 |
| **WASHC2A** | **T/S** | 0.21 | 0.19 | -0.10 | -0.10 | -0.05 | -0.21 | 0.115 |
| **WBP1** | **T209** | -0.13 | 0.18 | 0.20 | -0.36 | 0.15 | -0.13 | 0.336 |
| **WBP11** | **S361; S364** | -0.10 | -0.15 | -0.33 | 0.45 | -0.11 | 0.12 | 0.12 |
| **WBP11** | **S364; S** | -0.28 | -0.06 | -0.01 | -0.23 | 0.73 | -0.50 | 0.782 |
| **WBP11** | **S237** | -0.07 | -0.04 | 0.01 | -0.01 | 0.02 | 0.08 | 0.143 |
| **WBP11** | **S237** | -0.03 | -0.21 | 0.13 | 0.11 | -0.04 | 0.00 | 0.618 |
| **WBP11** | **S237** | -0.09 | -0.10 | 0.11 | 0.00 | -0.03 | 0.11 | 0.54 |
| **WBP11** | **S237** | 0.01 | 0.01 | -0.08 | -0.05 | -0.04 | 0.14 | 0.605 |
| **WBP11** | **S237** | -0.02 | 0.01 | -0.01 | -0.04 | -0.05 | 0.10 | 0.824 |
| **WBP11** | **S237** | -0.07 | 0.07 | 0.04 | -0.05 | -0.14 | 0.13 | 0.707 |
| **WBP11** | **S364** | 0.01 | 0.22 | -0.20 | -0.26 | 0.23 | -0.06 | 0.842 |
| **WBP11** | **S237** | 0.03 | 0.04 | 0.08 | 0.01 | -0.05 | -0.12 | 0.055 |
| **WBP11** | **S361** | 0.30 | 0.10 | 0.19 | 0.16 | -0.37 | -0.59 | 0.112 |
| **WDR12** | **T221** | -0.39 | -0.22 | -0.57 | 0.32 | -0.75 | 0.89 | 0.326 |
| **WDR13** | **S70** | -0.05 | 0.20 | -0.14 | 0.00 | 0.22 | -0.30 | 0.861 |
| **WDR20** | **S/T/Y** | -0.08 | -0.15 | -0.22 | 0.11 | 0.03 | 0.25 | 0.021 |
| **WDR20** | **S434** | 0.08 | 0.11 | 0.09 | 0.02 | -0.19 | -0.13 | 0.041 |
| **WDR24** | **T581** | 0.06 | 0.15 | 0.24 | -0.11 | -0.10 | -0.30 | 0.018 |
| **WDR26** | **S121** | -0.17 | 0.03 | -0.17 | 0.14 | -0.05 | 0.19 | 0.114 |
| **WDR3** | **S241** | -0.20 | -0.22 | -0.17 | 0.18 | -0.07 | 0.38 | 0.052 |
| **WDR3** | **S241** | -0.22 | -0.14 | -0.11 | 0.14 | -0.06 | 0.32 | 0.066 |
| **WDR3** | **T257; S/T** | -0.16 | -0.26 | 0.03 | 0.06 | -0.07 | 0.33 | 0.179 |
| **WDR3** | **S726** | 0.28 | -0.05 | -0.10 | -0.03 | -0.01 | -0.13 | 0.49 |
| **WDR3** | **S241** | 0.19 | -0.02 | 0.08 | 0.18 | -0.02 | -0.53 | 0.404 |
| **WDR3** | **S241** | 0.02 | 0.20 | 0.29 | -0.32 | 0.00 | -0.29 | 0.045 |

| | | | | | | | | |
|---|---|---|---|---|---|---|---|---|
| WDR33 | S1278 | 0.01 | -0.06 | -0.04 | -0.12 | 0.08 | 0.13 | 0.502 |
| WDR33 | S1210 | 0.11 | -0.05 | -0.05 | 0.14 | -0.19 | 0.01 | 0.879 |
| WDR4 | S391 | -0.13 | 0.00 | 0.05 | -0.04 | 0.17 | -0.07 | 0.652 |
| WDR43 | S431 | 0.04 | -0.22 | -0.33 | -0.11 | 0.56 | -0.12 | 0.325 |
| WDR43 | S77 | 0.26 | 0.22 | 0.18 | -0.26 | 0.07 | -0.71 | 0.086 |
| WDR44 | S561 | 0.39 | -0.24 | 0.19 | -0.24 | 0.09 | -0.34 | 0.29 |
| WDR46 | S41 | -0.11 | 0.05 | -0.06 | 0.28 | 0.09 | -0.33 | 0.793 |
| WDR46 | S41 | -0.12 | -0.04 | 0.38 | -0.06 | -0.03 | -0.21 | 0.358 |
| WDR55 | T31 | -0.11 | -0.56 | -0.31 | 0.24 | 0.29 | 0.25 | 0.011 |
| WDR62 | S33 | 0.10 | 0.04 | -0.32 | -0.18 | -0.06 | 0.33 | 0.678 |
| WDR7 | S935 | 0.24 | 0.14 | 0.15 | -1.64 | -0.09 | 0.43 | 0.384 |
| WDR70 | S638 | -0.09 | -0.03 | -0.08 | -0.05 | 0.09 | 0.15 | 0.11 |
| WDR75 | S779; S782 | 0.03 | -0.09 | -0.18 | 0.08 | 0.09 | 0.05 | 0.07 |
| WDR75 | S779; S782 | 0.04 | 0.00 | -0.13 | 0.06 | 0.01 | 0.01 | 0.327 |
| WDR75 | S779; S782 | 0.04 | -0.01 | -0.11 | 0.03 | 0.06 | -0.02 | 0.365 |
| WDR75 | S779; S782 | 0.12 | -0.07 | -0.09 | -0.05 | 0.06 | 0.02 | 0.798 |
| WDR75 | S/T | 0.18 | 0.27 | -0.28 | 0.06 | -0.01 | -0.31 | 0.531 |
| WDR75 | S796 | 0.19 | -0.16 | 0.26 | 0.02 | -0.13 | -0.24 | 0.23 |
| WDR75 | S782 | 0.04 | 0.25 | 0.01 | -0.04 | -0.10 | -0.20 | 0.07 |
| WDR75 | S/T | 0.26 | 0.35 | 0.06 | -0.13 | -0.03 | -0.77 | 0.095 |
| WDR75 | S782 | 0.30 | 0.28 | 0.17 | -0.20 | -0.05 | -0.75 | 0.053 |
| WDR77 | T5 | -0.08 | 0.02 | -0.05 | 0.10 | 0.17 | -0.19 | 0.606 |
| WDTC1 | S353 | -0.13 | -0.10 | 0.11 | 0.15 | -0.08 | 0.03 | 0.486 |
| WDTC1 | S353 | 0.06 | -0.12 | 0.06 | 0.07 | -0.20 | 0.10 | 0.934 |
| WEE1 | S165; T173 | -0.38 | -0.24 | -0.18 | 0.15 | 0.06 | 0.43 | 0.019 |
| WEE1 | S165 | -0.30 | -0.15 | 0.19 | 0.35 | -0.06 | -0.13 | 0.548 |
| WFS1 | S32 | -0.14 | 0.08 | 0.12 | -0.13 | 0.10 | -0.05 | 0.679 |
| WFS1 | S32 | 0.15 | -0.02 | 0.16 | -0.10 | 0.13 | -0.38 | 0.247 |
| WIPF2 | S267 | -0.06 | 0.33 | -0.83 | 0.47 | -0.27 | 0.01 | 0.558 |
| WIPI2 | S413 | 0.00 | 0.06 | -0.31 | 0.23 | -0.07 | 0.05 | 0.33 |
| WIZ | S1517 | -0.29 | -0.50 | -0.27 | 0.27 | 0.07 | 0.48 | 0.011 |
| WIZ | S1127; S1134 | -0.16 | -0.33 | -0.14 | 0.15 | 0.08 | 0.31 | 0.013 |
| WIZ | S1012; S1017 | 0.04 | -0.03 | -0.08 | 0.00 | 0.10 | -0.03 | 0.441 |
| WIZ | S983 | 0.09 | 0.04 | -0.02 | -0.05 | -0.05 | -0.02 | 0.067 |
| WIZ | S1146 | 0.03 | 0.07 | 0.05 | 0.33 | -0.36 | -0.22 | 0.558 |
| WIZ | S996 | 0.09 | 0.18 | 0.10 | -0.25 | 0.04 | -0.21 | 0.048 |
| WIZ | S1134 | 0.27 | -0.09 | 0.48 | -0.87 | -0.04 | -0.10 | 0.151 |
| WNK1 | S | 0.18 | 0.29 | -0.15 | -0.11 | 0.11 | -0.45 | 0.289 |
| WNK1 | S185 | 0.27 | 0.21 | 0.19 | -0.30 | -0.02 | -0.53 | 0.027 |
| WRAP53 | S85; S90 | -0.26 | -0.24 | -0.11 | 0.09 | -0.12 | 0.49 | 0.129 |
| WRAP53 | S26; S30 | 0.07 | -0.34 | -0.12 | 0.35 | -0.02 | -0.03 | 0.258 |
| WRAP53 | T102; S103 | -0.05 | -0.04 | 0.02 | 0.01 | -0.08 | 0.13 | 0.575 |
| WRAP53 | S54 | -0.03 | 0.04 | 0.03 | 0.02 | 0.09 | -0.17 | 0.681 |
| WRAP53 | S26; S30 | 0.37 | -0.17 | 0.00 | 0.08 | -0.17 | -0.20 | 0.423 |
| WRAP53 | S90 | 0.06 | 0.09 | 0.09 | 0.10 | -0.11 | -0.27 | 0.193 |
| WRAP53 | S/T | 0.31 | -0.63 | 0.49 | 0.00 | -0.23 | -0.22 | 0.592 |
| WRAP53 | S491 | 0.15 | 0.19 | -0.03 | -0.11 | 0.02 | -0.27 | 0.103 |
| WRAP53 | S/T | 0.37 | -0.01 | 0.37 | -0.25 | -0.20 | -0.50 | 0.023 |

| | | | | | | | | |
|---|---|---|---|---|---|---|---|---|
| **WRAP53** | **S114** | 0.39 | 0.12 | 0.53 | -0.54 | -0.34 | -0.55 | 0.004 |
| **WRN** | **S/T** | -0.39 | 0.01 | 0.15 | 0.18 | -0.18 | 0.14 | 0.572 |
| **WRN** | **S426; S440** | -0.10 | -0.07 | 0.17 | 0.14 | -0.18 | 0.00 | 0.941 |
| **WRN** | **S426; S440** | -0.10 | -0.07 | 0.17 | 0.14 | -0.18 | 0.00 | 0.941 |
| **WRN** | **S1133** | 0.08 | 0.08 | 0.01 | 0.09 | -0.33 | 0.03 | 0.409 |
| **WRNIP1** | **S65** | -0.33 | -0.43 | -0.42 | 0.43 | 0.07 | 0.41 | 0.005 |
| **WRNIP1** | **S65** | -0.02 | -0.36 | -0.58 | -0.07 | 0.47 | 0.30 | 0.074 |
| **WRNIP1** | **S153** | 0.01 | -0.36 | -0.11 | 0.33 | 0.07 | -0.04 | 0.155 |
| **WRNIP1** | **S75** | 0.09 | -0.03 | -0.11 | 0.11 | 0.10 | -0.18 | 0.816 |
| **WRNIP1** | **S470** | 0.09 | -0.01 | -0.07 | 0.04 | 0.10 | -0.16 | 0.883 |
| **WTAP** | **S306** | 0.20 | 0.20 | 0.15 | -0.09 | -0.09 | -0.48 | 0.035 |
| **WTAP** | **S306** | 0.31 | 0.19 | 0.36 | -0.27 | -0.20 | -0.65 | 0.011 |
| **WWC3** | **S479** | -0.25 | -0.15 | -0.43 | 0.29 | -0.23 | 0.52 | 0.118 |
| **WWC3** | **S927** | -0.09 | -0.09 | 0.08 | -0.08 | 0.07 | 0.09 | 0.487 |
| **WWC3** | **S911** | 0.04 | -0.05 | 0.05 | -0.15 | 0.27 | -0.21 | 0.784 |
| **WWOX** | **T12; S14** | -0.10 | 0.02 | -0.13 | -0.02 | 0.01 | 0.19 | 0.184 |
| **WWOX** | **S14** | -0.31 | 0.21 | 0.26 | -0.08 | -0.24 | 0.07 | 0.54 |
| **WWP2** | **S211** | 0.13 | 0.16 | 0.43 | -0.27 | -0.12 | -0.53 | 0.024 |
| **XAB2** | **S851** | -0.83 | -1.40 | -1.21 | 0.79 | 0.61 | 0.46 | 8E-04 |
| **XBP1** | **S47** | -0.23 | -0.34 | -0.10 | 0.15 | 0.12 | 0.30 | 0.009 |
| **XBP1** | **S68** | 0.07 | -0.17 | 0.07 | -0.10 | 0.11 | 0.00 | 0.874 |
| **XPA** | **S196** | -0.21 | 0.01 | -0.34 | 0.08 | 0.07 | 0.30 | 0.061 |
| **XPC** | **S496** | -0.21 | -0.42 | 0.06 | 0.37 | -0.13 | 0.19 | 0.173 |
| **XPC** | **S122** | 0.15 | -0.35 | -0.34 | 0.21 | 0.03 | 0.19 | 0.135 |
| **XPC** | **T/S** | -0.26 | -0.05 | -0.02 | -0.03 | 0.21 | 0.10 | 0.111 |
| **XPC** | **S129; S140** | -0.14 | 0.03 | -0.13 | -0.20 | 0.19 | 0.20 | 0.375 |
| **XPC** | **S122; S129** | -0.14 | 0.03 | -0.13 | -0.20 | 0.19 | 0.20 | 0.375 |
| **XPC** | **S94** | -0.01 | -0.04 | -0.13 | 0.01 | 0.00 | 0.16 | 0.131 |
| **XPC** | **S351** | -0.02 | -0.16 | 0.00 | -0.05 | 0.11 | 0.10 | 0.2 |
| **XPC** | **S129** | 0.09 | -0.10 | -0.15 | 0.08 | -0.11 | 0.16 | 0.436 |
| **XPC** | **S883; S884** | -0.03 | -0.11 | 0.15 | -0.20 | 0.13 | 0.03 | 0.926 |
| **XPC** | **S398; S399** | 0.15 | -0.02 | -0.08 | -0.17 | 0.01 | 0.09 | 0.728 |
| **XPC** | **S397; S398; S399** | 0.02 | 0.00 | 0.03 | -0.18 | -0.04 | 0.14 | 0.67 |
| **XPC** | **S122; S129** | 0.02 | 0.14 | 0.05 | 0.01 | -0.08 | -0.16 | 0.074 |
| **XPC** | **S397; S398; S399** | 0.18 | 0.11 | 0.03 | -0.17 | 0.07 | -0.28 | 0.1 |
| **XPC** | **S** | 0.12 | 0.08 | 0.17 | -0.21 | -0.22 | 0.01 | 0.029 |
| **XPC** | **S883** | 0.20 | 0.20 | 0.29 | -0.49 | -0.03 | -0.34 | 0.02 |
| **XPO4** | **S521** | 0.09 | 0.44 | 0.37 | -1.78 | -0.12 | 0.11 | 0.213 |
| **XPR1** | **T690** | 0.06 | -0.06 | 0.03 | -0.07 | -0.07 | 0.11 | 0.756 |
| **XRCC1** | **S461; S485; T488** | -0.09 | -0.28 | -0.07 | 0.15 | -0.01 | 0.24 | 0.048 |
| **XRCC1** | **S461** | -0.10 | -0.13 | -0.20 | 0.20 | -0.03 | 0.21 | 0.03 |
| **XRCC1** | **S/T** | -0.17 | -0.09 | -0.17 | 0.02 | 0.02 | 0.33 | 0.07 |
| **XRCC1** | **S475** | -0.50 | -0.09 | 0.14 | 0.21 | 0.03 | 0.09 | 0.249 |
| **XRCC1** | **S461** | -0.09 | -0.17 | -0.11 | 0.10 | -0.01 | 0.24 | 0.039 |
| **XRCC1** | **S461; S485; T488** | -0.20 | -0.15 | -0.01 | 0.05 | 0.01 | 0.26 | 0.081 |
| **XRCC1** | **S266** | -0.20 | 0.08 | -0.20 | -0.08 | 0.10 | 0.25 | 0.223 |
| **XRCC1** | **S461; S475; S485** | -0.12 | -0.17 | 0.00 | 0.35 | -0.21 | 0.08 | 0.369 |
| **XRCC1** | **S461; S475** | -0.02 | -0.05 | -0.15 | 0.12 | -0.11 | 0.19 | 0.229 |

| | | | | | | | | |
|---|---|---|---|---|---|---|---|---|
| XRCC1 | S475; S485; T488 | -0.04 | -0.08 | -0.07 | 0.21 | -0.18 | 0.12 | 0.38 |
| XRCC1 | S475; S485; T488 | -0.01 | -0.03 | -0.13 | 0.12 | -0.09 | 0.12 | 0.227 |
| XRCC1 | S241 | 0.07 | -0.04 | -0.09 | 0.11 | 0.06 | -0.12 | 0.673 |
| XRCC1 | S485; T488 | -0.05 | -0.01 | 0.01 | 0.07 | -0.01 | -0.01 | 0.36 |
| XRCC1 | S266 | -0.04 | -0.01 | 0.00 | 0.05 | -0.13 | 0.12 | 0.716 |
| XRCC1 | S461; S475 | 0.00 | 0.00 | -0.03 | 0.00 | 0.02 | 0.01 | 0.232 |
| XRCC1 | S461 | 0.11 | -0.02 | -0.10 | 0.04 | -0.01 | -0.03 | 0.978 |
| XRCC1 | S266 | 0.12 | 0.01 | -0.13 | -0.02 | 0.06 | -0.05 | 0.956 |
| XRCC1 | S461; S475 | 0.03 | -0.15 | 0.14 | 0.25 | -0.14 | -0.16 | 0.89 |
| XRCC1 | S461 | -0.08 | 0.09 | 0.05 | 0.01 | -0.02 | -0.07 | 0.442 |
| XRCC1 | S446; T453 | 0.01 | 0.06 | 0.00 | -0.09 | 0.11 | -0.09 | 0.51 |
| XRCC1 | S461; S475 | 0.07 | 0.10 | -0.03 | -0.06 | -0.12 | 0.03 | 0.183 |
| XRCC1 | S447; T453 | 0.03 | 0.10 | 0.03 | -0.06 | 0.11 | -0.23 | 0.319 |
| XRCC1 | S485; T488 | 0.25 | 0.02 | -0.10 | -0.43 | -0.24 | 0.35 | 0.559 |
| XRCC1 | S461; S485; T488 | 0.00 | -0.29 | 0.52 | -0.32 | 0.01 | -0.09 | 0.45 |
| XRCC1 | S461; S475 | 0.07 | 0.05 | 0.12 | 0.36 | -0.07 | -0.75 | 0.51 |
| XRCC1 | S/T | 0.27 | 0.27 | 0.15 | -0.32 | 0.05 | -0.64 | 0.059 |
| XRCC4 | S/T | 0.15 | -0.02 | 0.10 | -0.12 | -0.08 | -0.04 | 0.05 |
| XRCC5 | T472 | -1.26 | -0.95 | -1.24 | 0.59 | 0.65 | 0.65 | 6E-05 |
| XRCC6 | S560 | -2.21 | -2.14 | -1.90 | 0.87 | 0.86 | 0.72 | 1E-05 |
| XRCC6 | S520 | -2.09 | -1.78 | -2.25 | 0.82 | 1.00 | 0.58 | 1E-04 |
| XRCC6 | S477 | -0.87 | -0.57 | -0.40 | 0.45 | 0.12 | 0.65 | 0.008 |
| XRCC6 | S222 | -0.41 | -0.28 | -0.30 | 0.33 | -0.03 | 0.46 | 0.018 |
| XRCC6 | S27 | -0.17 | -0.23 | 0.06 | -0.08 | 0.30 | 0.06 | 0.22 |
| XRCC6 | S27 | -0.28 | -0.15 | 0.17 | 0.00 | 0.17 | 0.04 | 0.341 |
| XRCC6 | T455 | 0.12 | 0.04 | 0.01 | -0.08 | 0.11 | -0.23 | 0.289 |
| XRN2 | S678 | -0.41 | -0.08 | -0.05 | 0.27 | 0.20 | -0.03 | 0.088 |
| XRN2 | S499; S501 | 0.12 | 0.05 | 0.18 | -0.06 | -0.10 | -0.22 | 0.017 |
| XRN2 | S487; S/T | 0.06 | 0.46 | 0.10 | -0.18 | -0.48 | -0.14 | 0.049 |
| XRN2 | S448 | 0.18 | 0.07 | 0.42 | -0.27 | -0.05 | -0.57 | 0.046 |
| XRN2 | S470; S475 | -0.11 | 0.70 | 0.20 | -0.70 | -0.22 | -0.29 | 0.075 |
| YAP1 | S109 | -0.08 | -0.05 | -0.12 | 0.24 | 0.09 | -0.12 | 0.208 |
| YAP1 | S164 | 0.09 | 0.15 | 0.14 | -0.08 | -0.07 | -0.28 | 0.022 |
| YBX1 | S165 | -0.03 | -0.05 | -0.02 | 0.11 | -0.20 | 0.17 | 0.622 |
| YBX1 | S174 | 0.08 | 0.16 | -0.19 | 0.03 | 0.08 | -0.21 | 0.725 |
| YBX1 | S165 | -0.03 | 0.03 | 0.09 | -0.07 | -0.10 | 0.07 | 0.358 |
| YBX1 | S165 | 0.06 | 0.09 | 0.24 | -0.21 | -0.11 | -0.11 | 0.014 |
| YBX1 | S165; S174 | 0.04 | 0.22 | 0.23 | -0.11 | -0.15 | -0.30 | 0.015 |
| YBX1 | S174 | 0.33 | 0.08 | 0.15 | -0.22 | -0.07 | -0.38 | 0.023 |
| YBX1 | S/Y | 0.22 | 0.27 | 0.24 | -0.23 | -0.03 | -0.71 | 0.048 |
| YBX1 | S165; S167; S174 | 0.03 | 0.06 | 0.61 | -0.51 | -0.01 | -0.49 | 0.083 |
| YBX1 | S165; S174 | 0.16 | 0.24 | 0.36 | -0.39 | -0.07 | -0.50 | 0.015 |
| YBX1 | S165; S174 | 0.19 | 0.19 | 0.39 | -0.26 | -0.15 | -0.59 | 0.017 |
| YBX1 | S165 | 0.27 | 0.29 | 0.40 | -0.38 | -0.06 | -0.95 | 0.041 |
| YBX1 | S174 | 0.28 | 0.33 | 0.41 | -0.44 | -0.08 | -0.95 | 0.032 |
| YBX1 | S165; S167; S176 | 0.43 | 0.25 | 0.79 | -1.04 | -0.37 | -1.07 | 0.009 |
| YBX2 | T57 | -0.55 | 0.12 | 0.13 | 0.14 | 0.09 | -0.04 | 0.522 |
| YBX2 | T76 | -0.01 | 0.05 | 0.12 | -0.09 | -0.23 | 0.13 | 0.356 |

| | | | | | | | | |
|---|---|---|---|---|---|---|---|---|
| YEATS2 | S465 | 0.23 | -0.18 | 0.07 | 0.05 | -0.48 | 0.19 | 0.631 |
| YEATS2 | S/T | 0.03 | -0.04 | 0.14 | 0.39 | -0.46 | -0.21 | 0.631 |
| YEATS2 | S447 | 0.00 | 0.01 | 0.18 | 0.39 | -0.06 | -0.75 | 0.58 |
| YJU2 | S322 | 0.13 | 0.06 | -0.01 | -0.03 | -0.06 | -0.10 | 0.045 |
| YJU2B | S362; S367 | -0.23 | -0.44 | -0.21 | 0.27 | 0.27 | 0.19 | 0.002 |
| YJU2B | S332 | -0.06 | -0.06 | -0.06 | 0.09 | -0.15 | 0.21 | 0.341 |
| YJU2B | S332 | -0.03 | 0.03 | -0.05 | 0.05 | -0.05 | 0.05 | 0.453 |
| YJU2B | S362 | -0.02 | 0.26 | 0.22 | -0.06 | -0.25 | -0.24 | 0.037 |
| YJU2B | S362 | 0.15 | 0.41 | 0.33 | -0.34 | -0.32 | -0.49 | 0.002 |
| YLPM1 | S1100 | -0.21 | -0.16 | -0.05 | 0.20 | 0.02 | 0.15 | 0.021 |
| YLPM1 | S1100 | 0.12 | -0.30 | 0.22 | 0.11 | -0.17 | -0.05 | 0.787 |
| YLPM1 | S756 | 0.07 | -0.18 | 0.22 | -0.12 | 0.14 | -0.18 | 0.594 |
| YLPM1 | S829 | 0.18 | 0.47 | -0.56 | -0.01 | -0.28 | -0.02 | 0.704 |
| YME1L1 | S632 | -0.16 | 0.07 | -0.28 | -0.02 | -0.08 | 0.38 | 0.29 |
| YRDC | S37 | -0.02 | -0.10 | -0.01 | 0.18 | 0.02 | -0.10 | 0.403 |
| YTHDC1 | S515 | -0.26 | -0.15 | 0.12 | 0.03 | 0.34 | -0.16 | 0.421 |
| YTHDC1 | S146 | 0.00 | -0.10 | -0.09 | 0.11 | 0.03 | 0.03 | 0.047 |
| YTHDC1 | S146; T148 | 0.07 | -0.04 | -0.04 | 0.07 | -0.04 | -0.03 | 0.888 |
| YTHDC1 | S317; S318; S320 | 0.13 | -0.10 | 0.08 | -0.38 | 0.29 | -0.12 | 0.633 |
| YTHDC1 | S326 | 0.15 | 0.03 | -0.04 | -0.30 | 0.09 | 0.03 | 0.465 |
| YTHDC1 | S308 | 0.24 | 0.22 | 0.10 | -0.09 | -0.05 | -0.56 | 0.068 |
| YTHDC2 | S23 | 0.17 | -0.20 | -0.08 | -0.21 | 0.09 | 0.17 | 0.741 |
| YTHDC2 | S1269 | -0.03 | 0.16 | -0.07 | 0.08 | 0.11 | -0.30 | 0.722 |
| YTHDC2 | S23 | 0.04 | 0.06 | 0.15 | -0.28 | -0.26 | 0.23 | 0.343 |
| YTHDC2 | S1221 | 0.02 | 0.11 | 0.21 | -0.32 | -0.16 | 0.07 | 0.116 |
| YTHDC2 | S1090 | 0.06 | 0.11 | 0.18 | -0.29 | 0.10 | -0.22 | 0.108 |
| YTHDC2 | S1089; S1090 | 0.01 | 0.05 | 0.40 | -0.45 | -0.17 | 0.02 | 0.132 |
| YWHAQ | S/T | -0.03 | 0.02 | -0.10 | 0.09 | -0.11 | 0.11 | 0.438 |
| YWHAQ | T/S | -0.51 | 0.10 | 0.28 | 0.32 | 0.06 | -0.47 | 0.973 |
| YWHAQ | S64 | 0.10 | 0.06 | -0.03 | -0.07 | -0.09 | 0.02 | 0.144 |
| YY1 | T236 | 0.02 | -0.02 | 0.02 | 0.22 | -0.15 | -0.12 | 0.844 |
| YY1 | S/Y/T | 0.02 | -0.02 | 0.02 | 0.22 | -0.15 | -0.12 | 0.844 |
| ZBED6 | S381 | 0.05 | 0.11 | 0.16 | -0.11 | -0.19 | -0.05 | 0.013 |
| ZBTB1 | T169 | 0.14 | 0.23 | 0.29 | -0.34 | 0.01 | -0.50 | 0.034 |
| ZBTB1 | S355; T356 | 0.21 | 0.32 | 0.24 | -0.19 | -0.27 | -0.49 | 0.004 |
| ZBTB11 | S | 0.01 | 0.05 | -0.24 | 0.12 | -0.10 | 0.13 | 0.409 |
| ZBTB11 | S449; S450 | -0.14 | -0.02 | 0.00 | 0.25 | 0.10 | -0.25 | 0.585 |
| ZBTB11 | S449; S450 | -0.04 | -0.05 | 0.05 | 0.16 | -0.04 | -0.10 | 0.805 |
| ZBTB17 | S120 | -0.16 | -0.02 | -0.05 | 0.01 | -0.10 | 0.28 | 0.303 |
| ZBTB17 | T/S | 0.03 | 0.08 | -0.01 | -0.08 | 0.18 | -0.24 | 0.581 |
| ZBTB17 | T/S | 0.02 | 0.14 | 0.00 | -0.18 | 0.10 | -0.11 | 0.286 |
| ZBTB17 | T/S | 0.08 | 0.17 | 0.25 | -0.18 | -0.21 | -0.18 | 0.002 |
| ZBTB17 | S357 | 0.28 | 0.19 | 0.18 | -0.34 | -0.03 | -0.42 | 0.017 |
| ZBTB21 | S411 | -0.22 | -0.02 | -0.02 | 0.20 | -0.01 | 0.03 | 0.159 |
| ZBTB21 | S1003 | -0.06 | -0.02 | -0.04 | 0.01 | 0.05 | 0.05 | 0.015 |
| ZBTB21 | S/Y | -0.09 | -0.13 | -0.06 | 0.60 | 0.15 | -0.81 | 0.865 |
| ZBTB21 | S435 | 0.00 | 0.05 | 0.10 | -0.08 | -0.03 | -0.05 | 0.034 |
| ZBTB21 | S714 | 0.22 | 0.11 | 0.00 | 0.06 | -0.20 | -0.25 | 0.112 |

| | | | | | | | | |
|---|---|---|---|---|---|---|---|---|
| ZBTB21 | S/T | 0.15 | -0.07 | 0.33 | 0.16 | -0.11 | -0.66 | 0.276 |
| ZBTB3 | S362 | -0.19 | -0.12 | -0.22 | 0.14 | -0.03 | 0.33 | 0.038 |
| ZBTB3 | S369; S/T | 0.03 | 0.06 | -0.50 | 0.23 | -0.73 | 0.54 | 0.74 |
| ZBTB3 | S/T | 0.03 | 0.06 | -0.50 | 0.23 | -0.73 | 0.54 | 0.74 |
| ZBTB3 | S228 | 0.18 | 0.15 | -0.04 | -0.13 | -0.14 | -0.05 | 0.049 |
| ZBTB3 | S228 | 0.18 | 0.15 | -0.04 | -0.13 | -0.14 | -0.05 | 0.049 |
| ZBTB33 | S188 | -0.13 | -0.12 | -0.27 | 0.21 | -0.10 | 0.32 | 0.08 |
| ZBTB34 | S463 | 0.01 | 0.13 | 0.14 | -0.11 | 0.00 | -0.21 | 0.054 |
| ZBTB37 | S188; S195 | -0.12 | -0.24 | -0.47 | 0.20 | -0.11 | 0.52 | 0.085 |
| ZBTB37 | S188 | -0.16 | -0.14 | -0.08 | 0.05 | -0.04 | 0.32 | 0.097 |
| ZBTB37 | S480 | 0.09 | 0.01 | -0.56 | 0.17 | -0.49 | 0.50 | 0.592 |
| ZBTB37 | S188 | -0.13 | -0.32 | 0.09 | 0.11 | -0.29 | 0.41 | 0.454 |
| ZBTB37 | S480 | 0.19 | -0.11 | -0.04 | 0.03 | -0.35 | 0.21 | 0.794 |
| ZBTB37 | S480 | 0.26 | -0.19 | 0.04 | -0.20 | -0.34 | 0.31 | 0.661 |
| ZBTB37 | S465 | 0.02 | 0.14 | 0.06 | -0.51 | -0.04 | 0.22 | 0.446 |
| ZBTB38 | S947 | 0.13 | 0.04 | 0.07 | 0.06 | -0.33 | -0.02 | 0.219 |
| ZBTB40 | S703 | -0.65 | -0.61 | -0.60 | 0.05 | 0.51 | 0.66 | 0.005 |
| ZBTB40 | S/T | 0.10 | -1.73 | 0.22 | 0.51 | -0.28 | 0.28 | 0.397 |
| ZBTB40 | S181 | -0.05 | -1.01 | 0.12 | 0.33 | -0.12 | 0.34 | 0.262 |
| ZBTB40 | S/T | 0.09 | -0.25 | -0.18 | 0.19 | -0.12 | 0.21 | 0.237 |
| ZBTB40 | S190 | -0.12 | -0.03 | -0.06 | 0.01 | -0.13 | 0.29 | 0.381 |
| ZBTB40 | S728 | 0.10 | -0.03 | -0.03 | -0.08 | -0.01 | 0.04 | 0.656 |
| ZBTB40 | T221 | 0.05 | 0.19 | -0.06 | 0.25 | -0.13 | -0.39 | 0.486 |
| ZBTB40 | S214 | 0.08 | 0.08 | 0.26 | -0.21 | 0.03 | -0.32 | 0.066 |
| ZBTB40 | S728 | 0.10 | 0.08 | 0.36 | -0.25 | 0.15 | -0.64 | 0.155 |
| ZBTB41 | T266; S267 | -0.13 | 0.20 | 0.04 | 0.20 | 0.00 | -0.39 | 0.639 |
| ZBTB41 | S191 | 0.03 | 0.27 | 0.01 | -0.15 | 0.06 | -0.28 | 0.163 |
| ZBTB42 | S244 | 0.21 | 0.08 | -0.03 | -0.03 | -0.02 | -0.25 | 0.146 |
| ZBTB42 | S328 | -0.19 | 0.35 | 0.08 | -0.48 | 0.11 | -0.02 | 0.425 |
| ZBTB44 | S161 | -0.20 | -0.06 | -0.09 | -0.16 | -0.14 | 0.51 | 0.457 |
| ZBTB48 | S171 | -0.14 | -0.22 | -0.15 | 0.15 | -0.28 | 0.48 | 0.26 |
| ZBTB48 | S | -0.05 | -0.02 | -0.03 | 0.03 | -0.18 | 0.21 | 0.664 |
| ZBTB7A | S567 | -0.21 | -0.10 | 0.08 | 0.20 | -0.10 | 0.08 | 0.32 |
| ZBTB7A | S511; S526 | -0.18 | -0.15 | 0.16 | -0.07 | 0.08 | 0.11 | 0.472 |
| ZBTB7A | S526 | -0.12 | -0.14 | 0.22 | -0.09 | -0.07 | 0.16 | 0.915 |
| ZBTB7A | S526 | -0.07 | -0.17 | 0.23 | -0.10 | 0.07 | 0.00 | 0.952 |
| ZBTB7A | S549 | 0.04 | 0.00 | 0.05 | -0.08 | 0.01 | -0.04 | 0.109 |
| ZBTB7A | S337 | 0.07 | -0.11 | 0.17 | -0.05 | 0.02 | -0.11 | 0.381 |
| ZBTB7B | T232 | 0.20 | -0.72 | -0.25 | 0.13 | 0.05 | 0.34 | 0.196 |
| ZBTB7B | S487 | -0.45 | -0.42 | 0.15 | 0.04 | 0.03 | 0.44 | 0.162 |
| ZBTB7B | S487 | -0.45 | -0.42 | 0.15 | 0.04 | 0.03 | 0.44 | 0.162 |
| ZBTB7B | T529 | -0.07 | -0.12 | 0.19 | 0.06 | -0.06 | -0.01 | 0.967 |
| ZBTB7B | S/T/Y | 0.11 | 0.06 | -0.04 | -0.06 | -0.04 | -0.04 | 0.117 |
| ZBTB7B | S/T | 0.00 | 0.13 | 0.04 | 0.01 | -0.09 | -0.10 | 0.081 |
| ZBTB7C | S226 | -1.55 | -0.24 | 0.20 | 0.68 | 0.37 | -0.39 | 0.289 |
| ZBTB7C | S215 | -0.19 | -0.18 | 0.03 | 0.28 | 0.20 | -0.22 | 0.31 |
| ZC2HC1A | S292 | -0.14 | 0.06 | 0.06 | 0.12 | 0.03 | -0.17 | 0.983 |
| ZC3H11A | S781 | -0.20 | -0.46 | -0.37 | 0.48 | 0.03 | 0.27 | 0.015 |

| | | | | | | | | |
|---|---|---|---|---|---|---|---|---|
| **ZC3H11A** | **S758** | -0.18 | -0.23 | -0.25 | 0.29 | 0.09 | 0.18 | 0.003 |
| **ZC3H11A** | **S108** | -0.20 | -0.16 | -0.15 | 0.11 | 0.30 | 0.03 | 0.018 |
| **ZC3H11A** | **T185** | -0.28 | -0.15 | -0.09 | 0.02 | -0.09 | 0.47 | 0.163 |
| **ZC3H11A** | **S768** | -0.16 | 0.04 | -0.24 | 0.03 | 0.31 | -0.05 | 0.184 |
| **ZC3H11A** | **S758; S761** | -0.08 | -0.03 | 0.01 | -0.05 | 0.00 | 0.15 | 0.394 |
| **ZC3H11A** | **T321** | 0.05 | -0.14 | -0.03 | 0.46 | -0.29 | -0.18 | 0.888 |
| **ZC3H11A** | **T185** | -0.06 | 0.07 | -0.05 | -0.06 | -0.08 | 0.16 | 0.843 |
| **ZC3H11A** | **S290** | 0.00 | 0.15 | -0.12 | -0.17 | -0.18 | 0.26 | 0.825 |
| **ZC3H11A** | **S290** | -0.13 | 0.22 | -0.06 | 0.22 | 0.14 | -0.52 | 0.826 |
| **ZC3H11A** | **S494; S495** | -0.09 | -0.17 | 0.33 | -0.10 | -0.12 | 0.09 | 0.703 |
| **ZC3H11A** | **S494; S495** | -0.09 | -0.17 | 0.33 | -0.10 | -0.12 | 0.09 | 0.703 |
| **ZC3H11A** | **S171** | 0.02 | 0.15 | 0.12 | -0.09 | -0.07 | -0.15 | 0.011 |
| **ZC3H11A** | **S132** | 0.10 | 0.00 | 0.21 | -0.07 | -0.04 | -0.25 | 0.065 |
| **ZC3H11A** | **S738** | 0.08 | 0.13 | 0.11 | -0.15 | 0.09 | -0.31 | 0.121 |
| **ZC3H11A** | **S149; T151; S171** | 0.19 | 0.07 | 0.12 | -0.09 | -0.17 | -0.16 | 0.004 |
| **ZC3H11A** | **S149** | 0.13 | 0.13 | 0.13 | -0.26 | -0.01 | -0.18 | 0.02 |
| **ZC3H11A** | **S149; T151; S171** | 0.23 | 0.08 | 0.09 | 0.08 | -0.49 | -0.09 | 0.165 |
| **ZC3H11A** | **S/T** | 0.16 | 0.22 | 0.04 | -0.09 | 0.02 | -0.44 | 0.102 |
| **ZC3H11A** | **T151; S171** | 0.09 | 0.21 | 0.24 | -0.36 | 0.11 | -0.43 | 0.079 |
| **ZC3H11A** | **S/T** | 0.16 | -0.02 | 0.33 | 0.22 | -0.91 | -0.08 | 0.304 |
| **ZC3H11A** | **S171; S/T** | 0.11 | 0.22 | 0.31 | -0.43 | -0.11 | -0.24 | 0.012 |
| **ZC3H11A** | **T321** | 0.32 | 0.16 | 0.24 | -0.13 | -0.15 | -0.65 | 0.036 |
| **ZC3H11C** | **T/S** | 0.11 | 0.12 | 0.10 | 0.13 | -0.07 | -0.47 | 0.23 |
| **ZC3H13** | **S409** | -0.21 | 0.13 | -0.24 | -0.24 | 0.31 | 0.14 | 0.418 |
| **ZC3H13** | **S137** | -0.08 | -0.12 | -0.01 | 0.11 | -0.03 | 0.12 | 0.078 |
| **ZC3H13** | **S265** | -0.03 | 0.02 | -0.10 | 0.02 | 0.07 | 0.02 | 0.12 |
| **ZC3H13** | **S875; S877** | -0.26 | 0.04 | 0.15 | -0.03 | -0.28 | 0.29 | 0.936 |
| **ZC3H13** | **S/Y** | 0.13 | -0.09 | -0.01 | 0.06 | 0.02 | -0.12 | 0.788 |
| **ZC3H13** | **T263; S265** | 0.02 | 0.08 | -0.05 | 0.03 | -0.14 | 0.06 | 0.705 |
| **ZC3H13** | **S1065; S1068** | -0.44 | -0.18 | 0.50 | -0.06 | -0.55 | 0.40 | 0.939 |
| **ZC3H13** | **S1423** | 0.21 | 0.09 | -0.32 | 0.52 | -0.42 | -0.32 | 0.855 |
| **ZC3H13** | **S986; S993** | 0.01 | 0.10 | 0.00 | -0.01 | -0.08 | -0.02 | 0.138 |
| **ZC3H13** | **T1033** | 0.21 | 0.13 | -0.20 | 0.15 | -0.13 | -0.22 | 0.537 |
| **ZC3H13** | **S137** | -0.05 | 0.24 | -0.05 | 0.21 | -0.30 | -0.14 | 0.531 |
| **ZC3H13** | **S209** | 0.02 | 0.12 | 0.05 | -0.10 | -0.01 | -0.09 | 0.037 |
| **ZC3H13** | **S1465** | 0.14 | -0.09 | 0.15 | 0.02 | 0.02 | -0.29 | 0.322 |
| **ZC3H13** | **T263; S265** | 0.13 | 0.11 | -0.01 | 0.01 | -0.15 | -0.11 | 0.073 |
| **ZC3H13** | **S993** | 0.01 | 0.18 | 0.05 | -0.05 | -0.13 | -0.09 | 0.038 |
| **ZC3H13** | **S1364** | 0.05 | 0.12 | 0.11 | -0.11 | -0.11 | -0.08 | 0.001 |
| **ZC3H13** | **S77** | 0.08 | 0.11 | 0.11 | -0.06 | -0.11 | -0.16 | 0.003 |
| **ZC3H13** | **S877** | 0.18 | 0.07 | 0.05 | 0.00 | -0.24 | -0.09 | 0.056 |
| **ZC3H13** | **S845; S848; S853** | 0.22 | -0.02 | 0.11 | -0.17 | 0.02 | -0.20 | 0.081 |
| **ZC3H13** | **S356** | 0.08 | 0.10 | 0.13 | 0.02 | -0.06 | -0.31 | 0.095 |
| **ZC3H13** | **S207; S209** | 0.03 | 0.15 | 0.15 | -0.12 | -0.12 | -0.12 | 0.004 |
| **ZC3H13** | **S993** | 0.15 | 0.16 | 0.07 | -0.07 | -0.17 | -0.18 | 0.005 |
| **ZC3H13** | **S1366** | 0.14 | 0.03 | 0.25 | -0.01 | -0.17 | -0.32 | 0.049 |
| **ZC3H13** | **S1010; S1017** | 0.15 | 0.25 | 0.02 | -0.14 | 0.00 | -0.36 | 0.065 |
| **ZC3H13** | **T/S** | -0.06 | 0.30 | 0.19 | -0.20 | -0.10 | -0.22 | 0.047 |

| | | | | | | | | |
|---|---|---|---|---|---|---|---|---|
| ZC3H13 | S845; S848; S853 | 0.19 | -0.01 | 0.25 | -0.07 | 0.01 | -0.48 | 0.133 |
| ZC3H13 | S318; S325 | 0.05 | 0.15 | 0.26 | -0.13 | -0.07 | -0.34 | 0.032 |
| ZC3H13 | S1452; S1455; S1465 | 0.17 | 0.12 | 0.20 | -0.06 | -0.13 | -0.39 | 0.027 |
| ZC3H13 | S1068 | -0.03 | 0.37 | 0.02 | 0.13 | 0.18 | -1.02 | 0.431 |
| ZC3H13 | S198 | 0.21 | 0.21 | 0.10 | -0.13 | -0.08 | -0.40 | 0.024 |
| ZC3H13 | S370; S372; S381 | -0.05 | 0.08 | 0.50 | -0.16 | -0.24 | -0.29 | 0.076 |
| ZC3H13 | S77 | 0.17 | 0.20 | 0.19 | -0.18 | -0.09 | -0.39 | 0.01 |
| ZC3H13 | S | 0.17 | 0.21 | 0.20 | -0.15 | -0.07 | -0.49 | 0.028 |
| ZC3H13 | S993 | 0.39 | 0.20 | 0.00 | -0.22 | -0.14 | -0.37 | 0.027 |
| ZC3H13 | S316; S318; S325 | 0.26 | 0.22 | 0.15 | -0.24 | -0.13 | -0.36 | 0.004 |
| ZC3H13 | T1033 | 0.25 | 0.24 | 0.16 | -0.27 | -0.09 | -0.42 | 0.009 |
| ZC3H13 | S1191 | 0.22 | 0.29 | 0.18 | -0.21 | -0.08 | -0.58 | 0.027 |
| ZC3H13 | S64 | 0.29 | 0.14 | 0.32 | -0.34 | -0.26 | -0.32 | 7E-04 |
| ZC3H13 | S242 | 0.12 | 0.19 | 0.44 | -0.32 | -0.07 | -0.59 | 0.033 |
| ZC3H13 | S372; S/Y | -0.31 | 0.16 | 0.82 | -0.76 | -0.12 | -0.31 | 0.178 |
| ZC3H13 | S/Y | 0.29 | 0.27 | 0.29 | -0.20 | -0.19 | -0.74 | 0.022 |
| ZC3H13 | S853 | 0.30 | 0.39 | 0.27 | -0.30 | -0.24 | -0.74 | 0.01 |
| ZC3H13 | S1208 | 0.24 | 0.39 | 0.34 | -0.30 | -0.42 | -0.56 | 1E-03 |
| ZC3H13 | S1017 | 0.30 | 0.45 | 0.23 | -0.32 | -0.09 | -1.03 | 0.05 |
| ZC3H13 | S1240 | 0.28 | 0.49 | 0.35 | -0.38 | -0.24 | -1.01 | 0.021 |
| ZC3H13 | S/T | 0.33 | 0.34 | 0.49 | -0.49 | -0.15 | -1.10 | 0.027 |
| ZC3H13 | S848; S853 | 0.44 | 0.46 | 0.30 | -0.66 | -0.08 | -1.12 | 0.029 |
| ZC3H13 | S848; S853 | 0.51 | 0.55 | 0.24 | -0.72 | -0.21 | -1.11 | 0.016 |
| ZC3H13 | T364 | 0.43 | 0.31 | 0.51 | -0.57 | -0.02 | -1.60 | 0.07 |
| ZC3H13 | S/T | 0.79 | 0.77 | 0.30 | -1.02 | -0.26 | -1.50 | 0.017 |
| ZC3H14 | S343 | -0.24 | -0.22 | -0.19 | 0.29 | 0.20 | 0.06 | 0.004 |
| ZC3H14 | T/S | -0.15 | -0.14 | -0.22 | 0.27 | -0.25 | 0.37 | 0.198 |
| ZC3H14 | S579 | -0.17 | 0.12 | -0.10 | -0.07 | 0.53 | -0.50 | 0.911 |
| ZC3H14 | S409 | 0.05 | 0.00 | 0.03 | -0.01 | 0.03 | -0.12 | 0.285 |
| ZC3H14 | S579 | 0.23 | -0.22 | 0.07 | 0.31 | -0.03 | -0.52 | 0.714 |
| ZC3H14 | S515 | 0.17 | 0.09 | -0.04 | -0.04 | -0.11 | -0.09 | 0.08 |
| ZC3H14 | S620 | 0.08 | 0.11 | 0.04 | -0.01 | -0.19 | -0.05 | 0.059 |
| ZC3H14 | S285 | 0.07 | 0.14 | 0.16 | -0.20 | -0.14 | -0.07 | 0.005 |
| ZC3H14 | S132 | 0.16 | 0.14 | 0.11 | -0.19 | -0.16 | -0.11 | 6E-04 |
| ZC3H14 | S409 | 0.12 | 0.13 | 0.16 | -0.06 | 0.03 | -0.46 | 0.119 |
| ZC3H14 | S620 | 0.17 | 0.33 | 0.23 | -0.27 | -0.20 | -0.41 | 0.002 |
| ZC3H18 | S179 | -1.41 | -1.27 | -1.52 | 0.52 | 1.10 | 0.37 | 9E-04 |
| ZC3H18 | S173 | 0.02 | 0.08 | -0.16 | -0.09 | 0.13 | 0.01 | 0.71 |
| ZC3H18 | S842 | 0.09 | 0.06 | -0.07 | -0.01 | -0.06 | -0.02 | 0.347 |
| ZC3H18 | T677 | 0.05 | 0.01 | 0.06 | 0.05 | -0.05 | -0.14 | 0.207 |
| ZC3H18 | S95; T/S | 0.03 | 0.06 | 0.05 | 0.06 | -0.09 | -0.11 | 0.164 |
| ZC3H18 | T109 | 0.12 | -0.01 | 0.03 | 0.06 | -0.05 | -0.16 | 0.257 |
| ZC3H18 | S852 | 0.09 | 0.10 | 0.06 | -0.07 | -0.10 | -0.10 | 4E-04 |
| ZC3H18 | S46 | 0.12 | 0.12 | 0.19 | -0.25 | -0.07 | -0.15 | 0.006 |
| ZC3H18 | S868 | 0.14 | 0.13 | 0.31 | -0.07 | -0.13 | -0.50 | 0.045 |
| ZC3H18 | S842 | 0.26 | 0.23 | 0.11 | -0.34 | 0.01 | -0.41 | 0.031 |
| ZC3H18 | S842 | 0.26 | 0.21 | 0.22 | -0.20 | -0.06 | -0.62 | 0.038 |
| ZC3H18 | S534 | 0.25 | 0.25 | 0.25 | -0.31 | -0.01 | -0.68 | 0.041 |

| | | | | | | | | |
|---|---|---|---|---|---|---|---|---|
| ZC3H18 | T796 | 0.32 | 0.33 | 0.31 | -0.45 | -0.14 | -0.70 | 0.01 |
| ZC3H3 | S145 | -0.55 | -0.27 | -0.56 | 0.41 | -0.20 | 0.69 | 0.052 |
| ZC3H3 | S408 | -0.51 | -0.13 | -0.22 | 0.39 | -0.34 | 0.52 | 0.18 |
| ZC3H3 | S383 | 0.06 | -0.16 | -0.38 | -0.01 | -0.11 | 0.46 | 0.277 |
| ZC3H3 | S465 | 0.02 | -0.03 | 0.10 | -0.17 | 0.29 | -0.28 | 0.682 |
| ZC3H3 | S/T | -0.10 | 0.09 | 0.26 | -0.36 | 0.02 | 0.02 | 0.314 |
| ZC3H4 | S | -0.03 | -0.07 | -0.27 | -0.01 | -0.05 | 0.36 | 0.209 |
| ZC3H4 | S159 | 0.20 | 0.33 | -0.03 | -0.05 | -0.34 | -0.22 | 0.053 |
| ZC3H4 | S1114 | 0.39 | 0.12 | 0.03 | -0.04 | -0.09 | -0.57 | 0.109 |
| ZC3H4 | S159 | 0.04 | 0.58 | -0.06 | -0.24 | -0.10 | -0.44 | 0.114 |
| ZC3H4 | S1275 | 0.30 | 0.11 | 0.23 | -0.27 | 0.01 | -0.56 | 0.049 |
| ZC3H4 | S/Y/T | 0.23 | 0.34 | 0.09 | -0.30 | -0.15 | -0.35 | 0.006 |
| ZC3H4 | S1269; S1275 | 0.23 | 0.03 | 0.46 | -0.41 | -0.16 | -0.36 | 0.02 |
| ZC3H4 | S1275 | 0.18 | 0.31 | 0.26 | -0.51 | -0.14 | -0.30 | 0.008 |
| ZC3H7A | T210 | -0.28 | 0.47 | 0.06 | 0.55 | -1.50 | 0.36 | 0.704 |
| ZC3H8 | S77 | -0.07 | -0.21 | -0.22 | 0.22 | 0.02 | 0.20 | 0.017 |
| ZC3H8 | S77 | -0.12 | -0.20 | -0.04 | 0.22 | 0.05 | 0.06 | 0.035 |
| ZC3H8 | S77 | -0.19 | -0.01 | 0.03 | 0.02 | 0.05 | 0.09 | 0.212 |
| ZC3H8 | S59 | 0.14 | 0.21 | 0.06 | -0.26 | 0.04 | -0.26 | 0.054 |
| ZC3HAV1 | S526 | -0.09 | -0.43 | -0.56 | 0.34 | 0.42 | 0.05 | 0.025 |
| ZC3HAV1 | S387 | -0.32 | 0.13 | -0.19 | 0.33 | 0.07 | -0.11 | 0.297 |
| ZC3HAV1 | S275; S/T | -0.10 | 0.07 | 0.12 | 0.00 | -0.15 | 0.05 | 0.508 |
| ZC3HAV1 | S284 | 0.10 | 0.03 | 0.01 | -0.04 | -0.03 | -0.08 | 0.028 |
| ZC3HAV1 | S378 | 0.08 | 0.14 | -0.04 | 0.05 | -0.07 | -0.19 | 0.209 |
| ZC3HAV1 | S364 | 0.07 | 0.12 | 0.04 | -0.02 | 0.06 | -0.31 | 0.218 |
| ZC3HAV1 | S275 | -0.04 | 0.01 | 0.47 | -0.49 | -0.01 | -0.12 | 0.177 |
| ZC3HC1 | S24 | -0.10 | -0.17 | -0.18 | 0.17 | -0.04 | 0.26 | 0.036 |
| ZC3HC1 | S58 | -0.01 | 0.00 | -0.07 | 0.08 | 0.07 | -0.07 | 0.383 |
| ZC3HC1 | S395 | -0.23 | 0.20 | 0.02 | 0.20 | 0.23 | -0.60 | 0.875 |
| ZC3HC1 | S62 | 0.14 | -0.03 | 0.01 | -0.03 | -0.09 | -0.01 | 0.19 |
| ZC3HC1 | S62 | 0.21 | -0.02 | 0.07 | -0.11 | 0.02 | -0.20 | 0.121 |
| ZC3HC1 | S/T | 0.11 | 0.13 | -0.21 | 0.40 | 0.34 | -1.40 | 0.721 |
| ZC3HC1 | S344 | 0.05 | 0.25 | 0.01 | -0.02 | 0.06 | -0.43 | 0.236 |
| ZC3HC1 | S/T | 0.17 | 0.30 | -0.16 | -0.27 | 0.06 | -0.20 | 0.226 |
| ZC3HC1 | S410; S411 | 0.12 | 0.10 | 0.16 | 0.06 | -0.13 | -0.38 | 0.096 |
| ZC3HC1 | S370 | 0.13 | 0.19 | 0.15 | -0.20 | -0.05 | -0.28 | 0.009 |
| ZCCHC17 | S114 | 0.17 | -0.22 | -0.36 | 0.04 | 0.16 | 0.12 | 0.207 |
| ZCCHC17 | S170 | 0.29 | -0.05 | 0.02 | -0.12 | -0.10 | -0.08 | 0.151 |
| ZCCHC7 | S482 | 0.49 | 0.22 | 0.38 | -0.42 | -0.30 | -0.81 | 0.007 |
| ZCCHC8 | S/T | -0.43 | -0.72 | -0.68 | 0.67 | -0.39 | 0.75 | 0.065 |
| ZCCHC8 | S/T | -0.26 | -0.44 | -0.20 | 0.60 | -0.61 | 0.47 | 0.311 |
| ZCCHC8 | S598 | -0.13 | -0.30 | -0.31 | 0.52 | -0.43 | 0.37 | 0.253 |
| ZCCHC8 | S427 | -0.04 | -0.40 | -0.31 | 0.63 | -0.74 | 0.40 | 0.47 |
| ZCCHC8 | T479; T485 | -0.22 | 0.00 | -0.13 | 0.28 | -0.06 | 0.07 | 0.144 |
| ZCCHC8 | T479 | 0.17 | -0.08 | -0.16 | 0.07 | -0.18 | 0.15 | 0.815 |
| ZCCHC8 | S658 | 0.01 | -0.01 | 0.16 | 0.09 | -0.09 | -0.19 | 0.299 |
| ZCCHC8 | S500 | -0.10 | 0.07 | 0.20 | -0.05 | -0.16 | 0.01 | 0.273 |
| ZCCHC8 | S649 | 0.12 | -0.03 | 0.11 | 0.03 | -0.02 | -0.23 | 0.205 |

| | | | | | | | | |
|---|---|---|---|---|---|---|---|---|
| ZCCHC8 | S658 | 0.58 | -0.08 | 0.19 | -0.43 | 0.11 | -0.75 | 0.136 |
| ZCRB1 | S | 0.11 | -0.20 | -0.13 | 0.11 | -0.02 | 0.10 | 0.244 |
| ZCRB1 | S210; S216 | -0.02 | 0.10 | 0.01 | -0.19 | 0.07 | 0.01 | 0.479 |
| ZCRB1 | S210 | 0.15 | 0.27 | 0.25 | -0.37 | -0.07 | -0.38 | 0.01 |
| ZCWPW1 | S36 | 0.21 | 0.28 | 0.13 | -0.28 | -0.12 | -0.33 | 0.004 |
| ZDHHC18 | S19 | -0.26 | -0.05 | 0.14 | 0.05 | -0.14 | 0.21 | 0.573 |
| ZDHHC18 | S19 | -0.07 | -0.01 | 0.03 | 0.10 | -0.23 | 0.15 | 0.878 |
| ZDHHC18 | S366 | -0.45 | 0.37 | 0.30 | 0.27 | -0.48 | -0.27 | 0.54 |
| ZDHHC18 | S61 | 0.20 | -0.10 | 0.48 | -0.53 | -0.06 | -0.19 | 0.106 |
| ZDHHC20 | S305 | -0.44 | 0.05 | -0.33 | 0.21 | 0.08 | 0.28 | 0.056 |
| ZDHHC20 | S305 | -0.02 | 0.11 | -0.07 | 0.02 | -0.10 | 0.04 | 0.781 |
| ZDHHC5 | S380 | -0.31 | -0.39 | -0.52 | 0.30 | -0.01 | 0.59 | 0.019 |
| ZDHHC5 | S554 | -0.54 | 0.13 | -0.43 | 0.22 | -0.14 | 0.49 | 0.163 |
| ZDHHC5 | S621 | -0.11 | -0.25 | -0.17 | 0.19 | 0.02 | 0.24 | 0.014 |
| ZDHHC5 | S554 | -0.23 | 0.14 | -0.32 | 0.32 | -0.29 | 0.23 | 0.4 |
| ZDHHC5 | T348 | -0.07 | -0.04 | -0.03 | 0.02 | -0.10 | 0.20 | 0.359 |
| ZDHHC5 | S/T | -0.22 | 0.00 | 0.11 | 0.22 | 0.08 | -0.25 | 0.76 |
| ZDHHC5 | S455 | -0.06 | -0.12 | 0.10 | -0.01 | 0.23 | -0.17 | 0.754 |
| ZDHHC5 | S554 | -0.16 | 0.03 | 0.12 | -0.07 | 0.02 | 0.04 | 0.978 |
| ZDHHC5 | S299 | -0.18 | 0.03 | 0.15 | 0.10 | -0.20 | 0.07 | 0.946 |
| ZDHHC5 | S296 | -0.05 | 0.05 | 0.02 | 0.03 | 0.00 | -0.05 | 0.74 |
| ZDHHC5 | T348 | 0.26 | 0.30 | -0.21 | -0.11 | -0.02 | -0.33 | 0.224 |
| ZDHHC5 | S554 | 0.07 | 0.20 | 0.13 | -0.18 | 0.03 | -0.31 | 0.058 |
| ZDHHC5 | S/T | 0.04 | 0.06 | 0.35 | 0.02 | -0.28 | -0.28 | 0.078 |
| ZDHHC5 | S590 | 0.15 | 0.37 | 0.43 | -0.35 | -0.17 | -0.79 | 0.02 |
| ZFC3H1 | S655 | 0.01 | 0.22 | -0.27 | -0.05 | 0.02 | 0.04 | 0.898 |
| ZFC3H1 | S949 | 0.12 | -0.21 | 0.29 | -0.28 | -0.04 | 0.05 | 0.429 |
| ZFC3H1 | S655 | 0.20 | 0.17 | 0.01 | -0.52 | -0.19 | 0.21 | 0.253 |
| ZFHX3 | S1590 | -0.06 | 0.02 | -0.13 | 0.12 | -0.12 | 0.14 | 0.326 |
| ZFHX3 | S1590 | -0.06 | 0.07 | -0.03 | 0.03 | -0.15 | 0.13 | 0.929 |
| ZFHX3 | S3432 | 0.07 | 0.01 | -0.02 | -0.10 | -0.09 | 0.12 | 0.614 |
| ZFHX3 | S3409 | 0.11 | 0.07 | -0.08 | -0.05 | 0.04 | -0.10 | 0.385 |
| ZFHX3 | S2925 | 0.12 | -0.04 | 0.01 | -0.11 | 0.04 | -0.03 | 0.339 |
| ZFHX3 | S2925 | 0.10 | -0.06 | 0.08 | -0.12 | 0.05 | -0.06 | 0.29 |
| ZFHX3 | S3409 | 0.11 | 0.05 | -0.02 | 0.00 | -0.01 | -0.13 | 0.177 |
| ZFHX3 | S1189 | 0.17 | 0.06 | 0.03 | -0.05 | -0.26 | 0.02 | 0.12 |
| ZFHX3 | S2801 | 0.25 | 0.06 | -0.03 | -0.22 | 0.03 | -0.13 | 0.149 |
| ZFHX3 | S2230 | 0.16 | 0.16 | 0.27 | -0.47 | 0.00 | -0.26 | 0.035 |
| ZFHX3 | S3677 | -0.01 | 0.22 | 0.55 | -0.52 | -0.20 | -0.30 | 0.033 |
| ZFP3 | S63 | -0.10 | -0.11 | -0.42 | 0.12 | 0.00 | 0.38 | 0.07 |
| ZFP36 | S93 | -0.38 | 0.13 | -0.01 | -0.40 | 0.20 | 0.30 | 0.677 |
| ZFP36L2 | S438 | -0.30 | -0.19 | -0.10 | 0.11 | 0.23 | 0.16 | 0.006 |
| ZFP36L2 | S440 | -0.18 | -0.14 | -0.04 | 0.18 | -0.17 | 0.28 | 0.204 |
| ZFP36L2 | S125 | 0.22 | 0.03 | 0.26 | -0.23 | -0.24 | -0.12 | 0.009 |
| ZFP37 | S274 | 0.29 | 0.38 | 0.36 | -0.55 | -0.36 | -0.43 | 2E-04 |
| ZFP64 | T137 | 0.02 | 0.13 | -0.22 | 0.08 | -0.11 | 0.08 | 0.736 |
| ZFP90 | S/T | 0.05 | -0.32 | 0.04 | 0.11 | -0.07 | 0.15 | 0.366 |
| ZFP91 | S103 | -0.02 | -0.06 | -0.11 | 0.00 | -0.04 | 0.21 | 0.211 |

| | | | | | | | | |
|---|---|---|---|---|---|---|---|---|
| ZFP91 | S | 0.06 | 0.09 | 0.06 | -0.23 | 0.03 | -0.04 | 0.122 |
| ZFP91 | S | 0.24 | 0.17 | 0.24 | -0.40 | -0.26 | -0.11 | 0.006 |
| ZFP91 | S83 | 0.26 | 0.26 | 0.22 | -0.49 | -0.03 | -0.42 | 0.018 |
| ZFPM1 | S935 | -0.29 | -0.50 | 0.08 | 0.14 | -0.23 | 0.55 | 0.235 |
| ZFPM1 | S935 | -0.12 | -0.35 | 0.10 | 0.11 | 0.01 | 0.19 | 0.185 |
| ZFPM1 | S909 | 0.16 | -0.20 | -0.20 | 0.20 | -0.06 | 0.05 | 0.368 |
| ZFPM1 | S786 | 0.06 | 0.12 | -0.05 | -0.02 | -0.31 | 0.15 | 0.502 |
| ZFR | S1054 | 0.07 | -0.05 | -0.07 | 0.04 | -0.11 | 0.12 | 0.691 |
| ZFR | S1054 | 0.15 | -0.01 | -0.01 | -0.09 | -0.07 | 0.02 | 0.223 |
| ZFR | S1054 | 0.16 | 0.10 | 0.06 | -0.10 | -0.12 | -0.13 | 0.001 |
| ZFR | T550 | 0.03 | 0.02 | 0.30 | 0.20 | -0.92 | 0.08 | 0.422 |
| ZFX | S274 | -0.02 | -0.05 | -0.25 | 0.13 | -0.01 | 0.16 | 0.092 |
| ZFX | S274 | -0.15 | 0.15 | 0.07 | -0.08 | -0.14 | 0.12 | 0.648 |
| ZFYVE19 | S463 | 0.02 | -0.10 | 0.22 | -0.25 | -0.14 | 0.18 | 0.499 |
| ZFYVE19 | S144 | 0.38 | 0.15 | -0.06 | -0.31 | 0.03 | -0.32 | 0.103 |
| ZGRF1 | S880 | 0.00 | -0.22 | 0.00 | 0.26 | 0.00 | -0.09 | 0.371 |
| ZGRF1 | S793 | 0.04 | -0.01 | 0.05 | -0.11 | -0.13 | 0.15 | 0.573 |
| ZHX1 | S639; T/S | 0.19 | 0.20 | 0.04 | 0.10 | -0.44 | -0.20 | 0.12 |
| ZHX1 | T668 | 0.24 | 0.40 | -0.06 | -0.12 | -0.03 | -0.64 | 0.122 |
| ZHX3 | S946 | -0.28 | -0.21 | -0.12 | 0.06 | -0.07 | 0.48 | 0.105 |
| ZKSCAN1 | S471 | -0.75 | -1.02 | -1.21 | 0.60 | 0.64 | 0.48 | 4E-04 |
| ZKSCAN1 | T400 | -0.27 | -0.35 | -0.24 | 0.21 | 0.09 | 0.39 | 0.006 |
| ZKSCAN1 | S13 | -0.14 | -0.12 | -0.07 | 0.16 | 0.13 | 0.01 | 0.012 |
| ZKSCAN1 | S13 | -0.14 | -0.35 | 0.14 | 0.01 | -0.04 | 0.29 | 0.303 |
| ZKSCAN2 | S144 | -0.71 | -0.34 | -0.16 | 0.23 | 0.19 | 0.47 | 0.02 |
| ZKSCAN2 | S144 | -0.38 | -0.28 | -0.33 | 0.41 | 0.08 | 0.28 | 0.004 |
| ZKSCAN5 | T/S | -0.44 | 0.00 | -0.11 | 0.54 | -0.23 | 0.04 | 0.314 |
| ZKSCAN8 | S12 | -0.07 | -0.13 | -0.19 | 0.08 | 0.06 | 0.21 | 0.015 |
| ZKSCAN8 | S193 | 0.12 | 0.13 | -0.04 | -0.04 | 0.09 | -0.30 | 0.291 |
| ZKSCAN8 | S193 | 0.14 | 0.17 | 0.01 | 0.00 | -0.07 | -0.30 | 0.091 |
| ZMAT4 | S121 | 0.34 | 0.32 | 0.41 | -0.50 | -0.16 | -0.84 | 0.013 |
| ZMYM2 | S305 | 0.27 | -0.48 | 0.02 | 0.32 | -0.03 | -0.27 | 0.821 |
| ZMYM2 | S838 | -0.18 | -0.08 | 0.15 | -0.05 | 0.08 | 0.05 | 0.595 |
| ZMYM2 | S838 | -0.05 | 0.05 | 0.28 | -0.21 | -0.03 | -0.08 | 0.152 |
| ZMYM2 | S/Y | 0.20 | 0.24 | -0.08 | 0.04 | -0.18 | -0.29 | 0.129 |
| ZMYM2 | S1064 | 0.54 | 0.51 | 0.19 | -0.17 | -1.18 | -0.64 | 0.026 |
| ZMYM3 | S1045 | 0.05 | -0.10 | -0.26 | -0.27 | -0.23 | 0.60 | 0.674 |
| ZMYM3 | S950 | 0.05 | 0.06 | -0.17 | 0.12 | -0.01 | -0.06 | 0.703 |
| ZMYM3 | S51 | -0.10 | 0.10 | -0.05 | 0.16 | -0.13 | 0.00 | 0.809 |
| ZMYM3 | T817 | 0.00 | 0.10 | 0.16 | -0.12 | -0.06 | -0.11 | 0.022 |
| ZMYM3 | T795 | 0.05 | 0.08 | 0.22 | -0.23 | -0.01 | -0.16 | 0.041 |
| ZMYM3 | T795 | 0.06 | 0.11 | 0.28 | -0.03 | -0.19 | -0.31 | 0.035 |
| ZMYM3 | S/T | 0.32 | 0.20 | -0.08 | 0.32 | -0.30 | -0.73 | 0.305 |
| ZMYM3 | S464 | 0.11 | 0.20 | 0.26 | -0.37 | 0.07 | -0.42 | 0.056 |
| ZMYM3 | S774 | -0.29 | 0.46 | 0.38 | -0.57 | -0.18 | -0.08 | 0.177 |
| ZMYM4 | S1064 | -0.09 | -0.16 | -0.18 | 0.19 | -0.06 | 0.25 | 0.054 |
| ZMYM4 | S162 | -0.08 | -0.22 | -0.06 | 0.12 | 0.14 | 0.06 | 0.014 |
| ZMYM4 | T217 | 0.17 | 0.23 | -0.78 | -0.08 | 0.29 | -0.07 | 0.643 |

| | | | | | | | | |
|---|---|---|---|---|---|---|---|---|
| ZMYM4 | S1030 | -0.08 | -0.04 | -0.09 | 0.05 | 0.04 | 0.10 | 0.005 |
| ZMYM4 | S122 | -0.05 | -0.15 | 0.06 | 0.05 | 0.02 | 0.05 | 0.234 |
| ZMYM4 | S1434 | -0.09 | -0.38 | 0.35 | 0.32 | -0.38 | 0.00 | 0.941 |
| ZMYM4 | S1030 | 0.03 | 0.01 | -0.07 | 0.04 | 0.09 | -0.11 | 0.788 |
| ZMYM4 | S122 | -0.03 | -0.07 | 0.07 | 0.07 | -0.14 | 0.08 | 0.906 |
| ZMYM4 | S1104 | 0.24 | 0.12 | -0.18 | -0.03 | 0.13 | -0.37 | 0.49 |
| ZMYM4 | S1104 | 0.19 | 0.14 | -0.12 | -0.23 | 0.16 | -0.19 | 0.371 |
| ZMYM4 | S122 | 0.20 | 0.17 | 0.18 | -0.18 | -0.13 | -0.32 | 0.002 |
| ZMYM4 | S | 0.15 | 0.17 | 0.32 | -0.24 | -0.04 | -0.51 | 0.031 |
| ZMYND11 | S/T | -0.55 | -0.74 | -0.76 | 0.42 | 0.63 | 0.32 | 5E-04 |
| ZMYND11 | S/T | -0.37 | -0.77 | -0.19 | 0.13 | 0.46 | 0.37 | 0.018 |
| ZMYND11 | S/T | -0.30 | -0.26 | -0.50 | 0.18 | 0.38 | 0.28 | 0.003 |
| ZMYND11 | S | -0.23 | 0.30 | 0.12 | -0.04 | -0.52 | 0.22 | 0.547 |
| ZMYND11 | S403 | 0.22 | 0.12 | -0.04 | -0.24 | 0.00 | -0.12 | 0.1 |
| ZMYND11 | S/T | 0.20 | 0.09 | 0.17 | -0.35 | 0.13 | -0.34 | 0.104 |
| ZMYND11 | S380 | 0.23 | 0.24 | 0.18 | -0.57 | 0.20 | -0.54 | 0.107 |
| ZMYND8 | S682 | 0.00 | -0.14 | -0.33 | 0.12 | 0.06 | 0.22 | 0.056 |
| ZMYND8 | S1085 | 0.08 | -0.18 | -0.18 | 0.02 | 0.18 | 0.04 | 0.155 |
| ZMYND8 | S425 | -0.20 | -0.06 | -0.01 | 0.00 | 0.22 | 0.02 | 0.133 |
| ZMYND8 | S668 | 0.01 | -0.08 | -0.17 | 0.03 | -0.06 | 0.23 | 0.213 |
| ZMYND8 | S1141 | -0.17 | -0.12 | 0.06 | 0.07 | 0.10 | 0.04 | 0.105 |
| ZMYND8 | S668 | 0.04 | -0.04 | -0.18 | 0.09 | -0.15 | 0.20 | 0.427 |
| ZMYND8 | S444 | 0.25 | -0.50 | 0.06 | 0.03 | -0.09 | 0.14 | 0.714 |
| ZMYND8 | S547 | -0.11 | 0.06 | -0.10 | 0.09 | 0.26 | -0.25 | 0.645 |
| ZMYND8 | S425 | -0.03 | -0.06 | -0.02 | 0.08 | -0.06 | 0.08 | 0.211 |
| ZMYND8 | S/T | -0.31 | 0.01 | 0.21 | -0.04 | 0.08 | 0.00 | 0.799 |
| ZMYND8 | S12 | -0.03 | 0.07 | -0.02 | -0.02 | -0.01 | 0.01 | 0.67 |
| ZMYND8 | S668 | 0.18 | 0.05 | -0.21 | 0.00 | 0.03 | -0.07 | 0.877 |
| ZMYND8 | T746; S756 | 0.07 | -0.13 | 0.10 | -0.04 | -0.20 | 0.16 | 0.781 |
| ZMYND8 | T746; S756 | 0.02 | -0.16 | 0.22 | -0.16 | 0.09 | -0.06 | 0.61 |
| ZMYND8 | S797 | 0.24 | -0.20 | 0.05 | 0.10 | 0.04 | -0.30 | 0.646 |
| ZMYND8 | S406 | 0.36 | -0.62 | 0.30 | -0.26 | -0.33 | 0.27 | 0.768 |
| ZMYND8 | S756 | 0.04 | 0.05 | 0.07 | 0.08 | -0.25 | -0.02 | 0.295 |
| ZMYND8 | S547 | 0.08 | 0.07 | 0.03 | -0.11 | 0.03 | -0.11 | 0.062 |
| ZMYND8 | S475; S478 | 0.11 | -0.04 | 0.16 | -0.23 | 0.10 | -0.13 | 0.234 |
| ZMYND8 | S797 | 0.14 | 0.09 | 0.01 | 0.10 | -0.07 | -0.31 | 0.244 |
| ZMYND8 | S432; S444 | -0.08 | 0.30 | 0.02 | 0.28 | -0.55 | -0.13 | 0.462 |
| ZMYND8 | S406 | 0.16 | 0.03 | 0.12 | -0.16 | 0.04 | -0.24 | 0.077 |
| ZMYND8 | S432; S444; S/T | 0.12 | 0.05 | 0.17 | -0.08 | -0.26 | -0.03 | 0.038 |
| ZMYND8 | S432; S444; S/T | 0.32 | 0.10 | -0.10 | 0.19 | -0.31 | -0.33 | 0.282 |
| ZMYND8 | S650; S655 | 0.11 | 0.11 | 0.17 | -0.13 | -0.15 | -0.15 | 2E-04 |
| ZMYND8 | S444 | 0.24 | 0.19 | 0.00 | -0.05 | -0.35 | -0.11 | 0.059 |
| ZMYND8 | S475 | 0.17 | 0.09 | 0.22 | -0.24 | -0.03 | -0.28 | 0.016 |
| ZMYND8 | S425 | 0.07 | 0.25 | 0.32 | -0.25 | -0.17 | -0.35 | 0.007 |
| ZMYND8 | S/T | 0.29 | 0.29 | 0.39 | -0.25 | -0.19 | -0.94 | 0.032 |
| ZNF101 | S152 | 0.19 | -0.03 | -0.02 | -0.05 | -0.06 | -0.04 | 0.256 |
| ZNF106 | S1237 | -1.56 | -0.83 | -2.01 | 0.92 | 0.80 | 0.29 | 0.006 |
| ZNF106 | S1302 | -0.35 | -0.06 | -0.30 | 0.44 | 0.08 | 0.06 | 0.049 |

| | | | | | | | | |
|---|---|---|---|---|---|---|---|---|
| ZNF106 | S1328 | -0.23 | -0.07 | -0.02 | 0.14 | -0.10 | 0.24 | 0.174 |
| ZNF106 | S1370 | -0.13 | -0.02 | -0.02 | -0.03 | 0.01 | 0.18 | 0.213 |
| ZNF106 | S878 | 0.47 | -0.28 | -0.27 | -0.07 | -0.09 | 0.10 | 0.979 |
| ZNF106 | S641 | -0.11 | 0.02 | 0.08 | 0.07 | -0.03 | -0.04 | 0.957 |
| ZNF106 | T/S | -0.22 | 0.29 | -0.05 | -0.02 | -0.01 | -0.03 | 0.888 |
| ZNF106 | S861; S | 0.11 | -0.01 | -0.06 | -0.02 | -0.19 | 0.15 | 0.783 |
| ZNF106 | T1021 | 0.19 | -0.15 | -0.02 | -0.13 | -0.16 | 0.22 | 0.844 |
| ZNF106 | S1026 | 0.02 | 0.04 | 0.00 | 0.10 | -0.10 | -0.06 | 0.549 |
| ZNF106 | S1025; S1026 | 0.06 | 0.05 | 0.08 | -0.03 | -0.14 | -0.03 | 0.029 |
| ZNF106 | S1205 | -0.07 | 0.31 | -0.07 | -0.09 | 0.16 | -0.33 | 0.485 |
| ZNF106 | S1370 | 0.13 | 0.09 | 0.00 | -0.05 | -0.19 | -0.01 | 0.073 |
| ZNF106 | S1205 | 0.06 | 0.03 | 0.14 | -0.28 | -0.01 | 0.02 | 0.182 |
| ZNF106 | S861 | 0.07 | 0.05 | 0.11 | -0.04 | -0.12 | -0.10 | 0.005 |
| ZNF106 | S1205 | 0.23 | -0.10 | 0.10 | -0.18 | 0.19 | -0.33 | 0.37 |
| ZNF106 | T/S | -0.23 | 0.33 | 0.06 | -0.76 | 0.01 | 0.32 | 0.617 |
| ZNF106 | S1279 | 0.21 | 0.07 | 0.02 | -0.22 | -0.02 | -0.09 | 0.063 |
| ZNF106 | S861 | 0.15 | 0.11 | 0.06 | -0.01 | -0.07 | -0.27 | 0.06 |
| ZNF106 | T443 | 0.18 | 0.11 | 0.03 | -0.11 | -0.18 | -0.07 | 0.012 |
| ZNF106 | T/S | 0.27 | 0.15 | -0.09 | -0.07 | 0.00 | -0.33 | 0.177 |
| ZNF106 | S1205 | 0.09 | 0.13 | 0.19 | -0.10 | -0.01 | -0.37 | 0.054 |
| ZNF106 | S893 | 0.06 | 0.10 | 0.29 | -0.32 | -0.08 | -0.13 | 0.03 |
| ZNF106 | S1302 | 0.23 | 0.16 | 0.12 | -0.12 | -0.12 | -0.35 | 0.012 |
| ZNF106 | S1302 | 0.08 | 0.27 | 0.21 | -0.21 | -0.05 | -0.42 | 0.028 |
| ZNF106 | S1328 | -0.07 | 0.33 | 0.07 | 0.24 | 0.30 | -1.60 | 0.508 |
| ZNF106 | S1328 | -0.07 | 0.33 | 0.07 | 0.24 | 0.30 | -1.60 | 0.508 |
| ZNF106 | S1279 | 0.21 | 0.23 | 0.16 | -0.29 | 0.12 | -0.62 | 0.098 |
| ZNF106 | S1477 | 0.20 | 0.03 | 0.42 | -0.38 | -0.05 | -0.40 | 0.038 |
| ZNF106 | S1328 | 0.28 | 0.22 | 0.15 | -0.58 | 0.05 | -0.31 | 0.055 |
| ZNF136 | S292 | 0.13 | 0.37 | 0.12 | -0.53 | -0.14 | -0.10 | 0.043 |
| ZNF14 | T554 | 0.14 | -0.15 | -0.44 | 0.37 | -0.28 | 0.20 | 0.387 |
| ZNF142 | S/T | -1.15 | 0.22 | 0.16 | -0.32 | 0.37 | 0.24 | 0.515 |
| ZNF142 | S/T | -0.10 | 0.04 | -0.17 | -0.69 | 0.12 | 0.53 | 0.868 |
| ZNF142 | S1319 | -0.15 | -0.25 | 0.18 | -0.75 | 0.42 | 0.25 | 0.903 |
| ZNF142 | T/S | 0.03 | 0.04 | -0.05 | -0.07 | -0.01 | 0.05 | 0.707 |
| ZNF142 | T/S | 0.13 | 0.18 | -0.27 | -0.25 | -0.12 | 0.25 | 0.811 |
| ZNF142 | S1674 | 0.00 | 0.03 | 0.14 | -0.02 | -0.25 | 0.07 | 0.304 |
| ZNF142 | S/T | 0.13 | 0.38 | -0.34 | 0.02 | 0.06 | -0.39 | 0.566 |
| ZNF142 | T/S | 0.22 | -0.08 | 0.12 | -0.17 | 0.00 | -0.13 | 0.152 |
| ZNF142 | T/S | 0.09 | 0.38 | 0.01 | 0.28 | -0.16 | -0.94 | 0.316 |
| ZNF148 | S784 | -0.25 | -0.17 | -0.06 | 0.17 | 0.17 | 0.09 | 0.008 |
| ZNF148 | S306 | 0.02 | -0.16 | -0.12 | 0.00 | 0.06 | 0.18 | 0.088 |
| ZNF148 | T/S/Y | 0.03 | 0.13 | -0.29 | 0.11 | 0.06 | -0.08 | 0.646 |
| ZNF148 | S306 | -0.01 | 0.01 | -0.07 | -0.02 | 0.00 | 0.09 | 0.297 |
| ZNF148 | T410 | 0.10 | 0.11 | -0.16 | -0.02 | -0.09 | 0.03 | 0.654 |
| ZNF160 | S123 | 0.04 | -0.03 | 0.07 | -0.05 | -0.11 | 0.07 | 0.417 |
| ZNF165 | S153 | 0.33 | -0.73 | 0.03 | 0.08 | 0.13 | -0.03 | 0.595 |
| ZNF174 | S188 | 0.26 | -0.23 | 0.10 | -0.16 | -0.12 | 0.09 | 0.551 |
| ZNF174 | S188; S/Y | 0.46 | 0.17 | -0.02 | -0.19 | -0.21 | -0.37 | 0.039 |

| | | | | | | | | |
|---|---|---|---|---|---|---|---|---|
| **ZNF174** | **S287** | 0.11 | 0.46 | 0.14 | -0.41 | -0.18 | -0.30 | 0.015 |
| **ZNF175** | **S442** | 0.15 | 0.23 | 0.11 | -0.20 | -0.06 | -0.31 | 0.012 |
| **ZNF177** | **T/S** | -0.34 | 0.04 | 0.04 | 0.02 | 0.03 | 0.16 | 0.309 |
| **ZNF184** | **S199** | 0.16 | -0.40 | -0.60 | 0.43 | -0.06 | 0.20 | 0.153 |
| **ZNF184** | **S117** | 0.09 | 0.07 | 0.17 | -0.05 | -0.07 | -0.25 | 0.03 |
| **ZNF185** | **T513** | -0.47 | 0.04 | -0.05 | 0.36 | 0.15 | -0.16 | 0.268 |
| **ZNF185** | **T513** | -0.47 | 0.04 | -0.05 | 0.36 | 0.15 | -0.16 | 0.268 |
| **ZNF185** | **S465** | -0.18 | -0.06 | -0.14 | 0.16 | 0.07 | 0.10 | 0.006 |
| **ZNF185** | **S519; T/S** | 0.04 | -0.06 | -0.20 | -0.01 | -0.16 | 0.34 | 0.473 |
| **ZNF185** | **T505** | -0.05 | 0.10 | -0.10 | 0.00 | 0.12 | -0.10 | 0.812 |
| **ZNF185** | **S/T** | 0.00 | -0.01 | 0.01 | -0.24 | 0.03 | 0.18 | 0.931 |
| **ZNF185** | **T158** | -0.04 | 0.06 | -0.02 | -0.28 | 0.07 | 0.17 | 0.92 |
| **ZNF185** | **S453** | -0.15 | 0.19 | 0.07 | -0.13 | 0.19 | -0.21 | 0.622 |
| **ZNF202** | **T387** | -0.18 | -0.28 | -0.04 | 0.17 | -0.01 | 0.28 | 0.044 |
| **ZNF202** | **S33** | -0.05 | -0.07 | -0.12 | -0.10 | 0.15 | 0.17 | 0.149 |
| **ZNF205** | **S292** | -0.12 | 0.09 | 0.01 | 0.00 | -0.19 | 0.18 | 0.979 |
| **ZNF217** | **S441** | -0.08 | -0.30 | 0.09 | 0.03 | -0.09 | 0.29 | 0.341 |
| **ZNF217** | **S441; S445** | 0.06 | -0.05 | -0.20 | 0.02 | -0.07 | 0.20 | 0.352 |
| **ZNF217** | **S570** | 0.20 | -0.09 | -0.01 | -0.14 | -0.06 | 0.07 | 0.487 |
| **ZNF217** | **S662** | -0.32 | 0.30 | 0.11 | -0.29 | -0.09 | 0.18 | 0.688 |
| **ZNF219** | **S684** | -0.39 | -0.42 | -0.48 | 0.15 | 0.51 | 0.31 | 0.002 |
| **ZNF219** | **T257** | 0.13 | -0.13 | -0.50 | -0.37 | 0.60 | 0.00 | 0.511 |
| **ZNF219** | **T257** | 0.13 | -0.13 | -0.50 | -0.37 | 0.60 | 0.00 | 0.511 |
| **ZNF219** | **S114** | 0.11 | 0.07 | -0.30 | -0.16 | 0.07 | 0.16 | 0.694 |
| **ZNF219** | **S698** | -0.15 | 0.16 | -0.03 | 0.03 | -0.05 | 0.02 | 0.955 |
| **ZNF219** | **S698** | 0.25 | 0.00 | 0.14 | -0.31 | -0.24 | 0.08 | 0.111 |
| **ZNF22** | **S49** | -1.92 | -2.04 | -2.24 | 0.95 | 0.65 | 0.83 | 2E-05 |
| **ZNF239** | **T/S/Y** | -0.04 | 0.09 | 0.11 | -0.18 | -0.04 | 0.04 | 0.247 |
| **ZNF263** | **S153** | -0.05 | -0.03 | -0.05 | 0.05 | -0.03 | 0.10 | 0.099 |
| **ZNF263** | **S166** | -0.24 | 0.09 | 0.08 | -0.21 | -0.01 | 0.23 | 0.857 |
| **ZNF276** | **S/T** | -0.05 | -0.06 | -0.31 | 0.16 | 0.07 | 0.13 | 0.04 |
| **ZNF276** | **S357; S360; S366** | -0.05 | -0.06 | -0.04 | -0.05 | 0.16 | 0.04 | 0.168 |
| **ZNF276** | **S378; S379** | 0.18 | 0.07 | -0.12 | 0.06 | -0.20 | -0.01 | 0.462 |
| **ZNF276** | **S378; S379** | 0.18 | 0.07 | -0.12 | 0.06 | -0.20 | -0.01 | 0.462 |
| **ZNF280C** | **S227** | -0.18 | -0.36 | -0.25 | 0.19 | 0.15 | 0.32 | 0.003 |
| **ZNF280C** | **S80** | -0.01 | -0.19 | -0.09 | 0.11 | 0.00 | 0.15 | 0.052 |
| **ZNF280C** | **T/S** | 0.18 | -0.09 | 0.54 | -0.14 | -0.34 | -0.38 | 0.064 |
| **ZNF280D** | **S790** | -0.92 | -1.20 | -0.80 | 0.50 | 0.43 | 0.76 | 6E-04 |
| **ZNF280D** | **S179** | -0.19 | -0.35 | -0.33 | 0.36 | 0.17 | 0.18 | 0.002 |
| **ZNF280D** | **S179** | -0.11 | -0.28 | -0.23 | 0.26 | -0.13 | 0.36 | 0.083 |
| **ZNF280D** | **S545** | -0.09 | 0.11 | -0.38 | 0.19 | 0.15 | -0.06 | 0.263 |
| **ZNF280D** | **S806** | 0.02 | 0.10 | 0.14 | -0.20 | 0.07 | -0.16 | 0.113 |
| **ZNF280D** | **T/S** | 0.11 | 0.05 | 0.10 | -0.33 | 0.04 | -0.02 | 0.175 |
| **ZNF281** | **S620** | -1.08 | -0.75 | -0.80 | 0.53 | 0.48 | 0.60 | 2E-04 |
| **ZNF281** | **S/T** | -0.35 | -0.12 | -0.79 | 0.33 | -0.05 | 0.58 | 0.06 |
| **ZNF281** | **S807** | -0.41 | -0.29 | -0.19 | -0.04 | 0.27 | 0.46 | 0.03 |
| **ZNF282** | **S319** | -0.29 | -0.34 | -0.20 | 0.16 | -0.01 | 0.50 | 0.034 |
| **ZNF286A** | **S19** | 0.07 | 0.18 | -0.10 | -0.34 | -0.02 | 0.15 | 0.51 |

| | | | | | | | | |
|---|---|---|---|---|---|---|---|---|
| ZNF292 | S1992 | 0.16 | -0.09 | -0.03 | 0.21 | -0.08 | -0.21 | 0.791 |
| ZNF296 | S230 | -0.09 | -0.65 | -0.38 | 0.18 | 0.19 | 0.46 | 0.025 |
| ZNF296 | S230 | 0.03 | -0.11 | -0.14 | -0.17 | 0.04 | 0.29 | 0.433 |
| ZNF3 | S143 | -0.46 | 0.11 | -0.11 | -0.50 | 0.57 | 0.10 | 0.58 |
| ZNF316 | S112 | 0.16 | 0.04 | -0.21 | -0.16 | 0.02 | 0.12 | 0.978 |
| ZNF318 | S1420 | -0.44 | -0.27 | -0.44 | 0.23 | 0.40 | 0.27 | 1E-03 |
| ZNF318 | S2101 | 0.09 | -0.25 | -0.32 | 0.32 | 0.14 | -0.08 | 0.167 |
| ZNF318 | S79 | -0.37 | -0.07 | 0.00 | 0.20 | -0.16 | 0.30 | 0.224 |
| ZNF318 | S2035 | -0.05 | -0.28 | -0.12 | 0.07 | -0.35 | 0.54 | 0.419 |
| ZNF318 | S/T | 0.15 | -0.66 | 0.06 | -0.29 | 0.26 | 0.26 | 0.507 |
| ZNF318 | T/S | -0.59 | -0.12 | 0.30 | -0.65 | 0.07 | 0.59 | 0.768 |
| ZNF318 | S136 | 0.11 | 0.00 | -0.25 | 0.19 | -0.04 | -0.05 | 0.578 |
| ZNF318 | S89 | -0.21 | -0.11 | 0.21 | 0.01 | 0.05 | 0.02 | 0.643 |
| ZNF318 | S1971 | 0.14 | 0.01 | -0.20 | 0.12 | -0.09 | 0.00 | 0.815 |
| ZNF318 | S2091 | -0.02 | -0.08 | 0.05 | 0.20 | 0.21 | -0.46 | 0.984 |
| ZNF318 | S40 | 0.00 | 0.03 | -0.02 | 0.02 | -0.12 | 0.08 | 0.945 |
| ZNF318 | S173 | 0.09 | 0.04 | 0.06 | 0.02 | -0.02 | -0.21 | 0.149 |
| ZNF318 | S1037 | 0.16 | 0.12 | -0.05 | -0.15 | 0.08 | -0.19 | 0.206 |
| ZNF318 | S709 | 0.18 | -0.08 | 0.14 | -0.23 | -0.06 | 0.01 | 0.195 |
| ZNF318 | S1243 | 0.06 | 0.08 | 0.12 | -0.08 | 0.07 | -0.29 | 0.161 |
| ZNF318 | S1896 | 0.19 | -0.20 | 0.30 | 0.02 | -0.47 | 0.02 | 0.333 |
| ZNF318 | S | 0.27 | 0.19 | 0.24 | -0.29 | 0.08 | -0.72 | 0.079 |
| ZNF326 | S478 | -0.96 | -1.03 | -0.94 | 0.58 | 0.12 | 0.92 | 0.003 |
| ZNF326 | S478 | -0.59 | -0.77 | -0.75 | 0.42 | 0.31 | 0.66 | 6E-04 |
| ZNF326 | S478 | -0.60 | -0.74 | -0.68 | 0.51 | 0.22 | 0.62 | 9E-04 |
| ZNF326 | S118 | -0.15 | -0.26 | -0.08 | 0.06 | 0.07 | 0.30 | 0.031 |
| ZNF326 | S63 | 0.02 | -0.42 | -0.10 | 0.18 | -0.24 | 0.40 | 0.289 |
| ZNF326 | S63 | -0.05 | -0.15 | -0.10 | 0.08 | 0.08 | 0.12 | 0.004 |
| ZNF326 | S56 | -0.27 | -0.09 | 0.24 | 0.16 | -0.14 | 0.04 | 0.756 |
| ZNF326 | S56 | -0.30 | 0.04 | 0.42 | 0.00 | -0.19 | -0.07 | 0.534 |
| ZNF326 | S76 | 0.08 | 0.48 | 0.27 | -0.37 | -0.26 | -0.45 | 0.008 |
| ZNF335 | S1007 | -0.03 | -0.22 | -0.18 | 0.27 | 0.03 | 0.06 | 0.052 |
| ZNF34 | S113 | 0.21 | 0.05 | -0.08 | -0.35 | 0.32 | -0.27 | 0.517 |
| ZNF358 | S553 | -0.01 | -0.59 | 0.10 | -0.12 | -0.02 | 0.45 | 0.389 |
| ZNF358 | T485; S487 | 0.15 | -0.08 | 0.16 | 0.19 | -0.11 | -0.41 | 0.379 |
| ZNF358 | S487 | 0.14 | 0.11 | 0.31 | -0.19 | -0.23 | -0.22 | 0.003 |
| ZNF367 | S310 | 0.19 | -0.08 | -0.04 | -0.17 | 0.11 | -0.04 | 0.643 |
| ZNF367 | S310 | 0.02 | 0.18 | 0.04 | -0.11 | -0.20 | 0.04 | 0.119 |
| ZNF367 | S310 | 0.09 | 0.12 | 0.08 | -0.15 | -0.17 | 0.00 | 0.019 |
| ZNF384 | S214 | 0.18 | -0.12 | -0.72 | 0.02 | 0.11 | 0.33 | 0.255 |
| ZNF384 | S214 | -0.12 | 0.02 | -0.27 | 0.00 | 0.26 | 0.05 | 0.119 |
| ZNF384 | S214 | -0.02 | 0.26 | -0.41 | 0.21 | 0.12 | -0.30 | 0.799 |
| ZNF384 | S214 | -0.40 | 0.17 | 0.21 | -0.65 | 0.32 | 0.11 | 0.861 |
| ZNF384 | S214 | -0.58 | 0.40 | 0.16 | -0.38 | 0.30 | -0.16 | 0.855 |
| ZNF384 | S214 | -0.04 | 0.08 | 0.13 | -0.30 | -0.06 | 0.14 | 0.405 |
| ZNF384 | S234 | 0.23 | 0.02 | 0.08 | 0.16 | -0.03 | -0.60 | 0.323 |
| ZNF385A | T180 | 0.04 | -0.22 | 0.06 | 0.02 | 0.02 | 0.06 | 0.452 |
| ZNF385A | S160 | 0.08 | 0.10 | -0.04 | -0.17 | -0.11 | 0.11 | 0.339 |

| | | | | | | | | |
|---|---|---|---|---|---|---|---|---|
| ZNF385A | S/T | 0.59 | 0.20 | -0.21 | -1.50 | 0.46 | 0.15 | 0.493 |
| ZNF394 | S258 | -0.05 | 0.26 | -0.13 | 0.51 | -0.19 | -0.69 | 0.701 |
| ZNF397 | S142 | 0.13 | -0.03 | -0.11 | -0.02 | 0.01 | 0.01 | 0.997 |
| ZNF397 | S31 | -0.01 | -0.02 | 0.13 | -0.31 | 0.15 | 0.01 | 0.591 |
| ZNF397 | S142 | -0.07 | 0.18 | 0.23 | -0.02 | -0.66 | 0.16 | 0.343 |
| ZNF407 | S1262 | -0.03 | 0.07 | -0.02 | -0.21 | -0.01 | 0.17 | 0.849 |
| ZNF408 | S338 | 0.06 | 0.00 | 0.07 | 0.06 | -0.09 | -0.12 | 0.182 |
| ZNF428 | T108 | 0.02 | 0.07 | 0.01 | -0.14 | -0.17 | 0.18 | 0.516 |
| ZNF444 | S232 | 0.21 | 0.17 | 0.06 | -0.24 | -0.23 | -0.03 | 0.018 |
| ZNF444 | S162 | 0.44 | 0.07 | 0.10 | -0.28 | -0.34 | -0.14 | 0.028 |
| ZNF445 | S665 | -0.12 | -0.08 | -0.31 | 0.13 | 0.16 | 0.15 | 0.012 |
| ZNF445 | S665 | -0.08 | -0.09 | -0.04 | 0.08 | -0.19 | 0.28 | 0.397 |
| ZNF445 | S150 | -0.15 | -0.12 | -0.12 | 0.89 | -0.41 | -0.63 | 0.871 |
| ZNF445 | S665 | 0.09 | 0.02 | -0.23 | 0.41 | -0.28 | -0.13 | 0.881 |
| ZNF445 | S665 | -0.07 | -0.06 | 0.08 | 0.06 | -0.21 | 0.17 | 0.881 |
| ZNF446 | S137 | -0.19 | -0.29 | 0.02 | 0.00 | 0.11 | 0.28 | 0.084 |
| ZNF446 | S137; S141; S146 | -0.27 | 0.13 | 0.02 | 0.01 | 0.03 | 0.05 | 0.599 |
| ZNF446 | S137; S/T | -0.11 | 0.16 | 0.12 | -0.04 | -0.18 | 0.03 | 0.308 |
| ZNF446 | S137 | -0.10 | -0.01 | 0.24 | -0.56 | -0.06 | 0.33 | 0.644 |
| ZNF446 | S180; S181 | 0.10 | 0.23 | -0.06 | -0.07 | -0.12 | -0.10 | 0.091 |
| ZNF462 | S2172 | 0.09 | -0.06 | 0.08 | -0.09 | 0.15 | -0.21 | 0.481 |
| ZNF467 | T56 | 0.00 | -0.04 | 0.10 | 0.22 | -0.10 | -0.21 | 0.734 |
| ZNF48 | S416 | 0.23 | 0.24 | -0.27 | 0.26 | -0.03 | -0.65 | 0.548 |
| ZNF496 | S23 | -0.40 | -0.44 | -0.19 | 0.22 | 0.17 | 0.43 | 0.005 |
| ZNF496 | S23 | -0.12 | -0.23 | -0.14 | 0.15 | 0.33 | -0.06 | 0.058 |
| ZNF496 | S23 | -0.12 | -0.16 | 0.08 | -0.28 | 0.12 | 0.27 | 0.595 |
| ZNF496 | S23 | 0.06 | -0.02 | 0.07 | -0.13 | 0.00 | 0.02 | 0.252 |
| ZNF497 | S98 | 0.35 | -0.10 | 0.09 | 0.06 | -0.32 | -0.18 | 0.198 |
| ZNF507 | S95 | -0.02 | -0.38 | 0.26 | -0.18 | 0.00 | 0.22 | 0.796 |
| ZNF512B | S349 | -0.62 | 0.20 | -0.65 | 0.36 | 0.12 | 0.26 | 0.105 |
| ZNF512B | T318 | -0.20 | -0.25 | -0.09 | 0.41 | -0.15 | 0.16 | 0.136 |
| ZNF512B | S665 | -0.09 | -0.19 | -0.18 | 0.14 | -0.08 | 0.33 | 0.078 |
| ZNF512B | S63 | -0.16 | 0.03 | -0.33 | 0.19 | 0.20 | 0.01 | 0.079 |
| ZNF512B | T852 | -0.01 | -0.03 | -0.05 | 0.01 | 0.08 | 0.00 | 0.096 |
| ZNF513 | S96 | -0.28 | -0.24 | -0.10 | 0.08 | 0.03 | 0.41 | 0.046 |
| ZNF516 | S116 | 0.22 | 0.27 | 0.03 | -0.23 | -0.24 | -0.13 | 0.01 |
| ZNF518A | S655 | -0.27 | 0.04 | 0.01 | 0.16 | -0.06 | 0.08 | 0.315 |
| ZNF524 | S193 | 0.05 | -0.27 | 0.14 | -0.13 | 0.20 | -0.04 | 0.855 |
| ZNF532 | S314 | 0.05 | -0.17 | -0.11 | 0.17 | -0.21 | 0.21 | 0.428 |
| ZNF532 | S/T | -0.11 | 0.05 | -0.13 | 0.00 | -0.21 | 0.34 | 0.546 |
| ZNF551 | S126 | 0.08 | 0.04 | 0.06 | -0.07 | -0.07 | -0.06 | 4E-04 |
| ZNF574 | S717 | -0.06 | 0.08 | -0.23 | 0.19 | -0.12 | 0.09 | 0.374 |
| ZNF574 | S717; S728 | -0.42 | 0.07 | 0.34 | -0.24 | -0.06 | 0.18 | 0.894 |
| ZNF580 | S | -0.05 | -0.03 | 0.15 | -0.05 | -0.02 | -0.01 | 0.475 |
| ZNF580 | S15 | 0.13 | 0.07 | 0.08 | -0.24 | 0.02 | -0.10 | 0.059 |
| ZNF592 | S766 | -0.79 | -0.77 | 0.13 | 0.51 | -0.25 | 0.56 | 0.132 |
| ZNF592 | S348 | -0.17 | -0.06 | -0.28 | 0.29 | 0.05 | 0.10 | 0.029 |
| ZNF592 | S1205 | -0.38 | -0.14 | 0.08 | -0.37 | 0.15 | 0.47 | 0.459 |

| | | | | | | | | |
|---|---|---|---|---|---|---|---|---|
| ZNF592 | S1227 | -0.24 | -0.22 | 0.09 | 0.28 | -0.21 | 0.20 | 0.312 |
| ZNF592 | S368 | -0.13 | 0.01 | 0.06 | -0.07 | 0.24 | -0.14 | 0.861 |
| ZNF592 | S1089 | 0.13 | -0.06 | -0.02 | 0.00 | 0.13 | -0.21 | 0.732 |
| ZNF592 | S1264 | 0.04 | 0.15 | -0.05 | -0.10 | -0.12 | 0.05 | 0.252 |
| ZNF592 | T1024 | -0.07 | 0.10 | 0.19 | -0.22 | -0.11 | 0.07 | 0.235 |
| ZNF592 | S/T | 0.22 | -0.22 | 0.21 | 0.06 | -0.38 | 0.02 | 0.448 |
| ZNF592 | S1264 | 0.25 | 0.20 | 0.21 | -0.22 | -0.16 | -0.40 | 0.003 |
| ZNF593 | S/T | -0.34 | 0.16 | 0.11 | 0.15 | -0.25 | 0.09 | 0.93 |
| ZNF593 | S/T | -0.08 | 0.04 | 0.15 | -0.76 | 0.19 | 0.26 | 0.702 |
| ZNF597 | S326 | 0.29 | 0.21 | -0.03 | -0.27 | 0.01 | -0.32 | 0.07 |
| ZNF609 | S1285 | -0.19 | -0.59 | -0.21 | 0.24 | 0.25 | 0.30 | 0.011 |
| ZNF609 | S626 | -0.06 | 0.16 | -0.18 | 0.12 | -0.30 | 0.19 | 0.876 |
| ZNF609 | S626 | -0.06 | 0.16 | -0.18 | 0.12 | -0.30 | 0.19 | 0.876 |
| ZNF609 | T746 | -0.02 | 0.09 | -0.12 | -0.03 | -0.17 | 0.21 | 0.889 |
| ZNF609 | S758 | 0.21 | 0.04 | -0.24 | -0.08 | -0.05 | 0.09 | 0.891 |
| ZNF609 | S358 | 0.05 | 0.10 | -0.01 | 0.07 | -0.05 | -0.19 | 0.269 |
| ZNF609 | S491 | 0.05 | 0.04 | 0.12 | -0.25 | 0.24 | -0.26 | 0.4 |
| ZNF609 | S576 | 0.21 | 0.14 | 0.14 | -0.13 | 0.04 | -0.51 | 0.091 |
| ZNF622 | S143 | 0.03 | 0.01 | 0.00 | 0.12 | 0.10 | -0.31 | 0.765 |
| ZNF622 | S143 | -0.26 | -0.05 | 0.45 | -0.13 | 0.13 | -0.27 | 0.599 |
| ZNF638 | S1667 | -0.23 | -0.14 | -0.25 | 0.07 | 0.19 | 0.27 | 0.005 |
| ZNF638 | S885 | -0.18 | -0.06 | -0.22 | -0.03 | -0.13 | 0.50 | 0.263 |
| ZNF638 | S1401 | 0.02 | -0.07 | -0.17 | 0.13 | -0.12 | 0.19 | 0.267 |
| ZNF638 | S605 | -0.02 | -0.20 | -0.03 | 0.36 | -0.41 | 0.18 | 0.634 |
| ZNF638 | S585 | -0.13 | -0.05 | 0.12 | -0.24 | 0.09 | 0.17 | 0.857 |
| ZNF638 | S1913 | -0.29 | 0.15 | 0.09 | -0.05 | 0.04 | 0.03 | 0.869 |
| ZNF638 | S128 | -0.03 | 0.11 | 0.07 | -0.18 | 0.03 | -0.02 | 0.227 |
| ZNF638 | S128 | 0.06 | 0.15 | 0.02 | -0.33 | 0.10 | -0.05 | 0.263 |
| ZNF638 | S1100 | 0.13 | 0.11 | 0.05 | -0.01 | -0.07 | -0.24 | 0.051 |
| ZNF638 | S1100 | 0.39 | 0.23 | 0.51 | -0.11 | -0.82 | -0.73 | 0.017 |
| ZNF639 | S200 | -0.08 | -0.37 | 0.22 | 0.00 | 0.09 | 0.07 | 0.483 |
| ZNF641 | S426 | -0.33 | 0.23 | 0.16 | -0.39 | 0.26 | -0.07 | 0.762 |
| ZNF641 | S191 | -0.02 | 0.38 | 0.15 | -0.21 | -0.14 | -0.26 | 0.034 |
| ZNF644 | S1189 | 0.04 | -0.03 | 0.05 | -0.08 | -0.20 | 0.19 | 0.693 |
| ZNF644 | S1000 | 0.00 | 0.16 | 0.05 | -0.06 | 0.09 | -0.28 | 0.252 |
| ZNF644 | S1189 | 0.20 | 0.14 | 0.34 | -0.31 | -0.14 | -0.39 | 0.006 |
| ZNF646 | S725; S734 | 0.04 | -0.51 | 0.45 | 0.23 | 0.05 | -0.52 | 0.848 |
| ZNF646 | S725; S734 | 0.04 | -0.51 | 0.45 | 0.23 | 0.05 | -0.52 | 0.848 |
| ZNF646 | S725; S734 | 0.04 | -0.51 | 0.45 | 0.23 | 0.05 | -0.52 | 0.848 |
| ZNF646 | S930 | 0.13 | -0.15 | 0.15 | 0.03 | -0.28 | 0.08 | 0.544 |
| ZNF646 | S612 | 0.05 | 0.12 | 0.13 | -0.09 | 0.07 | -0.33 | 0.139 |
| ZNF646 | S | 0.35 | 0.06 | -0.11 | -0.29 | 0.03 | -0.13 | 0.233 |
| ZNF646 | S | 0.34 | 0.19 | -0.10 | -0.37 | 0.02 | -0.19 | 0.132 |
| ZNF649 | S474 | 0.34 | 0.10 | 0.37 | 0.01 | -0.47 | -0.63 | 0.04 |
| ZNF652 | S204 | -0.11 | -0.03 | -0.01 | 0.10 | 0.01 | 0.03 | 0.081 |
| ZNF652 | S57 | 0.14 | 0.04 | -0.15 | -0.05 | 0.09 | -0.09 | 0.797 |
| ZNF655 | S | 0.12 | -0.01 | 0.13 | 0.33 | -0.02 | -0.79 | 0.518 |
| ZNF658 | T821 | -0.33 | -0.48 | 0.11 | -0.12 | 0.41 | 0.21 | 0.168 |

| | | | | | | | | |
|---|---|---|---|---|---|---|---|---|
| **ZNF687** | **S271** | -0.52 | -0.33 | -0.70 | 0.37 | -0.22 | 0.80 | 0.056 |
| **ZNF687** | **S** | -0.26 | -0.41 | -0.67 | 0.39 | -0.21 | 0.68 | 0.063 |
| **ZNF687** | **S271** | -0.32 | -0.41 | -0.58 | 0.40 | -0.21 | 0.67 | 0.056 |
| **ZNF687** | **S1106** | -0.14 | -0.13 | -0.18 | 0.09 | 0.01 | 0.30 | 0.031 |
| **ZNF687** | **S1085** | -0.04 | 0.01 | -0.03 | -0.09 | -0.04 | 0.18 | 0.67 |
| **ZNF687** | **S266; S271** | 0.05 | -0.04 | -0.07 | -0.02 | 0.07 | 0.00 | 0.438 |
| **ZNF687** | **S433** | -0.05 | 0.01 | 0.00 | 0.07 | -0.18 | 0.14 | 0.844 |
| **ZNF687** | **S/T** | 0.00 | 0.02 | -0.06 | 0.05 | -0.17 | 0.14 | 0.846 |
| **ZNF687** | **S1211** | 0.03 | 0.03 | -0.05 | -0.20 | 0.05 | 0.12 | 0.904 |
| **ZNF687** | **S253** | 0.09 | 0.02 | -0.09 | 0.05 | -0.12 | 0.04 | 0.815 |
| **ZNF687** | **S1057** | -0.09 | -0.04 | 0.21 | -0.17 | 0.03 | 0.02 | 0.604 |
| **ZNF687** | **S140; S/T** | 0.36 | -0.25 | -0.05 | -0.38 | 0.13 | 0.06 | 0.75 |
| **ZNF687** | **S374** | 0.11 | 0.00 | 0.00 | -0.07 | 0.00 | -0.05 | 0.131 |
| **ZNF687** | **S266; S271; S275** | 0.11 | -0.01 | 0.05 | 0.00 | -0.08 | -0.09 | 0.07 |
| **ZNF687** | **S/T** | 0.15 | 0.23 | -0.22 | -0.47 | 0.02 | 0.16 | 0.566 |
| **ZNF687** | **S1191** | 0.11 | 0.13 | 0.09 | -0.18 | -0.01 | -0.18 | 0.016 |
| **ZNF687** | **S183** | 0.28 | 0.05 | 0.11 | -0.14 | -0.08 | -0.28 | 0.027 |
| **ZNF687** | **S242; S** | 0.20 | 0.07 | 0.20 | -0.18 | -0.22 | -0.12 | 0.003 |
| **ZNF687** | **S1146** | 0.36 | 0.00 | 0.11 | -0.32 | -0.18 | -0.07 | 0.054 |
| **ZNF687** | **S/T** | 0.17 | 0.06 | 0.36 | -0.43 | -0.14 | -0.15 | 0.026 |
| **ZNF687** | **S/T** | 0.29 | 0.16 | 0.19 | -0.38 | -0.25 | -0.14 | 0.004 |
| **ZNF687** | **S1082; S1083** | 0.23 | 0.14 | 0.27 | -0.22 | -0.14 | -0.41 | 0.006 |
| **ZNF687** | **S1057** | 0.10 | 0.31 | 0.34 | -0.41 | 0.02 | -0.61 | 0.043 |
| **ZNF687** | **S227; S** | 0.41 | 0.27 | 0.14 | -0.44 | -0.12 | -0.48 | 0.012 |
| **ZNF691** | **S39** | 0.04 | 0.01 | 0.02 | -0.01 | -0.08 | 0.02 | 0.209 |
| **ZNF691** | **S/T** | 0.05 | 0.05 | 0.10 | -0.19 | 0.02 | -0.06 | 0.09 |
| **ZNF692** | **S231** | 0.21 | 0.05 | -0.09 | -0.16 | 0.04 | -0.08 | 0.313 |
| **ZNF711** | **T221** | -0.19 | 0.01 | 0.06 | 0.04 | -0.20 | 0.24 | 0.662 |
| **ZNF711** | **S459** | 0.01 | 0.00 | 0.07 | -0.01 | -0.14 | 0.07 | 0.463 |
| **ZNF711** | **S/T/Y** | -0.02 | 0.11 | -0.01 | 0.08 | 0.05 | -0.24 | 0.596 |
| **ZNF740** | **S72** | -0.07 | -0.01 | -0.05 | -0.08 | 0.10 | 0.10 | 0.228 |
| **ZNF740** | **S44** | 0.00 | 0.02 | -0.05 | 0.03 | -0.04 | 0.04 | 0.625 |
| **ZNF768** | **S151** | -0.07 | 0.06 | -0.12 | 0.16 | 0.12 | -0.18 | 0.552 |
| **ZNF768** | **S97** | -0.07 | 0.05 | 0.00 | 0.02 | -0.11 | 0.10 | 0.836 |
| **ZNF768** | **S139** | 0.01 | -0.07 | 0.04 | -0.15 | 0.27 | -0.14 | 0.992 |
| **ZNF768** | **S90; S97** | 0.09 | 0.13 | -0.09 | 0.12 | -0.07 | -0.21 | 0.444 |
| **ZNF768** | **S23** | 0.09 | 0.28 | -0.01 | -0.07 | -0.25 | -0.10 | 0.059 |
| **ZNF768** | **S125** | 0.14 | 0.13 | 0.11 | -0.19 | -0.05 | -0.17 | 0.004 |
| **ZNF768** | **S160** | 0.10 | 0.09 | 0.22 | -0.04 | -0.04 | -0.41 | 0.089 |
| **ZNF768** | **S160** | 0.10 | 0.31 | 0.13 | -0.29 | -0.09 | -0.27 | 0.012 |
| **ZNF768** | **S83** | 0.16 | 0.23 | 0.26 | -0.24 | -0.12 | -0.40 | 0.006 |
| **ZNF777** | **S47** | -0.19 | 0.10 | 0.08 | 0.36 | -0.11 | -0.35 | 0.904 |
| **ZNF777** | **S7** | 0.12 | 0.06 | -0.01 | 0.04 | 0.02 | -0.27 | 0.293 |
| **ZNF791** | **S85** | 0.08 | -0.15 | -0.15 | 0.25 | -0.19 | 0.10 | 0.45 |
| **ZNF791** | **S85** | -0.13 | 0.05 | -0.08 | 0.35 | -0.07 | -0.18 | 0.634 |
| **ZNF800** | **S595** | 0.00 | -0.12 | -0.17 | 0.13 | -0.09 | 0.22 | 0.154 |
| **ZNF800** | **T157; T166** | -0.19 | -0.23 | 0.28 | 0.28 | 0.04 | -0.31 | 0.846 |
| **ZNF800** | **T157; T166** | -0.19 | -0.23 | 0.28 | 0.28 | 0.04 | -0.31 | 0.846 |

| | | | | | | | | |
|---|---|---|---|---|---|---|---|---|
| **ZNF800** | **S426** | 0.04 | -0.10 | 0.00 | 0.04 | -0.11 | 0.12 | 0.661 |
| **ZNF800** | **S419; S422; S426** | 0.05 | 0.02 | -0.06 | 0.03 | -0.01 | -0.04 | 0.823 |
| **ZNF800** | **S160; S/T** | -0.11 | 0.14 | 0.00 | -0.25 | 0.13 | 0.06 | 0.847 |
| **ZNF800** | **S422; S426** | -0.02 | -0.06 | 0.12 | 0.14 | -0.14 | -0.06 | 0.779 |
| **ZNF800** | **S647** | 0.00 | -0.07 | 0.18 | -0.05 | -0.02 | -0.05 | 0.337 |
| **ZNF800** | **S/T** | 0.13 | 0.08 | 0.17 | -0.13 | -0.03 | -0.27 | 0.023 |
| **ZNF800** | **S426** | 0.22 | -0.04 | 0.19 | -0.10 | -0.39 | 0.03 | 0.133 |
| **ZNF800** | **S336** | 0.07 | 0.36 | -0.03 | -0.28 | 0.19 | -0.48 | 0.229 |
| **ZNF800** | **S457** | 0.21 | 0.22 | 0.11 | -0.25 | 0.11 | -0.55 | 0.101 |
| **ZNF800** | **S595** | 0.29 | 0.28 | 0.23 | -0.15 | -0.27 | -0.60 | 0.011 |
| **ZNF830** | **S351** | -0.10 | 0.04 | -0.25 | 0.13 | -0.19 | 0.29 | 0.335 |
| **ZNF830** | **S351** | 0.05 | -0.02 | 0.01 | 0.03 | -0.08 | 0.01 | 0.535 |
| **ZNF830** | **S351** | 0.18 | 0.15 | 0.07 | -0.10 | 0.00 | -0.38 | 0.064 |
| **ZNF837** | **S351** | 0.13 | 0.06 | -0.09 | 0.06 | -0.17 | -0.01 | 0.5 |
| **ZNF84** | **S78** | -0.31 | -0.22 | -0.16 | 0.06 | 0.25 | 0.28 | 0.007 |
| **ZNF852** | **S145** | -0.21 | -0.06 | 0.16 | 0.04 | -0.15 | 0.17 | 0.723 |
| **ZNF875** | **S219; S220** | -0.25 | -0.01 | -0.07 | -0.05 | -0.03 | 0.35 | 0.249 |
| **ZNHIT6** | **S11** | 0.19 | 0.11 | -0.34 | -0.04 | 0.02 | 0.00 | 0.969 |
| **ZNRF2** | **S19** | 0.01 | -0.28 | 0.11 | 0.21 | -0.33 | 0.19 | 0.727 |
| **ZNRF2** | **S82** | 0.11 | -0.03 | -0.10 | 0.08 | -0.22 | 0.12 | 0.994 |
| **ZNRF2** | **S135** | -0.04 | 0.06 | 0.22 | 0.06 | -0.12 | -0.21 | 0.197 |
| **ZRANB2** | **S153** | 0.35 | 0.00 | -0.07 | -0.10 | -0.12 | -0.12 | 0.193 |
| **ZRANB2** | **S120** | 0.40 | 0.04 | 0.05 | -0.30 | 0.02 | -0.34 | 0.085 |
| **ZRANB2** | **S188** | 0.42 | 0.07 | 0.18 | -0.36 | -0.08 | -0.42 | 0.025 |
| **ZRANB2** | **S188** | 0.45 | 0.11 | 0.14 | -0.33 | -0.05 | -0.52 | 0.038 |
| **ZRANB2** | **S280** | 0.33 | 0.25 | 0.26 | -0.61 | 0.05 | -0.58 | 0.039 |
| **ZRSR2** | **S349** | 0.09 | 0.02 | -0.03 | 0.04 | -0.06 | -0.07 | 0.362 |
| **ZRSR2** | **S384** | 0.14 | 0.10 | -0.03 | 0.08 | -0.10 | -0.22 | 0.228 |
| **ZRSR2** | **S384** | 0.32 | 0.18 | 0.21 | -0.01 | -0.29 | -0.62 | 0.039 |
| **ZRSR2P1** | **S389** | -0.01 | 0.02 | -0.14 | 0.05 | 0.36 | -0.38 | 0.821 |
| **ZRSR2P1** | **S475** | 0.04 | -0.08 | 0.00 | 0.28 | 0.05 | -0.36 | 0.993 |
| **ZSCAN21** | **S157** | -0.10 | -0.18 | -0.41 | 0.30 | -0.29 | 0.46 | 0.192 |
| **ZSCAN21** | **S157** | -0.21 | -0.08 | 0.00 | 0.14 | -0.11 | 0.22 | 0.194 |
| **ZSCAN21** | **S/T** | 0.06 | -0.04 | -0.20 | 0.04 | 0.12 | 0.01 | 0.232 |
| **ZSCAN21** | **T136** | 0.08 | -0.02 | -0.22 | 0.21 | 0.05 | -0.16 | 0.578 |
| **ZSCAN21** | **S208** | 0.15 | 0.11 | -0.05 | -0.08 | -0.12 | -0.03 | 0.092 |
| **ZSCAN29** | **S153** | -0.05 | 0.03 | -0.12 | 0.11 | -0.01 | 0.03 | 0.17 |
| **ZSCAN29** | **S153** | -0.06 | 0.09 | -0.06 | 0.09 | -0.01 | -0.05 | 0.818 |
| **ZSCAN29** | **S153** | -0.04 | 0.08 | -0.08 | 0.27 | -0.31 | 0.03 | 0.961 |
| **ZSCAN29** | **S153** | 0.10 | 0.05 | -0.09 | 0.10 | -0.05 | -0.14 | 0.602 |
| **ZSCAN29** | **S/T** | -0.83 | 0.67 | 0.07 | -0.46 | -0.21 | 0.27 | 0.842 |
| **ZSCAN29** | **S537** | -0.83 | 0.67 | 0.07 | -0.46 | -0.21 | 0.27 | 0.842 |
| **ZSCAN29** | **S153** | 0.23 | -0.19 | 0.12 | 0.42 | -0.45 | -0.34 | 0.587 |
| **ZSCAN29** | **S405** | 0.34 | -0.21 | 0.08 | -0.18 | -0.28 | 0.14 | 0.424 |
| **ZSCAN29** | **T533; S537** | -0.18 | 0.37 | 0.52 | -0.27 | -0.17 | -0.56 | 0.079 |
| **ZSCAN5A** | **S296** | 0.05 | -0.06 | 0.03 | -0.02 | -0.11 | 0.10 | 0.771 |
| **ZSCAN5A** | **S245** | 0.30 | 0.13 | 0.16 | -0.33 | -0.19 | -0.17 | 0.004 |
| **ZSWIM3** | **S109** | -0.04 | -0.10 | -0.08 | 0.12 | -0.06 | 0.13 | 0.096 |

| | | | | | | | | |
|---|---|---|---|---|---|---|---|---|
| **ZSWIM8** | **S53** | -0.13 | -0.24 | -0.05 | -0.08 | 0.08 | 0.35 | 0.135 |
| **ZSWIM8** | **S567** | -0.04 | -0.05 | -0.01 | 0.10 | 0.07 | -0.07 | 0.274 |
| **ZZEF1** | **S1518** | 0.12 | 0.06 | -0.02 | -0.15 | -0.05 | 0.02 | 0.148 |
| **ZZZ3** | **S113** | 0.28 | 0.12 | 0.09 | 0.03 | -0.08 | -0.58 | 0.133 |
| **ZZZ3** | **S113** | 0.04 | 0.07 | 0.42 | -0.16 | -0.10 | -0.40 | 0.059 |

| Log2FC ATRi/UT |
| --- |
| 0.11 |
| 0.11 |
| -0.04 |
| -0.04 |
| -0.06 |
| -0.81 |
| -1.21 |
| 0.27 |
| -0.25 |
| 0.35 |
| 0.21 |
| -0.04 |
| -0.12 |
| -0.21 |
| 0.05 |
| 0.00 |
| -0.02 |
| -0.15 |
| -0.26 |
| -0.78 |
| -0.39 |
| -0.65 |
| -0.46 |
| 0.07 |
| -0.17 |
| -0.45 |
| 0.25 |
| -0.16 |
| -0.55 |
| -0.82 |
| -1.02 |
| -0.33 |
| -0.85 |
| 0.07 |
| -0.11 |
| -0.31 |
| 0.49 |
| 0.20 |

| |
|---|
| 0.19 |
| -0.05 |
| -0.08 |
| -0.10 |
| -0.12 |
| -0.32 |
| -0.33 |
| 0.38 |
| 0.30 |
| 0.02 |
| 0.01 |
| -0.08 |
| -0.23 |
| -0.32 |
| -0.37 |
| -0.48 |
| 0.56 |
| 0.45 |
| 0.09 |
| 0.06 |
| 0.22 |
| -0.46 |
| 0.18 |
| -0.08 |
| 0.00 |
| -0.01 |
| -0.15 |
| -0.28 |
| -0.28 |
| 1.47 |
| 0.51 |
| 0.44 |
| 0.21 |
| -0.08 |
| 1.55 |
| 0.73 |
| 0.54 |
| 0.30 |
| 0.25 |
| 0.23 |
| 0.13 |
| 0.11 |
| 0.03 |
| -0.06 |
| -0.09 |
| -0.09 |
| -0.14 |
| -0.19 |

-0.21
-0.22
-0.25
-0.26
-0.28
-0.28
-0.30
-0.33
-0.33
-0.33
-0.34
-0.36
-0.38
-0.38
-0.42
-0.47
-0.49
-0.54
-0.56
-0.66
-0.69
-0.72
-0.80
-0.30
-0.72
0.32
0.27
0.12
0.01
-0.31
-0.63
-0.17
0.21
0.45
0.22
-0.18
0.26
-0.79
0.17
-0.02
-0.46
-0.28
-0.06
0.42
0.15
0.11
-0.06
-0.06

| |
|---|
| -0.09 |
| -0.11 |
| -0.18 |
| -0.19 |
| -0.19 |
| -0.23 |
| -0.23 |
| -0.26 |
| -0.26 |
| -0.27 |
| -0.27 |
| -0.33 |
| -0.69 |
| -0.91 |
| -0.93 |
| -0.21 |
| -0.15 |
| -0.17 |
| -0.25 |
| -0.25 |
| -0.33 |
| 0.70 |
| 0.38 |
| 0.16 |
| 0.13 |
| 0.11 |
| 0.10 |
| -0.05 |
| 0.42 |
| -0.61 |
| 0.33 |
| 0.21 |
| 0.03 |
| -0.02 |
| -0.05 |
| -0.07 |
| -0.28 |
| -0.30 |
| -0.36 |
| -0.36 |
| -0.40 |
| -0.41 |
| -0.44 |
| -0.46 |
| -0.57 |
| 0.22 |
| 0.02 |
| -0.14 |

| |
|---|
| -0.42 |
| 0.01 |
| 0.68 |
| 0.57 |
| 0.57 |
| 0.39 |
| 0.33 |
| 0.31 |
| 0.29 |
| 0.27 |
| 0.25 |
| 0.21 |
| 0.16 |
| 0.15 |
| 0.15 |
| 0.04 |
| 0.04 |
| 0.03 |
| 0.01 |
| -0.02 |
| -0.03 |
| -0.07 |
| -0.10 |
| -0.15 |
| -0.15 |
| -0.16 |
| -0.26 |
| -0.31 |
| -0.34 |
| -0.35 |
| -0.60 |
| -0.70 |
| -0.72 |
| -0.88 |
| 0.19 |
| 0.19 |
| -0.01 |
| 0.47 |
| 0.38 |
| 0.13 |
| 0.13 |
| 0.12 |
| 0.08 |
| 0.08 |
| 0.08 |
| -0.01 |
| -0.18 |
| -0.20 |

| |
|---|
| -0.34 |
| 1.47 |
| 0.80 |
| 0.79 |
| 0.68 |
| 0.65 |
| 0.63 |
| 0.62 |
| 0.60 |
| 0.53 |
| 0.52 |
| 0.51 |
| 0.50 |
| 0.49 |
| 0.42 |
| 0.40 |
| 0.39 |
| 0.39 |
| 0.38 |
| 0.38 |
| 0.37 |
| 0.32 |
| 0.31 |
| 0.26 |
| 0.24 |
| 0.23 |
| 0.22 |
| 0.21 |
| 0.20 |
| 0.19 |
| 0.19 |
| 0.18 |
| 0.15 |
| 0.14 |
| 0.12 |
| 0.11 |
| 0.07 |
| 0.06 |
| 0.06 |
| 0.04 |
| 0.03 |
| 0.02 |
| 0.02 |
| 0.01 |
| 0.00 |
| -0.01 |
| -0.02 |
| -0.09 |

-0.10
-0.10
-0.11
-0.17
-0.17
-0.21
-0.22
-0.24
-0.41
-0.41
-0.43
-0.51
-0.52
-0.55
-0.83
0.14
0.06
0.41
-0.27
-0.33
0.25
0.49
0.35
0.32
0.29
0.28
0.17
0.12
-0.05
-0.13
-0.27
0.17
0.16
0.08
0.03
-0.04
-0.29
0.08
0.04
-0.12
-0.20
-0.45
0.56
0.12
-0.12
-0.13
-0.25
-0.40

| |
|---|
| -0.48 |
| 0.06 |
| 0.01 |
| 0.19 |
| 0.17 |
| 0.16 |
| 0.04 |
| -0.02 |
| -0.31 |
| -0.41 |
| 0.47 |
| -0.06 |
| -0.32 |
| 0.04 |
| 0.28 |
| -0.29 |
| -0.30 |
| 0.45 |
| 0.12 |
| 0.10 |
| -0.10 |
| -0.14 |
| -0.16 |
| -0.16 |
| -0.21 |
| -0.31 |
| -0.42 |
| 0.11 |
| 0.21 |
| 0.09 |
| -0.46 |
| -0.07 |
| 0.31 |
| 0.20 |
| 0.16 |
| -0.05 |
| -0.15 |
| -0.15 |
| -0.59 |
| -0.78 |
| 0.40 |
| 0.22 |
| -0.22 |
| -0.13 |
| -0.19 |
| -0.05 |
| 0.13 |
| -0.15 |

| |
|---|
| -0.04 |
| -0.35 |
| 0.12 |
| -0.39 |
| 0.31 |
| 0.12 |
| -0.27 |
| -0.30 |
| -0.03 |
| -0.25 |
| -0.52 |
| -0.53 |
| 0.58 |
| 0.23 |
| 0.00 |
| 0.40 |
| 0.15 |
| -0.27 |
| -0.30 |
| -0.44 |
| -0.07 |
| 0.32 |
| 0.08 |
| -0.26 |
| 0.11 |
| 0.06 |
| -0.05 |
| -0.11 |
| 0.22 |
| 0.12 |
| -0.23 |
| -0.23 |
| 0.00 |
| -0.46 |
| -0.46 |
| 0.25 |
| 0.13 |
| 0.13 |
| -0.03 |
| -0.31 |
| -0.69 |
| 0.15 |
| -0.01 |
| -0.10 |
| -0.39 |
| -0.53 |
| 0.02 |
| -0.09 |

0.41
0.02
0.00
-0.09
0.44
0.21
0.04
0.39
0.22
0.19
0.05
-0.08
-0.07
0.18
0.17
0.17
0.17
0.14
0.13
0.05
0.05
0.01
-0.04
-0.05
-0.09
-0.10
-0.22
0.10
-0.03
0.16
-0.04
0.61
0.29
0.29
0.25
0.22
0.69
-0.05
-0.04
-0.28
-0.64
-0.01
-0.02
0.11
0.44
0.32
0.14
0.14

-0.22
-0.06
-0.12
0.29
0.27
0.15
-0.28
0.08
-0.05
0.13
0.06
0.02
-0.01
-0.01
-0.02
-0.12
0.34
0.22
-0.24
-0.35
0.28
0.09
0.05
-0.47
0.32
0.17
0.12
0.02
0.01
-0.03
-0.07
-0.39
0.11
0.09
0.11
-0.06
-0.19
0.62
0.52
0.37
0.27
0.06
-0.07
-0.08
-0.10
-0.47
0.16
0.29

| |
|---|
| 0.01 |
| 0.00 |
| -0.14 |
| -0.23 |
| 0.19 |
| 0.11 |
| 0.11 |
| -0.01 |
| -0.24 |
| -0.26 |
| -0.40 |
| 0.23 |
| 0.16 |
| 0.12 |
| 0.12 |
| -0.04 |
| -0.08 |
| -0.13 |
| -0.23 |
| 0.04 |
| -0.61 |
| 0.06 |
| -0.06 |
| -0.12 |
| -0.24 |
| 0.04 |
| -0.10 |
| -0.18 |
| -0.28 |
| 0.21 |
| -0.29 |
| 0.06 |
| 0.03 |
| -0.07 |
| -0.26 |
| -0.34 |
| 0.07 |
| -0.01 |
| -0.04 |
| -0.16 |
| -0.34 |
| -0.38 |
| -0.32 |
| -0.38 |
| 0.20 |
| 0.26 |
| 0.02 |
| -0.43 |

| |
|---|
| 0.26 |
| 0.27 |
| 0.18 |
| -0.11 |
| 0.16 |
| 0.08 |
| -0.08 |
| -0.08 |
| -0.13 |
| 0.15 |
| 0.10 |
| -0.06 |
| -0.07 |
| -0.62 |
| -0.64 |
| 1.40 |
| 0.64 |
| 0.52 |
| 0.35 |
| 0.32 |
| 0.26 |
| 0.17 |
| 0.17 |
| 0.11 |
| 0.02 |
| -0.04 |
| -0.07 |
| -0.10 |
| -0.20 |
| -0.31 |
| 0.10 |
| -0.01 |
| 0.61 |
| 0.48 |
| 0.40 |
| 0.36 |
| 0.36 |
| 0.11 |
| 0.07 |
| 0.03 |
| 0.02 |
| 0.01 |
| -0.02 |
| -0.03 |
| -0.05 |
| -0.08 |
| -0.11 |
| -0.15 |

-0.18
-0.20
-0.25
-0.25
-0.28
-0.28
-0.30
-0.32
-0.32
-0.42
-0.46
-0.56
-0.63
0.66
0.44
0.38
0.37
0.29
0.25
0.24
0.24
0.19
0.17
0.16
0.12
-0.05
-0.05
-0.09
-0.10
-0.10
-0.11
-0.12
-0.41
-0.53
0.34
0.09
-0.05
-0.05
-0.07
-0.12
-0.16
-0.35
-0.41
-0.49
-0.65
0.40
0.25
0.09

-0.12
-0.21
-0.30
-0.40
-0.14
0.49
0.42
0.30
0.10
-0.34
-0.37
3.38
0.22
0.18
-0.09
-0.15
0.37
0.25
0.05
-0.39
-0.03
0.24
0.16
0.12
0.00
-0.23
-0.27
-0.29
-0.38
-0.21
0.03
-0.51
-0.29
-0.14
-0.21
-0.21
-0.45
-0.49
0.07
-0.07
-0.30
-0.67
0.42
0.11
-0.03
-0.03
-0.18
0.22

| |
|---|
| 0.12 |
| 0.06 |
| 0.66 |
| 0.50 |
| 0.28 |
| 0.22 |
| 0.22 |
| 0.19 |
| 0.19 |
| 0.17 |
| 0.14 |
| 0.09 |
| 0.09 |
| 0.08 |
| 0.07 |
| 0.06 |
| 0.06 |
| 0.05 |
| 0.04 |
| 0.04 |
| 0.04 |
| 0.03 |
| 0.02 |
| 0.02 |
| 0.01 |
| 0.01 |
| 0.00 |
| 0.00 |
| 0.00 |
| -0.01 |
| -0.03 |
| -0.06 |
| -0.08 |
| -0.09 |
| -0.09 |
| -0.11 |
| -0.14 |
| -0.14 |
| -0.18 |
| -0.19 |
| -0.21 |
| -0.24 |
| -0.24 |
| -0.27 |
| -0.27 |
| -0.31 |
| -0.35 |
| -0.39 |

| |
|---|
| -0.40 |
| -0.47 |
| -0.56 |
| -0.57 |
| -0.64 |
| -1.14 |
| 0.26 |
| 0.17 |
| 0.15 |
| 0.06 |
| -0.19 |
| -0.19 |
| -0.32 |
| -0.38 |
| 0.31 |
| 0.21 |
| 0.16 |
| -0.04 |
| -0.11 |
| -0.14 |
| -0.16 |
| -0.41 |
| -0.44 |
| -0.08 |
| 0.04 |
| -0.07 |
| 0.08 |
| -0.05 |
| 0.40 |
| 0.14 |
| 0.01 |
| -0.22 |
| -0.23 |
| 0.20 |
| -0.03 |
| -0.16 |
| 0.26 |
| -0.42 |
| -0.84 |
| 0.07 |
| 0.06 |
| -0.03 |
| -0.06 |
| -0.21 |
| -0.44 |
| 0.21 |
| -0.10 |
| 0.31 |

| |
|---|
| 0.23 |
| 0.13 |
| -0.06 |
| -0.28 |
| -0.47 |
| 0.59 |
| -0.12 |
| 0.06 |
| -0.21 |
| -0.25 |
| 0.05 |
| 0.01 |
| -0.38 |
| 1.16 |
| 0.90 |
| 0.64 |
| 0.40 |
| 0.37 |
| 0.21 |
| 0.16 |
| 0.14 |
| 0.03 |
| -0.08 |
| -0.13 |
| -0.16 |
| -0.37 |
| -0.61 |
| 0.43 |
| 0.43 |
| 0.37 |
| 0.22 |
| 0.22 |
| 0.21 |
| 0.15 |
| 0.12 |
| 0.11 |
| 0.05 |
| 0.04 |
| 0.02 |
| 0.01 |
| -0.01 |
| -0.03 |
| -0.03 |
| -0.04 |
| -0.04 |
| -0.05 |
| -0.07 |
| -0.11 |

| |
|---|
| -0.17 |
| -0.27 |
| -0.28 |
| -0.30 |
| -0.34 |
| -0.38 |
| -0.41 |
| -0.42 |
| -0.48 |
| -0.48 |
| -0.52 |
| -0.53 |
| -0.72 |
| -0.75 |
| -0.98 |
| -1.11 |
| 0.43 |
| 0.42 |
| 0.30 |
| 0.27 |
| 0.23 |
| 0.16 |
| 0.15 |
| 0.12 |
| 0.09 |
| 0.07 |
| 0.03 |
| 0.03 |
| 0.03 |
| 0.01 |
| 0.01 |
| 0.00 |
| -0.01 |
| -0.12 |
| -0.25 |
| -0.45 |
| -0.54 |
| 0.37 |
| 0.23 |
| 0.05 |
| -0.16 |
| 0.43 |
| 0.03 |
| -0.16 |
| -0.42 |
| -0.61 |
| 0.01 |
| 0.47 |

0.35
-0.22
-0.29
-0.21
-0.42
0.14
0.25
-0.08
-0.11
-0.23
-0.53
-0.54
-0.71
0.35
-0.43
0.27
0.37
0.26
-0.01
-0.07
0.63
0.53
0.27
0.16
0.11
-0.08
0.30
0.18
0.16
1.02
0.21
0.19
0.19
0.16
0.15
0.15
0.13
0.09
0.07
0.06
0.01
0.01
0.00
0.00
-0.01
-0.03
-0.09
-0.12

| |
|---|
| -0.12 |
| -0.13 |
| -0.14 |
| -0.16 |
| -0.17 |
| -0.17 |
| -0.18 |
| -0.19 |
| -0.20 |
| -0.26 |
| -0.27 |
| -0.27 |
| -0.28 |
| -0.28 |
| -0.29 |
| -0.30 |
| -0.32 |
| -0.33 |
| -0.34 |
| -0.35 |
| -0.38 |
| -0.40 |
| -0.41 |
| -0.48 |
| -0.52 |
| -0.72 |
| -0.90 |
| 0.11 |
| 0.00 |
| 0.13 |
| -0.04 |
| -0.07 |
| -0.24 |
| -0.26 |
| -0.28 |
| -0.29 |
| -0.35 |
| -0.37 |
| 0.24 |
| 0.07 |
| 0.00 |
| -0.04 |
| -0.12 |
| -0.16 |
| -0.37 |
| 0.35 |
| 0.29 |
| 0.28 |

| |
|---|
| 0.04 |
| 0.02 |
| -0.37 |
| -0.03 |
| -0.14 |
| 0.00 |
| -0.15 |
| -0.22 |
| 0.31 |
| 0.22 |
| 0.37 |
| 0.01 |
| 0.01 |
| -0.07 |
| -0.10 |
| -0.29 |
| -0.02 |
| 0.26 |
| 0.16 |
| -0.15 |
| -0.17 |
| -0.29 |
| -0.55 |
| -0.59 |
| -0.04 |
| -0.07 |
| -0.38 |
| 0.31 |
| 0.07 |
| -0.13 |
| -0.41 |
| 0.02 |
| -0.05 |
| -0.08 |
| -0.55 |
| 0.23 |
| 0.10 |
| 0.06 |
| -0.62 |
| 1.32 |
| 0.52 |
| 0.04 |
| 0.04 |
| 0.21 |
| 0.06 |
| 0.02 |
| 0.17 |
| 0.10 |

| |
|---|
| -0.12 |
| -0.46 |
| -0.24 |
| -0.22 |
| 2.60 |
| 0.63 |
| 0.53 |
| 0.49 |
| 0.45 |
| 0.39 |
| 0.31 |
| 0.29 |
| 0.29 |
| 0.28 |
| 0.27 |
| 0.18 |
| 0.17 |
| 0.15 |
| 0.14 |
| 0.12 |
| 0.12 |
| 0.03 |
| 0.03 |
| -0.02 |
| -0.02 |
| -0.05 |
| -0.05 |
| -0.06 |
| -0.07 |
| -0.09 |
| -0.16 |
| -0.17 |
| -0.23 |
| -0.26 |
| -0.31 |
| -0.50 |
| -0.54 |
| 0.01 |
| 0.07 |
| 0.70 |
| 0.48 |
| 0.01 |
| -0.04 |
| -0.55 |
| 0.59 |
| 0.28 |
| -0.07 |
| -0.13 |

| |
|---|
| -0.32 |
| -0.38 |
| -0.08 |
| -0.09 |
| -0.28 |
| -0.29 |
| 0.39 |
| 0.04 |
| 0.00 |
| -0.15 |
| -0.17 |
| -0.27 |
| -0.33 |
| -0.43 |
| 0.07 |
| 0.07 |
| 0.02 |
| 0.01 |
| -0.14 |
| -0.39 |
| -0.59 |
| -0.87 |
| 0.25 |
| 0.06 |
| 0.06 |
| -0.16 |
| -0.37 |
| 0.03 |
| 0.12 |
| 0.09 |
| -0.29 |
| 0.22 |
| 0.02 |
| -0.09 |
| -0.29 |
| -0.31 |
| 0.28 |
| 0.07 |
| -0.38 |
| -0.20 |
| -0.25 |
| 0.46 |
| 0.28 |
| 0.18 |
| 0.03 |
| 0.03 |
| -0.04 |
| -0.05 |

-0.07
-0.08
-0.08
-0.14
-0.15
-0.19
-0.20
-0.24
-0.45
0.41
0.21
0.16
0.15
0.08
0.05
0.04
-0.09
-0.12
-0.13
-0.18
-0.26
0.39
0.30
0.16
0.13
0.10
0.06
0.03
0.01
-0.03
-0.05
-0.07
-0.08
-0.08
-0.19
-0.26
-0.31
-0.33
-0.52
-0.62
0.03
-0.13
-0.14
-0.14
-0.22
-0.27
-0.69
-0.30

-0.03
-0.46
0.05
-0.24
-0.10
-0.21
0.34
0.30
0.29
0.16
0.10
0.06
0.04
-0.01
-0.07
-0.09
-0.09
-0.10
-0.13
-0.14
-0.14
-0.15
-0.19
-0.24
-0.24
-0.25
-0.32
-0.38
-0.49
-0.49
-0.52
-0.53
-0.80
-0.82
0.23
0.06
0.06
-0.10
-0.38
-0.03
-0.14
-0.37
-0.62
-0.38
0.54
-0.05
-0.07
-0.12

| |
|---|
| -0.16 |
| -0.45 |
| 0.41 |
| -0.01 |
| 0.76 |
| -0.44 |
| -0.51 |
| 0.19 |
| -0.02 |
| 0.18 |
| -0.12 |
| 0.20 |
| 0.11 |
| 0.13 |
| 0.01 |
| -0.05 |
| -0.09 |
| -0.14 |
| 0.13 |
| 0.09 |
| 0.03 |
| 0.02 |
| -0.01 |
| -0.25 |
| -0.30 |
| -0.44 |
| -0.59 |
| -0.14 |
| -0.23 |
| -0.37 |
| 0.41 |
| 0.75 |
| 0.30 |
| 0.21 |
| -0.02 |
| -0.45 |
| -0.39 |
| 0.20 |
| 0.12 |
| 0.03 |
| 0.17 |
| -0.16 |
| -0.50 |
| -0.55 |
| 0.34 |
| 0.20 |
| 0.14 |
| 0.06 |

-0.04
-0.09
-0.11
-0.18
-0.19
-0.25
1.18
0.84
0.68
0.13
-0.15
-0.15
-0.21
-0.28
0.14
0.02
-0.07
-0.14
-0.02
-0.04
-0.44
-0.33
-0.09
0.60
0.16
0.04
0.00
0.19
0.07
0.01
0.25
0.16
-0.71
-0.80
0.38
-0.02
-0.05
-0.24
-0.10
-0.29
-0.30
-0.39
-0.58
-0.71
0.24
0.15
-0.32
0.04

| |
|---|
| 0.12 |
| 0.12 |
| -0.15 |
| -0.26 |
| -0.45 |
| -0.58 |
| 0.29 |
| 0.16 |
| -0.23 |
| -0.49 |
| -0.46 |
| -0.46 |
| 0.27 |
| 0.15 |
| 0.13 |
| 0.11 |
| 0.10 |
| 0.05 |
| 0.03 |
| 0.01 |
| 0.01 |
| -0.03 |
| -0.05 |
| -0.05 |
| -0.07 |
| -0.08 |
| -0.09 |
| -0.14 |
| -0.18 |
| -0.19 |
| -0.19 |
| -0.21 |
| -0.34 |
| -0.87 |
| -0.28 |
| 0.73 |
| 0.07 |
| 0.05 |
| -0.03 |
| -0.17 |
| 1.69 |
| 0.99 |
| 0.15 |
| -0.04 |
| -0.14 |
| -0.14 |
| -0.26 |
| -0.55 |

| |
|---|
| -0.66 |
| -0.16 |
| -0.28 |
| -0.29 |
| -0.34 |
| -0.38 |
| 0.11 |
| 0.05 |
| -0.20 |
| -0.24 |
| -0.40 |
| 0.45 |
| 0.28 |
| 0.24 |
| 0.08 |
| -0.38 |
| -0.50 |
| 0.27 |
| -0.03 |
| -0.43 |
| -0.57 |
| -0.01 |
| -0.50 |
| 0.43 |
| 0.17 |
| 0.09 |
| 0.08 |
| -0.30 |
| 0.83 |
| 0.37 |
| -0.43 |
| -0.19 |
| -0.97 |
| -0.05 |
| -0.29 |
| 0.00 |
| 0.45 |
| -0.08 |
| -0.36 |
| -0.39 |
| -0.08 |
| -0.32 |
| -0.32 |
| -0.59 |
| -0.59 |
| 0.17 |
| 0.06 |
| 1.41 |

| |
|---|
| 0.41 |
| 0.30 |
| 0.25 |
| 0.21 |
| 0.21 |
| 0.15 |
| 0.09 |
| 0.09 |
| 0.07 |
| 0.07 |
| 0.06 |
| 0.04 |
| -0.01 |
| -0.01 |
| -0.01 |
| -0.03 |
| -0.04 |
| -0.04 |
| -0.05 |
| -0.09 |
| -0.17 |
| -0.17 |
| 0.35 |
| 0.32 |
| 0.28 |
| 0.21 |
| 0.19 |
| 0.16 |
| 0.05 |
| 0.02 |
| 0.00 |
| -0.12 |
| -0.25 |
| 0.16 |
| 0.42 |
| -0.02 |
| 0.31 |
| -0.28 |
| -0.52 |
| 0.15 |
| 0.12 |
| 0.10 |
| -0.02 |
| -0.07 |
| -0.62 |
| 0.70 |
| 0.09 |
| 0.04 |

-0.09
-0.17
-0.36
0.21
0.13
-0.10
-0.13
-0.17
0.38
-0.21
0.31
0.22
0.01
0.38
-0.17
0.16
-0.17
-0.28
-0.11
0.09
-0.14
-0.15
-0.36
-0.57
-0.20
-0.29
-0.34
-0.05
-0.05
-0.10
0.04
0.02
0.11
0.09
0.06
0.05
0.45
-0.07
0.38
0.15
0.08
-0.04
-0.07
-0.16
-0.24
-0.46
0.33
0.15

| |
|---|
| 0.06 |
| 0.07 |
| 0.45 |
| 0.00 |
| -0.01 |
| -0.03 |
| 0.33 |
| 0.00 |
| -0.07 |
| -0.46 |
| -0.58 |
| 0.59 |
| 0.43 |
| 0.32 |
| 0.19 |
| -0.21 |
| -0.02 |
| 0.68 |
| -0.10 |
| -0.18 |
| -0.23 |
| 0.06 |
| -0.64 |
| -0.17 |
| -0.82 |
| 0.20 |
| 0.50 |
| 0.37 |
| 0.22 |
| 0.21 |
| 0.18 |
| 0.13 |
| 0.10 |
| 0.03 |
| -0.32 |
| -0.50 |
| 0.79 |
| 0.19 |
| 0.15 |
| 0.11 |
| 0.11 |
| 0.09 |
| 0.08 |
| 0.07 |
| 0.06 |
| 0.04 |
| 0.03 |
| 0.01 |

-0.03
-0.03
-0.04
-0.04
-0.05
-0.05
-0.07
-0.08
-0.10
-0.11
-0.11
-0.13
-0.13
-0.13
-0.13
-0.14
-0.16
-0.17
-0.24
-0.24
-0.25
-0.32
-0.33
-0.39
-0.43
-0.44
-0.48
-0.54
0.43
0.30
0.10
0.04
-0.09
-0.09
-0.11
-0.15
-0.25
-0.29
-0.31
-0.35
-0.43
-0.45
-0.49
-0.50
-0.56
0.24
-0.30
0.17

| |
|---|
| 0.14 |
| 0.04 |
| 0.03 |
| 0.01 |
| -0.07 |
| -0.08 |
| -0.10 |
| -0.12 |
| -0.17 |
| -0.11 |
| -0.14 |
| -0.39 |
| 0.36 |
| 0.33 |
| -0.32 |
| 0.21 |
| -0.41 |
| -0.51 |
| -0.51 |
| 0.35 |
| 0.19 |
| -0.03 |
| -0.03 |
| -0.19 |
| -0.28 |
| 0.54 |
| 0.79 |
| 0.29 |
| 0.17 |
| 0.05 |
| 0.05 |
| 0.02 |
| 0.01 |
| -0.02 |
| -0.04 |
| -0.04 |
| 0.70 |
| 0.02 |
| -0.82 |
| 1.04 |
| 0.45 |
| 0.08 |
| 0.03 |
| 0.02 |
| -0.02 |
| 0.18 |
| -0.02 |
| -0.26 |

| |
|---|
| 0.41 |
| 0.35 |
| 0.34 |
| 0.17 |
| 0.17 |
| 0.12 |
| 0.12 |
| 0.07 |
| 0.00 |
| -0.04 |
| -0.08 |
| -0.10 |
| -0.15 |
| -0.15 |
| -0.17 |
| -0.17 |
| -0.17 |
| -0.23 |
| -0.32 |
| -0.44 |
| -0.45 |
| -0.45 |
| -0.52 |
| -0.68 |
| -0.78 |
| 0.17 |
| 0.60 |
| 0.42 |
| 0.41 |
| 0.35 |
| 0.31 |
| 0.30 |
| 0.23 |
| -0.02 |
| -0.03 |
| -0.05 |
| -0.05 |
| -0.14 |
| -0.16 |
| -0.17 |
| -0.17 |
| -0.26 |
| -0.43 |
| 0.21 |
| 0.01 |
| 0.15 |
| 0.00 |
| -0.06 |

-0.04
-0.06
-0.12
-0.31
-0.66
0.38
0.13
-0.36
0.13
-0.09
0.36
0.31
0.07
0.02
-0.08
-0.24
0.30
0.19
0.18
0.04
0.04
-0.02
-0.03
-0.13
-0.34
0.21
0.13
0.20
0.18
-0.01
-0.14
-0.29
-0.31
-0.35
-0.39
-0.46
-0.47
0.23
0.23
0.10
0.04
0.04
-0.05
-0.08
-0.09
-0.13
-0.17
-0.20

-0.31
-0.33
-0.34
-0.38
-0.48
-0.54
-0.58
-0.65
-0.70
0.32
0.18
0.11
-0.04
-0.09
-0.39
0.27
0.26
0.20
-0.29
0.15
0.00
-0.12
-0.40
0.98
0.77
0.45
0.35
0.08
-0.06
0.16
0.04
-0.18
0.56
-0.22
-0.63
-0.20
0.27
0.11
-0.07
-0.21
-0.27
-0.31
-0.53
-0.09
0.43
0.31
0.28
0.19

| |
|---|
| -0.13 |
| 0.85 |
| 0.23 |
| 0.07 |
| -0.31 |
| 0.16 |
| 0.10 |
| -0.06 |
| 0.19 |
| 0.12 |
| -0.27 |
| -0.44 |
| -0.18 |
| 0.36 |
| -0.39 |
| 0.01 |
| -0.65 |
| -0.11 |
| 0.04 |
| 0.00 |
| -0.17 |
| -0.19 |
| 0.39 |
| -0.14 |
| 0.16 |
| 0.11 |
| 0.00 |
| -0.06 |
| -0.08 |
| -0.08 |
| -0.16 |
| -0.23 |
| 0.51 |
| 0.47 |
| 0.45 |
| 0.38 |
| 0.34 |
| 0.33 |
| 0.19 |
| 0.05 |
| 0.03 |
| -0.02 |
| -0.03 |
| -0.07 |
| -0.09 |
| -0.39 |
| 0.59 |
| 0.24 |

| |
|---|
| 0.22 |
| 0.08 |
| -0.03 |
| -0.27 |
| -0.35 |
| -0.44 |
| 1.20 |
| 0.24 |
| 0.20 |
| 0.20 |
| 0.19 |
| 0.18 |
| 0.18 |
| 0.17 |
| 0.15 |
| 0.14 |
| 0.13 |
| 0.13 |
| 0.12 |
| 0.10 |
| 0.10 |
| 0.08 |
| 0.05 |
| 0.04 |
| 0.03 |
| 0.02 |
| 0.00 |
| -0.01 |
| -0.03 |
| -0.04 |
| -0.06 |
| -0.07 |
| -0.11 |
| -0.14 |
| -0.15 |
| -0.17 |
| -0.18 |
| -0.18 |
| -0.18 |
| -0.20 |
| -0.21 |
| -0.27 |
| -0.29 |
| -0.37 |
| -0.37 |
| -0.39 |
| -0.42 |
| -0.43 |

-0.49
-0.53
-0.54
0.45
-0.17
-0.28
0.22
0.05
-0.01
-0.04
-0.13
-0.25
-0.46
0.48
0.36
0.25
0.19
-0.13
-0.18
-0.21
-0.23
-0.46
-0.59
-0.67
0.26
0.21
0.14
0.12
0.11
0.04
0.03
0.01
0.01
-0.33
-0.37
-0.37
-0.41
-0.44
-0.44
-0.45
0.20
0.15
0.07
0.02
0.00
-0.03
-0.04
-0.08

| |
|---|
| -0.17 |
| 1.84 |
| 1.52 |
| 1.31 |
| 1.03 |
| 1.01 |
| 0.93 |
| 0.89 |
| 0.88 |
| 0.67 |
| 0.65 |
| 0.64 |
| 0.58 |
| 0.55 |
| 0.53 |
| 0.50 |
| 0.49 |
| 0.47 |
| 0.45 |
| 0.40 |
| 0.30 |
| 0.29 |
| 0.24 |
| 0.23 |
| 0.22 |
| 0.17 |
| 0.15 |
| 0.12 |
| 0.12 |
| 0.11 |
| 0.09 |
| 0.07 |
| 0.05 |
| 0.00 |
| 0.00 |
| -0.03 |
| -0.05 |
| -0.05 |
| -0.06 |
| -0.07 |
| -0.09 |
| -0.10 |
| -0.11 |
| -0.13 |
| -0.13 |
| -0.15 |
| -0.17 |
| -0.20 |

| |
|---|
| -0.21 |
| -0.25 |
| -0.29 |
| -0.37 |
| -0.39 |
| -0.46 |
| -0.46 |
| -0.50 |
| -0.66 |
| -0.82 |
| 0.52 |
| 0.38 |
| 0.21 |
| 0.28 |
| -0.03 |
| -0.25 |
| 0.29 |
| 0.22 |
| 0.13 |
| 0.11 |
| 0.08 |
| 0.00 |
| -0.01 |
| -0.37 |
| -0.37 |
| -0.56 |
| 0.08 |
| -0.09 |
| -0.12 |
| -0.18 |
| -0.26 |
| -0.34 |
| -0.36 |
| -0.48 |
| -0.56 |
| -0.57 |
| -0.70 |
| 0.24 |
| 0.05 |
| -0.07 |
| -0.58 |
| 0.95 |
| 0.48 |
| 0.30 |
| 0.57 |
| 0.07 |
| 0.06 |
| 0.03 |

| |
|---|
| -0.10 |
| -0.19 |
| 0.42 |
| 0.40 |
| 0.45 |
| 0.17 |
| 0.16 |
| 0.12 |
| -0.17 |
| -0.18 |
| 0.20 |
| 0.08 |
| 0.05 |
| 0.00 |
| -0.11 |
| -0.42 |
| -0.30 |
| 0.07 |
| -0.26 |
| -0.27 |
| -0.28 |
| -0.28 |
| -0.29 |
| -0.82 |
| -0.28 |
| -0.41 |
| 0.55 |
| -0.01 |
| -0.41 |
| 0.28 |
| 0.19 |
| 0.17 |
| 0.10 |
| -0.03 |
| -0.37 |
| 0.50 |
| 0.36 |
| 0.33 |
| 0.33 |
| 0.29 |
| 0.20 |
| 0.03 |
| 0.13 |
| -0.14 |
| 0.12 |
| -0.03 |
| -0.07 |
| -0.31 |

-0.44
0.17
0.16
-0.14
-0.15
-0.44
-0.67
-0.78
-0.28
-0.92
-0.12
-0.42
-0.38
0.44
0.15
-0.06
0.26
0.28
-0.35
0.05
0.55
0.43
0.02
-0.07
-0.59
-0.90
-0.32
-0.41
-0.50
-0.59
0.50
0.09
-0.08
0.14
0.11
0.06
0.04
-0.12
-0.24
-0.30
-0.56
-0.02
-0.09
0.45
0.45
0.22
0.20
0.20

| |
|---|
| 0.10 |
| 0.27 |
| 0.19 |
| -0.14 |
| -0.26 |
| -0.29 |
| -0.29 |
| -0.33 |
| -0.51 |
| 0.12 |
| 0.05 |
| -0.63 |
| -0.15 |
| -0.19 |
| 0.10 |
| -0.05 |
| 0.26 |
| 0.11 |
| 0.02 |
| -0.02 |
| 0.08 |
| 0.06 |
| 0.42 |
| 0.05 |
| -0.11 |
| -0.18 |
| -0.58 |
| -0.78 |
| 0.10 |
| 0.04 |
| -0.26 |
| -0.26 |
| -0.26 |
| -0.04 |
| -0.33 |
| -0.26 |
| 0.47 |
| -0.64 |
| 0.02 |
| 0.56 |
| 0.55 |
| 0.44 |
| 0.38 |
| -0.14 |
| -0.16 |
| -0.21 |
| 0.18 |
| 0.12 |

| |
|---|
| -0.09 |
| 0.16 |
| 0.32 |
| 0.04 |
| -0.01 |
| -0.07 |
| -0.15 |
| 0.15 |
| 0.12 |
| 0.08 |
| 0.07 |
| -0.17 |
| -0.19 |
| -0.22 |
| 0.16 |
| 0.10 |
| -0.08 |
| -0.23 |
| 0.39 |
| 0.35 |
| 0.00 |
| -0.21 |
| 0.49 |
| 0.32 |
| 0.29 |
| -0.05 |
| -0.62 |
| 0.07 |
| -0.07 |
| -0.13 |
| -0.56 |
| 0.55 |
| 0.02 |
| -0.30 |
| 0.27 |
| 0.47 |
| 1.41 |
| 0.14 |
| 0.14 |
| 0.09 |
| 0.05 |
| 1.08 |
| 1.08 |
| 0.24 |
| 0.15 |
| 0.14 |
| 0.04 |
| 0.00 |

| |
|---|
| -0.08 |
| -0.10 |
| 0.09 |
| 0.04 |
| 0.07 |
| 0.04 |
| 0.57 |
| -0.38 |
| 0.26 |
| 0.01 |
| -0.05 |
| -0.07 |
| -0.22 |
| -0.22 |
| -0.32 |
| -0.30 |
| 0.33 |
| 0.19 |
| 0.07 |
| 0.05 |
| 0.02 |
| -0.28 |
| -0.43 |
| -0.63 |
| 0.00 |
| -0.07 |
| -0.18 |
| 0.58 |
| 0.53 |
| 0.49 |
| 0.45 |
| 0.44 |
| 0.37 |
| 0.20 |
| 0.19 |
| 0.14 |
| 0.47 |
| -0.45 |
| 0.52 |
| 0.27 |
| 0.26 |
| 0.23 |
| 0.13 |
| 0.08 |
| -0.01 |
| -0.03 |
| -0.06 |
| -0.20 |

| |
|---|
| -0.44 |
| 0.14 |
| 0.13 |
| -0.06 |
| -0.11 |
| -0.23 |
| -0.32 |
| -0.44 |
| 0.41 |
| 0.17 |
| 0.18 |
| 1.00 |
| 0.51 |
| 0.45 |
| 0.06 |
| 0.34 |
| 0.33 |
| 0.06 |
| 0.02 |
| -0.22 |
| 0.18 |
| -0.35 |
| -0.65 |
| 0.12 |
| 0.08 |
| -0.11 |
| -0.18 |
| -0.19 |
| -0.22 |
| 0.30 |
| 0.07 |
| -0.23 |
| -0.26 |
| -0.27 |
| -0.35 |
| 0.11 |
| 0.01 |
| 0.01 |
| -0.42 |
| -0.49 |
| 0.02 |
| -0.12 |
| 0.13 |
| 0.08 |
| -0.28 |
| 0.12 |
| 0.11 |
| 0.11 |

| |
|---|
| -0.04 |
| -0.05 |
| -0.10 |
| -0.10 |
| -0.14 |
| -0.20 |
| -0.29 |
| -0.29 |
| -0.31 |
| -0.36 |
| -0.76 |
| -0.19 |
| 0.18 |
| 0.00 |
| -0.23 |
| 0.19 |
| 0.15 |
| 0.10 |
| 0.25 |
| 0.27 |
| 0.05 |
| -0.75 |
| 0.09 |
| 0.08 |
| -0.03 |
| -0.17 |
| 0.86 |
| 0.27 |
| 0.17 |
| 0.05 |
| -0.08 |
| -1.04 |
| -0.17 |
| -0.51 |
| 0.10 |
| 0.29 |
| 0.12 |
| -0.02 |
| -0.05 |
| -0.08 |
| 0.58 |
| 0.21 |
| -0.01 |
| -0.17 |
| -0.34 |
| -0.70 |
| -0.04 |
| -0.04 |

-0.08
-0.10
-0.15
-0.23
-0.26
-0.47
-0.75
-0.76
-0.07
-0.64
-0.01
-0.16
-0.33
0.64
-0.17
0.32
0.28
0.05
-0.02
-0.17
-0.49
0.27
0.27
-0.03
0.41
-0.05
0.01
0.13
0.02
-0.14
0.27
-0.12
-0.31
-0.31
-0.13
-0.33
0.19
-0.34
0.62
0.06
-0.14
-0.18
-0.50
0.40
0.11
0.19
0.13
0.09

| |
|---|
| 0.05 |
| 0.02 |
| 0.22 |
| 0.22 |
| 0.14 |
| 0.12 |
| 0.11 |
| 0.01 |
| -0.01 |
| -0.03 |
| -0.05 |
| -0.07 |
| -0.09 |
| -0.09 |
| -0.11 |
| -0.11 |
| -0.13 |
| -0.15 |
| -0.19 |
| -0.20 |
| -0.22 |
| -0.27 |
| -0.29 |
| -0.31 |
| -0.32 |
| -0.36 |
| -0.41 |
| -0.52 |
| -0.53 |
| 4.82 |
| -0.11 |
| 0.20 |
| 0.10 |
| -0.09 |
| -0.16 |
| 0.13 |
| 0.12 |
| -0.40 |
| 2.04 |
| 0.04 |
| -0.02 |
| -0.03 |
| -0.63 |
| 0.22 |
| -0.07 |
| -0.08 |
| -0.23 |
| -0.29 |

|  |
| --- |
| 0.03 |
| -0.05 |
| -0.05 |
| -0.06 |
| -0.15 |
| -0.18 |
| -0.22 |
| -0.27 |
| -0.30 |
| -0.32 |
| -0.37 |
| -0.41 |
| 1.62 |
| 0.24 |
| 0.07 |
| -0.18 |
| -0.22 |
| -0.51 |
| 0.52 |
| 0.40 |
| 0.23 |
| 0.11 |
| -0.27 |
| -0.30 |
| -0.30 |
| -0.34 |
| -0.46 |
| 0.11 |
| -0.31 |
| -0.39 |
| 0.41 |
| 0.05 |
| -0.58 |
| 0.36 |
| 0.13 |
| 0.04 |
| 0.03 |
| -0.03 |
| -0.04 |
| -0.14 |
| 0.92 |
| 0.13 |
| -0.10 |
| -0.23 |
| 0.16 |
| 0.08 |
| 0.14 |
| 0.06 |

| |
|---|
| 0.02 |
| -0.18 |
| -0.29 |
| -0.33 |
| -0.63 |
| -1.09 |
| -0.91 |
| 0.08 |
| -0.39 |
| -0.36 |
| -0.38 |
| -0.41 |
| -0.21 |
| -0.23 |
| -0.30 |
| 0.26 |
| 0.15 |
| 0.02 |
| -0.06 |
| -0.13 |
| 0.21 |
| -0.12 |
| -0.39 |
| -0.59 |
| -0.55 |
| 0.41 |
| 0.28 |
| 0.06 |
| -0.14 |
| -0.14 |
| -0.06 |
| -0.09 |
| 0.25 |
| 0.02 |
| -0.04 |
| -0.05 |
| -0.16 |
| -0.18 |
| -0.35 |
| -0.74 |
| 0.32 |
| 0.21 |
| 0.20 |
| 0.11 |
| 0.08 |
| 0.08 |
| 0.07 |
| 0.06 |

| |
|---|
| 0.05 |
| 0.01 |
| 0.00 |
| 0.00 |
| -0.01 |
| -0.07 |
| -0.09 |
| -0.70 |
| -0.02 |
| -0.22 |
| -0.62 |
| 0.18 |
| 0.34 |
| 0.12 |
| 0.04 |
| 0.01 |
| 0.26 |
| 0.21 |
| 0.12 |
| 0.02 |
| 0.01 |
| 0.00 |
| -0.02 |
| -0.04 |
| -0.19 |
| -0.30 |
| -0.73 |
| -0.80 |
| -1.03 |
| 0.14 |
| 0.14 |
| 0.16 |
| 0.11 |
| 0.11 |
| 0.07 |
| -0.16 |
| -0.27 |
| -0.41 |
| -0.59 |
| 0.09 |
| 0.06 |
| 0.04 |
| -0.56 |
| 0.18 |
| -0.10 |
| -0.11 |
| -0.19 |
| -0.37 |

| |
|---|
| 0.21 |
| -0.23 |
| 0.32 |
| 0.22 |
| -0.07 |
| 0.27 |
| 0.08 |
| 0.19 |
| -0.09 |
| -0.10 |
| -0.27 |
| 0.26 |
| 0.05 |
| -0.07 |
| -0.18 |
| -0.21 |
| -0.35 |
| -0.06 |
| -0.07 |
| -0.18 |
| 0.79 |
| 0.31 |
| 0.20 |
| 0.04 |
| -0.10 |
| -0.11 |
| -0.12 |
| -0.13 |
| -0.16 |
| -0.19 |
| -0.23 |
| -0.40 |
| 0.28 |
| 0.15 |
| 0.01 |
| -0.06 |
| -0.10 |
| -0.20 |
| 0.89 |
| 0.57 |
| 0.54 |
| 0.40 |
| 0.31 |
| 0.30 |
| 0.21 |
| 0.20 |
| 0.16 |
| 0.15 |

| |
|---|
| 0.13 |
| 0.08 |
| 0.08 |
| 0.05 |
| -0.02 |
| -0.05 |
| -0.07 |
| -0.07 |
| -0.08 |
| -0.14 |
| -0.23 |
| -0.51 |
| 0.00 |
| -0.06 |
| -0.16 |
| -0.09 |
| 0.60 |
| 0.42 |
| 0.29 |
| 0.09 |
| 0.11 |
| 0.11 |
| 0.16 |
| 0.00 |
| -0.01 |
| 1.13 |
| 0.23 |
| 0.19 |
| -0.01 |
| -0.13 |
| -0.14 |
| -0.14 |
| -0.23 |
| -0.23 |
| -0.25 |
| -0.35 |
| -0.43 |
| -0.53 |
| -0.67 |
| -0.82 |
| -0.92 |
| 1.70 |
| 0.19 |
| 0.17 |
| 0.08 |
| -0.29 |
| -0.07 |
| 0.26 |

| |
|---|
| 0.15 |
| 0.34 |
| 0.24 |
| 0.12 |
| 0.42 |
| 0.20 |
| 0.18 |
| 0.05 |
| -0.37 |
| -0.39 |
| 0.21 |
| 0.15 |
| 0.13 |
| 0.10 |
| 0.04 |
| -0.07 |
| -0.18 |
| -0.18 |
| -0.30 |
| 0.56 |
| 0.13 |
| -0.37 |
| 0.51 |
| 0.28 |
| 0.10 |
| -0.30 |
| 0.48 |
| 0.16 |
| -0.05 |
| 0.00 |
| 0.15 |
| 0.21 |
| 0.05 |
| -0.05 |
| -0.47 |
| -0.03 |
| -0.03 |
| -0.35 |
| -0.12 |
| -0.10 |
| 0.09 |
| -0.10 |
| -0.13 |
| -0.29 |
| -0.41 |
| -0.59 |
| -0.21 |
| -0.11 |

| |
|---|
| -0.11 |
| -0.21 |
| -0.22 |
| 0.83 |
| -0.48 |
| -0.06 |
| -0.33 |
| -0.36 |
| -0.46 |
| 0.20 |
| 0.18 |
| 0.04 |
| -0.13 |
| -0.16 |
| -0.24 |
| -0.39 |
| -0.45 |
| 0.55 |
| 0.24 |
| 0.07 |
| -0.51 |
| 0.61 |
| 0.16 |
| 0.15 |
| 0.15 |
| 0.03 |
| 0.02 |
| -0.02 |
| -0.17 |
| -0.28 |
| -0.08 |
| 0.29 |
| 0.06 |
| -0.55 |
| 0.05 |
| -0.18 |
| -0.24 |
| -0.63 |
| 0.25 |
| 1.04 |
| -0.41 |
| 0.51 |
| 0.47 |
| 0.00 |
| 0.84 |
| 0.36 |
| 0.23 |
| 0.02 |

| |
|---|
| -0.03 |
| -0.08 |
| 1.42 |
| 0.42 |
| 0.39 |
| 0.21 |
| 0.05 |
| -0.15 |
| -0.21 |
| -0.68 |
| -0.94 |
| -0.55 |
| -0.30 |
| 0.16 |
| 0.06 |
| -0.18 |
| 0.13 |
| -0.20 |
| -0.47 |
| -0.60 |
| 0.12 |
| -0.23 |
| 0.27 |
| 0.12 |
| 0.11 |
| 0.07 |
| -0.05 |
| 0.09 |
| -0.25 |
| -0.68 |
| 0.25 |
| 0.12 |
| 0.08 |
| 0.08 |
| -0.09 |
| 0.03 |
| -0.03 |
| -0.19 |
| -0.22 |
| 0.08 |
| -0.19 |
| 0.18 |
| 0.37 |
| 0.22 |
| 0.15 |
| 0.14 |
| 0.00 |
| 0.67 |

| |
|---|
| 0.03 |
| -0.20 |
| -0.47 |
| -0.51 |
| -0.51 |
| -0.70 |
| -0.78 |
| 0.02 |
| -0.22 |
| -0.30 |
| 0.12 |
| 0.07 |
| -0.14 |
| -0.15 |
| 0.15 |
| 0.11 |
| -0.14 |
| -0.21 |
| -0.25 |
| -0.30 |
| -0.20 |
| -0.23 |
| -1.09 |
| -0.03 |
| -0.12 |
| -0.40 |
| 0.21 |
| 0.02 |
| -0.02 |
| -0.04 |
| -0.26 |
| -0.28 |
| -0.31 |
| -0.35 |
| -0.37 |
| -0.44 |
| -0.47 |
| -0.48 |
| -0.53 |
| -0.56 |
| -0.57 |
| -0.66 |
| -0.72 |
| 0.83 |
| 0.02 |
| -0.17 |
| -0.23 |
| -0.33 |

| |
|---|
| -0.34 |
| 0.12 |
| -0.05 |
| -0.21 |
| -0.22 |
| -0.10 |
| -0.20 |
| 0.38 |
| -0.50 |
| -0.03 |
| -0.29 |
| -0.32 |
| 0.33 |
| -0.24 |
| -0.31 |
| 0.60 |
| -0.17 |
| 0.83 |
| 0.77 |
| 0.69 |
| 0.65 |
| 0.23 |
| 0.19 |
| 0.18 |
| 0.15 |
| 0.00 |
| -0.01 |
| -0.02 |
| -0.06 |
| 0.34 |
| 0.03 |
| 0.00 |
| -0.13 |
| 0.23 |
| 0.23 |
| 0.11 |
| 0.10 |
| 0.60 |
| 0.21 |
| 0.02 |
| -0.05 |
| -0.17 |
| -0.21 |
| -0.50 |
| -0.59 |
| 0.27 |
| 0.47 |
| 0.23 |

| |
|---|
| -0.11 |
| -0.12 |
| -0.14 |
| -0.34 |
| 1.15 |
| -0.07 |
| -0.53 |
| -0.69 |
| 0.04 |
| -0.32 |
| -0.39 |
| -0.02 |
| -0.72 |
| -0.07 |
| 0.95 |
| 0.46 |
| -0.03 |
| -0.05 |
| -0.06 |
| -0.18 |
| -0.22 |
| -0.28 |
| -0.40 |
| -0.64 |
| 0.25 |
| 0.10 |
| -0.26 |
| 0.19 |
| -0.04 |
| -0.01 |
| 0.00 |
| 0.59 |
| 0.30 |
| 0.24 |
| 1.01 |
| 0.24 |
| 0.15 |
| -0.15 |
| 0.51 |
| 0.33 |
| -0.02 |
| -0.08 |
| -0.22 |
| 0.20 |
| 0.15 |
| 0.12 |
| 0.11 |
| -0.10 |

-0.10
0.31
0.15
0.22
0.09
-0.12
-0.29
-0.34
0.20
0.56
0.22
0.19
0.18
0.18
0.17
0.11
0.10
0.09
0.08
0.02
-0.02
-0.11
-0.16
-0.24
0.11
-0.18
-0.27
0.61
0.59
0.28
0.27
0.18
0.08
0.08
-0.10
-0.14
-0.18
0.18
-0.11
0.35
0.32
0.11
0.09
-0.07
-0.13
-0.15
-0.28
-0.65

-0.21
0.21
0.12
-0.48
-0.18
-0.51
-0.25
-0.29
-0.34
0.14
0.11
-0.28
-0.34
0.37
0.35
0.05
-0.03
-0.34
0.54
0.25
0.15
0.11
0.04
-0.18
-0.30
-0.57
-1.04
-1.53
-0.25
0.42
0.05
-0.02
-0.24
-0.55
0.07
0.04
0.02
-0.04
-0.07
-0.08
-0.14
-0.22
-0.23
-0.29
-0.31
-0.46
-0.04
0.15

0.02
-0.02
0.40
0.34
0.18
-0.23
-0.25
0.05
-0.31
-0.27
0.30
0.53
-0.04
0.39
0.17
0.13
0.12
0.04
0.04
0.00
-0.01
-0.02
-0.04
-0.06
-0.09
-0.10
-0.11
-0.14
-0.18
-0.50
0.23
0.10
-0.05
0.42
0.32
0.31
0.18
-0.02
-0.05
-0.12
-0.18
-0.29
0.15
0.02
-0.06
0.10
0.01
-0.12

| |
|---|
| 0.01 |
| -0.13 |
| -0.13 |
| -0.35 |
| 0.32 |
| 0.31 |
| -0.27 |
| 0.11 |
| 0.08 |
| -0.70 |
| -0.58 |
| -0.66 |
| 0.09 |
| 0.28 |
| -0.12 |
| 0.02 |
| -0.02 |
| -0.62 |
| -0.65 |
| -0.02 |
| -0.08 |
| -0.08 |
| -0.15 |
| 0.12 |
| -0.06 |
| -0.11 |
| -0.17 |
| -0.24 |
| -0.32 |
| -0.34 |
| -0.37 |
| -0.07 |
| -0.09 |
| -0.21 |
| -0.21 |
| -0.25 |
| 0.97 |
| 0.10 |
| 0.05 |
| 0.21 |
| 0.13 |
| 0.12 |
| 0.10 |
| 2.27 |
| 1.71 |
| 0.63 |
| 0.61 |
| 0.59 |

| |
|---|
| 0.54 |
| 0.47 |
| 0.39 |
| 0.34 |
| 0.26 |
| 0.25 |
| 0.19 |
| 0.19 |
| 0.19 |
| 0.17 |
| 0.16 |
| 0.09 |
| 0.09 |
| 0.08 |
| 0.08 |
| 0.07 |
| 0.05 |
| 0.03 |
| 0.03 |
| 0.02 |
| -0.02 |
| -0.02 |
| -0.03 |
| -0.04 |
| -0.04 |
| -0.05 |
| -0.07 |
| -0.16 |
| -0.37 |
| -0.52 |
| -0.37 |
| 0.10 |
| 0.10 |
| 0.09 |
| -0.52 |
| 0.71 |
| 0.40 |
| 0.43 |
| 0.53 |
| 0.33 |
| -0.48 |
| 0.17 |
| -0.58 |
| -0.82 |
| -0.24 |
| -0.35 |
| -0.52 |
| -0.53 |

| |
|---|
| -0.04 |
| 0.08 |
| -0.05 |
| 0.56 |
| 0.52 |
| -0.69 |
| 0.02 |
| -0.19 |
| -0.59 |
| 0.52 |
| -0.23 |
| -0.33 |
| 0.25 |
| 0.05 |
| -0.05 |
| -0.05 |
| -0.02 |
| -0.06 |
| -0.10 |
| -0.26 |
| -0.76 |
| -0.72 |
| -0.48 |
| 0.18 |
| -0.27 |
| 0.15 |
| -0.54 |
| 0.19 |
| -0.19 |
| -0.24 |
| -0.59 |
| -0.72 |
| -0.21 |
| -0.31 |
| -0.33 |
| -0.17 |
| 0.23 |
| 0.17 |
| 0.14 |
| 0.14 |
| 0.08 |
| 0.07 |
| 0.07 |
| -0.10 |
| -0.74 |
| -1.18 |
| 0.25 |
| 0.07 |

-0.30
0.72
-0.28
0.10
-0.17
-0.27
-0.52
-0.53
-0.76
0.31
0.08
-0.01
-0.13
-0.21
-0.48
-0.53
-0.65
-0.76
0.88
0.42
-0.09
0.03
0.02
-0.06
-0.11
0.38
0.50
0.35
0.02
0.22
0.19
0.03
-0.37
0.09
0.32
0.11
0.11
0.02
0.46
-0.57
-0.39
-0.65
-0.04
-0.07
-0.17
-0.19
-0.21
-0.28

| |
|---|
| -0.34 |
| -0.36 |
| -0.44 |
| -0.52 |
| -0.63 |
| -0.04 |
| -0.17 |
| -0.20 |
| -0.23 |
| 0.16 |
| 0.20 |
| 0.40 |
| 0.14 |
| 0.13 |
| 0.10 |
| 0.09 |
| 0.01 |
| -0.04 |
| -0.15 |
| 0.28 |
| 0.27 |
| -0.05 |
| -0.16 |
| -0.18 |
| -0.19 |
| -0.21 |
| -0.24 |
| -0.29 |
| 0.07 |
| 0.07 |
| 0.10 |
| 0.06 |
| 0.70 |
| 0.15 |
| 0.10 |
| 0.09 |
| 0.06 |
| -0.04 |
| -0.05 |
| -0.06 |
| -0.11 |
| -0.13 |
| -0.17 |
| -0.23 |
| 0.44 |
| -0.06 |
| -0.09 |
| -0.12 |

| |
|---|
| -0.14 |
| -0.81 |
| -0.66 |
| 0.24 |
| 0.13 |
| 0.28 |
| 0.02 |
| -0.14 |
| -0.18 |
| -0.24 |
| -0.24 |
| -0.11 |
| 0.02 |
| -0.63 |
| -0.63 |
| 0.62 |
| 0.47 |
| 0.26 |
| 0.14 |
| 0.13 |
| 0.12 |
| 0.08 |
| 0.03 |
| 0.02 |
| -0.06 |
| -0.07 |
| -0.07 |
| -0.10 |
| -0.25 |
| -0.36 |
| -0.36 |
| -0.38 |
| 0.59 |
| 0.45 |
| -0.14 |
| -0.42 |
| 0.20 |
| 0.10 |
| 0.07 |
| -0.03 |
| -0.11 |
| -0.22 |
| -0.27 |
| -0.31 |
| -0.34 |
| -0.34 |
| -0.49 |
| 0.49 |

| |
|---|
| 0.32 |
| 0.26 |
| -0.17 |
| -0.20 |
| -0.59 |
| 0.09 |
| 0.22 |
| 0.04 |
| 0.04 |
| 0.23 |
| -0.12 |
| -0.29 |
| 0.03 |
| -0.01 |
| -0.11 |
| -0.12 |
| 0.15 |
| 0.00 |
| 0.11 |
| -0.22 |
| -0.29 |
| -0.45 |
| 0.12 |
| 0.49 |
| 0.20 |
| 0.13 |
| 0.01 |
| -0.14 |
| 0.18 |
| 0.09 |
| 0.07 |
| 0.83 |
| 0.80 |
| 0.01 |
| -0.37 |
| 0.46 |
| 0.19 |
| 0.13 |
| 0.08 |
| 0.01 |
| -0.08 |
| -0.25 |
| -0.46 |
| 0.16 |
| 0.08 |
| 0.03 |
| 0.01 |
| 0.00 |

| |
|---|
| -0.19 |
| -0.57 |
| 0.15 |
| 0.12 |
| 0.08 |
| 0.02 |
| 0.00 |
| -0.02 |
| -0.02 |
| -0.09 |
| 0.23 |
| -0.26 |
| -0.09 |
| 0.18 |
| 0.09 |
| -0.06 |
| -0.25 |
| 0.20 |
| -0.75 |
| 0.21 |
| 0.12 |
| -0.04 |
| -0.16 |
| 0.13 |
| -0.11 |
| -0.28 |
| -0.32 |
| -0.21 |
| -0.31 |
| -0.46 |
| 0.05 |
| -0.15 |
| -0.39 |
| 0.05 |
| -0.06 |
| -0.18 |
| -0.20 |
| -0.22 |
| 0.22 |
| -0.17 |
| -0.44 |
| 0.26 |
| 0.12 |
| -0.21 |
| 0.09 |
| -0.32 |
| -0.20 |
| -0.11 |

| |
|---|
| 0.35 |
| 0.33 |
| 0.05 |
| 0.03 |
| -0.50 |
| -0.55 |
| 0.20 |
| 0.20 |
| 0.82 |
| 0.58 |
| -0.03 |
| -0.05 |
| -0.05 |
| -0.08 |
| -0.13 |
| -0.23 |
| -0.45 |
| -0.45 |
| -0.58 |
| 0.31 |
| 0.27 |
| 0.01 |
| -0.12 |
| -0.20 |
| -0.52 |
| 0.14 |
| -0.14 |
| -0.19 |
| -0.24 |
| -0.24 |
| -0.42 |
| -0.86 |
| 0.13 |
| 0.26 |
| 0.18 |
| -0.15 |
| 0.10 |
| 0.13 |
| 0.06 |
| -0.12 |
| -0.19 |
| -0.30 |
| -0.83 |
| -0.37 |
| -0.37 |
| -0.43 |
| -0.54 |
| -0.59 |

| |
|---|
| 0.07 |
| 0.43 |
| 0.40 |
| -0.38 |
| 1.41 |
| 0.98 |
| 0.88 |
| 0.20 |
| 0.08 |
| 0.03 |
| 0.02 |
| -0.06 |
| -0.07 |
| -0.19 |
| -0.23 |
| -0.27 |
| -0.28 |
| -0.30 |
| 0.26 |
| -0.15 |
| 0.13 |
| 1.94 |
| 0.17 |
| 0.04 |
| -0.09 |
| -0.12 |
| -0.23 |
| -0.37 |
| -0.38 |
| -0.15 |
| -0.18 |
| -0.35 |
| -0.71 |
| 0.09 |
| 0.32 |
| 0.47 |
| -0.25 |
| -0.33 |
| -0.40 |
| -0.24 |
| 0.14 |
| -0.30 |
| -0.39 |
| -0.47 |
| 0.34 |
| 0.10 |
| 0.10 |
| 0.29 |

| |
|---|
| 0.21 |
| 0.04 |
| -0.16 |
| -0.10 |
| -0.14 |
| -0.31 |
| -0.33 |
| -0.34 |
| -0.40 |
| -0.67 |
| -0.73 |
| -0.66 |
| 0.37 |
| -0.01 |
| -0.01 |
| 0.00 |
| -0.11 |
| 0.32 |
| 0.14 |
| -0.16 |
| 0.25 |
| 0.14 |
| -0.12 |
| 0.36 |
| 0.13 |
| 0.06 |
| -0.02 |
| -0.07 |
| -0.13 |
| -0.15 |
| -0.53 |
| 0.47 |
| 0.22 |
| 0.11 |
| 0.10 |
| 0.09 |
| 0.05 |
| -0.23 |
| -0.35 |
| 0.00 |
| 1.07 |
| 0.73 |
| 0.66 |
| 0.33 |
| 0.29 |
| 0.29 |
| 0.29 |
| 0.22 |

| |
|---|
| 0.14 |
| 0.11 |
| 0.09 |
| 0.09 |
| 0.09 |
| 0.07 |
| 0.06 |
| 0.06 |
| 0.05 |
| 0.02 |
| 0.01 |
| -0.07 |
| -0.13 |
| -0.14 |
| -0.14 |
| -0.21 |
| -0.25 |
| -0.34 |
| -0.39 |
| 0.37 |
| -0.15 |
| 0.23 |
| 0.17 |
| 0.14 |
| 0.03 |
| 0.01 |
| -0.12 |
| -0.25 |
| -0.07 |
| -0.20 |
| -0.10 |
| -0.11 |
| -0.18 |
| -0.20 |
| -0.42 |
| -0.58 |
| 0.71 |
| 0.38 |
| 0.16 |
| 0.00 |
| -0.21 |
| -0.38 |
| 0.38 |
| -0.31 |
| -0.59 |
| 1.01 |
| 0.73 |
| 0.33 |

| |
|---|
| 0.30 |
| -0.07 |
| 0.63 |
| 0.36 |
| 0.34 |
| 0.80 |
| 0.75 |
| 0.67 |
| 0.61 |
| 0.40 |
| 0.11 |
| 0.09 |
| -0.08 |
| -0.37 |
| -1.22 |
| 0.36 |
| 0.35 |
| 0.25 |
| 0.45 |
| 0.22 |
| -0.11 |
| 0.52 |
| 0.47 |
| 0.21 |
| 0.13 |
| -0.52 |
| 0.45 |
| 0.04 |
| -0.50 |
| -0.44 |
| 0.48 |
| 0.16 |
| 0.09 |
| -0.03 |
| -0.10 |
| -0.34 |
| -0.44 |
| 1.38 |
| 1.11 |
| 0.35 |
| 1.43 |
| 1.15 |
| 0.21 |
| -0.01 |
| 0.08 |
| 0.21 |
| 0.02 |
| -0.03 |

| |
|---|
| -0.34 |
| -0.49 |
| -0.22 |
| -0.28 |
| -0.29 |
| -0.33 |
| -0.33 |
| -0.48 |
| 0.14 |
| 1.11 |
| 0.44 |
| 0.01 |
| -0.14 |
| 0.21 |
| 0.16 |
| 0.33 |
| 0.10 |
| -0.03 |
| -0.53 |
| 0.15 |
| 0.10 |
| 0.06 |
| -0.12 |
| -0.21 |
| -0.34 |
| -0.66 |
| 0.29 |
| 0.09 |
| 0.06 |
| 0.02 |
| 0.02 |
| -0.03 |
| -0.17 |
| -0.34 |
| 0.46 |
| 0.46 |
| -0.02 |
| 0.27 |
| 0.11 |
| -0.09 |
| -0.24 |
| -0.25 |
| -0.32 |
| -0.53 |
| 0.60 |
| 0.37 |
| 0.22 |
| 0.11 |

0.01
-0.01
-0.03
-0.04
-0.06
-0.07
-0.13
-0.22
-0.25
-0.34
-0.35
-0.46
-0.23
-0.71
0.22
0.08
0.47
0.42
0.27
0.30
0.30
0.15
0.13
-0.10
0.09
-0.05
0.32
0.10
0.02
-0.10
-0.48
0.57
0.02
0.02
-0.13
-0.44
-1.05
0.19
-0.44
0.16
-0.18
-0.64
0.62
-0.02
-0.09
-0.20
-0.35
0.44

| |
|---|
| -0.57 |
| 0.39 |
| 0.17 |
| -0.78 |
| 0.17 |
| -0.02 |
| -0.03 |
| -0.04 |
| -0.12 |
| -0.13 |
| -0.16 |
| -0.16 |
| -0.17 |
| -0.32 |
| -0.34 |
| -0.37 |
| -0.43 |
| -0.45 |
| -0.47 |
| -0.53 |
| -0.55 |
| -0.58 |
| -0.64 |
| -0.68 |
| -0.70 |
| -0.70 |
| -0.73 |
| -0.84 |
| -1.06 |
| 0.37 |
| 0.28 |
| -0.03 |
| -0.05 |
| -0.10 |
| -0.25 |
| -0.34 |
| -0.45 |
| -0.60 |
| 0.10 |
| -0.40 |
| 0.43 |
| 0.24 |
| 0.18 |
| -0.38 |
| -0.39 |
| -0.58 |
| 0.31 |
| 0.31 |

-0.48
-0.58
-0.17
-0.06
0.46
0.31
2.04
1.93
1.61
1.26
0.77
0.75
0.47
0.47
0.45
0.42
0.39
0.10
0.01
-0.01
-0.04
-0.07
-0.21
-0.24
-0.59
-0.33
0.21
-0.12
0.45
0.03
-0.02
-0.03
0.30
0.27
-0.09
-0.13
-0.18
-0.23
-0.34
1.92
0.48
0.03
-0.02
-0.03
-0.08
-0.15
-0.22
-0.76

-0.08
-0.13
-0.16
-0.60
-0.75
-0.89
-1.06
1.29
0.32
0.31
0.24
0.21
0.21
0.04
0.01
-0.12
-0.18
-0.29
-0.34
-0.36
-1.01
-0.08
-0.11
-0.16
0.81
0.61
0.57
0.55
0.53
0.19
0.08
0.29
0.23
0.21
0.20
0.18
0.16
0.13
0.11
0.08
0.03
0.00
-0.01
-0.03
-0.09
-0.13
-0.13
-0.16

-0.20
-0.22
-0.23
-0.24
-0.25
-0.27
-0.28
-0.28
-0.29
-0.30
-0.31
-0.31
-0.32
-0.33
-0.33
-0.48
-0.51
-0.56
-0.61
-0.18
-0.34
0.57
0.26
0.07
-0.08
-0.29
-0.34
-0.61
-0.64
0.43
0.32
0.03
0.00
1.64
-0.21
-0.31
0.26
0.16
0.15
-0.01
-0.24
0.03
1.21
0.95
0.10
-0.02
-0.29
-0.35

0.11
0.07
-0.06
-0.09
-0.19
-0.19
-0.22
-0.25
-0.87
0.17
0.33
0.22
-0.12
1.30
1.13
1.08
1.07
0.94
0.52
0.46
0.38
0.30
0.19
0.07
0.05
0.72
0.64
0.53
0.44
0.34
0.19
0.14
1.58
0.70
0.26
-0.02
-0.03
-0.03
-0.07
-0.12
-0.13
-0.20
-0.27
-0.73
0.09
0.28
0.22
0.20

| |
|---|
| 0.45 |
| -0.04 |
| -0.40 |
| 0.19 |
| 0.17 |
| 0.15 |
| 0.11 |
| 0.06 |
| -0.06 |
| -0.17 |
| -0.42 |
| 0.47 |
| 0.27 |
| 0.18 |
| 0.18 |
| 0.15 |
| 0.14 |
| 0.10 |
| -0.13 |
| -0.30 |
| -0.37 |
| -0.10 |
| -0.31 |
| 0.38 |
| 0.28 |
| -0.24 |
| -0.09 |
| -0.29 |
| -0.33 |
| -0.33 |
| 0.10 |
| -0.27 |
| -0.29 |
| -0.63 |
| 0.02 |
| -0.61 |
| 0.21 |
| -0.01 |
| 2.13 |
| 1.35 |
| 0.62 |
| 0.58 |
| -0.17 |
| -0.20 |
| -0.51 |
| 0.79 |
| 0.49 |
| 0.41 |

0.41
0.38
0.19
0.18
0.16
0.16
0.03
0.02
0.02
-0.04
-0.06
-0.07
-0.10
-0.11
-0.15
-0.15
-0.21
-0.23
-0.26
-0.28
-0.30
-0.36
-0.37
-0.37
-0.39
-0.45
-0.62
0.27
0.17
-0.13
-0.21
0.34
0.08
0.09
0.35
0.33
0.12
0.08
0.07
0.03
-0.09
-0.13
-0.16
-0.28
0.04
0.04
-0.05
0.24

0.13
-0.79
-0.22
0.27
0.29
-0.05
-0.35
-0.51
0.16
0.07
-0.01
-0.14
0.10
0.39
0.03
0.71
0.21
0.83
-0.12
-0.06
0.28
1.26
0.29
0.59
0.55
0.52
0.44
0.41
0.41
0.27
0.27
0.21
0.21
0.19
0.15
0.11
0.06
0.03
0.02
-0.05
-0.06
-0.13
-0.14
-0.14
-0.16
-0.16
-0.31
-0.34

| |
|---|
| -0.42 |
| -0.44 |
| -0.63 |
| 0.17 |
| 0.08 |
| 0.04 |
| -0.13 |
| -0.19 |
| -0.31 |
| 0.49 |
| 0.19 |
| 0.13 |
| 0.66 |
| 0.63 |
| 0.63 |
| 0.53 |
| 0.22 |
| 0.21 |
| 0.19 |
| 0.16 |
| 0.15 |
| 0.15 |
| 0.13 |
| 0.12 |
| 0.08 |
| 0.05 |
| 0.05 |
| 0.03 |
| -0.02 |
| -0.04 |
| -0.19 |
| -0.25 |
| -0.32 |
| -0.46 |
| -0.66 |
| -0.03 |
| -0.10 |
| -0.21 |
| -0.47 |
| -0.05 |
| 0.07 |
| -0.20 |
| -0.49 |
| -0.50 |
| -0.56 |
| 0.25 |
| 0.18 |
| 0.08 |

| |
|---|
| -0.06 |
| -0.18 |
| -0.24 |
| -0.43 |
| -0.13 |
| -0.17 |
| 0.29 |
| 0.15 |
| 0.16 |
| -0.32 |
| 0.57 |
| 0.49 |
| 0.37 |
| 0.35 |
| 0.07 |
| 0.01 |
| -0.25 |
| -0.41 |
| 0.24 |
| 0.07 |
| 0.00 |
| 0.00 |
| -0.13 |
| -0.35 |
| -0.55 |
| -0.61 |
| -0.63 |
| 0.22 |
| 0.04 |
| -0.15 |
| 0.07 |
| 0.06 |
| -0.22 |
| -0.28 |
| 0.40 |
| 0.20 |
| -0.06 |
| -0.19 |
| -0.20 |
| -0.74 |
| -0.02 |
| -0.23 |
| 0.26 |
| 0.19 |
| 0.08 |
| 0.06 |
| -0.08 |
| 0.05 |

| |
|---|
| 0.22 |
| 0.36 |
| 0.04 |
| -0.17 |
| -0.25 |
| -0.40 |
| -0.45 |
| 0.27 |
| 0.05 |
| -0.11 |
| -0.77 |
| 0.13 |
| -0.20 |
| -0.10 |
| -0.41 |
| -0.50 |
| 0.25 |
| 0.71 |
| 0.60 |
| -0.30 |
| -0.33 |
| -0.19 |
| 0.20 |
| 0.01 |
| 0.16 |
| -0.13 |
| 0.30 |
| 0.00 |
| -0.38 |
| -0.06 |
| -0.15 |
| 0.13 |
| 0.10 |
| 0.33 |
| 0.20 |
| 0.19 |
| 0.05 |
| 0.01 |
| 0.04 |
| 0.00 |
| -0.02 |
| -0.02 |
| -0.06 |
| -0.07 |
| -0.07 |
| -0.08 |
| -0.08 |
| -0.09 |

-0.13
-0.14
-0.17
-0.17
-0.22
-0.25
-0.25
-0.26
-0.26
-0.36
-0.39
-0.42
-0.50
-0.68
-0.76
-0.14
-0.30
0.43
0.39
0.38
0.23
0.19
0.16
0.15
0.13
0.12
0.08
0.08
0.06
0.02
-0.01
-0.08
-0.09
-0.10
-0.10
-0.13
-0.17
-0.22
-0.22
-0.22
-0.23
-0.24
-0.25
-0.28
-0.28
-0.33
-0.34
-0.38

| |
|---|
| -0.39 |
| -0.39 |
| -0.56 |
| -0.56 |
| -0.65 |
| -0.70 |
| -0.75 |
| 0.14 |
| -0.06 |
| -0.08 |
| -0.29 |
| -0.48 |
| 0.17 |
| 0.06 |
| -0.19 |
| -0.20 |
| -0.26 |
| 0.15 |
| -0.35 |
| -0.42 |
| -0.15 |
| -0.16 |
| -0.17 |
| -0.15 |
| -0.24 |
| -0.35 |
| 0.37 |
| 0.01 |
| -0.10 |
| -0.18 |
| -0.28 |
| -0.50 |
| 0.26 |
| 0.02 |
| -0.63 |
| -0.80 |
| 0.10 |
| -0.16 |
| 0.85 |
| 0.20 |
| 0.10 |
| 0.04 |
| 0.20 |
| 0.17 |
| 0.07 |
| 0.06 |
| 0.05 |
| 0.04 |

| |
|---|
| -0.16 |
| 0.56 |
| 0.04 |
| 0.04 |
| 0.01 |
| -0.02 |
| 0.11 |
| -0.64 |
| 0.29 |
| 0.18 |
| 0.08 |
| 0.01 |
| -0.04 |
| -0.11 |
| -0.15 |
| -0.15 |
| -0.15 |
| 0.07 |
| 0.08 |
| -0.16 |
| -0.31 |
| -0.33 |
| -0.13 |
| -0.62 |
| 0.54 |
| 0.36 |
| 0.35 |
| 0.34 |
| 0.29 |
| 0.24 |
| 0.14 |
| 0.11 |
| 0.03 |
| -0.04 |
| -0.08 |
| -0.10 |
| -0.33 |
| -0.34 |
| -0.53 |
| 0.01 |
| -0.14 |
| -0.17 |
| -0.23 |
| -0.37 |
| -0.43 |
| 0.73 |
| 0.11 |
| -0.04 |

| |
|---|
| -0.06 |
| -0.19 |
| -0.20 |
| -0.27 |
| -0.28 |
| -0.29 |
| -0.30 |
| -0.32 |
| -0.33 |
| -0.34 |
| -0.49 |
| -0.55 |
| -0.58 |
| -0.58 |
| -0.94 |
| 0.13 |
| -0.21 |
| -0.26 |
| 0.42 |
| 0.10 |
| -0.34 |
| -0.08 |
| -0.30 |
| 0.10 |
| 0.04 |
| 0.02 |
| -0.09 |
| -0.42 |
| 0.14 |
| -0.41 |
| -0.92 |
| 0.20 |
| -0.20 |
| -0.25 |
| 0.86 |
| 0.34 |
| 0.33 |
| 0.28 |
| 0.26 |
| 0.25 |
| 0.23 |
| 0.21 |
| 0.15 |
| 0.12 |
| 0.11 |
| 0.11 |
| 0.09 |
| 0.07 |

| |
|---|
| 0.01 |
| 0.00 |
| -0.05 |
| -0.12 |
| -0.13 |
| -0.21 |
| -0.25 |
| -0.52 |
| 0.01 |
| -0.05 |
| -0.30 |
| 0.20 |
| 0.12 |
| -0.02 |
| -0.03 |
| -0.04 |
| -0.05 |
| -0.05 |
| -0.05 |
| -0.06 |
| -0.15 |
| -0.16 |
| -0.23 |
| -0.41 |
| -0.59 |
| 0.79 |
| 0.32 |
| 0.18 |
| 0.02 |
| -0.08 |
| -0.25 |
| -0.30 |
| -0.32 |
| -0.40 |
| -0.50 |
| 0.11 |
| 0.11 |
| -0.31 |
| 0.57 |
| -0.04 |
| -0.36 |
| -0.01 |
| 0.24 |
| 0.08 |
| 0.06 |
| 0.04 |
| -0.01 |
| -0.01 |

| |
|---|
| -0.04 |
| -0.05 |
| -0.18 |
| -0.87 |
| -0.89 |
| 0.35 |
| -0.56 |
| -0.04 |
| -0.37 |
| 0.52 |
| 0.29 |
| 0.28 |
| 0.37 |
| 0.31 |
| 0.07 |
| -0.08 |
| -0.16 |
| 0.25 |
| 0.21 |
| 0.16 |
| 0.07 |
| 0.00 |
| -0.08 |
| -0.21 |
| 0.28 |
| -0.12 |
| -0.50 |
| -0.36 |
| -0.43 |
| -0.42 |
| -0.47 |
| 0.11 |
| -0.02 |
| -0.13 |
| -0.17 |
| -0.01 |
| -0.03 |
| -0.38 |
| -0.38 |
| 0.51 |
| 0.44 |
| 0.26 |
| 0.20 |
| 0.14 |
| 0.12 |
| 0.09 |
| 0.02 |
| -0.04 |

| |
|---|
| -0.07 |
| -0.17 |
| -0.23 |
| -0.24 |
| -0.36 |
| 0.52 |
| 0.43 |
| 0.41 |
| 0.21 |
| 0.19 |
| 0.13 |
| 0.10 |
| 0.05 |
| -0.03 |
| -0.09 |
| -0.11 |
| -0.21 |
| -0.27 |
| -0.29 |
| -0.28 |
| 0.12 |
| 0.11 |
| -0.02 |
| -0.05 |
| -0.13 |
| -0.22 |
| -0.23 |
| -0.23 |
| -0.28 |
| -0.30 |
| -0.31 |
| -0.43 |
| -0.44 |
| -0.51 |
| 0.17 |
| 0.57 |
| 0.20 |
| -0.38 |
| 0.26 |
| -0.16 |
| 0.21 |
| 0.18 |
| 0.18 |
| 0.11 |
| 0.08 |
| -0.05 |
| -0.07 |
| -0.09 |

| |
|---|
| -0.27 |
| 0.28 |
| 0.18 |
| 0.12 |
| 0.04 |
| -0.18 |
| -0.25 |
| -0.56 |
| 0.23 |
| -0.02 |
| -0.13 |
| -0.18 |
| -0.32 |
| -0.02 |
| -0.08 |
| -0.24 |
| -0.37 |
| 2.42 |
| 0.24 |
| 0.27 |
| 0.19 |
| 0.06 |
| 0.06 |
| -0.20 |
| -0.24 |
| -0.30 |
| -0.32 |
| -0.63 |
| 0.05 |
| -0.14 |
| 0.38 |
| 0.18 |
| 0.09 |
| 0.05 |
| 0.03 |
| -0.25 |
| -0.58 |
| 0.39 |
| 0.14 |
| 0.09 |
| 0.06 |
| -0.10 |
| -0.12 |
| 0.17 |
| 0.17 |
| 0.09 |
| -0.49 |
| -0.18 |

| |
|---|
| -0.12 |
| -0.14 |
| -0.25 |
| 0.10 |
| 0.04 |
| 0.01 |
| -0.14 |
| 0.57 |
| 0.10 |
| 0.05 |
| 0.16 |
| 0.00 |
| -0.16 |
| 0.46 |
| 0.23 |
| 0.31 |
| 0.28 |
| 0.28 |
| 0.27 |
| 0.24 |
| 0.19 |
| 0.17 |
| 0.11 |
| 0.08 |
| 0.08 |
| 0.07 |
| 0.06 |
| 0.05 |
| 0.02 |
| 0.01 |
| -0.02 |
| -0.03 |
| -0.08 |
| -0.13 |
| -0.15 |
| -0.27 |
| -0.30 |
| -0.30 |
| -0.36 |
| 0.34 |
| 0.25 |
| 0.24 |
| 0.24 |
| 0.20 |
| 0.15 |
| 0.11 |
| 0.09 |
| 0.07 |

| |
|---|
| 0.06 |
| 0.03 |
| 0.02 |
| 0.01 |
| 0.00 |
| 0.00 |
| 0.00 |
| -0.02 |
| -0.07 |
| -0.39 |
| 0.18 |
| 0.18 |
| 0.11 |
| 0.05 |
| 0.04 |
| 0.00 |
| -0.03 |
| -0.06 |
| -0.10 |
| -0.18 |
| -0.36 |
| 0.79 |
| 0.28 |
| 0.17 |
| 0.16 |
| 0.15 |
| 0.03 |
| 0.03 |
| -0.23 |
| -0.26 |
| -0.32 |
| -0.51 |
| -0.07 |
| 0.25 |
| 0.25 |
| -0.11 |
| -0.18 |
| -0.32 |
| 0.13 |
| -0.18 |
| -0.27 |
| 0.51 |
| 0.17 |
| 0.02 |
| -0.18 |
| 0.21 |
| -0.02 |
| -0.56 |

| |
|---|
| 0.20 |
| 0.04 |
| -0.01 |
| -0.11 |
| -0.24 |
| -0.08 |
| -0.07 |
| 0.35 |
| 0.34 |
| 0.19 |
| 0.18 |
| 0.17 |
| 0.15 |
| 0.10 |
| 0.42 |
| 0.01 |
| -0.03 |
| -0.05 |
| -0.19 |
| 0.41 |
| 0.31 |
| 0.30 |
| 0.23 |
| 0.22 |
| 0.17 |
| 0.09 |
| 0.01 |
| -0.02 |
| -0.05 |
| 0.34 |
| -0.04 |
| 0.52 |
| 1.93 |
| 0.52 |
| 0.33 |
| 0.32 |
| 0.27 |
| 0.22 |
| 0.11 |
| 0.08 |
| 0.05 |
| -0.08 |
| -0.06 |
| -0.01 |
| -0.11 |
| 0.11 |
| 0.00 |
| -0.01 |

-0.01
-0.04
-0.13
-0.15
-0.19
-0.20
-0.22
-0.24
-0.24
-0.39
0.04
-0.09
-0.34
0.34
0.16
0.13
0.11
-0.13
-0.18
-0.27
-0.29
0.31
0.28
-0.04
-0.23
-0.44
0.57
0.27
0.11
0.11
0.09
0.08
0.07
0.06
0.04
-0.09
-0.11
-0.14
-0.18
-0.21
-0.23
0.51
0.18
-0.14
-0.15
-0.25
-0.49
-0.02

-0.11
-0.19
-0.27
0.07
-0.01
-0.05
-0.18
-0.29
-0.33
0.66
0.41
0.26
0.25
0.24
0.11
0.09
0.02
0.00
-0.05
-0.06
0.18
0.63
0.14
0.21
-0.03
-0.06
-0.16
-0.19
-0.22
-0.24
-0.30
-0.40
-0.72
-0.75
-1.03
0.00
-0.06
-0.16
-0.10
0.68
0.26
0.24
-0.02
-0.19
-0.39
-0.52
0.35
0.27

| |
|---|
| 0.25 |
| 0.25 |
| 0.06 |
| 0.00 |
| -0.05 |
| -0.10 |
| 0.32 |
| -0.08 |
| -0.18 |
| -0.26 |
| -0.26 |
| -0.30 |
| -0.37 |
| -0.77 |
| 0.16 |
| 0.01 |
| -0.05 |
| -0.18 |
| -0.21 |
| -0.45 |
| -0.46 |
| -0.46 |
| -0.50 |
| 0.44 |
| 0.25 |
| 0.11 |
| 0.02 |
| -0.64 |
| -0.67 |
| 0.61 |
| 0.45 |
| 0.35 |
| 0.33 |
| 0.28 |
| 0.28 |
| 0.25 |
| 0.18 |
| 0.18 |
| 0.17 |
| 0.14 |
| 0.10 |
| 0.09 |
| 0.09 |
| 0.08 |
| 0.08 |
| 0.05 |
| 0.05 |
| 0.04 |

| |
|---|
| 0.01 |
| -0.01 |
| -0.18 |
| -0.24 |
| -0.39 |
| -0.64 |
| 0.32 |
| 0.32 |
| 0.29 |
| 0.11 |
| 0.07 |
| 0.04 |
| 0.02 |
| -0.02 |
| -0.07 |
| -0.14 |
| -0.17 |
| -0.26 |
| -0.29 |
| -0.33 |
| -0.34 |
| -0.41 |
| -0.41 |
| -0.54 |
| -0.83 |
| 0.75 |
| 0.51 |
| 0.19 |
| 0.15 |
| 0.12 |
| 0.11 |
| 0.09 |
| 0.02 |
| 0.00 |
| -0.07 |
| 0.27 |
| 0.12 |
| 0.08 |
| -0.01 |
| -0.11 |
| -0.05 |
| 0.64 |
| 0.41 |
| 0.34 |
| 0.30 |
| 0.27 |
| 0.26 |
| 0.19 |

| |
|---|
| 0.14 |
| 0.09 |
| 0.08 |
| 0.08 |
| 0.05 |
| 0.05 |
| 0.04 |
| 0.01 |
| 0.00 |
| 0.00 |
| -0.03 |
| -0.03 |
| -0.04 |
| -0.08 |
| -0.08 |
| -0.09 |
| -0.11 |
| -0.12 |
| -0.12 |
| -0.14 |
| -0.14 |
| -0.16 |
| -0.19 |
| -0.20 |
| -0.21 |
| -0.23 |
| -0.25 |
| -0.26 |
| -0.26 |
| -0.37 |
| -0.56 |
| 0.34 |
| 0.21 |
| 0.14 |
| 0.03 |
| 0.02 |
| 0.00 |
| -0.08 |
| -0.10 |
| -0.11 |
| -0.16 |
| -0.21 |
| -0.30 |
| -0.35 |
| -0.38 |
| -0.47 |
| -0.63 |
| 0.18 |

| |
|---|
| 0.13 |
| 0.12 |
| 0.10 |
| 0.05 |
| 0.03 |
| 0.00 |
| -0.12 |
| -0.13 |
| -0.23 |
| -0.27 |
| -0.30 |
| 0.49 |
| 0.12 |
| 0.11 |
| 0.11 |
| 0.08 |
| 0.00 |
| -0.04 |
| -0.06 |
| -0.12 |
| -0.13 |
| -0.28 |
| -0.67 |
| 0.13 |
| 0.09 |
| 0.08 |
| -0.36 |
| 0.15 |
| 0.02 |
| -0.55 |
| 0.22 |
| 0.22 |
| 0.21 |
| -0.08 |
| 1.09 |
| 0.13 |
| -0.01 |
| -0.12 |
| -0.14 |
| -0.17 |
| 0.37 |
| -0.21 |
| -0.27 |
| 0.23 |
| -0.13 |
| 0.12 |
| 0.01 |
| -0.09 |

| |
|---|
| -0.11 |
| -0.05 |
| -0.64 |
| 0.17 |
| 0.46 |
| 0.02 |
| -0.15 |
| -0.19 |
| -0.20 |
| -0.28 |
| -0.53 |
| -0.13 |
| -0.36 |
| -0.50 |
| -0.54 |
| 0.01 |
| -0.02 |
| -0.03 |
| -0.48 |
| -0.54 |
| -0.74 |
| 0.68 |
| 0.67 |
| 0.20 |
| 0.15 |
| 0.11 |
| 0.09 |
| 0.07 |
| 0.07 |
| 0.05 |
| -0.16 |
| 0.41 |
| 0.10 |
| -0.23 |
| -0.23 |
| -0.24 |
| -0.97 |
| -0.06 |
| 0.37 |
| 0.22 |
| 0.09 |
| -0.16 |
| -0.25 |
| -0.05 |
| 0.06 |
| -0.68 |
| 0.20 |
| 0.01 |

| |
|---|
| 0.37 |
| 0.35 |
| 0.22 |
| 0.09 |
| 0.04 |
| -0.03 |
| -0.03 |
| -0.15 |
| -0.38 |
| 0.19 |
| 0.05 |
| -0.09 |
| -0.20 |
| 0.22 |
| -0.27 |
| -0.29 |
| -0.34 |
| -0.45 |
| 0.72 |
| 0.55 |
| 0.43 |
| 0.31 |
| 0.28 |
| -0.09 |
| 0.22 |
| -0.31 |
| -0.39 |
| 0.18 |
| 0.15 |
| 0.12 |
| 0.57 |
| 0.32 |
| 0.22 |
| 0.04 |
| 0.00 |
| 0.07 |
| 0.06 |
| -0.04 |
| -0.20 |
| 0.05 |
| 0.10 |
| -0.04 |
| 2.15 |
| 1.46 |
| 0.25 |
| 0.18 |
| 0.17 |
| 0.11 |

| |
|---|
| 0.00 |
| 0.17 |
| 0.21 |
| 0.13 |
| 0.09 |
| -0.03 |
| -0.07 |
| 0.21 |
| 0.06 |
| -0.37 |
| -0.41 |
| 0.13 |
| -0.02 |
| -0.03 |
| 0.20 |
| 0.29 |
| -0.19 |
| -0.23 |
| -0.57 |
| 0.26 |
| 0.11 |
| 0.08 |
| 0.07 |
| 0.05 |
| -0.10 |
| -0.19 |
| -0.29 |
| -0.35 |
| -0.67 |
| 0.31 |
| 0.21 |
| 0.07 |
| 0.02 |
| 0.40 |
| 0.29 |
| 0.19 |
| 0.16 |
| 0.13 |
| 0.10 |
| 0.08 |
| 0.06 |
| -0.04 |
| -0.09 |
| -0.09 |
| -0.11 |
| -0.23 |
| -0.43 |
| -0.50 |

| |
|---|
| 0.30 |
| 0.28 |
| 0.04 |
| -0.33 |
| -0.47 |
| -0.52 |
| -0.52 |
| -0.72 |
| -0.25 |
| 0.36 |
| -0.05 |
| -0.16 |
| -0.17 |
| -0.18 |
| -0.18 |
| -0.23 |
| 0.28 |
| 0.11 |
| -0.15 |
| -0.17 |
| -0.40 |
| -0.20 |
| 0.22 |
| 0.15 |
| -0.11 |
| -0.21 |
| -0.13 |
| -0.14 |
| 0.01 |
| -0.15 |
| -0.15 |
| -0.44 |
| -0.04 |
| -0.45 |
| 2.20 |
| 0.38 |
| 0.29 |
| -0.34 |
| 0.98 |
| 0.40 |
| 0.23 |
| 0.22 |
| 0.15 |
| 0.10 |
| 0.05 |
| 0.02 |
| -0.05 |
| -0.05 |

| |
|---|
| -0.11 |
| -0.16 |
| -0.17 |
| -0.21 |
| -0.22 |
| -0.22 |
| -0.23 |
| -0.33 |
| 0.59 |
| 0.25 |
| 0.25 |
| 0.18 |
| 0.16 |
| 0.09 |
| 0.00 |
| 0.13 |
| 0.12 |
| 0.02 |
| -0.09 |
| -0.19 |
| 0.86 |
| 0.64 |
| 0.60 |
| 0.59 |
| 0.25 |
| 0.23 |
| 0.19 |
| 0.09 |
| 0.04 |
| -0.16 |
| -0.16 |
| -0.26 |
| 0.18 |
| 0.04 |
| -0.09 |
| -0.25 |
| -0.27 |
| -0.30 |
| 0.17 |
| -0.61 |
| -0.04 |
| 0.30 |
| 0.16 |
| 0.27 |
| -0.12 |
| -0.41 |
| -0.61 |
| 0.02 |

| |
|---|
| 0.16 |
| -0.02 |
| -0.07 |
| 0.50 |
| 0.40 |
| 0.16 |
| 0.16 |
| 0.00 |
| -0.14 |
| -0.18 |
| -0.19 |
| -0.19 |
| -0.24 |
| -0.66 |
| 0.10 |
| 0.03 |
| -0.08 |
| -0.12 |
| -0.32 |
| -0.37 |
| -0.37 |
| -0.42 |
| -0.53 |
| 0.17 |
| -0.33 |
| -0.35 |
| 0.47 |
| 0.13 |
| -0.06 |
| -0.21 |
| 0.13 |
| 0.10 |
| 1.03 |
| -0.02 |
| -0.35 |
| 0.52 |
| 0.07 |
| 0.03 |
| -0.21 |
| -0.10 |
| 0.08 |
| -0.04 |
| -0.29 |
| -0.18 |
| -0.01 |
| 0.21 |
| 0.10 |
| 0.02 |

0.08
-0.01
-0.07
-0.09
1.78
1.27
0.85
0.81
0.78
0.78
0.72
0.62
0.62
0.61
0.60
0.60
0.53
0.53
0.52
0.52
0.51
0.46
0.45
0.39
0.39
0.38
0.38
0.38
0.35
0.35
0.34
0.33
0.33
0.32
0.27
0.26
0.26
0.25
0.24
0.23
0.20
0.19
0.17
0.17
0.17
0.15
0.14
0.13

0.12
0.11
0.10
0.09
0.08
0.02
0.00
-0.02
-0.03
-0.05
-0.05
-0.07
-0.07
-0.08
-0.08
-0.09
-0.14
-0.15
-0.30
-0.37
-0.37
-0.46
-0.69
0.46
0.11
-0.54
0.29
0.05
-0.45
2.49
1.78
1.75
0.73
0.68
0.18
0.09
-0.16
-0.31
0.35
0.02
-0.05
-0.06
-0.10
-0.11
-0.12
-0.21
-0.21
-0.68

-0.91
-0.97
0.17
0.04
-0.02
-0.27
-0.27
0.22
0.35
-0.21
0.11
-0.26
0.13
0.33
0.02
-0.29
0.03
-0.01
0.25
0.23
0.05
0.03
-0.13
-0.14
-0.16
-0.43
-0.50
-0.54
0.63
1.50
-0.13
-0.11
-0.18
0.07
0.17
0.15
0.09
0.09
0.01
-0.08
-0.13
-0.18
-0.26
-0.29
-0.43
-0.48
-0.50
-0.06

| |
|---|
| 0.09 |
| -0.01 |
| -0.09 |
| -0.14 |
| -0.09 |
| -0.56 |
| -0.38 |
| 0.32 |
| 0.26 |
| 0.11 |
| -0.05 |
| -0.33 |
| 0.27 |
| 0.12 |
| 0.10 |
| 0.08 |
| -0.05 |
| -0.05 |
| 0.05 |
| -0.01 |
| -0.07 |
| -0.10 |
| 0.65 |
| 0.04 |
| 0.02 |
| 0.01 |
| 0.00 |
| -0.02 |
| -0.38 |
| -0.28 |
| -0.45 |
| 0.22 |
| 0.27 |
| 0.10 |
| -0.17 |
| -0.23 |
| 0.15 |
| 0.17 |
| 0.13 |
| 0.07 |
| 0.05 |
| -0.02 |
| -0.06 |
| -0.11 |
| -0.21 |
| -0.22 |
| -0.28 |
| -0.29 |

| |
|---|
| -0.40 |
| -0.41 |
| -0.41 |
| -0.72 |
| -0.25 |
| -0.38 |
| -0.19 |
| 0.01 |
| 0.05 |
| -0.52 |
| -0.24 |
| -0.16 |
| 0.92 |
| 0.50 |
| 0.36 |
| 0.32 |
| 0.30 |
| 0.29 |
| 0.26 |
| 0.26 |
| 0.14 |
| 0.00 |
| -0.02 |
| -0.03 |
| -0.06 |
| -0.07 |
| -0.17 |
| -0.43 |
| -0.44 |
| -0.46 |
| -0.50 |
| 0.51 |
| 0.42 |
| 0.26 |
| 0.00 |
| 0.17 |
| -0.10 |
| 1.41 |
| 1.15 |
| 1.03 |
| 0.81 |
| 0.79 |
| 0.78 |
| 0.76 |
| 0.74 |
| 0.74 |
| 0.74 |
| 0.71 |

0.70
0.67
0.63
0.63
0.62
0.57
0.54
0.54
0.52
0.52
0.50
0.49
0.49
0.48
0.42
0.39
0.38
0.38
0.38
0.38
0.35
0.33
0.33
0.32
0.32
0.32
0.32
0.31
0.30
0.29
0.24
0.24
0.20
0.19
0.18
0.14
0.13
0.10
0.10
0.09
0.08
0.07
0.05
0.02
0.01
0.00
-0.01
-0.02

-0.03
-0.03
-0.03
-0.04
-0.07
-0.08
-0.08
-0.08
-0.08
-0.09
-0.09
-0.10
-0.12
-0.18
-0.20
-0.21
-0.23
-0.26
-0.29
-0.38
0.11
0.03
-0.08
-0.02
-0.03
-0.05
-0.06
-0.13
-0.27
-0.30
-0.30
-0.44
-1.21
0.30
0.17
0.07
-0.19
-0.03
-0.11
-0.12
-0.25
-0.30
-0.43
0.72
0.38
0.14
0.14
0.09

0.04
0.00
-0.06
-0.07
-0.08
-0.08
-0.16
-0.16
-0.18
-0.18
-0.19
-0.21
-0.25
-0.25
-0.30
-0.31
-0.33
-0.35
-0.35
-0.37
-0.37
-0.46
0.10
0.06
0.04
0.00
-0.06
-0.33
0.07
0.01
0.35
-0.03
-0.06
-0.16
-0.26
-0.34
-0.61
0.15
0.14
0.01
-0.16
0.35
0.56
0.17
0.11
-0.02
-0.02
-0.21

| |
|---|
| 0.25 |
| 0.12 |
| 0.06 |
| 0.03 |
| -0.05 |
| 0.41 |
| 0.32 |
| 0.26 |
| 0.18 |
| 0.17 |
| 0.15 |
| 0.15 |
| 0.15 |
| 0.12 |
| 0.10 |
| 0.09 |
| 0.09 |
| 0.05 |
| 0.05 |
| 0.04 |
| -0.02 |
| -0.04 |
| -0.37 |
| 0.22 |
| 0.12 |
| 0.44 |
| 0.35 |
| 0.33 |
| 0.32 |
| 0.22 |
| 0.11 |
| 0.04 |
| 0.03 |
| -0.04 |
| -0.08 |
| -0.15 |
| -0.16 |
| -0.19 |
| -0.21 |
| -0.23 |
| -0.30 |
| -0.35 |
| -0.41 |
| -0.44 |
| -0.51 |
| -0.54 |
| -0.90 |
| 0.21 |

-0.40
0.39
0.21
0.04
-0.04
0.09
-1.01
-0.30
0.09
-0.36
-0.48
-0.12
-0.13
-0.06
0.05
-0.43
0.25
0.13
-0.23
0.13
-0.21
0.36
0.24
-0.07
-0.10
0.19
-0.15
-0.16
-0.24
-0.34
-0.39
-0.66
0.12
0.06
0.03
0.03
-0.01
-0.33
-0.77
0.12
-0.22
-0.28
-0.33
0.66
0.53
0.19
-0.04
-0.05

| |
|---|
| -0.12 |
| -0.14 |
| -0.15 |
| -0.16 |
| -0.18 |
| -0.34 |
| 0.23 |
| 0.19 |
| 0.05 |
| 0.31 |
| -0.02 |
| -0.03 |
| -0.06 |
| -0.43 |
| -0.85 |
| -0.15 |
| -0.13 |
| 0.35 |
| -0.07 |
| -0.10 |
| -0.15 |
| -0.17 |
| -0.24 |
| -0.71 |
| -0.80 |
| 0.25 |
| 0.23 |
| -0.01 |
| -0.42 |
| -0.20 |
| -0.44 |
| 0.06 |
| -0.13 |
| -0.13 |
| -0.18 |
| 0.14 |
| 0.12 |
| -0.03 |
| 0.16 |
| -0.01 |
| -0.36 |
| 0.14 |
| -0.03 |
| -0.07 |
| -0.07 |
| -0.29 |
| -0.46 |
| -0.07 |

| |
|---|
| -0.07 |
| 0.24 |
| 0.23 |
| 0.31 |
| 0.28 |
| 0.26 |
| 0.13 |
| 0.09 |
| 0.05 |
| 0.02 |
| 0.00 |
| -0.01 |
| -0.02 |
| -0.05 |
| -0.13 |
| -0.41 |
| -0.41 |
| -0.47 |
| -0.56 |
| -0.67 |
| 0.65 |
| 0.64 |
| 0.52 |
| 0.43 |
| 0.34 |
| -0.36 |
| -0.61 |
| 0.32 |
| 0.05 |
| 0.45 |
| 0.07 |
| -0.05 |
| -0.06 |
| -0.12 |
| -0.12 |
| -0.30 |
| -0.02 |
| -0.03 |
| -0.05 |
| -0.12 |
| -0.21 |
| -0.29 |
| -0.29 |
| -0.34 |
| 0.39 |
| 0.19 |
| -0.08 |
| -0.14 |

-0.17
-0.21
-0.21
-0.30
-0.50
-0.70
0.61
0.09
0.08
0.05
0.01
-0.02
-0.04
-0.04
-0.20
-0.26
-0.27
-0.36
-0.08
0.18
0.62
0.35
-0.15
-0.62
0.19
-0.19
-0.31
0.36
0.28
0.25
0.16
-0.04
-0.05
-0.14
-0.16
-0.19
-0.19
-0.23
-0.24
0.47
0.24
0.07
-0.30
0.51
0.24
0.21
-0.10
-0.57

| |
|---|
| -0.03 |
| 0.34 |
| 0.14 |
| 0.06 |
| -0.19 |
| -0.30 |
| -0.47 |
| -0.66 |
| 0.20 |
| 0.52 |
| 0.28 |
| 0.38 |
| 0.09 |
| 0.04 |
| -0.19 |
| 0.73 |
| -0.06 |
| -0.08 |
| -0.21 |
| -0.26 |
| -0.03 |
| -0.45 |
| -0.60 |
| -0.20 |
| 0.59 |
| 0.11 |
| 0.09 |
| -0.04 |
| -0.18 |
| 0.43 |
| 0.10 |
| 0.08 |
| 0.07 |
| -0.08 |
| 0.15 |
| 0.12 |
| -0.38 |
| 0.70 |
| 0.41 |
| 0.14 |
| 0.10 |
| 0.14 |
| 0.38 |
| 0.29 |
| 0.05 |
| -0.22 |
| 0.31 |
| -0.10 |

| |
|---|
| 0.67 |
| 0.67 |
| 0.66 |
| 0.49 |
| 0.45 |
| 0.29 |
| -0.12 |
| 0.57 |
| 0.51 |
| 0.50 |
| 0.40 |
| 0.38 |
| 0.26 |
| 0.23 |
| 0.36 |
| 0.17 |
| -0.24 |
| 0.03 |
| -0.02 |
| -0.09 |
| -0.12 |
| -0.19 |
| -0.31 |
| -0.32 |
| -0.61 |
| -0.37 |
| 0.11 |
| 0.81 |
| 0.71 |
| 0.29 |
| 0.28 |
| 0.13 |
| 0.11 |
| 0.04 |
| -0.02 |
| -0.14 |
| -0.15 |
| -0.16 |
| -0.18 |
| -0.22 |
| -0.26 |
| -0.28 |
| -0.36 |
| -0.37 |
| -0.38 |
| -0.53 |
| -0.69 |
| -1.25 |

| |
|---|
| -0.08 |
| 0.45 |
| 0.14 |
| 0.05 |
| 0.43 |
| 0.23 |
| 0.21 |
| 0.18 |
| 0.18 |
| 0.08 |
| 0.07 |
| -0.04 |
| -0.13 |
| -0.17 |
| -0.18 |
| 0.66 |
| 0.50 |
| 0.26 |
| 0.23 |
| 0.22 |
| 0.11 |
| 0.10 |
| 0.10 |
| 0.10 |
| -0.09 |
| -0.11 |
| -0.14 |
| -0.34 |
| 0.40 |
| 0.31 |
| 0.23 |
| 0.19 |
| 0.17 |
| 0.15 |
| 0.13 |
| 0.09 |
| 0.05 |
| 0.04 |
| -0.05 |
| -0.06 |
| -0.12 |
| -0.25 |
| 0.34 |
| 0.18 |
| -0.19 |
| -0.28 |
| -0.52 |
| -0.22 |

-0.02
-0.17
-0.44
0.29
-0.05
-0.15
0.24
0.04
0.04
-0.13
-0.32
-0.35
-0.38
0.00
-0.04
-0.19
-0.22
0.36
0.19
0.15
0.17
-0.11
-0.13
-0.22
0.15
0.29
-0.07
-0.16
-0.08
-0.12
0.60
0.29
0.14
0.12
0.06
-0.11
-0.18
-0.26
0.21
-0.36
-0.36
0.63
0.16
0.14
0.88
0.30
-0.07
-0.15

| |
|---|
| -0.17 |
| 0.00 |
| 0.18 |
| -0.22 |
| -0.41 |
| 0.21 |
| 0.17 |
| 0.20 |
| 0.12 |
| 0.10 |
| 0.51 |
| 0.41 |
| 0.23 |
| 0.16 |
| 0.09 |
| 0.07 |
| 0.02 |
| 0.00 |
| -0.18 |
| 0.30 |
| 0.28 |
| 0.09 |
| -0.04 |
| -0.05 |
| -0.22 |
| -0.30 |
| -0.39 |
| -0.43 |
| 0.26 |
| 0.07 |
| 0.04 |
| 0.01 |
| -0.11 |
| -0.13 |
| -0.13 |
| -0.13 |
| -0.13 |
| -0.24 |
| -0.25 |
| -0.27 |
| -0.27 |
| -0.33 |
| -0.36 |
| -0.42 |
| -0.50 |
| -0.63 |
| -0.64 |
| -0.64 |

0.10
0.04
-0.34
0.14
0.11
0.03
0.03
0.02
-0.01
-0.07
-0.09
0.48
0.00
-0.12
0.75
0.67
0.24
0.15
0.03
-0.12
-0.24
-0.57
0.29
0.24
0.07
-0.03
0.15
0.08
-0.04
-0.07
0.29
0.15
-0.10
-0.44
-0.48
-0.54
0.01
-0.08
-0.12
0.25
0.03
-0.11
-0.21
-0.22
-0.23
-0.21
-0.37
-0.45

| |
|---|
| 0.04 |
| -0.22 |
| -0.26 |
| -0.66 |
| -0.89 |
| -1.24 |
| -0.61 |
| 0.53 |
| 0.47 |
| 0.21 |
| 0.19 |
| 0.17 |
| 0.17 |
| 0.14 |
| 0.13 |
| 0.11 |
| 0.07 |
| 0.06 |
| -0.02 |
| -0.03 |
| -0.03 |
| -0.09 |
| -0.09 |
| -0.10 |
| -0.15 |
| -0.16 |
| -0.19 |
| -0.20 |
| -0.21 |
| -0.25 |
| -0.28 |
| -0.32 |
| -0.43 |
| -0.47 |
| -0.48 |
| -0.53 |
| -0.62 |
| -0.62 |
| -1.06 |
| -1.07 |
| -0.45 |
| -0.48 |
| -0.07 |
| -0.10 |
| -0.33 |
| -0.07 |
| 0.27 |
| 0.14 |

| |
|---|
| 0.09 |
| -0.06 |
| -0.07 |
| -0.09 |
| -0.15 |
| -0.20 |
| -0.25 |
| -0.28 |
| -0.35 |
| -0.36 |
| -0.63 |
| -0.17 |
| -0.22 |
| 0.29 |
| -0.10 |
| -0.33 |
| -0.36 |
| 0.08 |
| -0.17 |
| -0.23 |
| -0.23 |
| 0.00 |
| 0.44 |
| 0.09 |
| 0.07 |
| 0.04 |
| 0.04 |
| 0.03 |
| -0.03 |
| -0.05 |
| -0.13 |
| -0.52 |
| -0.42 |
| 0.18 |
| -0.52 |
| -0.89 |
| 0.29 |
| 0.29 |
| 0.11 |
| 0.09 |
| 0.35 |
| 0.33 |
| 0.19 |
| 0.09 |
| 0.07 |
| 0.04 |
| 0.04 |
| -0.03 |

-0.03
-0.10
-0.11
-0.14
-0.14
-0.16
-0.17
-0.19
-0.22
-0.24
-0.31
-0.32
-0.34
-0.67
-0.88
0.45
0.09
0.03
-0.01
-0.05
-0.05
-0.12
-0.18
-0.21
-0.23
-0.23
-0.35
-0.47
-0.72
-0.87
0.33
0.26
0.01
-0.73
-0.87
0.38
-0.26
-0.53
-0.29
-0.45
0.54
0.26
0.11
0.09
0.02
0.00
-0.02
-0.02

| |
|---|
| -0.05 |
| -0.06 |
| -0.07 |
| -0.12 |
| -0.22 |
| -0.24 |
| -0.31 |
| -1.18 |
| 1.24 |
| 0.54 |
| 0.52 |
| 0.04 |
| 0.00 |
| -0.07 |
| -0.10 |
| -0.12 |
| -0.13 |
| -0.15 |
| -0.16 |
| -0.16 |
| -0.28 |
| -0.42 |
| -0.78 |
| 0.24 |
| 0.07 |
| 0.02 |
| 0.00 |
| -0.01 |
| -0.05 |
| -0.26 |
| -0.39 |
| -0.42 |
| -0.17 |
| -0.22 |
| 0.15 |
| -0.09 |
| 0.48 |
| -0.24 |
| -0.49 |
| 0.17 |
| 0.05 |
| -0.16 |
| -0.25 |
| -0.26 |
| -0.05 |
| 0.28 |
| 0.21 |
| 0.19 |

| |
|---|
| 0.05 |
| 0.02 |
| -0.07 |
| -0.19 |
| 0.80 |
| 0.30 |
| 0.84 |
| 0.27 |
| 0.63 |
| 0.28 |
| 0.00 |
| -0.13 |
| -0.25 |
| -0.31 |
| -0.42 |
| -0.60 |
| -0.86 |
| 0.19 |
| 0.12 |
| 0.07 |
| 0.78 |
| 0.28 |
| -0.36 |
| 0.19 |
| -0.14 |
| 0.36 |
| 0.36 |
| 0.28 |
| 0.09 |
| -0.11 |
| -0.12 |
| -0.26 |
| -0.35 |
| -0.44 |
| -0.66 |
| 0.39 |
| 0.29 |
| 0.26 |
| 0.19 |
| 0.14 |
| 0.03 |
| -0.09 |
| -0.15 |
| -0.33 |
| -0.52 |
| 0.19 |
| 0.04 |
| 0.00 |

| |
|---|
| -0.02 |
| -0.07 |
| -0.18 |
| -0.19 |
| -0.19 |
| -0.35 |
| -0.38 |
| -0.45 |
| -0.52 |
| -0.53 |
| 0.80 |
| 0.05 |
| 0.04 |
| -0.21 |
| -0.21 |
| -0.27 |
| -0.42 |
| 0.11 |
| -0.07 |
| 0.21 |
| 0.15 |
| 0.15 |
| -0.02 |
| 0.52 |
| 0.24 |
| 0.04 |
| -0.05 |
| -0.16 |
| -0.17 |
| -0.19 |
| 0.07 |
| 0.06 |
| -0.10 |
| -0.21 |
| -0.65 |
| 0.32 |
| 0.30 |
| 0.28 |
| 0.11 |
| 0.05 |
| 0.01 |
| -0.04 |
| -0.38 |
| 0.05 |
| 0.00 |
| 0.16 |
| 0.05 |
| -0.10 |

| |
|---|
| 0.63 |
| 0.61 |
| 0.35 |
| 0.34 |
| 0.31 |
| 0.26 |
| 0.22 |
| 0.21 |
| 0.19 |
| 0.15 |
| 0.14 |
| 0.12 |
| 0.12 |
| 0.05 |
| 0.01 |
| -0.01 |
| -0.06 |
| -0.06 |
| -0.07 |
| -0.09 |
| -0.13 |
| -0.13 |
| -0.14 |
| -0.15 |
| -0.15 |
| -0.20 |
| -0.21 |
| -0.26 |
| -0.28 |
| -0.28 |
| -0.36 |
| -0.44 |
| -0.47 |
| -0.48 |
| -0.49 |
| -0.60 |
| -0.74 |
| 0.39 |
| -0.02 |
| -0.18 |
| -0.53 |
| 0.08 |
| -0.02 |
| -0.03 |
| -0.21 |
| -0.22 |
| -0.27 |
| -0.31 |

| |
|---|
| 1.68 |
| 1.52 |
| 1.39 |
| 0.62 |
| 0.58 |
| 0.54 |
| 0.52 |
| 0.50 |
| 0.37 |
| 0.25 |
| 0.19 |
| 0.17 |
| 0.06 |
| 0.04 |
| -0.18 |
| -0.18 |
| -0.36 |
| 0.06 |
| -0.09 |
| -0.26 |
| 0.50 |
| -0.17 |
| 0.48 |
| -0.03 |
| 0.60 |
| 0.50 |
| 0.30 |
| 0.24 |
| -0.05 |
| -0.09 |
| -0.11 |
| -0.17 |
| -0.19 |
| -0.27 |
| -0.33 |
| 0.44 |
| 0.30 |
| 0.29 |
| -0.26 |
| -0.03 |
| 0.31 |
| 0.31 |
| 0.21 |
| 0.13 |
| 0.10 |
| 0.01 |
| -0.14 |
| -0.22 |

| |
|---|
| -0.78 |
| 0.08 |
| 0.02 |
| -0.06 |
| -0.33 |
| -0.52 |
| -0.73 |
| 0.11 |
| -0.04 |
| -0.32 |
| 0.15 |
| 0.20 |
| -0.10 |
| 0.14 |
| 0.04 |
| -0.20 |
| -0.49 |
| -0.13 |
| -0.24 |
| -1.13 |
| 0.49 |
| 0.29 |
| 0.27 |
| 0.25 |
| 0.20 |
| 0.17 |
| 0.09 |
| -0.03 |
| -0.05 |
| -0.10 |
| -0.27 |
| -0.30 |
| -0.73 |
| 0.12 |
| 0.10 |
| -0.61 |
| -0.69 |
| 0.66 |
| 0.43 |
| 0.24 |
| 0.23 |
| 0.22 |
| 0.19 |
| 0.03 |
| -0.06 |
| -0.08 |
| -0.21 |
| 0.53 |

| |
|---|
| 0.49 |
| 0.34 |
| -0.23 |
| 0.30 |
| 0.81 |
| -0.33 |
| 0.11 |
| -0.20 |
| -0.22 |
| -0.41 |
| -0.48 |
| -0.49 |
| 0.34 |
| 0.15 |
| 0.08 |
| 0.07 |
| 0.05 |
| 0.02 |
| 0.00 |
| -0.02 |
| -0.04 |
| 0.06 |
| -0.01 |
| -0.12 |
| -0.24 |
| -0.37 |
| -0.14 |
| -0.32 |
| 0.24 |
| 0.17 |
| -0.16 |
| -0.29 |
| 1.50 |
| 0.76 |
| 0.04 |
| -0.45 |
| 0.03 |
| -0.07 |
| 0.40 |
| 0.26 |
| 0.23 |
| 0.02 |
| 0.00 |
| -0.03 |
| -0.22 |
| -0.34 |
| 0.06 |
| 0.02 |

-0.12
-0.11
-0.02
-0.30
-0.38
-0.43
-0.54
-0.29
0.34
0.11
-0.09
-0.11
-0.16
-0.32
-0.40
-0.86
-0.02
-0.15
-0.19
-0.28
-0.43
-0.90
0.05
-0.16
0.37
0.22
0.10
0.02
-0.02
-0.20
-0.27
-0.38
-0.64
-0.27
0.22
0.16
0.08
-0.06
-0.12
0.11
0.05
-0.48
-0.51
0.41
0.36
0.35
-0.15
-0.23

-0.31
-0.08
0.27
0.39
-0.09
-0.02
1.45
1.18
0.70
0.64
0.54
0.13
-0.18
0.49
0.13
0.09
0.04
-0.72
-0.18
-0.13
-0.64
0.08
-0.04
0.11
-0.50
-0.29
0.19
0.31
-0.09
-0.21
-0.22
-0.28
-0.40
0.64
-0.11
-0.30
-0.34
0.12
0.73
0.72
-1.01
0.17
0.24
0.08
0.00
-0.20
-0.16
-0.38

-0.05
0.00
0.54
0.40
0.15
0.11
0.07
-0.13
-0.16
-0.30
-0.38
0.71
0.34
-0.09
-0.12
-0.13
-0.30
-0.38
0.26
-0.02
0.33
0.27
-0.05
0.17
-0.25
-0.25
-0.44
-0.46
-0.52
-1.00
0.27
0.21
0.20
0.20
0.01
-0.05
-0.33
0.16
-0.22
-0.11
0.08
0.06
0.05
0.00
-0.21
-0.22
-0.27
-0.56

| |
|---|
| 0.40 |
| -0.31 |
| 0.97 |
| 0.24 |
| 0.19 |
| 0.15 |
| 0.14 |
| 0.13 |
| 0.09 |
| 0.08 |
| 0.05 |
| 0.04 |
| 0.03 |
| -0.10 |
| -0.12 |
| -0.15 |
| -0.26 |
| 0.40 |
| -0.36 |
| 0.42 |
| -0.53 |
| -0.72 |
| 0.13 |
| -0.63 |
| 0.08 |
| -0.05 |
| -0.05 |
| -0.07 |
| -0.09 |
| -0.10 |
| -0.16 |
| -0.23 |
| -0.40 |
| -0.05 |
| 1.17 |
| 0.44 |
| 0.30 |
| 0.29 |
| 0.19 |
| 0.14 |
| 0.02 |
| 0.01 |
| 0.00 |
| 0.00 |
| 0.00 |
| -0.05 |
| -0.07 |
| -0.09 |

-0.09
-0.09
-0.10
-0.17
-0.18
-0.18
-0.20
-0.21
-0.22
-0.22
-0.26
-0.26
-0.28
-0.31
-0.32
-0.34
-0.42
-0.44
-0.47
-0.47
-0.48
-0.60
-0.77
-0.78
-0.84
-0.85
0.29
0.07
-0.08
-0.20
-0.37
0.54
0.42
0.42
0.42
0.38
0.36
0.24
0.15
0.00
-0.03
-0.07
-0.10
-0.14
-0.15
-0.29
-0.42
0.36

| |
|---|
| 0.28 |
| 0.12 |
| 0.05 |
| -0.02 |
| -0.27 |
| -0.29 |
| -0.48 |
| 0.32 |
| 0.23 |
| -0.04 |
| -0.09 |
| 0.23 |
| 0.13 |
| 0.08 |
| -0.38 |
| -0.44 |
| 0.19 |
| 0.05 |
| -0.38 |
| -0.56 |
| -0.58 |
| 0.19 |
| 0.06 |
| 0.30 |
| 0.11 |
| -0.26 |
| -0.53 |
| -0.28 |
| 0.09 |
| 0.04 |
| 0.03 |
| -0.42 |
| 0.59 |
| 0.58 |
| 0.45 |
| 0.45 |
| 0.42 |
| 0.41 |
| 0.30 |
| 0.28 |
| 0.25 |
| 0.23 |
| 0.18 |
| 0.15 |
| 0.12 |
| 0.09 |
| 0.08 |
| 0.06 |

| |
|---|
| 0.06 |
| 0.04 |
| 0.02 |
| 0.01 |
| 0.01 |
| -0.01 |
| -0.01 |
| -0.02 |
| -0.02 |
| -0.04 |
| -0.04 |
| -0.04 |
| -0.08 |
| -0.10 |
| -0.10 |
| -0.15 |
| -0.16 |
| -0.16 |
| -0.17 |
| -0.17 |
| -0.17 |
| -0.27 |
| -0.27 |
| -0.34 |
| -0.38 |
| -0.39 |
| -0.41 |
| -0.41 |
| 0.27 |
| 0.28 |
| 0.24 |
| 0.20 |
| 0.01 |
| -0.22 |
| -0.26 |
| -0.25 |
| 0.56 |
| 0.51 |
| 0.39 |
| 0.37 |
| 0.31 |
| 0.30 |
| 0.18 |
| 0.08 |
| 0.05 |
| -0.07 |
| -0.08 |
| -0.25 |

| |
|---|
| 0.22 |
| -0.01 |
| 0.61 |
| 0.51 |
| 0.06 |
| -0.25 |
| -0.21 |
| -0.58 |
| 0.09 |
| 0.00 |
| -0.24 |
| -0.42 |
| 0.09 |
| 0.08 |
| 0.07 |
| 0.05 |
| 0.30 |
| 0.21 |
| -0.14 |
| -0.14 |
| -0.24 |
| -0.25 |
| 1.31 |
| 0.13 |
| -0.27 |
| -0.33 |
| -0.45 |
| -0.66 |
| -0.68 |
| -0.15 |
| -0.15 |
| -0.08 |
| 0.09 |
| -0.25 |
| 0.23 |
| 0.21 |
| -0.03 |
| -0.43 |
| -0.01 |
| -0.06 |
| -0.20 |
| 0.45 |
| 0.14 |
| 0.13 |
| -0.06 |
| -0.06 |
| -0.09 |
| -0.20 |

| |
|---|
| -0.30 |
| -0.31 |
| -0.32 |
| -0.54 |
| -0.59 |
| 0.15 |
| -0.24 |
| 0.19 |
| -0.58 |
| 0.33 |
| 0.19 |
| 0.06 |
| 0.02 |
| -0.04 |
| -0.18 |
| -0.31 |
| -0.45 |
| -1.15 |
| 1.56 |
| 1.48 |
| 1.27 |
| 0.50 |
| 0.32 |
| 0.20 |
| 0.19 |
| 0.14 |
| 0.06 |
| 0.06 |
| 0.00 |
| 0.00 |
| -0.01 |
| -0.02 |
| -0.02 |
| -0.08 |
| -0.09 |
| -0.10 |
| -0.10 |
| -0.11 |
| -0.22 |
| -0.27 |
| -0.27 |
| -0.29 |
| 0.44 |
| 0.41 |
| 0.30 |
| 0.27 |
| 0.19 |
| 0.19 |

| |
|---|
| 0.11 |
| 0.10 |
| 0.09 |
| 0.08 |
| 0.05 |
| 0.00 |
| -0.03 |
| -0.05 |
| -0.11 |
| -0.12 |
| -0.13 |
| -0.17 |
| -0.20 |
| -0.22 |
| -0.28 |
| -0.29 |
| -0.36 |
| 0.38 |
| 0.09 |
| 0.01 |
| -0.26 |
| -0.26 |
| -0.29 |
| -0.44 |
| 0.49 |
| -0.27 |
| -0.66 |
| -0.05 |
| -0.24 |
| 0.39 |
| 0.07 |
| 0.00 |
| -0.10 |
| -0.27 |
| -0.27 |
| -0.32 |
| -0.38 |
| 0.15 |
| 0.11 |
| -0.03 |
| 0.42 |
| 0.38 |
| 0.30 |
| 0.29 |
| 0.23 |
| 0.01 |
| -0.25 |
| -0.31 |

-0.33
-0.39
0.85
0.80
0.65
0.63
0.61
0.59
0.34
0.31
0.29
0.27
0.23
0.22
0.19
0.16
0.15
0.04
0.00
-0.07
-0.18
-0.21
-0.28
-0.36
-0.37
-0.38
-0.85
-1.35
-0.09
-0.24
-0.26
0.71
0.49
0.40
0.20
0.20
0.11
0.10
0.08
0.07
0.06
0.01
0.01
0.01
0.00
-0.01
-0.02
-0.06

| |
|---|
| -0.09 |
| -0.10 |
| -0.11 |
| -0.13 |
| -0.16 |
| -0.17 |
| -0.22 |
| -0.26 |
| -0.27 |
| -0.27 |
| -0.30 |
| -0.33 |
| -0.35 |
| -0.45 |
| -0.51 |
| -0.78 |
| -0.17 |
| 0.23 |
| -0.55 |
| 0.64 |
| 0.20 |
| 0.06 |
| 0.16 |
| 0.13 |
| -0.37 |
| -0.03 |
| -0.21 |
| -0.28 |
| 0.37 |
| 0.15 |
| 0.13 |
| -0.06 |
| -0.19 |
| -0.12 |
| -0.13 |
| -0.24 |
| -0.24 |
| -0.41 |
| 0.14 |
| -0.25 |
| 0.44 |
| 0.17 |
| -0.22 |
| 0.13 |
| -0.14 |
| -0.28 |
| 0.23 |
| 0.18 |

-0.01
-0.02
-0.13
-0.23
-0.08
1.00
0.90
0.57
0.51
0.48
0.30
0.09
0.08
0.06
-0.18
0.45
0.31
0.18
0.15
0.09
0.01
-0.01
-0.04
-0.13
-0.30
-0.40
0.16
0.05
-0.47
-0.07
0.05
-0.13
0.66
0.32
0.14
0.13
0.08
0.07
0.02
-0.01
-0.18
-0.20
-0.27
-0.27
0.22
0.40
0.08
0.01

-0.01
-0.10
-0.19
-0.41
-0.43
0.35
0.09
0.04
-0.10
-0.18
-0.09
0.15
0.10
0.07
0.05
0.04
0.03
-0.03
-0.05
-0.09
-0.12
-0.16
-0.20
-0.21
-0.71
-0.77
-0.79
0.45
0.43
0.29
-0.31
-0.15
0.04
-0.13
0.39
0.15
-0.19
0.54
0.44
-0.73
0.40
0.39
0.10
-0.22
1.06
0.39
0.16
-0.22

| |
|---|
| 0.93 |
| 0.87 |
| 0.58 |
| -0.04 |
| -0.18 |
| 0.87 |
| 0.79 |
| 0.78 |
| 0.71 |
| 0.50 |
| 0.48 |
| 0.43 |
| 0.29 |
| 0.25 |
| 0.22 |
| 0.21 |
| 0.21 |
| 0.21 |
| 0.20 |
| 0.16 |
| 0.11 |
| 0.06 |
| 0.05 |
| 0.01 |
| -0.02 |
| -0.02 |
| -0.03 |
| -0.05 |
| -0.06 |
| -0.09 |
| -0.13 |
| -0.21 |
| -0.27 |
| -0.29 |
| -0.32 |
| -0.33 |
| -0.37 |
| 0.05 |
| -0.08 |
| -0.13 |
| -0.47 |
| 0.35 |
| -0.40 |
| -0.09 |
| 0.12 |
| 0.15 |
| 0.08 |
| -0.09 |

-0.23
-0.33
-0.36
-0.43
0.35
0.38
0.25
-0.10
-0.14
-0.30
0.11
0.08
-0.12
-0.31
-0.57
0.61
0.34
-0.18
0.67
0.58
0.05
0.28
0.13
-0.15
-0.45
0.06
0.02
-0.10
-0.10
-0.17
-0.30
0.10
-0.19
-0.20
-0.23
-0.25
-0.26
-0.30
-0.31
-0.33
-0.38
-0.62
0.27
0.06
-0.11
0.10
-0.16
-0.16

| |
|---|
| 0.56 |
| 0.16 |
| 0.14 |
| 0.13 |
| 0.12 |
| 0.12 |
| 0.03 |
| 0.01 |
| -0.01 |
| -0.02 |
| -0.07 |
| -0.21 |
| -0.23 |
| -0.37 |
| -0.41 |
| -0.63 |
| 0.04 |
| 0.15 |
| -0.28 |
| 0.73 |
| -0.06 |
| -0.15 |
| -0.35 |
| 0.18 |
| 0.13 |
| 0.14 |
| -0.19 |
| -0.41 |
| -0.58 |
| 0.00 |
| -0.32 |
| 0.66 |
| 0.59 |
| -0.15 |
| 1.13 |
| 0.24 |
| 0.13 |
| 0.04 |
| 0.00 |
| -0.06 |
| -0.09 |
| -0.04 |
| -0.21 |
| -0.25 |
| -0.26 |
| -0.33 |
| -0.38 |
| -0.41 |

| |
|---|
| -0.42 |
| 0.10 |
| 1.08 |
| -0.03 |
| -0.06 |
| -0.07 |
| -0.11 |
| -0.13 |
| -0.13 |
| -0.13 |
| -0.14 |
| -0.17 |
| -0.19 |
| -0.19 |
| -0.19 |
| -0.23 |
| -0.25 |
| -0.25 |
| -0.28 |
| -0.29 |
| -0.30 |
| -0.34 |
| -0.35 |
| -0.37 |
| -0.41 |
| -0.44 |
| -0.44 |
| -0.44 |
| -0.47 |
| -0.49 |
| -0.54 |
| -0.55 |
| -0.55 |
| -0.59 |
| -0.69 |
| -0.72 |
| -0.78 |
| -0.82 |
| -0.87 |
| 1.21 |
| 1.20 |
| 0.21 |
| 0.06 |
| 0.50 |
| 0.29 |
| 0.25 |
| 0.06 |
| 0.06 |

| |
|---|
| -0.13 |
| -0.32 |
| 0.05 |
| -0.04 |
| 0.20 |
| 0.05 |
| 0.05 |
| -0.05 |
| -0.23 |
| 0.35 |
| 0.25 |
| -0.04 |
| -0.15 |
| -0.34 |
| -0.53 |
| -0.13 |
| -0.13 |
| 0.41 |
| 0.00 |
| 0.32 |
| 0.01 |
| 0.24 |
| 0.24 |
| 0.35 |
| 0.31 |
| 0.10 |
| -0.21 |
| -0.38 |
| 0.02 |
| 0.02 |
| 1.44 |
| 0.60 |
| 0.16 |
| 0.80 |
| 0.02 |
| 0.68 |
| 0.51 |
| 0.03 |
| -0.46 |
| 0.90 |
| 0.42 |
| 0.13 |
| 0.09 |
| -0.33 |
| 0.11 |
| -0.02 |
| -0.13 |
| -0.42 |

| |
|---|
| -0.50 |
| -0.47 |
| -0.05 |
| -0.15 |
| 0.11 |
| -0.18 |
| -0.38 |
| 0.49 |
| 0.49 |
| 0.16 |
| -0.52 |
| 0.36 |
| 0.24 |
| 0.14 |
| 0.14 |
| 0.11 |
| 0.08 |
| 0.02 |
| 0.00 |
| -0.04 |
| -0.11 |
| -0.14 |
| -0.34 |
| 0.32 |
| -0.01 |
| 0.48 |
| -0.16 |
| 0.52 |
| 0.63 |
| -0.30 |
| -0.47 |
| -0.58 |
| 0.08 |
| 0.30 |
| 0.20 |
| -0.49 |
| 0.01 |
| -0.13 |
| -0.34 |
| -0.58 |
| 0.70 |
| 0.67 |
| 0.51 |
| 0.13 |
| -0.05 |
| -0.22 |
| 0.35 |
| 0.05 |

| |
|---|
| 0.31 |
| 0.78 |
| 0.66 |
| 0.46 |
| 0.46 |
| 0.39 |
| 0.36 |
| 0.35 |
| 0.32 |
| 0.27 |
| 0.27 |
| 0.03 |
| -0.04 |
| -0.32 |
| 0.12 |
| 0.10 |
| 0.18 |
| 0.16 |
| -0.39 |
| 0.21 |
| -0.01 |
| -0.11 |
| -0.27 |
| -0.28 |
| -0.34 |
| -0.72 |
| 0.08 |
| -0.11 |
| -0.19 |
| -0.22 |
| -0.27 |
| 0.06 |
| -0.03 |
| -0.27 |
| -0.31 |
| -0.38 |
| -0.56 |
| -0.67 |
| 0.31 |
| -0.39 |
| -0.41 |
| -0.42 |
| -0.45 |
| -0.61 |
| -0.61 |
| -0.62 |
| -0.65 |
| -0.68 |

| |
|---|
| -0.79 |
| -0.95 |
| -0.96 |
| -1.25 |
| 0.13 |
| -0.32 |
| -0.35 |
| -0.38 |
| 0.95 |
| 0.28 |
| 0.23 |
| 0.15 |
| 0.13 |
| 0.11 |
| 0.10 |
| 0.08 |
| 0.07 |
| 0.06 |
| 0.06 |
| 0.05 |
| 0.04 |
| 0.02 |
| 0.01 |
| 0.01 |
| -0.04 |
| -0.04 |
| -0.06 |
| -0.06 |
| -0.06 |
| -0.06 |
| -0.06 |
| -0.11 |
| -0.12 |
| -0.13 |
| -0.15 |
| -0.17 |
| -0.18 |
| -0.19 |
| -0.20 |
| -0.23 |
| -0.25 |
| -0.26 |
| -0.27 |
| -0.31 |
| -0.31 |
| -0.31 |
| -0.31 |
| -0.31 |

-0.32
-0.35
-0.46
-0.50
-0.51
-0.54
-0.54
-0.58
-0.71
-0.97
0.41
0.03
-0.06
0.16
-0.48
0.37
0.25
0.23
0.16
0.09
0.01
-0.11
-0.12
-0.19
-0.32
-0.37
-0.42
0.38
0.11
-0.08
-0.10
-0.13
-0.24
0.27
-0.15
-0.16
-0.25
-0.22
0.61
0.60
0.47
0.38
0.36
0.28
0.22
0.16
0.15
0.09

| |
|---|
| 0.05 |
| 0.03 |
| 0.01 |
| 0.00 |
| 0.00 |
| 0.00 |
| -0.03 |
| -0.07 |
| -0.15 |
| -0.22 |
| -0.30 |
| -0.31 |
| -0.34 |
| -0.34 |
| -0.40 |
| 0.36 |
| 0.20 |
| 0.13 |
| 0.08 |
| -0.03 |
| -0.10 |
| -0.11 |
| 0.38 |
| 0.35 |
| 0.32 |
| 0.23 |
| 0.23 |
| 0.18 |
| 0.12 |
| 0.08 |
| -0.01 |
| -0.01 |
| -0.09 |
| -0.10 |
| -0.10 |
| -0.21 |
| -0.23 |
| -0.29 |
| -0.31 |
| -0.35 |
| -0.47 |
| 0.76 |
| -0.02 |
| 0.19 |
| 0.10 |
| 0.06 |
| -0.07 |
| -0.15 |

| |
|---|
| -0.26 |
| -0.32 |
| -0.65 |
| -0.66 |
| 0.57 |
| 0.16 |
| 0.08 |
| 0.07 |
| -0.10 |
| -0.13 |
| -0.14 |
| -0.24 |
| -0.27 |
| 0.74 |
| 0.49 |
| 0.38 |
| 0.21 |
| 0.37 |
| 0.33 |
| 0.28 |
| 0.47 |
| 0.31 |
| 0.30 |
| 0.03 |
| 0.02 |
| -0.16 |
| 0.29 |
| 0.48 |
| -0.01 |
| 0.00 |
| -0.02 |
| -0.12 |
| -0.25 |
| 0.32 |
| -0.49 |
| 0.33 |
| 0.25 |
| -0.16 |
| 0.39 |
| 0.24 |
| 0.06 |
| -0.04 |
| -0.08 |
| 0.21 |
| -0.41 |
| -0.69 |
| 0.33 |
| 0.33 |

-0.02
-0.52
-0.66
-0.25
-0.40
0.09
0.05
-0.17
0.21
-0.06
0.06
-0.09
0.16
0.12
0.07
0.07
0.36
0.20
0.34
0.29
-0.04
-0.61
-0.06
-0.26
-0.42
-0.72
0.25
0.16
-0.05
-0.17
-0.36
-0.04
0.17
0.15
-0.01
-0.29
-0.29
0.52
0.17
-0.20
0.05
0.24
0.17
-0.03
-0.05
-0.12
-0.12
-0.36

| |
|---|
| -0.45 |
| 0.04 |
| 0.01 |
| -0.07 |
| -0.86 |
| -0.28 |
| -0.29 |
| 0.06 |
| -0.45 |
| 0.29 |
| 0.04 |
| -0.41 |
| 0.34 |
| 0.13 |
| -0.47 |
| -0.17 |
| 0.32 |
| 0.36 |
| 0.30 |
| 0.28 |
| 0.24 |
| 0.09 |
| 0.06 |
| 0.01 |
| -0.06 |
| -0.07 |
| -0.10 |
| -0.17 |
| -0.22 |
| -0.31 |
| 0.05 |
| -0.16 |
| -0.31 |
| -0.21 |
| -0.13 |
| -0.34 |
| 0.51 |
| 0.13 |
| 0.00 |
| 0.00 |
| 0.08 |
| -0.09 |
| 0.33 |
| -0.19 |
| 0.80 |
| 0.13 |
| -0.03 |
| -0.18 |

| |
|---|
| 0.22 |
| 0.12 |
| -0.06 |
| 1.03 |
| 0.72 |
| 0.62 |
| 0.24 |
| 0.24 |
| 0.22 |
| 0.20 |
| 0.18 |
| 0.15 |
| 0.06 |
| 0.06 |
| 0.05 |
| 0.04 |
| 0.01 |
| -0.03 |
| -0.04 |
| -0.06 |
| -0.13 |
| -0.20 |
| -0.24 |
| -0.28 |
| -0.39 |
| -0.56 |
| -0.60 |
| -0.60 |
| -0.63 |
| 0.24 |
| 0.17 |
| -0.09 |
| -0.10 |
| 0.03 |
| -0.01 |
| -0.03 |
| -0.12 |
| -0.28 |
| -0.04 |
| -0.06 |
| -0.17 |
| -0.36 |
| -0.53 |
| 0.27 |
| -0.11 |
| -0.32 |
| -0.36 |
| -0.37 |

| |
|---|
| 0.21 |
| -0.38 |
| 0.38 |
| 0.42 |
| 0.23 |
| 0.04 |
| 0.01 |
| -0.04 |
| -0.09 |
| -0.14 |
| -0.15 |
| -0.17 |
| -0.27 |
| -0.46 |
| -0.47 |
| -0.58 |
| -0.81 |
| -0.03 |
| 0.31 |
| 0.30 |
| -0.09 |
| -0.10 |
| 0.33 |
| 0.12 |
| 0.06 |
| 0.03 |
| 0.00 |
| -0.08 |
| -0.08 |
| -0.09 |
| -0.10 |
| -0.11 |
| -0.13 |
| -0.13 |
| -0.14 |
| -0.14 |
| -0.15 |
| -0.19 |
| -0.22 |
| -0.23 |
| -0.28 |
| -0.31 |
| -0.33 |
| -0.36 |
| -0.48 |
| -0.54 |
| -0.63 |
| -0.74 |

-0.12
0.19
0.29
-0.15
0.07
0.46
0.26
0.07
0.06
-0.07
-0.20
-0.21
-0.23
-0.41
0.27
0.26
-0.05
-0.10
1.14
0.47
0.67
0.42
0.19
0.04
-0.04
-0.11
-0.18
-0.22
-0.25
-0.25
-0.25
-0.29
-0.34
-0.36
-0.44
-0.49
-0.54
-1.03
0.19
0.09
0.05
-0.05
-0.07
-0.17
-0.41
0.08
-1.18
0.50

| |
|---|
| 0.08 |
| 0.01 |
| -0.07 |
| -0.27 |
| -0.38 |
| -0.40 |
| 0.40 |
| 0.21 |
| 0.05 |
| 0.01 |
| -0.04 |
| -0.09 |
| -0.10 |
| -0.41 |
| -0.46 |
| -0.47 |
| 0.43 |
| 0.30 |
| 0.18 |
| 0.15 |
| 0.11 |
| 0.01 |
| -0.09 |
| -0.10 |
| -0.18 |
| -0.36 |
| 0.52 |
| 0.28 |
| 0.13 |
| 0.12 |
| 0.11 |
| 0.10 |
| 0.01 |
| -0.12 |
| -0.28 |
| 0.52 |
| 0.37 |
| 0.22 |
| 0.11 |
| 0.02 |
| 0.00 |
| -0.01 |
| -0.01 |
| -0.09 |
| -0.12 |
| -0.16 |
| -0.21 |
| -0.36 |

-0.36
-0.36
-0.41
-0.48
-0.50
-0.51
-0.53
-0.53
-0.57
-1.05
0.28
0.22
0.16
0.12
0.11
0.06
-0.01
-0.08
-0.10
-0.15
-0.22
-0.34
-0.49
-0.51
-0.03
-0.22
-0.29
-0.31
-0.35
-0.54
0.26
-0.25
-0.30
-0.28
-0.47
-0.57
-0.61
-0.62
-0.84
0.11
0.05
-0.11
-0.26
-0.22
-0.03
-0.01
0.26
0.00

-0.29
0.36
0.16
0.08
-0.05
-0.08
-0.39
-0.06
-0.11
-0.18
0.33
0.19
0.15
0.14
0.10
1.09
0.69
0.34
0.08
-0.21
-0.32
-0.36
-0.45
-0.45
0.43
0.04
0.33
0.17
0.17
0.10
-0.06
-0.08
-0.40
-0.02
-0.09
0.96
0.26
0.15
0.06
0.04
-0.02
-0.02
-0.06
-0.07
-0.10
-0.19
-0.22
-0.23

| |
|---|
| -0.23 |
| -0.33 |
| -0.34 |
| -0.49 |
| 0.56 |
| 0.27 |
| -0.26 |
| 0.07 |
| -0.06 |
| -0.43 |
| 0.15 |
| 0.14 |
| 0.08 |
| 0.05 |
| 0.00 |
| -0.05 |
| -0.20 |
| -0.46 |
| -0.21 |
| -0.39 |
| -0.44 |
| -0.49 |
| -0.04 |
| -0.27 |
| 0.01 |
| -0.02 |
| -0.07 |
| -0.17 |
| -0.20 |
| 0.46 |
| 0.11 |
| -0.12 |
| -0.15 |
| -0.16 |
| -0.61 |
| 0.43 |
| 0.24 |
| 0.21 |
| 0.20 |
| 0.09 |
| 0.05 |
| 0.04 |
| 0.01 |
| -0.01 |
| -0.01 |
| -0.11 |
| -0.14 |
| -0.17 |

| |
|---|
| -0.21 |
| 0.32 |
| 0.24 |
| 0.11 |
| 0.10 |
| 0.02 |
| -0.16 |
| -0.82 |
| -0.04 |
| -0.15 |
| 0.26 |
| 0.10 |
| 0.26 |
| -0.06 |
| -0.49 |
| 0.37 |
| 0.07 |
| -0.02 |
| -0.12 |
| -0.14 |
| -0.33 |
| -0.34 |
| -0.42 |
| -0.55 |
| 0.41 |
| 0.40 |
| 0.00 |
| -0.12 |
| 0.18 |
| -0.04 |
| 1.06 |
| 0.86 |
| 0.80 |
| 0.22 |
| 0.12 |
| -0.03 |
| -0.18 |
| -0.22 |
| -0.33 |
| 0.08 |
| -0.02 |
| -0.01 |
| 0.14 |
| -0.74 |
| 0.28 |
| 0.13 |
| 0.09 |
| 0.00 |

| |
|---|
| -0.10 |
| -0.10 |
| -0.16 |
| -0.62 |
| 0.39 |
| 0.28 |
| 0.10 |
| 0.15 |
| -0.21 |
| 0.45 |
| -0.33 |
| -0.08 |
| -0.01 |
| -0.01 |
| -0.24 |
| -0.31 |
| -0.02 |
| 0.50 |
| 0.38 |
| 0.04 |
| -0.06 |
| -0.07 |
| -0.25 |
| 1.34 |
| 0.40 |
| 0.36 |
| 0.36 |
| 0.23 |
| 0.18 |
| 0.15 |
| 0.11 |
| 0.09 |
| 0.01 |
| 0.00 |
| -0.07 |
| -0.13 |
| -0.13 |
| -0.22 |
| -0.23 |
| -0.30 |
| -0.36 |
| -0.48 |
| -1.01 |
| 0.39 |
| -0.16 |
| 0.34 |
| 0.26 |
| 0.25 |

| |
|---|
| 0.19 |
| 0.16 |
| 0.14 |
| 0.12 |
| 0.08 |
| -0.01 |
| -0.09 |
| -0.11 |
| -0.12 |
| -0.18 |
| -0.27 |
| 0.03 |
| 0.58 |
| -0.07 |
| 0.18 |
| 0.14 |
| 0.05 |
| 0.02 |
| 0.02 |
| -0.25 |
| -0.19 |
| 0.21 |
| 0.19 |
| -0.06 |
| -0.35 |
| 0.46 |
| -0.06 |
| 0.16 |
| 0.06 |
| 0.07 |
| 0.02 |
| -0.01 |
| -0.24 |
| 0.33 |
| 0.00 |
| -0.08 |
| -0.14 |
| -0.15 |
| 0.11 |
| -0.03 |
| -0.05 |
| -0.29 |
| -0.33 |
| 0.06 |
| 0.04 |
| 0.04 |
| 0.04 |
| -0.20 |

| |
|---|
| -0.29 |
| 0.17 |
| 0.65 |
| 0.21 |
| -0.08 |
| -0.15 |
| -0.11 |
| 0.35 |
| -0.36 |
| -0.63 |
| 0.10 |
| -0.23 |
| -0.28 |
| -0.10 |
| -0.10 |
| -0.17 |
| -0.27 |
| -0.73 |
| 0.25 |
| 0.16 |
| 0.38 |
| -0.18 |
| 0.05 |
| -0.19 |
| -0.23 |
| -0.61 |
| -0.93 |
| 0.20 |
| -0.08 |
| -0.44 |
| -0.50 |
| -0.10 |
| -0.20 |
| 0.00 |
| 0.00 |
| -0.16 |
| 0.31 |
| 0.16 |
| -0.02 |
| -0.11 |
| -0.14 |
| -0.31 |
| -0.43 |
| 0.11 |
| 0.07 |
| 0.00 |
| -0.03 |
| -0.37 |

| |
|---|
| -0.38 |
| -0.45 |
| -0.36 |
| -0.36 |
| -0.50 |
| -0.09 |
| 0.20 |
| -0.06 |
| 0.28 |
| -0.24 |
| -0.25 |
| -0.63 |
| 0.45 |
| 0.32 |
| 0.25 |
| 0.22 |
| 0.07 |
| 0.06 |
| 0.02 |
| -0.13 |
| -0.14 |
| -0.15 |
| -0.18 |
| -0.22 |
| -0.26 |
| -0.76 |
| 0.12 |
| 0.04 |
| 0.20 |
| 0.34 |
| -0.32 |
| 0.05 |
| -0.05 |
| 0.13 |
| 0.17 |
| 0.09 |
| -0.02 |
| -0.14 |
| -0.20 |
| 0.17 |
| 0.36 |
| 0.34 |
| -0.09 |
| -0.23 |
| 0.59 |
| 0.57 |
| 0.24 |
| 0.39 |

| |
|---|
| 0.08 |
| -0.25 |
| -0.43 |
| -0.46 |
| -0.31 |
| 0.03 |
| 0.03 |
| -0.01 |
| -0.30 |
| -0.02 |
| 0.22 |
| 0.03 |
| -0.05 |
| -0.11 |
| -0.17 |
| -0.18 |
| -0.28 |
| -0.31 |
| -0.31 |
| -0.33 |
| -0.37 |
| -0.44 |
| -0.50 |
| -0.70 |
| -0.76 |
| 0.28 |
| 0.28 |
| 0.20 |
| 0.17 |
| 0.17 |
| 0.15 |
| 0.13 |
| 0.12 |
| 0.07 |
| 0.05 |
| 0.02 |
| 0.01 |
| 0.01 |
| 0.00 |
| -0.01 |
| -0.01 |
| -0.01 |
| -0.01 |
| -0.02 |
| -0.03 |
| -0.03 |
| -0.04 |
| -0.05 |

-0.07
-0.13
-0.14
-0.18
-0.19
-0.29
-0.30
-0.30
-0.39
-0.43
-0.55
-0.63
-0.70
-0.81
0.27
0.89
-0.07
-0.23
0.41
0.21
0.10
0.01
0.37
0.26
0.23
0.09
0.92
-0.09
-0.72
0.48
0.33
-0.04
-0.05
-0.18
-0.20
-0.30
-0.52
-0.25
-0.57
0.23
0.17
0.02
-0.35
0.07
-0.04
-0.23
-0.79
-0.93

| |
|---|
| 0.05 |
| 0.07 |
| 1.48 |
| 0.59 |
| 0.50 |
| 0.50 |
| 0.46 |
| 0.46 |
| 0.31 |
| 0.27 |
| 0.24 |
| 0.21 |
| 0.19 |
| 0.18 |
| 0.09 |
| 0.09 |
| 0.08 |
| 0.08 |
| 0.07 |
| 0.01 |
| 0.00 |
| 0.00 |
| -0.01 |
| -0.03 |
| -0.03 |
| -0.03 |
| -0.05 |
| -0.05 |
| -0.06 |
| -0.06 |
| -0.07 |
| -0.08 |
| -0.11 |
| -0.11 |
| -0.12 |
| -0.12 |
| -0.13 |
| -0.16 |
| -0.17 |
| -0.23 |
| -0.24 |
| -0.31 |
| -0.42 |
| -0.44 |
| -0.57 |
| -1.67 |
| -1.70 |
| 0.76 |

| |
|---|
| 0.63 |
| 0.59 |
| 0.33 |
| 0.21 |
| 0.13 |
| 0.12 |
| 0.06 |
| 0.03 |
| 0.01 |
| -0.11 |
| -0.24 |
| -0.15 |
| -0.50 |
| -0.44 |
| -0.49 |
| -0.59 |
| -0.62 |
| 0.04 |
| 0.29 |
| 0.51 |
| 0.36 |
| 0.21 |
| 0.47 |
| -0.07 |
| -0.04 |
| 0.18 |
| -0.04 |
| -0.10 |
| -0.23 |
| 0.20 |
| 0.34 |
| -0.28 |
| 0.10 |
| 0.08 |
| -0.11 |
| -0.23 |
| -0.37 |
| -0.56 |
| -0.08 |
| -0.17 |
| -0.25 |
| 0.12 |
| -0.11 |
| -0.17 |
| 0.41 |
| -0.18 |
| -0.24 |
| 0.02 |

-0.09
0.27
0.21
0.16
0.12
0.08
0.03
0.02
-0.35
-0.53
2.49
1.31
0.77
0.59
0.53
0.42
0.29
0.27
0.24
0.23
0.22
0.19
0.18
0.13
0.13
0.01
0.01
0.00
-0.07
-0.15
-0.63
-0.80
0.24
0.14
0.11
0.06
-0.18
-0.59
-0.27
0.28
0.05
0.53
0.17
0.00
0.59
0.52
0.12
0.06

| |
|---|
| -0.24 |
| 0.01 |
| 0.49 |
| 0.33 |
| 0.33 |
| 0.32 |
| 0.11 |
| 0.08 |
| 0.03 |
| -0.01 |
| -0.05 |
| -0.06 |
| -0.06 |
| -0.07 |
| -0.10 |
| -0.13 |
| -0.14 |
| -0.15 |
| -0.17 |
| -0.18 |
| -0.20 |
| -0.23 |
| -0.27 |
| -0.34 |
| -0.36 |
| -0.40 |
| 0.38 |
| 0.17 |
| 0.02 |
| -0.05 |
| -0.10 |
| -0.16 |
| -0.20 |
| -0.29 |
| 0.14 |
| 0.77 |
| 0.53 |
| 0.40 |
| 0.26 |
| 0.22 |
| 0.18 |
| 0.16 |
| 0.15 |
| 0.14 |
| 0.12 |
| 0.12 |
| 0.07 |
| 0.07 |

| |
|---|
| 0.06 |
| 0.06 |
| 0.05 |
| -0.02 |
| -0.03 |
| -0.03 |
| -0.04 |
| -0.04 |
| -0.05 |
| -0.09 |
| -0.09 |
| -0.10 |
| -0.10 |
| -0.10 |
| -0.11 |
| -0.12 |
| -0.12 |
| -0.13 |
| -0.14 |
| -0.17 |
| -0.18 |
| -0.24 |
| -0.25 |
| -0.27 |
| -0.29 |
| -0.33 |
| -0.36 |
| -0.94 |
| -0.99 |
| 0.85 |
| 0.47 |
| 0.29 |
| 0.27 |
| 0.18 |
| 0.00 |
| -0.03 |
| -0.03 |
| -0.03 |
| -0.04 |
| -0.05 |
| -0.06 |
| -0.09 |
| -0.11 |
| -0.12 |
| -0.13 |
| -0.15 |
| -0.16 |
| -0.17 |

-0.18
-0.18
-0.22
-0.25
-0.25
-0.25
-0.28
-0.31
-0.38
-0.39
-0.48
-0.54
0.22
-0.51
0.22
0.01
-0.06
-0.16
-0.31
-0.45
-0.02
-0.03
-0.36
-0.09
-0.26
-0.60
-0.84
-0.49
0.25
0.01
-0.01
-0.07
-0.07
0.27
0.16
0.83
0.78
0.74
0.64
0.46
0.25
0.24
0.20
0.19
0.17
0.12
0.11
0.08

| |
|---|
| 0.04 |
| 0.04 |
| 0.02 |
| 0.01 |
| 0.00 |
| 0.00 |
| -0.01 |
| -0.01 |
| -0.03 |
| -0.04 |
| -0.04 |
| -0.06 |
| -0.09 |
| -0.15 |
| -0.22 |
| -0.61 |
| 0.19 |
| 1.78 |
| 0.06 |
| -0.23 |
| 0.17 |
| 0.12 |
| 0.10 |
| 0.04 |
| -0.08 |
| 0.25 |
| 0.22 |
| -0.05 |
| -0.29 |
| 0.82 |
| 0.73 |
| 0.24 |
| 0.30 |
| 0.30 |
| 0.21 |
| 0.20 |
| 0.17 |
| 0.15 |
| 0.14 |
| 0.12 |
| 0.10 |
| 0.09 |
| 0.08 |
| 0.08 |
| 0.07 |
| 0.07 |
| 0.07 |
| 0.05 |

| |
|---|
| 0.04 |
| 0.00 |
| 0.00 |
| -0.07 |
| -0.13 |
| -0.13 |
| -0.16 |
| -0.26 |
| -0.27 |
| -0.18 |
| -0.46 |
| 0.12 |
| 0.08 |
| -0.02 |
| 0.20 |
| -0.01 |
| -0.09 |
| -0.22 |
| -0.23 |
| -0.35 |
| -0.37 |
| -0.37 |
| -0.38 |
| -0.39 |
| -0.47 |
| -0.48 |
| -0.50 |
| -0.83 |
| -0.86 |
| 0.00 |
| 0.14 |
| 0.14 |
| -0.35 |
| -0.42 |
| 0.79 |
| 0.00 |
| 0.26 |
| 0.04 |
| 0.01 |
| 0.01 |
| -0.01 |
| -0.07 |
| -0.07 |
| -0.08 |
| -0.22 |
| -0.36 |
| -0.41 |
| 0.08 |

| |
|---|
| -0.34 |
| 0.62 |
| 0.32 |
| 0.07 |
| -0.07 |
| 0.24 |
| -0.09 |
| -0.10 |
| -0.39 |
| 0.14 |
| -0.04 |
| 0.65 |
| 0.40 |
| 0.08 |
| -0.17 |
| -0.24 |
| -0.24 |
| -0.06 |
| -0.04 |
| -0.15 |
| -0.20 |
| -0.33 |
| -0.17 |
| 0.10 |
| 0.06 |
| 2.73 |
| 0.34 |
| 0.15 |
| 0.14 |
| -0.01 |
| -0.05 |
| -0.05 |
| -0.06 |
| -0.07 |
| -0.10 |
| -0.14 |
| -0.18 |
| -0.29 |
| -0.35 |
| -0.36 |
| -0.40 |
| -0.41 |
| -0.45 |
| -0.57 |
| -0.76 |
| -0.76 |
| 0.39 |
| 0.16 |

| |
|---|
| -0.02 |
| -0.03 |
| -0.20 |
| -0.24 |
| 0.38 |
| -0.10 |
| 0.07 |
| 0.33 |
| 0.01 |
| -0.03 |
| 0.46 |
| 0.14 |
| -0.22 |
| -0.35 |
| 1.55 |
| 0.37 |
| 0.35 |
| 0.28 |
| 0.21 |
| 0.21 |
| 0.01 |
| -0.01 |
| -0.09 |
| -0.28 |
| -0.28 |
| -0.35 |
| -0.49 |
| 2.03 |
| -0.18 |
| -0.27 |
| 0.34 |
| 0.30 |
| 0.29 |
| 0.25 |
| 0.24 |
| 0.15 |
| 0.11 |
| 0.09 |
| 0.08 |
| 0.08 |
| 0.07 |
| 0.03 |
| -0.01 |
| -0.01 |
| -0.02 |
| -0.03 |
| -0.04 |
| -0.04 |

-0.08
-0.09
-0.10
-0.13
-0.15
-0.20
-0.21
-0.22
-0.26
-0.26
-0.27
-0.27
-0.28
-0.33
-0.38
-0.64
0.59
0.54
0.39
0.36
0.15
0.09
0.06
0.02
-0.08
-0.08
-0.19
-0.23
-0.24
-0.27
-0.29
-0.41
-0.63
-0.47
0.55
0.54
-0.23
-0.27
2.05
0.11
0.00
0.00
-0.08
-0.11
-0.44
-0.46
0.35
0.11

-0.23
0.17
0.13
0.24
-0.17
-0.20
-0.44
0.05
0.05
0.24
0.17
0.09
-0.53
-0.09
-0.16
0.46
0.77
0.41
0.01
0.01
-0.06
-0.09
-0.11
-0.13
0.14
0.12
0.05
-0.01
-0.09
-0.18
-0.29
-0.76
-0.13
0.18
0.09
0.07
-0.05
-0.17
-0.28
-0.36
-0.54
-0.67
0.29
0.04
0.19
0.17
0.16
0.15

| |
|---|
| 0.13 |
| 0.11 |
| 0.04 |
| 0.00 |
| -0.06 |
| -0.08 |
| -0.18 |
| -0.25 |
| -0.31 |
| -0.50 |
| -0.56 |
| -0.59 |
| 0.17 |
| -0.10 |
| 0.33 |
| 0.29 |
| 0.18 |
| 0.12 |
| -0.02 |
| -0.04 |
| -0.06 |
| -0.08 |
| -0.28 |
| -0.33 |
| -0.67 |
| 0.19 |
| 0.03 |
| -0.15 |
| -0.19 |
| -0.28 |
| -0.33 |
| -0.34 |
| 0.10 |
| 0.20 |
| 0.19 |
| 0.05 |
| -0.12 |
| -0.21 |
| 0.12 |
| -0.01 |
| -0.27 |
| -0.43 |
| -0.97 |
| -0.13 |
| -0.26 |
| -0.43 |
| -0.09 |
| -0.13 |

-0.41
-0.52
-0.14
-0.17
-0.31
-0.56
0.13
-0.18
-0.32
-0.18
-0.33
0.06
-0.41
-0.62
-0.18
-0.05
-0.39
-0.50
0.40
0.91
-0.49
0.07
0.13
0.02
-0.02
-0.14
-0.31
-0.40
0.05
0.20
0.34
0.31
-0.19
-0.31
0.22
0.04
-0.39
0.01
-0.09
-0.20
-0.22
-0.23
-0.34
-0.43
-0.45
-0.54
-0.54
-0.74

| |
|---|
| -0.07 |
| 0.02 |
| -0.08 |
| 0.02 |
| -0.66 |
| -0.44 |
| -0.05 |
| 0.28 |
| 0.04 |
| 0.12 |
| -0.10 |
| -0.25 |
| 0.87 |
| -0.11 |
| -0.14 |
| -0.35 |
| -0.35 |
| -0.50 |
| 0.58 |
| 0.30 |
| 0.23 |
| 0.18 |
| 0.05 |
| 0.03 |
| 0.02 |
| -0.18 |
| -0.30 |
| -0.36 |
| -0.54 |
| -0.54 |
| -0.06 |
| 0.13 |
| -0.29 |
| 0.24 |
| 0.04 |
| 0.22 |
| -0.06 |
| -0.16 |
| -0.31 |
| -0.40 |
| -0.21 |
| 0.18 |
| 0.16 |
| -0.13 |
| -0.02 |
| -0.03 |
| -0.16 |
| -0.39 |

| |
|---|
| 1.29 |
| 0.92 |
| 0.51 |
| 0.41 |
| 0.24 |
| 0.12 |
| 0.11 |
| 0.07 |
| 0.06 |
| 0.00 |
| -0.03 |
| -0.10 |
| -0.14 |
| -0.15 |
| -0.23 |
| -0.39 |
| -0.42 |
| -0.59 |
| -0.87 |
| -0.89 |
| 0.90 |
| -0.10 |
| -0.13 |
| -0.17 |
| -0.21 |
| 0.52 |
| 0.51 |
| 0.42 |
| 0.35 |
| 0.29 |
| 0.29 |
| 0.18 |
| 0.03 |
| 0.01 |
| -0.19 |
| -0.34 |
| 0.01 |
| -0.05 |
| -0.06 |
| -0.28 |
| -0.49 |
| 0.25 |
| -0.12 |
| -0.38 |
| 0.25 |
| 1.70 |
| 1.28 |
| 0.72 |

| |
|---|
| 0.22 |
| 0.12 |
| 0.11 |
| 0.08 |
| -0.04 |
| -0.14 |
| -0.50 |
| 0.57 |
| 0.33 |
| 0.27 |
| 0.22 |
| 0.16 |
| 0.15 |
| 0.13 |
| 0.06 |
| 0.05 |
| 0.04 |
| -0.01 |
| -0.08 |
| -0.09 |
| -0.11 |
| -0.11 |
| -0.14 |
| -0.15 |
| -0.33 |
| -0.35 |
| -0.45 |
| 0.89 |
| 0.60 |
| 0.52 |
| 0.13 |
| 0.09 |
| 0.02 |
| -0.01 |
| -0.02 |
| -0.16 |
| -0.25 |
| 0.62 |
| 0.33 |
| 0.31 |
| 0.23 |
| 0.04 |
| 0.00 |
| 0.00 |
| -0.02 |
| -0.05 |
| -0.34 |
| 0.93 |

| |
|---|
| 0.28 |
| 0.05 |
| 0.00 |
| 0.00 |
| -0.10 |
| -0.15 |
| -0.15 |
| -0.18 |
| -0.35 |
| -0.44 |
| -0.57 |
| -0.70 |
| 0.63 |
| 0.46 |
| 0.43 |
| 0.38 |
| 0.33 |
| 0.32 |
| 0.31 |
| 0.28 |
| 0.26 |
| 0.08 |
| 0.08 |
| 0.08 |
| 0.04 |
| 0.03 |
| 0.01 |
| 0.01 |
| 0.00 |
| -0.02 |
| -0.04 |
| -0.06 |
| -0.07 |
| -0.11 |
| -0.24 |
| -0.25 |
| -0.27 |
| -0.42 |
| 0.27 |
| 0.06 |
| -0.48 |
| 1.16 |
| 0.74 |
| 0.30 |
| 1.15 |
| 0.53 |
| 0.39 |
| 0.82 |

| |
|---|
| 0.03 |
| -0.18 |
| 0.17 |
| -0.29 |
| -0.41 |
| 0.07 |
| -0.15 |
| 0.03 |
| 0.00 |
| -0.14 |
| 0.73 |
| -0.09 |
| -0.17 |
| -0.02 |
| -0.08 |
| -0.46 |
| 0.02 |
| -0.09 |
| -0.10 |
| -0.43 |
| 0.17 |
| 0.00 |
| 0.00 |
| -0.19 |
| -0.38 |
| 0.18 |
| -0.26 |
| 0.09 |
| 0.08 |
| 0.01 |
| 0.60 |
| 0.06 |
| -0.06 |
| -0.10 |
| -0.12 |
| -0.14 |
| -0.19 |
| -0.21 |
| -0.44 |
| -0.57 |
| -1.13 |
| 1.46 |
| 0.30 |
| 0.30 |
| 0.27 |
| 0.14 |
| 0.08 |
| -0.02 |

-0.03
-0.06
-0.21
-0.12
0.16
0.13
0.13
0.05
-0.07
-0.45
-0.54
-0.62
0.27
-0.01
0.28
0.20
0.21
0.11
0.06
-0.18
0.48
0.45
0.11
-0.02
-0.07
0.23
0.14
0.00
-0.06
0.05
-0.07
-0.09
-0.13
-0.20
-0.18
0.56
-0.28
-0.17
0.00
-0.26
0.79
0.47
0.37
0.30
0.23
0.23
0.18
0.14

| |
|---|
| 0.14 |
| 0.12 |
| 0.07 |
| 0.04 |
| 0.00 |
| -0.06 |
| -0.13 |
| -0.13 |
| -0.14 |
| -0.21 |
| -0.22 |
| -0.23 |
| -0.28 |
| -0.44 |
| -0.60 |
| -0.62 |
| -0.67 |
| -0.69 |
| -0.70 |
| -0.71 |
| -0.78 |
| 0.22 |
| -0.10 |
| 0.08 |
| 0.08 |
| 0.04 |
| -0.02 |
| -0.14 |
| -0.18 |
| -0.41 |
| 0.32 |
| 0.16 |
| 0.03 |
| 0.40 |
| 0.34 |
| 0.21 |
| 0.13 |
| -0.13 |
| -0.18 |
| -0.21 |
| -0.42 |
| -0.47 |
| -0.62 |
| -0.30 |
| -0.32 |
| 0.16 |
| -0.37 |
| -0.37 |

| |
|---|
| 0.48 |
| 0.15 |
| -0.19 |
| -0.45 |
| 0.14 |
| -0.10 |
| -0.15 |
| -0.23 |
| -0.25 |
| -0.32 |
| -0.51 |
| -0.67 |
| 0.42 |
| 0.31 |
| 0.31 |
| 0.29 |
| 0.28 |
| 0.22 |
| 0.16 |
| 0.13 |
| 0.10 |
| 0.06 |
| 0.04 |
| -0.03 |
| -0.04 |
| -0.05 |
| -0.05 |
| -0.08 |
| -0.08 |
| -0.08 |
| -0.09 |
| -0.09 |
| -0.12 |
| -0.12 |
| -0.13 |
| -0.13 |
| -0.14 |
| -0.15 |
| -0.15 |
| -0.16 |
| -0.17 |
| -0.19 |
| -0.21 |
| -0.24 |
| -0.28 |
| -0.29 |
| -0.33 |
| -0.34 |

-0.34
-0.35
-0.39
-0.42
-0.45
-0.46
-0.47
-0.58
-0.58
-0.21
0.12
0.04
-0.18
0.45
0.27
0.57
0.45
0.05
0.01
-0.01
-0.48
0.02
-0.46
-0.30
0.20
0.20
0.19
0.72
0.09
0.03
0.02
-0.01
0.46
0.34
0.25
0.18
0.18
0.11
0.11
0.10
0.08
0.07
0.04
0.03
0.03
-0.02
-0.25
-0.27

| |
|---|
| -0.30 |
| -0.43 |
| 0.32 |
| 0.02 |
| -0.09 |
| 0.12 |
| 0.12 |
| -0.02 |
| -0.03 |
| 0.77 |
| 0.70 |
| 0.46 |
| 0.33 |
| 0.32 |
| 0.31 |
| 0.31 |
| 0.30 |
| 0.30 |
| 0.24 |
| 0.11 |
| 0.08 |
| -0.03 |
| -0.11 |
| 0.72 |
| 0.39 |
| 1.11 |
| 0.28 |
| 0.17 |
| 0.17 |
| 0.12 |
| 0.09 |
| 0.08 |
| 0.08 |
| 0.03 |
| 0.02 |
| 0.01 |
| 0.01 |
| 0.01 |
| -0.04 |
| -0.09 |
| -0.32 |
| -0.34 |
| 0.71 |
| 0.46 |
| 0.36 |
| 0.30 |
| 0.24 |
| 0.22 |

| |
|---|
| 0.17 |
| 0.14 |
| 0.08 |
| 0.07 |
| 0.06 |
| -0.01 |
| -0.04 |
| -0.29 |
| 0.05 |
| 0.01 |
| 0.03 |
| -0.17 |
| 0.18 |
| 0.15 |
| -0.01 |
| -0.10 |
| -0.30 |
| -0.42 |
| 0.06 |
| 0.53 |
| 0.39 |
| -0.03 |
| -0.11 |
| -0.15 |
| -0.45 |
| -0.13 |
| -0.34 |
| 0.00 |
| -0.01 |
| -0.14 |
| -0.16 |
| -0.20 |
| -0.22 |
| -0.30 |
| -0.31 |
| -0.47 |
| -0.68 |
| 0.73 |
| 0.27 |
| 0.27 |
| 0.21 |
| 0.20 |
| 0.20 |
| 0.18 |
| 0.16 |
| 0.14 |
| 0.13 |
| 0.13 |

0.11
0.10
0.08
0.08
0.07
0.05
0.04
0.03
0.03
0.03
0.03
0.02
0.02
0.02
0.02
0.01
0.01
0.00
0.00
0.00
0.00
-0.01
-0.01
-0.02
-0.02
-0.03
-0.03
-0.04
-0.05
-0.05
-0.05
-0.06
-0.06
-0.06
-0.07
-0.07
-0.08
-0.09
-0.09
-0.09
-0.09
-0.09
-0.10
-0.10
-0.10
-0.11
-0.11
-0.11

-0.12
-0.12
-0.12
-0.12
-0.12
-0.12
-0.12
-0.14
-0.14
-0.14
-0.15
-0.15
-0.15
-0.16
-0.16
-0.17
-0.17
-0.18
-0.18
-0.19
-0.19
-0.19
-0.20
-0.20
-0.21
-0.21
-0.21
-0.22
-0.22
-0.23
-0.23
-0.23
-0.24
-0.25
-0.25
-0.29
-0.30
-0.32
-0.33
-0.33
-0.33
-0.35
-0.36
-0.37
-0.40
-0.41
-0.41
-0.41

| |
|---|
| -0.43 |
| -0.44 |
| -0.47 |
| -0.48 |
| -0.49 |
| -0.49 |
| -0.54 |
| -0.54 |
| -0.54 |
| -0.56 |
| -0.58 |
| -0.59 |
| -0.75 |
| -0.83 |
| -0.84 |
| -0.94 |
| 1.16 |
| 0.78 |
| 0.62 |
| 0.62 |
| 0.58 |
| 0.55 |
| 0.54 |
| 0.36 |
| 0.36 |
| 0.36 |
| 0.33 |
| 0.33 |
| 0.31 |
| 0.27 |
| 0.27 |
| 0.26 |
| 0.26 |
| 0.22 |
| 0.18 |
| 0.18 |
| 0.14 |
| 0.14 |
| 0.14 |
| 0.13 |
| 0.13 |
| 0.12 |
| 0.11 |
| 0.10 |
| 0.10 |
| 0.09 |
| 0.09 |
| 0.08 |

0.08
0.08
0.08
0.07
0.06
0.05
0.05
0.05
0.05
0.04
0.03
0.03
0.03
0.03
0.03
0.02
0.01
0.00
0.00
0.00
0.00
0.00
0.00
0.00
-0.01
-0.01
-0.01
-0.01
-0.02
-0.02
-0.03
-0.03
-0.03
-0.04
-0.04
-0.05
-0.05
-0.05
-0.05
-0.05
-0.05
-0.05
-0.06
-0.06
-0.06
-0.07
-0.07
-0.07

-0.07
-0.08
-0.08
-0.09
-0.09
-0.09
-0.09
-0.09
-0.10
-0.10
-0.10
-0.11
-0.11
-0.11
-0.11
-0.12
-0.12
-0.12
-0.12
-0.12
-0.12
-0.13
-0.13
-0.14
-0.14
-0.14
-0.14
-0.15
-0.15
-0.15
-0.15
-0.15
-0.16
-0.16
-0.16
-0.17
-0.17
-0.18
-0.19
-0.19
-0.19
-0.20
-0.20
-0.20
-0.20
-0.20
-0.21
-0.21

-0.21
-0.21
-0.21
-0.21
-0.21
-0.21
-0.22
-0.23
-0.23
-0.24
-0.24
-0.24
-0.24
-0.24
-0.24
-0.24
-0.25
-0.26
-0.26
-0.26
-0.26
-0.26
-0.26
-0.27
-0.27
-0.27
-0.27
-0.27
-0.28
-0.28
-0.28
-0.29
-0.29
-0.29
-0.29
-0.29
-0.29
-0.29
-0.30
-0.30
-0.31
-0.31
-0.31
-0.31
-0.33
-0.34
-0.35
-0.35

-0.35
-0.35
-0.36
-0.36
-0.36
-0.36
-0.36
-0.36
-0.36
-0.37
-0.37
-0.38
-0.38
-0.39
-0.39
-0.40
-0.40
-0.41
-0.41
-0.41
-0.41
-0.42
-0.42
-0.43
-0.43
-0.43
-0.46
-0.46
-0.48
-0.48
-0.48
-0.48
-0.48
-0.49
-0.50
-0.52
-0.53
-0.54
-0.54
-0.54
-0.57
-0.58
-0.58
-0.60
-0.60
-0.60
-0.62
-0.62

| |
|---|
| -0.62 |
| -0.66 |
| -0.67 |
| -0.67 |
| -0.69 |
| -0.91 |
| -1.02 |
| -1.28 |
| -1.40 |
| -0.22 |
| -0.29 |
| -0.45 |
| -0.56 |
| -0.69 |
| -0.72 |
| -1.06 |
| -0.15 |
| -0.81 |
| 0.15 |
| 0.00 |
| -0.11 |
| -0.19 |
| -0.36 |
| -0.59 |
| -0.60 |
| -0.90 |
| -0.18 |
| -0.20 |
| -0.25 |
| -0.25 |
| -0.27 |
| -0.54 |
| -0.60 |
| 0.01 |
| -0.11 |
| -0.17 |
| -0.32 |
| 0.28 |
| 0.12 |
| -0.29 |
| -0.05 |
| -0.08 |
| -0.31 |
| -0.35 |
| -0.61 |
| 0.61 |
| 0.33 |
| -0.12 |

-0.20
-0.41
0.32
0.16
-0.01
-0.03
-0.19
-0.24
-0.30
-0.55
-0.28
0.03
-0.08
-0.35
0.03
-0.01
-0.07
-0.14
-0.16
-0.46
0.70
0.69
0.36
-0.18
-0.18
0.36
0.08
-0.10
-0.12
-0.23
-0.41
-0.65
-0.65
0.48
0.42
0.40
0.40
0.07
-0.50
1.06
1.00
0.38
0.08
0.02
-0.02
-0.10
-0.12
-0.13

| |
|---|
| -0.21 |
| -0.43 |
| -0.48 |
| -0.74 |
| 0.32 |
| 0.07 |
| -0.17 |
| 0.19 |
| 0.36 |
| -0.12 |
| -0.42 |
| -0.53 |
| -0.56 |
| -0.59 |
| 0.39 |
| 0.36 |
| 0.34 |
| 0.04 |
| 0.12 |
| 0.09 |
| -0.09 |
| -0.72 |
| 0.19 |
| 0.13 |
| 0.15 |
| -0.17 |
| 0.75 |
| 0.55 |
| 0.54 |
| 0.06 |
| 0.02 |
| -0.07 |
| -0.11 |
| -0.23 |
| -0.27 |
| -0.33 |
| 0.14 |
| 0.12 |
| -0.06 |
| -0.06 |
| -0.09 |
| -0.13 |
| -0.20 |
| -0.20 |
| 0.10 |
| -0.24 |
| -0.45 |
| -0.25 |

-0.18
-0.40
0.30
0.02
-0.16
0.15
0.12
0.01
-0.46
0.09
-0.18
-0.89
-0.25
0.17
0.06
-0.29
0.08
-0.05
-0.33
-0.35
-0.68
0.12
-0.06
-0.19
-0.44
-0.44
0.16
0.06
-0.02
-0.05
-0.52
-0.41
0.06
-0.03
-0.16
0.07
-0.08
-0.09
0.29
0.22
0.06
0.00
-0.11
-0.45
-0.15
-0.23
-0.37
-0.37

-0.56
-0.58
-0.75
-0.16
0.24
0.05
-0.21
-0.40
-0.20
0.08
-0.04
-0.15
-0.22
0.17
-0.01
-0.09
-0.15
-0.35
1.23
-0.06
-0.11
0.41
0.04
-0.05
-0.07
-0.09
-0.27
-0.27
-0.30
-0.44
0.50
0.18
0.11
0.04
-0.13
0.13
-0.01
-0.06
0.42
0.37
-0.25
0.15
0.12
0.03
-0.02
-0.05
-0.16
-0.16

-0.39
-0.46
0.11
-0.21
0.57
0.53
0.53
0.47
0.31
0.12
0.46
-0.05
-0.20
0.41
0.33
0.28
0.25
0.23
0.23
0.16
-0.06
0.21
0.15
0.12
0.03
0.03
-0.04
-0.08
-0.20
-0.21
-0.29
-0.32
-0.44
0.12
-0.27
0.14
0.45
0.21
0.18
-0.02
-0.04
-0.07
-0.17
-0.22
0.35
0.04
0.06
-0.01

| |
|---|
| 0.36 |
| 0.15 |
| 0.06 |
| 0.05 |
| 0.01 |
| 0.00 |
| 0.00 |
| -0.05 |
| -0.06 |
| -0.23 |
| -0.27 |
| -0.28 |
| -0.29 |
| -0.34 |
| -0.45 |
| -0.52 |
| -0.53 |
| -0.55 |
| 0.10 |
| -0.05 |
| -0.44 |
| 0.32 |
| 0.26 |
| 0.22 |
| 0.16 |
| 0.10 |
| -0.19 |
| 0.21 |
| 0.16 |
| 0.07 |
| 0.12 |
| 0.11 |
| -0.07 |
| -0.14 |
| -0.23 |
| -0.27 |
| -0.31 |
| 0.17 |
| 0.18 |
| 0.11 |
| -0.06 |
| -0.07 |
| -0.26 |
| -0.57 |
| 0.47 |
| -0.01 |
| -0.03 |
| -0.35 |

| |
|---|
| -0.42 |
| -0.48 |
| -0.33 |
| 0.06 |
| 0.14 |
| 0.03 |
| -0.04 |
| -0.08 |
| -0.77 |
| 0.22 |
| 0.14 |
| -0.53 |
| -0.67 |
| -0.12 |
| -0.24 |
| 0.40 |
| 0.28 |
| 0.70 |
| 0.33 |
| 0.31 |
| 0.25 |
| 0.11 |
| 0.03 |
| -0.02 |
| -0.12 |
| -0.21 |
| -0.29 |
| -0.35 |
| -0.08 |
| 0.08 |
| 0.04 |
| 0.03 |
| 0.00 |
| -0.03 |
| -0.05 |
| -0.08 |
| -0.17 |
| 0.41 |
| 0.21 |
| 0.04 |
| -0.01 |
| -0.19 |
| -0.48 |
| -0.01 |
| -0.15 |
| 0.29 |
| 0.34 |
| -0.02 |

0.10
0.07
-0.26
-0.27
-0.33
0.00
0.70
0.53
0.32
0.25
0.22
0.20
0.12
0.11
0.07
0.04
0.00
-0.09
-0.09
-0.13
-0.14
-0.19
-0.25
-0.26
-0.28
-0.29
-0.38
-0.44
-0.58
0.51
0.41
0.35
0.27
0.19
0.16
0.13
0.08
0.04
0.03
0.01
-0.06
-0.21
-0.33
-0.52
-0.85
0.17
0.16
0.07

| |
|---|
| 0.06 |
| -0.04 |
| -0.06 |
| -0.25 |
| -0.06 |
| -0.17 |
| -0.25 |
| -0.48 |
| -0.65 |
| 0.61 |
| 0.26 |
| 0.03 |
| 0.03 |
| 0.02 |
| -0.09 |
| -0.17 |
| -0.32 |
| -0.25 |
| -0.50 |
| 0.12 |
| 0.01 |
| -0.15 |
| 0.04 |
| -0.08 |
| -0.14 |
| -0.23 |
| 0.27 |
| -0.20 |
| -0.23 |
| -0.29 |
| -0.39 |
| -0.66 |
| -1.06 |
| 0.13 |
| 0.11 |
| 0.10 |
| -0.15 |
| -0.35 |
| 0.18 |
| 0.10 |
| -0.02 |
| 0.38 |
| -0.22 |
| -0.49 |
| -0.10 |
| 0.58 |
| 0.16 |
| 0.22 |

0.17
0.16
0.08
0.04
0.02
-0.01
-0.05
-0.09
-0.15
-0.26
-0.29
-0.45
0.35
0.00
0.47
-0.01
1.14
1.06
0.45
0.33
0.22
0.02
-0.15
-0.15
-0.18
-0.23
-0.27
-0.33
-0.35
-0.37
-0.38
-0.43
-0.46
-0.52
-0.60
-0.61
-0.71
-0.75
-0.88
0.21
-0.33
-0.43
-0.57
0.13
0.83
0.65
-0.01
-0.31

| |
|---|
| 1.29 |
| 0.01 |
| -0.07 |
| -0.31 |
| 0.23 |
| 1.49 |
| 1.33 |
| -0.28 |
| 0.29 |
| -0.12 |
| 0.26 |
| 0.33 |
| 0.28 |
| 0.10 |
| -0.11 |
| -0.34 |
| 0.03 |
| -0.23 |
| -0.24 |
| -0.04 |
| -0.16 |
| 0.23 |
| 0.00 |
| -0.10 |
| -0.16 |
| 0.15 |
| 0.07 |
| 0.07 |
| -0.64 |
| -0.66 |
| 0.07 |
| 0.38 |
| 0.36 |
| 0.28 |
| 0.22 |
| 0.08 |
| -0.06 |
| 0.25 |
| 0.21 |
| 0.12 |
| 0.09 |
| 0.08 |
| 0.03 |
| 0.00 |
| -0.07 |
| -0.21 |
| -0.28 |
| -0.51 |

| |
|---|
| 0.29 |
| 0.20 |
| 0.14 |
| 0.11 |
| 0.11 |
| 0.02 |
| -0.10 |
| -0.18 |
| 0.04 |
| -0.07 |
| -0.07 |
| -0.68 |
| 0.67 |
| 0.65 |
| -0.71 |
| -0.87 |
| -0.92 |
| 0.42 |
| -0.81 |
| 0.19 |
| 0.17 |
| -0.07 |
| -0.16 |
| 0.32 |
| 0.19 |
| -0.04 |
| -0.14 |
| -0.15 |
| -0.16 |
| -0.22 |
| -0.23 |
| -0.30 |
| -0.88 |
| 0.17 |
| 0.88 |
| 0.48 |
| -1.00 |
| -1.28 |
| 0.33 |
| 1.21 |
| 0.56 |
| 0.23 |
| 0.23 |
| 0.19 |
| 0.17 |
| 0.17 |
| 0.13 |
| 0.12 |

| |
|---|
| 0.12 |
| 0.12 |
| 0.10 |
| 0.07 |
| 0.05 |
| 0.04 |
| 0.03 |
| 0.01 |
| 0.00 |
| -0.01 |
| -0.04 |
| -0.04 |
| -0.05 |
| -0.10 |
| -0.11 |
| -0.11 |
| -0.13 |
| -0.13 |
| -0.15 |
| -0.19 |
| -0.20 |
| -0.20 |
| -0.24 |
| -0.27 |
| -0.29 |
| -0.30 |
| -0.32 |
| -0.34 |
| -0.34 |
| -0.41 |
| -0.44 |
| -0.46 |
| -0.54 |
| -0.60 |
| -0.68 |
| -0.68 |
| -0.71 |
| -0.33 |
| -0.50 |
| 0.40 |
| -0.48 |
| 0.33 |
| 0.14 |
| 0.05 |
| -0.02 |
| -0.09 |
| -0.11 |
| -0.14 |

-0.14
-0.16
-0.17
-0.22
-0.22
-0.25
-0.26
-0.36
-0.37
0.43
0.29
0.08
-0.05
0.20
0.19
-0.13
0.42
0.22
0.19
0.16
0.16
0.15
0.13
0.08
0.04
0.01
-0.01
-0.04
-0.05
-0.05
-0.09
-0.10
-0.13
-0.14
-0.19
-0.27
-0.29
-0.30
-0.30
-0.37
-0.72
0.33
0.32
0.29
0.24
0.23
0.11
0.07

| |
|---|
| 0.03 |
| -0.02 |
| -0.11 |
| -0.23 |
| -0.26 |
| -0.41 |
| 0.59 |
| 0.29 |
| 0.23 |
| 0.22 |
| 0.14 |
| 0.13 |
| 0.11 |
| 0.09 |
| 0.06 |
| 0.05 |
| 0.00 |
| -0.11 |
| -0.15 |
| -0.63 |
| 0.49 |
| 0.08 |
| -0.13 |
| -0.06 |
| 0.10 |
| -0.52 |
| 0.17 |
| -0.28 |
| 0.19 |
| -0.08 |
| -0.14 |
| 0.17 |
| -0.55 |
| -1.01 |
| 0.13 |
| -0.04 |
| -0.22 |
| 0.01 |
| 0.09 |
| -0.26 |
| -0.15 |
| -0.13 |
| 0.01 |
| 0.04 |
| -0.06 |
| -0.13 |
| -0.34 |
| -0.35 |

-0.25
-0.71
-0.75
-0.79
-0.36
-0.48
0.21
0.04
-0.50
0.05
-0.14
-0.19
-0.01
-0.08
0.22
0.11
-0.01
-0.15
0.34
0.03
-0.14
-0.28
-0.45
-0.05
-0.33
-0.39
0.23
-0.16
0.38
-0.10
-0.19
-0.11
-0.37
-0.60
0.11
-0.03
-0.12
0.73
0.43
0.29
0.11
0.11
0.07
0.03
0.02
0.00
-0.03
-0.03

-0.08
-0.14
-0.14
-0.17
-0.21
-0.22
-0.24
-0.26
-0.32
-0.33
-0.40
-0.41
-0.43
-0.51
-0.53
-0.56
-0.59
-0.80
-0.91
-0.94
-0.09
-0.29
-0.48
0.05
0.05
0.03
-0.17
-0.22
-0.23
-0.37
-0.39
0.16
0.15
0.11
0.05
0.05
-0.05
-0.20
-0.23
-0.26
-0.43
0.21
0.28
0.23
0.17
0.16
0.14
0.06

| |
|---|
| 0.04 |
| 0.03 |
| 0.03 |
| 0.03 |
| 0.00 |
| -0.01 |
| -0.07 |
| -0.07 |
| -0.09 |
| -0.13 |
| -0.16 |
| -0.16 |
| -0.17 |
| -0.17 |
| -0.21 |
| -0.28 |
| -0.37 |
| -0.39 |
| 0.28 |
| 0.16 |
| 0.15 |
| 0.11 |
| 0.08 |
| 0.02 |
| 0.02 |
| 0.01 |
| -0.03 |
| -0.09 |
| -0.14 |
| -0.18 |
| -0.18 |
| -0.21 |
| -0.39 |
| -0.46 |
| -0.61 |
| 0.11 |
| 0.08 |
| 0.02 |
| 0.29 |
| -0.14 |
| 0.02 |
| -0.12 |
| 0.71 |
| 0.69 |
| 0.29 |
| 0.23 |
| 0.10 |
| 0.25 |

| |
|---|
| 0.24 |
| 0.34 |
| -0.01 |
| -0.08 |
| -0.70 |
| 0.57 |
| 0.24 |
| 0.25 |
| 0.22 |
| 0.13 |
| -0.42 |
| 0.15 |
| -0.20 |
| -0.32 |
| -0.37 |
| 0.64 |
| 0.29 |
| 0.25 |
| 0.18 |
| 0.13 |
| 0.04 |
| -0.01 |
| -0.01 |
| -0.09 |
| -0.18 |
| -0.40 |
| 0.99 |
| 0.75 |
| 0.34 |
| 0.33 |
| 0.26 |
| 0.17 |
| 0.12 |
| 0.12 |
| 0.05 |
| 0.04 |
| 0.03 |
| 0.00 |
| -0.03 |
| -0.06 |
| -0.07 |
| -0.09 |
| -0.09 |
| -0.18 |
| -0.19 |
| -0.20 |
| -0.24 |
| -0.30 |

-0.39
-0.43
0.29
0.21
0.20
0.19
0.18
0.17
0.15
0.15
0.14
0.12
0.10
0.07
0.03
0.02
-0.01
-0.02
-0.03
-0.03
-0.07
-0.07
-0.11
-0.11
-0.11
-0.12
-0.12
-0.15
-0.20
-0.22
-0.22
-0.23
-0.24
-0.26
-0.27
-0.30
-0.32
-0.41
-0.73
-0.79
-0.98
0.28
0.13
0.06
-0.05
-0.10
-0.37
-0.39

| |
|---|
| -0.45 |
| -0.46 |
| 0.27 |
| 0.12 |
| 0.05 |
| 0.03 |
| 0.02 |
| -0.08 |
| -0.18 |
| -0.19 |
| -0.21 |
| -0.24 |
| -0.27 |
| -0.31 |
| -0.31 |
| -0.32 |
| -0.34 |
| -0.60 |
| -1.17 |
| -0.03 |
| -0.10 |
| -0.10 |
| 1.88 |
| 1.46 |
| 1.09 |
| 1.08 |
| 1.04 |
| 0.99 |
| 0.99 |
| 0.90 |
| 0.90 |
| 0.72 |
| 0.71 |
| 0.66 |
| 0.62 |
| 0.60 |
| 0.58 |
| 0.55 |
| 0.55 |
| 0.54 |
| 0.53 |
| 0.52 |
| 0.51 |
| 0.50 |
| 0.50 |
| 0.50 |
| 0.49 |
| 0.49 |

| |
|---|
| 0.48 |
| 0.45 |
| 0.43 |
| 0.40 |
| 0.36 |
| 0.36 |
| 0.35 |
| 0.31 |
| 0.30 |
| 0.29 |
| 0.28 |
| 0.26 |
| 0.26 |
| 0.23 |
| 0.22 |
| 0.22 |
| 0.21 |
| 0.20 |
| 0.19 |
| 0.17 |
| 0.17 |
| 0.17 |
| 0.16 |
| 0.16 |
| 0.15 |
| 0.14 |
| 0.14 |
| 0.13 |
| 0.13 |
| 0.12 |
| 0.11 |
| 0.11 |
| 0.11 |
| 0.07 |
| 0.07 |
| 0.06 |
| 0.04 |
| 0.04 |
| 0.03 |
| 0.03 |
| 0.03 |
| 0.02 |
| 0.01 |
| 0.01 |
| 0.00 |
| 0.00 |
| 0.00 |
| -0.01 |

-0.02
-0.03
-0.03
-0.03
-0.06
-0.06
-0.07
-0.07
-0.08
-0.08
-0.09
-0.09
-0.13
-0.14
-0.18
-0.18
-0.19
-0.29
-0.29
-0.35
-0.38
-0.40
-0.73
-0.94
-0.11
-0.23
-0.31
-0.30
-0.43
0.47
0.36
0.24
0.10
0.07
0.07
-0.03
0.69
0.61
0.39
0.27
0.22
0.17
0.06
0.05
-0.15
-0.54
-0.57
-0.06

| |
|---|
| -0.32 |
| 0.12 |
| -0.31 |
| -0.35 |
| -0.40 |
| -0.57 |
| -0.74 |
| -0.75 |
| 0.09 |
| 0.03 |
| 0.02 |
| -0.13 |
| -0.14 |
| -0.15 |
| -0.16 |
| -0.16 |
| -0.17 |
| -0.29 |
| -0.32 |
| -0.43 |
| -0.51 |
| -0.57 |
| -0.59 |
| -0.59 |
| -0.77 |
| 0.23 |
| -0.24 |
| 0.94 |
| 0.18 |
| 0.18 |
| -0.02 |
| 0.02 |
| -0.05 |
| -0.21 |
| -0.25 |
| -0.65 |
| -0.70 |
| -1.03 |
| -1.31 |
| 0.54 |
| 0.17 |
| -0.06 |
| 0.34 |
| 0.35 |
| 0.26 |
| 0.07 |
| -0.16 |
| -0.40 |

| |
|---|
| -0.47 |
| -0.60 |
| -0.71 |
| 0.00 |
| -0.06 |
| 0.84 |
| 0.49 |
| 0.26 |
| 0.16 |
| 0.03 |
| -0.11 |
| -0.13 |
| -0.39 |
| 0.06 |
| 0.00 |
| -0.07 |
| -0.10 |
| -0.12 |
| -0.24 |
| -0.28 |
| -0.03 |
| -0.01 |
| 0.16 |
| 0.14 |
| 0.10 |
| 0.02 |
| 0.01 |
| -0.04 |
| -0.07 |
| -0.11 |
| -0.13 |
| -0.13 |
| -0.15 |
| -0.19 |
| -0.21 |
| -0.24 |
| -0.25 |
| -0.27 |
| -0.32 |
| -0.46 |
| -0.65 |
| 0.04 |
| -0.23 |
| 0.56 |
| 0.27 |
| -0.04 |
| 0.12 |
| -0.20 |

| |
|---|
| -0.27 |
| 1.09 |
| 0.95 |
| 0.08 |
| 0.37 |
| 0.19 |
| 0.08 |
| 0.75 |
| 0.18 |
| 0.15 |
| 0.03 |
| -0.18 |
| -0.35 |
| -0.38 |
| 2.01 |
| 0.75 |
| 0.50 |
| 0.37 |
| 0.34 |
| 0.21 |
| 0.18 |
| 0.15 |
| 0.12 |
| 0.11 |
| 0.00 |
| -0.02 |
| -0.03 |
| -0.05 |
| -0.06 |
| -0.07 |
| -0.11 |
| -0.12 |
| -0.12 |
| -0.12 |
| -0.13 |
| -0.15 |
| -0.18 |
| -0.25 |
| -0.38 |
| -0.47 |
| -0.74 |
| -0.02 |
| -0.13 |
| 0.31 |
| 0.03 |
| -0.20 |
| 0.15 |
| 0.10 |

| |
|---|
| 0.13 |
| 0.49 |
| 0.05 |
| 0.10 |
| -0.14 |
| -0.28 |
| -0.13 |
| -0.23 |
| 0.09 |
| -0.06 |
| 0.57 |
| 0.51 |
| -0.41 |
| -1.30 |
| -0.09 |
| -0.41 |
| -0.07 |
| -0.22 |
| -0.03 |
| -0.08 |
| 0.33 |
| 0.19 |
| -0.05 |
| -0.49 |
| 0.59 |
| -0.08 |
| 0.02 |
| -0.09 |
| 0.86 |
| 0.31 |
| 0.15 |
| -0.02 |
| 0.33 |
| 0.04 |
| -0.12 |
| 0.06 |
| -0.02 |
| -0.07 |
| 0.42 |
| 0.27 |
| -0.28 |
| -0.30 |
| -0.39 |
| 0.14 |
| 1.22 |
| -0.12 |
| 0.07 |
| -0.01 |

-0.13
-0.31
-0.40
-0.44
0.04
0.20
-0.05
0.37
0.08
0.05
-0.13
-0.21
-0.27
-0.35
-0.36
-0.42
-0.42
-0.42
-0.51
-0.53
-0.15
-0.23
-0.26
-0.05
-0.09
-0.17
-0.28
-0.49
-0.55
-0.67
-1.28
0.30
0.15
0.08
0.00
-0.01
-0.01
-0.57
0.94
0.01
-0.04
-0.20
-0.15
0.28
0.20
0.07
-0.04
-0.34

| |
|---|
| 1.28 |
| 0.16 |
| 0.15 |
| 0.26 |
| 0.22 |
| -0.41 |
| 0.25 |
| 0.34 |
| 1.56 |
| -0.11 |
| 0.14 |
| -0.26 |
| -0.19 |
| -0.21 |
| -0.50 |
| 0.02 |
| 0.08 |
| 0.05 |
| 0.00 |
| -0.09 |
| -0.29 |
| -0.30 |
| -0.32 |
| -0.38 |
| -0.39 |
| 0.11 |
| 0.14 |
| -0.05 |
| -0.08 |
| 0.07 |
| -0.30 |
| 0.64 |
| 0.04 |
| -0.81 |
| 0.17 |
| 0.13 |
| -0.06 |
| -0.12 |
| 0.10 |
| -0.01 |
| -0.27 |
| 0.07 |
| -0.04 |
| 0.20 |
| 0.42 |
| 0.40 |
| -0.32 |
| -0.03 |

-0.04
0.33
0.25
-0.18
0.20
0.08
-0.26
-0.27
0.26
0.42
0.40
-0.35
-0.35
0.11
-0.09
-0.09
-0.47
-0.57
0.15
-0.02
0.42
0.27
0.27
0.41
0.38
0.01
0.29
0.29
0.16
0.11
0.08
0.06
0.06
0.04
0.03
-0.02
-0.13
-0.27
-0.55
0.67
0.24
0.18
0.12
0.07
0.03
-0.02
-0.07
-0.07

-0.09
-0.16
-0.17
-0.24
-0.27
-0.41
-0.83
-1.20
0.04
0.42
0.18
0.14
0.02
0.37
0.02
-0.31
-0.43
0.19
0.16
0.10
0.52
0.20
0.13
0.07
0.07
0.03
-0.06
-0.18
0.31
0.19
0.18
0.00
-0.57
-0.93
0.53
-0.28
-0.44
0.02
0.01
0.28
0.05
0.05
0.04
-0.05
-0.39
0.15
-0.13
-0.51

0.04
-0.14
-0.05
-0.12
-0.12
0.42
0.10
-0.04
-0.17
-0.20
0.46
0.10
-0.15
0.63
0.14
-0.09
-0.12
-0.24
-0.41
-0.41
0.14
0.11
-0.02
-0.38
-0.39
-0.43
-0.01
0.44
0.21
-0.13
-0.03
-0.11
-0.17
-0.38
-0.45
-0.45
-0.49
-0.50
-0.79
0.44
0.18
0.15
0.14
-0.02
-0.27
-0.72
0.41
0.01

-0.12
0.39
0.30
0.09
-0.02
0.24
0.03
-0.21
0.20
-0.12
-0.17
-0.36
-0.45
-0.57
-0.01
0.01
0.25
0.05
-0.04
-0.04
-0.05
0.03
-0.05
-0.22
-0.20
0.35
0.11
0.07
0.06
0.05
0.04
0.01
-0.04
-0.04
-0.10
-0.46
0.55
-0.03
0.28
-0.19
-0.32
0.20
0.36
0.29
0.24
-0.10
-0.21
-0.37

| |
|---|
| 0.06 |
| -0.02 |
| 0.05 |
| 0.28 |
| -0.52 |
| -0.28 |
| 0.05 |
| -0.17 |
| 0.59 |
| 0.09 |
| -0.61 |
| 0.13 |
| 0.15 |
| 0.06 |
| 0.05 |
| 0.02 |
| -0.14 |
| -0.21 |
| -0.22 |
| -0.54 |
| -0.59 |
| 0.06 |
| 0.08 |
| -0.01 |
| 0.48 |
| 0.14 |
| -0.05 |
| -0.21 |
| 0.26 |
| 0.16 |
| 0.63 |
| 0.39 |
| 0.05 |
| -0.08 |
| -0.13 |
| -0.27 |
| -0.56 |
| -0.26 |
| -0.51 |
| 0.36 |
| 0.23 |
| 0.04 |
| -0.04 |
| -0.16 |
| -0.17 |
| -0.21 |
| -0.23 |
| -0.56 |

-0.82
0.12
-0.01
-0.01
-0.12
0.69
0.55
0.27
0.03
-0.01
-0.40
-0.66
0.47
0.06
-0.05
0.13
-0.13
-0.55
1.76
0.42
0.02
0.33
0.33
0.32
0.21
0.14
0.14
0.12
0.11
0.09
-0.01
-0.04
-0.04
-0.15
-0.24
-0.26
-0.51
-0.90
-0.02
0.28
0.27
0.27
0.26
0.23
0.23
0.19
0.17
0.14

| |
|---|
| 0.11 |
| 0.11 |
| 0.04 |
| 0.04 |
| 0.03 |
| 0.02 |
| 0.00 |
| 0.00 |
| -0.02 |
| -0.05 |
| -0.05 |
| -0.09 |
| -0.11 |
| -0.16 |
| -0.21 |
| -0.23 |
| -0.53 |
| -0.16 |
| 1.78 |
| 2.90 |
| 2.84 |
| 1.02 |
| 0.58 |
| 0.20 |
| 0.15 |
| -0.13 |
| 0.33 |
| -0.24 |
| -0.47 |
| -0.52 |
| -0.67 |
| 0.16 |
| -0.27 |
| 0.06 |
| -0.05 |
| -0.07 |
| -0.27 |
| -0.35 |
| -0.41 |
| -0.57 |
| -0.57 |
| -0.57 |
| -0.59 |
| -0.79 |
| -0.83 |
| -1.31 |
| 0.16 |
| -0.12 |

-0.12
-0.13
-0.20
-0.12
0.53
0.11
0.03
-0.33
-0.68
0.26
-0.05
-0.09
-0.13
0.22
0.08
0.17
0.12
0.01
-0.11
-0.11
-0.42
0.06
-0.06
-0.18
-0.25
-0.26
-0.35
0.07
0.01
-0.09
-0.03
-0.03
-0.23
-0.50
-0.57
0.11
0.09
0.02
0.14
-0.08
-0.12
-0.35
-0.48
0.16
0.08
0.08
-0.10
-0.24

-0.34
0.32
0.15
0.15
-0.20
-0.20
0.31
-0.20
0.48
0.24
0.21
0.19
-0.05
-0.11
-0.18
-0.18
1.03
0.64
0.50
0.21
0.13
-0.03
-0.15
-0.31
-0.43
-0.10
-0.23
-0.19
-0.21
0.18
0.29
0.05
0.14
0.10
0.02
-0.01
-0.06
-0.09
0.43
0.41
0.41
0.00
-0.09
-0.11
0.75
0.20
0.00
0.61

| |
|---|
| 0.41 |
| 0.31 |
| 0.31 |
| 0.22 |
| 0.06 |
| 0.04 |
| 0.02 |
| -0.04 |
| -0.06 |
| -0.07 |
| -0.07 |
| -0.20 |
| -0.22 |
| -0.23 |
| -0.27 |
| -0.28 |
| -0.30 |
| -0.31 |
| -0.41 |
| -0.41 |
| -0.47 |
| -0.55 |
| -0.25 |
| 0.18 |
| 0.14 |
| 0.07 |
| 0.02 |
| -0.02 |
| -0.03 |
| -0.03 |
| -0.07 |
| -0.07 |
| -0.11 |
| -0.12 |
| -0.13 |
| -0.15 |
| -0.16 |
| -0.17 |
| -0.19 |
| -0.21 |
| -0.21 |
| -0.22 |
| -0.22 |
| -0.23 |
| -0.26 |
| -0.31 |
| -0.31 |
| -0.32 |

| |
|---|
| -0.32 |
| -0.33 |
| -0.35 |
| -0.36 |
| -0.38 |
| -0.41 |
| -0.41 |
| -0.43 |
| -0.44 |
| -0.45 |
| -0.48 |
| -0.52 |
| -0.56 |
| -0.58 |
| -0.62 |
| -0.66 |
| -0.74 |
| -0.75 |
| -0.81 |
| -0.91 |
| -0.97 |
| -1.02 |
| -1.12 |
| -1.15 |
| -1.55 |
| 0.40 |
| 0.30 |
| 0.04 |
| -0.06 |
| -0.11 |
| -0.15 |
| -0.16 |
| -0.26 |
| -0.29 |
| -0.30 |
| -0.54 |
| 2.06 |
| 0.04 |
| -0.05 |
| -0.09 |
| -0.09 |
| -0.10 |
| -0.17 |
| -0.30 |
| -0.43 |
| -0.45 |
| -0.52 |
| -0.58 |

-0.75
0.76
0.47
0.27
-0.08
-0.19
0.22
-0.37
-0.41
-0.45
-0.49
-0.49
-0.55
-0.57
-0.28
0.31
0.23
0.11
-0.30
0.63
0.22
-0.07
-0.10
-0.13
-0.17
-0.36
0.28
0.05
-0.05
-0.09
-0.18
-0.23
-0.24
-0.24
-0.28
-0.33
0.24
-0.18
-0.87
0.95
0.45
0.40
0.35
0.21
0.03
-0.12
-0.12
-0.14

| |
|---|
| -0.59 |
| 0.14 |
| -0.07 |
| -0.50 |
| -0.45 |
| 0.10 |
| 0.02 |
| -0.23 |
| -0.46 |
| 0.43 |
| -0.02 |
| 0.70 |
| 0.47 |
| 0.33 |
| 0.22 |
| 0.09 |
| 0.05 |
| 0.04 |
| 0.00 |
| -0.01 |
| -0.01 |
| -0.27 |
| -0.28 |
| -0.33 |
| -0.75 |
| 0.02 |
| -0.15 |
| -0.29 |
| 0.10 |
| 0.01 |
| -0.04 |
| -0.07 |
| -0.07 |
| -0.08 |
| -0.09 |
| -0.18 |
| -0.20 |
| -0.44 |
| -0.59 |
| 0.38 |
| 0.12 |
| 0.36 |
| 0.21 |
| -0.37 |
| -0.79 |
| 0.04 |
| 0.14 |
| 0.12 |

-0.15
-0.47
-0.56
0.39
0.23
0.14
-0.11
0.04
-0.09
-0.23
-0.33
0.20
-0.06
-0.12
-0.36
0.13
-0.06
-0.32
-0.46
0.36
1.57
0.52
0.22
0.20
0.70
0.59
0.30
0.24
-0.15
-0.23
-0.85
0.07
0.06
-0.20
-0.26
-1.08
0.14
0.04
0.03
-0.18
-0.25
-0.33
-0.39
-0.43
-0.46
0.27
0.23
0.17

| |
|---|
| 0.14 |
| 0.09 |
| 0.02 |
| 0.02 |
| 0.01 |
| -0.15 |
| -0.16 |
| -0.39 |
| -0.48 |
| 1.14 |
| 0.76 |
| 0.63 |
| -0.18 |
| -0.22 |
| -0.34 |
| -0.52 |
| 0.29 |
| 0.17 |
| 0.17 |
| 0.15 |
| 0.15 |
| 0.11 |
| 0.09 |
| 0.08 |
| 0.07 |
| 0.04 |
| -0.02 |
| -0.02 |
| -0.04 |
| -0.07 |
| -0.09 |
| -0.12 |
| -0.12 |
| -0.12 |
| -0.16 |
| -0.17 |
| -0.22 |
| -0.22 |
| -0.24 |
| -0.26 |
| -0.27 |
| -0.31 |
| -0.35 |
| -0.47 |
| -0.78 |
| -0.10 |
| 2.14 |
| 0.43 |

| |
|---|
| 0.20 |
| 0.11 |
| 0.01 |
| 0.00 |
| -0.02 |
| -0.03 |
| -0.03 |
| -0.04 |
| -0.13 |
| -0.15 |
| -0.16 |
| -0.17 |
| -0.17 |
| -0.18 |
| -0.20 |
| -0.21 |
| -0.22 |
| -0.23 |
| -0.24 |
| -0.30 |
| -0.33 |
| -0.36 |
| -0.41 |
| -0.46 |
| -0.46 |
| -0.46 |
| -0.49 |
| -0.50 |
| -0.46 |
| 0.25 |
| 0.35 |
| 0.06 |
| 0.05 |
| -0.02 |
| -0.05 |
| -0.12 |
| -0.16 |
| -0.18 |
| -0.43 |
| 0.30 |
| 0.17 |
| 0.07 |
| 0.05 |
| -0.05 |
| -0.05 |
| 0.18 |
| -0.11 |
| -0.46 |

| |
|---|
| -0.53 |
| -0.36 |
| 0.16 |
| 0.47 |
| -0.23 |
| 0.28 |
| 0.28 |
| 0.24 |
| 0.13 |
| 0.02 |
| -0.01 |
| -0.01 |
| -0.08 |
| 0.31 |
| 0.16 |
| 0.00 |
| 0.17 |
| 0.11 |
| -0.08 |
| -0.10 |
| 0.75 |
| 0.24 |
| 0.24 |
| 0.07 |
| 0.01 |
| -0.28 |
| 2.88 |
| -0.11 |
| 0.09 |
| 0.03 |
| 0.26 |
| 0.10 |
| -0.09 |
| -0.09 |
| 0.48 |
| 0.18 |
| -0.50 |
| 1.54 |
| 0.52 |
| 0.36 |
| 0.21 |
| -0.18 |
| -0.19 |
| 1.41 |
| 0.71 |
| 0.53 |
| 0.50 |
| -0.12 |

| |
|---|
| -0.04 |
| 0.65 |
| 0.13 |
| 0.21 |
| 0.00 |
| 0.68 |
| 0.29 |
| 0.26 |
| 0.24 |
| 0.23 |
| 0.14 |
| 0.08 |
| 0.06 |
| 0.03 |
| 0.00 |
| 0.00 |
| -0.13 |
| -0.16 |
| -0.17 |
| -0.19 |
| -0.24 |
| -0.54 |
| 1.52 |
| 1.16 |
| 1.12 |
| 0.31 |
| 0.28 |
| 0.19 |
| 0.06 |
| -0.14 |
| -0.63 |
| 0.26 |
| -0.16 |
| 0.27 |
| -0.19 |
| -0.40 |
| -0.06 |
| -0.17 |
| -0.20 |
| 0.37 |
| 0.23 |
| 0.07 |
| -0.07 |
| -0.07 |
| -0.13 |
| -0.27 |
| 0.07 |
| -0.10 |

-0.49
-0.15
0.00
-0.08
-0.29
-0.02
-0.10
-0.08
-0.32
-0.45
0.31
0.13
0.08
0.04
0.02
0.28
0.07
-0.12
-0.14
-0.19
-0.09
-0.05
-0.21
0.62
0.31
0.10
-0.08
-0.26
0.06
0.60
0.31
0.29
0.28
0.06
0.38
-0.37
0.13
0.03
0.13
0.11
-0.12
0.13
-0.04
-0.05
-0.20
0.75
0.32
0.23

| |
|---|
| 0.21 |
| 0.02 |
| -0.04 |
| -0.10 |
| -0.16 |
| -0.17 |
| -0.48 |
| 0.02 |
| -0.14 |
| -0.35 |
| 0.59 |
| 0.03 |
| 0.03 |
| 0.02 |
| -0.02 |
| -0.10 |
| -0.16 |
| -0.37 |
| -0.05 |
| -0.14 |
| 0.38 |
| 0.26 |
| 0.14 |
| 0.12 |
| 0.03 |
| 0.02 |
| -0.11 |
| -0.17 |
| -0.20 |
| -0.93 |
| 0.13 |
| -0.08 |
| -0.37 |
| -0.05 |
| -0.15 |
| -0.51 |
| -0.07 |
| -0.07 |
| -0.07 |
| -0.10 |
| -0.22 |
| -0.23 |
| -0.33 |
| -0.63 |
| 0.10 |
| -0.03 |
| -0.24 |
| 0.40 |

| |
|---|
| 0.83 |
| 0.73 |
| 0.72 |
| 0.28 |
| 0.04 |
| 0.04 |
| 0.02 |
| 0.02 |
| -0.01 |
| -0.02 |
| -0.06 |
| -0.08 |
| -0.08 |
| -0.11 |
| -0.15 |
| -0.23 |
| -0.31 |
| -0.33 |
| -0.35 |
| -0.44 |
| -0.47 |
| -0.47 |
| -0.58 |
| -0.62 |
| -0.05 |
| -0.14 |
| -0.12 |
| 0.07 |
| -0.05 |
| -0.06 |
| 0.09 |
| 0.02 |
| 0.08 |
| 0.02 |
| 0.00 |
| -0.10 |
| -0.26 |
| -0.26 |
| -0.30 |
| -0.39 |
| -0.47 |
| -0.03 |
| -0.13 |
| 0.13 |
| 0.09 |
| 0.19 |
| 0.05 |
| 0.05 |

| |
|---|
| 0.04 |
| -0.01 |
| -0.03 |
| -0.03 |
| -0.08 |
| -0.27 |
| -0.28 |
| -0.32 |
| -0.41 |
| -0.60 |
| 0.18 |
| -0.03 |
| -0.30 |
| -0.07 |
| 0.43 |
| 0.05 |
| 0.20 |
| 0.01 |
| 0.08 |
| 0.00 |
| -0.17 |
| -0.20 |
| -0.37 |
| -0.51 |
| -0.53 |
| -0.66 |
| -0.05 |
| -0.15 |
| -0.55 |
| 0.05 |
| 0.00 |
| 0.39 |
| 0.18 |
| 0.12 |
| 0.09 |
| -0.15 |
| 0.09 |
| 0.02 |
| 0.01 |
| -0.05 |
| -0.10 |
| -0.10 |
| -0.18 |
| -0.18 |
| -0.57 |
| -0.02 |
| -0.43 |
| 0.14 |

0.26
0.07
-0.11
-0.37
-0.40
